# Neurogenic colonic motility patterns and gut transit disrupted in a mouse model of Rett Syndrome

**DOI:** 10.64898/2026.09.29.755242

**Authors:** Nicole M. Procacci, Sushmita Debnath, Rajib Das, Emer Ni Bhraonain, Caroline A. Cobine, Thomas W. Gould

**Author notes:** Correspondence: Department of Cell Biology and Physiology University of Nevada School of Medicine 1664 N. Virginia St, Reno, NV 89554.

## Abstract

Gastrointestinal dysfunction is a prominent feature of neurodevelopmental disorders and contributes significantly to visceral discomfort, but the specific etiology is unclear. In particular, it is unknown which cell types within central, autonomic or enteric nervous systems, gut muscle, epithelium or immune system are the principal drivers of gut dysfunction. We report that the *Mecp2* mouse model of Rett syndrome, a neurodevelopmental disorder with profound gut dysmotility, exhibits impaired whole gut transit times and constipation *in vivo* as well as striking alterations to propagating neurogenic contractions in *ex vivo* colons. Selective elimination of *Mecp2* in neurons, but not glia or smooth muscle, recapitulates these dysmotility phenotypes. Bulk RNA sequencing of colonic external muscle, containing the myenteric compartment of the enteric nervous system, showed significant expression changes in specific neurochemical signaling pathways. Pharmacological modulation of one, the M1 acetylcholine receptor, modified the aberrant neurogenic contractile phenotype in mutant mice and reproduced it in wild-type mice. These results identify disruptions to neuronal muscarinic signaling in the enteric nervous system in a mouse model of Rett syndrome and provide an experimental strategy to identify translationally relevant cellular and molecular pathways underlying gastrointestinal dysmotility in neurodevelopmental disorders.

## Introduction

Rett syndrome (RTT) is an X-linked neurodevelopmental disorder (NDD) that is lethal in most boys and afflicts nearly 1:10,000 girls worldwide^1,2^. Girls develop normally until 6-18 months of age, when cognitive and motor skills begin to regress^1^. Although central nervous system pathology is prominent, most patients suffer from multisystem comorbidities, including seizures, breathing abnormalities, autonomic irregularities and gastrointestinal (GI) dysmotility^3^. GI issues cause considerable discomfort, compromising food intake, nutrition and health. While these observations suggest that GI dysmotility represents a key comorbidity of NDD, several studies implicate that such dysfunction itself may impact the core central cognitive and motor features of these disorders^4^. For example, manipulations of the gut microbiota appear sufficient to ameliorate the impaired social interactions in mouse models of and human patients with specific forms of autism spectrum disorder (ASD1)^5^. Together, these findings provide a compelling rationale to elucidate the mechanisms underlying GI dysmotility observed in distinct NDD: to remediate this dysfunction itself, and to improve the central components of these disorders.

GI motility is governed by myogenic and neurogenic mechanisms. Myogenic substrates include the SIP syncytium^6^, consisting of <u>s</u>mooth muscle cells (SMC), interstitial cells of Cajal (ICC), and <u>p</u>latelet-derived growth factor receptor-alpha (PDGFRα) cells functioning together as an electrically-coupled unit. Neurogenic control of GI motility is mediated by substrates intrinsic and extrinsic to the enteric nervous system (ENS). Intrinsic substrates within the myenteric plexus of the gut wall include intrinsic primary afferent neurons (intrinsic PANs or IPAN), which detect gut distention, interneurons that ascend and descend, and excitatory and inhibitory motor neurons^7^. Extrinsic autonomic substrates include efferent parasympathetic motor fibers from vagal and pelvic nuclei, which increase gut motility, efferent sympathetic motor fibers from prevertebral ganglia, which decrease gut motility, and extrinsic PAN residing within spinal and cranial ganglia. In addition to these core myogenic and neurogenic substrates, GI motility is also regulated by enteric submucosal neurons and bacterially derived activators of the innate immune system that regulate mucosal function and fecal pellet composition.

Most RTT cases are caused by loss-of-function mutations in the methyl CpG binding protein 2 (*MECP2* in human; *Mecp2* in mouse) gene^8^. This gene encodes an epigenetic transcriptional regulator (MeCP2 in human; Mecp2 in mouse^9^) that is abundant in the CNS, where it mediates synaptic development and plasticity^10^. Mecp2 is also found in many cells outside of the CNS, where less is known about its function^11^. For example, within the gut, Mecp2 expression has been detected in the mucosal epithelium in cells at the base of the crypts in the colon, but not small intestine^12^, in SMC^13^ and in neurons of the myenteric plexus^14^ throughout the entire intestine. The extent to which these expression patterns drive the GI dysfunction observed in RTT is unclear. Constipation is the most frequently reported GI symptom, occurring in 80% of patients with RTT, and dysbiosis of both bacterial and fungal gut microbiota, resulting in altered short chain fatty acid levels, has been detected^15^, implicating the colon as a potential source of GI dysmotility. A constitutive *Mecp2* null mouse model of RTT, in which the *Mecp2* gene is deleted in all cell types, recapitulates key features of RTT, including impaired GI transit^16,17^. Electrogenic absorption and secretion are not affected in the colonic epithelia of these mice, and selective ablation of the *Mecp2* gene in intestinal epithelial cells fails to reproduce delayed GI transit^12^, but the myenteric circuits underlying motility have not been examined.

In the present study, we observed delayed transit, signs of constipation, and alterations to neurogenic but not myogenic motility patterns in *ex vivo* preparations of colon derived from *Mecp2* null mutant mice. These patterns were also observed when *Mecp2* was selectively deleted from neurons. In colonic muscle tissue that includes the myenteric plexus, a number of differentially-expressed neuronal factors were detected by bulk RNA sequencing in *Mecp2* mutant vs. WT mice, including the muscarinic acetylcholine (ACh) receptor (M1R), the pharmacological modulation of which reproduced these aberrant colonic motility patterns in WT mice. Together, these studies shed light into the cellular and molecular features of a novel form of neurogenic dysmotility in an animal model of RTT, a NDD with prominent GI pathophysiology.

## Results

Male *Mecp2* null mutant mice (hemizygous; Mecp2^-/y^), which cannot breed, thus precluding the generation of female null mutant mice, begin to exhibit mortality at 8 weeks of age in the C57Black 6 (C57BL/6) strain^16^. We observed that null mutants were still alive at this timepoint, but all died by 12 weeks of age. Similar to a previously published report^17^, *Mecp2* null mutants exhibited a delay in whole gut transit time (WGTT) after intragastric gavage of Evans Blue dye, relative to littermate controls (***Figure 1A***). This delay occurred despite a significant reduction of overall intestinal length (***Figure 1A***). Typically, prolongation of transit is associated with enhanced water resorption and harder stools. However, the only report of fecal pellet composition in *Mecp2* null mutant mice anecdotally described watery stool^12^. Accordingly, we examined fecal pellet output and composition. Over the 12-hour dark period, the total number of fecal pellets was significantly higher, but the total dry weight of these pellets was similar, to those of control mice (***Figure 1B***). This pair of results produced a significant reduction in the weight of each individual dry pellet (9.1±2.9 vs. 5.7±1.3 mg, *Mecp2* wild-type (WT) vs. knockout (KO) mice, *p*=0.0001, *n*=16). These smaller fecal pellets produced by KO mice were narrower than those of WT mice. In addition, the gross appearance of fecal pellets derived from *Mecp2* KO mice was darker than that of those from control mice (***Figure 1C***). Together, the slower transit as well as smaller size and darker appearance of fecal pellets in *Mecp2* null mutants suggest a reduction of pellet water content due to prolonged water absorption. However, water content was higher in fecal pellets produced by *Mecp2* KO than those by WT mice (***Figure 1D***). For these experiments, we collected pellets immediately after singly caging mice. We observed that *Mecp2* null mutant mice frequently produced ∼5 wet pellets, of a consistency approaching diarrhea, in the first fifteen minutes, whereas control mice typically produced 0-1 pellets during this initial period. Therefore, we waited until the second hour to collect pellets. In these comparisons, the pellets derived from *Mecp2* KO mice contained significantly less water per pellet (***Figure 1D***). Together, these studies demonstrate a delay of intestinal motility, coupled with the eventual production of smaller, darker and less hydrated fecal pellets in symptomatic *Mecp2* null mutant mice.

**Figure 1.**
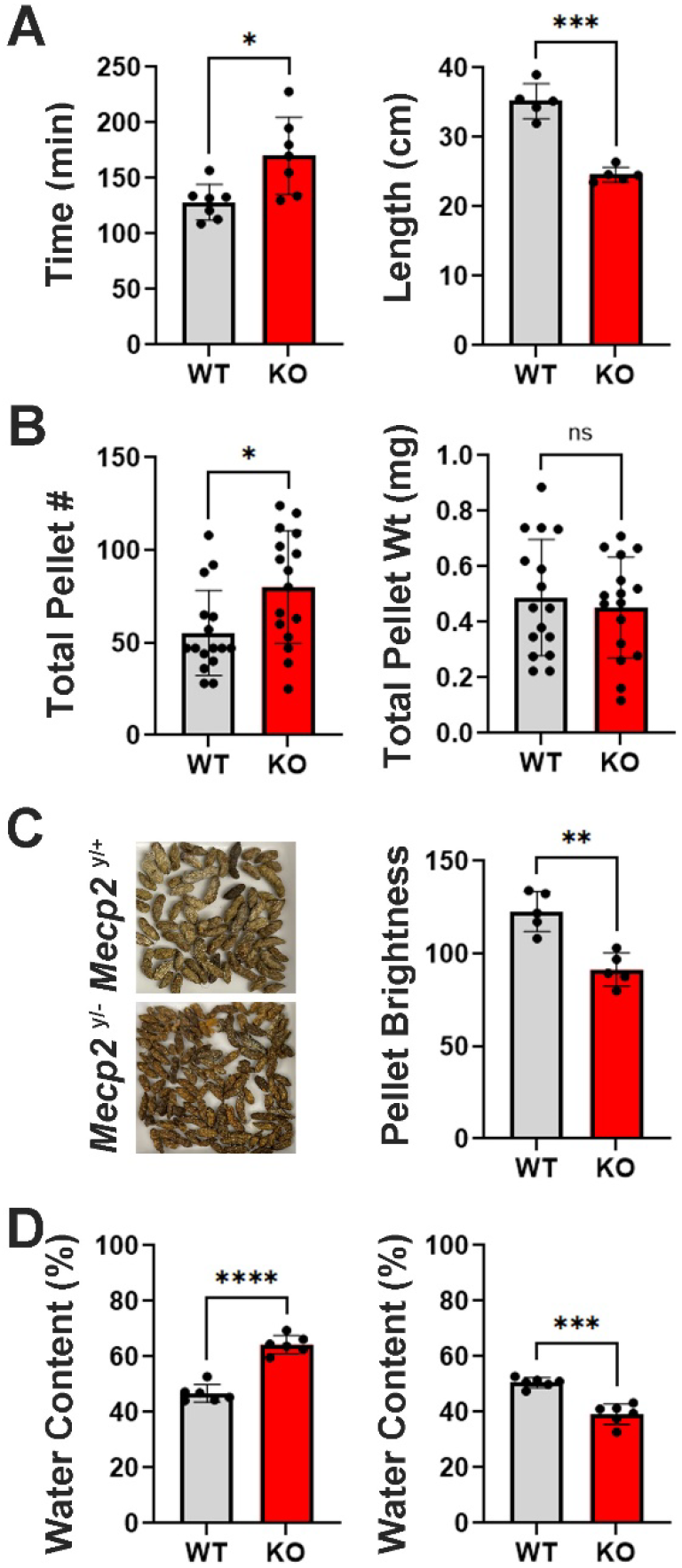
*Mecp2* null mutant (knockout; KO) mice display signs of impaired gastrointestinal function, relative to *Mecp2* wild-type (WT) mice. ***A*,** Left, Whole gut transit time was delayed in *Mecp2* KO vs. WT mice, \**p*=0.0188, *n*=7). Right, Intestinal length was shorter in *Mecp2* KO vs. WT mice, \*\*\**p*=0.0004, *n*=6). *B*, Total number of fecal pellets excreted in twelve-hour period was higher in KO vs. WT mice, \**p*=0.0147, *n*=16). However, total dry weight of these pellets was similar, *p*=0.6085, *n*=16). *C*, Fecal pellets were darker in KO vs. WT mice, \*\**p*=0.0011, *n*=5). *D*, In the first hour, fecal pellet water content was higher in KO than WT mice, \*\*\*\**p*=2.9*10e6, *n*=6), but in the second hour, water content was lower in KO than WT mice, \*\*\**p*=5.1*10e5, *n*=6).

A detailed Mecp2 description in the gut is lacking. We found that Hu C/D-immunoreactive myenteric neurons all expressed Mecp2 (99.1±1%, *n*=5; ***Figure 2A***). Smaller Mecp2-immunoreactive cells in myenteric ganglia that lacked Hu C/D also expressed Mecp2. Accordingly, we labeled enteric glial cells (EGC) genetically with *Kir4.1-CreERT2* mice and found that the vast majority expressed Mecp2 (95.6±1.2%, *n*=5). Enteric neurons and glia of the colonic submucosal ganglia also expressed Mecp2 (96.4±2.5%, *n*=5; ***Figure 2C***). We next examined neurons within the extrinsic circuitry that innervates the GI system. As Mecp2 expression has been reported in all spinal sensory neurons, including peptidergic and non-peptidergic subtypes^18^, and in cranial sensory neurons in the vagal nodose ganglia^19^, we focused on efferent gut-innervating motor neurons. We detected Mecp2 expression in all Hu C/D-positive prevertebral sympathetic neurons labeled genetically with *TH-Cre* mice (94±2.7%, *n*=5; ***Figure 3***). We also observed Mecp2 expression in nitrergic sacral spinal neurons (95±11%, *n*=5), which represent efferent neurons originally characterized as parasympathetic based on function but more recently defined as sympathetic based on developmental molecular specification^20^. Together, these studies show that Mecp2 expression is widespread in neurons and glia of the intrinsic and extrinsic neurogenic systems that drive GI motility.

**Figure 2.**
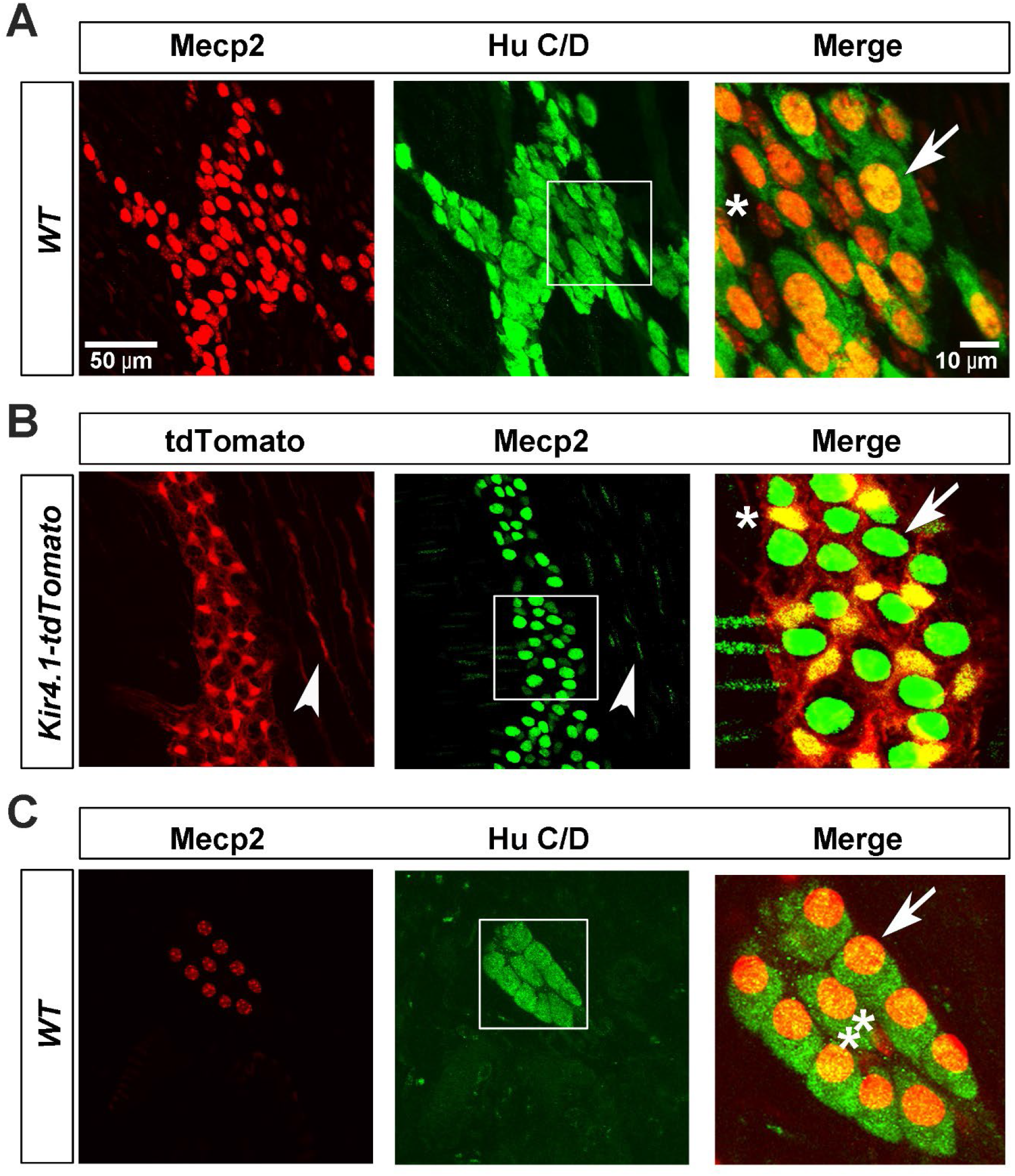
Neurons and glia of the colonic myenteric and submucosal plexi express Mecp2. ***A***, Mecp2 (red) is expressed by Hu C/D+ neurons (green) in myenteric plexus. Arrow in inset points to double-positive neuron, asterisk points to Mecp2-positive, non-neuron. ***B***, Mecp2 (green) is expressed by enteric glial cells (EGC) labeled genetically with *Kir4.1-tdTomato* mice (red). Arrow in inset points to Mecp2-positive, tdTomato-negative myenteric neuron, asterisk points to double-positive intraganglionic EGC. In left and middle images, arrowhead points to double-positive intramuscular EGC. ***C***, Mecp2 is expressed by Hu C/D+ neurons (arrow) as well as presumptive EGC (asterisks) in submucosal plexus. Percentages of labeled intrinsic neurons and glia in text. Images in left and middle columns, and enlarged images from inset boxes in right columns in ***B***,***C***, are at identical magnification to those in **A**.

**Figure 3.**
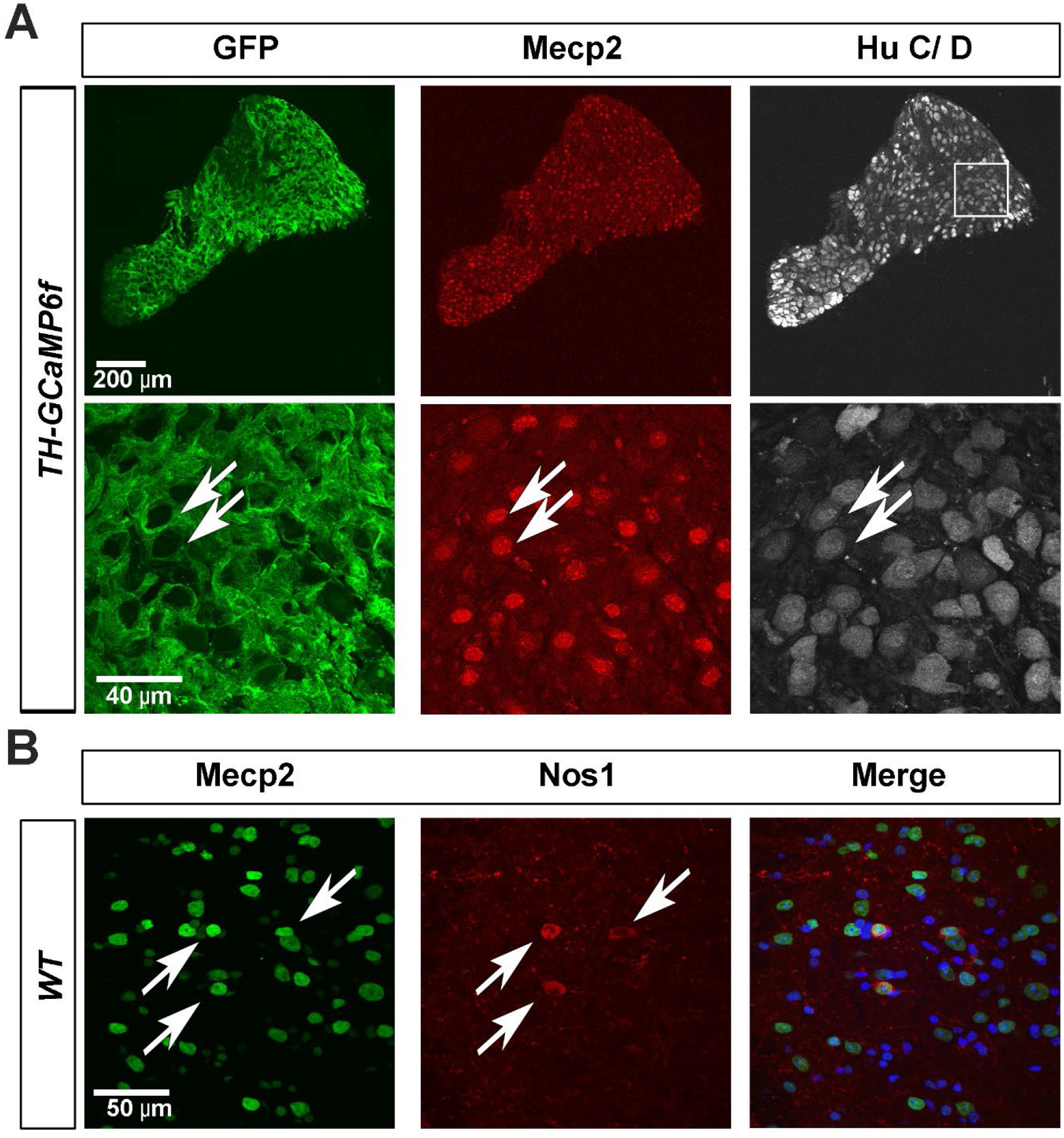
Neurons and glia of the sympathetic prevertebral ganglia and parasympathetic sacral intermediate gray matter express Mecp2. ***A***, Top row, representative image of a whole-mount preparation of a prevertebral ganglion taken from a *TH-Cre, Rosa26-LSL-GCaMP6f* (*TH-GCaMP6f*) mouse stained with antibodies against GFP to label GCaMP6f-expressing sympathetic neurons (left, green), Mecp2 (middle, red) and the pan-neuronal marker Hu C/D (right, white). Bottom row, in an amplified region of each of these images, the arrows point to a triple-positive neuron within this sympathetic ganglion. ***B***, Representative image of a cryosection of the sacral spinal cord from a WT mouse stained with antibodies against Mecp2 (left, green) and Nos1 (middle, red) to label colon-projecting parasympathetic neurons. Arrows point to double-positive neurons, and the merged image on the right shows the Hoechst-stained nuclei of these and nearby spinal cells. Percentages of labeled extrinsic neurons presented in text.

We then investigated expression of Mecp2 in myogenic substrates of colon, including ICC, PDGFRα-positive cells and SMC. We identified ICC in the myenteric and intramuscular layers (ICC-IM and ICC-My, respectively) by staining *Kit^CopGFP^* mice with GFP and Mecp2 antibodies. We used Mecp2-immunoreactive enteric neurons as a landmark to find ICC-My and arrays of cigar-shaped, Hoechst-labeled nuclei to label circular (CM) and longitudinal (LM) muscle layers. ICC-My did not express Mecp2 (1.7±1.5%, *n*=5), but a small population of ICC-IM were Mecp2-immunoreactive (11.5 ± 3.9%, *n*=5; ***Figure 4***). ICC along the submucosal surface of the CM (ICC-SM) also failed to express Mecp2 (*n*=5). Next, we identified PDGFRα-positive cells by staining *PDGFRα-nGFP* mice with GFP and Mecp2 antibodies. 35.5±6.4% of this cell type expressed Mecp2 within circular CM and LM smooth muscle layers. Finally, we identified SMC by staining *Myh11^Cre/GFP^*mice with GFP and Mecp2 antibodies. Mecp2 was expressed by nearly all SMC within CM and LM layers of external muscle (CM: 99±1.4%, LM: 97.4±2.7%, *n*=5). These experiments demonstrate that SMC are the predominant source of colonic muscle tissue-derived Mecp2 of the colon.

**Figure 4.**
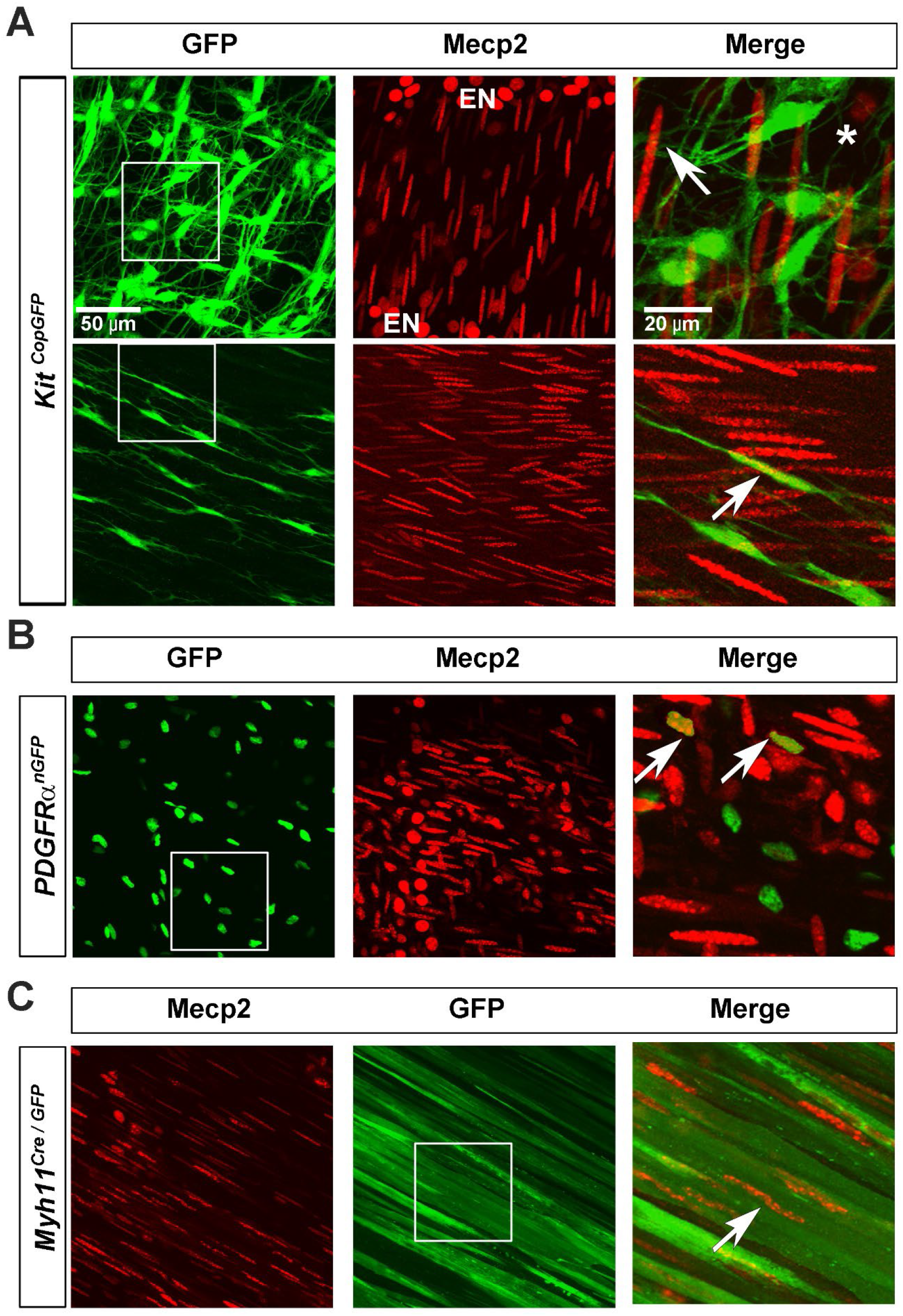
Colonic smooth muscle cells and a subpopulation of PDGFRα cells, but not ICC, express Mecp2. ***A***, Representative images of a whole-mount preparation of the external muscle layer taken from a *Kit^CopGFP^*mouse stained with antibodies against GFP to label GFP-expressing ICC (left, green) and Mecp2 (middle, red) in the region near enteric neurons (EN) of the myenteric ganglia that contain ICC of the myenteric plexus or ICC-My (top row) and in the region within the circular muscle (CM) that contain intramuscular ICC or ICC-IM (bottom row). In the amplified, merged image on the right of the top row, taken from the inset box shown in the left image, the arrow points to a Mecp2-positive, GFP-negative cell with a cigar-shaped immunoreactive nucleus (presumptive smooth muscle cell or SMC, see below), and the asterisk points to a Mecp2-positive, GFP-negative cell with an oval-shaped nucleus (presumptive PDGFRα cell; see below in ***B***), as well as GFP-positive, Mecp2-negative ICC-My cells. In the amplified, merged image on the right of the bottom row, taken from the inset box shown in the left image, the arrow points to a Mecp2-positive, GFP-positive ICC-IM cell as well as many Mecp2-positive, GFP-negative cells with cigar-shaped nuclei that are likely SMC (see below in ***C***). ***B***, Representative images of a whole-mount preparation of the external muscle layer taken from a *PDGFRα^nGFP^* mouse stained with antibodies against GFP to label GFP-expressing *PDGFRα* cells (left, green) and Mecp2 (middle, red). In the amplified, merged image on the right, taken from the inset box shown in the left image, the arrows point to Mecp2-positive, GFP-positive cells with oval-shaped immunoreactive nuclei. ***C***, Representative images of a whole-mount preparation of the muscularis externa taken from a *Myh11^Cre/GFP^* mouse stained with antibodies against GFP to label GFP-expressing SMC (middle, green) and Mecp2 (left, red). In the amplified, merged image on the right, taken from the inset box shown in the middle image, the arrow points to a Mecp2-positive, GFP-positive cell with a cigar-shaped immunoreactive nucleus. Percentages of labeled ICC subtypes, PDGFR*α* cells, and SMC presented in text. Images from lower row in left and middle columns of ***A***, as well as images in left and middle columns of ***B*** and ***C***, are at identical magnification to those in left and middle columns of upper row in ***A***. Enlarged image in right column of lower row in ***A***, as well as in right column of ***B*** and ***C***, are at identical magnification to that in right column of upper row in **A**.

Macrophages in the external muscle layer regulate GI motility through several signaling pathways including one involving macrophage-derived bone morphogenetic protein^21^. Mecp2 expression was observed in macrophages isolated by sorting from whole gut tissue^22^. When we stained colonic muscle with F4/80 antibodies to label macrophages, Mecp2 immunoreactivity was not detected within them (***Figure 5***). By contrast, at the base of the crypts of colonic mucosa, Mecp2-immunoreactive cells co-expressed F4/80. Mecp2 was also detected in other uncharacterized cells within crypts of Lieberkühn that include enterocytes, goblet cells and enteroendocrine cells (*n*=4). Finally, we noted Mecp2 expression in cells with spherical nuclei within serosa that likely represent epithelial cells (*n*=8). Collectively, these studies demonstrate widespread expression of Mecp2 in intrinsic and extrinsic neurons and glia as well as in SMC, in subtypes of PDGFRα cells and mucosal macrophages, in unidentified cells of the crypts, and in serosal epithelium.

**Figure 5.**
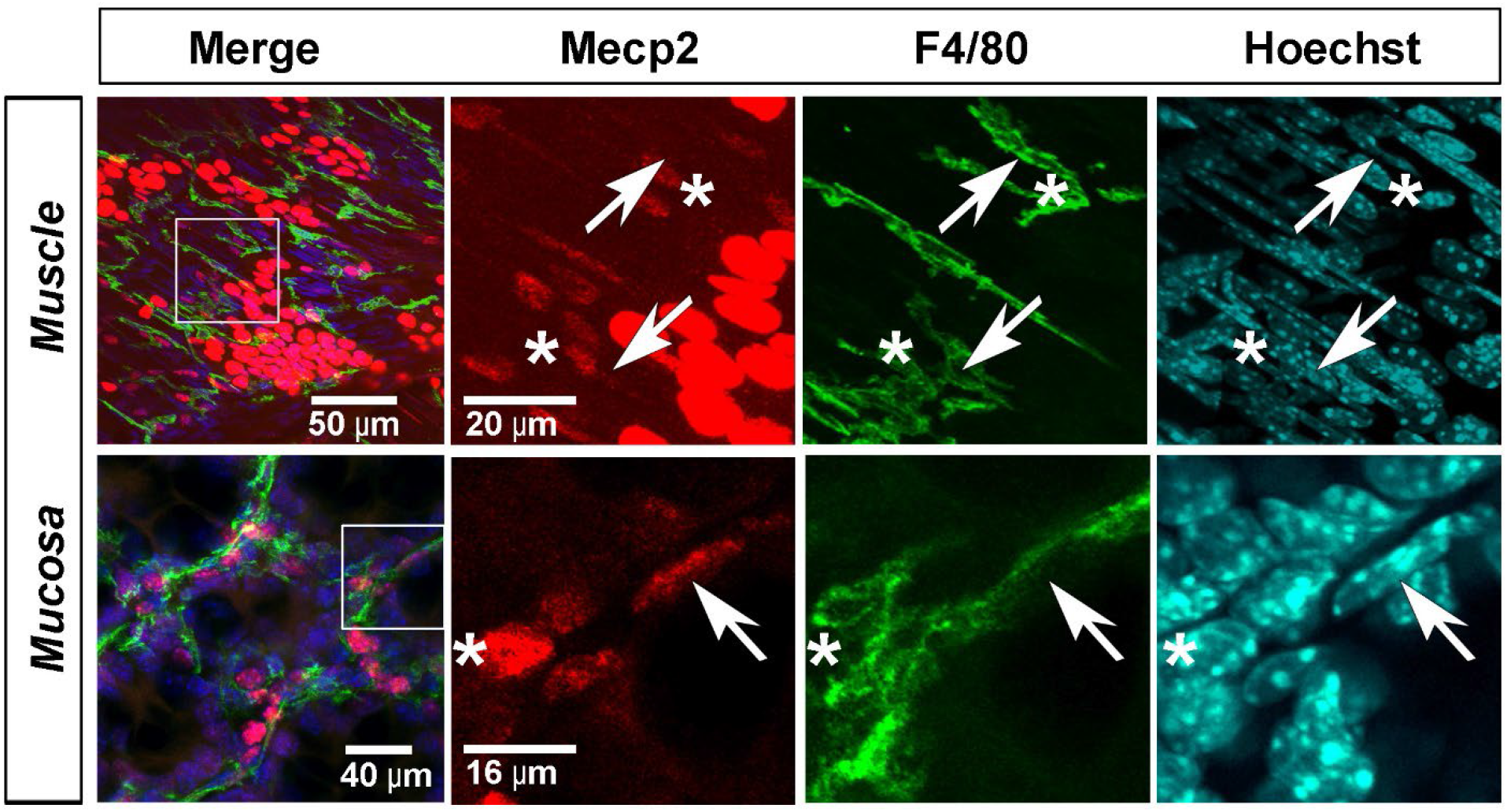
Mucosal but not muscularis externa-derived macrophages express Mecp2. ***A***, Representative image of a whole-mount preparation of the external muscle layer of the middle colon taken from a WT mouse stained with antibodies against Mecp2 (red) and the macrophage marker F4/80 (green) as well as with the nuclear stain Hoechst (light blue). In the merged image on the left, F4/80-immunoreactive macrophage can be seen running parallel to SMC in the general vicinity of intensely Mecp2-immunoreeactive myenteric ganglia. In the amplified images, taken from the inset box shown in the left image, the arrows point to the Hoechst-stained nuclei of F4/80-positive, Mecp2-negative macrophage (positive for green, negative for red), and the asterisks point to Mecp2-positive, F4/80-negative cells with oval-shaped nuclei that are likely PDGFRα cells. Note as well the presence of Mecp2-positive, F4/80-negative cells with cigar-shaped nuclei that are likely SMC. ***B***, Representative image of a whole-mount preparation of the submucosal surface of the mucosa, with the submucosa removed, a WT mouse stained with the same antibodies and Hoechst dye. In the merged image on the left, the Hoechst labeling shows the ring-shaped base of a colonic crypt of Lieberkuhn. Mecp2-immunoreactive cells are observed here, and the arrow in the amplified images, taken from the inset box shown in the left image, points to the Hoechst-stained nucleus of a Mecp2-positive, F4/80-positive macrophage, while the arrow points to the nucleus of a Mecp2-positive, F4/80-negative cell in the same vicinity.

Since Mecp2 expression was observed by both neurogenic and myogenic cell types that regulate GI motility, we examined motility patterns driven by each of them in *ex vivo* preparations of the colon. We first crossed these mutants to the transgenic *CAGGS-GCaMP3* line expressing the genetically-encoded calcium indicator GCaMP3 in all cell types to evaluate ripples, the phasic contractile pattern generated by ICC-SM that represent the predominant myogenic movement pattern observed in between neurogenic contractions in un-paralyzed imaging preparations of whole colon^23^. No differences between WT and KO were observed in ripple frequency or in the amplitudes of SMC Ca^2+^ responses during ripples (***Figure 6A,B***), despite the robust expression of Mecp2 in the SMC that produce them.

**Figure 6.**
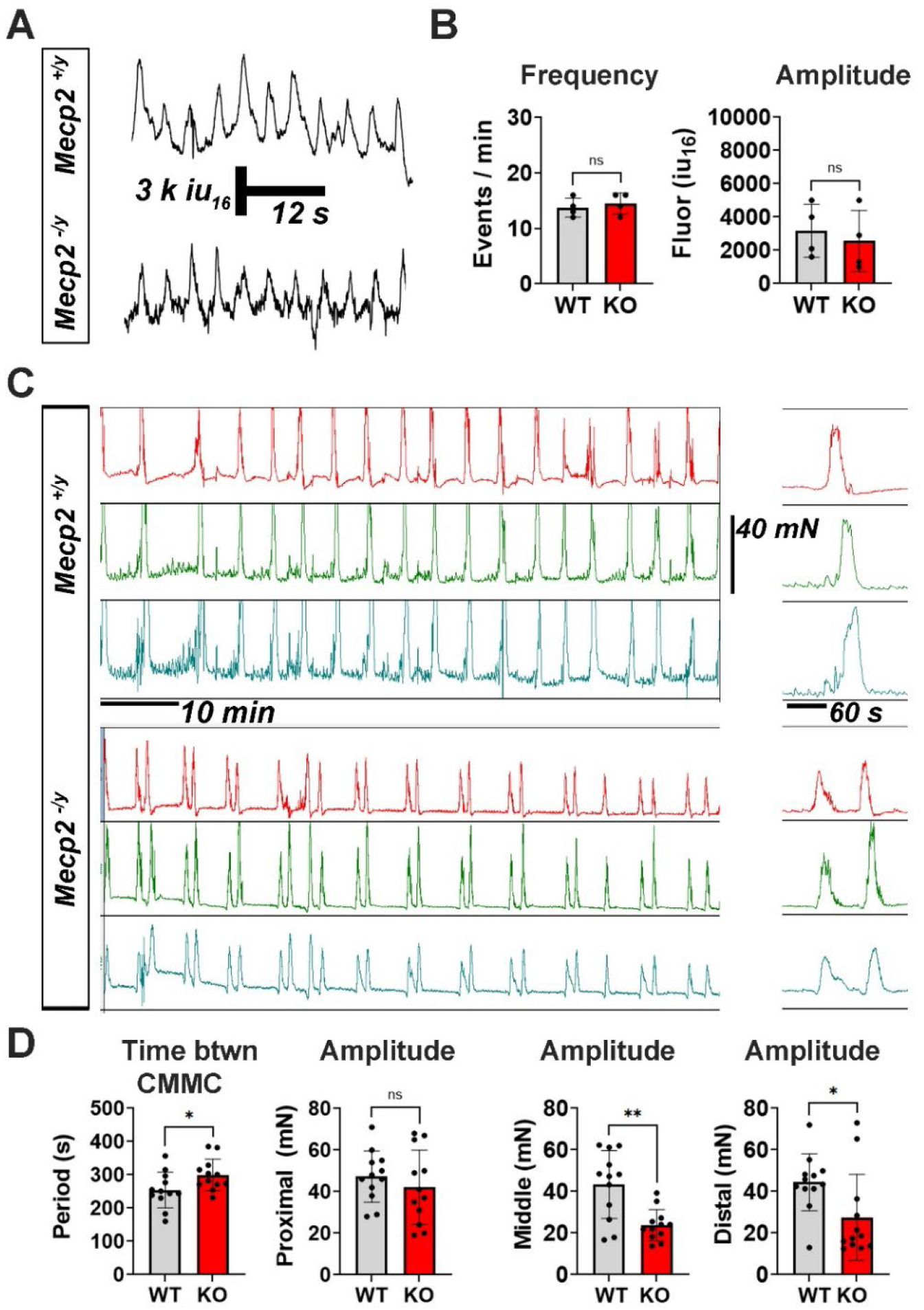
*Mecp2* KO mice exhibit alterations in neurogenic but not myogenic colonic motility patterns. ***A*,** Ca^2+^ transients within circular muscle of middle colon of *CAGGS-GCaMP3* mouse crossed to WT (upper trace) or KO mice (bottom trace) during myogenic ripple contractions. ***B***, Frequency and amplitude of myogenic Ca^2+^ responses were similar between genotypes (*p*=0.63, *n*=4). ***C***, Colonic migrating motor complexes (CMMC) of *Mecp2* WT (upper) and KO (lower) mice on left, individual (WT) or doublet contractions (KO) magnified on right. Note doublet CMMC in KO but not WT; Red top trace represents recording from chevron-containing region of proximal colon, green middle trace from middle colon, and blue trace from distal colon, of same animal. ***D***, If each doublet contraction-containing CMMC was measured as a single CMMC in KO, CMMC frequency was lower in KO than in WT (frequency measured by time period in between (btwn) CMMC); for these results, \**p*=0.0383, *n*=12. Middle (\*\**p*=0.001) and distal (\**p*=0.028) contractile amplitudes, but not proximal contractile amplitudes (*p*=0.429), were significantly lower in KO than in WT mice (*n*=12).

Next, we examined colonic migrating motor complexes (CMMC), a propagating neurogenic motility pattern thought to contribute to peristalsis *in vivo*. Strikingly, *Mecp2* null mutant mice exhibited a unique doublet pattern of these contractions (***Figure 6C***). The time between each propagating contraction within the doublet was 99.3±11.2 s (*n*=12). If these paired contractions were treated as a single CMMC, then the frequency of this colonic motility pattern was lower in KO vs. WT mice (***Figure 6D***). Each contraction within the doublet propagated the length of the colon and exhibited normal durations (54.6 ± 6.7 vs. 57.1 ± 7.4 s; WT vs. KO; *p*=0.399; *n*=12), proximal-to-middle (P-M) propagation times (22.1 ± 7.2 vs. 17.3 ± 6.7 s; WT vs. KO; *p*=0.109; *n*=12) and middle-to-distal (M-D) propagation times (14.2 ± 6.9 vs. 11.6 ± 4.9 s; WT vs. KO; *p*=0.297; *n*=12). However, although the proximal amplitudes of these contractions were similar, the middle and distal contractile amplitudes were significantly reduced in KO vs. WT mice (***Figure 6D***). All twelve mutant mice exhibited at least a single doublet. Most (8/12) exhibited doublets that occurred for over half of the two-hour recording period. In some cases, triplets and even quadruplets (i.e., contractions separated by ∼100 s intervals) were observed (***Supplemental Figure 1***). Doublets were abolished by blockade of intraganglionic nicotinic transmission with 100 μM hexamethonium (*n*=7; ***Supplemental Figure 1***). These studies suggest that intrinsic, myenteric neuronal circuits underlying CMMC may be altered in KO mice, as they were performed in *ex vivo* preparations of intact colon lacking extrinsic inputs.

Accordingly, we first characterized myenteric neuron numbers and found no difference between genotypes (534±185 vs. 544±187 neurons per mm^2^ when fields selected randomly, WT vs. KO, *p*=0.306; 645±70 vs. 695±74 neurons per mm^2^ when field percentage containing ganglia was normalized, WT vs. KO; *p*=0.306; *n*=5; see methods). Numbers of cholinergic and nitrergic neurons in *Mecp2* null mutant mice, which together account for over 90% of all colonic myenteric neurons, were also similar (***Figure 7A,B***). The overall average area of cell body or soma, and the frequency distributions of these areas, were also similar for both cholinergic and nitrergic subtypes (***Figure 7A,B***). This lack of effect of *Mecp2* deletion on neuronal size in the ENS contrasts with smaller-sized neurons observed in the CNS of these mice^24^.

**Figure 7.**
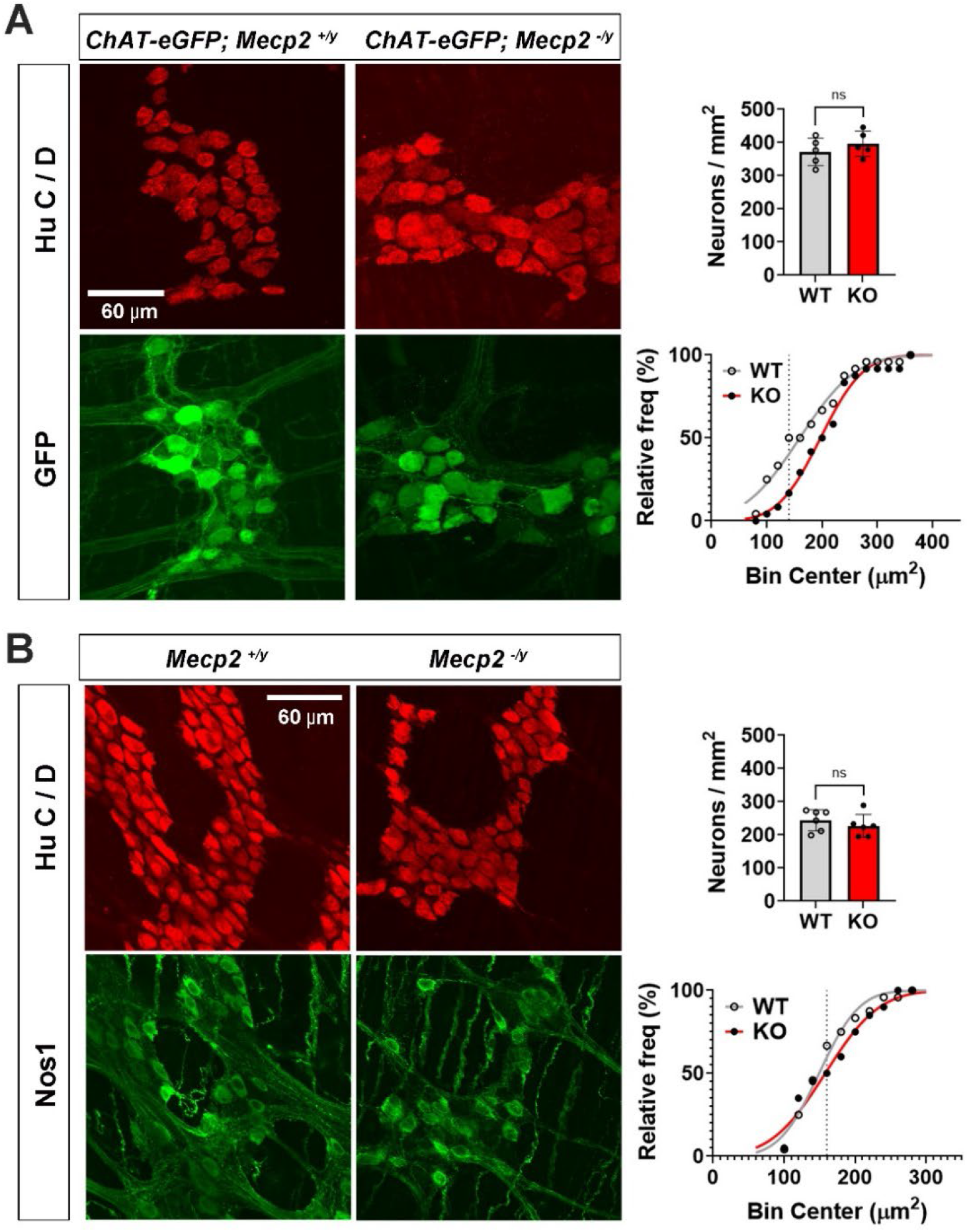
No effect on colonic myenteric neuronal number, size or subtype in *Mecp2* KO mice. ***A*,** Left, cholinergic neurons (green) and total Hu C/D+ neurons (red) in *Mecp2* WT (left) and KO (right) mice crossed to *ChAT-eGFP* mice. Right upper, neuron numbers were similar between genotypes, *p*=0.369, *n*=5. Right lower, distributions of cholinergic cell somal areas were also similar between genotypes: *p*=0.139, *d*=0.33 at 140 μm^2^; *c*=140 (WT) and *c*=96 (KO), *n*=7. Note trend toward larger cholinergic neurons in lower half of cumulative frequency distribution in KO vs. WT mice. ***B*,** Left, nitrergic neurons (green) and total Hu C/D+ neurons (red) in WT (left) and KO (right) mice. Right upper, nitrergic neuron numbers were similar between genotypes, *p*=0.397, *n*=5. Right lower, distributions of nitrergic cell somal areas were also similar between genotypes, *p*=0.775, *d*=0.2 at 160 μm^2^, *c*=110 (WT) and *c*=90 (KO), *n*=6. Images in right columns are at identical magnification to those in left columns.

Neurogenic but not myogenic motility patterns were altered in *Mecp2* null mutant mice, but whether these effects reflected disruptions to neuron- or muscle-derived signaling pathways was unclear. Indeed, a role for non-cell autonomous signaling from glia to neurons in *Mecp2* KO mice has been established^25^, and diffusible signals released by muscle regulate neurogenic motility patterns in C. Elegans^26^. We first crossed *Baf53b-Cre* mice to conditional *Mecp2* mutant mice (*Mecp2^Flox/y^*) in order to eliminate Mecp2 expression from neurons. In colonic muscle from these mice, Mecp2 was deleted from 92.7 ± 2.6% (*n*=5) Hu C/D-positive myenteric neurons but not from adjacent enteric glia or any other cell types in muscle (***Figure 8A***). Similar to constitutive KO mice, neuron-specific KO mice produced smaller fecal pellets (5.6±1.8 vs. 10±1.3 mg, *Baf53b-Mecp2^Flox/y^*vs. *Baf53b-Mecp2^+/y^*, *p*=0.013, *n*=4). However, whether these effects resulted from deletion in central and / or autonomic neurons, vs. enteric neurons, is unclear. Therefore, we examined CMMC in *ex vivo* preparations of intact colon, in which the only neuronal population present is enteric neurons. We observed doublet contractions in four of five *Baf53b-Mecp2^flox/y^* male mice (***Figure 8D***), suggesting that this alteration was triggered by loss of enteric neuronal-derived Mecp2. Middle (*p*=0.005) and distal (*p*=0.002), but not proximal (*p*=0.47), contractile amplitudes, were also reduced in these mice (*n*=5). By contrast, when *Mecp2* was genetically removed from glial cells or SMC in *Kir4.1*-*Mecp2^flox/y^* mice (*n*=3; recombination efficiency = 94.7 ± 2.5%) or *Myh11-Mecp2^flox/y^* mice (*n*=5; recombination efficiency = 81 ± 10.1%), these alterations to CMMC were rarely observed (***Figure 8B-D***).

**Figure 8.**
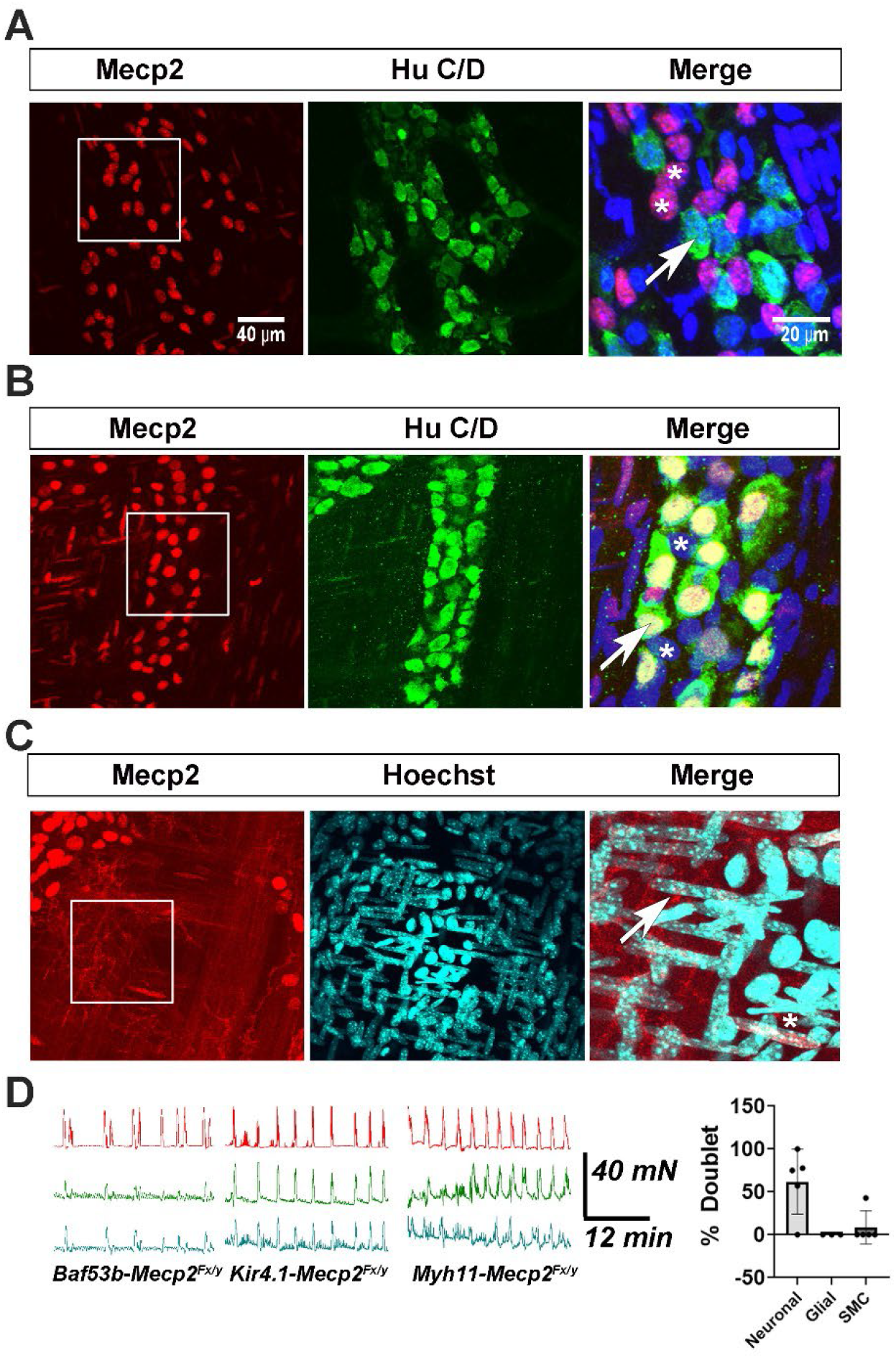
Genetic elimination of *Mecp2* within neurons but not within enteric glial or smooth muscle cells produces alterations of colonic neurogenic motility patterns. ***A***, *Baf53b-Mecp2^Flox/y^* mouse colon labeled with Mecp2 (red) and Hu C/D (green). In merged inset, arrow points to Hu C/D+, Mecp2-negative neuron, and asterisks denote Hu C/D-, Mecp2+ presumptive EGC. *B*, Similar image from a *Kir4.1-Mecp2^Flox/y^* mouse; arrow in right points to Hu C/D+, Mecp2+ neuron, asterisks denote Hu C/D-negative, Mecp2-negative EGC. ***C*,** Similar image from *Myh11-Mecp2^Flox/y^* mouse, stained with Mecp2 (red) and Hoechst. In merged inset, most cigar-shaped nuclei of presumptive SMC lack Mecp2 (e.g., arrow) although some SMC (e.g., asterisk) express Mecp2. Images in left and middle columns in ***B*** and ***C***, as well as enlarged images in right column in ***B*,** and ***C***, are at identical magnification as those in ***A*.*D***, Representative CMMC from *Baf53b*-*Mecp2^Flox/y^, Kir4.1*-*Mecp2^Flox/y^*, and *Myhll-Mecp2^Flox/y^* mice. Graph, CMMC % that exhibited doublets within 30-minute period in each line (*Baf53b*-*Mecp2^Flox/y^*(*n*=5), *Kir4.1*-*Mecp2^Flox/y^* (*n*=3), and *Myhll-Mecp2^Flox/y^* mice (*n*=5).

The demonstration that *Mecp2* deletion selectively from neurons produced the same alterations to motility in *ex vivo* colon preparations that were observed in response to constitutive *Mecp2* deletion, together with the absence of gross structural differences to myenteric neurons in constitutive KO mice, suggests that myenteric neuron circuits driving motility may be disrupted. In order to gain insight into these circuits, we probed the transcriptomes of the external muscle layer of proximal colon of KO and WT mice. The proximal colon was chosen simply because doublet contractions were observed in this region, where propagating neurogenic contractions initiate. If genes were considered statistically differentially expressed at a q-value < 0.05 and an absolute log2 fold change (lfc) ≥ 0.6, then 526 genes were downregulated in *Mecp2* KO samples, and 72 were upregulated (***Figure 9A,B*, *Figure 10, Table S1***). Using iPathwayGuide, we detected a variety of metabolic pathways, as well as the neuroactive ligand receptor interaction pathway, dysregulated in *Mecp2* KO mice (***Figure 9C***). Based on the genetic evidence above, we reasoned that the neuroactive ligand receptor interaction pathway might contribute to doublet CMMC (***Figure 9D***). Indeed, a number of genes from this pathway were within the most differentially regulated between genotypes, based on strength of adjusted p-values (***Figure 9E***) We also compared the dataset in this study with 1,261 genes identified as implicated in ASD by the Simons Foundation for Autism Research Initiative (SFARI) and found that 42 of the 598 differentially expressed (DE) genes are shared with this data set (***Figure 9F***).

**Figure 9.**
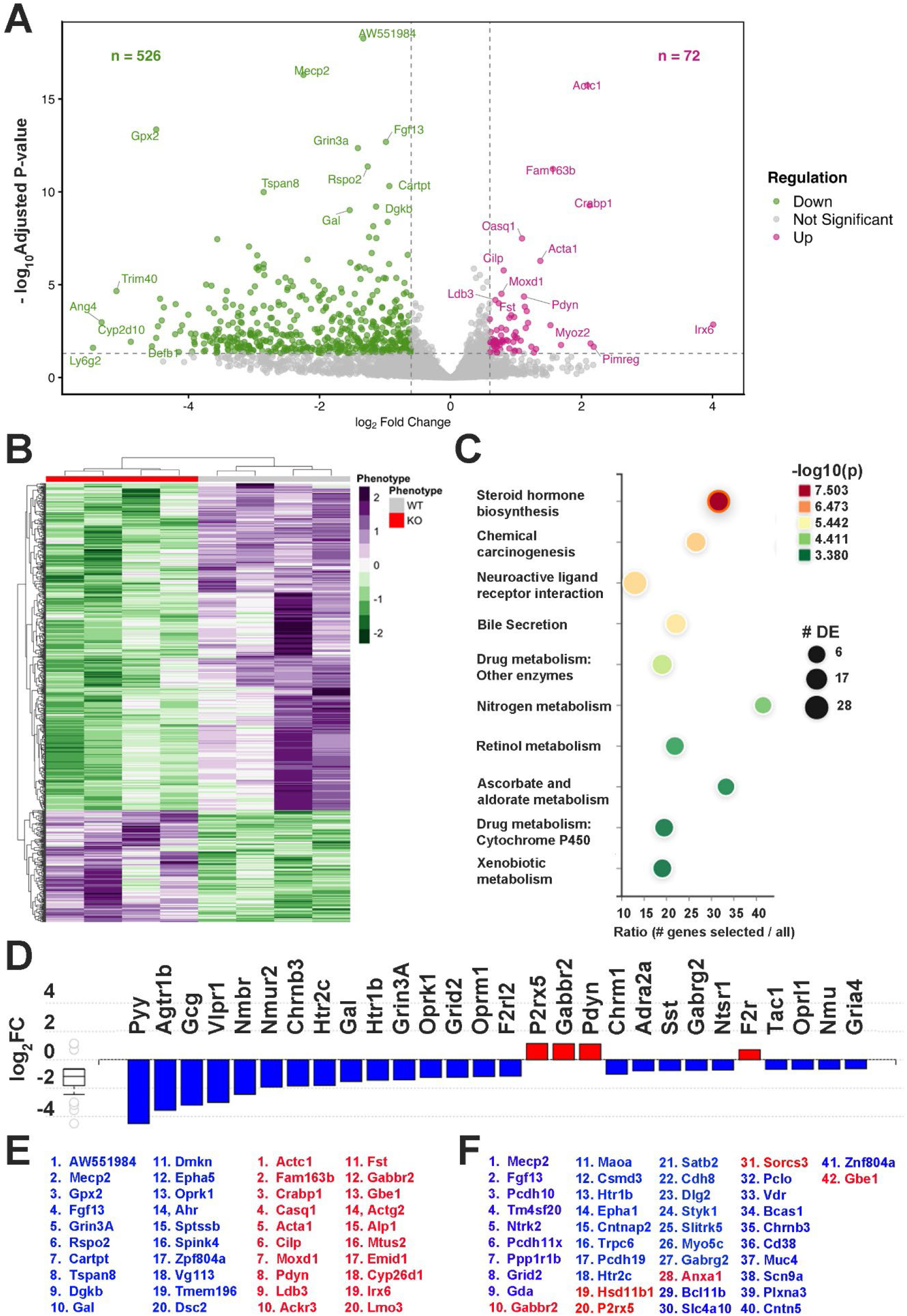
Transcriptomic changes in external muscle layer of *Mecp2* KO mice include neuronal-specific categories. ***A*,** 598 genes were statistically significant differentially expressed (DE; 526 downregulated, 72 upregulated in KO vs. WT; q-value <0.05; [log2FC] ≥ 0.6). ***B***, Heatmap showing z-scored scaled expression patterns of DE genes between KO and WT. ***C***, Pathways identified by iPathwayGuide as significantly represented by DE genes; circle size indicates DE gene number within pathway, and color indicates the significance of the DE levels of these genes. ***D***, Changes in expression (log_2_FC) of 24 downregulated (blue) and 4 upregulated (red) genes in the neuroactive ligand receptor interaction pathway. ***E***, Lists of overall top twenty downregulated (blue) and upregulated (red) genes in external muscle layer of proximal colon of KO vs. WT mice. ***F***, List of DE genes identified as dysregulated in autism spectrum disorder on SFARI website (downregulated blue, upregulated red).

**Figure 10.**
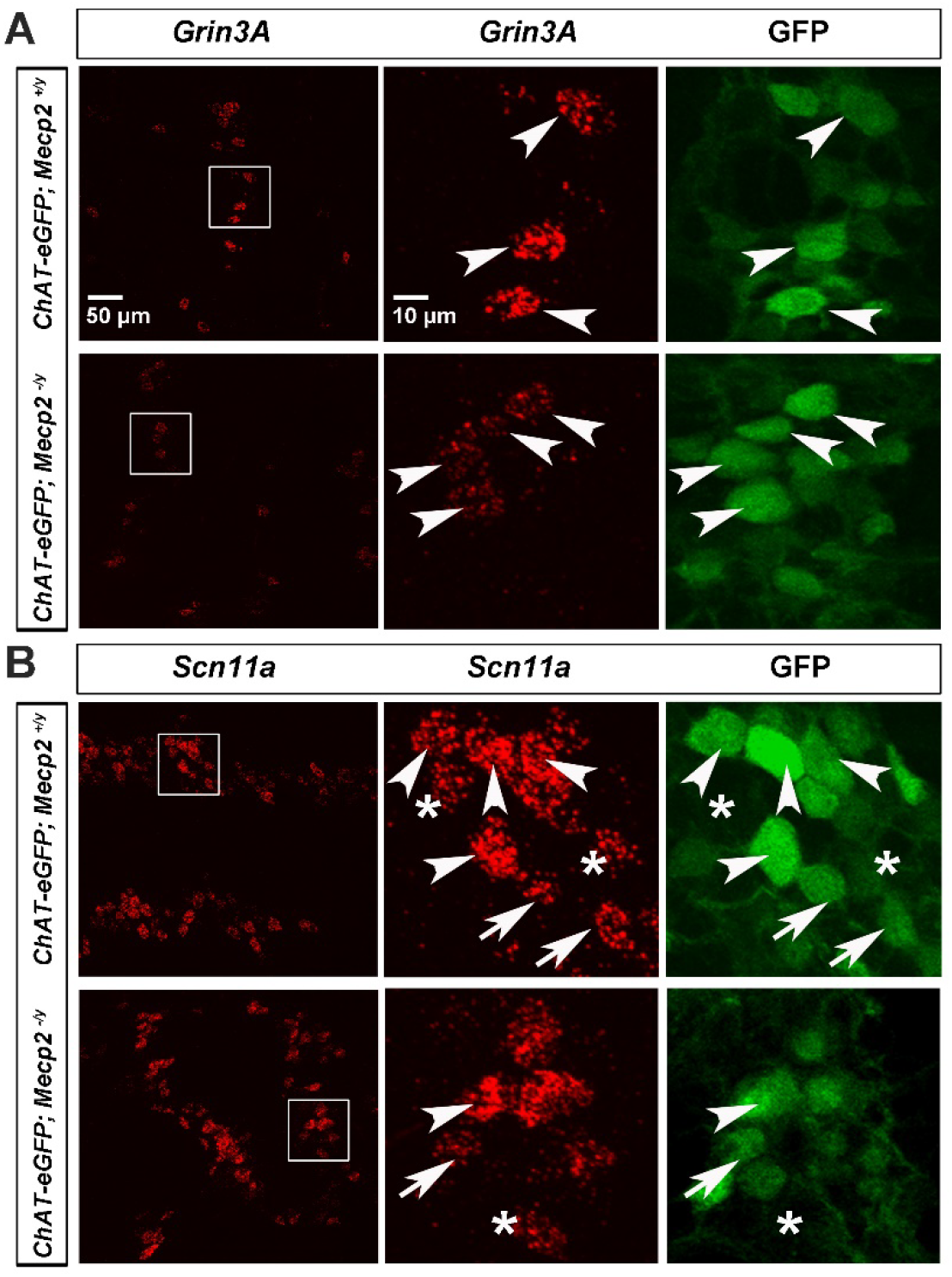
Downregulation of representative transcripts in myenteric neurons of *Mecp2* KO mice. ***A*,** Representative images of neurons of whole-mount preparations of the external muscle layer of the middle colon stained with fluorescently labeled riboprobes against *Grin3A* (red, left and middle columns), a transcript downregulated in this tissue in *Mecp2* KO mice, as well as antibodies against GFP to label cholinergic neurons (green, right column) in *Mecp2* WT (top row) and KO (*Mecp2^-/y^*; bottom row) mice crossed to *ChAT-eGFP* mice. The intensity of *Grin3A* signal within the nucleus was significantly reduced in myenteric neurons of *Mecp2* KO mice: (50.6 ± 12.4 vs. 26 ± 8.1 punctate dots, WT vs. KO, *p*=0.045, *c=4/n, n*=3). Arrowheads point to four large cholinergic neurons that express *Grin3A*. ***B*,** By contrast, the intensity of signal within the nucleus of *Scn11a*, a transcript not differentially expressed in this tissue, was similar between myenteric neurons of *Mecp2* WT and KO mice: (51.2 ± 2.3 vs. 53 ± 2.9 punctate dots, WT vs. KO, *p*=0.44, *c=4/n; n*=3). Arrowheads point to large cholinergic neurons that express *Scn11a*, arrows point to smaller cholinergic neurons that express *Scn11a*, and asterisks are just to the left of non-cholinergic neurons that express *Scn11a*. The left image, as well as the middle and right enlarged images, in the bottom row of ***A***, as well as the left images and middle and right images of ***B***, are at the same magnification as the left image, as well as the middle and right enlarged images, in the top row of ***A***.

In order to verify these changes in expression, we performed RNAscope on *Mecp2* KO and WT colonic muscle. *Grin3A*, which encodes the GluN3A subunit of the unconventional NMDA receptor that binds glycine and D-serine to convey an excitatory signal^27^, and whose expression in myenteric neurons regulates motility^28^, was the sixth most downregulated gene in KO mice based on strength of adjusted p-values (***Figure 9A,D,E).*** Expression of *Grin3A* in *ChAT-eGFP* mice was restricted to GFP-immunoreactive neurons with large somal areas, and was less expressed within these neurons in KO vs. WT mice (***Figure 10A***). The expression of *Grin3A* in large cholinergic myenteric neurons is consistent with its enrichment in IPAN identified by single-cell RNA-Seq^29^. By contrast, expression of *Scn11a*, which encodes the TTX-insensitive voltage-gated sodium channel Nav_1.9_, and which was not differentially expressed by RNA-Seq, was also not differentially expressed by RNAscope (***Figure 10B***).

The *Chrm1* gene encoding the muscarinic 1 AChR (M1R) was also downregulated in colonic muscle of *Mecp2* KO mice. We detected M1R expression in nearly all cholinergic and nitrergic myenteric neurons in WT mice by RNAscope (***Figure 11A***), with no signal detectable outside of them (i.e., in EGC, SMC, ICC). M1R was significantly less expressed within myenteric neurons in *Mecp2 KO* mice (***Figure 11B***). We first tested whether VU0357017 (10 μM), a positive allosteric modulator (PAM) of M1R that binds these receptors at a distinct site from that of ACh and works by enhancing the M1R response to ACh, could rescue the aberrant colonic motility pattern of *Mecp2* KO mice (i.e., eliminate doublet contractions). Immediately after perfusion of this drug, CMMCs were markedly increased in frequency (10.6±2.1 vs. 16.6±4.3 contractions per 30 minutes, pre-vs. post agonist, *p*=0.022), obscuring the doublet appearance. After 15-30 minutes, the doublet pattern returned (***Figure 11C***). Interestingly, when we added VU0357017 to WT mice, we failed to observe such an increase in the frequency of CMMC (9.4±1.7 vs. 10.4±1.7 contractions per 30 minutes, pre-vs. post agonist, *p*=0.372).

**Figure 11.**
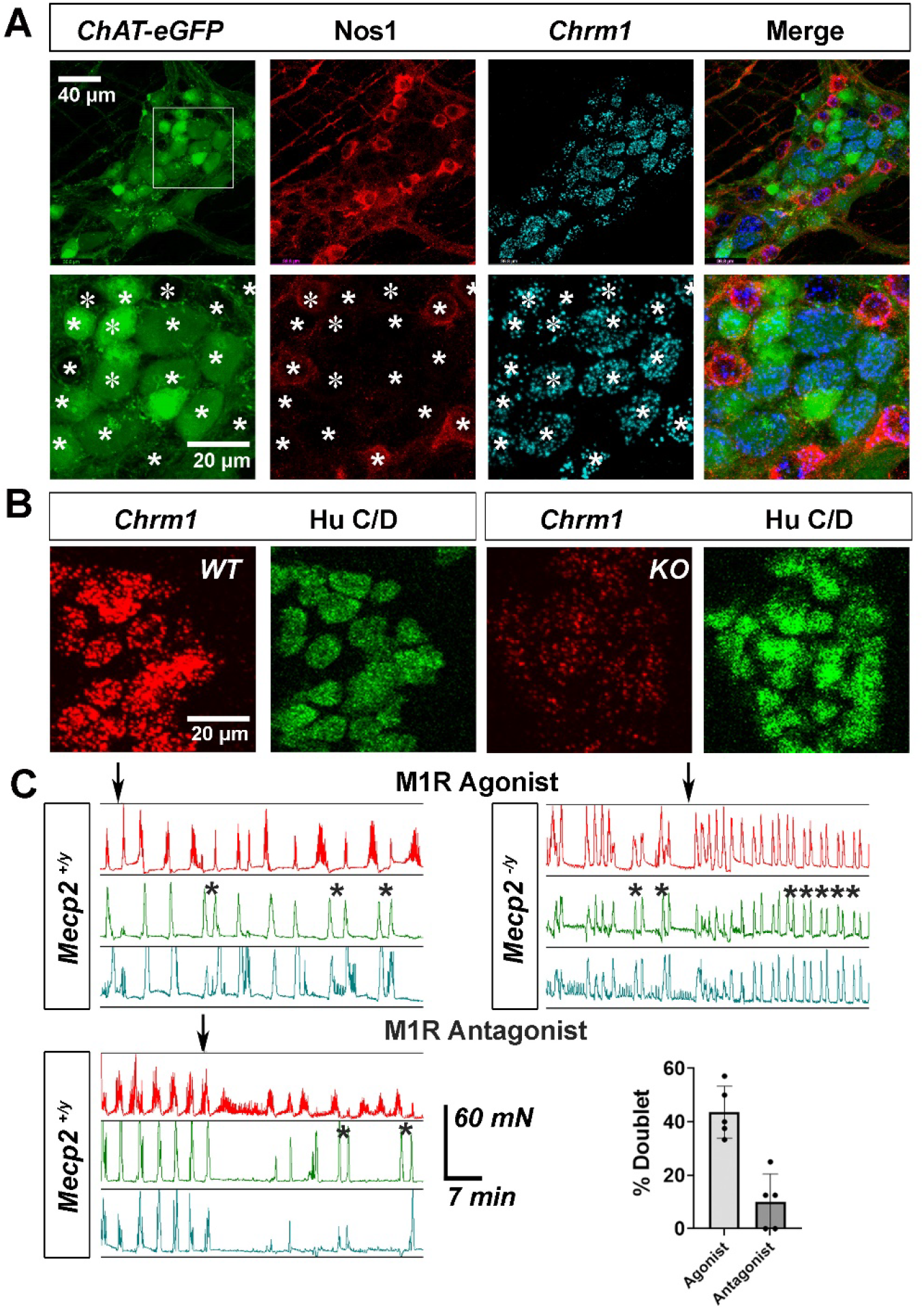
Downregulation of *Chrm1* in myenteric neurons of *Mecp2* KO mice, and effects of pharmacological manipulation of M1R on CMMC. ***A***, Cholinergic (green) and nitrergic neurons (red) each express *Chrm1* (light blue). *B*, Hu C/D+ neurons (green) of colonic myenteric plexus express *Chrm1* (red), a transcript downregulated in this tissue in KO vs. WT mice, (45.3 ± 4.2 vs. 21.9 ± 1.2 punctate dots, WT vs. KO, *p*=0.0007, *c=5/n, n*=3. *C*, Upper, CMMC after perfusion of M1R agonist VU 0357017 (10 μM), initiated at arrow. Note doublet contractions induced by treatment of WT with this drug (asterisks). Note also that doublet contractions observed in KO mice before drug re-appear after drug. Lower, CMMC in WT after perfusion of the M1R antagonist VU 0255035 (1 μM), initiated at arrow. Note occasional presence of doublets (asterisks). Right side, % of contractions that appeared as doublets within first half hour of adding M1R agonist or antagonist to WT (*n*=5). In addition to these effects of M1R PAM on doublets, the percentage increase of overall contractile frequency induced by M1R PAM was only enhanced in *Mecp2* KO mice (111±11.4 % vs. 177.5±31 %, WT vs. KO, \**p*=0.022, *n*=5).

Unexpectedly, we also observed that M1R PAM treatment of colons from *Mecp2* WT mice also exhibited doublet contractions, though not in as high a frequency as that observed in *Mecp2* KO mice before or long after M1R PAM (***Figure 11C***). Finally, we treated *Mecp2* WT mice with an M1R antagonist (VU0255035; 1 μM) whose affinity for M1R over M3R is much higher than other M1R antagonists like telenzepine^30^. Doublet contractions were observed, but in fewer mice and at a lower frequency than in response to M1R agonist (***Figure 11C***). Together, these results indicate that M1R dysregulation in *Mecp2 KO* mice may contribute to, if not underlie, aberrant GI motility.

## Discussion

Patients with NDD such as RTT exhibit prominent GI dysfunction, but whether specific motility patterns underlie this dysfunction has not been characterized. Utilizing a mouse model of RTT that features a deletion of the *Mecp2* gene, we observed a delay in intestinal transit, production of smaller, darker and more dehydrated fecal pellets, and smaller, fewer and aberrantly timed CMMC, a major neurogenic colonic motility pattern in constitutive *Mecp2* KO mice that lack Mecp2 expression in all cells. The reduction in frequency, together with the reduction in middle and distal contractile amplitudes of this *ex vivo* motor pattern in *Mecp2* KO mice, likely accounts for the delay WGTT *in vivo*. Although Mecp2 is robustly expressed by multiple cell types within the gut, deletion of this gene selectively from neurons recapitulated these phenotypes. Finally, we identified neuronal M1R signaling as a potential pathway to modulate GI dysmotility in RTT.

The presence of smaller, darker, less hydrated fecal pellets in *Mecp2* KO mice could result from impaired secretion or enhanced absorption. Alternatively, this phenotype could arise indirectly as a result of impaired motility: reduced contractile-mediated pellet propagation could prolong water absorption without any direct impairment of secretion or absorption. The absence of any alterations to electrogenic absorption or secretion in the colonic epithelia of *Mecp2* KO mice^12^ supports the idea that this phenotype derives from disruptions to motility. The initial presence of fecal pellets with high water content, followed by pellets with low water content, is consistent with surveys of patients with RTT, which found that while ∼70% of patients reported one of several signs of constipation, ∼35% of patients reported signs of diarrhea^31^. Similarly, in other GI conditions, such as subtypes of irritable bowel syndrome (IBS), patients report alternating bouts of constipation and diarrhea^32^. Alternatively, generation of these initial watery pellets may reflect stress triggered by handling. For example, some rodent models of ASD exhibit enhanced anxiety-like responses to distinct stressors^33^, and application of stressors increases both fecal output^34^ as well as fecal pellet water content^35^ in WT rodents.

Since altered CMMC (i.e., less frequent, doublet-containing CMMC with reduced middle and distal amplitudes) were observed in isolated colonic preparations from neuron-specific *Mecp2* conditional KO mice, and since absorption and secretion are unaffected in constitutive KO mice as described above, loss of Mecp2 expression in myenteric and not submucosal neurons likely accounts for this phenotype. Accordingly, we included this branch of the enteric nervous system in our bulk RNA-Seq studies. Interestingly, when the lfc was changed from 0.6 to 0.3, and the resulting DE genes compared to those marking such putative functional subclasses identified in one recent report^36^, it was noted that 8 of 10 IPAN genes were shared (*Nmu*, *Sst*, *Cdh8*, *Cbln2*, *Calcb*, *Pcdh10*, *Gpr149*, *Vgll3*), whereas just 2 excitatory motor neuron marker (eMN) genes (*Oprk1* and *Galntl6*), 3 inhibitory motor neuron (iMN) marker genes (*Dgkb*, *St18* and *Ptgir*), and 1 interneuron marker gene (*Penk*) were shared. Similarly, three other genes downregulated in the proximal colonic muscle of *Mecp2* KO mice, *Grin3A*, *Cntnap2*, and *Cdh6*, exhibit expression selectively within large cholinergic neurons (this study) or large NMU-positive neurons^37,38^ and thus presumptive IPAN. These findings indicate that colonic dysmotility may be altered in *Mecp2* KO mice as a result of transcriptomic dysregulation and subsequent dysfunction within IPAN.

Whether *Mecp2* deletion from enteric neurons mediates smaller fecal pellets is less clear because these studies occurred in living mice in which autonomic and central neurons also lacked Mecp2 expression; no genetic approach to selectively manipulate genes within enteric neurons currently exists. However, the cell-specific genetic approach alluded to above may help dissociate the effects of pellet composition from motility, if any, on GI transit. For example, if *Mecp2* deletion selectively in a neurochemical subtype (e.g., cholinergic or nitrergic) or transcriptomically-defined functional subclass (e.g., eMN or IPAN) delays GI transit and produces drier pellets but not altered CMMC, it would suggest that pellet composition is more integral to the delayed transit than altered CMMC. A similar approach to evaluating the relative effects of motility vs. fecal pellet composition on gut transit was recently demonstrated through the combined use of cell-specific genetic tools and selective pharmacological strategies to block events at the mucosal epithelium but not within the myenteric plexus^39^.

In addition to helping illuminate the cellular locus of GI dysfunction in *Mecp2* KO mice, the transcriptomic results indicated putative molecular mechanisms underlying this dysfunction. One such potential mechanism is signaling mediated by M1R, the expression of which was significantly downregulated in the proximal colon of these mutant mice. Although treatment of WT mice with the M1R antagonist VU0255035 produced doublet CMMC in WT mice, this occurred in some but not all samples and for limited duration, casting doubt that loss of M1R function alone is sufficient to induce these aberrant contractions. Interestingly, we also observed these altered neurogenic contractions in response to treatment of WT mice with the M1R PAM VU0357017. It is possible that the relatively high doses of this agonist triggered the internalization of M1R from the surface of enteric neurons by endocytosis^40^, leading to a state of reduced M1R availability that mimics that caused by downregulation of M1R expression in *Mecp2* KO mice. This interpretation is supported by the finding that M1R agonists, including PAM, exhibit a U-shaped curve of effects with increasing dose^41^.

On the other hand, treatment of *Mecp2* KO but not WT mice with VU0357017 increased the frequency of CMMC. Whether this enhanced response reflects a type of hypersensitivity to the effects of ACh on M1R triggered by reduced M1R expression in *Mecp2* KO vs. WT mice, or some compensatory downstream signaling pathway, is unclear. M1R activation on neurons typically triggers the activation of phospholipase C which in turn induces Ca^2+^ release from stores as well as closure of Kv7 channels and the associated M-current, leading to enhanced neuronal activity. In *Mecp2* KO mice, a reduction of M1R activation in all or in distinct myenteric subtypes or subclasses of neurons may in turn lead to reduced or dysregulated myenteric neuronal activity patterns and aberrant colonic motility. Interestingly, M1R agonists improve breathing deficits in these mice^42^, reduce repetitive behaviors and improve behavioral flexibility in the BTBR mouse model of ASD^43^, and enhance propulsive activity in several animal models of constipation^44^. However, it seems reasonable that diminished M1R expression mediates only a portion of the gut dysfunction observed in *Mecp2* KO mice. For example, *Grin3A* encodes the unconventional glutamate receptor subunit GluN3A, which converts the responsivity of receptors expressing this subunit from glutamate to glycine^27^, and positively regulates colonic motility^28^. Similarly, the *Gabrg2* gene encoding the γ2 subunit of GABA_A_ receptors, a subunit required for the clustering of these receptors, was downregulated in proximal colonic muscle of *Mecp2* KO mice, and a pair of recent studies demonstrate that GABA_A_ receptor antagonists reduce colonic motility in mouse models of ASD^45,46^. Together, these data support an association between altered M1R signaling and the altered motility pattern observed in RTT model mice, but do not establish causality.

Gut dysmotility is a prominent feature of NDD, but whether the type of dysmotility that occurs in *Mecp2* KO mice is shared by other mouse models of NDD is unclear. We note that at least 42 genes associated with autism were differentially expressed in the proximal colonic muscle of *Mecp2* KO vs. WT mice, suggesting the possibility of significant overlap. Several mouse models of ASD show distinct impairments of colonic activity as reported by spatiotemporal maps of video recordings of diameter changes during CMMC; for example, *Foxp1^+/-^*mice exhibit a higher percentage of retrograde CMMC^47^, *Shank3b^-/-^* mice display a fewer number of fully propagating CMMC^48^, and *Neuroligin3*^-/-^ mice show CMMC that travel more rapidly^49^. None of these three genes were differentially expressed in *Mecp2* KO vs. WT proximal colonic muscle. On the other hand, a recent study found that mutations of five distinct autism genes each triggered deficits in enteric neuron migration in *Xenopus*, which led to GI dysmotility^50^, demonstrating that comparative approaches may identify common abnormalities.

NDD may predispose patients to disorders of gut-brain interaction (DGBI), with evidence suggesting that dysbiosis in the gut lumen is responsible for at least some features of altered cognitive and motor behavior^4,5,51^. However, the cellular and molecular pathways underlying the DGBI may be distinct from those driving GI dysmotility, and identifying these latter mechanisms is imperative to produce treatment strategies aimed at ameliorating this significant symptom. In addition to providing such potential therapeutic targets, examination of the ENS in animal models of NDD may provide an alternative and more tractable methodology to identify the effects of such gene mutations on neuronal activity in the CNS, since many of the genes triggering this disorder are indeed expressed in the ENS and since population imaging of distinct ENS subtypes in intact tissue during contractile activity is possible^23^.

## Methods

### Sex as a Biological Variable

Sex was not considered as a biological variable because female *Mecp2* null mutant mice cannot be generated.

### Study Approval

Animal euthanasia, husbandry and experiments were performed in accordance with the National Institutes of Health Guide for the Care and Use of Laboratory Animals and the protocol approved by the Institutional Animal Care and Use Committee at the University of Nevada, Reno. (Protocol No. 20-08-1070-2; PHS Assurance No. D16-00311; USDA Research Registration No. 88-R-0005). Before experiments, mice were housed in groups of up to five, with standard rodent chow and water provided ad libitum. Mice were housed under a strict 12-h light/12-h dark (12 L:12D) cycle to ensure consistent circadian rhythmicity and standardized physiological responses.

### Mouse Lines

*Mecp2^tm^*^1^*^.1B^* mice, generated by crossing a mouse line harboring a *Mecp2* allele in which exons 3 and 4 are surrounded by loxP sites to a mouse line constitutively expressing Cre recombinase, exhibit no Mecp2 protein expression and are thus termed *Mecp2* null mutant or knockout (KO) mice (Jax #003890^15^). Male mice expressing one copy of this mutant allele are thus hemizygous (Mecp2^-/y^) were examined at a variety of ages. Some *Mecp2* null mutants were crossed to a *ChAT-eGFP* (Choline Acetlyltransferase-enhanced Green Fluorescent Protein) transgenic mouse line (Jax #007902^52^) in order to label cholinergic neurons in immunohistochemical experiments. Other *Mecp2* null mutants were crossed to a *CAGGS-GCaMP3* transgenic mouse line, which expresses the genetically-encoded calcium indicator GCaMP3 under the control of the CMV-chick β-actin hybrid promoter^53,54^, in order to examine the frequency and amplitude of myogenic contractions in intact colon. The *Kit-copGFP* knockin mouse line, in which GFP is inserted into the Kit promoter (Jax #015813)^55^, the *PDGFRα-H2B-eGFP* knockin mouse line, in which histone 2B-fused eGFP is inserted into the PDGFRα promoter (Jax #007669)^56^, and the *smMHC^Cre/eGFP^* transgenic mouse line, in which a bicistronic construct containing Cre and eGFP was placed under control of the myosin heavy chain 11 (Myh11) promoter (Jax #007742)^57^, were used to label ICC, nuclei of PDGFRα cells and SMC, respectively in immunohistochemical experiments. The tyrosine hydroxylase-Cre (*TH-Cre*) line (Jax #008601)^58^ was crossed to Cre-dependent *GCaMP6f* mice (Jax #028865)^59^ to label sympathetic neurons within the prevertebral ganglia. The *Kir4.1-CreER^T^*^2^ BAC transgenic mouse line, in which a tamoxifen-inducible Cre recombinase is inserted into the potassium inwardly-rectifying 4.1 channel (Kir4.1)^60^, was crossed to the Cre-dependent tdTomato Rosa26 knockin line of mice (Jax #007909) or the Mecp2 conditional mutant mice (Jax #07177) in order to label peripheral glial cells in immunohistochemical experiments or eliminate Mecp2 expression in them. The *Baf53b-Cre* mouse line (Jax #07826)^61^ and the *smMHC^Cre/eGFP^*(aka, *Myh11-Cre*) transgenic line discussed earlier were used to eliminate Mecp2 expression in neurons and smooth muscle cells, respectively. Tamoxifen-containing chow was fed to offspring of the *Kir4.1-tdTomato* or *Kir4.1-Mecp2* crosses for one week at 6 weeks of age. For all experiments, male mice were sacrificed between the ages 7-10 weeks.

### Drugs

Tamoxifen-containing chow was purchased from Envigo (TD.130855, 250mg/kg diet; Indianapolis, IN, USA). Hexamethonium bromide (100 mM stock dissolved in water) was purchased from Sigma-Aldrich (St. Louis, MO, USA). The M1R positive allosteric modulator VU0357017 and competitive orthosteric antagonist VU0255035 were purchased from MedChemExpress (Monmouth Junction, NJ, USA) and diluted in 100% DMSO as 10 mM stock solutions and thus DMSO was present at 0.1% in 10 μM VU0357017 and at 0.01% in 1 μM VU0255035. At these concentrations DMSO has no effect on CMMC measured by tension recording or by fluorescence imaging^23,63^.

### Tension recording of proximal, middle and distal intact colon

Whole colons were dissected and gently flushed. A 1.0-mm diameter fire-polished capillary tube, with artificial pellets placed and glued around the proximal, middle and distal regions of the tube, was then gently pushed through the colon. Protruding regions of the tube were fixed with metal clips to the bottom of an organ bath lined with Sylgard (Dow-Corning, Midland, MI, USA). Suture silk was attached via gluture just rostral to the proximal, middle and distal colon-placed pellets, and on the other side was attached to three force transducers (TST125C; Biopac Systems, Santa Barbara, CA, USA). Resting tension was set to 8 mN and monitored using an MP100 interface and recorded on a PC running Acqknowledge software 3.2.6 (Biopac Systems). Kreb’s Ringer’s Buffer (KRB; composition (mM): NaCl: 118; KCl: 4.7; NaHCO_3_: 25; MgCl_2_: 1.2; NaH_2_PO_4_: 1.2; glucose: 11; CaCl_2_: 2.5; gassed with 95% O_2_, 5% CO_2_) was perfused through the organ bath at a rate of 12ml/min and the chamber underlying the bath was heated to ∼35 °C. Frequencies of spontaneous colonic migrating motor complexes (CMMC), amplitudes and durations of proximal colonic contractions during a CMMC, and propagation times between peak proximal to peak middle contractions, and peak middle to peak distal contractions, were measured. At least three contractions were measured in every animal. Doublet contraction-containing CMMC were identified as those whose individual contractile amplitudes and durations were similar to those of singlet contraction-containing CMMC, but separated by fewer than 150s. For calculation of doublet contraction frequencies, the number of doublet contractions, as a percentage of the number of total contractions, within a 30-minute period was measured.

### Imaging of intact colon

A ventral midline incision was made and the entire colon was removed and placed in a 6-cm dish lined with Sylgard. The contents of the colon were gently flushed. Then, the middle of the intact colon (∼12mm in length) was split and pinned to a width of 6mm, mucosa-side down. This preparation was perfused at 6ml/min with KRB, heated before the dish by a temperature controller (TC-344C; Warner Instruments, Hamden, CT, USA) to ∼35 °C, for 45 and hour before imaging.

Two-minute Ca^2+^ imaging videos were performed on a Nikon FN-1 upright microscope using a 20X Fluor water-immersion lens. Tissue was illuminated with a Spectra X light engine (Lumencor, Beaverton, OR, USA). Image sequences were captured using a Prime 95B Camera (Photometrics, Tuscon, AZ, USA) and a Windows-based PC using Nikon NIS Elements 4.1. Image sequences were recorded at 20 frames per second and exported as 16-bit TIFF files into custom-written software (Volumetry G8d^62^). A region of interest (ROI) was placed over a region which stayed within focus during the entire movie, and this was captured as a track and used to measure the displacement of the tissue. This displacement track was then used as a template to subtract the motion that occurred during the contraction so that the onset and duration of SMC Ca^2+^ responses could be captured and compared with those of the movement. Background-subtracted traces of fluorescence intensity were generated from movies, and were reported as 16-bit intensity units (iu_16_). Large, tetrodotoxin-sensitive neurogenic contractile movements^23^ and associated fluorescence responses within SMC occurred every several minutes and could easily be distinguished from smaller, tetrodotoxin-resistant myogenic ripple contractile movements and associated responses that occurred during the tonic inhibitory period in between them.

### Intestinal length, gut transit and fecal pellet number and composition

Mice were fasted overnight and intragastrically gavaged with 100 μl of 1.5% methylcellulose containing 5% Evans Blue dye (Sigma-Aldrich) and then housed individually on white absorbent paper. WGTT was determined as the time of appearance of the first blue fecal pellet. In some cases, mice were sacrificed and the length of small and large intestine were measured. For fecal pellet analyses, mice were individually housed, and the total number and dry weight of fecal pellets collected over the 12-hour dark cycle were measured. For fecal pellet water content, individual pellets were collected immediately upon detection and pellet weight was measured immediately (wet weight) and after incubation for 12 hours at 65^0^ C (dry weight).

### Immunohistochemistry

Mice were euthanized via cervical dislocation and then decapitated. Colons were dissected, flushed, rinsed with phosphate-buffered saline (PBS) and then fixed in 4% paraformaldehyde dissolved in PBS for two hours at 4 °C. The mucosa and submucosa were removed from the muscle layer, and each of these layers was then incubated as whole-mount preparations at 4 °C with primary antibodies diluted 1:1000 in PBS containing 1% triton-X (PBST) and 10% fetal bovine serum (FBS). Specimens were confocally imaged with Fluoview software controlling an Olympus FV-1000 laser-scanning fluorescence microscope (Olympus, Tokyo, Japan). Primary antibodies were against Mecp2 (rabbit, D4F3, Cell Signaling Technology, Danvers, MA); GFP (goat, 600-101-215, Rockland Immunochemicals, Pottstown, PA); Nos1 (rabbit, PA1-033, ThermoFisher); F4/80 (rat, CI-A3-1, Novus Biologicals, Centennial, CO) and Hu C/D (human ANNA-1, kindly provided by Dr. Vanda Lennon, Mayo Clinic, Rochester, MN, USA). After extensive rinsing in PBST, fluorescently-conjugated secondary whole antibodies or Fab fragments (ThermoFisher or Jackson Immunoresearch, West Grove, PA, USA) were added to rinsed whole-mounts at 1/400 in 10% FBS/PBST overnight at 4^0^ C. Tissues were rinsed in PBST then PBS and mounted in 30% glycerol / PBS with the longitudinal muscle (for muscle whole-mounts) or muscularis mucosae (for mucosa whole-mounts) nearest to the coverslip.

### Myenteric neuronal measurements

In order to quantify numbers and somal areas of total, cholinergic and nitrergic myenteric neurons, we immunostained whole-mount preparations of the external muscle layers of colons of *Mecp2* null mutant and littermate control mice that had been crossed to *ChAT-eGFP* mice with antibodies against Hu C/D (to label all neurons), GFP (to label cholinergic neurons) and Nos1 (to label nitrergic neurons). For numbers, we imaged the middle colon at 10X and counted neurons in square millimeters (mm^2^). First, we selected 1-mm^2^ regions at random. Then, to minimize variability based on ganglia shape, number and size, in these low-power images, we selected 1-mm^2^ regions in which 25-30% of that are was covered by ganglia. For somal area, we imaged at 60X with oil immersion objectives and circled the cell body or soma of each cholinergic or nitrergic neuron based on Hu C/D staining in a separate fluorescence channel, which fills the soma. We used the measurement function under the image analysis dropdown in Adobe Photoshop to measure areas.

### Recombination efficiency

In order to measure the percentage of myenteric neurons in which Mecp2 was eliminated in *Baf53b-Mecp2^Flox/y^* mice, we stained these mice with Hu C/D and Mecp2 antibodies and counted Hu C/D-positive neurons that co-expressed or failed to express Mecp2 in 1-mm^2^ regions. In order to measure the percentage of myenteric intraganglionic glia in which Mecp2 was eliminated in *Kir4.1-Mecp2^Flox/y^*mice, we stained these mice with Hu C/D and Mecp2 antibodies and counted Hu C/D-negative cells within myenteric ganglia (presumptive enteric glial cells) that co-expressed or failed to express Mecp2 in 1-mm^2^ regions, since all Mecp2+-positive, Hu C/D-negative cells in these ganglia expressed tdTomato in *Kir4.1-tdtomato* mice (***Figure 2***) and all of these tdTomato-positive cells express the peripheral glial marker S100β^60^. In order to measure the percentage of SMC in which Mecp2 was eliminated in *Myhll-Mecp2^Flox/y^* mice, we stained these mice with Hu C/D antibodies and Hoechst 33342 dye and counted cigar-shaped, Hoechst-labeled nuclei (presumptive SMC) that co-expressed or failed to express Mecp2 in 50-μm^2^ regions.

### RNAscope

Colons were dissected, cut open, pinned flat and fixed in 4% paraformaldehyde at 4^0^ C for 24 hours. After rinsing and removal of mucosa and submucosa, small pieces of proximal andmiddle colon were dehydrated in 50%, 75% and 100% ethanol for five minutes each and allowed to dry. Tissues were then incubated in protease III solution (Advanced Cell Diagnostics (ACD), Newark, CA, USA) for 45 minutes at 40^0^ C, rinsed in 1X rinse buffer (ACD) five times for ten minutes each, then incubated with probe overnight at 40^0^ C. The next day tissue was rinsed twice for ten minutes in rinse buffer, followed by sequential 30-minute incubations at 40^0^ C and two 10-minute rinses at room temperature, with four solutions from the Multiplex Fluorescent Reagent Kit V2 (ACD): V2 AMP 1, V2 AMP2, V2 AMP3, V2 HRP-C1. Tissue was then incubated with TSA Vivid 570, diluted 1:1000 in TSA buffer (ACD) for 20 minutes at 40^0^ C. Tissue was then rinsed once in rinse buffer, once in 1% PBST, then incubated overnight at 4^0^ C with primary antibodies and blocking buffer in 1% PBST, rinsed again and then incubated overnight at 4^0^ C with fluorescently-conjugated secondary antibodies and Hoechst dye as described in immunohistochemistry section. Expression of *Grin3A*, *Scn11a* and *Chrm1* by RNAscope was examined both in the proximal and middle colon, since proximal colon was used for RNA sequencing and middle / distal colon exhibited functional impairments. The same differences in *Grin3A* and *Chrm1* expression, but not *Scn11a* expression, reported in the middle colon in ***Figures 10*** and ***11*** were also observed in proximal colon, but not shown.

### Transcriptomics

#### Dissection and RNA Isolation

Colons were dissected, with specific attention paid to the removal of the mesenteric fat. Colons were then flushed, rinsed with phosphate-buffered saline (PBS), and the mucosa and submucosa were removed from the muscle layer, whereas the serosal layer was left intact. A 0.5 cm region of the proximal colon muscle layer, over the area marked with submucosal chevrons, was dissected and minced into small pieces. These pieces were transferred into a tube containing 1 ml of Trizol (ThermoFisher). A syringe attached to an 18-guage needle pulled these pieces up and down ten times, and then the tissue was left to rest five minutes on ice. This process was repeated with 20-, 22-, 23- and 25-guage needles. RNA was then extracted and resuspended in nuclease-free water using standard Trizol purification methods.

#### Library Preparation

High-quality total RNA (500 ng; RIN > 8.4) was used as starting material for library construction. Messenger RNA (mRNA) was isolated via poly(A) selection and subsequently converted into sequencing-ready libraries using the Watchmaker mRNA Library Prep Kit (Watchmaker Genomics) according to the manufacturer’s protocol. Briefly, mRNA was fragmented and reverse-transcribed into cDNA. During the ligation step, xGen™ stubby adapters (Integrated DNA Technologies) were ligated to the cDNA fragments. Following ligation, the libraries were indexed and amplified via PCR using xGen™ UDI 10nt indexes (IDT).

#### RNA Sequencing

Final library yields were quantified and validated for size distribution. Libraries were pooled in equimolar ratios and sequenced on the Illumina NextSeq 2000 system using a paired-end 100 cycle (PE100) configuration, targeting a sequencing depth of >40 million reads per sample.

#### RNA-seq Read Processing, Alignment, Quantification, and Differential Expression Analysis

The Nevada Bioinformatics Center (RRID:SCR_017802) processed raw FASTQ files and conducted RNA-sequencing analyses. Adapter sequences and low-quality bases were trimmed using fastp (v0.20.0). Read quality was assessed before and after trimming using FastQC (v0.11.9), and quality control metrics were aggregated into a unified report with MultiQC (v1.12).

Trimmed reads were aligned to the mouse reference genome (GRCm39) using GENCODE release M36 gene annotations. Alignment was performed with STAR (v2.7). Comparative test alignments were conducted using both STAR and HISAT2, with STAR selected for downstream analyses based on superior read assignment rates. Gene-level counts were generated using featureCounts from the Subread package (v2.0). Only uniquely mapped reads were retained for quantification, and multi-mapping reads were excluded from counting.

Feature counts were further analyzed in R (v4.5.1) and initially underwent quality control procedures to ensure sample integrity. Gene-level filtering was performed using an in-house, group-aware approach that applies expression thresholds within experimental groups rather than across all samples collectively or per individual sample. Briefly, genes were retained if they demonstrated sufficient expression within at least one experimental group. Sufficient expression was defined as meeting both of the following criteria: (1) a minimum count ≥ 10 in the majority of samples within an experimental group and (2) an average count ≥ 10 within that experimental group. This filtering strategy ensures retention of genes with consistent, biologically meaningful expression patterns while removing low-abundance features unlikely to support robust statistical inference. Genes failing to meet these criteria in both WT and KO groups were removed from downstream analysis. Following filtering, analyses were restricted to protein-coding genes, and mitochondrial-encoded genes were excluded prior to differential expression testing.

Count data were variance-stabilizing transformed (VST) and assessed for overall dataset quality using principal component analysis (PCA). Variance-stabilized counts were used for all downstream exploratory visualization and investigative plotting.

Differential gene expression analysis was conducted using DESeq2 (v1.48.2). Prior to model fitting, exploratory analyses were performed to identify potential outlier transcripts exhibiting extreme expression differences driven by individual count measurements. The DESeq2 design formula was specified as ∼ genotype (KO vs WT). No additional covariates were included in the model. Each condition consisted of four biological replicates.

Genes were considered statistically differentially expressed at a *q*-value < 0.05 and an absolute log2 fold change ≥ 0.6. Q-values represent false discovery rate (FDR)–adjusted *p*-values calculated using the Benjamini–Hochberg multiple testing correction procedure.

### Statistics

Results are shown as the mean ± standard deviation (SD). *n* represents the number of mice and *c* the number of samples (i.e., cells, contractions). The minimum total number of animals chosen for any given experiment was determined by power analysis in G*power 3.010 (alpha=.05, power=80%), with an anticipated large effect size, based on our previous studies of contractile, Ca^2+^ activity and immunohistochemical phenomena^63^). When samples were compared between different animals, unpaired *t*-tests (e.g., frequency, amplitudes, durations, of CMMC) or mixed-effect two-way ANOVAs (e.g., propagation times of CMMC) were used. When the same samples were being compared, paired *t*-tests were used (e.g., CMMC pre- and post-drug). All *t*-tests were run with two tails. When populations were analyzed (e.g., cell size analysis), in addition to means ± SD, cumulative frequency distributions were generated and graphed, and the non-parametric Kolmogorov-Smirnov (K-S) test was run to evaluate these distributions. The K-S *d*-values were added manually to the cumulative frequency distribution plots. Differences in *p*-value were recognized as significant at *p* < 0.05 (*). Statistical data was generated by Graphpad Prism and is available upon request. All authors had access to the study data and have reviewed and approved the final manuscript.

## Abbreviations

(ACh): Acetylcholine
(ASD): Autism Spectrum Disorder
(Ca^2+^): Calcium
(ChAT): Choline acetyltransferase
(CMMC): Colonic Migrating Motor Complex
(ENS): Enteric Nervous System
(GI): Gastrointestinal
(KO): Knockout
(Mecp2): Methyl CpG binding protein 2
(M1R): Muscarinic ACh Receptor 1
(NDD): Neurodevelopmental Disorder
(Nos1): Nitric Oxide Synthase
(RTT): Rett Syndrome Wild-type (WT)

## Transcript profiling

Data uploaded onto NCBI sequence read archive (SRA) with BioProject ID PRJNA1446089.

## Data Availability

Bulk sequencing data uploaded onto NCBI sequence read archive (SRA) with BioProject ID PRJNA1446089. Other data available uploaded as a supporting data file. Graph pad files available upon request.

## Author Contributions

Designing research studies (NMP, TWG); Conducting experiments (NMP,SD,RD,ENB); Acquiring data (NMP,SD,RD,ENB); Analyzing data (NMP, TWG); Writing the manuscript (NMP); Editing the manuscript (CAC,TWG); Acquiring the funds (NMP,CAC,TWG)

## Acknowledgements

We are grateful to Dr. Andrew Gorzalski, Director of the Nevada Genomics Center, for library preparation and sequencing, Dr. Hans Vasquez-Gross, Bioinformatician and Dr. Julie Petereit, Director of the Nevada Bioinformatics Center, for bioinformatics.

## Figures, Legends

**Supplemental Figure 1.**
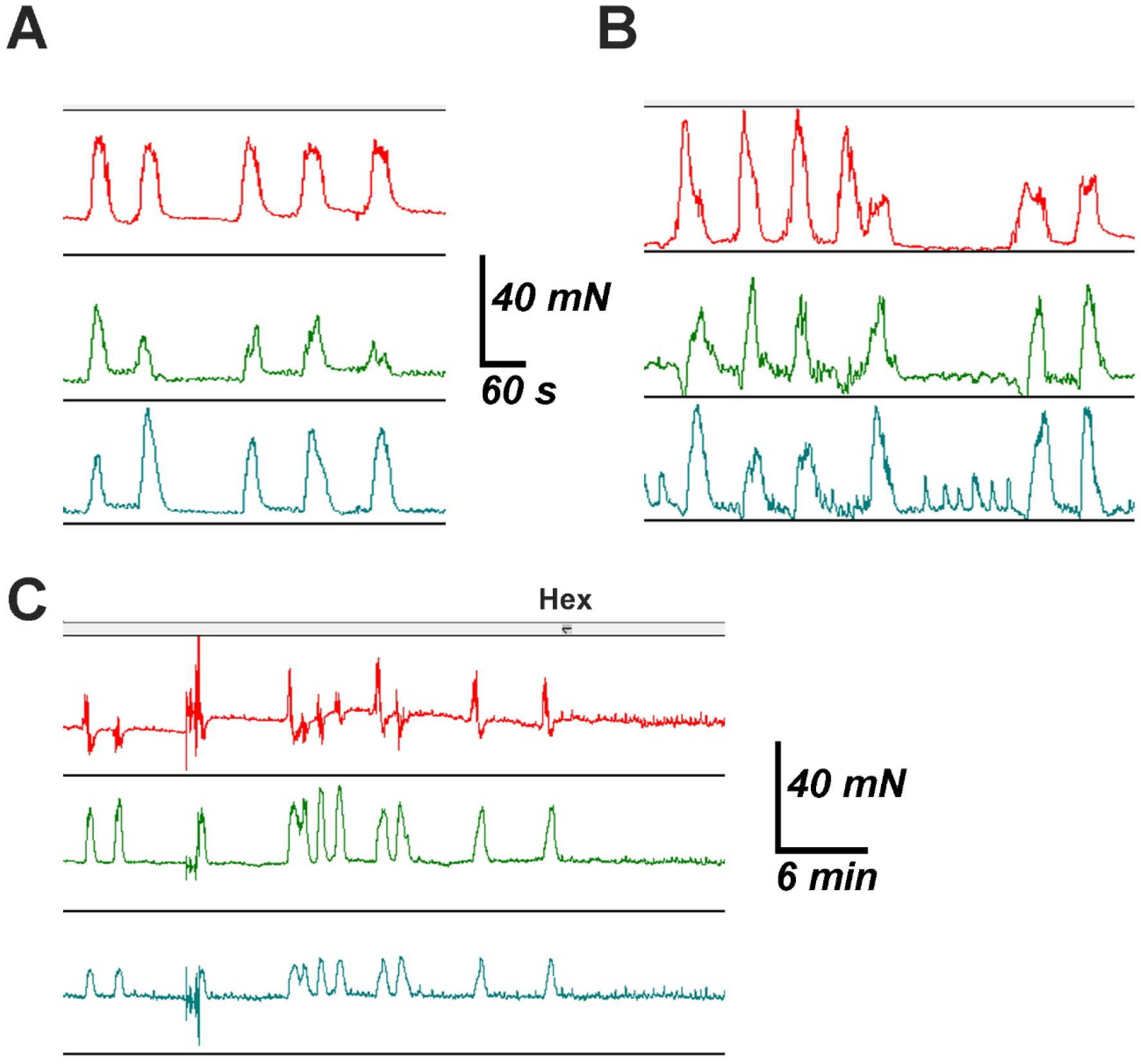
CMMC from *Mecp2* KO mice exhibit multiplets and are sensitive to hexamethonium. Examples of triplets (***A***) and quadruplets (***B***) in which each multiplet is defined by an inter-contraction interval of roughly 100 seconds. ***C***, Abolition of neurogenic contractions in proximal, middle and distal colon in response to perfusion of 100 μM hexamethonium (Hex). Representative example of *n*=7.

**Supplemental Figure 2.**
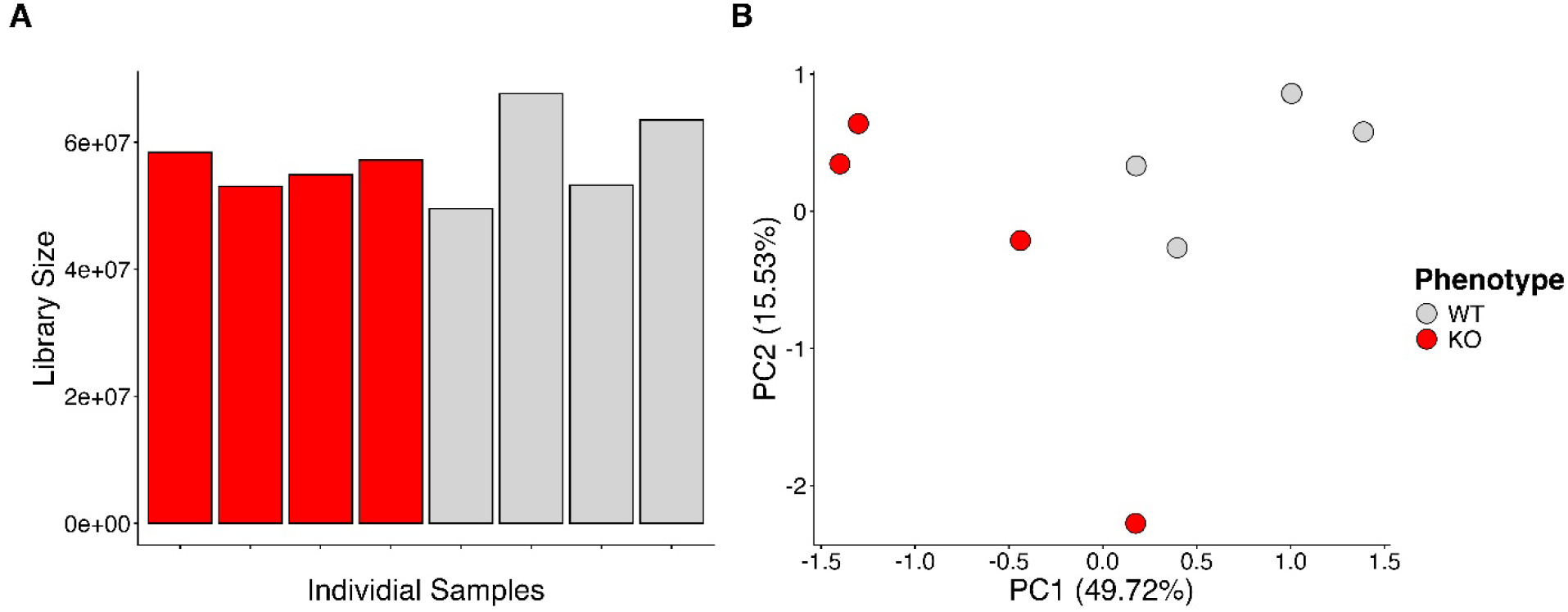
RNA Sequencing. ***A***, RNA sequencing uniquely mapped reads between 49.5 and 68.1 million reads per sample. Library sizes and overall sequencing depth were consistent across samples, indicating comparable data quality between experimental groups. Of the 78k annotated transcripts, 22,856 met the predefined expression thresholds. Subsequent restriction to protein-coding genes and exclusion of mitochondrial-encoded genes resulted in a final dataset of 15,293 genes for downstream analysis. ***B***, Principal component analysis (PCA) of variance-stabilized counts demonstrated clear separation between KO and WT samples, with genotype accounting for approximately 50% of the total variance across the dataset. A total of 29 transcripts were deemed outliers due to exhibiting extreme and biologically unexpected expression patterns and were removed prior to differential expression analysis, resulting in a final testing set of 15,264 genes.

**Supplemental Table 1.** Differential gene expression list. Ranked according to significance.

