## Supplemental Table 1 for "Neurogenic colonic motility patterns and gut transit disrupted in a mouse model of Rett Syndrome"

| Geneid | Chr | gene_name | log2FoldChange | padj |
|---|---|---|---|---|
| ENSMUSG0000003811 | chr9 | AW551984 | -1.332205107 | 5.44E-19 |
| ENSMUSG0000003139 | chrX | Mecp2 | -2.245940872 | 5.01E-17 |
| ENSMUSG0000006861 | chr2 | Actc1 | 2.086687529 | 1.83E-16 |
| ENSMUSG0000004280 | chr12 | Gpx2 | -4.492961823 | 4.35E-14 |
| ENSMUSG0000003113 | chrX | Fgf13 | -0.985909771 | 2.04E-13 |
| ENSMUSG0000003957 | chr4 | Grin3a | -1.413644444 | 4.38E-13 |
| ENSMUSG0000005192 | chr15 | Rspo2 | -1.264021532 | 4.32E-12 |
| ENSMUSG0000000921 | chr2 | Fam163b | 1.563379906 | 5.86E-12 |
| ENSMUSG0000002164 | chr13 | Cartpt | -0.933773152 | 4.86E-11 |
| ENSMUSG0000003412 | chr10 | Tspan8 | -2.852611464 | 1.03E-10 |
| ENSMUSG0000003229 | chr9 | Crabp1 | 2.123431872 | 5.34E-10 |
| ENSMUSG0000003609 | chr12 | Dgkb | -1.13637248 | 6.23E-10 |
| ENSMUSG0000002490 | chr19 | Gal | -1.539179952 | 9.66E-10 |
| ENSMUSG0000006096 | chr7 | Dmkn | -0.959395423 | 4.17E-09 |
| ENSMUSG0000002924 | chr5 | Epha5 | -1.179115854 | 7.09E-09 |
| ENSMUSG0000002590 | chr1 | Oprk1 | -1.243440647 | 2.74E-08 |
| ENSMUSG0000001925 | chr12 | Ahr | -1.132914169 | 3.10E-08 |
| ENSMUSG0000000712 | chr1 | Casq1 | 1.092381801 | 3.22E-08 |
| ENSMUSG0000004346 | chr3 | Sptssb | -3.560582198 | 3.54E-08 |
| ENSMUSG0000002841 | chr4 | Spink4 | -3.076821282 | 8.81E-08 |
| ENSMUSG0000007086 | chr2 | Zfp804a | -1.202970585 | 1.97E-07 |
| ENSMUSG0000009124 | chr16 | Vgll3 | -0.653030633 | 2.48E-07 |
| ENSMUSG0000007150 | chr6 | Tmem139 | -2.950699113 | 2.62E-07 |
| ENSMUSG0000002433 | chr18 | Dsc2 | -2.183240137 | 4.41E-07 |
| ENSMUSG0000003197 | chr8 | Acta1 | 1.371859946 | 5.21E-07 |
| ENSMUSG0000004800 | chr12 | Tmem196 | -1.528609553 | 6.45E-07 |
| ENSMUSG0000002783 | chr3 | Serpini1 | -0.937419849 | 7.70E-07 |
| ENSMUSG0000002878 | chr4 | Tmem54 | -2.851561071 | 7.95E-07 |
| ENSMUSG0000003549 | chr9 | Cdcp1 | -2.930687429 | 1.06E-06 |
| ENSMUSG0000003300 | chr1 | Asic4 | -1.394665205 | 1.06E-06 |
| ENSMUSG0000003913 | chr3 | Gipc2 | -2.962245736 | 1.26E-06 |
| ENSMUSG0000003321 | chr2 | AA467197 | -2.85581632 | 1.26E-06 |
| ENSMUSG0000004344 | chr3 | Gpr149 | -1.425068986 | 1.26E-06 |
| ENSMUSG0000006181 | chr1 | Myl1 | -1.308873527 | 1.26E-06 |
| ENSMUSG0000005291 | chr9 | Lamb2 | 0.356675302 | 1.36E-06 |
| ENSMUSG0000002745 | chr2 | Sdcbp2 | -2.934899808 | 1.61E-06 |
| ENSMUSG0000003374 | chr1 | St18 | -1.227866286 | 1.63E-06 |
| ENSMUSG0000004225 | chr9 | Cilp | 0.811642864 | 1.70E-06 |
| ENSMUSG0000003368 | chr1 | Qsox1 | 0.448657785 | 2.98E-06 |
| ENSMUSG0000003227 | chr9 | Paqr5 | -2.843391965 | 3.04E-06 |
| ENSMUSG0000002121 | chr13 | Akr1c13 | -2.51849879 | 4.13E-06 |
| ENSMUSG0000003277 | chr19 | Chrm1 | -1.017395772 | 4.32E-06 |
| ENSMUSG0000003033 | chr6 | Vamp1 | -0.827297307 | 4.32E-06 |
| ENSMUSG0000002281 | chr16 | Fstl1 | 0.389858348 | 4.32E-06 |
| ENSMUSG0000001989 | chr10 | Slc6a15 | -0.988625527 | 4.60E-06 |
| ENSMUSG0000002588 | chr9 | Casp1 | -1.374702516 | 4.96E-06 |
| ENSMUSG0000002797 | chr3 | Ndst4 | -1.101347774 | 5.25E-06 |

| | | | | |
|---|---|---|---|---|
| ENSMUSG0000001986 | chr10 | Nmbr | -2.440072763 | 6.38E-06 |
| ENSMUSG0000002839 | chr4 | Bspry | -2.091296052 | 6.58E-06 |
| ENSMUSG0000003237 | chr9 | Plod2 | 0.56118738 | 6.91E-06 |
| ENSMUSG0000006714 | chr5 | Jchain | -2.627933767 | 7.09E-06 |
| ENSMUSG0000004910 | chr3 | Pcdh10 | -0.703267605 | 7.97E-06 |
| ENSMUSG0000003266 | chr5 | Oas3 | -3.731264351 | 9.91E-06 |
| ENSMUSG0000000403 | chr3 | Gstm3 | -3.649364004 | 1.13E-05 |
| ENSMUSG0000002697 | chr2 | Psd4 | -2.012799839 | 1.13E-05 |
| ENSMUSG0000003491 | chr10 | Tjp3 | -2.406713906 | 1.18E-05 |
| ENSMUSG0000000020 | chr11 | Slfn4 | -2.986364368 | 1.23E-05 |
| ENSMUSG0000004482 | chr5 | Tlr1 | -2.486154255 | 1.47E-05 |
| ENSMUSG0000000680 | chr2 | Sulf2 | 0.49850664 | 1.49E-05 |
| ENSMUSG0000004718 | chr1 | Neurl3 | -1.670183562 | 1.61E-05 |
| ENSMUSG0000004130 | chr6 | Cftr | -2.491381048 | 1.94E-05 |
| ENSMUSG0000003739 | chr11 | Nmur2 | -1.920027671 | 1.94E-05 |
| ENSMUSG0000002822 | chr4 | Calb1 | -1.11919108 | 2.17E-05 |
| ENSMUSG0000002145 | chr13 | Fbp2 | -3.415213365 | 2.20E-05 |
| ENSMUSG0000007339 | chr17 | Trim40 | -5.100338559 | 2.20E-05 |
| ENSMUSG0000002614 | chr1 | Tm4sf20 | -3.054655461 | 2.59E-05 |
| ENSMUSG0000002663 | chr1 | Irf6 | -1.658362825 | 2.59E-05 |
| ENSMUSG0000002078 | chr11 | Atp2a3 | -0.676080418 | 2.59E-05 |
| ENSMUSG0000002447 | chr15 | Mal2 | -1.950142147 | 3.09E-05 |
| ENSMUSG0000001942 | chr7 | Ffar3 | -1.461704624 | 3.09E-05 |
| ENSMUSG0000002000 | chr10 | Moxd1 | 0.778061834 | 3.09E-05 |
| ENSMUSG0000004085 | chrX | Reps2 | -0.711446453 | 3.13E-05 |
| ENSMUSG0000005989 | chr18 | Dsc3 | -1.97753947 | 3.59E-05 |
| ENSMUSG0000005525 | chr13 | Ntrk2 | -1.190489224 | 3.61E-05 |
| ENSMUSG0000006021 | chr11 | Arrb2 | -0.604834514 | 3.81E-05 |
| ENSMUSG0000004837 | chr5 | Fgfbp1 | -2.26378732 | 3.86E-05 |
| ENSMUSG0000002740 | chr2 | Pdyn | 1.122285954 | 4.36E-05 |
| ENSMUSG0000003371 | chr13 | Akr1c14 | -2.303386189 | 4.45E-05 |
| ENSMUSG0000004577 | chr11 | Slc16a5 | -2.019498555 | 4.91E-05 |
| ENSMUSG0000005495 | chr17 | Moxd1 | -1.782266086 | 5.09E-05 |
| ENSMUSG0000006889 | chr3 | Sprr2a2 | -4.433709418 | 5.71E-05 |
| ENSMUSG0000003475 | chrX | Pcdh11x | -2.205084804 | 6.38E-05 |
| ENSMUSG0000009012 | chr1 | Ugt1a7c | -1.856058906 | 6.67E-05 |
| ENSMUSG0000002179 | chr14 | Ldb3 | 0.684221761 | 6.67E-05 |
| ENSMUSG0000004114 | chr7 | Pnma8a | -0.823809213 | 6.69E-05 |
| ENSMUSG0000002312 | chr17 | Sult1c2 | -3.434271093 | 6.73E-05 |
| ENSMUSG0000006171 | chr11 | Ppp1r1b | -1.99830884 | 7.13E-05 |
| ENSMUSG0000007142 | chr6 | Grid2 | -1.227553546 | 7.50E-05 |
| ENSMUSG0000005862 | chr19 | Gda | -0.849660137 | 8.18E-05 |
| ENSMUSG0000005802 | chr13 | Adtrp | -2.625820812 | 8.66E-05 |
| ENSMUSG0000004596 | chr2 | Gpr158 | -1.229395472 | 9.06E-05 |
| ENSMUSG0000007296 | chrX | Gprasp2 | -0.537290148 | 0.000100713 |
| ENSMUSG0000004433 | chr1 | Ackr3 | 0.597953048 | 0.000100713 |
| ENSMUSG0000002176 | chr13 | Fst | 0.736100861 | 0.000100713 |
| ENSMUSG0000002954 | chr5 | Cyp2w1 | -4.196614842 | 0.000111083 |

| | | | | |
|---|---|---|---|---|
| ENSMUSG0000002790 | chr3 | Dennd2d | -1.543237775 | 0.000116092 |
| ENSMUSG0000003516 | chr7 | Kcne3 | -2.543510298 | 0.000118949 |
| ENSMUSG0000002731 | chr2 | Ppp1r14d | -3.321393844 | 0.00012021 |
| ENSMUSG0000010371 | chr1 | Tstd1 | -2.622046963 | 0.000125647 |
| ENSMUSG0000000030 | chr8 | Cdh1 | -2.853346537 | 0.000136761 |
| ENSMUSG0000003624 | chr14 | Armh4 | -0.529594326 | 0.000138311 |
| ENSMUSG0000002729 | chr2 | Ehd4 | 0.4781085 | 0.000141582 |
| ENSMUSG0000003148 | chr8 | Rab11fip1 | -1.899807829 | 0.000153912 |
| ENSMUSG0000000039 | chr2 | Gcg | -3.196329664 | 0.000153912 |
| ENSMUSG0000002130 | chr13 | Gng4 | -0.768918002 | 0.000153912 |
| ENSMUSG0000003980 | chr4 | Gabbr2 | 1.135332438 | 0.000153912 |
| ENSMUSG0000002503 | chrX | Maoa | -1.041976686 | 0.000160663 |
| ENSMUSG0000002699 | chr2 | Galnt3 | -3.434545243 | 0.000164128 |
| ENSMUSG0000003714 | chr11 | Lypd8l | -4.376657256 | 0.000166498 |
| ENSMUSG0000002158 | chr13 | Pcsk1 | -1.043967082 | 0.000176597 |
| ENSMUSG0000005416 | chr8 | Spock3 | -0.831656423 | 0.000176721 |
| ENSMUSG0000003162 | chr8 | Isx | -1.904756805 | 0.000182395 |
| ENSMUSG0000000415 | chr12 | Etv1 | -0.452309626 | 0.000182395 |
| ENSMUSG0000005396 | chr7 | Lgals4 | -2.742804885 | 0.000183606 |
| ENSMUSG0000005463 | chr5 | Ugt2b5 | -3.68701104 | 0.000189161 |
| ENSMUSG0000002231 | chr15 | Csmd3 | -1.326309673 | 0.000189161 |
| ENSMUSG0000002270 | chr16 | Gbe1 | 0.582222522 | 0.000199045 |
| ENSMUSG0000000038 | chr16 | Tmprss2 | -2.129837547 | 0.000202534 |
| ENSMUSG0000001988 | chr10 | Mgat4c | -2.987602163 | 0.000208475 |
| ENSMUSG0000003991 | chr4 | Spsb1 | 0.4457426 | 0.000217088 |
| ENSMUSG0000002601 | chr1 | Ica1l | -0.851208544 | 0.000229656 |
| ENSMUSG0000008412 | chr8 | Esrp2 | -1.942933566 | 0.000251222 |
| ENSMUSG0000004951 | chr9 | Htr1b | -1.438192869 | 0.000251222 |
| ENSMUSG0000003150 | chr8 | Col4a1 | 0.445258203 | 0.000251222 |
| ENSMUSG0000002006 | chr10 | Mypn | 1.164394092 | 0.000264684 |
| ENSMUSG0000002112 | chr12 | Arg2 | -2.445507708 | 0.000273782 |
| ENSMUSG0000004669 | chrX | Slitrk4 | -0.842279589 | 0.000279207 |
| ENSMUSG0000003409 | chr3 | Fstl5 | -0.921474805 | 0.000306529 |
| ENSMUSG0000002735 | chr2 | Fermt1 | -3.018210986 | 0.000323491 |
| ENSMUSG0000002418 | chr17 | Pgap6 | -1.418736151 | 0.000323632 |
| ENSMUSG0000011057 | chr16 | Gm5485 | -3.239559591 | 0.000328867 |
| ENSMUSG0000007547 | chr14 | Slitrk1 | -0.585747377 | 0.000328867 |
| ENSMUSG0000000330 | chr9 | Ap1m2 | -3.608970443 | 0.000337632 |
| ENSMUSG0000006979 | chr11 | Slfn9 | -1.349158424 | 0.000348042 |
| ENSMUSG0000006335 | chr15 | Slc39a4 | -1.704085028 | 0.00038289 |
| ENSMUSG0000002944 | chr5 | Hpd | -4.002633494 | 0.000413329 |
| ENSMUSG0000001859 | chr11 | Sparc | 0.42974515 | 0.000413329 |
| ENSMUSG0000002067 | chr11 | Hnf1b | -3.194980799 | 0.00043674 |
| ENSMUSG0000002087 | chr11 | Hoxb9 | -1.676781208 | 0.000439869 |
| ENSMUSG0000002810 | chr3 | Ecm1 | 0.539515469 | 0.000439869 |
| ENSMUSG0000005943 | chr6 | Actg2 | 0.92729433 | 0.000439869 |
| ENSMUSG0000001931 | chr11 | Grb7 | -2.341658685 | 0.000454761 |
| ENSMUSG0000005096 | chr1 | Creg2 | -0.637919527 | 0.00046381 |

| | | | | |
|---|---|---|---|---|
| ENSMUSG0000002799 | chr3 | Trim2 | -0.479553258 | 0.00046381 |
| ENSMUSG0000005012 | chr1 | Vwa3b | -0.878239029 | 0.000479627 |
| ENSMUSG0000002985 | chr6 | Epha1 | -0.757831649 | 0.00048898 |
| ENSMUSG0000000860 | chr3 | Rab25 | -3.247897381 | 0.000501312 |
| ENSMUSG0000005272 | chr1 | Kcnt2 | -0.760448441 | 0.000509833 |
| ENSMUSG0000002077 | chr11 | Trim47 | 0.50022235 | 0.000521464 |
| ENSMUSG0000003284 | chr14 | Abcc4 | 0.530009025 | 0.000545242 |
| ENSMUSG0000002876 | chr4 | Alpl | 0.973004968 | 0.000553711 |
| ENSMUSG0000004750 | chr17 | Baiap3 | -1.779778131 | 0.000555303 |
| ENSMUSG0000002915 | chr5 | Cwh43 | -2.728864082 | 0.00059922 |
| ENSMUSG0000002086 | chr11 | Abcc3 | -1.554815704 | 0.000612577 |
| ENSMUSG0000004052 | chr7 | Cblc | -2.749707769 | 0.000617631 |
| ENSMUSG0000003941 | chr6 | Cntnap2 | -0.757970865 | 0.000617631 |
| ENSMUSG0000002965 | chr5 | Mtus2 | 0.898157213 | 0.000617631 |
| ENSMUSG0000001008 | chr11 | Rnf112 | -0.935406905 | 0.000670676 |
| ENSMUSG0000002768 | chr3 | Mecom | -1.804694676 | 0.000704249 |
| ENSMUSG0000003416 | chr11 | Emid1 | 0.600577739 | 0.000719739 |
| ENSMUSG0000003199 | chr9 | Trpc6 | -0.93556585 | 0.000759086 |
| ENSMUSG0000005745 | chr18 | Rit2 | -0.550649167 | 0.000759086 |
| ENSMUSG0000005382 | chr10 | Ppfia2 | -0.507540768 | 0.000759086 |
| ENSMUSG0000005132 | chrX | Pcdh19 | -1.02438917 | 0.000761957 |
| ENSMUSG0000002756 | chr3 | Car2 | -2.387078707 | 0.000791914 |
| ENSMUSG0000002641 | chr1 | Pkp1 | -1.366853214 | 0.000791914 |
| ENSMUSG0000003154 | chr8 | Ank1 | 0.583815352 | 0.000804618 |
| ENSMUSG0000004363 | chr4 | Fam221b | -0.941379859 | 0.000824832 |
| ENSMUSG0000002806 | chr3 | Sema4a | -1.578221089 | 0.000840259 |
| ENSMUSG0000005574 | chr15 | Gsdmc4 | -4.408728423 | 0.000844943 |
| ENSMUSG0000002920 | chr5 | Ugdh | -0.997012496 | 0.000914394 |
| ENSMUSG0000004138 | chrX | Htr2c | -1.813657649 | 0.000914394 |
| ENSMUSG0000002599 | chr1 | Ikzf2 | -0.94336147 | 0.000914394 |
| ENSMUSG0000003874 | chr7 | Nlrp6 | -1.296582404 | 0.000930474 |
| ENSMUSG0000002549 | chr7 | Irf7 | -1.53310128 | 0.00093605 |
| ENSMUSG0000004725 | chr2 | Ptgs1 | 0.437729224 | 0.00093605 |
| ENSMUSG0000000486 | chr17 | Mapk13 | -1.186460393 | 0.000948866 |
| ENSMUSG0000001365 | chr11 | 1810065E05Rik | -3.512434118 | 0.000953831 |
| ENSMUSG0000003218 | chr9 | Scg3 | -0.663265169 | 0.001012384 |
| ENSMUSG0000004870 | chr2 | Tprn | -0.96132121 | 0.001015559 |
| ENSMUSG0000006720 | chr5 | Lrrc66 | -3.202998224 | 0.001016576 |
| ENSMUSG0000003150 | chr8 | Col4a2 | 0.491805023 | 0.001045185 |
| ENSMUSG0000000030 | chr2 | Ckmt1 | -1.17102727 | 0.001054994 |
| ENSMUSG0000003114 | chrX | Plp2 | 0.449599708 | 0.00105769 |
| ENSMUSG0000002753 | chr3 | Chmp4c | -1.497969545 | 0.00105865 |
| ENSMUSG0000002339 | chr2 | Slc17a9 | -1.680598244 | 0.00105865 |
| ENSMUSG0000006061 | chr14 | Ang4 | -5.326928271 | 0.001063582 |
| ENSMUSG0000004080 | chrX | S100g | -3.32285726 | 0.001104847 |
| ENSMUSG0000002394 | chr17 | Slc5a7 | -0.728643932 | 0.001110328 |
| ENSMUSG0000003077 | chr7 | Parva | 0.317758751 | 0.001131993 |
| ENSMUSG0000004501 | chr17 | Acer1 | -3.187914469 | 0.001136696 |

| | | | | |
|---|---|---|---|---|
| ENSMUSG0000003680 | chr2 | Noxa1 | -2.951020287 | 0.001136696 |
| ENSMUSG0000007430 | chr8 | Smim31 | -2.64172666 | 0.001136696 |
| ENSMUSG0000006341 | chr6 | Cyp26b1 | 1.183964876 | 0.001136696 |
| ENSMUSG0000004750 | chr4 | Mroh7 | -0.932561047 | 0.001137768 |
| ENSMUSG0000003685 | chr7 | Hspb6 | 0.509479652 | 0.001287634 |
| ENSMUSG0000001845 | chr11 | 6330403K07Rik | -0.717026295 | 0.001370866 |
| ENSMUSG0000002932 | chr5 | Plac8 | -0.892208307 | 0.001379707 |
| ENSMUSG0000003040 | chr7 | Dmpk | 0.386997579 | 0.001381944 |
| ENSMUSG0000005099 | chr10 | Adgb | -1.186410151 | 0.001393799 |
| ENSMUSG0000003173 | chr8 | Irx6 | 4.011266467 | 0.001407687 |
| ENSMUSG0000002927 | chr5 | Sult1d1 | -3.389813527 | 0.001493146 |
| ENSMUSG0000003237 | chr9 | Car12 | -0.798826078 | 0.001522549 |
| ENSMUSG0000003022 | chr6 | Lmo3 | 1.525914384 | 0.001527156 |
| ENSMUSG0000005573 | chr8 | Ces2a | -4.09784972 | 0.001542911 |
| ENSMUSG0000007340 | chr17 | H2-T5 | -2.805979371 | 0.001542911 |
| ENSMUSG0000002268 | chr16 | Pla2g10 | -4.466649268 | 0.001733265 |
| ENSMUSG0000000393 | chr11 | Efnb3 | -1.133603333 | 0.001784231 |
| ENSMUSG0000007460 | chr3 | Mgst2 | -2.44359789 | 0.00189535 |
| ENSMUSG0000002857 | chr4 | Hook1 | -1.207852656 | 0.001896157 |
| ENSMUSG0000000514 | chr14 | Klf5 | -1.535080912 | 0.00191237 |
| ENSMUSG0000003063 | chr7 | Sh3gl3 | 0.436435023 | 0.00191237 |
| ENSMUSG0000006845 | chr2 | Duox2 | -1.367018518 | 0.002048863 |
| ENSMUSG0000003931 | chr9 | Minar1 | -1.039179931 | 0.002070967 |
| ENSMUSG0000006700 | chr1 | Serpinb5 | -1.963658562 | 0.00209413 |
| ENSMUSG0000003213 | chr9 | Abcg4 | -0.574942638 | 0.002108894 |
| ENSMUSG0000001619 | chr1 | Hsd11b1 | 0.776695602 | 0.002108894 |
| ENSMUSG0000000595 | chr11 | P2rx5 | 1.148856797 | 0.002151373 |
| ENSMUSG0000002647 | chr1 | Lamc2 | 0.542447913 | 0.002152899 |
| ENSMUSG0000003083 | chr7 | Ush1c | -1.652108134 | 0.002227016 |
| ENSMUSG0000003585 | chr10 | Misp | -3.251422269 | 0.002322289 |
| ENSMUSG0000002155 | chr13 | Golm1 | -1.13112901 | 0.002322289 |
| ENSMUSG0000004483 | chr1 | Ankrd45 | -0.802495955 | 0.002322289 |
| ENSMUSG0000002025 | chr9 | Glyctk | -1.609931313 | 0.002348704 |
| ENSMUSG0000007471 | chr13 | Ccl28 | -2.034989754 | 0.002450473 |
| ENSMUSG0000003108 | chrX | Slc6a14 | -3.532290404 | 0.00248519 |
| ENSMUSG0000002403 | chr17 | Cbs | -2.07159454 | 0.002494408 |
| ENSMUSG0000004750 | chr5 | Cldn4 | -2.734358767 | 0.002534524 |
| ENSMUSG0000003725 | chr2 | Dzank1 | -0.630319305 | 0.002534524 |
| ENSMUSG0000002727 | chr2 | Snap25 | -0.540225724 | 0.002534524 |
| ENSMUSG0000006723 | chr19 | Cyp2c65 | -3.57750249 | 0.002538485 |
| ENSMUSG0000004763 | chr10 | Nr1h4 | -2.704578168 | 0.002538485 |
| ENSMUSG0000006887 | chr3 | Selenbp1 | -1.351045756 | 0.002570035 |
| ENSMUSG0000002619 | chr1 | Fn1 | 0.849247668 | 0.002664688 |
| ENSMUSG0000003802 | chr11 | Rapgefl1 | -1.572244461 | 0.00271455 |
| ENSMUSG0000003041 | chr7 | Pglyrp1 | -1.805403694 | 0.002724899 |
| ENSMUSG0000003336 | chrX | Prrg3 | -0.543808314 | 0.002741828 |
| ENSMUSG0000004486 | chr9 | Gm1123 | -3.045831521 | 0.002781211 |
| ENSMUSG0000002145 | chr13 | Gadd45g | 0.705915365 | 0.002781211 |

| | | | | |
|---|---|---|---|---|
| ENSMUSG000000150 | chr11 | Col1a1 | 0.414575545 | 0.003087254 |
| ENSMUSG0000004680 | chr2 | Phgr1 | -2.663483253 | 0.003149683 |
| ENSMUSG0000004308 | chr4 | Tmem82 | -4.125396882 | 0.003149683 |
| ENSMUSG0000006876 | chr3 | Gstm6 | -1.66849795 | 0.003198477 |
| ENSMUSG0000002024 | chr10 | Col6a2 | 0.424942374 | 0.003396441 |
| ENSMUSG0000003050 | chr7 | Slc17a6 | -0.869624371 | 0.003512401 |
| ENSMUSG000000015 | chr7 | Slc22a18 | -1.999370092 | 0.003570083 |
| ENSMUSG0000003285 | chr3 | Ugt8a | -2.358132669 | 0.003604904 |
| ENSMUSG000000134 | chr9 | Cnn1 | 0.616053139 | 0.003604904 |
| ENSMUSG0000006937 | chr18 | Ctxn3 | 0.995095099 | 0.003604904 |
| ENSMUSG0000002519 | chr19 | Cpn1 | -2.940817609 | 0.003630977 |
| ENSMUSG000000386 | chr7 | Gys1 | 0.538497716 | 0.00366069 |
| ENSMUSG0000003023 | chr6 | Slco1a4 | -3.192770365 | 0.00366615 |
| ENSMUSG0000002663 | chr1 | Lamb3 | -2.024819335 | 0.003808575 |
| ENSMUSG0000008222 | chrX | Nap1l2 | -0.642524527 | 0.003808575 |
| ENSMUSG0000003795 | chr9 | A4gnt | -3.544709324 | 0.003829772 |
| ENSMUSG0000003224 | chr9 | Calml4 | -0.982073173 | 0.003829772 |
| ENSMUSG0000002616 | chr1 | Sphkap | -0.698652331 | 0.003829772 |
| ENSMUSG0000003254 | chr9 | Rab6b | -0.48439477 | 0.003847989 |
| ENSMUSG0000003371 | chr19 | Adra2a | -0.7715248 | 0.003887416 |
| ENSMUSG0000004889 | chr4 | Rimkla | -0.98950196 | 0.003887416 |
| ENSMUSG0000003833 | chr1 | Satb2 | -1.5247495 | 0.003920918 |
| ENSMUSG0000003938 | chr15 | Cdh6 | -0.630006106 | 0.00393582 |
| ENSMUSG0000002123 | chr12 | Entpd5 | -1.114395062 | 0.003962005 |
| ENSMUSG0000004779 | chrX | Gjb1 | -3.050428705 | 0.003996733 |
| ENSMUSG0000002147 | chr13 | Ctsl | 0.389079503 | 0.003996733 |
| ENSMUSG0000005597 | chr8 | Cldn23 | -3.06251415 | 0.004111324 |
| ENSMUSG0000003768 | chr5 | Atp8a1 | -0.559122879 | 0.004221124 |
| ENSMUSG0000002787 | chr3 | Hao2 | -3.962602102 | 0.004303612 |
| ENSMUSG0000004539 | chr17 | Epcam | -2.91681582 | 0.004317699 |
| ENSMUSG0000003657 | chr7 | Fxyd7 | -0.782328279 | 0.004317699 |
| ENSMUSG0000005642 | chr6 | Tgoln1 | -0.56859218 | 0.004317699 |
| ENSMUSG0000005664 | chr11 | Hoxb8 | -1.10888141 | 0.004372097 |
| ENSMUSG0000003127 | chrX | Col4a5 | 0.585907912 | 0.004372097 |
| ENSMUSG0000002908 | chr5 | Prom1 | -3.164330644 | 0.004403122 |
| ENSMUSG0000002926 | chr5 | Sult1b1 | -3.893532827 | 0.004403122 |
| ENSMUSG0000003523 | chrX | Pdk3 | 0.443627487 | 0.00440554 |
| ENSMUSG0000003925 | chr5 | Lgi2 | -0.543088589 | 0.004437531 |
| ENSMUSG0000003545 | chr12 | Foxa1 | -2.655148254 | 0.004502407 |
| ENSMUSG0000003473 | chrX | Pabpc5 | -1.68382942 | 0.004582053 |
| ENSMUSG0000004415 | chr6 | Hepacam2 | -2.441783091 | 0.004589454 |
| ENSMUSG0000005355 | chr7 | Shisa7 | -1.064482814 | 0.004589454 |
| ENSMUSG0000002461 | chr18 | Cdx1 | -2.999633209 | 0.004616491 |
| ENSMUSG0000003478 | chr4 | Dio1 | -4.204064258 | 0.004655713 |
| ENSMUSG0000003999 | chr1 | Ifi203 | -0.78310778 | 0.004655713 |
| ENSMUSG0000009170 | chr17 | H2-Q2 | -3.051270994 | 0.004736136 |
| ENSMUSG0000003998 | chr10 | Zc3h12d | -2.571332591 | 0.004798832 |
| ENSMUSG0000002453 | chr18 | Ppic | 0.470953326 | 0.004877188 |

| | | | | |
|---|---|---|---|---|
| ENSMUSG0000003184 | chr8 | Cdh13 | 0.543806654 | 0.004884583 |
| ENSMUSG0000004498 | chr15 | Tst | -1.473182515 | 0.004911811 |
| ENSMUSG0000002527 | chrX | Tro | -0.574085787 | 0.004941823 |
| ENSMUSG0000003020 | chr6 | Bcl2l14 | -2.417817856 | 0.004953262 |
| ENSMUSG0000003861 | chr7 | Rassf7 | -1.515526194 | 0.004953262 |
| ENSMUSG0000002604 | chr1 | Col5a2 | 0.400642148 | 0.004953262 |
| ENSMUSG0000003800 | chr15 | Dnajc22 | -3.587989245 | 0.005004022 |
| ENSMUSG0000001992 | chr10 | Lims1 | 0.31421789 | 0.005004022 |
| ENSMUSG0000003710 | chr15 | Fer1l6 | -2.487032269 | 0.005048599 |
| ENSMUSG0000009600 | chr14 | Gpr15lg | -3.724588401 | 0.005116267 |
| ENSMUSG0000002290 | chr16 | Ildr1 | -3.057475751 | 0.005116267 |
| ENSMUSG0000004439 | chr18 | Dsg2 | -2.288815055 | 0.005116267 |
| ENSMUSG0000006801 | chr6 | Mkrn2os | -2.347083508 | 0.005116267 |
| ENSMUSG0000004147 | chr14 | Syt15 | -0.808542922 | 0.005151894 |
| ENSMUSG0000002534 | chr10 | Tmt1b | -3.244042717 | 0.005168813 |
| ENSMUSG0000004057 | chr5 | Rundc3b | -0.765508464 | 0.005168813 |
| ENSMUSG0000002289 | chr16 | Adamts1 | -0.522903524 | 0.005182529 |
| ENSMUSG0000005035 | chr3 | Sprr1a | -2.545169848 | 0.005237121 |
| ENSMUSG0000000364 | chr4 | Rps6ka1 | -1.273873436 | 0.00528324 |
| ENSMUSG0000002610 | chr1 | Tmeff2 | -0.765670931 | 0.00528324 |
| ENSMUSG0000000222 | chr3 | Mov10 | -0.521995556 | 0.00528324 |
| ENSMUSG0000003651 | chr8 | Cdh8 | -0.760359207 | 0.005284324 |
| ENSMUSG0000007368 | chr4 | Tmem88b | -1.14682652 | 0.005284324 |
| ENSMUSG0000004047 | chr11 | Ggt6 | -3.052858236 | 0.005309233 |
| ENSMUSG0000004147 | chrX | Smpx | 0.772637518 | 0.005354985 |
| ENSMUSG0000002309 | chrX | Fhl1 | 0.969356083 | 0.005405736 |
| ENSMUSG0000003025 | chr6 | Sspn | 0.427859691 | 0.005476567 |
| ENSMUSG0000004204 | chr19 | 2010003K11Rik | -3.773575664 | 0.005633117 |
| ENSMUSG0000000027 | chr11 | Trim25 | -0.588178006 | 0.005909273 |
| ENSMUSG0000002451 | chr18 | Rab27b | -0.789356305 | 0.005944551 |
| ENSMUSG0000001998 | chr10 | Myb | -2.993576911 | 0.005948386 |
| ENSMUSG0000004635 | chr14 | Gjb2 | -2.486315123 | 0.005955097 |
| ENSMUSG0000005788 | chr16 | Abat | -0.557970078 | 0.005955097 |
| ENSMUSG0000003317 | chr6 | Mgll | 0.532376666 | 0.006280886 |
| ENSMUSG0000001364 | chr11 | Lypd8 | -3.730144914 | 0.006313867 |
| ENSMUSG0000003838 | chr7 | Rras | 0.447015484 | 0.006383744 |
| ENSMUSG0000006182 | chr8 | Ces2c | -3.705040678 | 0.006396373 |
| ENSMUSG0000002257 | chr15 | Ly6h | -0.718806026 | 0.006434975 |
| ENSMUSG0000007643 | chr13 | Sox4 | 0.460673116 | 0.006434975 |
| ENSMUSG0000004814 | chr5 | Nat8l | -0.555104316 | 0.006560094 |
| ENSMUSG0000003936 | chr11 | Sectm1b | -3.368969611 | 0.006612993 |
| ENSMUSG0000004002 | chr7 | Saa3 | -3.615695597 | 0.00666007 |
| ENSMUSG0000003185 | chr8 | Map3k21 | -1.880535459 | 0.006755425 |
| ENSMUSG0000005202 | chr18 | Slc6a7 | -3.461656763 | 0.006755425 |
| ENSMUSG0000009197 | chr17 | Hspa1a | 0.98923734 | 0.006827152 |
| ENSMUSG0000005386 | chr9 | Slc51b | -2.979111347 | 0.006895326 |
| ENSMUSG0000003786 | chr1 | Aim2 | -1.44556578 | 0.006895326 |
| ENSMUSG0000002174 | chr14 | Fam3d | -3.301132338 | 0.007124221 |

| | | | | |
|---|---|---|---|---|
| ENSMUSG0000001731 | chr11 | Pyy | -4.493151259 | 0.007348993 |
| ENSMUSG0000011034 | chr11 | Smim36 | 1.018169103 | 0.007397329 |
| ENSMUSG0000005892 | chr3 | Slc10a5 | -3.361764107 | 0.007417874 |
| ENSMUSG0000005215 | chr10 | Plpp2 | -1.056287534 | 0.007425779 |
| ENSMUSG0000007030 | chr9 | Scn2b | -0.572260657 | 0.007502492 |
| ENSMUSG0000000190 | chr11 | Kcnh6 | -1.142238442 | 0.007523171 |
| ENSMUSG0000003399 | chr8 | Kcnk1 | -1.418866148 | 0.007564972 |
| ENSMUSG0000004979 | chr4 | Lrrc19 | -1.868526349 | 0.00767871 |
| ENSMUSG0000004069 | chr1 | Slco4c1 | -3.917921801 | 0.007737764 |
| ENSMUSG0000002678 | chr2 | Gad2 | -1.569994588 | 0.007737764 |
| ENSMUSG0000005257 | chr7 | Dlg2 | -0.625427584 | 0.007737764 |
| ENSMUSG0000003870 | chr11 | Hoxb5 | -0.580757966 | 0.007782025 |
| ENSMUSG0000002426 | chr18 | Celf4 | -0.599054596 | 0.007821056 |
| ENSMUSG0000003443 | chr12 | Tmem30b | -2.747819983 | 0.00811747 |
| ENSMUSG0000002986 | chr6 | Fam131b | -0.585513533 | 0.00811747 |
| ENSMUSG0000003182 | chr8 | Crispld2 | 0.635634197 | 0.008213296 |
| ENSMUSG0000003117 | chrX | Otc | -3.25931307 | 0.008391085 |
| ENSMUSG0000006869 | chr6 | Flnc | 0.601774636 | 0.008391085 |
| ENSMUSG0000003289 | chr6 | Styk1 | -1.160132774 | 0.00886442 |
| ENSMUSG0000003155 | chr8 | Adam9 | 0.294498911 | 0.008906622 |
| ENSMUSG0000002403 | chr17 | Slc37a1 | -1.290407174 | 0.008907654 |
| ENSMUSG0000002078 | chr11 | Llgl2 | -1.105685712 | 0.008933972 |
| ENSMUSG0000002164 | chr13 | Naip1 | -3.565479361 | 0.009099857 |
| ENSMUSG0000003055 | chr7 | Lrrc28 | 0.481658978 | 0.009103676 |
| ENSMUSG0000002093 | chr11 | Plcd3 | 0.591222267 | 0.009103676 |
| ENSMUSG0000001773 | chr2 | Mmp9 | -1.419008245 | 0.009115077 |
| ENSMUSG0000004276 | chr9 | Mrap2 | -1.918294191 | 0.00914553 |
| ENSMUSG0000001804 | chr15 | Cyb5r3 | 0.312400882 | 0.009227051 |
| ENSMUSG0000005355 | chr15 | Smagp | -1.190122029 | 0.009271749 |
| ENSMUSG0000004528 | chrX | Dcaf12l1 | -0.89536787 | 0.009370601 |
| ENSMUSG0000003635 | chr3 | P2ry12 | -0.626916944 | 0.009370601 |
| ENSMUSG0000004041 | chr3 | Elapor1 | -1.120070415 | 0.009444044 |
| ENSMUSG0000003172 | chr8 | Ces1f | -2.862443296 | 0.009486433 |
| ENSMUSG0000003002 | chr6 | Adamts9 | 0.852623356 | 0.00955512 |
| ENSMUSG0000001585 | chr3 | Adamtsl4 | 0.523462864 | 0.009633258 |
| ENSMUSG0000003202 | chr9 | Rexo2 | 0.685360896 | 0.009711265 |
| ENSMUSG0000003160 | chr8 | Galnt7 | -0.67519826 | 0.009814108 |
| ENSMUSG0000002139 | chr13 | Ogn | 0.543106789 | 0.009814108 |
| ENSMUSG0000002948 | chr5 | Anxa3 | 0.543995229 | 0.009814108 |
| ENSMUSG0000002121 | chr13 | Akr1c12 | -3.508222465 | 0.009829937 |
| ENSMUSG0000005681 | chr18 | St8sia3 | -0.503756983 | 0.009866661 |
| ENSMUSG0000002120 | chr12 | Ifi27l2b | -2.413884813 | 0.009880137 |
| ENSMUSG0000003120 | chrX | Heph | -1.476359132 | 0.009880747 |
| ENSMUSG0000001786 | chr2 | Sgk2 | -3.509438875 | 0.01002951 |
| ENSMUSG0000009087 | chr17 | Hspa1b | 0.787828176 | 0.01002951 |
| ENSMUSG0000007436 | chr7 | Ehd2 | 0.448176674 | 0.010198832 |
| ENSMUSG0000002755 | chr3 | Car1 | -3.449207876 | 0.010199353 |
| ENSMUSG0000005466 | chrX | Irs4 | -1.418077803 | 0.010201698 |

| | | | | |
|---|---|---|---|---|
| ENSMUSG0000003294 | chr16 | Rbm11 | -0.906572693 | 0.010201698 |
| ENSMUSG0000002620 | chr1 | Des | 0.685805377 | 0.010309653 |
| ENSMUSG000000495 | chr5 | Hspb1 | 0.801337705 | 0.01032832 |
| ENSMUSG0000002898 | chr4 | Angptl7 | 1.071392091 | 0.01032832 |
| ENSMUSG0000004229 | chr15 | Mrtfa | 0.40567187 | 0.010365808 |
| ENSMUSG0000000436 | chr16 | Sst | -0.751175755 | 0.010373215 |
| ENSMUSG0000002661 | chr1 | Slc30a10 | -2.816528802 | 0.010391675 |
| ENSMUSG0000002977 | chr6 | Ahcyl2 | -1.008880232 | 0.010391675 |
| ENSMUSG0000003760 | chr4 | Kdf1 | -1.724109478 | 0.010399761 |
| ENSMUSG0000002751 | chr2 | Zbp1 | -1.80869394 | 0.01042094 |
| ENSMUSG0000004669 | chr4 | Tent5b | 0.878550948 | 0.010618251 |
| ENSMUSG0000004573 | chr7 | Sprn | -0.70022569 | 0.010633225 |
| ENSMUSG0000000076 | chr10 | Oprm1 | -1.184325276 | 0.010634296 |
| ENSMUSG0000003792 | chr8 | Ddx60 | -1.389986214 | 0.010772131 |
| ENSMUSG0000001816 | chr10 | Erbb3 | -0.772703452 | 0.010772131 |
| ENSMUSG0000002869 | chr4 | Tspan1 | -3.29566723 | 0.010903311 |
| ENSMUSG0000004443 | chr8 | Camsap3 | -0.822574636 | 0.010903311 |
| ENSMUSG0000003321 | chr14 | Slitrk5 | -0.648962512 | 0.010903311 |
| ENSMUSG0000005029 | chr1 | Abca12 | -3.673058291 | 0.010950764 |
| ENSMUSG0000002382 | chr17 | Slc22a1 | -2.822110014 | 0.011079109 |
| ENSMUSG0000008417 | chr7 | Sycn | -2.90720076 | 0.011079109 |
| ENSMUSG0000006176 | chr6 | Tac1 | -0.674640101 | 0.011079109 |
| ENSMUSG0000005022 | chr14 | Il17d | 0.659088935 | 0.011179506 |
| ENSMUSG0000003039 | chr7 | Ckm | 0.621931913 | 0.011354193 |
| ENSMUSG0000003359 | chr9 | Myo5c | -2.603429438 | 0.011359702 |
| ENSMUSG0000002251 | chr16 | Srl | 0.654755751 | 0.011359702 |
| ENSMUSG0000002168 | chr13 | Crhbp | -3.372555946 | 0.011439056 |
| ENSMUSG0000001749 | chr14 | Rarb | 0.538653295 | 0.011494377 |
| ENSMUSG0000002589 | chr9 | Gria4 | -0.632025054 | 0.011605413 |
| ENSMUSG0000002852 | chr4 | Sgip1 | -0.585631462 | 0.011619109 |
| ENSMUSG0000008996 | chr1 | Ugt1a1 | -3.333543779 | 0.011708423 |
| ENSMUSG0000009480 | chr15 | Cyp2d10 | -4.880932128 | 0.011857167 |
| ENSMUSG0000007476 | chr2 | Ism1 | 0.70121803 | 0.011868496 |
| ENSMUSG0000002639 | chr1 | Nr5a2 | -2.259912012 | 0.011870847 |
| ENSMUSG0000006269 | chr6 | Cav3 | 0.53108085 | 0.011925409 |
| ENSMUSG0000002043 | chr11 | Gabrg2 | -0.742653271 | 0.012006015 |
| ENSMUSG0000003555 | chr11 | Krt17 | 0.959730212 | 0.012073341 |
| ENSMUSG0000000080 | chr11 | Car4 | -3.146001335 | 0.012136207 |
| ENSMUSG0000001031 | chr1 | Optc | 0.651781036 | 0.012160108 |
| ENSMUSG0000002130 | chr13 | Hecw1 | -0.93421432 | 0.012205215 |
| ENSMUSG0000002846 | chr4 | Tpm2 | 0.565362657 | 0.012265453 |
| ENSMUSG0000003543 | chr8 | Haus8 | 0.463625657 | 0.012422876 |
| ENSMUSG0000000622 | chr4 | Hspb7 | 0.800145793 | 0.012422876 |
| ENSMUSG0000002412 | chr17 | Prss30 | -2.756256025 | 0.012456543 |
| ENSMUSG0000003248 | chr9 | Cspg5 | -1.026309755 | 0.012493483 |
| ENSMUSG0000003969 | chr10 | Ncoa7 | -0.480777318 | 0.01254949 |
| ENSMUSG0000002992 | chr6 | Smarcad1 | 0.334160399 | 0.01254949 |
| ENSMUSG0000006418 | chr10 | Rab3ip | 0.761750339 | 0.01254949 |

| | | | | |
|---|---|---|---|---|
| ENSMUSG0000005582 | chr15 | Gsdmc3 | -3.911029436 | 0.012642212 |
| ENSMUSG0000004436 | chr3 | Cxxc4 | -0.718703694 | 0.01267839 |
| ENSMUSG0000002464 | chr18 | Cbln2 | -0.481838747 | 0.01267839 |
| ENSMUSG0000001952 | chr3 | Gyg1 | 0.398442082 | 0.01267839 |
| ENSMUSG0000002465 | chr19 | Anxa1 | 0.60836852 | 0.01267839 |
| ENSMUSG0000002138 | chr13 | Aspn | 0.606209505 | 0.012752259 |
| ENSMUSG0000002683 | chr2 | Col5a1 | 0.48468097 | 0.012762012 |
| ENSMUSG0000002832 | chr4 | Stra6l | -2.028493052 | 0.012785127 |
| ENSMUSG0000009691 | chr8 | Galntl6 | -1.379934725 | 0.012788013 |
| ENSMUSG0000003080 | chr7 | Bckdk | 0.277392234 | 0.012912514 |
| ENSMUSG0000004825 | chr12 | Bcl11b | -2.313535259 | 0.013039556 |
| ENSMUSG0000001841 | chr1 | Myo1b | 0.478446196 | 0.013039556 |
| ENSMUSG0000002855 | chr4 | Atg4c | -0.396856229 | 0.013071464 |
| ENSMUSG0000003994 | chr2 | Plcb4 | 0.547228489 | 0.013071464 |
| ENSMUSG0000000703 | chr17 | Slc44a4 | -2.95673724 | 0.013088555 |
| ENSMUSG0000005466 | chr7 | Ano9 | -1.56500008 | 0.013088555 |
| ENSMUSG0000001522 | chr4 | Cyp2j9 | -0.406457146 | 0.013088555 |
| ENSMUSG0000005341 | chr16 | Hunk | -0.462529668 | 0.013183261 |
| ENSMUSG0000004462 | chr16 | Liph | -1.746917838 | 0.01320514 |
| ENSMUSG0000003457 | chr11 | Inpp5j | -0.884355159 | 0.01320514 |
| ENSMUSG0000007032 | chr11 | Rnf213 | -0.697831927 | 0.013313254 |
| ENSMUSG0000004773 | chr6 | Samd9l | -0.525737941 | 0.013512656 |
| ENSMUSG0000009820 | chr3 | Arl14 | -3.251023956 | 0.0135688 |
| ENSMUSG0000007056 | chr7 | Spaca4 | -3.634740297 | 0.013625414 |
| ENSMUSG0000002835 | chr4 | Kif12 | -2.582913877 | 0.013625414 |
| ENSMUSG0000003896 | chr2 | Slco4a1 | -1.668292809 | 0.013625414 |
| ENSMUSG0000003335 | chr16 | Rtp4 | -1.391427497 | 0.013958664 |
| ENSMUSG0000002758 | chr2 | Oprl1 | -0.664874052 | 0.01416589 |
| ENSMUSG0000007427 | chr7 | Ceacam1 | -3.197204649 | 0.014261516 |
| ENSMUSG0000003768 | chr12 | Aspg | -1.208298369 | 0.014302638 |
| ENSMUSG0000000284 | chr16 | Pla1a | 0.513637115 | 0.014362549 |
| ENSMUSG0000003420 | chr14 | Loxl2 | 0.691240776 | 0.014409092 |
| ENSMUSG0000002630 | chr1 | Mlph | -1.242292124 | 0.014476525 |
| ENSMUSG0000002933 | chr5 | Bmp3 | 0.513862164 | 0.014484273 |
| ENSMUSG0000002999 | chr6 | Gfpt1 | -0.847382863 | 0.014567737 |
| ENSMUSG0000003021 | chr6 | Plbd1 | -0.87580782 | 0.014601844 |
| ENSMUSG0000002811 | chr3 | Myoz2 | 2.139136015 | 0.014714095 |
| ENSMUSG0000002500 | chr19 | Cyp2c55 | -3.763464223 | 0.01475575 |
| ENSMUSG0000002631 | chr1 | Asb1 | -0.411508634 | 0.015054547 |
| ENSMUSG0000002191 | chr14 | Chat | -0.457580112 | 0.015149802 |
| ENSMUSG0000004462 | chr2 | Rnf208 | -0.613621787 | 0.015669757 |
| ENSMUSG0000004837 | chr13 | F2r | 0.71257738 | 0.015669757 |
| ENSMUSG0000003937 | chr1 | Hlx | 0.444499624 | 0.015683739 |
| ENSMUSG0000002167 | chr13 | F2rl2 | -1.15290679 | 0.015745838 |
| ENSMUSG0000004452 | chr3 | Tram1l1 | -0.456887389 | 0.015925224 |
| ENSMUSG0000002641 | chr1 | Pigr | -3.401772361 | 0.015982553 |
| ENSMUSG0000005629 | chr15 | Gsdmc2 | -3.904821653 | 0.015982553 |
| ENSMUSG0000004026 | chr1 | Dnm3 | -0.613893205 | 0.015982553 |

| | | | | |
|---|---|---|---|---|
| ENSMUSG0000002002 | chr10 | Cep83 | 0.422601401 | 0.016018126 |
| ENSMUSG000000430( | chr4 | Rasef | -2.240123677 | 0.016216212 |
| ENSMUSG000000268З | chr2 | Acvr1c | -1.722801906 | 0.016216212 |
| ENSMUSG000000260( | chr1 | Slc9a2 | -2.611015215 | 0.016255057 |
| ENSMUSG000000323( | chr9 | Tpm1 | 0.55935212 | 0.016255057 |
| ENSMUSG0000002731 | chr2 | Gfra4 | 0.388829798 | 0.016255057 |
| ENSMUSG0000006781 | chr2 | Myl9 | 0.523490323 | 0.016304822 |
| ENSMUSG000000207( | chr11 | Tmem132e | -0.689133168 | 0.016364403 |
| ENSMUSG000000333( | chr10 | Mif | 0.655110714 | 0.016372033 |
| ENSMUSG000000224( | chr15 | Slc38a2 | 0.247123764 | 0.016449359 |
| ENSMUSG0000002687 | chr2 | Cutal | -1.89470748 | 0.016502047 |
| ENSMUSG0000003357 | chr8 | Fa2h | -2.385161862 | 0.017231656 |
| ENSMUSG000000005Ŀ | chr15 | Itga5 | 0.752983601 | 0.017284726 |
| ENSMUSG000000541( | chr7 | Fam83e | -3.129541534 | 0.017510421 |
| ENSMUSG000000331Ś | chr11 | Myh2 | 1.687461468 | 0.017597629 |
| ENSMUSG000000161Ŀ | chrX | Tenm1 | -1.594583423 | 0.017756468 |
| ENSMUSG000000243Ą | chr17 | Btnl2 | -1.7337915 | 0.01795364 |
| ENSMUSG000000269( | chr2 | Slc4a10 | -1.178530529 | 0.01795364 |
| ENSMUSG000000240Ŀ | chr17 | Lpin2 | -0.621574694 | 0.018189517 |
| ENSMUSG000000172( | chr4 | Exosc10 | 0.273578932 | 0.018450686 |
| ENSMUSG000000174Ś | chr11 | Igfbp4 | -0.365495979 | 0.018470831 |
| ENSMUSG000000123Ŀ | chr2 | Ehf | -2.901510964 | 0.01851391 |
| ENSMUSG0000002131 | chr13 | Amph | -0.455783361 | 0.018686926 |
| ENSMUSG000000318Ŝ | chr8 | Ces2e | -3.403526838 | 0.01874847 |
| ENSMUSG000000106Ŀ | chr9 | Acaa1b | -2.638858315 | 0.01874847 |
| ENSMUSG000000900( | chr5 | Nwd2 | -1.220877883 | 0.01874847 |
| ENSMUSG000000480Ą | chrX | Arxes2 | -0.591857253 | 0.018862053 |
| ENSMUSG0000003577 | chr11 | Krt20 | -3.080473709 | 0.019131559 |
| ENSMUSG0000002252 | chr16 | Plaat1 | 0.517969838 | 0.019153681 |
| ENSMUSG000000468( | chr10 | Lrrc75b | -0.58416034 | 0.019494204 |
| ENSMUSG000000232Ą | chr4 | Guca2a | -2.931234674 | 0.01973113 |
| ENSMUSG000000608Ŝ | chr6 | Kcnd2 | -0.509763865 | 0.01973113 |
| ENSMUSG0000006241 | chr3 | Hsd3b3 | -3.076766848 | 0.01975496 |
| ENSMUSG0000001361 | chr15 | Snx31 | -1.425040469 | 0.020012786 |
| ENSMUSG000000392З | chr3 | Sec24d | 0.329315199 | 0.020054996 |
| ENSMUSG000000409( | chr3 | Arhgef38 | -2.902797757 | 0.020136257 |
| ENSMUSG000000283( | chr4 | Tnc | 0.495302372 | 0.020136257 |
| ENSMUSG000000789Ą | chr13 | Naip2 | -1.675575636 | 0.02026737 |
| ENSMUSG0000004557 | chr4 | Penk | -0.580237744 | 0.02026737 |
| ENSMUSG000000241( | chr17 | Themis3 | -2.908135705 | 0.020345672 |
| ENSMUSG0000001212 | chr4 | Crybg2 | -1.660265227 | 0.020345672 |
| ENSMUSG0000006292 | chr12 | Cfl2 | 0.389227943 | 0.020345672 |
| ENSMUSG0000002631 | chr1 | Serpinb8 | 0.65858507 | 0.020345672 |
| ENSMUSG000000489( | chr17 | Bnip5 | -3.075890174 | 0.020478939 |
| ENSMUSG000000379Ś | chr1 | Igsf9 | -1.353078008 | 0.020501943 |
| ENSMUSG000000320Ś | chr9 | Tmprss4 | -3.562077829 | 0.020517327 |
| ENSMUSG0000003837 | chr13 | Gmds | -1.176756473 | 0.020629629 |
| ENSMUSG000000316З | chr8 | Pdlim3 | 0.528435612 | 0.020657615 |

| | | | | |
|---|---|---|---|---|
| ENSMUSG0000001994 | chr10 | Rhobtb1 | 0.350695177 | 0.020713923 |
| ENSMUSG0000002976 | chr6 | Calu | 0.307863591 | 0.020752647 |
| ENSMUSG0000005707 | chr8 | Ces1g | -3.560007182 | 0.020765855 |
| ENSMUSG0000003222 | chr9 | Gcnt3 | -3.214888991 | 0.020765855 |
| ENSMUSG0000002776 | chr3 | Aadac | -2.907549673 | 0.020765855 |
| ENSMUSG0000003143 | chrX | Rnf128 | -1.293512218 | 0.020765855 |
| ENSMUSG0000004474 | chr8 | Defb1 | -4.557925366 | 0.020765855 |
| ENSMUSG0000006612 | chr7 | Kndc1 | -0.592115685 | 0.020765855 |
| ENSMUSG0000006259 | chr17 | Tubb4a | -0.492091134 | 0.020765855 |
| ENSMUSG0000001514 | chr12 | Actn1 | 0.428406414 | 0.020765855 |
| ENSMUSG0000003911 | chr9 | Itga9 | 0.472053936 | 0.020765855 |
| ENSMUSG0000003189 | chr8 | Hsd11b2 | -1.548299715 | 0.020790643 |
| ENSMUSG0000002343 | chr6 | Gnb3 | -0.833962562 | 0.021001585 |
| ENSMUSG0000002930 | chr5 | Sparcl1 | 0.413795738 | 0.021077526 |
| ENSMUSG0000004074 | chr4 | Slc25a34 | -1.871378979 | 0.021167209 |
| ENSMUSG0000003256 | chr9 | Cpne4 | -0.585029347 | 0.021268543 |
| ENSMUSG0000002003 | chr10 | Cry1 | 0.709589174 | 0.021402009 |
| ENSMUSG0000003235 | chr9 | Fam83b | -1.995466484 | 0.021659172 |
| ENSMUSG0000004256 | chr14 | Nek10 | 0.677784569 | 0.021858213 |
| ENSMUSG0000006343 | chr19 | Sorcs3 | 1.234954916 | 0.02192736 |
| ENSMUSG0000003703 | chr3 | Clca3b | -3.347020489 | 0.021928593 |
| ENSMUSG0000003650 | chrX | Tmem255a | -0.953046704 | 0.021932816 |
| ENSMUSG0000002080 | chr11 | Pimreg | 2.18632786 | 0.022022338 |
| ENSMUSG0000001998 | chr10 | Ahi1 | -0.460267018 | 0.022191208 |
| ENSMUSG0000003229 | chr9 | Pkm | 0.359505507 | 0.022264301 |
| ENSMUSG0000007950 | chr17 | H2-Q1 | -3.191804587 | 0.022326966 |
| ENSMUSG0000002855 | chr4 | Ttc39a | -1.307796374 | 0.022373452 |
| ENSMUSG0000002402 | chr17 | Tff3 | -3.161487338 | 0.022398406 |
| ENSMUSG0000006702 | chr1 | Cntnap5b | -1.844205176 | 0.022398406 |
| ENSMUSG0000006306 | chr7 | Mapk3 | -0.288361169 | 0.022460031 |
| ENSMUSG0000003068 | chr7 | Stard10 | -0.63130266 | 0.022554917 |
| ENSMUSG0000006160 | chr5 | Pclo | -0.699492564 | 0.022581297 |
| ENSMUSG0000002163 | chr13 | Marveld2 | -1.2989387 | 0.022705885 |
| ENSMUSG0000000219 | chr8 | Clgn | -0.705632307 | 0.022705885 |
| ENSMUSG0000002517 | chr19 | Ubtd1 | 0.545806312 | 0.022779054 |
| ENSMUSG0000004230 | chr3 | S100a14 | -3.257411202 | 0.022805501 |
| ENSMUSG0000004659 | chr16 | Bdh1 | 0.391085934 | 0.022805501 |
| ENSMUSG0000005521 | chr4 | Foxd2 | -2.18228198 | 0.02289654 |
| ENSMUSG0000002921 | chr5 | Slc10a4 | -0.540180688 | 0.023134901 |
| ENSMUSG0000002153 | chr13 | Adcy2 | -0.838227219 | 0.023310191 |
| ENSMUSG0000000678 | chr11 | Odad4 | -0.910089144 | 0.023471156 |
| ENSMUSG0000001973 | chr7 | Tmc4 | -2.180072068 | 0.023508743 |
| ENSMUSG0000002282 | chr16 | Muc13 | -3.248343726 | 0.023763727 |
| ENSMUSG0000002254 | chr15 | Gpt | -1.129181605 | 0.023763727 |
| ENSMUSG0000002180 | chr14 | Cdhr1 | -1.795554999 | 0.023763727 |
| ENSMUSG0000004932 | chr5 | Kmt5a | 0.385163955 | 0.023763727 |
| ENSMUSG0000002672 | chr2 | Rsu1 | 0.381663576 | 0.023997444 |
| ENSMUSG0000002247 | chr15 | Vdr | -2.52467581 | 0.024044692 |

| | | | | |
|---|---|---|---|---|
| ENSMUSG0000003581 | chr5 | Ugt2b35 | -2.418520395 | 0.024044692 |
| ENSMUSG0000003484 | chr5 | Art3 | 0.46576333 | 0.024044692 |
| ENSMUSG0000009251 | chr7 | Art2a | -2.688820204 | 0.024238482 |
| ENSMUSG0000003149 | chr8 | Chrnb3 | -1.846376137 | 0.024238482 |
| ENSMUSG0000004221 | chr5 | Sgsm1 | -0.570537905 | 0.024238482 |
| ENSMUSG0000005248 | chr13 | Tmem171 | -3.246616751 | 0.024392737 |
| ENSMUSG0000005667 | chr18 | Prelid2 | -1.503046324 | 0.024742681 |
| ENSMUSG0000003059 | chr7 | Ryr1 | -0.963836566 | 0.024834514 |
| ENSMUSG0000002604 | chr1 | Col3a1 | 0.371172078 | 0.024834514 |
| ENSMUSG0000003740 | chr1 | Cnnm4 | -1.218862133 | 0.024881907 |
| ENSMUSG0000003458 | chr9 | Exph5 | -1.324551941 | 0.024985611 |
| ENSMUSG0000004772 | chr15 | Ly6g2 | -5.458348438 | 0.025032482 |
| ENSMUSG0000007098 | chr4 | Acnat1 | -3.088127805 | 0.025032482 |
| ENSMUSG0000005058 | chr2 | Lrrc4c | -0.474416976 | 0.025032482 |
| ENSMUSG0000004411 | chr16 | Bmerb1 | -0.496514502 | 0.025032482 |
| ENSMUSG0000003781 | chr18 | Ctnna1 | 0.28000774 | 0.025032482 |
| ENSMUSG0000004603 | chrX | Snx12 | 0.347417476 | 0.025032482 |
| ENSMUSG0000003690 | chr8 | Neto2 | -0.449496408 | 0.025070838 |
| ENSMUSG0000001352 | chr2 | Bcas1 | -2.1082559 | 0.025206842 |
| ENSMUSG0000002907 | chr4 | Ttll10 | -2.196692662 | 0.025206842 |
| ENSMUSG0000005071 | chr1 | Scg2 | -0.458806492 | 0.025614147 |
| ENSMUSG0000002599 | chr1 | Cps1 | -1.947331797 | 0.025856413 |
| ENSMUSG0000000977 | chr1 | Nuak2 | -1.67201293 | 0.02593263 |
| ENSMUSG0000004821 | chr6 | Gpr85 | -0.513519416 | 0.02593263 |
| ENSMUSG0000001050 | chr2 | Myt1 | -0.733146669 | 0.026253557 |
| ENSMUSG0000007643 | chr11 | Acsf2 | -1.154818656 | 0.026315186 |
| ENSMUSG0000005966 | chr12 | Taf1b | 0.370120577 | 0.026315186 |
| ENSMUSG0000003625 | chr5 | Igfbp7 | 0.456133309 | 0.026315186 |
| ENSMUSG0000002817 | chr3 | Cth | -1.357357523 | 0.026503734 |
| ENSMUSG0000002133 | chr13 | Slc17a4 | -3.571737552 | 0.0268647 |
| ENSMUSG0000002323 | chr4 | Serinc2 | -1.332233639 | 0.027223823 |
| ENSMUSG0000000038 | chr16 | Mx1 | -2.19773038 | 0.027223823 |
| ENSMUSG0000007171 | chr15 | Mpst | -1.03602579 | 0.027272439 |
| ENSMUSG0000002420 | chr17 | Acsbg3 | 0.572603781 | 0.027272439 |
| ENSMUSG0000000664 | chr5 | Tcf23 | 0.898627934 | 0.027272439 |
| ENSMUSG0000002622 | chr1 | 2810459M11Rik | -2.488154331 | 0.027491175 |
| ENSMUSG0000004671 | chr11 | Nxph3 | -0.607835321 | 0.027491175 |
| ENSMUSG0000002244 | chr15 | Cyp2d26 | -3.058301538 | 0.027603717 |
| ENSMUSG0000002126 | chr12 | Degs2 | -2.442507242 | 0.027603717 |
| ENSMUSG0000003410 | chr1 | Ano7 | -2.258882012 | 0.027653945 |
| ENSMUSG0000010024 | chr14 | Slc18a3 | -0.725678364 | 0.027724431 |
| ENSMUSG0000002473 | chr19 | Ms4a8a | -2.533152884 | 0.027819823 |
| ENSMUSG0000002421 | chr17 | Spdef | -2.668591587 | 0.027827303 |
| ENSMUSG0000002182 | chr14 | Vcl | 0.435142748 | 0.028081386 |
| ENSMUSG0000002722 | chr2 | Mapk8ip1 | -0.330480444 | 0.028166063 |
| ENSMUSG0000003281 | chr9 | Igdcc4 | -0.752889526 | 0.028177848 |
| ENSMUSG0000003977 | chr4 | Galnt12 | -1.445787012 | 0.02839403 |
| ENSMUSG0000004043 | chr11 | Pitpnc1 | 0.294779057 | 0.02839403 |

| | | | | |
|---|---|---|---|---|
| ENSMUSG0000004063 | chr8 | Dok4 | -0.941872406 | 0.028900036 |
| ENSMUSG000000224€ | chr15 | Slc38a4 | 0.505731527 | 0.028919695 |
| ENSMUSG000000366ε | chr5 | Tmem184a | -1.845783467 | 0.028998754 |
| ENSMUSG0000002223 | chr15 | Cmbl | -0.446845776 | 0.028998754 |
| ENSMUSG0000002457 | chr18 | Il17b | 1.307273461 | 0.028998754 |
| ENSMUSG000000548ε | chr13 | Dsp | -2.290485099 | 0.029247903 |
| ENSMUSG0000006721 | chr5 | Nipal1 | -2.361058518 | 0.029324948 |
| ENSMUSG0000007444 | chr3 | Sprr2a3 | -3.217611221 | 0.029487764 |
| ENSMUSG0000002843 | chr4 | Epb41l4b | -0.647754621 | 0.029487764 |
| ENSMUSG0000003461 | chr10 | Ppm1h | -0.437710378 | 0.029487764 |
| ENSMUSG000000326€ | chr1 | 1700025G04Rik | 0.252370226 | 0.029487764 |
| ENSMUSG000000217E | chr13 | Plpp1 | 0.54797894 | 0.029539776 |
| ENSMUSG0000005454 | chr1 | Ugt1a6a | -1.768611272 | 0.029563654 |
| ENSMUSG0000001582 | chr1 | Tnr | 0.523468756 | 0.029563919 |
| ENSMUSG0000003937 | chr1 | Marchf4 | -0.753237305 | 0.029949254 |
| ENSMUSG000000353E | chr7 | Uvrag | 0.307681296 | 0.029949254 |
| ENSMUSG0000005997 | chr12 | Hspa2 | 0.338810796 | 0.029949254 |
| ENSMUSG0000002033 | chr11 | Acsl6 | -0.442343679 | 0.030219996 |
| ENSMUSG0000003897 | chr3 | Rabggtb | 0.321717746 | 0.030222002 |
| ENSMUSG000000297S | chr6 | Herc6 | -0.747004693 | 0.030269459 |
| ENSMUSG000000639C | chr7 | Klk1 | -3.398621266 | 0.030358434 |
| ENSMUSG000000211C | chr12 | Hif1a | 0.452837165 | 0.030358434 |
| ENSMUSG0000004221 | chr1 | Bag2 | 0.759650901 | 0.030503989 |
| ENSMUSG000000323€ | chr9 | Plscr1 | -0.901228104 | 0.030819961 |
| ENSMUSG0000002207 | chr14 | Klhl1 | -0.862145963 | 0.030878601 |
| ENSMUSG000000587C | chr17 | Memo1 | 0.417379426 | 0.030939528 |
| ENSMUSG0000003241 | chr9 | Me1 | 0.265774664 | 0.031001754 |
| ENSMUSG0000005794 | chr11 | Unc13d | -1.151992954 | 0.031155757 |
| ENSMUSG000000355S | chr9 | Chrna5 | -0.56581534 | 0.031155757 |
| ENSMUSG000000318C | chr8 | B3gnt3 | -2.072670138 | 0.031222747 |
| ENSMUSG000000391S | chr17 | Nlrc4 | -1.603523409 | 0.031526269 |
| ENSMUSG000001020S | chr2 | Or5m9 | -1.696827515 | 0.031526269 |
| ENSMUSG0000004277 | chr6 | Hebp1 | 0.492661435 | 0.031526269 |
| ENSMUSG000000431€ | chr3 | Loricrin | -1.674334755 | 0.031577162 |
| ENSMUSG0000004564 | chr1 | Vwc2l | -2.254832738 | 0.031624983 |
| ENSMUSG000000299S | chr6 | Tgfa | -1.041181015 | 0.031811329 |
| ENSMUSG0000004897 | chr13 | Nrsn1 | -0.606787845 | 0.031811329 |
| ENSMUSG000000581E | chr5 | Sez6l | 0.530058419 | 0.031890112 |
| ENSMUSG000000273S | chr2 | Slc20a1 | -1.016717234 | 0.032087928 |
| ENSMUSG0000002111 | chr12 | Plek2 | -2.125916962 | 0.032601043 |
| ENSMUSG0000005541 | chr11 | Atp10b | -3.015792858 | 0.03306589 |
| ENSMUSG000000306E | chr7 | Coq7 | 0.328804255 | 0.03306589 |
| ENSMUSG000000192E | chr7 | Ppp1r12c | 0.39905591 | 0.03306589 |
| ENSMUSG000000735E | chr1 | Ppp1r12b | 0.302024609 | 0.033405986 |
| ENSMUSG0000002492 | chr19 | Ovol1 | -2.491689454 | 0.033452051 |
| ENSMUSG000000227E | chr16 | Adgrg7 | -3.393887241 | 0.033853381 |
| ENSMUSG000000736C | chr1 | Gal3st2c | -3.744199663 | 0.034151686 |
| ENSMUSG0000004263 | chr6 | Gucy2c | -3.191284199 | 0.034151686 |

| | | | | |
|---|---|---|---|---|
| ENSMUSG0000001992 | chr10 | Zwint | -0.272540071 | 0.034313698 |
| ENSMUSG0000001812 | chr15 | Baiap2l2 | -1.948258862 | 0.034436792 |
| ENSMUSG0000002955 | chr5 | Hnf1a | -2.363035625 | 0.034571286 |
| ENSMUSG0000002833 | chr4 | Coro2a | -0.816187751 | 0.034735393 |
| ENSMUSG0000001620 | chr1 | Syt14 | -0.651442858 | 0.034735393 |
| ENSMUSG0000002081 | chr11 | Mxra7 | 0.327138299 | 0.034735393 |
| ENSMUSG0000002015 | chr11 | Kcnmb1 | 0.426788917 | 0.034735393 |
| ENSMUSG0000004250 | chr1 | Cpa6 | 0.560150972 | 0.034871293 |
| ENSMUSG0000003987 | chr10 | Slc39a5 | -2.199623688 | 0.035015028 |
| ENSMUSG0000002908 | chr5 | Cd38 | -1.208594178 | 0.035015028 |
| ENSMUSG0000004178 | chr1 | Lad1 | -2.875641841 | 0.035018163 |
| ENSMUSG0000002631 | chr1 | Cdh7 | -1.379485288 | 0.035018163 |
| ENSMUSG0000003985 | chr4 | Trim14 | -0.725861422 | 0.035018163 |
| ENSMUSG0000001763 | chr11 | Abr | -0.381610904 | 0.035018163 |
| ENSMUSG0000002332 | chr5 | Ache | -0.532406766 | 0.035026214 |
| ENSMUSG0000002276 | chr16 | Ccdc116 | -2.007896844 | 0.035085992 |
| ENSMUSG0000000621 | chr4 | Fblim1 | 0.482261131 | 0.035229451 |
| ENSMUSG0000002579 | chr11 | Slc25a10 | -1.042868986 | 0.03527226 |
| ENSMUSG0000002293 | chr16 | Fam3b | -3.5651389 | 0.035359645 |
| ENSMUSG0000002756 | chr2 | Ntsr1 | -0.713476052 | 0.035359645 |
| ENSMUSG0000006078 | chr6 | Lrrtm1 | -0.416905918 | 0.035441647 |
| ENSMUSG0000002422 | chr17 | Fkbp5 | 0.982101004 | 0.035444043 |
| ENSMUSG0000003252 | chr9 | Vipr1 | -3.007304801 | 0.03548098 |
| ENSMUSG0000002058 | chr12 | Agr2 | -2.506109872 | 0.03548098 |
| ENSMUSG0000006680 | chr1 | Rnasel | -0.971668055 | 0.03548098 |
| ENSMUSG0000002731 | chr2 | Spint1 | -2.256584386 | 0.035626449 |
| ENSMUSG0000006231 | chr11 | Erbb2 | -0.716532601 | 0.035752598 |
| ENSMUSG0000000288 | chr8 | Adgre5 | 0.401125261 | 0.035883475 |
| ENSMUSG0000002987 | chr15 | Ccdc184 | -0.772831659 | 0.036273326 |
| ENSMUSG0000006108 | chr11 | Myl4 | 1.046499935 | 0.036967608 |
| ENSMUSG0000004261 | chr3 | Pbxip1 | 0.282845515 | 0.037186856 |
| ENSMUSG0000004751 | chr7 | Dmbt1 | -3.296241071 | 0.03740596 |
| ENSMUSG0000002121 | chr12 | Rgs6 | 0.548096475 | 0.03740596 |
| ENSMUSG0000003090 | chr7 | Anks4b | -2.783498944 | 0.037422092 |
| ENSMUSG0000002121 | chr13 | Tubal3 | -2.547982284 | 0.037422092 |
| ENSMUSG0000003010 | chr6 | Usp18 | -1.614266291 | 0.037422092 |
| ENSMUSG0000003930 | chr3 | Tnfsf10 | -0.717533084 | 0.037422092 |
| ENSMUSG0000002149 | chr13 | Pdlim7 | 0.442408239 | 0.037422092 |
| ENSMUSG0000004301 | chr7 | Ptgir | -0.431213731 | 0.037632028 |
| ENSMUSG0000003350 | chr3 | Cdc14a | -0.393656491 | 0.037751937 |
| ENSMUSG0000004032 | chr3 | Slc25a24 | -0.455300172 | 0.03783476 |
| ENSMUSG0000003199 | chr9 | St14 | -2.671349068 | 0.038013854 |
| ENSMUSG0000004072 | chr9 | Cep126 | -0.673319209 | 0.038396579 |
| ENSMUSG0000003681 | chr2 | Entpd8 | -3.045235188 | 0.038569193 |
| ENSMUSG0000002107 | chr12 | Trim9 | -0.536476366 | 0.039236875 |
| ENSMUSG0000003615 | chr15 | Prickle1 | 0.341562897 | 0.039310148 |
| ENSMUSG0000003727 | chr2 | Ovol2 | -3.76641498 | 0.03941733 |
| ENSMUSG0000003071 | chr7 | Sult1a1 | -1.105043151 | 0.039422994 |

| | | | | |
|---|---|---|---|---|
| ENSMUSG0000002119 | chr12 | Lgmn | 0.249551226 | 0.039625088 |
| ENSMUSG0000002641 | chr1 | Tnnt2 | 0.980913594 | 0.039734959 |
| ENSMUSG0000000634 | chr10 | Susd2 | 0.373786688 | 0.039756718 |
| ENSMUSG0000011867 | chr16 | Muc4 | -3.944452643 | 0.039783843 |
| ENSMUSG0000006263 | chr17 | Btnl1 | -3.048377887 | 0.039783843 |
| ENSMUSG0000003054 | chr7 | Idh2 | 0.412131043 | 0.039783843 |
| ENSMUSG0000003208 | chr9 | Tagln | 0.500920305 | 0.039783843 |
| ENSMUSG0000002090 | chr11 | Usp43 | -1.858941831 | 0.039812871 |
| ENSMUSG0000003864 | chr3 | Ctss | -0.649303744 | 0.039920387 |
| ENSMUSG0000004326 | chr1 | Ifi209 | -1.309822371 | 0.040176966 |
| ENSMUSG0000002520 | chr19 | Sema4g | -1.68444059 | 0.040227693 |
| ENSMUSG0000001977 | chr10 | Vip | -0.52288482 | 0.040344986 |
| ENSMUSG0000002538 | chr10 | Il23a | -1.721305552 | 0.040513565 |
| ENSMUSG0000003024 | chr6 | Ldhb | 0.537810203 | 0.040693451 |
| ENSMUSG0000000111 | chr10 | Col6a1 | 0.434955969 | 0.040771467 |
| ENSMUSG0000003184 | chr8 | Hsd17b2 | -3.149115824 | 0.041194437 |
| ENSMUSG0000004881 | chr1 | Lonrf2 | -0.520963225 | 0.041194437 |
| ENSMUSG0000003015 | chr6 | Clec2e | -2.740628482 | 0.041279982 |
| ENSMUSG0000002923 | chr5 | Nmu | -0.655272364 | 0.041588271 |
| ENSMUSG0000002265 | chr16 | Nectin3 | 0.661203953 | 0.041588271 |
| ENSMUSG0000004715 | chr4 | Cyp4x1 | -0.75854974 | 0.041877819 |
| ENSMUSG0000004114 | chr1 | Dnah7b | -0.580658049 | 0.041877819 |
| ENSMUSG0000004676 | chr15 | Fam83h | -1.143355823 | 0.041902525 |
| ENSMUSG0000002117 | chr12 | Psmc1 | 0.377992755 | 0.041902525 |
| ENSMUSG0000003910 | chr3 | Nexn | 0.474657405 | 0.041902525 |
| ENSMUSG0000007531 | chr2 | Scn9a | -0.683340033 | 0.041952272 |
| ENSMUSG0000002758 | chr2 | Helz2 | -0.605695208 | 0.042267354 |
| ENSMUSG0000002077 | chr11 | Acox1 | -0.48389953 | 0.042267354 |
| ENSMUSG0000004173 | chr9 | Tmem45b | -3.149207627 | 0.042361738 |
| ENSMUSG0000003630 | chr16 | Zdhhc23 | -1.226373255 | 0.042361738 |
| ENSMUSG0000001795 | chr2 | Hnf4a | -3.035030014 | 0.042578497 |
| ENSMUSG0000005009 | chr8 | Ces2b | -4.145498339 | 0.042681833 |
| ENSMUSG0000003578 | chr19 | Acta2 | 0.603611827 | 0.042759052 |
| ENSMUSG0000008568 | chr11 | Tmem238l | -3.079309006 | 0.042818199 |
| ENSMUSG0000004930 | chr9 | Fut4 | -0.921919514 | 0.042818199 |
| ENSMUSG0000003137 | chrX | Bgn | 0.454058345 | 0.042991911 |
| ENSMUSG0000004704 | chr11 | Prr15l | -1.25226675 | 0.043407657 |
| ENSMUSG0000003139 | chrX | Plxna3 | -0.619587383 | 0.043508674 |
| ENSMUSG0000003808 | chr13 | Kdm1b | 0.489938151 | 0.043508674 |
| ENSMUSG0000004309 | chr15 | Tuba1c | 0.570979159 | 0.043508674 |
| ENSMUSG0000000054 | chr1 | Gpa33 | -3.167220362 | 0.043575238 |
| ENSMUSG0000005615 | chr11 | Car10 | -0.744083367 | 0.043575238 |
| ENSMUSG0000005498 | chr3 | Agtr1b | -3.559922972 | 0.043594706 |
| ENSMUSG0000003706 | chr7 | Cavin3 | 0.574801427 | 0.043607655 |
| ENSMUSG0000007406 | chr9 | Fbxw15 | -1.335644846 | 0.043616908 |
| ENSMUSG0000003162 | chr8 | Sorbs2 | 0.448600725 | 0.043784443 |
| ENSMUSG0000002009 | chr10 | Sgpl1 | -0.469280357 | 0.043815515 |
| ENSMUSG0000006352 | chr4 | Eno1 | 0.536538195 | 0.043854366 |

| | | | | |
|---|---|---|---|---|
| ENSMUSG0000002903 | chr4 | Arhgef16 | -1.043326335 | 0.043858049 |
| ENSMUSG0000002095 | chr12 | Coch | -1.043024999 | 0.044044294 |
| ENSMUSG0000003210 | chr9 | Nherf4 | -2.325710537 | 0.044214207 |
| ENSMUSG0000001777 | chr11 | Myo1c | 0.405465137 | 0.044279539 |
| ENSMUSG0000003125 | chrX | Srpx2 | 0.7039287 | 0.044279539 |
| ENSMUSG0000000155 | chr11 | Fkbp10 | 0.315967502 | 0.044319662 |
| ENSMUSG0000002986 | chr6 | Ephb6 | -0.631898517 | 0.044859816 |
| ENSMUSG0000003137 | chrX | Atp2b3 | -0.608695665 | 0.044859816 |
| ENSMUSG0000004390 | chr8 | Zfp469 | 1.273268162 | 0.045324025 |
| ENSMUSG0000002452 | chr18 | Lox | 0.712489432 | 0.045684628 |
| ENSMUSG0000000594 | chr11 | Itgae | -2.014025142 | 0.045917995 |
| ENSMUSG0000001856 | chr11 | Cldn7 | -2.765815307 | 0.04595286 |
| ENSMUSG0000003527 | chr4 | Raver2 | 0.360048124 | 0.04595286 |
| ENSMUSG0000001524 | chr4 | Nipsnap3b | -0.447950617 | 0.045970231 |
| ENSMUSG0000002871 | chr4 | Cyp4b1 | -0.6991539 | 0.046269064 |
| ENSMUSG0000003408 | chr1 | Hdlbp | 0.246179286 | 0.046296627 |
| ENSMUSG0000003550 | chr16 | Slc12a8 | -2.002535259 | 0.046468751 |
| ENSMUSG0000003206 | chr9 | Bco2 | -1.716634079 | 0.046468751 |
| ENSMUSG0000002336 | chr6 | Tmem176a | -0.380455003 | 0.046603244 |
| ENSMUSG0000002871 | chr4 | Pdzk1ip1 | -2.421447197 | 0.046895939 |
| ENSMUSG0000002986 | chr6 | Trpv6 | -2.204231003 | 0.04691422 |
| ENSMUSG0000003948 | chr9 | Cntn5 | -0.705326756 | 0.04691422 |
| ENSMUSG0000003852 | chr3 | Car14 | 0.9592259 | 0.04691422 |
| ENSMUSG0000003177 | chr8 | Pllp | -1.48448035 | 0.047118158 |
| ENSMUSG0000003193 | chr9 | Panx1 | -0.501746484 | 0.047118158 |
| ENSMUSG0000004333 | chr4 | Rhbdl2 | -1.872947363 | 0.047632384 |
| ENSMUSG0000004427 | chr17 | Crb3 | -1.571127537 | 0.047632384 |
| ENSMUSG0000003432 | chr18 | Slc26a2 | -1.547643286 | 0.047899799 |
| ENSMUSG0000000794 | chr7 | Phox2a | -0.60659048 | 0.047941425 |
| ENSMUSG0000005318 | chr16 | Cd200l1 | -2.594195298 | 0.047941537 |
| ENSMUSG0000002580 | chr8 | Itgb1 | 0.28424096 | 0.048088214 |
| ENSMUSG0000000673 | chr10 | B4galnt1 | -0.952804479 | 0.048262789 |
| ENSMUSG0000001985 | chr10 | Arfgef3 | -0.552316246 | 0.048311689 |
| ENSMUSG0000001341 | chr9 | Zbtb47 | 0.296325667 | 0.048428243 |
| ENSMUSG0000006254 | chr7 | Syt9 | -0.529657017 | 0.048511459 |
| ENSMUSG0000006146 | chr11 | Obscn | -0.88996293 | 0.048629994 |
| ENSMUSG0000003163 | chr8 | Slc25a4 | 0.443460211 | 0.048629994 |
| ENSMUSG0000000653 | chr1 | Ihh | -3.575425222 | 0.048783282 |
| ENSMUSG0000003226 | chr9 | Elovl4 | -0.499207268 | 0.048940349 |
| ENSMUSG0000004990 | chr5 | Rasl11b | 0.649847663 | 0.049147607 |
| ENSMUSG0000003891 | chr10 | Mtcl3 | -0.66415108 | 0.049204105 |
| ENSMUSG0000002703 | chr2 | Cers6 | -0.580968687 | 0.049706218 |
| ENSMUSG0000003669 | chrX | Zcchc12 | -0.502360506 | 0.049706218 |
| ENSMUSG0000003452 | chr5 | Hsd17b13 | -1.968595006 | 0.049856448 |
| ENSMUSG0000002137 | chr13 | Cap2 | 0.457006191 | 0.049988933 |
| ENSMUSG0000005445 | chr12 | Vsnl1 | -0.707759485 | 0.050290316 |
| ENSMUSG0000004725 | chr18 | Mc4r | -2.204986534 | 0.050312168 |
| ENSMUSG0000007315 | chr6 | Stmp1 | 0.368799138 | 0.050443955 |

| | | | | |
|---|---|---|---|---|
| ENSMUSG0000003481 | chr10 | Celf5 | -0.622090036 | 0.050465805 |
| ENSMUSG0000000489 | chr3 | Bcan | -0.7777439 | 0.050507908 |
| ENSMUSG0000001341 | chr11 | B4galnt2 | -3.167381463 | 0.050562154 |
| ENSMUSG0000003713 | chr1 | Aff3 | 0.570499475 | 0.050660562 |
| ENSMUSG0000004509 | chr6 | Magi1 | 0.338352088 | 0.050864334 |
| ENSMUSG0000002195 | chr14 | Eef1akmt1 | 0.756388294 | 0.050869553 |
| ENSMUSG0000007077 | chr7 | Ceacam20 | -3.148316156 | 0.050967709 |
| ENSMUSG0000005562 | chr7 | B4galnt4 | -0.653095268 | 0.050967709 |
| ENSMUSG0000003559 | chr7 | Mboat7 | -0.310453593 | 0.051331225 |
| ENSMUSG0000004275 | chrX | Bex2 | -0.424679973 | 0.051418612 |
| ENSMUSG0000000727 | chr7 | Scube2 | 0.506062872 | 0.051418612 |
| ENSMUSG0000002967 | chr5 | Eln | 0.794946075 | 0.051418612 |
| ENSMUSG0000007380 | chr4 | Cdkn2b | -1.37404427 | 0.051524362 |
| ENSMUSG0000004068 | chr17 | Kremen2 | -3.961659482 | 0.051547991 |
| ENSMUSG0000003998 | chr5 | Phtf2 | 0.455091637 | 0.051605585 |
| ENSMUSG0000002972 | chr5 | Cyp3a13 | -2.437481372 | 0.051735907 |
| ENSMUSG0000002660 | chr1 | Smyd2 | 0.359165847 | 0.051735907 |
| ENSMUSG0000008724 | chr1 | Alkal1 | 1.122359576 | 0.051735907 |
| ENSMUSG0000001531 | chr10 | Gadd45b | 0.745756911 | 0.051735907 |
| ENSMUSG0000004204 | chr9 | Sln | 0.957008291 | 0.052006891 |
| ENSMUSG0000007153 | chr14 | Gprin2 | -0.740421856 | 0.052060158 |
| ENSMUSG0000003820 | chr3 | Prkab2 | 0.260871427 | 0.052060158 |
| ENSMUSG0000003875 | chr2 | Ptk6 | -2.566905561 | 0.052078499 |
| ENSMUSG0000005690 | chr3 | Usp13 | -0.352515343 | 0.052078499 |
| ENSMUSG0000001714 | chr11 | Brca1 | -1.110678119 | 0.052250735 |
| ENSMUSG0000002796 | chr3 | Olfm3 | -0.493785307 | 0.052397524 |
| ENSMUSG0000002146 | chr13 | Auh | 0.325762772 | 0.052397524 |
| ENSMUSG0000009026 | chr18 | Eif4ebp3 | -1.840652247 | 0.052884063 |
| ENSMUSG0000004242 | chr15 | Mgat3 | -0.496268237 | 0.052884063 |
| ENSMUSG0000003481 | chr2 | Scn7a | -0.440462222 | 0.052884063 |
| ENSMUSG0000003159 | chr8 | Asah1 | 0.275789065 | 0.052884063 |
| ENSMUSG0000000341 | chr4 | Rab3b | 0.560768769 | 0.052884063 |
| ENSMUSG0000003011 | chr6 | Mfap5 | 0.535852443 | 0.052884063 |
| ENSMUSG0000004997 | chr5 | Glt1d1 | -1.531844354 | 0.052898269 |
| ENSMUSG0000000257 | chr10 | Ikzf4 | -0.466551434 | 0.052950172 |
| ENSMUSG0000003825 | chr6 | Hoxa5 | -0.560035581 | 0.053394289 |
| ENSMUSG0000006195 | chr17 | Ppp4r1 | 0.32020915 | 0.053394289 |
| ENSMUSG0000003770 | chr2 | Lzts3 | -0.32226673 | 0.05356198 |
| ENSMUSG0000003127 | chrX | Col4a6 | 0.533701394 | 0.053746054 |
| ENSMUSG0000003916 | chr15 | Dap | 0.327385706 | 0.05388014 |
| ENSMUSG0000002821 | chr4 | Cdh17 | -3.166642219 | 0.054284874 |
| ENSMUSG0000001890 | chr11 | P4ha2 | 0.475685505 | 0.054525711 |
| ENSMUSG0000003851 | chr8 | Grtp1 | -1.05429061 | 0.054588533 |
| ENSMUSG0000001158 | chr17 | Fsd1 | -0.634944183 | 0.054636112 |
| ENSMUSG0000000819 | chr7 | Spib | -3.378291233 | 0.054658559 |
| ENSMUSG0000004004 | chr7 | Tph1 | -3.070367045 | 0.055287222 |
| ENSMUSG0000002955 | chr6 | Tes | 0.546737188 | 0.055582222 |
| ENSMUSG0000002942 | chr5 | Rimbp2 | -0.544373269 | 0.055697432 |

| | | | | |
|---|---|---|---|---|
| ENSMUSG000000981ε | chr10 | Sowahc | -0.344033549 | 0.055697432 |
| ENSMUSG000000405ε | chr5 | Abcb1a | -1.873357175 | 0.055753333 |
| ENSMUSG0000003551 | chr2 | Ntng2 | -0.346278045 | 0.05593235 |
| ENSMUSG000000216ꜫ | chr13 | Ocln | -1.556277665 | 0.055968211 |
| ENSMUSG000000703ꜫ | chr11 | Gpr179 | -0.663777291 | 0.056208992 |
| ENSMUSG000000267C | chr1 | Prdx6 | -0.947157475 | 0.056303391 |
| ENSMUSG000000523Ꞩ | chr12 | Acot4 | -1.413272319 | 0.056303391 |
| ENSMUSG000000241Ꞩ | chr17 | Dusp1 | 0.628776807 | 0.056824607 |
| ENSMUSG000000250ℤ | chr19 | Xpnpep1 | 0.276991298 | 0.056921666 |
| ENSMUSG000000433ε | chrX | Gprasp1 | -0.378321253 | 0.057228085 |
| ENSMUSG000000444Ꞩ | chr10 | Hs3st5 | -0.696687051 | 0.057266282 |
| ENSMUSG000000293ꜫ | chr5 | Antxr2 | 0.294135054 | 0.057472568 |
| ENSMUSG000000306ϵ | chr7 | Calcb | -0.896322576 | 0.057733616 |
| ENSMUSG000000593ϵ | chr2 | Nrsn2 | -0.518500665 | 0.057888331 |
| ENSMUSG000000268ε | chr2 | Grb14 | 0.299917224 | 0.057962145 |
| ENSMUSG0000004531 | chr5 | Sowahb | -2.717948715 | 0.05820477 |
| ENSMUSG000000137ϵ | chr17 | Ly6g6e | -3.175454604 | 0.05820477 |
| ENSMUSG000000326ʇ | chr2 | Cd59a | 0.588604529 | 0.058433313 |
| ENSMUSG000000326Ꞩ | chr5 | Oas2 | -1.361095051 | 0.058543295 |
| ENSMUSG000000745ʇ | chr2 | Kcng1 | 0.564144018 | 0.059129308 |
| ENSMUSG000000712ϵ | chr17 | Zfp946 | -0.397440426 | 0.059152187 |
| ENSMUSG000000306C | chr7 | Mfge8 | 0.413301331 | 0.059152187 |
| ENSMUSG000000548ʇ | chr9 | Tmem158 | -0.555335376 | 0.059192343 |
| ENSMUSG000000008ε | chr2 | Dbh | 0.462418092 | 0.059192343 |
| ENSMUSG000000025ϵ | chr12 | Scin | -2.341934518 | 0.059213204 |
| ENSMUSG000000470ε | chr7 | Lrrc4b | -0.437916231 | 0.060165216 |
| ENSMUSG000000228Ꞩ | chr16 | Ets2 | 0.341376977 | 0.060165216 |
| ENSMUSG000000483C | chr3 | Slitrk3 | -0.599577059 | 0.060196936 |
| ENSMUSG000000335ʇ | chrX | Tmem35a | -0.482912486 | 0.06031244 |
| ENSMUSG000000197Ꞩ | chr10 | Stxbp5 | -0.394991857 | 0.0604913 |
| ENSMUSG000000358ϵ | chr10 | Syt1 | -0.498545996 | 0.060612854 |
| ENSMUSG000000662Ƽ | chr7 | Trim12a | -0.463511948 | 0.060612854 |
| ENSMUSG000000472ϵ | chr16 | Gap43 | -0.457260578 | 0.060638056 |
| ENSMUSG000000317Ꮞ | chr8 | St3gal2 | 0.277790209 | 0.060649036 |
| ENSMUSG000000296ꜫ | chr5 | Cdk8 | 0.33290808 | 0.06070504 |
| ENSMUSG0000001841 | chr11 | Mapt | -0.467848699 | 0.061115365 |
| ENSMUSG000000948Ꮞ | chr5 | Muc3a | -2.185974417 | 0.061403681 |
| ENSMUSG000001010ꜫ | chr19 | Ms4a12 | -3.033862948 | 0.061462207 |
| ENSMUSG000000715Ƽ | chr13 | Akr1c19 | -2.812737235 | 0.061531422 |
| ENSMUSG000000397ε | chr2 | Cercam | 0.341352584 | 0.061531422 |
| ENSMUSG000000007Ꞩ | chr8 | Slc5a5 | -0.838350143 | 0.061541663 |
| ENSMUSG000000004Ꮞ | chr6 | Raf1 | 0.279096012 | 0.061541663 |
| ENSMUSG000000204C | chr11 | Upp1 | -1.256051344 | 0.061955406 |
| ENSMUSG000000244ʇ | chr18 | Dcp2 | -0.307363969 | 0.062077535 |
| ENSMUSG000000056Ꞩ | chr8 | Pard6a | 0.609195993 | 0.062101962 |
| ENSMUSG000000176C | chr11 | Tns4 | -2.8743805 | 0.062361729 |
| ENSMUSG000000907C | chr17 | Cyp4f40 | -3.228521629 | 0.062361729 |
| ENSMUSG000000173ε | chr11 | Traf4 | -0.683878682 | 0.062361729 |

| | | | | |
|---|---|---|---|---|
| ENSMUSG000000408 | chr17 | Plekhh2 | 0.506606536 | 0.062361729 |
| ENSMUSG000000228 | chr16 | Clcn2 | -0.882970697 | 0.06240495 |
| ENSMUSG000000540 | chr7 | Pkp3 | -1.233744508 | 0.06240495 |
| ENSMUSG0000003911 | chr11 | Mycbpap | -1.147709654 | 0.062531234 |
| ENSMUSG0000004992 | chr2 | Slc35c1 | -0.758917438 | 0.062883489 |
| ENSMUSG0000005612 | chr17 | Fez2 | 0.481762471 | 0.063008527 |
| ENSMUSG000000159 | chr2 | Dstn | 0.462310237 | 0.063126238 |
| ENSMUSG0000001652 | chr5 | Lnx2 | 0.352210298 | 0.063126238 |
| ENSMUSG000000296 | chr6 | Col1a2 | 0.413598821 | 0.063497851 |
| ENSMUSG000000156 | chr17 | Srf | 0.31328002 | 0.063502107 |
| ENSMUSG000000290 | chr4 | Mmp23 | 0.466781217 | 0.063724692 |
| ENSMUSG0000001997 | chr10 | Ikbip | 0.301961774 | 0.063861086 |
| ENSMUSG000000510 | chr10 | Lingo3 | -1.789296503 | 0.064622723 |
| ENSMUSG000000308 | chr7 | Chp2 | -0.909645319 | 0.064734414 |
| ENSMUSG000000637 | chr3 | Hsd3b2 | -3.399744149 | 0.065877339 |
| ENSMUSG000000282 | chr3 | Clca1 | -2.800441548 | 0.066103463 |
| ENSMUSG0000005597 | chr7 | Fut2 | -2.910521283 | 0.066178465 |
| ENSMUSG000000378 | chr3 | Pcdh18 | 0.367420498 | 0.066324235 |
| ENSMUSG000000512 | chr10 | Msrb3 | 0.360110076 | 0.066334601 |
| ENSMUSG0000003347 | chr14 | Cysltr2 | -0.728422352 | 0.066504843 |
| ENSMUSG000000735 | chr18 | Iigp1c | -1.269226407 | 0.066609683 |
| ENSMUSG0000005714 | chr7 | Trim12c | -0.428780938 | 0.066609683 |
| ENSMUSG000000399 | chr19 | Dtx4 | -0.449334592 | 0.066681509 |
| ENSMUSG000000456 | chr18 | Pcdhb4 | -0.826079878 | 0.066682943 |
| ENSMUSG000000173 | chr11 | Aldoc | -0.523504148 | 0.066774668 |
| ENSMUSG0000006327 | chr2 | Hacd1 | 0.479435451 | 0.067785851 |
| ENSMUSG000000356 | chr10 | Kcnc2 | -0.553712199 | 0.067795321 |
| ENSMUSG0000001917 | chr5 | Mdh2 | 0.392292801 | 0.068456242 |
| ENSMUSG000000228 | chr16 | Lmln | -0.470331409 | 0.068539565 |
| ENSMUSG000000461 | chr9 | Cd109 | 0.519000349 | 0.068677642 |
| ENSMUSG000000176 | chr3 | Hnf4g | -2.137483818 | 0.06935396 |
| ENSMUSG0000003661 | chr2 | Etl4 | -0.411382741 | 0.06935396 |
| ENSMUSG000000229 | chr16 | Parp9 | -0.450242724 | 0.069366137 |
| ENSMUSG0000006251 | chr3 | Fabp4 | 0.599717779 | 0.069366137 |
| ENSMUSG000000445 | chrX | Tceal3 | -0.386406876 | 0.069496484 |
| ENSMUSG000000254 | chr10 | Gli1 | -0.827286914 | 0.069528272 |
| ENSMUSG000000492 | chr9 | Scn3b | -0.53234229 | 0.069528272 |
| ENSMUSG0000002682 | chr2 | Kcnj3 | -0.456890671 | 0.069528272 |
| ENSMUSG000000708 | chr7 | Pnma8b | -0.473405158 | 0.069694864 |
| ENSMUSG000000344 | chr13 | Edil3 | -0.431384432 | 0.069763059 |
| ENSMUSG000000712 | chr13 | Naip5 | -1.895720541 | 0.069964706 |
| ENSMUSG000000312 | chrX | Sytl4 | -0.968881322 | 0.069964706 |
| ENSMUSG0000003691 | chr8 | Trim67 | -0.711281299 | 0.069964706 |
| ENSMUSG000000360 | chr2 | Adamtsl2 | 0.853499114 | 0.069964706 |
| ENSMUSG000000303 | chr6 | Slc6a11 | 1.562768165 | 0.069964706 |
| ENSMUSG000000347 | chr2 | Nostrin | -1.188593731 | 0.07029868 |
| ENSMUSG000000668 | chr5 | Oas1e | -2.398394008 | 0.07029868 |
| ENSMUSG000000748 | chr19 | Hpse2 | 0.388219744 | 0.070507636 |

| | | | | |
|---|---|---|---|---|
| ENSMUSG0000003165 | chr8 | Adcy7 | 0.523136635 | 0.070869927 |
| ENSMUSG0000003657 | chr7 | Fxyd1 | 0.333786137 | 0.071537059 |
| ENSMUSG0000005256 | chr8 | Hook2 | -0.599488655 | 0.071856529 |
| ENSMUSG0000001902 | chr14 | Dnah1 | -0.568666796 | 0.072020603 |
| ENSMUSG0000003836 | chr8 | Fbxo25 | -0.277619686 | 0.072020603 |
| ENSMUSG0000003163 | chr8 | Tlr3 | -0.524662217 | 0.072023472 |
| ENSMUSG0000002954 | chr5 | Cabp1 | -0.616373425 | 0.072057263 |
| ENSMUSG0000002612 | chr1 | Ptpn18 | -1.230596698 | 0.072058653 |
| ENSMUSG0000007400 | chr7 | Omp | -1.099770959 | 0.072058653 |
| ENSMUSG0000004015 | chr11 | Tax1bp3 | 0.405879062 | 0.072058653 |
| ENSMUSG0000000305 | chr1 | Elf3 | -2.879996826 | 0.072560304 |
| ENSMUSG0000003199 | chr9 | Adamts8 | 0.438153985 | 0.072560304 |
| ENSMUSG0000002926 | chr5 | Ugt2b34 | -3.021159801 | 0.073578689 |
| ENSMUSG0000002921 | chr5 | Gabrb1 | -0.935886353 | 0.073578689 |
| ENSMUSG0000002577 | chr1 | Gdap1 | -0.466984689 | 0.073630701 |
| ENSMUSG0000006803 | chr17 | Tcp1 | 0.404164038 | 0.073829865 |
| ENSMUSG0000002622 | chr1 | Sp100 | -0.556299066 | 0.073871955 |
| ENSMUSG0000002795 | chr3 | Slc35a3 | -0.48812971 | 0.073962889 |
| ENSMUSG0000004961 | chr11 | Omg | -0.56788305 | 0.074023104 |
| ENSMUSG0000003111 | chrX | Gpc4 | 0.323624365 | 0.074102474 |
| ENSMUSG0000000115 | chr6 | Mxd1 | -0.90806797 | 0.07418654 |
| ENSMUSG0000006152 | chr15 | Krt5 | 1.844958099 | 0.075217415 |
| ENSMUSG0000002245 | chr15 | Septin3 | -0.469598102 | 0.075229172 |
| ENSMUSG0000003829 | chr5 | Atg9b | -0.670160412 | 0.075267553 |
| ENSMUSG0000002802 | chr3 | Aimp1 | 0.410965062 | 0.075267553 |
| ENSMUSG0000007570 | chr17 | Msrb1 | 0.496524811 | 0.075640144 |
| ENSMUSG0000003318 | chr16 | BC016579 | -2.718031312 | 0.075856458 |
| ENSMUSG0000002745 | chr2 | Fam110a | 0.416100146 | 0.075921675 |
| ENSMUSG0000005485 | chrX | Smim10l2a | -0.925499618 | 0.075940364 |
| ENSMUSG0000000147 | chr18 | Tubb6 | 0.578823119 | 0.076772798 |
| ENSMUSG0000002386 | chr17 | Mpc1 | 0.362909147 | 0.076867886 |
| ENSMUSG0000003850 | chr17 | Prr3 | -0.317721334 | 0.076890339 |
| ENSMUSG0000000281 | chr11 | Flii | 0.234436667 | 0.077263198 |
| ENSMUSG0000003253 | chr9 | Trak1 | 0.262028763 | 0.077447735 |
| ENSMUSG0000003721 | chrX | Syn1 | -0.469145897 | 0.077508781 |
| ENSMUSG0000004165 | chrX | Rragb | -0.402773847 | 0.07781997 |
| ENSMUSG0000003116 | chrX | Glod5 | -2.175798314 | 0.07827825 |
| ENSMUSG0000002542 | chr18 | Ark2c | 0.683984749 | 0.07827825 |
| ENSMUSG0000003065 | chr7 | Tmc5 | -3.254883533 | 0.078704501 |
| ENSMUSG0000004958 | chr7 | Tsku | -1.062529835 | 0.078704501 |
| ENSMUSG0000005109 | chr13 | Mblac2 | -0.467813197 | 0.078704501 |
| ENSMUSG0000004428 | chr4 | Cnr1 | -0.468905619 | 0.078803128 |
| ENSMUSG0000002964 | chr5 | Polr1d | 0.554622111 | 0.078847695 |
| ENSMUSG0000001479 | chr8 | Elmo3 | -0.614294208 | 0.079177468 |
| ENSMUSG0000002728 | chr2 | Snap23 | 0.34166176 | 0.079493144 |
| ENSMUSG0000003034 | chr6 | Cd9 | 0.459088136 | 0.079794801 |
| ENSMUSG0000006131 | chr10 | Naca | 0.426286767 | 0.080134204 |
| ENSMUSG0000005235 | chr7 | Cemip | 0.60799255 | 0.080382123 |

| | | | | |
|---|---|---|---|---|
| ENSMUSG0000003086 | chr7 | Ern2 | -3.503798863 | 0.080634964 |
| ENSMUSG0000009017 | chr11 | Fbxw10 | -0.794976026 | 0.080634964 |
| ENSMUSG0000000069 | chr11 | Hoxb6 | -0.628557127 | 0.080664731 |
| ENSMUSG0000004048 | chr11 | Xaf1 | -0.693372906 | 0.080681298 |
| ENSMUSG0000001932 | chr11 | Aoc3 | 0.31594121 | 0.080825776 |
| ENSMUSG0000005098 | chr4 | Selenon | 0.266600003 | 0.080825776 |
| ENSMUSG0000009203 | chr6 | Peg10 | 0.564791983 | 0.0808413 |
| ENSMUSG0000002977 | chr6 | Adcyap1r1 | -0.413732264 | 0.080888688 |
| ENSMUSG0000006050 | chr7 | Nlrp9b | -2.240861293 | 0.080895505 |
| ENSMUSG0000002566 | chrX | Rps6ka6 | -0.603859057 | 0.081029515 |
| ENSMUSG0000000178 | chr10 | Fbxo7 | -0.370965111 | 0.081309823 |
| ENSMUSG0000002146 | chr13 | Ror2 | 0.379744735 | 0.081309823 |
| ENSMUSG0000002803 | chr3 | Ptgfr | 0.847822615 | 0.081424296 |
| ENSMUSG0000002918 | chr5 | Pi4k2b | -0.82600715 | 0.081766892 |
| ENSMUSG0000006071 | chr12 | Plekhh1 | -0.983602514 | 0.082031446 |
| ENSMUSG0000009208 | chr1 | Kcnb2 | -0.486892631 | 0.082031446 |
| ENSMUSG0000003073 | chr7 | Syt3 | -0.604456015 | 0.082176753 |
| ENSMUSG0000001883 | chr16 | Myh11 | 0.362861961 | 0.082176753 |
| ENSMUSG0000003308 | chr15 | Triobp | 0.324644965 | 0.082176753 |
| ENSMUSG0000003470 | chr2 | Neurod1 | -3.921111884 | 0.082378964 |
| ENSMUSG0000002796 | chr3 | Vcam1 | -0.544938968 | 0.082378964 |
| ENSMUSG0000004159 | chr14 | Tmtc4 | -0.414896489 | 0.082378964 |
| ENSMUSG0000009151 | chr3 | Lamtor3 | 0.359769514 | 0.082378964 |
| ENSMUSG0000000380 | chr8 | Farsa | 0.284427832 | 0.082745179 |
| ENSMUSG0000003076 | chr7 | Aqp8 | -3.441358257 | 0.082921619 |
| ENSMUSG0000003233 | chr9 | Nptn | 0.285034311 | 0.083625626 |
| ENSMUSG0000003021 | chr6 | Mgp | 0.616157599 | 0.083625626 |
| ENSMUSG0000004504 | chr19 | Lrfn4 | -0.379791165 | 0.083837373 |
| ENSMUSG0000002576 | chr3 | Larp1b | 0.285786467 | 0.084751943 |
| ENSMUSG0000002741 | chr2 | Pcsk2 | -0.399626644 | 0.085439155 |
| ENSMUSG0000003786 | chr10 | Egr2 | 1.62005125 | 0.085439155 |
| ENSMUSG0000002692 | chr2 | Agpat2 | -0.53591971 | 0.08602073 |
| ENSMUSG0000002281 | chr16 | Itgb5 | 0.30644889 | 0.086062534 |
| ENSMUSG0000003129 | chrX | Slc7a3 | -0.712172347 | 0.086509578 |
| ENSMUSG0000007542 | chr11 | Smim6 | -1.545672898 | 0.086594873 |
| ENSMUSG0000003055 | chr7 | Synm | 0.485041823 | 0.086810195 |
| ENSMUSG0000003799 | chr5 | Arap2 | -0.523538258 | 0.087043478 |
| ENSMUSG0000002479 | chr19 | Htr7 | -0.760902025 | 0.087169838 |
| ENSMUSG0000001534 | chrX | Xk | 0.254724541 | 0.08756199 |
| ENSMUSG0000005753 | chr13 | Dtnbp1 | 0.357013943 | 0.087615961 |
| ENSMUSG0000005052 | chr16 | Cldn8 | -3.249184818 | 0.0881783 |
| ENSMUSG0000006721 | chr17 | H2-T23 | -0.709473298 | 0.088214791 |
| ENSMUSG0000002787 | chr3 | Reg4 | -3.547930435 | 0.088267592 |
| ENSMUSG0000002237 | chr15 | Asap1 | 0.31709534 | 0.088389434 |
| ENSMUSG0000003749 | chr1 | Nenf | 0.57586086 | 0.088389434 |
| ENSMUSG0000002871 | chr4 | Stil | -1.748645461 | 0.088403476 |
| ENSMUSG0000001521 | chrX | Mtmr1 | 0.336442487 | 0.088448236 |
| ENSMUSG0000005150 | chr5 | Ufsp1 | -0.466974831 | 0.088511019 |

| | | | | |
|---|---|---|---|---|
| ENSMUSG0000003382 | chr17 | Dnah8 | -1.615427016 | 0.088558284 |
| ENSMUSG0000005364 | chr9 | Plxnb1 | -0.432554769 | 0.088558284 |
| ENSMUSG0000006282 | chr11 | Actg1 | 0.379001705 | 0.088803585 |
| ENSMUSG0000009081 | chr12 | Samd15 | -1.478202006 | 0.089046436 |
| ENSMUSG0000002057 | chr12 | Tspan13 | -0.312798019 | 0.089046436 |
| ENSMUSG0000002847 | chr4 | Gne | -0.499768446 | 0.089065272 |
| ENSMUSG0000006113 | chr19 | Blnk | -0.946740353 | 0.089183095 |
| ENSMUSG0000002082 | chr11 | Nos2 | -2.166160756 | 0.089183095 |
| ENSMUSG0000003198 | chr8 | Capn9 | -1.735616172 | 0.090143794 |
| ENSMUSG0000002736 | chr2 | Sppl2a | -0.422785708 | 0.090227359 |
| ENSMUSG0000000593 | chr17 | Kctd20 | 0.252809971 | 0.090501733 |
| ENSMUSG0000003079 | chr7 | Pycard | -0.752889468 | 0.090508295 |
| ENSMUSG0000006215 | chr4 | Ifnlr1 | -0.722876499 | 0.090508295 |
| ENSMUSG0000005622 | chr1 | Pla2g4a | 0.353516859 | 0.090508295 |
| ENSMUSG0000004583 | chr2 | Ccdc9b | 0.376187176 | 0.090508295 |
| ENSMUSG0000002964 | chr5 | Cdx2 | -3.426435789 | 0.090753193 |
| ENSMUSG0000001977 | chr10 | Rgs17 | -0.507466859 | 0.090753193 |
| ENSMUSG0000001648 | chr6 | Ppfibp1 | 0.291873257 | 0.090753193 |
| ENSMUSG0000004420 | chrX | Vsig4 | 1.55695677 | 0.091204785 |
| ENSMUSG0000002609 | chr1 | Stk17b | 0.393580081 | 0.091207231 |
| ENSMUSG0000004899 | chr17 | Prss32 | -2.95510355 | 0.091255006 |
| ENSMUSG0000002205 | chr14 | Adamdec1 | -1.034226282 | 0.091479456 |
| ENSMUSG0000002748 | chr2 | Snta1 | 0.424673747 | 0.091763396 |
| ENSMUSG0000002837 | chr4 | Ptgr1 | 0.399937658 | 0.09191796 |
| ENSMUSG0000003845 | chr8 | Tmem255b | -0.578933476 | 0.092461348 |
| ENSMUSG0000002391 | chr17 | Mep1a | -2.812778354 | 0.092807397 |
| ENSMUSG0000002982 | chr6 | Osbpl3 | 0.425465638 | 0.093256438 |
| ENSMUSG0000004378 | chr17 | Bicdl2 | -1.585435557 | 0.093431455 |
| ENSMUSG0000002627 | chr1 | Septin2 | 0.228766316 | 0.094056842 |
| ENSMUSG0000004900 | chr6 | Ndnf | 0.606863539 | 0.095316438 |
| ENSMUSG0000000247 | chr18 | Abhd3 | -0.674306365 | 0.095536566 |
| ENSMUSG0000003569 | chr16 | Slc51a | -2.488115131 | 0.095803287 |
| ENSMUSG0000006153 | chr2 | Tmem236 | -2.599904072 | 0.095857471 |
| ENSMUSG0000003120 | chrX | Rab39b | -0.427017289 | 0.095857471 |
| ENSMUSG0000002698 | chr2 | Ly75 | -0.866959826 | 0.095990272 |
| ENSMUSG0000004072 | chr4 | Esrp1 | -3.378252754 | 0.09609638 |
| ENSMUSG0000002054 | chr12 | Bzw2 | 0.435420066 | 0.096119744 |
| ENSMUSG0000006096 | chr5 | Slc4a4 | -0.459455659 | 0.096177083 |
| ENSMUSG0000002228 | chr15 | Grhl2 | -1.430077715 | 0.097242672 |
| ENSMUSG0000003206 | chr9 | Cryab | 0.530578461 | 0.097278697 |
| ENSMUSG0000007956 | chr6 | Hoxa3 | -0.311299673 | 0.098886606 |
| ENSMUSG0000009094 | chr18 | F830016B08Rik | -1.596676751 | 0.099229024 |
| ENSMUSG0000002883 | chr4 | Ncdn | -0.326947602 | 0.099362476 |
| ENSMUSG0000003046 | chr8 | Psd3 | -0.320642846 | 0.099901903 |
| ENSMUSG0000005806 | chr17 | Trim31 | -3.197434383 | 0.100665991 |
| ENSMUSG0000003986 | chr3 | Slc44a3 | -1.254092478 | 0.100665991 |
| ENSMUSG0000005958 | chr2 | Calcrl | -0.393719931 | 0.100665991 |
| ENSMUSG0000002390 | chr17 | Paqr4 | -0.382434984 | 0.100665991 |

| | | | | |
|---|---|---|---|---|
| ENSMUSG0000005573 | chrX | Nap1l3 | -0.294636378 | 0.100665991 |
| ENSMUSG0000003635 | chr2 | Ubac1 | 0.29459211 | 0.100665991 |
| ENSMUSG0000000929 | chr10 | Ube2g2 | 0.366342464 | 0.100665991 |
| ENSMUSG0000004243 | chr11 | Mfap4 | 0.421886981 | 0.100665991 |
| ENSMUSG0000000130 | chr1 | Rrp15 | 0.451371063 | 0.100665991 |
| ENSMUSG0000003905 | chr8 | Myo16 | 1.084645436 | 0.100665991 |
| ENSMUSG0000005070 | chr2 | Emilin3 | -0.883592551 | 0.100975957 |
| ENSMUSG0000002803 | chr3 | Ifi44 | -1.301393585 | 0.100988202 |
| ENSMUSG0000000353 | chr16 | Dgcr6 | 0.425627049 | 0.100999516 |
| ENSMUSG0000002812 | chr3 | Abcd3 | -0.293362053 | 0.101297708 |
| ENSMUSG0000002506 | chr19 | Gsto1 | -0.582616385 | 0.10172109 |
| ENSMUSG0000003110 | chrX | Enox2 | -0.393265142 | 0.102019784 |
| ENSMUSG0000002729 | chr2 | Tyro3 | -0.483815721 | 0.102179069 |
| ENSMUSG0000003390 | chrX | Zdhhc15 | -0.393269507 | 0.102311536 |
| ENSMUSG0000003542 | chr7 | Ptprh | -2.895837204 | 0.102421009 |
| ENSMUSG0000002232 | chr15 | Cdh10 | -0.687564736 | 0.102615832 |
| ENSMUSG0000003188 | chr8 | Car7 | 0.838518995 | 0.103045261 |
| ENSMUSG0000003689 | chr2 | Gtdc1 | 0.34671427 | 0.103315855 |
| ENSMUSG0000003187 | chr9 | Pgr | -0.633963123 | 0.104059809 |
| ENSMUSG0000006123 | chr17 | H2-K1 | -0.39328607 | 0.104579557 |
| ENSMUSG0000003068 | chr7 | Sez6l2 | -0.511939323 | 0.104612601 |
| ENSMUSG0000002449 | chr18 | Lars1 | 0.278387523 | 0.104612601 |
| ENSMUSG0000002962 | chr5 | Zkscan14 | 0.474032264 | 0.104612601 |
| ENSMUSG0000004935 | chr7 | Zg16 | -2.78944611 | 0.105828295 |
| ENSMUSG0000005172 | chr7 | Kctd14 | -0.91670647 | 0.106018921 |
| ENSMUSG0000005047 | chr10 | Slc35d3 | -0.50855996 | 0.106070528 |
| ENSMUSG0000003252 | chr9 | Hhatl | 0.495908223 | 0.106070528 |
| ENSMUSG0000003417 | chr4 | Faah | -0.706240825 | 0.106091564 |
| ENSMUSG0000002806 | chr3 | Lmna | 0.443227453 | 0.106266919 |
| ENSMUSG0000002729 | chr2 | Ltk | -0.582191624 | 0.106399962 |
| ENSMUSG0000004045 | chr19 | Sgms1 | 0.387250648 | 0.106627635 |
| ENSMUSG0000004617 | chr7 | E2f8 | -1.412095302 | 0.107421091 |
| ENSMUSG0000004912 | chr4 | Frmd3 | -0.605092229 | 0.107421091 |
| ENSMUSG0000003922 | chr17 | Ppp1r10 | 0.299070463 | 0.107568286 |
| ENSMUSG0000002280 | chr16 | Nr1i2 | -2.869727292 | 0.108951284 |
| ENSMUSG0000004828 | chr12 | Frmd6 | 0.315058753 | 0.109125273 |
| ENSMUSG0000002281 | chr16 | Umps | 0.354012018 | 0.109125273 |
| ENSMUSG0000006073 | chr13 | Nsa2 | 0.391092431 | 0.109125273 |
| ENSMUSG0000007892 | chr11 | Ifi47 | -0.882682684 | 0.109174889 |
| ENSMUSG0000001986 | chr10 | Crybg1 | -0.752284206 | 0.109174889 |
| ENSMUSG0000003245 | chr9 | Mrps22 | 0.443078382 | 0.109740377 |
| ENSMUSG0000003554 | chr7 | Capn5 | -0.471339227 | 0.110086006 |
| ENSMUSG0000002696 | chr2 | Lrrc26 | -2.155148702 | 0.110177487 |
| ENSMUSG0000002242 | chr15 | Enpp2 | 0.256780145 | 0.110177487 |
| ENSMUSG0000001996 | chr10 | Kitl | 0.287878051 | 0.110177487 |
| ENSMUSG0000007030 | chr9 | Mpzl3 | -0.670439255 | 0.110556295 |
| ENSMUSG0000005436 | chr12 | Rhob | 0.308333091 | 0.110879009 |
| ENSMUSG0000003256 | chr9 | Acp3 | -1.626174881 | 0.110887356 |

| | | | | |
|---|---|---|---|---|
| ENSMUSG0000005233 | chr9 | Cx3cr1 | -0.784067641 | 0.110887356 |
| ENSMUSG0000003357 | chr15 | Apol6 | 0.650125306 | 0.110887356 |
| ENSMUSG0000001777 | chr13 | Cox7c | 0.507040742 | 0.111218874 |
| ENSMUSG0000003073 | chr7 | Chrdl2 | -1.151351889 | 0.111960644 |
| ENSMUSG0000006215 | chr9 | Unc13c | -0.457530476 | 0.111960644 |
| ENSMUSG0000003738 | chr15 | Rims2 | -0.418308839 | 0.111960644 |
| ENSMUSG0000002299 | chr1 | Enah | 0.471530759 | 0.111960644 |
| ENSMUSG0000004392 | chr4 | Ncmap | -1.528467314 | 0.11262578 |
| ENSMUSG0000001856 | chr11 | Elp5 | 0.321410118 | 0.112884653 |
| ENSMUSG0000001683 | chr5 | Mrps18c | 0.371098626 | 0.112884653 |
| ENSMUSG0000004373 | chr5 | Ptpn11 | 0.288632976 | 0.112944114 |
| ENSMUSG0000003114 | chrX | Syp | -0.5147619 | 0.113702533 |
| ENSMUSG0000007341 | chr17 | C4b | -0.579400764 | 0.113782592 |
| ENSMUSG0000005532 | chr7 | Tead1 | 0.399176995 | 0.113830021 |
| ENSMUSG0000002546 | chr7 | Prap1 | -2.010302843 | 0.114310287 |
| ENSMUSG0000007933 | chr19 | Ifit1bl1 | -0.919411062 | 0.11519774 |
| ENSMUSG0000002090 | chr11 | Myh3 | 1.789335398 | 0.115483971 |
| ENSMUSG0000002295 | chr16 | Atp5po | 0.462442134 | 0.115798108 |
| ENSMUSG0000007304 | chr6 | Atoh1 | -3.301999314 | 0.11600117 |
| ENSMUSG0000006191 | chr12 | Myt1l | -0.404133889 | 0.11600117 |
| ENSMUSG0000007359 | chr18 | Ecscr | 0.33986311 | 0.11600117 |
| ENSMUSG0000004053 | chr4 | Necab1 | -0.411729647 | 0.11600807 |
| ENSMUSG0000003077 | chr7 | Pak1 | -0.26576277 | 0.116362861 |
| ENSMUSG0000002439 | chr18 | Lims2 | 0.459508563 | 0.117004122 |
| ENSMUSG0000003072 | chr7 | Pgm2l1 | -0.513944952 | 0.117064171 |
| ENSMUSG0000003411 | chr8 | Atp2c2 | -1.706958912 | 0.117106305 |
| ENSMUSG0000003134 | chrX | Gabra3 | -0.54375124 | 0.117235691 |
| ENSMUSG0000003676 | chr9 | Kif15 | -1.404722574 | 0.117414993 |
| ENSMUSG0000000495 | chr5 | Rasa4 | -0.538859107 | 0.117583375 |
| ENSMUSG0000005230 | chr7 | Doc2a | -1.731033742 | 0.117937556 |
| ENSMUSG0000004949 | chr9 | Pls1 | -2.473064059 | 0.117962717 |
| ENSMUSG0000002407 | chr17 | Prkd3 | 0.268825999 | 0.117962717 |
| ENSMUSG0000002459 | chr18 | Fbn2 | 0.584383726 | 0.117962717 |
| ENSMUSG0000002808 | chr3 | Sh3d19 | 0.300834034 | 0.118139589 |
| ENSMUSG0000005131 | chr7 | Ffar2 | -3.039008927 | 0.118151578 |
| ENSMUSG0000005537 | chr4 | Fut9 | -1.082464305 | 0.118151578 |
| ENSMUSG0000002387 | chr17 | 1700010I14Rik | -0.935689632 | 0.118151578 |
| ENSMUSG0000007264 | chr11 | Lyrm9 | -0.313932598 | 0.118151578 |
| ENSMUSG0000002183 | chr14 | Samd4 | 0.41558768 | 0.118151578 |
| ENSMUSG0000004684 | chr8 | Vat1l | -0.389107897 | 0.119023624 |
| ENSMUSG0000002892 | chr4 | Padi2 | -0.831774842 | 0.119703371 |
| ENSMUSG0000001881 | chr7 | Lsp1 | 0.331538629 | 0.119893633 |
| ENSMUSG0000007400 | chr9 | Klhl40 | 1.210980545 | 0.120041828 |
| ENSMUSG0000004177 | chr16 | Ydjc | -0.803887415 | 0.120226372 |
| ENSMUSG0000005172 | chr12 | Kcnf1 | 0.818831719 | 0.120226372 |
| ENSMUSG0000004143 | chr1 | Mfsd6 | -0.256844564 | 0.120787856 |
| ENSMUSG0000001837 | chr11 | Cuedc1 | 0.309378599 | 0.120787856 |
| ENSMUSG0000002763 | chr2 | Rab5if | 0.575286163 | 0.120787856 |

| | | | | |
|---|---|---|---|---|
| ENSMUSG0000002390 | chr17 | Tnfrsf12a | 0.739642993 | 0.120787856 |
| ENSMUSG0000002875 | chr4 | Kif17 | -0.965644766 | 0.120798782 |
| ENSMUSG0000003129 | chrX | Pdha1 | 0.206179429 | 0.120910191 |
| ENSMUSG0000002408 | chr17 | Cyp1b1 | -0.617554913 | 0.121085443 |
| ENSMUSG0000002808 | chr3 | Fmo5 | -0.84915764 | 0.121265228 |
| ENSMUSG0000003543 | chr12 | Sstr1 | -0.719324127 | 0.121265228 |
| ENSMUSG0000005733 | chr10 | Chst3 | 0.95930545 | 0.122089869 |
| ENSMUSG0000002146 | chr13 | Ptch1 | -0.323913185 | 0.122132121 |
| ENSMUSG0000002006 | chr10 | Herc4 | 0.274097392 | 0.122132121 |
| ENSMUSG0000000484 | chr5 | Plod3 | 0.30997178 | 0.122132121 |
| ENSMUSG0000000411 | chr2 | Cacna1b | -0.527448585 | 0.122419459 |
| ENSMUSG0000011212 | chr10 | Pbld1 | -1.938236743 | 0.122419459 |
| ENSMUSG0000003679 | chrX | Slitrk2 | -0.71432435 | 0.122419459 |
| ENSMUSG0000002765 | chr2 | Fam83d | 0.461949959 | 0.122422747 |
| ENSMUSG0000002137 | chr13 | Kif13a | 0.271125153 | 0.122643512 |
| ENSMUSG0000004338 | chr5 | Tmem130 | -0.512047066 | 0.123009736 |
| ENSMUSG0000000813 | chr1 | Fhl2 | 0.473382212 | 0.123123832 |
| ENSMUSG0000005053 | chr11 | B230217C12Rik | -0.463518285 | 0.123432113 |
| ENSMUSG0000003968 | chr5 | Lap3 | 0.324077416 | 0.123432113 |
| ENSMUSG0000002979 | chr6 | Sspo | -0.910668986 | 0.123510508 |
| ENSMUSG0000003499 | chr14 | Htr2a | -0.815108175 | 0.123528403 |
| ENSMUSG0000004020 | chr9 | Pclaf | -1.019188907 | 0.123765713 |
| ENSMUSG0000004812 | chr1 | Col6a3 | 0.351067379 | 0.123776297 |
| ENSMUSG0000003149 | chr8 | Chrna6 | -1.246615241 | 0.123925413 |
| ENSMUSG0000003196 | chr8 | Aars1 | 0.209710314 | 0.123925413 |
| ENSMUSG0000002724 | chr2 | Arhgap1 | 0.255106472 | 0.123925413 |
| ENSMUSG0000002672 | chr2 | Vim | 0.321717868 | 0.123925413 |
| ENSMUSG0000004544 | chr6 | Gprin3 | -1.141290196 | 0.12516939 |
| ENSMUSG0000002864 | chr4 | P3h1 | 0.288161743 | 0.125239803 |
| ENSMUSG0000002929 | chr5 | Rufy3 | -0.307162147 | 0.125491922 |
| ENSMUSG0000004870 | chr4 | Lurap1l | 0.428516695 | 0.125766365 |
| ENSMUSG0000003808 | chr9 | Hspb2 | 0.78968637 | 0.126116753 |
| ENSMUSG0000002156 | chr13 | Nkd2 | 0.331932605 | 0.126304546 |
| ENSMUSG0000003460 | chrX | Pof1b | -2.806194302 | 0.126620337 |
| ENSMUSG0000000070 | chr17 | Kat2b | 0.308078463 | 0.126745355 |
| ENSMUSG0000003087 | chr7 | Cdr2 | 0.539626027 | 0.126745355 |
| ENSMUSG0000002737 | chr2 | Mal | -1.868251927 | 0.126760161 |
| ENSMUSG0000004832 | chr11 | Mfsd6l | -2.244751537 | 0.126760161 |
| ENSMUSG0000002194 | chr14 | Prkcd | -0.50080023 | 0.126886742 |
| ENSMUSG0000005133 | chr5 | 2900026A02Rik | -0.607801635 | 0.126948506 |
| ENSMUSG0000002240 | chr15 | Fam83f | -1.712758197 | 0.126948506 |
| ENSMUSG0000006216 | chrX | Ppef1 | -2.559870284 | 0.126948506 |
| ENSMUSG0000004233 | chr4 | Tnfrsf14 | -0.516599806 | 0.126948506 |
| ENSMUSG0000005157 | chrX | Tceal8 | 0.279010644 | 0.126948506 |
| ENSMUSG0000003320 | chr19 | Mamdc2 | 0.409320085 | 0.127013614 |
| ENSMUSG0000003836 | chr11 | Lasp1 | -0.458417067 | 0.127577789 |
| ENSMUSG0000003928 | chr3 | Fndc3b | 0.289892895 | 0.127577789 |
| ENSMUSG0000005014 | chrX | Ubqln2 | -0.203408263 | 0.127600527 |

| | | | | |
|---|---|---|---|---|
| ENSMUSG0000003335 | chr9 | Chst2 | -0.412099434 | 0.127812845 |
| ENSMUSG0000004082 | chr17 | Catsperd | -1.675899631 | 0.128270434 |
| ENSMUSG0000003425 | chr11 | Arhgap27 | -0.800970604 | 0.12879952 |
| ENSMUSG0000002877 | chr4 | Fabp3 | 0.465667582 | 0.12879952 |
| ENSMUSG0000003447 | chr8 | Rasd2 | -0.740258577 | 0.128842475 |
| ENSMUSG0000003210 | chr9 | Ddx25 | -0.529344111 | 0.128842475 |
| ENSMUSG0000002539 | chr10 | Atp5f1b | 0.347289957 | 0.129081885 |
| ENSMUSG0000002018 | chr11 | Ddc | 0.524305036 | 0.129081885 |
| ENSMUSG0000002981 | chr6 | Npy | 2.069211001 | 0.129081885 |
| ENSMUSG0000003858 | chr7 | Sct | -2.3305031 | 0.129164961 |
| ENSMUSG0000003556 | chr14 | Pcdh17 | -0.470899675 | 0.129164961 |
| ENSMUSG0000003520 | chr9 | Chrnb4 | -0.450771547 | 0.129164961 |
| ENSMUSG0000002635 | chr1 | Rgs1 | 0.473818164 | 0.129164961 |
| ENSMUSG0000003915 | chr2 | Eeig1 | -0.526314199 | 0.129207289 |
| ENSMUSG0000003493 | chr10 | Mrpl54 | 0.514414945 | 0.129410962 |
| ENSMUSG0000006165 | chr11 | Med9 | 0.26270498 | 0.129448013 |
| ENSMUSG0000007319 | chr12 | 5730507C01Rik | 0.875806085 | 0.129448013 |
| ENSMUSG0000003179 | chr8 | Mmp15 | -0.46648211 | 0.129640716 |
| ENSMUSG0000003346 | chr5 | Crlf2 | 0.445097107 | 0.130037573 |
| ENSMUSG0000000122 | chr12 | Slc26a3 | -3.367343863 | 0.130092282 |
| ENSMUSG0000010280 | chr6 | Mgam2-ps | -3.151284468 | 0.130376641 |
| ENSMUSG0000002315 | chr1 | Ivns1abp | -0.378058592 | 0.130501784 |
| ENSMUSG0000002636 | chr1 | Rgs2 | 0.285454266 | 0.130554039 |
| ENSMUSG0000003711 | chr2 | Ralgapa2 | -0.712602035 | 0.131318841 |
| ENSMUSG0000002438 | chr18 | Myo7b | -2.405605406 | 0.131361584 |
| ENSMUSG0000003869 | chr11 | Hoxb4 | -0.287062489 | 0.131361584 |
| ENSMUSG0000003223 | chr9 | Anxa2 | 0.541853173 | 0.131361584 |
| ENSMUSG0000002543 | chr3 | Usp33 | -0.243181324 | 0.131571443 |
| ENSMUSG0000004679 | chr3 | Gpr61 | -0.70905946 | 0.131707343 |
| ENSMUSG0000000765 | chr6 | Cav1 | 0.348204777 | 0.131981702 |
| ENSMUSG0000002981 | chr6 | Aoc1 | -2.827408074 | 0.132076191 |
| ENSMUSG0000002472 | chr19 | Trpm6 | -2.517596399 | 0.132076191 |
| ENSMUSG0000004182 | chr5 | Oasl1 | -0.860255885 | 0.132076191 |
| ENSMUSG0000007042 | chr7 | Il18bp | -0.468732369 | 0.132076191 |
| ENSMUSG0000002548 | chr7 | Urah | -2.830190309 | 0.132164621 |
| ENSMUSG0000002919 | chr5 | Cckar | -0.791176533 | 0.132385407 |
| ENSMUSG0000006141 | chr6 | Cracr2a | -1.099041756 | 0.132404951 |
| ENSMUSG0000002707 | chr2 | P2rx3 | -1.344908137 | 0.132404951 |
| ENSMUSG0000004778 | chr19 | Flrt1 | -0.421648258 | 0.132404951 |
| ENSMUSG0000002795 | chr3 | Chrnb2 | -0.433963044 | 0.132539214 |
| ENSMUSG0000003026 | chr6 | Mtmr14 | 0.291171454 | 0.132581949 |
| ENSMUSG0000002141 | chr13 | Eci3 | -2.536098418 | 0.132707985 |
| ENSMUSG0000002187 | chr14 | Slmap | 0.316508473 | 0.132707985 |
| ENSMUSG0000002761 | chr2 | Mmp24 | -0.525998436 | 0.133217842 |
| ENSMUSG0000002816 | chr3 | Ppp3ca | -0.229769323 | 0.133217842 |
| ENSMUSG0000000236 | chr17 | Snx9 | 0.235940966 | 0.133217842 |
| ENSMUSG0000003022 | chr6 | Ptpro | 0.536029641 | 0.133217842 |
| ENSMUSG0000003226 | chr9 | Tent5a | -0.419131364 | 0.133559376 |

| | | | | |
|---|---|---|---|---|
| ENSMUSG0000004189 | chr11 | Wipi1 | 0.338889261 | 0.133568426 |
| ENSMUSG0000004402 | chr19 | Slc35g1 | -0.508538378 | 0.134059077 |
| ENSMUSG0000004883 | chr11 | Vstm2a | -0.633087615 | 0.134085015 |
| ENSMUSG0000003875 | chr11 | Ttll6 | -0.668619624 | 0.134225148 |
| ENSMUSG0000000015 | chr16 | Itgb2l | -1.721954292 | 0.136816917 |
| ENSMUSG0000002634 | chr1 | Lypd1 | 0.720662258 | 0.136838554 |
| ENSMUSG0000004940 | chr6 | Prokr1 | -0.467714093 | 0.137093147 |
| ENSMUSG0000002634 | chr1 | Slc35f5 | -0.427132129 | 0.137107636 |
| ENSMUSG0000000122 | chr17 | Sema6b | -0.386453828 | 0.137107636 |
| ENSMUSG0000003837 | chr2 | Trp53inp2 | 0.280491572 | 0.137107636 |
| ENSMUSG0000003102 | chr7 | Akip1 | 0.449504054 | 0.137107636 |
| ENSMUSG0000002542 | chr18 | Atp5f1a | 0.295070117 | 0.137338645 |
| ENSMUSG0000006412 | chr8 | Prr36 | -0.601988348 | 0.137661462 |
| ENSMUSG0000004175 | chr1 | Plekha6 | -0.448353382 | 0.137985806 |
| ENSMUSG0000003230 | chr9 | Ulk3 | -0.381621466 | 0.137985806 |
| ENSMUSG0000004749 | chr8 | Dlgap2 | 0.497403525 | 0.137985806 |
| ENSMUSG0000009165 | chr15 | Apol11a | -4.387636926 | 0.138261841 |
| ENSMUSG0000007539 | chr15 | Hoxc4 | -0.386813724 | 0.138261841 |
| ENSMUSG0000001858 | chrX | Glra2 | -1.068870016 | 0.138693557 |
| ENSMUSG0000003089 | chr7 | Ilk | 0.5874568 | 0.13892317 |
| ENSMUSG0000003538 | chr11 | Ccl2 | 1.328377486 | 0.138971633 |
| ENSMUSG0000004336 | chr7 | Or51e2 | -0.843063264 | 0.139038008 |
| ENSMUSG0000004003 | chr10 | Stat2 | -0.536161464 | 0.140026453 |
| ENSMUSG0000002262 | chr15 | Kif21a | -0.408095249 | 0.140373899 |
| ENSMUSG0000002864 | chr4 | Rragc | 0.205137951 | 0.140985728 |
| ENSMUSG0000007078 | chr5 | Rbm47 | -1.384464393 | 0.141035979 |
| ENSMUSG0000002549 | chr7 | Sigirr | -0.871266869 | 0.141117228 |
| ENSMUSG0000005540 | chr7 | Map6 | -0.336669407 | 0.141117228 |
| ENSMUSG0000002137 | chr13 | Tpmt | 0.419970467 | 0.141117228 |
| ENSMUSG0000003829 | chr7 | Galnt18 | 0.509192369 | 0.141117228 |
| ENSMUSG0000002283 | chr16 | Mylk | 0.293609042 | 0.141372024 |
| ENSMUSG0000002618 | chr1 | Tmem169 | -0.492500826 | 0.141377043 |
| ENSMUSG0000000561 | chr16 | Pcyt1a | 0.22976993 | 0.141377043 |
| ENSMUSG0000004260 | chr4 | Stk40 | 0.460160298 | 0.141377043 |
| ENSMUSG0000002734 | chr2 | Slc23a2 | -0.362622455 | 0.141504624 |
| ENSMUSG0000002006 | chr10 | Slc5a8 | -3.116548126 | 0.141568147 |
| ENSMUSG0000002910 | chr5 | Hgfac | -2.551151912 | 0.142033576 |
| ENSMUSG0000005263 | chr6 | Sh2d6 | -1.660552793 | 0.142033576 |
| ENSMUSG0000003911 | chr10 | Adgrg6 | -0.304445338 | 0.142033576 |
| ENSMUSG0000002927 | chr5 | Gfi1 | -2.211363912 | 0.142323784 |
| ENSMUSG0000005805 | chr8 | Palld | 0.457148072 | 0.142323784 |
| ENSMUSG0000002751 | chr2 | Rbm38 | 0.399618223 | 0.142803044 |
| ENSMUSG0000002875 | chr4 | Cda | -1.572484997 | 0.142843551 |
| ENSMUSG0000006175 | chr16 | Kalrn | 0.329352987 | 0.143012745 |
| ENSMUSG0000006917 | chr13 | Nr2f1 | -0.721261132 | 0.143358604 |
| ENSMUSG0000005971 | chr4 | Rcan3 | -0.268798387 | 0.143358604 |
| ENSMUSG0000003140 | chrX | Tceal6 | -0.669378853 | 0.143774445 |
| ENSMUSG0000006722 | chr19 | Cyp2c66 | -2.889473315 | 0.144126139 |

| | | | | |
|---|---|---|---|---|
| ENSMUSG0000005291 | chr4 | Cyp2j6 | -0.356306774 | 0.144386771 |
| ENSMUSG0000006976 | chr11 | Tmem100 | 0.86610369 | 0.14505744 |
| ENSMUSG0000003715 | chr10 | Lrrc20 | 0.303736057 | 0.145290349 |
| ENSMUSG0000002119 | chr12 | Unc79 | -0.54623593 | 0.145420169 |
| ENSMUSG0000000493 | chr10 | Matk | -0.460488144 | 0.145420169 |
| ENSMUSG0000003687 | chr8 | Abcc12 | -1.987368814 | 0.145432283 |
| ENSMUSG0000004615 | chr12 | Tmem229b | -0.435009365 | 0.145900355 |
| ENSMUSG0000004250 | chr4 | Ago4 | -0.354558353 | 0.145965167 |
| ENSMUSG0000000210 | chr2 | Celf2 | 0.239330609 | 0.146385182 |
| ENSMUSG0000003659 | chr5 | Chst12 | 0.238107695 | 0.146520565 |
| ENSMUSG0000002571 | chr2 | Myo3a | -4.06738738 | 0.146727168 |
| ENSMUSG0000002186 | chr14 | Anxa11 | 0.241747906 | 0.146727168 |
| ENSMUSG0000002191 | chr14 | Ogdhl | -0.433322378 | 0.147391838 |
| ENSMUSG0000002976 | chr6 | Cald1 | 0.283673424 | 0.147391838 |
| ENSMUSG0000003000 | chr6 | Add2 | -0.494587939 | 0.149452077 |
| ENSMUSG0000005403 | chrX | Tceal5 | -0.607665062 | 0.149643326 |
| ENSMUSG0000002017 | chr10 | Yeats4 | 0.314642729 | 0.149643326 |
| ENSMUSG0000004060 | chr3 | Eps8l3 | -2.904210374 | 0.150410777 |
| ENSMUSG0000002542 | chr10 | Agap2 | -0.626497871 | 0.150410777 |
| ENSMUSG0000003002 | chr6 | Prickle2 | 0.410139929 | 0.150609705 |
| ENSMUSG0000003145 | chr8 | Rasa3 | 0.190568555 | 0.150845906 |
| ENSMUSG0000000163 | chr6 | Stk38l | 0.348185716 | 0.151512891 |
| ENSMUSG0000003498 | chr5 | Parm1 | -0.344053772 | 0.151792371 |
| ENSMUSG0000002879 | chr4 | Phc2 | 0.208438061 | 0.151792371 |
| ENSMUSG0000006227 | chr9 | Fbxw24 | -1.22980809 | 0.15181414 |
| ENSMUSG0000002642 | chr1 | Csrp1 | 0.515774405 | 0.152027862 |
| ENSMUSG0000000924 | chr7 | Ascl2 | -2.044112991 | 0.152091214 |
| ENSMUSG0000003986 | chr5 | Srrm3 | -0.401667827 | 0.152206015 |
| ENSMUSG0000004284 | chr13 | Serpinb6b | 0.814815799 | 0.152206015 |
| ENSMUSG0000004025 | chr6 | Ints13 | 0.295909764 | 0.152274316 |
| ENSMUSG0000003846 | chr1 | Olfml2b | 0.266588987 | 0.152351411 |
| ENSMUSG0000004646 | chr9 | Sh2d7 | -2.586120393 | 0.15276258 |
| ENSMUSG0000003932 | chr1 | Igfbp2 | 1.270224352 | 0.15276258 |
| ENSMUSG0000003426 | chr9 | Ints14 | 0.245155994 | 0.152860798 |
| ENSMUSG0000001100 | chr7 | Or51f23b | -1.756507804 | 0.153337696 |
| ENSMUSG0000001091 | chr2 | Apip | 0.421054774 | 0.153337696 |
| ENSMUSG0000002863 | chr4 | Ctps1 | 0.622023855 | 0.153556489 |
| ENSMUSG0000002063 | chr12 | Cmpk2 | -0.663840846 | 0.153871468 |
| ENSMUSG0000003826 | chr7 | Trpm4 | -0.530440805 | 0.153889689 |
| ENSMUSG0000007185 | chr18 | Ccdc112 | -0.442947676 | 0.154437616 |
| ENSMUSG0000006429 | chr6 | Cntn4 | -1.333879027 | 0.155112915 |
| ENSMUSG0000004181 | chr13 | Fam169a | -0.370093167 | 0.155254499 |
| ENSMUSG0000004637 | chr7 | Asphd1 | -0.926367107 | 0.155390696 |
| ENSMUSG0000003254 | chr9 | Slco2a1 | -0.805415417 | 0.155495941 |
| ENSMUSG0000004258 | chr5 | Cux2 | -0.464336183 | 0.155700939 |
| ENSMUSG0000007922 | chr9 | Ccr5 | -0.929837495 | 0.156248451 |
| ENSMUSG0000004169 | chr5 | Cox6a1 | 0.443768456 | 0.156248451 |
| ENSMUSG0000002334 | chr16 | Mx2 | -0.871016156 | 0.15641159 |

| | | | | |
|---|---|---|---|---|
| ENSMUSG0000004135 | chr4 | Rap1gap | -0.424311423 | 0.156748842 |
| ENSMUSG0000006372 | chr15 | Tnfrsf11b | 0.368894131 | 0.156858915 |
| ENSMUSG0000002547 | chr7 | Inpp5a | 0.330919013 | 0.156912143 |
| ENSMUSG0000002559 | chrX | Dach2 | -1.572981794 | 0.157096936 |
| ENSMUSG0000004994 | chr8 | BC030500 | -1.088626892 | 0.15754154 |
| ENSMUSG0000002253 | chr16 | Atp13a3 | 0.309324658 | 0.157603222 |
| ENSMUSG0000003633 | chr3 | Igsf10 | -0.448952708 | 0.157629607 |
| ENSMUSG0000007512 | chr16 | Cd80 | 0.866532041 | 0.157833022 |
| ENSMUSG0000005617 | chr4 | Col8a2 | 0.557079412 | 0.157833022 |
| ENSMUSG0000002324 | chr9 | Parp3 | 0.280111386 | 0.157977555 |
| ENSMUSG0000007103 | chr17 | Camkmt | -0.412116081 | 0.158101144 |
| ENSMUSG0000002401 | chr17 | Pi16 | 0.766440916 | 0.158101144 |
| ENSMUSG0000003078 | chr7 | Tgfb1i1 | 0.23989178 | 0.158947985 |
| ENSMUSG0000005842 | chr7 | Syt17 | -0.388509953 | 0.159311642 |
| ENSMUSG0000005158 | chr6 | Mical3 | 0.414991475 | 0.159311642 |
| ENSMUSG0000002550 | chr7 | Eps8l2 | -1.058137614 | 0.159568162 |
| ENSMUSG0000003259 | chr9 | Uba7 | -0.654766756 | 0.15993566 |
| ENSMUSG0000008978 | chr10 | Rdh1 | -2.847248762 | 0.16007747 |
| ENSMUSG0000001776 | chr8 | Slc12a4 | 0.231880168 | 0.160099316 |
| ENSMUSG0000003414 | chr12 | Tmem63c | -0.493637747 | 0.160752314 |
| ENSMUSG0000002380 | chr17 | Synj2 | 0.289643528 | 0.160752314 |
| ENSMUSG0000003766 | chr3 | Gpr160 | -0.905479185 | 0.160807268 |
| ENSMUSG0000006166 | chr11 | Gdpd1 | -0.450871657 | 0.160807268 |
| ENSMUSG0000000245 | chr1 | Rgs20 | -0.978505854 | 0.161226239 |
| ENSMUSG0000000086 | chr11 | Il4 | -0.647329345 | 0.161226239 |
| ENSMUSG0000003576 | chr16 | Fbxo45 | -0.332370617 | 0.161226239 |
| ENSMUSG0000006056 | chr1 | Fam78b | -0.488355634 | 0.161563426 |
| ENSMUSG0000002627 | chr1 | Gpr35 | -1.104150057 | 0.162716536 |
| ENSMUSG0000003212 | chr9 | Slc37a2 | -2.750850538 | 0.163124388 |
| ENSMUSG0000002008 | chr10 | Hkdc1 | -1.987381071 | 0.163310364 |
| ENSMUSG0000002072 | chr11 | Cacng4 | -0.410785474 | 0.163396875 |
| ENSMUSG0000001781 | chr2 | Jph2 | 0.297286159 | 0.163524927 |
| ENSMUSG0000003550 | chr10 | Reep6 | -0.718077029 | 0.164058766 |
| ENSMUSG0000000809 | chr5 | Fgfrl1 | 0.31916455 | 0.16418894 |
| ENSMUSG0000005412 | chr17 | H2-T3 | -2.73409229 | 0.164448993 |
| ENSMUSG0000003250 | chr9 | Stac | 0.607361416 | 0.165295176 |
| ENSMUSG0000002555 | chr14 | Farp1 | 0.27843552 | 0.165394764 |
| ENSMUSG0000006153 | chr5 | C1qtnf7 | 0.336691792 | 0.165394764 |
| ENSMUSG0000002528 | chrX | Acot9 | 0.469585526 | 0.165394764 |
| ENSMUSG0000002461 | chr18 | Cd74 | -0.565062947 | 0.165578656 |
| ENSMUSG0000002082 | chr11 | Kif1c | 0.274097231 | 0.165617527 |
| ENSMUSG0000001597 | chr14 | Chdh | -0.642282842 | 0.165652385 |
| ENSMUSG0000003202 | chr9 | Nxpe2 | -1.375091391 | 0.166201937 |
| ENSMUSG0000002795 | chr3 | Adar | -0.272992157 | 0.166201937 |
| ENSMUSG0000000062 | chr6 | Sema4f | -0.506259264 | 0.166899708 |
| ENSMUSG0000001778 | chr11 | Pitpna | 0.301338075 | 0.166899708 |
| ENSMUSG0000006180 | chr18 | Ttr | -1.531264179 | 0.16704229 |
| ENSMUSG0000002191 | chr14 | Spcs1 | 0.366501847 | 0.167711084 |

| | | | | |
|---|---|---|---|---|
| ENSMUSG0000004931 | chr9 | Sorl1 | -0.29540623 | 0.168089951 |
| ENSMUSG0000002261 | chr15 | Lmf2 | 0.171657354 | 0.168089951 |
| ENSMUSG0000002963 | chr6 | Ndufa4 | 0.423545565 | 0.16813917 |
| ENSMUSG0000000178 | chr10 | Pwp1 | 0.273422361 | 0.168404717 |
| ENSMUSG0000002630 | chr1 | Klhl30 | 0.365255141 | 0.168448606 |
| ENSMUSG0000002771 | chr3 | Exosc9 | 0.249295382 | 0.168464128 |
| ENSMUSG0000002417 | chr17 | St6gal2 | -0.806969765 | 0.16890498 |
| ENSMUSG0000000056 | chr11 | Sox9 | -1.004027039 | 0.169478106 |
| ENSMUSG0000002095 | chr12 | Scfd1 | 0.197470084 | 0.169478106 |
| ENSMUSG0000002813 | chr3 | Ptbp2 | 0.264588499 | 0.170376215 |
| ENSMUSG0000002653 | chr1 | Mptx1 | -3.532429789 | 0.170629498 |
| ENSMUSG0000003839 | chr6 | Gpr162 | -0.571269263 | 0.170629498 |
| ENSMUSG0000003120 | chrX | Msn | 0.236818925 | 0.170632419 |
| ENSMUSG0000003718 | chr15 | Krt80 | 0.495768197 | 0.171191272 |
| ENSMUSG0000004445 | chr12 | Rin3 | 0.54357353 | 0.171191272 |
| ENSMUSG0000006729 | chr19 | Ifit1bl2 | -0.684590634 | 0.171253236 |
| ENSMUSG0000002474 | chr19 | Ddb1 | 0.241053063 | 0.171253236 |
| ENSMUSG0000005540 | chr4 | Fbxo6 | 0.354711794 | 0.171253236 |
| ENSMUSG0000002179 | chr14 | Bmpr1a | 0.213250801 | 0.171544616 |
| ENSMUSG0000004875 | chr9 | Rpl29 | 0.370576543 | 0.172084579 |
| ENSMUSG0000008765 | chr10 | 1500009L16Rik | -0.552224372 | 0.172158126 |
| ENSMUSG0000008987 | chr11 | Tmem102 | -0.651259065 | 0.172158126 |
| ENSMUSG0000003966 | chr8 | Dusp26 | -0.365414077 | 0.172158126 |
| ENSMUSG0000003272 | chr2 | Abtb2 | 0.393061132 | 0.172158126 |
| ENSMUSG0000007434 | chr9 | Tnfaip8l3 | 0.429000537 | 0.172158126 |
| ENSMUSG0000004049 | chr2 | Chrm4 | -0.446069201 | 0.172500137 |
| ENSMUSG0000002159 | chr13 | Mctp1 | -0.456233316 | 0.172500137 |
| ENSMUSG0000002574 | chrX | Prps2 | -0.304630529 | 0.172500137 |
| ENSMUSG0000003087 | chr7 | Gga2 | -0.270446511 | 0.172500137 |
| ENSMUSG0000002647 | chr1 | Lamc1 | 0.426392542 | 0.172500137 |
| ENSMUSG0000004462 | chr11 | Cnrip1 | -0.35233918 | 0.172693294 |
| ENSMUSG0000002265 | chr16 | Gcsam | 0.81296513 | 0.173531743 |
| ENSMUSG0000001460 | chr1 | Kif1a | -0.459151813 | 0.17377013 |
| ENSMUSG0000004773 | chr7 | Fcgbp | -2.6180336 | 0.173859283 |
| ENSMUSG0000000843 | chr12 | Adam21 | -1.954793601 | 0.174935539 |
| ENSMUSG0000002638 | chr1 | Epb41l5 | -0.358282397 | 0.174935539 |
| ENSMUSG0000005155 | chr4 | Pusl1 | 0.45756459 | 0.174935539 |
| ENSMUSG0000002853 | chr4 | 2610528J11Rik | -2.027884011 | 0.175276012 |
| ENSMUSG0000005434 | chr7 | Kcnn4 | -0.814523076 | 0.175276012 |
| ENSMUSG0000004787 | chr15 | A4galt | 0.223684205 | 0.175502301 |
| ENSMUSG0000002221 | chr14 | Cideb | -2.046693806 | 0.175691241 |
| ENSMUSG0000002516 | chr11 | Sectm1a | -1.972640519 | 0.175697101 |
| ENSMUSG0000000068 | chr4 | Cd52 | -0.66502056 | 0.176383256 |
| ENSMUSG0000000210 | chr2 | Slc39a13 | 0.229088833 | 0.176447117 |
| ENSMUSG0000002906 | chr4 | Mrpl20 | 0.488740409 | 0.176447117 |
| ENSMUSG0000005007 | chrX | Bex1 | -0.396192957 | 0.177084189 |
| ENSMUSG0000001557 | chr17 | Atp6v0e | 0.372162532 | 0.177446892 |
| ENSMUSG0000005652 | chr4 | Ptafr | 0.41307754 | 0.178038855 |

| | | | | |
|---|---|---|---|---|
| ENSMUSG0000003235 | chr9 | Tinag | -2.436122652 | 0.178421411 |
| ENSMUSG0000002591 | chr1 | Cops5 | 0.274344083 | 0.178421411 |
| ENSMUSG0000007525 | chr16 | Heg1 | 0.355712893 | 0.178421411 |
| ENSMUSG0000002500 | chr19 | Sorbs1 | 0.409648006 | 0.178421411 |
| ENSMUSG0000003254 | chr9 | Ryk | 0.265486176 | 0.179507257 |
| ENSMUSG0000002148 | chr13 | Prelid1 | 0.312618972 | 0.179553901 |
| ENSMUSG0000006879 | chr6 | Col28a1 | -0.656405934 | 0.180004518 |
| ENSMUSG0000003860 | chr2 | Samd10 | -0.446294822 | 0.180393773 |
| ENSMUSG0000003491 | chr13 | Cdhr2 | -2.768888578 | 0.18063776 |
| ENSMUSG0000007411 | chr7 | Saa1 | -2.734483626 | 0.18063776 |
| ENSMUSG0000006073 | chr10 | Ipmk | -0.695646137 | 0.18063776 |
| ENSMUSG0000004279 | chr7 | Aqp11 | -0.880818924 | 0.18063776 |
| ENSMUSG0000003390 | chr3 | Map9 | -0.464467265 | 0.18063776 |
| ENSMUSG0000002884 | chr4 | Map7d1 | 0.23314552 | 0.18063776 |
| ENSMUSG0000002526 | chrX | Maged2 | 0.243975785 | 0.18063776 |
| ENSMUSG0000002483 | chr19 | Pola2 | 0.254270417 | 0.18063776 |
| ENSMUSG0000004143 | chr8 | Utp4 | 0.328831669 | 0.18063776 |
| ENSMUSG0000004699 | chr9 | Spsb4 | 0.698843223 | 0.18063776 |
| ENSMUSG0000000761 | chr4 | Tgfbr1 | 0.199819605 | 0.18068784 |
| ENSMUSG0000002407 | chr17 | Eif2ak2 | -0.481783338 | 0.180740018 |
| ENSMUSG0000003110 | chrX | Sash3 | -1.031487354 | 0.181310343 |
| ENSMUSG0000003131 | chrX | Itgb1bp2 | 0.646345854 | 0.181310343 |
| ENSMUSG0000002871 | chr4 | Cmpk1 | -0.309122522 | 0.18189325 |
| ENSMUSG0000000170 | chr18 | Gramd2b | -0.42563282 | 0.182443732 |
| ENSMUSG0000002660 | chr1 | Kctd3 | 0.181254931 | 0.182900124 |
| ENSMUSG0000007823 | chr4 | Fam43b | -1.822416213 | 0.183003241 |
| ENSMUSG0000003359 | chr17 | Caskin1 | -0.495025953 | 0.183054844 |
| ENSMUSG0000002601 | chr1 | Raph1 | -0.44227729 | 0.183120785 |
| ENSMUSG0000003164 | chr8 | Sh3rf1 | 0.43512796 | 0.183120785 |
| ENSMUSG0000003480 | chr8 | Colgalt1 | 0.200897375 | 0.183495969 |
| ENSMUSG0000002017 | chr10 | Rab36 | -0.467395089 | 0.18365477 |
| ENSMUSG0000000830 | chr4 | Tle1 | -0.267988911 | 0.18365477 |
| ENSMUSG0000003132 | chrX | Flna | 0.279717982 | 0.18365477 |
| ENSMUSG0000009027 | chr1 | Mndal | -0.57869687 | 0.184117674 |
| ENSMUSG0000002986 | chr6 | Zyx | 0.311838226 | 0.184311728 |
| ENSMUSG0000002409 | chr17 | Twsg1 | 0.221854248 | 0.184510413 |
| ENSMUSG0000003202 | chr9 | Clmp | 0.376034541 | 0.184566919 |
| ENSMUSG0000002596 | chr1 | Nrp2 | 0.370048588 | 0.184566919 |
| ENSMUSG0000000537 | chr5 | Mlxipl | -1.509128435 | 0.184581337 |
| ENSMUSG0000002855 | chr4 | Angptl3 | 1.083812295 | 0.185233367 |
| ENSMUSG0000007359 | chr18 | Smim33 | -1.888226795 | 0.185613843 |
| ENSMUSG0000004068 | chr2 | Madd | -0.388095937 | 0.185613843 |
| ENSMUSG0000001060 | chr15 | Apol7a | -0.712645495 | 0.185771101 |
| ENSMUSG0000004341 | chr11 | Rnf227 | -0.52467502 | 0.186517086 |
| ENSMUSG0000002549 | chr7 | Cdhr5 | -2.8493648 | 0.18680556 |
| ENSMUSG0000004723 | chrX | Cldn2 | -2.683743951 | 0.18680556 |
| ENSMUSG0000003952 | chr18 | Atp8b1 | -0.74621205 | 0.18680556 |
| ENSMUSG0000004644 | chrX | Nexmif | -0.698250195 | 0.18680556 |

| | | | | |
|---|---|---|---|---|
| ENSMUSG0000002755 | chr2 | Zfp64 | 0.259367749 | 0.18680556 |
| ENSMUSG0000002408 | chr17 | Qpct | 0.33345116 | 0.18680556 |
| ENSMUSG0000002220 | chr14 | Jph4 | -0.420901549 | 0.186959087 |
| ENSMUSG0000000503 | chr3 | Prkacb | 0.229978518 | 0.186959087 |
| ENSMUSG0000005865 | chr15 | Samd12 | -0.460587424 | 0.18704071 |
| ENSMUSG0000002551 | chr7 | Muc2 | -2.651392865 | 0.187088401 |
| ENSMUSG0000002487 | chr19 | Yif1a | 0.358700539 | 0.187398729 |
| ENSMUSG0000003069 | chr7 | Aldoa | 0.306900319 | 0.187569893 |
| ENSMUSG0000002825 | chr4 | Tstd3 | 0.429024846 | 0.188095423 |
| ENSMUSG0000003306 | chr2 | Entpd6 | -0.475086016 | 0.188212873 |
| ENSMUSG0000004343 | chr11 | Epop | -1.582375226 | 0.188601585 |
| ENSMUSG0000002269 | chr16 | Sidt1 | -0.708174683 | 0.189171396 |
| ENSMUSG0000004032 | chr3 | Il7 | -0.343022333 | 0.18937767 |
| ENSMUSG0000002575 | chr3 | Hspa4l | 0.300348708 | 0.189508824 |
| ENSMUSG0000004561 | chr6 | Chrm2 | 0.440151027 | 0.189508824 |
| ENSMUSG0000005259 | chr19 | A1cf | -2.801983686 | 0.191019949 |
| ENSMUSG0000002901 | chr5 | Orc5 | 0.274164315 | 0.191019949 |
| ENSMUSG0000002697 | chr2 | Itgb6 | -1.340781923 | 0.191448739 |
| ENSMUSG0000002496 | chr19 | Rcor2 | -0.492274354 | 0.192033982 |
| ENSMUSG0000003223 | chr9 | Cgnl1 | 0.349502814 | 0.192183342 |
| ENSMUSG0000002759 | chr2 | Raly | 0.412041807 | 0.19337191 |
| ENSMUSG0000004172 | chr9 | Coro2b | -0.407639781 | 0.193754114 |
| ENSMUSG0000006824 | chr14 | Phf11d | -0.709689571 | 0.193895133 |
| ENSMUSG0000003190 | chr8 | Smpd3 | -0.463830414 | 0.193895133 |
| ENSMUSG0000002496 | chr19 | Plcb3 | -0.265639528 | 0.193895133 |
| ENSMUSG0000001742 | chr11 | Zfp207 | 0.243969569 | 0.194137751 |
| ENSMUSG0000006907 | chr3 | Slc7a14 | -0.4037585 | 0.194889563 |
| ENSMUSG0000002804 | chr3 | Thbs3 | 0.324039065 | 0.195003769 |
| ENSMUSG0000002173 | chr14 | Psmd6 | 0.339357874 | 0.195003769 |
| ENSMUSG0000004195 | chr16 | Pkp2 | 0.362805291 | 0.195003769 |
| ENSMUSG0000002393 | chr17 | Mrpl14 | 0.576994998 | 0.195026488 |
| ENSMUSG0000003277 | chr7 | Mctp2 | -0.950917833 | 0.19509065 |
| ENSMUSG0000005600 | chr5 | Elapor2 | -0.34518787 | 0.195253164 |
| ENSMUSG0000007479 | chr13 | Arrdc3 | 0.287468125 | 0.196352806 |
| ENSMUSG0000002983 | chr6 | Ptn | 0.233018942 | 0.19696852 |
| ENSMUSG0000003482 | chr7 | Nrip3 | -0.473404034 | 0.197036123 |
| ENSMUSG0000002458 | chr18 | Grpel2 | -0.297916702 | 0.197036123 |
| ENSMUSG0000003217 | chr9 | Bmp5 | 0.652587264 | 0.197036123 |
| ENSMUSG0000002201 | chr14 | Tsc22d1 | 0.533619052 | 0.197036123 |
| ENSMUSG0000003619 | chr19 | Rorb | 1.194458882 | 0.197343077 |
| ENSMUSG0000004262 | chr15 | Maff | 0.458721333 | 0.198113404 |
| ENSMUSG0000000504 | chr4 | Chd5 | -0.42907381 | 0.19844126 |
| ENSMUSG0000003698 | chrX | Zdhhc9 | -0.292482277 | 0.19844126 |
| ENSMUSG0000002883 | chr4 | AU040320 | -0.322416079 | 0.198510648 |
| ENSMUSG0000006887 | chr3 | Selenbp2 | -1.522733361 | 0.198688327 |
| ENSMUSG0000002854 | chr4 | B4galt2 | 0.288047221 | 0.198900843 |
| ENSMUSG0000001915 | chr7 | Tmem160 | 0.674343825 | 0.198900843 |
| ENSMUSG0000003155 | chr8 | Ido1 | -3.26700758 | 0.198984121 |
| ENSMUSG0000005728 | chr11 | St6galnac2 | -0.946823201 | 0.198984121 |
| ENSMUSG0000003080 | chr7 | Prss8 | -1.323646468 | 0.198984121 |

| | | | | |
|---|---|---|---|---|
| ENSMUSG0000003070 | chr7 | Coro1a | -0.498008338 | 0.198984121 |
| ENSMUSG0000002804 | chr3 | Zbtb7b | -0.411051211 | 0.198984121 |
| ENSMUSG0000003128 | chrX | Pak3 | -0.358113231 | 0.198984121 |
| ENSMUSG0000004126 | chr3 | Rusc1 | -0.321869368 | 0.198984121 |
| ENSMUSG0000002769 | chr3 | Nceh1 | -0.320402983 | 0.198984121 |
| ENSMUSG0000005970 | chrX | Akap17b | -0.322509595 | 0.198984121 |
| ENSMUSG0000004102 | chrX | Map7d2 | -0.286541387 | 0.198984121 |
| ENSMUSG0000002650 | chr1 | Desi2 | 0.198330176 | 0.198984121 |
| ENSMUSG0000000290 | chr12 | Lamb1 | 0.234916661 | 0.198984121 |
| ENSMUSG0000003771 | chr14 | Fermt2 | 0.265840942 | 0.198984121 |
| ENSMUSG0000004667 | chr4 | Mtfr1l | 0.279230768 | 0.198984121 |
| ENSMUSG0000002784 | chr3 | Dclre1b | 0.346452369 | 0.198984121 |
| ENSMUSG0000002818 | chr3 | Adgrl2 | 0.334692193 | 0.198984121 |
| ENSMUSG0000006393 | chr7 | Pepd | 0.354400498 | 0.198984121 |
| ENSMUSG0000003529 | chr14 | Sgcg | 0.915892622 | 0.198984121 |
| ENSMUSG0000003378 | chr6 | Dysf | 0.448563233 | 0.198984121 |
| ENSMUSG0000003799 | chr11 | Rara | -0.293645012 | 0.199953219 |
| ENSMUSG0000002815 | chr3 | Tspan5 | 0.285411602 | 0.199953219 |
| ENSMUSG0000003613 | chr12 | Fam110c | -1.078593603 | 0.200154675 |
| ENSMUSG0000002565 | chr9 | Uqcrc1 | 0.260955248 | 0.20035484 |
| ENSMUSG0000004821 | chr11 | Nags | -3.16469984 | 0.200694725 |
| ENSMUSG0000003893 | chr7 | Rccd1 | 0.31121928 | 0.200694725 |
| ENSMUSG0000003890 | chr2 | Rpl12 | 0.51069454 | 0.200714612 |
| ENSMUSG0000003341 | chr9 | Snap91 | -0.435638137 | 0.200912157 |
| ENSMUSG0000002593 | chr1 | Tmem14a | 0.384715328 | 0.200912157 |
| ENSMUSG0000003251 | chr9 | Scn5a | -0.530329116 | 0.201448828 |
| ENSMUSG0000003565 | chr12 | Lrfn5 | -0.538085904 | 0.201448828 |
| ENSMUSG0000002845 | chr4 | Atp8b5 | -0.903619509 | 0.201448828 |
| ENSMUSG0000002978 | chr6 | Avl9 | -0.308363876 | 0.201448828 |
| ENSMUSG0000002780 | chr3 | Wwtr1 | 0.278161681 | 0.201448828 |
| ENSMUSG0000004153 | chr17 | H2-Ob | -1.729222537 | 0.201480573 |
| ENSMUSG0000007391 | chr4 | Mob3b | -0.415326839 | 0.201762932 |
| ENSMUSG0000000404 | chr11 | Stat3 | 0.24911334 | 0.201762932 |
| ENSMUSG0000003547 | chr17 | Galm | -0.682725847 | 0.201975756 |
| ENSMUSG0000003107 | chrX | Zfp300 | -0.552176576 | 0.201975756 |
| ENSMUSG0000004892 | chr7 | Fkrp | 0.194411093 | 0.201975756 |
| ENSMUSG0000003391 | chr16 | Parl | 0.221777109 | 0.201975756 |
| ENSMUSG0000003368 | chr7 | Ucp2 | 0.445744275 | 0.202386744 |
| ENSMUSG0000002533 | chr5 | Sbds | 0.316278064 | 0.202419868 |
| ENSMUSG0000002864 | chr4 | Svbp | 0.500223435 | 0.202419868 |
| ENSMUSG0000000463 | chr6 | Sgce | 0.31685757 | 0.202844492 |
| ENSMUSG0000004776 | chr9 | Lrrc49 | 0.355489632 | 0.203605205 |
| ENSMUSG0000003442 | chr16 | Parp14 | -0.337739045 | 0.203814134 |
| ENSMUSG0000009455 | chr15 | Cyp2d34 | -3.045016381 | 0.204159597 |
| ENSMUSG0000002061 | chr11 | Amz2 | 0.25669906 | 0.204279847 |
| ENSMUSG0000003179 | chr8 | Tmem38a | 0.311342651 | 0.204279847 |
| ENSMUSG0000004231 | chr3 | S100a13 | 0.381554124 | 0.204279847 |
| ENSMUSG0000002428 | chr18 | Ccny | 0.188175728 | 0.205732348 |
| ENSMUSG0000004500 | chr6 | Prrt3 | -0.525351753 | 0.207228502 |
| ENSMUSG0000005556 | chr1 | Unc80 | -0.449982212 | 0.20797253 |

| | | | | |
|---|---|---|---|---|
| ENSMUSG0000002471 | chr19 | Rfk | -0.368892825 | 0.20797253 |
| ENSMUSG0000002255 | chr15 | Cyc1 | 0.284426156 | 0.20797253 |
| ENSMUSG0000004051 | chr7 | Pvr | 0.377738365 | 0.20797253 |
| ENSMUSG0000000672 | chr10 | Cdk4 | 0.208538261 | 0.208385375 |
| ENSMUSG0000000924 | chr7 | Trpm5 | -1.808427213 | 0.208902838 |
| ENSMUSG0000004883 | chr12 | Slc39a9 | -0.216307708 | 0.209009571 |
| ENSMUSG0000003854 | chr7 | Tmc3 | -0.580365347 | 0.209384907 |
| ENSMUSG0000002980 | chr6 | Herc3 | -0.372051074 | 0.209384907 |
| ENSMUSG0000006229 | chr9 | Trank1 | -0.86619288 | 0.209384907 |
| ENSMUSG0000005467 | chr4 | Srsf12 | -0.418334167 | 0.209384907 |
| ENSMUSG0000004534 | chr5 | Nyap1 | -0.348436782 | 0.209384907 |
| ENSMUSG0000000260 | chr7 | Axl | 0.231121355 | 0.209384907 |
| ENSMUSG0000002940 | chr5 | Arl6ip4 | 0.278599969 | 0.209384907 |
| ENSMUSG0000003605 | chr4 | Dnajb5 | 0.32543124 | 0.209855922 |
| ENSMUSG0000002581 | chr8 | Nrp1 | 0.318353459 | 0.209855922 |
| ENSMUSG0000002578 | chr2 | Atp5f1c | 0.386249885 | 0.209855922 |
| ENSMUSG0000004516 | chr6 | Bola3 | 0.413675617 | 0.209855922 |
| ENSMUSG0000003703 | chr1 | Inhbb | 0.817120805 | 0.209855922 |
| ENSMUSG0000002548 | chr7 | Syce1 | 2.178553031 | 0.209855922 |
| ENSMUSG0000003171 | chr8 | Il15 | -0.533951673 | 0.21026795 |
| ENSMUSG0000006989 | chr11 | 9930111J21Rik1 | -0.832285616 | 0.21097668 |
| ENSMUSG0000004632 | chr19 | Ermp1 | -0.351777815 | 0.211027324 |
| ENSMUSG0000001977 | chr10 | Mtrf1l | 0.273852348 | 0.211027324 |
| ENSMUSG0000002280 | chr16 | Popdc2 | 0.472581561 | 0.211312231 |
| ENSMUSG0000002083 | chr11 | Coro6 | 0.711612245 | 0.211312231 |
| ENSMUSG0000002737 | chr2 | Mall | -0.833098316 | 0.211379139 |
| ENSMUSG0000003073 | chr7 | Myh14 | -1.027527955 | 0.211386205 |
| ENSMUSG0000005086 | chr7 | Clrn3 | -2.952756293 | 0.212293527 |
| ENSMUSG0000007492 | chr19 | Pabir1 | 0.296934898 | 0.212293527 |
| ENSMUSG0000002697 | chr2 | Arrdc1 | -0.471964415 | 0.212472291 |
| ENSMUSG0000002859 | chr4 | Gpx7 | 0.37417162 | 0.213041735 |
| ENSMUSG0000003511 | chr11 | Wnk4 | -0.611871159 | 0.213065538 |
| ENSMUSG0000002266 | chr16 | Cd200 | -0.280369115 | 0.213065538 |
| ENSMUSG0000003232 | chr9 | Rcn2 | 0.283140916 | 0.213065538 |
| ENSMUSG0000003584 | chrX | Ids | -0.289977812 | 0.213131274 |
| ENSMUSG0000002867 | chr4 | Gale | -0.803108063 | 0.213204768 |
| ENSMUSG0000003424 | chr6 | Hdac11 | -0.27072155 | 0.213262656 |
| ENSMUSG0000002676 | chr2 | Kif5c | -0.362875987 | 0.213892743 |
| ENSMUSG0000004148 | chr19 | Stx3 | -0.331376598 | 0.213892743 |
| ENSMUSG0000002182 | chr14 | Camk2g | 0.241296906 | 0.213892743 |
| ENSMUSG0000000384 | chr8 | Nob1 | 0.278481583 | 0.213892743 |
| ENSMUSG0000005801 | chr5 | Septin11 | 0.305437618 | 0.213892743 |
| ENSMUSG0000000584 | chr16 | Rsl1d1 | 0.273434798 | 0.213924313 |
| ENSMUSG0000003243 | chr9 | Cmtm7 | 0.417130504 | 0.213924313 |
| ENSMUSG0000003248 | chr1 | Ptgs2 | 0.45395724 | 0.213957351 |
| ENSMUSG0000002914 | chr5 | Snx17 | 0.308062836 | 0.21411813 |
| ENSMUSG0000002985 | chr6 | Tcaf2 | -0.411905178 | 0.214181378 |
| ENSMUSG0000002490 | chr19 | Efemp2 | 0.34215814 | 0.214181378 |
| ENSMUSG0000003816 | chr6 | Plekhg6 | -1.092159727 | 0.214218309 |
| ENSMUSG0000003796 | chr13 | Ninj1 | 0.248523101 | 0.214748279 |

| | | | | |
|---|---|---|---|---|
| ENSMUSG0000003629 | chr16 | Gramd1c | -0.624297186 | 0.21554638 |
| ENSMUSG0000006151 | chr1 | Cox5b | 0.420487085 | 0.21554638 |
| ENSMUSG0000002581 | chr7 | Homer2 | 0.442659008 | 0.21554638 |
| ENSMUSG0000002068 | chr11 | Itgb3 | 0.372569949 | 0.216223036 |
| ENSMUSG0000004136 | chr9 | Myzap | 0.284914097 | 0.216469215 |
| ENSMUSG0000007816 | chr3 | Erich3 | -0.455879772 | 0.216484163 |
| ENSMUSG0000002986 | chr6 | Gstk1 | -0.461908587 | 0.216484163 |
| ENSMUSG0000002790 | chr3 | S100a11 | 0.218185025 | 0.216484163 |
| ENSMUSG0000001829 | chr11 | Pfn1 | 0.35369247 | 0.216484163 |
| ENSMUSG0000003129 | chrX | Rs1 | 1.79841331 | 0.216484163 |
| ENSMUSG0000002625 | chr1 | Ngef | -0.762845685 | 0.216781857 |
| ENSMUSG0000002516 | chr11 | Slc16a3 | 0.500779803 | 0.217066099 |
| ENSMUSG0000002830 | chr4 | Aldob | -2.101719109 | 0.217074791 |
| ENSMUSG0000003270 | chr19 | Kank1 | 0.223705185 | 0.21713929 |
| ENSMUSG0000003713 | chr8 | Prmt9 | 0.303474065 | 0.217200269 |
| ENSMUSG0000006051 | chr5 | 0610040J01Rik | -0.848125224 | 0.217334495 |
| ENSMUSG0000004047 | chr5 | Cfap69 | -0.370856733 | 0.217572526 |
| ENSMUSG0000003012 | chr6 | Ptms | 0.28714205 | 0.217907712 |
| ENSMUSG0000006060 | chr11 | Eno3 | 0.666170975 | 0.217907712 |
| ENSMUSG0000004144 | chr9 | Arhgap32 | -0.292894642 | 0.217928213 |
| ENSMUSG0000002558 | chr7 | Cpeb1 | -0.394349175 | 0.217971308 |
| ENSMUSG0000002220 | chr15 | Golph3 | 0.220378972 | 0.217971308 |
| ENSMUSG0000002282 | chr16 | Stxbp5l | -0.570805912 | 0.217997227 |
| ENSMUSG0000001251 | chr8 | Mlkl | -0.756829442 | 0.218682419 |
| ENSMUSG0000003444 | chr11 | Dhrs11 | -0.813108861 | 0.218975083 |
| ENSMUSG0000003151 | chr8 | Asb5 | 0.397298049 | 0.218975083 |
| ENSMUSG0000000347 | chr6 | Inmt | -1.299577361 | 0.219035617 |
| ENSMUSG0000003519 | chr11 | Tubg1 | 0.273232379 | 0.219414681 |
| ENSMUSG0000003229 | chr9 | Celf6 | -0.40627391 | 0.219454704 |
| ENSMUSG0000003034 | chr6 | Tnfrsf1a | 0.231885743 | 0.219454704 |
| ENSMUSG0000006343 | chr5 | Wscd2 | 0.45468159 | 0.219454704 |
| ENSMUSG0000001511 | chr6 | Slc25a13 | -0.853548175 | 0.219628494 |
| ENSMUSG0000004074 | chrX | Siah1b | 0.462879459 | 0.22020816 |
| ENSMUSG0000003741 | chr9 | Endod1 | -0.436943629 | 0.220278622 |
| ENSMUSG0000000404 | chr11 | Cavin1 | 0.40759479 | 0.220573183 |
| ENSMUSG0000003793 | chr13 | Bicd2 | 0.28155294 | 0.220777723 |
| ENSMUSG0000006801 | chr14 | Lrch1 | 0.320056203 | 0.221478068 |
| ENSMUSG0000003282 | chr3 | Ank2 | -0.421189754 | 0.221505845 |
| ENSMUSG0000000167 | chr8 | Tat | -1.522779021 | 0.221519628 |
| ENSMUSG0000003093 | chr7 | Acsm3 | -1.372723655 | 0.222024768 |
| ENSMUSG0000005308 | chr19 | Zfta | -0.23721747 | 0.222024768 |
| ENSMUSG0000003926 | chr2 | Npepl1 | 0.235401003 | 0.222024768 |
| ENSMUSG0000004401 | chr5 | Adgrd1 | 0.332542622 | 0.222126103 |
| ENSMUSG0000003038 | chr7 | Zfp606 | -0.24957481 | 0.222292981 |
| ENSMUSG0000002693 | chr2 | Tmem141 | 0.465292818 | 0.222452284 |
| ENSMUSG0000002851 | chr4 | Prkaa2 | 0.326832251 | 0.222982387 |
| ENSMUSG0000003894 | chr1 | Cnst | 0.28647563 | 0.223653887 |
| ENSMUSG0000003361 | chr5 | Cplx1 | -0.433721595 | 0.223792585 |
| ENSMUSG0000009084 | chr10 | Myl6 | 0.505825326 | 0.223792585 |
| ENSMUSG0000002136 | chr13 | Edn1 | 0.667795846 | 0.223792585 |

| | | | | |
|---|---|---|---|---|
| ENSMUSG0000001533 | chr2 | Endog | 0.578484344 | 0.223949614 |
| ENSMUSG0000002088 | chr11 | Dlg4 | -0.282955953 | 0.224043253 |
| ENSMUSG0000004287 | chr6 | Lhfpl4 | -0.496215679 | 0.22420754 |
| ENSMUSG0000003198 | chr8 | Egln1 | 0.289698342 | 0.224595944 |
| ENSMUSG0000003739 | chr5 | Muc17 | -2.760299369 | 0.224650796 |
| ENSMUSG0000000598 | chr19 | Ankrd13d | -0.425534021 | 0.224650796 |
| ENSMUSG0000002894 | chr4 | Hes2 | -2.334662945 | 0.22467553 |
| ENSMUSG0000010342 | chr1 | Golt1a | -3.220109087 | 0.226031355 |
| ENSMUSG0000002756 | chr2 | Dok5 | 0.741099178 | 0.226968494 |
| ENSMUSG0000011797 | chr19 | Itprip | 0.315548413 | 0.227385613 |
| ENSMUSG0000003047 | chr7 | Ceacam18 | -2.364290409 | 0.227532578 |
| ENSMUSG0000002941 | chr2 | Ajm1 | -0.528791878 | 0.227615638 |
| ENSMUSG0000005473 | chr14 | Msra | 0.326181852 | 0.227971448 |
| ENSMUSG0000004768 | chrX | Rtl3 | -0.590075595 | 0.228016329 |
| ENSMUSG0000005098 | chr15 | Apol10a | -2.604264418 | 0.22828964 |
| ENSMUSG0000007917 | chr5 | Zan | -2.671648144 | 0.22828964 |
| ENSMUSG0000009409 | chr11 | Ankrd40cl | -1.627641007 | 0.22828964 |
| ENSMUSG0000000343 | chr7 | Supt5 | 0.174177688 | 0.22828964 |
| ENSMUSG0000007153 | chr6 | Klrg2 | 0.567921653 | 0.22828964 |
| ENSMUSG0000007894 | chr13 | Naip6 | -2.111559506 | 0.228313132 |
| ENSMUSG0000002540 | chr10 | Shmt2 | 0.259707381 | 0.229090371 |
| ENSMUSG0000011203 | chr10 | Brcc3dc | 1.969742751 | 0.229090371 |
| ENSMUSG0000003167 | chr8 | Got2 | 0.246070526 | 0.229272305 |
| ENSMUSG0000002657 | chr1 | Blzf1 | 0.292190714 | 0.229272305 |
| ENSMUSG0000001015 | chr8 | Spire2 | -0.466169912 | 0.229909898 |
| ENSMUSG0000002748 | chr2 | Cdk5rap1 | 0.422417753 | 0.230015489 |
| ENSMUSG0000002600 | chr1 | Acadl | 0.30931771 | 0.230044245 |
| ENSMUSG0000004089 | chr7 | Foxa3 | -2.068601124 | 0.23035042 |
| ENSMUSG0000004014 | chr9 | Rgl3 | -0.432249562 | 0.230911046 |
| ENSMUSG0000002423 | chr18 | Map3k8 | 0.44205557 | 0.230911046 |
| ENSMUSG0000005241 | chr3 | Tchh | -0.799322146 | 0.231168069 |
| ENSMUSG0000002426 | chr18 | Syt4 | -0.453091377 | 0.23128915 |
| ENSMUSG0000003801 | chr10 | Ccdc138 | -0.495807012 | 0.23128915 |
| ENSMUSG0000003824 | chr7 | Mical2 | 0.331446752 | 0.23128915 |
| ENSMUSG0000002170 | chr13 | Rasgrf2 | 0.381804724 | 0.23128915 |
| ENSMUSG0000000162 | chr12 | Ifrd1 | 0.443984744 | 0.23128915 |
| ENSMUSG0000001880 | chr11 | Abca5 | -0.447147433 | 0.231561901 |
| ENSMUSG0000002928 | chr5 | Cdc7 | 0.413743789 | 0.232845936 |
| ENSMUSG0000001666 | chr15 | Pacsin2 | 0.256091996 | 0.233067197 |
| ENSMUSG0000002972 | chr5 | Agfg2 | 0.259108577 | 0.233067197 |
| ENSMUSG0000003643 | chr17 | Tbcc | 0.37058615 | 0.233067197 |
| ENSMUSG0000004926 | chr5 | Kcnk3 | -0.316085995 | 0.23379806 |
| ENSMUSG0000003845 | chr7 | Abhd17c | -0.602379406 | 0.233799758 |
| ENSMUSG0000000335 | chr15 | Ccdc65 | 0.411088682 | 0.233927601 |
| ENSMUSG0000003238 | chr9 | Rbpms2 | 0.336547514 | 0.234395724 |
| ENSMUSG0000000848 | chr4 | Elavl2 | -0.521728315 | 0.234736842 |
| ENSMUSG0000005932 | chr15 | Tonsl | -0.866257832 | 0.23478422 |
| ENSMUSG0000003055 | chr7 | Ttc23 | 0.255479333 | 0.234931186 |
| ENSMUSG0000003177 | chr8 | Cx3cl1 | -0.633222082 | 0.235005874 |
| ENSMUSG0000002788 | chr3 | Gpsm2 | 0.159769549 | 0.235005874 |

| | | | | |
|---|---|---|---|---|
| ENSMUSG0000000654 | chr1 | Prkag3 | 0.81217164 | 0.235005874 |
| ENSMUSG0000003156 | chr8 | Fgfr1 | 0.2116065 | 0.236203088 |
| ENSMUSG0000003860 | chr1 | Dock10 | -0.390120645 | 0.23648929 |
| ENSMUSG0000005073 | chr6 | Vamp8 | 0.386513272 | 0.23648929 |
| ENSMUSG0000005256 | chr15 | Cpne8 | -0.360952473 | 0.236905104 |
| ENSMUSG0000003040 | chr7 | Qpctl | 0.230233909 | 0.236905104 |
| ENSMUSG0000004082 | chr7 | Snrpd2 | 0.648792241 | 0.236905104 |
| ENSMUSG0000002224 | chr15 | Tars1 | 0.249054432 | 0.237113316 |
| ENSMUSG0000009531 | chr2 | Zfp1001 | 0.580420286 | 0.237228216 |
| ENSMUSG0000002007 | chr10 | Ddx50 | 0.260907411 | 0.237327531 |
| ENSMUSG0000000505 | chr10 | Cstb | 0.344121187 | 0.237627286 |
| ENSMUSG0000003706 | chr3 | Sh3glb1 | 0.234135466 | 0.238169015 |
| ENSMUSG0000002650 | chr1 | Capn2 | 0.28121929 | 0.238173281 |
| ENSMUSG0000009199 | chr9 | Smim35 | -1.151987065 | 0.239321446 |
| ENSMUSG0000003655 | chr5 | Iqce | -0.328530086 | 0.23990401 |
| ENSMUSG0000003066 | chr14 | Ipo5 | 0.186001951 | 0.240135837 |
| ENSMUSG0000007955 | chr17 | Marchf2 | 0.265308419 | 0.240135837 |
| ENSMUSG0000007866 | chr3 | Sprr2a1 | -3.362586692 | 0.240284732 |
| ENSMUSG0000002966 | chr6 | Tfpi2 | -0.89433019 | 0.240284732 |
| ENSMUSG0000001114 | chr12 | Adss1 | 1.149768145 | 0.241466185 |
| ENSMUSG0000003427 | chr11 | Thoc5 | 0.255565627 | 0.242123758 |
| ENSMUSG0000011821 | chr1 | Gm29695 | -0.570972538 | 0.242272577 |
| ENSMUSG0000004250 | chr11 | Usp22 | -0.231263793 | 0.242386872 |
| ENSMUSG0000007528 | chr2 | Wipf1 | 0.230181599 | 0.242386872 |
| ENSMUSG0000004142 | chr3 | Paqr6 | -0.459961889 | 0.242458843 |
| ENSMUSG0000003149 | chr8 | Eif4ebp1 | 0.461540928 | 0.242565263 |
| ENSMUSG0000004909 | chr3 | Ankrd34a | -0.424908905 | 0.242775565 |
| ENSMUSG0000003194 | chr8 | Kars1 | 0.249910185 | 0.242975176 |
| ENSMUSG0000003306 | chr14 | Lmo7 | -0.674519395 | 0.242985349 |
| ENSMUSG0000000664 | chr5 | Slc5a6 | -0.472434423 | 0.243272591 |
| ENSMUSG0000005891 | chr15 | C1qtnf3 | 0.39851188 | 0.243272591 |
| ENSMUSG0000002878 | chr4 | Hpca | -0.524519287 | 0.24335137 |
| ENSMUSG0000002618 | chr1 | Pecr | 0.40174262 | 0.24335137 |
| ENSMUSG0000002803 | chr3 | Dnajb4 | 0.346434524 | 0.243432948 |
| ENSMUSG0000000987 | chr2 | Cox4i2 | 0.756044782 | 0.243567984 |
| ENSMUSG0000004346 | chrX | Rab9b | -0.371797204 | 0.243770972 |
| ENSMUSG0000002669 | chr1 | Myoc | 0.681820749 | 0.244063651 |
| ENSMUSG0000000533 | chr1 | Cadm3 | -0.329391717 | 0.244225933 |
| ENSMUSG0000004872 | chr11 | Zfp454 | 0.379240592 | 0.244225933 |
| ENSMUSG0000004253 | chr19 | Golga7b | -0.491738275 | 0.244556776 |
| ENSMUSG0000003491 | chr4 | Ttc22 | -0.696533975 | 0.244556776 |
| ENSMUSG0000002487 | chr19 | Rab1b | 0.215824992 | 0.244556776 |
| ENSMUSG0000004645 | chr7 | Tmem150b | -1.120425023 | 0.244946333 |
| ENSMUSG0000002750 | chr3 | Stmn2 | -0.315499496 | 0.245806658 |
| ENSMUSG0000004503 | chr13 | Ankrd34b | -1.283701961 | 0.245866577 |
| ENSMUSG0000004525 | chr1 | Klhdc9 | -0.851632229 | 0.245866577 |
| ENSMUSG0000004032 | chr11 | Or2v1 | -1.390414201 | 0.245866577 |
| ENSMUSG0000001634 | chr2 | Eef1a2 | -0.355894307 | 0.245866577 |
| ENSMUSG0000002346 | chr17 | Rab12 | 0.276176234 | 0.245866577 |
| ENSMUSG0000004968 | chr5 | Orai1 | 0.267676135 | 0.245866577 |

| | | | | |
|---|---|---|---|---|
| ENSMUSG0000002546 | chr7 | Caly | -0.389613199 | 0.246150339 |
| ENSMUSG0000003098 | chr7 | Knop1 | 0.263300804 | 0.246150339 |
| ENSMUSG0000002305 | chr3 | Fabp2 | -2.569849812 | 0.246438272 |
| ENSMUSG0000004216 | chr13 | Tent2 | 0.200788492 | 0.246438272 |
| ENSMUSG0000000581 | chr3 | Metap1 | 0.169288111 | 0.246703234 |
| ENSMUSG0000004676 | chr12 | Rhoj | 0.35307026 | 0.246703234 |
| ENSMUSG0000003520 | chr11 | Slfn8 | -0.997520619 | 0.247062793 |
| ENSMUSG0000006207 | chrX | Pgk1 | 0.360992281 | 0.247062793 |
| ENSMUSG0000002066 | chr12 | Sh3yl1 | -0.619711996 | 0.247173062 |
| ENSMUSG0000003106 | chrX | Usp11 | -0.242926036 | 0.247186143 |
| ENSMUSG0000003337 | chr3 | Palmd | 0.611825133 | 0.247664273 |
| ENSMUSG0000002951 | chr5 | Cit | 0.528286941 | 0.248861565 |
| ENSMUSG0000004473 | chr13 | Serpinb1a | -0.852927168 | 0.248924848 |
| ENSMUSG0000005117 | chr2 | Plcb1 | 0.255405282 | 0.250449065 |
| ENSMUSG0000005480 | chr7 | Actn4 | 0.331451227 | 0.250449065 |
| ENSMUSG0000003198 | chr8 | 2310022B05Rik | 0.369295344 | 0.250449065 |
| ENSMUSG0000008153 | chr15 | Slc48a1 | 0.206386948 | 0.25047266 |
| ENSMUSG0000003233 | chr9 | Cox7a2 | 0.426001823 | 0.25047266 |
| ENSMUSG0000007858 | chr4 | AU022252 | 0.208571667 | 0.250703869 |
| ENSMUSG0000006225 | chr17 | Lhfpl5 | -0.540941659 | 0.250938435 |
| ENSMUSG0000002109 | chr12 | Clmn | -0.453016882 | 0.250938435 |
| ENSMUSG0000007857 | chr4 | 1110065P20Rik | 0.528726658 | 0.250938435 |
| ENSMUSG0000003305 | chr2 | Pygb | 0.300543188 | 0.25140879 |
| ENSMUSG0000004370 | chr17 | Capn13 | -3.399452453 | 0.251784598 |
| ENSMUSG0000004155 | chrX | Ptchd1 | -0.483502175 | 0.251784598 |
| ENSMUSG0000002410 | chr17 | Nrxn1 | -0.416562945 | 0.251962451 |
| ENSMUSG0000003806 | chr8 | Cdkn2aip | 0.229046232 | 0.251962451 |
| ENSMUSG0000002195 | chr14 | Pinx1 | 0.441627633 | 0.251962451 |
| ENSMUSG0000004120 | chr4 | Pla2g2d | -1.332184896 | 0.252030482 |
| ENSMUSG0000004230 | chr15 | Sgsm3 | -0.313234759 | 0.252030482 |
| ENSMUSG0000009511 | chr7 | Itpripl2 | 0.20318971 | 0.252030482 |
| ENSMUSG0000003936 | chr7 | Picalm | 0.253883545 | 0.252030482 |
| ENSMUSG0000003181 | chr8 | Cox4i1 | 0.542590756 | 0.252030482 |
| ENSMUSG0000002009 | chr10 | Unc5b | -0.655248448 | 0.252135782 |
| ENSMUSG0000005641 | chr5 | Adap1 | -0.277064012 | 0.252135782 |
| ENSMUSG0000001987 | chr10 | Reep3 | 0.260628359 | 0.252135782 |
| ENSMUSG0000002984 | chr6 | Mtpn | 0.278004723 | 0.252135782 |
| ENSMUSG0000007491 | chr2 | Chst14 | 0.245964727 | 0.252135782 |
| ENSMUSG0000002799 | chr3 | Pla2g12a | 0.356826409 | 0.252135782 |
| ENSMUSG0000007422 | chr7 | Spint2 | -0.38827789 | 0.252316933 |
| ENSMUSG0000003860 | chr4 | Gng10 | 0.277785613 | 0.252316933 |
| ENSMUSG0000002534 | chr10 | Itga7 | 0.330119369 | 0.252316933 |
| ENSMUSG0000005251 | chr16 | Robo2 | -0.341501903 | 0.252779024 |
| ENSMUSG0000004938 | chr15 | Krt8 | -1.379981107 | 0.253874186 |
| ENSMUSG0000003161 | chr8 | Nr3c2 | -0.506514063 | 0.254146403 |
| ENSMUSG0000002689 | chr2 | Ifih1 | -0.374641101 | 0.254146403 |
| ENSMUSG0000003320 | chr17 | Tpsg1 | -1.457073465 | 0.254213657 |
| ENSMUSG0000002961 | chr5 | Aimp2 | 0.374099462 | 0.255188043 |
| ENSMUSG0000001705 | chrX | Il13ra1 | 0.273153825 | 0.255460939 |
| ENSMUSG0000004769 | chrX | Yipf6 | -0.246410115 | 0.255483379 |

| | | | | |
|---|---|---|---|---|
| ENSMUSG0000004444 | chr14 | Dock5 | -0.348539075 | 0.255558334 |
| ENSMUSG0000004632 | chr7 | Hs3st2 | -0.47409014 | 0.256093919 |
| ENSMUSG0000002306 | chr17 | Cdkn1a | 0.496366125 | 0.256406228 |
| ENSMUSG0000002937 | chr5 | Areg | -2.280732545 | 0.257223637 |
| ENSMUSG0000009788 | chr11 | Gsg1l2 | -0.655300147 | 0.257223637 |
| ENSMUSG0000004042 | chr7 | Plekha4 | -0.493011106 | 0.257223637 |
| ENSMUSG0000000426 | chr6 | Ptpn6 | -0.527349563 | 0.257223637 |
| ENSMUSG0000003700 | chr15 | Tns2 | -0.243747462 | 0.257223637 |
| ENSMUSG0000002169 | chr13 | Elovl7 | -0.54520993 | 0.25764245 |
| ENSMUSG0000002555 | chr14 | Fgf14 | -0.61928392 | 0.25764245 |
| ENSMUSG0000004064 | chr14 | Erc2 | -0.392298007 | 0.258306051 |
| ENSMUSG0000003118 | chrX | Aff2 | -0.927087784 | 0.258569713 |
| ENSMUSG0000002211 | chr14 | Sucla2 | 0.229706126 | 0.258634632 |
| ENSMUSG0000000476 | chr1 | Rab23 | 0.418937642 | 0.258634632 |
| ENSMUSG0000002971 | chr5 | Actl6b | -0.453127355 | 0.259037116 |
| ENSMUSG0000000620 | chr7 | Htra1 | 0.346853395 | 0.259037116 |
| ENSMUSG0000000127 | chr12 | Ckb | 0.450197243 | 0.259037116 |
| ENSMUSG0000005981 | chr17 | Atl2 | 0.223175335 | 0.259108628 |
| ENSMUSG0000003442 | chr11 | Myo15b | -2.60738155 | 0.259900843 |
| ENSMUSG0000002933 | chr5 | Cds1 | -0.596702102 | 0.259900843 |
| ENSMUSG0000003569 | chr4 | Isg15 | -1.48396 | 0.259900843 |
| ENSMUSG0000000354 | chr17 | Ier3 | 0.47057816 | 0.259900843 |
| ENSMUSG0000005736 | chr1 | Uxs1 | -0.213632942 | 0.259977238 |
| ENSMUSG0000003785 | chr8 | Cpe | 0.303927519 | 0.259977238 |
| ENSMUSG0000002477 | chr19 | Ehd1 | 0.355113989 | 0.259977238 |
| ENSMUSG0000006110 | chr8 | Sap18b | 1.100354124 | 0.259977238 |
| ENSMUSG0000000522 | chr6 | Plekha8 | -0.403864338 | 0.260356672 |
| ENSMUSG0000002607 | chr1 | Pdcl3 | 0.385334315 | 0.260356672 |
| ENSMUSG0000000849 | chr7 | Pou2f2 | -0.7060457 | 0.260634468 |
| ENSMUSG0000000403 | chr3 | Gstm5 | 0.333107244 | 0.260634468 |
| ENSMUSG0000005532 | chr1 | Tns1 | 0.289299446 | 0.260822343 |
| ENSMUSG0000002701 | chr2 | Cwc22 | 0.223685968 | 0.260822343 |
| ENSMUSG0000004764 | chr7 | Polr1g | 0.375174259 | 0.260822343 |
| ENSMUSG0000003385 | chr17 | Ston1 | 0.345249186 | 0.260822343 |
| ENSMUSG0000002859 | chr4 | Pramel13 | 1.294635498 | 0.260822343 |
| ENSMUSG0000005814 | chr7 | Adamts17 | 0.476379629 | 0.261334607 |
| ENSMUSG0000006268 | chr15 | Atp5mc2 | 0.337524057 | 0.261340781 |
| ENSMUSG0000005985 | chr18 | Kcng2 | 0.58405806 | 0.261340781 |
| ENSMUSG0000002032 | chr10 | Fstl3 | 0.591264109 | 0.261421547 |
| ENSMUSG0000002103 | chr12 | Ngb | -0.33194534 | 0.262002343 |
| ENSMUSG0000000798 | chr14 | Fzd3 | -0.272832096 | 0.262505153 |
| ENSMUSG0000003740 | chr3 | Atp11b | -0.316056344 | 0.262616467 |
| ENSMUSG0000001700 | chr2 | Sdc4 | 0.432701898 | 0.262939445 |
| ENSMUSG0000002289 | chr16 | App | 0.16486723 | 0.263654284 |
| ENSMUSG0000006258 | chr1 | Timm17a | 0.268386717 | 0.263654284 |
| ENSMUSG0000002588 | chr6 | Snca | -0.286542775 | 0.26377198 |
| ENSMUSG0000001877 | chr2 | Atp5mc3 | 0.364187542 | 0.26377198 |
| ENSMUSG0000003876 | chr15 | Trhr | 1.416669609 | 0.26377198 |
| ENSMUSG0000009003 | chr10 | Galnt4 | -0.43242691 | 0.264422309 |
| ENSMUSG0000004879 | chr18 | Cep120 | -0.227472312 | 0.264422309 |

| | | | | |
|---|---|---|---|---|
| ENSMUSG0000010377 | chr18 | Pcdha9 | -0.996968167 | 0.265335392 |
| ENSMUSG0000000303 | chr4 | Klf4 | -0.607873384 | 0.265335392 |
| ENSMUSG0000006190 | chr10 | Slc25a3 | 0.2650386 | 0.265335392 |
| ENSMUSG0000004875 | chr15 | Mcat | 0.376643097 | 0.265335392 |
| ENSMUSG0000003191 | chr8 | Nip7 | 0.414974295 | 0.265335392 |
| ENSMUSG0000007488 | chr2 | Serf2 | 0.479119888 | 0.265352786 |
| ENSMUSG0000002062 | chr11 | Abca8b | -0.457590757 | 0.265482771 |
| ENSMUSG0000003185 | chr8 | Ntpcr | 0.320732029 | 0.26555707 |
| ENSMUSG0000003854 | chr17 | Srd5a2 | -2.758097161 | 0.265682518 |
| ENSMUSG0000003405 | chr2 | Ypel4 | -0.460470586 | 0.265682518 |
| ENSMUSG0000002032 | chr11 | Ccng1 | 0.304002406 | 0.265682518 |
| ENSMUSG0000000432 | chr7 | Hif3a | 0.658257491 | 0.265682518 |
| ENSMUSG0000004363 | chr5 | Adamts3 | 0.493680313 | 0.267071773 |
| ENSMUSG0000003445 | chr14 | Gulo | -0.92410195 | 0.26708059 |
| ENSMUSG0000005229 | chr7 | Ppp6r1 | 0.204259044 | 0.26708059 |
| ENSMUSG0000001814 | chr5 | Mafk | 0.351358099 | 0.26708059 |
| ENSMUSG0000003218 | chr9 | Tmod2 | -0.350222357 | 0.268449788 |
| ENSMUSG0000009491 | chr12 | D430019H16Rik | -0.369969996 | 0.268614582 |
| ENSMUSG0000002348 | chr15 | Prph | -0.353442675 | 0.268671641 |
| ENSMUSG0000003643 | chr17 | Calm2 | 0.345719012 | 0.268795498 |
| ENSMUSG0000000418 | chr15 | Kifc2 | -0.387850952 | 0.269118592 |
| ENSMUSG0000003357 | chr9 | Myo6 | -0.323860472 | 0.269118592 |
| ENSMUSG0000002628 | chr1 | Atg16l1 | 0.184856756 | 0.269118592 |
| ENSMUSG0000002648 | chr1 | Rnf2 | 0.189069889 | 0.269118592 |
| ENSMUSG0000006195 | chr8 | Ces1e | -1.613436636 | 0.269190223 |
| ENSMUSG0000003331 | chr5 | Galnt9 | -0.52175604 | 0.269832464 |
| ENSMUSG0000004896 | chr8 | Yjefn3 | 0.846895477 | 0.269888883 |
| ENSMUSG0000002067 | chr12 | Rab10 | 0.21474178 | 0.271006308 |
| ENSMUSG0000002243 | chr15 | Samm50 | 0.262134473 | 0.271006308 |
| ENSMUSG0000002845 | chr4 | Stoml2 | 0.352091328 | 0.271006308 |
| ENSMUSG0000004633 | chr1 | Rpl37a | 0.501227827 | 0.271006308 |
| ENSMUSG0000002118 | chr12 | Tc2n | -0.731034819 | 0.271234531 |
| ENSMUSG0000005914 | chr1 | Mfsd4a | -0.269656381 | 0.271234531 |
| ENSMUSG0000000210 | chr2 | Psmc3 | 0.306323098 | 0.271234531 |
| ENSMUSG0000003862 | chr7 | Tm6sf1 | 0.411235423 | 0.271376723 |
| ENSMUSG0000003800 | chr8 | Acd | 0.225641096 | 0.271491765 |
| ENSMUSG0000005115 | chr2 | Commd3 | 0.320022394 | 0.271491765 |
| ENSMUSG0000000102 | chr3 | S100a6 | 0.564826467 | 0.271491765 |
| ENSMUSG0000000228 | chr17 | Angptl4 | 2.081999646 | 0.271517408 |
| ENSMUSG0000002763 | chr2 | Samhd1 | -0.282778658 | 0.271916075 |
| ENSMUSG0000006819 | chr16 | Col8a1 | 0.506791438 | 0.271916075 |
| ENSMUSG0000003675 | chr7 | Cox6b1 | 0.344270604 | 0.271975939 |
| ENSMUSG0000003300 | chr15 | Sox10 | -0.285708302 | 0.272043573 |
| ENSMUSG0000004466 | chr3 | Plppr4 | -0.53176952 | 0.272897077 |
| ENSMUSG0000000073 | chr10 | Icosl | -0.354097811 | 0.272897077 |
| ENSMUSG0000002480 | chr19 | Rpp30 | 0.361377894 | 0.272897077 |
| ENSMUSG0000011202 | chr10 | Lilrb4b | 0.8174744 | 0.272897077 |
| ENSMUSG0000003888 | chr9 | Shfl | -0.356709343 | 0.273363742 |
| ENSMUSG0000004493 | chr13 | Zfp367 | 0.253049776 | 0.273363742 |
| ENSMUSG0000000224 | chr15 | Adsl | 0.347709855 | 0.273363742 |

| | | | | |
|---|---|---|---|---|
| ENSMUSG0000002047 | chr11 | Mrps24 | 0.497363215 | 0.273363742 |
| ENSMUSG0000004840 | chr1 | Gli2 | 0.303730838 | 0.273662578 |
| ENSMUSG0000002617 | chr1 | Vil1 | -2.494748577 | 0.273706698 |
| ENSMUSG0000004276 | chr3 | Trim46 | -0.384718509 | 0.273706698 |
| ENSMUSG0000002043 | chr11 | Smtn | 0.339710805 | 0.274192711 |
| ENSMUSG0000000621 | chr4 | Zbtb17 | 0.264625694 | 0.274192711 |
| ENSMUSG0000000595 | chr11 | Shpk | -0.365880113 | 0.274229255 |
| ENSMUSG0000004038 | chr3 | Wdr47 | -0.276242882 | 0.274229255 |
| ENSMUSG0000002552 | chrX | Zfp711 | -0.679365143 | 0.2745688 |
| ENSMUSG0000005896 | chr7 | Tlcd3b | -0.379897454 | 0.274640704 |
| ENSMUSG0000003416 | chr17 | Ccnd3 | 0.387186146 | 0.274741849 |
| ENSMUSG0000007820 | chr2 | Nrarp | -1.139745163 | 0.274856009 |
| ENSMUSG0000003152 | chr8 | Dlc1 | 0.263985984 | 0.275092243 |
| ENSMUSG0000004167 | chr1 | Rims1 | 0.353094139 | 0.275092243 |
| ENSMUSG0000006356 | chr11 | Col23a1 | 0.382460904 | 0.275092243 |
| ENSMUSG0000004219 | chr5 | Cmklr1 | 0.384482459 | 0.275092243 |
| ENSMUSG0000009007 | chr1 | Cdk5r2 | -0.409683364 | 0.275183916 |
| ENSMUSG0000002080 | chr11 | Slc13a5 | -0.597741279 | 0.275616779 |
| ENSMUSG0000004399 | chr11 | Gpr75 | -0.505671068 | 0.275616779 |
| ENSMUSG0000002813 | chr3 | Adh5 | 0.278206063 | 0.275616779 |
| ENSMUSG0000002603 | chr1 | Ndufb3 | 0.334236422 | 0.275616779 |
| ENSMUSG0000001850 | chr11 | Trpv2 | 0.359352934 | 0.275616779 |
| ENSMUSG0000004728 | chr4 | Sfn | -0.847078334 | 0.275712466 |
| ENSMUSG0000002100 | chr12 | Ptpn21 | 0.261271634 | 0.275712466 |
| ENSMUSG0000003415 | chr3 | Zbbx | -1.227778328 | 0.275929108 |
| ENSMUSG0000007155 | chr6 | Cpa2 | -1.130436355 | 0.275929108 |
| ENSMUSG0000003085 | chr7 | Ptpn5 | -0.505836593 | 0.275929108 |
| ENSMUSG0000002205 | chr14 | Nefm | -0.310441049 | 0.275929108 |
| ENSMUSG0000000075 | chr11 | Rpa1 | 0.277167999 | 0.275929108 |
| ENSMUSG0000003198 | chr8 | Gnpat | 0.24249615 | 0.276088865 |
| ENSMUSG0000005148 | chr16 | Cbr1 | -0.279740517 | 0.276207117 |
| ENSMUSG0000002958 | chr5 | Zfp12 | 0.250390707 | 0.276207117 |
| ENSMUSG0000004371 | chr1 | Rpl7 | 0.36631977 | 0.276207117 |
| ENSMUSG0000002103 | chr12 | Sptlc2 | -0.272897756 | 0.276935812 |
| ENSMUSG0000003093 | chr7 | Oat | 0.326848948 | 0.27710338 |
| ENSMUSG0000004955 | chr9 | Tmie | -0.487380016 | 0.277150851 |
| ENSMUSG0000002430 | chr18 | Cdh2 | -0.316730164 | 0.277894684 |
| ENSMUSG0000002013 | chr11 | Peli1 | 0.239229012 | 0.277944151 |
| ENSMUSG0000004238 | chr4 | Smim12 | 0.398960321 | 0.277944151 |
| ENSMUSG0000002585 | chr5 | Prkar1b | -0.317651814 | 0.277945066 |
| ENSMUSG0000003142 | chrX | Plp1 | -0.255411552 | 0.277945066 |
| ENSMUSG0000005568 | chr8 | Cope | 0.346022869 | 0.277945066 |
| ENSMUSG0000001882 | chr19 | Avpi1 | 0.598636465 | 0.278067732 |
| ENSMUSG0000001982 | chr10 | Grm1 | -1.331058524 | 0.278080271 |
| ENSMUSG0000004512 | chr8 | Rpl18a | 0.280860606 | 0.278224033 |
| ENSMUSG0000002409 | chr17 | Ndufv2 | 0.352063823 | 0.278741751 |
| ENSMUSG0000007818 | chr1 | Chml | -0.310392584 | 0.278807065 |
| ENSMUSG0000002717 | chr2 | Eif3m | 0.360451717 | 0.27882809 |
| ENSMUSG0000005914 | chr7 | Ntrk3 | -0.390758478 | 0.279068197 |
| ENSMUSG0000007363 | chr1 | Fbxo36 | -0.760446288 | 0.279330426 |

| | | | | |
|---|---|---|---|---|
| ENSMUSG0000003416 | chr15 | Scx | 0.431634091 | 0.279388592 |
| ENSMUSG0000000354 | chr7 | Ercc1 | 0.29885497 | 0.27953 |
| ENSMUSG0000003901 | chr7 | Mtg1 | 0.328056325 | 0.27953 |
| ENSMUSG0000003815 | chr10 | Prdm1 | -1.587214978 | 0.280121212 |
| ENSMUSG0000004064 | chr6 | Rimklb | -0.451651626 | 0.280121212 |
| ENSMUSG0000002296 | chr16 | Tmem50b | 0.294797291 | 0.280121212 |
| ENSMUSG0000002178 | chr14 | Oxsm | -0.264836747 | 0.280284665 |
| ENSMUSG0000007088 | chr2 | Gad1 | -0.799031933 | 0.280693829 |
| ENSMUSG0000002701 | chr2 | Slc25a12 | 0.245834136 | 0.281591192 |
| ENSMUSG0000002969 | chr6 | Aass | -0.882060145 | 0.281656799 |
| ENSMUSG0000003431 | chr3 | Trim59 | -0.336396195 | 0.281656799 |
| ENSMUSG0000000199 | chr2 | Blvra | 0.36846976 | 0.281656799 |
| ENSMUSG0000007171 | chrX | Nalf2 | -0.532052323 | 0.28369875 |
| ENSMUSG0000001884 | chr11 | Pank3 | -0.256285412 | 0.284039226 |
| ENSMUSG0000004420 | chr18 | Cdc25c | -2.235657514 | 0.284228602 |
| ENSMUSG0000002863 | chr4 | Ccdc30 | -0.472222134 | 0.284550505 |
| ENSMUSG0000003060 | chr7 | Psmc4 | 0.322296257 | 0.284550505 |
| ENSMUSG0000002540 | chr10 | Myo1a | -2.231335242 | 0.284719358 |
| ENSMUSG0000003059 | chr7 | Psmd8 | 0.350383605 | 0.284719358 |
| ENSMUSG0000000699 | chr16 | Psmd2 | 0.157842372 | 0.284925898 |
| ENSMUSG0000002084 | chr11 | Nxn | 0.445149897 | 0.284925898 |
| ENSMUSG0000006174 | chr19 | Slc22a12 | 1.513241525 | 0.284925898 |
| ENSMUSG0000000344 | chr7 | Med29 | 0.391538016 | 0.284971049 |
| ENSMUSG0000003826 | chr13 | Slc22a23 | -0.564706054 | 0.285218861 |
| ENSMUSG0000002109 | chr12 | Ppm1a | 0.179211025 | 0.285218861 |
| ENSMUSG0000002064 | chr12 | Id2 | 0.437185592 | 0.285218861 |
| ENSMUSG0000000630 | chr1 | Arpc2 | 0.258386873 | 0.285439313 |
| ENSMUSG0000007916 | chr8 | Cd209g | -1.224525671 | 0.286183046 |
| ENSMUSG0000003020 | chr6 | Emp1 | -0.33253889 | 0.286183046 |
| ENSMUSG0000004784 | chr5 | Bri3 | 0.364765444 | 0.286183046 |
| ENSMUSG0000003867 | chr7 | Mybpc2 | 0.533242253 | 0.286183046 |
| ENSMUSG0000003495 | chr7 | Cebpa | -0.444336542 | 0.286325562 |
| ENSMUSG0000004609 | chr4 | Hpcal4 | -0.364332318 | 0.286745613 |
| ENSMUSG0000002707 | chr2 | Ube2l6 | -0.357068698 | 0.286934456 |
| ENSMUSG0000003984 | chr2 | Rapgef1 | 0.276067721 | 0.286934456 |
| ENSMUSG0000002910 | chr5 | Add1 | 0.130701295 | 0.286964425 |
| ENSMUSG0000000815 | chr6 | Clstn3 | -0.467974104 | 0.287589722 |
| ENSMUSG0000003402 | chr10 | Cct2 | 0.256380281 | 0.288173363 |
| ENSMUSG0000003058 | chr7 | Yif1b | 0.316405818 | 0.288247638 |
| ENSMUSG0000002943 | chr5 | Mmp17 | 0.314474495 | 0.288474146 |
| ENSMUSG0000005761 | chrX | Ldoc1 | -1.698752656 | 0.288817567 |
| ENSMUSG0000002284 | chr16 | Adcy5 | 0.186179677 | 0.288817567 |
| ENSMUSG0000002558 | chr7 | Pde8a | 0.337923999 | 0.288817567 |
| ENSMUSG0000003690 | chr19 | Unc93b1 | -0.321821378 | 0.289384344 |
| ENSMUSG0000003131 | chrX | Nono | 0.213531677 | 0.289419271 |
| ENSMUSG0000002094 | chr11 | Map3k14 | 0.293977521 | 0.289419271 |
| ENSMUSG0000003619 | chr8 | Ndufa13 | 0.409561675 | 0.28956501 |
| ENSMUSG0000005629 | chr14 | Synpr | -0.53968753 | 0.290027763 |
| ENSMUSG0000001828 | chr11 | Psmb6 | 0.428514343 | 0.290027763 |
| ENSMUSG0000003471 | chr11 | Ttyh2 | -0.297070807 | 0.290398994 |

| | | | | |
|---|---|---|---|---|
| ENSMUSG0000004674 | chr16 | Tmem45a2 | 1.947154117 | 0.290398994 |
| ENSMUSG0000001995 | chr10 | Poc1b | -0.231130569 | 0.290422945 |
| ENSMUSG0000006856 | chr7 | Myadm | 0.244167034 | 0.290422945 |
| ENSMUSG0000003182 | chr8 | Babam1 | 0.210550528 | 0.290422945 |
| ENSMUSG0000007834 | chr10 | Sf3b5 | 0.307250257 | 0.290422945 |
| ENSMUSG0000003866 | chr6 | Dgki | -0.484586008 | 0.290813225 |
| ENSMUSG0000005292 | chr18 | Ctif | 0.264845024 | 0.290813225 |
| ENSMUSG0000007277 | chr5 | Zfp951 | 0.375597452 | 0.290813225 |
| ENSMUSG0000001863 | chr11 | Crhr1 | 0.838172017 | 0.290813225 |
| ENSMUSG0000004068 | chr16 | Hmgn1 | 0.374373908 | 0.29108877 |
| ENSMUSG0000002940 | chr5 | Cdkl2 | -0.257135953 | 0.291889901 |
| ENSMUSG0000007243 | chr19 | Nanos1 | -0.505155976 | 0.292395165 |
| ENSMUSG0000002439 | chr17 | Ltb | -1.606208009 | 0.292395165 |
| ENSMUSG0000002897 | chr4 | Masp2 | 1.297402947 | 0.292561486 |
| ENSMUSG0000003967 | chr2 | Tbc1d13 | -0.385079598 | 0.292659992 |
| ENSMUSG0000002223 | chr15 | Cct5 | 0.328763319 | 0.292659992 |
| ENSMUSG0000006058 | chr17 | H2-Eb1 | -0.603135219 | 0.292722134 |
| ENSMUSG0000000488 | chr1 | Lbr | 0.197984188 | 0.293813366 |
| ENSMUSG0000002854 | chr4 | Elavl4 | -0.335834231 | 0.29416133 |
| ENSMUSG0000003218 | chr9 | Tmed1 | 0.350668442 | 0.29416133 |
| ENSMUSG0000003833 | chr10 | Sesn1 | 0.365575629 | 0.29416133 |
| ENSMUSG0000002927 | chr5 | Dipk1a | 0.377125337 | 0.29416133 |
| ENSMUSG0000003716 | chr3 | Mgarp | 0.94720434 | 0.294174191 |
| ENSMUSG0000004240 | chr15 | Atf4 | 0.318546155 | 0.294481689 |
| ENSMUSG0000003611 | chr13 | Slc17a2 | -2.641820768 | 0.29457561 |
| ENSMUSG0000004066 | chr19 | Clcf1 | -0.474682724 | 0.294985947 |
| ENSMUSG0000006157 | chr5 | Dpp6 | -0.32830212 | 0.294985947 |
| ENSMUSG0000003144 | chr8 | Cul4a | 0.158439952 | 0.295096268 |
| ENSMUSG0000003234 | chr9 | Ooep | 0.445870555 | 0.295096268 |
| ENSMUSG0000004264 | chr17 | Itpr3 | -0.434574247 | 0.295180434 |
| ENSMUSG0000006597 | chr16 | Cpped1 | 0.224467203 | 0.295180434 |
| ENSMUSG0000000198 | chr7 | Npas1 | -0.607164819 | 0.295264175 |
| ENSMUSG0000005567 | chr11 | Zzef1 | -0.26163133 | 0.295333893 |
| ENSMUSG0000003166 | chr8 | Aktip | 0.26389264 | 0.295333893 |
| ENSMUSG0000001126 | chr7 | Exoc3l2 | 0.937317015 | 0.295333893 |
| ENSMUSG0000002493 | chr19 | Ehbp1l1 | 0.220952665 | 0.295617524 |
| ENSMUSG0000003626 | chr11 | Fstl4 | 0.558959297 | 0.295617524 |
| ENSMUSG0000001522 | chr1 | Map2 | -0.370328845 | 0.296000724 |
| ENSMUSG0000003499 | chr11 | Vat1 | -0.292190846 | 0.296000724 |
| ENSMUSG0000000568 | chr10 | Cs | 0.184914074 | 0.296000724 |
| ENSMUSG0000005651 | chr17 | Rab31 | 0.344823997 | 0.296000724 |
| ENSMUSG0000000954 | chr7 | Kcnq1 | -0.988004955 | 0.296244822 |
| ENSMUSG0000005847 | chr19 | Gm8369 | 1.709278281 | 0.296337884 |
| ENSMUSG0000001989 | chr10 | Dcbld1 | -0.453477838 | 0.296417533 |
| ENSMUSG0000001973 | chr7 | Polr2i | 0.373203891 | 0.296417533 |
| ENSMUSG0000002102 | chr12 | Psma6 | 0.369372327 | 0.296883058 |
| ENSMUSG0000005760 | chr6 | Lmcd1 | 0.506470729 | 0.296976451 |
| ENSMUSG0000003701 | chr5 | Zscan21 | 0.261684968 | 0.297476947 |
| ENSMUSG0000002125 | chr12 | Tgfb3 | 0.352085335 | 0.297872773 |
| ENSMUSG0000009064 | chr13 | Zfp712 | -0.39756357 | 0.297919495 |

| | | | | |
|---|---|---|---|---|
| ENSMUSG0000003510 | chr16 | Dcbld2 | 0.218828224 | 0.297919495 |
| ENSMUSG0000001783 | chr11 | Dhx58 | -0.89264085 | 0.298036034 |
| ENSMUSG0000007349 | chr1 | Ifi213 | -1.207688042 | 0.298036034 |
| ENSMUSG0000006719 | chr19 | Frat1 | -0.67261182 | 0.298036034 |
| ENSMUSG0000000628 | chr14 | Tep1 | -0.357224213 | 0.298036034 |
| ENSMUSG0000002769 | chr3 | Pld1 | -0.405456843 | 0.298036034 |
| ENSMUSG0000005666 | chr6 | Retsat | -0.373568114 | 0.298036034 |
| ENSMUSG0000009965 | chr19 | 2310034G01Rik | -1.050055601 | 0.298036034 |
| ENSMUSG0000002631 | chr1 | Relch | -0.302759168 | 0.298036034 |
| ENSMUSG0000002248 | chr15 | Pde1b | -0.278096867 | 0.298036034 |
| ENSMUSG0000007879 | chr7 | Dact3 | 0.224096419 | 0.298036034 |
| ENSMUSG0000001091 | chr2 | Pdhx | 0.276074661 | 0.298036034 |
| ENSMUSG0000006003 | chr15 | Rpl3 | 0.373283951 | 0.298036034 |
| ENSMUSG0000006331 | chr11 | Rpl27 | 0.426271301 | 0.298036034 |
| ENSMUSG0000002265 | chr16 | Tagln3 | -0.386359445 | 0.298055134 |
| ENSMUSG0000002971 | chr5 | Tfr2 | -0.72793337 | 0.298055134 |
| ENSMUSG0000003534 | chr19 | Lzts2 | 0.188964859 | 0.298055134 |
| ENSMUSG0000003714 | chr8 | Arhgap10 | 0.320939278 | 0.298205895 |
| ENSMUSG0000002853 | chr4 | Artn | -0.35284139 | 0.298372021 |
| ENSMUSG0000003417 | chr11 | Rnf43 | -1.048179312 | 0.298692102 |
| ENSMUSG0000005094 | chr3 | Amigo1 | -0.259832837 | 0.298692102 |
| ENSMUSG0000009001 | chr5 | Gm15446 | -0.270333778 | 0.298692102 |
| ENSMUSG0000004307 | chr18 | Synpo | 0.288001625 | 0.298692102 |
| ENSMUSG0000002720 | chr2 | Fbn1 | 0.346339465 | 0.298692102 |
| ENSMUSG0000000693 | chr11 | Hap1 | -0.436869653 | 0.299067957 |
| ENSMUSG0000002895 | chr5 | Abcf2 | 0.230128099 | 0.299067957 |
| ENSMUSG0000002032 | chr10 | Polrmt | 0.240879679 | 0.299067957 |
| ENSMUSG0000002391 | chr17 | Tnfrsf21 | -0.381821868 | 0.299110558 |
| ENSMUSG0000004313 | chr6 | Mob1a | 0.195143726 | 0.299110558 |
| ENSMUSG0000000166 | chr10 | Gstt1 | 0.322306126 | 0.299273753 |
| ENSMUSG0000002793 | chr3 | Creb3l4 | -0.987126108 | 0.299339124 |
| ENSMUSG0000006762 | chr17 | Syngap1 | -0.306529273 | 0.299339124 |
| ENSMUSG0000002067 | chr12 | Sntg2 | 0.274534875 | 0.299339124 |
| ENSMUSG0000005273 | chr6 | Suclg1 | 0.288729292 | 0.299339124 |
| ENSMUSG0000002557 | chr11 | Cbx8 | 0.373736895 | 0.299339124 |
| ENSMUSG0000003999 | chr10 | Timeless | -0.538473506 | 0.299685128 |
| ENSMUSG0000004011 | chr9 | Gramd1b | -0.329730401 | 0.299685128 |
| ENSMUSG0000002012 | chr11 | Egfr | 0.271511645 | 0.299685128 |
| ENSMUSG0000007482 | chr9 | 2010315B03Rik | -0.265400932 | 0.299809331 |
| ENSMUSG0000006623 | chr7 | Ipo7 | 0.202586906 | 0.299971846 |
| ENSMUSG0000007225 | chr1 | Taf1a | 0.272897745 | 0.299971846 |
| ENSMUSG0000002605 | chr1 | Khdrbs2 | -0.482319326 | 0.30018622 |
| ENSMUSG0000007165 | chr19 | Gng3 | -0.224574797 | 0.300227535 |
| ENSMUSG0000002595 | chr1 | Idh1 | -0.205723626 | 0.300227535 |
| ENSMUSG0000002661 | chr1 | Eprs1 | 0.174607066 | 0.3004512 |
| ENSMUSG0000002950 | chr5 | Ddx51 | 0.215448354 | 0.3004512 |
| ENSMUSG0000002552 | chrX | Apool | 0.395578234 | 0.3004512 |
| ENSMUSG0000009813 | chr7 | Rassf10 | -0.761495069 | 0.300551853 |
| ENSMUSG0000002233 | chr15 | Osr2 | -2.143648693 | 0.301172589 |
| ENSMUSG0000005190 | chr8 | Cd209f | -0.962226168 | 0.301172589 |

| | | | | |
|---|---|---|---|---|
| ENSMUSG0000002160 | chr13 | Slc6a3 | -1.764031822 | 0.301172589 |
| ENSMUSG0000000232 | chr14 | Irf9 | -0.503661456 | 0.301172589 |
| ENSMUSG0000005552 | chr19 | Gucy2g | -1.068881538 | 0.301172589 |
| ENSMUSG0000003920 | chr16 | Atf7ip2 | -3.050383904 | 0.301172589 |
| ENSMUSG0000003133 | chrX | Mtm1 | -0.283429706 | 0.301172589 |
| ENSMUSG0000002074 | chr11 | Tmem94 | -0.267413557 | 0.301172589 |
| ENSMUSG0000001949 | chr5 | Cops6 | 0.323183844 | 0.301172589 |
| ENSMUSG0000001892 | chr11 | Med11 | 0.444446293 | 0.301172589 |
| ENSMUSG0000001858 | chr11 | Atox1 | 0.416773119 | 0.301172589 |
| ENSMUSG0000001239 | chr6 | Nanog | 1.838924558 | 0.301172589 |
| ENSMUSG0000005817 | chr3 | Gm5148 | -0.842972083 | 0.301596939 |
| ENSMUSG0000002522 | chr19 | Nfkb2 | 0.376135966 | 0.301654394 |
| ENSMUSG0000001935 | chrX | Gdpd2 | -2.267544335 | 0.301729799 |
| ENSMUSG0000003200 | chr9 | Birc3 | -0.244659621 | 0.301729799 |
| ENSMUSG0000001982 | chr10 | Smpd2 | 0.292746767 | 0.301729799 |
| ENSMUSG0000003459 | chr17 | Ppp1r18 | 0.310632363 | 0.301729799 |
| ENSMUSG0000002395 | chr17 | Vegfa | 0.258713326 | 0.301894477 |
| ENSMUSG0000002535 | chr10 | Rdh5 | 0.391008507 | 0.302393846 |
| ENSMUSG0000002026 | chr10 | Pofut2 | 0.273877448 | 0.302760836 |
| ENSMUSG0000003433 | chr19 | Ina | -0.385908725 | 0.303115402 |
| ENSMUSG0000007287 | chr6 | Gpr27 | -0.284627865 | 0.303115402 |
| ENSMUSG0000005021 | chr4 | Snip1 | 0.310992032 | 0.303115402 |
| ENSMUSG0000002245 | chr15 | Twf1 | 0.253883172 | 0.303300717 |
| ENSMUSG0000007871 | chr4 | Tmem8b | -0.315107226 | 0.303565115 |
| ENSMUSG0000002780 | chr3 | Tm4sf4 | -0.687442763 | 0.303957916 |
| ENSMUSG0000004035 | chr11 | Trim7 | -0.488683092 | 0.303957916 |
| ENSMUSG0000006155 | chr9 | Skic8 | 0.342227915 | 0.303957916 |
| ENSMUSG0000002786 | chr3 | Ptgfrn | 0.22426755 | 0.303965634 |
| ENSMUSG0000000019 | chr14 | Nalcn | -0.440378635 | 0.304207914 |
| ENSMUSG0000000224 | chr17 | Tead3 | 0.205480761 | 0.304207914 |
| ENSMUSG0000000202 | chr12 | Ltbp2 | 0.454008335 | 0.304207914 |
| ENSMUSG0000007843 | chr10 | Smim24 | -1.675221074 | 0.304256001 |
| ENSMUSG0000004425 | chr13 | Ctla2a | 0.455480221 | 0.30449052 |
| ENSMUSG0000005057 | chr9 | Mmp13 | -1.599527814 | 0.304562951 |
| ENSMUSG0000003724 | chr4 | Clic4 | 0.273516171 | 0.304562951 |
| ENSMUSG0000003053 | chr7 | Sema4b | -0.247661064 | 0.304613764 |
| ENSMUSG0000005277 | chr5 | Oas1a | -0.483946514 | 0.305121136 |
| ENSMUSG0000003863 | chr1 | Degs1 | 0.214579484 | 0.305121136 |
| ENSMUSG0000007031 | chr9 | Eif3g | 0.323225802 | 0.305121136 |
| ENSMUSG0000000334 | chr10 | Mob3a | 0.391555323 | 0.305121136 |
| ENSMUSG0000002222 | chr14 | Atp12a | 0.608106983 | 0.305121136 |
| ENSMUSG0000002518 | chr19 | Loxl4 | 0.289083532 | 0.305303363 |
| ENSMUSG0000003116 | chrX | Rbm3 | 0.399526674 | 0.30594673 |
| ENSMUSG0000002475 | chr19 | Atl3 | 0.203273847 | 0.306388098 |
| ENSMUSG0000006885 | chr3 | Sf3b4 | 0.222577829 | 0.306429266 |
| ENSMUSG0000000649 | chr2 | Pdk1 | 0.348871478 | 0.307107603 |
| ENSMUSG0000004026 | chr3 | Mrps28 | 0.48128351 | 0.307107603 |
| ENSMUSG0000003113 | chrX | Mcf2 | -1.509884721 | 0.307284233 |
| ENSMUSG0000003589 | chr5 | Uba6 | -0.209104298 | 0.307423356 |
| ENSMUSG0000003497 | chr10 | Dapk3 | 0.337646186 | 0.307717956 |

| | | | | |
|---|---|---|---|---|
| ENSMUSG0000003826 | chr11 | Ovca2 | 0.436065242 | 0.307717956 |
| ENSMUSG0000003990 | chr6 | Gpr37 | -0.508079811 | 0.307871764 |
| ENSMUSG0000002547 | chr7 | Dpysl4 | -0.375695838 | 0.308348824 |
| ENSMUSG0000002811 | chr3 | Ctsk | 0.46414913 | 0.308348824 |
| ENSMUSG0000004173 | chr3 | Kirrel1 | 0.21828482 | 0.309090636 |
| ENSMUSG0000001984 | chr10 | Prep | 0.255692134 | 0.309244729 |
| ENSMUSG0000009660 | chr7 | Tpbgl | -0.423721112 | 0.309311511 |
| ENSMUSG0000004446 | chr14 | Shisa2 | -0.319509069 | 0.309585746 |
| ENSMUSG0000003238 | chr9 | Ppib | 0.301986649 | 0.309707523 |
| ENSMUSG0000000003 | chrX | Scml2 | -0.559928039 | 0.309754793 |
| ENSMUSG0000002276 | chr16 | Ncam2 | -0.345487187 | 0.310038957 |
| ENSMUSG0000002307 | chrX | Mospd1 | 0.292182441 | 0.310038957 |
| ENSMUSG0000006822 | chr15 | Lgals1 | 0.47267084 | 0.310038957 |
| ENSMUSG0000003770 | chr7 | Cd81 | 0.208578869 | 0.31018151 |
| ENSMUSG0000007894 | chr13 | Ak6 | 0.344160823 | 0.31018151 |
| ENSMUSG0000002991 | chr6 | Mad2l1 | -0.588010043 | 0.310249536 |
| ENSMUSG0000002237 | chr15 | Mrpl13 | 0.359275504 | 0.310319411 |
| ENSMUSG0000002683 | chr2 | Cytip | -0.943361881 | 0.310465413 |
| ENSMUSG0000007052 | chr7 | Peg12 | -1.032212765 | 0.310465413 |
| ENSMUSG0000004407 | chr11 | Eml6 | -0.335406778 | 0.310465413 |
| ENSMUSG0000004558 | chr4 | Frrs1l | -0.382148098 | 0.31087407 |
| ENSMUSG0000000564 | chr3 | Insrr | -1.848029135 | 0.310940238 |
| ENSMUSG0000002125 | chr12 | Angel1 | -0.373314711 | 0.311071816 |
| ENSMUSG0000004366 | chr8 | Tox3 | -0.738674601 | 0.31109431 |
| ENSMUSG0000003969 | chr19 | Batf2 | -0.95909639 | 0.311476661 |
| ENSMUSG0000002160 | chr13 | Ndufs6 | 0.307243595 | 0.311476661 |
| ENSMUSG0000002052 | chr11 | Myo19 | -0.548971548 | 0.31159676 |
| ENSMUSG0000002459 | chr18 | Megf10 | -0.363143657 | 0.31159676 |
| ENSMUSG0000002880 | chr4 | Stpg1 | -0.979643915 | 0.312120396 |
| ENSMUSG0000003991 | chr2 | Pak5 | -0.447098055 | 0.31251864 |
| ENSMUSG0000002875 | chr4 | Ddost | 0.204442145 | 0.31251864 |
| ENSMUSG0000000489 | chr3 | Hdgf | 0.22494826 | 0.31251864 |
| ENSMUSG0000004021 | chr7 | Emp3 | 0.296961266 | 0.314549849 |
| ENSMUSG0000004325 | chr4 | Tmem64 | -0.196544914 | 0.314633311 |
| ENSMUSG0000002886 | chr4 | Meaf6 | -0.179031092 | 0.314633311 |
| ENSMUSG0000002879 | chr4 | Eif3i | 0.403946011 | 0.314664683 |
| ENSMUSG0000002127 | chr12 | Cinp | 0.279257855 | 0.315238762 |
| ENSMUSG0000007343 | chr17 | Arhgdig | -0.562631531 | 0.315622149 |
| ENSMUSG0000007000 | chr8 | Fcho1 | -0.540447839 | 0.316159139 |
| ENSMUSG0000003787 | chr1 | Ackr1 | -0.4075814 | 0.316159139 |
| ENSMUSG0000002007 | chr10 | Ddx21 | 0.2558943 | 0.316159139 |
| ENSMUSG0000004123 | chr3 | Pklr | -2.424569888 | 0.316388444 |
| ENSMUSG0000002902 | chr4 | Mfn2 | 0.19417111 | 0.316388444 |
| ENSMUSG0000000037 | chr11 | Ccm2 | 0.37864378 | 0.316569418 |
| ENSMUSG0000010340 | chr9 | Lsmem2 | 0.694427348 | 0.316569418 |
| ENSMUSG0000001992 | chr10 | Ube2d1 | 0.26723704 | 0.316650448 |
| ENSMUSG0000004833 | chr3 | Lhfpl6 | 0.290898601 | 0.31672384 |
| ENSMUSG0000002970 | chr5 | Cux1 | 0.274740832 | 0.316850061 |
| ENSMUSG0000005675 | chr10 | Hmga2 | -2.018008697 | 0.316901735 |
| ENSMUSG0000003287 | chr11 | Ccdc85a | -0.568109405 | 0.316901735 |

| | | | | |
|---|---|---|---|---|
| ENSMUSG0000002418 | chr17 | Rgs11 | -0.33545597 | 0.316901735 |
| ENSMUSG0000003396 | chr18 | Jcad | 0.28709613 | 0.316901735 |
| ENSMUSG0000000044 | chr6 | Pparg | -0.467423814 | 0.317256729 |
| ENSMUSG0000005479 | chr9 | Klhl18 | 0.261367938 | 0.317256729 |
| ENSMUSG0000003878 | chr17 | Stap2 | -0.775652096 | 0.317592143 |
| ENSMUSG0000002835 | chr1 | Fmn2 | -0.381414676 | 0.317844753 |
| ENSMUSG0000006861 | chr16 | Clxn | -0.381780781 | 0.318136771 |
| ENSMUSG0000007186 | chr11 | Ppia | 0.343016267 | 0.318136771 |
| ENSMUSG0000002224 | chr15 | Brix1 | 0.287828942 | 0.318216517 |
| ENSMUSG0000003762 | chr3 | Cldn11 | -0.737670259 | 0.3185204 |
| ENSMUSG0000000152 | chr17 | Tubb5 | 0.230322679 | 0.3185204 |
| ENSMUSG0000005770 | chr7 | Mex3b | -0.361417636 | 0.318995119 |
| ENSMUSG0000003369 | chr15 | Arhgap39 | -0.256033836 | 0.318995119 |
| ENSMUSG0000002124 | chr12 | Isca2 | 0.254149892 | 0.319295869 |
| ENSMUSG0000002652 | chr1 | Fh1 | 0.297965197 | 0.319295869 |
| ENSMUSG0000004621 | chr11 | Rprml | -0.843674619 | 0.319446607 |
| ENSMUSG0000003138 | chrX | Dusp9 | -2.518463411 | 0.319446607 |
| ENSMUSG0000002593 | chr1 | Msc | -1.264688002 | 0.319446607 |
| ENSMUSG0000002168 | chr13 | Scamp1 | -0.174249805 | 0.319446607 |
| ENSMUSG0000001642 | chrX | Ndufa1 | 0.3963853 | 0.319599303 |
| ENSMUSG0000003318 | chr6 | Kbtbd12 | 0.464831233 | 0.319618773 |
| ENSMUSG0000002493 | chr19 | Slc1a1 | 0.834680851 | 0.31978825 |
| ENSMUSG0000004159 | chr11 | Cdc42ep4 | -0.201267214 | 0.320751661 |
| ENSMUSG0000003857 | chr4 | Susd1 | -0.391518935 | 0.320893945 |
| ENSMUSG0000000110 | chr11 | Poldip2 | 0.171610397 | 0.32138326 |
| ENSMUSG0000002924 | chr5 | Paics | 0.23702985 | 0.32138326 |
| ENSMUSG0000004121 | chr4 | Clvs1 | -0.539190982 | 0.322011555 |
| ENSMUSG0000003033 | chr6 | Ltbr | 0.175265173 | 0.322011555 |
| ENSMUSG0000004354 | chr6 | Dnai7 | -0.560733543 | 0.32207466 |
| ENSMUSG0000003458 | chr11 | Hid1 | -0.344727358 | 0.322100227 |
| ENSMUSG0000007172 | chrX | Gspt2 | -0.282711396 | 0.322100227 |
| ENSMUSG0000002875 | chr4 | Pla2g2c | -2.758573691 | 0.32237967 |
| ENSMUSG0000003524 | chr10 | Oaz1 | 0.410976711 | 0.322709413 |
| ENSMUSG0000003206 | chr9 | Plet1 | -1.015894154 | 0.322979272 |
| ENSMUSG0000005438 | chr7 | Ceacam2 | -0.966661012 | 0.323383928 |
| ENSMUSG0000004355 | chr17 | Mdga1 | -0.32587606 | 0.323432057 |
| ENSMUSG0000002763 | chr2 | Ndrg3 | -0.216287995 | 0.323432057 |
| ENSMUSG0000000346 | chr1 | Pex19 | 0.196357331 | 0.323432057 |
| ENSMUSG0000003588 | chr19 | Cox8a | 0.312500442 | 0.323635877 |
| ENSMUSG0000003641 | chr18 | Arsi | 0.528377858 | 0.323911021 |
| ENSMUSG0000004493 | chr15 | Sstr3 | -0.915653245 | 0.324256179 |
| ENSMUSG0000004345 | chr7 | Zfp536 | -0.34116961 | 0.324256179 |
| ENSMUSG0000005694 | chr3 | Mab21l1 | -0.355434399 | 0.324256179 |
| ENSMUSG0000003393 | chr8 | Ndufb7 | 0.38025331 | 0.324256179 |
| ENSMUSG0000005496 | chr15 | Zfp647 | 0.340485229 | 0.324415778 |
| ENSMUSG0000002201 | chr14 | Enox1 | 0.400958428 | 0.324709906 |
| ENSMUSG0000003143 | chrX | Tsc22d3 | 0.553094607 | 0.324773953 |
| ENSMUSG0000002428 | chr18 | Thoc1 | 0.181000485 | 0.325016291 |
| ENSMUSG0000003198 | chr8 | 2810004N23Rik | 0.248401823 | 0.325016291 |
| ENSMUSG0000003805 | chr18 | Smim3 | 0.381875164 | 0.325607744 |

| | | | | |
|---|---|---|---|---|
| ENSMUSG0000003563 | chr4 | Grhpr | 0.35341749 | 0.326240853 |
| ENSMUSG0000005979 | chr11 | Eif4a1 | 0.207606061 | 0.326462824 |
| ENSMUSG0000006369 | chr6 | Cycs | 0.347021945 | 0.326462824 |
| ENSMUSG0000003981 | chr4 | Tbc1d2 | -0.791997285 | 0.327282897 |
| ENSMUSG0000003869 | chr5 | Atp5mf | 0.419233707 | 0.327299005 |
| ENSMUSG0000010308 | chr18 | Pcdhgb6 | -0.296623596 | 0.327379815 |
| ENSMUSG0000004929 | chr11 | Trappc1 | 0.183017916 | 0.327379815 |
| ENSMUSG0000002747 | chr2 | Mapre1 | 0.254345142 | 0.327379815 |
| ENSMUSG0000003537 | chr19 | 1810055G02Rik | 0.34291637 | 0.327379815 |
| ENSMUSG0000003904 | chr9 | Foxred1 | 0.232877246 | 0.327468226 |
| ENSMUSG0000007042 | chr5 | Zscan25 | 0.302745543 | 0.32811003 |
| ENSMUSG0000002548 | chr7 | Psmd13 | 0.305712435 | 0.32811003 |
| ENSMUSG0000002870 | chr4 | Pomgnt1 | 0.200336084 | 0.328387271 |
| ENSMUSG0000004838 | chr2 | Fam171b | -0.347856262 | 0.328655127 |
| ENSMUSG0000003176 | chr8 | Mt3 | -0.406837504 | 0.328933017 |
| ENSMUSG0000003824 | chr10 | Sobp | 0.262566494 | 0.329317003 |
| ENSMUSG0000005493 | chr10 | Kcnmb4 | 1.535665768 | 0.329732511 |
| ENSMUSG0000006784 | chr2 | Romo1 | 0.431173854 | 0.329732511 |
| ENSMUSG0000005712 | chr3 | Gja5 | 1.022881902 | 0.329782946 |
| ENSMUSG0000002916 | chr5 | Cgref1 | -0.491425556 | 0.329857562 |
| ENSMUSG0000004192 | chr18 | Nol4 | -0.582485695 | 0.329857562 |
| ENSMUSG0000000382 | chr8 | Syce2 | -0.726826182 | 0.330040135 |
| ENSMUSG0000002526 | chrX | Gnl3l | -0.209258154 | 0.330642369 |
| ENSMUSG0000002152 | chr13 | Uqcrb | 0.31765217 | 0.330642369 |
| ENSMUSG0000001588 | chr5 | Lcorl | -0.403945346 | 0.330912572 |
| ENSMUSG0000001066 | chr9 | Plcd1 | 0.201678426 | 0.330912572 |
| ENSMUSG0000006145 | chr12 | Nol10 | 0.230320935 | 0.330912572 |
| ENSMUSG0000002761 | chr2 | Gss | -0.323470676 | 0.33126911 |
| ENSMUSG0000009799 | chr1 | Ptprv | -0.906600155 | 0.331482816 |
| ENSMUSG0000003904 | chr2 | Usp6nl | 0.221428644 | 0.332053417 |
| ENSMUSG0000002480 | chr19 | Ankrd1 | 0.970182651 | 0.332053417 |
| ENSMUSG0000002815 | chr3 | Rorc | -0.512172718 | 0.33210955 |
| ENSMUSG0000003193 | chr9 | 4931406C07Rik | -0.366829305 | 0.33210955 |
| ENSMUSG0000000453 | chr5 | Coro1c | 0.257994496 | 0.33210955 |
| ENSMUSG0000010021 | chr13 | H3c7 | 0.971059884 | 0.33210955 |
| ENSMUSG0000001634 | chr2 | Kcnq2 | -0.366729005 | 0.332127933 |
| ENSMUSG0000003459 | chr9 | Myo5a | -0.328052102 | 0.332127933 |
| ENSMUSG0000002827 | chr4 | Rragd | -0.228113452 | 0.332127933 |
| ENSMUSG0000001716 | chr11 | Cntnap1 | -0.21385087 | 0.332127933 |
| ENSMUSG0000002179 | chr14 | Glud1 | 0.134748583 | 0.332127933 |
| ENSMUSG0000001450 | chr18 | Srp19 | 0.247646255 | 0.332127933 |
| ENSMUSG0000003782 | chr2 | Tgm2 | 0.33129408 | 0.332127933 |
| ENSMUSG0000005572 | chr7 | Rras2 | 0.410632289 | 0.332470179 |
| ENSMUSG0000005659 | chr11 | Drc3 | -0.463175613 | 0.332563147 |
| ENSMUSG0000003060 | chr7 | Pak4 | -0.307841186 | 0.332563147 |
| ENSMUSG0000006251 | chr6 | Zfp398 | -0.237246011 | 0.332563147 |
| ENSMUSG0000000200 | chrX | Pdzd4 | -0.226326646 | 0.332563147 |
| ENSMUSG0000002596 | chr1 | Ndufs1 | 0.186582005 | 0.332563147 |
| ENSMUSG0000002456 | chr18 | Cxxc1 | 0.252769475 | 0.332563147 |
| ENSMUSG0000006301 | chr2 | Manbal | 0.255683136 | 0.332563147 |

| | | | | |
|---|---|---|---|---|
| ENSMUSG0000007131 | chr10 | Bves | 0.296348999 | 0.332563147 |
| ENSMUSG0000006388 | chr17 | Rpl7l1 | 0.309711908 | 0.332563147 |
| ENSMUSG0000007422 | chr7 | Zfp568 | 0.296658163 | 0.332563147 |
| ENSMUSG0000002458 | chr18 | Nedd4l | 0.357916125 | 0.333000734 |
| ENSMUSG0000005301 | chr6 | Krcc1 | 0.241059411 | 0.33325218 |
| ENSMUSG0000003139 | chrX | L1cam | -0.37464454 | 0.333280714 |
| ENSMUSG0000003807 | chr5 | Galnt11 | 0.169761723 | 0.333280714 |
| ENSMUSG0000003169 | chr8 | Vps35 | 0.212206101 | 0.333280714 |
| ENSMUSG0000005091 | chr9 | Tmem123 | 0.220424113 | 0.333280714 |
| ENSMUSG0000002390 | chr17 | Hcfc1r1 | 0.231135036 | 0.333332099 |
| ENSMUSG0000003137 | chrX | Haus7 | 0.339799748 | 0.333332099 |
| ENSMUSG0000000605 | chr11 | Atp5mc1 | 0.338721698 | 0.333332099 |
| ENSMUSG0000007037 | chr3 | Capza1 | 0.177725253 | 0.333681104 |
| ENSMUSG0000000014 | chr5 | Brat1 | 0.210387934 | 0.333800233 |
| ENSMUSG0000009621 | chr16 | Smim22 | -1.801060321 | 0.334244212 |
| ENSMUSG0000009778 | chr7 | Ctxnd1 | -0.423697581 | 0.335172234 |
| ENSMUSG0000007291 | chr11 | Gm12258 | -0.291960664 | 0.335486503 |
| ENSMUSG0000004371 | chr9 | Col6a6 | -0.735273611 | 0.335963274 |
| ENSMUSG0000004917 | chrX | Frmpd4 | -0.385076191 | 0.335963274 |
| ENSMUSG0000000237 | chr17 | Ndufa11 | 0.41637031 | 0.336094444 |
| ENSMUSG0000000056 | chr3 | Atp5pb | 0.304987591 | 0.336172308 |
| ENSMUSG0000003116 | chrX | Was | -0.72218016 | 0.33666315 |
| ENSMUSG0000002413 | chr17 | Eci1 | 0.33816027 | 0.336678695 |
| ENSMUSG0000002414 | chr17 | Epas1 | 0.339908218 | 0.336678695 |
| ENSMUSG0000002010 | chr10 | Ddit4 | 0.371156037 | 0.336678695 |
| ENSMUSG0000000328 | chr2 | Hck | -0.573026049 | 0.337555397 |
| ENSMUSG0000003948 | chr10 | Tspyl4 | -0.258028101 | 0.337555397 |
| ENSMUSG0000002931 | chr5 | Klhl8 | 0.27196079 | 0.337555397 |
| ENSMUSG0000002572 | chr7 | Sec11a | 0.327653148 | 0.337555397 |
| ENSMUSG0000006365 | chr7 | Cnfn | 1.255085261 | 0.337555397 |
| ENSMUSG0000004673 | chr6 | Gprc5a | -0.831480181 | 0.3377752 |
| ENSMUSG0000007895 | chr15 | Arhgap8 | -0.5836305 | 0.338220113 |
| ENSMUSG0000004012 | chr6 | Rep15 | -0.881679921 | 0.338276708 |
| ENSMUSG0000002275 | chr16 | Tmem45a | 0.382641726 | 0.338276708 |
| ENSMUSG0000004773 | chr19 | Wbp1l | 0.202024696 | 0.338313036 |
| ENSMUSG0000003969 | chr4 | Msantd3 | 0.282326896 | 0.338313036 |
| ENSMUSG0000003183 | chr8 | Hsbp1 | 0.309641905 | 0.338313036 |
| ENSMUSG0000002214 | chr15 | Osmr | 0.239297375 | 0.338371428 |
| ENSMUSG0000002751 | chr2 | Pck1 | -1.683298052 | 0.338882865 |
| ENSMUSG0000005445 | chrX | Sytl5 | -0.647382932 | 0.338882865 |
| ENSMUSG0000003712 | chr19 | Psd | 0.204094686 | 0.338882865 |
| ENSMUSG0000003528 | chr19 | Adrb1 | 0.308230079 | 0.338882865 |
| ENSMUSG0000002006 | chr10 | Mybpc1 | 1.344045731 | 0.338882865 |
| ENSMUSG0000002960 | chr5 | Oas1b | -0.618139634 | 0.339400929 |
| ENSMUSG0000003217 | chr9 | Pde4a | -0.385481413 | 0.339400929 |
| ENSMUSG0000003096 | chr7 | Trim21 | -0.271867261 | 0.339400929 |
| ENSMUSG0000003489 | chr9 | Filip1 | 0.274007581 | 0.339462961 |
| ENSMUSG0000003030 | chr6 | Caprin2 | -0.43259991 | 0.339611891 |
| ENSMUSG0000003251 | chr9 | Ttc21a | -0.534113309 | 0.339611891 |
| ENSMUSG0000005593 | chr8 | Fto | 0.157019128 | 0.339611891 |

| | | | | |
|---|---|---|---|---|
| ENSMUSG0000002963 | chr5 | Rnf6 | 0.190717713 | 0.339611891 |
| ENSMUSG0000004701 | chr6 | Fbxo41 | -0.347490789 | 0.340260476 |
| ENSMUSG0000006785 | chr1 | Arfgef1 | -0.227317411 | 0.340260476 |
| ENSMUSG0000002089 | chr11 | Vamp2 | -0.209217164 | 0.340260476 |
| ENSMUSG0000000354 | chr17 | Klc4 | -0.338020225 | 0.340694191 |
| ENSMUSG0000010269 | chr18 | Pcdhac2 | -0.530406289 | 0.340905054 |
| ENSMUSG0000002263 | chr16 | Alcam | -0.334448238 | 0.340905054 |
| ENSMUSG0000002231 | chr15 | Eif3h | 0.338356617 | 0.340905054 |
| ENSMUSG0000002821 | chr4 | Gem | 0.41187483 | 0.340905054 |
| ENSMUSG0000001932 | chr17 | Noxo1 | -1.210454585 | 0.341192021 |
| ENSMUSG0000004531 | chr2 | Proser2 | 0.266317229 | 0.341440072 |
| ENSMUSG0000003731 | chr8 | Bag4 | -0.290778857 | 0.341444184 |
| ENSMUSG0000003973 | chr16 | Slx4 | 0.251440874 | 0.341444184 |
| ENSMUSG0000002837 | chr4 | Astn2 | 0.304601113 | 0.341444184 |
| ENSMUSG0000002579 | chr7 | Slco3a1 | 0.28496521 | 0.341772337 |
| ENSMUSG0000002245 | chr15 | Ndufa6 | 0.303237897 | 0.341837964 |
| ENSMUSG0000002106 | chr12 | Sav1 | 0.219023512 | 0.342209171 |
| ENSMUSG0000000551 | chr2 | Ndufs3 | 0.300289299 | 0.342314215 |
| ENSMUSG0000002013 | chr10 | Apc2 | -0.34996414 | 0.3423245 |
| ENSMUSG0000000142 | chr6 | Snd1 | 0.20594284 | 0.3423245 |
| ENSMUSG0000001905 | chr4 | Plod1 | 0.211144758 | 0.3423245 |
| ENSMUSG0000006276 | chr15 | Shisal1 | 0.25987642 | 0.3423245 |
| ENSMUSG0000004590 | chr19 | Npas4 | 0.46381101 | 0.3423245 |
| ENSMUSG0000003568 | chr7 | Thrsp | 0.778872535 | 0.3423245 |
| ENSMUSG0000004036 | chr5 | Lrrd1 | -1.523677626 | 0.342632039 |
| ENSMUSG0000004992 | chrX | Lpar4 | -0.421126572 | 0.342632039 |
| ENSMUSG0000009012 | chrX | Kcne5 | -0.721386984 | 0.342653576 |
| ENSMUSG0000001477 | chr8 | Nol3 | -0.353420815 | 0.342653576 |
| ENSMUSG0000002676 | chr2 | Mindy3 | 0.227949401 | 0.342653576 |
| ENSMUSG0000002311 | chr14 | Prmt5 | 0.202153325 | 0.342670924 |
| ENSMUSG0000007857 | chr11 | Ndufaf8 | 0.531121836 | 0.342670924 |
| ENSMUSG0000006329 | chr7 | Luzp2 | -0.434540017 | 0.342748064 |
| ENSMUSG0000003230 | chr9 | Chrna3 | -0.269664331 | 0.342748064 |
| ENSMUSG0000000263 | chr7 | Pdcd2l | 0.304437707 | 0.342985135 |
| ENSMUSG0000005626 | chr1 | Dennd1b | -0.373161036 | 0.342986128 |
| ENSMUSG0000002308 | chr6 | Ndufa5 | 0.295499307 | 0.342986128 |
| ENSMUSG0000002852 | chr4 | Dnajc6 | -0.346802814 | 0.34404551 |
| ENSMUSG0000003734 | chrX | Slc9a7 | -0.455978527 | 0.34404551 |
| ENSMUSG0000005010 | chr11 | Haspin | -1.322471957 | 0.34404551 |
| ENSMUSG0000009669 | chr17 | Zfp960 | 0.393947933 | 0.34404551 |
| ENSMUSG0000003465 | chr8 | Cacna1a | -0.287422135 | 0.34423525 |
| ENSMUSG0000002142 | chr13 | Fars2 | 0.278812015 | 0.344266541 |
| ENSMUSG0000001175 | chr19 | Pgam1 | 0.303361282 | 0.344266541 |
| ENSMUSG0000000039 | chr6 | Ndufa9 | 0.280619008 | 0.34455739 |
| ENSMUSG0000006736 | chrX | Trmt2b | 0.226590414 | 0.344561473 |
| ENSMUSG0000003553 | chr11 | Eif1 | 0.365993903 | 0.344610095 |
| ENSMUSG0000007994 | chr13 | Cox5b-ps | 0.644526748 | 0.345297591 |
| ENSMUSG0000004950 | chr16 | Dtx3l | -0.379345199 | 0.345502728 |
| ENSMUSG0000001564 | chr2 | Lama5 | 0.255882259 | 0.345502728 |
| ENSMUSG0000002129 | chr12 | Atp5mj | 0.511050834 | 0.345502728 |

| | | | | |
|---|---|---|---|---|
| ENSMUSG0000002854 | chr4 | Slc5a9 | -2.355672321 | 0.345525993 |
| ENSMUSG0000001985 | chr10 | Hebp2 | 0.507611246 | 0.345642755 |
| ENSMUSG0000004605 | chr7 | Sbsn | 0.473910777 | 0.345855394 |
| ENSMUSG0000005705 | chr11 | Skap1 | -0.523372388 | 0.346536121 |
| ENSMUSG0000003415 | chr1 | Mab21l4 | -1.647028599 | 0.347167211 |
| ENSMUSG0000005111 | chr13 | Sv2c | -0.427041247 | 0.347397972 |
| ENSMUSG0000007465 | chr10 | Kif5a | -0.355692051 | 0.347397972 |
| ENSMUSG0000002913 | chr5 | Rbks | -0.588743862 | 0.347397972 |
| ENSMUSG0000002822 | chr4 | Cpne3 | -0.174109346 | 0.347397972 |
| ENSMUSG0000007950 | chrX | Zfx | 0.151281749 | 0.347397972 |
| ENSMUSG0000003627 | chr11 | 9530068E07Rik | 0.274122319 | 0.347397972 |
| ENSMUSG0000007417 | chr7 | Plekhf1 | 0.464672034 | 0.347397972 |
| ENSMUSG0000002430 | chr17 | Pfdn6 | 0.37735785 | 0.347627395 |
| ENSMUSG0000003851 | chr10 | Rpf2 | 0.286362676 | 0.34804 |
| ENSMUSG0000002404 | chr17 | Myl12a | 0.38774962 | 0.34804 |
| ENSMUSG0000002884 | chr4 | Sh3bgrl3 | -0.290770547 | 0.348302774 |
| ENSMUSG0000003587 | chr10 | Pawr | 0.365062532 | 0.348302774 |
| ENSMUSG0000000337 | chr7 | Cd79a | -1.776705038 | 0.348496251 |
| ENSMUSG0000003461 | chr13 | Mtrr | 0.272098179 | 0.348561262 |
| ENSMUSG0000005116 | chr12 | Eml5 | -0.372886822 | 0.348736834 |
| ENSMUSG0000003232 | chr9 | Pstpip1 | 0.434007034 | 0.348736834 |
| ENSMUSG0000005059 | chr2 | Fam78a | -1.110997138 | 0.349035575 |
| ENSMUSG0000003041 | chr7 | Pdcd5 | 0.360332823 | 0.349035575 |
| ENSMUSG0000006837 | chr2 | D430041D05Rik | 0.908698631 | 0.349035575 |
| ENSMUSG0000002141 | chr13 | Pxdc1 | 0.421279482 | 0.349160729 |
| ENSMUSG0000002608 | chr1 | Mrpl30 | 0.315273316 | 0.349220452 |
| ENSMUSG0000000196 | chrX | Emd | 0.392788825 | 0.349382237 |
| ENSMUSG0000005512 | chr14 | Cgrrf1 | 0.309650949 | 0.349389786 |
| ENSMUSG0000006220 | chr1 | Erbb4 | -1.546560347 | 0.349477619 |
| ENSMUSG0000000053 | chr15 | Tamalin | -0.429163243 | 0.349477619 |
| ENSMUSG0000004838 | chr15 | Scrt1 | -0.545007303 | 0.349677736 |
| ENSMUSG0000004136 | chr19 | Shtn1 | -0.361140379 | 0.350196243 |
| ENSMUSG0000003132 | chrX | Rps4x | 0.319906053 | 0.350196243 |
| ENSMUSG0000003223 | chr9 | Rora | 0.334898831 | 0.350196243 |
| ENSMUSG0000002421 | chr17 | Taf11 | 0.265731599 | 0.350588146 |
| ENSMUSG0000000986 | chr4 | Sdhb | 0.348786533 | 0.350588146 |
| ENSMUSG0000003361 | chr16 | Map3k13 | -0.416615256 | 0.350596656 |
| ENSMUSG0000005685 | chr7 | Jakmip3 | -0.294908255 | 0.350596656 |
| ENSMUSG0000002257 | chr15 | Gpihbp1 | 0.480350005 | 0.351261838 |
| ENSMUSG0000003873 | chr7 | Shank1 | -0.285850475 | 0.351847897 |
| ENSMUSG0000004955 | chr5 | Clip1 | 0.377005759 | 0.35239421 |
| ENSMUSG0000002862 | chr4 | Col9a2 | -0.392017426 | 0.352455151 |
| ENSMUSG0000000510 | chr5 | Wdr1 | 0.332055809 | 0.352500856 |
| ENSMUSG0000003178 | chr8 | Drc7 | -0.638274342 | 0.352534961 |
| ENSMUSG0000007164 | chr19 | Tut1 | 0.221750006 | 0.352534961 |
| ENSMUSG0000006803 | chr17 | Afdn | -0.207526099 | 0.352585507 |
| ENSMUSG0000006244 | chr7 | Ap3b2 | -0.361042246 | 0.352767728 |
| ENSMUSG0000000562 | chr3 | Psmd4 | 0.290565385 | 0.352767728 |
| ENSMUSG0000002913 | chr5 | Plb1 | 0.377603097 | 0.35279374 |
| ENSMUSG0000004991 | chr16 | 2610318N02Rik | -2.861836293 | 0.353189981 |

| | | | | |
|---|---|---|---|---|
| ENSMUSG0000002626 | chr1 | Rnpepl1 | 0.173757051 | 0.353189981 |
| ENSMUSG0000004267 | chr2 | Immp1l | 0.464317593 | 0.353189981 |
| ENSMUSG0000003676 | chr10 | Dnajc12 | -0.316150463 | 0.353480997 |
| ENSMUSG0000002826 | chr3 | Lmo4 | 0.222776375 | 0.353968483 |
| ENSMUSG0000003730 | chr4 | Man1c1 | 0.227077355 | 0.353968483 |
| ENSMUSG0000003441 | chr11 | Tbc1d10a | 0.301070576 | 0.353968483 |
| ENSMUSG0000004772 | chr7 | Bola2 | 0.436606166 | 0.353968483 |
| ENSMUSG0000002968 | chr5 | Bcl7b | 0.245165688 | 0.354047898 |
| ENSMUSG0000004282 | chr11 | Fgf11 | -0.337814642 | 0.354464121 |
| ENSMUSG0000004025 | chr10 | Nxph4 | -0.536799078 | 0.354464121 |
| ENSMUSG0000010471 | chr5 | Gbp6 | -1.176584525 | 0.354466862 |
| ENSMUSG0000002068 | chr11 | Rasl10b | -0.430892502 | 0.354466862 |
| ENSMUSG0000004398 | chr9 | Cep164 | -0.329437766 | 0.354466862 |
| ENSMUSG0000004447 | chr17 | Zfand3 | 0.184061606 | 0.354466862 |
| ENSMUSG0000002630 | chr1 | Lrrfip1 | 0.229694824 | 0.354466862 |
| ENSMUSG0000010404 | chr3 | Gm6525 | -1.220393069 | 0.354647255 |
| ENSMUSG0000002582 | chr5 | Iscu | 0.318020395 | 0.354735356 |
| ENSMUSG0000002719 | chr2 | Ext2 | 0.144435845 | 0.354790195 |
| ENSMUSG0000002620 | chr1 | Tuba4a | 0.251013599 | 0.354898864 |
| ENSMUSG0000003212 | chr9 | Tmem218 | 0.290607042 | 0.354898864 |
| ENSMUSG0000007110 | chr8 | Cfap96 | -0.378893291 | 0.355602336 |
| ENSMUSG0000003960 | chr17 | Rcan2 | 0.25374587 | 0.355851667 |
| ENSMUSG0000002253 | chr16 | Lsg1 | 0.280384775 | 0.355851667 |
| ENSMUSG0000002667 | chr1 | Uap1 | -0.28704383 | 0.356232303 |
| ENSMUSG0000004173 | chr19 | Pgm5 | 0.287109261 | 0.356232303 |
| ENSMUSG0000003061 | chr7 | Mrps11 | 0.311787143 | 0.356232303 |
| ENSMUSG0000002532 | chr7 | Atp10a | 0.302416104 | 0.356232303 |
| ENSMUSG0000000901 | chr5 | Dynll1 | 0.482843204 | 0.356232303 |
| ENSMUSG0000005431 | chr3 | Mrps21 | 0.477438495 | 0.356320074 |
| ENSMUSG0000005085 | chr5 | Atp5me | 0.428768449 | 0.356320074 |
| ENSMUSG0000004262 | chr12 | Zfyve1 | 0.167011859 | 0.356788059 |
| ENSMUSG0000003686 | chr11 | Mrpl55 | 0.35616896 | 0.356788059 |
| ENSMUSG0000002737 | chr2 | Prom2 | -1.299083901 | 0.356925421 |
| ENSMUSG0000006153 | chr9 | Sec22c | -0.233075402 | 0.357104316 |
| ENSMUSG0000001431 | chr15 | Cox6c | 0.368357907 | 0.357104316 |
| ENSMUSG0000002106 | chr12 | Rab15 | -0.330072028 | 0.357315825 |
| ENSMUSG0000007379 | chr4 | Alg6 | -0.320950673 | 0.357315825 |
| ENSMUSG0000005137 | chr18 | Pcdh1 | -0.355056525 | 0.358143216 |
| ENSMUSG0000002956 | chr5 | Oasl2 | -0.567353078 | 0.358246341 |
| ENSMUSG0000002887 | chr4 | Epha10 | -0.973146607 | 0.358246341 |
| ENSMUSG0000002536 | chr10 | Esyt1 | 0.182660965 | 0.358246341 |
| ENSMUSG0000002032 | chr11 | Mdh1 | 0.281565263 | 0.358246341 |
| ENSMUSG0000002846 | chr4 | Creb3 | 0.284737301 | 0.358246341 |
| ENSMUSG0000002425 | chr18 | Polr2d | 0.312874724 | 0.358246341 |
| ENSMUSG0000005054 | chr11 | Adra1b | 0.993686963 | 0.358246341 |
| ENSMUSG0000002955 | chr5 | Mad1l1 | 0.250382567 | 0.358272729 |
| ENSMUSG0000002740 | chr2 | Snrpb | 0.362090821 | 0.358272729 |
| ENSMUSG0000002026 | chr10 | Adarb1 | 0.366718565 | 0.358744227 |
| ENSMUSG0000007351 | chr18 | Dok6 | -0.403442643 | 0.359077693 |
| ENSMUSG0000003702 | chr2 | Foxa2 | -1.96819432 | 0.359187198 |

| | | | | |
|---|---|---|---|---|
| ENSMUSG0000004248 | chr14 | Mustn1 | 0.449139181 | 0.359247792 |
| ENSMUSG0000004736 | chr1 | Gm973 | 0.803300849 | 0.360266841 |
| ENSMUSG0000002610 | chr1 | Stat1 | -0.435000155 | 0.360676313 |
| ENSMUSG0000004802 | chr17 | Rgmb | -0.24373859 | 0.360676313 |
| ENSMUSG0000002460 | chr18 | Rps14 | 0.582932733 | 0.360864904 |
| ENSMUSG0000000434 | chr6 | Pde1c | -0.383544897 | 0.361257952 |
| ENSMUSG0000002876 | chr4 | Hspg2 | 0.316446322 | 0.361257952 |
| ENSMUSG0000000444 | chr6 | Bid | -0.526787006 | 0.361721497 |
| ENSMUSG0000002303 | chr15 | Cela1 | 0.537355078 | 0.361721497 |
| ENSMUSG0000005151 | chr15 | Rps19bp1 | 0.285390126 | 0.36177467 |
| ENSMUSG0000005033 | chr14 | Lgals3 | -0.900812426 | 0.362280703 |
| ENSMUSG0000003750 | chr1 | Arhgef4 | -0.40439066 | 0.362280703 |
| ENSMUSG0000007450 | chr9 | Zfp558 | -0.797729065 | 0.362280703 |
| ENSMUSG0000002915 | chr5 | Ociad2 | -0.345033324 | 0.362280703 |
| ENSMUSG0000003026 | chr6 | Kras | 0.223677999 | 0.362280703 |
| ENSMUSG0000000307 | chr10 | Atp5f1d | 0.314101666 | 0.362280703 |
| ENSMUSG0000000074 | chr8 | Rpl13 | 0.387211685 | 0.362280703 |
| ENSMUSG0000000196 | chrX | Fam50a | 0.340461263 | 0.362280703 |
| ENSMUSG0000000494 | chr17 | Tmem242 | 0.381403702 | 0.362280703 |
| ENSMUSG0000009792 | chr12 | Tunar | 1.39832002 | 0.362280703 |
| ENSMUSG0000000291 | chr8 | Arrdc2 | 0.324981173 | 0.363120314 |
| ENSMUSG0000005471 | chr7 | Zfp771 | 0.312436541 | 0.363120314 |
| ENSMUSG0000005953 | chr11 | Uqcr10 | 0.456125848 | 0.363120314 |
| ENSMUSG0000005339 | chr7 | Cacng8 | -2.022634504 | 0.363568214 |
| ENSMUSG0000001769 | chr11 | Rhbdl3 | -0.298840495 | 0.363568214 |
| ENSMUSG0000003624 | chr4 | Ube2r2 | 0.208226298 | 0.363568214 |
| ENSMUSG0000000206 | chr11 | Sdf2 | 0.276568791 | 0.363568214 |
| ENSMUSG0000004489 | chr11 | Uqcrq | 0.474950214 | 0.363568214 |
| ENSMUSG0000002683 | chr2 | Olfm1 | 0.205475115 | 0.363716629 |
| ENSMUSG0000003109 | chr7 | Tnni2 | 0.766621602 | 0.363755618 |
| ENSMUSG0000002083 | chr11 | Tmigd1 | -2.273423944 | 0.364316208 |
| ENSMUSG0000000060 | chr7 | Clcn4 | -0.157552733 | 0.364368708 |
| ENSMUSG0000002517 | chr11 | Fn3k | -0.456198196 | 0.364918571 |
| ENSMUSG0000005151 | chr7 | Fam181b | -0.626769608 | 0.364918571 |
| ENSMUSG0000003568 | chr4 | Melk | -1.164612083 | 0.364957974 |
| ENSMUSG0000003133 | chrX | Abcb7 | -0.177145863 | 0.364957974 |
| ENSMUSG0000004462 | chr2 | Swi5 | 0.312915934 | 0.364957974 |
| ENSMUSG0000000893 | chr4 | Slc1a7 | 0.54242131 | 0.364957974 |
| ENSMUSG0000005500 | chr6 | Lrtm2 | -1.302483315 | 0.36504383 |
| ENSMUSG0000002807 | chr3 | Cd1d1 | -0.474717548 | 0.36504383 |
| ENSMUSG0000000665 | chr7 | Aplp1 | -0.349523878 | 0.36504383 |
| ENSMUSG0000004624 | chr9 | Hepacam | -0.323178587 | 0.36504383 |
| ENSMUSG0000004670 | chr8 | Csnk2a2 | 0.145391709 | 0.36504383 |
| ENSMUSG0000003025 | chr6 | Rassf8 | 0.18072211 | 0.36504383 |
| ENSMUSG0000002183 | chr14 | Psmc6 | 0.206694257 | 0.36504383 |
| ENSMUSG0000005778 | chr17 | Bak1 | 0.217941192 | 0.36504383 |
| ENSMUSG0000002894 | chr5 | Rheb | 0.303601144 | 0.36504383 |
| ENSMUSG0000002831 | chr4 | Polr1e | 0.387430434 | 0.36504383 |
| ENSMUSG0000005907 | chr7 | Rpl18 | 0.42470271 | 0.36504383 |
| ENSMUSG0000000204 | chr7 | Trappc6a | 0.41090371 | 0.36504383 |

| | | | | |
|---|---|---|---|---|
| ENSMUSG0000003124 | chrX | Hmgn5 | -0.338474399 | 0.365493965 |
| ENSMUSG0000001574 | chr3 | Plekho1 | 0.278304036 | 0.365493965 |
| ENSMUSG0000004004 | chr17 | Ndufb10 | 0.399342241 | 0.365493965 |
| ENSMUSG0000002220 | chr14 | Dhrs2 | -1.294406923 | 0.365813573 |
| ENSMUSG0000000651 | chr8 | Cyba | 0.454993388 | 0.365813573 |
| ENSMUSG0000000456 | chr8 | Pnpla6 | -0.29216105 | 0.365860054 |
| ENSMUSG0000003165 | chr8 | Heatr3 | 0.180082654 | 0.365873191 |
| ENSMUSG0000003370 | chr1 | Acbd6 | 0.189094505 | 0.365873191 |
| ENSMUSG0000004389 | chr9 | S1pr2 | 0.277762136 | 0.365965947 |
| ENSMUSG0000002118 | chr12 | Rps6ka5 | 0.411057614 | 0.365965947 |
| ENSMUSG0000000121 | chr10 | Agpat3 | 0.20126799 | 0.366154046 |
| ENSMUSG0000007165 | chr19 | Ubxn1 | 0.234323667 | 0.366179169 |
| ENSMUSG0000002213 | chr14 | Mbnl2 | 0.19256998 | 0.366347387 |
| ENSMUSG0000002438 | chr18 | Ercc3 | 0.1713698 | 0.366347387 |
| ENSMUSG0000006185 | chr4 | Patj | -0.512817607 | 0.366628872 |
| ENSMUSG0000004530 | chr5 | Preb | 0.182347833 | 0.366692023 |
| ENSMUSG0000004508 | chr4 | Lingo2 | -0.645074222 | 0.366733693 |
| ENSMUSG0000003485 | chr5 | Cxcl10 | -1.040848997 | 0.366751484 |
| ENSMUSG0000002219 | chr14 | Slc22a17 | -0.282709302 | 0.366751484 |
| ENSMUSG0000003714 | chr12 | Ddx1 | 0.190792689 | 0.366751484 |
| ENSMUSG0000003180 | chr8 | Pgls | 0.295093373 | 0.366751484 |
| ENSMUSG0000003987 | chr2 | Neurl2 | 0.423637112 | 0.366751484 |
| ENSMUSG0000003051 | chr7 | Snrpa1 | 0.33476102 | 0.366751484 |
| ENSMUSG0000002895 | chr5 | Fastk | 0.174980187 | 0.366869753 |
| ENSMUSG0000003795 | chr12 | Wdr20 | 0.219880822 | 0.366869753 |
| ENSMUSG0000005542 | chr14 | Pcdh9 | -0.344611799 | 0.367028699 |
| ENSMUSG0000006357 | chr17 | Klhdc3 | 0.155989295 | 0.367034424 |
| ENSMUSG0000005831 | chr14 | Ube2e2 | 0.209143119 | 0.367034424 |
| ENSMUSG0000003356 | chr1 | Adgrb3 | -0.34216355 | 0.367322876 |
| ENSMUSG0000006619 | chr4 | Spag8 | -0.656196849 | 0.367322876 |
| ENSMUSG0000004704 | chrX | Tmem164 | -0.254845603 | 0.367322876 |
| ENSMUSG0000003577 | chr2 | Mrps2 | 0.1958195 | 0.367322876 |
| ENSMUSG0000004517 | chr1 | Amer3 | -0.35048762 | 0.367349134 |
| ENSMUSG0000003487 | chr10 | Gipc3 | 0.361467253 | 0.367349134 |
| ENSMUSG0000004184 | chr15 | Rpl37 | 0.413243072 | 0.367349134 |
| ENSMUSG0000004160 | chr1 | Inava | -0.704773015 | 0.367445491 |
| ENSMUSG0000000973 | chrX | Kcnd1 | -0.412779458 | 0.367561266 |
| ENSMUSG0000004272 | chr19 | Wdr74 | 0.356327924 | 0.367695657 |
| ENSMUSG0000003817 | chr3 | Pde4dip | 0.205919826 | 0.368245696 |
| ENSMUSG0000001284 | chr7 | Rps5 | 0.34725876 | 0.368374915 |
| ENSMUSG0000000561 | chr7 | Eif4g2 | 0.207288461 | 0.368474323 |
| ENSMUSG0000003314 | chr16 | Phldb2 | 0.27867112 | 0.368671201 |
| ENSMUSG0000001981 | chr10 | Ltv1 | 0.313155523 | 0.368671201 |
| ENSMUSG0000003270 | chr12 | Exd2 | -0.175334839 | 0.368705146 |
| ENSMUSG0000003250 | chr9 | Dclk3 | -0.353755925 | 0.369073785 |
| ENSMUSG0000003974 | chr5 | Orai2 | -0.497957539 | 0.369349213 |
| ENSMUSG0000003974 | chr11 | Bahcc1 | -0.339581832 | 0.36939787 |
| ENSMUSG0000005807 | chr1 | Sdhc | 0.343092164 | 0.369555663 |
| ENSMUSG0000003504 | chr10 | Creb3l3 | -1.900154736 | 0.37036773 |
| ENSMUSG0000001999 | chr10 | Map7 | -0.363098939 | 0.37036773 |

| | | | | |
|---|---|---|---|---|
| ENSMUSG0000003044 | chr7 | Cyfip1 | 0.152692549 | 0.37036773 |
| ENSMUSG0000004015 | chr3 | Hs2st1 | 0.160271415 | 0.37036773 |
| ENSMUSG0000003793 | chr11 | Smarce1 | 0.204867803 | 0.37036773 |
| ENSMUSG0000003733 | chr5 | Fam53a | 0.204307571 | 0.37036773 |
| ENSMUSG0000000399 | chr13 | Ssbp2 | 0.256309803 | 0.37036773 |
| ENSMUSG0000003028 | chr6 | Creld1 | 0.268115827 | 0.37036773 |
| ENSMUSG0000000628 | chr14 | Osgep | 0.276347475 | 0.37036773 |
| ENSMUSG0000006298 | chr10 | Mrpl42 | 0.331209253 | 0.37036773 |
| ENSMUSG0000003029 | chr6 | Sec13 | 0.386094281 | 0.37036773 |
| ENSMUSG0000005442 | chr14 | Cadps | -0.315720323 | 0.370618422 |
| ENSMUSG0000002070 | chr11 | Ftsj3 | 0.174958095 | 0.370618422 |
| ENSMUSG0000007890 | chr2 | Gm14393 | 1.302581724 | 0.370644853 |
| ENSMUSG0000004632 | chr17 | Slc25a23 | 0.241622267 | 0.370715633 |
| ENSMUSG0000003434 | chr17 | Gtf2h5 | 0.233969905 | 0.370715633 |
| ENSMUSG0000002484 | chr19 | Tmem134 | 0.243562053 | 0.370715633 |
| ENSMUSG0000002070 | chr11 | Psmc5 | 0.301824766 | 0.371092342 |
| ENSMUSG0000003659 | chr17 | H2-Aa | -0.531991053 | 0.37109605 |
| ENSMUSG0000002877 | chr4 | Hcrtr1 | -0.426269543 | 0.37109605 |
| ENSMUSG0000003070 | chr7 | Mrpl48 | 0.281821552 | 0.37109605 |
| ENSMUSG0000003803 | chr1 | Igsf8 | -0.179678128 | 0.371237392 |
| ENSMUSG0000000194 | chr9 | Vsig2 | -0.88157268 | 0.371279702 |
| ENSMUSG0000007163 | chr16 | Cebpd | 0.399629308 | 0.371279702 |
| ENSMUSG0000002133 | chr13 | Slc17a1 | -2.143306812 | 0.372178445 |
| ENSMUSG0000003128 | chrX | Dcx | -0.54179384 | 0.372178445 |
| ENSMUSG0000003022 | chr6 | Dera | -0.398173581 | 0.372178445 |
| ENSMUSG0000001978 | chr10 | Rwdd1 | 0.289622403 | 0.372178445 |
| ENSMUSG0000004562 | chr9 | Cimap1c | 0.380679785 | 0.372178445 |
| ENSMUSG0000003036 | chr6 | Clec2h | -2.136977906 | 0.372402362 |
| ENSMUSG0000010133 | chr1 | Dnah7c | -0.894223955 | 0.372402362 |
| ENSMUSG0000001012 | chr11 | Slc47a1 | -1.046554744 | 0.372402362 |
| ENSMUSG0000003047 | chr7 | Zdhhc13 | -0.308484857 | 0.372402362 |
| ENSMUSG0000005825 | chrX | Tspan7 | -0.190362403 | 0.372402362 |
| ENSMUSG0000000352 | chr16 | Ess2 | 0.176976189 | 0.372402362 |
| ENSMUSG0000000388 | chr15 | Il7r | 1.109840477 | 0.372402362 |
| ENSMUSG0000000909 | chr10 | Derl3 | 0.853497873 | 0.372940733 |
| ENSMUSG0000004268 | chr18 | Garem1 | 0.292066917 | 0.37409866 |
| ENSMUSG0000003461 | chr19 | Ssh3 | -0.205779859 | 0.374424517 |
| ENSMUSG0000002746 | chr2 | Slc52a3 | -0.887978907 | 0.374846614 |
| ENSMUSG0000002318 | chr9 | Vwa5a | -0.17305449 | 0.374846614 |
| ENSMUSG0000003163 | chr8 | Ufsp2 | 0.206114286 | 0.374846614 |
| ENSMUSG0000002551 | chr7 | Cd151 | 0.29330999 | 0.374846614 |
| ENSMUSG0000003736 | chr15 | Ccn3 | -0.722097483 | 0.374959072 |
| ENSMUSG0000002486 | chr19 | Acy3 | -0.644735162 | 0.375144532 |
| ENSMUSG0000003557 | chr2 | L3mbtl1 | -0.765107 | 0.375144532 |
| ENSMUSG0000002599 | chr1 | Wdr75 | 0.196743305 | 0.37515615 |
| ENSMUSG0000004644 | chr11 | Ppm1e | 0.261993223 | 0.37573752 |
| ENSMUSG0000003062 | chr7 | Me3 | 0.428750273 | 0.375765473 |
| ENSMUSG0000003076 | chr1 | Disp1 | 0.28331143 | 0.375937392 |
| ENSMUSG0000002294 | chr16 | Clic6 | -2.539493847 | 0.375950981 |
| ENSMUSG0000003221 | chr9 | Nedd4 | 0.156344449 | 0.375950981 |

| | | | | |
|---|---|---|---|---|
| ENSMUSG0000005840 | chr1 | Txndc9 | 0.251849773 | 0.376217061 |
| ENSMUSG0000002752 | chr2 | Gnas | 0.18696096 | 0.376703574 |
| ENSMUSG0000003170 | chr8 | Rfx1 | -0.22814896 | 0.376741436 |
| ENSMUSG0000002333 | chr8 | Wfdc1 | 0.452004486 | 0.376741436 |
| ENSMUSG0000003031 | chr6 | Timp4 | 0.744951043 | 0.376741436 |
| ENSMUSG0000006887 | chr3 | Cgn | -0.673131707 | 0.376868437 |
| ENSMUSG0000004215 | chr2 | Klhl23 | 0.219966194 | 0.376868437 |
| ENSMUSG0000002938 | chr5 | Eif2b1 | 0.254594206 | 0.376868437 |
| ENSMUSG0000002121 | chr13 | Net1 | 0.294864355 | 0.376868437 |
| ENSMUSG0000002176 | chr13 | Gpx8 | 0.380656173 | 0.376868437 |
| ENSMUSG0000002052 | chr11 | Ppm1d | 0.242003443 | 0.377044581 |
| ENSMUSG0000005272 | chr13 | Map1b | -0.328619506 | 0.37708853 |
| ENSMUSG0000005104 | chr7 | Olfml1 | -0.178376716 | 0.37708853 |
| ENSMUSG0000000645 | chr19 | Rbm14 | 0.204675597 | 0.37708853 |
| ENSMUSG0000003340 | chr3 | Agl | 0.242172062 | 0.37708853 |
| ENSMUSG0000004814 | chr8 | Nwd1 | -0.482674053 | 0.377376775 |
| ENSMUSG0000007522 | chr2 | Lrrc55 | -0.404195691 | 0.377376775 |
| ENSMUSG0000004925 | chr2 | Lrp1b | -0.624188043 | 0.378188332 |
| ENSMUSG0000002685 | chr2 | Dolpp1 | -0.291444185 | 0.378479665 |
| ENSMUSG0000003475 | chr10 | Tle6 | -0.412635819 | 0.379525997 |
| ENSMUSG0000003526 | chr2 | Pkig | 0.305951562 | 0.379525997 |
| ENSMUSG0000002765 | chr2 | Ccn5 | 1.156626044 | 0.379525997 |
| ENSMUSG0000003078 | chr7 | Itgax | -0.825503393 | 0.37972042 |
| ENSMUSG0000002452 | chr18 | Srfbp1 | 0.24437419 | 0.37972042 |
| ENSMUSG0000002438 | chr17 | Csnk2b | 0.254755647 | 0.37972042 |
| ENSMUSG0000002324 | chr14 | Kcnk5 | -0.439661139 | 0.379916078 |
| ENSMUSG0000002408 | chr17 | Pja2 | 0.163638619 | 0.379916078 |
| ENSMUSG0000002767 | chr3 | Dnajc19 | 0.21544455 | 0.379916078 |
| ENSMUSG0000001991 | chr10 | Septin10 | 0.226463109 | 0.379916078 |
| ENSMUSG0000002682 | chr2 | Dnm1 | -0.310158341 | 0.380474253 |
| ENSMUSG0000000266 | chr17 | Dennd1c | -0.749523627 | 0.38052752 |
| ENSMUSG0000000636 | chr8 | Cbfa2t3 | -0.37657346 | 0.380602102 |
| ENSMUSG0000003803 | chr10 | Gcc2 | -0.255462521 | 0.380602102 |
| ENSMUSG0000002230 | chr15 | Angpt1 | -0.445174871 | 0.380975551 |
| ENSMUSG0000002436 | chr18 | Etf1 | 0.222310397 | 0.380975551 |
| ENSMUSG0000004620 | chr14 | Pnma2 | -0.335651565 | 0.381035084 |
| ENSMUSG0000003200 | chr9 | Dcun1d5 | 0.17811368 | 0.381035084 |
| ENSMUSG0000001843 | chr11 | Nol11 | 0.183803043 | 0.381035084 |
| ENSMUSG0000007570 | chr10 | Gpx4 | 0.412555145 | 0.381035084 |
| ENSMUSG0000002014 | chr10 | Lgr5 | -1.259781003 | 0.381149934 |
| ENSMUSG0000003702 | chr7 | Zfp146 | 0.1727937 | 0.381478221 |
| ENSMUSG0000001953 | chr7 | Rcn3 | 0.327354608 | 0.381920217 |
| ENSMUSG0000004518 | chrX | Shroom2 | -0.294723185 | 0.382108333 |
| ENSMUSG0000002033 | chr11 | Slc22a4 | -0.605746313 | 0.382180034 |
| ENSMUSG0000002125 | chr12 | Vash1 | -0.333212673 | 0.382180034 |
| ENSMUSG0000010356 | chr18 | Pcdhga5 | -0.411835076 | 0.38232421 |
| ENSMUSG0000002065 | chr12 | Slc26a4 | -0.717395769 | 0.382460108 |
| ENSMUSG0000002204 | chr14 | Dpysl2 | -0.477299898 | 0.382460108 |
| ENSMUSG0000002622 | chr1 | Slc16a14 | -1.673679552 | 0.382460108 |
| ENSMUSG0000003855 | chr5 | Fndc4 | -0.389119341 | 0.382460108 |

| | | | | |
|---|---|---|---|---|
| ENSMUSG0000003887 | chr15 | Nipal2 | -0.384117049 | 0.382460108 |
| ENSMUSG0000003207 | chr9 | Cadm1 | -0.275976581 | 0.382460108 |
| ENSMUSG0000003851 | chr11 | Tbkbp1 | -0.297325981 | 0.382460108 |
| ENSMUSG0000003193 | chr9 | Vstm5 | -0.281330713 | 0.382460108 |
| ENSMUSG0000004687 | chrX | Mbtps2 | -0.22856643 | 0.382460108 |
| ENSMUSG0000002720 | chr2 | Cops2 | 0.186238271 | 0.382460108 |
| ENSMUSG0000002660 | chr1 | Ptpn14 | 0.212924875 | 0.382460108 |
| ENSMUSG0000003674 | chr3 | Ttll7 | 0.217014074 | 0.382460108 |
| ENSMUSG0000002195 | chr14 | Anxa8 | 0.569147196 | 0.382460108 |
| ENSMUSG0000003482 | chr5 | Nup54 | 0.22594311 | 0.382706761 |
| ENSMUSG0000004718 | chr11 | Samd14 | -0.28332497 | 0.382905359 |
| ENSMUSG0000000212 | chr11 | Sf3a1 | 0.18488756 | 0.382905359 |
| ENSMUSG0000002648 | chr1 | Niban1 | 0.2166295 | 0.383225725 |
| ENSMUSG0000003086 | chr7 | Plk1 | -1.199962333 | 0.383523311 |
| ENSMUSG0000004176 | chr7 | Ppp2r2d | 0.203483273 | 0.383523311 |
| ENSMUSG0000005766 | chr6 | Gapdh | 0.319063987 | 0.383523311 |
| ENSMUSG0000002209 | chr14 | Ppp3cc | 0.339640337 | 0.383523311 |
| ENSMUSG0000006353 | chr7 | Zfp773 | -0.757409956 | 0.38366093 |
| ENSMUSG0000002467 | chr19 | Ms4a7 | -0.381044973 | 0.38366093 |
| ENSMUSG0000003012 | chr6 | Mlf2 | 0.249668335 | 0.38366093 |
| ENSMUSG0000009005 | chr4 | Pakap | 0.671549591 | 0.38366093 |
| ENSMUSG0000002099 | chr12 | Trappc6b | 0.199101774 | 0.384154812 |
| ENSMUSG0000003201 | chr9 | Thy1 | -0.267388098 | 0.384365189 |
| ENSMUSG0000003900 | chr2 | Rps21 | 0.425080175 | 0.384365189 |
| ENSMUSG0000004933 | chr11 | Tenm2 | 0.509772825 | 0.384365189 |
| ENSMUSG0000004164 | chr1 | Kif21b | -0.769280451 | 0.384416604 |
| ENSMUSG0000002453 | chr18 | Spire1 | 0.238144824 | 0.385157284 |
| ENSMUSG0000001857 | chr11 | Acadvl | 0.244627331 | 0.385157284 |
| ENSMUSG0000004618 | chr7 | Gsg1l | 0.264148405 | 0.385365881 |
| ENSMUSG0000000671 | chr13 | D130043K22Rik | -0.414337376 | 0.385535985 |
| ENSMUSG0000003887 | chr10 | Rnf146 | 0.222749596 | 0.385729563 |
| ENSMUSG0000009685 | chr15 | Cyp2d12 | -3.058674384 | 0.386251062 |
| ENSMUSG0000002581 | chr2 | Dhtkd1 | -0.634415371 | 0.386404334 |
| ENSMUSG0000006623 | chr9 | Pomgnt2 | -0.195586491 | 0.386404334 |
| ENSMUSG0000003674 | chr19 | Cuedc2 | 0.354553122 | 0.386404334 |
| ENSMUSG0000000958 | chr15 | Apobec3 | -0.363769659 | 0.386659698 |
| ENSMUSG0000002110 | chr12 | Mnat1 | 0.274977916 | 0.387405521 |
| ENSMUSG0000007481 | chr19 | Pdzd7 | -0.264364763 | 0.387447544 |
| ENSMUSG0000001995 | chr10 | Bltp3b | 0.208055242 | 0.387688956 |
| ENSMUSG0000005899 | chr14 | Vwa8 | -0.174313465 | 0.387792389 |
| ENSMUSG0000002964 | chr5 | Pomp | 0.312934364 | 0.387792389 |
| ENSMUSG0000002936 | chr5 | Nos1 | -0.334412109 | 0.387927195 |
| ENSMUSG0000007365 | chr1 | Catip | -0.507409265 | 0.387990147 |
| ENSMUSG0000001994 | chr10 | Cabcoco1 | -1.060822995 | 0.388272138 |
| ENSMUSG0000004169 | chr11 | Kcnj2 | -0.572432932 | 0.388272138 |
| ENSMUSG0000004003 | chr2 | Disp2 | -0.310079956 | 0.388272138 |
| ENSMUSG0000004070 | chr14 | Ap1g2 | -0.377769687 | 0.388272138 |
| ENSMUSG0000000394 | chr11 | Mmd | 0.186681818 | 0.388272138 |
| ENSMUSG0000002250 | chr16 | Litaf | 0.270499559 | 0.388272138 |
| ENSMUSG0000006200 | chr3 | Rpl34 | 0.424533477 | 0.388272138 |

| | | | | |
|---|---|---|---|---|
| ENSMUSG0000007897 | chr11 | Sec61g | 0.515178843 | 0.388272138 |
| ENSMUSG0000001789 | chr2 | Eya2 | 0.904121186 | 0.388272138 |
| ENSMUSG0000002865 | chr4 | Cap1 | 0.20145628 | 0.388558424 |
| ENSMUSG0000002602 | chr1 | Casp8 | -0.376210749 | 0.388607774 |
| ENSMUSG0000000598 | chr16 | Trap1 | 0.180995508 | 0.388607774 |
| ENSMUSG0000002111 | chr12 | Eif2s1 | 0.275142611 | 0.388607774 |
| ENSMUSG0000007465 | chr2 | Eif2s2 | 0.241992624 | 0.388670772 |
| ENSMUSG0000005083 | chr11 | Vwc2 | 0.476523741 | 0.388699369 |
| ENSMUSG0000001744 | chr11 | C1qtnf1 | -0.329285568 | 0.388718044 |
| ENSMUSG0000001655 | chr15 | Eif3d | 0.25446174 | 0.388718044 |
| ENSMUSG0000006388 | chr4 | Uqcrh | 0.401889223 | 0.388718044 |
| ENSMUSG0000002931 | chr5 | Nudt9 | 0.257230382 | 0.38882575 |
| ENSMUSG0000004417 | chr11 | Wfikkn2 | 0.286471085 | 0.389007965 |
| ENSMUSG0000000866 | chr17 | Rps18 | 0.464655748 | 0.389007965 |
| ENSMUSG0000004998 | chr13 | Ankrd55 | -0.948441258 | 0.389132585 |
| ENSMUSG0000009603 | chr14 | D830030K20Rik | -1.565523621 | 0.389280925 |
| ENSMUSG0000003807 | chr6 | Fkbp14 | 0.298909565 | 0.389756991 |
| ENSMUSG0000003982 | chr15 | Wdr70 | 0.213600297 | 0.389922085 |
| ENSMUSG0000002455 | chr18 | Mapk4 | 0.358183075 | 0.389922085 |
| ENSMUSG0000005134 | chr6 | Rab11fip5 | 0.209986604 | 0.390547108 |
| ENSMUSG0000002672 | chr2 | Trdmt1 | 0.222800419 | 0.390604696 |
| ENSMUSG0000006070 | chr2 | Cd302 | 0.264809752 | 0.391461314 |
| ENSMUSG0000007049 | chr5 | Chchd2 | 0.464148723 | 0.391584068 |
| ENSMUSG0000001513 | chr7 | Lrrk1 | 0.246102836 | 0.391795053 |
| ENSMUSG0000003845 | chr11 | Srcin1 | -0.337604942 | 0.39265495 |
| ENSMUSG0000003391 | chr7 | Chmp2a | 0.220510564 | 0.39265495 |
| ENSMUSG0000004239 | chr9 | Rbm7 | 0.220448333 | 0.393033799 |
| ENSMUSG0000004988 | chr12 | Vcpkmt | 0.45886051 | 0.39327967 |
| ENSMUSG0000004272 | chr12 | Map3k9 | -0.534364506 | 0.393421745 |
| ENSMUSG0000000062 | chr6 | Hk2 | -0.907168881 | 0.393717496 |
| ENSMUSG0000000158 | chr11 | Tnk1 | -1.308852715 | 0.393717496 |
| ENSMUSG0000003297 | chr4 | Guca2b | -1.729860108 | 0.393717496 |
| ENSMUSG0000002788 | chr3 | Clcc1 | 0.16410559 | 0.393717496 |
| ENSMUSG0000004000 | chr10 | Ginm1 | 0.214838535 | 0.393717496 |
| ENSMUSG0000003427 | chr12 | Jdp2 | 0.211719541 | 0.393717496 |
| ENSMUSG0000002141 | chr13 | Eci2 | 0.275611308 | 0.393717496 |
| ENSMUSG0000000316 | chr5 | Sri | -0.243360814 | 0.394129328 |
| ENSMUSG0000004587 | chr14 | Adra1a | 0.304127672 | 0.394215992 |
| ENSMUSG0000000265 | chr17 | Gtf2f1 | 0.199172962 | 0.394782625 |
| ENSMUSG0000004061 | chr6 | Apobec1 | -0.433727446 | 0.394798151 |
| ENSMUSG0000002072 | chr11 | Psmd12 | 0.223994852 | 0.395201275 |
| ENSMUSG0000003612 | chr13 | Slc9a3 | -2.152032578 | 0.395284602 |
| ENSMUSG0000002201 | chr14 | Epsti1 | -0.61000127 | 0.395284602 |
| ENSMUSG0000005591 | chr12 | Zfp277 | 0.186908005 | 0.395284602 |
| ENSMUSG0000000201 | chrX | Ssr4 | 0.247070259 | 0.395284602 |
| ENSMUSG0000003353 | chr9 | Casp4 | -0.32887747 | 0.395326412 |
| ENSMUSG0000002893 | chr4 | Kcnab2 | -0.260336288 | 0.395326412 |
| ENSMUSG0000002458 | chr18 | Nars1 | 0.156603406 | 0.396054727 |
| ENSMUSG0000006812 | chrX | Agtr2 | -0.686309926 | 0.396403425 |
| ENSMUSG0000005771 | chr18 | Sh3rf2 | -0.63877643 | 0.396403425 |

| | | | | |
|---|---|---|---|---|
| ENSMUSG0000002912 | chr5 | Crmp1 | -0.287166504 | 0.396403425 |
| ENSMUSG0000003249 | chr9 | Lrrc2 | -0.408165111 | 0.396403425 |
| ENSMUSG0000002658 | chr1 | Kifap3 | -0.176003237 | 0.396403425 |
| ENSMUSG0000002559 | chr8 | Tma16 | 0.238754537 | 0.396403425 |
| ENSMUSG0000002884 | chr4 | Trappc3 | 0.226326661 | 0.396403425 |
| ENSMUSG0000006264 | chr2 | Rpl7a | 0.298120903 | 0.396403425 |
| ENSMUSG0000007903 | chr8 | Zfp781a | 0.429672996 | 0.396403425 |
| ENSMUSG0000002466 | chr19 | Fads3 | 0.350898891 | 0.396447722 |
| ENSMUSG0000003637 | chr6 | Serbp1 | 0.207338115 | 0.396780346 |
| ENSMUSG0000002888 | chr5 | Sema3a | 0.239822928 | 0.396993745 |
| ENSMUSG0000005614 | chr7 | Trim34a | -0.439283507 | 0.397041486 |
| ENSMUSG0000002145 | chr13 | Sema4d | -0.321006738 | 0.397041486 |
| ENSMUSG0000005940 | chr10 | Tmprss9 | 0.37768675 | 0.397041486 |
| ENSMUSG0000005645 | chr13 | Prl2c3 | 0.924165289 | 0.397041486 |
| ENSMUSG0000002078 | chr11 | Srp68 | 0.20735074 | 0.397311051 |
| ENSMUSG0000003035 | chr6 | Tead4 | 0.638666253 | 0.397645054 |
| ENSMUSG0000004663 | chr4 | Ttc34 | -0.820179807 | 0.397919309 |
| ENSMUSG0000000138 | chr18 | Hars1 | 0.159677698 | 0.397919309 |
| ENSMUSG0000002946 | chr5 | Vps29 | 0.204238092 | 0.397919309 |
| ENSMUSG0000005440 | chr4 | Dnajc8 | 0.229776234 | 0.397919309 |
| ENSMUSG0000002832 | chr4 | Tmod1 | 0.216680363 | 0.397919309 |
| ENSMUSG0000009171 | chr16 | Sec14l5 | -1.353576492 | 0.398316404 |
| ENSMUSG0000004722 | chr6 | A2ml1 | -2.032615126 | 0.398316404 |
| ENSMUSG0000003801 | chr11 | Wipf2 | -0.21249143 | 0.398316404 |
| ENSMUSG0000003174 | chr8 | Gnao1 | -0.226550789 | 0.398316404 |
| ENSMUSG0000001847 | chr11 | Chd3 | -0.163835102 | 0.398316404 |
| ENSMUSG0000001783 | chr11 | Nkiras2 | 0.190833354 | 0.398316404 |
| ENSMUSG0000004975 | chr11 | Zfp672 | 0.260246669 | 0.398316404 |
| ENSMUSG0000002780 | chr3 | Serp1 | 0.28560615 | 0.398518994 |
| ENSMUSG0000003345 | chr17 | Tagap | 0.684278819 | 0.398518994 |
| ENSMUSG0000002754 | chr2 | Nfatc2 | 0.312322897 | 0.398828842 |
| ENSMUSG0000000033 | chr3 | Rtca | 0.253379833 | 0.399311369 |
| ENSMUSG0000004211 | chr1 | Klhdc8a | -0.700213941 | 0.39938632 |
| ENSMUSG0000004891 | chr17 | Efna5 | 0.22919136 | 0.39938632 |
| ENSMUSG0000002404 | chr17 | Myom1 | 0.318418312 | 0.39938632 |
| ENSMUSG0000003208 | chr9 | Dscaml1 | -0.389571196 | 0.399880016 |
| ENSMUSG0000004124 | chrX | Wnk3 | -0.439602864 | 0.399952437 |
| ENSMUSG0000003061 | chr7 | Mrpl46 | 0.275073016 | 0.399952437 |
| ENSMUSG0000004599 | chr9 | B3gat1 | 0.408503276 | 0.400160778 |
| ENSMUSG0000002725 | chr2 | Lrp4 | -0.471265369 | 0.400334937 |
| ENSMUSG0000002818 | chr3 | Rpf1 | 0.251150918 | 0.400334937 |
| ENSMUSG0000003263 | chr11 | Flcn | 0.257413594 | 0.400334937 |
| ENSMUSG0000004967 | chr19 | Morn4 | -0.323913183 | 0.400635548 |
| ENSMUSG0000002156 | chr13 | Trip13 | -0.941036472 | 0.401349408 |
| ENSMUSG0000002651 | chr1 | Cnih3 | -1.349311427 | 0.401349408 |
| ENSMUSG0000006809 | chr15 | Smim45 | -0.997629667 | 0.401349408 |
| ENSMUSG0000006252 | chr18 | Mppe1 | -0.233192801 | 0.401349408 |
| ENSMUSG0000003778 | chr12 | Coa8 | 0.245899962 | 0.401349408 |
| ENSMUSG0000003618 | chr13 | H1f2 | 0.346546824 | 0.401349408 |
| ENSMUSG0000000150 | chr11 | Sgca | 0.299328715 | 0.401437438 |

| | | | | |
|---|---|---|---|---|
| ENSMUSG0000005293[illegible] | chr8 | Fbxo31 | 0.17434344 | 0.40149794 |
| ENSMUSG0000005874[illegible] | chr2 | Kcnt1 | -0.430987293 | 0.40165204 |
| ENSMUSG0000005268[illegible] | chr10 | Rap1b | 0.231964179 | 0.401756488 |
| ENSMUSG0000004180[illegible] | chr1 | Phlda3 | 0.397645498 | 0.401878358 |
| ENSMUSG0000003178[illegible] | chr8 | Cngb1 | -0.333129033 | 0.402024666 |
| ENSMUSG0000001407[illegible] | chr2 | Chp1 | -0.210683499 | 0.402108764 |
| ENSMUSG0000009502[illegible] | chr8 | Gm3336 | -2.009772978 | 0.402166411 |
| ENSMUSG0000003286[illegible] | chr7 | P2ry2 | -0.746494852 | 0.402166411 |
| ENSMUSG0000004341[illegible] | chr2 | Otud1 | 0.189162464 | 0.402166411 |
| ENSMUSG0000004312[illegible] | chr12 | A530016L24Rik | 0.888321074 | 0.402166411 |
| ENSMUSG0000005786[illegible] | chr17 | Rpl36 | 0.430129446 | 0.402166411 |
| ENSMUSG0000002550[illegible] | chr7 | Lmntd2 | -1.678886436 | 0.403593214 |
| ENSMUSG0000007468[illegible] | chr2 | Zcchc3 | -0.228488133 | 0.40363033 |
| ENSMUSG0000002839[illegible] | chr4 | Ptprd | -0.223589039 | 0.403698101 |
| ENSMUSG0000003212[illegible] | chr9 | Robo3 | -1.050037394 | 0.403860519 |
| ENSMUSG0000001898[illegible] | chr4 | E2f2 | -0.575247927 | 0.404237175 |
| ENSMUSG0000004123[illegible] | chr4 | Chd7 | -0.339213732 | 0.404237175 |
| ENSMUSG0000000109[illegible] | chr5 | Kctd10 | 0.300265926 | 0.404237175 |
| ENSMUSG0000002016[illegible] | chr10 | Uqcr11 | 0.421732443 | 0.40447004 |
| ENSMUSG0000003415[illegible] | chr11 | Tspoap1 | -0.432571058 | 0.404735596 |
| ENSMUSG0000007467[illegible] | chr2 | Tspyl3 | -0.263820526 | 0.404735596 |
| ENSMUSG0000002250[illegible] | chr16 | Emp2 | 0.285949081 | 0.404766945 |
| ENSMUSG0000002694[illegible] | chr2 | Abca2 | -0.243644229 | 0.404994115 |
| ENSMUSG0000006074[illegible] | chr1 | H3f3a | 0.271091076 | 0.404994115 |
| ENSMUSG0000004778[illegible] | chr17 | Lix1 | -0.257389017 | 0.405315128 |
| ENSMUSG0000002919[illegible] | chr5 | Tbc1d14 | -0.173448693 | 0.405315128 |
| ENSMUSG0000003199[illegible] | chr9 | Aplp2 | 0.119852356 | 0.405315128 |
| ENSMUSG0000002228[illegible] | chr15 | Rnf19a | 0.164632998 | 0.405315128 |
| ENSMUSG0000002846[illegible] | chr4 | Tln1 | 0.252170042 | 0.405315128 |
| ENSMUSG0000002233[illegible] | chr15 | Emc2 | 0.2215267 | 0.405315128 |
| ENSMUSG0000007294[illegible] | chr5 | Sod3 | 0.222707132 | 0.405315128 |
| ENSMUSG0000006238[illegible] | chr8 | Tubb3 | -0.279646458 | 0.405642556 |
| ENSMUSG0000003184[illegible] | chr8 | 1700030J22Rik | -0.556292644 | 0.405916569 |
| ENSMUSG0000003256[illegible] | chr9 | Aste1 | 0.378680923 | 0.405916569 |
| ENSMUSG0000003053[illegible] | chr7 | Furin | 0.195256328 | 0.405960389 |
| ENSMUSG0000002015[illegible] | chr10 | Ndufs7 | 0.296702925 | 0.405960389 |
| ENSMUSG0000002450[illegible] | chr18 | Jakmip2 | -0.319893102 | 0.406231413 |
| ENSMUSG0000005541[illegible] | chr17 | H2-Q5 | -1.354251098 | 0.406312292 |
| ENSMUSG0000003411[illegible] | chr9 | Scn11a | -0.257016862 | 0.406312292 |
| ENSMUSG0000007858[illegible] | chr7 | Zfp764l1 | 0.305621877 | 0.4070118 |
| ENSMUSG0000005655[illegible] | chr12 | Ptprn2 | -0.321244279 | 0.407427607 |
| ENSMUSG0000001978[illegible] | chr10 | Clvs2 | -0.53796079 | 0.407541527 |
| ENSMUSG0000000053[illegible] | chr15 | Acvr1b | -0.339754997 | 0.407541527 |
| ENSMUSG0000001364[illegible] | chr11 | Sh3bp5l | 0.245362386 | 0.407541527 |
| ENSMUSG0000002762[illegible] | chr2 | Epb41l1 | -0.208988316 | 0.40816159 |
| ENSMUSG0000009380[illegible] | chr1 | Gal3st2b | -2.254627065 | 0.408382687 |
| ENSMUSG0000002487[illegible] | chr19 | Cnih2 | -0.455552496 | 0.408382687 |
| ENSMUSG0000004071[illegible] | chr1 | St8sia4 | -0.541606513 | 0.408382687 |
| ENSMUSG0000003764[illegible] | chr15 | Vps13b | -0.342953636 | 0.408382687 |
| ENSMUSG0000002133[illegible] | chr13 | Carmil1 | -0.296981804 | 0.408382687 |

| | | | | |
|---|---|---|---|---|
| ENSMUSG0000003766 | chr12 | Ldah | 0.147569333 | 0.408382687 |
| ENSMUSG0000002421 | chr17 | Snrpc | 0.349403544 | 0.408382687 |
| ENSMUSG0000002418 | chr17 | Mrpl28 | 0.397073301 | 0.408382687 |
| ENSMUSG0000003009 | chr6 | Nup210 | -0.511333514 | 0.40840485 |
| ENSMUSG0000004857 | chr5 | Mlec | 0.198845884 | 0.40840485 |
| ENSMUSG0000002767 | chr3 | Ndufb5 | 0.285841717 | 0.40840485 |
| ENSMUSG0000005208 | chr13 | Rgs14 | -0.759870301 | 0.408412835 |
| ENSMUSG0000001596 | chr1 | Adss2 | 0.16567868 | 0.408412835 |
| ENSMUSG0000005650 | chr2 | Cebpb | 0.449972925 | 0.408412835 |
| ENSMUSG0000009549 | chr14 | A630023A22Rik | 0.310712128 | 0.408414176 |
| ENSMUSG0000004035 | chr10 | Mars1 | 0.177799703 | 0.408561808 |
| ENSMUSG0000006422 | chr9 | Paqr9 | 0.490722613 | 0.4086442 |
| ENSMUSG0000002294 | chr16 | Pigp | 0.23073619 | 0.40886784 |
| ENSMUSG0000002209 | chr14 | Hr | 0.256868638 | 0.409007112 |
| ENSMUSG0000004204 | chr6 | Csgalnact2 | 0.18935516 | 0.409156364 |
| ENSMUSG0000002390 | chr17 | Pkmyt1 | -0.512487103 | 0.409842145 |
| ENSMUSG0000002812 | chr3 | Pip5k1a | 0.150332554 | 0.409842145 |
| ENSMUSG0000004300 | chr16 | Klhl6 | -0.602713504 | 0.409903532 |
| ENSMUSG0000003446 | chr5 | Pkd2 | 0.138909396 | 0.409903532 |
| ENSMUSG0000002114 | chr12 | Mta1 | 0.186158889 | 0.410085999 |
| ENSMUSG0000002033 | chr11 | Zfp354b | 0.374700841 | 0.410085999 |
| ENSMUSG0000003131 | chrX | Zmym3 | -0.259321596 | 0.410464679 |
| ENSMUSG0000003797 | chr5 | Ccdc92 | -0.244131581 | 0.410464679 |
| ENSMUSG0000002060 | chr12 | Smc6 | 0.195333616 | 0.410464679 |
| ENSMUSG0000003147 | chr8 | Vps36 | 0.177575919 | 0.41068331 |
| ENSMUSG0000005607 | chr5 | Eif3b | 0.247569289 | 0.41068331 |
| ENSMUSG0000002688 | chr2 | Mrrf | 0.292510302 | 0.411320854 |
| ENSMUSG0000003463 | chr6 | Setmar | 0.354559401 | 0.411422217 |
| ENSMUSG0000003457 | chr11 | Pla2g3 | -0.774115239 | 0.411432606 |
| ENSMUSG0000003610 | chr15 | Prr5 | -0.344264147 | 0.412356339 |
| ENSMUSG0000010308 | chr18 | Pcdhgb8 | 0.562364575 | 0.412356339 |
| ENSMUSG0000002062 | chr12 | Klhl29 | 0.22018297 | 0.412711627 |
| ENSMUSG0000002088 | chr11 | Cacnb1 | -0.29629259 | 0.412712314 |
| ENSMUSG0000004913 | chr7 | C5ar1 | 0.40367629 | 0.412712314 |
| ENSMUSG0000009164 | chr14 | Phf11b | -0.847529068 | 0.412797377 |
| ENSMUSG0000006388 | chr3 | Nlgn1 | -0.265429312 | 0.412797377 |
| ENSMUSG0000000548 | chr8 | Ddx39a | 0.235755339 | 0.412797377 |
| ENSMUSG0000002101 | chr12 | Npas3 | 0.528457288 | 0.413104077 |
| ENSMUSG0000004771 | chr10 | Ust | 0.300514323 | 0.413104077 |
| ENSMUSG0000003249 | chr9 | Pth1r | 0.32604154 | 0.413460049 |
| ENSMUSG0000003184 | chr8 | Rab3a | -0.204784071 | 0.413493769 |
| ENSMUSG0000004054 | chr2 | Ckap5 | -0.163288023 | 0.413550693 |
| ENSMUSG0000002716 | chr2 | Dnajc24 | 0.395553887 | 0.413550693 |
| ENSMUSG0000000152 | chr17 | Gtf2h4 | 0.27016749 | 0.413722395 |
| ENSMUSG0000004484 | chr11 | Lsm11 | -0.325125065 | 0.413816235 |
| ENSMUSG0000000029 | chr10 | Tpd52l1 | -0.272495424 | 0.414211614 |
| ENSMUSG0000002586 | chr13 | Higd2a | 0.29705842 | 0.414211614 |
| ENSMUSG0000001834 | chr11 | Anxa6 | 0.245054428 | 0.414230808 |
| ENSMUSG0000011390 | chr12 | Ndufb1 | 0.520423039 | 0.414230808 |
| ENSMUSG0000005145 | chr11 | Gm11437 | -1.573858406 | 0.414366122 |

| | | | | |
|---|---|---|---|---|
| ENSMUSG0000004496 | chr11 | Fbxo48 | -1.102256483 | 0.414366122 |
| ENSMUSG0000007941 | chrX | Atg4a | 0.199164046 | 0.414366122 |
| ENSMUSG0000003452 | chr11 | Gjc1 | 0.196598665 | 0.414918471 |
| ENSMUSG0000005843 | chr17 | Btnl4 | -2.833087118 | 0.415320144 |
| ENSMUSG0000001893 | chr11 | Natd1 | 0.286192616 | 0.415320144 |
| ENSMUSG0000003952 | chrX | Foxp3 | 1.123175675 | 0.415505055 |
| ENSMUSG0000002289 | chr16 | Atp5pf | 0.272055154 | 0.415552647 |
| ENSMUSG0000003575 | chr10 | Wdr18 | 0.270032907 | 0.415552647 |
| ENSMUSG0000006945 | chr10 | Rdh16 | -2.307427409 | 0.415585192 |
| ENSMUSG0000002519 | chr19 | Dnmbp | -0.23047273 | 0.416086082 |
| ENSMUSG0000002043 | chr11 | 4921536K21Rik | -0.757379247 | 0.416303365 |
| ENSMUSG0000002219 | chr14 | Psmb5 | 0.450103456 | 0.416303365 |
| ENSMUSG0000003105 | chrX | Ndufb11 | 0.339724603 | 0.416380872 |
| ENSMUSG0000003884 | chr11 | Phb1 | 0.378848407 | 0.416380872 |
| ENSMUSG0000001936 | chr8 | D8Ertd738e | 0.293637544 | 0.41639737 |
| ENSMUSG0000005553 | chr14 | Zcchc24 | 0.187868727 | 0.417261919 |
| ENSMUSG0000002664 | chr1 | Plxna2 | -0.354564331 | 0.41743196 |
| ENSMUSG0000005688 | chr10 | Glipr1 | 0.366225061 | 0.41743196 |
| ENSMUSG0000003780 | chr17 | Rpl10a | 0.311733292 | 0.418048744 |
| ENSMUSG0000002594 | chr1 | Tmem70 | 0.285548492 | 0.418364998 |
| ENSMUSG0000003195 | chr8 | Tmem170 | -0.262052869 | 0.41844017 |
| ENSMUSG0000003068 | chr7 | Ino80e | 0.235519156 | 0.418672353 |
| ENSMUSG0000001979 | chr10 | Lrp11 | -0.337312594 | 0.418799464 |
| ENSMUSG0000003310 | chr8 | Slc7a6os | 0.200223457 | 0.418799464 |
| ENSMUSG0000002107 | chr12 | Actr10 | 0.26421341 | 0.418799464 |
| ENSMUSG0000004957 | chr7 | Cfap46 | -0.447387989 | 0.41894142 |
| ENSMUSG0000002723 | chr2 | Eif3j1 | 0.249820246 | 0.419602302 |
| ENSMUSG0000009209 | chr5 | Zfp804b | -1.535393627 | 0.419727826 |
| ENSMUSG0000003686 | chr7 | Proser3 | -0.332490452 | 0.419727826 |
| ENSMUSG0000002771 | chr3 | Anxa5 | 0.287942785 | 0.419727826 |
| ENSMUSG0000000652 | chr14 | Itih3 | 0.488182207 | 0.419727826 |
| ENSMUSG0000005116 | chr6 | Rpusd3 | 0.498867541 | 0.419727826 |
| ENSMUSG0000003749 | chr10 | Slc2a12 | 0.323642236 | 0.419727826 |
| ENSMUSG0000003633 | chr14 | Tmem260 | -0.224645733 | 0.419972113 |
| ENSMUSG0000003049 | chr7 | Kctd15 | 0.239766804 | 0.419972113 |
| ENSMUSG0000002883 | chr4 | Psmb2 | 0.25695836 | 0.419972113 |
| ENSMUSG0000001476 | chr17 | Psmb1 | 0.287182759 | 0.419988061 |
| ENSMUSG0000000481 | chr5 | Dgkq | -0.320072132 | 0.420342147 |
| ENSMUSG0000010834 | chr7 | Pnma8c | -0.297562487 | 0.420723384 |
| ENSMUSG0000005307 | chr9 | Cfap300 | -0.384823714 | 0.420723384 |
| ENSMUSG0000003432 | chr19 | Exosc1 | 0.230526182 | 0.420748868 |
| ENSMUSG0000002038 | chr11 | Pdlim4 | 0.330418724 | 0.420922199 |
| ENSMUSG0000000541 | chr8 | Hmox1 | 0.493726391 | 0.420922199 |
| ENSMUSG0000002275 | chr16 | Slc7a4 | -0.432042031 | 0.421140642 |
| ENSMUSG0000002603 | chr1 | Orc2 | 0.244650788 | 0.421213287 |
| ENSMUSG0000002097 | chr12 | Pole2 | -0.701209943 | 0.421388171 |
| ENSMUSG0000000176 | chr2 | Crnkl1 | 0.20057862 | 0.421388171 |
| ENSMUSG0000002192 | chr14 | Kpna3 | 0.205915579 | 0.421388171 |
| ENSMUSG0000000657 | chr11 | Rundc3a | -0.291413533 | 0.421486348 |
| ENSMUSG0000002865 | chr4 | Mycl | -0.422300656 | 0.421619497 |

| | | | | |
|---|---|---|---|---|
| ENSMUSG0000003183 | chr8 | Necab2 | -0.36466512 | 0.421774 |
| ENSMUSG000000547C | chr1 | Ap1s3 | -0.798959247 | 0.421774 |
| ENSMUSG0000001647 | chr13 | E2f3 | -0.358623869 | 0.421774 |
| ENSMUSG0000003434 | chr9 | Cbl | -0.235681524 | 0.422224105 |
| ENSMUSG0000004982 | chr17 | Zbtb12 | -0.281119014 | 0.422224105 |
| ENSMUSG000000290E | chr4 | Cdk11b | 0.171726691 | 0.422224105 |
| ENSMUSG0000003022 | chr6 | Strap | 0.229052844 | 0.422224105 |
| ENSMUSG0000001111 | chr9 | Tbrg1 | 0.274058863 | 0.422224105 |
| ENSMUSG0000003061 | chr7 | Tmem126a | 0.322875424 | 0.422224105 |
| ENSMUSG0000004742 | chr17 | Dlk2 | -1.230559871 | 0.422621382 |
| ENSMUSG000000544C | chr15 | Tmem74 | -0.385964131 | 0.422621382 |
| ENSMUSG0000002694 | chr2 | Traf2 | 0.330859397 | 0.422621382 |
| ENSMUSG0000006411 | chr16 | Cadm2 | 0.360633303 | 0.422790306 |
| ENSMUSG0000002094 | chr11 | Efcab15 | -0.440365463 | 0.422840181 |
| ENSMUSG0000003791 | chr19 | Ndufv1 | 0.210698786 | 0.423194746 |
| ENSMUSG0000003524 | chrX | Pcyt1b | -0.362382076 | 0.423436363 |
| ENSMUSG0000000327 | chr7 | Sult2b1 | -1.070299364 | 0.42371854 |
| ENSMUSG0000003662 | chr11 | Mgat4b | -0.301648193 | 0.42371854 |
| ENSMUSG0000003424 | chr16 | Golgb1 | -0.205982784 | 0.42371854 |
| ENSMUSG000000537S | chr19 | Exoc6 | 0.156421239 | 0.42371854 |
| ENSMUSG0000003158 | chr8 | Rbpms | 0.199350266 | 0.42371854 |
| ENSMUSG0000004171 | chr12 | Ubr7 | 0.224571253 | 0.42371854 |
| ENSMUSG0000002651 | chr1 | Srp9 | 0.304300327 | 0.42371854 |
| ENSMUSG0000007063 | chr4 | Srarp | 1.026040037 | 0.42371854 |
| ENSMUSG000000785E | chr7 | Bnip3 | 0.221206622 | 0.423993718 |
| ENSMUSG000000748S | chr16 | B3galt5 | -0.918143915 | 0.424174943 |
| ENSMUSG0000002495 | chr19 | Bad | 0.331966434 | 0.424964564 |
| ENSMUSG000000749E | chr2 | Ano3 | -0.523824776 | 0.425019748 |
| ENSMUSG000000220S | chr14 | Dmtn | -0.287690311 | 0.425217405 |
| ENSMUSG0000005107 | chr1 | Rgs13 | -2.585521519 | 0.426459479 |
| ENSMUSG000000217E | chr13 | Cspg4b | -0.423508783 | 0.426459479 |
| ENSMUSG0000003945 | chr11 | Dcxr | -0.382714784 | 0.426459479 |
| ENSMUSG0000006141 | chr2 | Nol4l | -0.226592492 | 0.426459479 |
| ENSMUSG0000001652 | chr1 | Mapkapk2 | 0.273683802 | 0.426459479 |
| ENSMUSG000000491C | chr9 | Ccr2 | -0.405104645 | 0.426642995 |
| ENSMUSG0000006343 | chr7 | B9d2 | 0.309781592 | 0.427647103 |
| ENSMUSG0000005134 | chr17 | Zfp52 | -0.287478825 | 0.427879741 |
| ENSMUSG0000002052 | chr11 | Mfap3 | 0.169188532 | 0.427879741 |
| ENSMUSG000000150S | chr2 | Fbxw5 | 0.177164944 | 0.427879741 |
| ENSMUSG0000002295 | chr16 | Itsn1 | 0.212586316 | 0.427879741 |
| ENSMUSG0000002037 | chr11 | Rack1 | 0.303608381 | 0.427879741 |
| ENSMUSG0000002605 | chr1 | Ecrg4 | 0.451387077 | 0.427879741 |
| ENSMUSG0000001734 | chr11 | Vtn | 0.473908852 | 0.427879741 |
| ENSMUSG0000007043 | chr7 | Serpinh1 | 0.410781392 | 0.427879741 |
| ENSMUSG000000283E | chr3 | Slc44a5 | 0.35232556 | 0.427879741 |
| ENSMUSG0000006668 | chr9 | Zbtb16 | 0.945526744 | 0.427879741 |
| ENSMUSG000000353C | chr4 | Ror1 | 0.242330902 | 0.428115297 |
| ENSMUSG0000003247 | chr9 | Cdc25a | -0.300837547 | 0.428392988 |
| ENSMUSG000000210E | chr12 | Sptb | 0.19494716 | 0.428392988 |
| ENSMUSG0000002073 | chr11 | Sumo2 | 0.249888616 | 0.428392988 |

| | | | | |
|---|---|---|---|---|
| ENSMUSG000000316E | chr8 | Lsm6 | 0.263177719 | 0.428392988 |
| ENSMUSG000000272C | chr2 | Fgf7 | 0.585833944 | 0.428392988 |
| ENSMUSG00000002401 | chr17 | Cmtr1 | -0.425602962 | 0.428472097 |
| ENSMUSG000000352 | chr16 | Slc25a1 | -0.235333095 | 0.428472097 |
| ENSMUSG000000215E | chr13 | Rhobtb3 | -0.188298394 | 0.428472097 |
| ENSMUSG000000142S | chr18 | Ndufa2 | 0.433857053 | 0.428472097 |
| ENSMUSG000000200 | chr10 | Supv3l1 | 0.203614306 | 0.428844393 |
| ENSMUSG000000067 | chr11 | Krt23 | -1.115263843 | 0.428851138 |
| ENSMUSG00000002711 | chr2 | Itga6 | -0.401603511 | 0.42910534 |
| ENSMUSG000000704S | chr5 | Tmem132b | -0.974545257 | 0.42910534 |
| ENSMUSG000000027E | chr17 | Mea1 | 0.32504934 | 0.42910534 |
| ENSMUSG000000384E | chr3 | Sv2a | -0.311203149 | 0.429510239 |
| ENSMUSG000000289 | chr5 | Smarcd3 | 0.304245575 | 0.429779924 |
| ENSMUSG000000241 | chr17 | Slc3a1 | -1.239839185 | 0.429845482 |
| ENSMUSG000000198E | chr10 | Ptprk | 0.191688361 | 0.429857882 |
| ENSMUSG000000173 | chr11 | Nlk | 0.22022762 | 0.429991189 |
| ENSMUSG000000009E | chr14 | Mmp14 | 0.304261165 | 0.430015796 |
| ENSMUSG000000382S | chr3 | Pdzk1 | 1.998615814 | 0.430015796 |
| ENSMUSG000000415 | chr5 | Hspb8 | 0.315699926 | 0.430021673 |
| ENSMUSG000000503S | chr4 | Nhsl3 | -0.430189185 | 0.430086222 |
| ENSMUSG000000124E | chr6 | Rpa3 | 0.296656574 | 0.430086222 |
| ENSMUSG000000278C | chr3 | Tsc22d2 | 0.26620606 | 0.430715054 |
| ENSMUSG000000461E | chr9 | Gldn | -1.273617931 | 0.430762479 |
| ENSMUSG000000213 | chr13 | Scgn | -0.229510987 | 0.430762479 |
| ENSMUSG000000391C | chr15 | Marchf6 | -0.130730872 | 0.430762479 |
| ENSMUSG000000436 | chr10 | Diras1 | -0.368411182 | 0.43089091 |
| ENSMUSG00000005461 | chr7 | Mgmt | 0.35278068 | 0.43089759 |
| ENSMUSG000000288E | chr4 | Gpatch3 | 0.381032398 | 0.4311263 |
| ENSMUSG000000284 | chr4 | Hint2 | 0.406813607 | 0.43121954 |
| ENSMUSG000000793E | chr9 | Ackr4 | 0.401763832 | 0.43121954 |
| ENSMUSG000000407S | chr6 | Iqsec3 | -0.360046336 | 0.431243133 |
| ENSMUSG000000066S | chr4 | Cdc42 | 0.244267389 | 0.431243133 |
| ENSMUSG000000359E | chr8 | Tmem59l | -0.290249117 | 0.431429369 |
| ENSMUSG000000226E | chr16 | Retnlb | -2.64169623 | 0.431691547 |
| ENSMUSG000000301C | chr6 | Edem1 | -0.24471716 | 0.431691547 |
| ENSMUSG000000359 | chr17 | H2-Q4 | -0.339919345 | 0.431740612 |
| ENSMUSG000000523 | chrX | Il1rapl1 | -0.527745158 | 0.431951627 |
| ENSMUSG000000255 | chr14 | Tm9sf2 | -0.199708523 | 0.432079252 |
| ENSMUSG000000316E | chr8 | Snx20 | -0.572362054 | 0.432134126 |
| ENSMUSG000000195 | chr6 | Pdk4 | 0.574997906 | 0.432134126 |
| ENSMUSG00000001211 | chr16 | Med15 | 0.179430541 | 0.432410705 |
| ENSMUSG000000197E | chr10 | Ccdc170 | -0.557498738 | 0.432416008 |
| ENSMUSG000000049E | chr7 | Syt5 | -0.307632103 | 0.432552197 |
| ENSMUSG000000002E | chr13 | Gmpr | 0.257867443 | 0.432707593 |
| ENSMUSG000000222C | chr14 | Ngdn | 0.30571171 | 0.432707593 |
| ENSMUSG000000465E | chr8 | Lrrc8e | -0.837935656 | 0.432746096 |
| ENSMUSG000000531S | chr3 | Mllt11 | -0.233369256 | 0.432792768 |
| ENSMUSG000000253E | chr10 | Mmp19 | 0.221303945 | 0.433488522 |
| ENSMUSG000000110C | chr3 | Mcoln2 | -1.374525938 | 0.433624652 |
| ENSMUSG000000460 | chr7 | Stard5 | -0.407110318 | 0.433624652 |

| | | | | |
|---|---|---|---|---|
| ENSMUSG0000003092 | chr7 | Trim30a | -0.449080777 | 0.433745696 |
| ENSMUSG0000002983 | chr6 | Svopl | -0.414219959 | 0.43384917 |
| ENSMUSG0000004670 | chr5 | Mapk10 | -0.243614635 | 0.434243065 |
| ENSMUSG0000002234 | chr15 | Sybu | -0.238181764 | 0.434294934 |
| ENSMUSG0000002743 | chr2 | Napb | -0.207833231 | 0.434294934 |
| ENSMUSG0000002079 | chr11 | Ube2g1 | 0.202490878 | 0.434294934 |
| ENSMUSG0000004454 | chr12 | Dact1 | 0.233744712 | 0.434294934 |
| ENSMUSG0000003060 | chr7 | Aen | 0.268227411 | 0.434294934 |
| ENSMUSG0000000567 | chr1 | Tomm40l | 0.305286873 | 0.434294934 |
| ENSMUSG0000004268 | chr14 | Selenok | 0.303570015 | 0.434294934 |
| ENSMUSG0000002676 | chr2 | Mmadhc | 0.312808948 | 0.434294934 |
| ENSMUSG0000002286 | chr16 | Chodl | 0.363151501 | 0.434294934 |
| ENSMUSG0000007050 | chr7 | Rgma | 0.201779941 | 0.435060503 |
| ENSMUSG0000003644 | chr8 | Thap11 | 0.241630875 | 0.435060503 |
| ENSMUSG0000007149 | chr19 | Nutf2-ps1 | -1.21774639 | 0.435184636 |
| ENSMUSG0000005104 | chr11 | Gprc5c | -0.362792767 | 0.435184636 |
| ENSMUSG0000002731 | chr2 | Dll4 | -0.367648531 | 0.435184636 |
| ENSMUSG0000003339 | chr9 | Clasp2 | -0.227924287 | 0.435184636 |
| ENSMUSG0000002722 | chr2 | Chst1 | 0.251959273 | 0.435184636 |
| ENSMUSG0000002786 | chr3 | Gdap2 | 0.225450028 | 0.435184636 |
| ENSMUSG0000000334 | chr10 | Abhd17a | 0.233185017 | 0.435184636 |
| ENSMUSG0000002577 | chr1 | Crispld1 | -0.361536399 | 0.435581119 |
| ENSMUSG0000004904 | chrX | Armcx3 | 0.150470174 | 0.435760018 |
| ENSMUSG0000002553 | chr5 | Phkg1 | -0.666786049 | 0.435900933 |
| ENSMUSG0000004274 | chr13 | Sgtb | -0.304747526 | 0.435970605 |
| ENSMUSG0000002398 | chr17 | Bysl | 0.320949116 | 0.435970605 |
| ENSMUSG0000005023 | chr4 | Gja4 | 0.518814414 | 0.436034144 |
| ENSMUSG0000002838 | chr4 | Ptbp3 | -0.190746371 | 0.436312167 |
| ENSMUSG0000000117 | chrX | Ocrl | -0.203474598 | 0.436669428 |
| ENSMUSG0000007478 | chr10 | Plxnc1 | 0.374320086 | 0.436669428 |
| ENSMUSG0000003178 | chr8 | Polr2c | 0.356593218 | 0.437604208 |
| ENSMUSG0000007385 | chr7 | Iqck | -0.566523638 | 0.437650434 |
| ENSMUSG0000003412 | chr11 | Mks1 | -0.399338036 | 0.437650434 |
| ENSMUSG0000002067 | chr12 | Tpo | 1.287821319 | 0.437650434 |
| ENSMUSG0000004027 | chr5 | Cdk6 | 0.252528933 | 0.43774 |
| ENSMUSG0000001175 | chr7 | Sptbn4 | -0.365181801 | 0.438357195 |
| ENSMUSG0000003584 | chr17 | Ddx11 | -0.439799232 | 0.438357195 |
| ENSMUSG0000002778 | chr3 | Nmd3 | 0.195246535 | 0.438357195 |
| ENSMUSG0000002430 | chr18 | Dtna | 0.210909162 | 0.438357195 |
| ENSMUSG0000007500 | chr10 | Nrbf2 | 0.237465066 | 0.438357195 |
| ENSMUSG0000001520 | chr10 | Cnksr3 | 0.325929467 | 0.438357195 |
| ENSMUSG0000004214 | chr11 | Cox10 | 0.19708599 | 0.438694105 |
| ENSMUSG0000005355 | chr2 | Ebf4 | -0.287797298 | 0.438907991 |
| ENSMUSG0000002682 | chr2 | Galnt5 | -0.897202924 | 0.438930033 |
| ENSMUSG0000003544 | chr11 | Myo1d | -0.395283679 | 0.438930033 |
| ENSMUSG0000004972 | chr11 | Gal3st1 | -0.524420899 | 0.438930033 |
| ENSMUSG0000003886 | chr2 | Garnl3 | -0.278602459 | 0.438930033 |
| ENSMUSG0000005621 | chr6 | Lrguk | -0.398563912 | 0.438930033 |
| ENSMUSG0000004837 | chr8 | Pdp2 | -0.305497587 | 0.438930033 |
| ENSMUSG0000000939 | chr6 | Syn2 | -0.256307737 | 0.438930033 |

| | | | | |
|---|---|---|---|---|
| ENSMUSG00000002838 | chr4 | Ugcg | -0.246039287 | 0.438930033 |
| ENSMUSG00000002978 | chr6 | Fkbp9 | 0.157423793 | 0.438930033 |
| ENSMUSG00000003219 | chr9 | Kank2 | 0.220958715 | 0.438930033 |
| ENSMUSG00000003470 | chr11 | Grn | 0.230889233 | 0.438930033 |
| ENSMUSG00000004942 | chr7 | Zfp260 | 0.223686917 | 0.438930033 |
| ENSMUSG00000002585 | chr5 | Get4 | 0.219294571 | 0.438930033 |
| ENSMUSG00000003796 | chr13 | Card19 | 0.227859691 | 0.438930033 |
| ENSMUSG00000002918 | chr5 | Fam114a1 | 0.267382378 | 0.438930033 |
| ENSMUSG00000002316 | chr15 | Slc38a1 | 0.31161131 | 0.438930033 |
| ENSMUSG00000004555 | chr10 | Mettl24 | 0.323105187 | 0.438930033 |
| ENSMUSG00000002235 | chr15 | Ndufb9 | 0.364442169 | 0.438930033 |
| ENSMUSG00000007100 | chr4 | Hrct1 | 0.552771501 | 0.438930033 |
| ENSMUSG00000000786 | chr12 | Ift43 | 0.416127238 | 0.438930033 |
| ENSMUSG00000002407 | chr17 | Crim1 | 0.256852694 | 0.438946646 |
| ENSMUSG00000001765 | chr2 | Cd40 | 0.383366616 | 0.438946646 |
| ENSMUSG00000000489 | chr3 | Prcc | 0.211617068 | 0.439041711 |
| ENSMUSG00000003304 | chr15 | Eif3l | 0.232133233 | 0.439041711 |
| ENSMUSG00000002644 | chr1 | Nfasc | -0.319744595 | 0.439603027 |
| ENSMUSG00000006667 | chr1 | Ifi208 | -0.620893881 | 0.439603027 |
| ENSMUSG00000002261 | chr15 | Mapk8ip2 | -0.303374161 | 0.439603027 |
| ENSMUSG00000002205 | chr14 | Nefl | -0.275073368 | 0.439603027 |
| ENSMUSG00000002665 | chr1 | Fcgr2b | -0.247893055 | 0.439603027 |
| ENSMUSG00000006178 | chr7 | Rps17 | 0.295600827 | 0.439603027 |
| ENSMUSG00000006147 | chr13 | Mrps36 | 0.348155468 | 0.439603027 |
| ENSMUSG00000003175 | chr8 | Cenpn | 0.785226604 | 0.439603027 |
| ENSMUSG00000005675 | chr6 | Grm7 | -0.263487176 | 0.439740149 |
| ENSMUSG00000004097 | chr11 | Gm11992 | -0.402167559 | 0.439740149 |
| ENSMUSG00000002499 | chr19 | Plce1 | -0.204195473 | 0.439741715 |
| ENSMUSG00000005711 | chr11 | Npm1 | 0.284303187 | 0.439852513 |
| ENSMUSG00000004235 | chr14 | Gnl3 | 0.192278974 | 0.439969824 |
| ENSMUSG00000003088 | chr7 | Uqcrc2 | 0.258588878 | 0.439969824 |
| ENSMUSG00000003748 | chr2 | Erv3 | -0.707916968 | 0.440087382 |
| ENSMUSG00000002396 | chr17 | Mrps18a | 0.238734424 | 0.44023499 |
| ENSMUSG00000004951 | chr13 | Rps23 | 0.411562946 | 0.440651203 |
| ENSMUSG00000002722 | chr2 | Duoxa2 | -0.810021517 | 0.440838649 |
| ENSMUSG00000000595 | chr16 | Ephb3 | -0.266545896 | 0.440838649 |
| ENSMUSG00000000189 | chr11 | Ugp2 | 0.31929147 | 0.440838649 |
| ENSMUSG00000002451 | chr18 | Smad4 | 0.195481418 | 0.440945621 |
| ENSMUSG00000004105 | chr4 | Slc7a13 | -1.502776755 | 0.441280364 |
| ENSMUSG00000002025 | chr10 | Aldh1l2 | 0.432541814 | 0.441315731 |
| ENSMUSG00000002896 | chr4 | Park7 | 0.356037458 | 0.441315731 |
| ENSMUSG00000002054 | chr11 | Tom1l1 | -0.338767363 | 0.441507551 |
| ENSMUSG00000003112 | chr9 | Slc9a9 | -0.378780232 | 0.441697448 |
| ENSMUSG00000002733 | chr2 | Cdc25b | -0.28040363 | 0.441697448 |
| ENSMUSG00000000179 | chr7 | Capns1 | 0.199707099 | 0.441697448 |
| ENSMUSG00000002277 | chr16 | Dlg1 | 0.210813676 | 0.441697448 |
| ENSMUSG00000001964 | chr18 | Sema6a | -0.283881359 | 0.441766386 |
| ENSMUSG00000000319 | chr7 | Bcl2l12 | 0.267410689 | 0.441766386 |
| ENSMUSG00000002176 | chr13 | Ndufs4 | 0.310918919 | 0.442031607 |
| ENSMUSG00000002903 | chr4 | Ssu72 | 0.243034244 | 0.442163828 |

| | | | | |
|---|---|---|---|---|
| ENSMUSG0000003707 | chr19 | Scd1 | 0.362328511 | 0.442500769 |
| ENSMUSG0000000633 | chr7 | Rps9 | 0.362461658 | 0.442509613 |
| ENSMUSG0000004126 | chr4 | Car8 | -0.335687835 | 0.443148313 |
| ENSMUSG0000002435 | chr18 | Sil1 | 0.241660732 | 0.443148313 |
| ENSMUSG0000010691 | chr5 | Mrpl33 | 0.305503348 | 0.443148313 |
| ENSMUSG0000003799 | chr4 | Slc24a2 | 0.410962166 | 0.443258952 |
| ENSMUSG0000000346 | chr14 | Phyhip | -0.580838262 | 0.443280148 |
| ENSMUSG0000011427 | chr5 | Gm49027 | -0.38995998 | 0.443280148 |
| ENSMUSG0000001839 | chr11 | Kif3a | -0.215056955 | 0.443280148 |
| ENSMUSG0000003642 | chr7 | Gpi1 | 0.133825202 | 0.443280148 |
| ENSMUSG0000007375 | chr4 | Sh3d21 | 0.321936538 | 0.443280148 |
| ENSMUSG0000005409 | chr3 | 1810037I17Rik | 0.350407483 | 0.443280148 |
| ENSMUSG0000002451 | chr18 | Mbd2 | 0.163378116 | 0.443542955 |
| ENSMUSG0000005010 | chr12 | Agmo | -0.182530065 | 0.443995693 |
| ENSMUSG0000002564 | chr9 | Pfkfb4 | -0.366906735 | 0.444000731 |
| ENSMUSG0000005278 | chr5 | Grk4 | 0.390642595 | 0.444000731 |
| ENSMUSG0000001501 | chr8 | Trappc2l | 0.255992 | 0.444540013 |
| ENSMUSG0000004810 | chr6 | Tmem72 | 1.189551949 | 0.444624975 |
| ENSMUSG0000001856 | chr11 | Gabarap | 0.213750237 | 0.444779491 |
| ENSMUSG0000000789 | chr9 | Rplp1 | 0.316101325 | 0.445081394 |
| ENSMUSG0000002602 | chr1 | Trak2 | 0.190167531 | 0.445300023 |
| ENSMUSG0000000492 | chr10 | Thop1 | 0.249167805 | 0.445300023 |
| ENSMUSG0000003009 | chr6 | Grip2 | 0.264050561 | 0.445300023 |
| ENSMUSG0000002689 | chr2 | Ndufa8 | 0.294146754 | 0.445300023 |
| ENSMUSG0000004249 | chrX | Radx | -0.833249758 | 0.445624222 |
| ENSMUSG0000002065 | chr12 | Efr3b | -0.332977886 | 0.445914304 |
| ENSMUSG0000003732 | chr17 | Tap1 | -0.368308757 | 0.445914304 |
| ENSMUSG0000003797 | chr9 | 6430571L13Rik | -0.57456012 | 0.445914304 |
| ENSMUSG0000002941 | chr5 | Naaa | 0.199350943 | 0.445914304 |
| ENSMUSG0000002693 | chr2 | Nacc2 | 0.198120207 | 0.445980799 |
| ENSMUSG0000003906 | chr8 | Psmd7 | 0.240160045 | 0.445980799 |
| ENSMUSG0000003882 | chr4 | Abitram | 0.196986005 | 0.446425377 |
| ENSMUSG0000002044 | chr11 | Guk1 | 0.345941899 | 0.446743273 |
| ENSMUSG0000003608 | chr5 | Slain2 | 0.16738288 | 0.447053365 |
| ENSMUSG0000002063 | chr12 | Ubxn2a | 0.199349804 | 0.447053365 |
| ENSMUSG0000004236 | chr11 | Lgalsl | 0.221160927 | 0.447053365 |
| ENSMUSG0000002120 | chr12 | Asb2 | 0.258508446 | 0.447053365 |
| ENSMUSG0000002496 | chr19 | Dnajc4 | 0.251041581 | 0.447053365 |
| ENSMUSG0000002038 | chr11 | Sar1b | 0.296706127 | 0.447053365 |
| ENSMUSG0000001740 | chr11 | Stac2 | -0.571441827 | 0.447097703 |
| ENSMUSG0000005929 | chr4 | Rpl11 | 0.322480179 | 0.447596454 |
| ENSMUSG0000003620 | chr1 | Sh3bp4 | 0.197904486 | 0.447959366 |
| ENSMUSG0000002889 | chr4 | Taf12 | 0.291926041 | 0.448254101 |
| ENSMUSG0000010586 | chr5 | Tmem217rt | -0.405582974 | 0.448339244 |
| ENSMUSG0000005106 | chr11 | Fbll1 | -0.501549315 | 0.448339244 |
| ENSMUSG0000001359 | chr1 | Ndufs2 | 0.260421091 | 0.448339244 |
| ENSMUSG0000009026 | chr11 | Mettl23 | 0.429465205 | 0.448339244 |
| ENSMUSG0000003489 | chr12 | Rps29 | 0.417289974 | 0.448339244 |
| ENSMUSG0000009618 | chr8 | Cmtm4 | -0.264289011 | 0.448450176 |
| ENSMUSG0000003394 | chr18 | Trim36 | -0.430824501 | 0.448450541 |

| | | | | |
|---|---|---|---|---|
| ENSMUSG0000003433 | chr8 | Plcg2 | -0.483559262 | 0.448450541 |
| ENSMUSG0000003422 | chr2 | Rhov | -0.767276108 | 0.448450541 |
| ENSMUSG0000003242 | chr9 | Nt5e | 0.254035871 | 0.448450541 |
| ENSMUSG0000005067 | chr12 | Ism2 | 0.546795101 | 0.448450541 |
| ENSMUSG0000002230 | chr15 | Zfpm2 | 0.262236965 | 0.449073791 |
| ENSMUSG0000002922 | chr5 | Chic2 | 0.177013733 | 0.449098947 |
| ENSMUSG0000002498 | chr19 | Hhex | -0.477047278 | 0.449649779 |
| ENSMUSG0000007473 | chr19 | Zfp950 | -0.210225589 | 0.449649779 |
| ENSMUSG0000002163 | chr13 | Rad17 | 0.176601102 | 0.449649779 |
| ENSMUSG0000004964 | chr7 | 2310022A10Rik | 0.197036055 | 0.450087202 |
| ENSMUSG0000003184 | chr8 | Mphosph6 | 0.22588906 | 0.450356997 |
| ENSMUSG0000006387 | chr2 | Slc24a3 | 0.181690169 | 0.450468076 |
| ENSMUSG0000002635 | chr1 | Mcm6 | -0.364286182 | 0.450625199 |
| ENSMUSG0000003850 | chr6 | Parp12 | -0.18419741 | 0.450648265 |
| ENSMUSG0000005648 | chr2 | Chn1 | -0.213277733 | 0.450662523 |
| ENSMUSG0000002032 | chr11 | Nudcd2 | 0.251758408 | 0.450662523 |
| ENSMUSG0000002558 | chr11 | Nptx1 | 0.299505812 | 0.450662523 |
| ENSMUSG0000006198 | chr10 | Rps12 | 0.347632119 | 0.450662523 |
| ENSMUSG0000003088 | chr7 | Dnhd1 | -0.431372125 | 0.450761261 |
| ENSMUSG0000002642 | chr1 | Eif2d | 0.134173315 | 0.450761261 |
| ENSMUSG0000003421 | chr2 | Vps18 | 0.180470421 | 0.450761261 |
| ENSMUSG0000002959 | chr5 | Rbm19 | 0.230877771 | 0.450761261 |
| ENSMUSG0000003678 | chr9 | Rps27l | 0.419444273 | 0.451347635 |
| ENSMUSG0000004154 | chr6 | Sox5 | -0.240931833 | 0.451536612 |
| ENSMUSG0000002535 | chr10 | Cd63 | 0.300165878 | 0.451536612 |
| ENSMUSG0000003853 | chr11 | Osbpl7 | -0.415526457 | 0.451612921 |
| ENSMUSG0000003260 | chr9 | Qars1 | 0.216449276 | 0.451612921 |
| ENSMUSG0000002213 | chr14 | Gpr180 | 0.278266979 | 0.451612921 |
| ENSMUSG0000003716 | chr7 | Ppp1r14a | 0.398550335 | 0.451612921 |
| ENSMUSG0000002775 | chr3 | Postn | 0.318128078 | 0.451649086 |
| ENSMUSG0000003798 | chr13 | Wnk2 | 0.297524883 | 0.451649086 |
| ENSMUSG0000000658 | chr4 | Runx1t1 | 0.268251975 | 0.451649086 |
| ENSMUSG0000002635 | chr1 | Lct | 0.929432116 | 0.451649086 |
| ENSMUSG0000003410 | chr19 | Ccs | 0.33522792 | 0.451649086 |
| ENSMUSG0000002790 | chr3 | Dram2 | -0.180449198 | 0.451869782 |
| ENSMUSG0000003455 | chrX | Hdx | 0.444826362 | 0.451869782 |
| ENSMUSG0000007039 | chr11 | Nlrp1b | -1.32898261 | 0.451894667 |
| ENSMUSG0000002602 | chr1 | Stradb | 0.224944033 | 0.451894667 |
| ENSMUSG0000006127 | chrX | Mmgt1 | 0.150101162 | 0.452355419 |
| ENSMUSG0000007406 | chr8 | Mlycd | 0.28566344 | 0.452355419 |
| ENSMUSG0000002885 | chr4 | Nudc | 0.203232308 | 0.452472126 |
| ENSMUSG0000003815 | chr19 | Gstp2 | -0.874533707 | 0.452595838 |
| ENSMUSG0000003897 | chr5 | Lmtk2 | -0.209290552 | 0.452602104 |
| ENSMUSG0000006892 | chr3 | Syt11 | -0.267996974 | 0.452806735 |
| ENSMUSG0000003053 | chr7 | Cib1 | -0.368357413 | 0.452816254 |
| ENSMUSG0000006974 | chr11 | Psmb3 | 0.334833766 | 0.452816254 |
| ENSMUSG0000004176 | chr2 | Gpr155 | -0.252063037 | 0.452920505 |
| ENSMUSG0000006209 | chr2 | Btbd3 | 0.175912384 | 0.452920505 |
| ENSMUSG0000004455 | chr14 | 9930012K11Rik | -0.791858799 | 0.45338339 |
| ENSMUSG0000006908 | chr13 | Cdk7 | 0.207695169 | 0.45338339 |

| | | | | |
|---|---|---|---|---|
| ENSMUSG0000006879 | chr3 | Rap1a | 0.297258149 | 0.45338339 |
| ENSMUSG0000003694 | chr4 | Kdm1a | 0.138226208 | 0.453936616 |
| ENSMUSG0000004876 | chr11 | Hoxb3 | -0.274976062 | 0.4539667 |
| ENSMUSG0000006059 | chr9 | Layn | -0.267479881 | 0.454075862 |
| ENSMUSG0000001008 | chr11 | Epn3 | -0.603580385 | 0.454077194 |
| ENSMUSG0000003707 | chr3 | Selenof | 0.304157458 | 0.454077194 |
| ENSMUSG0000004965 | chr11 | Aftph | -0.142546385 | 0.454103913 |
| ENSMUSG0000002929 | chr5 | Gbp9 | -0.627657875 | 0.454499938 |
| ENSMUSG0000003995 | chr5 | Hip1 | 0.218155978 | 0.454672861 |
| ENSMUSG0000002531 | chr8 | Jph3 | -0.306945608 | 0.455132167 |
| ENSMUSG0000004953 | chr13 | H1f1 | -1.325886331 | 0.455132167 |
| ENSMUSG0000006980 | chr7 | Cacng7 | -0.209456435 | 0.455132167 |
| ENSMUSG0000005931 | chr2 | Slc27a4 | -0.211103269 | 0.455132167 |
| ENSMUSG0000003005 | chr6 | Copg1 | 0.172168943 | 0.455132167 |
| ENSMUSG0000002466 | chr19 | Fth1 | 0.272081244 | 0.455132167 |
| ENSMUSG0000002434 | chr18 | Pfdn1 | 0.26658536 | 0.455132167 |
| ENSMUSG0000004105 | chr17 | Wdr43 | 0.140395985 | 0.456425789 |
| ENSMUSG0000003149 | chr8 | Cd209a | -0.631358046 | 0.456456596 |
| ENSMUSG0000002204 | chr14 | Stmn4 | -0.279023844 | 0.456456596 |
| ENSMUSG0000000351 | chr11 | Dusp3 | 0.231956683 | 0.456456596 |
| ENSMUSG0000003627 | chr19 | Macrod1 | -0.226605686 | 0.456459474 |
| ENSMUSG0000002295 | chr16 | Rcan1 | 0.269036174 | 0.456459474 |
| ENSMUSG0000002955 | chr5 | Mrm2 | 0.325841422 | 0.456459474 |
| ENSMUSG0000003334 | chr3 | Plppr5 | -0.373458644 | 0.456759767 |
| ENSMUSG0000001548 | chr1 | Fam163a | -0.326113308 | 0.456759767 |
| ENSMUSG0000002612 | chr1 | Plekhb2 | 0.265574543 | 0.456759767 |
| ENSMUSG0000006221 | chr18 | Tnfaip8 | 0.294879837 | 0.456759767 |
| ENSMUSG0000004721 | chr5 | Rpl9 | 0.408728205 | 0.456759767 |
| ENSMUSG0000002113 | chr12 | Erh | 0.365597591 | 0.456759767 |
| ENSMUSG0000005856 | chr13 | Tmed9 | 0.282812791 | 0.456853237 |
| ENSMUSG0000002173 | chr13 | Hcn1 | 0.368158876 | 0.457626812 |
| ENSMUSG0000004022 | chr8 | Gas8 | 0.175858645 | 0.457634016 |
| ENSMUSG0000005031 | chr3 | Synpo2 | 0.253579607 | 0.457742628 |
| ENSMUSG0000002958 | chr5 | Actb | 0.294905083 | 0.458460411 |
| ENSMUSG0000002961 | chr5 | Rpl6 | 0.280234145 | 0.458984149 |
| ENSMUSG0000011572 | chr14 | Gm30083 | 0.381320988 | 0.458984149 |
| ENSMUSG0000002647 | chr1 | Mr1 | 0.229626018 | 0.45907481 |
| ENSMUSG0000007370 | chr1 | Rpl31 | 0.323032348 | 0.45907481 |
| ENSMUSG0000000539 | chr13 | Nid1 | 0.249194043 | 0.459200297 |
| ENSMUSG0000002800 | chr3 | Npy2r | -0.39886308 | 0.459338122 |
| ENSMUSG0000003272 | chr4 | Bmp8a | -2.08435557 | 0.459338122 |
| ENSMUSG0000003576 | chr9 | Xylb | -0.379253864 | 0.459338122 |
| ENSMUSG0000001917 | chr11 | Rab5c | 0.185364061 | 0.459338122 |
| ENSMUSG0000002879 | chr4 | Tmem234 | 0.217538141 | 0.459338122 |
| ENSMUSG0000002457 | chr18 | Rbfa | 0.234247344 | 0.459338122 |
| ENSMUSG0000002869 | chr4 | Prdx1 | 0.272456637 | 0.459338122 |
| ENSMUSG0000003354 | chr8 | Znrf1 | 0.225741264 | 0.459338122 |
| ENSMUSG0000002272 | chr16 | Crybg3 | 0.267790967 | 0.459338122 |
| ENSMUSG0000007165 | chr19 | Uqcc3 | 0.352454836 | 0.459338122 |
| ENSMUSG0000004188 | chr17 | Ndufa7 | 0.393691647 | 0.459338122 |

| | | | | |
|---|---|---|---|---|
| ENSMUSG0000004085 | chr12 | Dlk1 | 1.079944671 | 0.459338122 |
| ENSMUSG0000004904 | chr2 | Rapgef4 | -0.204955829 | 0.459441708 |
| ENSMUSG0000003927 | chrX | Pcsk1n | -0.29242857 | 0.459633727 |
| ENSMUSG0000002992 | chr6 | Slc37a3 | 0.213103942 | 0.459814768 |
| ENSMUSG0000000261 | chr17 | Zfp40 | -0.235196406 | 0.460109377 |
| ENSMUSG0000002586 | chr13 | Cplx2 | -0.211890182 | 0.460344576 |
| ENSMUSG0000002675 | chr2 | Psmb7 | 0.250284923 | 0.460344576 |
| ENSMUSG0000005060 | chr4 | Micos10 | 0.293740915 | 0.460344576 |
| ENSMUSG0000000090 | chr10 | Smarcb1 | 0.240575089 | 0.460422028 |
| ENSMUSG0000004848 | chr12 | Zdhhc22 | -0.722446946 | 0.460558222 |
| ENSMUSG0000005369 | chr8 | Mast1 | -0.344999271 | 0.460558222 |
| ENSMUSG0000003073 | chr7 | Eif3c | 0.1293687 | 0.460558222 |
| ENSMUSG0000000260 | chr7 | Ccdc97 | 0.140605283 | 0.460558222 |
| ENSMUSG0000002464 | chr18 | Cndp2 | 0.155159573 | 0.460558222 |
| ENSMUSG0000002835 | chr4 | Brinp1 | -0.35024503 | 0.460560989 |
| ENSMUSG0000003629 | chr14 | Ap5m1 | -0.178634987 | 0.460560989 |
| ENSMUSG0000003443 | chr1 | Cops8 | 0.218798146 | 0.460770489 |
| ENSMUSG0000002263 | chr15 | Zcrb1 | 0.246968696 | 0.460770489 |
| ENSMUSG0000003024 | chr6 | Kcnj8 | 0.339839938 | 0.460782223 |
| ENSMUSG0000003258 | chr9 | Bsn | -0.385949023 | 0.460942959 |
| ENSMUSG0000002277 | chr16 | Ypel1 | -0.355142042 | 0.460942959 |
| ENSMUSG0000002910 | chr5 | Lrpap1 | 0.175317249 | 0.460942959 |
| ENSMUSG0000004726 | chr8 | Zfp358 | 0.222171362 | 0.460942959 |
| ENSMUSG0000002632 | chr18 | Htr4 | -0.547609421 | 0.46139184 |
| ENSMUSG0000004860 | chr17 | Morc2b | -1.274289599 | 0.46164426 |
| ENSMUSG0000004276 | chr4 | Maneal | -0.402675496 | 0.46175983 |
| ENSMUSG0000003562 | chr10 | Midn | -0.250380637 | 0.46175983 |
| ENSMUSG0000005022 | chr1 | Pigm | -0.228457052 | 0.46175983 |
| ENSMUSG0000002053 | chr11 | Drg2 | 0.16108069 | 0.46175983 |
| ENSMUSG0000004667 | chr12 | Lyset | 0.201955305 | 0.46175983 |
| ENSMUSG0000000043 | chr6 | Mkrn2 | 0.185475673 | 0.46175983 |
| ENSMUSG0000005299 | chr7 | Uba2 | 0.212051395 | 0.46175983 |
| ENSMUSG0000003081 | chr7 | Bcl7c | 0.252960794 | 0.46175983 |
| ENSMUSG0000002558 | chr11 | Eif4a3 | 0.279099631 | 0.46175983 |
| ENSMUSG0000002756 | chr2 | Psma7 | 0.322383878 | 0.46175983 |
| ENSMUSG0000007501 | chr2 | Fjx1 | 0.41919286 | 0.46175983 |
| ENSMUSG0000002683 | chr2 | Acvr1 | 0.133694917 | 0.462144104 |
| ENSMUSG0000004182 | chr11 | Abca8a | -0.484700474 | 0.462350052 |
| ENSMUSG0000007959 | chr6 | Clec2l | -0.477635712 | 0.462350052 |
| ENSMUSG0000001341 | chr11 | Igf2bp1 | -0.793511726 | 0.462350052 |
| ENSMUSG0000009011 | chr17 | Usp49 | -0.311412179 | 0.462350052 |
| ENSMUSG0000004104 | chr1 | Fam117b | -0.208518486 | 0.462350052 |
| ENSMUSG0000001566 | chrX | Pdzd11 | 0.19098041 | 0.462350052 |
| ENSMUSG0000002931 | chr5 | Aff1 | 0.20855693 | 0.462350052 |
| ENSMUSG0000002931 | chr5 | Gpat3 | 0.265426015 | 0.462350052 |
| ENSMUSG0000003420 | chr11 | Rasl10a | 0.402651952 | 0.462350052 |
| ENSMUSG0000006818 | chr13 | Ndufaf2 | 0.352212309 | 0.462350052 |
| ENSMUSG0000003102 | chr7 | BC051019 | 1.783517339 | 0.462350052 |
| ENSMUSG0000000462 | chr8 | Stxbp2 | -0.550850619 | 0.462740109 |
| ENSMUSG0000002305 | chr15 | Map3k12 | -0.219532883 | 0.463519724 |

| | | | | |
|---|---|---|---|---|
| ENSMUSG0000002774 | chr3 | Ufm1 | 0.266193748 | 0.463519724 |
| ENSMUSG0000003881 | chr4 | Ctnnal1 | -0.186168619 | 0.464054219 |
| ENSMUSG0000006286 | chr9 | Impdh2 | 0.304533585 | 0.464402091 |
| ENSMUSG0000005789 | chr11 | Camk2b | -0.299643675 | 0.46539373 |
| ENSMUSG0000002513 | chr11 | P4hb | 0.242568365 | 0.46539373 |
| ENSMUSG0000002842 | chr4 | Tmem38b | 0.252439672 | 0.46539373 |
| ENSMUSG0000002419 | chr17 | Cuta | 0.216600958 | 0.465697289 |
| ENSMUSG0000003226 | chr9 | Sh3bgrl2 | -0.402289408 | 0.46577652 |
| ENSMUSG0000002982 | chr6 | Zc3hav1 | -0.238805123 | 0.46577652 |
| ENSMUSG0000007570 | chr11 | Selenom | 0.334258713 | 0.46577652 |
| ENSMUSG0000002707 | chr2 | Clp1 | 0.259879217 | 0.466323663 |
| ENSMUSG0000002446 | chr17 | Gabbr1 | -0.269077491 | 0.466771145 |
| ENSMUSG0000002307 | chr8 | Slc10a2 | -1.972877844 | 0.467264969 |
| ENSMUSG0000004740 | chr17 | Tgif1 | 0.264843251 | 0.467264969 |
| ENSMUSG0000003971 | chr9 | Dock3 | -0.365121559 | 0.467347669 |
| ENSMUSG0000005376 | chr6 | Chchd3 | 0.226387808 | 0.467446733 |
| ENSMUSG0000002554 | chr14 | Clybl | 0.31859652 | 0.467645415 |
| ENSMUSG0000004174 | chr5 | Rnf10 | 0.169088752 | 0.467690996 |
| ENSMUSG0000005402 | chr10 | Nt5dc3 | 0.254402012 | 0.467690996 |
| ENSMUSG0000009021 | chr2 | Peds1 | -0.203621721 | 0.467820827 |
| ENSMUSG0000004233 | chr11 | Specc1 | -0.192033808 | 0.468111946 |
| ENSMUSG0000001888 | chr11 | Mrpl45 | 0.211871026 | 0.468303482 |
| ENSMUSG0000007082 | chr7 | Zscan18 | -0.26856457 | 0.468315507 |
| ENSMUSG0000003315 | chr1 | Cnppd1 | 0.195624568 | 0.468315507 |
| ENSMUSG0000002542 | chr18 | Pstpip2 | -0.883504314 | 0.468580815 |
| ENSMUSG0000005355 | chr5 | 3110082I17Rik | 0.206076063 | 0.468740636 |
| ENSMUSG0000000192 | chrX | Uba1 | 0.123130879 | 0.468846655 |
| ENSMUSG0000002886 | chr4 | Wasf2 | 0.197086696 | 0.468846655 |
| ENSMUSG0000002919 | chr5 | Grpel1 | 0.32330332 | 0.469102149 |
| ENSMUSG0000003205 | chr9 | Fdx1 | 0.285388421 | 0.469276624 |
| ENSMUSG0000009227 | chr17 | Gm19684 | -1.110159558 | 0.469596143 |
| ENSMUSG0000003064 | chr7 | Ndufc2 | 0.227575398 | 0.469596143 |
| ENSMUSG0000002718 | chr2 | Pamr1 | 0.251048203 | 0.469596143 |
| ENSMUSG0000003756 | chr7 | Rps16 | 0.418160836 | 0.469596143 |
| ENSMUSG0000005908 | chr1 | Fcgr4 | -0.991794256 | 0.469843586 |
| ENSMUSG0000004608 | chr6 | Clec9a | -0.828616705 | 0.469843586 |
| ENSMUSG0000000012 | chr11 | Ngfr | -0.296625485 | 0.469843586 |
| ENSMUSG0000000138 | chr18 | Zmat2 | 0.138815656 | 0.469843586 |
| ENSMUSG0000003242 | chr9 | Syncrip | 0.181656002 | 0.469843586 |
| ENSMUSG0000003102 | chr7 | Eif3f | 0.218512256 | 0.469843586 |
| ENSMUSG0000002743 | chr2 | Gzf1 | 0.202662677 | 0.469843586 |
| ENSMUSG0000000494 | chr5 | Dtx2 | 0.254884532 | 0.469843586 |
| ENSMUSG0000002047 | chr11 | Zkscan17 | 0.287820192 | 0.469843586 |
| ENSMUSG0000000103 | chr11 | Epn2 | 0.295510297 | 0.469843586 |
| ENSMUSG0000003373 | chr14 | Egr3 | 1.005295412 | 0.469843586 |
| ENSMUSG0000002486 | chr19 | Pip5k1b | -0.3157925 | 0.469884994 |
| ENSMUSG0000003016 | chr6 | Klrd1 | -1.654992814 | 0.469915447 |
| ENSMUSG0000002474 | chr19 | Syt7 | -0.398325658 | 0.469915447 |
| ENSMUSG0000007525 | chr2 | Cerkl | -0.752812945 | 0.469915447 |
| ENSMUSG0000004503 | chr7 | Megf8 | 0.195750399 | 0.469915447 |

| | | | | |
|---|---|---|---|---|
| ENSMUSG0000002673 | chr2 | Pter | 0.228494064 | 0.469915447 |
| ENSMUSG0000002429 | chr17 | Cyp4f14 | -2.313792599 | 0.47031903 |
| ENSMUSG0000003182 | chr8 | Gse1 | -0.338934035 | 0.47031903 |
| ENSMUSG0000003173 | chr8 | Phlpp2 | -0.290894677 | 0.47031903 |
| ENSMUSG0000002475 | chr19 | Rtn3 | -0.132473832 | 0.47031903 |
| ENSMUSG0000003078 | chr7 | Rnf141 | 0.162327829 | 0.47031903 |
| ENSMUSG0000007164 | chr19 | Eef1g | 0.271436227 | 0.47031903 |
| ENSMUSG0000008478 | chr9 | Ubl5 | 0.267015909 | 0.47031903 |
| ENSMUSG0000006728 | chr17 | Rps28 | 0.294142758 | 0.47031903 |
| ENSMUSG0000002117 | chr12 | Esyt2 | 0.139507852 | 0.470379682 |
| ENSMUSG0000002183 | chr14 | Ero1a | 0.122666789 | 0.470477286 |
| ENSMUSG0000004754 | chr13 | Cltb | 0.212319678 | 0.470477286 |
| ENSMUSG0000002561 | chr16 | Map3k7cl | 0.752457499 | 0.470566217 |
| ENSMUSG0000000765 | chr11 | Gabrb2 | -1.007659745 | 0.470657636 |
| ENSMUSG0000003134 | chrX | Gpm6b | -0.16347445 | 0.470830197 |
| ENSMUSG0000002012 | chr10 | Usp15 | 0.137382855 | 0.470921524 |
| ENSMUSG0000006595 | chr8 | Tacc1 | 0.202906555 | 0.471273502 |
| ENSMUSG0000006161 | chr17 | U2af1 | 0.218240918 | 0.471273502 |
| ENSMUSG0000003629 | chr12 | Lrrn3 | -0.30062381 | 0.471394597 |
| ENSMUSG0000007361 | chr1 | Cops9 | 0.520268763 | 0.471394597 |
| ENSMUSG0000001117 | chr12 | Vipr2 | 0.270344866 | 0.471560544 |
| ENSMUSG0000003272 | chr13 | Mier3 | -0.320906439 | 0.471607924 |
| ENSMUSG0000002920 | chr5 | Nsun7 | -0.451627078 | 0.471607924 |
| ENSMUSG0000003790 | chr5 | Bri3bp | -0.267517453 | 0.471607924 |
| ENSMUSG0000002102 | chr12 | Ppp2r3c | 0.187343362 | 0.471607924 |
| ENSMUSG0000002716 | chr2 | Iftap | 0.292165024 | 0.471607924 |
| ENSMUSG0000000008 | chr9 | Cox5a | 0.280579091 | 0.471607924 |
| ENSMUSG0000007487 | chr13 | Ctla2b | 0.904811641 | 0.471607924 |
| ENSMUSG0000003527 | chr7 | Ssc5d | 0.353732739 | 0.471607924 |
| ENSMUSG0000004622 | chr2 | Scand1 | 0.380329206 | 0.471607924 |
| ENSMUSG0000003715 | chr3 | Ndufc1 | 0.401032743 | 0.471607924 |
| ENSMUSG0000002687 | chr2 | Gsn | 0.250176573 | 0.471801209 |
| ENSMUSG0000005565 | chr7 | Klhl25 | 0.269723125 | 0.471801209 |
| ENSMUSG0000005660 | chr5 | Fry | 0.262880152 | 0.471801209 |
| ENSMUSG0000001796 | chr2 | Ptgis | 0.38018638 | 0.471973369 |
| ENSMUSG0000001444 | chr8 | Piezo1 | 0.307693674 | 0.472201246 |
| ENSMUSG0000002868 | chr4 | Urod | 0.252341459 | 0.472313564 |
| ENSMUSG0000005298 | chr13 | Ube2ql1 | -0.331615331 | 0.472359294 |
| ENSMUSG0000002979 | chr6 | Cep41 | -0.287657254 | 0.472359294 |
| ENSMUSG0000000298 | chr7 | Apoe | -0.217555866 | 0.472445303 |
| ENSMUSG0000005506 | chr9 | Rab39 | -0.642941345 | 0.47260907 |
| ENSMUSG0000001082 | chr5 | Grid2ip | -0.832753201 | 0.473503538 |
| ENSMUSG0000003056 | chr7 | Ctsc | -0.3039186 | 0.473503538 |
| ENSMUSG0000003993 | chr4 | Pik3cd | -0.282046805 | 0.473503538 |
| ENSMUSG0000007141 | chr11 | Rpl23 | 0.277851977 | 0.473503538 |
| ENSMUSG0000002656 | chr1 | Styxl2 | 0.430795319 | 0.473503538 |
| ENSMUSG0000002918 | chr5 | Slc34a2 | -2.067407874 | 0.473670513 |
| ENSMUSG0000005281 | chr8 | Best2 | -1.914717691 | 0.473670513 |
| ENSMUSG0000008707 | chr12 | Lbhd2 | -1.415515818 | 0.473670513 |
| ENSMUSG0000002293 | chr16 | Grik1 | -0.377775578 | 0.473670513 |

| | | | | |
|---|---|---|---|---|
| ENSMUSG0000005543 | chr6 | Nap1l5 | -0.286415521 | 0.473670513 |
| ENSMUSG0000004126 | chr9 | Dmxl2 | -0.260095621 | 0.473670513 |
| ENSMUSG0000006044 | chr2 | Sycp2 | -0.714741606 | 0.473670513 |
| ENSMUSG0000004446 | chr7 | Fhip1b | -0.198577094 | 0.473670513 |
| ENSMUSG0000000007 | chr13 | Klf6 | -0.190184915 | 0.473670513 |
| ENSMUSG0000003577 | chr8 | Dync1li2 | 0.147126768 | 0.473670513 |
| ENSMUSG0000002499 | chr19 | Eif3a | 0.160048146 | 0.473670513 |
| ENSMUSG0000002622 | chr1 | Psmd1 | 0.170910318 | 0.473670513 |
| ENSMUSG0000002855 | chr4 | Osbpl9 | 0.184268193 | 0.473670513 |
| ENSMUSG0000000298 | chr7 | Tomm40 | 0.176940716 | 0.473670513 |
| ENSMUSG0000003584 | chr13 | Lysmd3 | 0.219285862 | 0.473670513 |
| ENSMUSG0000002936 | chr5 | Wsb2 | 0.228539485 | 0.473670513 |
| ENSMUSG0000004120 | chr8 | Trir | 0.247799614 | 0.473670513 |
| ENSMUSG0000002207 | chr14 | Tnfrsf10b | 0.299610589 | 0.473670513 |
| ENSMUSG0000005615 | chr8 | Nanos3 | 1.000349645 | 0.473670513 |
| ENSMUSG0000003639 | chr17 | Ppp1r11 | 0.312423259 | 0.473670513 |
| ENSMUSG0000001625 | chr2 | Atp5f1e | 0.529465121 | 0.473670513 |
| ENSMUSG0000003428 | chr11 | Evpl | -0.312868653 | 0.473700863 |
| ENSMUSG0000006360 | chr8 | Ccdc102a | 0.289053786 | 0.473700863 |
| ENSMUSG0000003493 | chr11 | Arl4d | 0.426195271 | 0.473700863 |
| ENSMUSG0000002749 | chr2 | Cstf1 | 0.159174096 | 0.474185644 |
| ENSMUSG0000002799 | chr3 | Sfrp2 | -0.464693799 | 0.474258115 |
| ENSMUSG0000002395 | chr17 | Gtpbp2 | -0.263761818 | 0.474258115 |
| ENSMUSG0000007353 | chr1 | Pappa2 | 1.09413597 | 0.474258115 |
| ENSMUSG0000000512 | chr15 | Ndrg1 | -0.226528298 | 0.474720758 |
| ENSMUSG0000002412 | chr17 | Prepl | -0.203459802 | 0.474720758 |
| ENSMUSG0000002480 | chr19 | Pcgf5 | 0.22158423 | 0.474746515 |
| ENSMUSG0000007036 | chr7 | Plpp4 | -0.463858407 | 0.474759961 |
| ENSMUSG0000002719 | chr2 | Gatm | -0.323193581 | 0.474759961 |
| ENSMUSG0000003265 | chr18 | Marchf3 | 0.415338861 | 0.474759961 |
| ENSMUSG0000003243 | chr9 | Stt3b | 0.155361695 | 0.475084592 |
| ENSMUSG0000002021 | chr10 | Jsrp1 | 1.293780671 | 0.475084592 |
| ENSMUSG0000003256 | chr9 | Gnai2 | 0.175238643 | 0.475107409 |
| ENSMUSG0000004565 | chr7 | Plekha7 | -0.348961821 | 0.475458225 |
| ENSMUSG0000002150 | chr13 | Slc25a48 | -0.756003902 | 0.475458225 |
| ENSMUSG0000002478 | chr19 | Snx15 | -0.229598003 | 0.475458225 |
| ENSMUSG0000003572 | chrX | Prkx | -0.206288574 | 0.475458225 |
| ENSMUSG0000006314 | chr5 | Clip2 | -0.174673156 | 0.475458225 |
| ENSMUSG0000003572 | chr10 | Abca7 | -0.165205103 | 0.475458225 |
| ENSMUSG0000002862 | chr4 | Mrpl37 | 0.183876472 | 0.475458225 |
| ENSMUSG0000001929 | chr14 | Nop9 | 0.216326941 | 0.475458225 |
| ENSMUSG0000001981 | chr10 | Plagl1 | 0.304873792 | 0.475458225 |
| ENSMUSG0000002816 | chr3 | Bdh2 | 0.367278158 | 0.475458225 |
| ENSMUSG0000003051 | chr7 | Entrep2 | 0.364687832 | 0.475458225 |
| ENSMUSG0000007294 | chr12 | Acot1 | 0.759079683 | 0.475458225 |
| ENSMUSG0000005393 | chr3 | Cnn3 | 0.233437405 | 0.475826633 |
| ENSMUSG0000005271 | chr18 | Zfp608 | 0.17708009 | 0.476456918 |
| ENSMUSG0000002123 | chr12 | Coq6 | 0.261098681 | 0.476456918 |
| ENSMUSG0000002523 | chr9 | Adpgk | 0.217402877 | 0.476506621 |
| ENSMUSG0000002868 | chr4 | Ptch2 | -0.703764092 | 0.477263374 |

| | | | | |
|---|---|---|---|---|
| ENSMUSG0000004237 | chr11 | Fam83g | -0.819606193 | 0.477263374 |
| ENSMUSG0000003254 | chr9 | Abhd5 | 0.201374011 | 0.477263374 |
| ENSMUSG0000003736 | chr12 | Sf3b6 | 0.25996697 | 0.477263374 |
| ENSMUSG0000005502 | chr7 | Gabrg3 | -0.305352127 | 0.477279474 |
| ENSMUSG0000000773 | chr11 | Cct4 | 0.272313394 | 0.477398929 |
| ENSMUSG0000004055 | chr4 | Otud6b | 0.19492762 | 0.478300138 |
| ENSMUSG0000002803 | chr1 | Kcnq5 | -0.270837761 | 0.478551503 |
| ENSMUSG0000001495 | chr5 | Ppp1cb | 0.219153467 | 0.478744557 |
| ENSMUSG0000002884 | chr4 | Cnksr1 | -0.93189891 | 0.478746786 |
| ENSMUSG0000003672 | chr13 | Zscan12 | 0.254062731 | 0.478746786 |
| ENSMUSG0000000271 | chr2 | Cse1l | 0.174175735 | 0.478779061 |
| ENSMUSG0000001130 | chr8 | Sugp1 | 0.147684005 | 0.478790288 |
| ENSMUSG0000000234 | chr8 | Borcs8 | 0.186420706 | 0.479352861 |
| ENSMUSG0000004137 | chr6 | B4galnt3 | -1.183725781 | 0.479493849 |
| ENSMUSG0000002452 | chr18 | Pmaip1 | -0.896669922 | 0.479493849 |
| ENSMUSG0000005304 | chr7 | Brsk2 | -0.342179166 | 0.479493849 |
| ENSMUSG0000003974 | chr1 | Exo1 | -1.08743376 | 0.479493849 |
| ENSMUSG0000006021 | chr8 | Pcnx2 | -0.282422176 | 0.479493849 |
| ENSMUSG0000002330 | chr19 | Marchf5 | 0.147476284 | 0.479493849 |
| ENSMUSG0000001553 | chr13 | Mocs2 | 0.171055255 | 0.479493849 |
| ENSMUSG0000004333 | chr16 | Filip1l | 0.254917519 | 0.479493849 |
| ENSMUSG0000006826 | chr2 | Ap5s1 | 0.238785161 | 0.479493849 |
| ENSMUSG0000002643 | chr1 | Rab29 | 0.256598019 | 0.479493849 |
| ENSMUSG0000004066 | chr16 | Sh3bgr | 0.29712825 | 0.479493849 |
| ENSMUSG0000003700 | chr11 | Zfp39 | 0.292927955 | 0.479493849 |
| ENSMUSG0000002579 | chr9 | Rpl14 | 0.326119403 | 0.479493849 |
| ENSMUSG0000001879 | chr8 | Acsl1 | 0.30927013 | 0.479493849 |
| ENSMUSG0000000868 | chr7 | Rps15a | 0.376715044 | 0.479493849 |
| ENSMUSG0000001730 | chr11 | Cd300lg | 0.511598832 | 0.479493849 |
| ENSMUSG0000002986 | chr6 | Clcn1 | -0.753983528 | 0.479499411 |
| ENSMUSG0000002626 | chr1 | Ndufa10 | 0.206697749 | 0.479499411 |
| ENSMUSG0000004771 | chr16 | Ppp1r2 | 0.242265082 | 0.479499411 |
| ENSMUSG0000001362 | chr5 | Atraid | 0.231902533 | 0.479499411 |
| ENSMUSG0000000397 | chr15 | Rpl8 | 0.344515429 | 0.479499411 |
| ENSMUSG0000003064 | chr7 | Rab30 | 0.201893153 | 0.479589259 |
| ENSMUSG0000002221 | chr14 | Carmil3 | -0.368219447 | 0.480153009 |
| ENSMUSG0000000567 | chr5 | Kit | -0.224763948 | 0.480153009 |
| ENSMUSG0000004414 | chrX | Nkrf | -0.221379709 | 0.480153009 |
| ENSMUSG0000000671 | chr13 | Acot13 | 0.329780747 | 0.480153009 |
| ENSMUSG0000001923 | chr12 | Rps6kl1 | -0.310397861 | 0.480328142 |
| ENSMUSG0000003610 | chrX | Mbnl3 | -0.924732467 | 0.480335696 |
| ENSMUSG0000005731 | chr5 | Arhgap24 | 0.371867485 | 0.480335696 |
| ENSMUSG0000003006 | chr6 | Uba3 | 0.173408589 | 0.480553373 |
| ENSMUSG0000007961 | chr1 | Ankrd39 | 0.238152298 | 0.480553373 |
| ENSMUSG0000005292 | chr19 | Prkg1 | 0.22884772 | 0.480553373 |
| ENSMUSG0000003112 | chrX | 3830403N18Rik | 0.270577267 | 0.480553373 |
| ENSMUSG0000001740 | chr11 | Rpl19 | 0.331139961 | 0.480553373 |
| ENSMUSG0000002905 | chr4 | Prkcz | -0.308311965 | 0.480780887 |
| ENSMUSG0000004191 | chr2 | Dlx1 | -0.434922982 | 0.481023742 |
| ENSMUSG0000005069 | chr15 | Prkaa1 | -0.124113754 | 0.481215966 |

| | | | | |
|---|---|---|---|---|
| ENSMUSG0000007342 | chr17 | H2-Ab1 | -0.487754463 | 0.481328147 |
| ENSMUSG0000003479 | chr8 | Unc13a | -0.406076634 | 0.481328147 |
| ENSMUSG0000005074 | chr17 | Trim15 | -0.959015473 | 0.481328147 |
| ENSMUSG0000006930 | chr13 | H2bc11 | -0.928067825 | 0.481328147 |
| ENSMUSG0000000557 | chr7 | Ube2m | 0.199168412 | 0.481328147 |
| ENSMUSG0000001844 | chr11 | C1qbp | 0.199557075 | 0.481328147 |
| ENSMUSG0000002402 | chr17 | Glo1 | 0.251372514 | 0.481328147 |
| ENSMUSG0000005331 | chr4 | Sec61b | 0.472622396 | 0.481328147 |
| ENSMUSG0000005013 | chr17 | Kcnk12 | 1.416287339 | 0.481474584 |
| ENSMUSG0000002241 | chr15 | Syngr1 | -0.2986477 | 0.481477125 |
| ENSMUSG0000004025 | chr3 | Gbp7 | -0.342285976 | 0.481672608 |
| ENSMUSG0000002417 | chr17 | Sox8 | -0.282690176 | 0.481672608 |
| ENSMUSG0000002805 | chr3 | Rit1 | 0.206883963 | 0.481672608 |
| ENSMUSG0000002623 | chr1 | Nmur1 | -0.870631009 | 0.481716367 |
| ENSMUSG0000004649 | chr11 | C1qtnf2 | 0.279879794 | 0.481716367 |
| ENSMUSG0000003124 | chrX | Chmp1b2 | -0.159991755 | 0.481737903 |
| ENSMUSG0000004976 | chr17 | Micos13 | 0.326062905 | 0.481737903 |
| ENSMUSG0000002002 | chr10 | Ndufa12 | 0.345153989 | 0.481737903 |
| ENSMUSG0000002041 | chr11 | Ascc2 | -0.163855083 | 0.48217473 |
| ENSMUSG0000009367 | chr10 | Rpl41 | 0.41062539 | 0.48255217 |
| ENSMUSG0000002716 | chr2 | Commd9 | 0.259241549 | 0.482603234 |
| ENSMUSG0000002517 | chr11 | Wdr45b | 0.168537703 | 0.482624367 |
| ENSMUSG0000005222 | chr12 | Ppp1r36 | 0.503361225 | 0.48263712 |
| ENSMUSG0000002167 | chr13 | Hmgcr | -0.426229168 | 0.482791117 |
| ENSMUSG0000002620 | chr1 | Ptprn | -0.30921369 | 0.482791117 |
| ENSMUSG0000002506 | chr19 | Sfr1 | 0.232549079 | 0.482791117 |
| ENSMUSG0000006345 | chr6 | D630045J12Rik | -0.234682167 | 0.482798704 |
| ENSMUSG0000007552 | chr2 | Malrd1 | -1.525362612 | 0.48313254 |
| ENSMUSG0000001649 | chr19 | Cd274 | -0.381397354 | 0.48313254 |
| ENSMUSG0000000194 | chr9 | Spa17 | -0.584248617 | 0.48313254 |
| ENSMUSG0000007943 | chrX | Rpl36a | 0.290598317 | 0.48313254 |
| ENSMUSG0000004470 | chr7 | Gemin7 | 0.369240288 | 0.48313254 |
| ENSMUSG0000003018 | chr6 | Ybx3 | 0.284198537 | 0.483343432 |
| ENSMUSG0000003193 | chr8 | Wwp2 | -0.240820133 | 0.483414897 |
| ENSMUSG0000003257 | chr9 | Manf | 0.457913136 | 0.483821524 |
| ENSMUSG0000007069 | chr1 | Cntnap5a | -0.288252494 | 0.484207356 |
| ENSMUSG0000003075 | chr7 | Thap12 | 0.200505638 | 0.484207356 |
| ENSMUSG0000004135 | chr3 | Ssr2 | 0.242982221 | 0.484207356 |
| ENSMUSG0000004082 | chr14 | 1700123O20Rik | 0.336012842 | 0.484207356 |
| ENSMUSG0000002492 | chr19 | Rnaseh2c | 0.357567259 | 0.484207356 |
| ENSMUSG0000009022 | chr16 | Pcp4 | 0.345617207 | 0.484207356 |
| ENSMUSG0000006037 | chr5 | Caln1 | 0.539217842 | 0.484207356 |
| ENSMUSG0000011866 | chr19 | Rps6ka4 | 0.174525634 | 0.484756975 |
| ENSMUSG0000000868 | chrX | Rpl10 | 0.32521332 | 0.484756975 |
| ENSMUSG0000002236 | chr15 | Atad2 | -0.463612785 | 0.485016907 |
| ENSMUSG0000006778 | chr1 | Pi15 | -0.489186338 | 0.485016907 |
| ENSMUSG0000003008 | chr6 | Aldh1l1 | -0.519920568 | 0.485016907 |
| ENSMUSG0000002042 | chr11 | Tns3 | -0.201050922 | 0.485104427 |
| ENSMUSG0000000535 | chr15 | Txn2 | 0.221454651 | 0.485104427 |
| ENSMUSG0000002939 | chr5 | Rilpl1 | 0.226196732 | 0.485104427 |

| | | | | |
|---|---|---|---|---|
| ENSMUSG0000001229 | chr17 | Tjap1 | 0.190119319 | 0.485364175 |
| ENSMUSG0000003884 | chr2 | Kif16b | -0.243978256 | 0.486017876 |
| ENSMUSG0000002561 | chr16 | Cct8 | 0.234331672 | 0.486017876 |
| ENSMUSG0000003121 | chrX | Gpr165 | -0.353799704 | 0.486073512 |
| ENSMUSG0000002224 | chr15 | Rad1 | 0.361315629 | 0.486413052 |
| ENSMUSG0000004753 | chr12 | Mis18bp1 | -0.685264178 | 0.48665265 |
| ENSMUSG0000002035 | chr11 | Hnrnpab | 0.201125326 | 0.48665265 |
| ENSMUSG0000002427 | chr18 | Slc39a6 | -0.229265763 | 0.486929321 |
| ENSMUSG0000004368 | chr17 | Fem1a | 0.134806124 | 0.486929321 |
| ENSMUSG0000003975 | chr3 | Dnttip2 | 0.233530024 | 0.486929321 |
| ENSMUSG0000003164 | chr8 | Cbr4 | 0.254292631 | 0.486929321 |
| ENSMUSG0000003203 | chr9 | Kirrel3 | 0.428288993 | 0.486929321 |
| ENSMUSG0000001440 | chr7 | Tsg101 | -0.163292402 | 0.487473182 |
| ENSMUSG0000006242 | chr11 | Arf2 | 0.210427386 | 0.487548783 |
| ENSMUSG0000007901 | chr12 | Ifi27l2a | -0.774768827 | 0.48755987 |
| ENSMUSG0000003239 | chr9 | Snapc5 | 0.227272205 | 0.487603973 |
| ENSMUSG0000004715 | chr14 | Khnyn | -0.135517405 | 0.487613761 |
| ENSMUSG0000002912 | chr5 | Ppp2r2c | -0.293027657 | 0.487789337 |
| ENSMUSG0000003135 | chrX | Rbbp7 | 0.216437268 | 0.487789337 |
| ENSMUSG0000005411 | chr15 | Skp2 | 0.275805115 | 0.487789337 |
| ENSMUSG0000003794 | chr13 | Fgd3 | 0.470129781 | 0.487789337 |
| ENSMUSG0000002114 | chr13 | Wdr37 | -0.184211512 | 0.487872063 |
| ENSMUSG0000002262 | chr15 | Arsa | -0.148723021 | 0.487872063 |
| ENSMUSG0000002255 | chr15 | Hsf1 | 0.133199667 | 0.487872063 |
| ENSMUSG0000002496 | chr19 | Vegfb | 0.217346773 | 0.487872063 |
| ENSMUSG0000003918 | chr13 | AW209491 | 0.241034841 | 0.487872063 |
| ENSMUSG0000000143 | chr10 | Col18a1 | 0.309414541 | 0.487872063 |
| ENSMUSG0000000334 | chr10 | Atp8b3 | -0.848494609 | 0.488126106 |
| ENSMUSG0000002959 | chr5 | Sdsl | -0.513656127 | 0.489350272 |
| ENSMUSG0000002840 | chr4 | Zdhhc21 | -0.230439951 | 0.489806136 |
| ENSMUSG0000003086 | chr7 | Acadsb | 0.175589506 | 0.489806136 |
| ENSMUSG0000000329 | chr9 | Mrpl4 | 0.21316945 | 0.489806136 |
| ENSMUSG0000002180 | chr14 | Rtraf | 0.229508109 | 0.489806136 |
| ENSMUSG0000005924 | chr11 | Septin9 | 0.248541633 | 0.489806136 |
| ENSMUSG0000003235 | chr9 | Tmed3 | 0.27432494 | 0.489806136 |
| ENSMUSG0000001240 | chr14 | Rpl15 | 0.330163059 | 0.489806136 |
| ENSMUSG0000002613 | chr1 | Prim2 | 0.28514898 | 0.489806136 |
| ENSMUSG0000002090 | chr11 | Pik3r5 | 0.607404923 | 0.489806136 |
| ENSMUSG0000002198 | chr14 | Amer2 | -0.418145672 | 0.489933695 |
| ENSMUSG0000000276 | chr17 | Gnmt | -1.043027855 | 0.490133559 |
| ENSMUSG0000002485 | chr19 | Grk2 | -0.14081847 | 0.490207852 |
| ENSMUSG0000003472 | chr2 | Tmx4 | -0.163268242 | 0.490309102 |
| ENSMUSG0000000180 | chr7 | Lrp3 | -0.208740024 | 0.490429485 |
| ENSMUSG0000002728 | chr2 | Zfp106 | 0.172717588 | 0.490617569 |
| ENSMUSG0000005346 | chr14 | Hs6st3 | 0.423916096 | 0.491131464 |
| ENSMUSG0000002105 | chr12 | Akap5 | 0.327470918 | 0.491131464 |
| ENSMUSG0000003342 | chr10 | Upb1 | -0.803012721 | 0.491639373 |
| ENSMUSG0000002760 | chr2 | Ggt7 | -0.214118309 | 0.491639373 |
| ENSMUSG0000002901 | chr5 | Pmpcb | 0.20832236 | 0.491639373 |
| ENSMUSG0000002173 | chr14 | Atxn7 | 0.268341 | 0.491639373 |

| | | | | |
|---|---|---|---|---|
| ENSMUSG0000003400 | chr18 | Slc66a2 | 0.192240219 | 0.491752643 |
| ENSMUSG0000000757 | chr17 | Fance | 0.211443106 | 0.491752643 |
| ENSMUSG0000001195 | chr9 | Bnip2 | 0.235173937 | 0.491752643 |
| ENSMUSG0000001630 | chrX | Ube2a | 0.280144709 | 0.491815519 |
| ENSMUSG0000005072 | chr9 | Plekho2 | 0.176634817 | 0.491861413 |
| ENSMUSG0000002949 | chr5 | Pxmp2 | 0.460378181 | 0.491861413 |
| ENSMUSG0000002947 | chr5 | Camkk2 | -0.245648249 | 0.491958376 |
| ENSMUSG0000001884 | chr11 | Rad51d | 0.216691777 | 0.491958376 |
| ENSMUSG0000005784 | chr6 | Rpl32 | 0.311931496 | 0.491958376 |
| ENSMUSG0000005960 | chr10 | Syn3 | -0.36718915 | 0.491965663 |
| ENSMUSG0000004346 | chr15 | Elfn2 | -0.319455351 | 0.491965663 |
| ENSMUSG0000001075 | chr7 | Cars1 | 0.171764252 | 0.491965663 |
| ENSMUSG0000002892 | chr5 | Cdk14 | 0.188462072 | 0.491965663 |
| ENSMUSG0000002478 | chr19 | Gpha2 | 1.301186046 | 0.491988308 |
| ENSMUSG0000002821 | chr4 | Cibar1 | 0.199066165 | 0.49206463 |
| ENSMUSG0000005271 | chr17 | BC004004 | 0.192627379 | 0.492200589 |
| ENSMUSG0000002095 | chr12 | Ap4s1 | 0.315326638 | 0.492200589 |
| ENSMUSG0000004095 | chr7 | Rps19 | 0.386766115 | 0.492200589 |
| ENSMUSG0000004255 | chr1 | Zp3r | 0.685534632 | 0.492200589 |
| ENSMUSG0000001370 | chr14 | Timm23 | 0.245178943 | 0.492672469 |
| ENSMUSG0000002275 | chr16 | Tfg | 0.196731622 | 0.493386038 |
| ENSMUSG0000002698 | chr2 | Il1rn | -1.616925663 | 0.493473423 |
| ENSMUSG0000004671 | chr8 | Bst2 | -0.597606001 | 0.493473423 |
| ENSMUSG0000006033 | chr2 | Zfp937 | -0.373692443 | 0.493473423 |
| ENSMUSG0000004971 | chr8 | Lig4 | -0.22984286 | 0.493473423 |
| ENSMUSG0000004953 | chr14 | Sall2 | -0.176303922 | 0.493473423 |
| ENSMUSG0000004266 | chr18 | Wdr55 | 0.252936644 | 0.493473423 |
| ENSMUSG0000005388 | chr8 | Sh2d4a | -0.647976787 | 0.493504928 |
| ENSMUSG0000002289 | chr16 | Slc15a2 | -0.5170801 | 0.493504928 |
| ENSMUSG0000002484 | chr19 | Cst6 | -0.644132354 | 0.493504928 |
| ENSMUSG0000002082 | chr11 | Slc46a1 | -0.233724622 | 0.493504928 |
| ENSMUSG0000004390 | chr2 | Trp53bp1 | -0.181314937 | 0.493504928 |
| ENSMUSG0000001804 | chr15 | Rrp7a | 0.189277257 | 0.493504928 |
| ENSMUSG0000002696 | chr2 | Ssna1 | 0.333895161 | 0.493504928 |
| ENSMUSG0000002805 | chr3 | Hcn3 | -0.356175376 | 0.493532571 |
| ENSMUSG0000003388 | chr11 | Lgals3bp | -0.32695423 | 0.493532571 |
| ENSMUSG0000004400 | chr10 | Gls2 | -0.451018699 | 0.493532571 |
| ENSMUSG0000002181 | chr14 | Anxa7 | -0.170665866 | 0.493532571 |
| ENSMUSG0000002286 | chr16 | Usp25 | 0.137628616 | 0.493532571 |
| ENSMUSG0000000172 | chr12 | Akt1 | 0.19489199 | 0.493532571 |
| ENSMUSG0000002710 | chr2 | Hoxd8 | 0.189168964 | 0.493532571 |
| ENSMUSG0000003204 | chr9 | Rpusd4 | 0.224553853 | 0.493532571 |
| ENSMUSG0000002155 | chr13 | Gkap1 | 0.225228018 | 0.493532571 |
| ENSMUSG0000003094 | chr7 | Lhpp | 0.306052008 | 0.493808804 |
| ENSMUSG0000001186 | chr2 | Afg2b | 0.291273639 | 0.493978673 |
| ENSMUSG0000003499 | chr10 | Eef2 | 0.189387244 | 0.493980356 |
| ENSMUSG0000002090 | chr11 | Stx8 | 0.246156435 | 0.494518664 |
| ENSMUSG0000004546 | chr6 | Zfp956 | -0.272598314 | 0.495636899 |
| ENSMUSG0000002708 | chr2 | Itgav | 0.174405926 | 0.495967129 |
| ENSMUSG0000002886 | chr4 | Map3k6 | 0.254532758 | 0.496035392 |

| | | | | |
|---|---|---|---|---|
| ENSMUSG0000003652 | chr5 | Card11 | -1.019658614 | 0.496498497 |
| ENSMUSG0000003647 | chr17 | Tbc1d24 | -0.235771486 | 0.497705501 |
| ENSMUSG0000004194 | chr1 | Mfsd9 | -0.538455305 | 0.497705501 |
| ENSMUSG0000002040 | chr11 | Vdac1 | 0.162935934 | 0.497705501 |
| ENSMUSG0000003864 | chr6 | Creb3l2 | 0.181083886 | 0.497705501 |
| ENSMUSG0000005326 | chr5 | Dspp | 0.631454468 | 0.497705501 |
| ENSMUSG0000002965 | chr5 | Medag | 0.292864736 | 0.497705501 |
| ENSMUSG0000004131 | chr17 | Thoc6 | 0.335125954 | 0.497705501 |
| ENSMUSG0000003359 | chr14 | Lgi3 | 0.595057289 | 0.497705501 |
| ENSMUSG0000007876 | chr11 | Slfn1 | -1.573610887 | 0.49782937 |
| ENSMUSG0000002117 | chr12 | Cdca7l | -0.629433289 | 0.49782937 |
| ENSMUSG0000007956 | chr5 | Maea | 0.218328783 | 0.49782937 |
| ENSMUSG0000002604 | chr1 | Poglut2 | 0.245901781 | 0.49782937 |
| ENSMUSG0000004145 | chr5 | Rpl21 | 0.296368147 | 0.49782937 |
| ENSMUSG0000000522 | chr5 | Corin | 0.64519942 | 0.497922076 |
| ENSMUSG0000003301 | chr18 | Nfatc1 | 0.3096331 | 0.498111452 |
| ENSMUSG0000005666 | chr15 | Them6 | -0.294897199 | 0.498188988 |
| ENSMUSG0000003765 | chr8 | Slc20a2 | 0.219789104 | 0.498188988 |
| ENSMUSG0000006225 | chr9 | Opcml | 0.465525443 | 0.498188988 |
| ENSMUSG0000001832 | chr2 | Tomm34 | -0.125648908 | 0.498293218 |
| ENSMUSG0000003547 | chr10 | Mbd3 | 0.257238045 | 0.498293218 |
| ENSMUSG0000002153 | chr13 | Cetn3 | 0.146167539 | 0.498518724 |
| ENSMUSG0000003879 | chr7 | Zscan2 | 0.253743273 | 0.4990088 |
| ENSMUSG0000003165 | chr8 | Cbln1 | 0.40357135 | 0.4990088 |
| ENSMUSG0000003201 | chr9 | Trim29 | 0.915132219 | 0.4990088 |
| ENSMUSG0000002120 | chr12 | Ppp4r4 | -0.552338945 | 0.499917958 |
| ENSMUSG0000002476 | chr19 | Cdc42bpg | -0.358109913 | 0.499917958 |
| ENSMUSG0000001655 | chr15 | Foxred2 | -0.385772562 | 0.499917958 |
| ENSMUSG0000004000 | chr10 | Gnaz | -0.248839154 | 0.499917958 |
| ENSMUSG0000002752 | chr2 | Edn3 | -0.299577098 | 0.499917958 |
| ENSMUSG0000009176 | chr8 | Zfp964 | -0.256920894 | 0.499917958 |
| ENSMUSG0000003899 | chr2 | Cables2 | -0.157001292 | 0.499917958 |
| ENSMUSG0000002655 | chr1 | Copa | 0.128896404 | 0.499917958 |
| ENSMUSG0000002782 | chr3 | Ssr3 | 0.156023638 | 0.499917958 |
| ENSMUSG0000004857 | chr19 | Tmem252 | 0.191404852 | 0.499917958 |
| ENSMUSG0000002305 | chr15 | Tarbp2 | 0.17967782 | 0.499917958 |
| ENSMUSG0000002139 | chr13 | Nol8 | 0.189747753 | 0.499917958 |
| ENSMUSG0000003818 | chr7 | Btbd10 | 0.196451661 | 0.499917958 |
| ENSMUSG0000003821 | chr8 | Cep44 | 0.214309785 | 0.499917958 |
| ENSMUSG0000003179 | chr8 | Tpm4 | 0.238727686 | 0.499917958 |
| ENSMUSG0000001694 | chr15 | Tmprss6 | 0.607886667 | 0.499917958 |
| ENSMUSG0000000428 | chr6 | Atp6v1f | 0.298035658 | 0.499917958 |
| ENSMUSG0000000128 | chr15 | Pfdn5 | 0.328416264 | 0.499917958 |
| ENSMUSG0000006359 | chr7 | Gng8 | 0.352882484 | 0.499917958 |
| ENSMUSG0000003291 | chr9 | Cspg4 | 0.353195024 | 0.499917958 |
| ENSMUSG0000004679 | chr8 | Ppp1r3b | 0.401332485 | 0.499917958 |
| ENSMUSG0000002652 | chr1 | Pycr2 | 0.367150532 | 0.500083963 |
| ENSMUSG0000002142 | chr13 | Ly86 | -0.50451352 | 0.500178914 |
| ENSMUSG0000002051 | chr11 | Tubd1 | 0.284663822 | 0.500344279 |
| ENSMUSG0000002842 | chr4 | Rad23b | 0.144075835 | 0.500799942 |

| | | | | |
|---|---|---|---|---|
| ENSMUSG0000004738 | chr8 | Atmin | -0.176345303 | 0.501273757 |
| ENSMUSG0000002945 | chr5 | Brap | 0.137256207 | 0.501362922 |
| ENSMUSG0000003205 | chr9 | Alg9 | -0.2039964 | 0.501684106 |
| ENSMUSG0000003027 | chr6 | C2cd5 | -0.184352043 | 0.501684106 |
| ENSMUSG0000002977 | chr6 | Hibadh | 0.188828554 | 0.501684106 |
| ENSMUSG0000003061 | chr7 | Det1 | 0.242279766 | 0.501912675 |
| ENSMUSG0000003518 | chr14 | Fam124a | 0.227823709 | 0.502031077 |
| ENSMUSG0000002482 | chr19 | Mrpl21 | 0.30122224 | 0.502031077 |
| ENSMUSG0000003644 | chr10 | Lum | 0.232958666 | 0.502098069 |
| ENSMUSG0000002242 | chr15 | Tomm22 | 0.287827742 | 0.502254429 |
| ENSMUSG0000000702 | chr17 | Syngr3 | -0.287080708 | 0.502811346 |
| ENSMUSG0000003324 | chr8 | Hsf4 | -0.388590678 | 0.502811346 |
| ENSMUSG0000002028 | chr11 | Pus10 | -0.242306896 | 0.502811346 |
| ENSMUSG0000002128 | chr12 | Tnfaip2 | 0.365258065 | 0.502811346 |
| ENSMUSG0000002937 | chr5 | Ereg | 0.679336152 | 0.502811346 |
| ENSMUSG0000000408 | chr2 | Map3k20 | 0.272442675 | 0.503043541 |
| ENSMUSG0000001947 | chr9 | Cdc37 | 0.221896358 | 0.50326395 |
| ENSMUSG0000005583 | chr17 | Elob | 0.311356175 | 0.503674607 |
| ENSMUSG0000004844 | chr17 | Cyp4f16 | -0.335265664 | 0.503720211 |
| ENSMUSG0000003475 | chr13 | Mast4 | 0.267325797 | 0.503720211 |
| ENSMUSG0000004453 | chr17 | Rps2 | 0.258297888 | 0.503728504 |
| ENSMUSG0000005860 | chr15 | Rpl30 | 0.247624317 | 0.503728504 |
| ENSMUSG0000003964 | chr11 | Mrpl12 | 0.32682906 | 0.503728504 |
| ENSMUSG0000001867 | chr11 | Copz2 | 0.276949523 | 0.503859605 |
| ENSMUSG0000002012 | chr10 | Avpr1a | -0.388946873 | 0.504621415 |
| ENSMUSG0000002452 | chr18 | Gnal | -0.363791012 | 0.504621415 |
| ENSMUSG0000009903 | chr1 | Tcf24 | -0.761198774 | 0.504621415 |
| ENSMUSG0000005267 | chrX | Zmat1 | -0.293444525 | 0.504621415 |
| ENSMUSG0000006630 | chr7 | Numa1 | -0.212521231 | 0.504621415 |
| ENSMUSG0000002494 | chr19 | Scyl1 | 0.131404681 | 0.504621415 |
| ENSMUSG0000002871 | chr4 | Atpaf1 | 0.157501183 | 0.504621415 |
| ENSMUSG0000003283 | chr9 | Trpc1 | 0.169521457 | 0.504621415 |
| ENSMUSG0000004827 | chr11 | Syngr2 | 0.183143249 | 0.504621415 |
| ENSMUSG0000003061 | chr7 | Eed | 0.185734703 | 0.504621415 |
| ENSMUSG0000005275 | chr17 | Traf7 | 0.18073483 | 0.504621415 |
| ENSMUSG0000003251 | chr9 | Rpsa | 0.237673552 | 0.504621415 |
| ENSMUSG0000002039 | chr11 | Kremen1 | 0.221765062 | 0.504621415 |
| ENSMUSG0000002025 | chr9 | Ppm1m | 0.223921258 | 0.504621415 |
| ENSMUSG0000002388 | chr17 | Smoc2 | 0.262107967 | 0.504621415 |
| ENSMUSG0000000276 | chr17 | Mrpl2 | 0.267337991 | 0.504621415 |
| ENSMUSG0000004636 | chr7 | Rpl27a | 0.338093819 | 0.504621415 |
| ENSMUSG0000005661 | chr19 | Ppp1r14b | 0.329093845 | 0.504621415 |
| ENSMUSG0000003718 | chr4 | Grhl3 | 0.974723212 | 0.504621415 |
| ENSMUSG0000006364 | chr5 | Jakmip1 | -0.485214418 | 0.504642045 |
| ENSMUSG0000006655 | chr5 | Hmgb1 | 0.166359031 | 0.504642045 |
| ENSMUSG0000002499 | chr19 | Rbp4 | 0.490391943 | 0.50526119 |
| ENSMUSG0000006055 | chr17 | H2-Q7 | -0.972363418 | 0.505687958 |
| ENSMUSG0000006808 | chr15 | Cyp2d9 | -2.271351412 | 0.505687958 |
| ENSMUSG0000002866 | chr4 | Ephb2 | -0.25413719 | 0.505687958 |
| ENSMUSG0000005879 | chr10 | Nap1l1 | 0.195157292 | 0.505687958 |

| | | | | |
|---|---|---|---|---|
| ENSMUSG0000003971 | chr3 | Ralyl | -0.265229119 | 0.505921142 |
| ENSMUSG0000000568 | chr7 | Ampd3 | -0.220424947 | 0.506500476 |
| ENSMUSG0000002207 | chr14 | Bora | -0.307309226 | 0.50664055 |
| ENSMUSG0000000772 | chr8 | Ccdc124 | 0.227710381 | 0.506741098 |
| ENSMUSG0000007101 | chr4 | Ndufb6 | 0.371231928 | 0.506741098 |
| ENSMUSG0000000320 | chr17 | Ebi3 | -1.195803601 | 0.506920499 |
| ENSMUSG0000003430 | chr18 | Fam53c | 0.113160259 | 0.506920499 |
| ENSMUSG0000003419 | chr14 | R3hcc1 | 0.20064863 | 0.506920499 |
| ENSMUSG0000000108 | chr4 | Luzp1 | 0.230481505 | 0.506920499 |
| ENSMUSG0000002645 | chr1 | Adipor1 | 0.148660375 | 0.507065174 |
| ENSMUSG0000004267 | chr7 | Ypel3 | -0.218922651 | 0.507118916 |
| ENSMUSG0000004803 | chr2 | Ccdc187 | -0.331772441 | 0.507631206 |
| ENSMUSG0000004741 | chr12 | Flrt2 | -0.272308854 | 0.507631206 |
| ENSMUSG0000002403 | chr17 | Ndufv3 | 0.25048826 | 0.507631206 |
| ENSMUSG0000003302 | chr18 | Cdo1 | 0.295928922 | 0.507631206 |
| ENSMUSG0000006661 | chr5 | Zfp932 | -0.467033627 | 0.507767283 |
| ENSMUSG0000005329 | chr6 | AI854703 | -0.317080833 | 0.508135313 |
| ENSMUSG0000003027 | chr6 | Ttll3 | -0.265350389 | 0.50816284 |
| ENSMUSG0000003142 | chrX | Zcchc18 | -0.283946202 | 0.508268178 |
| ENSMUSG0000003117 | chrX | Ftsj1 | 0.25055877 | 0.508268178 |
| ENSMUSG0000003225 | chr9 | Drd2 | -1.810452224 | 0.508412224 |
| ENSMUSG0000002285 | chr16 | Ehhadh | -0.253453296 | 0.508412224 |
| ENSMUSG0000002917 | chr5 | Tbc1d1 | 0.154412025 | 0.508704632 |
| ENSMUSG0000010939 | chr7 | Zfp582 | -0.512908792 | 0.508849335 |
| ENSMUSG0000002113 | chr12 | Smoc1 | 0.263537269 | 0.508849335 |
| ENSMUSG0000005243 | chr3 | Bmpr1b | 0.932156698 | 0.508859787 |
| ENSMUSG0000004280 | chr4 | Gpr153 | -0.192594403 | 0.508934727 |
| ENSMUSG0000006320 | chr13 | Nol7 | 0.21263289 | 0.509005627 |
| ENSMUSG0000000777 | chr11 | Trappc2b | 0.376942025 | 0.509181832 |
| ENSMUSG0000005682 | chr8 | Tsnax | 0.178528703 | 0.50936131 |
| ENSMUSG0000004006 | chr2 | Plcb2 | -0.795493082 | 0.509912819 |
| ENSMUSG0000004180 | chr1 | Efhc1 | -0.441501196 | 0.509912819 |
| ENSMUSG0000003074 | chr7 | Rps3 | 0.342643842 | 0.510444225 |
| ENSMUSG0000002902 | chr4 | Trp73 | 1.00832931 | 0.510444225 |
| ENSMUSG0000002977 | chr6 | Klhdc10 | -0.224568226 | 0.510455056 |
| ENSMUSG0000007320 | chrX | Ccdc160 | -0.445051266 | 0.510455056 |
| ENSMUSG0000005251 | chr7 | Nav2 | 0.23312434 | 0.510455056 |
| ENSMUSG0000002339 | chr2 | Dlx2 | -0.322282023 | 0.510486926 |
| ENSMUSG0000005352 | chr7 | Lgals7 | 0.998001129 | 0.510486926 |
| ENSMUSG0000005718 | chr2 | Scn3a | -0.280489096 | 0.510489932 |
| ENSMUSG0000004948 | chr8 | Ctu2 | 0.251672531 | 0.510593024 |
| ENSMUSG0000004240 | chr4 | Zmym6 | -0.184081729 | 0.511106408 |
| ENSMUSG0000001844 | chr11 | Rpain | 0.231837441 | 0.511200462 |
| ENSMUSG0000002931 | chr5 | Hsd17b11 | -0.238859731 | 0.512231829 |
| ENSMUSG0000006972 | chr7 | Zfp975 | 0.207582379 | 0.512356593 |
| ENSMUSG0000002718 | chr2 | Cat | 0.218224167 | 0.512356593 |
| ENSMUSG0000004684 | chr10 | Ckap4 | 0.247742557 | 0.512356593 |
| ENSMUSG0000007060 | chr4 | Zfp992 | 0.346662381 | 0.512356593 |
| ENSMUSG0000004330 | chr3 | B3galnt1 | 0.221226053 | 0.512708679 |
| ENSMUSG0000002766 | chr3 | Skil | -0.203267833 | 0.512904759 |

| | | | | |
|---|---|---|---|---|
| ENSMUSG0000002928 | chr5 | Tgfbr3 | 0.241418603 | 0.512904759 |
| ENSMUSG0000001959 | chr11 | Cyb561 | -0.35263503 | 0.512977668 |
| ENSMUSG0000005807 | chr12 | Eml1 | 0.229697517 | 0.512977668 |
| ENSMUSG0000006967 | chr10 | Nkain2 | -0.247353386 | 0.513126941 |
| ENSMUSG0000004758 | chr3 | Tyw3 | -0.251062261 | 0.513126941 |
| ENSMUSG0000006261 | chr17 | 2310039H08Rik | 0.311042796 | 0.513126941 |
| ENSMUSG0000002107 | chr12 | Timm9 | 0.209136193 | 0.51378945 |
| ENSMUSG0000002166 | chr13 | Btf3 | 0.24673825 | 0.51378945 |
| ENSMUSG0000004236 | chr4 | Gjb3 | -0.939605389 | 0.513839693 |
| ENSMUSG0000003110 | chrX | Elf4 | -0.387021758 | 0.513839693 |
| ENSMUSG0000004521 | chr18 | Asxl3 | -0.405079919 | 0.513839693 |
| ENSMUSG0000003367 | chr7 | Gabrb3 | -0.220083698 | 0.513839693 |
| ENSMUSG0000003210 | chr9 | Nlrx1 | -0.243949218 | 0.513839693 |
| ENSMUSG0000002519 | chr19 | Chuk | -0.124245905 | 0.513839693 |
| ENSMUSG0000003234 | chr9 | Mto1 | 0.174171622 | 0.513839693 |
| ENSMUSG0000003296 | chr2 | Fkbp1a | 0.231253883 | 0.513839693 |
| ENSMUSG0000003506 | chr7 | Eef2k | 0.214588451 | 0.513839693 |
| ENSMUSG0000002947 | chr5 | Rnf34 | 0.242936386 | 0.513839693 |
| ENSMUSG0000006108 | chr16 | Lsamp | 0.275630563 | 0.513839693 |
| ENSMUSG0000005622 | chr13 | Spock1 | 0.345207708 | 0.513839693 |
| ENSMUSG0000003784 | chr10 | Rtkn2 | 1.116612372 | 0.513839693 |
| ENSMUSG0000007094 | chr1 | Il1rl2 | 0.526791312 | 0.513839693 |
| ENSMUSG0000009261 | chr17 | Btnl6 | -2.629932617 | 0.514070602 |
| ENSMUSG0000003550 | chr5 | Cox18 | 0.27673385 | 0.514070602 |
| ENSMUSG0000005734 | chr15 | Apol9a | -0.649583248 | 0.514153209 |
| ENSMUSG0000002267 | chr16 | Cep20 | 0.271671134 | 0.514153209 |
| ENSMUSG0000006827 | chrX | Shroom4 | -0.271786425 | 0.514835459 |
| ENSMUSG0000006882 | chr3 | Csde1 | 0.231559919 | 0.514835459 |
| ENSMUSG0000002011 | chr11 | Pno1 | 0.319438326 | 0.514835459 |
| ENSMUSG0000000843 | chr7 | Rdh13 | 0.181414855 | 0.515110563 |
| ENSMUSG0000001990 | chr10 | Ppp1r12a | 0.211938347 | 0.515141729 |
| ENSMUSG0000001125 | chr4 | Pabpc4 | 0.127467606 | 0.515221099 |
| ENSMUSG0000002186 | chr14 | Ppif | -0.314740942 | 0.515397032 |
| ENSMUSG0000004594 | chr7 | Mrps12 | 0.367135812 | 0.516072492 |
| ENSMUSG0000002269 | chr16 | Naa50 | 0.158150144 | 0.51623798 |
| ENSMUSG0000002465 | chr19 | Asrgl1 | -0.175208297 | 0.516725492 |
| ENSMUSG0000005973 | chr19 | Ndufs8 | 0.255366722 | 0.516725492 |
| ENSMUSG0000002585 | chr5 | Fam20c | -0.206944911 | 0.516737099 |
| ENSMUSG0000002499 | chr19 | Prdx3 | 0.203151437 | 0.516843177 |
| ENSMUSG0000003519 | chr6 | Arl6ip5 | 0.219244583 | 0.516843177 |
| ENSMUSG0000000609 | chr7 | Tbcb | 0.28400503 | 0.516843177 |
| ENSMUSG0000007152 | chr19 | Atp5mk | 0.378243122 | 0.516843177 |
| ENSMUSG0000004049 | chr7 | Igsf23 | -1.56057392 | 0.516873984 |
| ENSMUSG0000002448 | chr18 | Hbegf | -0.532696863 | 0.516873984 |
| ENSMUSG0000004027 | chr17 | Pacsin1 | -0.302653853 | 0.516873984 |
| ENSMUSG0000008970 | chr8 | Galnt2 | 0.159411999 | 0.516873984 |
| ENSMUSG0000002829 | chr4 | Smim8 | 0.225531979 | 0.516873984 |
| ENSMUSG0000005902 | chr6 | Kcp | 0.484398671 | 0.516873984 |
| ENSMUSG0000004075 | chr14 | Cmtm5 | -0.234156734 | 0.516922632 |
| ENSMUSG0000004861 | chr8 | Rtbdn | -0.57003611 | 0.517226674 |

| | | | | |
|---|---|---|---|---|
| ENSMUSG0000007559 | chr14 | Nynrin | -0.176467944 | 0.517357914 |
| ENSMUSG0000000461 | chr7 | Etfb | 0.494812257 | 0.517357914 |
| ENSMUSG0000000022 | chrX | Drp2 | -0.268776303 | 0.517436949 |
| ENSMUSG0000003102 | chr7 | Tub | -0.305074724 | 0.517757429 |
| ENSMUSG0000000201 | chr17 | Fam98a | 0.239471188 | 0.517757429 |
| ENSMUSG0000003051 | chr7 | Pcsk6 | 0.23413721 | 0.517757429 |
| ENSMUSG0000000869 | chr15 | Ncaph2 | 0.176560289 | 0.517994607 |
| ENSMUSG0000009614 | chr7 | Kcnj11 | -0.27750327 | 0.518163165 |
| ENSMUSG0000005320 | chr14 | Styx | -0.197922518 | 0.518163165 |
| ENSMUSG0000004112 | chr9 | Msantd4 | 0.175071963 | 0.518163443 |
| ENSMUSG0000003853 | chr7 | Atf5 | 0.188807664 | 0.518345921 |
| ENSMUSG0000000525 | chr16 | Ripk4 | -0.789462343 | 0.518377092 |
| ENSMUSG0000003688 | chr7 | Arhgap33 | -0.350647829 | 0.518377092 |
| ENSMUSG0000004401 | chr8 | Npy5r | -0.644293659 | 0.518377092 |
| ENSMUSG0000000671 | chr13 | Gmnn | -0.5225697 | 0.518377092 |
| ENSMUSG0000003910 | chr5 | Htr5a | -0.700022693 | 0.518377092 |
| ENSMUSG0000003518 | chr2 | Slc24a5 | -0.454822154 | 0.518377092 |
| ENSMUSG0000002426 | chr18 | Tpgs2 | -0.170510616 | 0.518377092 |
| ENSMUSG0000002198 | chr14 | Cab39l | 0.167856342 | 0.518377092 |
| ENSMUSG0000002967 | chr6 | Ing3 | 0.217513325 | 0.518377092 |
| ENSMUSG0000001498 | chr1 | Tsen15 | 0.28158783 | 0.518377092 |
| ENSMUSG0000006232 | chr18 | Rpl17 | 0.359303081 | 0.518377092 |
| ENSMUSG0000004001 | chr8 | Slc7a5 | 0.314208579 | 0.518377092 |
| ENSMUSG0000004576 | chr13 | B230219D22Rik | 0.133045936 | 0.518728826 |
| ENSMUSG0000002845 | chr4 | Cd72 | -0.520214909 | 0.518862285 |
| ENSMUSG0000001495 | chr2 | Gorasp2 | 0.151801857 | 0.519136854 |
| ENSMUSG0000002197 | chr14 | Ints9 | 0.216976581 | 0.519136854 |
| ENSMUSG0000004162 | chr9 | Gucy1a2 | -0.51294621 | 0.519316215 |
| ENSMUSG0000004471 | chr18 | E230025N22Rik | -1.219920187 | 0.519939772 |
| ENSMUSG0000002803 | chr3 | Dkk2 | -0.326050829 | 0.519939772 |
| ENSMUSG0000003181 | chr8 | Map1lc3b | 0.249774283 | 0.520231037 |
| ENSMUSG0000002102 | chr12 | Nfkbia | 0.228873053 | 0.520231037 |
| ENSMUSG0000005910 | chr7 | Ifitm6 | -0.785212347 | 0.52031951 |
| ENSMUSG0000007521 | chr2 | Fads2b | -1.885860692 | 0.520456814 |
| ENSMUSG0000002916 | chr5 | Dpysl5 | -0.249574565 | 0.520456814 |
| ENSMUSG0000005282 | chrX | Cysltr1 | -0.361331428 | 0.520456814 |
| ENSMUSG0000003915 | chr17 | Runx2 | -0.58648876 | 0.520456814 |
| ENSMUSG0000002345 | chr5 | Pisd | -0.174768358 | 0.520456814 |
| ENSMUSG0000003381 | chr15 | Ppp1r16a | -0.168146633 | 0.520456814 |
| ENSMUSG0000002453 | chr18 | Snx24 | 0.21525679 | 0.520456814 |
| ENSMUSG0000001575 | chr14 | Cnih1 | 0.239785316 | 0.520456814 |
| ENSMUSG0000003199 | chr9 | Jam3 | 0.249905737 | 0.520456814 |
| ENSMUSG0000002088 | chr11 | Nr1d1 | -0.682819307 | 0.520473232 |
| ENSMUSG0000002818 | chr3 | Lrriq3 | 0.715556303 | 0.520473232 |
| ENSMUSG0000003567 | chr7 | Ndufa3 | 0.294922844 | 0.520473232 |
| ENSMUSG0000002623 | chr1 | Pde6d | 0.233267636 | 0.520554494 |
| ENSMUSG0000003880 | chr11 | Rap1gap2 | -0.253752643 | 0.520720297 |
| ENSMUSG0000007039 | chr11 | Tmem256 | 0.39305551 | 0.520720297 |
| ENSMUSG0000001980 | chr10 | Aig1 | 0.182446213 | 0.521087739 |
| ENSMUSG0000004270 | chr12 | Sipa1l1 | 0.181934335 | 0.521087739 |

| | | | | |
|---|---|---|---|---|
| ENSMUSG0000006290 | chr3 | Acadm | 0.206043739 | 0.521087739 |
| ENSMUSG0000004184 | chr18 | Haus1 | 0.234649234 | 0.521087739 |
| ENSMUSG0000003865 | chr7 | Rnh1 | 0.260504111 | 0.521087739 |
| ENSMUSG0000003510 | chr12 | Egln3 | 0.265777501 | 0.521087739 |
| ENSMUSG0000002468 | chr19 | Mrpl16 | 0.242917715 | 0.521087739 |
| ENSMUSG0000002202 | chr14 | Diaph3 | 0.393414088 | 0.521087739 |
| ENSMUSG0000000141 | chr3 | Cct3 | 0.290216332 | 0.521087739 |
| ENSMUSG0000002125 | chr12 | Esrrb | 0.748476999 | 0.521087739 |
| ENSMUSG0000000548 | chr8 | Dnajb1 | 0.315636692 | 0.521246559 |
| ENSMUSG0000001146 | chr3 | Cpb1 | -1.406071974 | 0.521352287 |
| ENSMUSG0000003807 | chr6 | Kcna6 | -0.225949966 | 0.521865315 |
| ENSMUSG0000003848 | chr8 | Tfdp1 | 0.164837445 | 0.52208595 |
| ENSMUSG0000002815 | chr3 | Eif4e | 0.228797087 | 0.52208595 |
| ENSMUSG0000006322 | chr7 | Ldha | 0.299036094 | 0.52208595 |
| ENSMUSG0000003858 | chr5 | Pptc7 | 0.127388655 | 0.522189522 |
| ENSMUSG0000002664 | chr2 | Nmt2 | -0.200741735 | 0.522433146 |
| ENSMUSG0000005776 | chr13 | Prp2rt | 0.590720998 | 0.522433146 |
| ENSMUSG0000002617 | chr1 | Plcd4 | -0.26089201 | 0.522846839 |
| ENSMUSG0000003486 | chr17 | Myl12b | -0.323102255 | 0.522846839 |
| ENSMUSG0000006380 | chr4 | Prpf38a | 0.160376403 | 0.522846839 |
| ENSMUSG0000002999 | chr6 | Nfu1 | 0.230589806 | 0.522846839 |
| ENSMUSG0000000640 | chr1 | Adamts4 | 1.451056649 | 0.522846839 |
| ENSMUSG0000003443 | chr5 | Gbp8 | -0.767422924 | 0.522883075 |
| ENSMUSG0000006254 | chr4 | Tlr12 | -1.602115575 | 0.522883075 |
| ENSMUSG0000005407 | chr18 | Iigp1 | -0.585427765 | 0.522883075 |
| ENSMUSG0000004538 | chr1 | Cxcr4 | -0.446658179 | 0.522883075 |
| ENSMUSG0000000112 | chr11 | Lgals9 | -0.275486452 | 0.522883075 |
| ENSMUSG0000004933 | chr1 | Retreg2 | -0.139658931 | 0.522883075 |
| ENSMUSG0000003352 | chr2 | Ppip5k1 | -0.135513674 | 0.522883075 |
| ENSMUSG0000003903 | chr2 | Tasp1 | 0.214799597 | 0.522883075 |
| ENSMUSG0000002415 | chr17 | Mcfd2 | 0.213479059 | 0.522883075 |
| ENSMUSG0000002017 | chr11 | Grb10 | 0.211110974 | 0.522883075 |
| ENSMUSG0000003525 | chr16 | Abi3bp | 0.260215824 | 0.522883075 |
| ENSMUSG0000002046 | chr11 | Rps27a | 0.294785469 | 0.522883075 |
| ENSMUSG0000002932 | chr5 | Slc10a6 | 0.549602312 | 0.522883075 |
| ENSMUSG0000003638 | chr3 | P2ry14 | 0.433914957 | 0.522883075 |
| ENSMUSG0000006015 | chr5 | Pop5 | 0.354269251 | 0.522992439 |
| ENSMUSG0000002282 | chr16 | Abcc5 | -0.147682027 | 0.523035037 |
| ENSMUSG0000003239 | chr9 | Rpl4 | 0.213775668 | 0.523217282 |
| ENSMUSG0000004675 | chr7 | Selenov | -0.617816802 | 0.523284723 |
| ENSMUSG0000003655 | chr17 | Ermard | 0.254015179 | 0.523284723 |
| ENSMUSG0000003685 | chr2 | Mrpl41 | 0.271026358 | 0.523284723 |
| ENSMUSG0000002078 | chr11 | Camkk1 | 0.256755438 | 0.523284723 |
| ENSMUSG0000002900 | chr4 | Draxin | -0.671537678 | 0.523590898 |
| ENSMUSG0000003087 | chr7 | Mrpl17 | 0.312133952 | 0.523590898 |
| ENSMUSG0000003968 | chr5 | Sdk1 | 0.354580603 | 0.523590898 |
| ENSMUSG0000003169 | chr8 | Orc6 | 0.192718722 | 0.523793269 |
| ENSMUSG0000004242 | chrX | Frmpd3 | 0.336888662 | 0.523864412 |
| ENSMUSG0000006869 | chr3 | Gpr88 | -0.687645001 | 0.523911004 |
| ENSMUSG0000002015 | chr10 | Ptprr | -0.318110604 | 0.523989793 |

| | | | | |
|---|---|---|---|---|
| ENSMUSG0000006913 | chr2 | Nxph2 | -1.132342646 | 0.524653142 |
| ENSMUSG0000004023 | chr6 | Tm7sf3 | -0.161968821 | 0.524653142 |
| ENSMUSG0000002500 | chr19 | Hells | -0.463112393 | 0.524961138 |
| ENSMUSG0000005001 | chr15 | Apol10b | -0.303052039 | 0.524961138 |
| ENSMUSG0000003766 | chr12 | Gdf7 | -0.616528663 | 0.525019292 |
| ENSMUSG0000002531 | chr2 | Ptprj | -0.211910991 | 0.525019292 |
| ENSMUSG0000003151 | chr8 | Dctn6 | 0.169485777 | 0.525019292 |
| ENSMUSG0000002162 | chr13 | Slc30a5 | 0.160134159 | 0.525019292 |
| ENSMUSG0000002541 | chr10 | Dctn2 | 0.126744932 | 0.525037833 |
| ENSMUSG0000003840 | chr2 | Pmepa1 | 0.286757318 | 0.525037833 |
| ENSMUSG0000002870 | chr4 | Lurap1 | -0.235624376 | 0.525251205 |
| ENSMUSG0000007312 | chrX | Xlr3b | 0.644348026 | 0.525251205 |
| ENSMUSG0000002652 | chr1 | Rgs7 | -0.2041613 | 0.525371525 |
| ENSMUSG0000002423 | chr18 | Svil | 0.177171418 | 0.525371525 |
| ENSMUSG0000002075 | chr11 | Recql5 | -0.218152297 | 0.525581946 |
| ENSMUSG0000002159 | chr13 | Med10 | 0.262104695 | 0.525581946 |
| ENSMUSG0000000233 | chr14 | Dhrs1 | 0.216833251 | 0.525977615 |
| ENSMUSG0000003846 | chr13 | Uqcrfs1 | 0.232386629 | 0.525977615 |
| ENSMUSG0000003637 | chr19 | Tmem258 | 0.276932282 | 0.525977615 |
| ENSMUSG0000002136 | chr13 | Tmem14c | 0.315887944 | 0.525977615 |
| ENSMUSG0000003013 | chr6 | Mug2 | 1.094490311 | 0.525977615 |
| ENSMUSG0000007262 | chr14 | Gm5460 | 0.616110159 | 0.526520224 |
| ENSMUSG0000005563 | chr7 | Zfp580 | 0.248646984 | 0.526728572 |
| ENSMUSG0000001576 | chr6 | Eps8 | -0.332650584 | 0.526733907 |
| ENSMUSG0000002823 | chr4 | Rps20 | 0.28157479 | 0.526746275 |
| ENSMUSG0000005572 | chr9 | Ubl7 | 0.2172223 | 0.52691333 |
| ENSMUSG0000000602 | chr7 | Kptn | 0.229182308 | 0.527192877 |
| ENSMUSG0000005033 | chrX | Amer1 | -0.186223119 | 0.527522419 |
| ENSMUSG0000003329 | chr5 | Noc4l | 0.202171389 | 0.527522419 |
| ENSMUSG0000003209 | chr9 | Cd3e | -1.745596214 | 0.527942374 |
| ENSMUSG0000006686 | chr5 | Oas1g | -0.511769104 | 0.527942374 |
| ENSMUSG0000003789 | chr12 | Rcor1 | -0.142202727 | 0.527945277 |
| ENSMUSG0000002438 | chr18 | Bin1 | -0.151362671 | 0.528028201 |
| ENSMUSG0000004190 | chr1 | Gpr45 | -0.322306895 | 0.528326328 |
| ENSMUSG0000002450 | chr18 | Hsd17b4 | 0.170149317 | 0.528419067 |
| ENSMUSG0000003916 | chr2 | Naif1 | -0.268592847 | 0.52853988 |
| ENSMUSG0000000703 | chr17 | Neu1 | -0.242201403 | 0.52853988 |
| ENSMUSG0000002295 | chr16 | Runx1 | -0.288658346 | 0.52853988 |
| ENSMUSG0000002780 | chr3 | Pfn2 | -0.195437982 | 0.52853988 |
| ENSMUSG0000003861 | chr3 | Mcl1 | 0.147016834 | 0.52853988 |
| ENSMUSG0000004228 | chr13 | Itga1 | 0.195156486 | 0.528549777 |
| ENSMUSG0000002498 | chr19 | Acsl5 | -0.150953103 | 0.528575926 |
| ENSMUSG0000003493 | chr6 | Rtkn | -0.298398374 | 0.52870848 |
| ENSMUSG0000004749 | chr1 | Rnf152 | -0.59095304 | 0.529370436 |
| ENSMUSG0000003195 | chr8 | Tmem231 | -0.174598603 | 0.529667138 |
| ENSMUSG0000004606 | chr1 | Ppp1r15b | -0.157717197 | 0.529667138 |
| ENSMUSG0000002114 | chr13 | Gtpbp4 | 0.176832069 | 0.529667138 |
| ENSMUSG0000007842 | chr10 | Sarnp | 0.250278771 | 0.529667138 |
| ENSMUSG0000002095 | chr12 | Dtd2 | 0.285059334 | 0.529667138 |
| ENSMUSG0000002796 | chr3 | Lrrc39 | 0.477065899 | 0.529667138 |

| | | | | |
|---|---|---|---|---|
| ENSMUSG0000001620 | chr17 | H2-M3 | -0.330882673 | 0.529833491 |
| ENSMUSG0000002260 | chr16 | Zbtb11 | -0.258474777 | 0.52985883 |
| ENSMUSG0000005484 | chr19 | Atrnl1 | -0.155854149 | 0.52985883 |
| ENSMUSG0000003920 | chr11 | Metrnl | -0.394182475 | 0.529900013 |
| ENSMUSG0000001587 | chr5 | Fam184b | -0.310555221 | 0.529900013 |
| ENSMUSG0000003835 | chr11 | Sgsm2 | -0.267174327 | 0.529900013 |
| ENSMUSG0000007398 | chr7 | Rhog | 0.18273071 | 0.529900013 |
| ENSMUSG0000004247 | chr12 | Zfp410 | 0.186955557 | 0.529900013 |
| ENSMUSG0000002522 | chr19 | Mfsd13a | -0.412850131 | 0.530043539 |
| ENSMUSG0000003845 | chr6 | Dennd2a | 0.185959215 | 0.530043539 |
| ENSMUSG0000000837 | chr7 | Prpf31 | 0.212073566 | 0.530043539 |
| ENSMUSG0000002608 | chr1 | Mitd1 | -0.20341012 | 0.530476012 |
| ENSMUSG0000010700 | chr16 | Ncbp2as2 | 0.207680543 | 0.530476012 |
| ENSMUSG0000002836 | chr4 | Txn1 | 0.247010077 | 0.530476012 |
| ENSMUSG0000001046 | chr10 | Eya4 | 0.326512451 | 0.530476012 |
| ENSMUSG0000002492 | chr19 | Vldlr | -0.289890802 | 0.530631665 |
| ENSMUSG0000007184 | chr18 | Apcdd1 | 0.268186665 | 0.530631665 |
| ENSMUSG0000007889 | chr2 | Gm4724 | 0.681165616 | 0.53089889 |
| ENSMUSG0000000907 | chr10 | Rsph14 | -1.617616783 | 0.530971725 |
| ENSMUSG0000005607 | chr10 | Grik2 | -0.335459358 | 0.530971725 |
| ENSMUSG0000005949 | chrX | Nhs | -0.331334331 | 0.530971725 |
| ENSMUSG0000004875 | chr9 | Prss50 | -0.857905492 | 0.530971725 |
| ENSMUSG0000004029 | chr4 | Rigi | -0.240965927 | 0.530971725 |
| ENSMUSG0000009380 | chr9 | Ppp2r3d | -0.15233856 | 0.530971725 |
| ENSMUSG0000002657 | chr1 | Dcaf6 | 0.146286178 | 0.530971725 |
| ENSMUSG0000002023 | chr10 | Hmg20b | 0.147352482 | 0.530971725 |
| ENSMUSG0000003256 | chr9 | Mrpl3 | 0.177651587 | 0.530971725 |
| ENSMUSG0000000839 | chr16 | Carhsp1 | 0.191215276 | 0.530971725 |
| ENSMUSG0000004599 | chr15 | Polr2k | 0.308428573 | 0.530971725 |
| ENSMUSG0000003940 | chr7 | Prss23 | 0.394234954 | 0.530971725 |
| ENSMUSG0000003125 | chrX | Nox1 | -1.577350009 | 0.531274221 |
| ENSMUSG0000002610 | chr1 | Nabp1 | -0.228204949 | 0.531274221 |
| ENSMUSG0000006026 | chr7 | Pwwp2b | 0.184829781 | 0.531274221 |
| ENSMUSG0000009468 | chr4 | Ccl21a | 0.226266408 | 0.531274221 |
| ENSMUSG0000001040 | chr14 | Mrpl52 | 0.359338163 | 0.531415662 |
| ENSMUSG0000002766 | chr3 | Zfp639 | 0.1760008 | 0.531425069 |
| ENSMUSG0000002154 | chr13 | Hnrnpk | 0.146701445 | 0.531456228 |
| ENSMUSG0000004273 | chr3 | Dpm3 | 0.40170126 | 0.531456228 |
| ENSMUSG0000002810 | chr3 | Nudt17 | -0.347007909 | 0.531786828 |
| ENSMUSG0000003812 | chr10 | Tbc1d32 | -0.273220614 | 0.531907609 |
| ENSMUSG0000001786 | chr2 | Mybl2 | -0.672294637 | 0.531907609 |
| ENSMUSG0000002983 | chr6 | Cbx3 | 0.186371626 | 0.531907609 |
| ENSMUSG0000006989 | chr11 | 9930111J21Rik2 | -0.381373161 | 0.532372985 |
| ENSMUSG0000006128 | chr7 | Exosc5 | 0.272455046 | 0.532582402 |
| ENSMUSG0000004749 | chr15 | Adamts12 | 0.369766579 | 0.532582402 |
| ENSMUSG0000005844 | chr15 | Panx2 | -0.345621869 | 0.532607214 |
| ENSMUSG0000007189 | chr6 | Vmn1r4 | -1.186263735 | 0.532607214 |
| ENSMUSG0000002648 | chr1 | Rgl1 | 0.129506638 | 0.532607214 |
| ENSMUSG0000004024 | chr6 | Fgfr1op2 | 0.155445347 | 0.532607214 |
| ENSMUSG0000002181 | chr14 | Ecd | 0.152152045 | 0.532607214 |

| | | | | |
|---|---|---|---|---|
| ENSMUSG0000001884 | chr11 | Rars1 | 0.183003562 | 0.532607214 |
| ENSMUSG0000002981 | chr6 | Malsu1 | 0.260628973 | 0.532607214 |
| ENSMUSG0000003549 | chr13 | Tgfbi | 0.331019922 | 0.532607214 |
| ENSMUSG0000000115 | chr6 | Snrnp27 | 0.286158531 | 0.532607214 |
| ENSMUSG0000003187 | chr8 | Ciao2b | 0.310478349 | 0.532607214 |
| ENSMUSG0000003504 | chr9 | Anapc13 | 0.29985841 | 0.532607214 |
| ENSMUSG0000004118 | chr19 | Hectd2 | 0.330778922 | 0.532623313 |
| ENSMUSG0000009086 | chr7 | Rps13 | 0.310272653 | 0.532665676 |
| ENSMUSG0000003988 | chr3 | Alg14 | 0.189780985 | 0.532855435 |
| ENSMUSG0000003890 | chr11 | Kat7 | -0.129053469 | 0.533032449 |
| ENSMUSG0000000109 | chr11 | Slc13a2 | -2.221925789 | 0.53304557 |
| ENSMUSG0000002300 | chr15 | Tuba1b | 0.269847748 | 0.533140681 |
| ENSMUSG0000000026 | chrX | Mid2 | 0.194181774 | 0.533731146 |
| ENSMUSG0000005162 | chr13 | H1f4 | -0.706811149 | 0.533960682 |
| ENSMUSG0000005150 | chr14 | Wdfy4 | -0.592223961 | 0.533960682 |
| ENSMUSG0000002237 | chr15 | Dnaaf11 | -0.737654935 | 0.533960682 |
| ENSMUSG0000001573 | chr6 | Capza2 | 0.196472564 | 0.533960682 |
| ENSMUSG0000002077 | chr11 | Mrpl38 | 0.172028575 | 0.533960682 |
| ENSMUSG0000006345 | chr10 | Rps15 | 0.324978613 | 0.533960682 |
| ENSMUSG0000000577 | chr3 | Psmb4 | 0.323895569 | 0.533960682 |
| ENSMUSG0000005091 | chr8 | Ankrd37 | 0.537014965 | 0.533960682 |
| ENSMUSG0000000568 | chr3 | Bcas2 | 0.129455005 | 0.534631034 |
| ENSMUSG0000002308 | chr16 | Abcc1 | 0.255956621 | 0.534697899 |
| ENSMUSG0000002586 | chr13 | Nop16 | 0.273184313 | 0.534697899 |
| ENSMUSG0000002919 | chr5 | Klb | -0.948735027 | 0.534919316 |
| ENSMUSG0000006326 | chr15 | Parp10 | -0.200745379 | 0.534950899 |
| ENSMUSG0000009100 | chr7 | Tcerg1l | 0.404132768 | 0.534950899 |
| ENSMUSG0000006299 | chr2 | Rpl35 | 0.302626525 | 0.534950899 |
| ENSMUSG0000005261 | chr10 | Pcdh15 | -0.42000056 | 0.535177972 |
| ENSMUSG0000003153 | chr8 | Dusp4 | -0.421659573 | 0.535177972 |
| ENSMUSG0000003005 | chr6 | Isy1 | 0.211112639 | 0.535177972 |
| ENSMUSG0000006211 | chr5 | Scfd2 | 0.238626228 | 0.535177972 |
| ENSMUSG0000004528 | chr7 | Tmem86b | 0.216676509 | 0.53542873 |
| ENSMUSG0000005990 | chr10 | Adamts14 | 0.506106565 | 0.53542873 |
| ENSMUSG0000006428 | chr2 | Tank | 0.165317917 | 0.535862374 |
| ENSMUSG0000004363 | chr13 | Ecm2 | 0.22160633 | 0.535862374 |
| ENSMUSG0000004408 | chr3 | S100a1 | 0.246282861 | 0.535878216 |
| ENSMUSG0000000605 | chr11 | Snf8 | 0.250234215 | 0.53600055 |
| ENSMUSG0000003888 | chr7 | Ctu1 | 0.235455062 | 0.53605885 |
| ENSMUSG0000002066 | chr12 | Dld | 0.143829087 | 0.536073626 |
| ENSMUSG0000004882 | chr17 | Dact2 | -0.624072288 | 0.536180926 |
| ENSMUSG0000002679 | chr2 | Stxbp1 | -0.248027058 | 0.53646575 |
| ENSMUSG0000002484 | chr19 | Aip | 0.301125096 | 0.53646575 |
| ENSMUSG0000000032 | chr16 | Comt | 0.264626557 | 0.536561454 |
| ENSMUSG0000002399 | chr17 | Tfeb | 0.182655084 | 0.536564803 |
| ENSMUSG0000003800 | chr4 | Acer2 | 0.3295238 | 0.536564803 |
| ENSMUSG0000004492 | chr6 | H1f10 | -0.271535132 | 0.536604556 |
| ENSMUSG0000007423 | chr3 | Ap1ar | -0.156939992 | 0.536604556 |
| ENSMUSG0000003286 | chr5 | Fbxw8 | 0.183316179 | 0.536604556 |
| ENSMUSG0000002104 | chr12 | Adck1 | 0.217845773 | 0.536604556 |

| | | | | |
|---|---|---|---|---|
| ENSMUSG0000004227 | chrX | Nxt2 | 0.198436162 | 0.536604556 |
| ENSMUSG0000006426 | chr7 | Zfp428 | 0.37389901 | 0.536604556 |
| ENSMUSG0000003186 | chr6 | Dctn1 | 0.101005018 | 0.536674514 |
| ENSMUSG0000006113 | chr17 | Ppm1b | 0.137821894 | 0.536674514 |
| ENSMUSG0000003106 | chr7 | Glrx3 | 0.265636532 | 0.536674514 |
| ENSMUSG0000002991 | chr6 | Clec5a | 0.567422085 | 0.536739163 |
| ENSMUSG0000002013 | chr10 | Pcsk4 | -0.580167233 | 0.536868265 |
| ENSMUSG0000005485 | chr15 | Rnd1 | 0.459346824 | 0.536868265 |
| ENSMUSG0000002030 | chr10 | Cdc34 | 0.235262039 | 0.536943909 |
| ENSMUSG0000001455 | chr6 | Mrps25 | 0.318401153 | 0.536943909 |
| ENSMUSG0000002662 | chr1 | Mark1 | 0.20511424 | 0.536952299 |
| ENSMUSG0000006004 | chr10 | Tmem26 | -0.376596374 | 0.536991023 |
| ENSMUSG0000001459 | chr4 | Camta1 | -0.171632567 | 0.536991023 |
| ENSMUSG0000002917 | chr5 | Anapc4 | 0.109335066 | 0.536991023 |
| ENSMUSG0000003835 | chr3 | Ankrd35 | -0.384384824 | 0.53735963 |
| ENSMUSG0000002294 | chr16 | Dop1b | -0.229714844 | 0.53735963 |
| ENSMUSG0000002090 | chr11 | Myh10 | 0.20475424 | 0.53735963 |
| ENSMUSG0000002457 | chr18 | Pcyox1l | 0.220383337 | 0.53735963 |
| ENSMUSG0000007256 | chr15 | Lratd2 | -0.398721257 | 0.537426871 |
| ENSMUSG0000004627 | chr11 | Trarg1 | 1.082949431 | 0.537854912 |
| ENSMUSG0000002420 | chr17 | Uqcc2 | 0.411770507 | 0.537854912 |
| ENSMUSG0000000245 | chr2 | Prpf6 | 0.135134576 | 0.537902834 |
| ENSMUSG0000002731 | chr2 | Adam33 | 0.167195281 | 0.537902834 |
| ENSMUSG0000005481 | chr5 | Usp46 | 0.198231973 | 0.537902834 |
| ENSMUSG0000004861 | chr19 | Myof | 0.249354781 | 0.537902834 |
| ENSMUSG0000003123 | chrX | Cox7b | 0.27325673 | 0.538068839 |
| ENSMUSG0000006431 | chr16 | Zpld1 | -1.061253579 | 0.538085353 |
| ENSMUSG0000002843 | chr4 | Fam219a | -0.252933761 | 0.538085353 |
| ENSMUSG0000000366 | chr17 | Has1 | 0.420572438 | 0.538085353 |
| ENSMUSG0000004194 | chr8 | Tango6 | 0.232316789 | 0.538361651 |
| ENSMUSG0000002167 | chr13 | Iqgap2 | -0.342398571 | 0.538370991 |
| ENSMUSG0000005554 | chr16 | Epha6 | -0.460647856 | 0.538370991 |
| ENSMUSG0000002975 | chr6 | Pon3 | -0.263569567 | 0.538483326 |
| ENSMUSG0000002499 | chr19 | Dennd10 | 0.154500022 | 0.538498042 |
| ENSMUSG0000002240 | chr15 | Slc25a17 | 0.186878817 | 0.539399374 |
| ENSMUSG0000004711 | chr6 | Fam221a | 0.283460737 | 0.539399374 |
| ENSMUSG0000001485 | chr13 | Msh3 | -0.19902696 | 0.539583539 |
| ENSMUSG0000002805 | chr3 | Arhgef2 | -0.163499686 | 0.539583539 |
| ENSMUSG0000007523 | chr10 | Amd1 | 0.256850876 | 0.539583539 |
| ENSMUSG0000002048 | chr11 | Supt4a | 0.269031613 | 0.539583539 |
| ENSMUSG0000001083 | chr15 | Kdelr3 | 0.316722413 | 0.539583539 |
| ENSMUSG0000004195 | chr3 | S100a10 | 0.322480959 | 0.539724835 |
| ENSMUSG0000007360 | chr1 | D2hgdh | -0.402089776 | 0.539744208 |
| ENSMUSG0000004977 | chrX | Tmsb4x | 0.303764766 | 0.539744208 |
| ENSMUSG0000002481 | chr19 | Slc25a45 | -0.378665958 | 0.539806548 |
| ENSMUSG0000005021 | chr4 | Eva1b | 0.292518897 | 0.539962031 |
| ENSMUSG0000002864 | chr4 | Slc2a1 | -0.345262385 | 0.540006976 |
| ENSMUSG0000003149 | chr7 | Ggn | 0.297400491 | 0.540006976 |
| ENSMUSG0000002119 | chr12 | Chga | -0.279467671 | 0.540021437 |
| ENSMUSG0000004028 | chr3 | Hey1 | 0.199330412 | 0.540021437 |

| | | | | |
|---|---|---|---|---|
| ENSMUSG0000004474 | chr11 | Mtfp1 | -0.439894982 | 0.5401316 |
| ENSMUSG0000004405 | chr11 | Ccr10 | -0.50072792 | 0.5401316 |
| ENSMUSG000000431 | chr8 | Clcn3 | -0.195812071 | 0.5401316 |
| ENSMUSG0000005185 | chr15 | Arf3 | -0.146970774 | 0.5401316 |
| ENSMUSG0000000493 | chr10 | Sgta | 0.121704493 | 0.5401316 |
| ENSMUSG0000005620 | chr19 | Cfl1 | 0.194521518 | 0.5401316 |
| ENSMUSG0000000105 | chr11 | Rmnd5b | 0.191971023 | 0.5401316 |
| ENSMUSG0000002454 | chr18 | Ldlrad4 | 0.216579718 | 0.5401316 |
| ENSMUSG0000003970 | chr3 | Usp53 | -0.301671252 | 0.540135274 |
| ENSMUSG0000002923 | chr5 | Pdgfra | 0.241880415 | 0.540135274 |
| ENSMUSG0000003245 | chr9 | Rbp2 | -1.294618867 | 0.540220645 |
| ENSMUSG0000006683 | chr9 | Ecsit | 0.157628209 | 0.540223419 |
| ENSMUSG0000003880 | chr5 | Ost4 | 0.325514556 | 0.540223419 |
| ENSMUSG0000005165 | chr10 | Lrrc3 | 0.194158514 | 0.540773016 |
| ENSMUSG0000003114 | chrX | Prickle3 | -0.189489798 | 0.541101651 |
| ENSMUSG0000005732 | chr11 | Rpl38 | 0.308533115 | 0.54143559 |
| ENSMUSG0000005055 | chr5 | Lamtor4 | 0.291161981 | 0.54143559 |
| ENSMUSG0000003107 | chr7 | Ano1 | -0.296863211 | 0.541684857 |
| ENSMUSG0000010977 | chr5 | Gm35315 | -0.574282758 | 0.541684857 |
| ENSMUSG0000002201 | chr14 | Akap11 | -0.168279077 | 0.541684857 |
| ENSMUSG0000001926 | chr8 | Map1s | 0.132263965 | 0.541684857 |
| ENSMUSG0000007947 | chr6 | Rab7 | 0.158150382 | 0.541684857 |
| ENSMUSG0000003151 | chr8 | Arhgef7 | 0.156652531 | 0.541684857 |
| ENSMUSG0000002865 | chr4 | Trit1 | 0.19317037 | 0.541684857 |
| ENSMUSG0000002860 | chr4 | Tnfrsf8 | 0.948255486 | 0.541684857 |
| ENSMUSG0000002520 | chr19 | Ndufb8 | 0.287964231 | 0.541684857 |
| ENSMUSG0000006757 | chr2 | Cbln4 | 0.597860188 | 0.541684857 |
| ENSMUSG0000002255 | chr15 | Bop1 | 0.250794678 | 0.541967258 |
| ENSMUSG0000002806 | chr3 | Gpatch4 | 0.304680075 | 0.541967258 |
| ENSMUSG0000002301 | chr15 | Racgap1 | -0.42553737 | 0.542133881 |
| ENSMUSG0000004911 | chr6 | Oxtr | -0.337928084 | 0.542133881 |
| ENSMUSG0000002160 | chr13 | Lpcat1 | -0.177193279 | 0.542133881 |
| ENSMUSG0000002959 | chr5 | Ddx54 | 0.191148533 | 0.542133881 |
| ENSMUSG0000006378 | chrX | Utp14a | 0.206577319 | 0.542133881 |
| ENSMUSG0000000493 | chr10 | Apba3 | 0.222669914 | 0.542133881 |
| ENSMUSG0000003523 | chr14 | Trim13 | 0.274280151 | 0.542133881 |
| ENSMUSG0000003264 | chr4 | Fhl3 | 0.521299298 | 0.542133881 |
| ENSMUSG0000003275 | chr5 | Slc8b1 | 0.217499089 | 0.542140963 |
| ENSMUSG0000003007 | chr6 | Ruvbl1 | 0.242684995 | 0.542213813 |
| ENSMUSG0000000312 | chr7 | Lipe | 0.341201613 | 0.542213813 |
| ENSMUSG0000001874 | chr11 | Slc25a35 | -0.24513197 | 0.542712717 |
| ENSMUSG0000003030 | chr6 | Ergic2 | 0.147023695 | 0.542712717 |
| ENSMUSG0000000558 | chr13 | Mef2c | 0.209683127 | 0.542822377 |
| ENSMUSG0000000341 | chr2 | St8sia6 | 0.515830837 | 0.543535532 |
| ENSMUSG0000002664 | chr2 | Dclre1c | -0.297872234 | 0.543990754 |
| ENSMUSG0000002829 | chr4 | Slc35a1 | -0.226294764 | 0.544021776 |
| ENSMUSG0000000478 | chr12 | Dlst | 0.146227352 | 0.544021776 |
| ENSMUSG0000002878 | chr5 | Sema3c | 0.153113994 | 0.544021776 |
| ENSMUSG0000004401 | chr4 | Mrpl50 | 0.155917762 | 0.544021776 |
| ENSMUSG0000006408 | chr6 | Fbln2 | 0.231035033 | 0.544021776 |

| | | | | |
|---|---|---|---|---|
| ENSMUSG0000002091 | chr11 | Top2a | -0.8230278 | 0.544209371 |
| ENSMUSG0000004972 | chr9 | Mmp12 | -1.035530084 | 0.544209371 |
| ENSMUSG0000003838 | chr6 | Pals2 | -0.518356654 | 0.544209371 |
| ENSMUSG0000009079 | chr14 | Klhl33 | -0.417389786 | 0.544209371 |
| ENSMUSG0000004915 | chr11 | Cfap144 | -0.515625327 | 0.544209371 |
| ENSMUSG0000003915 | chr17 | Shd | -0.430898514 | 0.544209371 |
| ENSMUSG0000002454 | chr18 | Cep192 | -0.269816413 | 0.544209371 |
| ENSMUSG0000003734 | chrX | Chst7 | -0.479162215 | 0.544209371 |
| ENSMUSG0000009687 | chr5 | Gm21083 | -1.072524738 | 0.544209371 |
| ENSMUSG0000005019 | chr3 | Eif5a2 | -0.240566558 | 0.544209371 |
| ENSMUSG0000003032 | chr6 | Rho | -0.589363585 | 0.544209371 |
| ENSMUSG0000003569 | chr10 | Arhgap45 | -0.322733234 | 0.544209371 |
| ENSMUSG0000000063 | chr11 | Myo18a | -0.215401579 | 0.544209371 |
| ENSMUSG0000009817 | chr15 | Ccdc166 | -0.292422761 | 0.544209371 |
| ENSMUSG0000005185 | chr6 | Mest | -0.207411499 | 0.544209371 |
| ENSMUSG0000002015 | chr10 | Gamt | -0.225714703 | 0.544209371 |
| ENSMUSG0000000079 | chr3 | Kcnn3 | -0.199162928 | 0.544209371 |
| ENSMUSG0000002098 | chr12 | L2hgdh | -0.193976033 | 0.544209371 |
| ENSMUSG0000002915 | chr5 | Sgcb | 0.121587573 | 0.544209371 |
| ENSMUSG0000003764 | chr3 | Prkci | 0.129129904 | 0.544209371 |
| ENSMUSG0000005648 | chr19 | Cd248 | 0.131208279 | 0.544209371 |
| ENSMUSG0000003031 | chr6 | Atg7 | 0.13409724 | 0.544209371 |
| ENSMUSG0000002765 | chr2 | Dhx35 | 0.140214174 | 0.544209371 |
| ENSMUSG0000003719 | chr2 | Rbm17 | 0.151006448 | 0.544209371 |
| ENSMUSG0000002766 | chr3 | Pik3ca | 0.151094487 | 0.544209371 |
| ENSMUSG0000003973 | chr5 | Prkrip1 | 0.15995325 | 0.544209371 |
| ENSMUSG0000000101 | chr3 | Snapin | 0.161038748 | 0.544209371 |
| ENSMUSG0000002587 | chr13 | Uimc1 | 0.18881171 | 0.544209371 |
| ENSMUSG0000007287 | chr6 | Rybp | 0.189896988 | 0.544209371 |
| ENSMUSG0000003900 | chr15 | Cpq | 0.195792025 | 0.544209371 |
| ENSMUSG0000002768 | chr3 | Fxr1 | 0.195000663 | 0.544209371 |
| ENSMUSG0000004177 | chr1 | Tram2 | 0.225862946 | 0.544209371 |
| ENSMUSG0000003230 | chr9 | Fbxo22 | 0.225527306 | 0.544209371 |
| ENSMUSG0000003343 | chr3 | Trmt13 | 0.272453665 | 0.544209371 |
| ENSMUSG0000002787 | chr3 | Notch2 | 0.222965623 | 0.544209371 |
| ENSMUSG0000002334 | chr5 | Trip6 | 0.256387236 | 0.544209371 |
| ENSMUSG0000004108 | chr3 | Ostc | 0.302472619 | 0.544209371 |
| ENSMUSG0000002026 | chr11 | Hint1 | 0.322464981 | 0.544209371 |
| ENSMUSG0000005631 | chr8 | Tcim | 0.291198663 | 0.544209371 |
| ENSMUSG0000009153 | chr9 | Tma7 | 0.332320395 | 0.544209371 |
| ENSMUSG0000004246 | chr7 | Dctpp1 | 0.436208052 | 0.544209371 |
| ENSMUSG0000005300 | chr6 | Creb5 | 0.349228274 | 0.544209371 |
| ENSMUSG0000002834 | chr4 | Nr4a3 | 1.085548711 | 0.544209371 |
| ENSMUSG0000002070 | chr11 | Asic2 | -0.239712844 | 0.544826443 |
| ENSMUSG0000003157 | chr8 | Star | -0.25189961 | 0.544826443 |
| ENSMUSG0000003260 | chr9 | Amt | -0.235084424 | 0.544826443 |
| ENSMUSG0000003140 | chrX | G6pdx | -0.171147932 | 0.544826443 |
| ENSMUSG0000002627 | chr1 | Stk25 | -0.136421309 | 0.544826443 |
| ENSMUSG0000002084 | chr11 | Blmh | 0.14006836 | 0.544826443 |
| ENSMUSG0000003947 | chr5 | Wfs1 | 0.149827914 | 0.544826443 |

| | | | | |
|---|---|---|---|---|
| ENSMUSG0000005673 | chr6 | Capg | 0.213539831 | 0.544826443 |
| ENSMUSG0000005696 | chr11 | Jmjd6 | 0.207737221 | 0.544826443 |
| ENSMUSG0000003480 | chr2 | Zfp661 | 0.283682038 | 0.544826443 |
| ENSMUSG0000003129 | chrX | Cdkl5 | 0.232184129 | 0.544826443 |
| ENSMUSG0000002406 | chr17 | Dpy30 | 0.297354489 | 0.544826443 |
| ENSMUSG0000002681 | chr2 | Eng | 0.221726361 | 0.545193325 |
| ENSMUSG0000004680 | chr19 | Mpeg1 | -0.327626834 | 0.54532865 |
| ENSMUSG0000002199 | chr14 | Cacna2d3 | 0.293683503 | 0.545484846 |
| ENSMUSG0000003800 | chr8 | Hpf1 | 0.147531255 | 0.545659161 |
| ENSMUSG0000000095 | chr14 | Slc7a7 | -0.167859202 | 0.546243202 |
| ENSMUSG0000002946 | chr7 | Fam168a | 0.130995581 | 0.546243202 |
| ENSMUSG0000004116 | chr17 | Lonp1 | 0.111312604 | 0.546748867 |
| ENSMUSG0000003695 | chrX | Bcorl1 | -0.25425642 | 0.546837477 |
| ENSMUSG0000009036 | chr19 | 4933411K16Rik | -0.726767334 | 0.547040726 |
| ENSMUSG0000004798 | chr8 | Palm3 | -0.484965385 | 0.547065723 |
| ENSMUSG0000002054 | chr11 | Stxbp4 | -0.169043274 | 0.547065723 |
| ENSMUSG0000004433 | chr2 | Aplnr | -0.430960552 | 0.547457782 |
| ENSMUSG0000004345 | chr18 | Pcdhb12 | -0.547392715 | 0.547457782 |
| ENSMUSG0000003682 | chr3 | Ssx2ip | -0.175651108 | 0.547457782 |
| ENSMUSG0000002127 | chr12 | Traf3 | 0.156208209 | 0.547457782 |
| ENSMUSG0000005237 | chr13 | Actn2 | 0.511442757 | 0.547457782 |
| ENSMUSG0000000352 | chr16 | Prodh | -0.415879553 | 0.547565269 |
| ENSMUSG0000004719 | chr9 | Dync2h1 | -0.259060178 | 0.547565269 |
| ENSMUSG0000004909 | chr18 | Ptgr3 | 0.170596706 | 0.547565269 |
| ENSMUSG0000000992 | chr9 | Rps25 | 0.30618746 | 0.547565269 |
| ENSMUSG0000009878 | chr2 | Jmjd7 | -0.97102752 | 0.547953365 |
| ENSMUSG0000002610 | chr1 | Gls | -0.158599537 | 0.547953365 |
| ENSMUSG0000004281 | chr8 | Foxf1 | 0.153046629 | 0.547953365 |
| ENSMUSG0000002890 | chr4 | Sf3a3 | 0.179955531 | 0.547953365 |
| ENSMUSG0000002010 | chr10 | Anapc16 | 0.224048041 | 0.547953365 |
| ENSMUSG0000002890 | chr4 | Utp11 | 0.23373513 | 0.547953365 |
| ENSMUSG0000002679 | chr2 | Med27 | 0.253450903 | 0.547953365 |
| ENSMUSG0000003147 | chr8 | Nek3 | -0.273792772 | 0.548153703 |
| ENSMUSG0000002724 | chr2 | Ell3 | -1.314417471 | 0.548163464 |
| ENSMUSG0000003273 | chr18 | Ablim3 | -0.206847477 | 0.548163464 |
| ENSMUSG0000002999 | chr6 | Anxa4 | -0.195262539 | 0.548425766 |
| ENSMUSG0000003774 | chr9 | Eef1a1 | 0.223190153 | 0.548675699 |
| ENSMUSG0000004422 | chr15 | Dnajc21 | 0.193138206 | 0.548928949 |
| ENSMUSG0000003590 | chr7 | Dennd5a | 0.127611425 | 0.549162153 |
| ENSMUSG0000003448 | chr9 | Uaca | 0.17052126 | 0.549162153 |
| ENSMUSG0000003488 | chr10 | Cactin | 0.173756085 | 0.549162153 |
| ENSMUSG0000003871 | chr9 | Atp5mg | 0.258177032 | 0.549162153 |
| ENSMUSG0000005870 | chr7 | Egln2 | 0.156745009 | 0.549166907 |
| ENSMUSG0000000005 | chr11 | Narf | 0.23025782 | 0.549166907 |
| ENSMUSG0000004225 | chr1 | Pm20d1 | -1.232825873 | 0.549182182 |
| ENSMUSG0000006824 | chr15 | Apol9b | -0.652380094 | 0.549182182 |
| ENSMUSG0000003606 | chr2 | Slc2a6 | -0.388990742 | 0.549182182 |
| ENSMUSG0000004093 | chr11 | Slc16a11 | -0.424768977 | 0.549182182 |
| ENSMUSG0000002288 | chr16 | Masp1 | -0.269430959 | 0.549182182 |
| ENSMUSG0000003733 | chr11 | Hand1 | 0.228568282 | 0.549182182 |

| | | | | |
|---|---|---|---|---|
| ENSMUSG0000002441 | chr11 | Mrpl27 | 0.269300517 | 0.549182182 |
| ENSMUSG000000211C | chr12 | Glrx5 | 0.2839169 | 0.549182182 |
| ENSMUSG0000007451 | chr2 | Zfp971 | 0.210586445 | 0.549322705 |
| ENSMUSG0000007931 | chrX | Rab9 | 0.236460648 | 0.549322705 |
| ENSMUSG000000330S | chr15 | Nol12 | 0.245743535 | 0.549322705 |
| ENSMUSG000000027E | chr2 | Fkbp7 | 0.26660456 | 0.549322705 |
| ENSMUSG000000794E | chr2 | Phyhd1 | -0.607813655 | 0.549446326 |
| ENSMUSG0000002627 | chr1 | Bok | 0.289898969 | 0.549446326 |
| ENSMUSG0000007467 | chr2 | Foxs1 | 1.409542793 | 0.549446326 |
| ENSMUSG000000326E | chr18 | Malt1 | -0.301463025 | 0.549452208 |
| ENSMUSG000000494C | chr3 | Rarres1 | -0.812750862 | 0.549452208 |
| ENSMUSG000000517S | chr11 | Nlgn2 | -0.243764468 | 0.549452208 |
| ENSMUSG000000521E | chr4 | Rbm12b2 | -0.206240595 | 0.549452208 |
| ENSMUSG0000002021 | chr10 | Sf3a2 | 0.134500396 | 0.549452208 |
| ENSMUSG000000065S | chr7 | Gtf2h1 | 0.202122658 | 0.549452208 |
| ENSMUSG000000136S | chr1 | Pea15a | 0.236563313 | 0.549452208 |
| ENSMUSG000000207S | chr11 | Tekt1 | 0.339650384 | 0.549452208 |
| ENSMUSG0000006232 | chr5 | Cytl1 | 1.207776459 | 0.549452208 |
| ENSMUSG000000274E | chr2 | Snph | -0.205640046 | 0.549573748 |
| ENSMUSG0000003902 | chr2 | Ttc16 | -0.485478958 | 0.550061844 |
| ENSMUSG0000000151 | chr11 | Dlx3 | -0.324090389 | 0.550518688 |
| ENSMUSG0000003441 | chr17 | Neurl1b | -0.283995532 | 0.550562582 |
| ENSMUSG000000630E | chr6 | Ctnna2 | -0.228646846 | 0.550562582 |
| ENSMUSG000000358E | chr13 | Polr3g | -0.395259788 | 0.550562582 |
| ENSMUSG0000003584 | chr11 | Krt222 | -0.189688026 | 0.550562582 |
| ENSMUSG000000262C | chr1 | Speg | 0.16601023 | 0.550562582 |
| ENSMUSG0000002864 | chr4 | Ndufs5 | 0.293380094 | 0.550562582 |
| ENSMUSG000000403E | chr19 | Ppp1ca | 0.216807119 | 0.550669072 |
| ENSMUSG000000901E | chr6 | BC035044 | -0.783412392 | 0.551395857 |
| ENSMUSG000000744C | chr7 | Zfp628 | -0.192629673 | 0.551400662 |
| ENSMUSG000000079E | chr8 | Abhd8 | 0.155886003 | 0.551400662 |
| ENSMUSG000000614E | chr5 | Smim20 | 0.200320661 | 0.551400662 |
| ENSMUSG000000793E | chr1 | Lemd1 | -1.046061357 | 0.551543294 |
| ENSMUSG000000223E | chr15 | Khdrbs3 | 0.219816051 | 0.551543294 |
| ENSMUSG000000328C | chr9 | Cdv3 | 0.122774114 | 0.551569667 |
| ENSMUSG000000307E | chr7 | Copb1 | 0.157701441 | 0.551635627 |
| ENSMUSG0000001142 | chr7 | Zfp790 | 0.169858034 | 0.551635627 |
| ENSMUSG000000322C | chr9 | Rab27a | -0.166859997 | 0.552101639 |
| ENSMUSG000000539E | chr1 | Stum | 0.176800414 | 0.552101639 |
| ENSMUSG0000002301 | chr15 | Tmbim6 | -0.216303613 | 0.552705907 |
| ENSMUSG0000006111 | chr11 | Mcrip1 | 0.282219412 | 0.553255356 |
| ENSMUSG0000003372 | chr6 | Sfxn5 | -0.228417648 | 0.553332587 |
| ENSMUSG000000211E | chr13 | Zmynd11 | 0.147001723 | 0.553332587 |
| ENSMUSG000000227E | chr16 | Aifm3 | -0.675254825 | 0.553524872 |
| ENSMUSG0000004157 | chr1 | Prelp | 0.159247864 | 0.553524872 |
| ENSMUSG0000003651 | chr3 | Commd2 | 0.263286588 | 0.553524872 |
| ENSMUSG000000319C | chr8 | Dus2 | 0.294146867 | 0.553524872 |
| ENSMUSG000000162E | chrX | Lonrf3 | 0.554057704 | 0.553524872 |
| ENSMUSG000000393S | chr8 | Neil3 | -1.209584588 | 0.553595336 |
| ENSMUSG000000574E | chrX | Phex | -0.498525303 | 0.553595336 |

| | | | | |
|---|---|---|---|---|
| ENSMUSG0000000341 | chr9 | Elavl3 | -0.236946221 | 0.553595336 |
| ENSMUSG0000004069 | chr11 | Limd2 | -0.218686281 | 0.553595336 |
| ENSMUSG0000002178 | chr14 | Ngly1 | 0.140962949 | 0.553595336 |
| ENSMUSG0000002310 | chr5 | Denr | 0.170288845 | 0.553595336 |
| ENSMUSG0000004767 | chr4 | Rps8 | 0.298177009 | 0.553595336 |
| ENSMUSG0000010367 | chr18 | Pcdhga4 | 0.409366221 | 0.553595336 |
| ENSMUSG0000003820 | chr11 | Pgap3 | -0.254811068 | 0.553641092 |
| ENSMUSG0000000544 | chr7 | Pafah1b3 | -0.396256774 | 0.55369478 |
| ENSMUSG0000003591 | chr13 | Dcdc2a | -0.478173053 | 0.55369478 |
| ENSMUSG0000002807 | chr3 | Dclk2 | -0.239799367 | 0.55369478 |
| ENSMUSG0000004072 | chr14 | Zfhx2 | -0.225655429 | 0.55369478 |
| ENSMUSG0000001890 | chr11 | Slc22a5 | -0.233041781 | 0.55369478 |
| ENSMUSG0000006966 | chr10 | Marcks | -0.18648935 | 0.55369478 |
| ENSMUSG0000003204 | chr9 | Acat1 | 0.168584541 | 0.55369478 |
| ENSMUSG0000003467 | chr13 | Dbn1 | 0.167266231 | 0.55369478 |
| ENSMUSG0000007965 | chr1 | Eloc | 0.197548452 | 0.55369478 |
| ENSMUSG0000004726 | chr11 | Emc6 | 0.260476046 | 0.55369478 |
| ENSMUSG0000002245 | chr15 | Smdt1 | 0.289276066 | 0.55369478 |
| ENSMUSG0000002058 | chr12 | Cyria | 0.297196913 | 0.55369478 |
| ENSMUSG0000004422 | chr9 | Nxpe4 | 0.458640965 | 0.55369478 |
| ENSMUSG0000004744 | chr14 | Antxrl | 0.370613992 | 0.55369478 |
| ENSMUSG0000008236 | chr5 | Btc | 0.516977777 | 0.55369478 |
| ENSMUSG0000002966 | chr6 | Tspan12 | 0.274995466 | 0.553722143 |
| ENSMUSG0000007028 | chr9 | Slc35g2 | -0.31744741 | 0.554024048 |
| ENSMUSG0000002735 | chr2 | Bmp2 | -0.422192573 | 0.554488748 |
| ENSMUSG0000004872 | chr11 | Fndc9 | -0.435606775 | 0.55472711 |
| ENSMUSG0000010932 | chr7 | Prmt1 | 0.229244409 | 0.554903473 |
| ENSMUSG0000005631 | chr5 | Tyw1 | 0.183804641 | 0.555066209 |
| ENSMUSG0000003691 | chr17 | Ttc7 | 0.199463353 | 0.555066209 |
| ENSMUSG0000006007 | chr12 | Psma3 | 0.23607456 | 0.555066209 |
| ENSMUSG0000004855 | chr2 | Thnsl1 | 0.238802165 | 0.555128821 |
| ENSMUSG0000002578 | chr9 | Exosc7 | 0.253159939 | 0.555298892 |
| ENSMUSG0000003053 | chr7 | Unc45a | 0.231991376 | 0.555322226 |
| ENSMUSG0000001424 | chr11 | Pigl | 0.24795465 | 0.555322226 |
| ENSMUSG0000005674 | chr13 | Nfil3 | 1.031749746 | 0.555322226 |
| ENSMUSG0000002860 | chr4 | Podn | 0.371035388 | 0.555414465 |
| ENSMUSG0000005475 | chr4 | Fsd1l | -0.221371576 | 0.555712638 |
| ENSMUSG0000004157 | chr7 | Selenow | 0.245667111 | 0.555743903 |
| ENSMUSG0000003065 | chr7 | Arl6ip1 | -0.14791001 | 0.555824465 |
| ENSMUSG0000009465 | chr1 | Gal3st2 | -2.805150975 | 0.555876482 |
| ENSMUSG0000002479 | chr19 | Kif20b | -0.621833666 | 0.555876482 |
| ENSMUSG0000003089 | chr7 | Tpp1 | 0.134836514 | 0.555876482 |
| ENSMUSG0000010201 | chr15 | Iqank1 | -0.537279571 | 0.556043291 |
| ENSMUSG0000008737 | chr13 | Tmem170b | -0.221060473 | 0.556043291 |
| ENSMUSG0000002299 | chr15 | Adcy6 | -0.188923087 | 0.556043291 |
| ENSMUSG0000004827 | chr14 | Sacs | 0.347870615 | 0.556043291 |
| ENSMUSG0000003804 | chr17 | Cntnap5c | -1.878037175 | 0.556306711 |
| ENSMUSG0000006365 | chr11 | Slc22a21 | -0.313391012 | 0.556415212 |
| ENSMUSG0000007367 | chr1 | Hspe1 | 0.308509148 | 0.556415212 |
| ENSMUSG0000002196 | chr14 | Dcp1a | 0.158789212 | 0.556474467 |

| | | | | |
|---|---|---|---|---|
| ENSMUSG0000006695 | chr5 | Myo1h | 0.433247077 | 0.556474467 |
| ENSMUSG0000000134 | chr9 | Acp5 | -0.624050089 | 0.556500794 |
| ENSMUSG0000005066 | chr10 | Trhde | -0.355792113 | 0.556500794 |
| ENSMUSG0000004271 | chrX | Tceal9 | 0.21113915 | 0.556601847 |
| ENSMUSG0000002872 | chr4 | Ebna1bp2 | 0.133183548 | 0.556704697 |
| ENSMUSG0000000781 | chr9 | Rhoa | 0.177585488 | 0.556909727 |
| ENSMUSG0000002913 | chr5 | Rnf32 | -0.289560034 | 0.55740338 |
| ENSMUSG0000004932 | chr11 | Zfp2 | -0.227782533 | 0.55740338 |
| ENSMUSG0000004694 | chr13 | Nqo2 | -0.173436337 | 0.55740338 |
| ENSMUSG0000004745 | chr2 | Dynlrb1 | 0.244036366 | 0.55740338 |
| ENSMUSG0000002764 | chr2 | Rbl1 | -0.259836746 | 0.557512363 |
| ENSMUSG0000003872 | chr5 | Bud31 | 0.267437483 | 0.557512363 |
| ENSMUSG0000002149 | chr13 | Pcbd2 | 0.360170836 | 0.557512363 |
| ENSMUSG0000003080 | chr7 | Stx1b | -0.289817774 | 0.558121929 |
| ENSMUSG0000002760 | chr2 | Map1lc3a | 0.263887687 | 0.558121929 |
| ENSMUSG0000003564 | chr10 | Cbarp | -0.304704856 | 0.558125302 |
| ENSMUSG0000001516 | chr7 | Hnrnpl | 0.100941832 | 0.558125302 |
| ENSMUSG0000002421 | chr17 | Mllt1 | 0.145937953 | 0.558125302 |
| ENSMUSG0000004960 | chr7 | Zbtb45 | 0.186246678 | 0.558125302 |
| ENSMUSG0000002137 | chr13 | Mcur1 | 0.183579237 | 0.558125302 |
| ENSMUSG0000002822 | chr4 | Decr1 | 0.214087649 | 0.558125302 |
| ENSMUSG0000002307 | chr4 | Akirin1 | 0.234506511 | 0.558125302 |
| ENSMUSG0000004355 | chr15 | Fbxl7 | 0.256719897 | 0.558125302 |
| ENSMUSG0000006147 | chr12 | Rps7 | 0.335186332 | 0.558125302 |
| ENSMUSG0000002971 | chr5 | Pop7 | 0.305304706 | 0.558125302 |
| ENSMUSG0000003267 | chr11 | Prorsd1 | 0.324535018 | 0.558125302 |
| ENSMUSG0000009025 | chr12 | Churc1 | 0.358892106 | 0.558125302 |
| ENSMUSG0000001722 | chr11 | Psmd3 | 0.156705649 | 0.558605449 |
| ENSMUSG0000003159 | chr8 | Pdgfrl | 0.441507957 | 0.558724477 |
| ENSMUSG0000003032 | chr6 | Ift122 | -0.14612634 | 0.558996023 |
| ENSMUSG0000003735 | chr1 | Actr1b | 0.162726587 | 0.559076045 |
| ENSMUSG0000002458 | chr18 | Fech | 0.130785713 | 0.559098164 |
| ENSMUSG0000000704 | chr17 | Clic1 | 0.190056742 | 0.559099531 |
| ENSMUSG0000004144 | chr14 | Mmrn2 | 0.279768944 | 0.559099531 |
| ENSMUSG0000002508 | chr19 | Gfra1 | -0.228892749 | 0.559420925 |
| ENSMUSG0000002788 | chr3 | Ampd2 | -0.162222331 | 0.559420925 |
| ENSMUSG0000005869 | chr14 | Ccser2 | 0.143178092 | 0.559420925 |
| ENSMUSG0000001051 | chr4 | Faf1 | 0.202132367 | 0.559420925 |
| ENSMUSG0000009532 | chr17 | Zfp870 | 0.212426663 | 0.559420925 |
| ENSMUSG0000002683 | chr2 | Upp2 | 0.299002048 | 0.559420925 |
| ENSMUSG0000003183 | chr8 | Mast3 | -0.19045009 | 0.560338961 |
| ENSMUSG0000000741 | chr12 | Mark3 | 0.11090335 | 0.560448148 |
| ENSMUSG0000002251 | chr16 | Trp63 | -0.64572923 | 0.56062622 |
| ENSMUSG0000007329 | chrX | Nudt10 | -0.275100135 | 0.560703955 |
| ENSMUSG0000002161 | chr13 | Atg10 | 0.229213733 | 0.560871167 |
| ENSMUSG0000003579 | chr10 | Zdhhc17 | -0.134931189 | 0.560960279 |
| ENSMUSG0000003868 | chr13 | Pak1ip1 | 0.19028539 | 0.560960279 |
| ENSMUSG0000004764 | chr10 | Fbxo30 | 0.231038958 | 0.560960279 |
| ENSMUSG0000002060 | chr12 | Apob | -1.948508555 | 0.561094063 |
| ENSMUSG0000000516 | chr8 | Prdx2 | 0.255009685 | 0.561765311 |

| | | | | |
|---|---|---|---|---|
| ENSMUSG0000002226 | chr15 | Ank | 0.209873661 | 0.562015959 |
| ENSMUSG0000007086 | chr4 | Skint3 | -1.060343748 | 0.562331746 |
| ENSMUSG0000007881 | chr11 | Eif5a | 0.259106469 | 0.562331746 |
| ENSMUSG0000005977 | chr7 | Slx1b | -0.314443243 | 0.562651109 |
| ENSMUSG0000002532 | chr12 | Sp4 | -0.240347995 | 0.562651109 |
| ENSMUSG0000002089 | chr11 | Aurkb | -0.854164489 | 0.562841216 |
| ENSMUSG0000003094 | chr7 | Thumpd1 | 0.127613727 | 0.562890836 |
| ENSMUSG0000002440 | chr18 | Riok3 | 0.127588064 | 0.563123554 |
| ENSMUSG0000002081 | chr11 | Mfsd11 | 0.158745126 | 0.563123554 |
| ENSMUSG0000002008 | chr10 | Aifm2 | 0.220679795 | 0.563123554 |
| ENSMUSG0000005319 | chr7 | Prx | 0.263526366 | 0.563123554 |
| ENSMUSG0000000865 | chr16 | Rbfox1 | 0.280087957 | 0.563123554 |
| ENSMUSG0000012160 | chr7 | Ahsp | 1.455475257 | 0.563123554 |
| ENSMUSG0000002826 | chr3 | Gbp3 | -0.388440002 | 0.563797795 |
| ENSMUSG0000007930 | chr6 | Tex52 | -0.545862045 | 0.563797795 |
| ENSMUSG0000007329 | chrX | Nudt11 | -0.219152822 | 0.563797795 |
| ENSMUSG0000002256 | chr15 | Grina | -0.149313037 | 0.563797795 |
| ENSMUSG0000009147 | chr1 | Catspere2 | -0.828199865 | 0.564494921 |
| ENSMUSG0000006406 | chr16 | Dzip3 | -0.167368911 | 0.564494921 |
| ENSMUSG0000002934 | chr5 | Tpst2 | 0.194770869 | 0.564494921 |
| ENSMUSG0000006279 | chr7 | Hikeshi | 0.298720315 | 0.56495623 |
| ENSMUSG0000002209 | chr14 | Pdlim2 | -0.367277717 | 0.565069705 |
| ENSMUSG0000005164 | chr8 | Kctd19 | 0.9360719 | 0.565069705 |
| ENSMUSG0000004470 | chr1 | Kcnj10 | -0.255033882 | 0.565365907 |
| ENSMUSG0000003837 | chr1 | Pcp4l1 | 0.297916076 | 0.565530598 |
| ENSMUSG0000005580 | chr11 | Fmnl1 | -0.276400159 | 0.566016569 |
| ENSMUSG0000002638 | chr1 | Tfcp2l1 | -0.352687607 | 0.566237773 |
| ENSMUSG0000000386 | chr7 | Ppfia3 | -0.261068818 | 0.566237773 |
| ENSMUSG0000009904 | chr2 | Gm28035 | 0.942308218 | 0.566237773 |
| ENSMUSG0000002724 | chr2 | Harbi1 | 0.255715562 | 0.566237773 |
| ENSMUSG0000003461 | chr10 | Chst11 | 0.212586553 | 0.566237773 |
| ENSMUSG0000002380 | chr17 | Rps6ka2 | 0.281195404 | 0.566237773 |
| ENSMUSG0000002814 | chr3 | Them4 | -0.189260381 | 0.566375835 |
| ENSMUSG0000003465 | chr11 | Cd300a | 0.658345995 | 0.566386225 |
| ENSMUSG0000000015 | chr16 | Igsf5 | -0.890754141 | 0.566426914 |
| ENSMUSG0000003596 | chr4 | Rusc2 | 0.248680389 | 0.566819206 |
| ENSMUSG0000002827 | chr3 | Gbp2 | 0.224177137 | 0.566819206 |
| ENSMUSG0000004048 | chr17 | Dxo | 0.188289482 | 0.567069164 |
| ENSMUSG0000004059 | chr2 | Cstpp1 | -0.112725403 | 0.567159491 |
| ENSMUSG0000000232 | chr14 | Gmpr2 | 0.225866749 | 0.567159491 |
| ENSMUSG0000005383 | chr11 | Nudcd3 | 0.143006958 | 0.567621962 |
| ENSMUSG0000002902 | chr4 | Miip | -0.166268088 | 0.567672115 |
| ENSMUSG0000000016 | chr9 | Dlat | 0.142832222 | 0.567672115 |
| ENSMUSG0000005332 | chr10 | Gatd3a | 0.16294292 | 0.567828143 |
| ENSMUSG0000005027 | chr16 | Dscam | -0.299458204 | 0.567987548 |
| ENSMUSG0000002890 | chr4 | Epb41 | -0.198924162 | 0.567987548 |
| ENSMUSG0000002229 | chr15 | Slc25a32 | -0.163341448 | 0.567987548 |
| ENSMUSG0000004942 | chr10 | Chchd10 | 0.241424022 | 0.567987548 |
| ENSMUSG0000002184 | chr14 | Peli2 | 0.31525792 | 0.568142981 |
| ENSMUSG0000003386 | chr19 | Klf9 | 0.310078306 | 0.568142981 |

| | | | | |
|---|---|---|---|---|
| ENSMUSG000001856 | chr11 | Slc2a4 | 0.138243589 | 0.568187776 |
| ENSMUSG000002939 | chr5 | Cdk2ap1 | 0.20051869 | 0.568572039 |
| ENSMUSG000002487 | chr19 | Doc2g | 0.59326442 | 0.568572039 |
| ENSMUSG000005586 | chr10 | Izumo4 | -0.263617281 | 0.568834475 |
| ENSMUSG000002405 | chr17 | Clip4 | 0.134443137 | 0.56904173 |
| ENSMUSG000002242 | chr15 | Cby1 | 0.252124005 | 0.56904173 |
| ENSMUSG000004173 | chr5 | Coq5 | 0.137524116 | 0.56944203 |
| ENSMUSG000004084 | chr1 | Sft2d2 | -0.144886088 | 0.569462167 |
| ENSMUSG000004251 | chr1 | AA986860 | -0.506742147 | 0.569645607 |
| ENSMUSG000002536 | chr10 | Pa2g4 | 0.153112298 | 0.569723491 |
| ENSMUSG000005140 | chr18 | Kctd16 | -0.979179565 | 0.570519417 |
| ENSMUSG000003226 | chr9 | Usp28 | 0.171244999 | 0.570606109 |
| ENSMUSG000006719 | chrX | Eif1ax | 0.196264825 | 0.570606109 |
| ENSMUSG000001717 | chr11 | Nt5c3b | 0.20493958 | 0.570613888 |
| ENSMUSG000006828 | chr16 | Usf3 | -0.265986714 | 0.570644161 |
| ENSMUSG000002419 | chr17 | Phf1 | -0.195701469 | 0.570644161 |
| ENSMUSG000009140 | chr12 | Rd3l | 0.808557478 | 0.570664421 |
| ENSMUSG000002899 | chr5 | Tomm7 | 0.320550325 | 0.570938248 |
| ENSMUSG000006030 | chr10 | 2610008E11Rik | 0.166113961 | 0.57097359 |
| ENSMUSG000004987 | chr10 | Calhm5 | 0.263798386 | 0.57097359 |
| ENSMUSG000004675 | chr11 | Mrps7 | 0.270759727 | 0.57097359 |
| ENSMUSG000003128 | chrX | Chrdl1 | 0.321344384 | 0.57097359 |
| ENSMUSG000005401 | chr12 | Tmem179 | -0.297039257 | 0.571182226 |
| ENSMUSG000002008 | chr10 | Sar1a | 0.200281206 | 0.571632197 |
| ENSMUSG000002408 | chr17 | Ndufaf7 | 0.240023278 | 0.571632197 |
| ENSMUSG000000283 | chr17 | Chaf1a | -0.514052205 | 0.57165781 |
| ENSMUSG000003167 | chr8 | Cdh11 | -0.177093481 | 0.571674748 |
| ENSMUSG000002161 | chr13 | Vcan | 0.246513864 | 0.571769571 |
| ENSMUSG000003918 | chr17 | Nubp2 | 0.271612722 | 0.572144528 |
| ENSMUSG000002596 | chr1 | Eef1b2 | 0.300612237 | 0.572172411 |
| ENSMUSG000003135 | chrX | Syap1 | 0.185800878 | 0.572232781 |
| ENSMUSG000000831 | chr7 | Relt | 0.562976322 | 0.572232781 |
| ENSMUSG000003022 | chr6 | Rerg | 0.32466982 | 0.572571744 |
| ENSMUSG000007310 | chr5 | Drc1 | -0.336954109 | 0.572919856 |
| ENSMUSG000010986 | chr2 | Hspa14 | 0.18624532 | 0.572937663 |
| ENSMUSG000003028 | chr6 | St8sia1 | 0.328834864 | 0.572937663 |
| ENSMUSG000005256 | chr13 | H1f3 | -0.485238783 | 0.573028126 |
| ENSMUSG000009017 | chr1 | Ugt1a2 | -1.291535027 | 0.573028126 |
| ENSMUSG000004187 | chr5 | Ankrd13a | 0.143381723 | 0.573028126 |
| ENSMUSG000002291 | chr16 | Psmg1 | 0.283265687 | 0.573028126 |
| ENSMUSG000005789 | chr9 | Zfp105 | 0.190577622 | 0.573082951 |
| ENSMUSG000000205 | chr11 | Rab34 | 0.198898265 | 0.573082951 |
| ENSMUSG000002860 | chr4 | Scp2 | 0.209648906 | 0.573332794 |
| ENSMUSG000002685 | chr2 | Miga2 | 0.184644799 | 0.573332794 |
| ENSMUSG000002785 | chr3 | Tspan2 | 0.175167602 | 0.573480666 |
| ENSMUSG000000820 | chr8 | Cers4 | -0.204980958 | 0.573626636 |
| ENSMUSG000002944 | chr5 | Cct6a | 0.232909122 | 0.573685712 |
| ENSMUSG000003515 | chrX | Eif2s3x | 0.125862758 | 0.573906055 |
| ENSMUSG000004474 | chr11 | Abca6 | -0.533814014 | 0.573926349 |
| ENSMUSG000002237 | chr15 | Adcy8 | -1.512848243 | 0.574191704 |

| | | | | |
|---|---|---|---|---|
| ENSMUSG0000004550 | chr13 | Gpr150 | -1.176626131 | 0.574237895 |
| ENSMUSG0000002889 | chr4 | Sesn2 | -0.268702103 | 0.574237895 |
| ENSMUSG0000002898 | chr4 | Ctnnbip1 | 0.206225702 | 0.574237895 |
| ENSMUSG0000003201 | chr9 | Oaf | 0.218491203 | 0.574237895 |
| ENSMUSG0000002924 | chr5 | Ppat | 0.180468967 | 0.574358133 |
| ENSMUSG0000000588 | chr2 | Uqcc1 | 0.162876098 | 0.574431436 |
| ENSMUSG0000002868 | chr4 | Plk3 | -0.3546712 | 0.574547469 |
| ENSMUSG0000003380 | chr2 | Tmem87a | -0.142122182 | 0.574547469 |
| ENSMUSG0000002278 | chr16 | Wdr53 | 0.264513506 | 0.574547469 |
| ENSMUSG0000003143 | chrX | Rbm41 | 0.247270052 | 0.574547469 |
| ENSMUSG0000002848 | chr4 | Glipr2 | 0.350322985 | 0.574547469 |
| ENSMUSG0000002070 | chr11 | Adap2 | 0.58630774 | 0.574547469 |
| ENSMUSG0000003153 | chr8 | Smim19 | 0.17603632 | 0.575003862 |
| ENSMUSG0000004848 | chr7 | Mypop | -0.203119501 | 0.575300087 |
| ENSMUSG0000002191 | chr14 | Nek4 | -0.188791815 | 0.575300087 |
| ENSMUSG0000001955 | chrX | Slc6a8 | 0.138317134 | 0.575300087 |
| ENSMUSG0000007348 | chr1 | Mtarc2 | 0.175500876 | 0.575300087 |
| ENSMUSG0000003847 | chr1 | Nos1ap | 0.297389368 | 0.575300087 |
| ENSMUSG0000002851 | chr4 | Plpp3 | 0.159618907 | 0.575454983 |
| ENSMUSG0000002713 | chr2 | Lpcat4 | -0.231619423 | 0.575616569 |
| ENSMUSG0000002072 | chr11 | Cep112 | -0.379033454 | 0.575636488 |
| ENSMUSG0000001801 | chr11 | Rac3 | -0.268063873 | 0.575977466 |
| ENSMUSG0000004339 | chr16 | 2510009E07Rik | 0.13895572 | 0.575994858 |
| ENSMUSG0000004562 | chr2 | Esf1 | 0.188783149 | 0.575994858 |
| ENSMUSG0000007491 | chr2 | Inafm2 | 0.193573721 | 0.575994858 |
| ENSMUSG0000009024 | chr10 | Bloc1s1 | 0.283582213 | 0.575994858 |
| ENSMUSG0000005254 | chr3 | St6galnac3 | 0.217295526 | 0.576298728 |
| ENSMUSG0000002264 | chr16 | Bbx | -0.245487172 | 0.576792838 |
| ENSMUSG0000003008 | chr6 | Sec61a1 | 0.16236263 | 0.576792838 |
| ENSMUSG0000001969 | chr1 | Akt3 | 0.174438976 | 0.576792838 |
| ENSMUSG0000002785 | chr3 | Ngf | 0.33865235 | 0.576793638 |
| ENSMUSG0000003251 | chr9 | Csrnp1 | -0.304926502 | 0.577238501 |
| ENSMUSG0000003667 | chr10 | Tmtc3 | 0.1840025 | 0.577238501 |
| ENSMUSG0000003456 | chr11 | Atp5pd | 0.327090881 | 0.577238501 |
| ENSMUSG0000002122 | chr12 | Acot2 | 0.406495002 | 0.577238501 |
| ENSMUSG0000003009 | chr6 | Tmem43 | 0.201114841 | 0.577263965 |
| ENSMUSG0000001528 | chrX | Lage3 | 0.240713281 | 0.577287283 |
| ENSMUSG0000004162 | chr11 | Vcf1 | 0.245967893 | 0.577331045 |
| ENSMUSG0000005114 | chr16 | Camk2n2 | -0.248237772 | 0.577452602 |
| ENSMUSG0000003410 | chr2 | Ctnnd1 | -0.139138564 | 0.577452602 |
| ENSMUSG0000000252 | chr15 | Puf60 | 0.194133401 | 0.577452602 |
| ENSMUSG0000002380 | chr17 | Rsph3b | 0.22686278 | 0.577452602 |
| ENSMUSG0000003720 | chr15 | Atg101 | 0.255466826 | 0.577452602 |
| ENSMUSG0000002526 | chrX | Hsd17b10 | 0.263843176 | 0.577452602 |
| ENSMUSG0000002256 | chr15 | Plec | 0.199870568 | 0.577510168 |
| ENSMUSG0000004059 | chr11 | Mis12 | 0.171332214 | 0.577567051 |
| ENSMUSG0000002080 | chr11 | Txndc17 | 0.242363039 | 0.577567051 |
| ENSMUSG0000002669 | chr1 | Mettl13 | 0.186543316 | 0.577792291 |
| ENSMUSG0000001991 | chr10 | P4ha1 | 0.34063648 | 0.577792291 |
| ENSMUSG0000002655 | chr1 | Uck2 | 0.224633918 | 0.577863668 |

| | | | | |
|---|---|---|---|---|
| ENSMUSG0000004148 | chr18 | Piezo2 | -0.199441474 | 0.578131663 |
| ENSMUSG0000004001 | chr3 | Ptger3 | -0.167022757 | 0.578131663 |
| ENSMUSG0000003091 | chr7 | Ldaf1 | 0.202211522 | 0.578131663 |
| ENSMUSG0000002944 | chr5 | Rhof | -0.4043147 | 0.578172217 |
| ENSMUSG0000002950 | chr5 | Pgam5 | 0.182767982 | 0.578318326 |
| ENSMUSG0000004653 | chrX | Ar | -0.380609909 | 0.57842228 |
| ENSMUSG0000002489 | chr19 | Apba1 | -0.211908196 | 0.57842228 |
| ENSMUSG0000003430 | chr8 | Sdr42e1 | -0.2405371 | 0.57842228 |
| ENSMUSG0000000005 | chr6 | Cav2 | 0.162348269 | 0.57842228 |
| ENSMUSG0000003284 | chr13 | Zswim6 | 0.218312816 | 0.57842228 |
| ENSMUSG0000002106 | chr12 | Nin | 0.235930344 | 0.57842228 |
| ENSMUSG0000001818 | chr1 | Uchl5 | 0.256584784 | 0.57842228 |
| ENSMUSG0000002815 | chr3 | Dapp1 | 0.313335236 | 0.57842228 |
| ENSMUSG0000004490 | chr18 | 4930503L19Rik | -0.330080606 | 0.578463108 |
| ENSMUSG0000002630 | chr1 | Scly | 0.193362064 | 0.578463108 |
| ENSMUSG0000003161 | chr8 | Hpgd | -0.305820234 | 0.578534469 |
| ENSMUSG0000001853 | chr11 | Pcgf2 | -0.26239655 | 0.578534469 |
| ENSMUSG0000002537 | chr11 | Aatk | -0.243953021 | 0.578534469 |
| ENSMUSG0000002834 | chr4 | Erp44 | 0.164930804 | 0.578534469 |
| ENSMUSG0000001999 | chr10 | Pde7b | 0.224269263 | 0.579070223 |
| ENSMUSG0000003464 | chr6 | Lrrn1 | -0.242672221 | 0.579561693 |
| ENSMUSG0000003904 | chr2 | Adrm1 | 0.138601718 | 0.579561693 |
| ENSMUSG0000005084 | chr1 | Cdh20 | 0.436778146 | 0.579561693 |
| ENSMUSG0000002242 | chr15 | Nptxr | -0.213992222 | 0.579883929 |
| ENSMUSG0000003578 | chr10 | R3hdm4 | -0.180002039 | 0.579883929 |
| ENSMUSG0000004422 | chr19 | Nkx2-3 | 0.220765181 | 0.579883929 |
| ENSMUSG0000000794 | chr7 | Ttc9b | -0.331695603 | 0.579950635 |
| ENSMUSG0000003219 | chr9 | Polr2m | 0.190836054 | 0.579950635 |
| ENSMUSG0000006299 | chr8 | Nrg1 | -0.255883042 | 0.580139928 |
| ENSMUSG0000003411 | chr17 | Vav1 | -0.454196294 | 0.580139928 |
| ENSMUSG0000005082 | chr5 | Slc29a4 | -0.23572416 | 0.580139928 |
| ENSMUSG0000003710 | chr5 | Zcwpw1 | -0.413395232 | 0.580139928 |
| ENSMUSG0000002252 | chr15 | Zfp251 | 0.134103409 | 0.580139928 |
| ENSMUSG0000001109 | chr7 | Akt1s1 | 0.183898286 | 0.580139928 |
| ENSMUSG0000002155 | chr13 | Dapk1 | 0.197309497 | 0.580139928 |
| ENSMUSG0000003394 | chr6 | Brk1 | 0.247647801 | 0.580139928 |
| ENSMUSG0000002018 | chr10 | Csrp2 | 0.264378652 | 0.580139928 |
| ENSMUSG0000007964 | chrX | Rpl39 | 0.346997545 | 0.580139928 |
| ENSMUSG0000007129 | chr13 | Zfp58 | -0.309948895 | 0.580505091 |
| ENSMUSG0000002703 | chr2 | Stk39 | -0.162610972 | 0.580505091 |
| ENSMUSG0000003611 | chr10 | Metap2 | 0.119976363 | 0.580505091 |
| ENSMUSG0000001211 | chr4 | Dhdds | 0.13493736 | 0.580505091 |
| ENSMUSG0000005849 | chr4 | Rnf207 | 0.209934996 | 0.580505091 |
| ENSMUSG0000002052 | chr11 | Gria1 | 0.236098582 | 0.580505091 |
| ENSMUSG0000001983 | chr10 | Slc16a10 | 0.268993836 | 0.580505091 |
| ENSMUSG0000004714 | chr10 | Tet1 | -0.333938907 | 0.580622213 |
| ENSMUSG0000005321 | chr10 | Raet1e | -1.658902159 | 0.580622213 |
| ENSMUSG0000003384 | chr1 | B3galt2 | -0.344462833 | 0.580622213 |
| ENSMUSG0000005621 | chr7 | Cebpg | -0.14422618 | 0.580622213 |
| ENSMUSG0000002787 | chr3 | Sec22b | 0.178232023 | 0.580622213 |

| | | | | |
|---|---|---|---|---|
| ENSMUSG0000001599 | chr8 | Fnta | 0.244164962 | 0.580622213 |
| ENSMUSG0000002667 | chr2 | Ccdc3 | 0.254933376 | 0.580622213 |
| ENSMUSG0000001905 | chr5 | Fis1 | 0.258055407 | 0.580622213 |
| ENSMUSG0000002930 | chr5 | Dmp1 | 1.235843024 | 0.580622213 |
| ENSMUSG0000002992 | chr6 | Rab19 | -0.677177776 | 0.580662472 |
| ENSMUSG0000005848 | chr5 | Kl | -0.383121926 | 0.580662472 |
| ENSMUSG0000006937 | chr18 | Prdm6 | 0.31967655 | 0.580736738 |
| ENSMUSG0000000158 | chr11 | Acap1 | -0.753842271 | 0.580818441 |
| ENSMUSG0000002808 | chr3 | Fbxw7 | -0.134080461 | 0.580818441 |
| ENSMUSG0000001416 | chr13 | Klhl3 | -0.336106315 | 0.581011731 |
| ENSMUSG0000007905 | chr2 | Kcnip3 | -0.212374716 | 0.581011731 |
| ENSMUSG0000002982 | chr6 | Gsdme | -0.397656519 | 0.581047364 |
| ENSMUSG0000005024 | chr16 | Hic2 | -0.351122706 | 0.581047364 |
| ENSMUSG0000002766 | chr3 | Gnb4 | -0.153502109 | 0.581047364 |
| ENSMUSG0000003532 | chr12 | Fbxo33 | 0.267635028 | 0.581047364 |
| ENSMUSG0000002744 | chr2 | Cst3 | 0.23656614 | 0.581212805 |
| ENSMUSG0000000690 | chr6 | Stambp | 0.192207032 | 0.58164138 |
| ENSMUSG0000011851 | chr15 | Cyp2d13 | -1.807302116 | 0.581843365 |
| ENSMUSG0000006980 | chr11 | Tlcd3a | -0.199676732 | 0.581843365 |
| ENSMUSG0000005001 | chr5 | Pitpnb | 0.162958529 | 0.581843365 |
| ENSMUSG0000006403 | chr5 | Gpn1 | 0.186660305 | 0.581843365 |
| ENSMUSG0000005623 | chr14 | Ncoa4 | 0.234470993 | 0.581843365 |
| ENSMUSG0000000342 | chr7 | Fcgrt | 0.202608104 | 0.582325519 |
| ENSMUSG0000001565 | chr9 | Hspa8 | 0.325733792 | 0.582325519 |
| ENSMUSG0000004101 | chr9 | Cmtm8 | -0.736503173 | 0.582812785 |
| ENSMUSG0000004137 | chr16 | Cldn5 | -0.233205518 | 0.582812785 |
| ENSMUSG0000002718 | chr2 | Nat10 | 0.116937957 | 0.582812785 |
| ENSMUSG0000002538 | chr10 | Cnpy2 | 0.243917549 | 0.582812785 |
| ENSMUSG0000002814 | chr3 | Mrpl9 | 0.207756623 | 0.582958874 |
| ENSMUSG0000001165 | chr7 | Fuz | -0.143587277 | 0.583188777 |
| ENSMUSG0000003725 | chr18 | Mex3c | 0.125711668 | 0.583188777 |
| ENSMUSG0000005024 | chr5 | Evc2 | 0.142378916 | 0.583188777 |
| ENSMUSG0000003171 | chr8 | Zfp330 | 0.152601348 | 0.583188777 |
| ENSMUSG0000005935 | chr8 | Wdr83os | 0.174006669 | 0.583188777 |
| ENSMUSG0000004481 | chr4 | Shb | 0.178463489 | 0.583188777 |
| ENSMUSG0000009073 | chr3 | Rps27 | 0.342712511 | 0.583188777 |
| ENSMUSG0000000536 | chr6 | Crbn | 0.163265002 | 0.583348648 |
| ENSMUSG0000004167 | chr14 | Lrrc18 | -0.383390073 | 0.583834384 |
| ENSMUSG0000003913 | chr4 | Whrn | 0.207245797 | 0.584279671 |
| ENSMUSG0000002125 | chr12 | Cyp46a1 | -0.425652737 | 0.584368227 |
| ENSMUSG0000003370 | chr15 | Lrrc24 | -0.499690018 | 0.584368227 |
| ENSMUSG0000004207 | chr5 | Svop | -0.262789555 | 0.584368227 |
| ENSMUSG0000004858 | chr3 | E130311K13Rik | -0.307459818 | 0.584368227 |
| ENSMUSG0000002243 | chr15 | Parvb | -0.235142551 | 0.584368227 |
| ENSMUSG0000002732 | chr2 | Spef1 | -0.264201021 | 0.584368227 |
| ENSMUSG0000003125 | chrX | Xkrx | -0.709668723 | 0.584368227 |
| ENSMUSG0000004666 | chr18 | Cxxc5 | -0.224351096 | 0.584368227 |
| ENSMUSG0000005097 | chr7 | Gdpgp1 | -0.257705148 | 0.584368227 |
| ENSMUSG0000004861 | chr11 | Nog | -0.249356791 | 0.584368227 |
| ENSMUSG0000002495 | chr19 | Esrra | -0.162347061 | 0.584368227 |

| | | | | |
|---|---|---|---|---|
| ENSMUSG0000003072 | chr7 | Pold3 | -0.158054244 | 0.584368227 |
| ENSMUSG0000003211 | chr9 | Fez1 | -0.146319363 | 0.584368227 |
| ENSMUSG0000002247 | chr15 | Aco2 | 0.120249149 | 0.584368227 |
| ENSMUSG0000003211 | chr9 | Stt3a | 0.132523453 | 0.584368227 |
| ENSMUSG0000004072 | chr1 | Rcsd1 | 0.154848276 | 0.584368227 |
| ENSMUSG0000000017 | chr9 | Sdhd | 0.166309814 | 0.584368227 |
| ENSMUSG0000001508 | chr2 | Rabl6 | 0.174766911 | 0.584368227 |
| ENSMUSG0000001979 | chr10 | Katna1 | 0.188951866 | 0.584368227 |
| ENSMUSG0000012160 | chrX | Akap17a | 0.165452145 | 0.584368227 |
| ENSMUSG0000003860 | chr8 | Ripor1 | 0.172478619 | 0.584368227 |
| ENSMUSG0000000323 | chr16 | Eif2b5 | 0.17619055 | 0.584368227 |
| ENSMUSG0000001946 | chr11 | Plscr3 | 0.159975862 | 0.584368227 |
| ENSMUSG0000002209 | chr14 | Slc39a14 | 0.189631195 | 0.584368227 |
| ENSMUSG0000003931 | chr17 | Rftn1 | 0.224875651 | 0.584368227 |
| ENSMUSG0000002086 | chr11 | Lrrc59 | 0.222995769 | 0.584368227 |
| ENSMUSG0000001842 | chr11 | Ypel2 | 0.22758198 | 0.584368227 |
| ENSMUSG0000001382 | chr9 | Elof1 | 0.225889244 | 0.584368227 |
| ENSMUSG0000002549 | chr7 | Hras | 0.229699636 | 0.584368227 |
| ENSMUSG0000002794 | chr3 | Il6ra | 0.227519116 | 0.584368227 |
| ENSMUSG0000004204 | chr13 | Tbca | 0.233557978 | 0.584368227 |
| ENSMUSG0000007028 | chr9 | Ndufaf3 | 0.241209592 | 0.584368227 |
| ENSMUSG0000002713 | chr2 | Emc4 | 0.229759075 | 0.584368227 |
| ENSMUSG0000004093 | chr9 | Ulk4 | 0.294615298 | 0.584368227 |
| ENSMUSG0000010389 | chr18 | Pcdhga8 | 0.303119258 | 0.584368227 |
| ENSMUSG0000002795 | chr3 | Pmvk | 0.236990568 | 0.584368227 |
| ENSMUSG0000007958 | chr1 | Tmem182 | 0.859810454 | 0.584368227 |
| ENSMUSG0000004913 | chr19 | Nrap | 0.518894869 | 0.584368227 |
| ENSMUSG0000002019 | chr11 | Zpbp | 0.644841465 | 0.584368227 |
| ENSMUSG0000003654 | chr11 | Adamts2 | 0.262323722 | 0.584368227 |
| ENSMUSG0000003182 | chr8 | Gins2 | 0.610916958 | 0.584368227 |
| ENSMUSG0000003701 | chr3 | Frem2 | 0.284191462 | 0.584368227 |
| ENSMUSG0000002413 | chr17 | Six2 | 0.990738729 | 0.584368227 |
| ENSMUSG0000003416 | chr10 | Zfc3h1 | -0.169885446 | 0.584547502 |
| ENSMUSG0000002418 | chr17 | Axin1 | 0.140846423 | 0.584547502 |
| ENSMUSG0000007046 | chr7 | Tlnrd1 | -0.173762461 | 0.584600029 |
| ENSMUSG0000003306 | chr15 | Pfkm | 0.140891703 | 0.584936956 |
| ENSMUSG0000004595 | chr1 | Cavin2 | 0.170848023 | 0.584936956 |
| ENSMUSG0000002112 | chr12 | Vti1b | 0.277890701 | 0.584936956 |
| ENSMUSG0000011792 | chr19 | Tmem223 | 0.249004347 | 0.585387024 |
| ENSMUSG0000007370 | chr4 | Cenps | -0.707389881 | 0.585429018 |
| ENSMUSG0000003854 | chr13 | Ranbp9 | 0.157129596 | 0.585429018 |
| ENSMUSG0000002657 | chr1 | Dpt | 0.202736104 | 0.585429018 |
| ENSMUSG0000002136 | chr13 | Gcnt2 | 0.187484364 | 0.585429018 |
| ENSMUSG0000002490 | chr19 | Mrpl11 | 0.219726161 | 0.585429018 |
| ENSMUSG0000002889 | chr4 | Trnau1ap | 0.235189204 | 0.585429018 |
| ENSMUSG0000003130 | chrX | Nlgn3 | -0.279825972 | 0.585537382 |
| ENSMUSG0000003701 | chr2 | Sstr4 | -0.271394753 | 0.585537382 |
| ENSMUSG0000003021 | chr6 | Wbp11 | 0.107740103 | 0.585537382 |
| ENSMUSG0000005723 | chr2 | Mettl15 | 0.254050182 | 0.585537382 |
| ENSMUSG0000004765 | chr19 | Trpt1 | 0.270383322 | 0.586134446 |

| | | | | |
|---|---|---|---|---|
| ENSMUSG0000004034 | chr10 | Arhgap9 | -0.473991061 | 0.586705936 |
| ENSMUSG0000003943 | chr8 | Mtmr7 | -0.196277466 | 0.586705936 |
| ENSMUSG0000002602 | chr1 | Nop58 | 0.137759095 | 0.586705936 |
| ENSMUSG0000002624 | chr1 | Mrpl44 | 0.233254343 | 0.586705936 |
| ENSMUSG0000002169 | chr13 | Pde4d | 0.25344828 | 0.586782992 |
| ENSMUSG0000003109 | chrX | Smarca1 | -0.168683229 | 0.586926513 |
| ENSMUSG0000002233 | chr15 | Eif3e | 0.219323193 | 0.586932096 |
| ENSMUSG0000002617 | chr1 | Cnot9 | 0.145191631 | 0.587190199 |
| ENSMUSG0000004935 | chr18 | Brd8dc | 0.513522054 | 0.587190199 |
| ENSMUSG0000002513 | chr11 | Anapc11 | 0.266504012 | 0.587190199 |
| ENSMUSG0000001183 | chr8 | Snapc2 | 0.195380599 | 0.587383294 |
| ENSMUSG0000000343 | chr7 | Timm50 | 0.272303347 | 0.587391113 |
| ENSMUSG0000006689 | chr9 | Fbxl12 | -0.200233978 | 0.587406158 |
| ENSMUSG0000005951 | chr5 | Znhit1 | 0.279783733 | 0.587532714 |
| ENSMUSG0000005993 | chr4 | 9430015G10Rik | -0.30195648 | 0.58790454 |
| ENSMUSG0000004283 | chr18 | Nrep | -0.258769377 | 0.58790454 |
| ENSMUSG0000002992 | chr6 | Tbxas1 | -0.354349343 | 0.58790454 |
| ENSMUSG0000003717 | chr6 | Dennd11 | -0.221526986 | 0.58790454 |
| ENSMUSG0000001772 | chr11 | Etv4 | -0.419660395 | 0.58790454 |
| ENSMUSG0000007029 | chr9 | Ddx43 | -0.861072051 | 0.58790454 |
| ENSMUSG0000003075 | chr7 | Far1 | -0.159594318 | 0.58790454 |
| ENSMUSG0000003821 | chr13 | Mfsd14b | -0.144084336 | 0.58790454 |
| ENSMUSG0000003985 | chr16 | Epcip | 0.433274766 | 0.58790454 |
| ENSMUSG0000005883 | chr8 | Rex1bd | 0.222784535 | 0.58790454 |
| ENSMUSG0000000694 | chr9 | Eif1b | 0.250193902 | 0.58790454 |
| ENSMUSG0000001125 | chr11 | Adam19 | 0.254610425 | 0.58790454 |
| ENSMUSG0000004615 | chr13 | Chrm3 | 0.265493879 | 0.58790454 |
| ENSMUSG0000004368 | chr8 | 1190005I06Rik | 0.717274951 | 0.58790454 |
| ENSMUSG0000010244 | chr18 | Pcdhga9 | 0.35703976 | 0.58790454 |
| ENSMUSG0000004028 | chr10 | Ndufa4l2 | 0.504472131 | 0.58790454 |
| ENSMUSG0000002786 | chr3 | Casq2 | 0.446399455 | 0.58806241 |
| ENSMUSG0000003715 | chr4 | Il22ra1 | -0.614242511 | 0.588153172 |
| ENSMUSG0000002633 | chr1 | Pam | 0.115525234 | 0.588153172 |
| ENSMUSG0000003084 | chr7 | Lamtor1 | 0.132367399 | 0.588153172 |
| ENSMUSG0000002472 | chr19 | Carnmt1 | 0.180878978 | 0.588153172 |
| ENSMUSG0000002853 | chr4 | St3gal3 | 0.209098713 | 0.588153172 |
| ENSMUSG0000002638 | chr1 | 3110009E18Rik | 0.420880332 | 0.588153172 |
| ENSMUSG0000004554 | chr11 | Krt14 | 0.98515613 | 0.588153172 |
| ENSMUSG0000006432 | chr12 | Siva1 | 0.297447866 | 0.588154125 |
| ENSMUSG0000002886 | chr5 | Hgf | -0.279257772 | 0.588195688 |
| ENSMUSG0000002145 | chr13 | Aopep | 0.16693893 | 0.588278824 |
| ENSMUSG0000004911 | chr13 | Agtr1a | 0.184581876 | 0.588278824 |
| ENSMUSG0000003878 | chr6 | Cnot4 | 0.186366893 | 0.588278824 |
| ENSMUSG0000003965 | chr17 | Rxrb | 0.175922534 | 0.5886453 |
| ENSMUSG0000002611 | chr1 | Cnga3 | -0.305824012 | 0.588967643 |
| ENSMUSG0000005968 | chr6 | Zfp637 | 0.244390256 | 0.588967643 |
| ENSMUSG0000002822 | chr4 | Mmp16 | -0.361605891 | 0.589071833 |
| ENSMUSG0000002857 | chr4 | Eqtn | -1.0593834 | 0.589071833 |
| ENSMUSG0000004035 | chr5 | Ankib1 | -0.17558702 | 0.589071833 |
| ENSMUSG0000012159 | chr5 | AB010352 | 0.337102099 | 0.589072915 |

| | | | | |
|---|---|---|---|---|
| ENSMUSG0000003289 | chr4 | Rims3 | -0.280104227 | 0.589084789 |
| ENSMUSG0000003000 | chr6 | Nat8 | -0.5667196 | 0.589084789 |
| ENSMUSG0000002416 | chr17 | Mapk8ip3 | -0.180441756 | 0.589084789 |
| ENSMUSG0000004721 | chr3 | Ythdf3 | 0.117637468 | 0.589116066 |
| ENSMUSG0000003944 | chr2 | Prpf18 | 0.146351004 | 0.589292566 |
| ENSMUSG0000002350 | chr6 | Cdca3 | -0.764504799 | 0.589532448 |
| ENSMUSG0000000914 | chr6 | Dqx1 | -0.264270738 | 0.589532448 |
| ENSMUSG0000003397 | chr17 | Zfp944 | -0.154247273 | 0.589532448 |
| ENSMUSG0000003162 | chr8 | Sin3b | 0.159926025 | 0.589532448 |
| ENSMUSG0000005445 | chr10 | Tle5 | 0.193390341 | 0.589532448 |
| ENSMUSG0000002397 | chr17 | Ptk7 | 0.162625903 | 0.589532448 |
| ENSMUSG0000002474 | chr19 | Fen1 | 0.81170545 | 0.589532448 |
| ENSMUSG0000002514 | chr7 | Mob2 | 0.293696765 | 0.589532448 |
| ENSMUSG0000004071 | chr11 | Sh3pxd2b | 0.306979034 | 0.589532448 |
| ENSMUSG0000002737 | chr2 | Bub1 | -1.185539463 | 0.589638269 |
| ENSMUSG0000004952 | chr15 | Cdc42ep1 | 0.156807661 | 0.589638269 |
| ENSMUSG0000002991 | chr6 | Ssbp1 | 0.184260985 | 0.589638269 |
| ENSMUSG0000003908 | chr14 | Zfp503 | 0.196970992 | 0.589638269 |
| ENSMUSG0000007948 | chrX | Pin4 | 0.280842628 | 0.589638269 |
| ENSMUSG0000003983 | chr3 | Arhgap29 | 0.300605923 | 0.589638269 |
| ENSMUSG0000006618 | chr7 | Cacng3 | 0.796148902 | 0.589638269 |
| ENSMUSG0000007892 | chr11 | Tgtp1 | -0.6110534 | 0.590155474 |
| ENSMUSG0000003114 | chrX | Magix | -0.741600612 | 0.590167731 |
| ENSMUSG0000002917 | chr5 | Sepsecs | -0.170015594 | 0.590167731 |
| ENSMUSG0000006329 | chr15 | Tmem117 | 0.14466412 | 0.590493687 |
| ENSMUSG0000006742 | chr17 | Zfp563 | 0.172360105 | 0.590493687 |
| ENSMUSG0000000426 | chr6 | Eno2 | -0.2817347 | 0.590887509 |
| ENSMUSG0000000551 | chr5 | Por | 0.158924875 | 0.590887509 |
| ENSMUSG0000007815 | chr11 | Cdk2ap1rt | -0.59088217 | 0.590988132 |
| ENSMUSG0000002416 | chr17 | Ift140 | -0.141751622 | 0.590988132 |
| ENSMUSG0000002474 | chr19 | Tmc1 | -1.713978116 | 0.591267405 |
| ENSMUSG0000007421 | chr7 | Sdhaf1 | 0.159427867 | 0.591267405 |
| ENSMUSG0000000478 | chr12 | Eif2b2 | 0.190906155 | 0.591559725 |
| ENSMUSG0000002383 | chr17 | Igf2r | 0.129320286 | 0.591587691 |
| ENSMUSG0000002597 | chr1 | Mob4 | 0.177851828 | 0.591634615 |
| ENSMUSG0000002453 | chr18 | Ptpn2 | 0.159094617 | 0.591953022 |
| ENSMUSG0000002172 | chr13 | Rnf180 | -0.333921466 | 0.592189621 |
| ENSMUSG0000009094 | chr12 | Ccdc71l | -0.242671062 | 0.592519947 |
| ENSMUSG0000002681 | chr2 | Slc25a25 | -0.199302286 | 0.592519947 |
| ENSMUSG0000004892 | chr7 | Zfp689 | -0.22504726 | 0.592519947 |
| ENSMUSG0000005544 | chr16 | Cd47 | -0.110587678 | 0.592519947 |
| ENSMUSG0000008968 | chr14 | Bcl2l2 | 0.102375137 | 0.592519947 |
| ENSMUSG0000003517 | chr19 | 1110059E24Rik | 0.197539295 | 0.592519947 |
| ENSMUSG0000002005 | chr10 | Ascl1 | -0.690226449 | 0.592670882 |
| ENSMUSG0000004132 | chr12 | Ak7 | -0.777221211 | 0.592670882 |
| ENSMUSG0000006981 | chr11 | Ccdc92b | -0.280704308 | 0.592670882 |
| ENSMUSG0000001470 | chr6 | Hoxa2 | -0.280252841 | 0.592670882 |
| ENSMUSG0000004085 | chr7 | Erf | -0.200898933 | 0.592670882 |
| ENSMUSG0000001930 | chr11 | Atp6v0a1 | -0.19101497 | 0.592670882 |
| ENSMUSG0000002761 | chr2 | Nfs1 | 0.133545994 | 0.592670882 |

| | | | | |
|---|---|---|---|---|
| ENSMUSG0000002894 | chr5 | Prkag2 | 0.185076174 | 0.592670882 |
| ENSMUSG0000002409 | chr17 | Vapa | 0.19097159 | 0.592670882 |
| ENSMUSG0000003193 | chr9 | Ankrd49 | 0.176054993 | 0.592670882 |
| ENSMUSG0000002617 | chr1 | Bcs1l | 0.211292905 | 0.592670882 |
| ENSMUSG0000003227 | chr9 | Idh3a | 0.211337535 | 0.592670882 |
| ENSMUSG0000002127 | chr12 | Hsp90aa1 | 0.27220325 | 0.592670882 |
| ENSMUSG0000007004 | chr8 | Fat1 | 0.218983429 | 0.592670882 |
| ENSMUSG0000002550 | chr7 | Rplp2 | 0.290633969 | 0.592670882 |
| ENSMUSG0000005659 | chr4 | Trnp1 | 0.298115972 | 0.592670882 |
| ENSMUSG0000003812 | chr18 | Camk4 | 0.403557115 | 0.592670882 |
| ENSMUSG0000004567 | chr4 | Col27a1 | 0.297695368 | 0.592670882 |
| ENSMUSG0000003007 | chr6 | Chl1 | -0.241033877 | 0.592866695 |
| ENSMUSG0000002535 | chr10 | Dgka | -0.2332931 | 0.593019653 |
| ENSMUSG0000003312 | chr10 | Ybey | -0.265935319 | 0.593019653 |
| ENSMUSG0000002277 | chr16 | Senp5 | 0.135567648 | 0.593019653 |
| ENSMUSG0000003040 | chr7 | Rtn2 | 0.159062493 | 0.593019653 |
| ENSMUSG0000004719 | chr11 | Gjd3 | 0.702653604 | 0.593019653 |
| ENSMUSG0000004084 | chr7 | Six5 | 0.199609042 | 0.593052004 |
| ENSMUSG0000002079 | chr11 | Ankfy1 | -0.09974966 | 0.593068654 |
| ENSMUSG0000003183 | chr8 | Taf1c | 0.20910412 | 0.593068654 |
| ENSMUSG0000003019 | chr6 | Etv6 | 0.196219572 | 0.594194427 |
| ENSMUSG0000007475 | chr2 | Smim26 | 0.341068372 | 0.594368307 |
| ENSMUSG0000002487 | chr19 | Zng1 | 0.207779133 | 0.594401252 |
| ENSMUSG0000008568 | chr11 | 4930469K13Rik | -0.456828675 | 0.594405832 |
| ENSMUSG0000003491 | chr9 | Pygo1 | -0.172161824 | 0.594405832 |
| ENSMUSG0000002972 | chr5 | Tsc22d4 | 0.165520674 | 0.594405832 |
| ENSMUSG0000002030 | chr10 | Tpgs1 | 0.186316283 | 0.594405832 |
| ENSMUSG0000002953 | chr5 | Srsf9 | 0.180567364 | 0.594405832 |
| ENSMUSG0000001037 | chr14 | Nedd8 | 0.216960878 | 0.594405832 |
| ENSMUSG0000005198 | chr16 | Smim11 | 0.30929399 | 0.594405832 |
| ENSMUSG0000005131 | chr2 | Mtln | 0.360608453 | 0.594405832 |
| ENSMUSG0000003870 | chr7 | Aspdh | -0.679180973 | 0.594759657 |
| ENSMUSG0000001583 | chr11 | Sqstm1 | 0.159266581 | 0.594759657 |
| ENSMUSG0000002515 | chr19 | Zdhhc16 | 0.179641699 | 0.594996759 |
| ENSMUSG0000003182 | chr8 | Klhl36 | -0.186924234 | 0.595661154 |
| ENSMUSG0000005932 | chr19 | Csf2ra | -0.126263228 | 0.595832912 |
| ENSMUSG0000002435 | chr18 | Hspa9 | 0.12507091 | 0.59650117 |
| ENSMUSG0000005771 | chr1 | A830018L16Rik | -0.324112833 | 0.596506794 |
| ENSMUSG0000003065 | chr7 | Smg1 | -0.13723822 | 0.596859188 |
| ENSMUSG0000002891 | chr4 | Mecr | 0.20259303 | 0.596859188 |
| ENSMUSG0000003327 | chr17 | Ptprm | 0.26400541 | 0.597126965 |
| ENSMUSG0000002832 | chr4 | Xpa | 0.23116592 | 0.597128246 |
| ENSMUSG0000001774 | chr11 | Ghdc | 0.232045828 | 0.597128246 |
| ENSMUSG0000004780 | chr11 | Akap10 | 0.149504949 | 0.597208587 |
| ENSMUSG0000004050 | chr10 | Marchf9 | -0.172384561 | 0.597940958 |
| ENSMUSG0000002818 | chr3 | Spata1 | -0.318868733 | 0.597952366 |
| ENSMUSG0000002146 | chr13 | Sptlc1 | 0.20466032 | 0.597996893 |
| ENSMUSG0000002710 | chr2 | Chrna1 | -1.278850779 | 0.598050957 |
| ENSMUSG0000006365 | chr1 | Zbtb18 | 0.14700863 | 0.598050957 |
| ENSMUSG0000002866 | chr4 | Epha8 | 0.405725346 | 0.598050957 |

| | | | | |
|---|---|---|---|---|
| ENSMUSG0000003757 | chr15 | Kcnh3 | -0.581237256 | 0.598190419 |
| ENSMUSG0000001886 | chr15 | Sult4a1 | -0.242866503 | 0.598190419 |
| ENSMUSG0000005667 | chrX | Gpr173 | -0.255974573 | 0.598190419 |
| ENSMUSG0000000635 | chr12 | Crip2 | -0.194274222 | 0.598190419 |
| ENSMUSG0000001860 | chr5 | Tbx3 | -0.119814663 | 0.598190419 |
| ENSMUSG0000000071 | chr10 | Rab5b | 0.115430996 | 0.598190419 |
| ENSMUSG0000002456 | chr18 | Mbd1 | 0.145395111 | 0.598190419 |
| ENSMUSG0000002806 | chr3 | Khdc4 | 0.145902707 | 0.598190419 |
| ENSMUSG0000000466 | chr10 | Cnn2 | 0.208743257 | 0.598190419 |
| ENSMUSG0000002685 | chr2 | Ntmt1 | 0.186140996 | 0.598190419 |
| ENSMUSG0000003092 | chr7 | Lyrm1 | 0.264970401 | 0.598190419 |
| ENSMUSG0000004127 | chr11 | Ttc1 | 0.22031117 | 0.598190419 |
| ENSMUSG0000006293 | chr4 | Mtap | 0.242682823 | 0.598190419 |
| ENSMUSG0000005285 | chr6 | Reep1 | -0.213381205 | 0.598209003 |
| ENSMUSG0000003532 | chr5 | Sec31a | 0.102799125 | 0.598209003 |
| ENSMUSG0000003353 | chr12 | Ttc7b | 0.149702767 | 0.598402413 |
| ENSMUSG0000007052 | chr7 | Nsmce3 | 0.155769574 | 0.598458801 |
| ENSMUSG0000004582 | chr13 | Serpinb9 | 0.240099175 | 0.598458801 |
| ENSMUSG0000003174 | chr8 | Mmp2 | 0.268160868 | 0.598458801 |
| ENSMUSG0000002780 | chr3 | Tm4sf1 | 0.347501545 | 0.598458801 |
| ENSMUSG0000005317 | chr7 | Bcl3 | -0.438987489 | 0.598491965 |
| ENSMUSG0000002171 | chr13 | Srek1ip1 | 0.204797415 | 0.598642168 |
| ENSMUSG0000003302 | chr15 | Polr2f | 0.315889274 | 0.598642168 |
| ENSMUSG0000002431 | chr18 | Mep1b | -1.858972185 | 0.598759613 |
| ENSMUSG0000002916 | chr5 | Emilin1 | 0.141763622 | 0.598759613 |
| ENSMUSG0000003434 | chr3 | Smc4 | 0.181487819 | 0.598793746 |
| ENSMUSG0000002122 | chr12 | Papln | -0.37882616 | 0.598841036 |
| ENSMUSG0000007396 | chr7 | Trim68 | -0.242218208 | 0.59914541 |
| ENSMUSG0000004760 | chr19 | Frat2 | -0.466877565 | 0.59914541 |
| ENSMUSG0000003031 | chr6 | Cand2 | -0.162194104 | 0.599145488 |
| ENSMUSG0000003688 | chr3 | Arhgef26 | 0.258917399 | 0.599336488 |
| ENSMUSG0000002696 | chr2 | Anapc2 | 0.117594293 | 0.599464893 |
| ENSMUSG0000000222 | chr5 | Paxip1 | -0.178560392 | 0.599767234 |
| ENSMUSG0000002063 | chr12 | Fkbp1b | -0.23748 | 0.599816829 |
| ENSMUSG0000001574 | chr3 | Anp32e | 0.112362341 | 0.599829991 |
| ENSMUSG0000005724 | chr17 | Tsbp1 | -0.608148685 | 0.599872225 |
| ENSMUSG0000003890 | chr18 | Ccdc68 | -0.324712424 | 0.600362234 |
| ENSMUSG0000000024 | chr7 | Tspan32 | -0.583396331 | 0.600494986 |
| ENSMUSG0000007527 | chr2 | Ift70b | -0.171296222 | 0.600549819 |
| ENSMUSG0000005858 | chr9 | Tmod3 | 0.112229207 | 0.600549819 |
| ENSMUSG0000002730 | chr2 | Dnaaf9 | 0.22952367 | 0.600549819 |
| ENSMUSG0000002284 | chr16 | Ece2 | -0.339759024 | 0.600712614 |
| ENSMUSG0000003883 | chr6 | Agbl3 | -0.261156709 | 0.600712614 |
| ENSMUSG0000007356 | chr18 | Csnk1g3 | -0.122355715 | 0.600712614 |
| ENSMUSG0000003751 | chr2 | Pank2 | 0.137655407 | 0.600712614 |
| ENSMUSG0000002286 | chr16 | Dgkg | -0.248222513 | 0.600769862 |
| ENSMUSG0000009625 | chr17 | Dynlt1b | 0.360922213 | 0.600937874 |
| ENSMUSG0000001943 | chr8 | Gipc1 | 0.193735463 | 0.601038643 |
| ENSMUSG0000006323 | chr15 | 1110038F14Rik | 0.1649851 | 0.601171869 |
| ENSMUSG0000005283 | chr8 | Junb | -0.383481503 | 0.601519861 |

| | | | | |
|---|---|---|---|---|
| ENSMUSG0000003132 | chrX | Chic1 | -0.175017295 | 0.601760465 |
| ENSMUSG0000001971 | chr2 | Gle1 | 0.16404247 | 0.601760465 |
| ENSMUSG0000002904 | chr4 | Rer1 | 0.200120983 | 0.601760465 |
| ENSMUSG0000002064 | chr12 | Rnf144a | 0.241886185 | 0.601760465 |
| ENSMUSG0000002817 | chr3 | Tacr3 | -0.321954169 | 0.60192599 |
| ENSMUSG0000004249 | chr9 | Prdm10 | -0.180723422 | 0.60192599 |
| ENSMUSG0000002614 | chr1 | Rhbdd1 | 0.122006948 | 0.60192599 |
| ENSMUSG0000007343 | chr17 | Wdr90 | -0.205000382 | 0.60234701 |
| ENSMUSG0000004753 | chr1 | Fbxo28 | 0.105776227 | 0.602471755 |
| ENSMUSG0000002053 | chr11 | Shmt1 | -0.522596729 | 0.602693316 |
| ENSMUSG0000002687 | chr2 | Zeb2 | -0.211877155 | 0.602773988 |
| ENSMUSG0000001984 | chr10 | Tube1 | -0.392599125 | 0.602773988 |
| ENSMUSG0000007313 | chrX | Vma21 | 0.132479461 | 0.602773988 |
| ENSMUSG0000007955 | chr1 | Mpp4 | -0.824686426 | 0.602971232 |
| ENSMUSG0000004687 | chr11 | Irgm1 | -0.234318533 | 0.603303943 |
| ENSMUSG0000007296 | chrX | Bhlhb9 | -0.134648312 | 0.603339938 |
| ENSMUSG0000005620 | chr19 | Npm3 | 0.390675466 | 0.603339938 |
| ENSMUSG0000003148 | chr8 | Slc25a15 | -0.233155009 | 0.603356616 |
| ENSMUSG0000002136 | chr13 | Elovl2 | 0.274951547 | 0.603356616 |
| ENSMUSG0000004852 | chr5 | Fbxl13 | -0.872921289 | 0.603640708 |
| ENSMUSG0000003018 | chr6 | Klra2 | -1.000999324 | 0.603640708 |
| ENSMUSG0000003025 | chr6 | Srgap3 | -0.24061307 | 0.603640708 |
| ENSMUSG0000006922 | chr13 | Gprin1 | -0.263146134 | 0.603640708 |
| ENSMUSG0000006810 | chr15 | Cenpm | -0.994551016 | 0.603640708 |
| ENSMUSG0000006203 | chr7 | Pgghg | -0.1726918 | 0.603640708 |
| ENSMUSG0000002812 | chr3 | Gclm | -0.17877201 | 0.603640708 |
| ENSMUSG0000002031 | chr11 | Sptbn1 | -0.123548654 | 0.603640708 |
| ENSMUSG0000002869 | chr4 | Mmachc | 0.206388436 | 0.603640708 |
| ENSMUSG0000004841 | chr3 | Mlf1 | -0.616796541 | 0.603826754 |
| ENSMUSG0000004031 | chr2 | Alx4 | -0.918209411 | 0.603826754 |
| ENSMUSG0000003463 | chr4 | Zyg11b | -0.132472409 | 0.603826754 |
| ENSMUSG0000003370 | chr8 | Fcsk | 0.162668888 | 0.603826754 |
| ENSMUSG0000008726 | chr3 | Lamtor5 | 0.245160022 | 0.603826754 |
| ENSMUSG0000002372 | chr11 | Mrps23 | 0.255089394 | 0.603826754 |
| ENSMUSG0000002506 | chr19 | Col17a1 | 0.415087775 | 0.603826754 |
| ENSMUSG0000005240 | chr4 | Ccdc171 | -0.480660351 | 0.603840457 |
| ENSMUSG0000007936 | chr5 | Gbp4 | -0.443032758 | 0.603840457 |
| ENSMUSG0000003782 | chr14 | Tspan14 | -0.171203701 | 0.603840457 |
| ENSMUSG0000004910 | chr12 | Dcaf5 | -0.116835767 | 0.603840457 |
| ENSMUSG0000003462 | chr10 | Rxylt1 | 0.17997349 | 0.603840457 |
| ENSMUSG0000004004 | chr10 | Rbms2 | 0.173875278 | 0.603840457 |
| ENSMUSG0000003034 | chr6 | D6Wsu163e | 0.192155547 | 0.603840457 |
| ENSMUSG0000004042 | chr7 | Hipk4 | 0.262480476 | 0.603840457 |
| ENSMUSG0000002043 | chr11 | Pes1 | 0.125994158 | 0.604087455 |
| ENSMUSG0000004842 | chr9 | Timm29 | 0.13947034 | 0.604087455 |
| ENSMUSG0000000298 | chr7 | Bcam | 0.317610581 | 0.604087455 |
| ENSMUSG0000006427 | chr1 | Gpbar1 | 0.939205389 | 0.604175014 |
| ENSMUSG0000007388 | chr4 | Il11ra1 | 0.278857953 | 0.604220515 |
| ENSMUSG0000002424 | chr17 | Sos1 | -0.163273071 | 0.604320503 |
| ENSMUSG0000002489 | chr19 | Minpp1 | 0.142343686 | 0.60454309 |

| | | | | |
|---|---|---|---|---|
| ENSMUSG0000002841 | chr4 | Bag1 | 0.243127386 | 0.60454309 |
| ENSMUSG0000011521 | chr16 | Eef1akmt4 | 0.560634837 | 0.60454309 |
| ENSMUSG0000006406 | chr3 | Mtx1 | 0.17173567 | 0.60460032 |
| ENSMUSG0000002860 | chr4 | Cpt2 | -0.209148316 | 0.604651756 |
| ENSMUSG0000004835 | chrX | Arxes1 | -0.380352593 | 0.604718703 |
| ENSMUSG0000006425 | chr7 | Ethe1 | -0.316359029 | 0.604718703 |
| ENSMUSG0000006053 | chr18 | Dcc | 0.374969381 | 0.604718703 |
| ENSMUSG0000004979 | chr12 | Bag5 | 0.182589143 | 0.604865986 |
| ENSMUSG0000003701 | chr18 | Ss18 | 0.133505501 | 0.604959484 |
| ENSMUSG0000002840 | chr4 | Aco1 | 0.131552907 | 0.605635293 |
| ENSMUSG0000009672 | chr17 | Psmb9 | -0.417940993 | 0.605834435 |
| ENSMUSG0000007367 | chr1 | Pgap1 | -0.209462519 | 0.605834435 |
| ENSMUSG0000002620 | chr1 | Glb1l | -0.193616149 | 0.605834435 |
| ENSMUSG0000004575 | chr7 | Tssc4 | 0.162768458 | 0.605834435 |
| ENSMUSG0000002232 | chr15 | Stk3 | 0.194200057 | 0.605834435 |
| ENSMUSG0000002847 | chr4 | Reck | 0.201451702 | 0.605834435 |
| ENSMUSG0000003181 | chr8 | Mvb12a | 0.219318174 | 0.605834435 |
| ENSMUSG0000002275 | chr16 | Nit2 | 0.250391826 | 0.605875055 |
| ENSMUSG0000007175 | chr15 | Zhx2 | 0.198789264 | 0.605897737 |
| ENSMUSG0000002171 | chr13 | Trim23 | 0.132577268 | 0.606157297 |
| ENSMUSG0000005095 | chr13 | Zfp169 | -0.289088304 | 0.606232344 |
| ENSMUSG0000004010 | chr19 | Plpp6 | -0.203660418 | 0.606232344 |
| ENSMUSG0000001521 | chrX | Hmgb3 | -0.160269825 | 0.606232344 |
| ENSMUSG0000005199 | chr1 | Lax1 | 0.351927333 | 0.606232344 |
| ENSMUSG0000004059 | chr1 | Pogk | -0.16863202 | 0.606557104 |
| ENSMUSG0000001546 | chr17 | Notch4 | 0.460158469 | 0.606623597 |
| ENSMUSG0000003253 | chr9 | Ephb1 | 0.351965406 | 0.606629302 |
| ENSMUSG0000003138 | chrX | Plxnb3 | -0.235147052 | 0.606713601 |
| ENSMUSG0000002803 | chr3 | Efna3 | -0.792286394 | 0.606713601 |
| ENSMUSG0000000692 | chr11 | Ezh1 | -0.144741744 | 0.606713601 |
| ENSMUSG0000002668 | chr1 | Nuf2 | -0.61692699 | 0.606725882 |
| ENSMUSG0000000223 | chr3 | Rhoc | 0.249145994 | 0.606725882 |
| ENSMUSG0000003420 | chr16 | Polq | -0.536710306 | 0.606758745 |
| ENSMUSG0000000526 | chr15 | Prlr | -0.266672671 | 0.606801175 |
| ENSMUSG0000005531 | chr2 | Them7 | -1.012885078 | 0.606866239 |
| ENSMUSG0000001836 | chr11 | Smurf2 | 0.129655452 | 0.606866239 |
| ENSMUSG0000001509 | chr2 | Clic3 | 0.564085064 | 0.606866239 |
| ENSMUSG0000004165 | chr11 | Slc39a11 | -0.316851745 | 0.606915665 |
| ENSMUSG0000003675 | chr2 | Tubb4b | 0.159675995 | 0.606915665 |
| ENSMUSG0000003078 | chr7 | Rusf1 | -0.145618763 | 0.607314178 |
| ENSMUSG0000004197 | chr3 | Arhgef11 | -0.175616895 | 0.607374205 |
| ENSMUSG0000005135 | chr4 | Zfp46 | 0.244435088 | 0.607546551 |
| ENSMUSG0000005892 | chr18 | Ccdc192 | 0.452216915 | 0.607849556 |
| ENSMUSG0000003429 | chr12 | Nek9 | 0.142481738 | 0.608046222 |
| ENSMUSG0000003264 | chr7 | Chsy1 | 0.169602558 | 0.608046222 |
| ENSMUSG0000005504 | chr19 | Pdlim1 | 0.237574614 | 0.608046222 |
| ENSMUSG0000003731 | chrX | Jade3 | -0.227612287 | 0.608288561 |
| ENSMUSG0000006282 | chr13 | 4833420G17Rik | -0.378267161 | 0.608311661 |
| ENSMUSG0000000661 | chr13 | Hfe | 0.167786276 | 0.608311661 |
| ENSMUSG0000007080 | chr4 | Cited4 | 0.282370036 | 0.608929943 |

| | | | | |
|---|---|---|---|---|
| ENSMUSG0000000013 | chrX | Tfe3 | 0.132143038 | 0.608967793 |
| ENSMUSG0000001980 | chr10 | Snx3 | 0.226236394 | 0.609251435 |
| ENSMUSG0000003144 | chr8 | Mcf2l | -0.25475035 | 0.609261706 |
| ENSMUSG0000007946 | chr9 | Pigb | -0.190870042 | 0.609261706 |
| ENSMUSG0000003248 | chr9 | Dhx30 | 0.101738562 | 0.609261706 |
| ENSMUSG0000004202 | chr12 | Ncapg2 | -0.281442576 | 0.609335796 |
| ENSMUSG0000002755 | chr3 | Car13 | -0.336079766 | 0.609335796 |
| ENSMUSG0000007069 | chr5 | Utp3 | 0.173915591 | 0.609335796 |
| ENSMUSG0000005754 | chr5 | Pus7 | 0.147514632 | 0.609387413 |
| ENSMUSG0000002136 | chr13 | Tbc1d7 | 0.19671992 | 0.60951282 |
| ENSMUSG0000002073 | chr11 | Nherf1 | -0.377299772 | 0.609658724 |
| ENSMUSG0000004321 | chr6 | Hoxa6 | -0.290239972 | 0.609658724 |
| ENSMUSG0000004598 | chr16 | Eif4g1 | 0.110832349 | 0.609658724 |
| ENSMUSG0000002284 | chr16 | Ap2m1 | 0.148071993 | 0.609658724 |
| ENSMUSG0000002280 | chr16 | Fyttd1 | 0.14142998 | 0.609658724 |
| ENSMUSG0000003373 | chr17 | Fkbpl | 0.214783071 | 0.609658724 |
| ENSMUSG0000003089 | chr7 | Cckbr | -0.546148704 | 0.609741082 |
| ENSMUSG0000001892 | chr11 | Pelp1 | 0.151054019 | 0.609741082 |
| ENSMUSG0000003697 | chr1 | Tmem177 | 0.175095486 | 0.609741082 |
| ENSMUSG0000005309 | chr5 | Tmem248 | 0.098899123 | 0.609812212 |
| ENSMUSG0000003746 | chr1 | Ints7 | -0.156780534 | 0.610103754 |
| ENSMUSG0000002078 | chr11 | Tsen54 | 0.166686837 | 0.610103754 |
| ENSMUSG0000002166 | chr13 | Ankra2 | 0.161033161 | 0.610103754 |
| ENSMUSG0000004690 | chr11 | Tefm | 0.255627411 | 0.610103754 |
| ENSMUSG0000002888 | chr4 | Smpdl3b | 1.085798487 | 0.610103754 |
| ENSMUSG0000002127 | chr12 | Amn | -1.109829518 | 0.610218915 |
| ENSMUSG0000004238 | chr4 | Dlgap3 | -0.241302022 | 0.610218915 |
| ENSMUSG0000004143 | chr13 | Ccnb1 | -0.845653803 | 0.610509164 |
| ENSMUSG0000002347 | chr9 | Celsr3 | -0.297639672 | 0.610509164 |
| ENSMUSG0000000128 | chr15 | Myg1 | 0.163636172 | 0.610509164 |
| ENSMUSG0000003227 | chr9 | Nnmt | 0.362942651 | 0.610509164 |
| ENSMUSG0000003434 | chr1 | Ube2f | 0.169509357 | 0.610535058 |
| ENSMUSG0000007272 | chr5 | Myo18b | 0.72775012 | 0.610535058 |
| ENSMUSG0000002170 | chr13 | Rab3c | -0.218399004 | 0.610809349 |
| ENSMUSG0000001535 | chr9 | Pcolce2 | 0.350729243 | 0.610809349 |
| ENSMUSG0000002674 | chr2 | Dnajc1 | 0.11632638 | 0.611374026 |
| ENSMUSG0000005835 | chr8 | Abce1 | 0.15086133 | 0.61143489 |
| ENSMUSG0000003733 | chr11 | Larp1 | -0.155809554 | 0.611470855 |
| ENSMUSG0000007892 | chr11 | Tgtp2 | -0.660967617 | 0.611535117 |
| ENSMUSG0000004403 | chr9 | Als2cl | -0.369797977 | 0.611535117 |
| ENSMUSG0000005381 | chr3 | Camk2d | 0.125395269 | 0.611535117 |
| ENSMUSG0000004065 | chr9 | Oaz2 | 0.147018367 | 0.611535117 |
| ENSMUSG0000001873 | chr11 | Ndel1 | 0.14586023 | 0.611535117 |
| ENSMUSG0000002812 | chr3 | F3 | 0.166489419 | 0.611535117 |
| ENSMUSG0000004425 | chr4 | Pcsk9 | 0.513562938 | 0.611535117 |
| ENSMUSG0000003978 | chr1 | Kmo | -0.643215629 | 0.611774022 |
| ENSMUSG0000001676 | chr15 | Scube1 | -0.209242742 | 0.611804277 |
| ENSMUSG0000003814 | chr8 | Stox2 | -0.20813469 | 0.611886355 |
| ENSMUSG0000003448 | chr18 | Snx2 | 0.165240128 | 0.611946143 |
| ENSMUSG0000002899 | chr4 | Rbp7 | 0.883143669 | 0.611946143 |

| | | | | |
|---|---|---|---|---|
| ENSMUSG0000001559 | chr17 | Ttbk1 | -0.268128677 | 0.612066514 |
| ENSMUSG0000001524 | chr4 | Abca1 | -0.39866381 | 0.61213505 |
| ENSMUSG0000002476 | chr19 | Naa40 | -0.144134641 | 0.61213505 |
| ENSMUSG0000009601 | chr8 | Sox1 | -0.541496623 | 0.612256563 |
| ENSMUSG0000003746 | chr12 | Tedc1 | -0.681065507 | 0.612301975 |
| ENSMUSG0000003348 | chr14 | Fndc3a | -0.137706073 | 0.612515627 |
| ENSMUSG0000002515 | chr11 | Cbr2 | 0.273261529 | 0.612515627 |
| ENSMUSG0000002677 | chr2 | Pfkfb3 | -0.294813856 | 0.612655472 |
| ENSMUSG0000003170 | chr8 | Dnaja2 | 0.12198805 | 0.612655472 |
| ENSMUSG0000003520 | chr10 | Sppl2b | -0.148516166 | 0.612974154 |
| ENSMUSG0000003290 | chr1 | Sgpp2 | -0.263488682 | 0.613015952 |
| ENSMUSG0000002051 | chr11 | Cnot8 | 0.16127628 | 0.613015952 |
| ENSMUSG0000007288 | chr5 | Nfxl1 | 0.162311721 | 0.613015952 |
| ENSMUSG0000003391 | chr7 | Gde1 | 0.182091273 | 0.613015952 |
| ENSMUSG0000003907 | chr6 | Cpa4 | -0.838128644 | 0.613417583 |
| ENSMUSG0000003002 | chr6 | Lrig1 | 0.196826982 | 0.613522732 |
| ENSMUSG0000003859 | chr1 | Capn8 | -1.6470343 | 0.614101726 |
| ENSMUSG0000004193 | chr4 | Agrn | -0.224152274 | 0.614101726 |
| ENSMUSG0000002092 | chr11 | Eftud2 | 0.112123593 | 0.614101726 |
| ENSMUSG0000002420 | chr17 | Kdm4b | 0.120608746 | 0.614101726 |
| ENSMUSG0000003729 | chr4 | Ldlrap1 | 0.194326189 | 0.614101726 |
| ENSMUSG0000000703 | chr17 | Vwa7 | 0.34462093 | 0.614101726 |
| ENSMUSG0000005713 | chr6 | Tmem140 | -0.376228996 | 0.614194546 |
| ENSMUSG0000003960 | chr9 | Rbms3 | -0.141492337 | 0.614194546 |
| ENSMUSG0000001943 | chr17 | Ddx39b | 0.097472279 | 0.614194546 |
| ENSMUSG0000010933 | chr7 | Samd4b | 0.148857048 | 0.614194546 |
| ENSMUSG0000003850 | chr8 | Dcun1d2 | 0.140144123 | 0.614194546 |
| ENSMUSG0000004955 | chr5 | Fzd9 | 0.441027607 | 0.614194546 |
| ENSMUSG0000002707 | chr2 | Timm10 | 0.332990309 | 0.614194546 |
| ENSMUSG0000002684 | chr2 | Abl1 | 0.155079938 | 0.614210591 |
| ENSMUSG0000004226 | chr8 | Cibar2 | -0.317388834 | 0.614238874 |
| ENSMUSG0000000693 | chr9 | Ctnnb1 | 0.143764257 | 0.614238874 |
| ENSMUSG0000004135 | chrX | Vcf2 | 0.170295918 | 0.614245132 |
| ENSMUSG0000002663 | chr1 | Batf3 | 0.764237022 | 0.614487509 |
| ENSMUSG0000005044 | chr9 | Cyp8b1 | -0.802179318 | 0.614520817 |
| ENSMUSG0000002019 | chr10 | Ap3d1 | 0.106109703 | 0.614520817 |
| ENSMUSG0000002874 | chr4 | Slc66a1 | 0.192762042 | 0.614520817 |
| ENSMUSG0000004131 | chr5 | Slc7a1 | 0.232891539 | 0.614520817 |
| ENSMUSG0000002981 | chr6 | Gpnmb | -0.401129554 | 0.614525086 |
| ENSMUSG0000003724 | chr11 | Zfp692 | -0.159692595 | 0.614525086 |
| ENSMUSG0000002573 | chr17 | Jmjd8 | 0.115260286 | 0.614525086 |
| ENSMUSG0000003875 | chr6 | Nup205 | 0.125756233 | 0.614525086 |
| ENSMUSG0000002124 | chr12 | Fcf1 | 0.164343245 | 0.614525086 |
| ENSMUSG0000006317 | chr7 | Pstk | 0.18381882 | 0.614525086 |
| ENSMUSG0000001117 | chr12 | Odc1 | 0.268629146 | 0.614525086 |
| ENSMUSG0000009013 | chr8 | Uba52 | 0.373724333 | 0.614525086 |
| ENSMUSG0000002494 | chr19 | Map4k2 | -0.225480418 | 0.614572834 |
| ENSMUSG0000002786 | chr3 | Vangl1 | 0.137826546 | 0.614572834 |
| ENSMUSG0000000395 | chr16 | Fam162a | 0.323052704 | 0.614582941 |
| ENSMUSG0000005000 | chr13 | Idnk | -0.16102006 | 0.61505504 |

| | | | | |
|---|---|---|---|---|
| ENSMUSG0000002662 | chr1 | Nek2 | -1.231186711 | 0.61512056 |
| ENSMUSG0000002737 | chr2 | Mrps5 | 0.10676442 | 0.61512056 |
| ENSMUSG0000004712 | chr10 | 1700113H08Rik | 0.746593224 | 0.61512056 |
| ENSMUSG0000000764 | chr11 | Rad51c | 0.403785079 | 0.61512056 |
| ENSMUSG0000000834 | chr5 | Ubc | 0.322220608 | 0.61512056 |
| ENSMUSG0000005304 | chr9 | Aph1c | -0.230516965 | 0.615716264 |
| ENSMUSG0000002219 | chr15 | 6030458C11Rik | 0.148408192 | 0.616194144 |
| ENSMUSG0000002806 | chr3 | Lamtor2 | 0.270080113 | 0.616194144 |
| ENSMUSG0000007266 | chr6 | Mansc4 | 0.491118567 | 0.61629736 |
| ENSMUSG0000002218 | chr15 | Fbxo4 | 0.232600825 | 0.616404268 |
| ENSMUSG0000007343 | chr17 | Eme2 | 0.199795815 | 0.616633226 |
| ENSMUSG0000003901 | chr9 | Timm8b | 0.348101848 | 0.616659386 |
| ENSMUSG0000003277 | chr7 | Gtf3c1 | -0.108529655 | 0.617179099 |
| ENSMUSG0000002691 | chr2 | Wdr5 | 0.13260474 | 0.61754439 |
| ENSMUSG0000002902 | chr4 | Lrrc47 | -0.126882732 | 0.617584496 |
| ENSMUSG0000002783 | chr3 | Pdcd10 | 0.163831634 | 0.617584496 |
| ENSMUSG0000003035 | chr6 | Tspan9 | 0.27400783 | 0.617584496 |
| ENSMUSG0000002141 | chr13 | Prpf4b | -0.123011036 | 0.617674479 |
| ENSMUSG0000002590 | chr1 | Sntg1 | -0.249277415 | 0.617761228 |
| ENSMUSG0000003794 | chr9 | Ano10 | 0.118240412 | 0.617881518 |
| ENSMUSG0000006026 | chr5 | Gtf2i | 0.104147489 | 0.618340327 |
| ENSMUSG0000004703 | chr2 | Spata2 | -0.163376429 | 0.618370639 |
| ENSMUSG0000006137 | chr10 | Zfp873 | 0.197590313 | 0.618370639 |
| ENSMUSG0000002505 | chrX | Gk | -0.254661279 | 0.618454398 |
| ENSMUSG0000001060 | chr9 | Pigyl | 0.229509169 | 0.618454398 |
| ENSMUSG0000000410 | chr2 | Angptl2 | 0.24192393 | 0.618454398 |
| ENSMUSG0000000781 | chr5 | Zfp655 | 0.123336764 | 0.618520311 |
| ENSMUSG0000000789 | chr7 | Ctsd | 0.11553754 | 0.618980346 |
| ENSMUSG0000002054 | chr11 | Myocd | 0.307857708 | 0.618980346 |
| ENSMUSG0000002091 | chr11 | Krt19 | -0.735367628 | 0.619363585 |
| ENSMUSG0000005383 | chr17 | H2-T24 | -0.401566819 | 0.619363585 |
| ENSMUSG0000004101 | chr14 | Nrg3 | -0.304809529 | 0.619363585 |
| ENSMUSG0000005629 | chr19 | Ms4a4b | -1.440537934 | 0.619363585 |
| ENSMUSG0000003872 | chr11 | Hoxb7 | -0.314348701 | 0.619363585 |
| ENSMUSG0000002297 | chr16 | Synj1 | -0.203124115 | 0.619363585 |
| ENSMUSG0000003656 | chr7 | Lgi4 | -0.215234171 | 0.619363585 |
| ENSMUSG0000004037 | chr11 | Cacng5 | -0.354723977 | 0.619363585 |
| ENSMUSG0000000456 | chr8 | Mcoln1 | -0.194717336 | 0.619363585 |
| ENSMUSG0000003834 | chrX | Txlng | -0.182628188 | 0.619363585 |
| ENSMUSG0000002883 | chr4 | Stmn1 | -0.137490782 | 0.619363585 |
| ENSMUSG0000001837 | chr11 | Srsf1 | -0.09380923 | 0.619363585 |
| ENSMUSG0000001201 | chr16 | Scarf2 | 0.137195668 | 0.619363585 |
| ENSMUSG0000008767 | chr2 | Tmem250 | 0.138957561 | 0.619363585 |
| ENSMUSG0000002834 | chr4 | Tex10 | 0.159329167 | 0.619363585 |
| ENSMUSG0000002961 | chr5 | Erp29 | 0.15684851 | 0.619363585 |
| ENSMUSG0000002189 | chr14 | Mettl6 | 0.168807745 | 0.619363585 |
| ENSMUSG0000001567 | chr13 | Psma2 | 0.199767069 | 0.619363585 |
| ENSMUSG0000003014 | chr6 | Zfp248 | 0.189113362 | 0.619363585 |
| ENSMUSG0000002553 | chr5 | Crcp | 0.210860805 | 0.619363585 |
| ENSMUSG0000002016 | chr10 | Best3 | 0.526207247 | 0.619363585 |

| | | | | |
|---|---|---|---|---|
| ENSMUSG0000000761 | chr13 | Homer1 | 0.209340964 | 0.619363585 |
| ENSMUSG0000002051 | chr11 | Mrpl22 | 0.28714556 | 0.619363585 |
| ENSMUSG0000009403 | chr4 | Ldlrad2 | 0.744313165 | 0.619363585 |
| ENSMUSG0000007412 | chr7 | Rpl13a | 0.2831301 | 0.619363585 |
| ENSMUSG0000003776 | chr10 | Slc16a9 | -0.418512469 | 0.619486217 |
| ENSMUSG0000003624 | chr8 | Gmip | -0.256909874 | 0.619486217 |
| ENSMUSG0000004392 | chr9 | Cenatac | -0.204210388 | 0.619655421 |
| ENSMUSG0000003156 | chr8 | Tenm3 | 0.246733481 | 0.619699669 |
| ENSMUSG0000007887 | chr2 | Gm14410 | 0.216918622 | 0.619824372 |
| ENSMUSG0000000273 | chr2 | Prkra | 0.170227192 | 0.619841732 |
| ENSMUSG0000003061 | chr7 | Sytl2 | -0.373974065 | 0.620188402 |
| ENSMUSG0000001580 | chr5 | Qdpr | -0.251732843 | 0.620469962 |
| ENSMUSG0000002733 | chr2 | Ivd | -0.146269248 | 0.620469962 |
| ENSMUSG0000002959 | chr5 | Plbd2 | 0.161088465 | 0.620469962 |
| ENSMUSG0000002546 | chr7 | Fuom | 0.162241486 | 0.620469962 |
| ENSMUSG0000003924 | chr13 | B3galnt2 | 0.185504549 | 0.620469962 |
| ENSMUSG0000004734 | chrX | Lancl3 | 0.237821735 | 0.620469962 |
| ENSMUSG0000001691 | chr1 | Sulf1 | 0.211455338 | 0.620469962 |
| ENSMUSG0000003373 | chrX | Fndc3c1 | -0.899885088 | 0.620581683 |
| ENSMUSG0000003217 | chr9 | Olfm2 | -0.256154052 | 0.620581683 |
| ENSMUSG0000005845 | chr7 | Dhcr7 | -0.225753683 | 0.620581683 |
| ENSMUSG0000003125 | chrX | Cstf2 | -0.143111642 | 0.620581683 |
| ENSMUSG0000000627 | chr8 | Atp6v1b2 | -0.151164034 | 0.620581683 |
| ENSMUSG0000001899 | chr7 | Nars2 | 0.200517191 | 0.620702135 |
| ENSMUSG0000003913 | chr6 | Zc3hc1 | 0.261559577 | 0.620702135 |
| ENSMUSG0000002228 | chr15 | Pabpc1 | 0.170452775 | 0.620905584 |
| ENSMUSG0000003197 | chr8 | Ccsap | -0.319491427 | 0.621042691 |
| ENSMUSG0000005636 | chr5 | Actr3b | -0.221500855 | 0.621042691 |
| ENSMUSG0000003271 | chr17 | Mdfi | -0.277390871 | 0.621042691 |
| ENSMUSG0000001392 | chr7 | Clip3 | -0.133217954 | 0.621042691 |
| ENSMUSG0000003113 | chrX | Arhgef6 | 0.205026299 | 0.621042691 |
| ENSMUSG0000006009 | chr8 | Prmt7 | 0.227623593 | 0.621042691 |
| ENSMUSG0000001864 | chr11 | Dusp14 | 0.264128281 | 0.621042691 |
| ENSMUSG0000002286 | chr16 | Btg3 | 0.325513311 | 0.621042691 |
| ENSMUSG0000002922 | chr5 | Lnx1 | -0.414442234 | 0.621061701 |
| ENSMUSG0000003052 | chr7 | Crtc3 | 0.131876703 | 0.621061701 |
| ENSMUSG0000001486 | chr2 | Surf4 | 0.161599787 | 0.621061701 |
| ENSMUSG0000004870 | chr10 | Ccdc6 | 0.149902478 | 0.621061701 |
| ENSMUSG0000003413 | chr9 | Sik3 | 0.149350208 | 0.621061701 |
| ENSMUSG0000000426 | chr6 | Phb2 | 0.144142342 | 0.621061701 |
| ENSMUSG0000003230 | chr9 | Psma4 | 0.172700631 | 0.621061701 |
| ENSMUSG0000004771 | chr4 | Ubiad1 | 0.178576544 | 0.621061701 |
| ENSMUSG0000005018 | chr4 | Lsm10 | 0.258918893 | 0.621061701 |
| ENSMUSG0000002716 | chr2 | Elp4 | 0.175644186 | 0.621208841 |
| ENSMUSG0000003233 | chr9 | Loxl1 | 0.199236702 | 0.621516935 |
| ENSMUSG0000000277 | chr7 | Grin2d | -0.26833027 | 0.621953282 |
| ENSMUSG0000002151 | chr13 | Ptdss1 | -0.148120249 | 0.621982257 |
| ENSMUSG0000002883 | chr4 | Extl1 | -0.556366472 | 0.622147964 |
| ENSMUSG0000002684 | chr2 | Fibcd1 | -0.310681048 | 0.622147964 |
| ENSMUSG0000003115 | chrX | Timm17b | 0.203354395 | 0.622147964 |

| | | | | |
|---|---|---|---|---|
| ENSMUSG0000002440 | chr17 | Atp6v1g2 | -0.189095492 | 0.622152531 |
| ENSMUSG0000007467 | chr2 | Ttll9 | -0.70445486 | 0.622152531 |
| ENSMUSG0000002038 | chr11 | Clk4 | 0.134320622 | 0.622152531 |
| ENSMUSG0000004246 | chr6 | C1galt1 | -0.143541335 | 0.622180293 |
| ENSMUSG0000004077 | chr3 | Cept1 | -0.192145145 | 0.62233098 |
| ENSMUSG0000002170 | chr13 | Plk2 | 0.161750162 | 0.622472469 |
| ENSMUSG0000004575 | chr7 | Zfp764 | 0.193927068 | 0.622472469 |
| ENSMUSG0000007871 | chr4 | Tomm5 | 0.258788649 | 0.622472469 |
| ENSMUSG0000004628 | chrX | Pnma3 | -0.387277 | 0.622485503 |
| ENSMUSG0000001794 | chr2 | Gdap1l1 | -0.20523939 | 0.622485503 |
| ENSMUSG0000002882 | chr4 | Sfpq | 0.159278952 | 0.622485503 |
| ENSMUSG0000003098 | chr7 | Dhx32 | 0.171344092 | 0.622485503 |
| ENSMUSG0000000225 | chr17 | Ppard | -0.226952202 | 0.622775005 |
| ENSMUSG0000001253 | chr6 | Tnpo3 | 0.118012019 | 0.622775005 |
| ENSMUSG0000002915 | chr5 | Ociad1 | 0.174534124 | 0.622775005 |
| ENSMUSG0000003545 | chr5 | Prdm8 | 0.564097331 | 0.622899605 |
| ENSMUSG0000010550 | chr3 | Gbp5 | -0.260297977 | 0.622928148 |
| ENSMUSG0000002194 | chr14 | Gdf10 | -0.335372869 | 0.622973395 |
| ENSMUSG0000005093 | chr8 | Map10 | 0.254507469 | 0.623145058 |
| ENSMUSG0000003823 | chr9 | Gask1a | -0.29672236 | 0.623824662 |
| ENSMUSG0000005214 | chr17 | Rps10 | -0.38609834 | 0.623824662 |
| ENSMUSG0000002189 | chr14 | Arhgef3 | -0.127362944 | 0.623824662 |
| ENSMUSG0000002239 | chr15 | Rangap1 | -0.158049 | 0.624262135 |
| ENSMUSG0000003662 | chr4 | Atp13a2 | -0.164325706 | 0.624269761 |
| ENSMUSG0000004316 | chr6 | Pyurf | 0.201683139 | 0.624528217 |
| ENSMUSG0000001432 | chr10 | Bicc1 | 0.229561079 | 0.624835208 |
| ENSMUSG0000002056 | chr12 | Efcab10 | -1.271420638 | 0.624863327 |
| ENSMUSG0000007360 | chr18 | Prob1 | -0.29332445 | 0.624863327 |
| ENSMUSG0000002734 | chr2 | Rasgrp1 | -0.500972198 | 0.624863327 |
| ENSMUSG0000005055 | chr2 | Prokr2 | -0.84317069 | 0.624863327 |
| ENSMUSG0000005408 | chr7 | Capn12 | -0.460517782 | 0.624863327 |
| ENSMUSG0000007460 | chr2 | Tox2 | -0.228987531 | 0.624863327 |
| ENSMUSG0000002753 | chr3 | Snx16 | -0.181258268 | 0.624863327 |
| ENSMUSG0000004056 | chr18 | Wdr7 | -0.155981398 | 0.624863327 |
| ENSMUSG0000003075 | chr7 | Nsmce1 | 0.197722535 | 0.624863327 |
| ENSMUSG0000002267 | chr16 | Mzt2 | 0.204081942 | 0.624863327 |
| ENSMUSG0000004631 | chr18 | Ccbe1 | 0.255986158 | 0.624863327 |
| ENSMUSG0000002873 | chr4 | Cfap57 | -0.823684832 | 0.625058009 |
| ENSMUSG0000002730 | chr2 | Ubox5 | -0.174129121 | 0.625058009 |
| ENSMUSG0000004507 | chr5 | Rnf216 | 0.131485595 | 0.625058009 |
| ENSMUSG0000003160 | chr8 | Klhl2 | 0.140699942 | 0.625058009 |
| ENSMUSG0000002893 | chr4 | Rpl22 | 0.239856916 | 0.625058009 |
| ENSMUSG0000002484 | chr19 | Chka | -0.128517005 | 0.625762754 |
| ENSMUSG0000004790 | chr11 | Sstr2 | -0.440752061 | 0.625837053 |
| ENSMUSG0000002934 | chr5 | Srrd | 0.251762003 | 0.625913228 |
| ENSMUSG0000003347 | chr19 | Fhip2a | 0.132152376 | 0.625991082 |
| ENSMUSG0000003593 | chr12 | Cog5 | 0.124985768 | 0.626154909 |
| ENSMUSG0000001857 | chr11 | 2810408A11Rik | -0.639868832 | 0.626301625 |
| ENSMUSG0000002301 | chr15 | Faim2 | -0.227712981 | 0.626720238 |
| ENSMUSG0000005502 | chr15 | Cntn1 | -0.152640329 | 0.626735612 |

| | | | | |
|---|---|---|---|---|
| ENSMUSG0000007117 | chr8 | Arhgef10 | 0.13876484 | 0.626735612 |
| ENSMUSG0000006024 | chr7 | Cend1 | -0.185926554 | 0.626776532 |
| ENSMUSG0000000555 | chr7 | Atp4a | -0.888701443 | 0.626832549 |
| ENSMUSG0000002232 | chr15 | Matn2 | 0.136960432 | 0.626882607 |
| ENSMUSG0000003027 | chr6 | Camk1 | 0.144726987 | 0.627256401 |
| ENSMUSG0000007073 | chr4 | Tmem35b | -0.265791976 | 0.62746732 |
| ENSMUSG0000003189 | chr8 | Tsnaxip1 | -0.636570354 | 0.62746732 |
| ENSMUSG0000003164 | chr8 | Nek1 | -0.179245989 | 0.62746732 |
| ENSMUSG0000002557 | chr11 | Cant1 | -0.158135424 | 0.62746732 |
| ENSMUSG0000002846 | chr4 | Gba2 | -0.126110398 | 0.62746732 |
| ENSMUSG0000001601 | chr13 | Mtrex | 0.116962298 | 0.62746732 |
| ENSMUSG0000002940 | chr5 | Uso1 | 0.124122584 | 0.62746732 |
| ENSMUSG0000002655 | chr1 | Dcaf8 | 0.145518813 | 0.62746732 |
| ENSMUSG0000003142 | chrX | Psmd10 | 0.25617724 | 0.62746732 |
| ENSMUSG0000002935 | chr5 | Tesc | 0.392951617 | 0.62746732 |
| ENSMUSG0000000175 | chr19 | Tcirg1 | -0.215586242 | 0.627669414 |
| ENSMUSG0000003921 | chr9 | Il18 | -0.504058586 | 0.627732657 |
| ENSMUSG0000002442 | chr18 | Ttc39c | -0.261328204 | 0.627844408 |
| ENSMUSG0000002667 | chr1 | Rgs5 | 0.147670002 | 0.627844408 |
| ENSMUSG0000003468 | chr17 | Rnps1 | 0.115682738 | 0.627844408 |
| ENSMUSG0000002663 | chr1 | Traf5 | 0.183504038 | 0.627844408 |
| ENSMUSG0000002405 | chr17 | Ndc80 | -0.867751263 | 0.627930514 |
| ENSMUSG0000003226 | chr9 | Tmprss5 | -0.219701331 | 0.627930514 |
| ENSMUSG0000008768 | chr8 | Pet100 | 0.317182243 | 0.628027881 |
| ENSMUSG0000005900 | chr2 | Hnrnpa3 | 0.138803862 | 0.628051745 |
| ENSMUSG0000003250 | chr9 | Arpp21 | -0.280243219 | 0.628671621 |
| ENSMUSG0000007529 | chr19 | Aldh3b2 | -0.980950456 | 0.628786293 |
| ENSMUSG0000002741 | chr2 | Otor | -0.473966977 | 0.628786293 |
| ENSMUSG0000004765 | chr19 | Gal3st3 | -0.500662742 | 0.628786293 |
| ENSMUSG0000007085 | chr5 | Nicol1 | -0.29243369 | 0.628786293 |
| ENSMUSG0000004046 | chr11 | Mybbp1a | 0.111770102 | 0.628786293 |
| ENSMUSG0000002771 | chr3 | Acad9 | 0.136059839 | 0.628786293 |
| ENSMUSG0000003711 | chr9 | Sik2 | 0.204933216 | 0.628786293 |
| ENSMUSG0000003249 | chr9 | Nradd | 0.301732865 | 0.628786293 |
| ENSMUSG0000004450 | chr11 | Bod1 | 0.213495136 | 0.628795767 |
| ENSMUSG0000000787 | chr4 | Id3 | 0.364089151 | 0.628993803 |
| ENSMUSG0000002221 | chr14 | Cpne6 | -0.481275164 | 0.629105241 |
| ENSMUSG0000006873 | chr3 | Tmem167b | 0.124821502 | 0.62917632 |
| ENSMUSG0000001093 | chr8 | Vac14 | 0.138271939 | 0.62917632 |
| ENSMUSG0000000445 | chr1 | Ralb | 0.140717097 | 0.62917632 |
| ENSMUSG0000004148 | chr1 | Zfp281 | 0.15021016 | 0.62917632 |
| ENSMUSG0000001965 | chr9 | Ccdc12 | 0.163380075 | 0.62917632 |
| ENSMUSG0000004553 | chr8 | Ddx28 | 0.211117325 | 0.62917632 |
| ENSMUSG0000004072 | chr9 | Scamp5 | -0.218768524 | 0.629377303 |
| ENSMUSG0000003882 | chr10 | Hace1 | -0.161375188 | 0.629393245 |
| ENSMUSG0000002848 | chr4 | Sh3gl2 | 0.152008058 | 0.62948761 |
| ENSMUSG0000004084 | chr4 | Szrd1 | 0.136753242 | 0.629606189 |
| ENSMUSG0000003502 | chr10 | Map2k2 | 0.177452188 | 0.629606189 |
| ENSMUSG0000006157 | chr5 | Ksr2 | -0.298441029 | 0.629858444 |
| ENSMUSG0000002889 | chr4 | Mtf1 | 0.157050955 | 0.62998917 |

| | | | | |
|---|---|---|---|---|
| ENSMUSG0000005313 | chr15 | Mapk11 | -0.241623065 | 0.630150102 |
| ENSMUSG0000003193 | chr9 | Med17 | 0.130871089 | 0.630150102 |
| ENSMUSG0000003615 | chr8 | Tm6sf2 | -0.579367658 | 0.630476317 |
| ENSMUSG0000002103 | chr12 | Ahsa1 | 0.241162414 | 0.630476317 |
| ENSMUSG0000002639 | chr1 | Ptprc | -0.334656424 | 0.630848015 |
| ENSMUSG0000011855 | chr16 | A930007A09Rik | -0.339092533 | 0.630848015 |
| ENSMUSG0000006726 | chr4 | Foxd3 | -0.238125563 | 0.630848015 |
| ENSMUSG0000002525 | chr3 | Zfhx4 | -0.243010673 | 0.630848015 |
| ENSMUSG0000002574 | chr5 | Tyms | -0.237690255 | 0.630848015 |
| ENSMUSG0000002415 | chr17 | Msh2 | -0.174420233 | 0.630848015 |
| ENSMUSG0000002311 | chr7 | Sympk | 0.092580073 | 0.630848015 |
| ENSMUSG0000005214 | chr6 | Ppp4r2 | 0.102109096 | 0.630848015 |
| ENSMUSG0000007643 | chr12 | Ywhaq | 0.182068709 | 0.630848015 |
| ENSMUSG0000002124 | chr12 | Npc2 | 0.167877765 | 0.630848015 |
| ENSMUSG0000000702 | chr17 | Vars1 | 0.155120711 | 0.630848015 |
| ENSMUSG0000003423 | chr14 | Usp54 | 0.16960128 | 0.630848015 |
| ENSMUSG0000003043 | chr7 | Rpl28 | 0.215628546 | 0.630848015 |
| ENSMUSG0000002631 | chr1 | Hdac4 | 0.20244326 | 0.630848015 |
| ENSMUSG0000003375 | chr8 | Gadd45gip1 | 0.279317154 | 0.630848015 |
| ENSMUSG0000002502 | chr19 | Slit1 | 0.280924073 | 0.630848015 |
| ENSMUSG0000007925 | chr9 | Trim71 | 1.415599887 | 0.630848015 |
| ENSMUSG0000005862 | chr17 | Capn11 | 1.923986201 | 0.630848015 |
| ENSMUSG0000008168 | chr5 | Fzd10 | -0.630716057 | 0.630863474 |
| ENSMUSG0000001385 | chr10 | Tmem259 | 0.106410681 | 0.630863474 |
| ENSMUSG0000005296 | chr6 | Mrpl35 | 0.161812537 | 0.630865325 |
| ENSMUSG0000002405 | chr17 | Cyp4f13 | -0.241650661 | 0.631058053 |
| ENSMUSG0000004615 | chr8 | Fut10 | -0.179047005 | 0.631058053 |
| ENSMUSG0000006388 | chr6 | Ptcd3 | 0.097071137 | 0.631058053 |
| ENSMUSG0000003084 | chr7 | Bag3 | 0.110103199 | 0.631058053 |
| ENSMUSG0000002408 | chr17 | Cebpz | 0.115950795 | 0.631058053 |
| ENSMUSG0000000320 | chr17 | Sh3gl1 | 0.124705499 | 0.631058053 |
| ENSMUSG0000003422 | chr1 | Gpc1 | 0.14372174 | 0.631058053 |
| ENSMUSG0000002750 | chr2 | Rtf2 | 0.145534107 | 0.631058053 |
| ENSMUSG0000002464 | chr18 | Timm21 | 0.151564108 | 0.631058053 |
| ENSMUSG0000003207 | chr9 | Zpr1 | 0.175907371 | 0.631058053 |
| ENSMUSG0000001449 | chr14 | Ankrd28 | 0.172972198 | 0.631058053 |
| ENSMUSG0000002946 | chr5 | Fam216a | 0.185342068 | 0.631058053 |
| ENSMUSG0000003777 | chr7 | Mrpl23 | 0.218103251 | 0.631058053 |
| ENSMUSG0000000206 | chr7 | Ccne1 | 0.332047297 | 0.631058053 |
| ENSMUSG0000002033 | chr10 | Hcn2 | -0.246262093 | 0.631355345 |
| ENSMUSG0000007262 | chr11 | Slfn2 | -0.348651626 | 0.631355345 |
| ENSMUSG0000002951 | chr5 | Rab35 | 0.115127217 | 0.631355345 |
| ENSMUSG0000004078 | chr1 | Cop1 | 0.163631894 | 0.631355345 |
| ENSMUSG0000006839 | chr15 | Chrac1 | 0.194830864 | 0.631355345 |
| ENSMUSG0000002219 | chr15 | Drosha | 0.111025565 | 0.631399367 |
| ENSMUSG0000000055 | chr15 | Zfp385a | 0.154024755 | 0.631935727 |
| ENSMUSG0000003096 | chr7 | Abraxas2 | 0.126548631 | 0.632004641 |
| ENSMUSG0000002894 | chr4 | Nol9 | 0.142522292 | 0.632004641 |
| ENSMUSG0000002528 | chrX | Prdx4 | 0.19928956 | 0.632004641 |
| ENSMUSG0000003419 | chr6 | Lsm3 | 0.226859622 | 0.632004641 |

| | | | | |
|---|---|---|---|---|
| ENSMUSG0000003490 | chr2 | Cobll1 | 0.179234606 | 0.632160638 |
| ENSMUSG0000002501 | chr19 | Pik3ap1 | -0.383203679 | 0.632231653 |
| ENSMUSG0000005346 | chr15 | Tg | 0.841979052 | 0.632277134 |
| ENSMUSG0000004359 | chr17 | Unc5cl | -0.91716306 | 0.632373974 |
| ENSMUSG0000003500 | chr11 | Rundc1 | -0.160488637 | 0.632373974 |
| ENSMUSG0000002869 | chr4 | Akr1a1 | 0.195079568 | 0.632373974 |
| ENSMUSG0000002092 | chr11 | Adam11 | -0.276944639 | 0.633333616 |
| ENSMUSG0000002326 | chr9 | Acy1 | -0.234117382 | 0.633443488 |
| ENSMUSG0000002187 | chr14 | Arf4 | 0.203585403 | 0.633720702 |
| ENSMUSG0000001977 | chr10 | Frk | -0.206058639 | 0.633864133 |
| ENSMUSG0000002788 | chr3 | Stxbp3 | 0.121837156 | 0.633864133 |
| ENSMUSG0000003420 | chr11 | Gas2l1 | 0.16170462 | 0.633864133 |
| ENSMUSG0000004761 | chr4 | A430005L14Rik | 0.270678224 | 0.633864133 |
| ENSMUSG0000001561 | chr2 | Gata3 | -0.976231713 | 0.634004704 |
| ENSMUSG0000002949 | chr5 | Slc49a3 | -0.261987184 | 0.634004704 |
| ENSMUSG0000002499 | chr19 | Noc3l | 0.119624808 | 0.634004704 |
| ENSMUSG0000001985 | chr10 | Tnfaip3 | -0.266715191 | 0.635228588 |
| ENSMUSG0000003431 | chr6 | Iqsec1 | 0.120373717 | 0.635228588 |
| ENSMUSG0000003935 | chr1 | Smarcal1 | 0.116190114 | 0.635228588 |
| ENSMUSG0000003447 | chr16 | Sec22a | 0.163942643 | 0.635228588 |
| ENSMUSG0000000898 | chr5 | Rasgef1b | -0.317625871 | 0.63523574 |
| ENSMUSG0000002443 | chr17 | Mrps18b | 0.230513987 | 0.635377274 |
| ENSMUSG0000002547 | chr7 | Adgra1 | -0.337496054 | 0.635576768 |
| ENSMUSG0000000523 | chr11 | Dnah2 | -0.3077032 | 0.635576768 |
| ENSMUSG0000003209 | chr9 | Arcn1 | 0.121837824 | 0.635576768 |
| ENSMUSG0000003717 | chr3 | Elf2 | 0.117214923 | 0.635576768 |
| ENSMUSG0000004875 | chr10 | Foxo3 | 0.176575503 | 0.635576768 |
| ENSMUSG0000002513 | chr11 | Sirt7 | -0.168165559 | 0.635889966 |
| ENSMUSG0000004122 | chr18 | Arhgap12 | -0.148138516 | 0.636090136 |
| ENSMUSG0000010275 | chr1 | Gm7694 | 0.157502699 | 0.636090136 |
| ENSMUSG0000003188 | chr8 | Phaf1 | 0.134918516 | 0.636328395 |
| ENSMUSG0000003342 | chr4 | Eri3 | 0.142720644 | 0.636509785 |
| ENSMUSG0000004899 | chr3 | Atxn7l2 | -0.239826679 | 0.636624524 |
| ENSMUSG0000004244 | chr6 | Mios | 0.146302298 | 0.636624524 |
| ENSMUSG0000007815 | chr11 | Psme2b | 0.310427478 | 0.636624524 |
| ENSMUSG0000004712 | chr11 | Cltc | -0.093974501 | 0.636948561 |
| ENSMUSG0000002948 | chr5 | Aacs | -0.276559339 | 0.637544638 |
| ENSMUSG0000002185 | chr14 | Ccdc198 | -1.020688629 | 0.637843627 |
| ENSMUSG0000002504 | chr19 | Nt5c2 | 0.156767785 | 0.637887199 |
| ENSMUSG0000000649 | chr10 | Ptbp1 | 0.07876618 | 0.63797062 |
| ENSMUSG0000003045 | chr7 | Nipa2 | 0.143839791 | 0.63797062 |
| ENSMUSG0000003854 | chr3 | BC028528 | 0.414417285 | 0.63797062 |
| ENSMUSG0000006380 | chr7 | Hspbp1 | 0.230541076 | 0.638137178 |
| ENSMUSG0000004434 | chr2 | Snhg11 | -0.256038354 | 0.638485275 |
| ENSMUSG0000003150 | chr8 | Rab20 | -0.486234574 | 0.638485275 |
| ENSMUSG0000002272 | chr16 | Arl6 | -0.133645234 | 0.638485275 |
| ENSMUSG0000002103 | chr12 | Vipas39 | 0.130618519 | 0.638485275 |
| ENSMUSG0000002460 | chr18 | Rbm22 | 0.132427271 | 0.638485275 |
| ENSMUSG0000002042 | chr11 | Castor1 | 0.240015917 | 0.638485275 |
| ENSMUSG0000003086 | chr7 | Ndufab1 | 0.238854061 | 0.638485275 |

| | | | | |
|---|---|---|---|---|
| ENSMUSG0000004500 | chr1 | Fzd5 | -0.226433761 | 0.638591813 |
| ENSMUSG0000002177 | chr14 | Vdac2 | 0.173825188 | 0.638707056 |
| ENSMUSG0000004843 | chr5 | Nup42 | 0.196568633 | 0.638771215 |
| ENSMUSG0000002892 | chr4 | Necap2 | 0.144920522 | 0.638963651 |
| ENSMUSG0000002608 | chr1 | Eif5b | 0.144511979 | 0.639178134 |
| ENSMUSG0000002205 | chr14 | Ebf2 | 0.452455118 | 0.639178134 |
| ENSMUSG0000002874 | chr4 | Capzb | 0.200583177 | 0.639396579 |
| ENSMUSG0000003700 | chrX | Xpnpep2 | -0.643913768 | 0.639589839 |
| ENSMUSG0000003042 | chr7 | Ttyh1 | -0.227902778 | 0.639589839 |
| ENSMUSG0000002691 | chr2 | Strbp | -0.171269207 | 0.639693284 |
| ENSMUSG0000002808 | chr3 | Rps3a1 | 0.307524737 | 0.639693284 |
| ENSMUSG0000007530 | chr2 | Erich2 | -0.432595396 | 0.639823873 |
| ENSMUSG0000000327 | chr7 | Car11 | -0.246875347 | 0.640184614 |
| ENSMUSG0000000619 | chr13 | Cdkal1 | 0.210075821 | 0.640184614 |
| ENSMUSG0000002196 | chr14 | Mrpl57 | 0.235561307 | 0.640184614 |
| ENSMUSG0000004423 | chr13 | Nhlrc1 | 0.322687075 | 0.640184614 |
| ENSMUSG0000002066 | chr12 | Dnmt3a | -0.190191252 | 0.640398119 |
| ENSMUSG0000002946 | chr5 | Atp2a2 | 0.115744619 | 0.640398119 |
| ENSMUSG0000007270 | chr6 | Smim10l1 | 0.160620573 | 0.640398119 |
| ENSMUSG0000007424 | chr8 | Dda1 | 0.162966221 | 0.640398119 |
| ENSMUSG0000006739 | chr9 | Trim43c | 0.502032456 | 0.640398119 |
| ENSMUSG0000007415 | chr8 | Nlrc5 | -0.503829956 | 0.640776499 |
| ENSMUSG0000002306 | chr10 | Nus1 | -0.108771143 | 0.640776499 |
| ENSMUSG0000003922 | chr3 | Rpl22l1 | 0.255668761 | 0.640776499 |
| ENSMUSG0000004889 | chr11 | Cdk5r1 | -0.262926038 | 0.640820518 |
| ENSMUSG0000005985 | chr7 | Kmt5c | -0.206425822 | 0.640820518 |
| ENSMUSG0000002507 | chr19 | Casp7 | -0.25045935 | 0.640820518 |
| ENSMUSG0000006201 | chr14 | Gmfb | -0.142679559 | 0.640820518 |
| ENSMUSG0000002817 | chr3 | Wls | 0.139723966 | 0.640820518 |
| ENSMUSG0000002647 | chr1 | Stx6 | 0.13669132 | 0.640820518 |
| ENSMUSG0000000845 | chr8 | Nutf2 | 0.1709364 | 0.640820518 |
| ENSMUSG0000002414 | chr17 | Mlst8 | -0.160205509 | 0.641076862 |
| ENSMUSG0000003522 | chr2 | Rims4 | 0.343630483 | 0.641076862 |
| ENSMUSG0000005535 | chrX | Capza1b | 0.840352761 | 0.641367797 |
| ENSMUSG0000004183 | chr9 | Pdcd7 | -0.179608375 | 0.641460925 |
| ENSMUSG0000003771 | chr10 | Cisd1 | 0.242255329 | 0.64156593 |
| ENSMUSG0000007497 | chr2 | Fibin | -0.200673759 | 0.64169036 |
| ENSMUSG0000005877 | chr13 | H1f5 | -0.647017775 | 0.64169036 |
| ENSMUSG0000006656 | chr7 | Lsm14a | 0.139914274 | 0.64169036 |
| ENSMUSG0000004238 | chr6 | Tsen2 | 0.165370861 | 0.64169036 |
| ENSMUSG0000003539 | chr2 | Oser1 | 0.16003425 | 0.64169036 |
| ENSMUSG0000004494 | chr19 | Cfap43 | 0.280347968 | 0.64169036 |
| ENSMUSG0000004012 | chr4 | Pnrc1 | 0.196125556 | 0.64169036 |
| ENSMUSG0000000377 | chr18 | Kif20a | 0.418585465 | 0.64169036 |
| ENSMUSG0000007546 | chr2 | Dnlz | 0.186371422 | 0.64213548 |
| ENSMUSG0000002929 | chr5 | Abcg3 | -0.618748496 | 0.642219867 |
| ENSMUSG0000005696 | chr5 | Gjc3 | -0.323875842 | 0.642431427 |
| ENSMUSG0000002593 | chr1 | Lactb2 | 0.145949076 | 0.642693475 |
| ENSMUSG0000004695 | chr5 | Slc26a1 | -0.509491751 | 0.642796121 |
| ENSMUSG0000009353 | chr7 | Smim17 | -0.389945423 | 0.642796121 |

| | | | | |
|---|---|---|---|---|
| ENSMUSG0000001865 | chr11 | Pnpo | -0.26414908 | 0.642796121 |
| ENSMUSG0000009010 | chr2 | Ttbk2 | -0.175325623 | 0.642796121 |
| ENSMUSG0000003286 | chr2 | Psmf1 | 0.125821119 | 0.642796121 |
| ENSMUSG0000004596 | chr8 | Ing1 | 0.138194237 | 0.642796121 |
| ENSMUSG0000002893 | chr5 | Psmc2 | 0.164059976 | 0.642796121 |
| ENSMUSG0000000343 | chr7 | Paf1 | 0.169786847 | 0.642796121 |
| ENSMUSG0000000968 | chr7 | Fxyd5 | 0.182215004 | 0.642796121 |
| ENSMUSG0000003758 | chr14 | Gch1 | 0.378012907 | 0.642946399 |
| ENSMUSG0000002119 | chr12 | Golga5 | 0.120585517 | 0.642981208 |
| ENSMUSG0000002144 | chr13 | Shc3 | -0.385450129 | 0.643198473 |
| ENSMUSG0000002967 | chr6 | Wnt16 | 0.836703416 | 0.643245969 |
| ENSMUSG0000006336 | chr2 | 3300002I08Rik | -0.483736327 | 0.643332665 |
| ENSMUSG0000004828 | chr13 | Zfp738 | -0.139585619 | 0.643332665 |
| ENSMUSG0000002793 | chr3 | Slc27a3 | 0.331824384 | 0.643332665 |
| ENSMUSG0000003673 | chr2 | Cysrt1 | -0.596535522 | 0.643435105 |
| ENSMUSG0000002464 | chr18 | Cyb5a | 0.167571189 | 0.643473909 |
| ENSMUSG0000002845 | chr4 | Vcp | 0.108007809 | 0.643543209 |
| ENSMUSG0000004098 | chr7 | Mill2 | -0.137904925 | 0.643689099 |
| ENSMUSG0000002300 | chr15 | Fmnl3 | 0.229253377 | 0.643857578 |
| ENSMUSG0000000350 | chr6 | Impdh1 | -0.156039397 | 0.643936026 |
| ENSMUSG0000005145 | chr7 | Crebzf | -0.150111885 | 0.643943804 |
| ENSMUSG0000005676 | chr1 | Cspp1 | -0.139003537 | 0.644373573 |
| ENSMUSG0000005458 | chr2 | Pla2r1 | 0.181401023 | 0.644373573 |
| ENSMUSG0000002687 | chr2 | Traf1 | -0.268822191 | 0.644843231 |
| ENSMUSG0000003426 | chr17 | Zdhhc14 | 0.182559718 | 0.644843231 |
| ENSMUSG0000005181 | chr2 | Sox12 | -0.215193245 | 0.645151135 |
| ENSMUSG0000005752 | chr11 | Spop | 0.135399868 | 0.645151135 |
| ENSMUSG0000000674 | chr18 | Kif5b | 0.154550894 | 0.645151135 |
| ENSMUSG0000004425 | chr18 | Osbpl1a | 0.147282452 | 0.645151135 |
| ENSMUSG0000002877 | chr4 | Tinagl1 | 0.237646309 | 0.645151135 |
| ENSMUSG0000003183 | chr8 | Ifi30 | -0.217086982 | 0.645770575 |
| ENSMUSG0000003303 | chr19 | Calhm2 | 0.20305549 | 0.645770575 |
| ENSMUSG0000002668 | chr1 | Aldh9a1 | 0.180956069 | 0.645770575 |
| ENSMUSG0000000227 | chr17 | Lmf1 | 0.155400095 | 0.645916694 |
| ENSMUSG0000003991 | chr5 | Rhbdd2 | -0.173214246 | 0.645968741 |
| ENSMUSG0000005989 | chr15 | Ptp4a3 | 0.23718394 | 0.646402861 |
| ENSMUSG0000006428 | chr5 | Ccdc146 | -0.379194537 | 0.646427658 |
| ENSMUSG0000000128 | chr15 | Sp1 | -0.1607031 | 0.646427658 |
| ENSMUSG0000004734 | chr11 | Zfp286 | 0.148145338 | 0.646427658 |
| ENSMUSG0000003255 | chr9 | Uba5 | 0.181077895 | 0.646427658 |
| ENSMUSG0000004414 | chrX | 1810030O07Rik | 0.115793693 | 0.646892912 |
| ENSMUSG0000003098 | chr7 | Bccip | 0.217635974 | 0.646892912 |
| ENSMUSG0000002894 | chr4 | Espn | -0.611783363 | 0.647548762 |
| ENSMUSG0000002537 | chr10 | Rnf41 | -0.097809789 | 0.647548762 |
| ENSMUSG0000000008 | chr4 | Scmh1 | 0.153530345 | 0.647548762 |
| ENSMUSG0000003535 | chr10 | Nup37 | 0.253025166 | 0.647548762 |
| ENSMUSG0000002391 | chr17 | Cenpq | 0.236844389 | 0.647548762 |
| ENSMUSG0000002076 | chr11 | Galk1 | 0.247507583 | 0.647548762 |
| ENSMUSG0000004229 | chr9 | Gm5617 | 0.475705705 | 0.647548762 |
| ENSMUSG0000000220 | chr7 | Napsa | -0.592775715 | 0.647824487 |

| | | | | |
|---|---|---|---|---|
| ENSMUSG0000004370 | chr14 | Pde12 | 0.146550899 | 0.647824487 |
| ENSMUSG0000004018 | chr9 | Scamp2 | -0.207715804 | 0.647836771 |
| ENSMUSG0000003849 | chr3 | Otud7b | 0.114110712 | 0.647836771 |
| ENSMUSG0000003373 | chr6 | Spr | 0.284059976 | 0.647836771 |
| ENSMUSG0000002069 | chr11 | Tlk2 | 0.144091568 | 0.648012406 |
| ENSMUSG0000002299 | chr15 | Lmbr1l | -0.213861597 | 0.648080095 |
| ENSMUSG0000006342 | chr10 | Ddo | 0.153319168 | 0.648122447 |
| ENSMUSG0000007165 | chr19 | Ganab | -0.11393887 | 0.648233038 |
| ENSMUSG0000003184 | chr8 | Lsm4 | 0.188522784 | 0.648233038 |
| ENSMUSG0000000631 | chr7 | Tmem147 | 0.214754185 | 0.648233038 |
| ENSMUSG0000004254 | chr6 | Sem1 | 0.214092464 | 0.64825135 |
| ENSMUSG0000003354 | chr1 | Angptl1 | 0.207180378 | 0.64825135 |
| ENSMUSG0000004026 | chr8 | Klhdc4 | 0.203643035 | 0.648261404 |
| ENSMUSG0000000550 | chr2 | Celf1 | 0.113328141 | 0.648294869 |
| ENSMUSG0000000183 | chr9 | Septin7 | 0.163565821 | 0.648294869 |
| ENSMUSG0000003071 | chr7 | Ppme1 | 0.194116234 | 0.648294869 |
| ENSMUSG0000000012 | chr17 | Fer | 0.190783954 | 0.648294869 |
| ENSMUSG0000000962 | chr8 | Tex15 | 0.381762912 | 0.648294869 |
| ENSMUSG0000005193 | chr15 | Spats2 | 0.138133124 | 0.648316052 |
| ENSMUSG0000001982 | chr10 | Utrn | 0.160106209 | 0.648316052 |
| ENSMUSG0000004847 | chr8 | Spata33 | -0.234648724 | 0.648915033 |
| ENSMUSG0000000203 | chr9 | Tmem25 | -0.198057983 | 0.648942047 |
| ENSMUSG0000007877 | chr7 | Zfp59 | 0.184428463 | 0.648942047 |
| ENSMUSG0000004458 | chrX | Tlr7 | 0.334676803 | 0.648942047 |
| ENSMUSG0000004532 | chr3 | Fndc7 | 1.135661502 | 0.648942047 |
| ENSMUSG0000000375 | chr7 | Itpkc | -0.280670312 | 0.648970828 |
| ENSMUSG0000004882 | chr8 | Pkd1l3 | -0.481062857 | 0.648970828 |
| ENSMUSG0000002102 | chr12 | Ralgapa1 | -0.129287527 | 0.648970828 |
| ENSMUSG0000006341 | chr14 | Stk24 | 0.11882034 | 0.648970828 |
| ENSMUSG0000005471 | chr8 | Hmgb2 | -0.167717982 | 0.649216894 |
| ENSMUSG0000003297 | chr10 | Slx9 | 0.165579298 | 0.649567761 |
| ENSMUSG0000004802 | chr6 | Tmem229a | -0.258619896 | 0.649580289 |
| ENSMUSG0000000467 | chr8 | Myo9b | 0.169161154 | 0.649580289 |
| ENSMUSG0000006205 | chr12 | Iah1 | 0.198967225 | 0.649580289 |
| ENSMUSG0000003486 | chr7 | Ankrd27 | 0.090341973 | 0.64971006 |
| ENSMUSG0000003284 | chr17 | Abcc10 | -0.294132348 | 0.650060065 |
| ENSMUSG0000003722 | chr5 | Zfyve28 | -0.242205862 | 0.650060065 |
| ENSMUSG0000003494 | chr11 | Tmem106a | -0.161455319 | 0.650060065 |
| ENSMUSG0000002048 | chr11 | Xbp1 | 0.287998853 | 0.650060065 |
| ENSMUSG0000005094 | chr11 | Efcab5 | -0.588526888 | 0.650577568 |
| ENSMUSG0000007457 | chr2 | Mocs3 | 0.211453579 | 0.650577568 |
| ENSMUSG0000000556 | chr7 | Trim28 | 0.151841418 | 0.650763672 |
| ENSMUSG0000002183 | chr14 | Bmp4 | 0.20297323 | 0.650951011 |
| ENSMUSG0000006298 | chr6 | Cped1 | 0.175184529 | 0.651009649 |
| ENSMUSG0000003861 | chr3 | Ensa | 0.1361113 | 0.651162047 |
| ENSMUSG0000003361 | chr19 | Pank1 | -0.220069275 | 0.651192341 |
| ENSMUSG0000003818 | chr2 | Spag4 | -0.438191058 | 0.651442764 |
| ENSMUSG0000000420 | chr10 | Psap | 0.094088483 | 0.651457703 |
| ENSMUSG0000004222 | chr4 | Lyn | -0.2491707 | 0.651582832 |
| ENSMUSG0000005291 | chr16 | Senp7 | -0.167852692 | 0.651582832 |

| | | | | |
|---|---|---|---|---|
| ENSMUSG0000002798 | chr3 | Hadh | -0.204481855 | 0.652021636 |
| ENSMUSG0000002439 | chr17 | Bag6 | 0.098194373 | 0.652021636 |
| ENSMUSG0000003853 | chr1 | Rgs4 | -0.344102035 | 0.652044725 |
| ENSMUSG0000004903 | chr12 | Tmem121 | -0.303932627 | 0.652044725 |
| ENSMUSG0000000600 | chr1 | Tpr | 0.120752594 | 0.652044725 |
| ENSMUSG0000002315 | chr14 | Rpp14 | 0.155135171 | 0.652044725 |
| ENSMUSG0000000803 | chr7 | Ap2s1 | 0.196589307 | 0.652044725 |
| ENSMUSG0000000466 | chr10 | Polr2e | 0.188241382 | 0.652044725 |
| ENSMUSG0000003295 | chr5 | Pebp1 | 0.25707358 | 0.652044725 |
| ENSMUSG0000001838 | chr11 | Shroom1 | 0.41170931 | 0.652177024 |
| ENSMUSG0000002737 | chr2 | Nphp1 | 0.157692132 | 0.652218076 |
| ENSMUSG0000002943 | chr5 | Ran | 0.245354627 | 0.652316958 |
| ENSMUSG0000003821 | chr6 | Tapbpl | -0.176246908 | 0.652896283 |
| ENSMUSG0000006098 | chr2 | Galnt13 | -0.252114913 | 0.653330412 |
| ENSMUSG0000002593 | chr1 | Eya1 | 0.380256518 | 0.653458141 |
| ENSMUSG0000007823 | chr4 | Klhdc7a | -0.642723578 | 0.653912779 |
| ENSMUSG0000004790 | chr2 | Tshz2 | 0.127692361 | 0.653912779 |
| ENSMUSG0000004826 | chr13 | Dip2c | 0.159551079 | 0.653912779 |
| ENSMUSG0000003840 | chr13 | Foxf2 | 0.203987902 | 0.653912779 |
| ENSMUSG0000004446 | chr17 | Tnfaip8l1 | -0.315465005 | 0.653980119 |
| ENSMUSG0000002005 | chr10 | Washc3 | 0.224707841 | 0.653980119 |
| ENSMUSG0000002658 | chr1 | Astn1 | -0.237733646 | 0.654073311 |
| ENSMUSG0000004509 | chr18 | Arhgef37 | -0.287632816 | 0.654231893 |
| ENSMUSG0000002848 | chr4 | Psip1 | -0.098832167 | 0.654365411 |
| ENSMUSG0000004002 | chr6 | Ipo8 | -0.112697747 | 0.654564454 |
| ENSMUSG0000005134 | chr1 | Plekhm3 | -0.146320599 | 0.654591786 |
| ENSMUSG0000002457 | chr18 | Naa12 | 0.146868747 | 0.654591786 |
| ENSMUSG0000004005 | chr10 | Baz2a | -0.201458135 | 0.65480475 |
| ENSMUSG0000006991 | chr11 | Insyn2b | -0.358177544 | 0.654966497 |
| ENSMUSG0000001978 | chr10 | Hey2 | -0.21174531 | 0.654966497 |
| ENSMUSG0000003885 | chr1 | Itpkb | 0.131762306 | 0.654966497 |
| ENSMUSG0000002466 | chr19 | Sdhaf2 | 0.167381392 | 0.654966497 |
| ENSMUSG0000005091 | chr13 | Zfp366 | 0.203708074 | 0.654966497 |
| ENSMUSG0000003888 | chr17 | Mrps34 | 0.209544357 | 0.654966497 |
| ENSMUSG0000009637 | chr19 | Gm21992 | 0.357792745 | 0.654966497 |
| ENSMUSG0000002807 | chr3 | Naxe | 0.187168182 | 0.655343806 |
| ENSMUSG0000002455 | chr18 | Me2 | -0.219490014 | 0.655463738 |
| ENSMUSG0000001886 | chr11 | Fdxr | -0.213481686 | 0.655466663 |
| ENSMUSG0000003411 | chr5 | Tpst1 | 0.172195688 | 0.655623374 |
| ENSMUSG0000007445 | chr3 | S100a16 | 0.2531821 | 0.655623374 |
| ENSMUSG0000000741 | chr5 | Gatad1 | 0.144915163 | 0.656006771 |
| ENSMUSG0000007107 | chr17 | Lrrc73 | -0.311395473 | 0.656092735 |
| ENSMUSG0000003085 | chr7 | Tacc2 | 0.200844335 | 0.656092735 |
| ENSMUSG0000007878 | chr3 | Rbis | 0.185975818 | 0.656092735 |
| ENSMUSG0000003115 | chrX | Pqbp1 | 0.188727746 | 0.656651393 |
| ENSMUSG0000000470 | chr1 | Cd244a | -1.003729809 | 0.657039072 |
| ENSMUSG0000005082 | chr16 | Fam131a | -0.151784163 | 0.657039072 |
| ENSMUSG0000002018 | chr10 | Snrpd3 | 0.188710605 | 0.657039072 |
| ENSMUSG0000004267 | chr1 | Rgs8 | -0.316893052 | 0.657087313 |
| ENSMUSG0000004686 | chr7 | Fbl | 0.163918348 | 0.657212114 |

| | | | | |
|---|---|---|---|---|
| ENSMUSG0000005757 | chr4 | Zbtb8os | 0.194235883 | 0.65733431 |
| ENSMUSG0000004406 | chr12 | Gpr22 | -0.186041591 | 0.657644504 |
| ENSMUSG0000004919 | chrX | Rtl5 | -0.18015993 | 0.657644504 |
| ENSMUSG0000002445 | chr17 | Trim26 | -0.114398127 | 0.657644504 |
| ENSMUSG0000002090 | chr11 | Ntn1 | 0.118810077 | 0.657644504 |
| ENSMUSG0000002905 | chr4 | Ski | 0.119917552 | 0.657644504 |
| ENSMUSG0000003755 | chr7 | Plekhg2 | 0.165586773 | 0.657801085 |
| ENSMUSG0000004968 | chr15 | Pheta2 | -0.327866827 | 0.657915829 |
| ENSMUSG0000002839 | chr4 | Wdr31 | -0.273373431 | 0.658043953 |
| ENSMUSG0000004281 | chr5 | Flt3 | -0.720494702 | 0.658043953 |
| ENSMUSG0000004151 | chr8 | Irf8 | -0.225833864 | 0.658043953 |
| ENSMUSG0000007163 | chr16 | 2510002D24Rik | 0.186189468 | 0.658043953 |
| ENSMUSG0000002159 | chr13 | Slf1 | -0.189344746 | 0.658291719 |
| ENSMUSG0000002380 | chr17 | Tiam2 | 0.236843744 | 0.658291719 |
| ENSMUSG0000002495 | chr19 | Gpr137 | -0.167888579 | 0.658601188 |
| ENSMUSG0000003028 | chr6 | Emc3 | 0.133800861 | 0.65869976 |
| ENSMUSG0000004330 | chr16 | Kcnj6 | 0.390692821 | 0.65869976 |
| ENSMUSG0000002911 | chr5 | Nelfa | 0.116788439 | 0.659077607 |
| ENSMUSG0000002853 | chr4 | Cachd1 | -0.174206746 | 0.659189955 |
| ENSMUSG0000004220 | chr4 | Slc35e2 | -0.121191577 | 0.659189955 |
| ENSMUSG0000004818 | chr2 | Bend7 | 0.170635039 | 0.659189955 |
| ENSMUSG0000001553 | chr13 | Itga2 | -0.269471109 | 0.659351903 |
| ENSMUSG0000002394 | chr17 | Sgo1 | -0.564086135 | 0.659351903 |
| ENSMUSG0000002061 | chr11 | Gna13 | 0.114904098 | 0.659351903 |
| ENSMUSG0000002289 | chr16 | Vps26c | 0.169546668 | 0.659351903 |
| ENSMUSG0000002725 | chr2 | Pacsin3 | 0.157044123 | 0.659351903 |
| ENSMUSG0000002789 | chr3 | Kcnc4 | 0.235267672 | 0.659351903 |
| ENSMUSG0000005309 | chr14 | Myh7 | 0.665931357 | 0.659351903 |
| ENSMUSG0000004490 | chr7 | Psg22 | -1.151507905 | 0.659737389 |
| ENSMUSG0000002538 | chr11 | Faap100 | -0.14288056 | 0.659790941 |
| ENSMUSG0000002149 | chr13 | Fam193b | -0.158933453 | 0.660224072 |
| ENSMUSG0000004121 | chr2 | Arhgap11a | -0.493215948 | 0.661332243 |
| ENSMUSG0000003180 | chr8 | Slc27a1 | -0.237391281 | 0.661332243 |
| ENSMUSG0000003121 | chrX | Efnb1 | -0.259111596 | 0.661332243 |
| ENSMUSG0000003915 | chr4 | Akna | -0.225527534 | 0.661332243 |
| ENSMUSG0000002257 | chr15 | Gsdmd | -0.149930626 | 0.661332243 |
| ENSMUSG0000005037 | chr2 | Snx21 | 0.116404107 | 0.661332243 |
| ENSMUSG0000002684 | chr2 | Tor1b | 0.115210591 | 0.661332243 |
| ENSMUSG0000002296 | chr16 | Ifnar1 | 0.124566487 | 0.661332243 |
| ENSMUSG0000002086 | chr11 | Ankrd40 | 0.129904618 | 0.661332243 |
| ENSMUSG0000001349 | chr5 | Tmem175 | 0.13800572 | 0.661332243 |
| ENSMUSG0000000765 | chr2 | Bcl2l1 | 0.14622155 | 0.661332243 |
| ENSMUSG0000003348 | chr2 | Catsper2 | 0.3553817 | 0.661332243 |
| ENSMUSG0000002860 | chr4 | Magoh | 0.191763043 | 0.661332243 |
| ENSMUSG0000004242 | chr13 | Dhx29 | 0.19936313 | 0.661332243 |
| ENSMUSG0000006852 | chr3 | Gng5 | 0.252189443 | 0.661332243 |
| ENSMUSG0000005165 | chr11 | B3gnt2 | 0.282431433 | 0.661332243 |
| ENSMUSG0000003071 | chr7 | Nupr1 | 0.324018582 | 0.661332243 |
| ENSMUSG0000002615 | chr1 | Fam135a | -0.189677534 | 0.661350894 |
| ENSMUSG0000004541 | chr7 | 2410002F23Rik | -0.165567678 | 0.661350894 |

| | | | | |
|---|---|---|---|---|
| ENSMUSG0000004678 | chr9 | Epm2aip1 | -0.144875226 | 0.661350894 |
| ENSMUSG0000000167 | chr1 | Ddx18 | 0.158905429 | 0.661350894 |
| ENSMUSG0000002235 | chr15 | Ntaq1 | 0.229665386 | 0.661350894 |
| ENSMUSG0000005707 | chr1 | Spata45 | -0.785144894 | 0.661379006 |
| ENSMUSG0000003426 | chr6 | Setd5 | -0.169346745 | 0.661379006 |
| ENSMUSG0000005185 | chrX | Rtl8c | -0.131314917 | 0.661379006 |
| ENSMUSG0000000438 | chr8 | Large1 | 0.132444805 | 0.661379006 |
| ENSMUSG0000002810 | chr3 | Tars2 | 0.133606626 | 0.661379006 |
| ENSMUSG0000002869 | chr4 | Pik3r3 | 0.198238468 | 0.661379006 |
| ENSMUSG0000003196 | chr9 | Bmper | 0.271190221 | 0.661379006 |
| ENSMUSG0000005123 | chr9 | Swsap1 | -0.19490794 | 0.661899584 |
| ENSMUSG0000002029 | chr11 | Nsg2 | -0.151093603 | 0.661899584 |
| ENSMUSG0000002719 | chr2 | Api5 | 0.11690961 | 0.661899584 |
| ENSMUSG0000004072 | chr4 | Virma | 0.123192864 | 0.661899584 |
| ENSMUSG0000003066 | chr7 | Calca | -0.758150771 | 0.662930103 |
| ENSMUSG0000007530 | chr2 | Klhl41 | 1.051770853 | 0.662988707 |
| ENSMUSG0000003694 | chr9 | Rab8b | -0.121498182 | 0.66299637 |
| ENSMUSG0000002736 | chr2 | Gabpb1 | 0.119833581 | 0.66299637 |
| ENSMUSG0000003806 | chr6 | Mturn | 0.134794194 | 0.66299637 |
| ENSMUSG0000003024 | chr6 | Golt1b | 0.187687586 | 0.663440585 |
| ENSMUSG0000003676 | chrX | Wdr44 | -0.142911379 | 0.663885012 |
| ENSMUSG0000004702 | chr1 | Cfap65 | -0.462153916 | 0.66389413 |
| ENSMUSG0000002157 | chr13 | Ahrr | -0.468247416 | 0.66389413 |
| ENSMUSG0000003970 | chr15 | Lmbrd2 | -0.172400826 | 0.66389413 |
| ENSMUSG0000003109 | chr7 | Nadsyn1 | -0.233798843 | 0.66389413 |
| ENSMUSG0000010750 | chr9 | Gm35549 | -0.584216487 | 0.66389413 |
| ENSMUSG0000002181 | chr14 | Ppp3cb | 0.093575124 | 0.66389413 |
| ENSMUSG0000003239 | chr9 | Dpp8 | 0.108361545 | 0.66389413 |
| ENSMUSG0000000335 | chr15 | Cacnb3 | 0.104969361 | 0.66389413 |
| ENSMUSG0000002058 | chr12 | Laptm4a | 0.161589205 | 0.66389413 |
| ENSMUSG0000000830 | chr18 | Phax | 0.136384889 | 0.66389413 |
| ENSMUSG0000003524 | chr6 | Eogt | 0.132519202 | 0.66389413 |
| ENSMUSG0000002920 | chr5 | Ube2k | 0.146398658 | 0.66389413 |
| ENSMUSG0000004358 | chr9 | Pxylp1 | 0.167952933 | 0.66389413 |
| ENSMUSG0000002572 | chr7 | Wdr73 | 0.167418988 | 0.66389413 |
| ENSMUSG0000004557 | chr3 | St7l | 0.178149361 | 0.66389413 |
| ENSMUSG0000001484 | chr8 | Tppp3 | 0.274489892 | 0.66389413 |
| ENSMUSG0000000621 | chr4 | Fam131c | 0.607895248 | 0.66389413 |
| ENSMUSG0000003184 | chr8 | Bco1 | 0.566524892 | 0.66389413 |
| ENSMUSG0000004296 | chr15 | Egflam | -0.156288414 | 0.664282696 |
| ENSMUSG0000002678 | chr1 | Abi2 | -0.099939492 | 0.664358144 |
| ENSMUSG0000004116 | chr4 | Otud3 | 0.186069192 | 0.664358144 |
| ENSMUSG0000006326 | chr15 | Syt10 | 0.423977874 | 0.664648845 |
| ENSMUSG0000003557 | chr16 | Iqcg | -0.246505706 | 0.665001728 |
| ENSMUSG0000002130 | chr13 | Gpr137b | -0.179046449 | 0.665001728 |
| ENSMUSG0000003317 | chr16 | Tmprss7 | -0.80231862 | 0.665001728 |
| ENSMUSG0000002536 | chr10 | Rps26 | 0.192819074 | 0.665001728 |
| ENSMUSG0000004679 | chr12 | Riox1 | 0.307115451 | 0.665001728 |
| ENSMUSG0000005717 | chr7 | Cfap119 | -0.228413496 | 0.665383628 |
| ENSMUSG0000000140 | chr2 | Ube2c | -0.735687065 | 0.665728889 |

| | | | | |
|---|---|---|---|---|
| ENSMUSG0000002096 | chr12 | Gtf2a1 | -0.185591151 | 0.665728889 |
| ENSMUSG0000000720 | chr5 | Stx1a | -0.174272581 | 0.665728889 |
| ENSMUSG0000000587 | chr18 | Reep5 | -0.136521691 | 0.665728889 |
| ENSMUSG0000002957 | chr5 | Ube3b | 0.131600977 | 0.665728889 |
| ENSMUSG0000005451 | chr11 | Zfp867 | 0.142158254 | 0.665728889 |
| ENSMUSG0000002838 | chr4 | Hsdl2 | 0.162431587 | 0.665728889 |
| ENSMUSG0000009008 | chr17 | Rnf8 | 0.193203958 | 0.665728889 |
| ENSMUSG0000002473 | chr19 | Ccdc86 | 0.193800919 | 0.665728889 |
| ENSMUSG0000002214 | chr15 | Gdnf | 0.209951618 | 0.665728889 |
| ENSMUSG0000007819 | chr1 | Rpl35rt | 0.286178168 | 0.665728889 |
| ENSMUSG0000003506 | chr14 | Xkr6 | -0.244912519 | 0.665924969 |
| ENSMUSG0000002464 | chr19 | Tle4 | -0.129650917 | 0.665924969 |
| ENSMUSG0000005010 | chr11 | Tmc8 | -0.465699616 | 0.665939769 |
| ENSMUSG0000011616 | chr15 | Pdxp | -0.32027432 | 0.665939769 |
| ENSMUSG0000003287 | chr7 | Arhgef17 | -0.119997272 | 0.665939769 |
| ENSMUSG0000001885 | chr11 | Mrpl58 | 0.193607077 | 0.665939769 |
| ENSMUSG0000000673 | chr10 | Mettl1 | 0.293346035 | 0.665939769 |
| ENSMUSG0000005276 | chr6 | Zfp212 | 0.148355374 | 0.666439939 |
| ENSMUSG0000002262 | chr15 | Acr | -0.663811756 | 0.666453452 |
| ENSMUSG0000002503 | chrX | Efhc2 | -0.474085653 | 0.666886811 |
| ENSMUSG0000002044 | chr11 | Rnf185 | 0.119061723 | 0.666886811 |
| ENSMUSG0000006428 | chr15 | Cdpf1 | 0.237563707 | 0.666886811 |
| ENSMUSG0000003142 | chrX | Morf4l2 | 0.155469947 | 0.667003525 |
| ENSMUSG0000005245 | chr8 | Get3 | 0.143766761 | 0.667003525 |
| ENSMUSG0000005445 | chr2 | Vapb | 0.135733901 | 0.667050677 |
| ENSMUSG0000003827 | chr6 | Iffo1 | -0.19376643 | 0.667130933 |
| ENSMUSG0000002611 | chr1 | Coa5 | -0.118441945 | 0.667130933 |
| ENSMUSG0000002557 | chr11 | Tk1 | 0.203181776 | 0.667130933 |
| ENSMUSG0000002867 | chr4 | Kif2c | -0.930520418 | 0.667481514 |
| ENSMUSG0000002543 | chr10 | Atp23 | -0.305736248 | 0.667613096 |
| ENSMUSG0000005225 | chr15 | Zfp622 | 0.138964008 | 0.667613096 |
| ENSMUSG0000006874 | chr3 | Psma5 | 0.183914263 | 0.667613096 |
| ENSMUSG0000007257 | chr14 | Tmem253 | -0.39196368 | 0.667689965 |
| ENSMUSG0000003320 | chr10 | S100b | -0.141982337 | 0.667714715 |
| ENSMUSG0000002023 | chr10 | Ncln | 0.102801971 | 0.667714715 |
| ENSMUSG0000002524 | chr9 | Fyco1 | 0.149655715 | 0.667714715 |
| ENSMUSG0000002017 | chr10 | Frs2 | 0.174476066 | 0.667779485 |
| ENSMUSG0000003257 | chr9 | Cish | -0.419272186 | 0.667841397 |
| ENSMUSG0000006428 | chr13 | H4c12 | -0.711119932 | 0.667862968 |
| ENSMUSG0000000812 | chr8 | Brme1 | -0.544948461 | 0.667862968 |
| ENSMUSG0000003241 | chr9 | Rasa2 | -0.182741451 | 0.667862968 |
| ENSMUSG0000002577 | chr1 | Ly96 | -0.257204559 | 0.667862968 |
| ENSMUSG0000006984 | chr11 | Sco1 | -0.173369019 | 0.667862968 |
| ENSMUSG0000002976 | chr6 | Ccdc136 | -0.153245604 | 0.667862968 |
| ENSMUSG0000004360 | chr11 | Zfp3 | -0.187958659 | 0.667862968 |
| ENSMUSG0000004220 | chr11 | Sanbr | -0.143291114 | 0.667862968 |
| ENSMUSG0000005954 | chr2 | Tcea2 | -0.16635085 | 0.667862968 |
| ENSMUSG0000003332 | chr4 | Kdm4a | 0.096159919 | 0.667862968 |
| ENSMUSG0000002494 | chr19 | Ltbp3 | 0.128515541 | 0.667862968 |
| ENSMUSG0000004354 | chr3 | Zc2hc1a | 0.123052948 | 0.667862968 |

| | | | | |
|---|---|---|---|---|
| ENSMUSG0000000963 | chr8 | Ppp2cb | 0.166975891 | 0.667862968 |
| ENSMUSG0000002674 | chr2 | Nek6 | 0.165338923 | 0.667862968 |
| ENSMUSG0000000272 | chr2 | Naa20 | 0.176340759 | 0.667862968 |
| ENSMUSG0000002603 | chr1 | Nif3l1 | 0.188084502 | 0.667862968 |
| ENSMUSG0000002266 | chr16 | Ccdc80 | 0.191581674 | 0.667862968 |
| ENSMUSG0000002270 | chr16 | Qtrt2 | 0.192279564 | 0.667862968 |
| ENSMUSG0000001968 | chr6 | Fmc1 | 0.411957801 | 0.667862968 |
| ENSMUSG0000004784 | chr13 | Diras2 | 0.298427315 | 0.667862968 |
| ENSMUSG0000002849 | chr4 | Rps6 | 0.234000034 | 0.667973384 |
| ENSMUSG0000005813 | chr3 | Gstm1 | 0.185518955 | 0.668003692 |
| ENSMUSG0000005588 | chr12 | Fancm | -0.295602296 | 0.668083598 |
| ENSMUSG0000002466 | chr19 | Rab3il1 | -0.187330279 | 0.668692943 |
| ENSMUSG0000003235 | chr9 | Gclc | -0.22301212 | 0.669010049 |
| ENSMUSG0000003476 | chr4 | Glis1 | -0.854628049 | 0.669010049 |
| ENSMUSG0000006858 | chr2 | Zfyve19 | -0.143702739 | 0.669010049 |
| ENSMUSG0000002112 | chr12 | Zfp36l1 | 0.140771345 | 0.669010049 |
| ENSMUSG0000003278 | chr10 | Pdxk | -0.234196321 | 0.669135054 |
| ENSMUSG0000004525 | chr17 | Morn2 | -0.285140614 | 0.66954404 |
| ENSMUSG0000005844 | chr9 | Rpl10-ps3 | 0.325838827 | 0.66954404 |
| ENSMUSG0000009211 | chr7 | Fancf | -0.323710059 | 0.669982753 |
| ENSMUSG0000003327 | chr1 | Stk36 | -0.238329692 | 0.669982753 |
| ENSMUSG0000006429 | chr1 | Aox3 | -0.794136782 | 0.669982753 |
| ENSMUSG0000004591 | chr4 | Tmem268 | -0.182413223 | 0.669982753 |
| ENSMUSG0000002573 | chr17 | Wdr24 | -0.124780156 | 0.669982753 |
| ENSMUSG0000001002 | chr11 | Aldh3a2 | -0.133792916 | 0.669982753 |
| ENSMUSG0000001775 | chr13 | Slc12a7 | 0.091300804 | 0.669982753 |
| ENSMUSG0000002574 | chr4 | Sdc3 | 0.161038726 | 0.670015352 |
| ENSMUSG0000002443 | chr18 | Nr3c1 | -0.15751546 | 0.670115889 |
| ENSMUSG0000002036 | chr11 | Gfpt2 | 0.228593831 | 0.670158603 |
| ENSMUSG0000003159 | chr8 | Slc7a2 | -0.27159726 | 0.670382075 |
| ENSMUSG0000004831 | chr8 | Pskh1 | 0.141757669 | 0.670913537 |
| ENSMUSG0000000110 | chr11 | Ift20 | 0.266752042 | 0.670913537 |
| ENSMUSG0000002813 | chr3 | Celf3 | -0.234849535 | 0.670980992 |
| ENSMUSG0000004746 | chr18 | Ark2n | 0.122281086 | 0.671001453 |
| ENSMUSG0000006362 | chr8 | Unc5d | -0.237891164 | 0.671291163 |
| ENSMUSG0000007887 | chr2 | Gm14305 | -0.187647177 | 0.671291163 |
| ENSMUSG0000003819 | chr8 | Hand2 | -0.092330522 | 0.671291163 |
| ENSMUSG0000007527 | chr2 | Pde11a | 0.549443872 | 0.671291163 |
| ENSMUSG0000005489 | chr12 | Dmac2l | -0.173848503 | 0.671394804 |
| ENSMUSG0000002697 | chr2 | Rbms1 | 0.100017791 | 0.671740815 |
| ENSMUSG0000003372 | chr15 | Lrrc14 | 0.157319809 | 0.671740815 |
| ENSMUSG0000009614 | chr17 | Ankrd66 | 0.529791837 | 0.671950706 |
| ENSMUSG0000004713 | chr10 | Cd24a | 0.201484609 | 0.672022552 |
| ENSMUSG0000009138 | chr13 | Gcnt4 | -0.735332291 | 0.672437332 |
| ENSMUSG0000009635 | chr4 | Samd11 | -0.540689901 | 0.672437332 |
| ENSMUSG0000004807 | chr11 | Pirt | -0.189275165 | 0.672437332 |
| ENSMUSG0000002779 | chr3 | Smad9 | -0.141762154 | 0.672437332 |
| ENSMUSG0000002946 | chr5 | Ift81 | -0.114127496 | 0.672437332 |
| ENSMUSG0000004221 | chr18 | Fbxo38 | 0.087324683 | 0.672437332 |
| ENSMUSG0000002651 | chr1 | Tmem63a | 0.101914582 | 0.672437332 |

| | | | | |
|---|---|---|---|---|
| ENSMUSG0000006956 | chr10 | Dazap1 | 0.112291474 | 0.672437332 |
| ENSMUSG0000003178 | chr8 | Katnb1 | 0.116882227 | 0.672437332 |
| ENSMUSG0000002602 | chr1 | Sumo1 | 0.144548156 | 0.672437332 |
| ENSMUSG0000006173 | chr15 | Ext1 | 0.154688316 | 0.672437332 |
| ENSMUSG0000002317 | chr11 | Gps2 | 0.162464057 | 0.672437332 |
| ENSMUSG0000002495 | chr19 | Prdx5 | 0.229636266 | 0.672437332 |
| ENSMUSG0000000239 | chr8 | Use1 | 0.250723251 | 0.672437332 |
| ENSMUSG0000003743 | chr1 | Fer1l5 | -0.189332308 | 0.672926729 |
| ENSMUSG0000004342 | chr18 | Eif3j2 | 0.517315694 | 0.672926729 |
| ENSMUSG0000002204 | chr14 | Chrna2 | 0.790459566 | 0.672926729 |
| ENSMUSG0000003308 | chr14 | Tbc1d4 | -0.218813771 | 0.673014778 |
| ENSMUSG0000005989 | chr9 | Ube4a | -0.137988188 | 0.673014778 |
| ENSMUSG0000000144 | chr11 | Kpnb1 | 0.122449208 | 0.673014778 |
| ENSMUSG0000004248 | chr9 | Leo1 | 0.38674386 | 0.673014778 |
| ENSMUSG0000002251 | chr16 | Mgrn1 | 0.093533374 | 0.673198523 |
| ENSMUSG0000003230 | chr9 | Fam219b | -0.122007885 | 0.673242648 |
| ENSMUSG0000005313 | chr5 | Supt7l | 0.138121973 | 0.673242648 |
| ENSMUSG0000002802 | chr3 | Glrb | -0.190009905 | 0.673274998 |
| ENSMUSG0000006690 | chr5 | Suds3 | 0.135650653 | 0.67349925 |
| ENSMUSG0000006095 | chr12 | Trmt61a | 0.270624597 | 0.67349925 |
| ENSMUSG0000004252 | chr11 | Kcnj12 | -0.155229286 | 0.673841109 |
| ENSMUSG0000004267 | chr11 | Myo15a | -0.856514711 | 0.674162541 |
| ENSMUSG0000005114 | chr2 | Adnp | 0.257299007 | 0.674162541 |
| ENSMUSG0000002164 | chr13 | Smn1 | 0.169132616 | 0.674213013 |
| ENSMUSG0000003487 | chr7 | Nudt19 | 0.164151269 | 0.674213013 |
| ENSMUSG0000005027 | chr11 | Tmem220 | -0.303130086 | 0.674242082 |
| ENSMUSG0000002440 | chr18 | Wdr33 | 0.105600527 | 0.674242082 |
| ENSMUSG0000005253 | chr2 | Nup188 | 0.110011831 | 0.674242082 |
| ENSMUSG0000002701 | chr2 | Ube2e3 | 0.163941062 | 0.674242082 |
| ENSMUSG0000002713 | chr2 | Nop10 | 0.243553083 | 0.674242082 |
| ENSMUSG0000003761 | chr3 | Kcnmb2 | 0.711247346 | 0.674242082 |
| ENSMUSG0000003545 | chr10 | Stab2 | 0.328408534 | 0.674242082 |
| ENSMUSG0000007869 | chr11 | Cisd3 | 0.21180605 | 0.674274561 |
| ENSMUSG0000005357 | chr11 | 4930563E22Rik | -0.329482989 | 0.674490145 |
| ENSMUSG0000002919 | chr5 | Tada2b | 0.127007371 | 0.674490145 |
| ENSMUSG0000002460 | chr18 | Dctn4 | 0.14005808 | 0.674490145 |
| ENSMUSG0000003909 | chr11 | Rsad1 | -0.293391868 | 0.674570388 |
| ENSMUSG0000004385 | chr11 | Mgat5b | -0.377575805 | 0.6745838 |
| ENSMUSG0000004148 | chr12 | Serpina3g | -0.746552882 | 0.6745838 |
| ENSMUSG0000007482 | chr13 | Ramacl | -0.283685592 | 0.6745838 |
| ENSMUSG0000000082 | chr2 | Tpd52l2 | 0.163438996 | 0.6745838 |
| ENSMUSG0000002656 | chr1 | Mpc2 | 0.227511723 | 0.6745838 |
| ENSMUSG0000007570 | chr16 | Txnrd2 | 0.17706242 | 0.674733406 |
| ENSMUSG0000005523 | chr6 | Kcmf1 | 0.160773524 | 0.675319281 |
| ENSMUSG0000002095 | chr12 | Strn3 | 0.155223393 | 0.675597448 |
| ENSMUSG0000000954 | chr2 | Srp14 | 0.232205863 | 0.675953343 |
| ENSMUSG0000002424 | chr17 | Thumpd2 | 0.210710289 | 0.675953673 |
| ENSMUSG0000002879 | chr4 | Ccdc28b | -0.214094515 | 0.676072819 |
| ENSMUSG0000002816 | chr3 | Nfkb1 | 0.079080936 | 0.676072819 |
| ENSMUSG0000002857 | chr4 | Plaa | 0.094100684 | 0.676072819 |

| | | | | |
|---|---|---|---|---|
| ENSMUSG0000007198 | chr17 | Fndc1 | 0.253251846 | 0.676072819 |
| ENSMUSG0000002842 | chr4 | Aqp7 | 0.494984899 | 0.676101193 |
| ENSMUSG0000002953 | chr6 | St7 | 0.137374301 | 0.676290462 |
| ENSMUSG0000002852 | chr4 | Mier1 | 0.10423369 | 0.676448621 |
| ENSMUSG0000002452 | chr18 | Impa2 | 0.236651014 | 0.676448621 |
| ENSMUSG0000003784 | chr2 | Vstm2l | -0.201430089 | 0.676735573 |
| ENSMUSG0000007492 | chr2 | Pak6 | -0.305613164 | 0.676735573 |
| ENSMUSG0000002692 | chr2 | Card9 | -0.511661754 | 0.676735573 |
| ENSMUSG0000002053 | chr11 | Srebf1 | -0.169458038 | 0.676735573 |
| ENSMUSG0000002433 | chr17 | Tap2 | -0.157772683 | 0.676735573 |
| ENSMUSG0000001986 | chr10 | Rtn4ip1 | -0.159502878 | 0.676735573 |
| ENSMUSG0000003027 | chr6 | Etnk1 | -0.144467017 | 0.676735573 |
| ENSMUSG0000002726 | chr2 | Tubgcp4 | -0.137439331 | 0.676735573 |
| ENSMUSG0000003708 | chr17 | Slc35b2 | 0.117732889 | 0.676735573 |
| ENSMUSG0000002808 | chr3 | Gatb | 0.18486864 | 0.676735573 |
| ENSMUSG0000003354 | chr9 | Gtf2a2 | 0.15130019 | 0.676735573 |
| ENSMUSG0000001626 | chr1 | Sertad4 | 0.14995663 | 0.676735573 |
| ENSMUSG0000007900 | chr8 | Samd1 | 0.177847944 | 0.676735573 |
| ENSMUSG0000003619 | chr17 | Slc26a8 | 0.402710408 | 0.676735573 |
| ENSMUSG0000002879 | chr4 | Ak2 | 0.18186634 | 0.676735573 |
| ENSMUSG0000002414 | chr17 | Cript | 0.232310181 | 0.676735573 |
| ENSMUSG0000005493 | chr13 | Zkscan4 | 0.351521813 | 0.676735573 |
| ENSMUSG0000003035 | chr6 | Prmt8 | 0.353191641 | 0.676735573 |
| ENSMUSG0000004537 | chr11 | Tmem88 | 0.346261001 | 0.676735573 |
| ENSMUSG0000006093 | chr11 | Rpl26 | 0.340511 | 0.676735573 |
| ENSMUSG0000002170 | chr13 | Thbs4 | 0.329016491 | 0.676735573 |
| ENSMUSG0000004092 | chr19 | Rfx3 | -0.167088732 | 0.676927804 |
| ENSMUSG0000003376 | chr19 | Nrxn2 | -0.181759407 | 0.676927804 |
| ENSMUSG0000003355 | chr3 | Dph5 | 0.164353235 | 0.676927804 |
| ENSMUSG0000003899 | chr13 | Txndc5 | 0.145292761 | 0.676927804 |
| ENSMUSG0000000381 | chr4 | Mast2 | 0.171759723 | 0.676927804 |
| ENSMUSG0000000284 | chr16 | Adprh | 0.143962054 | 0.676942583 |
| ENSMUSG0000005032 | chr18 | Neto1 | -0.251487864 | 0.677155436 |
| ENSMUSG0000000143 | chr10 | Slc19a1 | -0.185368688 | 0.677155436 |
| ENSMUSG0000003948 | chr10 | Nt5dc1 | 0.160148282 | 0.677155436 |
| ENSMUSG0000002891 | chr4 | Arhgef19 | -0.164566064 | 0.677173555 |
| ENSMUSG0000002075 | chr11 | Sap30bp | 0.09537949 | 0.677173555 |
| ENSMUSG0000003762 | chr6 | Atoh8 | 0.229547726 | 0.677173555 |
| ENSMUSG0000000636 | chr12 | Crip1 | 0.324236311 | 0.677173555 |
| ENSMUSG0000003417 | chr5 | Zbed5 | -0.294974619 | 0.677315849 |
| ENSMUSG0000001529 | chrX | Ubl4a | 0.172868258 | 0.677315849 |
| ENSMUSG0000000756 | chr17 | Ppp2r1a | 0.108666159 | 0.677494588 |
| ENSMUSG0000002903 | chr4 | Tprg1l | 0.142215863 | 0.677494588 |
| ENSMUSG0000002087 | chr11 | Scrn2 | -0.235064891 | 0.677507487 |
| ENSMUSG0000001810 | chr13 | H2bc4 | 0.285982219 | 0.677507487 |
| ENSMUSG0000000283 | chr17 | Plin4 | 0.232505635 | 0.677552665 |
| ENSMUSG0000003497 | chr9 | Dop1a | -0.130097644 | 0.678010905 |
| ENSMUSG0000003505 | chr17 | Dhx57 | -0.13673923 | 0.678041106 |
| ENSMUSG0000005992 | chr16 | 4930453N24Rik | 0.147593438 | 0.678153932 |
| ENSMUSG0000000957 | chr15 | Cbx5 | -0.10591758 | 0.678297753 |

| | | | | |
|---|---|---|---|---|
| ENSMUSG0000003465 | chr19 | Tmem109 | 0.169320609 | 0.678297753 |
| ENSMUSG0000003858 | chr10 | Pln | 0.190745954 | 0.678297753 |
| ENSMUSG0000002022 | chr4 | Vps13d | -0.166981121 | 0.678457935 |
| ENSMUSG0000005173 | chr2 | Mettl5 | 0.189324637 | 0.678457935 |
| ENSMUSG0000006824 | chr4 | Uba52rt | 0.348922257 | 0.678457935 |
| ENSMUSG0000003229 | chr9 | Ireb2 | -0.132721776 | 0.678462065 |
| ENSMUSG0000002523 | chr9 | Bbs4 | -0.131653845 | 0.678555788 |
| ENSMUSG0000002137 | chr13 | Dek | 0.104339291 | 0.678555788 |
| ENSMUSG0000003204 | chr9 | Tirap | 0.133036404 | 0.678555788 |
| ENSMUSG0000006599 | chr4 | Aurkaip1 | 0.180965778 | 0.678555788 |
| ENSMUSG0000007868 | chr7 | Tm2d3 | 0.220570937 | 0.678555788 |
| ENSMUSG0000005511 | chr15 | Anxa13 | -1.089155705 | 0.67930148 |
| ENSMUSG0000002145 | chr13 | Syk | -0.295110963 | 0.67930148 |
| ENSMUSG0000003799 | chr6 | Parp11 | -0.164071382 | 0.67930148 |
| ENSMUSG0000002938 | chr5 | Ddx55 | -0.150234648 | 0.67930148 |
| ENSMUSG0000002209 | chr14 | Bmp1 | 0.129173302 | 0.67930148 |
| ENSMUSG0000003669 | chr5 | Nop14 | 0.129480776 | 0.67930148 |
| ENSMUSG0000003380 | chr16 | Alg3 | 0.156836619 | 0.67930148 |
| ENSMUSG0000002682 | chr2 | Nr4a2 | 0.445390932 | 0.67930148 |
| ENSMUSG0000003633 | chr8 | Slc18a1 | -0.784824722 | 0.67978505 |
| ENSMUSG0000002087 | chr11 | Slc35b1 | 0.126641435 | 0.67978505 |
| ENSMUSG0000004996 | chr14 | Mrps16 | 0.232451356 | 0.67978505 |
| ENSMUSG0000002211 | chr14 | Spry2 | -0.186747654 | 0.67987458 |
| ENSMUSG0000003455 | chr4 | Zfyve9 | -0.151905542 | 0.68057541 |
| ENSMUSG0000003217 | chr9 | Pin1 | 0.215964911 | 0.68057541 |
| ENSMUSG0000005039 | chr4 | Tnfsf15 | -0.81081682 | 0.680592235 |
| ENSMUSG0000006299 | chr6 | Ica1 | -0.185766353 | 0.680689116 |
| ENSMUSG0000002084 | chr11 | Rph3al | 0.116778929 | 0.680744401 |
| ENSMUSG0000003729 | chr8 | Lsm1 | 0.214771863 | 0.681313615 |
| ENSMUSG0000004042 | chr15 | Cdh18 | -0.359907098 | 0.681404825 |
| ENSMUSG0000005023 | chrX | Cxcr3 | -0.922611533 | 0.681404825 |
| ENSMUSG0000000854 | chr6 | Mgst1 | -0.261512619 | 0.681404825 |
| ENSMUSG0000003202 | chr9 | Jhy | -0.615478065 | 0.681404825 |
| ENSMUSG0000002488 | chr19 | Aldh3b1 | -0.209154499 | 0.681404825 |
| ENSMUSG0000002526 | chrX | Fam120c | -0.170815084 | 0.681404825 |
| ENSMUSG0000003122 | chrX | Magee2 | -0.449117906 | 0.681404825 |
| ENSMUSG0000002091 | chr11 | Acly | -0.139835236 | 0.681404825 |
| ENSMUSG0000006804 | chr2 | Tm9sf4 | -0.113810382 | 0.681404825 |
| ENSMUSG0000004807 | chr11 | Arf1 | 0.168249666 | 0.681404825 |
| ENSMUSG0000007559 | chr15 | Nrbp2 | 0.178190399 | 0.681404825 |
| ENSMUSG0000004308 | chr6 | Il17re | 0.261798604 | 0.681404825 |
| ENSMUSG0000002564 | chr9 | Shisa5 | -0.12023371 | 0.681807057 |
| ENSMUSG0000004075 | chr14 | Myh6 | -0.394618087 | 0.681913827 |
| ENSMUSG0000002017 | chr10 | Adora2a | -0.259203413 | 0.682267191 |
| ENSMUSG0000002045 | chr11 | Eif4enif1 | 0.087018456 | 0.682322863 |
| ENSMUSG0000002937 | chr5 | Pf4 | 0.161848657 | 0.682322863 |
| ENSMUSG0000002424 | chr17 | Cox7a2l | 0.20601417 | 0.682322863 |
| ENSMUSG0000005672 | chr9 | Nbeal2 | -0.222744181 | 0.682916633 |
| ENSMUSG0000002917 | chr5 | Cenpa | -0.430108557 | 0.68292808 |
| ENSMUSG0000004829 | chr19 | Or10v5 | -0.911223865 | 0.68292808 |

| | | | | |
|---|---|---|---|---|
| ENSMUSG0000002926 | chr5 | Mtf2 | -0.167896898 | 0.68292808 |
| ENSMUSG0000002643 | chr1 | Nucks1 | -0.090498542 | 0.68292808 |
| ENSMUSG0000005060 | chr7 | Zfp61 | 0.149798379 | 0.682937276 |
| ENSMUSG0000005565 | chrX | Gpc3 | -0.252290871 | 0.683099454 |
| ENSMUSG0000000479 | chr12 | Pgf | -0.437879531 | 0.683099454 |
| ENSMUSG0000002642 | chr1 | Gpr37l1 | -0.187461359 | 0.683099454 |
| ENSMUSG0000003137 | chrX | Zfp92 | -0.377798946 | 0.683099454 |
| ENSMUSG0000005139 | chr17 | Zbtb22 | -0.161731044 | 0.683099454 |
| ENSMUSG0000002947 | chr5 | Anapc5 | 0.09628956 | 0.683099454 |
| ENSMUSG0000005056 | chr11 | Maml1 | 0.136861682 | 0.683099454 |
| ENSMUSG0000003327 | chr18 | Slc35a4 | 0.127337466 | 0.683099454 |
| ENSMUSG0000007430 | chr9 | Peak1 | 0.146774628 | 0.683099454 |
| ENSMUSG0000002920 | chr5 | Gnpda2 | 0.181793698 | 0.683099454 |
| ENSMUSG0000000928 | chr6 | Rarres2 | 0.289398915 | 0.683099454 |
| ENSMUSG0000006803 | chr17 | Mas1 | 0.403925821 | 0.683099454 |
| ENSMUSG0000003569 | chr10 | Caps2 | 0.448015156 | 0.683099454 |
| ENSMUSG0000004258 | chr1 | Thsd7b | -0.300964419 | 0.683100633 |
| ENSMUSG0000002498 | chr19 | Tcf7l2 | -0.211796581 | 0.683100633 |
| ENSMUSG0000002294 | chr16 | Setd4 | -0.30471893 | 0.683100633 |
| ENSMUSG0000003340 | chr14 | Nudt15 | -0.335591665 | 0.683100633 |
| ENSMUSG0000003616 | chr12 | Sostdc1 | -0.170132614 | 0.683100633 |
| ENSMUSG0000002675 | chr2 | Golga1 | -0.114008934 | 0.683100633 |
| ENSMUSG0000004473 | chr4 | Airim | 0.098653976 | 0.683100633 |
| ENSMUSG0000002247 | chr15 | Pmm1 | 0.114557953 | 0.683100633 |
| ENSMUSG0000005283 | chr7 | Sae1 | 0.156895505 | 0.683100633 |
| ENSMUSG0000002547 | chr7 | Cyp2e1 | 0.808952356 | 0.683100633 |
| ENSMUSG0000002444 | chr17 | Rpp21 | 0.224435445 | 0.683100633 |
| ENSMUSG0000002728 | chr2 | Mtch2 | 0.091332937 | 0.683143628 |
| ENSMUSG0000003939 | chr1 | Mreg | 0.405808563 | 0.683143628 |
| ENSMUSG0000002900 | chr4 | Mad2l2 | 0.157231429 | 0.683277066 |
| ENSMUSG0000000456 | chr8 | Arhgef18 | 0.165106594 | 0.683288311 |
| ENSMUSG0000005492 | chr5 | Klhl5 | 0.164307196 | 0.683484232 |
| ENSMUSG0000005738 | chr17 | Mrpl18 | 0.244203909 | 0.683484232 |
| ENSMUSG0000003906 | chr2 | Echdc3 | -0.277105705 | 0.683499472 |
| ENSMUSG0000000112 | chr10 | Pcbp3 | -0.13642167 | 0.683499472 |
| ENSMUSG0000002920 | chr5 | Rhoh | -0.610229953 | 0.683675224 |
| ENSMUSG0000003778 | chr6 | Vopp1 | -0.14872079 | 0.683675224 |
| ENSMUSG0000003131 | chrX | Taf1 | -0.129158611 | 0.683675224 |
| ENSMUSG0000003004 | chr6 | Mrpl19 | 0.229454462 | 0.683675224 |
| ENSMUSG0000004799 | chrX | Prrg1 | -0.301708057 | 0.683894477 |
| ENSMUSG0000004531 | chr17 | Fahd1 | -0.181836975 | 0.683894477 |
| ENSMUSG0000005677 | chr12 | Setd3 | 0.158733923 | 0.683894477 |
| ENSMUSG0000004274 | chr3 | Krtcap2 | 0.21687778 | 0.683894477 |
| ENSMUSG0000002761 | chr2 | Procr | 0.373357761 | 0.683894477 |
| ENSMUSG0000003671 | chr5 | Micall2 | -0.43792196 | 0.684371577 |
| ENSMUSG0000000384 | chr8 | Nqo1 | -0.200678175 | 0.684542589 |
| ENSMUSG0000003366 | chr15 | Zfp7 | -0.211846097 | 0.684826499 |
| ENSMUSG0000003694 | chr18 | Elac1 | -0.107867331 | 0.684826499 |
| ENSMUSG0000002845 | chr4 | Tesk1 | 0.111896688 | 0.684826499 |
| ENSMUSG0000005763 | chr4 | Prdm2 | 0.091095311 | 0.68486051 |

| | | | | |
|---|---|---|---|---|
| ENSMUSG0000003067 | chr7 | Qprt | 0.328747461 | 0.684942611 |
| ENSMUSG0000003885 | chr5 | Baiap2l1 | -0.203400587 | 0.685184068 |
| ENSMUSG0000002442 | chr18 | Lama3 | -0.677390054 | 0.685466142 |
| ENSMUSG0000003102 | chr7 | Dennd2b | 0.134960586 | 0.685793617 |
| ENSMUSG0000000366 | chr2 | Ciao1 | 0.170840275 | 0.685907402 |
| ENSMUSG0000002647 | chr1 | Glul | 0.184941434 | 0.685907402 |
| ENSMUSG0000002032 | chr10 | Fgf22 | -0.591194112 | 0.686062916 |
| ENSMUSG0000006902 | chr2 | Urm1 | 0.154149041 | 0.686062916 |
| ENSMUSG0000002155 | chr13 | Qng1 | 0.162737781 | 0.686062916 |
| ENSMUSG0000002519 | chr19 | Got1 | 0.167086816 | 0.686062916 |
| ENSMUSG0000001450 | chr18 | Pkd2l2 | 0.165083999 | 0.686062916 |
| ENSMUSG0000002057 | chr12 | Pdia6 | 0.324259921 | 0.686062916 |
| ENSMUSG0000002009 | chr10 | Pald1 | 0.097315397 | 0.686131054 |
| ENSMUSG0000005128 | chr1 | Pcmtd1 | 0.108947763 | 0.686131054 |
| ENSMUSG0000002278 | chr16 | Fgd4 | -0.119690022 | 0.686224238 |
| ENSMUSG0000003785 | chr13 | Iars1 | 0.133740794 | 0.686224238 |
| ENSMUSG0000002635 | chr1 | Rgs18 | 0.364913897 | 0.686224238 |
| ENSMUSG0000005198 | chr19 | Sec31b | 0.271909221 | 0.686224238 |
| ENSMUSG0000002108 | chr12 | Rtn1 | -0.171068594 | 0.686318776 |
| ENSMUSG0000002274 | chr16 | Cmss1 | -0.226818199 | 0.686318776 |
| ENSMUSG0000002914 | chr5 | Krtcap3 | -0.20803831 | 0.686318776 |
| ENSMUSG0000002804 | chr3 | Adam15 | 0.138700303 | 0.686583672 |
| ENSMUSG0000002547 | chr7 | Adam8 | -0.356530379 | 0.686598636 |
| ENSMUSG0000004095 | chr18 | Cables1 | -0.330365932 | 0.686598636 |
| ENSMUSG0000003860 | chr10 | Slc35f1 | -0.202040605 | 0.686598636 |
| ENSMUSG0000003394 | chr4 | Zswim5 | -0.274175513 | 0.686598636 |
| ENSMUSG0000003550 | chr13 | Fbxl21 | -0.296845122 | 0.686598636 |
| ENSMUSG0000003731 | chr1 | Traf3ip3 | -0.577738658 | 0.686598636 |
| ENSMUSG0000003529 | chr2 | Wdr38 | -0.542492828 | 0.686598636 |
| ENSMUSG0000003478 | chr13 | Rab24 | -0.236995739 | 0.686598636 |
| ENSMUSG0000007346 | chr17 | Sft2d1 | -0.167018559 | 0.686598636 |
| ENSMUSG0000002046 | chr11 | Pnpt1 | -0.104494903 | 0.686598636 |
| ENSMUSG0000002193 | chr14 | Ctsb | 0.084148629 | 0.686598636 |
| ENSMUSG0000002513 | chr7 | Tollip | 0.078935435 | 0.686598636 |
| ENSMUSG0000002079 | chr11 | Exoc7 | 0.094040799 | 0.686598636 |
| ENSMUSG0000000330 | chr9 | Keap1 | 0.115017674 | 0.686598636 |
| ENSMUSG0000002062 | chr12 | Trappc12 | 0.114508257 | 0.686598636 |
| ENSMUSG0000002286 | chr16 | D16Ertd472e | 0.139428002 | 0.686598636 |
| ENSMUSG0000003075 | chr7 | Psma1 | 0.151236114 | 0.686598636 |
| ENSMUSG0000002921 | chr5 | Commd8 | 0.148209676 | 0.686598636 |
| ENSMUSG0000007942 | chr6 | Arpc4 | 0.152520161 | 0.686598636 |
| ENSMUSG0000007942 | chr9 | Mthfsl | 0.196852638 | 0.686598636 |
| ENSMUSG0000002880 | chr4 | Zbtb8a | 0.196721564 | 0.686598636 |
| ENSMUSG0000006059 | chr7 | Ifitm2 | 0.168908535 | 0.686598636 |
| ENSMUSG0000005549 | chr19 | Pprc1 | 0.207656923 | 0.686598636 |
| ENSMUSG0000002202 | chr14 | Mtrf1 | 0.208770019 | 0.686598636 |
| ENSMUSG0000008740 | chrX | Kantr | -0.146888488 | 0.686886233 |
| ENSMUSG0000005520 | chr7 | Sertad3 | -0.178897258 | 0.686961318 |
| ENSMUSG0000005237 | chr11 | Mpp3 | -0.235377177 | 0.687027741 |
| ENSMUSG0000010406 | chr18 | Pcdhgb7 | -0.228874814 | 0.687509956 |

| | | | | |
|---|---|---|---|---|
| ENSMUSG0000006672 | chr13 | Zfp184 | -0.252493532 | 0.687509956 |
| ENSMUSG0000006727 | chr19 | Ppp1r3c | 0.313611877 | 0.687509956 |
| ENSMUSG000000234 | chr8 | Armc6 | 0.248184973 | 0.687524434 |
| ENSMUSG0000005405 | chr14 | Ercc6 | -0.164245895 | 0.687897885 |
| ENSMUSG0000004485 | chr17 | Lemd2 | 0.10796524 | 0.687897885 |
| ENSMUSG0000003287 | chr9 | Cyb5r4 | 0.154065439 | 0.687897885 |
| ENSMUSG0000009848 | chr2 | Pla2g4b | 0.516847595 | 0.687897885 |
| ENSMUSG0000003237 | chr9 | Lactb | 0.163927531 | 0.688049117 |
| ENSMUSG0000002736 | chr2 | Usp50 | 0.499412731 | 0.688049117 |
| ENSMUSG0000002007 | chr10 | Srgn | 0.208333004 | 0.688371441 |
| ENSMUSG0000002609 | chr1 | Osgepl1 | 0.228062927 | 0.688371441 |
| ENSMUSG0000003022 | chr6 | Arhgdib | 0.201250506 | 0.688371441 |
| ENSMUSG000000342 | chr7 | Rps11 | 0.259835473 | 0.688371441 |
| ENSMUSG0000001996 | chr12 | Psen1 | -0.125718786 | 0.689009614 |
| ENSMUSG0000003726 | chr2 | Kin | 0.19283134 | 0.689085072 |
| ENSMUSG0000001483 | chr8 | Matcap1 | -0.269481531 | 0.689145115 |
| ENSMUSG0000004388 | chr14 | Kbtbd7 | -0.11317973 | 0.689145115 |
| ENSMUSG000000295 | chr7 | Ap2a2 | -0.098110853 | 0.689145115 |
| ENSMUSG000000237 | chr17 | Ranbp3 | 0.102473846 | 0.689145115 |
| ENSMUSG0000004023 | chr8 | Trappc5 | 0.165229707 | 0.689145115 |
| ENSMUSG0000003280 | chr2 | Srxn1 | 0.146616846 | 0.689145115 |
| ENSMUSG0000006602 | chr4 | Dhrs3 | 0.176889477 | 0.689145115 |
| ENSMUSG0000002864 | chr4 | Ermap | 0.67655579 | 0.689145115 |
| ENSMUSG0000003188 | chr8 | Rrad | 0.242078448 | 0.689145115 |
| ENSMUSG0000002379 | chr16 | Pigx | 0.190142684 | 0.689324057 |
| ENSMUSG0000001550 | chr10 | Hivep2 | 0.213683359 | 0.689324057 |
| ENSMUSG0000002844 | chr4 | Enho | 0.25958126 | 0.689324057 |
| ENSMUSG000000167 | chr8 | Marveld3 | -0.291253103 | 0.689517748 |
| ENSMUSG0000002255 | chr15 | Adck5 | -0.181958627 | 0.689517748 |
| ENSMUSG0000002086 | chr11 | Luc7l3 | -0.130154011 | 0.689517748 |
| ENSMUSG0000006051 | chr1 | Tor3a | -0.147286504 | 0.689728116 |
| ENSMUSG0000002777 | chr3 | Ift80 | -0.103149411 | 0.689728116 |
| ENSMUSG0000006324 | chr4 | Zfp993 | 0.479491866 | 0.689728116 |
| ENSMUSG0000004102 | chr4 | Iffo2 | 0.138456826 | 0.689758997 |
| ENSMUSG0000009956 | chr9 | Gm28729 | -0.364479488 | 0.689896026 |
| ENSMUSG0000002197 | chr14 | Fgf9 | -0.436948188 | 0.689896026 |
| ENSMUSG0000000030 | chr2 | Cdh4 | -0.279578008 | 0.689896026 |
| ENSMUSG0000005356 | chr8 | Ier2 | -0.259851763 | 0.689896026 |
| ENSMUSG0000007394 | chr7 | Hbb-bt | -0.715680652 | 0.689896026 |
| ENSMUSG0000002986 | chr6 | Casp2 | -0.163030841 | 0.689896026 |
| ENSMUSG0000002299 | chr15 | Spmip11 | -0.737815878 | 0.689896026 |
| ENSMUSG0000002992 | chr6 | Mkrn1 | -0.15222394 | 0.689896026 |
| ENSMUSG0000004403 | chr7 | Irf2bp1 | -0.113583412 | 0.689896026 |
| ENSMUSG0000002068 | chr11 | Taf15 | -0.089856515 | 0.689896026 |
| ENSMUSG0000001832 | chr2 | Ywhab | 0.086809893 | 0.689896026 |
| ENSMUSG0000001514 | chr7 | Sirt2 | 0.089002041 | 0.689896026 |
| ENSMUSG0000004189 | chr18 | Lman1 | 0.091877029 | 0.689896026 |
| ENSMUSG0000002431 | chr17 | Wdr46 | 0.140150943 | 0.689896026 |
| ENSMUSG0000009674 | chr19 | Lbhd1 | 0.219670108 | 0.689896026 |
| ENSMUSG000000267 | chr12 | Med6 | 0.156596979 | 0.689896026 |

| | | | | |
|---|---|---|---|---|
| ENSMUSG0000003278 | chr9 | Alas1 | 0.143966098 | 0.689896026 |
| ENSMUSG0000005133 | chr6 | Cacna1c | 0.206604259 | 0.689896026 |
| ENSMUSG0000004315 | chr9 | Ppp2r3a | 0.173257852 | 0.689896026 |
| ENSMUSG0000000573 | chr16 | Ranbp1 | 0.208804801 | 0.689896026 |
| ENSMUSG0000002847 | chr4 | Spaar | 0.294434235 | 0.689896026 |
| ENSMUSG0000003345 | chr9 | Adamts15 | 0.306270359 | 0.690173788 |
| ENSMUSG0000002807 | chr3 | Sh2d2a | -1.541408455 | 0.690398319 |
| ENSMUSG0000004122 | chr3 | Elovl6 | -0.238863619 | 0.690398319 |
| ENSMUSG0000003823 | chr7 | Hrc | -0.542513659 | 0.690398319 |
| ENSMUSG0000005889 | chr3 | Col25a1 | -0.210847391 | 0.690398319 |
| ENSMUSG0000003092 | chr7 | Rexo5 | -0.413915439 | 0.690398319 |
| ENSMUSG0000005719 | chr9 | AB124611 | -0.428476886 | 0.690398319 |
| ENSMUSG0000005007 | chr3 | Gpr171 | -0.691085694 | 0.690398319 |
| ENSMUSG0000005955 | chr10 | Ccdc28a | -0.194648818 | 0.690398319 |
| ENSMUSG0000006139 | chr9 | Acvr2b | -0.237885353 | 0.690398319 |
| ENSMUSG0000000242 | chr3 | Hltf | -0.179895148 | 0.690398319 |
| ENSMUSG0000002573 | chr17 | Fbxl16 | -0.168598028 | 0.690398319 |
| ENSMUSG0000006874 | chr3 | Celsr2 | -0.156423793 | 0.690398319 |
| ENSMUSG0000004040 | chr5 | Akap9 | -0.166534121 | 0.690398319 |
| ENSMUSG0000003954 | chr14 | Cfap70 | -0.436600871 | 0.690398319 |
| ENSMUSG0000002959 | chr5 | Sds | -0.625685879 | 0.690398319 |
| ENSMUSG0000007851 | chr4 | Ddi2 | -0.111764542 | 0.690398319 |
| ENSMUSG0000003393 | chr8 | Rbm34 | -0.119614694 | 0.690398319 |
| ENSMUSG0000002700 | chr2 | Dnajc10 | 0.109744394 | 0.690398319 |
| ENSMUSG0000006239 | chr15 | Zfp706 | 0.128202998 | 0.690398319 |
| ENSMUSG0000005911 | chr7 | Nap1l4 | 0.152917533 | 0.690398319 |
| ENSMUSG0000003752 | chr2 | Mavs | 0.149349996 | 0.690398319 |
| ENSMUSG0000002117 | chr12 | Tdp1 | 0.144628362 | 0.690398319 |
| ENSMUSG0000005333 | chr5 | Ficd | 0.150921981 | 0.690398319 |
| ENSMUSG0000002827 | chr3 | Pdlim5 | 0.155076146 | 0.690398319 |
| ENSMUSG0000003864 | chr7 | Ramac | 0.163322388 | 0.690398319 |
| ENSMUSG0000003906 | chr2 | Mtg2 | 0.163438595 | 0.690398319 |
| ENSMUSG0000005855 | chr5 | Rpl5 | 0.200288677 | 0.690398319 |
| ENSMUSG0000005824 | chr16 | Cryzl1 | 0.1760445 | 0.690398319 |
| ENSMUSG0000003975 | chr5 | Alkbh4 | 0.191131601 | 0.690398319 |
| ENSMUSG0000002794 | chr3 | Hax1 | 0.208103078 | 0.690398319 |
| ENSMUSG0000003283 | chr10 | Pwp2 | 0.192974303 | 0.690398319 |
| ENSMUSG0000003683 | chr7 | Psenen | 0.198021781 | 0.690398319 |
| ENSMUSG0000004510 | chr14 | Saysd1 | 0.270337632 | 0.690398319 |
| ENSMUSG0000002042 | chr1 | Btg2 | -0.263989607 | 0.691050007 |
| ENSMUSG0000003099 | chr7 | Pgap2 | 0.174291013 | 0.691050007 |
| ENSMUSG0000003963 | chr4 | Zfp189 | 0.207653296 | 0.691050007 |
| ENSMUSG0000000269 | chr11 | Lcp2 | -0.280063293 | 0.691445733 |
| ENSMUSG0000007294 | chr12 | Ptgr2 | 0.1424195 | 0.691500026 |
| ENSMUSG0000007057 | chr4 | Gm572 | 0.721347635 | 0.691500026 |
| ENSMUSG0000002005 | chr10 | Dram1 | 0.231318558 | 0.691536769 |
| ENSMUSG0000001989 | chr10 | Lrriq1 | -0.374153835 | 0.691571563 |
| ENSMUSG0000003053 | chr7 | Iqgap1 | 0.117132052 | 0.691700305 |
| ENSMUSG0000002534 | chr5 | Rabgef1 | 0.13645213 | 0.691715243 |
| ENSMUSG0000002849 | chr4 | Plin2 | 0.193648767 | 0.691778308 |

| | | | | |
|---|---|---|---|---|
| ENSMUSG0000002043 | chr11 | Myo1g | 0.515813549 | 0.691835031 |
| ENSMUSG0000003376 | chr15 | Recql4 | -0.526931083 | 0.691856504 |
| ENSMUSG0000003635 | chr8 | Csgalnact1 | 0.270132824 | 0.691856504 |
| ENSMUSG0000000571 | chr16 | Tfap4 | -0.334335715 | 0.692208652 |
| ENSMUSG0000002945 | chr5 | Mapkapk5 | -0.654052637 | 0.692208652 |
| ENSMUSG0000004171 | chr12 | Gm20604 | -0.20929704 | 0.692208652 |
| ENSMUSG0000004024 | chr10 | Lrp1 | -0.157280477 | 0.692208652 |
| ENSMUSG0000002406 | chr17 | Slc30a6 | 0.153933884 | 0.692208652 |
| ENSMUSG0000005756 | chr18 | Eif1a | 0.164549312 | 0.692208652 |
| ENSMUSG0000003014 | chr6 | Klrk1 | -0.692201903 | 0.692244945 |
| ENSMUSG0000002755 | chr3 | Lrrcc1 | -0.138059393 | 0.692244945 |
| ENSMUSG0000003250 | chr9 | Pdcd6ip | 0.104996819 | 0.692244945 |
| ENSMUSG0000006012 | chr14 | Tpt1 | 0.236870328 | 0.692244945 |
| ENSMUSG0000002395 | chr17 | Clic5 | -0.290418036 | 0.692266102 |
| ENSMUSG0000002757 | chr2 | Chrna4 | -0.444276197 | 0.692266102 |
| ENSMUSG0000002818 | chr3 | Zranb2 | 0.110336546 | 0.692266102 |
| ENSMUSG0000002267 | chr16 | Mpv17l | -0.200523598 | 0.692387697 |
| ENSMUSG0000002729 | chr2 | Itpka | -0.788384303 | 0.692418624 |
| ENSMUSG0000000615 | chr7 | Eps8l1 | -0.415572136 | 0.692418624 |
| ENSMUSG0000003188 | chr8 | Tradd | -0.276915067 | 0.692418624 |
| ENSMUSG0000004980 | chrX | Armcx4 | -0.202871384 | 0.692418624 |
| ENSMUSG0000002227 | chr15 | Retreg1 | -0.177439836 | 0.692418624 |
| ENSMUSG0000003911 | chr2 | Taf4 | -0.134896707 | 0.692418624 |
| ENSMUSG0000001474 | chr6 | Tex261 | -0.11396015 | 0.692418624 |
| ENSMUSG0000003117 | chrX | Dynlt3 | 0.12877817 | 0.692418624 |
| ENSMUSG0000002242 | chr15 | Dnal4 | 0.120975938 | 0.692418624 |
| ENSMUSG0000002226 | chr15 | Sdc2 | 0.153506842 | 0.692418624 |
| ENSMUSG0000002548 | chr7 | Bet1l | 0.184046266 | 0.692418624 |
| ENSMUSG0000006378 | chr14 | Chchd1 | 0.276827301 | 0.692418624 |
| ENSMUSG0000002863 | chr4 | Ppcs | -0.247271018 | 0.692426763 |
| ENSMUSG0000003923 | chrX | Suv39h1 | -0.150929077 | 0.692519566 |
| ENSMUSG0000002327 | chr15 | Creld2 | 0.327222347 | 0.692519566 |
| ENSMUSG0000003114 | chrX | Praf2 | -0.185228643 | 0.692687905 |
| ENSMUSG0000002579 | chr10 | Rassf3 | 0.253619668 | 0.692687905 |
| ENSMUSG0000003853 | chr9 | Zfp280d | 0.173926852 | 0.692687905 |
| ENSMUSG0000002821 | chr4 | Trp53inp1 | 0.206468404 | 0.692687905 |
| ENSMUSG0000002682 | chr2 | Lcn2 | 0.182688443 | 0.692687905 |
| ENSMUSG0000002327 | chr9 | Twf2 | 0.216845299 | 0.692687905 |
| ENSMUSG0000006930 | chr13 | H4c17 | 1.400641248 | 0.692687905 |
| ENSMUSG0000004520 | chr4 | Dpy19l4 | -0.1177651 | 0.692862348 |
| ENSMUSG0000002638 | chr1 | Sctr | -0.771589795 | 0.69319099 |
| ENSMUSG0000004344 | chr17 | Pgp | 0.10581707 | 0.69319099 |
| ENSMUSG0000004700 | chr15 | Zfp41 | -0.140888043 | 0.69351917 |
| ENSMUSG0000005809 | chr13 | Zfp729b | -0.11811715 | 0.69351917 |
| ENSMUSG0000002899 | chr4 | Nmnat1 | 0.199887804 | 0.69351917 |
| ENSMUSG0000002662 | chr1 | Lpgat1 | 0.111657249 | 0.693647954 |
| ENSMUSG0000003778 | chr9 | Dzip1l | 0.130883303 | 0.693667054 |
| ENSMUSG0000002908 | chr5 | Kcnip4 | -0.182263685 | 0.693765059 |
| ENSMUSG0000005563 | chr2 | Hmcn2 | 0.23365108 | 0.693765059 |
| ENSMUSG0000007643 | chr11 | Cep295nl | 0.382893737 | 0.693765059 |

| | | | | |
|---|---|---|---|---|
| ENSMUSG0000003033 | chr6 | Mrpl51 | 0.200526468 | 0.6938554 |
| ENSMUSG0000007904 | chr8 | Apela | -0.848776444 | 0.693864188 |
| ENSMUSG0000000205 | chr11 | Spag5 | -0.565470806 | 0.693864188 |
| ENSMUSG0000011020 | chr7 | Flt3l | -0.226913305 | 0.693864188 |
| ENSMUSG0000002275 | chr16 | Tmem30c | -0.793285732 | 0.693864188 |
| ENSMUSG0000004185 | chr15 | Tcf20 | -0.250937814 | 0.693864188 |
| ENSMUSG0000006896 | chr7 | Zfp114 | -0.405804446 | 0.693864188 |
| ENSMUSG0000003197 | chr8 | Dbndd1 | -0.199431832 | 0.693864188 |
| ENSMUSG0000000280 | chr13 | Epdr1 | -0.132447393 | 0.693864188 |
| ENSMUSG0000002392 | chr17 | Mmut | -0.097191877 | 0.693864188 |
| ENSMUSG0000006664 | chr12 | Wdr35 | -0.101108211 | 0.693864188 |
| ENSMUSG0000006892 | chr3 | Msto1 | 0.106634482 | 0.693864188 |
| ENSMUSG0000002052 | chr11 | Rnft1 | 0.100648748 | 0.693864188 |
| ENSMUSG0000003681 | chr8 | Cnep1r1 | 0.107712153 | 0.693864188 |
| ENSMUSG0000002138 | chr13 | Ippk | 0.11959662 | 0.693864188 |
| ENSMUSG0000004124 | chr4 | Mul1 | 0.124351926 | 0.693864188 |
| ENSMUSG0000002450 | chr18 | Ppp2r2b | 0.150216672 | 0.693864188 |
| ENSMUSG0000005826 | chr1 | Mrps14 | 0.216227397 | 0.693864188 |
| ENSMUSG0000006129 | chr4 | Agbl4 | -0.35561396 | 0.693937293 |
| ENSMUSG0000006414 | chr13 | Trim38 | 0.595172513 | 0.693937293 |
| ENSMUSG0000003523 | chr7 | Neu3 | 0.232943583 | 0.693982655 |
| ENSMUSG0000002854 | chr4 | Slc6a9 | -0.263456725 | 0.693991892 |
| ENSMUSG0000002123 | chr12 | Fam161b | -0.166835293 | 0.693991892 |
| ENSMUSG0000002121 | chr13 | Gdi2 | 0.104563407 | 0.693991892 |
| ENSMUSG0000003453 | chr7 | Zfp418 | -0.297195401 | 0.69404592 |
| ENSMUSG0000005848 | chr6 | Wdr91 | 0.20043746 | 0.69413378 |
| ENSMUSG0000002382 | chr17 | Agpat4 | 0.170354211 | 0.694159839 |
| ENSMUSG0000003170 | chr8 | Gpt2 | -0.142041978 | 0.694229059 |
| ENSMUSG0000002537 | chr15 | Cdh9 | -0.545560913 | 0.694394373 |
| ENSMUSG0000001570 | chr7 | Arnt2 | 0.226016632 | 0.694394373 |
| ENSMUSG0000002881 | chr4 | Yars1 | 0.131486361 | 0.694465473 |
| ENSMUSG0000002067 | chr11 | Ddx52 | 0.131439007 | 0.694465473 |
| ENSMUSG0000000676 | chr7 | Saal1 | 0.147056628 | 0.694465473 |
| ENSMUSG0000004760 | chr7 | Zfp235 | 0.162541581 | 0.694465473 |
| ENSMUSG0000005389 | chr7 | Ech1 | 0.239895247 | 0.694465473 |
| ENSMUSG0000003325 | chr2 | Shf | -0.228349041 | 0.694737697 |
| ENSMUSG0000000641 | chr1 | Nectin4 | -0.569062376 | 0.694737697 |
| ENSMUSG0000003116 | chrX | Wdr13 | 0.093437849 | 0.694737697 |
| ENSMUSG0000002127 | chr12 | Cdc42bpb | 0.11255257 | 0.694737697 |
| ENSMUSG0000003594 | chr10 | Acss3 | -0.415235206 | 0.694933042 |
| ENSMUSG0000002909 | chr5 | Ablim2 | -0.199067169 | 0.694933042 |
| ENSMUSG0000002045 | chr11 | Limk2 | -0.1607112 | 0.694933042 |
| ENSMUSG0000007862 | chr11 | Ccdc47 | 0.124337607 | 0.694933042 |
| ENSMUSG0000003097 | chr7 | Ctbp2 | 0.14739869 | 0.694933042 |
| ENSMUSG0000001946 | chr8 | Ptger1 | 0.249285174 | 0.694933042 |
| ENSMUSG0000002288 | chr16 | St6gal1 | -0.198735618 | 0.694967798 |
| ENSMUSG0000000102 | chr11 | Icam2 | -0.344518564 | 0.694967798 |
| ENSMUSG0000003396 | chr7 | Zfp446 | -0.140668052 | 0.694967798 |
| ENSMUSG0000003124 | chrX | Sh3bgrl | 0.132320335 | 0.694967798 |
| ENSMUSG0000003824 | chr2 | Cep250 | -0.182347138 | 0.695098678 |

| | | | | |
|---|---|---|---|---|
| ENSMUSG0000003650 | chr3 | Rnf13 | 0.09547975 | 0.695098678 |
| ENSMUSG0000000177 | chr9 | Chordc1 | 0.193794043 | 0.695494455 |
| ENSMUSG0000002270 | chr16 | Ccdc191 | -0.199489467 | 0.695602879 |
| ENSMUSG0000002074 | chr11 | Cluh | 0.11735998 | 0.695602879 |
| ENSMUSG0000003425 | chr9 | Senp6 | 0.098755167 | 0.695784989 |
| ENSMUSG0000004112 | chr4 | Nbl1 | 0.149797565 | 0.695784989 |
| ENSMUSG0000001650 | chr5 | Gtf3a | 0.169230636 | 0.695784989 |
| ENSMUSG0000003078 | chr7 | Slc5a2 | 0.802559299 | 0.695784989 |
| ENSMUSG0000004669 | chr8 | Chtf8 | 0.433513321 | 0.695784989 |
| ENSMUSG0000001766 | chr2 | Zfp334 | -0.129159434 | 0.695870244 |
| ENSMUSG0000003558 | chr7 | Tsen34 | 0.163966114 | 0.695870244 |
| ENSMUSG0000004844 | chr11 | Smim5 | 0.381282033 | 0.695870244 |
| ENSMUSG0000003785 | chr10 | Zfp365 | 0.227810444 | 0.695870244 |
| ENSMUSG0000003051 | chr7 | Tars3 | -0.186114911 | 0.695928696 |
| ENSMUSG0000003811 | chr6 | Ano2 | -0.178389499 | 0.695928696 |
| ENSMUSG0000007457 | chr3 | Lekr1 | -0.243400633 | 0.696398895 |
| ENSMUSG0000005521 | chr1 | Pld5 | -0.204569741 | 0.696414743 |
| ENSMUSG0000002170 | chr13 | Mtx3 | -0.16337149 | 0.696414743 |
| ENSMUSG0000002247 | chr15 | Xrcc6 | 0.157558689 | 0.696414743 |
| ENSMUSG0000002781 | chr3 | Eif2a | 0.208978581 | 0.696414743 |
| ENSMUSG0000002016 | chr10 | Cnot2 | 0.117756196 | 0.696466363 |
| ENSMUSG0000006787 | chr19 | Ccdc87 | -0.47770264 | 0.696674716 |
| ENSMUSG0000003101 | chrX | Usp9x | -0.168773224 | 0.696674716 |
| ENSMUSG0000002036 | chr11 | Mapk9 | -0.100843286 | 0.697340835 |
| ENSMUSG0000002964 | chr5 | Flt1 | 0.251516305 | 0.697897385 |
| ENSMUSG0000002769 | chr3 | Ect2 | -0.47651413 | 0.698046661 |
| ENSMUSG0000003338 | chr3 | Frrs1 | -0.318832465 | 0.698046661 |
| ENSMUSG0000007221 | chr16 | Septin5 | -0.168714781 | 0.698046661 |
| ENSMUSG0000002610 | chr1 | Inpp1 | -0.148376682 | 0.698046661 |
| ENSMUSG0000002942 | chr5 | Scarb2 | -0.081798024 | 0.698046661 |
| ENSMUSG0000003182 | chr8 | Usp10 | 0.123019074 | 0.698046661 |
| ENSMUSG0000003472 | chr17 | Mrps10 | 0.145603122 | 0.698046661 |
| ENSMUSG0000002799 | chr3 | Plrg1 | 0.160335145 | 0.698046661 |
| ENSMUSG0000002497 | chr19 | Gpam | -0.117723754 | 0.698118141 |
| ENSMUSG0000002317 | chr10 | Bsg | 0.17036149 | 0.698118141 |
| ENSMUSG0000000338 | chr7 | Rabac1 | 0.185101788 | 0.698118141 |
| ENSMUSG0000001598 | chr7 | Stk32c | -0.262297233 | 0.698425409 |
| ENSMUSG0000004785 | chr16 | Stx19 | -0.480945929 | 0.698566743 |
| ENSMUSG0000001509 | chr2 | Edf1 | 0.173206584 | 0.698566743 |
| ENSMUSG0000005345 | chr14 | Thoc7 | 0.175932853 | 0.698566743 |
| ENSMUSG0000005356 | chr7 | Eif3k | 0.212850915 | 0.698566743 |
| ENSMUSG0000005955 | chr2 | Tor4a | -0.28201716 | 0.698687936 |
| ENSMUSG0000000436 | chr10 | 9330159F19Rik | -0.132889035 | 0.698802636 |
| ENSMUSG0000004651 | chr16 | Cox17 | 0.218180599 | 0.698802636 |
| ENSMUSG0000001649 | chr1 | Cd46 | -0.311329788 | 0.699045967 |
| ENSMUSG0000004434 | chrX | Slc25a53 | -0.195819012 | 0.699045967 |
| ENSMUSG0000003195 | chr8 | Cfdp1 | 0.135960417 | 0.699045967 |
| ENSMUSG0000002526 | chrX | Apex2 | 0.19416603 | 0.699045967 |
| ENSMUSG0000004184 | chr19 | Rhod | 0.353801229 | 0.699109827 |
| ENSMUSG0000004099 | chr3 | Npnt | -0.172995933 | 0.699249271 |

| | | | | |
|---|---|---|---|---|
| ENSMUSG0000005351 | chr4 | Nrdc | 0.074779406 | 0.699249271 |
| ENSMUSG0000002688 | chr2 | Rbm18 | 0.141677664 | 0.699249271 |
| ENSMUSG0000007886 | chr2 | Zfp970 | 0.184270138 | 0.699249271 |
| ENSMUSG0000002865 | chr4 | Mfsd2a | 0.399395653 | 0.699249271 |
| ENSMUSG0000006776 | chrX | Xlr4b | 1.11753425 | 0.699249271 |
| ENSMUSG0000005824 | chr13 | Krbox5 | -0.663513572 | 0.699360479 |
| ENSMUSG0000004511 | chr7 | Prrt2 | -0.738832705 | 0.699360479 |
| ENSMUSG0000002173 | chr14 | Slc4a7 | 0.183895369 | 0.699360479 |
| ENSMUSG0000003038 | chr7 | Mzf1 | -0.230954129 | 0.699387074 |
| ENSMUSG0000005723 | chr4 | Rbbp4 | 0.136009665 | 0.699387074 |
| ENSMUSG0000003234 | chr9 | Elovl5 | 0.135496114 | 0.699387074 |
| ENSMUSG0000002920 | chr5 | Apbb2 | 0.136948053 | 0.699387074 |
| ENSMUSG0000002656 | chr1 | Mpzl1 | 0.128437004 | 0.699387074 |
| ENSMUSG0000004809 | chr1 | Lmod1 | 0.22802185 | 0.699387074 |
| ENSMUSG0000002089 | chr11 | Gucy2e | -0.675221583 | 0.699570418 |
| ENSMUSG0000002478 | chr19 | Rcl1 | 0.183259385 | 0.699570418 |
| ENSMUSG0000001545 | chr17 | Ager | -0.492893941 | 0.699762208 |
| ENSMUSG0000007482 | chr2 | Itpripl1 | 0.254477197 | 0.699762208 |
| ENSMUSG0000002609 | chr1 | Cracdl | -0.256241234 | 0.699918376 |
| ENSMUSG0000001419 | chr11 | Dnajc7 | -0.096860608 | 0.70019929 |
| ENSMUSG0000002447 | chr18 | Ik | 0.112740786 | 0.70019929 |
| ENSMUSG0000002209 | chr14 | Sorbs3 | 0.165496267 | 0.70019929 |
| ENSMUSG0000004244 | chr9 | Mindy2 | -0.137423844 | 0.700244715 |
| ENSMUSG0000005233 | chr6 | Immt | 0.088511969 | 0.700289226 |
| ENSMUSG0000002201 | chr14 | Rgcc | 0.240252859 | 0.700289226 |
| ENSMUSG0000003453 | chr14 | Fbxo16 | -0.338673741 | 0.700695607 |
| ENSMUSG0000008621 | chr19 | A330040F15Rik | -0.665078791 | 0.70072114 |
| ENSMUSG0000007249 | chr14 | Ppp1r3e | -0.208085888 | 0.70072114 |
| ENSMUSG0000000294 | chr8 | Timm44 | 0.103708797 | 0.70072114 |
| ENSMUSG0000006675 | chr1 | Tnfsf18 | 0.837811381 | 0.70072114 |
| ENSMUSG0000002638 | chr1 | Dbi | 0.196853309 | 0.70072114 |
| ENSMUSG0000003029 | chr6 | Med21 | 0.18512759 | 0.70072114 |
| ENSMUSG0000003330 | chr16 | Lpp | 0.185667291 | 0.70072114 |
| ENSMUSG0000002502 | chr19 | Mxi1 | 0.152840051 | 0.701197666 |
| ENSMUSG0000002623 | chr1 | Epha4 | 0.188320665 | 0.701197666 |
| ENSMUSG0000002898 | chr4 | Slc25a33 | 0.226058999 | 0.701197666 |
| ENSMUSG0000003809 | chr5 | Sbno1 | -0.141353791 | 0.701297297 |
| ENSMUSG0000002961 | chr5 | Ccz1 | 0.111423814 | 0.701512103 |
| ENSMUSG0000011345 | chr13 | Zfp935 | 0.224662785 | 0.701512103 |
| ENSMUSG0000003387 | chr18 | Ppargc1b | -0.253118095 | 0.701573661 |
| ENSMUSG0000007479 | chr2 | Itpa | 0.185338362 | 0.701573661 |
| ENSMUSG0000007277 | chr6 | Grcc10 | 0.191528415 | 0.701573661 |
| ENSMUSG0000002020 | chr10 | Phlda1 | 0.269711089 | 0.701610192 |
| ENSMUSG0000001986 | chr10 | Qrsl1 | 0.174916043 | 0.702032791 |
| ENSMUSG0000000484 | chr5 | Ap1s1 | 0.197752363 | 0.702032791 |
| ENSMUSG0000007813 | chr2 | Ankrd63 | 0.523827714 | 0.702032791 |
| ENSMUSG0000001047 | chr7 | Ebf3 | -0.229059879 | 0.702078319 |
| ENSMUSG0000000519 | chr11 | Polr2a | -0.176102467 | 0.702225835 |
| ENSMUSG0000006172 | chr7 | Tnnt3 | 0.568560898 | 0.702225835 |
| ENSMUSG0000006230 | chr7 | Nectin2 | -0.239782649 | 0.702278926 |

| | | | | |
|---|---|---|---|---|
| ENSMUSG0000002066 | chr12 | Kif3c | -0.167522207 | 0.702278926 |
| ENSMUSG0000000016 | chr9 | Pih1d2 | -0.285029361 | 0.702278926 |
| ENSMUSG0000003774 | chr10 | Phyhipl | -0.204210291 | 0.702278926 |
| ENSMUSG0000003618 | chr12 | Ankmy2 | -0.134969527 | 0.702278926 |
| ENSMUSG0000004589 | chr6 | Paip2b | -0.118505396 | 0.702278926 |
| ENSMUSG0000003995 | chr9 | Dag1 | 0.092654303 | 0.702278926 |
| ENSMUSG0000001655 | chr11 | H3f3b | 0.109155791 | 0.702278926 |
| ENSMUSG0000005311 | chr6 | Chmp3 | 0.109691624 | 0.702278926 |
| ENSMUSG0000003867 | chr2 | Arfrp1 | 0.105429938 | 0.702278926 |
| ENSMUSG0000002037 | chr11 | Rnf130 | 0.116278117 | 0.702278926 |
| ENSMUSG0000008488 | chr12 | Ccdc85c | 0.12322057 | 0.702278926 |
| ENSMUSG0000007478 | chr10 | Ube2n | 0.165382886 | 0.702278926 |
| ENSMUSG0000002855 | chr4 | Rnf11 | 0.173463338 | 0.702278926 |
| ENSMUSG0000002182 | chr14 | Plau | 0.194919971 | 0.702278926 |
| ENSMUSG0000000426 | chr6 | Emg1 | 0.199857273 | 0.702278926 |
| ENSMUSG0000001556 | chr8 | Lpl | 0.221669467 | 0.702278926 |
| ENSMUSG0000009230 | chr11 | Cdk3 | -0.811108146 | 0.702341669 |
| ENSMUSG0000004298 | chr11 | Notum | -0.663115593 | 0.702341669 |
| ENSMUSG0000004284 | chr10 | Lrrtm3 | -0.273166832 | 0.702341669 |
| ENSMUSG0000009544 | chr15 | Fignl2 | -0.282605216 | 0.702341669 |
| ENSMUSG0000007147 | chr13 | H2ac7 | -0.532135818 | 0.702341669 |
| ENSMUSG0000003235 | chr9 | Rasgrf1 | -0.434448974 | 0.702341669 |
| ENSMUSG0000002308 | chr17 | H2-M10.2 | -0.691232973 | 0.702341669 |
| ENSMUSG0000003331 | chr10 | Gstt2 | -0.182650147 | 0.702341669 |
| ENSMUSG0000001990 | chr10 | Lin7a | -0.205963443 | 0.702341669 |
| ENSMUSG0000007047 | chr2 | Fam217b | -0.164222999 | 0.702341669 |
| ENSMUSG0000005832 | chr7 | Dock1 | -0.142526637 | 0.702341669 |
| ENSMUSG0000004644 | chr4 | Camk2n1 | -0.13836705 | 0.702341669 |
| ENSMUSG0000002575 | chr3 | Mfsd8 | -0.132456908 | 0.702341669 |
| ENSMUSG0000003515 | chr8 | Elmod2 | -0.117593142 | 0.702341669 |
| ENSMUSG0000002692 | chr2 | Pmpca | -0.095644723 | 0.702341669 |
| ENSMUSG0000002524 | chr9 | Lztfl1 | -0.087710584 | 0.702341669 |
| ENSMUSG0000002615 | chr1 | Agfg1 | -0.088246061 | 0.702341669 |
| ENSMUSG0000000990 | chr1 | Vps4b | -0.08294426 | 0.702341669 |
| ENSMUSG0000002119 | chr13 | Pitrm1 | 0.088767132 | 0.702341669 |
| ENSMUSG0000003037 | chr7 | Strn4 | 0.087086889 | 0.702341669 |
| ENSMUSG0000001865 | chr11 | Tada2a | 0.114322688 | 0.702341669 |
| ENSMUSG0000002418 | chr17 | Fam234a | 0.111324308 | 0.702341669 |
| ENSMUSG0000002149 | chr13 | Txndc15 | 0.128261312 | 0.702341669 |
| ENSMUSG0000003889 | chr9 | Zfp653 | 0.150561935 | 0.702341669 |
| ENSMUSG0000002840 | chr4 | Mpdz | 0.135812898 | 0.702341669 |
| ENSMUSG0000003924 | chr5 | Ints15 | 0.140676251 | 0.702341669 |
| ENSMUSG0000002856 | chr4 | Txndc12 | 0.180849165 | 0.702341669 |
| ENSMUSG0000002952 | chr5 | Pxn | 0.173258232 | 0.702341669 |
| ENSMUSG0000002564 | chr9 | Ccdc51 | 0.230199109 | 0.702341669 |
| ENSMUSG0000007919 | chr14 | Psme2 | 0.215939427 | 0.702341669 |
| ENSMUSG0000005458 | chr2 | Pabpc1l | 0.590130272 | 0.702341669 |
| ENSMUSG0000002771 | chr3 | Trpc3 | 0.229288751 | 0.702341669 |
| ENSMUSG0000002485 | chr19 | Cdk2ap2 | 0.265662901 | 0.702474457 |
| ENSMUSG0000003856 | chr5 | Rad9b | -0.275652641 | 0.702849668 |

| | | | | |
|---|---|---|---|---|
| ENSMUSG0000002612 | chr1 | Sema4c | 0.096975837 | 0.702849668 |
| ENSMUSG0000003356 | chr15 | Rbfox2 | 0.110658751 | 0.702879221 |
| ENSMUSG0000000907 | chr11 | Zmat5 | 0.193474253 | 0.702879221 |
| ENSMUSG0000002420 | chr17 | Rfx2 | 0.275710107 | 0.702879221 |
| ENSMUSG0000009066 | chr6 | Mdfic2 | -0.599885572 | 0.703076755 |
| ENSMUSG0000002750 | chr3 | Pag1 | 0.165406684 | 0.703076755 |
| ENSMUSG0000003629 | chr15 | Slc2a13 | -0.188441531 | 0.703414365 |
| ENSMUSG0000002151 | chr13 | Mterf3 | 0.139656268 | 0.703414365 |
| ENSMUSG0000001595 | chr7 | Wnt11 | -0.244660083 | 0.70367026 |
| ENSMUSG0000001459 | chr3 | Csf1 | -0.154488813 | 0.704074444 |
| ENSMUSG0000006227 | chr9 | Morf4l1 | 0.124016928 | 0.704074444 |
| ENSMUSG0000003956 | chr16 | Ubald1 | 0.122768065 | 0.704074444 |
| ENSMUSG0000002008 | chr10 | Tysnd1 | 0.149382375 | 0.704074444 |
| ENSMUSG0000003827 | chr19 | Fau | 0.2456738 | 0.704074444 |
| ENSMUSG0000002406 | chr17 | Lbh | 0.240656589 | 0.704074444 |
| ENSMUSG0000004568 | chr10 | Tcf21 | 0.218955115 | 0.704074444 |
| ENSMUSG0000002345 | chr6 | Tpi1 | 0.168473665 | 0.704412438 |
| ENSMUSG0000002274 | chr16 | Cldnd1 | 0.182659464 | 0.704412438 |
| ENSMUSG0000003173 | chr8 | Dhodh | 0.192922137 | 0.704412438 |
| ENSMUSG0000003252 | chr9 | Ss18l2 | 0.175958344 | 0.704475615 |
| ENSMUSG0000004911 | chr4 | Fam110b | 0.180131503 | 0.704791497 |
| ENSMUSG0000006278 | chr7 | Kcnc3 | -0.243202208 | 0.705493459 |
| ENSMUSG0000000116 | chr5 | Oas1h | -0.73615295 | 0.705493459 |
| ENSMUSG0000003544 | chr9 | Thyn1 | 0.140313499 | 0.705493459 |
| ENSMUSG0000003834 | chr5 | Mlxip | 0.175996301 | 0.705493459 |
| ENSMUSG0000002491 | chr19 | Drap1 | 0.206529585 | 0.705493459 |
| ENSMUSG0000005467 | chr5 | Tmem119 | 0.273758712 | 0.705493459 |
| ENSMUSG0000002976 | chr6 | Akr1b8 | -0.233315481 | 0.705604292 |
| ENSMUSG0000006860 | chr18 | Gm4841 | -0.931818587 | 0.705604292 |
| ENSMUSG0000002925 | chr5 | Stap1 | -0.498499091 | 0.705604292 |
| ENSMUSG0000006672 | chr7 | Zfp575 | -0.299318348 | 0.705604292 |
| ENSMUSG0000002729 | chr2 | Vps39 | -0.133089764 | 0.705604292 |
| ENSMUSG0000004987 | chr4 | Rlf | 0.096266994 | 0.705604292 |
| ENSMUSG0000007488 | chr13 | Grk6 | 0.094122784 | 0.705604292 |
| ENSMUSG0000004672 | chr3 | Cdc42se1 | 0.112430028 | 0.705604292 |
| ENSMUSG0000005221 | chr7 | Opa3 | 0.129416996 | 0.705604292 |
| ENSMUSG0000002093 | chr11 | Dcakd | 0.127628707 | 0.705604292 |
| ENSMUSG0000002124 | chr12 | Mlh3 | 0.13779767 | 0.705604292 |
| ENSMUSG0000005740 | chr17 | Zfp53 | 0.208995755 | 0.705604292 |
| ENSMUSG0000005016 | chr15 | Mchr1 | 0.484428732 | 0.705604292 |
| ENSMUSG0000003358 | chr16 | Igf2bp2 | 0.187340617 | 0.705604292 |
| ENSMUSG0000002044 | chr11 | Npc1l1 | 1.019716147 | 0.705604292 |
| ENSMUSG0000003219 | chr9 | Gnb5 | -0.171656069 | 0.705892391 |
| ENSMUSG0000003683 | chr2 | Pnpla7 | -0.274277962 | 0.705923796 |
| ENSMUSG0000007848 | chr4 | Plekhn1 | -0.218377284 | 0.705923796 |
| ENSMUSG0000003731 | chr5 | Tacc3 | -0.291760405 | 0.705923796 |
| ENSMUSG0000006065 | chr16 | Marf1 | -0.125218701 | 0.705923796 |
| ENSMUSG0000002617 | chr1 | Rnf25 | 0.108430839 | 0.705923796 |
| ENSMUSG0000002235 | chr15 | Washc5 | 0.130892462 | 0.705923796 |
| ENSMUSG0000002268 | chr16 | Boc | 0.142580665 | 0.705923796 |

| | | | | |
|---|---|---|---|---|
| ENSMUSG0000005695 | chr6 | Tatdn2 | 0.133582495 | 0.705923796 |
| ENSMUSG0000004256 | chr15 | Fam227a | 0.179519162 | 0.705923796 |
| ENSMUSG0000001979 | chr10 | Mtres1 | 0.156974238 | 0.705923796 |
| ENSMUSG0000005380 | chr7 | Grwd1 | 0.199716855 | 0.705923796 |
| ENSMUSG0000005125 | chr6 | Jagn1 | 0.18879126 | 0.705923796 |
| ENSMUSG0000010314 | chr18 | Pcdhga1 | 0.278081736 | 0.705923796 |
| ENSMUSG0000000298 | chr7 | Relb | 0.243111478 | 0.705923796 |
| ENSMUSG0000001773 | chr2 | Dbndd2 | 0.280374703 | 0.705923796 |
| ENSMUSG0000009000 | chr18 | Ier3ip1 | 0.144953305 | 0.706068828 |
| ENSMUSG0000002760 | chr3 | Mtfr1 | 0.167253169 | 0.706068828 |
| ENSMUSG0000005984 | chr2 | Zfp341 | 0.236872443 | 0.706068828 |
| ENSMUSG0000002041 | chr11 | Hus1 | 0.10982965 | 0.706135809 |
| ENSMUSG0000002453 | chr18 | Psmg2 | 0.154532331 | 0.706135809 |
| ENSMUSG0000002245 | chr15 | Nell2 | 0.211303301 | 0.706175009 |
| ENSMUSG0000003535 | chr6 | Pdzrn3 | 0.236421277 | 0.706247407 |
| ENSMUSG0000002696 | chr2 | Arl6ip6 | 0.155394081 | 0.70634313 |
| ENSMUSG0000001761 | chr11 | Tnfaip1 | 0.099235345 | 0.706368207 |
| ENSMUSG0000003982 | chr2 | Trub2 | 0.144207518 | 0.706368207 |
| ENSMUSG0000002243 | chr15 | Sh3bp1 | -0.23039676 | 0.706600315 |
| ENSMUSG0000004199 | chr2 | Tlk1 | 0.093570013 | 0.706717234 |
| ENSMUSG0000004071 | chr11 | Camta2 | -0.136902896 | 0.706934292 |
| ENSMUSG0000002710 | chr2 | Ola1 | 0.122757012 | 0.706934292 |
| ENSMUSG0000000733 | chr19 | Mrpl49 | 0.161992188 | 0.706934292 |
| ENSMUSG0000003993 | chr5 | Gsap | -0.149070282 | 0.706986697 |
| ENSMUSG0000005555 | chr8 | Kxd1 | 0.146121336 | 0.706986697 |
| ENSMUSG0000004045 | chr15 | Cdh12 | -0.337322814 | 0.707117244 |
| ENSMUSG0000001997 | chr10 | Sgk1 | -0.298725114 | 0.707117244 |
| ENSMUSG0000002045 | chr11 | Ogdh | 0.086025225 | 0.707117244 |
| ENSMUSG0000004739 | chr15 | Cimap1b | -0.437434218 | 0.707372825 |
| ENSMUSG0000007644 | chr2 | Ass1 | -0.182574766 | 0.707372825 |
| ENSMUSG0000004638 | chr18 | Pcdhb17 | 0.194097406 | 0.707564382 |
| ENSMUSG0000006820 | chr15 | Pick1 | -0.156857387 | 0.707743469 |
| ENSMUSG0000005840 | chr7 | Zfp420 | 0.184690642 | 0.707743469 |
| ENSMUSG0000003693 | chr8 | 4921524J17Rik | 0.150380516 | 0.708060687 |
| ENSMUSG0000002423 | chr18 | Mtpap | 0.129294487 | 0.708233068 |
| ENSMUSG0000002069 | chr11 | Mrc2 | 0.199654949 | 0.708489774 |
| ENSMUSG0000002101 | chr12 | Zc3h14 | 0.107679797 | 0.708583701 |
| ENSMUSG0000002585 | chr5 | Dnaaf5 | 0.129056179 | 0.708583701 |
| ENSMUSG0000002425 | chr17 | Dync2li1 | 0.150989507 | 0.708583701 |
| ENSMUSG0000002407 | chr17 | Ttc27 | 0.184426054 | 0.708583701 |
| ENSMUSG0000003231 | chr9 | Etfa | 0.161113066 | 0.70858871 |
| ENSMUSG0000003853 | chr2 | Cbfa2t2 | 0.101052926 | 0.708713984 |
| ENSMUSG0000006958 | chr10 | Tspear | -0.480186029 | 0.709034453 |
| ENSMUSG0000002865 | chr4 | Ppt1 | 0.103209531 | 0.709034453 |
| ENSMUSG0000005241 | chr6 | Smim30 | 0.138371641 | 0.709034453 |
| ENSMUSG0000004076 | chr11 | Snrnp25 | 0.285354622 | 0.709034453 |
| ENSMUSG0000002286 | chr16 | Cxadr | -0.556668467 | 0.709044079 |
| ENSMUSG0000007464 | chr2 | 6430550D23Rik | -0.362825446 | 0.709044079 |
| ENSMUSG0000002640 | chr1 | Pfkfb2 | -0.196488498 | 0.709044079 |
| ENSMUSG0000005914 | chr17 | Zfp945 | -0.136303975 | 0.709044079 |

| | | | | |
|---|---|---|---|---|
| ENSMUSG0000003287 | chr4 | Smap2 | 0.081099696 | 0.709044079 |
| ENSMUSG0000001115 | chr12 | Brf1 | 0.110914275 | 0.709044079 |
| ENSMUSG0000003152 | chr8 | Eri1 | 0.14801507 | 0.709044079 |
| ENSMUSG0000003238 | chr9 | Trip4 | 0.090061917 | 0.709098612 |
| ENSMUSG0000007944 | chr1 | Alpi | -1.708172051 | 0.709109933 |
| ENSMUSG0000003014 | chr6 | Clec4b1 | -1.088435245 | 0.709139123 |
| ENSMUSG0000000115 | chr6 | Gmcl1 | -0.124979481 | 0.709191887 |
| ENSMUSG0000002267 | chr16 | Nde1 | 0.093692568 | 0.709191887 |
| ENSMUSG0000005639 | chr7 | Lig1 | -0.241935318 | 0.709943988 |
| ENSMUSG0000001594 | chr5 | Castor2 | -0.118882669 | 0.710043129 |
| ENSMUSG0000002886 | chr4 | Mrps15 | 0.082279301 | 0.710043129 |
| ENSMUSG0000003087 | chr7 | Scnn1b | -0.910003345 | 0.710175796 |
| ENSMUSG0000002916 | chr5 | Dhx15 | 0.096032097 | 0.710175796 |
| ENSMUSG0000000550 | chr2 | Kbtbd4 | 0.183094587 | 0.710175796 |
| ENSMUSG0000007420 | chr3 | Adh1 | 0.507113395 | 0.710175796 |
| ENSMUSG0000002540 | chr10 | Nab2 | 0.232104641 | 0.710465012 |
| ENSMUSG0000003176 | chr8 | Mt2 | -0.563621319 | 0.710573641 |
| ENSMUSG0000002518 | chr19 | Hps1 | -0.177257235 | 0.710661715 |
| ENSMUSG0000002975 | chr6 | Dync1i1 | -0.200748536 | 0.710661715 |
| ENSMUSG0000003200 | chr9 | Sesn3 | -0.137406551 | 0.710661715 |
| ENSMUSG0000004111 | chrX | Iqsec2 | -0.136480692 | 0.710661715 |
| ENSMUSG0000007341 | chr17 | Mpig6b | -0.751633743 | 0.710661715 |
| ENSMUSG0000003346 | chrX | Armcx1 | -0.102460139 | 0.710661715 |
| ENSMUSG0000000520 | chr11 | Senp3 | 0.080399873 | 0.710661715 |
| ENSMUSG0000002216 | chr14 | Mettl3 | 0.126169647 | 0.710661715 |
| ENSMUSG0000005331 | chr9 | Nrgn | 0.173301609 | 0.710661715 |
| ENSMUSG0000002272 | chr16 | Riox2 | 0.214727114 | 0.710661715 |
| ENSMUSG0000002250 | chr16 | Bcl6 | 0.27543985 | 0.710661715 |
| ENSMUSG0000003821 | chr10 | Bend3 | -0.196469949 | 0.710725013 |
| ENSMUSG0000007457 | chr2 | Ripor3 | -0.285779653 | 0.710923331 |
| ENSMUSG0000003656 | chr17 | Bicral | -0.149379133 | 0.710923331 |
| ENSMUSG0000002199 | chr14 | Esd | 0.157637021 | 0.711368885 |
| ENSMUSG0000004289 | chr15 | Abra | 0.417570815 | 0.71171253 |
| ENSMUSG0000004766 | chr10 | Msl3l2 | -0.227057525 | 0.712027802 |
| ENSMUSG0000003873 | chr13 | Mboat1 | -0.250485642 | 0.71203102 |
| ENSMUSG0000003402 | chr18 | Cd226 | -0.420834934 | 0.71203102 |
| ENSMUSG0000003273 | chr7 | Inppl1 | -0.144425252 | 0.71203102 |
| ENSMUSG0000002508 | chr19 | Tdrd1 | -1.038112008 | 0.71203102 |
| ENSMUSG0000002239 | chr15 | L3mbtl2 | -0.150718636 | 0.71203102 |
| ENSMUSG0000002814 | chr3 | Rap1gds1 | -0.135553177 | 0.71203102 |
| ENSMUSG0000001584 | chr2 | Rxra | -0.095908629 | 0.71203102 |
| ENSMUSG0000001746 | chr11 | Timp2 | 0.07709809 | 0.71203102 |
| ENSMUSG0000005131 | chr18 | Taf7 | 0.163016279 | 0.71203102 |
| ENSMUSG0000001987 | chr10 | Hsf2 | 0.116389269 | 0.712389192 |
| ENSMUSG0000003618 | chr2 | Dipk1b | -0.197108946 | 0.712564259 |
| ENSMUSG0000005764 | chr13 | Brd9 | -0.089529729 | 0.712687602 |
| ENSMUSG0000004320 | chr4 | Zmpste24 | 0.094666568 | 0.712687602 |
| ENSMUSG0000002662 | chr1 | Atf3 | 0.478143777 | 0.712687602 |
| ENSMUSG0000002700 | chr2 | Nckap1 | 0.076723307 | 0.713004382 |
| ENSMUSG0000004585 | chr4 | Lyrm2 | 0.12975401 | 0.713004382 |

| | | | | |
|---|---|---|---|---|
| ENSMUSG0000003761 | chr15 | Spag1 | 0.160621793 | 0.713004382 |
| ENSMUSG0000004179 | chr9 | Myrip | -0.106235133 | 0.713342005 |
| ENSMUSG0000004252 | chr15 | Sun2 | 0.132256266 | 0.713342005 |
| ENSMUSG0000000035 | chrX | Mcts1 | 0.134313181 | 0.713342005 |
| ENSMUSG0000005450 | chr14 | Parp4 | -0.180146365 | 0.713624032 |
| ENSMUSG0000003150 | chr8 | Naxd | 0.148427682 | 0.713774462 |
| ENSMUSG0000003485 | chr9 | Atosa | 0.240713596 | 0.713774462 |
| ENSMUSG0000000658 | chr8 | Cdt1 | -0.39795802 | 0.713816434 |
| ENSMUSG0000002496 | chr19 | Stip1 | 0.22772947 | 0.714004414 |
| ENSMUSG0000003006 | chr6 | Foxp1 | 0.132272659 | 0.714048305 |
| ENSMUSG0000007224 | chr7 | Trim6 | -0.653745338 | 0.714320709 |
| ENSMUSG0000004064 | chr1 | Ppip5k2 | -0.183863796 | 0.714320709 |
| ENSMUSG0000002479 | chr19 | Zfpl1 | -0.196610902 | 0.714320709 |
| ENSMUSG0000004563 | chr8 | Mtus1 | 0.089611348 | 0.714320709 |
| ENSMUSG0000005066 | chr17 | Gpatch11 | 0.123499554 | 0.714320709 |
| ENSMUSG0000004696 | chr16 | Zbtb21 | 0.140636351 | 0.714320709 |
| ENSMUSG0000002450 | chr18 | Dtwd2 | 0.260520114 | 0.714320709 |
| ENSMUSG0000002649 | chr1 | Tfb2m | 0.174866447 | 0.714320709 |
| ENSMUSG0000003594 | chr15 | Ttc38 | -0.222257373 | 0.714604856 |
| ENSMUSG0000010431 | chr18 | Pcdha7 | -0.62951907 | 0.714617496 |
| ENSMUSG0000001729 | chr2 | Dnttip1 | 0.125210785 | 0.714784266 |
| ENSMUSG0000002280 | chr16 | Cfap91 | -0.280118906 | 0.71492653 |
| ENSMUSG0000002082 | chr11 | Mink1 | -0.074728334 | 0.71492653 |
| ENSMUSG0000003009 | chr6 | Cntn6 | -0.708297524 | 0.715094344 |
| ENSMUSG0000002285 | chr16 | Tra2b | 0.131171967 | 0.715340377 |
| ENSMUSG0000004747 | chr7 | Zfp30 | 0.170108114 | 0.715340377 |
| ENSMUSG0000003138 | chrX | Renbp | 0.176566783 | 0.715767618 |
| ENSMUSG0000005543 | chr8 | Maf | 0.214596181 | 0.716489033 |
| ENSMUSG0000000582 | chr17 | Gpr108 | -0.109266061 | 0.716514327 |
| ENSMUSG0000002751 | chr2 | Rab22a | 0.06687117 | 0.716613459 |
| ENSMUSG0000003041 | chr7 | Nova2 | -0.261061536 | 0.716653099 |
| ENSMUSG0000002246 | chr15 | Srebf2 | 0.211895765 | 0.716690097 |
| ENSMUSG0000002417 | chr17 | Pot1b | -0.187435127 | 0.716859376 |
| ENSMUSG0000004069 | chr4 | Col16a1 | 0.187544709 | 0.717024401 |
| ENSMUSG0000000298 | chr7 | Clptm1 | -0.092858334 | 0.717087248 |
| ENSMUSG0000007294 | chrX | Nup62cl | -0.719173814 | 0.717116412 |
| ENSMUSG0000000918 | chr11 | Ccl8 | -0.491901578 | 0.717116412 |
| ENSMUSG0000004696 | chr16 | Gpr156 | -0.486276973 | 0.717116412 |
| ENSMUSG0000005645 | chr12 | Mok | -0.17326506 | 0.717116412 |
| ENSMUSG0000000080 | chr11 | Usp32 | -0.127367256 | 0.717116412 |
| ENSMUSG0000004000 | chr2 | Bahd1 | -0.106524762 | 0.717116412 |
| ENSMUSG0000000580 | chr2 | Slc30a4 | -0.100511595 | 0.717116412 |
| ENSMUSG0000001571 | chr3 | Prune1 | -0.09055232 | 0.717116412 |
| ENSMUSG0000001742 | chr11 | Psmd11 | 0.076530151 | 0.717116412 |
| ENSMUSG0000003476 | chr12 | Map4k5 | 0.096344291 | 0.717116412 |
| ENSMUSG0000003508 | chr11 | Becn1 | 0.10407552 | 0.717116412 |
| ENSMUSG0000002195 | chr14 | Xpo4 | 0.122876311 | 0.717116412 |
| ENSMUSG0000003020 | chr6 | Ddx47 | 0.130691776 | 0.717116412 |
| ENSMUSG0000002529 | chr14 | Rps24 | 0.210083681 | 0.717116412 |
| ENSMUSG0000005078 | chr6 | Ccdc126 | 0.193649729 | 0.717116412 |

| | | | | |
|---|---|---|---|---|
| ENSMUSG0000009241 | chr17 | Gpank1 | 0.199245312 | 0.717116412 |
| ENSMUSG0000007042 | chr7 | Or51e1 | 0.570395048 | 0.717116412 |
| ENSMUSG0000002106 | chr12 | Pygl | 0.21810545 | 0.717116412 |
| ENSMUSG0000003268 | chr2 | 6820408C15Rik | -0.514899856 | 0.717316031 |
| ENSMUSG0000001980 | chr10 | Pex3 | -0.114677928 | 0.717316031 |
| ENSMUSG0000003303 | chr15 | Micall1 | 0.148324819 | 0.717316031 |
| ENSMUSG0000007844 | chr10 | Dohh | 0.258303438 | 0.717525071 |
| ENSMUSG0000005420 | chr12 | Alkal2 | -0.603378183 | 0.71759601 |
| ENSMUSG0000002852 | chr4 | Ak4 | 0.114751529 | 0.71759601 |
| ENSMUSG0000003229 | chr9 | Ptpn9 | -0.15488854 | 0.717762843 |
| ENSMUSG0000005835 | chr14 | Uqcc5 | -0.206270528 | 0.717762843 |
| ENSMUSG0000003461 | chr4 | Tut4 | -0.128612348 | 0.717762843 |
| ENSMUSG0000003293 | chr16 | Hspa13 | -0.093601421 | 0.717762843 |
| ENSMUSG0000004272 | chr5 | Trafd1 | 0.093386073 | 0.717762843 |
| ENSMUSG0000000113 | chrX | Timp1 | 0.242590153 | 0.717762843 |
| ENSMUSG0000005197 | chr8 | Erich1 | 0.207103742 | 0.717762843 |
| ENSMUSG0000002404 | chr17 | Arhgap28 | 0.243267946 | 0.717762843 |
| ENSMUSG0000003487 | chr4 | Fam151a | -0.865568971 | 0.71836483 |
| ENSMUSG0000006417 | chr6 | Ghrl | -0.497070379 | 0.71836483 |
| ENSMUSG0000002721 | chr2 | Slc28a2 | -0.355063211 | 0.71836483 |
| ENSMUSG0000003115 | chrX | Ccdc120 | -0.190859655 | 0.71836483 |
| ENSMUSG0000004157 | chr1 | Camsap2 | 0.111224676 | 0.71836483 |
| ENSMUSG0000002162 | chr13 | Zcchc9 | 0.124071659 | 0.71836483 |
| ENSMUSG0000002527 | chr14 | Flnb | 0.144191507 | 0.71836483 |
| ENSMUSG0000003837 | chr3 | Rbm8a | 0.138861553 | 0.71836483 |
| ENSMUSG0000002730 | chr2 | Ndufaf1 | 0.174524958 | 0.71836483 |
| ENSMUSG0000002607 | chr1 | Il1r1 | 0.168591614 | 0.71836483 |
| ENSMUSG0000003432 | chr5 | Tmem132c | 0.278330236 | 0.71836483 |
| ENSMUSG0000002535 | chr10 | Dnajc14 | 0.087397805 | 0.718392677 |
| ENSMUSG0000005214 | chr17 | Rasal3 | -0.581856256 | 0.718405328 |
| ENSMUSG0000002654 | chr1 | Dusp23 | -0.213374034 | 0.718545686 |
| ENSMUSG0000002918 | chr5 | Sel1l3 | -0.210538237 | 0.718561009 |
| ENSMUSG0000004553 | chr6 | Kcna5 | -0.14219587 | 0.718561009 |
| ENSMUSG0000007342 | chr17 | Zfp414 | -0.127919601 | 0.718561009 |
| ENSMUSG0000003197 | chr8 | Cog2 | 0.122309443 | 0.718561009 |
| ENSMUSG0000002411 | chr17 | Cacna1h | 0.130618432 | 0.718561009 |
| ENSMUSG0000005430 | chr12 | Eapp | 0.140928688 | 0.718561009 |
| ENSMUSG0000002085 | chr11 | Nme2 | 0.328884055 | 0.718561009 |
| ENSMUSG0000003898 | chr2 | Cfap157 | -0.300510176 | 0.718599432 |
| ENSMUSG0000003072 | chr7 | Cln3 | -0.199725127 | 0.718599432 |
| ENSMUSG0000004160 | chr18 | Mbp | -0.134539862 | 0.718599432 |
| ENSMUSG0000003668 | chr8 | Cc2d1a | -0.114696851 | 0.718599432 |
| ENSMUSG0000003144 | chr8 | Lamp1 | 0.073106751 | 0.718599432 |
| ENSMUSG0000003586 | chr10 | Palm | 0.089731754 | 0.718599432 |
| ENSMUSG0000001770 | chr2 | Serinc3 | 0.113947766 | 0.718599432 |
| ENSMUSG0000003182 | chr8 | Zdhhc7 | 0.104191495 | 0.718599432 |
| ENSMUSG0000003807 | chr1 | Tmem237 | 0.10148316 | 0.718599432 |
| ENSMUSG0000000136 | chr9 | Fbxo9 | 0.122375054 | 0.718599432 |
| ENSMUSG0000000381 | chr8 | Calr | 0.257741067 | 0.718599432 |
| ENSMUSG0000003264 | chr19 | Pygm | 0.173682971 | 0.718599432 |

| | | | | |
|---|---|---|---|---|
| ENSMUSG0000003354 | chr5 | Idua | 0.145532013 | 0.718645783 |
| ENSMUSG0000002482 | chr19 | Dpf2 | 0.087041731 | 0.718835996 |
| ENSMUSG0000004166 | chr12 | Prima1 | 0.18295072 | 0.718835996 |
| ENSMUSG0000002166 | chr13 | Gfm2 | -0.115191372 | 0.719332661 |
| ENSMUSG0000008500 | chr3 | Smim43 | 0.609502157 | 0.719332661 |
| ENSMUSG0000003713 | chr4 | Myom3 | -0.419311753 | 0.719712523 |
| ENSMUSG0000002302 | chr15 | Atf1 | 0.140964964 | 0.719870533 |
| ENSMUSG0000003827 | chr5 | Asic3 | 0.546034854 | 0.71990985 |
| ENSMUSG0000002802 | chr3 | Enpep | 0.171184139 | 0.71990985 |
| ENSMUSG0000003667 | chr15 | Aaas | -0.230158899 | 0.719936578 |
| ENSMUSG0000000166 | chr10 | Gstt3 | -0.158175513 | 0.719936578 |
| ENSMUSG0000002402 | chr17 | Glp1r | -0.253102426 | 0.720170864 |
| ENSMUSG0000005833 | chr13 | Zfp85 | 0.172984981 | 0.720170864 |
| ENSMUSG0000002717 | chr2 | Prrg4 | -0.242637534 | 0.720293269 |
| ENSMUSG0000002494 | chr19 | Men1 | -0.103662065 | 0.720293269 |
| ENSMUSG0000006991 | chr11 | Hba-a1 | -0.499514948 | 0.720386882 |
| ENSMUSG0000003757 | chr11 | Tob1 | -0.162572062 | 0.720386882 |
| ENSMUSG0000002018 | chr10 | Mdm2 | 0.103381191 | 0.720386882 |
| ENSMUSG0000002596 | chr1 | Mdh1b | -1.184878655 | 0.720502716 |
| ENSMUSG0000003005 | chr6 | Cnbp | 0.134272772 | 0.720502716 |
| ENSMUSG0000004506 | chr18 | Pcdhb7 | -0.323570242 | 0.720629116 |
| ENSMUSG0000002220 | chr14 | Efs | -0.186822708 | 0.720994634 |
| ENSMUSG0000003767 | chr9 | Rfx7 | -0.1718412 | 0.721079463 |
| ENSMUSG0000001777 | chr11 | Crk | 0.101127735 | 0.721079463 |
| ENSMUSG0000003483 | chr9 | Larp6 | 0.211196789 | 0.721079463 |
| ENSMUSG0000000328 | chr4 | Plag1 | -0.254878081 | 0.721236963 |
| ENSMUSG0000007434 | chr3 | Ovgp1 | -0.3668674 | 0.721236963 |
| ENSMUSG0000008759 | chr7 | Zfp111 | -0.148834692 | 0.721236963 |
| ENSMUSG0000002067 | chr12 | Pxdn | -0.101187842 | 0.721236963 |
| ENSMUSG0000003595 | chr14 | Pip4p1 | -0.088831951 | 0.721236963 |
| ENSMUSG0000006216 | chr1 | Cnih4 | 0.129935295 | 0.721236963 |
| ENSMUSG0000000201 | chrX | Bcap31 | 0.158951233 | 0.721236963 |
| ENSMUSG0000006927 | chr13 | H2ac8 | 0.757907663 | 0.721236963 |
| ENSMUSG0000002717 | chr2 | Tcp11l1 | -0.215943659 | 0.721468919 |
| ENSMUSG0000001103 | chr5 | Slc5a1 | -1.023445699 | 0.721687997 |
| ENSMUSG0000006352 | chr15 | Ly6m | -1.19534012 | 0.721687997 |
| ENSMUSG0000002301 | chr15 | Asic1 | -0.228656196 | 0.721687997 |
| ENSMUSG0000005440 | chr8 | Cklf | -0.211681469 | 0.721687997 |
| ENSMUSG0000004798 | chr18 | Ino80c | -0.15345464 | 0.721687997 |
| ENSMUSG0000003378 | chr13 | Asb13 | -0.14167404 | 0.721687997 |
| ENSMUSG0000000313 | chr9 | Pafah1b2 | -0.1095497 | 0.721687997 |
| ENSMUSG0000002189 | chr14 | Capn7 | -0.0900354 | 0.721687997 |
| ENSMUSG0000003681 | chr5 | Sun1 | 0.091982375 | 0.721687997 |
| ENSMUSG0000004994 | chr3 | Pgrmc2 | 0.107925568 | 0.721687997 |
| ENSMUSG0000004399 | chr12 | Mgat2 | 0.137271239 | 0.721687997 |
| ENSMUSG0000000323 | chr16 | Dvl3 | 0.167175429 | 0.721687997 |
| ENSMUSG0000007097 | chr4 | Dnajc25 | 0.14969499 | 0.721687997 |
| ENSMUSG0000004533 | chr8 | Zfp423 | 0.16409272 | 0.721687997 |
| ENSMUSG0000007300 | chr6 | Vamp5 | 0.259642509 | 0.721687997 |
| ENSMUSG0000002846 | chr4 | Ccdc107 | 0.251944112 | 0.721687997 |

| | | | | |
|---|---|---|---|---|
| ENSMUSG0000006412 | chr17 | Mocs1 | -0.135023961 | 0.721876964 |
| ENSMUSG0000002909 | chr5 | Adgra3 | 0.09340438 | 0.72190465 |
| ENSMUSG0000002650 | chr1 | Cox20 | -1.112019309 | 0.721915787 |
| ENSMUSG0000003799 | chr3 | Slc9b2 | -0.29651812 | 0.721915787 |
| ENSMUSG0000003149 | chr8 | Tnfsf13b | -0.252794432 | 0.721915787 |
| ENSMUSG0000002049 | chr11 | Rnf187 | 0.134052753 | 0.721915787 |
| ENSMUSG0000003241 | chr9 | Ube3d | -0.438363709 | 0.7219176 |
| ENSMUSG0000001477 | chr17 | Dll1 | -0.288252972 | 0.721949782 |
| ENSMUSG0000000647 | chr2 | Nsmf | -0.112016074 | 0.722016863 |
| ENSMUSG0000003972 | chr2 | Trp53rka | 0.180047793 | 0.722016863 |
| ENSMUSG0000004199 | chr13 | Zbed3 | 0.17142775 | 0.722051009 |
| ENSMUSG0000004067 | chr10 | Mthfd1l | 0.510550454 | 0.722060654 |
| ENSMUSG0000004412 | chr11 | Proca1 | -0.254337724 | 0.722068025 |
| ENSMUSG0000007560 | chr15 | Ly6a | -0.178655535 | 0.722068025 |
| ENSMUSG0000003397 | chr5 | Rfc3 | -0.204406211 | 0.722068025 |
| ENSMUSG0000006211 | chr11 | Rai1 | -0.145039501 | 0.722068025 |
| ENSMUSG0000004111 | chr17 | Pde9a | -0.159623239 | 0.722068025 |
| ENSMUSG0000003994 | chr15 | Ptger4 | -0.139671854 | 0.722068025 |
| ENSMUSG0000006219 | chr12 | 2700097O09Rik | -0.147053074 | 0.722068025 |
| ENSMUSG0000003655 | chr8 | Cnot1 | -0.112149152 | 0.722068025 |
| ENSMUSG0000002127 | chr12 | Tecpr2 | -0.109664575 | 0.722068025 |
| ENSMUSG0000002634 | chr1 | Actr3 | 0.113461148 | 0.722068025 |
| ENSMUSG0000004232 | chr14 | Pbrm1 | 0.132519371 | 0.722068025 |
| ENSMUSG0000003735 | chr15 | Letmd1 | 0.145452199 | 0.722068025 |
| ENSMUSG0000002128 | chr12 | Zfyve21 | 0.19987037 | 0.722068025 |
| ENSMUSG0000002452 | chr18 | Afg3l2 | -0.102698573 | 0.722492751 |
| ENSMUSG0000002789 | chr3 | Slc6a17 | 0.177933536 | 0.722492751 |
| ENSMUSG0000002401 | chr17 | Pim1 | 0.21530157 | 0.722492751 |
| ENSMUSG0000005689 | chr11 | H2bc27 | -0.34127213 | 0.722539374 |
| ENSMUSG0000005548 | chr7 | Ano5 | -0.213846419 | 0.722539374 |
| ENSMUSG0000007849 | chr4 | Zfp982 | -0.214962098 | 0.722539374 |
| ENSMUSG0000000783 | chr7 | Prrg2 | -0.166081262 | 0.722539374 |
| ENSMUSG0000002001 | chr10 | Cdk17 | 0.098530648 | 0.722539374 |
| ENSMUSG0000001487 | chr2 | Surf2 | 0.135990141 | 0.722539374 |
| ENSMUSG0000003789 | chr3 | H2az1 | 0.170400152 | 0.722539374 |
| ENSMUSG0000004783 | chr12 | Cdca4 | 0.194905293 | 0.722539374 |
| ENSMUSG0000004447 | chr10 | Ascc1 | 0.172045353 | 0.722539374 |
| ENSMUSG0000005135 | chr11 | Commd1 | 0.170187271 | 0.722635639 |
| ENSMUSG0000007426 | chr3 | Amy1 | -0.297750697 | 0.722873989 |
| ENSMUSG0000000152 | chr6 | Tulp3 | 0.147865586 | 0.722873989 |
| ENSMUSG0000002249 | chr16 | Txndc11 | 0.106797637 | 0.723115053 |
| ENSMUSG0000002947 | chr5 | P2rx4 | -0.158728434 | 0.723125148 |
| ENSMUSG0000003052 | chr7 | Chrna7 | 0.274588879 | 0.723125148 |
| ENSMUSG0000003100 | chr7 | Mki67 | -0.781571274 | 0.723561836 |
| ENSMUSG0000005569 | chr16 | Tmem191 | -0.377210267 | 0.723561836 |
| ENSMUSG0000003762 | chr15 | Rgs22 | -0.508889065 | 0.723561836 |
| ENSMUSG0000004903 | chr6 | Clec4a1 | -0.284302732 | 0.723561836 |
| ENSMUSG0000003122 | chrX | Magee1 | -0.135093447 | 0.723561836 |
| ENSMUSG0000003042 | chr7 | Pop4 | 0.138248724 | 0.723561836 |
| ENSMUSG0000003990 | chr11 | Slc26a11 | 0.200202126 | 0.723561836 |

| | | | | |
|---|---|---|---|---|
| ENSMUSG0000003132 | chrX | Dmrtc1a | -0.47099853 | 0.723694723 |
| ENSMUSG0000001946 | chr10 | Arhgef25 | 0.089826961 | 0.723694723 |
| ENSMUSG0000003001 | chr6 | Fbxl14 | -0.175264088 | 0.724002476 |
| ENSMUSG0000002916 | chr5 | Agbl5 | -0.160955201 | 0.724002476 |
| ENSMUSG0000002150 | chr13 | B4galt7 | 0.135877062 | 0.724002476 |
| ENSMUSG0000002878 | chr4 | Adgrb2 | -0.232875824 | 0.72401261 |
| ENSMUSG0000002758 | chr2 | Pcmtd2 | -0.15316791 | 0.724184163 |
| ENSMUSG0000002040 | chr11 | Slu7 | 0.122692482 | 0.724184163 |
| ENSMUSG0000002825 | chr4 | Ccnc | 0.108547283 | 0.724264334 |
| ENSMUSG0000002863 | chr4 | Ybx1 | 0.157113194 | 0.724264334 |
| ENSMUSG0000002740 | chr2 | Cpxm1 | -0.325524606 | 0.724398264 |
| ENSMUSG0000004082 | chr11 | Zmynd15 | -0.3005776 | 0.724398264 |
| ENSMUSG0000003929 | chr4 | Cdk5rap2 | -0.151314996 | 0.724398264 |
| ENSMUSG0000003523 | chr5 | Abraxas1 | 0.145332621 | 0.724398264 |
| ENSMUSG0000006611 | chr4 | Adamtsl1 | 0.145086783 | 0.724398264 |
| ENSMUSG0000006885 | chr3 | H2bc21 | -0.126858288 | 0.724597906 |
| ENSMUSG0000002045 | chr11 | Drg1 | 0.133771282 | 0.724597906 |
| ENSMUSG0000002167 | chr13 | F2rl1 | -0.280867118 | 0.724654565 |
| ENSMUSG0000004209 | chr6 | Zfp239 | -0.377023999 | 0.724654565 |
| ENSMUSG0000003481 | chr10 | Grip1 | -0.237686202 | 0.724654565 |
| ENSMUSG0000005883 | chr2 | Abi1 | 0.074251827 | 0.724654565 |
| ENSMUSG0000002615 | chr1 | Smap1 | 0.103679616 | 0.724654565 |
| ENSMUSG0000002220 | chr15 | Zfr | 0.120128531 | 0.724654565 |
| ENSMUSG0000005844 | chr6 | Nrf1 | 0.122075858 | 0.724654565 |
| ENSMUSG0000002448 | chr18 | Yipf5 | 0.124589161 | 0.724654565 |
| ENSMUSG0000002857 | chr4 | Caap1 | 0.149156906 | 0.724654565 |
| ENSMUSG0000002357 | chr4 | C1qtnf12 | 0.136027002 | 0.724654565 |
| ENSMUSG0000006235 | chr12 | Itgb1bp1 | 0.181039345 | 0.724654565 |
| ENSMUSG0000003804 | chr11 | Mrm3 | 0.151926117 | 0.724898028 |
| ENSMUSG0000003873 | chr1 | Wdr26 | 0.106385022 | 0.724913619 |
| ENSMUSG0000005850 | chr5 | Fam133b | 0.147500328 | 0.725555405 |
| ENSMUSG0000003000 | chr6 | Pradc1 | -0.20319963 | 0.725695717 |
| ENSMUSG0000003883 | chr17 | Vars2 | 0.126548597 | 0.725695717 |
| ENSMUSG0000002015 | chr10 | Pwwp3a | -0.126999296 | 0.72608386 |
| ENSMUSG0000003935 | chr14 | Fut11 | 0.090355615 | 0.726207518 |
| ENSMUSG0000005167 | chr8 | Coa6 | 0.206561626 | 0.726331764 |
| ENSMUSG0000003003 | chr6 | 1700003E16Rik | -0.291071821 | 0.726360953 |
| ENSMUSG0000003567 | chr8 | Zswim4 | -0.153334537 | 0.726360953 |
| ENSMUSG0000007493 | chr2 | Grem1 | 0.148960007 | 0.726360953 |
| ENSMUSG0000002657 | chr1 | Atp1b1 | -0.155720801 | 0.726622519 |
| ENSMUSG0000002510 | chr7 | 3110040N11Rik | 0.179287774 | 0.726622519 |
| ENSMUSG0000002787 | chr3 | Hmgcs2 | -0.590068506 | 0.726745704 |
| ENSMUSG0000002916 | chr5 | Ppargc1a | -0.230656112 | 0.726745704 |
| ENSMUSG0000003854 | chr8 | Pcid2 | 0.141590247 | 0.726745704 |
| ENSMUSG0000002720 | chr2 | Galk2 | -0.127732383 | 0.726768215 |
| ENSMUSG0000001454 | chr14 | Wdfy2 | -0.132650443 | 0.726848258 |
| ENSMUSG0000003232 | chr9 | Stra6 | -0.287964035 | 0.726899356 |
| ENSMUSG0000005824 | chr1 | Kcnh1 | -0.236119192 | 0.726899356 |
| ENSMUSG0000004210 | chr7 | Inpp5f | -0.139673793 | 0.726899356 |
| ENSMUSG0000002442 | chr17 | Atat1 | -0.144984574 | 0.726899356 |

| | | | | |
|---|---|---|---|---|
| ENSMUSG0000003425 | chr15 | Exosc4 | 0.153309623 | 0.726899356 |
| ENSMUSG0000003371 | chr12 | Foxn3 | 0.169698237 | 0.726899356 |
| ENSMUSG0000006689 | chr5 | Vsig10 | -0.24447566 | 0.727134313 |
| ENSMUSG0000003049 | chr7 | Gas2 | -0.292775244 | 0.727135729 |
| ENSMUSG0000004209 | chr17 | Kank3 | 0.186641171 | 0.727257703 |
| ENSMUSG0000004274 | chr5 | Hectd4 | -0.169587943 | 0.727273019 |
| ENSMUSG0000002607 | chr1 | Map4k4 | 0.110960532 | 0.727273197 |
| ENSMUSG0000001036 | chr11 | Rdm1 | 0.21785434 | 0.727273197 |
| ENSMUSG0000003659 | chr8 | Ccdc113 | -0.336662831 | 0.727327899 |
| ENSMUSG0000009007 | chr2 | Lime1 | -0.678064038 | 0.727774993 |
| ENSMUSG0000002611 | chr1 | Unc50 | 0.163776226 | 0.727892693 |
| ENSMUSG0000004013 | chrX | Ndp | -0.424718081 | 0.728128713 |
| ENSMUSG0000003107 | chr7 | Fadd | -0.133639596 | 0.728128713 |
| ENSMUSG0000005987 | chr6 | Zfp422 | 0.123398571 | 0.728155194 |
| ENSMUSG0000001559 | chr17 | Zfp318 | -0.17813142 | 0.728286075 |
| ENSMUSG0000002565 | chrX | Arhgef9 | -0.1019396 | 0.728286075 |
| ENSMUSG0000003857 | chr1 | Susd4 | 0.161813297 | 0.728286075 |
| ENSMUSG0000007107 | chr8 | Jund | 0.143138449 | 0.728286075 |
| ENSMUSG0000003377 | chrX | Tlr13 | 0.401017233 | 0.728286075 |
| ENSMUSG0000000637 | chr15 | Gcat | 0.416562041 | 0.728286075 |
| ENSMUSG0000002886 | chr4 | Gnl2 | 0.086037601 | 0.728312269 |
| ENSMUSG0000002026 | chr10 | Sumo3 | 0.157125251 | 0.728312269 |
| ENSMUSG0000005440 | chr11 | Slfn5 | -0.256962851 | 0.728385329 |
| ENSMUSG0000005858 | chr10 | Anks1b | 0.149013808 | 0.72869355 |
| ENSMUSG0000002262 | chr15 | Rabl2 | 0.105442754 | 0.728699486 |
| ENSMUSG0000004234 | chr1 | Ikbke | -0.244843452 | 0.728725959 |
| ENSMUSG0000002218 | chr15 | Oxct1 | 0.120130236 | 0.728725959 |
| ENSMUSG0000003000 | chr6 | Cct7 | 0.168353794 | 0.728725959 |
| ENSMUSG0000003178 | chr8 | Coq9 | 0.138686801 | 0.728725959 |
| ENSMUSG0000006201 | chr17 | Zfp13 | 0.163191735 | 0.728725959 |
| ENSMUSG0000001979 | chr10 | Pcmt1 | 0.136782417 | 0.728945126 |
| ENSMUSG0000001004 | chr9 | Hyal2 | 0.115150799 | 0.728945126 |
| ENSMUSG0000004008 | chr3 | Tnni3k | 0.573649062 | 0.729250107 |
| ENSMUSG0000001994 | chr10 | Arid5b | 0.125300146 | 0.729316556 |
| ENSMUSG0000002523 | chr9 | Parp6 | -0.090423205 | 0.729401367 |
| ENSMUSG0000001011 | chr19 | Stx5a | 0.127635167 | 0.729401367 |
| ENSMUSG0000003616 | chr10 | Ccdc38 | -0.558695663 | 0.729513769 |
| ENSMUSG0000002094 | chr11 | Gosr2 | 0.085611217 | 0.729513769 |
| ENSMUSG0000002425 | chr18 | Slc25a46 | -0.066936515 | 0.729612223 |
| ENSMUSG0000004261 | chr4 | Oscp1 | -0.136531147 | 0.729790881 |
| ENSMUSG0000003468 | chr5 | Fras1 | 0.159385013 | 0.729790881 |
| ENSMUSG0000003726 | chr4 | Rsrp1 | -0.116090722 | 0.729938122 |
| ENSMUSG0000002077 | chr11 | Aspa | -0.226596876 | 0.730070721 |
| ENSMUSG0000002298 | chr16 | Sod1 | 0.173455173 | 0.730555862 |
| ENSMUSG0000003951 | chr3 | Cyp7b1 | 0.228039912 | 0.730772852 |
| ENSMUSG0000004166 | chr2 | Bbox1 | -0.286778664 | 0.730896138 |
| ENSMUSG0000002062 | chr11 | Map2k6 | -0.189496013 | 0.731053598 |
| ENSMUSG0000002608 | chr1 | Chst10 | -0.185464805 | 0.731053598 |
| ENSMUSG0000004661 | chr2 | Olfml2a | -0.152120622 | 0.731053598 |
| ENSMUSG0000003808 | chr16 | Opa1 | -0.098204695 | 0.731053598 |

| | | | | |
|---|---|---|---|---|
| ENSMUSG0000002064 | chr12 | Itsn2 | 0.107026191 | 0.731053598 |
| ENSMUSG0000002051 | chr11 | Rps6kb1 | 0.112323301 | 0.731053598 |
| ENSMUSG0000002948 | chr5 | Mrpl1 | 0.144218486 | 0.731053598 |
| ENSMUSG0000002220 | chr15 | Sub1 | 0.154660268 | 0.731053598 |
| ENSMUSG0000001957 | chr17 | Mydgf | 0.151533122 | 0.731053598 |
| ENSMUSG0000007436 | chr7 | C5ar2 | -0.524399605 | 0.73108854 |
| ENSMUSG0000007340 | chr17 | H2-Q6 | -0.622894371 | 0.73108854 |
| ENSMUSG0000003799 | chr16 | Rmi2 | -1.25717223 | 0.731127537 |
| ENSMUSG0000003391 | chr3 | Gucy1a1 | -0.10942811 | 0.731127537 |
| ENSMUSG0000002943 | chr5 | Nipsnap2 | 0.096105018 | 0.731127537 |
| ENSMUSG0000004620 | chr17 | Scaf8 | 0.112207166 | 0.731127537 |
| ENSMUSG0000004189 | chr5 | Git2 | 0.134390789 | 0.731127537 |
| ENSMUSG0000002676 | chr2 | Itga8 | 0.143644722 | 0.731127537 |
| ENSMUSG0000002294 | chr16 | Cbr3 | -0.254933486 | 0.731246889 |
| ENSMUSG0000002720 | chr2 | Sema6d | -0.151739451 | 0.731320376 |
| ENSMUSG0000007481 | chr19 | Hps6 | 0.136715256 | 0.731320376 |
| ENSMUSG0000004135 | chr17 | Rgl2 | 0.156055879 | 0.731320376 |
| ENSMUSG0000004130 | chr8 | Sntb2 | 0.135339276 | 0.731363074 |
| ENSMUSG0000003896 | chr16 | Ube2l3 | 0.086868549 | 0.731560868 |
| ENSMUSG0000005599 | chr5 | Zkscan5 | 0.135057657 | 0.731560868 |
| ENSMUSG0000002775 | chr3 | Exosc8 | -0.153188115 | 0.731816827 |
| ENSMUSG0000003096 | chr7 | Zranb1 | 0.096204139 | 0.731816827 |
| ENSMUSG0000002680 | chr2 | Ak8 | -0.634919693 | 0.73248816 |
| ENSMUSG0000003072 | chr7 | Rabep2 | -0.179903901 | 0.73248816 |
| ENSMUSG0000002563 | chrX | Hprt1 | 0.151830683 | 0.73248816 |
| ENSMUSG0000006350 | chr14 | Arhgap22 | -0.20208144 | 0.732586656 |
| ENSMUSG0000007857 | chr3 | Ube2d3 | 0.116328176 | 0.732686131 |
| ENSMUSG0000006714 | chr17 | Polr1c | 0.155824446 | 0.732686131 |
| ENSMUSG0000003510 | chr2 | Shc4 | -0.17654015 | 0.732696275 |
| ENSMUSG0000004077 | chr17 | Oard1 | 0.126373367 | 0.732798472 |
| ENSMUSG0000000633 | chr7 | Tfpt | 0.191012966 | 0.732798472 |
| ENSMUSG0000004513 | chr13 | Tubb2b | -0.144337644 | 0.733465435 |
| ENSMUSG0000002026 | chr11 | Slc36a1 | -0.154209732 | 0.733719484 |
| ENSMUSG0000005709 | chr7 | Fxyd3 | -0.275576297 | 0.733719484 |
| ENSMUSG0000003049 | chr7 | Faap24 | 0.210220302 | 0.733719484 |
| ENSMUSG0000007368 | chr4 | Faap20 | 0.128894972 | 0.733855395 |
| ENSMUSG0000004867 | chr7 | Tpcn2 | 0.165971939 | 0.733855395 |
| ENSMUSG0000003700 | chr5 | Zfp113 | -0.150452104 | 0.733859351 |
| ENSMUSG0000002960 | chr5 | Ankrd61 | -0.377469966 | 0.734126902 |
| ENSMUSG0000000137 | chr6 | Vps50 | 0.117480562 | 0.734126902 |
| ENSMUSG0000003814 | chr17 | Notch3 | 0.366937393 | 0.734126902 |
| ENSMUSG0000002472 | chr19 | Ostf1 | 0.163492736 | 0.73421422 |
| ENSMUSG0000000226 | chr7 | Peg3 | -0.211015215 | 0.734400385 |
| ENSMUSG0000006688 | chr8 | Zfp617 | 0.129828364 | 0.734400385 |
| ENSMUSG0000003727 | chr3 | Bltp1 | -0.165436818 | 0.73446759 |
| ENSMUSG0000005295 | chr13 | Gas1 | 0.173354308 | 0.73446759 |
| ENSMUSG0000003905 | chr3 | Ak5 | 0.344067187 | 0.73446759 |
| ENSMUSG0000002966 | chr5 | Tex26 | 0.451786848 | 0.734547667 |
| ENSMUSG0000006366 | chrX | Brwd3 | -0.199644698 | 0.734603617 |
| ENSMUSG0000000186 | chr2 | Aif1l | -0.21977933 | 0.734603617 |

| | | | | |
|---|---|---|---|---|
| ENSMUSG0000003121 | chrX | Ophn1 | -0.121075181 | 0.734603617 |
| ENSMUSG0000002236 | chr15 | Derl1 | 0.139782318 | 0.734603617 |
| ENSMUSG0000009008 | chrX | Srpx | 0.215193771 | 0.734603617 |
| ENSMUSG0000004219 | chr4 | Chchd7 | 0.169372449 | 0.734658774 |
| ENSMUSG0000000040 | chrX | Egfl6 | -0.396695095 | 0.734846193 |
| ENSMUSG0000005051 | chr4 | Oprd1 | -0.248502308 | 0.734846193 |
| ENSMUSG0000002645 | chr1 | Ppfia4 | -0.227396766 | 0.734846193 |
| ENSMUSG0000003026 | chr6 | Irag2 | -0.394525832 | 0.734846193 |
| ENSMUSG0000002573 | chr17 | Mcrip2 | -0.216508107 | 0.734846193 |
| ENSMUSG0000003070 | chr7 | Plekhb1 | -0.167322789 | 0.734846193 |
| ENSMUSG0000006721 | chrX | Usp51 | -0.49948306 | 0.734846193 |
| ENSMUSG0000007927 | chr2 | Hoxd3 | -0.256582552 | 0.734846193 |
| ENSMUSG0000004908 | chr2 | Bmyc | -0.17267791 | 0.734846193 |
| ENSMUSG0000002780 | chr3 | Etfdh | 0.098263022 | 0.734846193 |
| ENSMUSG0000002148 | chr13 | Lman2 | 0.134018196 | 0.734846193 |
| ENSMUSG0000003201 | chr9 | Nectin1 | 0.133863159 | 0.734846193 |
| ENSMUSG0000000027 | chr11 | Scpep1 | 0.13681628 | 0.734846193 |
| ENSMUSG0000002417 | chr17 | Nme4 | 0.263953518 | 0.734846193 |
| ENSMUSG0000002670 | chr2 | Nsun6 | 0.177965147 | 0.734846193 |
| ENSMUSG0000003078 | chr7 | Lyve1 | 0.156544745 | 0.734846193 |
| ENSMUSG0000004782 | chr11 | Trim16 | -0.175200217 | 0.734858594 |
| ENSMUSG0000005420 | chr19 | Ffar4 | -0.495303242 | 0.734943637 |
| ENSMUSG0000003110 | chrX | Slc25a14 | -0.139059888 | 0.734943637 |
| ENSMUSG0000002388 | chr17 | Phf10 | 0.106685376 | 0.734943637 |
| ENSMUSG0000004280 | chr1 | Spata46 | 0.456331774 | 0.734943637 |
| ENSMUSG0000002080 | chr11 | Ube2o | -0.124215261 | 0.735050923 |
| ENSMUSG0000002822 | chr4 | Nbn | 0.133185058 | 0.735050923 |
| ENSMUSG0000002667 | chr2 | Optn | 0.132810906 | 0.735277773 |
| ENSMUSG0000003232 | chr9 | Tmem30a | -0.082751475 | 0.735370203 |
| ENSMUSG0000002671 | chr2 | Stam | 0.086796935 | 0.735370203 |
| ENSMUSG0000002957 | chr6 | Tmem106b | 0.099998245 | 0.735370203 |
| ENSMUSG0000003365 | chr8 | Ddx19b | 0.142104598 | 0.735370203 |
| ENSMUSG0000000941 | chr1 | Nav1 | 0.176310263 | 0.735370203 |
| ENSMUSG0000003761 | chr7 | Tnfrsf23 | 0.391854305 | 0.735370203 |
| ENSMUSG0000002489 | chr19 | Pcx | -0.170658152 | 0.735459606 |
| ENSMUSG0000002481 | chr19 | Uhrf2 | 0.112821964 | 0.736597845 |
| ENSMUSG0000004043 | chr2 | Large2 | -0.506585081 | 0.736793767 |
| ENSMUSG0000009504 | JH584304.1 | ENSMUSG000000 | -0.220806116 | 0.736793767 |
| ENSMUSG0000000624 | chr9 | Ccdc159 | -0.191488473 | 0.736793767 |
| ENSMUSG0000004065 | chr14 | Tasor | -0.117311944 | 0.736793767 |
| ENSMUSG0000003732 | chr3 | Bbs7 | 0.099839011 | 0.736793767 |
| ENSMUSG0000002707 | chr2 | Slc43a3 | 0.160724782 | 0.736793767 |
| ENSMUSG0000003058 | chr7 | Dpf1 | 0.274809608 | 0.736793767 |
| ENSMUSG0000002770 | chr3 | Lrrc34 | -0.830651586 | 0.737096226 |
| ENSMUSG0000004027 | chr2 | Accs | -0.212630621 | 0.737096226 |
| ENSMUSG0000003527 | chr16 | Impg2 | -0.554810099 | 0.737096226 |
| ENSMUSG0000005294 | chr19 | Glis3 | -0.215546254 | 0.737096226 |
| ENSMUSG0000000505 | chr5 | Sh2b2 | -0.336353149 | 0.737096226 |
| ENSMUSG0000000638 | chr4 | Tek | -0.160917075 | 0.737096226 |
| ENSMUSG0000001889 | chr11 | Irf1 | -0.160241363 | 0.737096226 |

| | | | | |
|---|---|---|---|---|
| ENSMUSG00000523€ | chrX | B630019K06Rik | -0.208372296 | 0.737096226 |
| ENSMUSG0000002388 | chr17 | Zfp54 | -0.210453727 | 0.737096226 |
| ENSMUSG0000004249 | chr7 | Tbc1d10b | -0.114225385 | 0.737096226 |
| ENSMUSG0000002228 | chr15 | Ywhaz | 0.087635753 | 0.737096226 |
| ENSMUSG0000004107 | chr1 | Fzd7 | 0.081399828 | 0.737096226 |
| ENSMUSG0000002437 | chr18 | Cdc23 | 0.101694027 | 0.737096226 |
| ENSMUSG0000002125 | chr12 | Gpatch2l | 0.1013352 | 0.737096226 |
| ENSMUSG0000002263 | chr15 | Yaf2 | 0.108004012 | 0.737096226 |
| ENSMUSG0000002011 | chr10 | Tbk1 | 0.105957599 | 0.737096226 |
| ENSMUSG0000000705 | chr17 | Lsm2 | 0.145598435 | 0.737096226 |
| ENSMUSG0000003946 | chr9 | Tcta | 0.127462095 | 0.737096226 |
| ENSMUSG0000003850 | chr7 | Mesd | 0.142294734 | 0.737096226 |
| ENSMUSG0000003707 | chr8 | Rbmxl1 | 0.158717726 | 0.737096226 |
| ENSMUSG0000002669 | chr1 | Pigc | 0.15672721 | 0.737096226 |
| ENSMUSG0000003599 | chr11 | Fnip1 | 0.165050666 | 0.737096226 |
| ENSMUSG0000005576 | chr15 | Eef1d | 0.174655425 | 0.737096226 |
| ENSMUSG0000003756 | chr1 | Vash2 | 0.482125467 | 0.737096226 |
| ENSMUSG0000003827 | chr6 | Nop2 | 0.185971625 | 0.737096226 |
| ENSMUSG0000003135 | chrX | Arhgap6 | 0.161385261 | 0.737096226 |
| ENSMUSG0000000258 | chr11 | Mien1 | 0.250488754 | 0.737096226 |
| ENSMUSG0000000093 | chr15 | Top1mt | 0.177961703 | 0.737096226 |
| ENSMUSG0000002249 | chr16 | Shisa9 | 0.560185814 | 0.737096226 |
| ENSMUSG0000002738 | chr2 | Ndufaf5 | 0.230143534 | 0.737096226 |
| ENSMUSG0000003737 | chr5 | Spon2 | 0.734149755 | 0.737096226 |
| ENSMUSG0000005656 | chr1 | Mpz | -0.555695394 | 0.737261944 |
| ENSMUSG0000008994 | chr1 | Ugt1a5 | -0.87295036 | 0.737261944 |
| ENSMUSG0000003208 | chr9 | Il10ra | -0.365025021 | 0.737261944 |
| ENSMUSG0000000537 | chr17 | Fbxo11 | -0.10321266 | 0.737261944 |
| ENSMUSG0000002169 | chr13 | Kif2a | -0.094117735 | 0.737261944 |
| ENSMUSG0000003988 | chr5 | Tmem120a | 0.14860738 | 0.737261944 |
| ENSMUSG0000003721 | chr1 | Lipt1 | 0.204989632 | 0.737261944 |
| ENSMUSG0000003686 | chr3 | Syde2 | 0.136692865 | 0.737261944 |
| ENSMUSG0000004053 | chr5 | Adam22 | -0.180736601 | 0.737291899 |
| ENSMUSG0000002682 | chr2 | Ralgds | -0.153609354 | 0.737385972 |
| ENSMUSG0000002229 | chr15 | Fzd6 | 0.153146565 | 0.73773469 |
| ENSMUSG0000005123 | chr9 | Rnf7 | 0.17173331 | 0.73773469 |
| ENSMUSG0000003952 | chr4 | Cep104 | 0.095422289 | 0.737774493 |
| ENSMUSG0000003434 | chr11 | Wbp2 | 0.11883671 | 0.737774493 |
| ENSMUSG0000000294 | chr5 | Cd36 | 0.182232595 | 0.737774493 |
| ENSMUSG0000004193 | chr5 | Mvk | 0.201086592 | 0.737774493 |
| ENSMUSG0000002246 | chr15 | Rapgef3 | -0.257696972 | 0.738121089 |
| ENSMUSG0000003031 | chr6 | Dennd5b | 0.159791376 | 0.738121089 |
| ENSMUSG0000001718 | chr11 | Coa3 | 0.212770551 | 0.738331347 |
| ENSMUSG0000000566 | chr6 | Mthfd2 | 0.261334018 | 0.738399323 |
| ENSMUSG0000006636 | chr12 | Serpina3c | -0.533743249 | 0.738856782 |
| ENSMUSG0000003003 | chr6 | Mrpl53 | 0.24639001 | 0.738856782 |
| ENSMUSG0000002057 | chr12 | Sypl1 | 0.090905405 | 0.738959874 |
| ENSMUSG0000003559 | chr10 | Fam174c | 0.316491551 | 0.739106138 |
| ENSMUSG0000003984 | chr8 | Mcph1 | -0.189301332 | 0.739748111 |
| ENSMUSG0000003122 | chrX | Pbdc1 | 0.160620591 | 0.739748111 |

| | | | | |
|---|---|---|---|---|
| ENSMUSG0000005198 | chr16 | Casr | 0.302769686 | 0.739748111 |
| ENSMUSG0000006204 | chr7 | Lmtk3 | -0.256011192 | 0.740257866 |
| ENSMUSG0000002025 | chr9 | Wdr82 | 0.081973142 | 0.740360879 |
| ENSMUSG0000002459 | chr18 | Lmnb1 | -0.191567996 | 0.740873799 |
| ENSMUSG0000007118 | chr13 | Smim15 | 0.127677447 | 0.741370807 |
| ENSMUSG0000006237 | chr19 | Borcs7 | 0.164706504 | 0.741370807 |
| ENSMUSG0000002119 | chr13 | Pfkp | -0.158136399 | 0.741483227 |
| ENSMUSG0000002686 | chr2 | Hspa5 | 0.294438549 | 0.741483227 |
| ENSMUSG0000005825 | chr13 | Idi1 | 0.311843539 | 0.741487053 |
| ENSMUSG0000002630 | chr1 | Rab17 | -0.418642578 | 0.741590325 |
| ENSMUSG0000002782 | chr3 | Slc33a1 | -0.146396515 | 0.74176185 |
| ENSMUSG0000003076 | chr7 | Acer3 | -0.17733762 | 0.741804927 |
| ENSMUSG0000002573 | chr17 | Mettl26 | -0.162487843 | 0.742109288 |
| ENSMUSG0000004253 | chr15 | Gtpbp1 | 0.087880105 | 0.74220579 |
| ENSMUSG0000003821 | chr6 | Hoxa11 | 0.144099011 | 0.74220579 |
| ENSMUSG0000006598 | chr8 | Cd209b | -0.516384156 | 0.742238787 |
| ENSMUSG0000009023 | chr17 | Cfb | -0.660174726 | 0.742241626 |
| ENSMUSG0000003477 | chr10 | Tle2 | -0.207298375 | 0.742241626 |
| ENSMUSG0000002014 | chr11 | Slc1a4 | -0.199059067 | 0.742241626 |
| ENSMUSG0000003764 | chr17 | H2-DMa | -0.244989271 | 0.742241626 |
| ENSMUSG0000004247 | chr5 | Abcb4 | -0.505921227 | 0.742241626 |
| ENSMUSG0000003259 | chr9 | Apeh | 0.105704084 | 0.742241626 |
| ENSMUSG0000002196 | chr14 | Sap18 | 0.134360623 | 0.742241626 |
| ENSMUSG0000000833 | chr2 | Snrpb2 | 0.132028515 | 0.742241626 |
| ENSMUSG0000001188 | chr5 | Gltp | 0.133017784 | 0.742241626 |
| ENSMUSG0000004043 | chr9 | Zbtb38 | 0.139145298 | 0.742241626 |
| ENSMUSG0000001567 | chr13 | Mrpl32 | 0.177075593 | 0.742241626 |
| ENSMUSG0000002027 | chr10 | Pfkl | 0.174467935 | 0.742241626 |
| ENSMUSG0000002727 | chr2 | Mkks | 0.216248694 | 0.742241626 |
| ENSMUSG0000003574 | chr10 | Grin3b | 0.379735511 | 0.742241626 |
| ENSMUSG0000000703 | chr17 | Ddah2 | 0.142183056 | 0.742437104 |
| ENSMUSG0000003336 | chr4 | Ipo13 | 0.088251309 | 0.742804035 |
| ENSMUSG0000009540 | chr17 | Tmem200c | 0.281770362 | 0.742817651 |
| ENSMUSG0000002652 | chr1 | Opn3 | -0.356239642 | 0.742870478 |
| ENSMUSG0000002750 | chr2 | Rae1 | 0.111472073 | 0.7429664 |
| ENSMUSG0000004616 | chr14 | Kcnrg | -0.545496957 | 0.743001857 |
| ENSMUSG0000000260 | chr7 | Tgfb1 | 0.15064316 | 0.743103268 |
| ENSMUSG0000005486 | chr15 | Tafa5 | -0.125458775 | 0.743163553 |
| ENSMUSG0000000386 | chr7 | Ruvbl2 | 0.145480316 | 0.743163553 |
| ENSMUSG0000003276 | chr6 | Gng11 | 0.191730937 | 0.743163553 |
| ENSMUSG0000004362 | chr4 | Ubxn10 | -0.26609471 | 0.743214962 |
| ENSMUSG0000003385 | chr12 | Kcnk10 | -0.256171691 | 0.74325136 |
| ENSMUSG0000003379 | chr7 | Tubgcp5 | 0.094476976 | 0.74325136 |
| ENSMUSG0000002226 | chr15 | Marchf11 | 0.649918597 | 0.743274251 |
| ENSMUSG0000004216 | chr3 | Adrm1b | 0.366329389 | 0.743542446 |
| ENSMUSG0000002081 | chr11 | Wscd1 | -0.259989788 | 0.74400982 |
| ENSMUSG0000004909 | chr7 | Sephs2 | -0.174593427 | 0.74400982 |
| ENSMUSG0000000373 | chr4 | Kpna6 | 0.099908707 | 0.74400982 |
| ENSMUSG0000004136 | chr19 | Pum3 | 0.111197955 | 0.74400982 |
| ENSMUSG0000002709 | chr2 | Mtx2 | 0.118237109 | 0.74400982 |

| | | | | |
|---|---|---|---|---|
| ENSMUSG0000002734 | chr2 | Pcna | 0.137229785 | 0.74400982 |
| ENSMUSG0000006799 | chr14 | Gtf2f2 | 0.154835367 | 0.74400982 |
| ENSMUSG0000004788 | chr5 | Rell1 | 0.154850352 | 0.74400982 |
| ENSMUSG0000002940 | chr5 | Rilpl2 | 0.173975453 | 0.74400982 |
| ENSMUSG0000005319 | chr9 | Arhgap20 | 0.170806994 | 0.74400982 |
| ENSMUSG0000007845 | chr10 | Abracl | 0.182572049 | 0.74400982 |
| ENSMUSG0000007878 | chr7 | Actmap | -0.15133043 | 0.744061952 |
| ENSMUSG0000004147 | chr6 | Dcp1b | 0.15098117 | 0.744061952 |
| ENSMUSG0000002225 | chr15 | Laptm4b | 0.18533282 | 0.744061952 |
| ENSMUSG0000005118 | chr7 | Zfp524 | 0.313198494 | 0.744061952 |
| ENSMUSG0000003166 | chr8 | Sall1 | 0.3538432 | 0.744061952 |
| ENSMUSG0000000570 | chr8 | Agrp | -0.962600156 | 0.744069747 |
| ENSMUSG0000006403 | chr10 | Pym1 | 0.120313838 | 0.744069747 |
| ENSMUSG0000002555 | chr14 | Dock9 | 0.187022392 | 0.744328641 |
| ENSMUSG0000002656 | chr1 | Pou2f1 | -0.175795413 | 0.744380476 |
| ENSMUSG0000003491 | chr12 | Mdga2 | -0.171945198 | 0.744380476 |
| ENSMUSG0000004062 | chrX | Gemin8 | -0.169715946 | 0.744380476 |
| ENSMUSG0000003849 | chr8 | Tmco3 | 0.143722697 | 0.745443518 |
| ENSMUSG0000001840 | chr11 | Mrm1 | -0.176135707 | 0.745541689 |
| ENSMUSG0000001490 | chr8 | Naf1 | -0.145842923 | 0.745541689 |
| ENSMUSG0000002765 | chr2 | Ralgapb | -0.095796897 | 0.745541689 |
| ENSMUSG0000005615 | chr18 | Socs6 | 0.09674226 | 0.745541689 |
| ENSMUSG0000000128 | chr15 | Rarg | -0.083606874 | 0.745730279 |
| ENSMUSG0000000202 | chr9 | Kmt2a | -0.179069776 | 0.745957373 |
| ENSMUSG0000010990 | chr18 | Chmp1b | -0.130365333 | 0.745957373 |
| ENSMUSG0000005004 | chr2 | Tmx2 | -0.132874854 | 0.745957373 |
| ENSMUSG0000002181 | chr14 | Dnajc9 | -0.137628298 | 0.745957373 |
| ENSMUSG0000004425 | chr15 | Pced1b | -0.128328094 | 0.745957373 |
| ENSMUSG0000001768 | chr11 | Rhot1 | 0.105918329 | 0.745957373 |
| ENSMUSG0000004479 | chr13 | Isca1 | 0.11917562 | 0.745957373 |
| ENSMUSG0000004043 | chr7 | Ppp1r15a | 0.138694486 | 0.745957373 |
| ENSMUSG0000002950 | chr5 | Pus1 | 0.182527667 | 0.745957373 |
| ENSMUSG0000007526 | chr10 | Cenpw | 0.401832682 | 0.745957373 |
| ENSMUSG0000000232 | chr14 | Rec8 | 0.391265815 | 0.745957373 |
| ENSMUSG0000003595 | chr12 | Dock4 | 0.160937749 | 0.745959486 |
| ENSMUSG0000002833 | chr4 | Col15a1 | 0.159392945 | 0.746017889 |
| ENSMUSG0000002645 | chr1 | Syt2 | -0.17340105 | 0.746089156 |
| ENSMUSG0000003841 | chr9 | Higd1a | 0.244691213 | 0.746089156 |
| ENSMUSG0000008659 | chr9 | Susd5 | 0.237302015 | 0.74615103 |
| ENSMUSG0000009578 | chr5 | Nupr2 | -0.633968739 | 0.746183397 |
| ENSMUSG0000005203 | chr2 | Pfdn4 | 0.19822679 | 0.746183397 |
| ENSMUSG0000001772 | chr2 | Wfdc2 | -0.628515088 | 0.746227493 |
| ENSMUSG0000003025 | chr6 | Rad18 | -0.324503101 | 0.746227493 |
| ENSMUSG0000002814 | chr3 | Oaz3 | -0.734164427 | 0.746227493 |
| ENSMUSG0000002969 | chr5 | Ssc4d | -0.490916684 | 0.746227493 |
| ENSMUSG0000005191 | chr7 | Sox6 | -0.207895966 | 0.746227493 |
| ENSMUSG0000002641 | chr1 | Tnni1 | -0.433787632 | 0.746227493 |
| ENSMUSG0000006841 | chr14 | Pnp2 | -0.538016234 | 0.746227493 |
| ENSMUSG0000004848 | chr6 | Depp1 | -0.232990972 | 0.746227493 |
| ENSMUSG0000003224 | chr9 | Fem1b | -0.125193438 | 0.746227493 |

| | | | | |
|---|---|---|---|---|
| ENSMUSG0000006603 | chr4 | Ubr4 | -0.130830298 | 0.746227493 |
| ENSMUSG0000003854 | chr17 | Cul7 | -0.099827135 | 0.746227493 |
| ENSMUSG0000003594 | chr2 | Fbxw2 | 0.092662818 | 0.746227493 |
| ENSMUSG0000003979 | chr3 | Zfand1 | 0.127801859 | 0.746227493 |
| ENSMUSG0000004656 | chr3 | Arsj | 0.324739311 | 0.746227493 |
| ENSMUSG0000001950 | chr11 | Ubb | 0.165481004 | 0.746247838 |
| ENSMUSG0000002334 | chr9 | Poc1a | -0.191802686 | 0.746415085 |
| ENSMUSG0000001638 | chr2 | Mpped2 | 0.19317291 | 0.74643035 |
| ENSMUSG0000002711 | chr2 | Kif18a | -0.300006671 | 0.746755098 |
| ENSMUSG0000003122 | chrX | Awat2 | -0.36948471 | 0.746755098 |
| ENSMUSG0000003187 | chr8 | Nae1 | 0.086411735 | 0.746789992 |
| ENSMUSG0000005657 | chr11 | Tug1 | -0.103888249 | 0.746962697 |
| ENSMUSG0000004149 | chr1 | Kif14 | 0.242838434 | 0.746962697 |
| ENSMUSG0000002124 | chr12 | Tmed10 | 0.093568038 | 0.747092703 |
| ENSMUSG0000005904 | chr4 | Frem1 | -0.487134151 | 0.747092759 |
| ENSMUSG0000007004 | chr8 | Fam149a | -0.175203956 | 0.747092759 |
| ENSMUSG0000004325 | chr4 | Pigv | -0.171835063 | 0.747092759 |
| ENSMUSG0000004187 | chr1 | Ipo9 | 0.075761053 | 0.747092759 |
| ENSMUSG0000002799 | chr3 | Tlr2 | 0.162541381 | 0.747092759 |
| ENSMUSG0000005936 | chr19 | Fxn | 0.203552056 | 0.747092759 |
| ENSMUSG0000008740 | chr8 | Cers1 | -0.184073629 | 0.747239886 |
| ENSMUSG0000002834 | chr4 | Invs | -0.130162263 | 0.74731497 |
| ENSMUSG0000002856 | chr4 | Tm2d1 | 0.108294493 | 0.74731497 |
| ENSMUSG0000003829 | chr8 | Snx25 | 0.123522746 | 0.74731497 |
| ENSMUSG0000005630 | chr4 | Zfp981 | 0.683104191 | 0.74731497 |
| ENSMUSG0000002620 | chr1 | Stk16 | 0.129598929 | 0.747623866 |
| ENSMUSG0000002951 | chr5 | Gpc2 | -0.282836728 | 0.747707634 |
| ENSMUSG0000003012 | chr6 | Plxnd1 | 0.167605283 | 0.747707634 |
| ENSMUSG0000005933 | chr18 | Slc14a1 | 0.506120508 | 0.747707634 |
| ENSMUSG0000002175 | chr13 | Il6st | 0.100001825 | 0.747867688 |
| ENSMUSG0000002619 | chr1 | Abcb6 | -0.448920401 | 0.748077222 |
| ENSMUSG0000000973 | chr13 | Pou6f2 | 0.442317999 | 0.748077222 |
| ENSMUSG0000003244 | chr9 | Slc25a36 | -0.08804876 | 0.748163029 |
| ENSMUSG0000006380 | chr7 | Gpatch1 | 0.094019945 | 0.748478944 |
| ENSMUSG0000004559 | chr9 | Glb1 | -0.143557902 | 0.748558721 |
| ENSMUSG0000000493 | chr10 | Pias4 | 0.114183015 | 0.748558721 |
| ENSMUSG0000002970 | chr5 | Lrwd1 | 0.122362204 | 0.748558721 |
| ENSMUSG0000001490 | chr12 | Dnajb9 | 0.141656105 | 0.748558721 |
| ENSMUSG0000003004 | chr6 | Pole4 | 0.180471955 | 0.748558721 |
| ENSMUSG0000002615 | chr1 | Sdhaf4 | 0.192175111 | 0.748560638 |
| ENSMUSG0000003637 | chr13 | Abt1 | 0.176905838 | 0.748560638 |
| ENSMUSG0000004613 | chr19 | Patl1 | 0.120782813 | 0.748620462 |
| ENSMUSG0000002114 | chr12 | Pacs2 | -0.113515635 | 0.748627728 |
| ENSMUSG0000002561 | chr16 | Usp16 | 0.104161821 | 0.748627728 |
| ENSMUSG0000006895 | chr7 | Zfp619 | 0.18461585 | 0.748627728 |
| ENSMUSG0000004709 | chr14 | Rnf31 | -0.139209088 | 0.748652803 |
| ENSMUSG0000003929 | chr11 | Cybc1 | 0.124354715 | 0.748711297 |
| ENSMUSG0000002972 | chr5 | Gm20605 | -0.538924593 | 0.748768067 |
| ENSMUSG0000003750 | chr1 | Fam168b | 0.076236725 | 0.748871042 |
| ENSMUSG0000002171 | chr13 | Trappc13 | 0.115897979 | 0.748871042 |

| | | | | |
|---|---|---|---|---|
| ENSMUSG0000003697 | chr8 | Anapc10 | 0.12090474 | 0.748871042 |
| ENSMUSG0000004850 | chr9 | Tlcd5 | 0.119646431 | 0.748923571 |
| ENSMUSG0000003117 | chrX | Slc38a5 | -0.568775303 | 0.748940668 |
| ENSMUSG0000002296 | chr16 | Donson | -0.128780456 | 0.748940668 |
| ENSMUSG0000003222 | chr9 | Tcf12 | 0.097992919 | 0.748940668 |
| ENSMUSG0000003945 | chr9 | Snx22 | 0.404808996 | 0.748940668 |
| ENSMUSG0000000313 | chr1 | Tbc1d8 | 0.104448793 | 0.749324015 |
| ENSMUSG0000000956 | chr2 | Fpgs | 0.129816483 | 0.749453598 |
| ENSMUSG0000002522 | chr19 | Gbf1 | -0.081512453 | 0.749530887 |
| ENSMUSG0000003773 | chr5 | Limch1 | 0.098198342 | 0.749530887 |
| ENSMUSG0000007556 | chr5 | Rsph10b | -0.287751221 | 0.749550586 |
| ENSMUSG0000004736 | chr1 | Dnah14 | -0.574912357 | 0.749550586 |
| ENSMUSG0000000550 | chr1 | Usp40 | -0.138479829 | 0.749553503 |
| ENSMUSG0000002845 | chr4 | Pigo | -0.15415158 | 0.749683372 |
| ENSMUSG0000005767 | chr8 | Pkn1 | 0.1092099 | 0.749752458 |
| ENSMUSG0000002248 | chr15 | Hoxc10 | 0.323417903 | 0.749752458 |
| ENSMUSG0000004156 | chr17 | L3mbtl4 | 0.301933246 | 0.749856793 |
| ENSMUSG0000000282 | chr9 | Atg4d | -0.121809568 | 0.750504059 |
| ENSMUSG0000002820 | chr4 | Asph | 0.095519733 | 0.750504059 |
| ENSMUSG0000006892 | chr3 | Dap3 | 0.128856197 | 0.750504059 |
| ENSMUSG0000002912 | chr5 | Stx18 | 0.136665504 | 0.750504059 |
| ENSMUSG0000006657 | chr7 | Garre1 | 0.133794199 | 0.750504059 |
| ENSMUSG0000005066 | chr14 | Vstm4 | 0.191182453 | 0.750565962 |
| ENSMUSG0000004002 | chr10 | Lats1 | -0.118362656 | 0.750635934 |
| ENSMUSG0000004776 | chr7 | Atg16l2 | -0.118601739 | 0.750635934 |
| ENSMUSG0000000514 | chr8 | Man2b1 | 0.1583317 | 0.750828795 |
| ENSMUSG0000003535 | chr11 | Ccl12 | -0.454268322 | 0.750896964 |
| ENSMUSG0000001748 | chr14 | Top2b | 0.08063069 | 0.750896964 |
| ENSMUSG0000002902 | chr4 | Wrap73 | 0.137110616 | 0.750896964 |
| ENSMUSG0000005391 | chr9 | Kdm4d | -0.577504244 | 0.750943266 |
| ENSMUSG0000002234 | chr15 | Kcnv1 | -0.298587001 | 0.750956622 |
| ENSMUSG0000002081 | chr11 | Cygb | -0.144758033 | 0.750956622 |
| ENSMUSG0000005713 | chr2 | Chd6 | -0.143625964 | 0.750956622 |
| ENSMUSG0000005047 | chr9 | Fam118b | 0.10722374 | 0.750956622 |
| ENSMUSG0000002620 | chr1 | Dnpep | 0.122143415 | 0.750956622 |
| ENSMUSG0000005717 | chr7 | Gsk3a | 0.084543261 | 0.751085681 |
| ENSMUSG0000002038 | chr11 | Mrnip | -0.31502643 | 0.751098527 |
| ENSMUSG0000006707 | chr1 | Hes6 | -0.153216495 | 0.751098527 |
| ENSMUSG0000005400 | chr12 | Tdrd9 | -0.631217335 | 0.751105518 |
| ENSMUSG0000004068 | chr17 | Tbl3 | 0.119924025 | 0.751105518 |
| ENSMUSG0000000884 | chr6 | Cd163 | 0.318419373 | 0.751105518 |
| ENSMUSG0000007548 | chr14 | Commd6 | 0.160774178 | 0.751127325 |
| ENSMUSG0000003917 | chr5 | Tbc1d19 | 0.11593643 | 0.751212874 |
| ENSMUSG0000004191 | chr3 | Tdrkh | -0.109968825 | 0.751423942 |
| ENSMUSG0000005050 | chr14 | Pcdh20 | -0.215293147 | 0.751517918 |
| ENSMUSG0000002408 | chr17 | Man2a1 | -0.111292002 | 0.751662051 |
| ENSMUSG0000002897 | chr4 | Pex14 | 0.128405455 | 0.751662051 |
| ENSMUSG0000002248 | chr15 | Nckap1l | -0.187237736 | 0.7517971 |
| ENSMUSG0000005530 | chr5 | Mrfap1 | 0.124045657 | 0.751829453 |
| ENSMUSG0000002304 | chr15 | Prr13 | -0.173887202 | 0.751841929 |

| | | | | |
|---|---|---|---|---|
| ENSMUSG0000006336 | chr8 | Alg11 | -0.091563486 | 0.751841929 |
| ENSMUSG0000003175 | chr8 | Cog4 | 0.124682982 | 0.751841929 |
| ENSMUSG0000002467 | chr19 | Ms4a6d | -0.223206849 | 0.751925246 |
| ENSMUSG0000006198 | chr11 | Flot2 | 0.080781987 | 0.751925246 |
| ENSMUSG0000003148 | chr8 | Brf2 | 0.127812129 | 0.751925246 |
| ENSMUSG0000002210 | chr14 | Rcbtb2 | 0.115801407 | 0.751925246 |
| ENSMUSG0000003274 | chr4 | Heyl | 0.222703932 | 0.751925246 |
| ENSMUSG0000002946 | chr5 | Arpc3 | 0.137417842 | 0.752022488 |
| ENSMUSG0000003607 | chr4 | Sigmar1 | 0.108640003 | 0.752033008 |
| ENSMUSG0000003947 | chr8 | Zdhhc2 | -0.169304761 | 0.752107546 |
| ENSMUSG0000002244 | chr15 | C1qtnf6 | 0.175859254 | 0.752107546 |
| ENSMUSG0000002695 | chr2 | Neb | -0.329562934 | 0.752317537 |
| ENSMUSG0000007522 | chr10 | Ccdc162 | -0.384656512 | 0.752446883 |
| ENSMUSG0000002038 | chr11 | Cdkl3 | -0.16754645 | 0.752446883 |
| ENSMUSG0000002148 | chr13 | Zfp346 | 0.095919979 | 0.752446883 |
| ENSMUSG0000002914 | chr5 | Eif2b4 | 0.132020236 | 0.752446883 |
| ENSMUSG0000003239 | chr9 | Tipin | 0.122372288 | 0.752446883 |
| ENSMUSG0000005642 | chr11 | Slit3 | 0.160597926 | 0.752641699 |
| ENSMUSG0000002867 | chr4 | Lypla2 | -0.120254776 | 0.752722718 |
| ENSMUSG0000002727 | chr2 | Ubr1 | -0.106419634 | 0.752722718 |
| ENSMUSG0000003710 | chr8 | Dcaf15 | -0.144954882 | 0.752731733 |
| ENSMUSG0000003698 | chr9 | Pml | -0.109600494 | 0.752731733 |
| ENSMUSG0000006421 | chr15 | Ano6 | 0.078829078 | 0.752731733 |
| ENSMUSG0000002184 | chr14 | Ktn1 | 0.087233594 | 0.752746668 |
| ENSMUSG0000003977 | chr5 | Polr2j | 0.186756644 | 0.752746668 |
| ENSMUSG0000004088 | chr9 | Tmem205 | 0.198851442 | 0.752746668 |
| ENSMUSG0000005715 | chr14 | Homez | -0.138331244 | 0.752784748 |
| ENSMUSG0000003228 | chr9 | Acsbg1 | 0.288982059 | 0.753839242 |
| ENSMUSG0000002404 | chr17 | Sik1 | 0.186315826 | 0.753976879 |
| ENSMUSG0000005507 | chr7 | Gabra5 | -0.706269049 | 0.754335195 |
| ENSMUSG0000005495 | chr4 | Nt5c1a | -0.254615687 | 0.754335195 |
| ENSMUSG0000001115 | chr7 | Cfap161 | -0.473564821 | 0.754335195 |
| ENSMUSG0000005292 | chr11 | Arhgef15 | -0.257406194 | 0.754668005 |
| ENSMUSG0000001906 | chr9 | Rab3d | -0.101414643 | 0.754668005 |
| ENSMUSG0000005122 | chr1 | Bzw1 | 0.095697433 | 0.754668005 |
| ENSMUSG0000003876 | chr7 | Alpk3 | 0.515561227 | 0.754668005 |
| ENSMUSG0000011479 | chr14 | Gm49336 | 0.100773952 | 0.754668302 |
| ENSMUSG0000005343 | chr17 | Mapk14 | 0.107183781 | 0.755269261 |
| ENSMUSG0000002255 | chr15 | Dgat1 | -0.129694365 | 0.755398127 |
| ENSMUSG0000003184 | chr8 | Pde4c | -0.282102848 | 0.75560027 |
| ENSMUSG0000002815 | chr3 | Mttp | -0.283750379 | 0.755790843 |
| ENSMUSG0000007137 | chr12 | Hpcal1 | -0.144074996 | 0.755815846 |
| ENSMUSG0000003915 | chr5 | Stim2 | 0.12397628 | 0.755815846 |
| ENSMUSG0000004801 | chr7 | Zfp473 | -0.418122733 | 0.756091529 |
| ENSMUSG0000002221 | chr14 | Psme1 | 0.15328495 | 0.7563073 |
| ENSMUSG0000006888 | chr2 | Ssb | 0.103485702 | 0.756477515 |
| ENSMUSG0000000839 | chr10 | Elk3 | 0.185923634 | 0.756511187 |
| ENSMUSG0000001580 | chr5 | Med28 | 0.204009727 | 0.756633951 |
| ENSMUSG0000004521 | chr14 | Nudt18 | 0.173266982 | 0.756805097 |
| ENSMUSG0000003221 | chr9 | Ccnb2 | -0.564627046 | 0.756973381 |

| | | | | |
|---|---|---|---|---|
| ENSMUSG0000002301 | chr15 | Gpd1 | -0.374542198 | 0.756973381 |
| ENSMUSG000000455C | chr2 | Sys1 | 0.161662296 | 0.756973381 |
| ENSMUSG000000004E | chr7 | Septin1 | -0.223344739 | 0.757197145 |
| ENSMUSG000000562E | chr15 | Kcnq3 | 0.170973034 | 0.757600456 |
| ENSMUSG000000214C | chr13 | Ripk1 | 0.121153659 | 0.757756772 |
| ENSMUSG000000243 | chr18 | Psd2 | 0.224721317 | 0.757756772 |
| ENSMUSG0000004021 | chr9 | Ttc12 | -0.153139574 | 0.757794056 |
| ENSMUSG000000206E | chr12 | Bcap29 | 0.07543539 | 0.757794056 |
| ENSMUSG000000391C | chr5 | Rbpj | 0.094652602 | 0.757794056 |
| ENSMUSG000000281 | chr3 | Snx27 | -0.102350641 | 0.757965087 |
| ENSMUSG000000324E | chr9 | Nmnat3 | 0.206141508 | 0.758265732 |
| ENSMUSG000000185C | chr11 | Adora2b | 0.458095716 | 0.758265732 |
| ENSMUSG0000002477 | chr19 | Fas | 0.239297109 | 0.758335487 |
| ENSMUSG0000002952 | chr5 | Sirt4 | 0.170757663 | 0.758382156 |
| ENSMUSG000000461 | chr7 | Nkg7 | -0.706245277 | 0.758609734 |
| ENSMUSG000000271E | chr2 | Trim44 | -0.074659673 | 0.758609734 |
| ENSMUSG0000002661 | chr1 | Cr2 | -0.551399175 | 0.758625393 |
| ENSMUSG0000004337 | chr11 | Hexim2 | -0.175471825 | 0.758625393 |
| ENSMUSG000000416C | chr9 | Entpd3 | -0.211022003 | 0.758695473 |
| ENSMUSG000000386E | chr7 | Fsd2 | -0.465841984 | 0.758695473 |
| ENSMUSG000000047C | chr1 | Ly9 | -0.49064768 | 0.758695473 |
| ENSMUSG0000003201 | chr9 | Sc5d | -0.146724191 | 0.758695473 |
| ENSMUSG000000441 | chr5 | Col26a1 | -0.571023233 | 0.758695473 |
| ENSMUSG0000005647 | chr3 | Med12l | -0.168586127 | 0.758695473 |
| ENSMUSG000000229E | chr16 | Il10rb | -0.36561881 | 0.758695473 |
| ENSMUSG000000525E | chr6 | Lrrtm4 | -0.202361076 | 0.758695473 |
| ENSMUSG000000334 | chr8 | Terf2ip | -0.130436076 | 0.758695473 |
| ENSMUSG000000441 | chr5 | Pheta1 | -0.185195204 | 0.758695473 |
| ENSMUSG0000010117 | chr2 | Hoxd4 | -0.14182324 | 0.758695473 |
| ENSMUSG000000204 | chr11 | Tcn2 | -0.122438665 | 0.758695473 |
| ENSMUSG000000420E | chr1 | Tmcc2 | -0.111978829 | 0.758695473 |
| ENSMUSG000000326E | chr3 | Entrep3 | -0.110712111 | 0.758695473 |
| ENSMUSG000000271E | chr2 | Lin7c | -0.096769833 | 0.758695473 |
| ENSMUSG000000687 | chr3 | Sort1 | -0.095926953 | 0.758695473 |
| ENSMUSG000000215 | chr13 | Smad5 | 0.07907251 | 0.758695473 |
| ENSMUSG0000002717 | chr2 | Hipk3 | 0.097390762 | 0.758695473 |
| ENSMUSG000000207E | chr11 | Ncbp3 | 0.105804697 | 0.758695473 |
| ENSMUSG0000000657 | chr1 | Slc4a3 | 0.11020604 | 0.758695473 |
| ENSMUSG000000251C | chr7 | Btbd1 | 0.101402644 | 0.758695473 |
| ENSMUSG0000002092 | chr11 | Ccdc43 | 0.115862813 | 0.758695473 |
| ENSMUSG000000060E | chr18 | Sra1 | 0.127765356 | 0.758695473 |
| ENSMUSG0000003331 | chr18 | Fem1c | 0.139277718 | 0.758695473 |
| ENSMUSG000000313E | chrX | Msl3 | 0.135901883 | 0.758695473 |
| ENSMUSG000000198C | chr10 | Adat2 | 0.205173975 | 0.758695473 |
| ENSMUSG0000002897 | chr4 | Dffa | 0.116762702 | 0.758695473 |
| ENSMUSG000000287 | chr4 | Mrto4 | 0.154831139 | 0.758695473 |
| ENSMUSG0000007881 | chr7 | Leng1 | 0.151534094 | 0.758695473 |
| ENSMUSG000000579E | chr12 | Itpk1 | 0.161176964 | 0.758695473 |
| ENSMUSG000000757C | chr7 | Selenos | 0.161994458 | 0.758695473 |
| ENSMUSG000000598 | chr13 | Zfp874b | 0.172097582 | 0.758695473 |

| | | | | |
|---|---|---|---|---|
| ENSMUSG0000002839 | chr4 | Dmac1 | 0.204296313 | 0.758695473 |
| ENSMUSG0000000882 | chr12 | Acyp1 | 0.200661208 | 0.758695473 |
| ENSMUSG0000002079 | chr11 | Spns3 | 0.43404553 | 0.758695473 |
| ENSMUSG0000002201 | chr14 | Dnajc15 | 0.196685715 | 0.758695473 |
| ENSMUSG0000004510 | chrX | Dmd | 0.158727369 | 0.758695473 |
| ENSMUSG0000004245 | chr5 | Reln | 0.246877714 | 0.758695473 |
| ENSMUSG0000002837 | chr4 | Pappa | 0.195912483 | 0.758695473 |
| ENSMUSG0000003772 | chr8 | Ckap2 | 0.611955659 | 0.758695473 |
| ENSMUSG0000002460 | chr18 | Slc27a6 | 0.25913745 | 0.758695473 |
| ENSMUSG0000003138 | chrX | Asb11 | 0.625106186 | 0.758695473 |
| ENSMUSG0000004742 | chr6 | Fam180a | 0.382079225 | 0.758695473 |
| ENSMUSG0000000303 | chr8 | Fam32a | 0.099200439 | 0.758743519 |
| ENSMUSG0000003253 | chr9 | Amotl2 | 0.155612762 | 0.758743519 |
| ENSMUSG0000003843 | chr11 | Mllt6 | -0.135657839 | 0.758760578 |
| ENSMUSG0000002588 | chr18 | Myo5b | -0.48792613 | 0.758762975 |
| ENSMUSG0000002479 | chr19 | Vps51 | -0.126109219 | 0.758762975 |
| ENSMUSG0000003782 | chr5 | Smim14 | 0.080879682 | 0.758762975 |
| ENSMUSG0000003848 | chr7 | Polr2l | 0.234171701 | 0.758762975 |
| ENSMUSG0000003781 | chr2 | D630003M21Rik | -0.246330962 | 0.75881702 |
| ENSMUSG0000002767 | chr3 | Ccdc39 | -0.154213279 | 0.758986626 |
| ENSMUSG0000001540 | chrX | Ace2 | 1.300618633 | 0.758986626 |
| ENSMUSG0000007929 | chr6 | Clec7a | 0.326849873 | 0.758986626 |
| ENSMUSG0000001816 | chr15 | Mfng | -0.278839695 | 0.758993316 |
| ENSMUSG0000002591 | chr1 | Ppp1r42 | -0.585389558 | 0.758993316 |
| ENSMUSG0000002306 | chr14 | Sncg | -0.137956506 | 0.758993316 |
| ENSMUSG0000002477 | chr19 | Atg2a | -0.125741273 | 0.758993316 |
| ENSMUSG0000005626 | chr3 | Lrif1 | -0.10420394 | 0.758993316 |
| ENSMUSG0000000644 | chr4 | Srm | 0.122615461 | 0.758993316 |
| ENSMUSG0000003211 | chr9 | Hinfp | 0.129322488 | 0.758993316 |
| ENSMUSG0000006013 | chr2 | Atp8b4 | 0.46590951 | 0.758993316 |
| ENSMUSG0000002855 | chr4 | Cdkn2c | 0.216165601 | 0.758993316 |
| ENSMUSG0000005713 | chr18 | Txnl4a | 0.207201661 | 0.758993316 |
| ENSMUSG0000002198 | chr14 | Atp8a2 | -0.199103793 | 0.759166853 |
| ENSMUSG0000002519 | chr19 | Erlin1 | -0.185567211 | 0.759294121 |
| ENSMUSG0000002513 | chr11 | Arhgdia | 0.102798281 | 0.759294121 |
| ENSMUSG0000002597 | chr1 | Maip1 | 0.141188072 | 0.759294121 |
| ENSMUSG0000003074 | chr7 | Il4ra | 0.147818712 | 0.759308271 |
| ENSMUSG0000002240 | chr15 | Xpnpep3 | 0.083668648 | 0.759625043 |
| ENSMUSG0000002266 | chr16 | Gtpbp8 | -0.103768856 | 0.759677157 |
| ENSMUSG0000009189 | chr18 | Ube2d2a | 0.105382364 | 0.759908115 |
| ENSMUSG0000004820 | chr7 | Cracr2b | -0.19435865 | 0.759954372 |
| ENSMUSG0000007947 | chr1 | Utp14b | -0.234705739 | 0.760063076 |
| ENSMUSG0000002905 | chr4 | Plch2 | -0.159780002 | 0.760063076 |
| ENSMUSG0000002868 | chr4 | Eif2b3 | 0.098957626 | 0.760214322 |
| ENSMUSG0000009055 | chr1 | Snrpe | 0.210397185 | 0.76052202 |
| ENSMUSG0000005142 | chr11 | Ccdc157 | -0.13784688 | 0.76074626 |
| ENSMUSG0000002846 | chr4 | Npr2 | -0.12781805 | 0.760925125 |
| ENSMUSG0000002096 | chr12 | Tshr | 0.366893492 | 0.760925125 |
| ENSMUSG0000007910 | chr15 | C7 | -0.241126353 | 0.761166744 |
| ENSMUSG0000003863 | chr7 | Lrrc56 | -0.22312695 | 0.761551264 |

| | | | | |
|---|---|---|---|---|
| ENSMUSG0000002010 | chr10 | Vsir | -0.118141844 | 0.761551264 |
| ENSMUSG0000002073 | chr11 | Nt5c | 0.12958887 | 0.761551264 |
| ENSMUSG0000001447 | chr8 | Rnf166 | 0.133676296 | 0.761551264 |
| ENSMUSG0000000670 | chr17 | Pknox1 | 0.140018582 | 0.761551264 |
| ENSMUSG0000003070 | chr7 | Dnajb13 | 0.284123272 | 0.761551264 |
| ENSMUSG0000007961 | chr18 | Seh1l | 0.088402216 | 0.762324562 |
| ENSMUSG0000003911 | chr13 | Nrn1 | -0.238384958 | 0.762401302 |
| ENSMUSG0000002891 | chr4 | Plekhm2 | 0.108619661 | 0.762458774 |
| ENSMUSG0000005476 | chr8 | Mtnr1a | 0.722330996 | 0.762458774 |
| ENSMUSG0000007458 | chr2 | Arfgef2 | -0.125805232 | 0.762865652 |
| ENSMUSG0000003442 | chr8 | Gcsh | 0.169370201 | 0.762948664 |
| ENSMUSG0000002819 | chr3 | Ddah1 | -0.163480637 | 0.763247681 |
| ENSMUSG0000002732 | chr2 | Siglec1 | -0.472918714 | 0.763571568 |
| ENSMUSG0000002732 | chr2 | Rad51 | -0.463802544 | 0.763571568 |
| ENSMUSG0000002833 | chr4 | Nans | -0.184595961 | 0.763571568 |
| ENSMUSG0000002804 | chr3 | Efna4 | -0.248429448 | 0.763571568 |
| ENSMUSG0000003010 | chr6 | Arl8b | 0.075944372 | 0.763571568 |
| ENSMUSG0000002114 | chr12 | Pcnx1 | 0.120577312 | 0.763571568 |
| ENSMUSG0000002077 | chr11 | Unk | 0.12141981 | 0.763571568 |
| ENSMUSG0000002779 | chr3 | Sis | 1.472621899 | 0.763571568 |
| ENSMUSG0000002290 | chr16 | Kpna1 | 0.122093424 | 0.763614713 |
| ENSMUSG0000007915 | chr8 | Nalf1 | -0.169916042 | 0.763925406 |
| ENSMUSG0000003887 | chr6 | Bpgm | 0.141862336 | 0.763925406 |
| ENSMUSG0000005054 | chr3 | Fam241a | 0.150632474 | 0.763925406 |
| ENSMUSG0000005315 | chr7 | Fes | -0.186215431 | 0.764006053 |
| ENSMUSG0000002136 | chr13 | Nedd9 | -0.133282666 | 0.764006053 |
| ENSMUSG0000003790 | chr2 | Sirpa | -0.091905095 | 0.764006053 |
| ENSMUSG0000006826 | chr2 | Cenpb | 0.087331325 | 0.764006053 |
| ENSMUSG0000003136 | chrX | Ctps2 | 0.088524988 | 0.764006053 |
| ENSMUSG0000003723 | chr5 | Mxd4 | 0.097011036 | 0.764006053 |
| ENSMUSG0000001983 | chr10 | Gtf3c6 | 0.116174171 | 0.764006053 |
| ENSMUSG0000004432 | chr11 | Trp53i13 | 0.132007477 | 0.764006053 |
| ENSMUSG0000002907 | chr4 | Tas1r3 | 0.235634788 | 0.764006053 |
| ENSMUSG0000003576 | chr18 | Dym | -0.124980632 | 0.764013573 |
| ENSMUSG0000002423 | chr18 | Zeb1 | 0.121937936 | 0.764177757 |
| ENSMUSG0000002297 | chr16 | Mis18a | 0.222187865 | 0.764177757 |
| ENSMUSG0000002045 | chr11 | Patz1 | -0.132394562 | 0.764278251 |
| ENSMUSG0000003501 | chr10 | Zbtb7a | -0.128823857 | 0.764278251 |
| ENSMUSG0000002662 | chr1 | Ppp2r5a | 0.108399217 | 0.764278251 |
| ENSMUSG0000002867 | chr4 | Hmgcl | 0.154614212 | 0.764278251 |
| ENSMUSG0000002501 | chr19 | Lcor | -0.23617933 | 0.764413649 |
| ENSMUSG0000002230 | chr15 | Lrp12 | 0.087683569 | 0.764413649 |
| ENSMUSG0000005038 | chr7 | Kif7 | 0.141142754 | 0.764501262 |
| ENSMUSG0000007853 | chr4 | Nkain1 | -0.326254981 | 0.764848959 |
| ENSMUSG0000007053 | chr1 | Ccdc190 | -0.59828175 | 0.764848959 |
| ENSMUSG0000006395 | chr17 | Brpf3 | -0.159230778 | 0.764848959 |
| ENSMUSG0000003582 | chr15 | Pim3 | -0.183321138 | 0.764848959 |
| ENSMUSG0000002863 | chr4 | Edn2 | -0.630356635 | 0.764848959 |
| ENSMUSG0000005567 | chr8 | Kbtbd11 | -0.148418511 | 0.764848959 |
| ENSMUSG0000007003 | chr1 | Sp110 | -0.204261079 | 0.764848959 |

| | | | | |
|---|---|---|---|---|
| ENSMUSG0000002824 | chr4 | Ubxn2b | -0.120095602 | 0.764848959 |
| ENSMUSG0000005605 | chr1 | Mia3 | -0.089094345 | 0.764848959 |
| ENSMUSG0000001307 | chr9 | Amotl1 | 0.084043258 | 0.764848959 |
| ENSMUSG0000001316 | chr8 | Atp6v0d1 | 0.095864443 | 0.764848959 |
| ENSMUSG0000002560 | chr6 | Mkln1 | 0.114747038 | 0.764848959 |
| ENSMUSG0000007911 | chr5 | Kdelr2 | 0.121390759 | 0.764848959 |
| ENSMUSG0000004434 | chr1 | Phlpp1 | 0.119864562 | 0.764848959 |
| ENSMUSG0000004048 | chr11 | Sox30 | 0.499348943 | 0.764848959 |
| ENSMUSG0000002028 | chr11 | Nprl3 | 0.126208182 | 0.764848959 |
| ENSMUSG0000004431 | chr3 | Mab21l3 | 0.607024623 | 0.764848959 |
| ENSMUSG0000004740 | chr9 | Ctdspl | 0.142119287 | 0.764848959 |
| ENSMUSG0000004838 | chr12 | Osr1 | 0.163991777 | 0.764848959 |
| ENSMUSG0000002964 | chr5 | Rasl11a | 0.625166215 | 0.764848959 |
| ENSMUSG0000004227 | chr2 | Sestd1 | 0.119835494 | 0.764859149 |
| ENSMUSG0000003645 | chr18 | Arhgap26 | -0.160123855 | 0.765084195 |
| ENSMUSG0000004845 | chr5 | Msx1 | -0.462578098 | 0.765348237 |
| ENSMUSG0000003817 | chr8 | Enpp6 | -0.188354442 | 0.765686826 |
| ENSMUSG0000001847 | chr11 | Kcnab3 | -0.322122463 | 0.765686826 |
| ENSMUSG0000003861 | chr11 | Nfe2l1 | -0.085742329 | 0.765686826 |
| ENSMUSG0000002001 | chr10 | Ntn4 | 0.155866203 | 0.765686826 |
| ENSMUSG0000005238 | chr16 | Nrros | 0.24404681 | 0.765755321 |
| ENSMUSG0000002276 | chr16 | Thap7 | 0.145090609 | 0.765787125 |
| ENSMUSG0000005348 | chr1 | Usp21 | 0.105797097 | 0.765814026 |
| ENSMUSG0000002260 | chr15 | Arc | -0.42991377 | 0.766091958 |
| ENSMUSG0000002804 | chr3 | Scamp3 | 0.084468998 | 0.766091958 |
| ENSMUSG0000005037 | chr13 | Il31ra | -0.515238678 | 0.76613166 |
| ENSMUSG0000004134 | chr12 | Bdkrb1 | -0.415139428 | 0.76613166 |
| ENSMUSG0000003659 | chr2 | Arhgap21 | -0.140988568 | 0.76613166 |
| ENSMUSG0000002448 | chr18 | Slc4a9 | -0.583262658 | 0.76613166 |
| ENSMUSG0000002821 | chr4 | Ccne2 | -0.196720447 | 0.76613166 |
| ENSMUSG0000011767 | chr18 | Apbb3 | -0.127676777 | 0.76613166 |
| ENSMUSG0000003349 | chr13 | Larp4b | -0.111641895 | 0.76613166 |
| ENSMUSG0000002678 | chr2 | Apbb1ip | 0.136849278 | 0.76613166 |
| ENSMUSG0000006778 | chr2 | Nnat | 0.132183971 | 0.76613166 |
| ENSMUSG0000007950 | chrX | Apoo | 0.167021986 | 0.76613166 |
| ENSMUSG0000004196 | chr11 | Znrf3 | 0.147907377 | 0.76613166 |
| ENSMUSG0000000842 | chr9 | Herpud2 | 0.095194514 | 0.766160731 |
| ENSMUSG0000002015 | chr10 | Ptprb | 0.214787019 | 0.766284745 |
| ENSMUSG0000003937 | chr8 | Wdr17 | -0.261942335 | 0.766433704 |
| ENSMUSG0000002520 | chr19 | Scd2 | -0.185972313 | 0.766539878 |
| ENSMUSG0000006911 | chr3 | Zbtb10 | -0.146027156 | 0.766539878 |
| ENSMUSG0000002410 | chr17 | Washc1 | 0.088088272 | 0.766671649 |
| ENSMUSG0000002436 | chr18 | Gfra3 | -0.429687929 | 0.766801351 |
| ENSMUSG0000001612 | chr5 | Stard13 | 0.102954347 | 0.766801351 |
| ENSMUSG0000003363 | chr18 | AW554918 | -0.181661155 | 0.766817039 |
| ENSMUSG0000002725 | chr2 | Map1a | -0.137178127 | 0.766817039 |
| ENSMUSG0000002503 | chr19 | Arl3 | 0.148494624 | 0.766817039 |
| ENSMUSG0000002956 | chr6 | Tmem168 | 0.073458938 | 0.766860311 |
| ENSMUSG0000002070 | chr11 | Map3k3 | 0.080015468 | 0.766860311 |
| ENSMUSG0000002058 | chr12 | Rock2 | 0.097986117 | 0.766860311 |

| | | | | |
|---|---|---|---|---|
| ENSMUSG0000002574 | chr5 | Hadha | 0.111620298 | 0.766860311 |
| ENSMUSG0000002177 | chr14 | Ube2e1 | 0.110496664 | 0.766860311 |
| ENSMUSG0000001004 | chr9 | Tmem115 | 0.127654333 | 0.766860311 |
| ENSMUSG0000004702 | chr12 | Mipol1 | 0.15892288 | 0.766860311 |
| ENSMUSG0000002103 | chr12 | Snw1 | 0.123519443 | 0.766860311 |
| ENSMUSG0000006328 | chr18 | Zfp35 | 0.183590959 | 0.766860311 |
| ENSMUSG0000005713 | chr14 | Rpgrip1 | 0.478038159 | 0.766860311 |
| ENSMUSG0000007849 | chr4 | Zfp984 | 0.183384286 | 0.766860311 |
| ENSMUSG0000005036 | chr2 | Hoxd10 | 0.171163459 | 0.766860311 |
| ENSMUSG0000009189 | chr5 | Gm17019 | -0.515402394 | 0.766897425 |
| ENSMUSG0000003637 | chr5 | Cracd | -0.14903258 | 0.766897425 |
| ENSMUSG0000003192 | chr9 | Maml2 | 0.127680275 | 0.766897425 |
| ENSMUSG0000001855 | chr11 | Ctdnep1 | 0.075809535 | 0.766931088 |
| ENSMUSG0000004387 | chr4 | Zmym1 | 0.137470921 | 0.766931088 |
| ENSMUSG0000002279 | chr16 | Igsf11 | -0.13813333 | 0.766994606 |
| ENSMUSG0000000693 | chr11 | P3h4 | 0.158976523 | 0.766994606 |
| ENSMUSG0000005726 | chr12 | Bbof1 | -0.273764746 | 0.767363891 |
| ENSMUSG0000006307 | chr4 | Kif1b | -0.12791451 | 0.767363891 |
| ENSMUSG0000002275 | chr16 | Tomm70a | 0.074753952 | 0.767363891 |
| ENSMUSG0000002675 | chr2 | Arpc5l | 0.117138595 | 0.767363891 |
| ENSMUSG0000001755 | chr11 | Atad5 | -0.218940834 | 0.767671862 |
| ENSMUSG0000004424 | chr9 | Il20rb | 0.245990756 | 0.767972145 |
| ENSMUSG0000002737 | chr2 | Fahd2a | 0.186280423 | 0.767972145 |
| ENSMUSG0000003503 | chr9 | Nek11 | -0.267900978 | 0.768061441 |
| ENSMUSG0000004613 | chr19 | Brd10 | -0.130252412 | 0.768061441 |
| ENSMUSG0000002764 | chr2 | Src | -0.098524896 | 0.768061441 |
| ENSMUSG0000007418 | chr3 | Znhit6 | 0.117356066 | 0.768061441 |
| ENSMUSG0000003205 | chr9 | Ppp2r1b | 0.106085428 | 0.768061441 |
| ENSMUSG0000002801 | chr3 | Ctso | 0.118091864 | 0.768061441 |
| ENSMUSG0000005032 | chr4 | Ndufaf6 | 0.206114043 | 0.768061441 |
| ENSMUSG0000005227 | chr5 | Bhlha15 | -0.981347744 | 0.768219906 |
| ENSMUSG0000002527 | chrX | Alas2 | 0.458268788 | 0.768219906 |
| ENSMUSG0000004435 | chrX | P2ry4 | -1.132530189 | 0.768270406 |
| ENSMUSG0000002838 | chr4 | Snx30 | -0.169887201 | 0.768747245 |
| ENSMUSG0000003148 | chr8 | Erlin2 | -0.092935132 | 0.768747245 |
| ENSMUSG0000006333 | chr10 | Krr1 | 0.092483734 | 0.768747245 |
| ENSMUSG0000000277 | chr7 | Kdelr1 | 0.112152079 | 0.768747245 |
| ENSMUSG0000002977 | chr6 | Irf5 | 0.186403129 | 0.768963977 |
| ENSMUSG0000002797 | chr3 | Ndst3 | -0.306965952 | 0.76906806 |
| ENSMUSG0000003101 | chr7 | Swap70 | -0.147280943 | 0.76906806 |
| ENSMUSG0000001315 | chr8 | Gfod2 | -0.146612335 | 0.76906806 |
| ENSMUSG0000007259 | chr14 | Ear2 | -0.376356307 | 0.76906806 |
| ENSMUSG0000004503 | chr3 | Cetn4 | -0.137143102 | 0.76906806 |
| ENSMUSG0000002708 | chr2 | Tfpi | 0.109978176 | 0.76906806 |
| ENSMUSG0000002873 | chr4 | Aldh4a1 | 0.152031743 | 0.76906806 |
| ENSMUSG0000005932 | chr5 | Hopx | 0.192648349 | 0.769158369 |
| ENSMUSG0000004898 | chr5 | Elfn1 | -0.302650551 | 0.7691685 |
| ENSMUSG0000000628 | chr14 | Ttc5 | -0.092018437 | 0.7691685 |
| ENSMUSG0000004551 | chr9 | Zfp560 | 0.129716926 | 0.7691685 |
| ENSMUSG0000000885 | chr13 | Rala | 0.121049335 | 0.7691685 |

| | | | | |
|---|---|---|---|---|
| ENSMUSG0000003802 | chr5 | Atp6v0a2 | -0.101842801 | 0.769452085 |
| ENSMUSG0000004928 | chr3 | Lix1l | 0.114656726 | 0.769452085 |
| ENSMUSG0000003010 | chr6 | Itpr1 | 0.117608205 | 0.769452085 |
| ENSMUSG0000004872 | chr19 | Tbc1d12 | -0.090308488 | 0.769505221 |
| ENSMUSG0000002940 | chr5 | Pitpnm2 | 0.123531652 | 0.769563528 |
| ENSMUSG0000003908 | chr2 | Ss18l1 | -0.140717161 | 0.769666524 |
| ENSMUSG0000006237 | chr15 | Tmem65 | 0.12512968 | 0.769685707 |
| ENSMUSG0000003058 | chr7 | Sipa1l3 | -0.134981568 | 0.769770083 |
| ENSMUSG0000002811 | chr3 | Mettl14 | 0.089011331 | 0.769770083 |
| ENSMUSG0000003817 | chr13 | Mylip | -0.207514245 | 0.769966193 |
| ENSMUSG0000002083 | chr11 | Dhrs13 | -0.267459684 | 0.769966193 |
| ENSMUSG0000000590 | chr5 | Pex1 | -0.142427516 | 0.769966193 |
| ENSMUSG0000001821 | chr11 | Pmp22 | -0.10653217 | 0.769966193 |
| ENSMUSG0000002431 | chr17 | Vps52 | 0.090508653 | 0.769966193 |
| ENSMUSG0000005317 | chr5 | Mterf1b | 0.370424215 | 0.770021176 |
| ENSMUSG0000004129 | chr13 | Cdk13 | -0.115555659 | 0.770132788 |
| ENSMUSG0000002478 | chr19 | Cdc37l1 | 0.087975349 | 0.770132788 |
| ENSMUSG0000005958 | chr15 | Nsmce2 | 0.112203523 | 0.770144458 |
| ENSMUSG0000002484 | chr19 | Cabp4 | -0.412332386 | 0.77024142 |
| ENSMUSG0000003660 | chr15 | Plxnb2 | -0.068734577 | 0.77024142 |
| ENSMUSG0000000073 | chr8 | Spg7 | 0.108359746 | 0.77024142 |
| ENSMUSG0000005242 | chr1 | Tmco1 | 0.154151148 | 0.77024142 |
| ENSMUSG0000005102 | chr5 | Hs3st1 | 0.184923533 | 0.77024142 |
| ENSMUSG0000003640 | chr6 | Gng12 | 0.11027324 | 0.770412375 |
| ENSMUSG0000004918 | chr8 | Purg | 0.134384718 | 0.770663123 |
| ENSMUSG0000003605 | chr2 | Fmnl2 | 0.111432854 | 0.77083444 |
| ENSMUSG0000006319 | chr11 | Cd300lb | 0.333259025 | 0.77083444 |
| ENSMUSG0000004686 | chr4 | Hectd3 | -0.137687928 | 0.770966726 |
| ENSMUSG0000000059 | chr11 | Gm2a | 0.103622077 | 0.770966726 |
| ENSMUSG0000003218 | chr9 | Carm1 | 0.079599234 | 0.771048614 |
| ENSMUSG0000006614 | chr4 | Cdc26 | 0.161810801 | 0.771048614 |
| ENSMUSG0000010269 | chr3 | Dchs2 | -0.305066794 | 0.771112324 |
| ENSMUSG0000005253 | chr3 | Magi3 | 0.117672917 | 0.771112324 |
| ENSMUSG0000002212 | chr14 | Dct | -0.721411046 | 0.771136119 |
| ENSMUSG0000006086 | chr4 | Zbtb40 | -0.103435148 | 0.771136119 |
| ENSMUSG0000005275 | chr6 | Repin1 | 0.083732254 | 0.771136119 |
| ENSMUSG0000003241 | chr9 | Rwdd2a | -0.257176698 | 0.771639447 |
| ENSMUSG0000000481 | chr5 | Ccl24 | -0.228456455 | 0.771639447 |
| ENSMUSG0000002170 | chr13 | Serinc5 | -0.142930073 | 0.771639447 |
| ENSMUSG0000000083 | chrX | Fmr1 | -0.077898224 | 0.771639447 |
| ENSMUSG0000003043 | chr7 | U2af2 | 0.078583734 | 0.771639447 |
| ENSMUSG0000003773 | chr3 | Mynn | 0.085658763 | 0.771639447 |
| ENSMUSG0000001997 | chr10 | Hbs1l | 0.082409817 | 0.771639447 |
| ENSMUSG0000003419 | chr14 | Chmp7 | 0.103533478 | 0.771639447 |
| ENSMUSG0000004782 | chr3 | Pygo2 | 0.10839377 | 0.771639447 |
| ENSMUSG0000002047 | chr11 | Aebp1 | 0.228352478 | 0.771639447 |
| ENSMUSG0000007421 | chr7 | Cox7a1 | 0.541443886 | 0.771639447 |
| ENSMUSG0000003823 | chr1 | F11r | -0.262283023 | 0.771646951 |
| ENSMUSG0000000268 | chr12 | Prkd1 | 0.132832098 | 0.771646951 |
| ENSMUSG0000006296 | chr1 | Ufc1 | 0.162645252 | 0.771646951 |

| | | | | |
|---|---|---|---|---|
| ENSMUSG0000005497 | chr1 | Nyap2 | -0.261397755 | 0.771798356 |
| ENSMUSG0000004817 | chr7 | Mcmbp | 0.087324139 | 0.771798356 |
| ENSMUSG0000008527 | chr7 | Sbk3 | -0.400970911 | 0.771961955 |
| ENSMUSG0000005514 | chr8 | Klf2 | 0.214804771 | 0.771961955 |
| ENSMUSG0000004002 | chr8 | Elavl1 | 0.12721948 | 0.772078582 |
| ENSMUSG0000003582 | chr2 | Ppp1r26 | -0.121747215 | 0.772137449 |
| ENSMUSG0000002955 | chr5 | 2210016L21Rik | 0.09792936 | 0.772137449 |
| ENSMUSG0000003813 | chr13 | Rbm24 | 0.144397111 | 0.772137449 |
| ENSMUSG0000000512 | chr15 | Ccn4 | 0.179211613 | 0.772137449 |
| ENSMUSG0000002045 | chr11 | Rtn4 | 0.09142403 | 0.77238508 |
| ENSMUSG0000003946 | chr2 | Slc9a8 | -0.127334172 | 0.772410519 |
| ENSMUSG0000001867 | chr11 | Slc25a39 | 0.111419519 | 0.772410519 |
| ENSMUSG0000000078 | chr11 | Tcf7 | 0.129573864 | 0.772410519 |
| ENSMUSG0000006210 | chr17 | Zfp119b | 0.200780823 | 0.772445435 |
| ENSMUSG0000003126 | chrX | Cenpi | 0.545289321 | 0.77297576 |
| ENSMUSG0000003744 | chr10 | Vnn1 | -0.324259816 | 0.77309879 |
| ENSMUSG0000002543 | chr10 | Avil | -0.134570714 | 0.77309879 |
| ENSMUSG0000002153 | chr13 | Fastkd3 | -0.163238037 | 0.77309879 |
| ENSMUSG0000004416 | chr3 | Foxo1 | -0.112882951 | 0.77309879 |
| ENSMUSG0000007153 | chr16 | Pcnp | 0.117700774 | 0.77309879 |
| ENSMUSG0000003315 | chr6 | Podxl2 | 0.105483138 | 0.77309879 |
| ENSMUSG0000002956 | chr5 | Snx8 | 0.133688279 | 0.77309879 |
| ENSMUSG0000002524 | chr9 | Slc6a20b | 0.406793094 | 0.77309879 |
| ENSMUSG0000004297 | chr7 | Sbk1 | -0.145561664 | 0.773102994 |
| ENSMUSG0000003444 | chr12 | Trmt5 | 0.192159412 | 0.773102994 |
| ENSMUSG0000005027 | chr8 | Prag1 | 0.147086003 | 0.773343548 |
| ENSMUSG0000006807 | chr2 | Tcf15 | 0.714168409 | 0.773343548 |
| ENSMUSG0000004967 | chr7 | Catsperg1 | -0.333817349 | 0.77344118 |
| ENSMUSG0000002773 | chr3 | Rab33b | 0.090881079 | 0.77344118 |
| ENSMUSG0000003179 | chr8 | Usb1 | 0.102978644 | 0.77344118 |
| ENSMUSG0000004485 | chr1 | 1700056E22Rik | 0.43842596 | 0.77344118 |
| ENSMUSG0000004118 | chr11 | Chrnb1 | 0.12185899 | 0.77344118 |
| ENSMUSG0000004017 | chr2 | Alkbh3 | 0.13678064 | 0.77344118 |
| ENSMUSG0000004188 | chr1 | Shisa4 | 0.158217802 | 0.77344118 |
| ENSMUSG0000000029 | chr10 | Hddc2 | 0.182010493 | 0.77344118 |
| ENSMUSG0000004291 | chr7 | Mamstr | -0.308437961 | 0.77385121 |
| ENSMUSG0000002255 | chr15 | Fbxl6 | 0.132027552 | 0.773905765 |
| ENSMUSG0000005274 | chr1 | Swt1 | 0.144957607 | 0.773905765 |
| ENSMUSG0000002779 | chr3 | Bche | -0.136474086 | 0.774092777 |
| ENSMUSG0000006044 | chr7 | Trim5 | -0.227709055 | 0.774335254 |
| ENSMUSG0000003814 | chr17 | Tmem181a | -0.132664296 | 0.774335254 |
| ENSMUSG0000002052 | chr11 | Galnt10 | -0.094623495 | 0.774335254 |
| ENSMUSG0000002946 | chr5 | Anapc7 | 0.098267885 | 0.774335254 |
| ENSMUSG0000003912 | chr19 | Prune2 | 0.147085123 | 0.774335254 |
| ENSMUSG0000003557 | chr11 | Utp6 | 0.090097539 | 0.774579803 |
| ENSMUSG0000005438 | chr12 | Pnma1 | -0.262895541 | 0.774798211 |
| ENSMUSG0000007486 | chr13 | Zfp934 | -0.185381436 | 0.775001654 |
| ENSMUSG0000004721 | chr1 | Cdh19 | -0.224400004 | 0.775214432 |
| ENSMUSG0000002747 | chr2 | Ccm2l | -0.189976549 | 0.775214432 |
| ENSMUSG0000002638 | chr1 | Ptpn4 | -0.110935308 | 0.775214432 |

| | | | | |
|---|---|---|---|---|
| ENSMUSG0000000280 | chr12 | Nudt14 | -0.184667743 | 0.775214432 |
| ENSMUSG0000002753 | chr3 | Impa1 | 0.109809841 | 0.775214432 |
| ENSMUSG0000003692 | chr10 | Stox1 | 0.245818593 | 0.775214432 |
| ENSMUSG0000002169 | chr13 | Ercc8 | 0.160721924 | 0.775214432 |
| ENSMUSG0000002280 | chr16 | Osbpl11 | 0.137106309 | 0.775271492 |
| ENSMUSG0000002109 | chr12 | 4930447C04Rik | -0.356757941 | 0.775562238 |
| ENSMUSG0000003129 | chrX | Lrch2 | 0.101141961 | 0.775562238 |
| ENSMUSG0000002287 | chr16 | Fetub | 0.390253013 | 0.775562238 |
| ENSMUSG0000006053 | chr7 | Tmem219 | 0.122996458 | 0.775562238 |
| ENSMUSG0000005750 | chr19 | Bloc1s2 | 0.151343678 | 0.775562238 |
| ENSMUSG0000002435 | chr18 | Slc23a1 | 0.396811725 | 0.775562238 |
| ENSMUSG0000002754 | chr2 | Ptpn1 | 0.098142304 | 0.775629209 |
| ENSMUSG0000002802 | chr3 | Pitx2 | -1.102251247 | 0.775891013 |
| ENSMUSG0000003075 | chr7 | Zkscan2 | -0.182122716 | 0.775891013 |
| ENSMUSG0000004587 | chr14 | Slitrk6 | -0.215681279 | 0.775891013 |
| ENSMUSG0000002425 | chr17 | Adcyap1 | -0.227836588 | 0.775891013 |
| ENSMUSG0000006426 | chr6 | Gimap8 | -0.197721313 | 0.775891013 |
| ENSMUSG0000005470 | chr10 | Ankrd24 | -0.161195868 | 0.775891013 |
| ENSMUSG0000002193 | chr14 | Gucy1b2 | -0.410460072 | 0.775891013 |
| ENSMUSG0000002825 | chr3 | Odf2l | -0.121460583 | 0.775891013 |
| ENSMUSG0000007908 | chr9 | Jrkl | -0.120547091 | 0.775891013 |
| ENSMUSG0000007354 | chr18 | Cep76 | -0.112562326 | 0.775891013 |
| ENSMUSG0000002202 | chr14 | Sugt1 | 0.082687215 | 0.775891013 |
| ENSMUSG0000003984 | chr2 | Pcif1 | 0.08627908 | 0.775891013 |
| ENSMUSG0000002905 | chr4 | Pank4 | 0.097412483 | 0.775891013 |
| ENSMUSG0000002767 | chr3 | Actl6a | 0.123841989 | 0.775891013 |
| ENSMUSG0000004211 | chr1 | Ccdc115 | 0.118363426 | 0.775891013 |
| ENSMUSG0000002099 | chr12 | Cdkl1 | 0.125305022 | 0.775891013 |
| ENSMUSG0000006430 | chr7 | Lrrc51 | 0.203576573 | 0.775891013 |
| ENSMUSG0000006054 | chr14 | Tnfrsf19 | 0.13302496 | 0.775891013 |
| ENSMUSG0000001485 | chr8 | Tmem208 | 0.162428965 | 0.775891013 |
| ENSMUSG0000007058 | chr4 | Fv1 | 0.178308537 | 0.775891013 |
| ENSMUSG0000002972 | chr5 | Ppp1r35 | 0.17936741 | 0.775891013 |
| ENSMUSG0000000077 | chr14 | Polr3d | 0.155157418 | 0.775891013 |
| ENSMUSG0000000601 | chr1 | Prg4 | 0.305910163 | 0.775891013 |
| ENSMUSG0000007830 | chr9 | AI593442 | 0.328250826 | 0.775891013 |
| ENSMUSG0000002024 | chr10 | Hcfc2 | 0.112596836 | 0.775968955 |
| ENSMUSG0000010325 | chr18 | Pcdhac1 | -0.424361775 | 0.776058015 |
| ENSMUSG0000002199 | chr14 | Mipep | 0.139911562 | 0.776160439 |
| ENSMUSG0000003048 | chr7 | Zfp108 | 0.200807302 | 0.776160439 |
| ENSMUSG0000006251 | chr1 | Nsl1 | -0.716610854 | 0.776326292 |
| ENSMUSG0000004956 | chr19 | Ap5b1 | -0.132196085 | 0.776507387 |
| ENSMUSG0000002384 | chr17 | Lnpep | -0.111264642 | 0.776721168 |
| ENSMUSG0000007447 | chr9 | Spc24 | -0.514904142 | 0.777288043 |
| ENSMUSG0000007307 | chrX | Cfap47 | -0.35147949 | 0.777288043 |
| ENSMUSG0000001419 | chr11 | Zfp385c | -0.399361659 | 0.777288043 |
| ENSMUSG0000003900 | chr5 | Ube3c | -0.090408141 | 0.777288043 |
| ENSMUSG0000002623 | chr1 | Ncl | 0.113363928 | 0.777288043 |
| ENSMUSG0000007379 | chr4 | Efcab7 | 0.156412629 | 0.777288043 |
| ENSMUSG0000003016 | chr6 | Rad52 | 0.118725354 | 0.777288043 |

| | | | | |
|---|---|---|---|---|
| ENSMUSG0000005665 | chr15 | Apol8 | 0.398164431 | 0.777288043 |
| ENSMUSG0000005997 | chr9 | Ntm | 0.239917448 | 0.777288043 |
| ENSMUSG0000002991 | chr6 | Qrfprl | 0.297988079 | 0.777288043 |
| ENSMUSG000000029€ | chr7 | Pnkp | 0.132275139 | 0.777440721 |
| ENSMUSG0000006615 | chr4 | Fkbp15 | -0.102312709 | 0.777681345 |
| ENSMUSG0000003483 | chr6 | Tet3 | -0.175059848 | 0.777739047 |
| ENSMUSG000000204€ | chr11 | Efemp1 | -0.190125826 | 0.777739047 |
| ENSMUSG000000553C | chr3 | Adh7 | 0.615786856 | 0.777739047 |
| ENSMUSG0000002069 | chr11 | Efcab3 | 0.479748299 | 0.777739047 |
| ENSMUSG0000005173 | chr10 | Fam229b | -0.239869271 | 0.777903746 |
| ENSMUSG000000306C | chr7 | Lrfn1 | -0.155396865 | 0.778003991 |
| ENSMUSG0000006682 | chr9 | Zfp810 | 0.139412084 | 0.778003991 |
| ENSMUSG0000006858 | chr6 | Mgam | -0.810425633 | 0.77835617 |
| ENSMUSG0000004439 | chr19 | Tigd3 | -0.482757969 | 0.77835617 |
| ENSMUSG000000227C | chr16 | Mrpl40 | -0.168671874 | 0.77835617 |
| ENSMUSG0000004161 | chr16 | Ccdc74a | -0.229602244 | 0.77835617 |
| ENSMUSG0000004083 | chr1 | Gpr161 | -0.150057437 | 0.77835617 |
| ENSMUSG0000002664 | chr2 | Suv39h2 | -0.210167991 | 0.77835617 |
| ENSMUSG000000212€ | chr12 | Evl | 0.098756357 | 0.77835617 |
| ENSMUSG0000000561 | chr7 | Irag1 | 0.118103543 | 0.77835617 |
| ENSMUSG000000201C | chr10 | Dnajb12 | 0.119576746 | 0.77835617 |
| ENSMUSG0000004623 | chr19 | Vps13a | 0.212006563 | 0.77835617 |
| ENSMUSG000000315C | chr8 | Ankrd10 | 0.082492382 | 0.778582596 |
| ENSMUSG0000004207 | chr6 | Hnrnpf | 0.092601173 | 0.77862002 |
| ENSMUSG0000002793 | chr3 | Nup210l | 0.316069545 | 0.778621587 |
| ENSMUSG0000002758 | chr2 | Stmn3 | -0.162138031 | 0.778927249 |
| ENSMUSG0000000847 | chr1 | Arpc5 | 0.084046955 | 0.778927249 |
| ENSMUSG0000002593 | chr1 | Slco5a1 | 0.335212713 | 0.778927249 |
| ENSMUSG0000000541 | chr8 | Mcm5 | -0.325120082 | 0.77919253 |
| ENSMUSG0000011137 | chr5 | Btbd8 | -0.163087892 | 0.77919253 |
| ENSMUSG0000003582 | chr8 | Tk2 | -0.125022352 | 0.77919253 |
| ENSMUSG0000005151 | chr11 | Mafg | -0.111190794 | 0.77919253 |
| ENSMUSG000000600C | chr10 | Chpt1 | 0.131050822 | 0.77919253 |
| ENSMUSG000000208C | chr11 | Med31 | 0.193349701 | 0.77919253 |
| ENSMUSG0000007917 | chr12 | Fam228a | -0.614456416 | 0.779840586 |
| ENSMUSG0000004173 | chr15 | Tspo | 0.217430628 | 0.779840586 |
| ENSMUSG0000006787 | chr1 | Vxn | -0.400429509 | 0.780093976 |
| ENSMUSG0000005464 | chr8 | Zfp869 | -0.116908483 | 0.780093976 |
| ENSMUSG0000000631 | chr7 | Upk1a | -0.451496611 | 0.780093976 |
| ENSMUSG0000004174 | chr13 | Utp15 | 0.091265423 | 0.780093976 |
| ENSMUSG0000002109 | chr12 | Dhrs7 | 0.125109676 | 0.780093976 |
| ENSMUSG0000000239 | chr8 | Nr2f6 | 0.180126489 | 0.780093976 |
| ENSMUSG0000003438 | chr6 | Ssu2 | 0.313097425 | 0.780093976 |
| ENSMUSG0000002138 | chr13 | Susd3 | 0.405172649 | 0.780093976 |
| ENSMUSG000000016C | chr4 | Tcea3 | -0.191818551 | 0.780176816 |
| ENSMUSG0000000587 | chr18 | Apc | -0.144640125 | 0.780176816 |
| ENSMUSG000001186€ | chr5 | Lin54 | -0.154581936 | 0.780176816 |
| ENSMUSG0000001745 | chr11 | Pipox | -0.546061006 | 0.780176816 |
| ENSMUSG000000340€ | chr6 | 4930590J08Rik | -0.341178965 | 0.780176816 |
| ENSMUSG0000002175 | chr14 | Acox2 | -0.525032606 | 0.780176816 |

| | | | | |
|---|---|---|---|---|
| ENSMUSG0000005777 | chr11 | Cyb5d2 | -0.117328965 | 0.780176816 |
| ENSMUSG0000001870 | chr12 | Dync1h1 | -0.113366314 | 0.780176816 |
| ENSMUSG0000009647 | chr9 | Cdkn2d | 0.15034489 | 0.780176816 |
| ENSMUSG0000009011 | chr10 | Shprh | -0.101712508 | 0.780537146 |
| ENSMUSG0000002396 | chr17 | Rsph9 | 0.124068674 | 0.780537146 |
| ENSMUSG0000009093 | chr12 | Synj2bp | 0.139034246 | 0.780537146 |
| ENSMUSG0000002279 | chr16 | Yars2 | 0.153436809 | 0.780537146 |
| ENSMUSG0000001649 | chr1 | Cd34 | 0.151004382 | 0.780537146 |
| ENSMUSG0000005947 | chr7 | B3gnt8 | 0.16059764 | 0.780537146 |
| ENSMUSG0000004488 | chr7 | Coa4 | 0.234499298 | 0.780537146 |
| ENSMUSG0000004531 | chr5 | Adra2c | 0.23740313 | 0.780537146 |
| ENSMUSG0000003003 | chr6 | Wbp1 | 0.143596188 | 0.780652724 |
| ENSMUSG0000000144 | chr11 | Npepps | 0.076701955 | 0.781039043 |
| ENSMUSG0000005100 | chr7 | Gatd1 | 0.146266225 | 0.781039043 |
| ENSMUSG0000008222 | chrX | Gm715 | -0.512518036 | 0.781813177 |
| ENSMUSG0000003390 | chr7 | Ccp110 | -0.11551723 | 0.781953802 |
| ENSMUSG0000004617 | chr6 | Nxph1 | -0.293868444 | 0.782013459 |
| ENSMUSG0000002646 | chr1 | Atp2b4 | 0.119916342 | 0.782117119 |
| ENSMUSG0000003795 | chr11 | Nsrp1 | 0.137676727 | 0.782229568 |
| ENSMUSG0000002706 | chr2 | Dhrs9 | -0.490687635 | 0.782340756 |
| ENSMUSG0000002657 | chr1 | F5 | -0.556140458 | 0.782340756 |
| ENSMUSG0000001984 | chr10 | Lama4 | 0.135056473 | 0.782340756 |
| ENSMUSG0000002221 | chr14 | Dhrs4 | 0.149716907 | 0.782340756 |
| ENSMUSG0000005054 | chr12 | Fam228b | -0.337542692 | 0.782385367 |
| ENSMUSG0000009255 | chr17 | Med20 | -0.114793903 | 0.782385367 |
| ENSMUSG0000002404 | chr17 | Epb41l3 | -0.103849186 | 0.782385367 |
| ENSMUSG0000002581 | chr2 | Sec61a2 | -0.091685249 | 0.782385367 |
| ENSMUSG0000002673 | chr2 | Bmi1 | 0.104234194 | 0.782385367 |
| ENSMUSG0000002296 | chr16 | Gart | 0.124284754 | 0.782385367 |
| ENSMUSG0000002210 | chr14 | Dok2 | 0.232516792 | 0.782385367 |
| ENSMUSG0000006018 | chr10 | Lrrc10 | 0.534796645 | 0.782385367 |
| ENSMUSG0000004203 | chr3 | Igsf3 | 0.095986194 | 0.782502391 |
| ENSMUSG0000003043 | chr7 | Sbk2 | -0.515539249 | 0.782580196 |
| ENSMUSG0000008977 | chr16 | Slc5a3 | -0.173651113 | 0.782605574 |
| ENSMUSG0000003742 | chr5 | Vgf | -0.263160682 | 0.782646843 |
| ENSMUSG0000003276 | chr10 | Ilvbl | -0.182876295 | 0.782646843 |
| ENSMUSG0000002938 | chr5 | Ccng2 | -0.202902694 | 0.782646843 |
| ENSMUSG0000002485 | chr19 | Pitpnm1 | -0.107650389 | 0.782646843 |
| ENSMUSG0000004351 | chr5 | Hscb | 0.161240054 | 0.782646843 |
| ENSMUSG0000002779 | chr3 | Nbea | -0.136706877 | 0.782976362 |
| ENSMUSG0000002903 | chr4 | Atad3a | -0.103452632 | 0.783139705 |
| ENSMUSG0000003485 | chr2 | Tmem127 | -0.063572553 | 0.783139705 |
| ENSMUSG0000002730 | chr2 | Rtf1 | 0.07925038 | 0.783139705 |
| ENSMUSG0000000142 | chr3 | Tmem79 | -0.294851601 | 0.783212759 |
| ENSMUSG0000002284 | chr16 | Hspbap1 | 0.164525026 | 0.783247465 |
| ENSMUSG0000009145 | chr10 | Otogl | 0.156562355 | 0.783247465 |
| ENSMUSG0000005983 | chr3 | Sclt1 | -0.230197358 | 0.78356826 |
| ENSMUSG0000000131 | chr11 | Rnd2 | -0.169438625 | 0.78356826 |
| ENSMUSG0000003069 | chr7 | Fchsd2 | -0.122102767 | 0.78356826 |
| ENSMUSG0000010274 | chr18 | Pcdhga11 | -0.16441472 | 0.78356826 |

| | | | | |
|---|---|---|---|---|
| ENSMUSG0000002280 | chr16 | Snx4 | 0.077631547 | 0.78356826 |
| ENSMUSG0000002203 | chr14 | Elp3 | 0.073099543 | 0.78356826 |
| ENSMUSG0000003841 | chr8 | Cdc16 | 0.094891049 | 0.78356826 |
| ENSMUSG0000003688 | chr18 | Acaa2 | 0.189171214 | 0.78356826 |
| ENSMUSG0000001407 | chr16 | Dynlt2b | 0.142772721 | 0.78356826 |
| ENSMUSG0000007119 | chr17 | Wfikkn1 | 0.326486982 | 0.78356826 |
| ENSMUSG0000003008 | chr6 | Slc41a3 | 0.210075337 | 0.78356826 |
| ENSMUSG0000002593 | chr1 | Gsta3 | 0.486379199 | 0.78356826 |
| ENSMUSG0000002490 | chr19 | Mus81 | 0.105466576 | 0.783950331 |
| ENSMUSG0000009813 | chr12 | Rnf113a2 | 0.141513724 | 0.783950331 |
| ENSMUSG0000003224 | chr9 | Itga11 | 0.160533082 | 0.78400771 |
| ENSMUSG0000003837 | chr9 | Ttk | -0.553984182 | 0.784054406 |
| ENSMUSG0000004993 | chr9 | H2ax | 0.140973829 | 0.78420678 |
| ENSMUSG0000003812 | chr16 | Ccdc50 | 0.070564392 | 0.784406738 |
| ENSMUSG0000004432 | chr2 | 1700001O22Rik | -0.592925115 | 0.784769398 |
| ENSMUSG0000003478 | chr10 | Gna11 | -0.077840615 | 0.785027232 |
| ENSMUSG0000002003 | chr10 | Tcp11l2 | 0.156400391 | 0.785283115 |
| ENSMUSG0000002103 | chr12 | Gstz1 | -0.075895767 | 0.785329733 |
| ENSMUSG0000002118 | chr12 | Fbln5 | 0.107327756 | 0.785329733 |
| ENSMUSG0000004281 | chr2 | Mcts2 | 0.143776576 | 0.785515437 |
| ENSMUSG0000002057 | chr12 | Pik3cg | -0.19573405 | 0.785616141 |
| ENSMUSG0000005796 | chr9 | Sema3b | -0.144193447 | 0.785642641 |
| ENSMUSG0000003628 | chr14 | Naa30 | -0.077955911 | 0.785675893 |
| ENSMUSG0000005347 | chr2 | Tnfaip6 | 0.221056132 | 0.785675893 |
| ENSMUSG0000003225 | chr9 | Kif23 | -0.520863446 | 0.785889809 |
| ENSMUSG0000001014 | chr11 | Tnfrsf13b | -0.478284853 | 0.785889809 |
| ENSMUSG0000009021 | chr3 | Itga10 | -0.245346266 | 0.785889809 |
| ENSMUSG0000004451 | chr14 | Zfp488 | -0.175513614 | 0.785889809 |
| ENSMUSG0000002401 | chr17 | Fgd2 | -0.236573469 | 0.785889809 |
| ENSMUSG0000002000 | chr10 | Il20ra | -0.227460742 | 0.785889809 |
| ENSMUSG0000003852 | chr18 | Fchsd1 | -0.104706342 | 0.785889809 |
| ENSMUSG0000006018 | chr10 | Slc35e3 | 0.09469023 | 0.785889809 |
| ENSMUSG0000001937 | chr11 | Cops3 | 0.120905021 | 0.785889809 |
| ENSMUSG0000003167 | chr8 | Setd6 | 0.111430166 | 0.785889809 |
| ENSMUSG0000004546 | chr7 | Ttll13 | 0.357712612 | 0.785889809 |
| ENSMUSG0000002171 | chr13 | Cwc27 | 0.134787757 | 0.785889809 |
| ENSMUSG0000006604 | chr4 | Med18 | 0.252301317 | 0.785889809 |
| ENSMUSG0000002308 | chr3 | Lrrc71 | 0.467794627 | 0.785889809 |
| ENSMUSG0000004674 | chr3 | Fat4 | 0.214956059 | 0.785889809 |
| ENSMUSG0000003086 | chr7 | Cpxm2 | -0.099681377 | 0.785980043 |
| ENSMUSG0000004955 | chr6 | Polr1a | 0.075154083 | 0.785980043 |
| ENSMUSG0000003373 | chr8 | Sf3b3 | 0.057664038 | 0.786007384 |
| ENSMUSG0000003260 | chr9 | Klhdc8b | 0.091865072 | 0.786013836 |
| ENSMUSG0000005201 | chr16 | Btla | -0.666530818 | 0.78606981 |
| ENSMUSG0000004072 | chr7 | Hnrnpul1 | 0.089312775 | 0.78606981 |
| ENSMUSG0000002576 | chr3 | D3Ertd751e | -0.12846901 | 0.786129098 |
| ENSMUSG0000002551 | chr7 | Chid1 | 0.157729809 | 0.786129098 |
| ENSMUSG0000004004 | chr14 | Ift88 | 0.102778138 | 0.786184117 |
| ENSMUSG0000004383 | chr7 | Lysmd4 | 0.09549179 | 0.786184117 |
| ENSMUSG0000004842 | chr15 | Ranbp3l | -0.556036612 | 0.786226121 |

| | | | | |
|---|---|---|---|---|
| ENSMUSG0000002062 | chr12 | Adi1 | -0.120406663 | 0.786226121 |
| ENSMUSG0000002660 | chr1 | Soat1 | -0.11856356 | 0.786226121 |
| ENSMUSG0000003301 | chr3 | Trim33 | -0.093991854 | 0.786226121 |
| ENSMUSG0000002256 | chr15 | Slc52a2 | -0.117750555 | 0.786226121 |
| ENSMUSG0000004220 | chr1 | Kdm5b | 0.101394512 | 0.786226121 |
| ENSMUSG0000004177 | chr12 | Slc24a4 | 0.229562738 | 0.786226121 |
| ENSMUSG0000002962 | chr5 | Cpsf4 | 0.124943171 | 0.786226121 |
| ENSMUSG0000003259 | chr9 | Nckipsd | -0.100363207 | 0.786278844 |
| ENSMUSG0000003291 | chr3 | Lrig2 | -0.112985509 | 0.786307365 |
| ENSMUSG0000003685 | chr4 | Wnt4 | 0.493912148 | 0.786649273 |
| ENSMUSG0000006640 | chr7 | Akap13 | -0.120986344 | 0.786687788 |
| ENSMUSG0000000639 | chr4 | Elovl1 | 0.088668549 | 0.786687788 |
| ENSMUSG0000007477 | chr2 | Ankef1 | -0.459695367 | 0.786773075 |
| ENSMUSG0000004201 | chr13 | Wdr41 | -0.095756965 | 0.786773075 |
| ENSMUSG0000006927 | chr13 | H2ac6 | 0.380562696 | 0.786773075 |
| ENSMUSG0000003055 | chr7 | Rab38 | 0.45202682 | 0.786857622 |
| ENSMUSG0000004366 | chr8 | Tmem221 | -0.429144147 | 0.786880892 |
| ENSMUSG0000001060 | chr1 | Psen2 | 0.105374578 | 0.786880892 |
| ENSMUSG0000002277 | chr16 | Ncbp2 | 0.135773837 | 0.786880892 |
| ENSMUSG0000003889 | chr8 | Irs2 | 0.167059663 | 0.786880892 |
| ENSMUSG0000006756 | chrX | Hdac8 | -0.187285297 | 0.786994787 |
| ENSMUSG0000002724 | chr2 | Hypk | -0.226763478 | 0.786994787 |
| ENSMUSG0000003971 | chr4 | Plekhg5 | -0.143386443 | 0.786994787 |
| ENSMUSG0000005002 | chrX | Rap2c | -0.079208981 | 0.786994787 |
| ENSMUSG0000002463 | chr19 | Gnaq | 0.094516007 | 0.786994787 |
| ENSMUSG0000003204 | chr9 | Srpra | 0.106711859 | 0.786994787 |
| ENSMUSG0000002826 | chr4 | Ndufaf4 | 0.122296643 | 0.786994787 |
| ENSMUSG0000002557 | chr11 | Cbx2 | 0.130094412 | 0.786994787 |
| ENSMUSG0000000170 | chr13 | Eef1e1 | 0.211729179 | 0.786994787 |
| ENSMUSG0000003828 | chr13 | Bphl | 0.161213212 | 0.787039137 |
| ENSMUSG0000007474 | chr10 | Atxn7l3b | -0.080648042 | 0.787105827 |
| ENSMUSG0000007133 | chr6 | Tia1 | -0.117617675 | 0.787294013 |
| ENSMUSG0000003023 | chr6 | Plcz1 | -0.454871845 | 0.787424198 |
| ENSMUSG0000002888 | chr4 | Yrdc | 0.166381595 | 0.787424198 |
| ENSMUSG0000001972 | chr13 | Lyst | -0.133620932 | 0.78750218 |
| ENSMUSG0000003537 | chr16 | Hacd2 | 0.129135681 | 0.78750218 |
| ENSMUSG0000004306 | chr8 | Fam89a | -0.192726797 | 0.787565597 |
| ENSMUSG0000003805 | chr5 | Kmt2c | -0.145444468 | 0.787855056 |
| ENSMUSG0000009510 | chr13 | Edaradd | -0.545289846 | 0.787855056 |
| ENSMUSG0000003218 | chr9 | Smarca4 | 0.080638262 | 0.788073146 |
| ENSMUSG0000006262 | chr4 | Mysm1 | -0.108143854 | 0.788284536 |
| ENSMUSG0000003784 | chr1 | Ifi206 | -0.488608402 | 0.78838122 |
| ENSMUSG0000004521 | chr1 | Hs6st1 | -0.117600675 | 0.78838122 |
| ENSMUSG0000002504 | chr19 | Taf5 | 0.142298727 | 0.78838122 |
| ENSMUSG0000001698 | chr11 | Etaa1 | -0.120198096 | 0.788382553 |
| ENSMUSG0000005137 | chr2 | Plpp7 | 0.149555264 | 0.788382553 |
| ENSMUSG0000002943 | chr5 | Vps33a | -0.069682359 | 0.788876555 |
| ENSMUSG0000002782 | chr3 | Ccnl1 | -0.07293279 | 0.788987194 |
| ENSMUSG0000002870 | chr4 | Lrrc41 | 0.089093873 | 0.788987194 |
| ENSMUSG0000003716 | chr12 | Mycn | 0.327170783 | 0.789520444 |

| | | | | |
|---|---|---|---|---|
| ENSMUSG0000002006 | chr10 | Sirt1 | 0.091881537 | 0.789634078 |
| ENSMUSG0000003337 | chr4 | Atp6v0b | 0.112493639 | 0.789634078 |
| ENSMUSG0000000105 | chr3 | Sec24b | -0.098920034 | 0.789963877 |
| ENSMUSG0000005524 | chr17 | Zfp101 | -0.115107335 | 0.790043792 |
| ENSMUSG0000005067 | chr5 | Ccdc96 | 0.158530984 | 0.790043792 |
| ENSMUSG0000003794 | chr8 | Inpp4b | 0.215824222 | 0.790043792 |
| ENSMUSG0000002405 | chr17 | Smchd1 | -0.09535155 | 0.790205774 |
| ENSMUSG0000005346 | chr6 | Ggcx | 0.111236573 | 0.790287971 |
| ENSMUSG0000004257 | chr3 | Ube2q1 | -0.076410193 | 0.790534956 |
| ENSMUSG0000002421 | chr6 | Grm8 | -0.638166895 | 0.790726072 |
| ENSMUSG0000002600 | chr1 | Rpe | 0.106399461 | 0.790763661 |
| ENSMUSG0000004250 | chr12 | Mideas | -0.11835123 | 0.790790335 |
| ENSMUSG0000002592 | chr1 | Terf1 | -0.121786993 | 0.790790335 |
| ENSMUSG0000004542 | chrX | Hnrnph2 | 0.06838379 | 0.790790335 |
| ENSMUSG0000004058 | chrX | Ofd1 | 0.123276203 | 0.790814102 |
| ENSMUSG0000004362 | chr1 | 1700019D03Rik | -0.301734706 | 0.791143561 |
| ENSMUSG0000002748 | chr2 | Necab3 | -0.475110864 | 0.791143561 |
| ENSMUSG0000004334 | chr2 | Hoxd9 | 0.121645707 | 0.791143561 |
| ENSMUSG0000007479 | chr2 | Hspa12b | -0.151920649 | 0.791187144 |
| ENSMUSG0000007128 | chr13 | Zfp65 | 0.103861842 | 0.791187144 |
| ENSMUSG0000006137 | chr7 | Fiz1 | 0.119467159 | 0.791187144 |
| ENSMUSG0000001989 | chr10 | Nts | -0.345313794 | 0.791810523 |
| ENSMUSG0000000141 | chr3 | Smg5 | 0.078029254 | 0.791810523 |
| ENSMUSG0000007452 | chr2 | Zfp1003 | 0.106477449 | 0.791810523 |
| ENSMUSG0000000760 | chr17 | Dus3l | 0.087065131 | 0.791810523 |
| ENSMUSG0000003945 | chr15 | Mtmr12 | 0.115470083 | 0.791810523 |
| ENSMUSG0000001004 | chr9 | Ifrd2 | 0.155047839 | 0.791810523 |
| ENSMUSG0000006048 | chr10 | Samd5 | 0.209890178 | 0.791810523 |
| ENSMUSG0000003934 | chr1 | C130074G19Rik | -0.222334985 | 0.792301078 |
| ENSMUSG0000002667 | chr1 | Ddr2 | 0.104145757 | 0.792345643 |
| ENSMUSG0000003285 | chr17 | Pkd1 | 0.153009517 | 0.792345643 |
| ENSMUSG0000001982 | chr10 | Zbtb24 | 0.08582338 | 0.79280282 |
| ENSMUSG0000003533 | chr9 | Uchl4 | 0.45095855 | 0.79280282 |
| ENSMUSG0000005358 | chr5 | Zfand2a | 0.102951282 | 0.792835098 |
| ENSMUSG0000007229 | chr14 | Klf12 | 0.20977434 | 0.792922392 |
| ENSMUSG0000006063 | chr16 | Rpl35a | 0.41545117 | 0.792922392 |
| ENSMUSG0000002466 | chr19 | Fads2 | -0.085044788 | 0.793150735 |
| ENSMUSG0000005941 | chr9 | Fxyd2 | -0.215044566 | 0.793208557 |
| ENSMUSG0000002937 | chr5 | Mthfd2l | 0.123915039 | 0.793219416 |
| ENSMUSG0000003445 | chrX | Eda2r | 0.176299962 | 0.793219416 |
| ENSMUSG0000000093 | chr6 | Hoxa10 | 0.134049677 | 0.793410782 |
| ENSMUSG0000003342 | chr7 | Mcee | 0.168181647 | 0.793410782 |
| ENSMUSG0000002733 | chr2 | Knstrn | -0.517423216 | 0.793489135 |
| ENSMUSG0000002800 | chr3 | Snx7 | -0.107783961 | 0.793489135 |
| ENSMUSG0000001597 | chr14 | Actr8 | 0.082127077 | 0.793489135 |
| ENSMUSG0000003696 | chr11 | Trim17 | 0.246547941 | 0.793637318 |
| ENSMUSG0000004242 | chr1 | Adora1 | -0.160317425 | 0.793743467 |
| ENSMUSG0000004263 | chr15 | Pla2g6 | -0.143035457 | 0.793743467 |
| ENSMUSG0000000526 | chr16 | Ufd1 | 0.06272208 | 0.793743467 |
| ENSMUSG0000003681 | chr11 | Jmjd4 | 0.113152032 | 0.793743467 |

| | | | | |
|---|---|---|---|---|
| ENSMUSG0000004416 | chr13 | Rnf182 | 0.238903385 | 0.793743467 |
| ENSMUSG0000005331 | chr1 | Slamf8 | -0.546410839 | 0.793781313 |
| ENSMUSG0000004591 | chr10 | C2cd4c | -0.280980779 | 0.794335157 |
| ENSMUSG0000005429 | chrX | P2ry10b | -0.403976533 | 0.794785123 |
| ENSMUSG0000002456 | chr18 | Atp9b | -0.086415934 | 0.794785123 |
| ENSMUSG0000001713 | chr11 | Cyth1 | -0.091468962 | 0.794785123 |
| ENSMUSG0000004324 | chr2 | Upf2 | 0.102980172 | 0.794785123 |
| ENSMUSG0000004384 | chr7 | Tmem145 | -0.250279405 | 0.795039108 |
| ENSMUSG0000006813 | chr2 | Zfp442 | -0.271748962 | 0.795039108 |
| ENSMUSG0000005122 | chrX | Ercc6l | -0.371721354 | 0.795039108 |
| ENSMUSG0000000767 | chr17 | Khsrp | 0.083140232 | 0.795039108 |
| ENSMUSG0000000191 | chr8 | Nacc1 | 0.088095814 | 0.795039108 |
| ENSMUSG0000003991 | chr10 | Coq10a | 0.10217718 | 0.795039108 |
| ENSMUSG0000006845 | chrY | Uty | 0.103978085 | 0.795039108 |
| ENSMUSG0000002519 | chr19 | Cutc | 0.223970056 | 0.795039108 |
| ENSMUSG0000011019 | chr7 | Pde2a | 0.400100718 | 0.795039108 |
| ENSMUSG0000004013 | chr2 | Gpr176 | 0.207874349 | 0.795039108 |
| ENSMUSG0000004070 | chr7 | Cyp2s1 | -0.397863722 | 0.795043759 |
| ENSMUSG0000002066 | chr12 | Pomc | -0.702102823 | 0.795043759 |
| ENSMUSG0000000056 | chr5 | Gabra2 | -0.694278979 | 0.795043759 |
| ENSMUSG0000003770 | chr6 | Fam13a | -0.1729237 | 0.795043759 |
| ENSMUSG0000002170 | chr13 | Dhfr | -0.194440156 | 0.795043759 |
| ENSMUSG0000011667 | chr16 | A630089N07Rik | -0.185860685 | 0.795043759 |
| ENSMUSG0000004167 | chr6 | Pyroxd1 | -0.145554869 | 0.795043759 |
| ENSMUSG0000000466 | chr9 | Arid3b | -0.141929243 | 0.795043759 |
| ENSMUSG0000003138 | chrX | Piga | -0.152722601 | 0.795043759 |
| ENSMUSG0000001973 | chr8 | Slc35e1 | -0.102396442 | 0.795043759 |
| ENSMUSG0000002143 | chr13 | Slc35b3 | -0.125599022 | 0.795043759 |
| ENSMUSG0000004018 | chr7 | Odad1 | -0.125416228 | 0.795043759 |
| ENSMUSG0000003136 | chrX | Zfp275 | -0.107218428 | 0.795043759 |
| ENSMUSG0000003248 | chr9 | Smarcc1 | -0.096629429 | 0.795043759 |
| ENSMUSG0000003802 | chr4 | Dennd4c | -0.09675347 | 0.795043759 |
| ENSMUSG0000006771 | chr6 | Lpar5 | -0.667701592 | 0.795043759 |
| ENSMUSG0000002182 | chr14 | Ap3m1 | 0.070688667 | 0.795043759 |
| ENSMUSG0000000178 | chr10 | Rtcb | 0.082522553 | 0.795043759 |
| ENSMUSG0000002260 | chr15 | Ptk2 | 0.092338724 | 0.795043759 |
| ENSMUSG0000002923 | chr5 | Tmem165 | 0.087536093 | 0.795043759 |
| ENSMUSG0000002845 | chr4 | Unc13b | 0.097404832 | 0.795043759 |
| ENSMUSG0000003997 | chr11 | Tbc1d16 | 0.097380034 | 0.795043759 |
| ENSMUSG0000002414 | chr17 | Rhoq | 0.119528925 | 0.795043759 |
| ENSMUSG0000002762 | chr2 | Aar2 | 0.136121069 | 0.795043759 |
| ENSMUSG0000003802 | chr6 | Tigar | 0.148418731 | 0.795043759 |
| ENSMUSG0000003919 | chr2 | Bbln | 0.146341185 | 0.795043759 |
| ENSMUSG0000004014 | chrX | Maob | 0.146889224 | 0.795043759 |
| ENSMUSG0000007886 | chr2 | Gm14412 | 0.376085074 | 0.795043759 |
| ENSMUSG0000007160 | chr19 | Entrep1 | 0.179269892 | 0.795043759 |
| ENSMUSG0000005455 | chr7 | Adam12 | 0.309887771 | 0.79513005 |
| ENSMUSG0000006659 | chr1 | Flvcr1 | -0.110992096 | 0.795335193 |
| ENSMUSG0000002159 | chr13 | Srd5a1 | -0.221370337 | 0.795565723 |
| ENSMUSG0000002891 | chr4 | Casp9 | -0.126666731 | 0.795565723 |

| | | | | |
|---|---|---|---|---|
| ENSMUSG0000001913 | chr13 | BC005537 | -0.082232855 | 0.795565723 |
| ENSMUSG0000003139 | chrX | Irak1 | 0.113403028 | 0.795565723 |
| ENSMUSG0000007073 | chr5 | Fryl | -0.145902023 | 0.79629748 |
| ENSMUSG0000005240 | chr2 | Rexo4 | 0.079085503 | 0.796394032 |
| ENSMUSG0000009039 | chr1 | 4930523C07Rik | 0.152325624 | 0.796482372 |
| ENSMUSG0000004478 | chr1 | Hjurp | -0.169668616 | 0.796507853 |
| ENSMUSG0000002294 | chr16 | Chaf1b | -0.330979225 | 0.796507853 |
| ENSMUSG0000006109 | chr7 | Gapdhs | 0.23704679 | 0.796507853 |
| ENSMUSG0000007366 | chr1 | Nbeal1 | -0.122814909 | 0.7965482 |
| ENSMUSG0000002169 | chr13 | Depdc1b | -0.699279323 | 0.796695808 |
| ENSMUSG0000005465 | chr4 | Pm20d2 | -0.293003028 | 0.796695808 |
| ENSMUSG0000006264 | chr2 | Ganc | -0.109366901 | 0.796695808 |
| ENSMUSG0000003082 | chr7 | Nucb1 | 0.072532724 | 0.796695808 |
| ENSMUSG0000002484 | chr19 | Banf1 | 0.132936659 | 0.796695808 |
| ENSMUSG0000005208 | chr19 | Dock8 | -0.248511046 | 0.796906162 |
| ENSMUSG0000002910 | chr5 | Htt | -0.116331931 | 0.796906162 |
| ENSMUSG0000002096 | chr12 | Ston2 | -0.120193959 | 0.796906162 |
| ENSMUSG0000002600 | chr1 | Kansl1l | -0.129468568 | 0.796906162 |
| ENSMUSG0000004039 | chr8 | Abhd13 | 0.071193556 | 0.796906162 |
| ENSMUSG0000003927 | chr11 | Foxk2 | 0.096285871 | 0.796906162 |
| ENSMUSG0000002141 | chr13 | Rpp40 | 0.133437444 | 0.796906162 |
| ENSMUSG0000000365 | chr8 | Calb2 | 0.117675237 | 0.796906162 |
| ENSMUSG0000003249 | chr9 | Mlh1 | 0.103138105 | 0.796906162 |
| ENSMUSG0000002240 | chr15 | Rbx1 | 0.125537969 | 0.796906162 |
| ENSMUSG0000004593 | chr3 | Mtmr11 | 0.116093116 | 0.796906162 |
| ENSMUSG0000003504 | chr19 | Rrp12 | 0.146982167 | 0.796906162 |
| ENSMUSG0000002886 | chr4 | Sytl1 | 0.316810339 | 0.796906162 |
| ENSMUSG0000002445 | chr17 | H2-M5 | -0.599039841 | 0.796981095 |
| ENSMUSG0000005139 | chr6 | Tacstd2 | -0.404368014 | 0.797370239 |
| ENSMUSG0000004903 | chr10 | Mterf2 | -0.142503534 | 0.797370239 |
| ENSMUSG0000004699 | chr1 | Mars2 | -0.129125113 | 0.797370239 |
| ENSMUSG0000003897 | chr11 | Ppp1r9b | -0.114277562 | 0.797370239 |
| ENSMUSG0000000287 | chr6 | Tpra1 | -0.086659464 | 0.797370239 |
| ENSMUSG0000001873 | chr11 | Pex12 | 0.105866763 | 0.797370239 |
| ENSMUSG0000001999 | chr10 | Stx7 | 0.107294648 | 0.797370239 |
| ENSMUSG0000004053 | chr5 | Abhd11 | 0.106497499 | 0.797370239 |
| ENSMUSG0000007952 | chr6 | Tmsb10 | 0.172570858 | 0.797370239 |
| ENSMUSG0000006323 | chr2 | Ptpmt1 | 0.130869536 | 0.797370239 |
| ENSMUSG0000002661 | chr1 | Esrrg | 0.338912633 | 0.797370239 |
| ENSMUSG0000003796 | chr5 | Rflna | 0.339751797 | 0.797370239 |
| ENSMUSG0000003488 | chr10 | Tbxa2r | 0.203519982 | 0.797370239 |
| ENSMUSG0000001617 | chr1 | Camk1g | 0.348004323 | 0.797370239 |
| ENSMUSG0000000303 | chr8 | Ap1m1 | 0.076648897 | 0.797708919 |
| ENSMUSG0000005759 | chr7 | Trim30d | -0.189866988 | 0.79791103 |
| ENSMUSG0000006724 | chr19 | Lgi1 | -0.248447867 | 0.797988901 |
| ENSMUSG0000002588 | chr18 | Smad7 | -0.19657054 | 0.797988901 |
| ENSMUSG0000003255 | chr9 | Nphp3 | -0.164861047 | 0.797988901 |
| ENSMUSG0000004169 | chr9 | Rasl12 | -0.216994713 | 0.797988901 |
| ENSMUSG0000003203 | chr9 | Kcnj5 | -0.160119876 | 0.797988901 |
| ENSMUSG0000003643 | chr8 | Npy1r | -0.153403865 | 0.797988901 |

| | | | | |
|---|---|---|---|---|
| ENSMUSG0000006026 | chr4 | Armh1 | -0.339815495 | 0.797988901 |
| ENSMUSG0000003030 | chr6 | Far2 | -0.188497284 | 0.797988901 |
| ENSMUSG0000003439 | chr8 | Cmip | -0.127914901 | 0.797988901 |
| ENSMUSG0000005572 | chr5 | Paqr3 | -0.190106511 | 0.797988901 |
| ENSMUSG0000002897 | chr4 | Slc2a5 | -0.604726706 | 0.797988901 |
| ENSMUSG0000005141 | chr2 | Plagl2 | -0.12610544 | 0.797988901 |
| ENSMUSG0000004791 | chr14 | Npm2 | -0.558714835 | 0.797988901 |
| ENSMUSG0000002694 | chr2 | Mamdc4 | -0.170353577 | 0.797988901 |
| ENSMUSG0000001212 | chr4 | Ubxn11 | -0.18337151 | 0.797988901 |
| ENSMUSG0000001876 | chr11 | Mpdu1 | -0.110078039 | 0.797988901 |
| ENSMUSG0000003595 | chr13 | Tdp2 | -0.113287297 | 0.797988901 |
| ENSMUSG0000003234 | chr9 | Neo1 | -0.101201489 | 0.797988901 |
| ENSMUSG0000000695 | chr16 | Chrd | -0.106909921 | 0.797988901 |
| ENSMUSG0000002212 | chr14 | Fbxl3 | -0.116117275 | 0.797988901 |
| ENSMUSG0000005213 | chr5 | Babam2 | -0.093892195 | 0.797988901 |
| ENSMUSG0000007165 | chr19 | Bscl2 | -0.073291068 | 0.797988901 |
| ENSMUSG0000001980 | chr10 | Sec63 | 0.072786645 | 0.797988901 |
| ENSMUSG0000004141 | chr13 | Pik3r1 | 0.082135391 | 0.797988901 |
| ENSMUSG0000003645 | chr19 | Hif1an | 0.076973675 | 0.797988901 |
| ENSMUSG0000006687 | chr1 | Nck2 | 0.084063133 | 0.797988901 |
| ENSMUSG0000004073 | chr5 | Eif4h | 0.089523176 | 0.797988901 |
| ENSMUSG0000005253 | chr1 | Pbx1 | 0.086291165 | 0.797988901 |
| ENSMUSG0000000427 | chr6 | Lpcat3 | 0.08295182 | 0.797988901 |
| ENSMUSG0000005732 | chr1 | Bcl2 | 0.09350776 | 0.797988901 |
| ENSMUSG0000002172 | chr13 | Parp8 | 0.095594604 | 0.797988901 |
| ENSMUSG0000003811 | chr2 | Phf20 | 0.098416315 | 0.797988901 |
| ENSMUSG0000000409 | chr9 | Cwc15 | 0.111749215 | 0.797988901 |
| ENSMUSG0000003949 | chr10 | Dse | 0.097292153 | 0.797988901 |
| ENSMUSG0000003871 | chr2 | Pbx3 | 0.104000338 | 0.797988901 |
| ENSMUSG0000000164 | chr6 | Akr1b1 | 0.116139689 | 0.797988901 |
| ENSMUSG0000003485 | chr10 | Mfsd12 | 0.117613378 | 0.797988901 |
| ENSMUSG0000006016 | chr16 | Zdhhc8 | 0.118967584 | 0.797988901 |
| ENSMUSG0000002553 | chr5 | Sumf2 | 0.137902266 | 0.797988901 |
| ENSMUSG0000003510 | chr6 | Eva1a | 0.150683403 | 0.797988901 |
| ENSMUSG0000005662 | chr19 | Fkbp2 | 0.134646537 | 0.797988901 |
| ENSMUSG0000007317 | chrX | Zfp449 | 0.114572533 | 0.797988901 |
| ENSMUSG0000003158 | chr8 | Gtf2e2 | 0.139029668 | 0.797988901 |
| ENSMUSG0000002001 | chr10 | Snrpf | 0.274816973 | 0.797988901 |
| ENSMUSG0000003420 | chr6 | Chchd4 | 0.13735577 | 0.797988901 |
| ENSMUSG0000005469 | chr3 | Emcn | 0.161417498 | 0.797988901 |
| ENSMUSG0000005464 | chr6 | Mmrn1 | 0.162655502 | 0.797988901 |
| ENSMUSG0000005670 | chr1 | Ier5 | 0.167507283 | 0.797988901 |
| ENSMUSG0000003727 | chr11 | Tmem97 | 0.192595674 | 0.797988901 |
| ENSMUSG0000004046 | chr7 | Blvrb | 0.17417645 | 0.797988901 |
| ENSMUSG0000000068 | chr11 | Abhd15 | -0.467714134 | 0.798202018 |
| ENSMUSG0000008998 | chr5 | Fbxo24 | -0.418696386 | 0.798202018 |
| ENSMUSG0000002497 | chr19 | Lgals12 | -0.988812536 | 0.798717539 |
| ENSMUSG0000001436 | chr2 | Mertk | -0.252492861 | 0.798717539 |
| ENSMUSG0000005227 | chr7 | Dnah3 | -0.484641002 | 0.798717539 |
| ENSMUSG0000006170 | chr7 | Tmem91 | -0.462814178 | 0.798717539 |

| | | | | |
|---|---|---|---|---|
| ENSMUSG0000004456 | chr13 | Cage1 | -0.219124316 | 0.798717539 |
| ENSMUSG0000002464 | chr19 | Psat1 | -0.143706594 | 0.798717539 |
| ENSMUSG0000003630 | chr8 | Lzts1 | -0.203313577 | 0.798717539 |
| ENSMUSG0000006412 | chr14 | Cenpj | -0.123418272 | 0.798717539 |
| ENSMUSG0000003102 | chr7 | Tmem9b | -0.098033265 | 0.798717539 |
| ENSMUSG0000005943 | chr12 | Max | -0.089042498 | 0.798717539 |
| ENSMUSG0000004479 | chr11 | Cyb5d1 | 0.092968584 | 0.798717539 |
| ENSMUSG0000002203 | chr14 | Ccdc25 | 0.112980827 | 0.798717539 |
| ENSMUSG0000004237 | chr13 | Esm1 | 0.415456258 | 0.798717539 |
| ENSMUSG0000002637 | chr1 | Nifk | 0.125150832 | 0.798717539 |
| ENSMUSG0000003518 | chr12 | Heatr5a | 0.12520682 | 0.798717539 |
| ENSMUSG0000003314 | chr3 | Slc22a15 | 0.125647941 | 0.798717539 |
| ENSMUSG0000004112 | chr11 | H2az2 | 0.145681042 | 0.798717539 |
| ENSMUSG0000003903 | chr3 | St6galnac5 | 0.23925692 | 0.798717539 |
| ENSMUSG0000006117 | chr3 | Fnip2 | 0.193210733 | 0.798717539 |
| ENSMUSG0000003916 | chr3 | Adgrl4 | 0.18869286 | 0.798717539 |
| ENSMUSG0000003008 | chr6 | Chchd6 | 0.185474186 | 0.798717539 |
| ENSMUSG0000007865 | chr11 | Vps25 | 0.369253066 | 0.798717539 |
| ENSMUSG0000004453 | chr9 | Ackr2 | 0.187893855 | 0.798717539 |
| ENSMUSG0000005502 | chr6 | Smyd1 | 0.422146976 | 0.798717539 |
| ENSMUSG0000002284 | chr16 | Slc49a4 | 0.075725632 | 0.799260804 |
| ENSMUSG0000002018 | chr10 | Nav3 | -0.163899624 | 0.799338949 |
| ENSMUSG0000006992 | chr8 | B3gnt9 | -0.179359611 | 0.799338949 |
| ENSMUSG0000007473 | chr4 | Fndc10 | -0.148873716 | 0.799338949 |
| ENSMUSG0000002788 | chr3 | Sypl2 | -0.165994179 | 0.799338949 |
| ENSMUSG0000001424 | chr11 | Zswim7 | -0.169527349 | 0.799338949 |
| ENSMUSG0000003830 | chr6 | Snx10 | -0.090708133 | 0.799338949 |
| ENSMUSG0000002867 | chr4 | Pnrc2 | 0.095866778 | 0.799338949 |
| ENSMUSG0000000228 | chr17 | Ciao3 | 0.089637741 | 0.799338949 |
| ENSMUSG0000007107 | chr17 | Yipf3 | 0.109788013 | 0.799338949 |
| ENSMUSG0000002793 | chr3 | Jtb | 0.113792935 | 0.799338949 |
| ENSMUSG0000007035 | chr11 | Evi2 | 0.509866853 | 0.799338949 |
| ENSMUSG0000001908 | chr15 | Mfsd3 | 0.210015993 | 0.799338949 |
| ENSMUSG0000000490 | chr6 | Slc25a18 | 0.370692394 | 0.799338949 |
| ENSMUSG0000006385 | chr9 | Gpx1 | 0.214388213 | 0.799338949 |
| ENSMUSG0000003830 | chr3 | Cd160 | 0.471653057 | 0.799338949 |
| ENSMUSG0000003600 | chr13 | Ripor2 | 0.21782638 | 0.799338949 |
| ENSMUSG0000004264 | chr3 | Flad1 | -0.101669221 | 0.799372377 |
| ENSMUSG0000001569 | chr3 | Setdb1 | -0.069324085 | 0.799372377 |
| ENSMUSG0000006825 | chr6 | Amn1 | -0.10643303 | 0.799443853 |
| ENSMUSG0000003415 | chr13 | Exoc3 | -0.07927872 | 0.799443853 |
| ENSMUSG0000003850 | chr7 | Ptov1 | 0.085798203 | 0.799443853 |
| ENSMUSG0000002014 | chr11 | Rab1a | 0.093473489 | 0.799443853 |
| ENSMUSG0000004434 | chr19 | Marveld1 | 0.108175427 | 0.799443853 |
| ENSMUSG0000003197 | chr8 | Urb2 | 0.114727799 | 0.799443853 |
| ENSMUSG0000002962 | chr5 | Ptcd1 | 0.121228589 | 0.799443853 |
| ENSMUSG0000004328 | chr11 | Tmem11 | 0.135021385 | 0.799443853 |
| ENSMUSG0000010242 | chr18 | Pcdhga12 | 0.182745976 | 0.799443853 |
| ENSMUSG0000004016 | chr2 | Kcns1 | 0.217473792 | 0.799443853 |
| ENSMUSG0000003231 | chr9 | Nrg4 | 0.379084356 | 0.799443853 |

| | | | | |
|---|---|---|---|---|
| ENSMUSG0000002113 | chr12 | Galnt16 | 0.154560587 | 0.799443853 |
| ENSMUSG0000002203 | chr14 | Scara5 | 0.23189874 | 0.799443853 |
| ENSMUSG0000005920 | chr6 | Lep | 0.755080295 | 0.799518178 |
| ENSMUSG0000003611 | chr7 | Lmo1 | 0.267366133 | 0.799518178 |
| ENSMUSG0000003404 | chr8 | Lyl1 | 0.205207711 | 0.799518178 |
| ENSMUSG0000003955 | chr10 | Rsph4a | -0.20055324 | 0.7995264 |
| ENSMUSG0000004027 | chr4 | Bach2 | -0.177131419 | 0.799648534 |
| ENSMUSG0000002120 | chr12 | Otub2 | 0.104091741 | 0.799648534 |
| ENSMUSG0000003109 | chrX | Cul4b | 0.09630174 | 0.799648534 |
| ENSMUSG0000002827 | chr3 | Gtf2b | 0.111205211 | 0.799648534 |
| ENSMUSG0000004089 | chr3 | Kcnd3 | 0.112636943 | 0.799648534 |
| ENSMUSG0000002863 | chr10 | Dyrk2 | 0.104796735 | 0.799743277 |
| ENSMUSG0000004528 | chr15 | Gpr20 | 0.138358878 | 0.799743277 |
| ENSMUSG0000003330 | chr3 | Dpyd | -0.137731443 | 0.799901334 |
| ENSMUSG0000002411 | chr17 | Tedc2 | -0.132844802 | 0.799901334 |
| ENSMUSG0000002908 | chr5 | Bst1 | 0.321361637 | 0.799901334 |
| ENSMUSG0000002466 | chr19 | Incenp | -0.230561903 | 0.799998586 |
| ENSMUSG0000000816 | chr8 | Fbxw9 | -0.162602794 | 0.799998586 |
| ENSMUSG0000002069 | chr11 | Cct6b | -0.532902984 | 0.799998586 |
| ENSMUSG0000004024 | chr19 | Tbc1d10c | -0.582364595 | 0.799998586 |
| ENSMUSG0000002724 | chr2 | Pdia3 | 0.147729925 | 0.799998586 |
| ENSMUSG0000005944 | chr5 | Hadhb | 0.110003629 | 0.799998586 |
| ENSMUSG0000002708 | chr2 | Phospho2 | 0.139834025 | 0.799998586 |
| ENSMUSG0000000422 | chrX | Ikbkg | -0.077451746 | 0.800134269 |
| ENSMUSG0000003934 | chr6 | Atp6v0e2 | 0.068095504 | 0.800160605 |
| ENSMUSG0000004755 | chr3 | Lxn | 0.132091827 | 0.800311054 |
| ENSMUSG0000004722 | chr12 | Gm527 | 0.160295416 | 0.800311054 |
| ENSMUSG0000002243 | chr15 | Parvg | 0.26130419 | 0.800311054 |
| ENSMUSG0000003259 | chr9 | Amigo3 | -0.17812404 | 0.800610401 |
| ENSMUSG0000003021 | chr6 | Atf7ip | 0.093001427 | 0.800637923 |
| ENSMUSG0000001776 | chr2 | Ctsa | 0.098753187 | 0.800637923 |
| ENSMUSG0000007463 | chr13 | Tmem267 | -0.281611154 | 0.800728982 |
| ENSMUSG0000003053 | chr7 | Vps33b | -0.117044734 | 0.800728982 |
| ENSMUSG0000003285 | chr5 | Rnft2 | -0.202229642 | 0.80079941 |
| ENSMUSG0000003155 | chr8 | Plekha2 | -0.120560954 | 0.80079941 |
| ENSMUSG0000002695 | chr2 | Dpp7 | -0.134199538 | 0.80097126 |
| ENSMUSG0000005085 | chr4 | Tmem125 | -0.628870351 | 0.801477833 |
| ENSMUSG0000002148 | chr13 | Nsd1 | -0.120018205 | 0.801524208 |
| ENSMUSG0000002862 | chr4 | Exo5 | 0.113006513 | 0.801724374 |
| ENSMUSG0000000580 | chr2 | Sqor | -0.127166076 | 0.801988418 |
| ENSMUSG0000003890 | chr3 | Pogz | -0.115580274 | 0.802044547 |
| ENSMUSG0000005327 | chr19 | Aldh1a1 | 0.172245342 | 0.802280251 |
| ENSMUSG0000002648 | chr1 | Ncf2 | 0.169431499 | 0.802316085 |
| ENSMUSG0000003923 | chr11 | Zfp750 | -0.322303866 | 0.802600967 |
| ENSMUSG0000002621 | chr1 | Stk11ip | -0.139861231 | 0.802600967 |
| ENSMUSG0000000910 | chr3 | Gnat2 | 0.586909127 | 0.802600967 |
| ENSMUSG0000006641 | chr9 | Msl2 | -0.084443575 | 0.802716397 |
| ENSMUSG0000003069 | chr7 | Tbx6 | 0.269792056 | 0.802816966 |
| ENSMUSG0000002540 | chr10 | Mbd6 | -0.18019194 | 0.802907066 |
| ENSMUSG0000003402 | chr6 | Fancd2 | 0.379495562 | 0.802907066 |

| | | | | |
|---|---|---|---|---|
| ENSMUSG0000002816 | chr3 | Manba | 0.090560817 | 0.8030471 |
| ENSMUSG0000002942 | chr5 | Stx2 | -0.079406799 | 0.803730767 |
| ENSMUSG0000002578 | chr2 | Taf3 | 0.105338614 | 0.803730767 |
| ENSMUSG0000002515 | chr19 | Arhgap19 | -0.219943759 | 0.803852841 |
| ENSMUSG0000000289 | chr6 | Il17ra | -0.200096373 | 0.803852841 |
| ENSMUSG0000002175 | chr14 | Kctd6 | 0.100493314 | 0.803852841 |
| ENSMUSG0000002912 | chr5 | Rab28 | 0.133915543 | 0.803852841 |
| ENSMUSG0000002276 | chr16 | Sdf2l1 | 0.287444736 | 0.803852841 |
| ENSMUSG0000002619 | chr1 | Bard1 | -0.252961972 | 0.80398001 |
| ENSMUSG0000003961 | chr4 | Pgap4 | -0.135400055 | 0.80398001 |
| ENSMUSG0000002482 | chr19 | Gldc | 0.252173803 | 0.80398001 |
| ENSMUSG0000002521 | chr19 | Kazald1 | -0.197172549 | 0.804029544 |
| ENSMUSG0000003245 | chr9 | Copb2 | 0.07820669 | 0.804029544 |
| ENSMUSG0000004673 | chr11 | Kctd11 | 0.125203772 | 0.804121468 |
| ENSMUSG0000002179 | chr14 | Prxl2a | 0.235600917 | 0.80418241 |
| ENSMUSG0000001892 | chr11 | Cxcl16 | -0.18863745 | 0.804191198 |
| ENSMUSG0000003012 | chr6 | Lrrc23 | -0.254142926 | 0.804191198 |
| ENSMUSG0000001675 | chr13 | Cmah | -0.153630505 | 0.804191198 |
| ENSMUSG0000004168 | chrX | Amot | -0.126743411 | 0.804191198 |
| ENSMUSG0000002906 | chr4 | Mib2 | -0.073974138 | 0.804191198 |
| ENSMUSG0000003987 | chr10 | Heca | 0.067800901 | 0.804191198 |
| ENSMUSG0000004667 | chr6 | C87436 | 0.125017072 | 0.804191198 |
| ENSMUSG0000002643 | chr1 | Slc45a3 | 0.195323508 | 0.804191198 |
| ENSMUSG0000002981 | chr6 | Tmem176b | -0.109338815 | 0.804537465 |
| ENSMUSG0000000154 | chr13 | Ell2 | 0.151900574 | 0.804537465 |
| ENSMUSG0000002798 | chr3 | Lef1 | 0.437867291 | 0.804537465 |
| ENSMUSG0000006874 | chr3 | Psrc1 | 0.335835829 | 0.804537465 |
| ENSMUSG0000002303 | chr15 | Slc4a8 | -0.119315152 | 0.804539848 |
| ENSMUSG0000002546 | chr7 | Echs1 | 0.094050112 | 0.804574355 |
| ENSMUSG0000000058 | chr11 | C1d | 0.126100324 | 0.804587379 |
| ENSMUSG0000001986 | chr10 | Vta1 | 0.113179621 | 0.804835239 |
| ENSMUSG0000005737 | chr2 | Ryr3 | 0.183220409 | 0.804835239 |
| ENSMUSG0000006077 | chr1 | Tsga10 | -0.118275086 | 0.80492926 |
| ENSMUSG0000002039 | chr11 | Nefh | -0.186797661 | 0.805057926 |
| ENSMUSG0000002741 | chr2 | Lpin3 | -0.251550639 | 0.805057926 |
| ENSMUSG0000001741 | chr2 | Cacnb4 | -0.160782511 | 0.805057926 |
| ENSMUSG0000002334 | chr6 | Clec4n | -0.339669345 | 0.805057926 |
| ENSMUSG0000003495 | chr10 | Atcay | -0.146265163 | 0.805057926 |
| ENSMUSG0000002027 | chr11 | Rel | -0.167677544 | 0.805057926 |
| ENSMUSG0000003954 | chr9 | Ncam1 | -0.131765236 | 0.805057926 |
| ENSMUSG0000011728 | chr5 | Gm1043 | -0.159712946 | 0.805057926 |
| ENSMUSG0000002084 | chr11 | Cpd | -0.133549073 | 0.805057926 |
| ENSMUSG0000008714 | chr16 | Plcxd2 | -0.167746736 | 0.805057926 |
| ENSMUSG0000006031 | chr7 | Zfp941 | -0.153645243 | 0.805057926 |
| ENSMUSG0000003523 | chr8 | Lcat | -0.218644268 | 0.805057926 |
| ENSMUSG0000002824 | chr4 | Pnisr | -0.121080815 | 0.805057926 |
| ENSMUSG0000002486 | chr19 | Klc2 | -0.116701533 | 0.805057926 |
| ENSMUSG0000002044 | chr6 | Arf5 | -0.109258159 | 0.805057926 |
| ENSMUSG0000003857 | chr7 | Saxo2 | -0.181749339 | 0.805057926 |
| ENSMUSG0000003669 | chr15 | Ago2 | -0.118271651 | 0.805057926 |

| | | | | |
|---|---|---|---|---|
| ENSMUSG0000002031 | chr11 | Wdpcp | -0.124491887 | 0.805057926 |
| ENSMUSG0000002950 | chr5 | Ep400 | -0.106973267 | 0.805057926 |
| ENSMUSG0000004197 | chr16 | Spidr | -0.116233792 | 0.805057926 |
| ENSMUSG0000003359 | chr8 | Spata2l | -0.14180812 | 0.805057926 |
| ENSMUSG0000006001 | chr14 | Kif13b | -0.09882779 | 0.805057926 |
| ENSMUSG0000002100 | chr12 | Galc | -0.131802145 | 0.805057926 |
| ENSMUSG0000002349 | chr9 | Pcbp4 | -0.103532413 | 0.805057926 |
| ENSMUSG0000005819 | chr9 | Zfp846 | -0.092262643 | 0.805057926 |
| ENSMUSG0000003137 | chrX | Zrsr2 | -0.080885147 | 0.805057926 |
| ENSMUSG0000001187 | chr11 | Git1 | -0.08304993 | 0.805057926 |
| ENSMUSG0000000037 | chr10 | Trappc10 | -0.086259143 | 0.805057926 |
| ENSMUSG0000002732 | chr2 | Adissp | -0.086977369 | 0.805057926 |
| ENSMUSG0000002128 | chr12 | Eif5 | 0.072023352 | 0.805057926 |
| ENSMUSG0000004081 | chr14 | Dennd6a | 0.075788705 | 0.805057926 |
| ENSMUSG0000000873 | chr7 | Gprc5b | 0.087762002 | 0.805057926 |
| ENSMUSG0000003052 | chr7 | Mphosph10 | 0.089828667 | 0.805057926 |
| ENSMUSG0000001921 | chr6 | Atp6v1e1 | 0.099302744 | 0.805057926 |
| ENSMUSG0000002925 | chr5 | Polr2b | 0.089735462 | 0.805057926 |
| ENSMUSG0000006391 | chr5 | Srrm4 | 0.094556986 | 0.805057926 |
| ENSMUSG0000004129 | chr5 | Katnal1 | 0.08747711 | 0.805057926 |
| ENSMUSG0000002492 | chr19 | Rela | 0.088007064 | 0.805057926 |
| ENSMUSG0000000439 | chr11 | Ddx56 | 0.100738968 | 0.805057926 |
| ENSMUSG0000004162 | chr14 | Ggact | 0.104230377 | 0.805057926 |
| ENSMUSG0000004422 | chr5 | Grsf1 | 0.104546487 | 0.805057926 |
| ENSMUSG0000003819 | chr11 | Rilp | 0.164930439 | 0.805057926 |
| ENSMUSG0000003243 | chr9 | Crtap | 0.122179086 | 0.805057926 |
| ENSMUSG0000002847 | chr4 | Clta | 0.108027628 | 0.805057926 |
| ENSMUSG0000003560 | chr4 | Trmt10b | 0.107261766 | 0.805057926 |
| ENSMUSG0000003228 | chr9 | Dnaja4 | 0.113181914 | 0.805057926 |
| ENSMUSG0000002305 | chr15 | Calcoco1 | 0.140805128 | 0.805057926 |
| ENSMUSG0000003836 | chr4 | Focad | 0.119795811 | 0.805057926 |
| ENSMUSG0000007474 | chr2 | Thbd | 0.173201357 | 0.805057926 |
| ENSMUSG0000005396 | chr3 | Pde5a | 0.14422425 | 0.805057926 |
| ENSMUSG0000008979 | chr5 | 1700028K03Rik | 0.335966441 | 0.805057926 |
| ENSMUSG0000006009 | chr13 | H4c1 | 0.559370877 | 0.805057926 |
| ENSMUSG0000003900 | chr4 | Tlr4 | 0.172277989 | 0.805057926 |
| ENSMUSG0000007489 | chr19 | Ifit3 | -0.62635552 | 0.8050715 |
| ENSMUSG0000003310 | chr13 | Cdc14b | 0.135875893 | 0.8050715 |
| ENSMUSG0000003848 | chr10 | Cdk19 | -0.160194639 | 0.805178567 |
| ENSMUSG0000003054 | chr7 | Plin1 | 0.603424903 | 0.805178567 |
| ENSMUSG0000002513 | chr7 | Ints4 | 0.066674579 | 0.805178567 |
| ENSMUSG0000003989 | chr5 | Fgl2 | 0.128010768 | 0.805178567 |
| ENSMUSG0000003039 | chr7 | Zik1 | 0.158399931 | 0.805178567 |
| ENSMUSG0000002422 | chr17 | Nudt12 | -0.147804238 | 0.805227376 |
| ENSMUSG0000004123 | chr18 | Rbbp8 | 0.156788176 | 0.805227376 |
| ENSMUSG0000001052 | chr12 | Gm266 | -0.463229756 | 0.805448554 |
| ENSMUSG0000005316 | chr1 | Daw1 | -0.446223462 | 0.805448554 |
| ENSMUSG0000002960 | chr5 | Rph3a | -0.162132675 | 0.805497181 |
| ENSMUSG0000001585 | chr3 | Fcrl2 | -0.165149574 | 0.805521748 |
| ENSMUSG0000003212 | chr9 | C2cd2l | -0.105392012 | 0.805521748 |

| | | | | |
|---|---|---|---|---|
| ENSMUSG0000005060 | chr2 | Zfp831 | -0.694307925 | 0.805855828 |
| ENSMUSG0000004893 | chr6 | Tada3 | 0.12230573 | 0.805855828 |
| ENSMUSG0000002413 | chr17 | Srbd1 | -0.143001849 | 0.80585838 |
| ENSMUSG0000005487 | chr19 | Pcnx3 | -0.099433215 | 0.80585838 |
| ENSMUSG0000003874 | chr2 | Mvb12b | 0.068005248 | 0.80585838 |
| ENSMUSG0000005857 | chr14 | Gpc6 | -0.167376664 | 0.805867951 |
| ENSMUSG0000003187 | chr8 | Bean1 | -0.156921254 | 0.805895288 |
| ENSMUSG0000006639 | chr12 | Nrxn3 | 0.146378207 | 0.805926537 |
| ENSMUSG0000003513 | chr13 | Secisbp2 | 0.09632567 | 0.805941459 |
| ENSMUSG0000001366 | chr19 | Atad1 | 0.072196684 | 0.805986724 |
| ENSMUSG0000002396 | chr17 | Fbxl17 | -0.084274516 | 0.806055253 |
| ENSMUSG0000003070 | chr7 | Rab6a | -0.07520528 | 0.806055253 |
| ENSMUSG0000001819 | chr1 | Ro60 | 0.08969903 | 0.806055253 |
| ENSMUSG0000003261 | chr11 | Nt5m | 0.132058377 | 0.806055253 |
| ENSMUSG0000003445 | chr9 | Slc24a1 | 0.482261711 | 0.806055253 |
| ENSMUSG0000003488 | chr8 | Mrpl34 | 0.493032799 | 0.806145147 |
| ENSMUSG0000001774 | chr2 | Slc12a5 | -0.27949513 | 0.806491221 |
| ENSMUSG0000002195 | chr14 | Lats2 | 0.067457649 | 0.806739166 |
| ENSMUSG0000002038 | chr11 | Jade2 | -0.194121951 | 0.806838417 |
| ENSMUSG0000002943 | chr5 | Bcl7a | -0.170945074 | 0.806838417 |
| ENSMUSG0000004633 | chr2 | Gpat2 | -0.39569161 | 0.806838417 |
| ENSMUSG0000010374 | chr18 | Pcdhgb5 | -0.226775841 | 0.806838417 |
| ENSMUSG0000000198 | chr4 | Grik3 | -0.131268062 | 0.806838417 |
| ENSMUSG0000004132 | chr13 | Inhba | -0.260085163 | 0.806838417 |
| ENSMUSG0000004324 | chr8 | Niban3 | -0.261756536 | 0.806838417 |
| ENSMUSG0000001836 | chr11 | Kpna2 | -0.155820688 | 0.806838417 |
| ENSMUSG0000002401 | chr17 | Ccdc167 | -0.155491958 | 0.806838417 |
| ENSMUSG0000002922 | chr5 | Uchl1 | -0.116535073 | 0.806838417 |
| ENSMUSG0000002858 | chr4 | Cc2d1b | -0.097919063 | 0.806838417 |
| ENSMUSG0000001956 | chr10 | Arid3a | -0.121054149 | 0.806838417 |
| ENSMUSG0000002760 | chr2 | Acss2 | -0.097221643 | 0.806838417 |
| ENSMUSG0000002147 | chr13 | Sfxn1 | -0.08134012 | 0.806838417 |
| ENSMUSG0000003333 | chr9 | Dnm2 | -0.083607037 | 0.806838417 |
| ENSMUSG0000000667 | chr9 | Qrich1 | 0.061004637 | 0.806838417 |
| ENSMUSG0000000459 | chr3 | Pkn2 | 0.06661598 | 0.806838417 |
| ENSMUSG0000001987 | chr10 | Serinc1 | 0.085012228 | 0.806838417 |
| ENSMUSG0000003531 | chr10 | Gnptab | 0.082469637 | 0.806838417 |
| ENSMUSG0000000407 | chr16 | Hmox2 | 0.086414463 | 0.806838417 |
| ENSMUSG0000002615 | chr1 | Mff | 0.093633297 | 0.806838417 |
| ENSMUSG0000002276 | chr16 | Snap29 | 0.099535681 | 0.806838417 |
| ENSMUSG0000004543 | chr5 | Cox19 | 0.090469198 | 0.806838417 |
| ENSMUSG0000003628 | chr5 | Noa1 | 0.107575211 | 0.806838417 |
| ENSMUSG0000003006 | chr6 | Rpn1 | 0.144019848 | 0.806838417 |
| ENSMUSG0000006410 | chr19 | Cnnm2 | 0.132072814 | 0.806838417 |
| ENSMUSG0000002251 | chr16 | Il1rap | 0.116143713 | 0.806838417 |
| ENSMUSG0000003629 | chr18 | BC031181 | 0.141868376 | 0.806838417 |
| ENSMUSG0000002581 | chr2 | Nudt5 | 0.127786361 | 0.806838417 |
| ENSMUSG0000002909 | chr5 | Trmt44 | 0.153623824 | 0.806838417 |
| ENSMUSG0000004394 | chr9 | Naalad2 | 0.138581161 | 0.806838417 |
| ENSMUSG0000003393 | chr6 | Vhl | 0.134779359 | 0.806838417 |

| | | | | |
|---|---|---|---|---|
| ENSMUSG0000004081 | chr7 | Eml2 | 0.13243195 | 0.806838417 |
| ENSMUSG0000002063 | chr12 | Rnaseh1 | 0.162004626 | 0.806838417 |
| ENSMUSG0000002634 | chr1 | Gpr39 | -0.474395145 | 0.806916567 |
| ENSMUSG0000000435 | chr10 | Utp20 | -0.113590797 | 0.806916567 |
| ENSMUSG0000005013 | chr11 | Sarm1 | -0.134604473 | 0.806916567 |
| ENSMUSG0000002429 | chr17 | Zfp871 | -0.103302424 | 0.806916567 |
| ENSMUSG0000001512 | chr17 | Ube2i | 0.059135449 | 0.806916567 |
| ENSMUSG0000004072 | chr3 | Kcna2 | -0.169524659 | 0.806968251 |
| ENSMUSG0000001455 | chr6 | Rbsn | 0.068402362 | 0.807174053 |
| ENSMUSG0000007877 | chr4 | Rad54b | 0.46459178 | 0.807174053 |
| ENSMUSG0000002580 | chr8 | Ccdc7a | -0.606520533 | 0.807254451 |
| ENSMUSG0000000565 | chr12 | Snx6 | 0.100503864 | 0.807354394 |
| ENSMUSG0000004090 | chr7 | Atp1a3 | -0.166786583 | 0.807656649 |
| ENSMUSG0000011714 | chr17 | Gng5c | -0.414893915 | 0.807656649 |
| ENSMUSG0000007282 | chr12 | Cep170b | -0.097061185 | 0.807656649 |
| ENSMUSG0000004444 | chr16 | N6amt1 | 0.096852152 | 0.807656649 |
| ENSMUSG0000002164 | chr13 | Mccc2 | 0.087586536 | 0.807656649 |
| ENSMUSG0000002971 | chr5 | Gnb2 | 0.107498793 | 0.807656649 |
| ENSMUSG0000000355 | chr19 | As3mt | 0.111070089 | 0.807656649 |
| ENSMUSG0000003805 | chr6 | Nod1 | 0.105339248 | 0.807792995 |
| ENSMUSG0000002546 | chr7 | Paox | 0.137599996 | 0.807847554 |
| ENSMUSG0000003762 | chr14 | Cdkn3 | -0.371061218 | 0.807948355 |
| ENSMUSG0000003258 | chr9 | Traip | -0.314592386 | 0.807948355 |
| ENSMUSG0000000210 | chr2 | Ddb2 | -0.166911995 | 0.807948355 |
| ENSMUSG0000003964 | chr16 | Vasn | -0.131911349 | 0.807948355 |
| ENSMUSG0000002308 | chr3 | Noct | -0.209120773 | 0.807948355 |
| ENSMUSG0000003146 | chr8 | Agpat5 | -0.07734583 | 0.807948355 |
| ENSMUSG0000000199 | chr2 | Ap4e1 | 0.166729588 | 0.807948355 |
| ENSMUSG0000000315 | chr6 | Slc2a3 | -0.141854182 | 0.80813743 |
| ENSMUSG0000004640 | chr1 | Yod1 | -0.125161093 | 0.80813743 |
| ENSMUSG0000003381 | chr1 | Tcea1 | 0.077254716 | 0.80813743 |
| ENSMUSG0000002913 | chr5 | Dnajb6 | 0.096666387 | 0.80813743 |
| ENSMUSG0000003148 | chr8 | Plpbp | 0.091923976 | 0.80813743 |
| ENSMUSG0000000895 | chr3 | Vps72 | 0.099380168 | 0.80813743 |
| ENSMUSG0000002537 | chr10 | Nabp2 | 0.109029114 | 0.80813743 |
| ENSMUSG0000004877 | chr6 | Pthlh | 0.185746891 | 0.80813743 |
| ENSMUSG0000000165 | chr15 | Hoxc8 | 0.31496779 | 0.80813743 |
| ENSMUSG0000002658 | chr1 | Sec16b | -0.306899463 | 0.808257389 |
| ENSMUSG0000002695 | chr2 | Grin1 | -0.168328334 | 0.808257389 |
| ENSMUSG0000003785 | chr11 | Nufip2 | -0.132454025 | 0.808257389 |
| ENSMUSG0000004641 | chr7 | Kcnk6 | -0.230010687 | 0.808257389 |
| ENSMUSG0000001771 | chr11 | Afmid | -0.189822933 | 0.808257389 |
| ENSMUSG0000002211 | chr14 | Rbm26 | -0.102316676 | 0.808257389 |
| ENSMUSG0000004001 | chr19 | Cox15 | -0.080833821 | 0.808257389 |
| ENSMUSG0000001857 | chr11 | Phf23 | 0.080493639 | 0.808257389 |
| ENSMUSG0000002833 | chr4 | Ncbp1 | 0.084251513 | 0.808257389 |
| ENSMUSG0000004755 | chr7 | Tmem41b | 0.080125488 | 0.808257389 |
| ENSMUSG0000002656 | chr1 | Tada1 | 0.113751945 | 0.808257389 |
| ENSMUSG0000004710 | chr15 | Dnajb7 | 0.612656678 | 0.808257389 |
| ENSMUSG0000009366 | chr13 | Pou5f2 | 0.588449553 | 0.808257389 |

| | | | | |
|---|---|---|---|---|
| ENSMUSG0000009023 | chr16 | Car15 | -0.320153647 | 0.80841328 |
| ENSMUSG0000002123 | chr12 | Aldh6a1 | -0.120863003 | 0.808756208 |
| ENSMUSG0000011040 | chr4 | Rnf223 | -0.601673039 | 0.808756208 |
| ENSMUSG0000005390 | chr6 | Mat2a | -0.085067294 | 0.808756208 |
| ENSMUSG0000002434 | chr18 | Sting1 | -0.092014307 | 0.808756208 |
| ENSMUSG0000006301 | chr5 | Ccni | 0.082157629 | 0.808756208 |
| ENSMUSG0000001500 | chr15 | Efr3a | -0.072289774 | 0.808781018 |
| ENSMUSG0000001653 | chrX | Lamp2 | 0.078653822 | 0.808781018 |
| ENSMUSG0000002778 | chr3 | Kpna4 | 0.100376221 | 0.808781018 |
| ENSMUSG0000004587 | chr18 | Pcdhb8 | -0.21658884 | 0.808903796 |
| ENSMUSG0000007965 | chr17 | Rab26 | -0.386969663 | 0.808923156 |
| ENSMUSG0000002678 | chr2 | Pkn3 | -0.207947838 | 0.809019689 |
| ENSMUSG0000003203 | chr9 | Cul5 | 0.074615516 | 0.809019689 |
| ENSMUSG0000001546 | chr17 | Atf6b | 0.095299587 | 0.809019689 |
| ENSMUSG0000003560 | chr9 | Ky | 0.379951161 | 0.809210197 |
| ENSMUSG0000000334 | chr10 | Csnk1g2 | 0.092059567 | 0.809210197 |
| ENSMUSG0000004102 | chr14 | Ghitm | -0.074829895 | 0.809354111 |
| ENSMUSG0000002322 | chr2 | Serping1 | 0.130608074 | 0.809354111 |
| ENSMUSG0000003188 | chr8 | Cbfb | -0.097247237 | 0.809581895 |
| ENSMUSG0000004009 | chr2 | Bmf | 0.133357306 | 0.809621543 |
| ENSMUSG0000005816 | chr11 | Gm5431 | -0.339118769 | 0.80987742 |
| ENSMUSG0000002275 | chr16 | Lrrc74b | -0.34673185 | 0.809934382 |
| ENSMUSG0000002404 | chr17 | Akap8 | -0.114434929 | 0.809934382 |
| ENSMUSG0000002644 | chr1 | Pik3c2b | -0.095596439 | 0.809934382 |
| ENSMUSG0000004042 | chr5 | Mterf1a | 0.187534849 | 0.81003993 |
| ENSMUSG0000003438 | chr17 | AI661453 | -0.22166851 | 0.81017418 |
| ENSMUSG0000003005 | chr6 | Rab43 | -0.277496303 | 0.81017418 |
| ENSMUSG0000004510 | chr2 | Ccdc73 | -0.176667392 | 0.81017418 |
| ENSMUSG0000003653 | chr8 | Slc38a7 | -0.099571389 | 0.81017418 |
| ENSMUSG0000004299 | chr3 | Nhlrc3 | -0.10316594 | 0.81017418 |
| ENSMUSG0000005347 | chr6 | Kdm3a | 0.093144899 | 0.81017418 |
| ENSMUSG0000006670 | chr9 | Fxyd6 | 0.122874348 | 0.81017418 |
| ENSMUSG0000002084 | chr11 | Timm22 | 0.133514324 | 0.81017418 |
| ENSMUSG0000003959 | chr14 | Fam149b | 0.123124357 | 0.81017418 |
| ENSMUSG0000011413 | chr13 | Btf3l4b | 0.625307611 | 0.81017418 |
| ENSMUSG0000002200 | chr14 | Nufip1 | 0.111085405 | 0.810408787 |
| ENSMUSG0000006103 | chr9 | Or8b53 | -0.292706955 | 0.810487283 |
| ENSMUSG0000000546 | chr8 | Il27ra | -0.432718451 | 0.810536741 |
| ENSMUSG0000003974 | chr4 | Alg2 | -0.072640245 | 0.810536741 |
| ENSMUSG0000006949 | chr2 | Epc2 | 0.084747275 | 0.810536741 |
| ENSMUSG0000001985 | chr10 | Asf1a | 0.132024264 | 0.810536741 |
| ENSMUSG0000007155 | chr16 | Cfap44 | -0.232429374 | 0.810613701 |
| ENSMUSG0000002002 | chr10 | Tmcc3 | -0.16841624 | 0.810613701 |
| ENSMUSG0000003165 | chr8 | N4bp1 | -0.076589979 | 0.810613701 |
| ENSMUSG0000001552 | chr3 | Arnt | 0.081424265 | 0.810613701 |
| ENSMUSG0000003083 | chr7 | Nomo1 | 0.09289266 | 0.810613701 |
| ENSMUSG0000002895 | chr5 | Tmub1 | -0.117393917 | 0.810641846 |
| ENSMUSG0000003034 | chr6 | Scnn1a | -0.235921759 | 0.810676209 |
| ENSMUSG0000006286 | chr10 | Phactr2 | 0.086066396 | 0.810705636 |
| ENSMUSG0000002281 | chr16 | Gsk3b | -0.111781755 | 0.81095518 |

| | | | | |
|---|---|---|---|---|
| ENSMUSG0000004339 | chr12 | Gpr135 | -0.177391294 | 0.810980888 |
| ENSMUSG0000002961 | chr5 | Eif2ak1 | -0.070299827 | 0.811016867 |
| ENSMUSG0000005432 | chr8 | Lrrc36 | -0.557463426 | 0.811029193 |
| ENSMUSG0000005784 | chr13 | Zfp595 | -0.149148281 | 0.811029193 |
| ENSMUSG0000003822 | chr8 | Primpol | -0.164227381 | 0.811029193 |
| ENSMUSG0000009033 | chr8 | Cfap97d2 | -0.170314003 | 0.811029193 |
| ENSMUSG0000003208 | chr9 | Bace1 | -0.094604834 | 0.811029193 |
| ENSMUSG0000003248 | chr9 | Scap | -0.08531178 | 0.811029193 |
| ENSMUSG0000004736 | chr19 | Abhd17b | -0.099694031 | 0.811029193 |
| ENSMUSG0000001711 | chr11 | Nbr1 | -0.06883793 | 0.811029193 |
| ENSMUSG0000002013 | chr10 | Rab21 | 0.064038246 | 0.811029193 |
| ENSMUSG0000006060 | chr7 | Nr1h2 | 0.08567208 | 0.811029193 |
| ENSMUSG0000002877 | chr5 | Ptpn12 | 0.086454876 | 0.811029193 |
| ENSMUSG0000001423 | chr16 | Cluap1 | 0.135915711 | 0.811029193 |
| ENSMUSG0000002611 | chr1 | Inpp4a | 0.112393703 | 0.811029193 |
| ENSMUSG0000002333 | chr2 | Dtwd1 | 0.159722352 | 0.811029193 |
| ENSMUSG0000002747 | chr2 | Pdrg1 | 0.140517697 | 0.811047277 |
| ENSMUSG0000002399 | chr17 | Tspo2 | -0.694236381 | 0.811085352 |
| ENSMUSG0000002227 | chr15 | Myo10 | 0.092301588 | 0.811085352 |
| ENSMUSG0000009506 | chr7 | E030018B13Rik | 0.488148562 | 0.811085352 |
| ENSMUSG0000006952 | chr10 | Tmem19 | -0.088744915 | 0.811711524 |
| ENSMUSG0000001942 | chr8 | Fkbp8 | 0.091699373 | 0.811842927 |
| ENSMUSG0000003636 | chr17 | Rmdn2 | 0.143841861 | 0.812077521 |
| ENSMUSG0000000588 | chr1 | Ncoa2 | -0.087040441 | 0.812090067 |
| ENSMUSG0000000094 | chr6 | Hoxa4 | -0.193178435 | 0.812140573 |
| ENSMUSG0000004164 | chrX | Klf8 | -0.180336209 | 0.812140573 |
| ENSMUSG0000003211 | chr9 | Chek1 | -0.392004133 | 0.812140573 |
| ENSMUSG0000003191 | chr9 | Mtmr2 | 0.069091311 | 0.812140573 |
| ENSMUSG0000006338 | chr9 | Bcl9l | 0.134698232 | 0.812140573 |
| ENSMUSG0000002888 | chr4 | Ppp1r8 | 0.114239184 | 0.812140573 |
| ENSMUSG0000003429 | chr18 | Fhod3 | -0.133856862 | 0.812484686 |
| ENSMUSG0000002217 | chr14 | Lrp10 | -0.074423969 | 0.812484686 |
| ENSMUSG0000003829 | chr18 | Wdr36 | 0.08147965 | 0.812484686 |
| ENSMUSG0000003379 | chrX | Atp7a | 0.087478011 | 0.812484686 |
| ENSMUSG0000005085 | chr7 | Zfp940 | -0.141694028 | 0.812617593 |
| ENSMUSG0000003858 | chr10 | Akap12 | -0.084894403 | 0.812617593 |
| ENSMUSG0000003184 | chr8 | Comp | -0.247227258 | 0.812745146 |
| ENSMUSG0000003190 | chr8 | Nfatc3 | -0.089871068 | 0.812745146 |
| ENSMUSG0000001987 | chr10 | Smpdl3a | -0.091400003 | 0.812745146 |
| ENSMUSG0000000101 | chr3 | Ilf2 | 0.067755314 | 0.812773514 |
| ENSMUSG0000002685 | chr2 | Crat | 0.092879329 | 0.812774726 |
| ENSMUSG0000003020 | chr6 | Fam234b | -0.124016207 | 0.813249247 |
| ENSMUSG0000002442 | chr18 | Zfp521 | 0.114549499 | 0.813249247 |
| ENSMUSG0000000412 | chr3 | Trmt10a | 0.133615249 | 0.813249247 |
| ENSMUSG0000002451 | chr18 | Sec11c | 0.119007228 | 0.813249247 |
| ENSMUSG0000004144 | chr9 | Gk5 | 0.162325923 | 0.813249247 |
| ENSMUSG0000002083 | chr11 | Slc6a4 | 0.261511104 | 0.813249247 |
| ENSMUSG0000003838 | chr2 | Pigu | 0.074245031 | 0.813336924 |
| ENSMUSG0000002083 | chr11 | 0610010K14Rik | 0.308147354 | 0.813479417 |
| ENSMUSG0000003193 | chr9 | Taf1d | -0.11546745 | 0.813596569 |

| | | | | |
|---|---|---|---|---|
| ENSMUSG0000006045 | chr16 | Kng2 | -0.477323313 | 0.81371297 |
| ENSMUSG0000002281 | chr16 | Cyp2ab1 | -0.639562888 | 0.81371297 |
| ENSMUSG0000002474 | chr19 | Aldh1a7 | -0.382283467 | 0.81371297 |
| ENSMUSG0000003656 | chr5 | Ttyh3 | -0.144140089 | 0.81371297 |
| ENSMUSG0000000357 | chr8 | Crtc1 | -0.117166971 | 0.81371297 |
| ENSMUSG0000001547 | chr17 | Prrt1 | -0.1177459 | 0.81371297 |
| ENSMUSG0000005325 | chr14 | Ndfip2 | -0.08172497 | 0.81371297 |
| ENSMUSG0000003244 | chr9 | Tgfbr2 | 0.057417846 | 0.81371297 |
| ENSMUSG0000006047 | chr17 | Wtap | 0.061878287 | 0.81371297 |
| ENSMUSG0000002488 | chr19 | Rce1 | 0.114923638 | 0.81371297 |
| ENSMUSG0000006086 | chr7 | Ube2s | 0.121682089 | 0.81371297 |
| ENSMUSG0000004106 | chr17 | Mslnl | 0.239016765 | 0.81371297 |
| ENSMUSG0000003251 | chr9 | Slc25a38 | 0.145373626 | 0.81371297 |
| ENSMUSG0000002598 | chr1 | Ccdc150 | -0.556944151 | 0.813722689 |
| ENSMUSG0000001884 | chr11 | Wwc1 | -0.267961585 | 0.813878403 |
| ENSMUSG0000001281 | chr10 | Cdh23 | -0.284321392 | 0.813920994 |
| ENSMUSG0000006904 | chrY | Eif2s3y | 0.074639288 | 0.814095302 |
| ENSMUSG0000006348 | chr9 | Zkscan7 | -0.15279142 | 0.814164714 |
| ENSMUSG0000003867 | chr17 | Scube3 | -0.199540682 | 0.814272943 |
| ENSMUSG0000002884 | chr4 | Ago3 | -0.127044358 | 0.814299324 |
| ENSMUSG0000007745 | chr17 | Rab11b | 0.083616675 | 0.814299324 |
| ENSMUSG0000004268 | chr9 | Mapk6 | -0.119078677 | 0.814870618 |
| ENSMUSG0000002250 | chr16 | Ciita | -0.228110434 | 0.814870618 |
| ENSMUSG0000002843 | chr4 | Ubap2 | 0.085434456 | 0.814870618 |
| ENSMUSG0000005017 | chr3 | Nudt6 | 0.239686029 | 0.814870618 |
| ENSMUSG0000002267 | chr16 | Mcm4 | -0.106622894 | 0.814940185 |
| ENSMUSG0000011728 | chr17 | Gm7072 | 0.088042369 | 0.815039484 |
| ENSMUSG0000005442 | chr4 | Atp5if1 | 0.110642315 | 0.815039484 |
| ENSMUSG0000004529 | chr5 | Insig1 | -0.188530219 | 0.815288305 |
| ENSMUSG0000003822 | chr6 | Hoxa9 | -0.124542924 | 0.815302656 |
| ENSMUSG0000002016 | chr10 | Tcf3 | 0.082681772 | 0.815370566 |
| ENSMUSG0000009948 | chr7 | Xndc1 | -0.182800317 | 0.815473059 |
| ENSMUSG0000003646 | chr9 | Megf11 | -0.194329162 | 0.815473059 |
| ENSMUSG0000003349 | chr2 | Strc | -0.852919978 | 0.815473059 |
| ENSMUSG0000005531 | chr13 | Pgbd1 | -0.243581918 | 0.815473059 |
| ENSMUSG0000001535 | chr9 | Clpx | 0.087030597 | 0.815473059 |
| ENSMUSG0000004603 | chr15 | Otulin | 0.091375475 | 0.815473059 |
| ENSMUSG0000004652 | chr14 | Kctd4 | 0.120663035 | 0.815473059 |
| ENSMUSG0000000963 | chr1 | G0s2 | 0.109285478 | 0.815473059 |
| ENSMUSG0000000636 | chr15 | Fbln1 | 0.126403184 | 0.815473059 |
| ENSMUSG0000002005 | chr10 | Sycp3 | 0.378618701 | 0.815473059 |
| ENSMUSG0000005245 | chr16 | Atp6v1a | -0.065100939 | 0.815594775 |
| ENSMUSG0000009733 | chr13 | Zfp87 | -0.099790255 | 0.815603366 |
| ENSMUSG0000000583 | chr18 | Gata6 | -0.298486038 | 0.815646338 |
| ENSMUSG0000002448 | chr18 | Ap3s1 | 0.100884051 | 0.815721838 |
| ENSMUSG0000003107 | chr7 | LTO1 | 0.103541468 | 0.815721838 |
| ENSMUSG0000005213 | chr16 | Sema5b | 0.388953273 | 0.815798806 |
| ENSMUSG0000002483 | chr19 | Coro1b | -0.102626689 | 0.815834201 |
| ENSMUSG0000002396 | chr17 | Crip3 | -0.353523775 | 0.815915544 |
| ENSMUSG0000003696 | chr3 | Clca2 | -0.5067467 | 0.815915544 |

| | | | | |
|---|---|---|---|---|
| ENSMUSG0000003495 | chr7 | Cog7 | -0.087685568 | 0.815915544 |
| ENSMUSG0000004804 | chrX | Zbtb33 | -0.076605901 | 0.81597436 |
| ENSMUSG0000005161 | chr14 | Rap2a | -0.096066535 | 0.81601509 |
| ENSMUSG0000001505 | chr6 | Gata2 | 0.253749964 | 0.81605364 |
| ENSMUSG0000003689 | chr3 | Rap2b | -0.128546895 | 0.816100344 |
| ENSMUSG0000003006 | chr6 | Frmd4b | 0.129874677 | 0.816100344 |
| ENSMUSG0000003743 | chr1 | Slc30a1 | -0.159333483 | 0.816151254 |
| ENSMUSG0000004834 | chr18 | Pcdhb18 | -0.162154167 | 0.816217526 |
| ENSMUSG0000004787 | chr4 | Gpr157 | -0.148071019 | 0.816534996 |
| ENSMUSG0000002865 | chr4 | Ppie | 0.166083751 | 0.816534996 |
| ENSMUSG0000003995 | chr16 | Mrap | 0.632906781 | 0.816534996 |
| ENSMUSG0000010956 | chr9 | Muc16 | -0.366149915 | 0.816682822 |
| ENSMUSG0000002467 | chr19 | Ms4a4c | -0.503457882 | 0.816682822 |
| ENSMUSG0000002416 | chr17 | Spsb3 | 0.179996338 | 0.816682822 |
| ENSMUSG0000009202 | chr5 | Gbp11 | -0.63951397 | 0.816751112 |
| ENSMUSG0000007910 | chr5 | Pms2 | -0.115072501 | 0.816751112 |
| ENSMUSG0000002237 | chr15 | Cyrib | 0.070417162 | 0.816751112 |
| ENSMUSG0000003398 | chr11 | Coil | 0.126338743 | 0.816751112 |
| ENSMUSG0000009530 | chr14 | Plac9 | 0.225516308 | 0.816751112 |
| ENSMUSG0000004541 | chr9 | Dipk2a | -0.095172155 | 0.817019006 |
| ENSMUSG0000004616 | chr13 | Adamts6 | -0.219255951 | 0.817135398 |
| ENSMUSG0000004087 | chr12 | Wdr25 | -0.155513529 | 0.817135398 |
| ENSMUSG0000004792 | chr15 | Trappc9 | -0.085006103 | 0.817135398 |
| ENSMUSG0000003049 | chr7 | Rhpn2 | -0.310567147 | 0.817311435 |
| ENSMUSG0000004079 | chr2 | C1qtnf4 | -0.300153677 | 0.817311435 |
| ENSMUSG0000002606 | chr1 | Ptp4a1 | -0.580038805 | 0.817311435 |
| ENSMUSG0000002921 | chr5 | Tec | -0.218839199 | 0.817311435 |
| ENSMUSG0000002749 | chr2 | E2f1 | -0.209942802 | 0.817311435 |
| ENSMUSG0000002829 | chr4 | Cfap206 | -0.260262407 | 0.817311435 |
| ENSMUSG0000003396 | chrX | Slc16a2 | -0.127047611 | 0.817311435 |
| ENSMUSG0000003095 | chr7 | Fam53b | -0.106579837 | 0.817311435 |
| ENSMUSG0000002279 | chr16 | Arhgap31 | -0.10756559 | 0.817311435 |
| ENSMUSG0000002800 | chr3 | Lrat | -0.323721118 | 0.817311435 |
| ENSMUSG0000005737 | chr4 | Yipf1 | -0.1095502 | 0.817311435 |
| ENSMUSG0000002485 | chr19 | Pacs1 | -0.098187147 | 0.817311435 |
| ENSMUSG0000002950 | chr5 | P2rx2 | -0.096682889 | 0.817311435 |
| ENSMUSG0000002823 | chr4 | Tmem68 | -0.099790409 | 0.817311435 |
| ENSMUSG0000002852 | chr4 | Slc35d1 | -0.117596744 | 0.817311435 |
| ENSMUSG0000006219 | chr6 | Lancl2 | -0.094471756 | 0.817311435 |
| ENSMUSG0000000781 | chr14 | Zmiz1 | -0.085790432 | 0.817311435 |
| ENSMUSG0000003775 | chr11 | Fam222b | -0.078643797 | 0.817311435 |
| ENSMUSG0000002721 | chr2 | Tspan18 | -0.076694979 | 0.817311435 |
| ENSMUSG0000002045 | chr11 | Trim11 | -0.081395303 | 0.817311435 |
| ENSMUSG0000003814 | chr9 | Snrk | -0.079699441 | 0.817311435 |
| ENSMUSG0000004935 | chr11 | Dcaf7 | -0.071136178 | 0.817311435 |
| ENSMUSG0000003470 | chr10 | Gns | 0.088090631 | 0.817311435 |
| ENSMUSG0000002406 | chr17 | Spast | 0.076758981 | 0.817311435 |
| ENSMUSG0000002663 | chr1 | Angel2 | 0.084372614 | 0.817311435 |
| ENSMUSG0000007105 | chr17 | Safb | 0.082852429 | 0.817311435 |
| ENSMUSG0000003196 | chr9 | Acad8 | 0.080229912 | 0.817311435 |

| | | | | |
|---|---|---|---|---|
| ENSMUSG0000004439 | chr11 | Tmed4 | 0.093807525 | 0.817311435 |
| ENSMUSG0000003454 | chr3 | Rsrc1 | 0.088954681 | 0.817311435 |
| ENSMUSG0000002919 | chr5 | Lias | 0.089250699 | 0.817311435 |
| ENSMUSG0000007375 | chr4 | 5730409E04Rik | 0.096622415 | 0.817311435 |
| ENSMUSG0000002132 | chr13 | Trim27 | 0.093559307 | 0.817311435 |
| ENSMUSG0000002166 | chr13 | Polk | 0.100441891 | 0.817311435 |
| ENSMUSG0000002944 | chr5 | Psmd9 | 0.115377652 | 0.817311435 |
| ENSMUSG0000006868 | chr2 | Cd59b | 0.138512059 | 0.817311435 |
| ENSMUSG0000003821 | chr11 | Tlcd2 | 0.152052008 | 0.817311435 |
| ENSMUSG0000003457 | chr5 | Ptpn13 | 0.135570478 | 0.817311435 |
| ENSMUSG0000006787 | chrX | Map7d3 | 0.194147731 | 0.817311435 |
| ENSMUSG0000003763 | chrX | Slc25a43 | 0.335205889 | 0.817311435 |
| ENSMUSG0000003744 | chr1 | Arid5a | 0.210425282 | 0.817311435 |
| ENSMUSG0000002391 | chr17 | Slc25a27 | -0.148970769 | 0.817383801 |
| ENSMUSG0000005152 | chr7 | Usp29 | -0.161748658 | 0.817383801 |
| ENSMUSG0000004267 | chr4 | Zc3h12a | -0.175370847 | 0.817383801 |
| ENSMUSG0000003916 | chr10 | Akap7 | -0.10981719 | 0.817383801 |
| ENSMUSG0000002071 | chr11 | Nf1 | -0.134735301 | 0.817383801 |
| ENSMUSG0000002297 | chr16 | Paxbp1 | -0.105261533 | 0.817383801 |
| ENSMUSG0000006352 | chr13 | Stmnd1 | -0.132436443 | 0.817383801 |
| ENSMUSG0000002628 | chr1 | Inpp5d | -0.140831221 | 0.817383801 |
| ENSMUSG0000006022 | chr2 | Golm2 | -0.099797652 | 0.817383801 |
| ENSMUSG0000002896 | chr4 | Pgd | -0.109418196 | 0.817383801 |
| ENSMUSG0000003211 | chr9 | Slc37a4 | -0.108565477 | 0.817383801 |
| ENSMUSG0000001072 | chr5 | Lmbr1 | -0.088603432 | 0.817383801 |
| ENSMUSG0000005830 | chr8 | Upf1 | 0.059443102 | 0.817383801 |
| ENSMUSG0000001407 | chr16 | Rnf168 | 0.069937074 | 0.817383801 |
| ENSMUSG0000004300 | chr14 | Gng2 | 0.086595048 | 0.817383801 |
| ENSMUSG0000005248 | chr8 | Cherp | 0.077510735 | 0.817383801 |
| ENSMUSG0000001434 | chr11 | Ube2z | 0.076590504 | 0.817383801 |
| ENSMUSG0000002159 | chr13 | Nsun2 | 0.079737201 | 0.817383801 |
| ENSMUSG0000003143 | chrX | Prps1 | 0.090521256 | 0.817383801 |
| ENSMUSG0000003293 | chr8 | Nup93 | 0.08223819 | 0.817383801 |
| ENSMUSG0000002213 | chr14 | Dnajc3 | 0.106939256 | 0.817383801 |
| ENSMUSG0000003787 | chr10 | Jmjd1c | 0.096716081 | 0.817383801 |
| ENSMUSG0000005464 | chr17 | Slc8a1 | 0.103255758 | 0.817383801 |
| ENSMUSG0000002962 | chr5 | Pdap1 | 0.096376832 | 0.817383801 |
| ENSMUSG0000001631 | chrX | Slc25a5 | 0.123841208 | 0.817383801 |
| ENSMUSG0000003823 | chr6 | Hoxa7 | 0.089079666 | 0.817383801 |
| ENSMUSG0000004586 | chr10 | Cradd | 0.120607421 | 0.817383801 |
| ENSMUSG0000003255 | chr9 | 1110059G10Rik | 0.108392364 | 0.817383801 |
| ENSMUSG0000002202 | chr14 | Naa16 | 0.125909229 | 0.817383801 |
| ENSMUSG0000002032 | chr10 | Prss57 | 0.473409087 | 0.817383801 |
| ENSMUSG0000002069 | chr11 | Nle1 | 0.142159526 | 0.817383801 |
| ENSMUSG0000003305 | chr15 | Ankrd54 | 0.11522002 | 0.817383801 |
| ENSMUSG0000006306 | chr14 | Sox7 | 0.23174451 | 0.817383801 |
| ENSMUSG0000003042 | chr7 | Lilra6 | 0.435835303 | 0.817383801 |
| ENSMUSG0000006643 | chr12 | Plekhd1 | 0.35947284 | 0.817383801 |
| ENSMUSG0000001565 | chr5 | Steap1 | 0.594911033 | 0.817383801 |
| ENSMUSG0000002598 | chr1 | Coq10b | 0.186248156 | 0.817383801 |

| | | | | |
|---|---|---|---|---|
| ENSMUSG0000003912 | chr2 | Cdc123 | 0.090393854 | 0.817742137 |
| ENSMUSG0000004580 | chrX | Hsf3 | -0.857568116 | 0.817787514 |
| ENSMUSG0000003169 | chr8 | Mylk3 | -0.377560564 | 0.817843419 |
| ENSMUSG0000003010 | chr6 | Bhlhe40 | -0.25583266 | 0.817949047 |
| ENSMUSG0000001244 | chr19 | Kif11 | -0.32594524 | 0.818242279 |
| ENSMUSG0000006104 | chr8 | Cdh3 | 0.36542482 | 0.818408432 |
| ENSMUSG0000002173 | chr13 | Fgf10 | -0.570398309 | 0.818517156 |
| ENSMUSG0000004260 | chr7 | Hirip3 | -0.11234997 | 0.818517156 |
| ENSMUSG0000001728 | chr11 | Vps53 | -0.085277954 | 0.818517156 |
| ENSMUSG0000004047 | chr14 | Rabggta | -0.104479098 | 0.818517156 |
| ENSMUSG0000000313 | chr1 | Cnot11 | 0.085390363 | 0.818517156 |
| ENSMUSG0000002479 | chr19 | Sac3d1 | 0.138282403 | 0.818517156 |
| ENSMUSG0000004662 | chr4 | Gjb4 | 0.62193069 | 0.818517156 |
| ENSMUSG0000002884 | chr4 | Zfp593 | 0.48254494 | 0.818542192 |
| ENSMUSG0000002678 | chr2 | Pdss1 | -0.145231608 | 0.818675041 |
| ENSMUSG0000003856 | chr5 | Ift172 | -0.073329671 | 0.818675041 |
| ENSMUSG0000006042 | chr8 | Zfp868 | 0.100191792 | 0.818675041 |
| ENSMUSG0000002526 | chrX | Tsr2 | 0.08939374 | 0.818943197 |
| ENSMUSG0000005685 | chrX | Pou3f4 | 0.507750694 | 0.818943197 |
| ENSMUSG0000001252 | chr5 | Phox2b | -0.120286195 | 0.8190415 |
| ENSMUSG0000003098 | chr7 | Stim1 | 0.083624768 | 0.8190415 |
| ENSMUSG0000000035 | chr2 | Rem1 | 0.170657044 | 0.8190415 |
| ENSMUSG0000002603 | chr1 | Sgo2a | 0.531502616 | 0.819062118 |
| ENSMUSG0000004869 | chr10 | Mex3d | 0.110774974 | 0.819153041 |
| ENSMUSG0000002296 | chr16 | Son | -0.112541665 | 0.819197938 |
| ENSMUSG0000003694 | chr15 | Tmem71 | -0.231110322 | 0.81923242 |
| ENSMUSG0000003175 | chr8 | Bbs2 | -0.112523544 | 0.81923242 |
| ENSMUSG0000003746 | chr7 | Fbxo27 | -0.181190814 | 0.81923242 |
| ENSMUSG0000005150 | chr5 | Ccdc121rt3 | -0.56579108 | 0.81923242 |
| ENSMUSG0000002761 | chr3 | Hps3 | -0.087907985 | 0.81923242 |
| ENSMUSG0000004951 | chr7 | Spty2d1 | -0.077284776 | 0.81923242 |
| ENSMUSG0000001399 | chr1 | Nit1 | 0.093251121 | 0.81923242 |
| ENSMUSG0000004205 | chr12 | Dync2i1 | 0.090966136 | 0.81923242 |
| ENSMUSG0000002938 | chr5 | Gtf2h3 | 0.114431817 | 0.81923242 |
| ENSMUSG0000004505 | chr19 | Prlhr | 0.511596828 | 0.81923242 |
| ENSMUSG0000002635 | chr1 | Dars1 | 0.13693939 | 0.81923242 |
| ENSMUSG0000003074 | chr7 | Spns1 | 0.098914669 | 0.81923242 |
| ENSMUSG0000003874 | chr9 | Angptl6 | 0.231727888 | 0.81923242 |
| ENSMUSG0000003844 | chr10 | Cdc40 | 0.094165678 | 0.819247459 |
| ENSMUSG0000002034 | chr11 | Ppp2ca | 0.092724013 | 0.819295116 |
| ENSMUSG0000002771 | chr3 | Ccna2 | -0.25857338 | 0.819384676 |
| ENSMUSG0000002684 | chr2 | Lamc3 | -0.133796617 | 0.819384676 |
| ENSMUSG0000000510 | chr5 | Slc2a9 | -0.216228003 | 0.819384676 |
| ENSMUSG0000003512 | chr14 | Neil2 | -0.307636097 | 0.819384676 |
| ENSMUSG0000000990 | chr1 | Kdsr | -0.112527854 | 0.819384676 |
| ENSMUSG0000006861 | chr2 | Gjd2 | -0.234528396 | 0.819384676 |
| ENSMUSG0000000032 | chr11 | Alox12 | -0.292882891 | 0.819384676 |
| ENSMUSG0000000316 | chr16 | Dgcr2 | -0.092559797 | 0.819384676 |
| ENSMUSG0000003191 | chr8 | Cog8 | -0.094218237 | 0.819384676 |
| ENSMUSG0000002627 | chr1 | Thap4 | -0.097203208 | 0.819384676 |

| | | | | |
|---|---|---|---|---|
| ENSMUSG0000004660 | chr9 | Tcaim | -0.081302636 | 0.819384676 |
| ENSMUSG0000001529 | chrX | Gdi1 | -0.063021186 | 0.819384676 |
| ENSMUSG0000007117 | chr17 | Srsf3 | 0.103746651 | 0.819384676 |
| ENSMUSG0000003246 | chr9 | Dbr1 | 0.099116426 | 0.819384676 |
| ENSMUSG0000002181 | chr14 | Mss51 | 0.291030947 | 0.819384676 |
| ENSMUSG0000001072 | chr5 | Mpv17 | 0.128253114 | 0.81940949 |
| ENSMUSG0000005749 | chr6 | Fam136a | 0.128282988 | 0.819519397 |
| ENSMUSG0000005286 | chr6 | Dnah6 | 0.40763622 | 0.819586259 |
| ENSMUSG0000002161 | chr13 | Hapln1 | -0.184767973 | 0.819859279 |
| ENSMUSG0000003502 | chr9 | Ncapd3 | -0.160733588 | 0.819937902 |
| ENSMUSG0000005145 | chr7 | Spn | -0.439539024 | 0.819944381 |
| ENSMUSG0000005448 | chr2 | Tmem62 | -0.142173603 | 0.82027755 |
| ENSMUSG0000002012 | chr10 | Srgap1 | 0.173408694 | 0.82027755 |
| ENSMUSG0000003965 | chr19 | Cpeb3 | -0.149450326 | 0.82039414 |
| ENSMUSG0000009009 | chr2 | Gm14399 | -0.171859905 | 0.82039414 |
| ENSMUSG0000002633 | chr1 | Gin1 | -0.147845305 | 0.82039414 |
| ENSMUSG0000000031 | chr11 | Clec10a | -0.313715878 | 0.82039414 |
| ENSMUSG0000002064 | chr12 | Mboat2 | -0.111370828 | 0.82039414 |
| ENSMUSG0000005831 | chr2 | Phf21a | -0.097297693 | 0.82039414 |
| ENSMUSG0000000535 | chr10 | Slc1a6 | -0.305257778 | 0.82039414 |
| ENSMUSG0000003466 | chr10 | Xpot | -0.073034045 | 0.82039414 |
| ENSMUSG0000002291 | chr16 | Arl13b | -0.068990719 | 0.82039414 |
| ENSMUSG0000002241 | chr15 | Mief1 | 0.07746259 | 0.82039414 |
| ENSMUSG0000002082 | chr11 | Sec14l1 | 0.080467671 | 0.82039414 |
| ENSMUSG0000007166 | chr19 | Polr2g | 0.160211942 | 0.82039414 |
| ENSMUSG0000005901 | chr2 | Sh2d3c | -0.17669155 | 0.820434617 |
| ENSMUSG0000009855 | chr14 | Kctd12 | -0.077941049 | 0.820614915 |
| ENSMUSG0000002643 | chr1 | Rassf5 | 0.117102063 | 0.820614915 |
| ENSMUSG0000006027 | chr18 | Nrg2 | -0.189293619 | 0.820990229 |
| ENSMUSG0000003403 | chr6 | Fgd5 | -0.156061765 | 0.82113635 |
| ENSMUSG0000001039 | chr11 | Gosr1 | -0.062215664 | 0.82113635 |
| ENSMUSG0000005883 | chr6 | Opn1sw | 0.279310333 | 0.82113635 |
| ENSMUSG0000007388 | chr4 | Ccl27a | -0.144949027 | 0.821372088 |
| ENSMUSG0000004566 | chr15 | Mfsd5 | 0.071895904 | 0.821461168 |
| ENSMUSG0000001892 | chr11 | Ccl6 | -0.1079576 | 0.821505022 |
| ENSMUSG0000003160 | chr8 | Cnot7 | 0.066438003 | 0.821505022 |
| ENSMUSG0000003300 | chr14 | Mycbp2 | -0.101048026 | 0.821518216 |
| ENSMUSG0000003839 | chr8 | Upf3a | 0.088718139 | 0.821531576 |
| ENSMUSG0000000122 | chr17 | Dpp9 | 0.060449932 | 0.82166001 |
| ENSMUSG0000005195 | chr5 | B3glct | -0.097500745 | 0.821758874 |
| ENSMUSG0000003962 | chr2 | Prex1 | 0.15871849 | 0.82181797 |
| ENSMUSG0000007448 | chr3 | Bglap3 | -0.780087169 | 0.822019037 |
| ENSMUSG0000000337 | chr7 | Grik5 | -0.112395894 | 0.822019037 |
| ENSMUSG0000006073 | chr15 | Rxfp3 | -0.236974726 | 0.822019037 |
| ENSMUSG0000003820 | chr6 | Hoxa13 | 0.190651022 | 0.822271092 |
| ENSMUSG0000004778 | chr13 | Slc38a9 | -0.096241795 | 0.822291482 |
| ENSMUSG0000004346 | chr1 | Zbtb37 | -0.119098299 | 0.822401846 |
| ENSMUSG0000003526 | chr5 | Helq | 0.085793617 | 0.822417766 |
| ENSMUSG0000005019 | chr2 | Lgr4 | 0.068737784 | 0.822461848 |
| ENSMUSG0000006991 | chr11 | Spdl1 | -0.527008126 | 0.822667972 |

| | | | | |
|---|---|---|---|---|
| ENSMUSG0000006022 | chr19 | Pyroxd2 | -0.130666903 | 0.822790328 |
| ENSMUSG0000002004 | chr10 | Timp3 | 0.087423711 | 0.822790328 |
| ENSMUSG0000003953 | chr2 | Stau1 | 0.081772088 | 0.822790328 |
| ENSMUSG0000008985 | chr8 | Zfp882 | -0.176931748 | 0.822794161 |
| ENSMUSG0000002654 | chr1 | Cfap45 | -0.216088473 | 0.822794161 |
| ENSMUSG0000004132 | chr7 | Pcf11 | -0.110252842 | 0.822794161 |
| ENSMUSG0000002476 | chr19 | Otub1 | 0.071456589 | 0.822794161 |
| ENSMUSG0000003995 | chr18 | Stk32a | 0.109338239 | 0.822794161 |
| ENSMUSG0000005815 | chr18 | Chsy3 | 0.279465954 | 0.822794161 |
| ENSMUSG0000004885 | chr12 | Slc25a47 | 0.229619464 | 0.822794161 |
| ENSMUSG0000000181 | chr2 | Hoxd13 | 0.468326694 | 0.822903708 |
| ENSMUSG0000003802 | chr1 | Kcnj9 | -0.272604377 | 0.82292062 |
| ENSMUSG0000003228 | chr9 | Tle3 | 0.058566246 | 0.82292062 |
| ENSMUSG0000003730 | chr8 | Ttc13 | 0.081013388 | 0.82292062 |
| ENSMUSG0000004494 | chr11 | Ubtd2 | 0.110820409 | 0.82292062 |
| ENSMUSG0000005134 | chr10 | Sprydd4 | 0.125572442 | 0.82292062 |
| ENSMUSG0000002526 | chrX | Fgd1 | -0.124693416 | 0.822973109 |
| ENSMUSG0000004332 | chr5 | Fbrsl1 | -0.094504493 | 0.822973109 |
| ENSMUSG0000001625 | chr2 | Nelfcd | -0.079834 | 0.822973109 |
| ENSMUSG0000002101 | chr12 | Ttc8 | -0.081099644 | 0.822973109 |
| ENSMUSG0000006980 | chr11 | Gm10277 | -0.40042489 | 0.82316739 |
| ENSMUSG0000004013 | chr7 | Abcc8 | -0.138749649 | 0.82316739 |
| ENSMUSG0000003864 | chr7 | Pold1 | -0.096794319 | 0.82316739 |
| ENSMUSG0000003834 | chr6 | Zfp384 | -0.089760183 | 0.82316739 |
| ENSMUSG0000002102 | chr12 | Mbip | -0.09469595 | 0.82316739 |
| ENSMUSG0000006039 | chr7 | Zfp128 | -0.09889552 | 0.82316739 |
| ENSMUSG0000002283 | chr16 | Eaf2 | -0.356216903 | 0.82316739 |
| ENSMUSG0000002940 | chr5 | G3bp2 | 0.07621479 | 0.82316739 |
| ENSMUSG0000003931 | chr1 | Rab3gap2 | 0.074042402 | 0.82316739 |
| ENSMUSG0000002856 | chr4 | Btf3l4 | 0.071156868 | 0.82316739 |
| ENSMUSG0000002770 | chr3 | Dcun1d1 | 0.081977631 | 0.82316739 |
| ENSMUSG0000002466 | chr19 | Cd5 | 0.473399398 | 0.82316739 |
| ENSMUSG0000002169 | chr13 | Dimt1 | 0.124593506 | 0.82316739 |
| ENSMUSG0000002902 | chr4 | Dffb | 0.184141655 | 0.82316739 |
| ENSMUSG0000004155 | chr1 | Fmod | 0.215127921 | 0.82316739 |
| ENSMUSG0000004572 | chr6 | Prr15 | -0.356279632 | 0.823271134 |
| ENSMUSG0000002795 | chr3 | Sass6 | -0.162154041 | 0.823271134 |
| ENSMUSG0000002086 | chr11 | Xylt2 | -0.107607284 | 0.823271134 |
| ENSMUSG0000000999 | chrX | Tafazzin | -0.088095173 | 0.823271134 |
| ENSMUSG0000004450 | chr3 | Lingo4 | -0.539267145 | 0.823425514 |
| ENSMUSG0000002749 | chr2 | Aurka | 0.23554842 | 0.823425514 |
| ENSMUSG0000002793 | chr3 | Npr1 | -0.179502609 | 0.823547073 |
| ENSMUSG0000001638 | chrX | Pls3 | -0.080585396 | 0.823547073 |
| ENSMUSG0000000306 | chr10 | Stk11 | 0.069667379 | 0.823547073 |
| ENSMUSG0000003494 | chr11 | Synrg | -0.074712323 | 0.823600194 |
| ENSMUSG0000004192 | chr11 | Slc16a6 | 0.165513424 | 0.823600194 |
| ENSMUSG0000005501 | chr1 | Agap1 | 0.102101171 | 0.823766983 |
| ENSMUSG0000003250 | chr9 | Fbxl2 | -0.117044125 | 0.823767035 |
| ENSMUSG0000000529 | chr5 | Letm1 | -0.073177137 | 0.823969782 |
| ENSMUSG0000007372 | chr1 | Lmbrd1 | -0.072314558 | 0.823969782 |

| | | | | |
|---|---|---|---|---|
| ENSMUSG0000002728 | chr2 | Haus2 | 0.075028915 | 0.823969782 |
| ENSMUSG0000004113 | chr4 | Ripk2 | 0.083927265 | 0.823969782 |
| ENSMUSG0000010189 | chr15 | 9130401M01Rik | 0.166958749 | 0.823969782 |
| ENSMUSG0000002238 | chr15 | Gtse1 | 0.389263628 | 0.823969782 |
| ENSMUSG0000004030 | chr5 | Rbm48 | -0.117675313 | 0.824048159 |
| ENSMUSG0000002727 | chr2 | Jag1 | 0.103737213 | 0.824075697 |
| ENSMUSG0000003630 | chr11 | Skp1 | 0.110750433 | 0.824100825 |
| ENSMUSG0000002303 | chr15 | Nr4a1 | -0.521554761 | 0.824265125 |
| ENSMUSG0000007532 | chr2 | Fign | -0.123295141 | 0.824265125 |
| ENSMUSG0000003059 | chr7 | Nfkbib | 0.107587436 | 0.824265125 |
| ENSMUSG0000004981 | chr11 | Fam161a | 0.276347455 | 0.824265125 |
| ENSMUSG0000002676 | chr2 | Lypd6b | -0.361727064 | 0.824384768 |
| ENSMUSG0000002655 | chr1 | Vangl2 | -0.138938464 | 0.824384768 |
| ENSMUSG0000002809 | chr3 | Acp6 | -0.102729116 | 0.824384768 |
| ENSMUSG0000004349 | chr6 | Tril | -0.097098528 | 0.824384768 |
| ENSMUSG0000003757 | chr15 | Mcrs1 | -0.104581137 | 0.824384768 |
| ENSMUSG0000003920 | chr7 | Abhd2 | -0.104102632 | 0.824384768 |
| ENSMUSG0000003257 | chr9 | Atp2c1 | 0.052598403 | 0.824384768 |
| ENSMUSG0000000019 | chr2 | Gpr107 | 0.053978164 | 0.824384768 |
| ENSMUSG0000002394 | chr17 | Hsp90ab1 | 0.116810174 | 0.824384768 |
| ENSMUSG0000003177 | chr8 | Psme3ip1 | 0.0877138 | 0.824384768 |
| ENSMUSG0000002661 | chr1 | Bpnt1 | 0.128964831 | 0.824384768 |
| ENSMUSG0000002302 | chr15 | Cox14 | 0.153319622 | 0.824384768 |
| ENSMUSG0000003884 | chr2 | Birc7 | 0.498980233 | 0.824384768 |
| ENSMUSG0000005311 | chr11 | Socs3 | 0.151532832 | 0.824384768 |
| ENSMUSG0000002063 | chr12 | Pfn4 | 0.362327288 | 0.824384768 |
| ENSMUSG0000010303 | chr18 | Pcdhgb1 | 0.189214338 | 0.824384768 |
| ENSMUSG0000002022 | chr10 | Tmbim4 | -0.083043092 | 0.824494369 |
| ENSMUSG0000001030 | chr7 | Tmem86a | -0.189765294 | 0.824622603 |
| ENSMUSG0000001994 | chr10 | Cdk1 | -0.311428844 | 0.824622603 |
| ENSMUSG0000002898 | chr5 | Klhl7 | 0.079082609 | 0.824622603 |
| ENSMUSG0000005311 | chr9 | Yap1 | 0.082167872 | 0.824622603 |
| ENSMUSG0000005746 | chr12 | E2f6 | 0.108739213 | 0.824622603 |
| ENSMUSG0000004206 | chr12 | Zfp386 | 0.099721364 | 0.824622603 |
| ENSMUSG0000005777 | chr3 | Mab21l2 | 0.097210535 | 0.824622603 |
| ENSMUSG0000004103 | chr7 | Irgq | 0.070905969 | 0.824713692 |
| ENSMUSG0000006420 | chr19 | Spata6l | 0.310972783 | 0.824713692 |
| ENSMUSG0000002864 | chr4 | Macf1 | 0.159463218 | 0.824713692 |
| ENSMUSG0000003231 | chr9 | Csk | -0.08703329 | 0.824838152 |
| ENSMUSG0000003839 | chr3 | Txnip | 0.196777349 | 0.824838152 |
| ENSMUSG0000005653 | chr5 | Ccdc18 | -0.509095036 | 0.824869974 |
| ENSMUSG0000003296 | chr16 | Ift57 | 0.078961992 | 0.824869974 |
| ENSMUSG0000003237 | chr9 | Plscr2 | -0.18843382 | 0.825059536 |
| ENSMUSG0000005364 | chr18 | Aldh7a1 | 0.103773285 | 0.825059536 |
| ENSMUSG0000002034 | chr11 | Cyfip2 | -0.096730149 | 0.825232043 |
| ENSMUSG0000002087 | chr11 | Lrrc46 | -0.313438987 | 0.82542009 |
| ENSMUSG0000006220 | chr16 | Gspt1 | 0.08158154 | 0.82542009 |
| ENSMUSG0000002699 | chr2 | Pkp4 | 0.080352718 | 0.82605062 |
| ENSMUSG0000000409 | chr9 | Dnmt1 | -0.087821979 | 0.826150902 |
| ENSMUSG0000004408 | chr2 | Efcab8 | 0.431657958 | 0.826367258 |

| | | | | |
|---|---|---|---|---|
| ENSMUSG00000062960 chr12 | | Ccdc177 | -0.631641555 | 0.826453614 |
| ENSMUSG0000006370 chr15 | | Mapk15 | -0.252821352 | 0.826453614 |
| ENSMUSG0000004648 chr9 | | Scn4b | -0.227793163 | 0.826453614 |
| ENSMUSG0000004671 chr17 | | Hmga1 | -0.097107891 | 0.826453614 |
| ENSMUSG0000004094 chr4 | | Rcc2 | -0.069395445 | 0.826453614 |
| ENSMUSG0000002978 chr6 | | Tmem209 | -0.061566584 | 0.826453614 |
| ENSMUSG0000001747 chr8 | | Zc3h18 | 0.061571871 | 0.826453614 |
| ENSMUSG0000003555 chr8 | | Mpv17l2 | 0.093753906 | 0.826453614 |
| ENSMUSG0000007069 chr7 | | Sars2 | 0.098711973 | 0.826453614 |
| ENSMUSG0000002469 chr19 | | Gna14 | 0.175866567 | 0.826453614 |
| ENSMUSG0000002617 chr1 | | Slc11a1 | 0.154031072 | 0.826453614 |
| ENSMUSG0000006875 chr14 | | Il3ra | 0.133376232 | 0.826495795 |
| ENSMUSG0000003160 chr8 | | Vps37a | 0.06866953 | 0.826726997 |
| ENSMUSG0000009802 chr7 | | Zfp82 | -0.538860915 | 0.826814709 |
| ENSMUSG0000009390 chr8 | | Tomm20 | 0.067561134 | 0.826814709 |
| ENSMUSG0000007407 chr7 | | Fam169b | 0.384320825 | 0.826814709 |
| ENSMUSG0000000639 chr4 | | Cdc20 | -0.302352689 | 0.826937371 |
| ENSMUSG0000002786 chr3 | | Cd2 | 0.262294753 | 0.826937371 |
| ENSMUSG0000002927 chr5 | | Glmn | -0.126677384 | 0.826984481 |
| ENSMUSG0000005075 chr8 | | Pgbd5 | -0.061842758 | 0.826997029 |
| ENSMUSG0000002433 chr17 | | Psmb8 | -0.192139702 | 0.827005073 |
| ENSMUSG0000002065 chr12 | | Adcy3 | -0.098307801 | 0.827005073 |
| ENSMUSG0000001918 chr2 | | H13 | -0.067108956 | 0.827005073 |
| ENSMUSG0000003084 chr7 | | Rgs10 | 0.121271133 | 0.827005073 |
| ENSMUSG0000005393 chr11 | | Shisa6 | -0.314884941 | 0.827143606 |
| ENSMUSG0000004503 chr17 | | Prkce | -0.107471209 | 0.827143606 |
| ENSMUSG0000003733 chr12 | | Mfsd2b | 0.292490091 | 0.827320618 |
| ENSMUSG0000007064 chr1 | | Etnk2 | 0.250623825 | 0.827489712 |
| ENSMUSG0000002297 chr16 | | Ifnar2 | 0.088856618 | 0.827644966 |
| ENSMUSG0000002757 chr2 | | Gid8 | 0.076509727 | 0.827658744 |
| ENSMUSG0000003187 chr8 | | Cmtm3 | -0.079038291 | 0.827833367 |
| ENSMUSG0000004011 chr6 | | Mrps35 | 0.100005801 | 0.827833367 |
| ENSMUSG0000000250 chr17 | | Nherf2 | 0.071470664 | 0.827883734 |
| ENSMUSG0000002514 chr11 | | Cenpx | 0.123221471 | 0.827883734 |
| ENSMUSG0000000601 chr1 | | Odr4 | 0.061694703 | 0.828055149 |
| ENSMUSG0000004119 chr17 | | Rpusd1 | 0.096201723 | 0.828055149 |
| ENSMUSG0000011022 chr7 | | Gm36210 | 0.350795153 | 0.828055149 |
| ENSMUSG0000003816 chr16 | | P3h2 | 0.110225845 | 0.828055149 |
| ENSMUSG0000003437 chr17 | | Tulp4 | 0.110302459 | 0.828055149 |
| ENSMUSG0000005605 chr1 | | Sag | -0.20244543 | 0.828278629 |
| ENSMUSG0000003976 chr4 | | Dnajc11 | 0.060860897 | 0.828278629 |
| ENSMUSG0000004528 chr11 | | Rtn4rl1 | 0.100063759 | 0.828278629 |
| ENSMUSG0000005839 chr17 | | Rrp1b | 0.096718084 | 0.828278629 |
| ENSMUSG0000007383 chr7 | | Tufm | 0.102740572 | 0.828278629 |
| ENSMUSG0000003881 chr19 | | Trmt112 | 0.113304338 | 0.828278629 |
| ENSMUSG0000003082 chr7 | | Zfp747l1 | 0.135577649 | 0.828278629 |
| ENSMUSG0000003752 chr14 | | Atg14 | 0.131034388 | 0.828278629 |
| ENSMUSG0000004274 chr2 | | Id1 | 0.184250387 | 0.828278629 |
| ENSMUSG0000002777 chr3 | | Gfm1 | 0.070012552 | 0.82837777 |
| ENSMUSG0000000703 chr17 | | Abhd16a | 0.075459014 | 0.828380826 |

| | | | | |
|---|---|---|---|---|
| ENSMUSG0000003134 | chrX | Gabre | -0.186747353 | 0.828423083 |
| ENSMUSG0000002589 | chr9 | Kbtbd3 | -0.13438604 | 0.828423083 |
| ENSMUSG0000003949 | chr2 | Cdnf | 0.171668684 | 0.828499086 |
| ENSMUSG0000004325 | chr8 | Exoc3l | -0.356852256 | 0.828681419 |
| ENSMUSG0000006331 | chr7 | Usp31 | -0.113553929 | 0.828702232 |
| ENSMUSG0000002508 | chr19 | Afap1l2 | 0.127754847 | 0.828843074 |
| ENSMUSG0000001242 | chr13 | Tmem167 | 0.087316045 | 0.82904664 |
| ENSMUSG0000006301 | chr17 | Msln | 0.403532426 | 0.82904664 |
| ENSMUSG0000003440 | chr4 | Spata6 | -0.064164707 | 0.829351847 |
| ENSMUSG0000002118 | chr12 | Atxn3 | 0.074635719 | 0.829373474 |
| ENSMUSG0000002849 | chr4 | Hacd4 | 0.090847612 | 0.829373474 |
| ENSMUSG0000002126 | chr12 | Wars1 | 0.101403551 | 0.829373474 |
| ENSMUSG0000003394 | chr2 | Mga | -0.101941148 | 0.829473281 |
| ENSMUSG0000002553 | chr5 | Gusb | 0.064544332 | 0.829696505 |
| ENSMUSG0000004524 | chr8 | Med26 | 0.092010011 | 0.829696505 |
| ENSMUSG0000002500 | chr19 | Aldh18a1 | 0.129499383 | 0.829696505 |
| ENSMUSG0000007093 | chr4 | Rraga | 0.097233476 | 0.829744127 |
| ENSMUSG0000008678 | chr7 | Isoc2a | 0.128532481 | 0.829744127 |
| ENSMUSG0000003406 | chr1 | Farp2 | -0.155694274 | 0.829806129 |
| ENSMUSG0000002587 | chr13 | Arl10 | -0.122653996 | 0.829806129 |
| ENSMUSG0000005710 | chr7 | Zfp180 | -0.078573993 | 0.829806129 |
| ENSMUSG0000004399 | chr18 | Pura | -0.081081827 | 0.829806129 |
| ENSMUSG0000006632 | chr4 | Bpnt2 | -0.070441862 | 0.829806129 |
| ENSMUSG0000012158 | chr15 | Gm57852 | 0.067144582 | 0.829806129 |
| ENSMUSG0000002191 | chr14 | Parg | 0.070933617 | 0.829806129 |
| ENSMUSG0000003001 | chr6 | Zfp638 | 0.075340798 | 0.829806129 |
| ENSMUSG0000005395 | chr18 | Adnp2 | 0.076937625 | 0.829806129 |
| ENSMUSG0000003377 | chr5 | Rpap2 | 0.086975553 | 0.829806129 |
| ENSMUSG0000008991 | chr2 | Uckl1 | 0.08774069 | 0.829806129 |
| ENSMUSG0000002892 | chr4 | Fbxo42 | 0.085878952 | 0.829806129 |
| ENSMUSG0000004268 | chr1 | Jph1 | 0.152929301 | 0.829820381 |
| ENSMUSG0000003284 | chr2 | Prr5l | -0.213646338 | 0.829853543 |
| ENSMUSG0000005523 | chr5 | Wdr86 | -0.553100075 | 0.82986296 |
| ENSMUSG0000004751 | chr10 | Tspyl1 | 0.086274132 | 0.82986296 |
| ENSMUSG0000004660 | chr11 | B3gntl1 | 0.14110409 | 0.82986296 |
| ENSMUSG0000000326 | chr7 | Cyth2 | 0.06908783 | 0.830375301 |
| ENSMUSG0000005740 | chr5 | Nsd2 | -0.099887156 | 0.830828065 |
| ENSMUSG0000002132 | chr13 | Zkscan3 | 0.074758675 | 0.831000455 |
| ENSMUSG0000000450 | chr7 | Zfp324 | 0.10174492 | 0.831000455 |
| ENSMUSG0000007908 | chr9 | Ccdc82 | 0.099134721 | 0.831000455 |
| ENSMUSG0000006788 | chr19 | Sptbn2 | -0.118183169 | 0.831413198 |
| ENSMUSG0000003126 | chrX | Gla | -0.138279087 | 0.831413198 |
| ENSMUSG0000004869 | chr15 | Krt90 | -0.413095791 | 0.831413198 |
| ENSMUSG0000002833 | chr4 | Anp32b | 0.084695655 | 0.831413198 |
| ENSMUSG0000003406 | chr16 | Poglut1 | 0.099941228 | 0.831413198 |
| ENSMUSG0000003154 | chr8 | Gins4 | 0.129958123 | 0.831413198 |
| ENSMUSG0000003176 | chr8 | Slc12a3 | -0.493117404 | 0.831427843 |
| ENSMUSG0000002643 | chr1 | Rbbp5 | -0.0635653 | 0.831427843 |
| ENSMUSG0000001875 | chr11 | Zbtb4 | 0.092105124 | 0.831427843 |
| ENSMUSG0000003628 | chr2 | Snapc4 | 0.101584671 | 0.831427843 |

| | | | | |
|---|---|---|---|---|
| ENSMUSG0000002750 | chr3 | Tpd52 | 0.086600773 | 0.831456899 |
| ENSMUSG0000004003 | chr10 | Nup43 | 0.14350624 | 0.831494477 |
| ENSMUSG0000004171 | chrX | Trpc5 | -0.46034091 | 0.831661509 |
| ENSMUSG0000002679 | chr2 | Coq4 | -0.107234953 | 0.831661509 |
| ENSMUSG0000003137 | chrX | Pir | 0.158718047 | 0.831661509 |
| ENSMUSG0000005002 | chr5 | Amz1 | 0.242153302 | 0.831661509 |
| ENSMUSG0000005044 | chr2 | Lypd6 | 0.281620645 | 0.831661509 |
| ENSMUSG0000003011 | chr6 | A2m | -0.445854012 | 0.831676078 |
| ENSMUSG0000003290 | chr3 | Slc16a1 | -0.149936562 | 0.831676078 |
| ENSMUSG0000003869 | chr8 | Taf5l | 0.081609653 | 0.831676078 |
| ENSMUSG0000002223 | chr15 | Ropn1l | 0.465653754 | 0.831676078 |
| ENSMUSG0000004235 | chr8 | Frem3 | 0.517191882 | 0.831676078 |
| ENSMUSG0000003788 | chr7 | Dusp8 | -0.136037639 | 0.831677377 |
| ENSMUSG0000006080 | chr19 | Gstp1 | 0.138238388 | 0.831677377 |
| ENSMUSG0000000456 | chr14 | Arhgef40 | 0.100379573 | 0.831677377 |
| ENSMUSG0000003325 | chr4 | Szt2 | -0.103511327 | 0.831705657 |
| ENSMUSG0000003003 | chr6 | Wdr54 | -0.137507034 | 0.831705657 |
| ENSMUSG0000002830 | chr4 | Rnf20 | -0.049536901 | 0.831705657 |
| ENSMUSG0000000630 | chr1 | Tmbim1 | 0.085169787 | 0.831705657 |
| ENSMUSG0000004210 | chr14 | Uggt2 | 0.120065998 | 0.831705657 |
| ENSMUSG0000001663 | chr15 | Ift27 | 0.10890518 | 0.831705657 |
| ENSMUSG0000007877 | chr11 | Evi2a | 0.228857385 | 0.831705657 |
| ENSMUSG0000005124 | chr5 | Msantd1 | 0.300501794 | 0.831924134 |
| ENSMUSG0000004892 | chr14 | Cdca2 | -0.320883073 | 0.831945013 |
| ENSMUSG0000004573 | chr18 | Adrb2 | -0.182291388 | 0.831964884 |
| ENSMUSG0000002980 | chr6 | Abcg2 | -0.131207493 | 0.831964884 |
| ENSMUSG0000002682 | chr2 | Gpd2 | -0.12062545 | 0.831964884 |
| ENSMUSG0000006259 | chr1 | Armc9 | -0.074430191 | 0.831964884 |
| ENSMUSG0000000210 | chr2 | Acp2 | -0.058925542 | 0.831964884 |
| ENSMUSG0000003055 | chr7 | Pgpep1l | 0.298023194 | 0.831964884 |
| ENSMUSG0000002666 | chr2 | Phyh | 0.099666346 | 0.831964884 |
| ENSMUSG0000003661 | chr3 | Rfxap | 0.098752076 | 0.831964884 |
| ENSMUSG0000006147 | chr7 | Snrpa | 0.120808867 | 0.831964884 |
| ENSMUSG0000000133 | chr4 | Fndc5 | 0.115072402 | 0.831964884 |
| ENSMUSG0000001565 | chr17 | Serac1 | 0.17019743 | 0.831964884 |
| ENSMUSG0000002859 | chr4 | Tnfrsf1b | 0.179724231 | 0.831964884 |
| ENSMUSG0000003665 | chr12 | Colec11 | -0.488392234 | 0.832047307 |
| ENSMUSG0000003241 | chr9 | Xrn1 | -0.113101994 | 0.832047307 |
| ENSMUSG0000002535 | chr10 | Ormdl2 | -0.156217218 | 0.832047307 |
| ENSMUSG0000003355 | chr1 | Fam20b | -0.06363481 | 0.832047307 |
| ENSMUSG0000004807 | chr7 | Tenm4 | 0.135522792 | 0.832047307 |
| ENSMUSG0000006369 | chr19 | Sfxn4 | -0.186555103 | 0.83206237 |
| ENSMUSG0000005029 | chr13 | Foxc1 | -0.288658731 | 0.83206237 |
| ENSMUSG0000002875 | chr4 | Vwa5b1 | -0.13585654 | 0.83206237 |
| ENSMUSG0000002490 | chr19 | Cpt1a | -0.1462594 | 0.83206237 |
| ENSMUSG0000004416 | chr6 | Tnip3 | -0.484901086 | 0.83206237 |
| ENSMUSG0000003248 | chr9 | Kif9 | -0.130224526 | 0.83206237 |
| ENSMUSG0000002210 | chr14 | Gfra2 | -0.107549926 | 0.83206237 |
| ENSMUSG0000011812 | chr19 | Gm50387 | -0.220281449 | 0.83206237 |
| ENSMUSG0000004879 | chr3 | Cyb561d1 | -0.094631414 | 0.83206237 |

| | | | | |
|---|---|---|---|---|
| ENSMUSG0000007341 | chr17 | H2-D1 | -0.103680156 | 0.83206237 |
| ENSMUSG0000003195 | chr8 | Bcar1 | -0.089349416 | 0.83206237 |
| ENSMUSG0000003218 | chr9 | Lysmd2 | -0.081587539 | 0.83206237 |
| ENSMUSG0000002200 | chr14 | Gpalpp1 | -0.062017866 | 0.83206237 |
| ENSMUSG0000000112 | chrX | Araf | -0.051226145 | 0.83206237 |
| ENSMUSG0000000602 | chr7 | Napa | 0.049402497 | 0.83206237 |
| ENSMUSG0000003480 | chr12 | Sos2 | 0.101275439 | 0.83206237 |
| ENSMUSG0000003341 | chr2 | Ctdspl2 | 0.080228437 | 0.83206237 |
| ENSMUSG0000005419 | chr3 | Gon4l | 0.092788363 | 0.83206237 |
| ENSMUSG0000006037 | chr7 | Bckdha | 0.126924208 | 0.83206237 |
| ENSMUSG0000003900 | chr13 | Bmp6 | 0.133464925 | 0.83206237 |
| ENSMUSG0000003923 | chr10 | Stx11 | 0.217553296 | 0.83206237 |
| ENSMUSG0000003212 | chr9 | Hmbs | -0.089151889 | 0.832354952 |
| ENSMUSG0000000309 | chr7 | Ppp5c | 0.067542093 | 0.832413897 |
| ENSMUSG0000007527 | chr2 | Ift70a1 | -0.17451674 | 0.832760392 |
| ENSMUSG0000002177 | chr14 | Nkiras1 | -0.084350083 | 0.832760392 |
| ENSMUSG0000005037 | chrX | Septin6 | 0.101501021 | 0.832760392 |
| ENSMUSG0000010787 | chr11 | 4933427D14Rik | -0.236476319 | 0.832768474 |
| ENSMUSG0000004467 | chr8 | Zfp612 | -0.081135648 | 0.832857377 |
| ENSMUSG0000002457 | chr18 | Csnk1a1 | 0.053632399 | 0.832989578 |
| ENSMUSG0000003157 | chr8 | Tti2 | 0.073438266 | 0.832989578 |
| ENSMUSG0000005229 | chr11 | Cdc42se2 | 0.079121953 | 0.832989578 |
| ENSMUSG0000001648 | chr1 | Cr1l | 0.082810575 | 0.832992369 |
| ENSMUSG0000002901 | chr4 | Clcn6 | -0.133151069 | 0.833061726 |
| ENSMUSG0000007208 | chr17 | Ccnf | -0.301906196 | 0.833061726 |
| ENSMUSG0000000072 | chr11 | Cryba1 | -0.386908529 | 0.833061726 |
| ENSMUSG0000003411 | chr5 | Kctd7 | -0.096716659 | 0.833061726 |
| ENSMUSG0000002651 | chr1 | Trp53bp2 | 0.078580277 | 0.833061726 |
| ENSMUSG0000005638 | chr7 | AI987944 | 0.103549104 | 0.833061726 |
| ENSMUSG0000004631 | chr4 | Myorg | 0.099762405 | 0.833061726 |
| ENSMUSG0000005149 | chr6 | Zfp786 | 0.291363667 | 0.833061726 |
| ENSMUSG0000002535 | chr10 | Gdf11 | -0.167816424 | 0.833168353 |
| ENSMUSG0000004052 | chr11 | Milr1 | -0.324642465 | 0.833168353 |
| ENSMUSG0000005982 | chr7 | Dbp | -0.766931322 | 0.833280181 |
| ENSMUSG0000003554 | chr7 | Leng8 | -0.138507599 | 0.833280181 |
| ENSMUSG0000003608 | chr1 | Zranb3 | -0.15445662 | 0.833280181 |
| ENSMUSG0000009233 | chr7 | Zfp977 | -0.173076654 | 0.833280181 |
| ENSMUSG0000000238 | chr4 | Bmp8b | -0.54587271 | 0.833280181 |
| ENSMUSG0000003416 | chrX | Ogt | -0.105697532 | 0.833280181 |
| ENSMUSG0000005167 | chr5 | Dcun1d4 | -0.09903047 | 0.833280181 |
| ENSMUSG0000002795 | chr3 | Efna1 | -0.128225929 | 0.833280181 |
| ENSMUSG0000003166 | chr8 | Rbl2 | -0.092820977 | 0.833280181 |
| ENSMUSG0000002810 | chr3 | Polr3gl | -0.107727745 | 0.833280181 |
| ENSMUSG0000007428 | chr7 | Zfp109 | -0.122983408 | 0.833280181 |
| ENSMUSG0000001441 | chr7 | Hps5 | -0.093942805 | 0.833280181 |
| ENSMUSG0000006953 | chr10 | Scyl2 | -0.075877612 | 0.833280181 |
| ENSMUSG0000005591 | chr6 | Tmem150a | -0.086971393 | 0.833280181 |
| ENSMUSG0000003252 | chr9 | Pccb | -0.064117551 | 0.833280181 |
| ENSMUSG0000005773 | chr2 | Sptan1 | 0.061518284 | 0.833280181 |
| ENSMUSG0000002736 | chr2 | Usp8 | 0.056753638 | 0.833280181 |

| | | | | |
|---|---|---|---|---|
| ENSMUSG0000003781 | chr13 | Fbxw17 | 0.085955109 | 0.833280181 |
| ENSMUSG0000004817 | chr15 | Asb8 | 0.073198262 | 0.833280181 |
| ENSMUSG0000003065 | chr7 | Nucb2 | 0.075870936 | 0.833280181 |
| ENSMUSG0000002943 | chr5 | Sfswap | 0.087329131 | 0.833280181 |
| ENSMUSG0000002691 | chr2 | Psmd14 | 0.099838889 | 0.833280181 |
| ENSMUSG0000003457 | chr12 | Daam1 | 0.085140142 | 0.833280181 |
| ENSMUSG0000004559 | chr7 | Zfp553 | 0.088495578 | 0.833280181 |
| ENSMUSG0000006067 | chr19 | Plaat3 | 0.099614736 | 0.833280181 |
| ENSMUSG0000003152 | chr8 | Aga | 0.107075842 | 0.833280181 |
| ENSMUSG0000007377 | chr4 | Kti12 | 0.128150515 | 0.833280181 |
| ENSMUSG0000004091 | chr19 | Fbxw4 | 0.105579208 | 0.833280181 |
| ENSMUSG0000008105 | chr3 | H3c15 | 0.196192834 | 0.833280181 |
| ENSMUSG0000006388 | chr18 | Crem | 0.139011518 | 0.833280181 |
| ENSMUSG0000005095 | chr10 | Gja1 | 0.12885236 | 0.833280181 |
| ENSMUSG0000007452 | chr2 | Gm14296 | 0.326120092 | 0.833280181 |
| ENSMUSG0000004014 | chr7 | Hmx3 | 0.142921267 | 0.833280181 |
| ENSMUSG0000005609 | chr6 | St3gal5 | 0.13912261 | 0.833280181 |
| ENSMUSG0000009619 | chr12 | Ptrhd1 | 0.188967896 | 0.833280181 |
| ENSMUSG0000002052 | chr11 | Znhit3 | 0.13872933 | 0.833303338 |
| ENSMUSG0000002492 | chr19 | Smarca2 | -0.123983629 | 0.833491907 |
| ENSMUSG0000002073 | chr11 | Nup85 | 0.077904046 | 0.833491907 |
| ENSMUSG0000002524 | chr9 | Sacm1l | 0.072859225 | 0.833491907 |
| ENSMUSG0000005518 | chr1 | Fam72a | 0.278065291 | 0.833491907 |
| ENSMUSG0000003768 | chr9 | Esyt3 | 0.250285631 | 0.833491907 |
| ENSMUSG0000007897 | chr11 | Dnaaf10 | 0.115466457 | 0.833700463 |
| ENSMUSG0000004100 | chr4 | Trim62 | -0.112819134 | 0.833845878 |
| ENSMUSG0000004243 | chr18 | Zfp532 | 0.076193216 | 0.83393182 |
| ENSMUSG0000002728 | chr2 | Slx4ip | -0.130198341 | 0.833951426 |
| ENSMUSG0000002177 | chr14 | Comtd1 | -0.29062503 | 0.833987293 |
| ENSMUSG0000007960 | chr18 | Stard6 | -0.246171068 | 0.834193027 |
| ENSMUSG0000001858 | chr11 | G3bp1 | 0.077134805 | 0.8342244 |
| ENSMUSG0000002393 | chr17 | Spats1 | 0.294938324 | 0.834267984 |
| ENSMUSG0000003345 | chr7 | Fan1 | 0.127335634 | 0.834332064 |
| ENSMUSG0000004610 | chr1 | Mcmdc2 | -0.473185341 | 0.834467067 |
| ENSMUSG0000003390 | chr2 | Mapkbp1 | 0.112556636 | 0.834467067 |
| ENSMUSG0000003166 | chr8 | Nkd1 | 0.141599212 | 0.834467067 |
| ENSMUSG0000008513 | chrX | Sertm2 | -0.157806184 | 0.834658567 |
| ENSMUSG0000002490 | chr19 | Peli3 | -0.112804035 | 0.834658567 |
| ENSMUSG0000005205 | chr2 | Zfp217 | -0.102637137 | 0.834658567 |
| ENSMUSG0000002147 | chr13 | Ercc6l2 | -0.070140298 | 0.834658567 |
| ENSMUSG0000002180 | chr14 | Nudt13 | 0.089382742 | 0.834710938 |
| ENSMUSG0000005511 | chr7 | Bmal1 | 0.640532025 | 0.834710938 |
| ENSMUSG0000007425 | chr9 | Gramd2a | -0.414184735 | 0.834781136 |
| ENSMUSG0000003010 | chr6 | Slc6a13 | -0.50355097 | 0.834781136 |
| ENSMUSG0000005341 | chr15 | Cbx7 | -0.101586603 | 0.834781136 |
| ENSMUSG0000006743 | chr17 | Zfp763 | -0.133531368 | 0.834781136 |
| ENSMUSG0000001772 | chr2 | Pigt | -0.103730671 | 0.834781136 |
| ENSMUSG0000002127 | chr12 | Zfp839 | -0.093186672 | 0.834781136 |
| ENSMUSG0000002260 | chr16 | Cep97 | -0.098363869 | 0.834781136 |
| ENSMUSG0000003203 | chr9 | Ets1 | -0.077385657 | 0.834781136 |

| | | | | |
|---|---|---|---|---|
| ENSMUSG0000002964 | chr5 | Usp12 | -0.079067795 | 0.834781136 |
| ENSMUSG0000006376 | chr10 | Rnf217 | -0.07767979 | 0.834781136 |
| ENSMUSG0000000947 | chr13 | Tnpo1 | 0.058733603 | 0.834781136 |
| ENSMUSG0000002061 | chr11 | Fam20a | 0.096733553 | 0.834781136 |
| ENSMUSG0000000088 | chrX | Dlg3 | 0.087167541 | 0.834781136 |
| ENSMUSG0000004698 | chr18 | Tshz1 | 0.104565191 | 0.834781136 |
| ENSMUSG0000002459 | chr18 | Slc12a2 | 0.095971476 | 0.834781136 |
| ENSMUSG0000007398 | chr4 | Ggh | 0.093228205 | 0.834781136 |
| ENSMUSG0000002074 | chr11 | Mif4gd | 0.117912314 | 0.834781136 |
| ENSMUSG0000003916 | chr9 | Cmc1 | 0.116756137 | 0.834781136 |
| ENSMUSG0000002516 | chr11 | Ogfod3 | 0.114771513 | 0.834781136 |
| ENSMUSG0000002122 | chr12 | Dpf3 | 0.135419819 | 0.834781136 |
| ENSMUSG0000002479 | chr4 | Tnfrsf25 | -0.219406332 | 0.835008999 |
| ENSMUSG0000003153 | chr8 | Mrps31 | 0.087028158 | 0.835056486 |
| ENSMUSG0000005425 | chr5 | Msi1 | -0.143549281 | 0.835228614 |
| ENSMUSG0000001866 | chr11 | Cog1 | -0.082743408 | 0.835228614 |
| ENSMUSG0000002210 | chr14 | Rb1 | -0.074291688 | 0.835228614 |
| ENSMUSG0000001731 | chr11 | Mpp2 | -0.120203043 | 0.835683206 |
| ENSMUSG0000002775 | chr3 | Supt20 | -0.066339944 | 0.835683206 |
| ENSMUSG0000003258 | chr9 | Rbm5 | 0.080231655 | 0.835683206 |
| ENSMUSG0000002146 | chr13 | Fancc | 0.12332562 | 0.835683206 |
| ENSMUSG0000007063 | chr5 | Lrrc8b | -0.16609988 | 0.835751571 |
| ENSMUSG0000006397 | chr2 | Nr6a1 | -0.143846632 | 0.835751571 |
| ENSMUSG0000003059 | chr7 | Fam98c | -0.139545973 | 0.835751571 |
| ENSMUSG0000005669 | chr6 | Elmod3 | 0.106810389 | 0.835751571 |
| ENSMUSG0000005106 | chr16 | Mb21d2 | 0.103065493 | 0.835751571 |
| ENSMUSG0000009688 | chr15 | Shisa8 | 0.34494253 | 0.835751571 |
| ENSMUSG0000002515 | chr11 | Gps1 | 0.093419776 | 0.835896554 |
| ENSMUSG0000002559 | chrX | Klhl4 | -0.109493808 | 0.835970167 |
| ENSMUSG0000003428 | chr11 | Nipsnap1 | -0.114291177 | 0.836021805 |
| ENSMUSG0000004850 | chr14 | Duxbl1 | -0.176179524 | 0.836034498 |
| ENSMUSG0000000678 | chr11 | Cnp | 0.120669902 | 0.836153688 |
| ENSMUSG0000006697 | chr7 | Bub3 | 0.087899995 | 0.836286935 |
| ENSMUSG0000000598 | chr16 | 1700037C18Rik | 0.14722885 | 0.836428082 |
| ENSMUSG0000003640 | chr5 | Cep135 | -0.134778042 | 0.836728617 |
| ENSMUSG0000005947 | chr11 | Mbtd1 | 0.090857736 | 0.836728617 |
| ENSMUSG0000002515 | chr11 | Rfng | -0.073044925 | 0.836765886 |
| ENSMUSG0000002904 | chr4 | Pex10 | 0.101456373 | 0.836765886 |
| ENSMUSG0000003264 | chr6 | Gpr19 | -0.133179001 | 0.837073224 |
| ENSMUSG0000003281 | chr7 | Arap1 | -0.073758398 | 0.837073224 |
| ENSMUSG0000003602 | chr17 | Tmem63b | -0.073455825 | 0.837073224 |
| ENSMUSG0000002190 | chr14 | Btd | 0.07175147 | 0.837073224 |
| ENSMUSG0000002030 | chr11 | Asb3 | 0.112502575 | 0.837073224 |
| ENSMUSG0000005037 | chr19 | Ch25h | 0.381432934 | 0.837073224 |
| ENSMUSG0000001973 | chr8 | Calr3 | 0.272318289 | 0.837073224 |
| ENSMUSG0000001478 | chr8 | Slc9a5 | -0.196712954 | 0.83714001 |
| ENSMUSG0000000964 | chr4 | Dmap1 | 0.089249753 | 0.83714001 |
| ENSMUSG0000003102 | chr7 | Trim66 | 0.238407596 | 0.837186304 |
| ENSMUSG0000004256 | chr11 | Dhrs7b | 0.110037739 | 0.837268417 |
| ENSMUSG0000004396 | chr7 | Orai3 | 0.104842562 | 0.837337319 |

| | | | | |
|---|---|---|---|---|
| ENSMUSG0000006874 | chr2 | Cry2 | -0.134988538 | 0.837356029 |
| ENSMUSG0000004220 | chr17 | Tbc1d22b | -0.095212459 | 0.837459368 |
| ENSMUSG0000003263 | chr7 | Atxn2l | -0.089442472 | 0.837459368 |
| ENSMUSG0000001517 | chr19 | Nolc1 | -0.05304547 | 0.837459368 |
| ENSMUSG0000003249 | chr9 | Lrrfip2 | 0.053934463 | 0.837459368 |
| ENSMUSG0000000282 | chr9 | Qtrt1 | 0.100333278 | 0.837459368 |
| ENSMUSG0000002075 | chr11 | Itgb4 | -0.181270517 | 0.837562739 |
| ENSMUSG0000004417 | chr18 | Spink10 | 0.395431544 | 0.837581764 |
| ENSMUSG0000000876 | chr3 | Man1a2 | 0.052862793 | 0.837810806 |
| ENSMUSG0000002193 | chr14 | Pspc1 | 0.084458605 | 0.837810806 |
| ENSMUSG0000002944 | chr5 | Psph | 0.146160926 | 0.837853515 |
| ENSMUSG0000002248 | chr15 | Hoxc5 | -0.217235717 | 0.837879636 |
| ENSMUSG0000004638 | chr15 | Jrk | -0.134646188 | 0.837879636 |
| ENSMUSG0000003149 | chr8 | Cd209d | -0.458273143 | 0.837879636 |
| ENSMUSG0000003202 | chr9 | Crtam | -0.44197655 | 0.837879636 |
| ENSMUSG0000002135 | chr13 | Exoc2 | -0.070597087 | 0.837879636 |
| ENSMUSG0000002015 | chr11 | Actr2 | -0.055439857 | 0.837879636 |
| ENSMUSG0000000436 | chr1 | Cul3 | 0.063781906 | 0.837879636 |
| ENSMUSG0000003065 | chr7 | Xylt1 | 0.104794227 | 0.837879636 |
| ENSMUSG0000002162 | chr13 | Cd180 | 0.272228309 | 0.837879636 |
| ENSMUSG0000006363 | chr12 | Sox11 | -0.211456089 | 0.837935201 |
| ENSMUSG0000007820 | chr2 | Tmem203 | -0.188932454 | 0.838064233 |
| ENSMUSG0000007174 | chrX | Styx-ps | -0.463262328 | 0.838064233 |
| ENSMUSG0000004436 | chr11 | Slc16a13 | -0.177079494 | 0.838203714 |
| ENSMUSG0000002742 | chr2 | Bfsp1 | -0.227660737 | 0.838203714 |
| ENSMUSG0000003316 | chr14 | Dis3 | 0.076624233 | 0.838203714 |
| ENSMUSG0000001728 | chr11 | Glod4 | 0.111585941 | 0.838203714 |
| ENSMUSG0000000360 | chr2 | Aven | 0.107468952 | 0.838203714 |
| ENSMUSG0000002092 | chr11 | Lsm12 | 0.080220532 | 0.838220541 |
| ENSMUSG0000005102 | chr13 | Serpinb1b | -0.39482886 | 0.838331589 |
| ENSMUSG0000000962 | chr2 | Vav2 | 0.086031805 | 0.838331589 |
| ENSMUSG0000005259 | chr12 | Adam17 | 0.099508642 | 0.838394274 |
| ENSMUSG0000005706 | chr13 | Ero1b | -0.117493874 | 0.838533533 |
| ENSMUSG0000003693 | chr7 | Nfkbid | -0.256425627 | 0.838533533 |
| ENSMUSG0000002183 | chr14 | Txndc16 | 0.172655782 | 0.838533533 |
| ENSMUSG0000004225 | chr17 | Ptchd4 | 0.385942464 | 0.838533533 |
| ENSMUSG0000003075 | chr7 | Kdm8 | -0.134262053 | 0.838625608 |
| ENSMUSG0000000113 | chr1 | Cnnm3 | -0.078261721 | 0.838625608 |
| ENSMUSG0000007948 | chrX | Nhsl2 | -0.132179431 | 0.838713837 |
| ENSMUSG0000003643 | chr3 | Siah2 | 0.101233766 | 0.838713837 |
| ENSMUSG0000004811 | chr12 | Arid4a | 0.105399687 | 0.838713837 |
| ENSMUSG0000001554 | chr11 | Nat9 | 0.13794175 | 0.838713837 |
| ENSMUSG0000002088 | chr11 | Dvl2 | 0.117338176 | 0.838713837 |
| ENSMUSG0000003753 | chr3 | Mrpl47 | 0.106893784 | 0.839039198 |
| ENSMUSG0000005601 | chr8 | Zfp709 | -0.118784271 | 0.839075011 |
| ENSMUSG0000000685 | chr18 | Tmco6 | -0.123667114 | 0.839403562 |
| ENSMUSG0000002376 | chr11 | Sfi1 | -0.1019822 | 0.839467971 |
| ENSMUSG0000004948 | chr4 | Tmem67 | -0.084105486 | 0.839467971 |
| ENSMUSG0000002764 | chr2 | Rpn2 | 0.078140854 | 0.839514531 |
| ENSMUSG0000004525 | chr7 | Zfp688 | 0.150178984 | 0.839514531 |

| | | | | |
|---|---|---|---|---|
| ENSMUSG0000004153 | chr4 | Ago1 | -0.07089775 | 0.839671297 |
| ENSMUSG0000004682 | chr10 | Slc39a3 | 0.086736007 | 0.839671297 |
| ENSMUSG0000002279 | chr16 | B4galt4 | -0.123007214 | 0.839692671 |
| ENSMUSG0000006737 | chrX | Tspan6 | -0.100334743 | 0.839692671 |
| ENSMUSG0000003336 | chr1 | Usp37 | -0.078346801 | 0.839692671 |
| ENSMUSG0000002247 | chr15 | Polr3h | -0.080557123 | 0.839692671 |
| ENSMUSG0000002158 | chr13 | Cast | 0.052389922 | 0.839692671 |
| ENSMUSG0000006103 | chr10 | Rrp1 | 0.055937998 | 0.839692671 |
| ENSMUSG0000003084 | chr7 | Fgfr2 | 0.077029359 | 0.839692671 |
| ENSMUSG0000003030 | chr6 | Atp2b2 | -0.141880279 | 0.839727315 |
| ENSMUSG0000002649 | chr1 | Kif26b | -0.122748478 | 0.839727315 |
| ENSMUSG0000003650 | chr18 | Fam13b | 0.082563998 | 0.839727315 |
| ENSMUSG0000003016 | chr6 | Olr1 | -0.424524005 | 0.839789682 |
| ENSMUSG0000002442 | chr18 | Spry4 | -0.110985237 | 0.839789682 |
| ENSMUSG0000004193 | chr5 | Fam222a | -0.198188732 | 0.839789682 |
| ENSMUSG0000003968 | chr2 | Zer1 | -0.087588352 | 0.839789682 |
| ENSMUSG0000005759 | chr11 | Arl16 | 0.113581675 | 0.839789682 |
| ENSMUSG0000003521 | chr10 | Lsm7 | 0.121751407 | 0.839789682 |
| ENSMUSG0000001539 | chr13 | Cd83 | 0.147124763 | 0.839789682 |
| ENSMUSG0000001771 | chr11 | Birc5 | -0.410808344 | 0.839855351 |
| ENSMUSG0000004271 | chr5 | Naa25 | -0.069901581 | 0.840534548 |
| ENSMUSG0000002953 | chr5 | Triap1 | 0.14041914 | 0.840534548 |
| ENSMUSG0000001005 | chr9 | Hyal1 | 0.099025495 | 0.840832324 |
| ENSMUSG0000010309 | chr18 | Pcdha5 | 0.222380429 | 0.84103709 |
| ENSMUSG0000002829 | chr4 | Akirin2 | 0.096761592 | 0.84103709 |
| ENSMUSG0000002074 | chr11 | Slc25a19 | 0.10859857 | 0.84103709 |
| ENSMUSG0000003354 | chr6 | Arhgef5 | 0.096800765 | 0.84103709 |
| ENSMUSG0000003877 | chr9 | Vill | -0.235760463 | 0.841130452 |
| ENSMUSG0000003619 | chrX | Arhgap36 | -0.462883724 | 0.841130452 |
| ENSMUSG0000003115 | chrX | Slc35a2 | -0.099629795 | 0.841130452 |
| ENSMUSG0000002893 | chr4 | Acot7 | -0.105825752 | 0.841138035 |
| ENSMUSG0000004703 | chr9 | Zfp445 | -0.090378389 | 0.841138035 |
| ENSMUSG0000002442 | chr18 | Impact | -0.07358173 | 0.841138035 |
| ENSMUSG0000005213 | chr4 | Foxo6 | -0.146503721 | 0.841239012 |
| ENSMUSG0000003807 | chr4 | Cntln | -0.11989902 | 0.841239012 |
| ENSMUSG0000004914 | chr15 | Plcxd3 | -0.198064702 | 0.841239012 |
| ENSMUSG0000002841 | chr4 | Slc44a1 | -0.091624183 | 0.841239012 |
| ENSMUSG0000003649 | chr10 | Eea1 | -0.076925406 | 0.841239012 |
| ENSMUSG0000004521 | chr1 | Vcpip1 | -0.071023562 | 0.841239012 |
| ENSMUSG0000003947 | chr8 | Micu3 | -0.064540166 | 0.841239012 |
| ENSMUSG0000003338 | chr11 | Arhgap44 | 0.061069587 | 0.841239012 |
| ENSMUSG0000003098 | chr7 | Vps35l | 0.06397566 | 0.841239012 |
| ENSMUSG0000000392 | chr10 | Tfam | 0.082989127 | 0.841239012 |
| ENSMUSG0000003522 | chr7 | Spcs2 | 0.114977045 | 0.841239012 |
| ENSMUSG0000003737 | chr10 | Enpp1 | 0.082322277 | 0.841239012 |
| ENSMUSG0000000303 | chr8 | Rab8a | 0.100224741 | 0.841239012 |
| ENSMUSG0000001596 | chr14 | Cacna1d | 0.130690832 | 0.841239012 |
| ENSMUSG0000004980 | chr11 | Arhgap23 | 0.102487263 | 0.841239012 |
| ENSMUSG0000005170 | chr9 | Senp8 | 0.160164865 | 0.841239012 |
| ENSMUSG0000006057 | chr4 | Mfap2 | -0.128801423 | 0.841256648 |

| | | | | |
|---|---|---|---|---|
| ENSMUSG0000003577 | chr10 | Kiss1r | -0.175329372 | 0.841256648 |
| ENSMUSG0000004563 | chr7 | Zfp629 | 0.09392071 | 0.841256648 |
| ENSMUSG0000006406 | chr10 | Ipcef1 | -0.338498949 | 0.841810501 |
| ENSMUSG0000003772 | chr5 | Tmem33 | 0.084129276 | 0.841810501 |
| ENSMUSG0000002962 | chr5 | Arpc1a | 0.082242922 | 0.841810501 |
| ENSMUSG0000001455 | chr6 | Dguok | 0.090051754 | 0.841810501 |
| ENSMUSG0000006012 | chr12 | Gemin2 | 0.143426004 | 0.841810501 |
| ENSMUSG0000002092 | chr11 | Higd1b | -0.422216058 | 0.842114314 |
| ENSMUSG0000003157 | chr8 | Kcnu1 | -0.271184519 | 0.842114314 |
| ENSMUSG0000005375 | chr14 | Chd8 | -0.091751826 | 0.842114314 |
| ENSMUSG0000003656 | chr8 | Ndrg4 | -0.098660349 | 0.842114314 |
| ENSMUSG0000002111 | chr12 | Papola | 0.077868295 | 0.842114314 |
| ENSMUSG0000003710 | chr17 | Socs5 | 0.072529545 | 0.842114314 |
| ENSMUSG0000002735 | chr2 | Chgb | -0.158080692 | 0.842286291 |
| ENSMUSG0000002215 | chr15 | Ttc33 | 0.075369775 | 0.842286291 |
| ENSMUSG0000002173 | chr13 | Mrps30 | 0.095098262 | 0.842286291 |
| ENSMUSG0000003410 | chr8 | Meak7 | 0.094856442 | 0.842359135 |
| ENSMUSG0000004974 | chr2 | Arhgap15 | -0.111499623 | 0.842386146 |
| ENSMUSG0000001080 | chr11 | Gabra1 | 0.449194877 | 0.842386146 |
| ENSMUSG0000005329 | chr5 | Pom121 | -0.089170795 | 0.842484589 |
| ENSMUSG0000001828 | chr11 | Spag7 | 0.070904837 | 0.842605409 |
| ENSMUSG0000002960 | chr5 | Dtx1 | -0.112610809 | 0.842890631 |
| ENSMUSG0000001820 | chr2 | Stk4 | 0.059781336 | 0.842929099 |
| ENSMUSG0000003618 | chr8 | Gatad2a | 0.072253401 | 0.842929099 |
| ENSMUSG0000004004 | chr4 | Orc3 | 0.072004617 | 0.842929099 |
| ENSMUSG0000002601 | chr1 | Wdr12 | 0.080357166 | 0.842929099 |
| ENSMUSG0000002801 | chr3 | Ints12 | 0.086771468 | 0.842929099 |
| ENSMUSG0000001242 | chr5 | Steap4 | -0.161095716 | 0.842944311 |
| ENSMUSG0000003540 | chr4 | Kank4 | -0.146389231 | 0.842944311 |
| ENSMUSG0000003982 | chr10 | Myl6b | -0.194481314 | 0.842944311 |
| ENSMUSG0000006207 | chr17 | Qki | 0.067436786 | 0.842944311 |
| ENSMUSG0000002917 | chr5 | Zcchc4 | 0.072939898 | 0.842944311 |
| ENSMUSG0000002725 | chr2 | Arfgap2 | 0.071697082 | 0.842944311 |
| ENSMUSG0000002016 | chr11 | Meis1 | 0.090716516 | 0.842944311 |
| ENSMUSG0000004046 | chr5 | Gtpbp10 | 0.087556126 | 0.842944311 |
| ENSMUSG0000003294 | chr19 | Rasgrp2 | 0.111039799 | 0.842944311 |
| ENSMUSG0000003255 | chr9 | Trf | 0.166593965 | 0.842944311 |
| ENSMUSG0000003092 | chr7 | Eri2 | 0.127475754 | 0.842944311 |
| ENSMUSG0000009671 | chr10 | Zfp781b | 0.325511852 | 0.842944311 |
| ENSMUSG0000007899 | chr13 | Zfp456 | 0.354909798 | 0.842944311 |
| ENSMUSG0000004629 | chr8 | Ankle1 | -0.468460719 | 0.843031622 |
| ENSMUSG0000001913 | chr8 | Isyna1 | -0.14234154 | 0.843033997 |
| ENSMUSG0000006909 | chr3 | Pde7a | -0.096704768 | 0.843033997 |
| ENSMUSG0000003593 | chr13 | Aldh5a1 | -0.088976328 | 0.843033997 |
| ENSMUSG0000001594 | chr3 | Bola1 | 0.146302439 | 0.843033997 |
| ENSMUSG0000007211 | chr14 | Ang | -0.185629161 | 0.843135736 |
| ENSMUSG0000002199 | chr14 | Spata13 | 0.092199482 | 0.843135736 |
| ENSMUSG0000002214 | chr15 | Nup155 | 0.078549878 | 0.843234553 |
| ENSMUSG0000003663 | chr7 | Mag | -0.517240406 | 0.843273853 |
| ENSMUSG0000009684 | chr17 | Tmem151b | -0.158582156 | 0.843289925 |

| | | | | |
|---|---|---|---|---|
| ENSMUSG0000000229 | chr5 | Dbf4 | -0.234847778 | 0.843532576 |
| ENSMUSG0000004353 | chr19 | Sorcs1 | -0.132230248 | 0.843621889 |
| ENSMUSG0000005472 | chr5 | Lrrc8c | -0.11172423 | 0.843621889 |
| ENSMUSG0000004198 | chr9 | Elmod1 | 0.104873147 | 0.843621889 |
| ENSMUSG0000011840 | chr1 | Gpr52 | 0.30218302 | 0.843621889 |
| ENSMUSG0000001797 | chr6 | Cadps2 | -0.13527157 | 0.843712173 |
| ENSMUSG0000001933 | chr3 | Zfp687 | -0.101466776 | 0.843712173 |
| ENSMUSG0000007344 | chr17 | Gm10509 | -0.100422873 | 0.843712173 |
| ENSMUSG0000004833 | chr7 | Ric3 | -0.056659578 | 0.843712173 |
| ENSMUSG0000003031 | chr6 | Vgll4 | 0.055243601 | 0.843712173 |
| ENSMUSG0000002587 | chr13 | 4833439L19Rik | 0.076839213 | 0.843712173 |
| ENSMUSG0000000299 | chr12 | Hbp1 | 0.113641763 | 0.843712173 |
| ENSMUSG0000003515 | chr11 | Ap2b1 | -0.069432591 | 0.843746681 |
| ENSMUSG0000002161 | chr13 | Tert | -0.226729657 | 0.843874778 |
| ENSMUSG0000002222 | chr14 | Adcy4 | -0.172864684 | 0.843874778 |
| ENSMUSG0000003924 | chr1 | Lyplal1 | -0.129040654 | 0.843874778 |
| ENSMUSG0000005888 | chr7 | Deaf1 | -0.085542671 | 0.843874778 |
| ENSMUSG0000004236 | chr13 | Snx18 | 0.070153408 | 0.843874778 |
| ENSMUSG0000004073 | chr7 | Ppp1r13l | 0.116981753 | 0.843874778 |
| ENSMUSG0000002968 | chr6 | Ezh2 | 0.140414956 | 0.843874778 |
| ENSMUSG0000002898 | chr4 | H6pd | 0.117432843 | 0.843874778 |
| ENSMUSG0000003035 | chr6 | Tspan11 | 0.200338882 | 0.843874778 |
| ENSMUSG0000004671 | chr8 | Foxc2 | 0.26589252 | 0.843874778 |
| ENSMUSG0000009366 | chr6 | Eif4e3 | 0.069291764 | 0.844156623 |
| ENSMUSG0000009991 | chr1 | Gm28551 | -0.409624399 | 0.844363081 |
| ENSMUSG0000003921 | chr17 | Srrm2 | -0.131521969 | 0.844531958 |
| ENSMUSG0000005239 | chr7 | Mogat2 | -0.671832804 | 0.844561018 |
| ENSMUSG0000007313 | chrX | Tmem185a | -0.066306126 | 0.844561018 |
| ENSMUSG0000002960 | chr5 | Iqcd | -0.220666961 | 0.845164569 |
| ENSMUSG0000002840 | chr4 | Smu1 | 0.0855582 | 0.845197353 |
| ENSMUSG0000002341 | chr14 | Nfatc4 | -0.071680871 | 0.845209018 |
| ENSMUSG0000003830 | chr1 | Spats2l | 0.092914572 | 0.84522284 |
| ENSMUSG0000003670 | chr1 | Cab39 | -0.053342594 | 0.845304809 |
| ENSMUSG0000001908 | chrX | Dnase1l1 | -0.116279838 | 0.845306785 |
| ENSMUSG0000000652 | chr14 | Stimate | -0.077717955 | 0.845306785 |
| ENSMUSG0000010994 | chr2 | Brd3os | -0.090908322 | 0.845306785 |
| ENSMUSG0000002793 | chr3 | Ints3 | -0.062771832 | 0.845306785 |
| ENSMUSG0000000141 | chr3 | Glmp | 0.075269458 | 0.845306785 |
| ENSMUSG0000004972 | chr7 | Zfp668 | 0.104113999 | 0.845306785 |
| ENSMUSG0000002550 | chr7 | Pidd1 | 0.219696255 | 0.845306785 |
| ENSMUSG0000002797 | chr3 | Prss12 | 0.302716406 | 0.845306785 |
| ENSMUSG0000005927 | chr11 | Naa38 | 0.166319829 | 0.84531592 |
| ENSMUSG0000002666 | chr1 | Uhmk1 | -0.069366384 | 0.845429036 |
| ENSMUSG0000004306 | chr9 | Dpy19l1 | -0.068493552 | 0.845622878 |
| ENSMUSG0000004507 | chr4 | E130308A19Rik | -0.13413915 | 0.845855683 |
| ENSMUSG0000004018 | chr4 | Ankrd6 | 0.08168968 | 0.846124976 |
| ENSMUSG0000002251 | chr16 | Nudt16l1 | 0.101824891 | 0.846209815 |
| ENSMUSG0000005137 | chr11 | Kif18b | -0.484162017 | 0.846364073 |
| ENSMUSG0000005985 | chr8 | Hydin | -0.38185818 | 0.846364073 |
| ENSMUSG0000004270 | chr1 | Shcbp1l | -0.533535265 | 0.846364073 |

| | | | | |
|---|---|---|---|---|
| ENSMUSG000001093C | chr7 | Smim38 | -0.577365589 | 0.846364073 |
| ENSMUSG0000001992 | chr10 | Dcn | 0.11428193 | 0.846364073 |
| ENSMUSG0000006317 | chr6 | Rps4l | 0.141812317 | 0.846364073 |
| ENSMUSG0000003404 | chr5 | Galnt17 | -0.100563624 | 0.846529705 |
| ENSMUSG000000259C | chr1 | Rb1cc1 | 0.080435587 | 0.846529705 |
| ENSMUSG0000003023 | chr6 | Aebp2 | 0.066566015 | 0.84677247 |
| ENSMUSG0000007342 | chr10 | Gm4924 | 0.14663049 | 0.84677247 |
| ENSMUSG0000002537 | chr11 | Baiap2 | -0.136562548 | 0.84679886 |
| ENSMUSG0000001865 | chr11 | Ikzf1 | -0.206281223 | 0.846964627 |
| ENSMUSG0000007363 | chr18 | Rab18 | 0.075695607 | 0.846964627 |
| ENSMUSG0000002692 | chr2 | Egfl7 | -0.110622102 | 0.846966101 |
| ENSMUSG0000002254 | chr16 | Rogdi | -0.076282799 | 0.846966101 |
| ENSMUSG0000000387 | chr7 | Bax | 0.103094029 | 0.846966101 |
| ENSMUSG0000000643 | chr19 | Neurl1a | -0.136369398 | 0.847134231 |
| ENSMUSG0000009015 | chr9 | Acad11 | -0.120066288 | 0.847134231 |
| ENSMUSG0000003666 | chr6 | Tcaf1 | -0.065161044 | 0.847134231 |
| ENSMUSG0000002126 | chr12 | Yy1 | -0.043269878 | 0.847134231 |
| ENSMUSG0000000155 | chr11 | Klhl10 | 0.335722238 | 0.847134231 |
| ENSMUSG0000002129 | chr12 | Gpr132 | -0.703376811 | 0.847204575 |
| ENSMUSG0000003748 | chr12 | Asxl2 | -0.12107562 | 0.847318503 |
| ENSMUSG000000448C | chr11 | Zfp354c | -0.085856467 | 0.847318503 |
| ENSMUSG0000002239 | chr15 | Zc3h7b | 0.06357696 | 0.847318503 |
| ENSMUSG0000003673 | chr9 | Oxsr1 | 0.062232966 | 0.847318503 |
| ENSMUSG0000003372 | chr3 | Vav3 | 0.0681875 | 0.847318503 |
| ENSMUSG0000003221 | chr9 | Rnf111 | 0.086142618 | 0.847318503 |
| ENSMUSG0000003504 | chr9 | Kri1 | 0.089916439 | 0.847318503 |
| ENSMUSG0000000266 | chr17 | Clpp | 0.114433367 | 0.847318503 |
| ENSMUSG0000006389 | chr13 | Zkscan8 | -0.084658163 | 0.8473258 |
| ENSMUSG0000002041 | chr11 | Nipal4 | 0.13051612 | 0.8473258 |
| ENSMUSG0000005569 | chrX | Gja6 | 0.404276342 | 0.847391441 |
| ENSMUSG000000307C | chr7 | Gdpd3 | -0.247262403 | 0.847708927 |
| ENSMUSG0000003074 | chr7 | Il21r | -0.408147772 | 0.847708927 |
| ENSMUSG0000007886 | chr2 | Gm14322 | -0.206737031 | 0.847708927 |
| ENSMUSG0000002382 | chr17 | Slc22a3 | -0.23060997 | 0.847708927 |
| ENSMUSG0000003498 | chr13 | Hrh2 | -0.360290021 | 0.847708927 |
| ENSMUSG0000002888 | chr4 | Rpa2 | -0.120084754 | 0.847708927 |
| ENSMUSG0000003586 | chr17 | Zfp983 | -0.115507973 | 0.847708927 |
| ENSMUSG0000003721 | chr5 | Fam193a | -0.091327033 | 0.847708927 |
| ENSMUSG0000004965 | chr4 | Zbtb5 | -0.10036775 | 0.847708927 |
| ENSMUSG0000004012 | chr7 | Gpr26 | -0.140017232 | 0.847708927 |
| ENSMUSG0000002526 | chrX | Huwe1 | -0.09195822 | 0.847708927 |
| ENSMUSG0000005242 | chr1 | B4galt3 | -0.082649724 | 0.847708927 |
| ENSMUSG0000005139 | chr5 | Ywhag | -0.065426386 | 0.847708927 |
| ENSMUSG0000007459 | chr2 | Wfdc6a | -0.586683518 | 0.847708927 |
| ENSMUSG0000003664 | chr2 | Man1b1 | 0.063275059 | 0.847708927 |
| ENSMUSG0000000014 | chr5 | Gna12 | 0.065560773 | 0.847708927 |
| ENSMUSG0000002166 | chr13 | Cert1 | 0.064758852 | 0.847708927 |
| ENSMUSG000000609C | chr10 | Arl1 | 0.103576194 | 0.847708927 |
| ENSMUSG0000004212 | chr5 | Ssh1 | 0.083233065 | 0.847708927 |
| ENSMUSG000000259C | chr1 | Lypla1 | 0.090352817 | 0.847708927 |

| | | | | |
|---|---|---|---|---|
| ENSMUSG0000002759 | chr3 | Armc1 | 0.089333836 | 0.847708927 |
| ENSMUSG0000004107 | chr11 | Nacad | 0.093436898 | 0.847708927 |
| ENSMUSG0000001588 | chr10 | Lta4h | 0.122667469 | 0.847708927 |
| ENSMUSG0000003116 | chrX | Ebp | 0.101790292 | 0.847708927 |
| ENSMUSG0000005649 | chr17 | Adgrf5 | 0.142324965 | 0.847708927 |
| ENSMUSG0000005839 | chr10 | Gpr182 | 0.127832183 | 0.847708927 |
| ENSMUSG0000004795 | chr3 | Kcna3 | 0.3700581 | 0.847708927 |
| ENSMUSG0000007935 | chr14 | Gm11100 | -0.415325491 | 0.847881391 |
| ENSMUSG0000002280 | chr16 | Lrch3 | -0.082083739 | 0.848094702 |
| ENSMUSG0000004235 | chr4 | Gjb5 | 0.529191295 | 0.848141416 |
| ENSMUSG0000004569 | chr12 | Wdr89 | -0.23891771 | 0.848569505 |
| ENSMUSG0000002513 | chr11 | Pcyt2 | -0.132790224 | 0.848569505 |
| ENSMUSG0000007885 | chr11 | H2ac25 | -0.137558886 | 0.848569505 |
| ENSMUSG0000007448 | chr3 | Mex3a | -0.169352729 | 0.848569505 |
| ENSMUSG0000005438 | chr1 | Mdm4 | -0.090349953 | 0.848569505 |
| ENSMUSG0000003652 | chr12 | Greb1 | -0.105876032 | 0.848569505 |
| ENSMUSG0000006272 | chr1 | Ppox | -0.090971088 | 0.848569505 |
| ENSMUSG0000001780 | chr11 | Mlx | -0.089563304 | 0.848569505 |
| ENSMUSG0000000897 | chr16 | Gabpa | -0.085202204 | 0.848569505 |
| ENSMUSG0000004205 | chr7 | Wdr11 | -0.08028767 | 0.848569505 |
| ENSMUSG0000004415 | chrX | Bclaf3 | -0.088490855 | 0.848569505 |
| ENSMUSG0000002156 | chr13 | Slc6a19 | 0.347159879 | 0.848569505 |
| ENSMUSG0000004602 | chr2 | Pofut1 | 0.070366804 | 0.848569505 |
| ENSMUSG0000000163 | chr6 | Brpf1 | 0.078847318 | 0.848569505 |
| ENSMUSG0000006348 | chr15 | Snu13 | 0.095747636 | 0.848569505 |
| ENSMUSG0000002766 | chr3 | Zmat3 | 0.084764878 | 0.848569505 |
| ENSMUSG0000003157 | chr8 | Plpp5 | 0.079372671 | 0.848569505 |
| ENSMUSG0000004666 | chr4 | Rbm12b1 | 0.098291439 | 0.848569505 |
| ENSMUSG0000002403 | chr17 | Rsph1 | 0.113374078 | 0.848569505 |
| ENSMUSG0000004171 | chrX | Alg13 | 0.122514295 | 0.848569505 |
| ENSMUSG0000002734 | chr2 | Gpcpd1 | 0.105475079 | 0.848569505 |
| ENSMUSG0000004477 | chr10 | Scml4 | 0.169572908 | 0.848569505 |
| ENSMUSG0000003763 | chr12 | Zbtb42 | -0.074824398 | 0.848577213 |
| ENSMUSG0000002625 | chr1 | Efhd1 | -0.138872578 | 0.848659216 |
| ENSMUSG0000003238 | chr9 | Snx1 | 0.087840587 | 0.849131572 |
| ENSMUSG0000000322 | chr10 | Ranbp2 | 0.079266208 | 0.849131572 |
| ENSMUSG0000003030 | chr6 | Tmtc1 | 0.098238675 | 0.849131572 |
| ENSMUSG0000004139 | chr1 | Mettl18 | 0.146992332 | 0.849131572 |
| ENSMUSG0000004092 | chr4 | S100pbp | -0.075267791 | 0.849171184 |
| ENSMUSG0000003841 | chr18 | Egr1 | -0.383889965 | 0.8492075 |
| ENSMUSG0000002934 | chr5 | Asphd2 | -0.189963611 | 0.8492075 |
| ENSMUSG0000002033 | chr11 | Hmmr | -0.325708975 | 0.8492075 |
| ENSMUSG0000002025 | chr10 | Glt8d2 | -0.219231595 | 0.8492075 |
| ENSMUSG0000003814 | chr1 | Cd84 | -0.168185069 | 0.8492075 |
| ENSMUSG0000004532 | chr9 | Tlr9 | -0.201353507 | 0.8492075 |
| ENSMUSG0000004259 | chr6 | Kdm7a | -0.108160371 | 0.8492075 |
| ENSMUSG0000002416 | chr17 | Jpt2 | -0.101665929 | 0.8492075 |
| ENSMUSG0000005090 | chr16 | Tvp23a | -0.143631551 | 0.8492075 |
| ENSMUSG0000003741 | chr9 | Tbc1d2b | -0.106947535 | 0.8492075 |
| ENSMUSG0000010686 | chr5 | Gtf3c2 | -0.095836251 | 0.8492075 |

| | | | | |
|---|---|---|---|---|
| ENSMUSG0000002593 | chr1 | Paqr8 | -0.093614267 | 0.8492075 |
| ENSMUSG0000005829 | chr15 | Espl1 | -0.19833727 | 0.8492075 |
| ENSMUSG0000003189 | chr8 | Psmb10 | -0.090394639 | 0.8492075 |
| ENSMUSG0000002071 | chr11 | Ern1 | -0.091536024 | 0.8492075 |
| ENSMUSG0000003817 | chr1 | Slamf7 | -0.391345155 | 0.8492075 |
| ENSMUSG0000002734 | chr2 | Fsip1 | -0.359510409 | 0.8492075 |
| ENSMUSG0000006683 | chr7 | Zfp772 | -0.092998869 | 0.8492075 |
| ENSMUSG0000005683 | chr6 | Ift56 | -0.08509704 | 0.8492075 |
| ENSMUSG0000002767 | chr3 | Ttc14 | -0.08297365 | 0.8492075 |
| ENSMUSG0000002570 | chr6 | Marchf8 | -0.070792548 | 0.8492075 |
| ENSMUSG0000001767 | chr2 | Elmo2 | -0.067545667 | 0.8492075 |
| ENSMUSG0000002496 | chr19 | Mark2 | -0.071252025 | 0.8492075 |
| ENSMUSG0000002506 | chr19 | Gsto2 | -0.485214984 | 0.8492075 |
| ENSMUSG0000006223 | chr5 | Gak | -0.050665062 | 0.8492075 |
| ENSMUSG0000000115 | chr10 | Mcm3ap | 0.06049188 | 0.8492075 |
| ENSMUSG0000002976 | chr6 | Exoc4 | 0.064234243 | 0.8492075 |
| ENSMUSG0000001640 | chrX | Nkap | 0.070772491 | 0.8492075 |
| ENSMUSG0000003521 | chr4 | Leprot | 0.074759132 | 0.8492075 |
| ENSMUSG0000006344 | chr5 | Ube2d2b | 0.628411456 | 0.8492075 |
| ENSMUSG0000004081 | chr9 | Tex264 | 0.072409485 | 0.8492075 |
| ENSMUSG0000002867 | chr4 | Srsf10 | 0.071769503 | 0.8492075 |
| ENSMUSG0000005713 | chr10 | Ado | 0.076238406 | 0.8492075 |
| ENSMUSG0000002046 | chr11 | Ppp4r3b | 0.077211575 | 0.8492075 |
| ENSMUSG0000000091 | chr5 | Nsun5 | 0.08275185 | 0.8492075 |
| ENSMUSG0000000201 | chrX | Idh3g | 0.08375533 | 0.8492075 |
| ENSMUSG0000003168 | chr8 | Slc10a7 | 0.078814092 | 0.8492075 |
| ENSMUSG0000005981 | chr4 | Rgs3 | 0.084938457 | 0.8492075 |
| ENSMUSG0000003941 | chr17 | Heatr5b | 0.084701368 | 0.8492075 |
| ENSMUSG0000004221 | chr9 | Abhd14a | 0.088708125 | 0.8492075 |
| ENSMUSG0000003335 | chr15 | Pus7l | 0.104035908 | 0.8492075 |
| ENSMUSG0000007000 | chr8 | Ell | 0.079943225 | 0.8492075 |
| ENSMUSG0000003852 | chr5 | Armc10 | 0.092717491 | 0.8492075 |
| ENSMUSG0000004305 | chr5 | Zfp513 | 0.086370644 | 0.8492075 |
| ENSMUSG0000003910 | chr4 | Atp6v1g1 | 0.120872303 | 0.8492075 |
| ENSMUSG0000002048 | chr11 | Septin4 | 0.095540586 | 0.8492075 |
| ENSMUSG0000004848 | chr2 | Fitm2 | 0.121916643 | 0.8492075 |
| ENSMUSG0000002649 | chr1 | Efcab2 | 0.102886341 | 0.8492075 |
| ENSMUSG0000002578 | chr7 | St8sia2 | 0.206727422 | 0.8492075 |
| ENSMUSG0000003005 | chr6 | Tmf1 | 0.088104935 | 0.8492075 |
| ENSMUSG0000002595 | chr1 | Mettl21a | 0.104349549 | 0.8492075 |
| ENSMUSG0000002222 | chr14 | Sdr39u1 | 0.094345298 | 0.8492075 |
| ENSMUSG0000002187 | chr14 | Rnase4 | 0.101540687 | 0.8492075 |
| ENSMUSG0000002795 | chr3 | Gask1b | 0.105273523 | 0.8492075 |
| ENSMUSG0000004656 | chr5 | Unc119b | 0.109173322 | 0.8492075 |
| ENSMUSG0000004170 | chr15 | Mpped1 | 0.292006934 | 0.8492075 |
| ENSMUSG0000002963 | chr6 | Glcci1 | 0.161462013 | 0.8492075 |
| ENSMUSG0000003966 | chr2 | Spout1 | 0.122553519 | 0.8492075 |
| ENSMUSG0000000018 | chr6 | Fgf23 | 0.437017199 | 0.8492075 |
| ENSMUSG0000002909 | chr5 | Htra3 | 0.141097763 | 0.8492075 |
| ENSMUSG0000006811 | chr2 | Ninl | 0.194056748 | 0.8492075 |

| | | | | |
|---|---|---|---|---|
| ENSMUSG0000011445 | chr13 | H2bc9 | 0.324238168 | 0.8492075 |
| ENSMUSG0000006101 | chr18 | Mkx | 0.156086402 | 0.8492075 |
| ENSMUSG0000007276 | chr5 | 4930522L14Rik | 0.173311509 | 0.8492075 |
| ENSMUSG0000005699 | chr19 | Ide | 0.144844696 | 0.8492075 |
| ENSMUSG0000000335 | chr15 | Fkbp11 | 0.285072883 | 0.8492075 |
| ENSMUSG0000002399 | chr17 | Nfya | 0.060350827 | 0.849230759 |
| ENSMUSG0000007942 | chr1 | Mroh2a | -0.156590382 | 0.849261278 |
| ENSMUSG0000003649 | chr17 | Rnf39 | -0.139792082 | 0.849261278 |
| ENSMUSG0000002664 | chr1 | Cfap126 | -0.212328466 | 0.849261278 |
| ENSMUSG0000004409 | chr5 | C130050O18Rik | 0.307927064 | 0.849261278 |
| ENSMUSG0000004478 | chr11 | Fads6 | -0.102119393 | 0.849400486 |
| ENSMUSG0000002972 | chr5 | Zkscan1 | -0.072304059 | 0.849400486 |
| ENSMUSG0000002970 | chr6 | Gcc1 | -0.06571298 | 0.849400486 |
| ENSMUSG0000002669 | chr1 | Vamp4 | -0.059197085 | 0.849400486 |
| ENSMUSG0000005204 | chr7 | Klf13 | 0.073684635 | 0.849400486 |
| ENSMUSG0000002008 | chr10 | Ppa1 | 0.145905576 | 0.849549661 |
| ENSMUSG0000004575 | chr4 | Mms22l | -0.331812842 | 0.84955983 |
| ENSMUSG0000002009 | chr10 | Npffr1 | -0.427203113 | 0.84955983 |
| ENSMUSG0000001303 | chr8 | Adgrl1 | -0.099395632 | 0.84955983 |
| ENSMUSG0000003865 | chr19 | Ric1 | 0.088286086 | 0.84955983 |
| ENSMUSG0000007218 | chr5 | Speer4e2 | -0.403690613 | 0.849835775 |
| ENSMUSG0000000057 | chr17 | Dynlt1c | -0.158101547 | 0.850085861 |
| ENSMUSG0000005530 | chr7 | Zfp93 | 0.101361048 | 0.850097283 |
| ENSMUSG0000005949 | chr9 | Arhgef12 | 0.080214366 | 0.850097283 |
| ENSMUSG0000002627 | chr1 | Mterf4 | 0.113659956 | 0.850097283 |
| ENSMUSG0000011695 | chr17 | Ermardl2 | 0.248339389 | 0.850173097 |
| ENSMUSG0000002196 | chr14 | Ska3 | -0.377560878 | 0.850201674 |
| ENSMUSG0000000929 | chr10 | Trpm2 | -0.158296771 | 0.850351335 |
| ENSMUSG0000011447 | chr13 | Gm49395 | -0.2793141 | 0.850351335 |
| ENSMUSG0000003245 | chr9 | Clstn2 | -0.122309205 | 0.850351335 |
| ENSMUSG0000005561 | chr2 | Cdca7 | -0.105284449 | 0.850351335 |
| ENSMUSG0000001649 | chr19 | Plgrkt | -0.096683912 | 0.850351335 |
| ENSMUSG0000006778 | chr2 | Blcap | 0.0739375 | 0.850351335 |
| ENSMUSG0000008982 | chr2 | Rbm12 | 0.094948247 | 0.850351335 |
| ENSMUSG0000003991 | chr10 | Cited2 | 0.076686208 | 0.850351335 |
| ENSMUSG0000005755 | chr13 | Lgals8 | -0.072618604 | 0.850423193 |
| ENSMUSG0000002732 | chr2 | Knl1 | -0.515882745 | 0.850982916 |
| ENSMUSG0000002692 | chr2 | Notch1 | -0.155995114 | 0.850982916 |
| ENSMUSG0000002599 | chr1 | Slc40a1 | -0.182838876 | 0.850982916 |
| ENSMUSG0000003701 | chrX | Apln | -0.381595586 | 0.850982916 |
| ENSMUSG0000001976 | chr10 | Iyd | -0.725571254 | 0.850982916 |
| ENSMUSG0000003962 | chr5 | Ap5z1 | -0.120996123 | 0.850982916 |
| ENSMUSG0000003311 | chr13 | Slc35d2 | -0.168170211 | 0.850982916 |
| ENSMUSG0000000909 | chr16 | Tbx1 | -0.186736167 | 0.850982916 |
| ENSMUSG0000003963 | chr16 | Coro7 | -0.098576426 | 0.850982916 |
| ENSMUSG0000002453 | chr18 | Prelid3a | -0.185925406 | 0.850982916 |
| ENSMUSG0000003687 | chr10 | Dna2 | -0.239645167 | 0.850982916 |
| ENSMUSG0000005339 | chr3 | Phgdh | -0.097481607 | 0.850982916 |
| ENSMUSG0000002148 | chr13 | Cdk20 | -0.114197211 | 0.850982916 |
| ENSMUSG0000000287 | chr6 | Mcm2 | -0.103725631 | 0.850982916 |

| | | | | |
|---|---|---|---|---|
| ENSMUSG0000003111 | chrX | Igsf1 | -0.141119968 | 0.850982916 |
| ENSMUSG0000001994 | chr10 | Atp2b1 | -0.070034537 | 0.850982916 |
| ENSMUSG0000003479 | chr11 | G6pc3 | -0.069637645 | 0.850982916 |
| ENSMUSG0000002889 | chr4 | Inpp5b | 0.052810042 | 0.850982916 |
| ENSMUSG0000003728 | chr9 | Tbcel | 0.072281749 | 0.850982916 |
| ENSMUSG0000003302 | chr1 | Gmppa | 0.066450078 | 0.850982916 |
| ENSMUSG0000002097 | chr12 | Klhdc2 | 0.071595359 | 0.850982916 |
| ENSMUSG0000004121 | chr16 | Yeats2 | 0.081165806 | 0.850982916 |
| ENSMUSG0000002407 | chr17 | Yipf4 | 0.0733231 | 0.850982916 |
| ENSMUSG0000003309 | chr2 | Apmap | 0.065137339 | 0.850982916 |
| ENSMUSG0000002069 | chr11 | Rffl | 0.079922592 | 0.850982916 |
| ENSMUSG0000006067 | chr1 | Mrps9 | 0.077244042 | 0.850982916 |
| ENSMUSG0000002399 | chr17 | Trem2 | 0.315895253 | 0.850982916 |
| ENSMUSG0000003663 | chr3 | Alg5 | 0.094060461 | 0.850982916 |
| ENSMUSG0000005948 | chr6 | Kbtbd2 | 0.072288412 | 0.850982916 |
| ENSMUSG0000002713 | chr2 | Katnbl1 | 0.088536048 | 0.850982916 |
| ENSMUSG0000003719 | chr9 | Cyb561d2 | 0.139809221 | 0.850982916 |
| ENSMUSG0000007003 | chr1 | Sp140 | 0.189263164 | 0.850982916 |
| ENSMUSG0000000222 | chr3 | Ppm1j | 0.123832324 | 0.850982916 |
| ENSMUSG0000003310 | chr18 | Rnf125 | 0.1787178 | 0.850982916 |
| ENSMUSG0000005073 | chr2 | Ptges | 0.164855441 | 0.850982916 |
| ENSMUSG0000002707 | chr2 | Slc43a1 | 0.198040365 | 0.850982916 |
| ENSMUSG0000002089 | chr11 | Aloxe3 | 0.340167695 | 0.850982916 |
| ENSMUSG0000003156 | chr8 | Dctd | 0.282757949 | 0.850982916 |
| ENSMUSG0000005534 | chr13 | Zfp457 | -0.318360858 | 0.851058069 |
| ENSMUSG0000003003 | chr6 | Kbtbd8 | -0.180056646 | 0.851058069 |
| ENSMUSG0000003054 | chr7 | Pex11a | -0.078553903 | 0.851058069 |
| ENSMUSG0000002654 | chr1 | Slamf9 | 0.239129668 | 0.851058069 |
| ENSMUSG0000003376 | chr6 | Exoc6b | -0.071938198 | 0.851246264 |
| ENSMUSG0000003882 | chr5 | Tmem214 | 0.056112265 | 0.851246264 |
| ENSMUSG0000003870 | chr2 | Gmeb2 | 0.086816034 | 0.851246264 |
| ENSMUSG0000003773 | chr8 | Nek5 | 0.165090273 | 0.851246264 |
| ENSMUSG0000004073 | chr4 | Ints8 | 0.098300554 | 0.851246264 |
| ENSMUSG0000002306 | chr18 | Rttn | -0.140738804 | 0.851524882 |
| ENSMUSG0000000645 | chr19 | Actn3 | -0.201031856 | 0.851666525 |
| ENSMUSG0000001840 | chr11 | Mtmr4 | -0.089986428 | 0.851703635 |
| ENSMUSG0000002742 | chr2 | Rbbp9 | 0.082147308 | 0.851703635 |
| ENSMUSG0000002977 | chr6 | Gars1 | 0.093251324 | 0.851703635 |
| ENSMUSG0000002527 | chr14 | Abhd6 | 0.087183066 | 0.851703635 |
| ENSMUSG0000011533 | chr14 | Pnp | 0.145013134 | 0.851703635 |
| ENSMUSG0000003178 | chr8 | Adgrg1 | -0.112144445 | 0.851746286 |
| ENSMUSG0000002693 | chr2 | Camsap1 | 0.078437423 | 0.851746286 |
| ENSMUSG0000003677 | chr17 | Decr2 | 0.074317345 | 0.851746286 |
| ENSMUSG0000005222 | chr18 | Gpr17 | 0.13368219 | 0.851746286 |
| ENSMUSG0000002023 | chr10 | Tektip1 | 0.582924858 | 0.851746286 |
| ENSMUSG0000002880 | chr4 | Nipal3 | -0.113441861 | 0.851838814 |
| ENSMUSG0000002026 | chr10 | Appl2 | -0.112550701 | 0.851838814 |
| ENSMUSG0000005103 | chr13 | Zfp455 | 0.15821884 | 0.851838814 |
| ENSMUSG0000003491 | chr8 | Ushbp1 | -0.150943938 | 0.851862715 |
| ENSMUSG0000006414 | chr17 | Rab44 | -0.278331421 | 0.851862715 |

| | | | | |
|---|---|---|---|---|
| ENSMUSG0000004139 | chr6 | Mdfic | 0.084740158 | 0.851862715 |
| ENSMUSG0000003878 | chr7 | 1700028J19Rik | 0.415242048 | 0.851862715 |
| ENSMUSG0000005149 | chr11 | Kcnj16 | 0.642502419 | 0.851862715 |
| ENSMUSG0000004104 | chr11 | Ramp3 | 0.156383261 | 0.851867895 |
| ENSMUSG0000004998 | chr8 | Lrrc25 | -0.201908761 | 0.851902022 |
| ENSMUSG0000003237 | chr9 | Usp3 | -0.102728717 | 0.851902022 |
| ENSMUSG0000002914 | chr5 | Nrbp1 | 0.05220189 | 0.851902022 |
| ENSMUSG0000002098 | chr12 | Nemf | 0.05782732 | 0.851902022 |
| ENSMUSG0000009448 | chr11 | Purb | 0.074757207 | 0.851902022 |
| ENSMUSG0000004548 | chr5 | Trrap | -0.090372851 | 0.852328602 |
| ENSMUSG0000002578 | chr9 | Tgm4 | -0.229622419 | 0.852328602 |
| ENSMUSG0000002784 | chr3 | Wnt2b | -0.128890509 | 0.852328602 |
| ENSMUSG0000007547 | chr15 | Alg10b | -0.074143071 | 0.852328602 |
| ENSMUSG0000003524 | chr13 | Tut7 | -0.074772707 | 0.852328602 |
| ENSMUSG0000003782 | chr6 | Ppm1k | -0.07386964 | 0.852328602 |
| ENSMUSG0000003223 | chr9 | Ice2 | 0.081608579 | 0.852328602 |
| ENSMUSG0000012192 | chr16 | ENSMUSG000001 | 0.510461411 | 0.852328602 |
| ENSMUSG0000001608 | chr9 | Fli1 | -0.157023273 | 0.852423018 |
| ENSMUSG0000002944 | chr5 | Cfap251 | -0.167572981 | 0.852423018 |
| ENSMUSG0000007427 | chr7 | Phldb3 | -0.145984887 | 0.852423018 |
| ENSMUSG0000007861 | chr4 | Fyb2 | -0.305322715 | 0.852423018 |
| ENSMUSG0000011850 | chr3 | Gm53012 | -0.199068179 | 0.852423018 |
| ENSMUSG0000005344 | chr18 | Adamts19 | -0.103565834 | 0.852423018 |
| ENSMUSG0000007359 | chr18 | Pcdhb22 | -0.102505981 | 0.852423018 |
| ENSMUSG0000002965 | chr5 | N4bp2l2 | 0.06178356 | 0.852423018 |
| ENSMUSG0000003191 | chr8 | Vps4a | 0.082836008 | 0.852423018 |
| ENSMUSG0000002670 | chr1 | Dars2 | 0.08878437 | 0.852423018 |
| ENSMUSG0000003362 | chr18 | Pik3c3 | 0.086421866 | 0.852423018 |
| ENSMUSG0000006791 | chr4 | Zfp991 | 0.128273159 | 0.852423018 |
| ENSMUSG0000004657 | chr7 | Prr12 | -0.10587915 | 0.852608366 |
| ENSMUSG0000007415 | chr7 | Zfp976 | -0.123104655 | 0.852608366 |
| ENSMUSG0000002441 | chr18 | Npc1 | -0.075748974 | 0.852608366 |
| ENSMUSG0000002795 | chr3 | Slc50a1 | -0.064415838 | 0.852608366 |
| ENSMUSG0000002277 | chr16 | Top3b | 0.054113139 | 0.852608366 |
| ENSMUSG0000007965 | chr5 | Tmem243 | 0.074590777 | 0.852608366 |
| ENSMUSG0000000150 | chr11 | Itga3 | 0.086418063 | 0.852608366 |
| ENSMUSG0000003012 | chr6 | Lag3 | 0.181957424 | 0.852608366 |
| ENSMUSG0000002882 | chr4 | Tmem50a | 0.093737335 | 0.852659312 |
| ENSMUSG0000002165 | chr13 | Ptcd2 | 0.115365611 | 0.852659312 |
| ENSMUSG0000006204 | chr7 | Zfp27 | 0.104405159 | 0.852659312 |
| ENSMUSG0000002030 | chr11 | Stc2 | 0.210793399 | 0.852659312 |
| ENSMUSG0000002948 | chr5 | Dhx37 | -0.07321452 | 0.852687326 |
| ENSMUSG0000009069 | chr9 | Trim43a | -0.371372909 | 0.852802633 |
| ENSMUSG0000000639 | chr4 | Med8 | 0.067167876 | 0.852802633 |
| ENSMUSG0000005601 | chr5 | A430033K04Rik | -0.078110452 | 0.852970502 |
| ENSMUSG0000000765 | chr9 | Arpp19 | 0.070407221 | 0.852970502 |
| ENSMUSG0000004619 | chr18 | Pcdhb20 | -0.142948695 | 0.853004115 |
| ENSMUSG0000004506 | chr12 | Zc2hc1c | 0.118595812 | 0.853004115 |
| ENSMUSG0000003898 | chr15 | Tspyl5 | -0.086483385 | 0.85301882 |
| ENSMUSG0000003990 | chr19 | Armh3 | 0.091369756 | 0.85301882 |

| | | | | |
|---|---|---|---|---|
| ENSMUSG0000004254 | chr12 | Acot5 | -0.325339132 | 0.853071892 |
| ENSMUSG0000002834 | chr4 | Tmeff1 | 0.107932386 | 0.853156166 |
| ENSMUSG0000004924 | chr5 | Hcar1 | -0.119533287 | 0.85321199 |
| ENSMUSG0000004679 | chr5 | Cldn12 | -0.084993514 | 0.853316257 |
| ENSMUSG0000003854 | chr4 | Inip | -0.065510984 | 0.853316257 |
| ENSMUSG0000002094 | chr12 | Klhl28 | -0.070033301 | 0.853316257 |
| ENSMUSG0000006794 | chr17 | Zfp160 | 0.056573471 | 0.853316257 |
| ENSMUSG0000002502 | chr19 | Smndc1 | 0.084978997 | 0.853316257 |
| ENSMUSG0000004958 | chr11 | Ccdc69 | 0.136988713 | 0.853316257 |
| ENSMUSG0000000157 | chr17 | Ergic1 | 0.103687778 | 0.853389619 |
| ENSMUSG0000002813 | chr3 | Tlcd4 | -0.102305362 | 0.853487444 |
| ENSMUSG0000004601 | chr11 | Zfp830 | 0.09513278 | 0.853487444 |
| ENSMUSG0000002414 | chr17 | Pigf | 0.138753415 | 0.853487444 |
| ENSMUSG0000002124 | chr12 | Ylpm1 | -0.092054788 | 0.853547455 |
| ENSMUSG0000002070 | chr11 | Ddx42 | -0.065172215 | 0.853547455 |
| ENSMUSG0000002709 | chr2 | Zc3h15 | 0.07264605 | 0.853547455 |
| ENSMUSG0000002238 | chr15 | Trmu | 0.0956493 | 0.853547455 |
| ENSMUSG0000002188 | chr12 | Gpr65 | -0.323492742 | 0.853614745 |
| ENSMUSG0000002393 | chr17 | Aars2 | -0.091158067 | 0.853803838 |
| ENSMUSG0000002557 | chr11 | Rbfox3 | -0.09349462 | 0.854142939 |
| ENSMUSG0000002694 | chr2 | Nmi | -0.075921017 | 0.854142939 |
| ENSMUSG0000002677 | chr2 | Prkcq | 0.118086935 | 0.854142939 |
| ENSMUSG0000002270 | chr16 | Hira | 0.111763888 | 0.85414554 |
| ENSMUSG0000005249 | chr15 | Pkdrej | -0.143656389 | 0.854267184 |
| ENSMUSG0000001138 | chr7 | Dhdh | -0.100210242 | 0.854267184 |
| ENSMUSG0000002888 | chr4 | Eya3 | -0.076274553 | 0.854267184 |
| ENSMUSG0000002968 | chr6 | Wasl | -0.061849277 | 0.854267184 |
| ENSMUSG0000004995 | chr11 | Ccdc137 | 0.096875731 | 0.854267184 |
| ENSMUSG0000002298 | chr15 | Zfp641 | 0.148457447 | 0.854267184 |
| ENSMUSG0000003935 | chr2 | Exosc2 | 0.103665248 | 0.854267184 |
| ENSMUSG0000003090 | chr7 | Crym | 0.220493174 | 0.854267184 |
| ENSMUSG0000003296 | chr1 | Inha | -0.152276983 | 0.854478042 |
| ENSMUSG0000008301 | chr5 | Fam220a | -0.434811863 | 0.854478042 |
| ENSMUSG0000002064 | chr12 | Ncoa1 | -0.089030411 | 0.854478042 |
| ENSMUSG0000003571 | chr7 | Usp35 | -0.094695878 | 0.854478042 |
| ENSMUSG0000002254 | chr16 | Apod | -0.172295806 | 0.854478042 |
| ENSMUSG0000004661 | chr16 | Vwa5b2 | -0.093920014 | 0.854478042 |
| ENSMUSG0000003460 | chr10 | Mon2 | -0.057984152 | 0.854478042 |
| ENSMUSG0000000968 | chr10 | Bcr | -0.050407365 | 0.854478042 |
| ENSMUSG0000003840 | chr7 | Scaf1 | 0.066591785 | 0.854478042 |
| ENSMUSG0000002877 | chr4 | Pef1 | 0.090607055 | 0.854478042 |
| ENSMUSG0000002230 | chr15 | Dcaf13 | 0.076696114 | 0.854478042 |
| ENSMUSG0000004495 | chr11 | Pwwp2a | 0.086667782 | 0.854478042 |
| ENSMUSG0000003172 | chr8 | Hp | 0.238261943 | 0.854478042 |
| ENSMUSG0000002817 | chr3 | Lrrc7 | 0.24770158 | 0.854478042 |
| ENSMUSG0000003067 | chr7 | Cyp2r1 | -0.278222535 | 0.854495603 |
| ENSMUSG0000004120 | chr16 | Map6d1 | -0.222389924 | 0.854495603 |
| ENSMUSG0000003004 | chr6 | Tacr1 | -0.115975977 | 0.854495603 |
| ENSMUSG0000003282 | chr6 | Ppp1r9a | -0.093818585 | 0.854495603 |
| ENSMUSG0000004651 | chr3 | Golph3l | -0.094164648 | 0.854495603 |

| | | | | |
|---|---|---|---|---|
| ENSMUSG0000005167 | chr4 | Trim32 | -0.079965311 | 0.854495603 |
| ENSMUSG0000001309 | chr7 | Tmem190 | -0.259121909 | 0.854495603 |
| ENSMUSG0000003577 | chr2 | Ggta1 | -0.063703726 | 0.854495603 |
| ENSMUSG0000002742 | chr2 | Polr3f | 0.056104566 | 0.854495603 |
| ENSMUSG0000002912 | chr5 | Zbtb49 | 0.095828369 | 0.854495603 |
| ENSMUSG0000005982 | chr9 | Nkapd1 | 0.072996783 | 0.854495603 |
| ENSMUSG0000000184 | chr5 | Zdhhc4 | 0.079831506 | 0.854495603 |
| ENSMUSG0000003061 | chr7 | Tmem126b | 0.089363858 | 0.854495603 |
| ENSMUSG0000007504 | chr2 | Zfp408 | 0.11504244 | 0.854495603 |
| ENSMUSG0000002699 | chr2 | Nup35 | 0.128067717 | 0.854495603 |
| ENSMUSG0000002959 | chr5 | Ung | 0.216638983 | 0.854495603 |
| ENSMUSG0000000060 | chr5 | Krit1 | 0.062540762 | 0.854541757 |
| ENSMUSG0000002617 | chr1 | Cyp27a1 | 0.118068936 | 0.85471349 |
| ENSMUSG0000002739 | chr2 | Ttl | -0.069170864 | 0.854721048 |
| ENSMUSG0000005689 | chr12 | Immp2l | -0.174739159 | 0.854726002 |
| ENSMUSG0000006237 | chr5 | Otof | 0.24535466 | 0.854822576 |
| ENSMUSG0000005265 | chr6 | Rnf103 | -0.054762511 | 0.854829371 |
| ENSMUSG0000003097 | chr7 | Uros | 0.116480541 | 0.855201883 |
| ENSMUSG0000002949 | chr5 | Pde6b | 0.293062787 | 0.855201883 |
| ENSMUSG0000007890 | chr15 | Fam186b | -0.390126528 | 0.855224189 |
| ENSMUSG0000004712 | chr17 | Ticam1 | -0.105280971 | 0.855276959 |
| ENSMUSG0000000348 | chr8 | Cyp4f18 | -0.277175528 | 0.855276959 |
| ENSMUSG0000000306 | chr13 | Stard3nl | -0.08649337 | 0.855276959 |
| ENSMUSG0000002421 | chr17 | Anks1 | -0.062232728 | 0.855276959 |
| ENSMUSG0000000323 | chr16 | Abcf3 | 0.058148536 | 0.855276959 |
| ENSMUSG0000000168 | chr5 | Ubl3 | 0.074843141 | 0.855276959 |
| ENSMUSG0000002406 | chr17 | Ehd3 | 0.065044238 | 0.855276959 |
| ENSMUSG0000003182 | chr8 | Cotl1 | 0.070178614 | 0.855276959 |
| ENSMUSG0000002314 | chr16 | Nagpa | 0.094739106 | 0.855276959 |
| ENSMUSG0000005590 | chr4 | Tmem69 | 0.093064874 | 0.855276959 |
| ENSMUSG0000003250 | chr9 | Myd88 | 0.102234693 | 0.855276959 |
| ENSMUSG0000002547 | chr7 | Zfp511 | 0.083958296 | 0.855276959 |
| ENSMUSG0000005121 | chr14 | Gpr183 | 0.171843074 | 0.855276959 |
| ENSMUSG0000002139 | chr13 | Nxnl2 | 0.441933133 | 0.855276959 |
| ENSMUSG0000006296 | chr5 | Kdr | 0.110372355 | 0.855276959 |
| ENSMUSG0000004836 | chr13 | Omd | 0.389669731 | 0.855276959 |
| ENSMUSG0000003293 | chr9 | Camkv | 0.365839068 | 0.855430933 |
| ENSMUSG0000003747 | chr1 | Dtl | 0.211690497 | 0.855430933 |
| ENSMUSG0000002035 | chr11 | Sgcd | 0.144106271 | 0.855430933 |
| ENSMUSG0000000073 | chr10 | Dnmt3l | 0.38754093 | 0.855430933 |
| ENSMUSG0000004810 | chr3 | Rbm15 | -0.128369494 | 0.855449304 |
| ENSMUSG0000005691 | chr9 | Cep162 | -0.064390038 | 0.855449304 |
| ENSMUSG0000000124 | chr7 | Gramd1a | -0.054176287 | 0.855487545 |
| ENSMUSG0000000547 | chr8 | Asf1b | -0.257690453 | 0.85565253 |
| ENSMUSG0000003576 | chr13 | Tmem161b | 0.071366133 | 0.85565253 |
| ENSMUSG0000003737 | chr5 | Ctbp1 | 0.09485564 | 0.855976227 |
| ENSMUSG0000003408 | chr6 | Ccdc174 | 0.093651668 | 0.855976227 |
| ENSMUSG0000006432 | chr8 | Hhip | -0.247983214 | 0.856058714 |
| ENSMUSG0000006619 | chr4 | Anks6 | -0.113595057 | 0.856058714 |
| ENSMUSG0000004259 | chrX | Fam199x | -0.089042447 | 0.856058714 |

| | | | | |
|---|---|---|---|---|
| ENSMUSG0000004184 | chr3 | Fhdc1 | -0.1196265 | 0.856058714 |
| ENSMUSG0000002299 | chr15 | Kansl2 | 0.062505996 | 0.856058714 |
| ENSMUSG0000001125 | chr11 | Thg1l | 0.126299615 | 0.856058714 |
| ENSMUSG0000003178 | chr8 | Kifc3 | 0.121624319 | 0.856058714 |
| ENSMUSG0000003389 | chr1 | Cfhr2 | 0.568150143 | 0.856072834 |
| ENSMUSG0000002846 | chr4 | Car9 | -0.129472274 | 0.856635508 |
| ENSMUSG0000006873 | chr3 | Sars1 | 0.057407875 | 0.856635508 |
| ENSMUSG0000003055 | chr7 | Mef2a | 0.09020896 | 0.856635508 |
| ENSMUSG0000002609 | chr1 | Ormdl1 | 0.114693828 | 0.856635508 |
| ENSMUSG0000002284 | chr16 | Thpo | -0.172285288 | 0.856730473 |
| ENSMUSG0000004140 | chr1 | Firrm | -0.320649728 | 0.856983134 |
| ENSMUSG0000009968 | chr7 | Zfp383 | 0.096911942 | 0.856983134 |
| ENSMUSG0000002159 | chr13 | Arsk | -0.065538517 | 0.857023013 |
| ENSMUSG0000002951 | chr5 | Prkab1 | -0.09559426 | 0.857153908 |
| ENSMUSG0000000231 | chr14 | Ipo4 | 0.097198188 | 0.857153908 |
| ENSMUSG0000002175 | chr13 | Map3k1 | -0.086633796 | 0.857207035 |
| ENSMUSG0000003905 | chr2 | Osbpl2 | -0.076035208 | 0.857207035 |
| ENSMUSG0000003923 | chr13 | Tbce | -0.067977676 | 0.857207035 |
| ENSMUSG0000003711 | chr3 | Setd7 | 0.071729964 | 0.857207035 |
| ENSMUSG0000003178 | chr8 | Ciapin1 | 0.089660656 | 0.857207035 |
| ENSMUSG0000002150 | chr13 | Cxcl14 | -0.172463821 | 0.857356032 |
| ENSMUSG0000003856 | chr7 | Efl1 | -0.064147217 | 0.857356032 |
| ENSMUSG0000002724 | chr2 | Wdr76 | 0.099648003 | 0.857391314 |
| ENSMUSG0000006160 | chr17 | Mdc1 | -0.085165208 | 0.857485557 |
| ENSMUSG0000002650 | chr1 | Sdccag8 | -0.083095678 | 0.857485557 |
| ENSMUSG0000006070 | chr5 | Bloc1s4 | 0.135617674 | 0.857838156 |
| ENSMUSG0000003513 | chr12 | Arhgap5 | 0.063404401 | 0.857907114 |
| ENSMUSG0000002007 | chr10 | Ccar1 | 0.059780432 | 0.858037981 |
| ENSMUSG0000003003 | chr6 | Mogs | -0.101042862 | 0.858278366 |
| ENSMUSG0000003274 | chr7 | Katnip | -0.068431572 | 0.858278366 |
| ENSMUSG0000004527 | chr16 | Lca5l | 0.141829076 | 0.858278366 |
| ENSMUSG0000002838 | chr4 | Zfp37 | -0.087278709 | 0.858540137 |
| ENSMUSG0000004463 | chr15 | Csrnp2 | -0.107234118 | 0.858642846 |
| ENSMUSG0000002497 | chr19 | Spindoc | 0.069454991 | 0.858642846 |
| ENSMUSG0000003891 | chr17 | Plcl2 | -0.111945273 | 0.858711245 |
| ENSMUSG0000000239 | chr8 | Ocel1 | -0.073145984 | 0.858711245 |
| ENSMUSG0000003365 | chr16 | Vps8 | 0.055953675 | 0.858711245 |
| ENSMUSG0000003773 | chr4 | Themis2 | -0.180686117 | 0.858713019 |
| ENSMUSG0000004302 | chr11 | Trpv3 | -0.408453458 | 0.858713019 |
| ENSMUSG0000002618 | chr1 | Xrcc5 | 0.081781938 | 0.858713019 |
| ENSMUSG0000002392 | chr17 | Satb1 | 0.111143431 | 0.858713019 |
| ENSMUSG0000002668 | chr1 | Mgst3 | 0.113783053 | 0.858713019 |
| ENSMUSG0000004990 | chr11 | Tmem17 | -0.124700436 | 0.858759111 |
| ENSMUSG0000004849 | chr16 | Nrip1 | -0.08336936 | 0.858759111 |
| ENSMUSG0000004309 | chr11 | Hic1 | -0.092769859 | 0.858759111 |
| ENSMUSG0000002770 | chr3 | Mccc1 | -0.079302741 | 0.858759111 |
| ENSMUSG0000002822 | chr4 | Pip4p2 | -0.066439625 | 0.858759111 |
| ENSMUSG0000000357 | chr8 | Homer3 | -0.088895038 | 0.858759111 |
| ENSMUSG0000003232 | chr9 | Hmg20a | 0.049077074 | 0.858759111 |
| ENSMUSG0000002999 | chr6 | Pcyox1 | 0.051738076 | 0.858759111 |

| | | | | |
|---|---|---|---|---|
| ENSMUSG0000003921 | chr13 | Arid4b | 0.06271953 | 0.858759111 |
| ENSMUSG0000003230 | chr9 | Ube2q2 | 0.069676725 | 0.858759111 |
| ENSMUSG0000003284 | chr5 | Spring1 | 0.083860142 | 0.858759111 |
| ENSMUSG0000001006 | chr9 | Cacna2d2 | 0.108366162 | 0.858759111 |
| ENSMUSG0000001463 | chr8 | Cmc2 | 0.114674576 | 0.858759111 |
| ENSMUSG0000002885 | chr4 | Tmem222 | 0.101947768 | 0.858759111 |
| ENSMUSG0000002021 | chr10 | Timm13 | 0.210374254 | 0.858759111 |
| ENSMUSG0000006758 | chr13 | S1pr3 | -0.099465226 | 0.858965536 |
| ENSMUSG0000000345 | chr6 | Bicd1 | -0.073315286 | 0.858965536 |
| ENSMUSG0000004009 | chr17 | Flywch1 | -0.066870486 | 0.858965536 |
| ENSMUSG0000004889 | chr7 | Zfp710 | -0.063595104 | 0.858965536 |
| ENSMUSG0000003348 | chr1 | Cryzl2 | 0.114914276 | 0.858965536 |
| ENSMUSG0000003564 | chr7 | Aamdc | 0.120758014 | 0.858965536 |
| ENSMUSG0000002287 | chr16 | Samsn1 | 0.226390147 | 0.859088065 |
| ENSMUSG0000002700 | chr2 | Itprid2 | 0.09739941 | 0.859093035 |
| ENSMUSG0000002111 | chr12 | Atp6v1d | 0.055414034 | 0.859425661 |
| ENSMUSG0000000537 | chr5 | Bud23 | 0.086809172 | 0.859425661 |
| ENSMUSG0000000102 | chr11 | Scn4a | -0.411703733 | 0.859874424 |
| ENSMUSG0000001780 | chr11 | Retreg3 | 0.052186946 | 0.859874424 |
| ENSMUSG0000000053 | chr15 | Acvrl1 | 0.080628059 | 0.860015589 |
| ENSMUSG0000002304 | chr15 | Krt18 | -0.34422041 | 0.860227826 |
| ENSMUSG0000002052 | chr11 | Prpsap2 | -0.093516656 | 0.860227826 |
| ENSMUSG0000002553 | chr5 | Asl | -0.096555507 | 0.860227826 |
| ENSMUSG0000007428 | chr7 | Zfp94 | -0.113380745 | 0.860227826 |
| ENSMUSG0000011689 | chr17 | Ermardl1 | -0.379855576 | 0.860227826 |
| ENSMUSG0000002909 | chr5 | Sorcs2 | -0.07546769 | 0.860227826 |
| ENSMUSG0000003396 | chr1 | Zbtb41 | -0.072385776 | 0.860227826 |
| ENSMUSG0000001981 | chr10 | Fuca2 | -0.103551768 | 0.860227826 |
| ENSMUSG0000003920 | chrX | Tbc1d25 | -0.08666928 | 0.860227826 |
| ENSMUSG0000007185 | chr18 | Mcc | -0.081850981 | 0.860227826 |
| ENSMUSG0000002925 | chr5 | Cenpc1 | -0.055695425 | 0.860227826 |
| ENSMUSG0000000283 | chr17 | Hdgfl2 | 0.04606345 | 0.860227826 |
| ENSMUSG0000002515 | chr11 | Dus1l | 0.079014762 | 0.860227826 |
| ENSMUSG0000004471 | chr12 | Gskip | 0.08011431 | 0.860227826 |
| ENSMUSG0000003775 | chr14 | Ptger2 | 0.126496223 | 0.860227826 |
| ENSMUSG0000003006 | chr6 | Gm20696 | 0.273158438 | 0.860227826 |
| ENSMUSG0000003205 | chr9 | Rdx | 0.064179844 | 0.860231761 |
| ENSMUSG0000005667 | chrY | Kdm5d | -0.067927715 | 0.860234663 |
| ENSMUSG0000002007 | chr10 | Pbld2 | -0.183350551 | 0.86039091 |
| ENSMUSG0000003262 | chr6 | Thsd7a | -0.088148696 | 0.86039091 |
| ENSMUSG0000002603 | chr1 | Cflar | -0.052971755 | 0.86039091 |
| ENSMUSG0000002981 | chr6 | Tra2a | 0.075225791 | 0.86039091 |
| ENSMUSG0000003396 | chr17 | Cdkl4 | -0.225221936 | 0.86041554 |
| ENSMUSG0000004788 | chr15 | Tnrc6b | -0.112343778 | 0.860482027 |
| ENSMUSG0000003485 | chr4 | Acot11 | 0.086071455 | 0.860482027 |
| ENSMUSG0000004179 | chr11 | Abca9 | -0.11565616 | 0.860874414 |
| ENSMUSG0000003842 | chr6 | Usp5 | 0.046671993 | 0.860874414 |
| ENSMUSG0000002283 | chr16 | Iqcb1 | 0.077589249 | 0.860874414 |
| ENSMUSG0000000641 | chr1 | Pfdn2 | 0.083859237 | 0.860874414 |
| ENSMUSG0000002785 | chr3 | Nras | -0.072749044 | 0.860885489 |

| | | | | |
|---|---|---|---|---|
| ENSMUSG0000003396 | chr7 | Rnf225 | -0.342859709 | 0.860965218 |
| ENSMUSG0000001771 | chr11 | Pgs1 | 0.101063539 | 0.860965218 |
| ENSMUSG0000002192 | chr14 | Ebpl | 0.10038492 | 0.860965218 |
| ENSMUSG0000002190 | chr14 | Phf7 | 0.097859857 | 0.861038695 |
| ENSMUSG0000002755 | chr3 | E2f5 | 0.079693499 | 0.861150449 |
| ENSMUSG0000002406 | chr17 | Xdh | -0.168358158 | 0.861202887 |
| ENSMUSG0000002009 | chr10 | Eif4ebp2 | -0.051162767 | 0.861202887 |
| ENSMUSG0000009247 | chr16 | Gm20518 | 0.445652981 | 0.861202887 |
| ENSMUSG0000003734 | chr19 | Nudt22 | 0.149996353 | 0.861202887 |
| ENSMUSG0000002912 | chr5 | Evc | -0.063991211 | 0.861288078 |
| ENSMUSG0000003349 | chr9 | Prss35 | -0.179111986 | 0.861568993 |
| ENSMUSG0000002021 | chr10 | Mdm1 | -0.09598298 | 0.861568993 |
| ENSMUSG0000003747 | chr1 | Uggt1 | 0.050010391 | 0.861568993 |
| ENSMUSG0000002689 | chr2 | Gca | -0.131911328 | 0.861590027 |
| ENSMUSG0000002249 | chr15 | Ppp1r1a | 0.153041322 | 0.861590027 |
| ENSMUSG0000002177 | chr14 | Nr1d2 | -0.373328079 | 0.861614552 |
| ENSMUSG0000004476 | chr1 | Macir | -0.092313766 | 0.86164144 |
| ENSMUSG0000003115 | chrX | Gripap1 | -0.072366037 | 0.86164144 |
| ENSMUSG0000007406 | chr8 | Osgin1 | -0.229618766 | 0.861653375 |
| ENSMUSG0000002397 | chr17 | Guca1b | -0.156397006 | 0.861653375 |
| ENSMUSG0000006046 | chr9 | Dppa5a | 0.379084051 | 0.861653375 |
| ENSMUSG0000002631 | chr8 | Cln8 | 0.06392031 | 0.861699147 |
| ENSMUSG0000007170 | chrX | Sms | -0.068776711 | 0.86183862 |
| ENSMUSG0000004014 | chr17 | Tdrd6 | -0.368531818 | 0.8618497 |
| ENSMUSG0000003154 | chr8 | Sfrp1 | -0.096943305 | 0.8618497 |
| ENSMUSG0000005216 | chr12 | Pld4 | -0.157909213 | 0.8618497 |
| ENSMUSG0000003380 | chr5 | Ephx4 | -0.135511943 | 0.8618497 |
| ENSMUSG0000007342 | chr17 | Hsd17b8 | -0.342389673 | 0.8618497 |
| ENSMUSG0000004395 | chr9 | Ccrl2 | -0.093550657 | 0.8618497 |
| ENSMUSG0000006815 | chr2 | Insm1 | -0.161379735 | 0.8618497 |
| ENSMUSG0000005733 | chr1 | Cep170 | -0.07599595 | 0.8618497 |
| ENSMUSG0000007815 | chr11 | Fem1al | -0.49636909 | 0.8618497 |
| ENSMUSG0000002777 | chr3 | Dhx36 | -0.058947836 | 0.8618497 |
| ENSMUSG0000000078 | chrX | Ddx3x | 0.05024732 | 0.8618497 |
| ENSMUSG0000003689 | chr2 | Ehmt1 | 0.056063441 | 0.8618497 |
| ENSMUSG0000003704 | chr7 | Smpd1 | 0.061540717 | 0.8618497 |
| ENSMUSG0000004943 | chr1 | Cyp20a1 | 0.066946327 | 0.8618497 |
| ENSMUSG0000000907 | chr11 | Ewsr1 | 0.091194281 | 0.8618497 |
| ENSMUSG0000002698 | chr2 | Wdsub1 | 0.069116094 | 0.8618497 |
| ENSMUSG0000000681 | chr17 | Sod2 | 0.069188071 | 0.8618497 |
| ENSMUSG0000001442 | chr17 | Map3k4 | 0.067682634 | 0.8618497 |
| ENSMUSG0000004142 | chr1 | Hibch | 0.082648823 | 0.8618497 |
| ENSMUSG0000003646 | chr8 | Marchf1 | 0.125438121 | 0.8618497 |
| ENSMUSG0000007887 | chr2 | Gm14295 | 0.100607981 | 0.8618497 |
| ENSMUSG0000003584 | chr15 | Alg12 | 0.08510039 | 0.8618497 |
| ENSMUSG0000000464 | chr5 | Slbp | 0.091576318 | 0.8618497 |
| ENSMUSG0000007267 | chr14 | Tmem254 | 0.123265965 | 0.8618497 |
| ENSMUSG0000000166 | chr10 | Ddt | 0.134138934 | 0.8618497 |
| ENSMUSG0000005139 | chr2 | Msantd7 | 0.123724132 | 0.8618497 |
| ENSMUSG0000011214 | chr10 | Lilrb4a | 0.372553447 | 0.8618497 |

| | | | | |
|---|---|---|---|---|
| ENSMUSG0000005979 | chr17 | Nrm | -0.325734784 | 0.86213463 |
| ENSMUSG0000003906 | chr3 | Zzz3 | -0.046167883 | 0.86213463 |
| ENSMUSG0000003961 | chr17 | Stub1 | 0.067729028 | 0.862319342 |
| ENSMUSG0000003433 | chr13 | Fam151b | 0.101863499 | 0.862603974 |
| ENSMUSG0000002722 | chr2 | Syt13 | -0.111643082 | 0.862624867 |
| ENSMUSG0000003473 | chr15 | Adgrb1 | -0.087380566 | 0.862624867 |
| ENSMUSG0000004113 | chr16 | Cyyr1 | -0.114800188 | 0.862624867 |
| ENSMUSG0000002939 | chr5 | Tmed2 | 0.101576469 | 0.86272483 |
| ENSMUSG0000002861 | chr4 | Ndc1 | -0.121865066 | 0.862788731 |
| ENSMUSG0000006220 | chr17 | Btbd9 | -0.083557493 | 0.862824546 |
| ENSMUSG0000002274 | chr16 | Tbc1d23 | 0.077415696 | 0.862824546 |
| ENSMUSG0000002064 | chr12 | Dus4l | 0.137516644 | 0.862824546 |
| ENSMUSG0000002964 | chr5 | Pan3 | -0.090926908 | 0.863159446 |
| ENSMUSG0000002383 | chr17 | Acat2 | 0.134349651 | 0.863159446 |
| ENSMUSG0000001855 | chr11 | Ybx2 | -0.164559254 | 0.863239995 |
| ENSMUSG0000008738 | chr4 | Frg2f1 | -0.121659528 | 0.863239995 |
| ENSMUSG0000002073 | chr11 | Rab37 | -0.148783704 | 0.863239995 |
| ENSMUSG0000004235 | chr15 | Grap2 | -0.368199765 | 0.863239995 |
| ENSMUSG0000002170 | chr13 | Zfyve16 | -0.069332252 | 0.863239995 |
| ENSMUSG0000003116 | chrX | Hdac6 | -0.079639486 | 0.863239995 |
| ENSMUSG0000000114 | chr1 | Lman2l | -0.06535029 | 0.863239995 |
| ENSMUSG0000001819 | chr1 | Glrx2 | -0.064543289 | 0.863239995 |
| ENSMUSG0000002971 | chr5 | Ephb4 | -0.061176943 | 0.863239995 |
| ENSMUSG0000002860 | chr4 | Czib | 0.064942733 | 0.863239995 |
| ENSMUSG0000003828 | chr10 | Ostm1 | 0.063947527 | 0.863239995 |
| ENSMUSG0000009052 | chr18 | Gypc | 0.077088524 | 0.863239995 |
| ENSMUSG0000002582 | chr6 | Pdia4 | 0.102665255 | 0.863239995 |
| ENSMUSG0000003080 | chr7 | Kat8 | 0.116310629 | 0.863239995 |
| ENSMUSG0000003234 | chr9 | Gsta4 | 0.102831399 | 0.863239995 |
| ENSMUSG0000003975 | chr4 | Thap3 | 0.092955888 | 0.863239995 |
| ENSMUSG0000000403 | chr1 | Brinp2 | 0.154025126 | 0.863239995 |
| ENSMUSG0000005496 | chr6 | Lmntd1 | -0.409001871 | 0.86333078 |
| ENSMUSG0000002627 | chr1 | Pask | -0.142675892 | 0.86333078 |
| ENSMUSG0000000745 | chr6 | M6pr | -0.064479953 | 0.86333078 |
| ENSMUSG0000005189 | chr12 | Klhdc1 | 0.091963412 | 0.86333078 |
| ENSMUSG0000002955 | chr5 | Psmg3 | 0.166600171 | 0.86333078 |
| ENSMUSG0000002683 | chr2 | Pierce1 | -0.296869884 | 0.863396373 |
| ENSMUSG0000002174 | chr14 | Cfap20dc | -0.232283117 | 0.863418952 |
| ENSMUSG0000002635 | chr1 | Ubxn4 | 0.050970811 | 0.863418952 |
| ENSMUSG0000004176 | chr14 | Ubac2 | 0.065245969 | 0.863418952 |
| ENSMUSG0000002018 | chr10 | Osbpl8 | 0.070613718 | 0.863418952 |
| ENSMUSG0000004048 | chr11 | Bptf | 0.08211404 | 0.863418952 |
| ENSMUSG0000002609 | chr1 | Pms1 | 0.122315381 | 0.863418952 |
| ENSMUSG0000005556 | chr13 | Zfp459 | 0.43514486 | 0.863418952 |
| ENSMUSG0000003079 | chr7 | Tead2 | -0.089770214 | 0.863572535 |
| ENSMUSG0000004681 | chr3 | Ddit4l | -0.142769916 | 0.863603396 |
| ENSMUSG0000003241 | chr9 | Tfdp2 | -0.083975485 | 0.863603396 |
| ENSMUSG0000000445 | chr5 | Ppp1cc | 0.056918047 | 0.863603396 |
| ENSMUSG0000000507 | chr12 | Jkamp | 0.069796387 | 0.863619156 |
| ENSMUSG0000003202 | chr9 | Ubash3b | -0.121929286 | 0.863747653 |

| | | | | |
|---|---|---|---|---|
| ENSMUSG0000003152 | chr8 | Tnks | -0.070739168 | 0.863747653 |
| ENSMUSG0000011018 | chr18 | Igip | -0.060515193 | 0.863747653 |
| ENSMUSG0000003872 | chr4 | Pakap | 0.294622364 | 0.863747653 |
| ENSMUSG0000002739 | chr2 | Polr1b | 0.081823668 | 0.863747653 |
| ENSMUSG0000003529 | chr7 | Klhl35 | 0.262021232 | 0.863747653 |
| ENSMUSG0000003946 | chr10 | Col10a1 | 0.342547121 | 0.863747653 |
| ENSMUSG0000003060 | chr7 | Zfp626 | -0.09225937 | 0.863785401 |
| ENSMUSG0000003620 | chr2 | Rif1 | -0.08972239 | 0.863785401 |
| ENSMUSG0000002874 | chr4 | Htr6 | -0.340362372 | 0.863800447 |
| ENSMUSG0000009091 | chr3 | Pabpc4l | -0.174148876 | 0.863894535 |
| ENSMUSG0000004745 | chr12 | Gphn | -0.065511851 | 0.863912639 |
| ENSMUSG0000002024 | chr10 | Nfyb | 0.073554457 | 0.864166912 |
| ENSMUSG0000007136 | chr10 | Map3k5 | 0.068186908 | 0.864186294 |
| ENSMUSG0000004203 | chr11 | Mat2b | 0.085590121 | 0.86420884 |
| ENSMUSG0000007533 | chr2 | Rprm | -0.143929917 | 0.864463801 |
| ENSMUSG0000006376 | chr15 | Chadl | -0.125808398 | 0.864463801 |
| ENSMUSG0000003150 | chr1 | Ptpn7 | -0.278410443 | 0.864463801 |
| ENSMUSG0000002508 | chr19 | Ablim1 | -0.080769837 | 0.864463801 |
| ENSMUSG0000002039 | chr11 | Itk | -0.370941165 | 0.864463801 |
| ENSMUSG0000002955 | chr5 | Sppl3 | -0.053486473 | 0.864463801 |
| ENSMUSG0000001854 | chr11 | Cwc25 | -0.059798445 | 0.864463801 |
| ENSMUSG0000001977 | chr10 | Hdac2 | 0.051921779 | 0.864463801 |
| ENSMUSG0000002523 | chr9 | Arih1 | 0.051745991 | 0.864463801 |
| ENSMUSG0000001565 | chr5 | Steap2 | 0.056134476 | 0.864463801 |
| ENSMUSG0000002649 | chr1 | Parp1 | 0.064852761 | 0.864463801 |
| ENSMUSG0000002288 | chr16 | Mrpl39 | 0.076600099 | 0.864463801 |
| ENSMUSG0000002515 | chr19 | Mms19 | 0.067966362 | 0.864463801 |
| ENSMUSG0000002494 | chr19 | Arl2 | 0.089538966 | 0.864463801 |
| ENSMUSG0000003435 | chr1 | Ramp1 | 0.120257425 | 0.864463801 |
| ENSMUSG0000004741 | chr13 | Cmya5 | 0.17965741 | 0.864463801 |
| ENSMUSG0000002868 | chr4 | Toe1 | 0.119723231 | 0.864463801 |
| ENSMUSG0000002802 | chr3 | Alpk1 | 0.187487336 | 0.864463801 |
| ENSMUSG0000002243 | chr15 | Ribc2 | 0.433902423 | 0.864463801 |
| ENSMUSG0000005062 | chr9 | Rps27rt | 0.225233794 | 0.864672252 |
| ENSMUSG0000000247 | chr18 | Snrpd1 | 0.114003582 | 0.864736306 |
| ENSMUSG0000000406 | chr16 | Dnaja3 | -0.064319305 | 0.864816067 |
| ENSMUSG0000005095 | chr19 | Insl6 | -0.224243618 | 0.864823953 |
| ENSMUSG0000004803 | chr3 | Isg20l2 | 0.064908617 | 0.864823953 |
| ENSMUSG0000006904 | chrY | Ddx3y | 0.065220209 | 0.864823953 |
| ENSMUSG0000006736 | chr5 | Lyar | 0.092155581 | 0.864823953 |
| ENSMUSG0000005829 | chr10 | Spock2 | -0.111007681 | 0.864884311 |
| ENSMUSG0000003317 | chr15 | Card10 | -0.106846298 | 0.864884311 |
| ENSMUSG0000002746 | chr2 | Tbc1d20 | 0.04887418 | 0.864884311 |
| ENSMUSG0000002492 | chr19 | Kat5 | 0.059285324 | 0.864884311 |
| ENSMUSG0000007492 | chr19 | Ptar1 | 0.067058879 | 0.86490214 |
| ENSMUSG0000000322 | chr19 | Grk5 | -0.104399241 | 0.86491957 |
| ENSMUSG0000002822 | chr4 | Rmdn1 | -0.096054072 | 0.86503872 |
| ENSMUSG0000002801 | chr3 | Tdo2 | -0.303708138 | 0.865191958 |
| ENSMUSG0000000166 | chr15 | Hoxc6 | -0.22852401 | 0.865191958 |
| ENSMUSG0000002175 | chr13 | Ddx4 | -0.387017855 | 0.865191958 |

| | | | | |
|---|---|---|---|---|
| ENSMUSG0000002275 | chr16 | P2rx6 | -0.113585096 | 0.865191958 |
| ENSMUSG0000004883 | chr19 | Vps37c | -0.06559514 | 0.865191958 |
| ENSMUSG0000008991 | chr3 | Mfsd14a | 0.0566059 | 0.865191958 |
| ENSMUSG0000006327 | chr3 | Naa15 | 0.06272715 | 0.865191958 |
| ENSMUSG0000004743 | chr16 | Xxylt1 | 0.067485978 | 0.865191958 |
| ENSMUSG0000007473 | chr2 | Syndig1 | 0.088158802 | 0.865191958 |
| ENSMUSG0000004159 | chr11 | Sdk2 | 0.097946822 | 0.865191958 |
| ENSMUSG0000002420 | chr17 | Acsbg2 | 0.35530098 | 0.865191958 |
| ENSMUSG0000004682 | chr7 | Fam187b | -0.332748597 | 0.865424819 |
| ENSMUSG0000004704 | chr19 | D030056L22Rik | 0.103428933 | 0.865424819 |
| ENSMUSG0000003096 | chr7 | Eef1akmt2 | 0.091349698 | 0.865424819 |
| ENSMUSG0000003610 | chr18 | Colec12 | 0.064388942 | 0.865449424 |
| ENSMUSG0000003088 | chr7 | Vwa3a | -0.23047842 | 0.865505572 |
| ENSMUSG0000002796 | chr3 | Col11a1 | -0.132587838 | 0.865505572 |
| ENSMUSG0000003957 | chr6 | Ccser1 | -0.166699235 | 0.865505572 |
| ENSMUSG0000007171 | chr15 | Ncf4 | -0.166367356 | 0.865505572 |
| ENSMUSG0000002789 | chr3 | Ahcyl1 | 0.042777754 | 0.865505572 |
| ENSMUSG0000005691 | chr19 | Sipa1 | 0.051557993 | 0.865505572 |
| ENSMUSG0000002053 | chr11 | Llgl1 | 0.060481165 | 0.865505572 |
| ENSMUSG0000005091 | chr11 | Cdr2l | -0.085172915 | 0.865737439 |
| ENSMUSG0000003385 | chr11 | Engase | -0.161139891 | 0.8657791 |
| ENSMUSG0000002100 | chr12 | Spata7 | -0.0863251 | 0.8657791 |
| ENSMUSG0000002161 | chr13 | Clptm1l | 0.075328856 | 0.8657791 |
| ENSMUSG0000007918 | chr14 | Mphosph8 | 0.068020013 | 0.8657791 |
| ENSMUSG0000004164 | chr12 | Ddx24 | 0.073181902 | 0.8657791 |
| ENSMUSG0000002410 | chr6 | Washc2 | 0.083258207 | 0.8657791 |
| ENSMUSG0000002971 | chr5 | Pcolce | 0.081498377 | 0.8657791 |
| ENSMUSG0000006356 | chr6 | Jazf1 | 0.075655127 | 0.8657791 |
| ENSMUSG0000009260 | chr3 | Scnm1 | 0.093118756 | 0.8657791 |
| ENSMUSG0000003893 | chr2 | Tcfl5 | 0.271093068 | 0.8657791 |
| ENSMUSG0000003460 | chr12 | 2700049A03Rik | 0.10682836 | 0.8657791 |
| ENSMUSG0000007934 | chr6 | C1s2 | 0.808734138 | 0.8657791 |
| ENSMUSG0000001004 | chr9 | Zmynd10 | -0.360275717 | 0.865922584 |
| ENSMUSG0000002586 | chrX | Stag2 | 0.065395743 | 0.866044131 |
| ENSMUSG0000002491 | chr19 | Lrp5 | 0.056778432 | 0.866044131 |
| ENSMUSG0000003376 | chr3 | Tmem131l | 0.137666798 | 0.866398605 |
| ENSMUSG0000000148 | chr8 | Def8 | 0.060980395 | 0.866775914 |
| ENSMUSG0000004597 | chr4 | Slc25a51 | 0.04879145 | 0.866811942 |
| ENSMUSG0000000479 | chr11 | Ulk2 | 0.059223859 | 0.866934039 |
| ENSMUSG0000002266 | chr16 | Atg3 | 0.080861566 | 0.866934039 |
| ENSMUSG0000002388 | chr17 | Thbs2 | 0.097190768 | 0.866937495 |
| ENSMUSG0000006166 | chr17 | Cd2ap | -0.099601056 | 0.867061537 |
| ENSMUSG0000005473 | chrX | Zfp182 | -0.089416853 | 0.867061537 |
| ENSMUSG0000000790 | chr9 | Hmgcll1 | 0.099747698 | 0.867105599 |
| ENSMUSG0000002326 | chr17 | Frs3 | -0.079132659 | 0.86720043 |
| ENSMUSG0000002198 | chr14 | Cdadc1 | 0.065520968 | 0.86720043 |
| ENSMUSG0000011010 | chr7 | Gm45844 | -0.15814216 | 0.867276882 |
| ENSMUSG0000003757 | chr14 | Wdhd1 | -0.174898874 | 0.867276882 |
| ENSMUSG0000004061 | chr4 | Tmem51 | -0.110076379 | 0.867276882 |
| ENSMUSG0000002885 | chr4 | Slc9a1 | -0.08051936 | 0.867276882 |

| | | | | |
|---|---|---|---|---|
| ENSMUSG0000004823 | chr1 | Rnf149 | -0.101333388 | 0.867276882 |
| ENSMUSG0000005506 | chr15 | Ddx17 | -0.050418012 | 0.867276882 |
| ENSMUSG0000002254 | chr16 | Eef2kmt | 0.090771948 | 0.867276882 |
| ENSMUSG0000002049 | chr11 | Prr11 | -0.169171296 | 0.867283032 |
| ENSMUSG0000010957 | chr5 | Cfap99 | -0.356111529 | 0.867283032 |
| ENSMUSG0000002776 | chr3 | P2ry1 | -0.101913917 | 0.867283032 |
| ENSMUSG0000003264 | chr1 | Colgalt2 | -0.095988344 | 0.867283032 |
| ENSMUSG0000007419 | chr8 | Zfp791 | -0.127053736 | 0.867283032 |
| ENSMUSG0000002112 | chr12 | Rdh12 | -0.21780691 | 0.867283032 |
| ENSMUSG0000002733 | chr2 | Smox | -0.07876551 | 0.867283032 |
| ENSMUSG0000002528 | chrX | Sat1 | -0.05123283 | 0.867283032 |
| ENSMUSG0000003087 | chr7 | Mfsd13b | -0.43999836 | 0.867283032 |
| ENSMUSG0000006787 | chrX | Htatsf1 | 0.050278572 | 0.867283032 |
| ENSMUSG0000002651 | chr1 | Nvl | 0.057100602 | 0.867283032 |
| ENSMUSG0000005778 | chr8 | Ddx49 | 0.062466014 | 0.867283032 |
| ENSMUSG0000003271 | chr10 | Syde1 | 0.069818614 | 0.867283032 |
| ENSMUSG0000005312 | chr9 | Rnf26 | 0.175631313 | 0.867283032 |
| ENSMUSG0000004269 | chr19 | Stn1 | 0.096057691 | 0.867283032 |
| ENSMUSG0000007297 | chr12 | Gm4787 | 0.205890577 | 0.867283032 |
| ENSMUSG0000003231 | chr9 | Cyp1a1 | 0.677054562 | 0.867283032 |
| ENSMUSG0000004901 | chr3 | Prss48 | 0.279400269 | 0.867367919 |
| ENSMUSG0000003028 | chr6 | Itpr2 | -0.137750712 | 0.867800644 |
| ENSMUSG0000003012 | chr6 | Tmcc1 | -0.076882545 | 0.867800644 |
| ENSMUSG0000002674 | chr2 | Mllt10 | -0.068673028 | 0.867800644 |
| ENSMUSG0000002282 | chr16 | Gtf2e1 | 0.082133813 | 0.867800644 |
| ENSMUSG0000002148 | chr13 | Mxd3 | 0.436441126 | 0.867800644 |
| ENSMUSG0000000105 | chr11 | N4bp3 | 0.095914343 | 0.867800644 |
| ENSMUSG0000003877 | chr1 | Ephx1 | 0.13147417 | 0.867800644 |
| ENSMUSG0000002251 | chr16 | 4930562C15Rik | -0.16931831 | 0.868303074 |
| ENSMUSG0000007313 | chrX | Pasd1 | -0.339997818 | 0.868303074 |
| ENSMUSG0000002202 | chr14 | Cnmd | -0.301428081 | 0.868303074 |
| ENSMUSG0000001662 | chr15 | Phf21b | -0.149222938 | 0.868303074 |
| ENSMUSG0000003465 | chr18 | Ythdc2 | -0.08176606 | 0.868303074 |
| ENSMUSG0000002442 | chr18 | Ndfip1 | -0.077205442 | 0.868303074 |
| ENSMUSG0000000589 | chr6 | Nr2c2 | -0.075847319 | 0.868303074 |
| ENSMUSG0000002503 | chr19 | Trim8 | -0.061339407 | 0.868303074 |
| ENSMUSG0000002106 | chr12 | Fut8 | -0.062909703 | 0.868303074 |
| ENSMUSG0000002202 | chr14 | Wbp4 | 0.044762726 | 0.868303074 |
| ENSMUSG0000002628 | chr1 | Atg4b | 0.058128803 | 0.868303074 |
| ENSMUSG0000003212 | chr9 | Vps11 | 0.061329362 | 0.868303074 |
| ENSMUSG0000002856 | chr4 | Usp1 | 0.067576255 | 0.868303074 |
| ENSMUSG0000002429 | chr18 | Rock1 | 0.06707355 | 0.868303074 |
| ENSMUSG0000004597 | chr16 | C2cd2 | 0.066912196 | 0.868303074 |
| ENSMUSG0000002619 | chr1 | Ankzf1 | 0.07144021 | 0.868303074 |
| ENSMUSG0000003793 | chr2 | Chchd5 | 0.106843179 | 0.868303074 |
| ENSMUSG0000003849 | chr1 | Slc19a3 | 0.239548839 | 0.868303074 |
| ENSMUSG0000004979 | chr7 | Fzd4 | 0.137752437 | 0.868303074 |
| ENSMUSG0000004017 | chr1 | Fmo2 | 0.302200526 | 0.868303074 |
| ENSMUSG0000002824 | chr4 | Faxc | 0.0957485 | 0.868385002 |
| ENSMUSG0000002611 | chr1 | Mgat4a | -0.084537312 | 0.868385055 |

| | | | | |
|---|---|---|---|---|
| ENSMUSG0000000149 | chr11 | Meox1 | 0.148611598 | 0.868385055 |
| ENSMUSG0000003645 | chr7 | Wtip | 0.108459238 | 0.868570397 |
| ENSMUSG0000003622 | chr18 | Kctd1 | 0.087004417 | 0.868753216 |
| ENSMUSG0000004790 | chr2 | Ankrd16 | -0.118703079 | 0.869336183 |
| ENSMUSG0000005346 | chr1 | Hhipl2 | -0.355530761 | 0.869336183 |
| ENSMUSG0000005809 | chr15 | Nfam1 | -0.183870927 | 0.869336183 |
| ENSMUSG0000002125 | chr12 | Fos | -0.190828212 | 0.869336183 |
| ENSMUSG0000003209 | chr9 | Mpzl2 | -0.204864188 | 0.869336183 |
| ENSMUSG0000000387 | chr7 | Lin7b | -0.12501844 | 0.869336183 |
| ENSMUSG0000005362 | chr8 | Tll1 | -0.118996563 | 0.869336183 |
| ENSMUSG0000008484 | chr4 | Tmem240 | -0.178106354 | 0.869336183 |
| ENSMUSG0000001060 | chr12 | Rbm25 | -0.076538117 | 0.869336183 |
| ENSMUSG0000006175 | chr5 | Bod1l | -0.088570883 | 0.869336183 |
| ENSMUSG0000001183 | chr8 | Evi5l | -0.074234931 | 0.869336183 |
| ENSMUSG0000001957 | chr17 | Ubxn6 | -0.07178347 | 0.869336183 |
| ENSMUSG0000004402 | chr18 | Rell2 | -0.195853813 | 0.869336183 |
| ENSMUSG0000005494 | chr3 | Miga1 | -0.06372074 | 0.869336183 |
| ENSMUSG0000002598 | chr1 | Slc39a10 | -0.056821557 | 0.869336183 |
| ENSMUSG0000004250 | chr5 | Dmtf1 | -0.050596058 | 0.869336183 |
| ENSMUSG0000000546 | chr8 | Prkaca | -0.051074484 | 0.869336183 |
| ENSMUSG0000003970 | chr11 | Nploc4 | 0.042178939 | 0.869336183 |
| ENSMUSG0000006128 | chr5 | Taok3 | 0.054097015 | 0.869336183 |
| ENSMUSG0000002213 | chr14 | Tgds | 0.075860905 | 0.869336183 |
| ENSMUSG0000002147 | chr13 | Habp4 | 0.059438891 | 0.869336183 |
| ENSMUSG0000002841 | chr4 | Aptx | 0.063690882 | 0.869336183 |
| ENSMUSG0000002745 | chr2 | Nsfl1c | 0.069692599 | 0.869336183 |
| ENSMUSG0000002049 | chr11 | Smg8 | 0.069398991 | 0.869336183 |
| ENSMUSG0000005997 | chr7 | Zfp74 | 0.068222549 | 0.869336183 |
| ENSMUSG0000002757 | chr2 | Nkain4 | 0.176663726 | 0.869336183 |
| ENSMUSG0000000106 | chr8 | Vps9d1 | 0.062799875 | 0.869336183 |
| ENSMUSG0000002160 | chr13 | Mrpl36 | 0.112211946 | 0.869336183 |
| ENSMUSG0000003135 | chrX | Hccs | 0.081681537 | 0.869336183 |
| ENSMUSG0000003172 | chr8 | Txnl4b | 0.075850293 | 0.869336183 |
| ENSMUSG0000002945 | chr5 | Aldh2 | 0.09092253 | 0.869336183 |
| ENSMUSG0000006918 | chr13 | Zfp72 | 0.098761196 | 0.869336183 |
| ENSMUSG0000002806 | chr3 | Pmf1 | 0.115132662 | 0.869336183 |
| ENSMUSG0000004509 | chr3 | S1pr1 | 0.091229096 | 0.869336183 |
| ENSMUSG0000004015 | chr2 | Thbs1 | 0.243079801 | 0.869336183 |
| ENSMUSG0000002131 | chr13 | Ryr2 | 0.115615752 | 0.869336183 |
| ENSMUSG0000002858 | chr4 | Laptm5 | 0.108212313 | 0.869336183 |
| ENSMUSG0000004178 | chr12 | Cpsf2 | 0.047656257 | 0.869410388 |
| ENSMUSG0000000318 | chr7 | Irf3 | 0.071010176 | 0.869410388 |
| ENSMUSG0000000473 | chr17 | Adgre1 | -0.11256153 | 0.869449489 |
| ENSMUSG0000003175 | chr8 | Nudt21 | 0.062701286 | 0.869449489 |
| ENSMUSG0000000273 | chr2 | Plekha3 | 0.078053438 | 0.869449489 |
| ENSMUSG0000000200 | chrX | Srpk3 | 0.09217696 | 0.869449489 |
| ENSMUSG0000002104 | chr12 | Mthfd1 | 0.084249004 | 0.869530506 |
| ENSMUSG0000006727 | chr5 | Rplp0 | 0.280617887 | 0.869876282 |
| ENSMUSG0000002681 | chr2 | Gtf3c5 | 0.051251377 | 0.869900863 |
| ENSMUSG0000004461 | chr10 | Zbtb39 | 0.060720325 | 0.869900863 |

| | | | | |
|---|---|---|---|---|
| ENSMUSG0000004796 | chr5 | Stbd1 | 0.11668225 | 0.869900863 |
| ENSMUSG0000002855 | chr4 | Calr4 | -0.448414156 | 0.86998259 |
| ENSMUSG0000007033 | chr11 | Fbxo47 | -0.337849578 | 0.86998259 |
| ENSMUSG0000003362 | chr8 | Pdpr | -0.104402253 | 0.86998259 |
| ENSMUSG0000003573 | chr19 | Dagla | -0.10761252 | 0.86998259 |
| ENSMUSG0000003064 | chr7 | Anapc15 | -0.113724629 | 0.86998259 |
| ENSMUSG0000002846 | chr4 | Rgp1 | -0.078371667 | 0.86998259 |
| ENSMUSG0000005694 | chr2 | Commd7 | -0.068245592 | 0.86998259 |
| ENSMUSG0000003855 | chr18 | Reep2 | -0.08002795 | 0.86998259 |
| ENSMUSG0000002532 | chr7 | Ube3a | -0.06238885 | 0.86998259 |
| ENSMUSG0000000646 | chr19 | Zdhhc24 | -0.067296199 | 0.86998259 |
| ENSMUSG0000002433 | chr17 | Col11a2 | -0.110641578 | 0.86998259 |
| ENSMUSG0000007154 | chr14 | 3425401B19Rik | 0.372252598 | 0.86998259 |
| ENSMUSG0000001837 | chr11 | Vezf1 | 0.05853648 | 0.86998259 |
| ENSMUSG0000001430 | chr16 | Glis2 | 0.059909012 | 0.86998259 |
| ENSMUSG0000003516 | chr2 | Tanc1 | 0.066704483 | 0.86998259 |
| ENSMUSG0000004878 | chr7 | Dcun1d3 | 0.0739174 | 0.86998259 |
| ENSMUSG0000004235 | chr2 | Osbpl6 | 0.079903576 | 0.86998259 |
| ENSMUSG0000001415 | chr5 | Trpv4 | 0.127373876 | 0.86998259 |
| ENSMUSG0000007899 | chr13 | Zfp429 | -0.157453879 | 0.870023353 |
| ENSMUSG0000003676 | chr15 | Kcnk9 | -0.177476281 | 0.870206457 |
| ENSMUSG0000003521 | chr7 | Xrra1 | -0.270272068 | 0.870206457 |
| ENSMUSG0000002400 | chr17 | Brd4 | -0.083401785 | 0.8702615 |
| ENSMUSG0000002793 | chr3 | Rab13 | 0.135591915 | 0.8702615 |
| ENSMUSG0000004197 | chr2 | Mettl8 | 0.091006394 | 0.8702615 |
| ENSMUSG0000003088 | chr7 | Pdzd9 | -0.257507061 | 0.870399481 |
| ENSMUSG0000000940 | chrX | Elk1 | 0.065817859 | 0.870399481 |
| ENSMUSG0000002159 | chr13 | Glrx | -0.10771953 | 0.870448218 |
| ENSMUSG0000003886 | chr1 | Zcchc2 | -0.085688163 | 0.870448218 |
| ENSMUSG0000003815 | chr11 | Ormdl3 | 0.056016875 | 0.870448218 |
| ENSMUSG0000001749 | chr11 | Cdc6 | -0.301369594 | 0.870456548 |
| ENSMUSG0000001979 | chr10 | Trmt11 | 0.073111772 | 0.870456548 |
| ENSMUSG0000003996 | chr5 | Rsbn1l | 0.06397254 | 0.870456548 |
| ENSMUSG0000002113 | chr12 | S2bpcox16 | 0.319070391 | 0.870456548 |
| ENSMUSG0000004470 | chr7 | Palb2 | 0.19369974 | 0.870456548 |
| ENSMUSG0000002068 | chr11 | Gas2l2 | 0.154152862 | 0.870456548 |
| ENSMUSG0000001783 | chr17 | Rab5a | 0.052395589 | 0.870515678 |
| ENSMUSG0000004960 | chr1 | Gpr55 | -0.321309085 | 0.87059209 |
| ENSMUSG0000002461 | chr18 | Tmx3 | -0.050646208 | 0.87059209 |
| ENSMUSG0000003824 | chr10 | Pdss2 | 0.09090432 | 0.87059209 |
| ENSMUSG0000011021 | chr17 | Tincr | 0.210450535 | 0.87059209 |
| ENSMUSG0000002445 | chr18 | Arap3 | 0.122141776 | 0.87059209 |
| ENSMUSG0000002801 | chr3 | Pdgfc | -0.065449496 | 0.870694813 |
| ENSMUSG0000002047 | chr11 | Pgam2 | 0.17750137 | 0.870694813 |
| ENSMUSG0000002723 | chr2 | Creb3l1 | -0.279594279 | 0.87070379 |
| ENSMUSG0000002900 | chr4 | Fbxo44 | -0.090392596 | 0.87070379 |
| ENSMUSG0000003196 | chr8 | Afg3l1 | 0.054084102 | 0.87070379 |
| ENSMUSG0000004115 | chr4 | Osgin2 | 0.075011477 | 0.87070379 |
| ENSMUSG0000003942 | chr7 | Tmem135 | -0.051151117 | 0.870740897 |
| ENSMUSG0000002211 | chr14 | Gpc5 | -0.64568818 | 0.870767344 |

| | | | | |
|---|---|---|---|---|
| ENSMUSG0000001378 | chr17 | Ehmt2 | -0.059420017 | 0.870767344 |
| ENSMUSG0000004353 | chr2 | Setx | -0.074416373 | 0.87076952 |
| ENSMUSG0000006402 | chr7 | Klk8 | 0.2897099 | 0.870955085 |
| ENSMUSG0000003771 | chr9 | Ccdc33 | -0.349387212 | 0.870956223 |
| ENSMUSG0000000380 | chr8 | Gcdh | -0.091449734 | 0.871003381 |
| ENSMUSG0000004086 | chr1 | Ino80d | -0.082970077 | 0.871052875 |
| ENSMUSG0000004471 | chr12 | Slc38a6 | -0.086814611 | 0.871052875 |
| ENSMUSG0000006258 | chr4 | Cnr2 | -0.428426614 | 0.871101545 |
| ENSMUSG0000003422 | chr8 | Slc38a8 | -0.231580234 | 0.871101545 |
| ENSMUSG0000002346 | chr7 | Tulp2 | 0.399516617 | 0.871101545 |
| ENSMUSG0000002037 | chr11 | Ltc4s | 0.207438731 | 0.871101545 |
| ENSMUSG0000004741 | chr12 | Gpr68 | 0.21622207 | 0.871101545 |
| ENSMUSG0000006644 | chr9 | Mthfs | -0.132682236 | 0.87119356 |
| ENSMUSG0000003341 | chr19 | Cacul1 | -0.05509858 | 0.87119356 |
| ENSMUSG0000002870 | chr4 | Mknk1 | 0.072975574 | 0.87119356 |
| ENSMUSG0000002305 | chr15 | Npff | 0.229259274 | 0.87119356 |
| ENSMUSG0000003031 | chr6 | Slc6a1 | 0.234797792 | 0.87119356 |
| ENSMUSG0000004540 | chr12 | Kcnk13 | -0.182905008 | 0.871650357 |
| ENSMUSG0000003869 | chr7 | Josd2 | 0.072655984 | 0.871650357 |
| ENSMUSG0000002223 | chr15 | Sema5a | -0.148940138 | 0.871683382 |
| ENSMUSG0000004525 | chr7 | Zfp574 | 0.054852907 | 0.871683382 |
| ENSMUSG0000003663 | chr5 | Nudt1 | 0.129091504 | 0.871683382 |
| ENSMUSG0000002826 | chr3 | Clca3a2 | 0.134402785 | 0.871683382 |
| ENSMUSG0000006615 | chr4 | Slc31a2 | -0.090371769 | 0.871745916 |
| ENSMUSG0000003424 | chr11 | Plekhm1 | -0.066933361 | 0.871745916 |
| ENSMUSG0000003198 | chr9 | Vps26b | -0.063193516 | 0.871745916 |
| ENSMUSG0000009193 | chr12 | Gon7 | 0.150799611 | 0.871745916 |
| ENSMUSG0000004957 | chr8 | Zfpm1 | 0.108178632 | 0.871745916 |
| ENSMUSG0000006380 | chr10 | Lin28b | -0.239505094 | 0.87189602 |
| ENSMUSG0000002238 | chr15 | Ppara | -0.119008731 | 0.87189602 |
| ENSMUSG0000004520 | chr14 | Lrrc3b | 0.173874054 | 0.87189602 |
| ENSMUSG0000002557 | chr11 | Tmc6 | -0.143338489 | 0.872099748 |
| ENSMUSG0000007139 | chr12 | 2410004P03Rik | 0.186373623 | 0.87212778 |
| ENSMUSG0000002127 | chr14 | Fdft1 | -0.108739979 | 0.872129614 |
| ENSMUSG0000002532 | chr12 | Itgb8 | -0.101208218 | 0.872129614 |
| ENSMUSG0000003351 | chr7 | Otud7a | -0.147055487 | 0.872129614 |
| ENSMUSG0000003240 | chr9 | U2surp | -0.067575069 | 0.872129614 |
| ENSMUSG0000004252 | chr12 | Dnal1 | -0.068737272 | 0.872129614 |
| ENSMUSG0000004094 | chr7 | Arhgef1 | -0.067078725 | 0.872129614 |
| ENSMUSG0000001866 | chr11 | Cbx1 | -0.062783919 | 0.872129614 |
| ENSMUSG0000002291 | chr16 | Pros1 | -0.064121382 | 0.872129614 |
| ENSMUSG0000001976 | chr10 | Rmnd1 | -0.06120509 | 0.872129614 |
| ENSMUSG0000005669 | chr17 | Ilrun | 0.041890304 | 0.872129614 |
| ENSMUSG0000002740 | chr2 | Idh3b | 0.055324129 | 0.872129614 |
| ENSMUSG0000002255 | chr15 | Sharpin | 0.048595582 | 0.872129614 |
| ENSMUSG0000005053 | chr2 | Fam171a1 | 0.069749976 | 0.872129614 |
| ENSMUSG0000004056 | chr3 | Gstm2 | 0.109336684 | 0.872129614 |
| ENSMUSG0000002124 | chr12 | Abcd4 | 0.082824528 | 0.872129614 |
| ENSMUSG0000002753 | chr3 | Fabp5 | 0.119936306 | 0.872129614 |
| ENSMUSG0000002897 | chr5 | Abcb8 | 0.087705735 | 0.872129614 |

| | | | | |
|---|---|---|---|---|
| ENSMUSG0000002469 | chr19 | Lpxn | 0.178839595 | 0.872129614 |
| ENSMUSG0000004260 | chr6 | Asb4 | 0.276715928 | 0.872129614 |
| ENSMUSG0000003234 | chr9 | Cgas | 0.198863003 | 0.872129614 |
| ENSMUSG0000007860 | chr7 | Gvin2 | 0.480634171 | 0.872129614 |
| ENSMUSG0000012160 | chrX | Nlgn4l | 0.062483065 | 0.872131469 |
| ENSMUSG0000000336 | chr15 | Ddx23 | 0.047747155 | 0.872291241 |
| ENSMUSG0000002026 | chr11 | Lyrm7 | -0.12751268 | 0.872500064 |
| ENSMUSG0000002895 | chr4 | Vamp3 | 0.055430509 | 0.872500064 |
| ENSMUSG0000004325 | chr10 | Fam13c | 0.068114717 | 0.872500064 |
| ENSMUSG0000002743 | chr2 | Dtd1 | -0.123147545 | 0.872606584 |
| ENSMUSG0000001384 | chr15 | St3gal1 | -0.104258688 | 0.872606584 |
| ENSMUSG0000003260 | chr9 | Slc25a20 | -0.105691115 | 0.872606584 |
| ENSMUSG0000005592 | chr2 | Gm14137 | -0.178338637 | 0.872606584 |
| ENSMUSG0000003198 | chr8 | Sprtn | -0.0667469 | 0.872606584 |
| ENSMUSG0000002462 | chr18 | Hmgxb3 | -0.065305742 | 0.872606584 |
| ENSMUSG0000002502 | chr19 | Add3 | -0.062460419 | 0.872606584 |
| ENSMUSG0000001536 | chr15 | Trabd | -0.06770633 | 0.872606584 |
| ENSMUSG0000002967 | chr6 | Fam3c | 0.054309552 | 0.872606584 |
| ENSMUSG0000003204 | chr9 | Dcps | 0.084310539 | 0.872606584 |
| ENSMUSG0000001005 | chr9 | Nprl2 | 0.074304628 | 0.872606584 |
| ENSMUSG0000001596 | chr14 | Il17rb | 0.11299298 | 0.872606584 |
| ENSMUSG0000004852 | chr9 | Cxcr6 | 0.235768684 | 0.872606584 |
| ENSMUSG0000000307 | chr10 | Efna2 | 0.108882815 | 0.872606584 |
| ENSMUSG0000010138 | chr19 | Ms4a4a | 0.208941253 | 0.872606584 |
| ENSMUSG0000004744 | chr1 | Erfe | 0.298135673 | 0.872606584 |
| ENSMUSG0000004802 | chr19 | Eno4 | 0.205539359 | 0.872606584 |
| ENSMUSG0000003013 | chr6 | Tuba8 | 0.336330305 | 0.872606584 |
| ENSMUSG0000001652 | chr1 | Dyrk3 | 0.1553273 | 0.872632962 |
| ENSMUSG0000003872 | chr15 | Pkhd1l1 | -0.132131599 | 0.87270575 |
| ENSMUSG0000001066 | chr19 | Fads1 | -0.073683142 | 0.87270575 |
| ENSMUSG0000003472 | chr5 | Cnot6l | -0.069483053 | 0.87270575 |
| ENSMUSG0000004952 | chr9 | Tmem202 | -0.251738995 | 0.87270575 |
| ENSMUSG0000004659 | chr7 | Ticrr | -0.464334367 | 0.87270575 |
| ENSMUSG0000002288 | chr16 | Eif4a2 | 0.065996704 | 0.87270575 |
| ENSMUSG0000002903 | chr4 | Ints11 | 0.07247251 | 0.87270575 |
| ENSMUSG0000005061 | chr2 | Zscan29 | 0.060681771 | 0.87270575 |
| ENSMUSG0000005573 | chr15 | Ghr | 0.086192505 | 0.87270575 |
| ENSMUSG0000002302 | chr15 | Cers5 | 0.055456491 | 0.87270575 |
| ENSMUSG0000002840 | chr4 | Smim27 | -0.18494359 | 0.872712145 |
| ENSMUSG0000002780 | chr3 | Ppid | -0.062000673 | 0.872712145 |
| ENSMUSG0000003636 | chr3 | P2ry13 | 0.141257861 | 0.872712145 |
| ENSMUSG0000005333 | chr1 | Dis3l2 | -0.082770045 | 0.872793229 |
| ENSMUSG0000000184 | chr5 | Rac1 | 0.049594464 | 0.872793229 |
| ENSMUSG0000009599 | chr17 | Zfp97 | 0.103579782 | 0.872997781 |
| ENSMUSG0000002220 | chr15 | Npr3 | 0.186448824 | 0.872997781 |
| ENSMUSG0000003052 | chr7 | Blm | -0.214955908 | 0.873278782 |
| ENSMUSG0000004791 | chr18 | Pcdhb16 | -0.139473554 | 0.873278782 |
| ENSMUSG0000004086 | chr4 | Crocc | -0.089510875 | 0.873278782 |
| ENSMUSG0000003440 | chrX | Pja1 | -0.073510618 | 0.873278782 |
| ENSMUSG0000006353 | chr5 | Sema3e | -0.076586599 | 0.873278782 |

| | | | | |
|---|---|---|---|---|
| ENSMUSG0000003888 | chr7 | Man2a2 | -0.056857873 | 0.873278782 |
| ENSMUSG0000002942 | chr5 | Rsrc2 | 0.089307309 | 0.873278782 |
| ENSMUSG0000002263 | chr16 | Cblb | 0.069163704 | 0.873278782 |
| ENSMUSG0000002746 | chr2 | Angpt4 | 0.330386597 | 0.873278782 |
| ENSMUSG0000005728 | chr4 | Musk | 0.125637969 | 0.873278782 |
| ENSMUSG0000003192 | chr8 | Cyb5b | -0.072492184 | 0.873480817 |
| ENSMUSG0000003831 | chr2 | Edem2 | -0.065049904 | 0.873480817 |
| ENSMUSG0000002458 | chr18 | Txnl1 | 0.054723792 | 0.873480817 |
| ENSMUSG0000004744 | chr12 | Arl4a | 0.085730775 | 0.873480817 |
| ENSMUSG0000000198 | chr11 | Taco1 | 0.099306742 | 0.873480817 |
| ENSMUSG0000003203 | chr9 | Barx2 | 0.267812305 | 0.873518211 |
| ENSMUSG0000003646 | chr14 | Elf1 | 0.051511566 | 0.873518211 |
| ENSMUSG0000004278 | chr9 | Exog | 0.114634564 | 0.873518211 |
| ENSMUSG0000008983 | chr7 | Shkbp1 | -0.064843034 | 0.873668807 |
| ENSMUSG0000003328 | chr3 | Wdr3 | 0.056416232 | 0.873893744 |
| ENSMUSG0000005031 | chr15 | Rictor | -0.061563482 | 0.87395493 |
| ENSMUSG0000002000 | chr10 | Pex7 | -0.083582681 | 0.874082856 |
| ENSMUSG0000004985 | chr10 | Suox | -0.0547355 | 0.874082856 |
| ENSMUSG0000003600 | chr4 | Atosb | 0.063579352 | 0.874082856 |
| ENSMUSG0000004748 | chrX | Klhl34 | -0.185422393 | 0.874195779 |
| ENSMUSG0000002861 | chr4 | Lrrc42 | -0.098977587 | 0.874195779 |
| ENSMUSG0000005800 | chr4 | Mdn1 | -0.091132068 | 0.874195779 |
| ENSMUSG0000003155 | chr8 | Adam5 | -0.297701494 | 0.874195779 |
| ENSMUSG0000002763 | chr3 | Tbl1xr1 | -0.069662036 | 0.874195779 |
| ENSMUSG0000004172 | chr16 | Pi4ka | -0.064067586 | 0.874195779 |
| ENSMUSG0000002869 | chr4 | Ipp | -0.076323986 | 0.874195779 |
| ENSMUSG0000003539 | chr7 | Brsk1 | -0.069290758 | 0.874195779 |
| ENSMUSG0000000405 | chr7 | Akt2 | 0.060460125 | 0.874195779 |
| ENSMUSG0000002764 | chr2 | Ctnnbl1 | 0.057373037 | 0.874195779 |
| ENSMUSG0000005008 | chr17 | 1600012H06Rik | 0.070764127 | 0.874195779 |
| ENSMUSG0000000176 | chr6 | Smo | 0.075212731 | 0.874195779 |
| ENSMUSG0000003230 | chr9 | 1700017B05Rik | 0.068376388 | 0.874195779 |
| ENSMUSG0000004013 | chr7 | 9430038I01Rik | 0.140368289 | 0.874195779 |
| ENSMUSG0000001512 | chr17 | Tsr3 | 0.080044944 | 0.874375759 |
| ENSMUSG0000005114 | chr8 | Nat2 | -0.131425555 | 0.874535818 |
| ENSMUSG0000006003 | chr8 | Dhps | 0.075248598 | 0.874535818 |
| ENSMUSG0000003656 | chr15 | Ppp6r2 | -0.051573384 | 0.874717942 |
| ENSMUSG0000006218 | chrX | Hs6st2 | 0.265010946 | 0.874717942 |
| ENSMUSG0000006003 | chr6 | H2aj | -0.098117942 | 0.874990904 |
| ENSMUSG0000002938 | chr5 | Shroom3 | -0.112305998 | 0.874990904 |
| ENSMUSG0000001763 | chr11 | Rab11fip4 | 0.131939587 | 0.874990904 |
| ENSMUSG0000003274 | chr5 | Tpcn1 | 0.06003658 | 0.875028419 |
| ENSMUSG0000002240 | chr15 | St13 | 0.104015533 | 0.875028419 |
| ENSMUSG0000003528 | chr9 | Vps13c | -0.095221203 | 0.875125135 |
| ENSMUSG0000003908 | chr13 | Rreb1 | -0.08941604 | 0.875125135 |
| ENSMUSG0000004028 | chr11 | Btnl9 | -0.152108183 | 0.875125135 |
| ENSMUSG0000004774 | chr8 | Rnf150 | -0.077541564 | 0.875125135 |
| ENSMUSG0000005932 | chrX | Eda | -0.092183119 | 0.875125135 |
| ENSMUSG0000003077 | chr7 | Dkk3 | 0.041452425 | 0.875125135 |
| ENSMUSG0000003073 | chr7 | Sh2b1 | 0.056314437 | 0.875125135 |

| | | | | |
|---|---|---|---|---|
| ENSMUSG0000007851 | chr4 | Emc1 | 0.063341377 | 0.875125135 |
| ENSMUSG0000006168 | chr2 | Dlgap4 | 0.06314206 | 0.875125135 |
| ENSMUSG0000003219 | chr9 | Dock6 | 0.074163136 | 0.875125135 |
| ENSMUSG0000004016 | chr7 | Ikzf5 | 0.116183489 | 0.875125135 |
| ENSMUSG0000003148 | chr8 | Adrb3 | 0.137848083 | 0.875125135 |
| ENSMUSG0000007465 | chr3 | Lrrc31 | -0.633035395 | 0.875143417 |
| ENSMUSG0000000210 | chr2 | Nr1h3 | -0.113171951 | 0.875143417 |
| ENSMUSG0000003221 | chr9 | Sltm | -0.066618512 | 0.875143417 |
| ENSMUSG0000004290 | chr1 | Aida | 0.046434484 | 0.875143417 |
| ENSMUSG0000000973 | chr15 | Pou6f1 | 0.083417786 | 0.875143417 |
| ENSMUSG0000000673 | chr10 | Tspan31 | 0.083184565 | 0.875143417 |
| ENSMUSG0000002632 | chr1 | Tnfrsf11a | 0.086873908 | 0.875143417 |
| ENSMUSG0000003417 | chr11 | Rhbdd3 | 0.088688527 | 0.875143417 |
| ENSMUSG0000001989 | chr10 | Lama2 | 0.164219509 | 0.875143417 |
| ENSMUSG0000003953 | chr8 | Sgcz | 0.259103431 | 0.875143417 |
| ENSMUSG0000001288 | chr8 | Podnl1 | -0.278590085 | 0.875247941 |
| ENSMUSG0000010749 | chr6 | Ccdc142 | -0.116496765 | 0.875247941 |
| ENSMUSG0000002522 | chr19 | Actr1a | -0.038139629 | 0.875247941 |
| ENSMUSG0000002429 | chr18 | Esco1 | 0.049116274 | 0.875247941 |
| ENSMUSG0000002743 | chr2 | Xrn2 | 0.063278917 | 0.875247941 |
| ENSMUSG0000003764 | chr7 | Zfp60 | 0.07359135 | 0.875247941 |
| ENSMUSG0000005613 | chr9 | Pgm3 | 0.092975408 | 0.875247941 |
| ENSMUSG0000005876 | chr7 | Rnf169 | 0.075706868 | 0.875247941 |
| ENSMUSG0000005035 | chr14 | Gpr18 | 0.26229 | 0.875247941 |
| ENSMUSG0000003358 | chr7 | Ndn | -0.082674942 | 0.875422209 |
| ENSMUSG0000003748 | chr15 | Ubr5 | -0.058047484 | 0.875422209 |
| ENSMUSG0000002498 | chr19 | Vti1a | 0.051195493 | 0.875422209 |
| ENSMUSG0000004686 | chr4 | Pramel12 | 0.077515923 | 0.875422209 |
| ENSMUSG0000009126 | chr13 | Smim13 | -0.059863213 | 0.875619718 |
| ENSMUSG0000004941 | chr18 | Tmem241 | -0.143401977 | 0.875730799 |
| ENSMUSG0000002943 | chr5 | Diablo | -0.107050004 | 0.875755804 |
| ENSMUSG0000005460 | chr16 | Cggbp1 | 0.058397121 | 0.875755804 |
| ENSMUSG0000001625 | chr2 | Prelid3b | -0.071624789 | 0.875816522 |
| ENSMUSG0000003919 | chr4 | Orm1 | -0.404617563 | 0.875907169 |
| ENSMUSG0000001729 | chr11 | Taok1 | -0.079230455 | 0.875907169 |
| ENSMUSG0000002591 | chr1 | Sgk3 | 0.049722841 | 0.875907169 |
| ENSMUSG0000003312 | chr1 | Atg9a | 0.055798117 | 0.875907169 |
| ENSMUSG0000007087 | chr1 | Ccnyl1 | 0.092974051 | 0.875907169 |
| ENSMUSG0000003784 | chr19 | Nmrk1 | 0.141057606 | 0.875907169 |
| ENSMUSG0000012151 | chr14 | Ptprg | 0.063031985 | 0.876321215 |
| ENSMUSG0000003253 | chr9 | Cep63 | -0.072362433 | 0.876429324 |
| ENSMUSG0000002645 | chr1 | Cyb5r1 | 0.087509457 | 0.876429324 |
| ENSMUSG0000006028 | chr4 | Ppih | 0.098392574 | 0.876507634 |
| ENSMUSG0000004509 | chr19 | Kmt5b | -0.073069705 | 0.876632249 |
| ENSMUSG0000003376 | chr8 | Mtss2 | 0.082082626 | 0.876632249 |
| ENSMUSG0000000284 | chr16 | Tmem39a | 0.067297093 | 0.876632249 |
| ENSMUSG0000003968 | chr16 | Mrps6 | 0.131900867 | 0.876632249 |
| ENSMUSG0000000658 | chr8 | Aprt | 0.086115808 | 0.876632249 |
| ENSMUSG0000000392 | chr17 | Zfp81 | -0.062850414 | 0.876878699 |
| ENSMUSG0000003679 | chr2 | Mbd5 | -0.103905576 | 0.876997201 |

| | | | | |
|---|---|---|---|---|
| ENSMUSG0000002004 | chr10 | Abtb3 | -0.111430998 | 0.876997201 |
| ENSMUSG0000001971 | chr12 | L3hypdh | -0.113092077 | 0.876997201 |
| ENSMUSG0000004207 | chr9 | Abhd14b | -0.088378122 | 0.876997201 |
| ENSMUSG0000003137 | chrX | Car5b | 0.080976239 | 0.876997201 |
| ENSMUSG0000002721 | chr2 | Meis2 | 0.085309622 | 0.876997201 |
| ENSMUSG0000009131 | chr1 | Semp2l1 | 0.290377561 | 0.876997201 |
| ENSMUSG0000003853 | chr6 | Ubn2 | -0.084123987 | 0.877315357 |
| ENSMUSG0000002139 | chr13 | Cenpp | -0.191271837 | 0.877326273 |
| ENSMUSG0000003335 | chr11 | Map2k4 | 0.053028141 | 0.877326273 |
| ENSMUSG0000004134 | chr12 | Atg2b | -0.067733413 | 0.877470891 |
| ENSMUSG0000002541 | chr10 | Pip4k2c | -0.046563534 | 0.877470891 |
| ENSMUSG0000003618 | chr17 | Sapcd1 | 0.398026028 | 0.877470891 |
| ENSMUSG0000002497 | chr19 | Shoc2 | 0.050242808 | 0.877470891 |
| ENSMUSG0000002139 | chr13 | Spin1 | 0.050071121 | 0.877470891 |
| ENSMUSG0000003862 | chr2 | Polr3k | 0.077318395 | 0.877470891 |
| ENSMUSG0000000175 | chr11 | Naglu | 0.083287233 | 0.877470891 |
| ENSMUSG0000005906 | chr12 | Rad51b | -0.268931659 | 0.877518804 |
| ENSMUSG0000003587 | chr9 | Hykk | -0.114254026 | 0.877518804 |
| ENSMUSG0000002111 | chr12 | Pals1 | -0.065915442 | 0.877518804 |
| ENSMUSG0000002221 | chr14 | Dcaf11 | -0.049656613 | 0.877518804 |
| ENSMUSG0000005926 | chr7 | Usp47 | 0.041779631 | 0.877518804 |
| ENSMUSG0000004543 | chr5 | Tmem60 | 0.083815426 | 0.877578862 |
| ENSMUSG0000003687 | chr3 | 2410004B18Rik | 0.079302459 | 0.877841459 |
| ENSMUSG0000004724 | chrX | Taf9b | 0.067248101 | 0.878036285 |
| ENSMUSG0000005306 | chr6 | Clec12a | -0.249600155 | 0.878061212 |
| ENSMUSG0000003591 | chr9 | Bbs9 | -0.104027433 | 0.878061212 |
| ENSMUSG0000002905 | chr4 | Prxl2b | 0.110997388 | 0.87806652 |
| ENSMUSG0000002736 | chr2 | Dusp2 | 0.352414602 | 0.87806652 |
| ENSMUSG0000006413 | chr13 | Arb2a | -0.058356176 | 0.878102407 |
| ENSMUSG0000002190 | chr14 | Sema3g | 0.185822019 | 0.878102407 |
| ENSMUSG0000003259 | chr9 | Ip6k1 | -0.061993327 | 0.878206661 |
| ENSMUSG0000000489 | chr3 | Mettl25b | 0.063613257 | 0.878228497 |
| ENSMUSG0000003225 | chr9 | Irak1bp1 | 0.080600191 | 0.878228497 |
| ENSMUSG0000003831 | chr5 | Kcnh2 | -0.070637948 | 0.878335125 |
| ENSMUSG0000001847 | chr11 | Kdm6b | -0.14249309 | 0.878346525 |
| ENSMUSG0000000536 | chr15 | Slc1a3 | -0.140342558 | 0.878346525 |
| ENSMUSG0000002954 | chr5 | Acads | -0.104818136 | 0.878346525 |
| ENSMUSG0000002059 | chr12 | Sdc1 | -0.129024153 | 0.878346525 |
| ENSMUSG0000003726 | chr8 | Hgsnat | 0.042988745 | 0.878346525 |
| ENSMUSG0000000814 | chr7 | Emc10 | 0.055233995 | 0.878346525 |
| ENSMUSG0000002174 | chr14 | Pdhb | 0.047400071 | 0.878346525 |
| ENSMUSG0000004017 | chr7 | 2310057M21Rik | 0.058778471 | 0.878346525 |
| ENSMUSG0000004386 | chr7 | Taf10 | 0.09783739 | 0.878346525 |
| ENSMUSG0000002628 | chr1 | Dtymk | 0.082608359 | 0.878346525 |
| ENSMUSG0000002960 | chr5 | Rita1 | 0.095205478 | 0.878346525 |
| ENSMUSG0000002895 | chr4 | Zbtb48 | 0.089153502 | 0.878354922 |
| ENSMUSG0000003759 | chr12 | Clba1 | -0.123892122 | 0.878487742 |
| ENSMUSG0000004074 | chr11 | Rnf167 | -0.074900493 | 0.878487742 |
| ENSMUSG0000002155 | chr13 | Agtpbp1 | 0.054110111 | 0.878487742 |
| ENSMUSG0000004830 | chr15 | Ankrd46 | 0.056170725 | 0.878633437 |

| | | | | |
|---|---|---|---|---|
| ENSMUSG0000004022 | chr1 | Prrc2c | -0.081411156 | 0.878723787 |
| ENSMUSG0000001877 | chr11 | Cd68 | 0.149229885 | 0.878723787 |
| ENSMUSG0000006381 | chr6 | Alms1 | -0.179057799 | 0.87874557 |
| ENSMUSG0000002301 | chr15 | Smarcd1 | -0.089686265 | 0.87874557 |
| ENSMUSG0000003467 | chr17 | Pbx2 | -0.060610503 | 0.87874557 |
| ENSMUSG0000002686 | chr2 | Sh3glb2 | -0.064768468 | 0.87874557 |
| ENSMUSG0000002428 | chr18 | Wac | 0.053141162 | 0.87874557 |
| ENSMUSG0000003230 | chr9 | Mpi | 0.082679864 | 0.87874557 |
| ENSMUSG0000002278 | chr16 | Spag6l | -0.206913524 | 0.878772828 |
| ENSMUSG0000004274 | chr6 | Bmt2 | 0.050877907 | 0.87879605 |
| ENSMUSG0000001546 | chr17 | Egfl8 | -0.269191785 | 0.878855623 |
| ENSMUSG0000004577 | chr7 | Ifitm10 | -0.234244266 | 0.878915874 |
| ENSMUSG0000002257 | chr15 | Naprt | -0.099053531 | 0.878915874 |
| ENSMUSG0000002219 | chr15 | Pdzd2 | 0.087422696 | 0.878915874 |
| ENSMUSG0000000320 | chr17 | Yju2 | 0.088175179 | 0.878985839 |
| ENSMUSG0000005443 | chr5 | Tmem120b | 0.095193678 | 0.878985839 |
| ENSMUSG0000001323 | chr17 | Ptprs | -0.068207486 | 0.879271387 |
| ENSMUSG0000003806 | chr13 | Rnf144b | 0.050248608 | 0.879271387 |
| ENSMUSG0000005328 | chr9 | Ddx10 | 0.088005018 | 0.879271387 |
| ENSMUSG0000002303 | chr18 | Pcdhgc4 | -0.127129511 | 0.879288082 |
| ENSMUSG0000000737 | chr3 | Dennd2c | -0.111070261 | 0.879288082 |
| ENSMUSG0000006111 | chr7 | Prcp | -0.069912561 | 0.879288082 |
| ENSMUSG0000002112 | chr12 | Pigh | -0.072985294 | 0.879288082 |
| ENSMUSG0000003760 | chr7 | Osbpl5 | 0.050718437 | 0.879288082 |
| ENSMUSG0000002624 | chr1 | Serpine2 | 0.080540228 | 0.879288082 |
| ENSMUSG0000003516 | chr14 | Ints6 | 0.063049115 | 0.879288082 |
| ENSMUSG0000006110 | chr16 | Retnla | 0.111473952 | 0.879288082 |
| ENSMUSG0000007502 | chr2 | Prdm11 | 0.099633421 | 0.879288082 |
| ENSMUSG0000003765 | chr5 | Kctd8 | -0.120042092 | 0.879294508 |
| ENSMUSG0000002907 | chr4 | Dvl1 | 0.064976857 | 0.879303135 |
| ENSMUSG0000004069 | chr4 | Dnajc16 | 0.05153399 | 0.879546272 |
| ENSMUSG0000003624 | chr9 | Tbc1d21 | -0.282542246 | 0.879624125 |
| ENSMUSG0000002426 | chr18 | Sap130 | -0.069051685 | 0.879680148 |
| ENSMUSG0000002740 | chr2 | Nop56 | 0.068862315 | 0.879816185 |
| ENSMUSG0000004784 | chrX | Bex4 | -0.136048267 | 0.879833265 |
| ENSMUSG0000004109 | chrX | Tspyl2 | -0.070381921 | 0.880071365 |
| ENSMUSG0000002515 | chrX | Maged1 | 0.050365434 | 0.880196272 |
| ENSMUSG0000005772 | chr4 | Lepr | -0.195555737 | 0.880223663 |
| ENSMUSG0000001898 | chr11 | Slfn3 | -0.173186811 | 0.880333055 |
| ENSMUSG0000003122 | chrX | Atrx | -0.074691336 | 0.880338064 |
| ENSMUSG0000002877 | chr4 | Zcchc17 | 0.071976311 | 0.880338064 |
| ENSMUSG0000006057 | chr14 | Fhit | -0.155479628 | 0.880524681 |
| ENSMUSG0000003950 | chr8 | Nup133 | -0.058915559 | 0.880524681 |
| ENSMUSG0000002950 | chr5 | Golga3 | 0.053130765 | 0.880524681 |
| ENSMUSG0000010333 | chr18 | Pcdhga2 | 0.1106042 | 0.880524681 |
| ENSMUSG0000003130 | chrX | Rps6ka3 | 0.068381389 | 0.880628151 |
| ENSMUSG0000005974 | chr2 | Kcnh7 | 0.096199072 | 0.880628151 |
| ENSMUSG0000001983 | chr10 | Rab32 | 0.080083967 | 0.880628151 |
| ENSMUSG0000004622 | chr7 | Plaur | 0.130617715 | 0.880628151 |
| ENSMUSG0000002298 | chr16 | Scaf4 | 0.074540149 | 0.880657544 |

| | | | | |
|---|---|---|---|---|
| ENSMUSG0000003269 | chr11 | Nlrp3 | -0.30624769 | 0.880666947 |
| ENSMUSG0000002634 | chr1 | Ccnt2 | -0.064048967 | 0.880666947 |
| ENSMUSG0000004116 | chr11 | Spem1 | -0.359883897 | 0.880712754 |
| ENSMUSG0000003172 | chr8 | Ist1 | -0.051401395 | 0.880712754 |
| ENSMUSG0000002244 | chr15 | Adamts20 | -0.140549713 | 0.880739251 |
| ENSMUSG0000003229 | chr9 | Neil1 | -0.096890479 | 0.880739251 |
| ENSMUSG0000006269 | chr17 | Cebpzos | 0.123665473 | 0.880739251 |
| ENSMUSG0000002927 | chr5 | Brdt | 0.082622997 | 0.880786697 |
| ENSMUSG0000003694 | chr2 | Slc39a12 | -0.359889078 | 0.880833705 |
| ENSMUSG0000003951 | chr2 | Ptpa | 0.071870034 | 0.880833705 |
| ENSMUSG0000000768 | chr12 | Dio2 | 0.166345295 | 0.880924215 |
| ENSMUSG0000003183 | chr8 | Mbtps1 | 0.048871355 | 0.881087817 |
| ENSMUSG0000002587 | chr13 | Tspan17 | -0.080750973 | 0.881198653 |
| ENSMUSG0000003439 | chr11 | Lif | -0.20994076 | 0.881370187 |
| ENSMUSG0000006177 | chrX | Mospd2 | 0.064097079 | 0.881370187 |
| ENSMUSG0000003506 | chrX | Zc4h2 | 0.064223893 | 0.881370187 |
| ENSMUSG0000002869 | chr4 | Nasp | 0.081450197 | 0.881370187 |
| ENSMUSG0000005315 | chr1 | Spag16 | 0.277441913 | 0.881370187 |
| ENSMUSG0000000234 | chr8 | Slc25a42 | 0.100790851 | 0.881370187 |
| ENSMUSG0000000254 | chr2 | Golga2 | 0.050216038 | 0.881380809 |
| ENSMUSG0000006874 | chr6 | Ptprz1 | 0.096312913 | 0.881454428 |
| ENSMUSG0000002616 | chr1 | Nhej1 | 0.125441146 | 0.881454428 |
| ENSMUSG0000003703 | chr7 | Apbb1 | -0.080741281 | 0.88220363 |
| ENSMUSG0000002752 | chr1 | Zdbf2 | 0.095651277 | 0.882204469 |
| ENSMUSG0000005290 | chr8 | Ubxn8 | 0.057702808 | 0.882243276 |
| ENSMUSG0000002134 | chr13 | Gpld1 | -0.165093305 | 0.882335991 |
| ENSMUSG0000000094 | chr7 | Gm38393 | -0.110634971 | 0.88236996 |
| ENSMUSG0000004986 | chr1 | Arl4c | 0.099888493 | 0.88236996 |
| ENSMUSG0000002806 | chr3 | Iqgap3 | 0.391161276 | 0.88236996 |
| ENSMUSG0000005267 | chr7 | Zfp112 | -0.079085951 | 0.88238285 |
| ENSMUSG0000002597 | chr1 | Rftn2 | -0.061290044 | 0.88239038 |
| ENSMUSG0000002520 | chr19 | Cwf19l1 | 0.070674609 | 0.88239038 |
| ENSMUSG0000009597 | chr14 | Cphx1 | -0.460998944 | 0.88245925 |
| ENSMUSG0000002247 | chr15 | Hdac7 | 0.083005885 | 0.88253601 |
| ENSMUSG0000003570 | chr7 | Alg8 | 0.07489014 | 0.88253601 |
| ENSMUSG0000003531 | chr7 | Gdpd5 | -0.111491899 | 0.882771309 |
| ENSMUSG0000003063 | chr7 | Fah | -0.087072541 | 0.88290506 |
| ENSMUSG0000005490 | chr17 | Arhgef33 | -0.321117255 | 0.88290506 |
| ENSMUSG0000004021 | chr3 | Kyat3 | -0.110198411 | 0.88290506 |
| ENSMUSG0000002229 | chr15 | Baalc | -0.109808788 | 0.88290506 |
| ENSMUSG0000002736 | chr2 | Trpm7 | 0.076447733 | 0.88290506 |
| ENSMUSG0000002477 | chr19 | Ppp2r5b | -0.076391995 | 0.882923037 |
| ENSMUSG0000003157 | chr8 | Mak16 | 0.099714839 | 0.882923037 |
| ENSMUSG0000007984 | chrX | Xlr4a | 0.643058574 | 0.882923037 |
| ENSMUSG0000003067 | chr7 | Kif22 | -0.193766884 | 0.882977143 |
| ENSMUSG0000002742 | chr2 | Snx5 | 0.063219545 | 0.882977143 |
| ENSMUSG0000004657 | chr13 | Lyrm4 | 0.079866407 | 0.882977143 |
| ENSMUSG0000001903 | chr9 | Dalrd3 | -0.095761531 | 0.883194851 |
| ENSMUSG0000005039 | chrX | Armcx6 | -0.132927628 | 0.883194851 |
| ENSMUSG0000003308 | chr6 | Clec1a | -0.146880202 | 0.883194851 |

| | | | | |
|---|---|---|---|---|
| ENSMUSG0000003523 | chr2 | Scai | -0.077718422 | 0.883194851 |
| ENSMUSG0000002268 | chr16 | Pdxdc1 | 0.048969259 | 0.883194851 |
| ENSMUSG000000342 | chr7 | Pih1d1 | 0.082419757 | 0.883194851 |
| ENSMUSG0000003176 | chr8 | Mt1 | 0.155759172 | 0.883194851 |
| ENSMUSG0000009560 | chr13 | Anxa2r1 | -0.359665991 | 0.883199951 |
| ENSMUSG0000003280 | chrX | Slc10a3 | -0.113884885 | 0.883495382 |
| ENSMUSG0000002592 | chr1 | Stau2 | -0.073902316 | 0.88361998 |
| ENSMUSG0000007944 | chr1 | B3gnt7 | -0.208092104 | 0.883849557 |
| ENSMUSG0000007531 | chr2 | Scn2a | -0.109971565 | 0.883849557 |
| ENSMUSG0000003130 | chrX | Il2rg | -0.142661728 | 0.883849557 |
| ENSMUSG0000004759 | chr15 | Mafa | -0.130197955 | 0.883849557 |
| ENSMUSG0000002418 | chr17 | Pdia2 | -0.299729933 | 0.883849557 |
| ENSMUSG0000004620 | chr11 | Pik3r6 | -0.201519257 | 0.883849557 |
| ENSMUSG0000001403 | chr16 | Prdm15 | -0.08379762 | 0.883849557 |
| ENSMUSG0000002097 | chr12 | Dnaaf2 | -0.252967947 | 0.883849557 |
| ENSMUSG0000005014 | chr8 | F2rl3 | -0.30543951 | 0.883849557 |
| ENSMUSG0000002257 | chr15 | Gfus | -0.089744848 | 0.883849557 |
| ENSMUSG0000004244 | chr4 | Zmym4 | -0.069945001 | 0.883849557 |
| ENSMUSG0000004306 | chr16 | Spice1 | -0.06072536 | 0.883849557 |
| ENSMUSG0000003806 | chr9 | Dlec1 | -0.154743646 | 0.883849557 |
| ENSMUSG0000002560 | chr9 | Zfp202 | -0.093770841 | 0.883849557 |
| ENSMUSG0000006124 | chr14 | Exoc5 | -0.04670569 | 0.883849557 |
| ENSMUSG0000004038 | chr2 | Aqr | 0.043285107 | 0.883849557 |
| ENSMUSG0000007944 | chr2 | St6galnac4 | 0.057142693 | 0.883849557 |
| ENSMUSG0000005506 | chr1 | Smyd3 | 0.071637972 | 0.883849557 |
| ENSMUSG0000002476 | chr19 | Lipo3 | 0.065194667 | 0.883849557 |
| ENSMUSG0000003247 | chr9 | Mras | 0.060770745 | 0.883849557 |
| ENSMUSG0000001919 | chr7 | Scn1b | 0.085869176 | 0.883849557 |
| ENSMUSG0000003677 | chr9 | Anln | 0.098492806 | 0.883849557 |
| ENSMUSG0000003583 | chr10 | Plppr3 | 0.097901331 | 0.883849557 |
| ENSMUSG0000003974 | chr7 | Htatip2 | 0.108815355 | 0.883849557 |
| ENSMUSG0000003259 | chr9 | Ip6k2 | 0.05927876 | 0.883973978 |
| ENSMUSG0000000838 | chr7 | Sertad1 | 0.085088094 | 0.883973978 |
| ENSMUSG0000003058 | chr7 | 2200002D01Rik | 0.128254265 | 0.883973978 |
| ENSMUSG0000002286 | chr16 | Ahsg | -0.398142251 | 0.883980274 |
| ENSMUSG0000002210 | chr14 | Med4 | 0.062364298 | 0.884058798 |
| ENSMUSG0000005426 | chr15 | Lifr | 0.089332622 | 0.884058798 |
| ENSMUSG0000004824 | chr17 | Crebrf | -0.102402925 | 0.884160875 |
| ENSMUSG0000004278 | chr3 | Muc1 | -0.157279958 | 0.884160875 |
| ENSMUSG0000011866 | chr16 | Arvcf | -0.07908692 | 0.884160875 |
| ENSMUSG000000336 | chr7 | Pld3 | -0.054337786 | 0.884160875 |
| ENSMUSG0000003338 | chr18 | Trappc8 | -0.052261202 | 0.884160875 |
| ENSMUSG0000003120 | chrX | Brcc3 | 0.079449952 | 0.884264044 |
| ENSMUSG0000004201 | chr5 | Acacb | 0.071772847 | 0.884264044 |
| ENSMUSG0000003007 | chr6 | Cntn3 | -0.104563265 | 0.88431409 |
| ENSMUSG0000004710 | chr6 | Pbp2 | -0.509385473 | 0.88431409 |
| ENSMUSG0000006120 | chr17 | Stk19 | -0.056948261 | 0.88431409 |
| ENSMUSG0000002679 | chr2 | Niban2 | -0.061269744 | 0.88431409 |
| ENSMUSG0000005398 | chr7 | Zfp14 | -0.059831687 | 0.88431409 |
| ENSMUSG0000007903 | chr2 | Prnp | 0.052784733 | 0.88431409 |

| | | | | |
|---|---|---|---|---|
| ENSMUSG0000003605 | chr9 | Ptpn23 | 0.059412272 | 0.88431409 |
| ENSMUSG0000003162 | chr8 | Irf2 | 0.058228894 | 0.88431409 |
| ENSMUSG0000003613 | chrX | Frmd7 | 0.280989051 | 0.88431409 |
| ENSMUSG0000007910 | chr12 | Srp54c | 0.064534493 | 0.88431409 |
| ENSMUSG0000002695 | chr2 | Uap1l1 | 0.080142386 | 0.88431409 |
| ENSMUSG0000002582 | chr6 | Zfp282 | 0.071048347 | 0.88431409 |
| ENSMUSG0000004210 | chr15 | Csdc2 | 0.078877217 | 0.88431409 |
| ENSMUSG0000004654 | chr16 | Fam43a | 0.100697572 | 0.88431409 |
| ENSMUSG0000003138 | chrX | Vegfd | 0.22817752 | 0.88431409 |
| ENSMUSG0000000447 | chr7 | Clec11a | 0.114312934 | 0.884344259 |
| ENSMUSG0000003059 | chr7 | Fbxo17 | -0.095056937 | 0.884734764 |
| ENSMUSG0000004837 | chr14 | Socs4 | 0.04798176 | 0.884734764 |
| ENSMUSG0000009095 | chr7 | Lrrc32 | 0.139369012 | 0.884734764 |
| ENSMUSG0000003403 | chr4 | Ccdc17 | -0.126250324 | 0.884993615 |
| ENSMUSG0000001005 | chr9 | Tusc2 | -0.092751198 | 0.884993615 |
| ENSMUSG0000005791 | chr2 | Cacnb2 | -0.060753166 | 0.884993615 |
| ENSMUSG0000005452 | chr5 | Sh3bp2 | -0.102778942 | 0.885055417 |
| ENSMUSG0000005829 | chr10 | Mcm9 | -0.066190478 | 0.885055417 |
| ENSMUSG0000002679 | chr2 | Lrsam1 | -0.052926891 | 0.885055417 |
| ENSMUSG0000001985 | chr10 | Reps1 | 0.060603084 | 0.885055417 |
| ENSMUSG0000002701 | chr2 | Zfp385b | 0.142868621 | 0.885055417 |
| ENSMUSG0000003989 | chr10 | Txlnb | 0.184220738 | 0.885055417 |
| ENSMUSG0000000469 | chr12 | Hdac9 | 0.08692694 | 0.885055417 |
| ENSMUSG0000002454 | chr18 | Setbp1 | 0.089827414 | 0.885055417 |
| ENSMUSG0000005900 | chr16 | Grin2a | 0.282841785 | 0.885055417 |
| ENSMUSG0000003067 | chr7 | Maz | -0.054833489 | 0.88516965 |
| ENSMUSG0000005920 | chr17 | Hnrnpm | 0.063124747 | 0.88516965 |
| ENSMUSG0000006603 | chr4 | Hnrnpr | 0.05762333 | 0.88516965 |
| ENSMUSG0000003774 | chr2 | Mrps26 | 0.072678726 | 0.88516965 |
| ENSMUSG0000003541 | chr11 | Tmem98 | -0.149590286 | 0.885275727 |
| ENSMUSG0000006340 | chr5 | Tmed5 | 0.0510283 | 0.885275727 |
| ENSMUSG0000002962 | chr6 | Phf14 | 0.076490562 | 0.885275727 |
| ENSMUSG0000004993 | chr6 | Lrrc4 | 0.10119748 | 0.885275727 |
| ENSMUSG0000009189 | chr14 | Tnnc1 | 0.210783891 | 0.885275727 |
| ENSMUSG0000004123 | chr11 | Ublcp1 | 0.086412548 | 0.885281655 |
| ENSMUSG0000002586 | chrX | Xiap | -0.04964322 | 0.885481075 |
| ENSMUSG0000004270 | chr11 | Atpaf2 | 0.056491779 | 0.885621092 |
| ENSMUSG0000002578 | chr9 | Clec3b | 0.059767428 | 0.885621092 |
| ENSMUSG0000002309 | chr2 | Msrb2 | 0.097605354 | 0.885621092 |
| ENSMUSG0000010379 | chr18 | Pcdhga6 | 0.103713697 | 0.885625146 |
| ENSMUSG0000004759 | chr5 | Nxpe5 | -0.216594011 | 0.885633667 |
| ENSMUSG0000002785 | chr3 | Tshb | -0.108317605 | 0.885633667 |
| ENSMUSG0000003192 | chr8 | Terf2 | -0.048226223 | 0.885633667 |
| ENSMUSG0000007315 | chr12 | 9030624G23Rik | 0.231304469 | 0.885633667 |
| ENSMUSG0000001912 | chr11 | Ccl9 | 0.097353639 | 0.885633667 |
| ENSMUSG0000002493 | chr19 | Kcnk7 | 0.240681676 | 0.885633667 |
| ENSMUSG0000002659 | chr1 | Abl2 | 0.070993263 | 0.885781656 |
| ENSMUSG0000003271 | chr1 | Sbspon | -0.189160838 | 0.885787739 |
| ENSMUSG0000003182 | chr8 | 6430548M08Rik | 0.086019833 | 0.885787739 |
| ENSMUSG0000003977 | chr17 | Ypel5 | 0.064215141 | 0.885855344 |

| | | | | |
|---|---|---|---|---|
| ENSMUSG0000004041 | chr4 | Fbxl4 | -0.078849877 | 0.885863172 |
| ENSMUSG0000004327 | chr5 | Trim56 | -0.055938275 | 0.885863172 |
| ENSMUSG0000001242 | chr13 | Mplkip | -0.074391178 | 0.886052158 |
| ENSMUSG0000004193 | chr15 | Rimoc1 | 0.053366553 | 0.886087303 |
| ENSMUSG0000001053 | chr3 | Tsacc | -0.260072933 | 0.886121945 |
| ENSMUSG0000003836 | chr2 | Ncoa6 | -0.074340905 | 0.886121945 |
| ENSMUSG0000000537 | chr5 | Tbl2 | -0.070153751 | 0.886121945 |
| ENSMUSG0000002214 | chr15 | Nipbl | -0.07847318 | 0.886121945 |
| ENSMUSG0000003081 | chr7 | Fbxl19 | -0.087595323 | 0.886121945 |
| ENSMUSG0000001260 | chr12 | Ttll5 | -0.067289253 | 0.886121945 |
| ENSMUSG0000004210 | chr9 | Inka1 | -0.133243418 | 0.886121945 |
| ENSMUSG0000002794 | chr3 | 4933434E20Rik | -0.066088807 | 0.886121945 |
| ENSMUSG0000005300 | chr6 | Hrh1 | -0.213493032 | 0.886121945 |
| ENSMUSG0000001981 | chr10 | Cd164 | 0.047067274 | 0.886121945 |
| ENSMUSG0000000108 | chr5 | Mfsd10 | 0.04965495 | 0.886121945 |
| ENSMUSG0000006355 | chr1 | Aox1 | 0.128321673 | 0.886121945 |
| ENSMUSG0000003632 | chr2 | Qsox2 | 0.053893519 | 0.886121945 |
| ENSMUSG0000004887 | chr1 | Phf3 | 0.058958322 | 0.886121945 |
| ENSMUSG0000002697 | chr2 | Zmynd19 | 0.059780108 | 0.886121945 |
| ENSMUSG0000002535 | chr10 | Cdk2 | 0.071649983 | 0.886121945 |
| ENSMUSG0000004633 | chr1 | Fam178b | 0.194047444 | 0.886121945 |
| ENSMUSG0000004923 | chr3 | Myef2l | 0.164952407 | 0.886121945 |
| ENSMUSG0000005515 | chr7 | Zfp78 | 0.1295338 | 0.886121945 |
| ENSMUSG0000003891 | chr2 | Dido1 | -0.063017084 | 0.886238942 |
| ENSMUSG0000003478 | chr17 | Gpsm3 | -0.111719379 | 0.886238942 |
| ENSMUSG0000003407 | chr2 | Zdhhc5 | 0.053969201 | 0.886238942 |
| ENSMUSG0000002810 | chr3 | Pias3 | 0.054513567 | 0.886238942 |
| ENSMUSG0000003976 | chr16 | Dnajc28 | 0.101656613 | 0.886238942 |
| ENSMUSG0000004071 | chr7 | Klc3 | 0.164007765 | 0.886238942 |
| ENSMUSG0000007489 | chr2 | Lcmt2 | 0.10577819 | 0.886291932 |
| ENSMUSG0000003876 | chr4 | Ptpn3 | -0.109869373 | 0.886344993 |
| ENSMUSG0000002501 | chr19 | Tm9sf3 | -0.035178635 | 0.886344993 |
| ENSMUSG0000002396 | chr17 | Enpp4 | 0.057895704 | 0.886344993 |
| ENSMUSG0000007914 | chr1 | A130010J15Rik | 0.097490741 | 0.886344993 |
| ENSMUSG0000007047 | chr3 | Erich6 | 0.247520591 | 0.886357646 |
| ENSMUSG0000003507 | chr14 | Mtmr9 | -0.060118978 | 0.88635821 |
| ENSMUSG0000002890 | chr4 | Gmeb1 | 0.061406821 | 0.88635821 |
| ENSMUSG0000000039 | chr2 | Fap | 0.280733119 | 0.88635821 |
| ENSMUSG0000005420 | chr10 | Gzmm | -0.242939767 | 0.886392892 |
| ENSMUSG0000003981 | chr10 | Zc3h10 | -0.083438434 | 0.886392892 |
| ENSMUSG0000002816 | chr3 | Cisd2 | -0.060058812 | 0.886392892 |
| ENSMUSG0000005579 | chr6 | Tcf7l1 | 0.07486541 | 0.886392892 |
| ENSMUSG0000003384 | chr1 | Mrpl15 | 0.090897808 | 0.886392892 |
| ENSMUSG0000005123 | chr11 | Tmem199 | 0.078034941 | 0.886392892 |
| ENSMUSG0000002984 | chr6 | Slc23a4 | 0.222753356 | 0.886392892 |
| ENSMUSG0000001866 | chr11 | Cdk5rap3 | 0.082561371 | 0.886392892 |
| ENSMUSG0000007452 | chr2 | Zfp972 | -0.406757291 | 0.886635092 |
| ENSMUSG0000007307 | chr12 | Srp54a | 0.186694492 | 0.886650058 |
| ENSMUSG0000001976 | chr10 | Esr1 | -0.130717502 | 0.886668176 |
| ENSMUSG0000005718 | chr11 | 5730455P16Rik | -0.058029356 | 0.886668176 |

| | | | | |
|---|---|---|---|---|
| ENSMUSG0000002045 | chr11 | Mtif2 | 0.057207115 | 0.886668176 |
| ENSMUSG0000006294 | chrX | Atp11c | -0.09354581 | 0.886678785 |
| ENSMUSG0000002508 | chr19 | Trub1 | -0.080111867 | 0.886678785 |
| ENSMUSG0000005302 | chr7 | Sv2b | 0.086808484 | 0.886693718 |
| ENSMUSG0000011033 | chr8 | Gm19935 | -0.274046588 | 0.886877741 |
| ENSMUSG0000002253 | chr16 | Tmem44 | -0.105026504 | 0.886877741 |
| ENSMUSG0000003290 | chr18 | Atg12 | 0.054015399 | 0.887179847 |
| ENSMUSG0000000405 | chr19 | Map3k11 | -0.088433671 | 0.88724062 |
| ENSMUSG0000003005 | chr6 | Aplf | -0.087473187 | 0.88724062 |
| ENSMUSG0000002947 | chr5 | Kdm2b | 0.072766631 | 0.887306991 |
| ENSMUSG0000003018 | chr6 | Magohb | 0.101826456 | 0.887306991 |
| ENSMUSG0000000112 | chrX | Cfp | -0.085303225 | 0.887364026 |
| ENSMUSG0000003421 | chr5 | Mrps17 | 0.079497182 | 0.887364026 |
| ENSMUSG0000003238 | chr9 | Csnk1g1 | 0.069817464 | 0.88737748 |
| ENSMUSG0000002507 | chr19 | Nhlrc2 | 0.069932244 | 0.88737748 |
| ENSMUSG0000003695 | chr10 | Kifbp | -0.044689373 | 0.887434463 |
| ENSMUSG0000003664 | chr2 | Ccdc148 | -0.113373662 | 0.887679489 |
| ENSMUSG0000007421 | chr8 | Gm10643 | -0.11977892 | 0.887679489 |
| ENSMUSG0000006635 | chr9 | Wdr6 | -0.055993167 | 0.887679489 |
| ENSMUSG0000002593 | chr1 | Tram1 | 0.055916694 | 0.887679489 |
| ENSMUSG0000002782 | chr3 | Vmn2r1 | 0.327857604 | 0.887679489 |
| ENSMUSG0000001055 | chr11 | Mettl16 | 0.083940867 | 0.887679489 |
| ENSMUSG0000002452 | chr18 | Cidea | 0.152429856 | 0.887679489 |
| ENSMUSG0000003186 | chr8 | Pbx4 | 0.172070181 | 0.887679489 |
| ENSMUSG0000002261 | chr15 | Chkb | -0.159921309 | 0.887700992 |
| ENSMUSG0000003906 | chr15 | Atpsckmt | 0.064054137 | 0.887794553 |
| ENSMUSG0000002757 | chr2 | Arfgap1 | 0.050111553 | 0.888153483 |
| ENSMUSG0000000463 | chr8 | Wwox | -0.089272389 | 0.888198803 |
| ENSMUSG0000002595 | chr1 | Klf7 | -0.07031766 | 0.888198803 |
| ENSMUSG0000003817 | chr1 | Hycc2 | -0.071425663 | 0.888198803 |
| ENSMUSG0000001882 | chr19 | Zfyve27 | -0.051766591 | 0.888198803 |
| ENSMUSG0000003170 | chr8 | Itfg1 | -0.041285831 | 0.888198803 |
| ENSMUSG0000002093 | chr11 | Gfap | 0.218027715 | 0.888198803 |
| ENSMUSG0000002936 | chr5 | Rfc5 | -0.097320445 | 0.888206434 |
| ENSMUSG0000001006 | chr9 | Rassf1 | -0.10019735 | 0.888206434 |
| ENSMUSG0000001800 | chr5 | Cyth3 | 0.054501215 | 0.888206434 |
| ENSMUSG0000003138 | chrX | Naa10 | 0.072182095 | 0.888206434 |
| ENSMUSG0000004445 | chr7 | Zfp507 | 0.046218209 | 0.888343724 |
| ENSMUSG0000003160 | chr8 | Msmo1 | -0.119546925 | 0.888611777 |
| ENSMUSG0000002658 | chr1 | Prrx1 | 0.328597583 | 0.888611777 |
| ENSMUSG0000002093 | chr11 | Nmt1 | 0.031037537 | 0.888611777 |
| ENSMUSG0000003980 | chr2 | Ncoa5 | 0.064971038 | 0.888611777 |
| ENSMUSG0000002527 | chrX | Pfkfb1 | 0.268576092 | 0.888778152 |
| ENSMUSG0000002801 | chr3 | Gstcd | 0.099103244 | 0.888778152 |
| ENSMUSG0000004607 | chr5 | Lrrc8d | 0.09435167 | 0.889115595 |
| ENSMUSG0000005126 | chr6 | Nat8f3 | -0.305362726 | 0.889134589 |
| ENSMUSG0000004417 | chr17 | Ptx4 | -0.249615853 | 0.889134589 |
| ENSMUSG0000001533 | chr2 | Zdhhc12 | -0.085202938 | 0.889134589 |
| ENSMUSG0000002027 | chr11 | Papolg | 0.06082227 | 0.889134589 |
| ENSMUSG0000004730 | chr18 | Pcdhb13 | 0.154811734 | 0.889134589 |

| | | | | |
|---|---|---|---|---|
| ENSMUSG0000000116 | chr5 | Oas1c | 0.103756231 | 0.889134589 |
| ENSMUSG0000003797 | chr16 | Snn | 0.060793119 | 0.889134589 |
| ENSMUSG0000004382 | chr10 | Adamtsl5 | 0.118540064 | 0.889134589 |
| ENSMUSG0000004073 | chr16 | Erg | -0.112052718 | 0.889421801 |
| ENSMUSG0000003115 | chrX | Otud5 | -0.04201989 | 0.889421801 |
| ENSMUSG0000000117 | chr12 | Calm1 | 0.052674186 | 0.889512321 |
| ENSMUSG0000004177 | chr13 | Enc1 | 0.086886583 | 0.889512321 |
| ENSMUSG0000002899 | chr5 | Hycc1 | -0.075520223 | 0.88964697 |
| ENSMUSG0000003246 | chr9 | Pik3cb | -0.093238623 | 0.88964697 |
| ENSMUSG0000002478 | chr19 | Lipa | -0.077459766 | 0.88964697 |
| ENSMUSG0000002271 | chr16 | Pmm2 | -0.065839024 | 0.88964697 |
| ENSMUSG0000001700 | chr2 | Slpi | 0.375038885 | 0.88964697 |
| ENSMUSG0000002206 | chr14 | Pibf1 | 0.06006211 | 0.88964697 |
| ENSMUSG0000005739 | chr13 | Zfp759 | 0.083247855 | 0.88964697 |
| ENSMUSG0000003015 | chr6 | Clec2d | 0.086344194 | 0.88964697 |
| ENSMUSG0000001766 | chr2 | Slc35c2 | -0.052772921 | 0.889778316 |
| ENSMUSG0000002970 | chr6 | Slc13a1 | -0.956657407 | 0.889804597 |
| ENSMUSG0000002831 | chr4 | Smc2 | -0.159636467 | 0.889804597 |
| ENSMUSG0000002839 | chr4 | Alad | -0.116891367 | 0.889804597 |
| ENSMUSG0000004265 | chr7 | Arrdc4 | -0.112476856 | 0.889804597 |
| ENSMUSG0000002443 | chr18 | Cabyr | -0.213077026 | 0.889804597 |
| ENSMUSG0000003955 | chrX | Ppp1r3f | -0.165897023 | 0.889804597 |
| ENSMUSG0000002779 | chr3 | Dclk1 | -0.079507765 | 0.889804597 |
| ENSMUSG0000004315 | chr12 | Crppa | -0.075119387 | 0.889804597 |
| ENSMUSG0000002911 | chr5 | Rnf4 | 0.037596141 | 0.889804597 |
| ENSMUSG0000002947 | chr5 | Ncor2 | 0.088381292 | 0.889804597 |
| ENSMUSG0000003616 | chr15 | Pphln1 | 0.047450856 | 0.889804597 |
| ENSMUSG0000005693 | chr11 | Acbd4 | 0.047708722 | 0.889804597 |
| ENSMUSG0000002681 | chr2 | Tsc1 | 0.070520733 | 0.889804597 |
| ENSMUSG0000002385 | chr17 | Chd1 | 0.073156377 | 0.889804597 |
| ENSMUSG0000006290 | chr16 | Klhl24 | 0.089071423 | 0.889804597 |
| ENSMUSG0000004849 | chr1 | Tyw5 | 0.063812643 | 0.889804597 |
| ENSMUSG0000002738 | chr2 | Fbln7 | 0.094244643 | 0.889804597 |
| ENSMUSG0000006896 | chr2 | Zbtb34 | 0.082046349 | 0.889804597 |
| ENSMUSG0000000358 | chr11 | Rnf215 | 0.075006411 | 0.889804597 |
| ENSMUSG0000006006 | chr5 | Alox5ap | 0.130080547 | 0.889804597 |
| ENSMUSG0000003328 | chr15 | Kctd17 | 0.071460739 | 0.889804597 |
| ENSMUSG0000003134 | chrX | Gabrq | 0.251157346 | 0.889804597 |
| ENSMUSG0000003134 | chrX | Nsdhl | -0.100320701 | 0.889815022 |
| ENSMUSG0000004593 | chr12 | Clec14a | -0.107690969 | 0.889815022 |
| ENSMUSG0000001575 | chr10 | Tab2 | 0.055943226 | 0.889815022 |
| ENSMUSG0000004685 | chr1 | Cmklr2 | -0.359899271 | 0.889872371 |
| ENSMUSG0000002427 | chr18 | 2700062C07Rik | -0.070986339 | 0.889872371 |
| ENSMUSG0000004499 | chr2 | Shld1 | -0.104821475 | 0.889904232 |
| ENSMUSG0000001002 | chr11 | Kif19a | -0.126008625 | 0.889904232 |
| ENSMUSG0000000411 | chr1 | Cacna1e | -0.098839831 | 0.889904232 |
| ENSMUSG0000003376 | chr19 | Rbm4b | -0.069501806 | 0.889904232 |
| ENSMUSG0000004724 | chr7 | C2cd3 | -0.062249905 | 0.889904232 |
| ENSMUSG0000007070 | chr7 | Zfp974 | -0.055968575 | 0.889904232 |
| ENSMUSG0000003703 | chr10 | Tspan15 | 0.058723254 | 0.889904232 |

| | | | | |
|---|---|---|---|---|
| ENSMUSG0000002030 | chr9 | Ift46 | 0.061888553 | 0.889904232 |
| ENSMUSG0000005239 | chr14 | Rft1 | 0.06348402 | 0.889904232 |
| ENSMUSG0000003631 | chr17 | Polr1h | 0.098827793 | 0.889904232 |
| ENSMUSG0000008328 | chr19 | Ctsf | 0.101302427 | 0.889904232 |
| ENSMUSG0000002213 | chr14 | Cldn10 | 0.181529832 | 0.889904232 |
| ENSMUSG0000002789 | chr3 | Slc16a4 | 0.110789178 | 0.890121614 |
| ENSMUSG0000004409 | chr3 | Rsbn1 | -0.071957721 | 0.890151722 |
| ENSMUSG0000001998 | chr10 | Arg1 | -0.336327423 | 0.890285019 |
| ENSMUSG0000002728 | chr2 | Lrrc57 | -0.064624643 | 0.890285019 |
| ENSMUSG0000003066 | chr7 | 1110004F10Rik | 0.063964609 | 0.890366059 |
| ENSMUSG0000005270 | chr7 | Tnrc6a | 0.063967872 | 0.890366059 |
| ENSMUSG0000006424 | chr5 | Plcxd1 | 0.1976188 | 0.890414594 |
| ENSMUSG0000002041 | chr11 | Pttg1 | 0.152091897 | 0.890414594 |
| ENSMUSG0000004150 | chr9 | Rrp9 | 0.085178496 | 0.890549217 |
| ENSMUSG0000002131 | chr13 | Gli3 | 0.085987578 | 0.890549217 |
| ENSMUSG0000009723 | chr11 | Gm27029 | 0.255782932 | 0.890640891 |
| ENSMUSG0000011216 | chr10 | BC024063 | 0.128532402 | 0.890743447 |
| ENSMUSG0000010937 | chr7 | Gm49396 | -0.144302472 | 0.890874373 |
| ENSMUSG0000007364 | chr1 | Wdfy1 | -0.172186549 | 0.890883138 |
| ENSMUSG0000002601 | chr1 | Cd28 | -0.249817644 | 0.890883138 |
| ENSMUSG0000000885 | chr11 | Hdac5 | 0.056181597 | 0.890883138 |
| ENSMUSG0000002967 | chr6 | Pot1a | 0.061331826 | 0.890970119 |
| ENSMUSG0000002258 | chr15 | Ly6e | -0.107391956 | 0.891180373 |
| ENSMUSG0000207608 | chr4 | Klhl17 | -0.049395629 | 0.891180373 |
| ENSMUSG0000004643 | chr15 | Hnrnpa1 | 0.062222156 | 0.891180373 |
| ENSMUSG0000002140 | chr13 | Serpinb9b | 0.275246337 | 0.891180373 |
| ENSMUSG0000003639 | chr9 | Glb1l2 | 0.088597462 | 0.891180373 |
| ENSMUSG0000002937 | chr5 | Rassf6 | 0.115598261 | 0.891180373 |
| ENSMUSG0000003028 | chr6 | Cmas | -0.055085622 | 0.891192826 |
| ENSMUSG0000003444 | chr19 | Cyb561a3 | -0.07624302 | 0.891406973 |
| ENSMUSG0000002304 | chr15 | Igfbp6 | 0.238652099 | 0.891406973 |
| ENSMUSG0000002642 | chr1 | Ube2t | 0.225412056 | 0.891500993 |
| ENSMUSG0000002615 | chr1 | Ogfrl1 | -0.079656241 | 0.891531081 |
| ENSMUSG0000007552 | chr11 | Aarsd1 | -0.218598182 | 0.891531081 |
| ENSMUSG0000002074 | chr11 | Pafah1b1 | -0.049346608 | 0.891531081 |
| ENSMUSG0000002205 | chr14 | Ppp2r2a | 0.039688916 | 0.891531081 |
| ENSMUSG0000002878 | chr4 | Ptp4a2 | 0.07134647 | 0.891531081 |
| ENSMUSG0000005294 | chr11 | Rnf157 | 0.082039955 | 0.891531081 |
| ENSMUSG0000002092 | chr11 | Tmem101 | 0.093252253 | 0.891531081 |
| ENSMUSG0000002620 | chr1 | Dnajb2 | 0.074184376 | 0.891531081 |
| ENSMUSG0000000667 | chr9 | P4htm | 0.082730719 | 0.891531081 |
| ENSMUSG0000003281 | chr8 | Fanca | 0.160971018 | 0.891531081 |
| ENSMUSG0000002500 | chr19 | Tctn3 | -0.060325701 | 0.891562854 |
| ENSMUSG0000002765 | chr2 | Rprd1b | -0.047720866 | 0.891562854 |
| ENSMUSG0000003084 | chr7 | Tial1 | -0.045646946 | 0.891562854 |
| ENSMUSG0000001841 | chr11 | Kansl1 | 0.046672006 | 0.891562854 |
| ENSMUSG0000002906 | chr4 | Nadk | 0.066072667 | 0.891562854 |
| ENSMUSG0000004026 | chr7 | Plekha1 | 0.060734984 | 0.891562854 |
| ENSMUSG0000009827 | chr16 | Rpl24 | 0.201069037 | 0.891562854 |
| ENSMUSG0000001075 | chr7 | Tnfrsf22 | 0.099057153 | 0.891562854 |

| | | | | |
|---|---|---|---|---|
| ENSMUSG0000000130 | chr8 | Efnb2 | 0.062011087 | 0.891570423 |
| ENSMUSG0000002684 | chr2 | Fubp3 | 0.064209478 | 0.891578388 |
| ENSMUSG0000003415 | chr2 | Ino80 | 0.073176613 | 0.891578388 |
| ENSMUSG0000006610 | chr7 | Muc5b | -0.385300754 | 0.891618521 |
| ENSMUSG0000004815 | chr15 | Kmt2d | -0.101303091 | 0.891618521 |
| ENSMUSG0000000128 | chr15 | Itgb7 | -0.179619024 | 0.891618521 |
| ENSMUSG0000002587 | chr13 | Unc5a | -0.082292111 | 0.891618521 |
| ENSMUSG0000003229 | chr9 | Man2c1 | -0.063175057 | 0.891618521 |
| ENSMUSG0000002413 | chr17 | Abca3 | -0.064565293 | 0.891618521 |
| ENSMUSG0000003251 | chr9 | Gorasp1 | -0.059929178 | 0.891618521 |
| ENSMUSG0000005621 | chr18 | Pard6g | -0.064639142 | 0.891618521 |
| ENSMUSG0000003538 | chr9 | Pcsk7 | -0.069806664 | 0.891618521 |
| ENSMUSG0000004054 | chr11 | Pitpnm3 | -0.057183333 | 0.891618521 |
| ENSMUSG0000004011 | chr5 | Cacna2d1 | 0.055513672 | 0.891618521 |
| ENSMUSG0000002319 | chr6 | P3h3 | 0.060436348 | 0.891618521 |
| ENSMUSG0000000362 | chr5 | Crot | 0.064138132 | 0.891618521 |
| ENSMUSG0000004537 | chr11 | Wdr81 | 0.080281433 | 0.891618521 |
| ENSMUSG0000002027 | chr11 | Stk10 | 0.074902846 | 0.891640539 |
| ENSMUSG0000006951 | chr10 | Lyz1 | 0.08372588 | 0.891670083 |
| ENSMUSG0000001987 | chr10 | Fabp7 | -0.31840069 | 0.891680641 |
| ENSMUSG0000002919 | chr5 | Rfc1 | 0.05017597 | 0.891680641 |
| ENSMUSG0000003433 | chr15 | Zbed4 | 0.097872961 | 0.891722446 |
| ENSMUSG0000003826 | chr9 | Sema7a | 0.132445035 | 0.891722446 |
| ENSMUSG0000000133 | chr4 | Sync | -0.095363684 | 0.891861242 |
| ENSMUSG0000006248 | chr19 | Ifit3b | -0.382825468 | 0.891999459 |
| ENSMUSG0000006286 | chr7 | Zfp28 | -0.102068676 | 0.891999459 |
| ENSMUSG0000003990 | chr16 | Eva1c | -0.088956358 | 0.891999459 |
| ENSMUSG0000003411 | chr12 | Tmed8 | -0.059069022 | 0.891999459 |
| ENSMUSG0000005804 | chr4 | 4933430I17Rik | -0.248340875 | 0.891999459 |
| ENSMUSG0000003832 | chr2 | Trpc4ap | -0.035101567 | 0.891999459 |
| ENSMUSG0000005062 | chr11 | Ubald2 | 0.066812138 | 0.891999459 |
| ENSMUSG0000002136 | chr13 | Hivep1 | 0.079949185 | 0.891999459 |
| ENSMUSG0000003877 | chr3 | Sema6c | -0.103978669 | 0.89217513 |
| ENSMUSG0000000759 | chr14 | Tssk4 | -0.256868392 | 0.89217513 |
| ENSMUSG0000002236 | chr15 | Mtbp | 0.097442645 | 0.892330758 |
| ENSMUSG0000002879 | chr4 | Zfp362 | 0.073798522 | 0.892454515 |
| ENSMUSG0000004481 | chr11 | Cd300c2 | -0.21927083 | 0.892656322 |
| ENSMUSG0000002281 | chr16 | Zfp148 | -0.049539698 | 0.892867238 |
| ENSMUSG0000003344 | chr14 | Lpar6 | -0.074328532 | 0.893650792 |
| ENSMUSG0000000234 | chr8 | Tmem161a | 0.047637418 | 0.893656608 |
| ENSMUSG0000002746 | chr2 | Tpx2 | 0.223381877 | 0.894149315 |
| ENSMUSG0000002587 | chr13 | Thoc3 | 0.064727208 | 0.89432435 |
| ENSMUSG0000006018 | chr7 | Cxcl17 | 0.32427077 | 0.894408166 |
| ENSMUSG0000003904 | chr3 | Pigk | -0.049974125 | 0.894654226 |
| ENSMUSG0000002957 | chr5 | Radil | 0.083560422 | 0.894877293 |
| ENSMUSG0000002267 | chr16 | Snai2 | -0.10484049 | 0.894892119 |
| ENSMUSG0000004683 | chr15 | Krt1 | -0.122778059 | 0.894892119 |
| ENSMUSG0000002268 | chr16 | Rrn3 | 0.037136107 | 0.894892119 |
| ENSMUSG0000002197 | chr14 | Hmbox1 | -0.059105909 | 0.894942986 |
| ENSMUSG0000003799 | chr10 | Sh3rf3 | -0.107180801 | 0.894965006 |

| | | | | |
|---|---|---|---|---|
| ENSMUSG0000003587 | chr2 | Zhx3 | 0.095756728 | 0.894965006 |
| ENSMUSG0000002827 | chr4 | Ube2j1 | 0.041645687 | 0.895003599 |
| ENSMUSG0000003540 | chr7 | Emsy | 0.052756673 | 0.895003599 |
| ENSMUSG0000002153 | chr13 | Cfap90 | 0.333518893 | 0.895003599 |
| ENSMUSG0000008495 | chr19 | Bbip1 | 0.072097737 | 0.895062378 |
| ENSMUSG0000007855 | chr4 | Dcdc2b | -0.114850549 | 0.895238563 |
| ENSMUSG0000002151 | chr13 | Zfp729a | 0.066066899 | 0.895238563 |
| ENSMUSG0000002017 | chr11 | Cobl | 0.099031909 | 0.895285435 |
| ENSMUSG0000002614 | chr1 | Col19a1 | -0.547129126 | 0.895871007 |
| ENSMUSG0000003898 | chr13 | Bloc1s5 | -0.077731693 | 0.895871007 |
| ENSMUSG0000011044 | chr8 | Gm10033 | -0.068734417 | 0.895918437 |
| ENSMUSG0000003602 | chr14 | Parp2 | 0.064198704 | 0.896214981 |
| ENSMUSG0000000409 | chr9 | Col5a3 | -0.103553321 | 0.896359007 |
| ENSMUSG0000005055 | chr9 | Hyls1 | 0.107859932 | 0.896359007 |
| ENSMUSG0000002517 | chr19 | Pi4k2a | 0.061889433 | 0.896658107 |
| ENSMUSG0000002923 | chr5 | Srd5a3 | 0.059399896 | 0.896791229 |
| ENSMUSG0000004733 | chr1 | Kcne4 | 0.116056179 | 0.896817812 |
| ENSMUSG0000004845 | chr3 | Inka2 | 0.057236712 | 0.896817812 |
| ENSMUSG0000002952 | chr5 | Chek2 | 0.099821667 | 0.896884451 |
| ENSMUSG0000006979 | chr11 | Wfdc17 | 0.124826344 | 0.896884451 |
| ENSMUSG0000003032 | chr6 | Necap1 | -0.04999028 | 0.896932458 |
| ENSMUSG0000002008 | chr10 | Tacr2 | 0.077748808 | 0.896932458 |
| ENSMUSG0000003120 | chrX | Mtcp1 | -0.081784663 | 0.897013844 |
| ENSMUSG0000003053 | chr7 | Hddc3 | 0.092599526 | 0.897033194 |
| ENSMUSG0000003166 | chr8 | Gins3 | 0.128921528 | 0.897204704 |
| ENSMUSG0000003682 | chr17 | Amdhd2 | -0.095525945 | 0.897317509 |
| ENSMUSG0000005671 | chr2 | Gm13199 | 0.235036262 | 0.897317509 |
| ENSMUSG0000002917 | chr5 | Slc35f6 | -0.043022321 | 0.897348674 |
| ENSMUSG0000002909 | chr5 | Afap1 | 0.048037681 | 0.897430305 |
| ENSMUSG0000005177 | chr3 | Iqcj | -0.340481056 | 0.897496023 |
| ENSMUSG0000006310 | chr9 | Zfp26 | -0.071867628 | 0.897496023 |
| ENSMUSG0000000124 | chr7 | Hpn | -0.343842934 | 0.897496023 |
| ENSMUSG0000005880 | chr10 | Col13a1 | -0.087499194 | 0.897496023 |
| ENSMUSG0000001435 | chr2 | Tmem87b | -0.049788078 | 0.897496023 |
| ENSMUSG0000003971 | chr2 | Dync2i2 | 0.074922512 | 0.897496023 |
| ENSMUSG0000004087 | chr9 | Osbpl10 | 0.076697103 | 0.897496023 |
| ENSMUSG0000000073 | chr8 | Sult5a1 | 0.175182432 | 0.897496023 |
| ENSMUSG0000002238 | chr15 | Tef | -0.365872348 | 0.897868995 |
| ENSMUSG0000002799 | chr3 | Mcub | -0.112866991 | 0.897868995 |
| ENSMUSG0000007080 | chr7 | Bicra | -0.081169665 | 0.897868995 |
| ENSMUSG0000006104 | chr17 | Haghl | -0.09734641 | 0.897868995 |
| ENSMUSG0000003474 | chr6 | Nagk | -0.0711246 | 0.897868995 |
| ENSMUSG0000000303 | chr6 | Cdkn1b | -0.062765222 | 0.897868995 |
| ENSMUSG0000003609 | chr19 | Slf2 | -0.054236399 | 0.897868995 |
| ENSMUSG0000000499 | chr8 | Yju2b | -0.061276351 | 0.897868995 |
| ENSMUSG0000006323 | chr17 | Grm4 | -0.19251171 | 0.897868995 |
| ENSMUSG0000002866 | chr4 | Pithd1 | -0.050971004 | 0.897868995 |
| ENSMUSG0000004179 | chr11 | Gck | 0.350959761 | 0.897868995 |
| ENSMUSG0000003632 | chr5 | Srp72 | 0.045773699 | 0.897868995 |
| ENSMUSG0000002639 | chr1 | Cd55 | 0.064570174 | 0.897868995 |

| | | | | |
|---|---|---|---|---|
| ENSMUSG0000002667 | chr2 | Enkur | 0.070226139 | 0.897868995 |
| ENSMUSG0000004496 | chr5 | Napepld | 0.061570484 | 0.897868995 |
| ENSMUSG0000009444 | chr17 | Zfp955a | 0.066019577 | 0.897868995 |
| ENSMUSG0000000437 | chr7 | Il11 | 0.272783726 | 0.897868995 |
| ENSMUSG0000006663 | chr12 | Ttc32 | 0.124695472 | 0.897868995 |
| ENSMUSG0000002615 | chr1 | B3gat2 | 0.138214625 | 0.897868995 |
| ENSMUSG0000011865 | chr9 | Ube2srt | 0.260551796 | 0.897868995 |
| ENSMUSG0000005706 | chr8 | Slc35f3 | 0.247719622 | 0.897868995 |
| ENSMUSG0000005389 | chr3 | Slc39a8 | 0.267313827 | 0.897868995 |
| ENSMUSG0000004577 | chr14 | Lrtm1 | 0.19776139 | 0.897868995 |
| ENSMUSG0000000284 | chr16 | Timmdc1 | 0.077684634 | 0.898215569 |
| ENSMUSG0000002862 | chr4 | Cyb5rl | 0.077106892 | 0.898238582 |
| ENSMUSG0000007005 | chr8 | Mfhas1 | -0.056102027 | 0.898328886 |
| ENSMUSG0000003709 | chr17 | Rab11fip3 | 0.061119754 | 0.898328886 |
| ENSMUSG0000003129 | chrX | Adgrg2 | 0.100574965 | 0.898328886 |
| ENSMUSG0000003446 | chr14 | Scara3 | -0.126062773 | 0.898338609 |
| ENSMUSG0000001034 | chr11 | Tex14 | -0.357914534 | 0.898338609 |
| ENSMUSG0000003919 | chr7 | Rlbp1 | -0.27130012 | 0.898338609 |
| ENSMUSG0000002617 | chr1 | Ctdsp1 | 0.040200627 | 0.898338609 |
| ENSMUSG0000002589 | chr9 | Aasdhppt | 0.064574822 | 0.898338609 |
| ENSMUSG0000002166 | chr13 | Hexb | 0.072456584 | 0.898338609 |
| ENSMUSG0000002813 | chr3 | Rwdd3 | 0.077808389 | 0.898338609 |
| ENSMUSG0000004996 | chr4 | Plekhf2 | 0.064154291 | 0.898338609 |
| ENSMUSG0000007164 | chr19 | Rom1 | 0.078292239 | 0.898338609 |
| ENSMUSG0000003372 | chr1 | BC034090 | -0.070264097 | 0.898363139 |
| ENSMUSG0000003762 | chr1 | Kcnk2 | -0.049951875 | 0.898363139 |
| ENSMUSG0000002645 | chr1 | Klhl12 | -0.047333645 | 0.898363139 |
| ENSMUSG0000005917 | chr2 | Pde1a | 0.101399317 | 0.898363139 |
| ENSMUSG0000004222 | chr1 | Rabif | 0.044331639 | 0.898363139 |
| ENSMUSG0000006614 | chr4 | Prpf4 | 0.054759741 | 0.898363139 |
| ENSMUSG0000002048 | chr11 | Ccdc117 | 0.067146095 | 0.898363139 |
| ENSMUSG0000003146 | chr8 | Myom2 | 0.140580428 | 0.898363139 |
| ENSMUSG0000007461 | chr3 | Fundc2b | 0.329661714 | 0.898544605 |
| ENSMUSG0000002545 | chr13 | Paip1 | 0.053673237 | 0.898579971 |
| ENSMUSG0000003535 | chr11 | Kcnh4 | -0.272336685 | 0.898606841 |
| ENSMUSG0000002630 | chr1 | Iqca1 | -0.32355877 | 0.898606841 |
| ENSMUSG0000003926 | chr6 | Gimap3 | -0.195413187 | 0.898606841 |
| ENSMUSG0000002587 | chr13 | Hk3 | -0.203311661 | 0.898606841 |
| ENSMUSG0000003421 | chr4 | Efcab14 | -0.036363182 | 0.898606841 |
| ENSMUSG0000003658 | chr18 | Fgf1 | 0.074126336 | 0.898606841 |
| ENSMUSG0000005118 | chr1 | Fam174a | 0.068059093 | 0.898606841 |
| ENSMUSG0000003494 | chr10 | Zfr2 | 0.048180012 | 0.89873365 |
| ENSMUSG0000002191 | chr14 | Nisch | 0.041628141 | 0.898853297 |
| ENSMUSG0000003653 | chr17 | Cdc42ep3 | 0.079010246 | 0.898853297 |
| ENSMUSG0000003011 | chr6 | Ret | -0.080801666 | 0.899044421 |
| ENSMUSG0000001594 | chr3 | Fcgr1 | -0.21062673 | 0.899044421 |
| ENSMUSG0000002758 | chr2 | Zbtb46 | -0.088476836 | 0.899044421 |
| ENSMUSG0000002087 | chr11 | Snx11 | -0.064464665 | 0.899044421 |
| ENSMUSG0000002204 | chr14 | Ephx2 | -0.052840522 | 0.899044421 |
| ENSMUSG0000000082 | chr2 | Dnajc5 | -0.046543389 | 0.899044421 |

| | | | | |
|---|---|---|---|---|
| ENSMUSG0000003177 | chr8 | Arl2bp | 0.038982277 | 0.899044421 |
| ENSMUSG0000002767 | chr2 | Ncoa3 | 0.061261369 | 0.899044421 |
| ENSMUSG0000002863 | chr4 | Hivep3 | 0.088910118 | 0.899044421 |
| ENSMUSG0000004656 | chr10 | Rlig1 | 0.087694704 | 0.899044421 |
| ENSMUSG0000000803 | chrX | Mid1ip1 | 0.093758586 | 0.899044421 |
| ENSMUSG0000004765 | chr8 | Tssk6 | -0.292812063 | 0.899323189 |
| ENSMUSG0000002482 | chr19 | Rad9a | -0.063345768 | 0.899323189 |
| ENSMUSG0000003190 | chr8 | Zfp90 | 0.056230025 | 0.899323189 |
| ENSMUSG0000002166 | chr13 | Arhgef28 | -0.067936622 | 0.899434961 |
| ENSMUSG0000004066 | chr6 | Phc1 | -0.060785015 | 0.899434961 |
| ENSMUSG0000001029 | chr1 | AI597479 | -0.052241901 | 0.899434961 |
| ENSMUSG0000000338 | chr3 | Etv3 | -0.049175842 | 0.899434961 |
| ENSMUSG0000003306 | chr11 | Gas7 | 0.070531684 | 0.899434961 |
| ENSMUSG0000004224 | chr7 | Tmc7 | -0.106492515 | 0.899499632 |
| ENSMUSG0000003561 | chr12 | Togaram1 | -0.05109672 | 0.899499632 |
| ENSMUSG0000003725 | chr2 | Itih2 | 0.236990192 | 0.899499632 |
| ENSMUSG0000007072 | chr4 | Tmem200b | -0.077453971 | 0.899500637 |
| ENSMUSG0000004078 | chr16 | Ttc3 | -0.046004637 | 0.899500637 |
| ENSMUSG0000002710 | chr2 | Sp3 | 0.038562422 | 0.899589676 |
| ENSMUSG0000010534 | chr5 | Gm42878 | 0.11667833 | 0.899589676 |
| ENSMUSG0000002688 | chr2 | Ttll11 | 0.089780057 | 0.899589676 |
| ENSMUSG0000002923 | chr5 | Clock | 0.082895201 | 0.899589676 |
| ENSMUSG0000007135 | chr14 | Setdb2 | 0.074794709 | 0.899677149 |
| ENSMUSG0000001534 | chrX | Cybb | -0.167940379 | 0.899715116 |
| ENSMUSG0000002488 | chr19 | Asah2 | -0.077735906 | 0.899715116 |
| ENSMUSG0000005697 | chr8 | Ces1d | -0.105271045 | 0.899715116 |
| ENSMUSG0000002164 | chr13 | Serf1 | -0.113834584 | 0.899715116 |
| ENSMUSG0000002647 | chr1 | Rgs16 | -0.169458497 | 0.899715116 |
| ENSMUSG0000000455 | chr14 | Ndrg2 | -0.043624557 | 0.899715116 |
| ENSMUSG0000003482 | chr19 | Cpsf7 | -0.036544832 | 0.899715116 |
| ENSMUSG0000003427 | chr9 | Igsf9b | 0.079229264 | 0.899715116 |
| ENSMUSG0000004774 | chr16 | Fbxo40 | 0.294433336 | 0.899715116 |
| ENSMUSG0000003723 | chr7 | Spred3 | -0.098607376 | 0.899870081 |
| ENSMUSG0000002550 | chr7 | Taldo1 | 0.077597946 | 0.899870081 |
| ENSMUSG0000006631 | chr9 | Rtp3 | -0.288573937 | 0.90003184 |
| ENSMUSG0000007301 | chrX | Uprt | -0.093666657 | 0.90003184 |
| ENSMUSG0000004510 | chr6 | Slc25a26 | -0.088474661 | 0.90003184 |
| ENSMUSG0000002007 | chr10 | Rufy2 | -0.06082445 | 0.90003184 |
| ENSMUSG0000003719 | chr17 | Pacrg | -0.1007747 | 0.90003184 |
| ENSMUSG0000002665 | chr2 | Fam107b | -0.063856532 | 0.90003184 |
| ENSMUSG0000000410 | chr9 | Ppan | -0.059259204 | 0.90003184 |
| ENSMUSG0000000613 | chr16 | Crkl | -0.034003046 | 0.90003184 |
| ENSMUSG0000002061 | chr11 | Prkar1a | 0.06271204 | 0.90003184 |
| ENSMUSG0000003052 | chr7 | Mtmr10 | 0.049037896 | 0.90003184 |
| ENSMUSG0000003010 | chr6 | Sumf1 | 0.054325656 | 0.90003184 |
| ENSMUSG0000003678 | chrX | Klhl13 | 0.066003737 | 0.90003184 |
| ENSMUSG0000006772 | chr17 | Uqcc4 | 0.077375579 | 0.90003184 |
| ENSMUSG0000003239 | chr9 | Igdcc3 | 0.205158331 | 0.90003184 |
| ENSMUSG0000002607 | chr1 | Npas2 | 0.371306584 | 0.90003184 |
| ENSMUSG0000002235 | chr15 | Sqle | 0.124098767 | 0.900063766 |

| | | | | |
|---|---|---|---|---|
| ENSMUSG0000002910 | chr5 | Pcdh7 | 0.061360484 | 0.900112497 |
| ENSMUSG0000005531 | chr7 | Sec23ip | 0.047119147 | 0.900288024 |
| ENSMUSG0000004281 | chr18 | Gpr151 | 0.301459576 | 0.900288024 |
| ENSMUSG0000000279 | chr6 | Ggct | 0.122098401 | 0.900288024 |
| ENSMUSG0000002046 | chr11 | Cfap36 | 0.044558673 | 0.900709311 |
| ENSMUSG0000003725 | chr9 | Aagab | -0.050210227 | 0.900782463 |
| ENSMUSG0000002068 | chr11 | Ace | 0.131553424 | 0.900782463 |
| ENSMUSG0000004227 | chr13 | Pelo | 0.083771475 | 0.900782463 |
| ENSMUSG0000003886 | chr3 | Pi4kb | 0.043162954 | 0.900782463 |
| ENSMUSG0000002578 | chr9 | Zdhhc3 | 0.04288871 | 0.900782463 |
| ENSMUSG0000004718 | chr5 | Irs3 | 0.200909398 | 0.900782463 |
| ENSMUSG0000004268 | chr1 | Npl | 0.1172187 | 0.900782463 |
| ENSMUSG0000007831 | chrX | F8a | 0.062858595 | 0.900782463 |
| ENSMUSG0000004929 | chr14 | Zfp219 | 0.062579181 | 0.900782463 |
| ENSMUSG0000004782 | chr9 | Angptl8 | 0.155179374 | 0.900782463 |
| ENSMUSG0000002933 | chr5 | Fgf5 | 0.25923482 | 0.900782463 |
| ENSMUSG0000005540 | chr7 | Nell1 | 0.076483445 | 0.900783123 |
| ENSMUSG0000009140 | chr3 | H4c14 | -0.195066552 | 0.900915345 |
| ENSMUSG0000002031 | chr11 | Erlec1 | 0.068695717 | 0.900915345 |
| ENSMUSG0000003130 | chrX | Map3k15 | -0.200145956 | 0.900959818 |
| ENSMUSG0000004132 | chr11 | Atp1b2 | -0.050854169 | 0.900959818 |
| ENSMUSG0000003300 | chr8 | Ogfod1 | -0.041706181 | 0.900959818 |
| ENSMUSG0000002004 | chr10 | Hsp90b1 | 0.095048767 | 0.900959818 |
| ENSMUSG0000002409 | chr17 | Hnrnpll | 0.051351665 | 0.900959818 |
| ENSMUSG0000003375 | chr3 | Mnd1 | 0.172628534 | 0.900959818 |
| ENSMUSG0000008604 | chr6 | Wipf3 | 0.1026029 | 0.900959818 |
| ENSMUSG0000003923 | chr7 | Isg20 | 0.078471915 | 0.900959818 |
| ENSMUSG0000002883 | chr4 | Slc30a2 | -0.275767019 | 0.901261348 |
| ENSMUSG0000004464 | chr2 | Pard6b | -0.206174649 | 0.901261348 |
| ENSMUSG0000002047 | chr11 | Polm | -0.09741997 | 0.901261348 |
| ENSMUSG0000003686 | chr7 | Dchs1 | -0.079831122 | 0.901261348 |
| ENSMUSG0000005578 | chr15 | Abcd2 | -0.094171029 | 0.901261348 |
| ENSMUSG0000003442 | chr15 | Zfp707 | -0.071835388 | 0.901261348 |
| ENSMUSG0000002818 | chr3 | Ctbs | -0.05903279 | 0.901261348 |
| ENSMUSG0000009643 | chr17 | Zfp994 | -0.071718625 | 0.901261348 |
| ENSMUSG0000002071 | chr11 | Ddx5 | -0.040956125 | 0.901261348 |
| ENSMUSG0000000588 | chr2 | Ergic3 | 0.071059008 | 0.901261348 |
| ENSMUSG0000002507 | chr19 | Dclre1a | 0.067466324 | 0.901261348 |
| ENSMUSG0000004326 | chr7 | Uevld | 0.060213587 | 0.901261348 |
| ENSMUSG0000003893 | chr1 | Sccpdh | 0.057495176 | 0.901261348 |
| ENSMUSG0000002268 | chr16 | Parn | 0.062912915 | 0.901261348 |
| ENSMUSG0000005854 | chr11 | Rpl23a | 0.089174025 | 0.901261348 |
| ENSMUSG0000004028 | chr2 | Cdin1 | 0.11247169 | 0.901261348 |
| ENSMUSG0000000368 | chr19 | Taf6l | 0.098644382 | 0.901261348 |
| ENSMUSG0000003012 | chr6 | Cops7a | 0.038246975 | 0.901298399 |
| ENSMUSG0000002243 | chr15 | Csnk1e | 0.052617378 | 0.901298399 |
| ENSMUSG0000003755 | chr4 | Zdhhc18 | 0.050753995 | 0.901298399 |
| ENSMUSG0000006266 | chr2 | Ncs1 | -0.079508832 | 0.901504884 |
| ENSMUSG0000003256 | chr9 | Nudt16 | -0.065404879 | 0.901504884 |
| ENSMUSG0000004962 | chr13 | Tifab | -0.132733089 | 0.901504884 |

| | | | | |
|---|---|---|---|---|
| ENSMUSG0000000187 | chr17 | Ltbp1 | 0.059498548 | 0.901504884 |
| ENSMUSG0000003237 | chr9 | Plscr4 | -0.087424932 | 0.902136341 |
| ENSMUSG0000000088 | chr16 | Gnb1l | -0.11168491 | 0.902136341 |
| ENSMUSG0000007166 | chr19 | Ttc9c | -0.044998702 | 0.902136341 |
| ENSMUSG0000005685 | chr15 | Pcbp2 | 0.045925756 | 0.902229725 |
| ENSMUSG0000001882 | chr19 | Sfrp5 | 0.120740504 | 0.902229725 |
| ENSMUSG0000009958 | chr13 | H3c4 | -0.145589495 | 0.902392344 |
| ENSMUSG0000004567 | chr11 | Spred2 | -0.065193752 | 0.902392344 |
| ENSMUSG0000006174 | chr15 | Cyp2d22 | -0.063126752 | 0.902392344 |
| ENSMUSG0000002957 | chr5 | Lfng | -0.065807026 | 0.902392344 |
| ENSMUSG0000002247 | chr15 | Desi1 | -0.05901296 | 0.902392344 |
| ENSMUSG0000006383 | chr7 | Cdc42ep5 | -0.199362899 | 0.902392344 |
| ENSMUSG0000004470 | chr4 | Tmem201 | -0.051347237 | 0.902392344 |
| ENSMUSG0000007842 | chr10 | Ctdsp2 | 0.054659626 | 0.902392344 |
| ENSMUSG0000002013 | chr10 | 2310011J03Rik | 0.078941787 | 0.902392344 |
| ENSMUSG0000002135 | chr13 | Tfap2a | 0.20891391 | 0.902392344 |
| ENSMUSG0000007850 | chr4 | Zfp268 | 0.194487078 | 0.902392344 |
| ENSMUSG0000003515 | chr6 | Mitf | 0.089139448 | 0.902392344 |
| ENSMUSG0000000086 | chr11 | Bcl11a | 0.091699558 | 0.902392344 |
| ENSMUSG0000002303 | chr15 | Krt7 | 0.272928684 | 0.902392344 |
| ENSMUSG0000002687 | chr2 | Rab14 | 0.03513021 | 0.902545899 |
| ENSMUSG0000002913 | chr5 | Fosl2 | -0.075879548 | 0.902573764 |
| ENSMUSG0000002718 | chr2 | Fbxo3 | 0.045947431 | 0.902573764 |
| ENSMUSG0000000541 | chr11 | Mprip | 0.0600421 | 0.902573764 |
| ENSMUSG0000004163 | chr13 | Mrps27 | 0.070696555 | 0.902573764 |
| ENSMUSG0000002572 | chr7 | Nmb | -0.206087074 | 0.90274064 |
| ENSMUSG0000005989 | chr8 | Zfp930 | -0.074353776 | 0.90274064 |
| ENSMUSG0000007848 | chr4 | Perm1 | -0.106411483 | 0.90274064 |
| ENSMUSG0000003450 | chr12 | Pcnx4 | -0.058417983 | 0.90274064 |
| ENSMUSG0000003869 | chr2 | Mapkap1 | 0.03808416 | 0.90274064 |
| ENSMUSG0000002262 | chr15 | Shank3 | 0.099606351 | 0.90274064 |
| ENSMUSG0000004003 | chr3 | Negr1 | 0.061157698 | 0.9027506 |
| ENSMUSG0000003985 | chr11 | Endov | -0.071603351 | 0.902940576 |
| ENSMUSG0000003612 | chr8 | Rfxank | -0.072399709 | 0.902940576 |
| ENSMUSG0000002438 | chr18 | Iws1 | -0.048545771 | 0.902940576 |
| ENSMUSG0000000334 | chr10 | Btbd2 | -0.051115531 | 0.902940576 |
| ENSMUSG0000002610 | chr1 | Mstn | -0.268732017 | 0.902940576 |
| ENSMUSG0000003299 | chr4 | Foxj3 | -0.041441336 | 0.902940576 |
| ENSMUSG0000002036 | chr11 | Canx | 0.047791299 | 0.902940576 |
| ENSMUSG0000003173 | chr8 | Ap1g1 | -0.045842836 | 0.902945294 |
| ENSMUSG0000004267 | chr3 | Dcst1 | -0.182117106 | 0.903085076 |
| ENSMUSG0000003279 | chr17 | Lama1 | -0.119432291 | 0.903085076 |
| ENSMUSG0000002000 | chr10 | Ifngr1 | -0.063137902 | 0.903085076 |
| ENSMUSG0000001806 | chr11 | Ints2 | 0.055380256 | 0.903085076 |
| ENSMUSG0000005127 | chr3 | Zgrf1 | 0.121725009 | 0.903085076 |
| ENSMUSG0000003275 | chrX | Gab3 | -0.216103038 | 0.903177198 |
| ENSMUSG0000004149 | chr19 | Cep78 | -0.059519051 | 0.903177198 |
| ENSMUSG0000002843 | chr4 | Elp1 | -0.037687293 | 0.903177198 |
| ENSMUSG0000005616 | chr9 | Cnot10 | 0.035759868 | 0.903177198 |
| ENSMUSG0000007505 | chr13 | Yae1d1 | 0.056385429 | 0.903177198 |

| | | | | |
|---|---|---|---|---|
| ENSMUSG0000002251 | chr16 | Cldn1 | 0.221566598 | 0.903177198 |
| ENSMUSG0000003919 | chr8 | Zdhhc1 | -0.036824456 | 0.903365472 |
| ENSMUSG0000003395 | chr1 | Aspm | 0.288521728 | 0.903486786 |
| ENSMUSG0000004055 | chr5 | Mettl27 | -0.107931818 | 0.903572623 |
| ENSMUSG0000002085 | chr11 | Prpf8 | -0.033493476 | 0.903572623 |
| ENSMUSG0000000788 | chr4 | Arid1a | -0.077610464 | 0.903633385 |
| ENSMUSG0000002479 | chr19 | Tm7sf2 | -0.091004489 | 0.903633385 |
| ENSMUSG0000002104 | chr12 | Nova1 | -0.084629066 | 0.903633385 |
| ENSMUSG0000003910 | chr2 | Lsm14b | -0.048104665 | 0.903633385 |
| ENSMUSG0000003837 | chr7 | Sbf2 | 0.06085115 | 0.903633385 |
| ENSMUSG0000003246 | chr9 | Faim | 0.055761313 | 0.903633385 |
| ENSMUSG0000003226 | chr9 | Zw10 | 0.058959618 | 0.903633385 |
| ENSMUSG0000003222 | chr9 | Myo1e | 0.073358549 | 0.903633385 |
| ENSMUSG0000004361 | chr9 | Mmp3 | -0.189286993 | 0.903638893 |
| ENSMUSG0000005477 | chr1 | Kctd18 | -0.059736381 | 0.903638893 |
| ENSMUSG0000003322 | chr15 | Rac2 | -0.114715466 | 0.903638893 |
| ENSMUSG0000004722 | chr5 | Fam185a | 0.063638305 | 0.903952667 |
| ENSMUSG0000005653 | chr1 | Pign | -0.052003407 | 0.904035857 |
| ENSMUSG0000005838 | chr13 | H2bc8 | -0.135994787 | 0.904098103 |
| ENSMUSG0000002788 | chr3 | Cfap276 | -0.153223001 | 0.90416134 |
| ENSMUSG0000002505 | chr19 | Pcgf6 | -0.05593385 | 0.904189107 |
| ENSMUSG0000002036 | chr11 | Hspa4 | 0.05765962 | 0.904189107 |
| ENSMUSG0000003101 | chr7 | Wee1 | 0.304053606 | 0.904189107 |
| ENSMUSG0000006230 | chr9 | Rpp25 | 0.176267481 | 0.904198438 |
| ENSMUSG0000000454 | chr7 | Psg17 | 0.350655133 | 0.904198438 |
| ENSMUSG0000002524 | chrX | Tbl1x | 0.055592629 | 0.904202368 |
| ENSMUSG0000003870 | chr1 | Dsel | -0.048610879 | 0.9042835 |
| ENSMUSG0000003172 | chr8 | Zfp821 | 0.058670321 | 0.9042835 |
| ENSMUSG0000002681 | chr2 | St6galnac6 | -0.0622145 | 0.9044925 |
| ENSMUSG0000002412 | chr17 | Pdpk1 | -0.04662924 | 0.9044925 |
| ENSMUSG0000009621 | chr15 | H1f0 | 0.040346923 | 0.9044925 |
| ENSMUSG0000004045 | chr4 | Usp45 | 0.051592702 | 0.9044925 |
| ENSMUSG0000002747 | chr2 | Dnmt3b | 0.101066209 | 0.9044925 |
| ENSMUSG0000001174C | chr17 | Ntn3 | 0.060495126 | 0.9044925 |
| ENSMUSG0000005479 | chr7 | Cadm4 | 0.068139163 | 0.9044925 |
| ENSMUSG0000000639 | chr4 | Hyi | 0.074887848 | 0.9044925 |
| ENSMUSG0000004257 | chr10 | Mier2 | 0.062459516 | 0.9044925 |
| ENSMUSG0000002423 | chr18 | Bambi | 0.116384625 | 0.9044925 |
| ENSMUSG0000003996 | chr11 | Ccdc40 | -0.122868634 | 0.904510863 |
| ENSMUSG0000005092 | chrX | P2ry10 | -0.285922058 | 0.904510863 |
| ENSMUSG0000002983 | chr6 | Trim24 | -0.064245558 | 0.904510863 |
| ENSMUSG0000002503 | chr19 | Sfxn2 | -0.095028126 | 0.904510863 |
| ENSMUSG0000005890 | chr13 | Rsl1 | -0.083226608 | 0.904510863 |
| ENSMUSG0000002489 | chr19 | Slc29a2 | -0.084624139 | 0.904510863 |
| ENSMUSG0000002576 | chr3 | Jade1 | -0.054055823 | 0.904510863 |
| ENSMUSG0000004032 | chr2 | Zfp770 | 0.046824415 | 0.904510863 |
| ENSMUSG0000000095 | chr14 | Oxa1l | 0.064815454 | 0.904510863 |
| ENSMUSG0000003017 | chr6 | Ccdc77 | 0.090544658 | 0.904510863 |
| ENSMUSG0000003992 | chr16 | Urb1 | 0.067832745 | 0.904510863 |
| ENSMUSG0000004037 | chr3 | Pex2 | 0.067499272 | 0.904510863 |

| | | | | |
|---|---|---|---|---|
| ENSMUSG0000006723 | chr17 | H2-Q10 | 0.121151037 | 0.904510863 |
| ENSMUSG0000003195 | chr8 | Chst5 | 0.108481142 | 0.904510863 |
| ENSMUSG0000002430 | chr17 | Kifc5b | 0.255230131 | 0.904510863 |
| ENSMUSG0000002275 | chr16 | Klhl22 | -0.047174827 | 0.904526796 |
| ENSMUSG0000003211 | chr9 | Hyou1 | 0.086335425 | 0.904587458 |
| ENSMUSG0000003288 | chr1 | Acsl3 | -0.06572827 | 0.904625554 |
| ENSMUSG0000002552 | chr8 | Tmem192 | -0.084036303 | 0.904706958 |
| ENSMUSG0000002573 | chr17 | Rhbdl1 | 0.173572 | 0.904706958 |
| ENSMUSG0000000597 | chr2 | Rcn1 | 0.044894173 | 0.90475706 |
| ENSMUSG0000005285 | chrX | Nrk | 0.223716531 | 0.90475706 |
| ENSMUSG0000002461 | chr18 | Tcof1 | -0.036664145 | 0.904893976 |
| ENSMUSG0000002642 | chr1 | Arl8a | -0.045701177 | 0.905057523 |
| ENSMUSG0000003175 | chr8 | Il34 | -0.154388162 | 0.905089163 |
| ENSMUSG0000004736 | chr2 | Cstad | -0.402646295 | 0.905089163 |
| ENSMUSG0000000647 | chr2 | Ndor1 | -0.076289455 | 0.905182874 |
| ENSMUSG0000005443 | chr6 | Gimap4 | -0.077168696 | 0.905182874 |
| ENSMUSG0000003016 | chr6 | Adipor2 | 0.034084952 | 0.905182874 |
| ENSMUSG0000004096 | chr4 | Arhgef10l | 0.059575509 | 0.905182874 |
| ENSMUSG0000005143 | chr18 | Cd14 | 0.104607797 | 0.905182874 |
| ENSMUSG0000005089 | chr2 | Rtn4rl2 | -0.221265195 | 0.905235787 |
| ENSMUSG0000001930 | chr11 | Hsd17b1 | 0.279386044 | 0.905398648 |
| ENSMUSG0000002742 | chr2 | Kat14 | 0.047987142 | 0.905648816 |
| ENSMUSG0000003332 | chr18 | Ctdp1 | 0.059918882 | 0.905648816 |
| ENSMUSG0000000198 | chrX | Gria3 | 0.065667046 | 0.905743816 |
| ENSMUSG0000002975 | chr6 | Asns | 0.066661165 | 0.906024981 |
| ENSMUSG0000003271 | chr6 | Mansc1 | -0.078942717 | 0.906060931 |
| ENSMUSG0000004255 | chr9 | Sin3a | -0.046752702 | 0.906060931 |
| ENSMUSG0000009029 | chr8 | Tarbp1 | 0.063319801 | 0.906060931 |
| ENSMUSG0000000299 | chr12 | Prkar2b | 0.07421159 | 0.906060931 |
| ENSMUSG0000003932 | chr8 | Rnf122 | 0.094791479 | 0.906060931 |
| ENSMUSG0000006245 | chr19 | Rpl9-ps6 | 0.109117123 | 0.906060931 |
| ENSMUSG0000003663 | chr17 | Clcn7 | 0.04340353 | 0.906344946 |
| ENSMUSG0000002400 | chr17 | Ppil1 | 0.096412159 | 0.906344946 |
| ENSMUSG0000002417 | chr17 | Telo2 | -0.058029635 | 0.90646617 |
| ENSMUSG0000003078 | chr7 | Itgam | -0.188541018 | 0.906575203 |
| ENSMUSG0000007257 | chr14 | Slc39a2 | -0.137670946 | 0.907095719 |
| ENSMUSG0000003569 | chr4 | Rnf38 | -0.067384433 | 0.907128055 |
| ENSMUSG0000002573 | chr17 | Rhot2 | -0.049575897 | 0.907128055 |
| ENSMUSG0000003224 | chr9 | Cln6 | 0.075455572 | 0.907128055 |
| ENSMUSG0000000029 | chr10 | Itgb2 | 0.079384304 | 0.907128055 |
| ENSMUSG0000007052 | chr7 | Mkrn3 | 0.171150546 | 0.907128055 |
| ENSMUSG0000002548 | chr7 | Ifitm5 | 0.297462439 | 0.907128055 |
| ENSMUSG0000004531 | chr13 | Lhfpl2 | -0.062179554 | 0.907202288 |
| ENSMUSG0000003721 | chr3 | Spry1 | 0.062575704 | 0.907202288 |
| ENSMUSG0000003915 | chr6 | Ube2h | 0.044041818 | 0.907240968 |
| ENSMUSG0000003235 | chr9 | Lrrc1 | -0.106945653 | 0.907288174 |
| ENSMUSG0000003222 | chr9 | Mns1 | -0.078307808 | 0.907307235 |
| ENSMUSG0000000124 | chr7 | Lsr | -0.175196546 | 0.907324178 |
| ENSMUSG0000005010 | chr10 | Bpifc | -0.352510678 | 0.907324178 |
| ENSMUSG0000002079 | chr11 | Galr2 | -0.100755888 | 0.907324178 |

| | | | | |
|---|---|---|---|---|
| ENSMUSG0000005151 | chr4 | Arhgef39 | -0.119854864 | 0.907324178 |
| ENSMUSG0000002199 | chr14 | Lcp1 | -0.084903245 | 0.907324178 |
| ENSMUSG0000003745 | chr10 | Slc18b1 | -0.064892805 | 0.907324178 |
| ENSMUSG0000005988 | chr15 | Irak4 | 0.06287773 | 0.907324178 |
| ENSMUSG0000004071 | chr1 | Creg1 | 0.090462749 | 0.907324178 |
| ENSMUSG0000003004 | chr6 | Arhgap25 | 0.138569153 | 0.907324178 |
| ENSMUSG0000002723 | chr2 | Mdk | 0.072016904 | 0.907324178 |
| ENSMUSG0000001823 | chr11 | Gdf9 | 0.168989118 | 0.907324178 |
| ENSMUSG0000000191 | chr7 | Slc1a5 | 0.095740109 | 0.907324178 |
| ENSMUSG0000002518 | chr19 | Cnnm1 | -0.119258406 | 0.90741512 |
| ENSMUSG0000003425 | chr17 | Agpat1 | 0.042521795 | 0.907464638 |
| ENSMUSG0000002700 | chr2 | Itga4 | -0.099454897 | 0.907887297 |
| ENSMUSG0000003877 | chr10 | Ascc3 | -0.058150731 | 0.907887297 |
| ENSMUSG0000003166 | chr6 | Eif2ak3 | 0.054442371 | 0.907920281 |
| ENSMUSG0000002303 | chr15 | Slc11a2 | 0.06065907 | 0.907920281 |
| ENSMUSG0000004930 | chr3 | Prmt6 | -0.068291372 | 0.907981246 |
| ENSMUSG0000002445 | chr18 | Hdac3 | -0.042253397 | 0.908400978 |
| ENSMUSG0000002903 | chr4 | Acap3 | -0.067242287 | 0.90871587 |
| ENSMUSG0000003677 | chr8 | Tent4b | -0.049677945 | 0.90871587 |
| ENSMUSG0000002389 | chr17 | Zfp51 | 0.071972743 | 0.90871587 |
| ENSMUSG0000000859 | chr9 | Htr3b | 0.070858147 | 0.908780543 |
| ENSMUSG0000003176 | chr8 | Nudt7 | -0.097580129 | 0.908819727 |
| ENSMUSG0000002622 | chr1 | Itm2c | -0.068339976 | 0.908819727 |
| ENSMUSG0000007269 | chr5 | Gm57857 | -0.104685529 | 0.908819727 |
| ENSMUSG0000001618 | chr1 | Utp25 | -0.046723772 | 0.908819727 |
| ENSMUSG0000002510 | chr7 | Hdgfl3 | -0.048897036 | 0.908819727 |
| ENSMUSG0000000042 | chr18 | Galnt1 | 0.035866549 | 0.908819727 |
| ENSMUSG0000006014 | chr13 | Serpinb6a | 0.054921368 | 0.908819727 |
| ENSMUSG0000003537 | chr6 | Shq1 | 0.113461389 | 0.908819727 |
| ENSMUSG0000005094 | chr18 | Zfp438 | 0.079311536 | 0.908819727 |
| ENSMUSG0000002039 | chr11 | Med7 | 0.058178487 | 0.908819727 |
| ENSMUSG0000007945 | chrX | Cldn34c1 | 0.088224714 | 0.908819727 |
| ENSMUSG0000005329 | chr7 | Rab4b | -0.104859409 | 0.908863092 |
| ENSMUSG0000008723 | chr1 | Mroh3 | -0.153267554 | 0.908866409 |
| ENSMUSG0000000553 | chr7 | Igf1r | 0.058724385 | 0.908866409 |
| ENSMUSG0000002022 | chr10 | Llph | 0.061561272 | 0.908943934 |
| ENSMUSG0000002255 | chr15 | Maf1 | 0.065687996 | 0.908943934 |
| ENSMUSG0000004706 | chr1 | Dusp28 | 0.149259003 | 0.908943934 |
| ENSMUSG0000003158 | chr8 | Gsr | -0.063629667 | 0.909079267 |
| ENSMUSG0000002370 | chr5 | Ogfod2 | -0.08334529 | 0.909100938 |
| ENSMUSG0000010951 | chr7 | Nup62 | 0.036114043 | 0.909100938 |
| ENSMUSG0000003705 | chr18 | Paip2 | 0.068323664 | 0.909100938 |
| ENSMUSG0000003135 | chrX | Amelx | 0.306635711 | 0.909100938 |
| ENSMUSG0000000820 | chr2 | Fnbp4 | -0.074195774 | 0.909151163 |
| ENSMUSG0000004079 | chr4 | Iqcc | -0.06173831 | 0.909151163 |
| ENSMUSG0000000141 | chr3 | Mef2d | 0.075526661 | 0.909151163 |
| ENSMUSG0000003998 | chr6 | Sinhcaf | -0.105586878 | 0.909187879 |
| ENSMUSG0000002105 | chr12 | Ppp2r5e | -0.034625807 | 0.909187879 |
| ENSMUSG0000000075 | chr11 | Serpinf1 | 0.082278508 | 0.909187879 |
| ENSMUSG0000005645 | chr12 | Zbtb25 | -0.056439203 | 0.909443617 |

| | | | | |
|---|---|---|---|---|
| ENSMUSG0000004050 | chr17 | Abcg5 | 0.259471666 | 0.90963041 |
| ENSMUSG0000005144 | chr3 | Bbs12 | -0.09650798 | 0.909685811 |
| ENSMUSG0000000544 | chr7 | Cic | -0.061402808 | 0.909685811 |
| ENSMUSG0000003236 | chr9 | Adamts7 | -0.097121262 | 0.909685811 |
| ENSMUSG0000002407 | chr17 | Birc6 | -0.064640728 | 0.909685811 |
| ENSMUSG0000004361 | chr5 | Vps37d | -0.066835826 | 0.909685811 |
| ENSMUSG0000003984 | chr6 | Zfp800 | -0.057383816 | 0.909685811 |
| ENSMUSG0000000955 | chr2 | Cdk9 | 0.031474672 | 0.909685811 |
| ENSMUSG0000000562 | chr7 | Zfp592 | 0.046831005 | 0.909685811 |
| ENSMUSG0000003278 | chr15 | Troap | 0.837003592 | 0.909685811 |
| ENSMUSG0000002804 | chr3 | Gba1 | 0.046225777 | 0.909685811 |
| ENSMUSG0000002662 | chr1 | Pacc1 | 0.072293723 | 0.909685811 |
| ENSMUSG0000003399 | chr13 | Skic3 | 0.042362376 | 0.909816444 |
| ENSMUSG0000002224 | chr15 | Rai14 | -0.054545139 | 0.909905021 |
| ENSMUSG0000003914 | chr19 | Sart1 | 0.040965064 | 0.909905021 |
| ENSMUSG0000000626 | chr5 | Mob1b | 0.08506082 | 0.910086418 |
| ENSMUSG0000005233 | chr1 | Ankrd44 | 0.048344388 | 0.910086418 |
| ENSMUSG0000003238 | chr9 | Ciao2a | 0.085746764 | 0.910086418 |
| ENSMUSG0000009939 | chr19 | Ms4a14 | -0.247057622 | 0.910178039 |
| ENSMUSG0000003769 | chr4 | Ahdc1 | -0.055135969 | 0.910183656 |
| ENSMUSG0000002557 | chr11 | Gaa | 0.039813886 | 0.910276592 |
| ENSMUSG0000004476 | chr16 | Trmt10c | 0.054748853 | 0.910276592 |
| ENSMUSG0000004492 | chr10 | Rassf9 | 0.100798287 | 0.910276592 |
| ENSMUSG0000000241 | chr6 | Ndufb2 | -0.074605139 | 0.910291215 |
| ENSMUSG0000004394 | chr5 | Wdfy3 | -0.074284756 | 0.910291215 |
| ENSMUSG0000004928 | chr11 | Iba57 | -0.094452817 | 0.910291215 |
| ENSMUSG0000003752 | chr15 | Bcdin3d | -0.073976117 | 0.910291215 |
| ENSMUSG0000004350 | chr6 | Gimap5 | 0.158125685 | 0.910291215 |
| ENSMUSG0000004741 | chr9 | Zbtb44 | 0.056530653 | 0.910291215 |
| ENSMUSG0000000266 | chr17 | Alkbh7 | 0.110430874 | 0.910291215 |
| ENSMUSG0000003209 | chr9 | Ddx6 | -0.041273401 | 0.910351999 |
| ENSMUSG0000003658 | chr3 | Spart | 0.042569074 | 0.910351999 |
| ENSMUSG0000002013 | chr10 | Tbc1d15 | -0.073779844 | 0.910449902 |
| ENSMUSG0000001574 | chr3 | Prpf3 | 0.056690428 | 0.910449902 |
| ENSMUSG0000007926 | chr9 | Tmppe | -0.088870559 | 0.910456263 |
| ENSMUSG0000002157 | chr13 | Sdha | 0.035413627 | 0.910456263 |
| ENSMUSG0000002514 | chr11 | Lrrc45 | -0.040672813 | 0.910609919 |
| ENSMUSG0000002469 | chr19 | Fam111a | 0.057374389 | 0.910830302 |
| ENSMUSG0000000146 | chr5 | Cyp51 | 0.193723631 | 0.910857645 |
| ENSMUSG0000003367 | chr1 | Cep350 | -0.063114144 | 0.910863541 |
| ENSMUSG0000004279 | chr9 | Rnf214 | -0.056627077 | 0.910863541 |
| ENSMUSG0000003306 | chr1 | Resp18 | -0.073207002 | 0.911190074 |
| ENSMUSG0000000343 | chr7 | Dll3 | -0.316722852 | 0.91126749 |
| ENSMUSG0000003793 | chr5 | Scarb1 | -0.092132131 | 0.91126749 |
| ENSMUSG0000001387 | chr8 | Rnf170 | -0.056147067 | 0.91126749 |
| ENSMUSG0000003342 | chr6 | Antxr1 | 0.042148983 | 0.91126749 |
| ENSMUSG0000004270 | chr18 | Commd10 | 0.053890344 | 0.91126749 |
| ENSMUSG0000003817 | chr11 | Slc43a2 | 0.078502104 | 0.91126749 |
| ENSMUSG0000005409 | chr5 | Slc25a40 | -0.050924963 | 0.911268435 |
| ENSMUSG0000007073 | chr2 | Rmdn3 | -0.062214009 | 0.911268435 |

| | | | | |
|---|---|---|---|---|
| ENSMUSG0000002013 | chr10 | Thap2 | -0.047326306 | 0.911268435 |
| ENSMUSG0000000034 | chr3 | Dbt | 0.041571089 | 0.911268435 |
| ENSMUSG0000005650 | chr12 | 1700001K19Rik | 0.144808509 | 0.911268435 |
| ENSMUSG0000002035 | chr11 | Flt4 | 0.100995482 | 0.911268435 |
| ENSMUSG0000006143 | chr6 | Hipk2 | -0.099047139 | 0.91133409 |
| ENSMUSG0000002870 | chr4 | Mob3c | -0.073198745 | 0.91133409 |
| ENSMUSG0000001478 | chr8 | Plekhg4 | 0.295701947 | 0.911445687 |
| ENSMUSG0000002157 | chr13 | Ccdc127 | 0.048381849 | 0.911445687 |
| ENSMUSG0000004235 | chr12 | Arel1 | -0.05494533 | 0.911577597 |
| ENSMUSG0000003914 | chr2 | Camk1d | -0.067818435 | 0.911763777 |
| ENSMUSG0000010231 | chr18 | Pcdha3 | -0.165881626 | 0.911897631 |
| ENSMUSG0000003639 | chr6 | Gadd45a | 0.093028442 | 0.911897631 |
| ENSMUSG0000002627 | chr1 | Capn10 | -0.065227931 | 0.911946343 |
| ENSMUSG0000002782 | chr3 | Gmps | -0.038413764 | 0.911966087 |
| ENSMUSG0000005255 | chr13 | Adarb2 | -0.137283753 | 0.912048784 |
| ENSMUSG0000003606 | chr15 | Smug1 | -0.056955346 | 0.912048784 |
| ENSMUSG0000002430 | chr17 | Tapbp | -0.041579843 | 0.912048784 |
| ENSMUSG0000005231 | chr3 | Slc39a1 | 0.034499616 | 0.912048784 |
| ENSMUSG0000003255 | chr9 | Srprb | 0.074089381 | 0.912048784 |
| ENSMUSG0000002001 | chr10 | Cfap54 | -0.173094592 | 0.912079318 |
| ENSMUSG0000003459 | chr10 | Slc41a2 | -0.096131601 | 0.912079318 |
| ENSMUSG0000004500 | chr11 | Tubg2 | -0.056627568 | 0.912079318 |
| ENSMUSG0000004824 | chr10 | Gng7 | -0.075479893 | 0.912079318 |
| ENSMUSG0000002536 | chr10 | Smarcc2 | -0.047859314 | 0.912079318 |
| ENSMUSG0000002658 | chr1 | Scyl3 | -0.06400637 | 0.912079318 |
| ENSMUSG0000004218 | chr1 | Bend6 | -0.05454132 | 0.912079318 |
| ENSMUSG0000000175 | chr11 | Coasy | -0.047602988 | 0.912079318 |
| ENSMUSG0000003362 | chr5 | Pcgf3 | -0.039467192 | 0.912079318 |
| ENSMUSG0000007921 | chr5 | Zfp664 | 0.031939659 | 0.912079318 |
| ENSMUSG0000002858 | chr4 | Pdpn | 0.086728752 | 0.912079318 |
| ENSMUSG0000001422 | chr1 | Cacybp | 0.095328639 | 0.912079318 |
| ENSMUSG0000003722 | chr3 | Fgf2 | 0.041193468 | 0.912079318 |
| ENSMUSG0000003247 | chr9 | Map4 | 0.051699136 | 0.912079318 |
| ENSMUSG0000002624 | chr1 | Farsb | 0.050075763 | 0.912079318 |
| ENSMUSG0000001848 | chr11 | Appbp2 | 0.046285117 | 0.912079318 |
| ENSMUSG0000002516 | chr11 | Csnk1d | 0.040807392 | 0.912079318 |
| ENSMUSG0000007107 | chr10 | Ptges3 | 0.081772464 | 0.912079318 |
| ENSMUSG0000000404 | chr11 | Stat5a | 0.056034897 | 0.912079318 |
| ENSMUSG0000004976 | chr10 | Zfp280b | 0.045464466 | 0.912079318 |
| ENSMUSG0000002533 | chr4 | Padi4 | 0.183952039 | 0.912079318 |
| ENSMUSG0000009069 | chr6 | Apold1 | 0.119471653 | 0.912079318 |
| ENSMUSG0000001984 | chr10 | Rev3l | 0.061791836 | 0.912253707 |
| ENSMUSG0000006736 | chr5 | Tmem128 | 0.069054756 | 0.912253707 |
| ENSMUSG0000002201 | chr14 | Tnfsf11 | -0.578326801 | 0.912332977 |
| ENSMUSG0000004779 | chr1 | Sned1 | -0.108404781 | 0.912332977 |
| ENSMUSG0000002515 | chr11 | Fasn | -0.088337732 | 0.912332977 |
| ENSMUSG0000002237 | chr15 | Col14a1 | -0.123144149 | 0.912332977 |
| ENSMUSG0000005148 | chr18 | Pcdhb11 | -0.141416303 | 0.912332977 |
| ENSMUSG0000006092 | chr8 | Csmd1 | -0.100513771 | 0.912332977 |
| ENSMUSG0000007169 | chr19 | Top6bl | -0.218233372 | 0.912332977 |

| | | | | |
|---|---|---|---|---|
| ENSMUSG0000004234 | chr7 | Ctf1 | -0.128579144 | 0.912332977 |
| ENSMUSG0000002871 | chr4 | Tal1 | -0.127372688 | 0.912332977 |
| ENSMUSG0000003527 | chr9 | Tpbg | -0.087761223 | 0.912332977 |
| ENSMUSG0000002232 | chr15 | Pop1 | -0.087015523 | 0.912332977 |
| ENSMUSG0000003014 | chr6 | Clec4a2 | -0.129885959 | 0.912332977 |
| ENSMUSG0000001872 | chr11 | Cpsf4l | -0.112159795 | 0.912332977 |
| ENSMUSG0000001997 | chr10 | Cep290 | -0.05515857 | 0.912332977 |
| ENSMUSG0000002514 | chr11 | Pycr1 | -0.091147222 | 0.912332977 |
| ENSMUSG0000005462 | chrX | Xlr | -0.126896266 | 0.912332977 |
| ENSMUSG0000003257 | chr9 | Mapkapk3 | -0.076270598 | 0.912332977 |
| ENSMUSG0000005467 | chr7 | 1600014C10Rik | -0.049067258 | 0.912332977 |
| ENSMUSG0000007125 | chr10 | Slc25a16 | -0.0483176 | 0.912332977 |
| ENSMUSG0000004864 | chr8 | Ctxn1 | -0.050097007 | 0.912332977 |
| ENSMUSG0000006293 | chr10 | Zfp938 | -0.0526933 | 0.912332977 |
| ENSMUSG0000002642 | chr1 | Srgap2 | -0.051853514 | 0.912332977 |
| ENSMUSG0000003358 | chr14 | Reep4 | -0.07155128 | 0.912332977 |
| ENSMUSG0000003711 | chr15 | Fam91a1 | -0.040127954 | 0.912332977 |
| ENSMUSG0000003016 | chr6 | Gabarapl1 | 0.066999086 | 0.912332977 |
| ENSMUSG0000002600 | chr1 | Lancl1 | 0.041483973 | 0.912332977 |
| ENSMUSG0000002424 | chr18 | Epc1 | 0.039308066 | 0.912332977 |
| ENSMUSG0000002866 | chr4 | Eloa | 0.039774627 | 0.912332977 |
| ENSMUSG0000000982 | chr9 | Cilk1 | 0.046669814 | 0.912332977 |
| ENSMUSG0000002233 | chr15 | Ebag9 | 0.060756339 | 0.912332977 |
| ENSMUSG0000002039 | chr11 | Cdkn2aipnl | 0.064689046 | 0.912332977 |
| ENSMUSG0000002570 | chr6 | Alox5 | 0.110259578 | 0.912332977 |
| ENSMUSG0000002035 | chr11 | Phykpl | 0.066144314 | 0.912332977 |
| ENSMUSG0000003286 | chr2 | Rag2 | 0.219635798 | 0.912332977 |
| ENSMUSG0000000322 | chr10 | Edar | 0.194818295 | 0.912332977 |
| ENSMUSG0000003616 | chr2 | Surf6 | 0.048022605 | 0.912332977 |
| ENSMUSG0000002531 | chr8 | Banp | 0.095605664 | 0.912332977 |
| ENSMUSG0000002241 | chr15 | Cacna1i | 0.167936399 | 0.912332977 |
| ENSMUSG0000002113 | chr12 | Slc10a1 | 0.209501106 | 0.912332977 |
| ENSMUSG0000004551 | chr1 | Pou3f3 | -0.071786403 | 0.912399565 |
| ENSMUSG0000003514 | chr12 | Nubpl | -0.083556383 | 0.912399565 |
| ENSMUSG0000002072 | chr11 | Helz | -0.062710298 | 0.912399565 |
| ENSMUSG0000001264 | chr7 | Zfp715 | -0.048695346 | 0.912399565 |
| ENSMUSG0000004032 | chr9 | Dcaf1 | -0.044421936 | 0.912399565 |
| ENSMUSG0000001045 | chr1 | Kansl3 | 0.034645862 | 0.912399565 |
| ENSMUSG0000003543 | chr2 | Rabgap1 | 0.034618734 | 0.912399565 |
| ENSMUSG0000005107 | chr3 | Vtcn1 | 0.245163002 | 0.912399565 |
| ENSMUSG0000003762 | chr4 | Wdtc1 | 0.04278472 | 0.912399565 |
| ENSMUSG0000000453 | chr6 | Tax1bp1 | 0.04476158 | 0.912399565 |
| ENSMUSG0000003106 | chrX | Rbm10 | 0.039096061 | 0.912399565 |
| ENSMUSG0000003082 | chr7 | Bcat2 | 0.078150912 | 0.912399565 |
| ENSMUSG0000007165 | chr19 | Ints5 | 0.045825581 | 0.912399565 |
| ENSMUSG0000004763 | chr5 | Mtrfr | 0.084273589 | 0.912399565 |
| ENSMUSG0000003247 | chr9 | Nck1 | 0.057409359 | 0.912399565 |
| ENSMUSG0000003698 | chr7 | Trim3 | 0.048326968 | 0.912399565 |
| ENSMUSG0000007291 | chr12 | Noxred1 | 0.171330772 | 0.912399565 |
| ENSMUSG0000003050 | chr7 | Prmt3 | -0.049329406 | 0.912417101 |

| | | | | |
|---|---|---|---|---|
| ENSMUSG0000007901 | chr15 | Ly6c1 | 0.03922536 | 0.912417101 |
| ENSMUSG0000005548 | chr13 | Zfp458 | -0.11719956 | 0.912431499 |
| ENSMUSG0000002827 | chr4 | Rngtt | -0.047839879 | 0.912459023 |
| ENSMUSG0000003671 | chr8 | Cyld | -0.035259538 | 0.912719072 |
| ENSMUSG0000000358 | chr11 | Sec14l2 | -0.116886239 | 0.912885111 |
| ENSMUSG0000003745 | chr15 | Azin1 | 0.064422127 | 0.91296129 |
| ENSMUSG0000004094 | chr3 | Tet2 | -0.066463547 | 0.912987563 |
| ENSMUSG0000003009 | chr6 | Xpc | 0.050633497 | 0.912987563 |
| ENSMUSG0000002523 | chr9 | Hexa | -0.047073191 | 0.91312962 |
| ENSMUSG0000004891 | chr2 | Rnf24 | 0.072690456 | 0.91312962 |
| ENSMUSG0000002314 | chr16 | Get1 | 0.056493817 | 0.91312962 |
| ENSMUSG0000003415 | chr16 | Lrrc58 | -0.065369849 | 0.913165088 |
| ENSMUSG0000003025 | chr6 | Bhlhe41 | -0.26575497 | 0.913326855 |
| ENSMUSG0000002549 | chr7 | Ifitm1 | -0.122344745 | 0.913326855 |
| ENSMUSG0000007459 | chr3 | Ankub1 | -0.280691071 | 0.913326855 |
| ENSMUSG0000000074 | chr8 | Chmp1a | -0.054946746 | 0.913326855 |
| ENSMUSG0000005612 | chr18 | B4galt6 | -0.04709862 | 0.913326855 |
| ENSMUSG0000003402 | chr5 | Pds5b | -0.044910328 | 0.913326855 |
| ENSMUSG0000005992 | chr11 | Grb2 | 0.039685237 | 0.913326855 |
| ENSMUSG0000000484 | chr16 | Chmp2b | 0.05653606 | 0.913326855 |
| ENSMUSG0000006951 | chr10 | Lyz2 | 0.050733007 | 0.913326855 |
| ENSMUSG0000003001 | chr6 | Reg3g | 0.732775312 | 0.913326855 |
| ENSMUSG0000002067 | chr11 | Ccl11 | -0.119971965 | 0.913430608 |
| ENSMUSG0000006424 | chr1 | Chi3l1 | -0.13596437 | 0.913430608 |
| ENSMUSG0000004419 | chr5 | Gpr146 | -0.084007106 | 0.913430608 |
| ENSMUSG0000003818 | chr5 | Chpf2 | -0.070182552 | 0.913430608 |
| ENSMUSG0000002083 | chr11 | Eral1 | -0.058594249 | 0.913430608 |
| ENSMUSG0000001841 | chr11 | Gid4 | -0.052656307 | 0.913430608 |
| ENSMUSG0000004569 | chr14 | Thtpa | -0.065143314 | 0.913430608 |
| ENSMUSG0000000331 | chr8 | Glg1 | -0.045279216 | 0.913430608 |
| ENSMUSG0000003566 | chr2 | Gtf3c4 | -0.037530566 | 0.913430608 |
| ENSMUSG0000002598 | chr1 | Sf3b1 | 0.037665354 | 0.913430608 |
| ENSMUSG0000002880 | chr4 | Csmd2 | 0.125519004 | 0.913430608 |
| ENSMUSG0000003042 | chr7 | Uri1 | 0.046152163 | 0.913430608 |
| ENSMUSG0000004999 | chr2 | Ppp1r3d | 0.062224041 | 0.913430608 |
| ENSMUSG0000002844 | chr4 | Dctn3 | 0.073509243 | 0.913430608 |
| ENSMUSG0000000788 | chr8 | Crlf1 | 0.136884658 | 0.913430608 |
| ENSMUSG0000006345 | chr12 | Syne2 | 0.066187473 | 0.913430608 |
| ENSMUSG0000003201 | chr9 | Usp2 | 0.244890314 | 0.913606089 |
| ENSMUSG0000007166 | chr16 | Snx29 | -0.075879196 | 0.913608731 |
| ENSMUSG0000001886 | chr11 | Otop3 | -0.124783172 | 0.913608731 |
| ENSMUSG0000002896 | chr5 | Slc4a2 | -0.039129457 | 0.913608731 |
| ENSMUSG0000007541 | chr11 | Prcd | 0.286423832 | 0.913608731 |
| ENSMUSG0000007462 | chr2 | Mafb | 0.103000998 | 0.913608731 |
| ENSMUSG0000005775 | chr4 | Megf6 | 0.083351403 | 0.913608731 |
| ENSMUSG0000004806 | chr7 | Cyb5r2 | -0.090378749 | 0.913706188 |
| ENSMUSG0000004596 | chr1 | Teddm2 | -0.211529147 | 0.913706188 |
| ENSMUSG0000003516 | chr9 | Zc3h12c | 0.071240693 | 0.913706188 |
| ENSMUSG0000004163 | chrX | Kctd12b | -0.079130835 | 0.913757721 |
| ENSMUSG0000002770 | chr3 | Sec62 | 0.035663527 | 0.914022682 |

| | | | | |
|---|---|---|---|---|
| ENSMUSG0000001476 | chr17 | Fam120b | -0.052235262 | 0.914078574 |
| ENSMUSG0000008622 | chr9 | Ubap1l | -0.154447686 | 0.914084179 |
| ENSMUSG0000004287 | chr8 | Tom1 | -0.067352277 | 0.914195505 |
| ENSMUSG0000002019 | chr10 | Mknk2 | 0.087240204 | 0.914195505 |
| ENSMUSG0000001767 | chr2 | Ttpal | 0.037356214 | 0.91441617 |
| ENSMUSG0000007447 | chr9 | Zfp872 | -0.240689062 | 0.914645553 |
| ENSMUSG0000004137 | chr7 | Ccdc9 | -0.064877939 | 0.914709156 |
| ENSMUSG0000003723 | chr8 | Hook3 | -0.056341449 | 0.914709156 |
| ENSMUSG0000005710 | chr6 | Nat8f1 | 0.116960995 | 0.91482176 |
| ENSMUSG0000002669 | chr1 | Fmo3 | 0.372965324 | 0.914965536 |
| ENSMUSG0000002224 | chr15 | Ctnnd2 | -0.06687007 | 0.914992373 |
| ENSMUSG0000002565 | chr9 | Col7a1 | -0.121118914 | 0.914992373 |
| ENSMUSG0000000598 | chr16 | Dnase1 | -0.086524622 | 0.914992373 |
| ENSMUSG0000000964 | chr10 | Mcu | -0.049403452 | 0.914992373 |
| ENSMUSG0000004858 | chr7 | Igf2 | -0.164183712 | 0.914992373 |
| ENSMUSG0000002665 | chr2 | Meig1 | 0.184078491 | 0.914992373 |
| ENSMUSG0000002983 | chr6 | Nfe2l3 | 0.150492215 | 0.914992373 |
| ENSMUSG0000002068 | chr11 | Mmp28 | 0.102363205 | 0.914992373 |
| ENSMUSG0000004379 | chr7 | Prr33 | -0.08777632 | 0.914997456 |
| ENSMUSG0000002803 | chr3 | Papss1 | 0.053308125 | 0.914997456 |
| ENSMUSG0000004523 | chrX | Eola1 | 0.073753365 | 0.914997456 |
| ENSMUSG0000002211 | chr14 | Uchl3 | 0.070123822 | 0.914997456 |
| ENSMUSG0000001937 | chr7 | Calm3 | -0.048110699 | 0.915005098 |
| ENSMUSG0000003783 | chr14 | Sh2d4b | -0.267567337 | 0.915021689 |
| ENSMUSG0000007164 | chr19 | Eml3 | -0.07580214 | 0.915021689 |
| ENSMUSG0000002221 | chr14 | Emc9 | -0.084371012 | 0.915021689 |
| ENSMUSG0000003647 | chr10 | Btg1 | -0.052383064 | 0.915021689 |
| ENSMUSG0000005897 | chr6 | Hdhd5 | -0.047068491 | 0.915021689 |
| ENSMUSG0000005229 | chr16 | Ltn1 | -0.040044319 | 0.915021689 |
| ENSMUSG0000004598 | chr11 | Tmem104 | -0.043102383 | 0.915021689 |
| ENSMUSG0000004301 | chr1 | Edem3 | -0.045263846 | 0.915021689 |
| ENSMUSG0000002851 | chr4 | Usp24 | -0.03739455 | 0.915021689 |
| ENSMUSG0000003593 | chr8 | Chst4 | 0.271500868 | 0.915021689 |
| ENSMUSG0000002177 | chr14 | Thrb | 0.106361119 | 0.915021689 |
| ENSMUSG0000004213 | chr2 | Ppig | 0.042915099 | 0.915021689 |
| ENSMUSG0000005447 | chr6 | Thnsl2 | 0.064536142 | 0.915021689 |
| ENSMUSG0000009614 | chr7 | Vkorc1 | 0.087523285 | 0.915021689 |
| ENSMUSG0000009536 | chr2 | Gm14325 | 0.054236031 | 0.915021689 |
| ENSMUSG0000004031 | chr17 | Cchcr1 | 0.05656475 | 0.915021689 |
| ENSMUSG0000000744 | chr18 | Pcdha10 | 0.208329184 | 0.915021689 |
| ENSMUSG0000004039 | chr7 | Map3k10 | 0.061133591 | 0.915021689 |
| ENSMUSG0000003754 | chr14 | Dlgap5 | 0.266845712 | 0.915021689 |
| ENSMUSG0000005084 | chr15 | Zfp623 | -0.068977684 | 0.91505339 |
| ENSMUSG0000006305 | chr3 | Lrrc40 | -0.046388032 | 0.91505339 |
| ENSMUSG0000003325 | chr1 | Ttll4 | 0.053602927 | 0.91505339 |
| ENSMUSG0000003490 | chr2 | Ncaph | 0.15845886 | 0.91505339 |
| ENSMUSG0000002051 | chr11 | Sap30l | 0.059116869 | 0.91505339 |
| ENSMUSG0000007380 | chr7 | Insyn2a | 0.13660957 | 0.915263758 |
| ENSMUSG0000003908 | chr11 | Chad | -0.300411969 | 0.915320228 |
| ENSMUSG0000011017 | chr11 | St6galnac2 | -0.264300782 | 0.915320228 |

| | | | | |
|---|---|---|---|---|
| ENSMUSG0000003252 | chr9 | Nktr | -0.072106108 | 0.915320228 |
| ENSMUSG0000006390 | chr19 | Dpp3 | -0.049783748 | 0.915320228 |
| ENSMUSG0000004204 | chr1 | Dstyk | -0.043143534 | 0.915320228 |
| ENSMUSG0000002302 | chr15 | Lima1 | -0.050247736 | 0.915320228 |
| ENSMUSG0000002497 | chr19 | Smc3 | 0.036770506 | 0.915320228 |
| ENSMUSG0000005269 | chr9 | Tln2 | 0.058070055 | 0.915320228 |
| ENSMUSG0000000506 | chr6 | Pex5 | 0.035329654 | 0.915320228 |
| ENSMUSG0000003841 | chr10 | Fig4 | 0.051251185 | 0.915320228 |
| ENSMUSG0000003166 | chr8 | Brd7 | 0.047210228 | 0.915320228 |
| ENSMUSG0000007878 | chr7 | Cyp2t4 | 0.158374208 | 0.915320228 |
| ENSMUSG0000001756 | chr11 | Crlf3 | 0.052946904 | 0.915320228 |
| ENSMUSG0000002763 | chr2 | Dsn1 | 0.140901716 | 0.915320228 |
| ENSMUSG0000006087 | chr19 | Or10v1 | 0.264098889 | 0.915320228 |
| ENSMUSG0000001460 | chr11 | Slc25a11 | 0.039429812 | 0.91566768 |
| ENSMUSG0000003211 | chr9 | Trappc4 | 0.076642396 | 0.91566768 |
| ENSMUSG0000002808 | chr3 | Lrba | 0.049055535 | 0.91566768 |
| ENSMUSG0000007130 | chr8 | Zfp1006 | 0.060206192 | 0.91566768 |
| ENSMUSG0000003200 | chr9 | Pdgfd | 0.058087756 | 0.91566768 |
| ENSMUSG0000002483 | chr19 | Rps6kb2 | 0.062805814 | 0.91566768 |
| ENSMUSG0000001721 | chr11 | Med24 | 0.048178115 | 0.91566768 |
| ENSMUSG0000006972 | chr17 | Arid1b | 0.059285545 | 0.91566768 |
| ENSMUSG0000002692 | chr2 | Entr1 | -0.033495746 | 0.91570743 |
| ENSMUSG0000007370 | chr4 | Klhl21 | 0.045110665 | 0.91570743 |
| ENSMUSG0000003587 | chr2 | AI182371 | -0.218762867 | 0.915714744 |
| ENSMUSG0000004392 | chrX | Klhl15 | -0.072383656 | 0.915834832 |
| ENSMUSG0000002697 | chr2 | Marchf7 | -0.038727687 | 0.916020416 |
| ENSMUSG0000002018 | chr10 | E2f7 | 0.384535857 | 0.916020416 |
| ENSMUSG0000002717 | chr2 | Cstf3 | -0.052106335 | 0.916032542 |
| ENSMUSG0000003072 | chr7 | Nfatc2ip | -0.055786946 | 0.916081921 |
| ENSMUSG0000000082 | chr2 | Zfp512b | -0.05280686 | 0.91617975 |
| ENSMUSG0000001588 | chr5 | Ncapg | -0.365988681 | 0.91617975 |
| ENSMUSG0000002190 | chr14 | Oxnad1 | 0.052470966 | 0.916226376 |
| ENSMUSG0000003473 | chr9 | Mfrp | 0.203831801 | 0.916226376 |
| ENSMUSG0000005440 | chr8 | Spcs3 | -0.047970053 | 0.916255138 |
| ENSMUSG0000003652 | chr7 | Ppfibp2 | 0.075550029 | 0.916255138 |
| ENSMUSG0000002073 | chr11 | Jpt1 | 0.088670537 | 0.916255138 |
| ENSMUSG0000003741 | chr5 | Serpine1 | -0.130410885 | 0.916520751 |
| ENSMUSG0000004059 | chr11 | Ranbp17 | -0.091232143 | 0.916520751 |
| ENSMUSG0000002396 | chr17 | Enpp5 | 0.024515491 | 0.916520751 |
| ENSMUSG0000003619 | chr15 | Gxylt1 | 0.049830492 | 0.916520751 |
| ENSMUSG0000002481 | chr19 | Tjp2 | -0.066000415 | 0.91659213 |
| ENSMUSG0000003430 | chr9 | Ccdc15 | -0.103833661 | 0.916621245 |
| ENSMUSG0000003171 | chr8 | Smarca5 | -0.04397467 | 0.916621245 |
| ENSMUSG0000002953 | chr5 | Gatc | 0.039120606 | 0.916621245 |
| ENSMUSG0000003467 | chr10 | Tdg | -0.117179201 | 0.916636219 |
| ENSMUSG0000004524 | chr8 | Kcng4 | -0.076645563 | 0.916636219 |
| ENSMUSG0000001183 | chr5 | Evi5 | 0.034700557 | 0.916649241 |
| ENSMUSG0000004168 | chr1 | Bivm | -0.047843843 | 0.917018088 |
| ENSMUSG0000002965 | chr5 | Wdr95 | -0.312405284 | 0.917032975 |
| ENSMUSG0000002203 | chr14 | Pbk | -0.240616395 | 0.917095311 |

| | | | | |
|---|---|---|---|---|
| ENSMUSG0000002890 | chr4 | Ptpru | -0.079821113 | 0.917095311 |
| ENSMUSG0000005949 | chr1 | Fcgr3 | -0.072024856 | 0.917095311 |
| ENSMUSG0000005082 | chr17 | Sstr5 | -0.192608687 | 0.917095311 |
| ENSMUSG0000003695 | chr7 | Lrfn3 | -0.045783905 | 0.917095311 |
| ENSMUSG0000001845 | chr2 | Slc13a3 | -0.170528745 | 0.917095311 |
| ENSMUSG0000003217 | chr9 | Icam5 | -0.258861639 | 0.917095311 |
| ENSMUSG0000005955 | chr11 | Trp53 | 0.037782147 | 0.917095311 |
| ENSMUSG0000002407 | chr17 | Strn | 0.040674169 | 0.917095311 |
| ENSMUSG0000002060 | chr11 | Arsg | 0.121840759 | 0.917095311 |
| ENSMUSG0000002758 | chr2 | Zgpat | 0.040576799 | 0.917095311 |
| ENSMUSG0000009077 | chr16 | Ccdc188 | 0.209649715 | 0.917095311 |
| ENSMUSG0000009221 | chr9 | Slc22a13b | 0.212068404 | 0.917095311 |
| ENSMUSG0000005041 | chr17 | Tcf19 | -0.130588121 | 0.917120169 |
| ENSMUSG0000003083 | chr7 | Abcc6 | -0.177341425 | 0.917120169 |
| ENSMUSG0000005461 | chr15 | Tmt1a | -0.090121155 | 0.917120169 |
| ENSMUSG0000002743 | chr2 | Cd93 | -0.078834914 | 0.917120169 |
| ENSMUSG0000006042 | chr15 | Sntb1 | -0.060645138 | 0.917120169 |
| ENSMUSG0000003490 | chr10 | Pip5k1c | -0.030410215 | 0.917120169 |
| ENSMUSG0000005092 | chrX | Dcaf12l2 | -0.335756611 | 0.917120169 |
| ENSMUSG0000002195 | chr14 | Tkt | 0.036482649 | 0.917120169 |
| ENSMUSG0000006351 | chr7 | Snrnp70 | 0.062990039 | 0.917120169 |
| ENSMUSG0000004154 | chr4 | Disp3 | 0.156225423 | 0.917120169 |
| ENSMUSG0000005766 | chr7 | Bloc1s3 | 0.084940831 | 0.917120169 |
| ENSMUSG0000002294 | chr16 | Ripply3 | 0.164625923 | 0.917120169 |
| ENSMUSG0000001126 | chr7 | Zfp296 | 0.199814383 | 0.917120169 |
| ENSMUSG0000009847 | chr6 | C1rb | 0.551236044 | 0.917120169 |
| ENSMUSG0000004081 | chr2 | Agbl2 | -0.102894348 | 0.917188412 |
| ENSMUSG0000002734 | chr2 | Tmem230 | -0.046057648 | 0.917188412 |
| ENSMUSG0000002828 | chr4 | Map3k7 | 0.034411524 | 0.917190644 |
| ENSMUSG0000011451 | chr7 | Gm57854 | -0.173933571 | 0.917224885 |
| ENSMUSG0000005742 | chrX | Las1l | 0.049766703 | 0.917224885 |
| ENSMUSG0000004103 | chr19 | Dpcd | 0.060053422 | 0.917224885 |
| ENSMUSG0000003140 | chrX | Mpp1 | 0.050800538 | 0.917224885 |
| ENSMUSG0000000081 | chr6 | Txnrd3 | 0.087759614 | 0.917236167 |
| ENSMUSG0000003529 | chr5 | Cops4 | 0.051980123 | 0.917251402 |
| ENSMUSG0000002558 | chr8 | Nat1 | -0.346685025 | 0.917447597 |
| ENSMUSG0000004281 | chr6 | Krba1 | -0.067109058 | 0.917447597 |
| ENSMUSG0000007474 | chr2 | Kiz | -0.067203184 | 0.917447597 |
| ENSMUSG0000002512 | chr11 | Gcgr | -0.100903678 | 0.917447597 |
| ENSMUSG0000003371 | chr14 | Ccar2 | -0.046551768 | 0.917447597 |
| ENSMUSG0000005875 | chr11 | Thra | -0.046862111 | 0.917447597 |
| ENSMUSG0000002882 | chr4 | Maco1 | 0.083829435 | 0.917447597 |
| ENSMUSG0000004152 | chr9 | Rnf123 | 0.035134735 | 0.917447597 |
| ENSMUSG0000003577 | chrX | Cd99l2 | 0.040888577 | 0.917447597 |
| ENSMUSG0000006080 | chr2 | B2m | 0.051638067 | 0.917447597 |
| ENSMUSG0000004168 | chr13 | Fcho2 | 0.047218582 | 0.917447597 |
| ENSMUSG0000002265 | chr16 | Morc1 | 0.171050278 | 0.917447597 |
| ENSMUSG0000006421 | chr12 | Ifi27 | 0.064265255 | 0.917447597 |
| ENSMUSG0000001875 | chr11 | Tnfsfm13 | 0.130969279 | 0.917447597 |
| ENSMUSG0000004435 | chr11 | Sowaha | -0.169015961 | 0.917476398 |

| | | | | |
|---|---|---|---|---|
| ENSMUSG0000002242 | chr15 | Dscc1 | -0.274819477 | 0.917476398 |
| ENSMUSG0000001927 | chr8 | Dpep1 | -0.094682449 | 0.917476398 |
| ENSMUSG000000022C | chr7 | Vrk3 | -0.060222374 | 0.917476398 |
| ENSMUSG0000002785 | chr3 | Sike1 | 0.027948177 | 0.917476398 |
| ENSMUSG0000003364 | chr14 | Piwil2 | 0.172901561 | 0.917476398 |
| ENSMUSG0000003804 | chr13 | Ptpdc1 | 0.055946472 | 0.917476398 |
| ENSMUSG0000003177 | chr8 | Herpud1 | 0.102689622 | 0.917476398 |
| ENSMUSG0000002246 | chr15 | Rpap3 | 0.051646101 | 0.917517686 |
| ENSMUSG000000440C | chr8 | Cilp2 | -0.19613723 | 0.917630538 |
| ENSMUSG0000005602 | chr3 | Clca3a1 | -0.107553344 | 0.917630538 |
| ENSMUSG0000002245 | chr15 | Naga | -0.057057732 | 0.917630538 |
| ENSMUSG000000749C | chr19 | Ranbp6 | -0.037043705 | 0.917630538 |
| ENSMUSG0000002475 | chr19 | Zfand5 | 0.038982719 | 0.917630538 |
| ENSMUSG0000000553 | chr8 | Insr | 0.046024386 | 0.917630538 |
| ENSMUSG0000002735 | chr2 | Crls1 | 0.071105394 | 0.917630538 |
| ENSMUSG0000001994 | chr10 | Actr6 | 0.082896787 | 0.917630538 |
| ENSMUSG0000002157 | chr13 | Cep72 | 0.096130552 | 0.917630538 |
| ENSMUSG0000002991 | chr6 | Agk | -0.048337332 | 0.917711526 |
| ENSMUSG0000003259 | chr9 | Cdhr4 | -0.27454205 | 0.917725077 |
| ENSMUSG0000000493 | chr9 | Map2k1 | -0.04999666 | 0.917725077 |
| ENSMUSG0000004415 | chr6 | Lsm8 | 0.065023365 | 0.917725077 |
| ENSMUSG0000002717 | chr2 | Depdc7 | 0.111634055 | 0.91783886 |
| ENSMUSG0000002126 | chr12 | Hhipl1 | -0.134648889 | 0.917942241 |
| ENSMUSG0000003742 | chr5 | Depdc5 | -0.063205044 | 0.917984079 |
| ENSMUSG0000004055 | chr6 | C3ar1 | -0.097858868 | 0.917984079 |
| ENSMUSG0000005062 | chr19 | Catsperz | -0.246001436 | 0.917984079 |
| ENSMUSG0000005419 | chr15 | Cthrc1 | -0.084856984 | 0.917984079 |
| ENSMUSG000000219C | chr14 | Bap1 | 0.028367353 | 0.917984079 |
| ENSMUSG0000000248 | chr5 | Tchp | 0.079671573 | 0.917984079 |
| ENSMUSG0000002573 | chr17 | Rab40c | -0.047831947 | 0.918024357 |
| ENSMUSG0000003068 | chr7 | Mvp | -0.050962073 | 0.918024357 |
| ENSMUSG0000000446 | chr16 | Dnajb11 | 0.065067249 | 0.918024357 |
| ENSMUSG0000001017 | chr1 | Prox1 | 0.068388671 | 0.918174538 |
| ENSMUSG0000002914 | chr5 | Ppm1g | 0.03535684 | 0.918260227 |
| ENSMUSG0000002701 | chr2 | Cybrd1 | -0.223170827 | 0.918551113 |
| ENSMUSG0000002154 | chr13 | Trpc7 | -0.159851495 | 0.918551113 |
| ENSMUSG0000008968 | chr4 | Rab42 | -0.230790992 | 0.918551113 |
| ENSMUSG0000004071 | chr14 | Il17rd | -0.062767009 | 0.918551113 |
| ENSMUSG0000007132 | chr10 | Armc2 | -0.063637103 | 0.918551113 |
| ENSMUSG0000003168 | chr8 | Smad1 | -0.043728418 | 0.918551113 |
| ENSMUSG0000002621 | chr1 | Obsl1 | -0.039757107 | 0.918551113 |
| ENSMUSG0000005594 | chr2 | Emc7 | 0.04874028 | 0.918551113 |
| ENSMUSG0000003792 | chr11 | Ssh2 | 0.060279667 | 0.918551113 |
| ENSMUSG0000004596 | chr6 | Wnk1 | 0.046280618 | 0.918551113 |
| ENSMUSG000000264C | chr1 | Ddx59 | 0.052708005 | 0.918551113 |
| ENSMUSG000000252C | chr19 | Twnk | 0.055702717 | 0.918551113 |
| ENSMUSG0000002494 | chr19 | Smc5 | 0.032287638 | 0.918553649 |
| ENSMUSG0000002789 | chr3 | Gstm4 | 0.047661476 | 0.918553649 |
| ENSMUSG0000003044 | chr13 | Zfp273 | -0.072570287 | 0.91884526 |
| ENSMUSG0000002671 | chr1 | Serpinc1 | 0.238181995 | 0.91884526 |

| | | | | |
|---|---|---|---|---|
| ENSMUSG0000002137 | chr13 | Nup153 | 0.031697577 | 0.91884526 |
| ENSMUSG0000003609 | chr9 | Hyal3 | 0.145805433 | 0.91884526 |
| ENSMUSG0000003026 | chr6 | Bcat1 | 0.116462388 | 0.91884526 |
| ENSMUSG0000003343 | chr5 | Gtpbp6 | 0.064531585 | 0.91884526 |
| ENSMUSG0000003789 | chr5 | Wdr19 | -0.053528209 | 0.918859784 |
| ENSMUSG0000002933 | chr5 | Prkg2 | -0.08750764 | 0.91906024 |
| ENSMUSG0000008700 | chr2 | Gm13889 | 0.093004227 | 0.919133441 |
| ENSMUSG0000007905 | chr2 | Vinac1 | -0.246751274 | 0.919159936 |
| ENSMUSG0000003318 | chr18 | Tmed7 | -0.02943727 | 0.919159936 |
| ENSMUSG0000002226 | chr15 | Trio | 0.058049852 | 0.919159936 |
| ENSMUSG0000002686 | chr2 | Psmd5 | 0.035065897 | 0.919171514 |
| ENSMUSG0000002188 | chr14 | Rnase6 | -0.202248273 | 0.919207134 |
| ENSMUSG0000007886 | chr2 | Gm14418 | -0.122299828 | 0.919207134 |
| ENSMUSG0000003796 | chr16 | Zc3h7a | -0.03371315 | 0.919207134 |
| ENSMUSG0000000860 | chr3 | Ubqln4 | -0.03880562 | 0.919207134 |
| ENSMUSG0000002225 | chr15 | Nadk2 | -0.054740385 | 0.919230221 |
| ENSMUSG0000003195 | chr8 | Ldhd | -0.053739027 | 0.919230221 |
| ENSMUSG0000003655 | chr5 | Sh3tc1 | -0.082984494 | 0.919230221 |
| ENSMUSG0000002790 | chr3 | Ddx20 | 0.055961287 | 0.919230221 |
| ENSMUSG0000002129 | chr12 | Kif26a | 0.108229052 | 0.919230221 |
| ENSMUSG0000003247 | chr9 | Nme6 | -0.079945627 | 0.919375312 |
| ENSMUSG0000005566 | chr17 | Mettl4 | -0.046040824 | 0.919375312 |
| ENSMUSG0000002678 | chr2 | Acbd5 | -0.028879901 | 0.919517142 |
| ENSMUSG0000003707 | chr15 | Rnf139 | 0.040824306 | 0.919517142 |
| ENSMUSG0000000372 | chr1 | Insig2 | 0.039140194 | 0.919517142 |
| ENSMUSG0000003938 | chrX | Wdr45 | -0.07126308 | 0.919632146 |
| ENSMUSG0000003115 | chrX | Pim2 | -0.052469841 | 0.919632146 |
| ENSMUSG0000002193 | chr14 | Mapk8 | -0.04385917 | 0.919632146 |
| ENSMUSG0000004177 | chr2 | Cir1 | 0.071319517 | 0.919632146 |
| ENSMUSG0000005197 | chr17 | Prdm9 | -0.132422216 | 0.919670096 |
| ENSMUSG0000005942 | chr4 | Zfp933 | -0.047787062 | 0.919909533 |
| ENSMUSG0000004853 | chr9 | Phldb1 | 0.051387843 | 0.919909533 |
| ENSMUSG0000002030 | chr11 | Cpeb4 | -0.0463307 | 0.92003653 |
| ENSMUSG0000002939 | chr5 | Rchy1 | 0.051358845 | 0.92003653 |
| ENSMUSG0000003069 | chr7 | Ppp4c | 0.059040889 | 0.92003653 |
| ENSMUSG0000007306 | chrX | Zxdb | -0.077663726 | 0.920044662 |
| ENSMUSG0000004805 | chr2 | Ldlrad3 | 0.044598863 | 0.920044662 |
| ENSMUSG0000003222 | chr9 | Fam81a | -0.156976422 | 0.920078876 |
| ENSMUSG0000005130 | chr5 | Usp42 | -0.053298308 | 0.920078876 |
| ENSMUSG0000004301 | chr1 | Nemp2 | -0.058107254 | 0.920078876 |
| ENSMUSG0000007890 | chr8 | Mon1b | -0.046069484 | 0.920078876 |
| ENSMUSG0000003225 | chr9 | Glce | -0.042354887 | 0.920078876 |
| ENSMUSG0000003876 | chr17 | Abcf1 | 0.036943322 | 0.920078876 |
| ENSMUSG0000004283 | chr7 | Alkbh6 | 0.055470386 | 0.920078876 |
| ENSMUSG0000006765 | chr1 | Ankrd23 | 0.112738364 | 0.920078876 |
| ENSMUSG0000003478 | chr2 | B3galt1 | 0.078776099 | 0.920078876 |
| ENSMUSG0000004038 | chr4 | Klhl32 | -0.060545455 | 0.920098929 |
| ENSMUSG0000005103 | chr5 | Zfp11 | -0.049489412 | 0.920098929 |
| ENSMUSG0000002926 | chr5 | Dr1 | -0.033830188 | 0.920098929 |
| ENSMUSG0000002648 | chr1 | Coq8a | 0.134557626 | 0.920098929 |

| | | | | |
|---|---|---|---|---|
| ENSMUSG0000002613 | chr1 | Zfp142 | 0.064545336 | 0.920098929 |
| ENSMUSG0000002403 | chr17 | Abcg1 | 0.056890819 | 0.920098929 |
| ENSMUSG0000004156 | chr7 | Nop53 | 0.064661376 | 0.920098929 |
| ENSMUSG0000002391 | chr17 | Adgrf4 | 0.193346274 | 0.920098929 |
| ENSMUSG0000005586 | chr3 | Tafa3 | -0.185118298 | 0.920198285 |
| ENSMUSG0000002218 | chr14 | Acin1 | 0.030815762 | 0.920198285 |
| ENSMUSG0000002658 | chr1 | Selp | 0.206700729 | 0.920198285 |
| ENSMUSG0000002391 | chr17 | Pla2g7 | -0.076547249 | 0.920376339 |
| ENSMUSG0000004074 | chr3 | Cd53 | 0.056999108 | 0.920410706 |
| ENSMUSG0000003212 | chr9 | Robo4 | -0.080441018 | 0.920552824 |
| ENSMUSG0000003788 | chr2 | Stk35 | 0.059933445 | 0.920552824 |
| ENSMUSG0000006260 | chr5 | Srpk2 | 0.041950839 | 0.92065492 |
| ENSMUSG0000000477 | chr9 | Rab11a | 0.043627121 | 0.92065492 |
| ENSMUSG0000001816 | chr11 | Stard3 | 0.064487806 | 0.92065492 |
| ENSMUSG0000008031 | chr17 | Spaca6 | -0.069143593 | 0.920758172 |
| ENSMUSG0000004331 | chr18 | Pcdhb19 | -0.090560438 | 0.920758172 |
| ENSMUSG0000000276 | chr17 | Pex6 | -0.052741736 | 0.920758172 |
| ENSMUSG0000007858 | chr4 | Ccdc24 | 0.156127905 | 0.920758172 |
| ENSMUSG0000005736 | chr9 | Birc2 | 0.045403148 | 0.920758172 |
| ENSMUSG0000002887 | chr4 | Fam76a | 0.063109563 | 0.920758172 |
| ENSMUSG0000002128 | chr12 | Exoc3l4 | -0.099991615 | 0.920818936 |
| ENSMUSG0000002889 | chr4 | Rcc1 | 0.072786431 | 0.920818936 |
| ENSMUSG0000006068 | chrX | Slc9a6 | -0.040497109 | 0.920907561 |
| ENSMUSG0000000989 | chr11 | Snap47 | 0.060327314 | 0.92095378 |
| ENSMUSG0000002437 | chr18 | Stard4 | -0.104132342 | 0.921035395 |
| ENSMUSG0000003121 | chrX | Stard8 | -0.062985049 | 0.921035395 |
| ENSMUSG0000004848 | chr2 | Bdnf | -0.175231988 | 0.921035395 |
| ENSMUSG0000006197 | chr5 | Rcc1l | -0.06682104 | 0.921035395 |
| ENSMUSG0000005428 | chr5 | Prr14l | -0.057622438 | 0.921035395 |
| ENSMUSG0000002688 | chr2 | Stom | -0.049238072 | 0.921035395 |
| ENSMUSG0000002617 | chr1 | Pnkd | -0.04598581 | 0.921035395 |
| ENSMUSG0000002197 | chr14 | Micu2 | -0.049099644 | 0.921035395 |
| ENSMUSG0000002449 | chr18 | Rbm27 | -0.043156783 | 0.921035395 |
| ENSMUSG0000002481 | chr19 | Tnks2 | -0.038894936 | 0.921035395 |
| ENSMUSG0000002965 | chr5 | Hsph1 | 0.174799235 | 0.921035395 |
| ENSMUSG0000008971 | chr15 | Cbx6 | 0.032111723 | 0.921035395 |
| ENSMUSG0000003947 | chr5 | Tnrc18 | 0.055723816 | 0.921035395 |
| ENSMUSG0000002661 | chr1 | Iars2 | 0.033883651 | 0.921035395 |
| ENSMUSG0000002880 | chr4 | Srrm1 | 0.055313726 | 0.921035395 |
| ENSMUSG0000002449 | chr18 | Tcerg1 | 0.050667044 | 0.921035395 |
| ENSMUSG0000004240 | chr3 | Dennd4b | 0.067359345 | 0.921035395 |
| ENSMUSG0000003500 | chr7 | Igsf6 | 0.097866811 | 0.921035395 |
| ENSMUSG0000002636 | chr1 | Cdc73 | 0.043016596 | 0.921035395 |
| ENSMUSG0000003127 | chrX | Acsl4 | 0.05538764 | 0.921035395 |
| ENSMUSG0000003233 | chr9 | Stoml1 | 0.045930972 | 0.921035395 |
| ENSMUSG0000003008 | chr6 | Abtb1 | 0.064534179 | 0.921035395 |
| ENSMUSG0000011781 | chr17 | Gm50253 | 0.185727738 | 0.921035395 |
| ENSMUSG0000002099 | chr12 | Pnn | -0.052819863 | 0.921177896 |
| ENSMUSG0000003395 | chr11 | Ppp3r1 | -0.031215335 | 0.921177896 |
| ENSMUSG0000005971 | chr17 | Flot1 | 0.032428769 | 0.921276697 |

| | | | | |
|---|---|---|---|---|
| ENSMUSG0000000056 | chr5 | Hnrnpd | -0.036313835 | 0.921307265 |
| ENSMUSG0000003539 | chr2 | Dennd1a | 0.040690574 | 0.921307265 |
| ENSMUSG0000004742 | chr19 | AI837181 | 0.041336065 | 0.921307265 |
| ENSMUSG0000003524 | chr12 | Hectd1 | 0.045280821 | 0.921307265 |
| ENSMUSG0000004060 | chr4 | Kazn | 0.068504503 | 0.921307265 |
| ENSMUSG0000003502 | chr12 | Baz1a | 0.071773797 | 0.921307265 |
| ENSMUSG0000001741 | chr11 | Plxdc1 | -0.06727981 | 0.921402636 |
| ENSMUSG0000004775 | chr11 | Hs3st3a1 | -0.125593697 | 0.921402636 |
| ENSMUSG0000005089 | chr4 | Pdik1l | -0.054953448 | 0.921402636 |
| ENSMUSG0000003194 | chr8 | Adat1 | 0.070774374 | 0.921402636 |
| ENSMUSG0000005351 | chr11 | Kcnip1 | 0.060598658 | 0.921602184 |
| ENSMUSG0000002250 | chr16 | Hapstr1 | -0.03429242 | 0.921627061 |
| ENSMUSG0000003684 | chr7 | Lin37 | -0.075212884 | 0.921632136 |
| ENSMUSG0000002841 | chr4 | Dnaja1 | -0.055588237 | 0.921632136 |
| ENSMUSG0000003079 | chr7 | Fus | -0.054741166 | 0.921632136 |
| ENSMUSG0000002653 | chr1 | Ifi211 | -0.080024933 | 0.921632136 |
| ENSMUSG0000001996 | chr10 | Tmpo | -0.028999098 | 0.921632136 |
| ENSMUSG0000003684 | chr8 | Siah1a | 0.039950976 | 0.921632136 |
| ENSMUSG0000003321 | chr6 | Eefsec | 0.047260393 | 0.921632136 |
| ENSMUSG0000001998 | chr10 | Nedd1 | 0.053454088 | 0.921632136 |
| ENSMUSG0000009509 | chr19 | Ccdc85b | 0.068752063 | 0.921632136 |
| ENSMUSG0000002259 | chr15 | Lynx1 | 0.039451418 | 0.921679836 |
| ENSMUSG0000003817 | chr4 | Ttc39b | -0.067089012 | 0.921718769 |
| ENSMUSG0000002375 | chr15 | Rhebl1 | -0.092383589 | 0.921718769 |
| ENSMUSG0000002602 | chr1 | Als2 | 0.049219713 | 0.921718769 |
| ENSMUSG0000002413 | chr17 | Dnase1l2 | -0.104023142 | 0.921895204 |
| ENSMUSG0000002409 | chr17 | Srsf7 | -0.044911579 | 0.921895204 |
| ENSMUSG0000003073 | chr7 | Slco2b1 | -0.062854255 | 0.921895204 |
| ENSMUSG0000007264 | chr5 | Adam1a | -0.122015279 | 0.922048441 |
| ENSMUSG0000003077 | chr7 | Rbbp6 | -0.05310747 | 0.922048441 |
| ENSMUSG0000000504 | chr11 | Sgsh | -0.047322881 | 0.922048441 |
| ENSMUSG0000003962 | chr6 | Strip2 | 0.10142344 | 0.922048441 |
| ENSMUSG0000003088 | chr7 | Rrp8 | 0.055358522 | 0.922048441 |
| ENSMUSG0000005387 | chr3 | Fpgt | 0.053118449 | 0.922048441 |
| ENSMUSG0000006989 | chr11 | Sft2d1rt | 0.19141957 | 0.922048441 |
| ENSMUSG0000002030 | chr11 | Chac2 | 0.086581202 | 0.922048441 |
| ENSMUSG0000003749 | chr8 | Zmat4 | 0.350932249 | 0.922048441 |
| ENSMUSG0000002587 | chr13 | Faf2 | 0.027018127 | 0.9220656 |
| ENSMUSG0000001315 | chr8 | Enkd1 | 0.068083844 | 0.9220656 |
| ENSMUSG0000005155 | chr7 | Zfp579 | 0.079530887 | 0.9220656 |
| ENSMUSG0000002170 | chr13 | Erbin | -0.047419513 | 0.922090844 |
| ENSMUSG0000000493 | chr10 | Nmrk2 | 0.131138361 | 0.922107351 |
| ENSMUSG0000009739 | chr5 | Thoc2l | 0.059499095 | 0.922162579 |
| ENSMUSG0000004907 | chr16 | Acap2 | 0.042071342 | 0.922243831 |
| ENSMUSG0000004066 | chr9 | Rad54l2 | -0.045595739 | 0.922261536 |
| ENSMUSG0000005407 | chr11 | Utp18 | 0.042719479 | 0.922261536 |
| ENSMUSG0000002899 | chr4 | Lzic | 0.04933362 | 0.922261536 |
| ENSMUSG0000001548 | chr2 | Cacfd1 | -0.03884839 | 0.92239005 |
| ENSMUSG0000005230 | chr10 | Tbc1d30 | -0.075310545 | 0.922593509 |
| ENSMUSG0000003765 | chr3 | Phc3 | 0.044082719 | 0.922593509 |

| | | | | |
|---|---|---|---|---|
| ENSMUSG0000002769 | chr3 | Tnik | 0.059791237 | 0.922593509 |
| ENSMUSG0000002636 | chr1 | Cfh | 0.110894729 | 0.922593509 |
| ENSMUSG0000001921 | chr17 | Chtf18 | -0.168022972 | 0.922634089 |
| ENSMUSG0000002858 | chr4 | Orc1 | -0.139888443 | 0.922634089 |
| ENSMUSG0000002010 | chr10 | Slc29a3 | -0.051214131 | 0.922634089 |
| ENSMUSG0000003101 | chrX | Cask | -0.040766749 | 0.922634089 |
| ENSMUSG0000003810 | chr8 | Trappc11 | -0.031484221 | 0.922634089 |
| ENSMUSG0000001983 | chr10 | Wasf1 | 0.043340854 | 0.922634089 |
| ENSMUSG0000001634 | chr2 | Ppdpf | 0.067781868 | 0.922634089 |
| ENSMUSG0000005203 | chr17 | Tagap1 | 0.064413284 | 0.922634089 |
| ENSMUSG0000002521 | chr19 | Sfxn3 | 0.055173769 | 0.922634089 |
| ENSMUSG0000004139 | chr4 | 1700013G24Rik | 0.243292804 | 0.922634089 |
| ENSMUSG0000002483 | chr19 | Ighmbp2 | 0.048703458 | 0.922646603 |
| ENSMUSG0000005948 | chr4 | Cfap418 | 0.042274764 | 0.922646603 |
| ENSMUSG0000005798 | chr9 | Zfp809 | 0.056944922 | 0.922646603 |
| ENSMUSG0000004242 | chr7 | Fbrs | -0.052346552 | 0.922862785 |
| ENSMUSG0000002560 | chr6 | Copg2 | -0.032080752 | 0.922862785 |
| ENSMUSG0000004310 | chr2 | Qrfp | 0.270331583 | 0.922862785 |
| ENSMUSG0000002734 | chr2 | Fam98b | 0.02918128 | 0.922862785 |
| ENSMUSG0000003769 | chr14 | Ddhd1 | -0.059767185 | 0.922870375 |
| ENSMUSG0000001373 | chr6 | Trnt1 | 0.036075301 | 0.922870375 |
| ENSMUSG0000007037 | chr7 | Prss36 | -0.067633308 | 0.922955045 |
| ENSMUSG0000003179 | chr8 | Cfap20 | -0.039417874 | 0.923046132 |
| ENSMUSG0000003405 | chr10 | Myrfl | 0.274158439 | 0.923046132 |
| ENSMUSG0000005024 | chr13 | Heatr1 | -0.051469111 | 0.923152215 |
| ENSMUSG0000004565 | chr1 | Pid1 | -0.05475933 | 0.923177754 |
| ENSMUSG0000002687 | chr2 | Hc | -0.30349818 | 0.92333158 |
| ENSMUSG0000008999 | chr6 | Cfap92 | -0.254322276 | 0.92333158 |
| ENSMUSG0000007849 | chr4 | Cfap74 | -0.099932863 | 0.92333158 |
| ENSMUSG0000002774 | chr3 | Trpc4 | -0.049631985 | 0.92333158 |
| ENSMUSG0000009521 | chr13 | H2bc15 | -0.20181866 | 0.92333158 |
| ENSMUSG0000003594 | chr9 | Ibtk | -0.040001102 | 0.92333158 |
| ENSMUSG0000005483 | chr9 | Elp6 | -0.085395487 | 0.92333158 |
| ENSMUSG0000000903 | chr15 | Tmem184b | -0.038303919 | 0.92333158 |
| ENSMUSG0000003246 | chr9 | Armc8 | -0.033004978 | 0.92333158 |
| ENSMUSG0000004781 | chr19 | Ccdc88b | -0.122774559 | 0.92333158 |
| ENSMUSG0000004097 | chr4 | Igsf21 | 0.115857276 | 0.92333158 |
| ENSMUSG0000003418 | chr8 | Hsdl1 | 0.03919275 | 0.92333158 |
| ENSMUSG0000002839 | chr4 | Pole3 | 0.045507855 | 0.92333158 |
| ENSMUSG0000003825 | chr2 | Gdf5 | 0.12492775 | 0.92333158 |
| ENSMUSG0000003528 | chr7 | Nat14 | 0.070646367 | 0.92333158 |
| ENSMUSG0000006833 | chr6 | Dok1 | 0.078225981 | 0.92333158 |
| ENSMUSG0000004391 | chr5 | Ccdc60 | 0.083919339 | 0.92333158 |
| ENSMUSG0000006158 | chr10 | Dot1l | 0.086832327 | 0.923354837 |
| ENSMUSG0000003749 | chr9 | Cib2 | 0.057409618 | 0.923479753 |
| ENSMUSG0000006927 | chr13 | H3c6 | 0.22945734 | 0.923691547 |
| ENSMUSG0000002151 | chr13 | Zfp369 | 0.051309543 | 0.923691547 |
| ENSMUSG0000006414 | chr4 | Zfp69 | -0.169140918 | 0.923692866 |
| ENSMUSG0000002959 | chr5 | Usp30 | -0.049627486 | 0.923692866 |
| ENSMUSG0000003691 | chr9 | Piwil4 | -0.265166832 | 0.923692866 |

| | | | | |
|---|---|---|---|---|
| ENSMUSG0000003448 | chrX | Diaph2 | 0.051106714 | 0.923692866 |
| ENSMUSG0000003613 | chr9 | Acaa1a | 0.051086609 | 0.923692866 |
| ENSMUSG0000007164 | chr19 | B3gat3 | 0.053840797 | 0.923692866 |
| ENSMUSG0000003447 | chr11 | Caskin2 | 0.050045298 | 0.923692866 |
| ENSMUSG0000002077 | chr11 | Ten1 | 0.07053072 | 0.923692866 |
| ENSMUSG0000002310 | chr5 | Rfc2 | 0.047840335 | 0.923742128 |
| ENSMUSG0000003307 | chr15 | Senp1 | 0.040282591 | 0.923766976 |
| ENSMUSG0000003262 | chr17 | Eml4 | 0.053880483 | 0.923998371 |
| ENSMUSG0000002788 | chr3 | Prpf38b | -0.042757046 | 0.924019554 |
| ENSMUSG0000003958 | chr9 | Myo9a | -0.056772307 | 0.924019554 |
| ENSMUSG0000002593 | chr1 | Ube2w | -0.041959491 | 0.924019554 |
| ENSMUSG0000003195 | chr8 | Gabarapl2 | 0.057233586 | 0.924019554 |
| ENSMUSG0000000240 | chr7 | Dyrk1b | 0.081313805 | 0.924190787 |
| ENSMUSG0000008984 | chr1 | Sp140l1 | -0.127065471 | 0.924272024 |
| ENSMUSG0000003867 | chr15 | Trps1 | 0.066103579 | 0.924292972 |
| ENSMUSG0000000115 | chr10 | Pcnt | 0.082756477 | 0.924442125 |
| ENSMUSG0000006145 | chr4 | Stx17 | 0.049097799 | 0.924532533 |
| ENSMUSG0000004922 | chr4 | Pdp1 | -0.052179734 | 0.924980384 |
| ENSMUSG0000002028 | chr11 | Rhbdf1 | -0.041380632 | 0.924980384 |
| ENSMUSG0000003801 | chr5 | Dnah10 | -0.146972911 | 0.925126988 |
| ENSMUSG0000005007 | chr9 | Spink8 | -0.227480392 | 0.925126988 |
| ENSMUSG0000004294 | chr18 | Greb1l | -0.071866894 | 0.925126988 |
| ENSMUSG0000003186 | chr8 | Lpar2 | -0.109493448 | 0.925126988 |
| ENSMUSG0000002682 | chr2 | Ptges2 | -0.058001016 | 0.925126988 |
| ENSMUSG0000003295 | chr3 | Ap4b1 | -0.117638271 | 0.925126988 |
| ENSMUSG0000003452 | chr13 | Ice1 | -0.047463294 | 0.925126988 |
| ENSMUSG0000001493 | chr5 | Yes1 | -0.04682087 | 0.925126988 |
| ENSMUSG0000004155 | chr4 | Fbxo2 | -0.070033605 | 0.925126988 |
| ENSMUSG0000005007 | chr8 | Rspry1 | -0.038843614 | 0.925126988 |
| ENSMUSG0000001654 | chr15 | Atxn10 | -0.035868707 | 0.925126988 |
| ENSMUSG0000002431 | chr18 | Rnf138 | -0.034932554 | 0.925126988 |
| ENSMUSG0000002445 | chr18 | Diaph1 | -0.032550531 | 0.925126988 |
| ENSMUSG0000001481 | chr14 | Stc1 | 0.109605539 | 0.925126988 |
| ENSMUSG0000002891 | chr4 | Srsf4 | 0.034473164 | 0.925126988 |
| ENSMUSG0000005400 | chr18 | Ndst1 | 0.047041386 | 0.925126988 |
| ENSMUSG0000002302 | chr15 | Larp4 | 0.045476346 | 0.925126988 |
| ENSMUSG0000002588 | chr9 | Casp12 | 0.054339745 | 0.925126988 |
| ENSMUSG0000004282 | chr2 | Snai1 | 0.075305382 | 0.925126988 |
| ENSMUSG0000004106 | chr9 | Pif1 | 0.345322846 | 0.925126988 |
| ENSMUSG0000002126 | chr12 | Slc25a29 | 0.092996654 | 0.925126988 |
| ENSMUSG0000011613 | chr15 | C030006K11Rik | 0.062275048 | 0.925126988 |
| ENSMUSG0000002940 | chr5 | Snrnp35 | 0.063537894 | 0.9253379 |
| ENSMUSG0000005263 | chr12 | Asap2 | 0.061574033 | 0.9253379 |
| ENSMUSG0000009525 | chr17 | Zfp799 | 0.044743332 | 0.925443186 |
| ENSMUSG0000002251 | chr16 | Anks3 | 0.029152917 | 0.925703043 |
| ENSMUSG0000000560 | chr7 | Ctr9 | 0.033760825 | 0.925943392 |
| ENSMUSG0000003024 | chr6 | Recql | -0.046134262 | 0.925948626 |
| ENSMUSG0000003970 | chr5 | Ldb2 | 0.047466662 | 0.925948626 |
| ENSMUSG0000006920 | chr13 | Zfp874a | 0.058513161 | 0.925948626 |
| ENSMUSG0000003967 | chr11 | Oxld1 | 0.09796176 | 0.926019693 |

| | | | | |
|---|---|---|---|---|
| ENSMUSG0000005358 | chr11 | Tanc2 | -0.064985614 | 0.926132763 |
| ENSMUSG0000005279 | chr10 | Nup107 | -0.040793575 | 0.926132763 |
| ENSMUSG0000004909 | chr14 | Gpr137c | -0.102473791 | 0.926235006 |
| ENSMUSG0000002596 | chr1 | Adam23 | -0.062186987 | 0.926562718 |
| ENSMUSG0000002436 | chr17 | Nelfe | -0.04727784 | 0.926562718 |
| ENSMUSG0000003337 | chr12 | Fntb | -0.048225512 | 0.92658476 |
| ENSMUSG0000005291 | chr11 | Msl1 | -0.029525947 | 0.92658476 |
| ENSMUSG0000002941 | chr5 | Ppef2 | 0.131951492 | 0.92658476 |
| ENSMUSG0000001979 | chr10 | Hint3 | 0.062558437 | 0.92658476 |
| ENSMUSG0000005447 | chr18 | Kcnn2 | 0.068216076 | 0.92658476 |
| ENSMUSG0000000281 | chr11 | Top3a | 0.064716173 | 0.92658476 |
| ENSMUSG0000003768 | chr2 | Armc3 | -0.190305808 | 0.926592916 |
| ENSMUSG0000004440 | chr12 | Sptssa | 0.057113774 | 0.926592916 |
| ENSMUSG0000005549 | chr10 | Epm2a | 0.064891031 | 0.926931587 |
| ENSMUSG0000003830 | chr10 | Afg1l | -0.05769487 | 0.926963512 |
| ENSMUSG0000005874 | chr8 | Zfp958 | -0.05072357 | 0.926963512 |
| ENSMUSG0000004012 | chr1 | Gorab | -0.053485968 | 0.926963512 |
| ENSMUSG0000006360 | chr3 | Egfem1 | 0.11414921 | 0.926963512 |
| ENSMUSG0000003023 | chr6 | Slco1c1 | 0.166261649 | 0.926963512 |
| ENSMUSG0000004763 | chr19 | Fgfbp3 | 0.134832408 | 0.926963512 |
| ENSMUSG0000002657 | chr1 | Nme7 | -0.06004076 | 0.927066972 |
| ENSMUSG0000002193 | chr14 | Rnaseh2b | -0.091649058 | 0.92708022 |
| ENSMUSG0000004655 | chrX | Spin2c | -0.098425814 | 0.92708022 |
| ENSMUSG0000007147 | chr6 | Zfp777 | 0.041363023 | 0.92708022 |
| ENSMUSG0000007042 | chr7 | Rnf121 | 0.046223386 | 0.92708022 |
| ENSMUSG0000003298 | chr15 | Slc16a8 | 0.146201821 | 0.92708022 |
| ENSMUSG0000004436 | chr7 | Ccdc89 | -0.154262024 | 0.927119259 |
| ENSMUSG0000004127 | chr4 | Tox | -0.06126416 | 0.927240989 |
| ENSMUSG0000003242 | chr9 | Zfp949 | -0.03956381 | 0.927240989 |
| ENSMUSG0000000531 | chr13 | Ubqln1 | -0.030263551 | 0.927240989 |
| ENSMUSG0000002085 | chr11 | Spag9 | 0.040131963 | 0.927240989 |
| ENSMUSG0000003274 | chr11 | Ccdc88a | 0.052974614 | 0.927240989 |
| ENSMUSG0000005571 | chr14 | Slain1 | 0.052086836 | 0.92747019 |
| ENSMUSG0000002279 | chr16 | Tfrc | -0.080760518 | 0.927517155 |
| ENSMUSG0000000974 | chr9 | Ubp1 | -0.045319393 | 0.927551425 |
| ENSMUSG0000004534 | chr4 | Sh2d5 | -0.141764879 | 0.927551425 |
| ENSMUSG0000002710 | chr2 | Atf2 | -0.038333025 | 0.927551425 |
| ENSMUSG0000002100 | chr12 | Mia2 | -0.046268806 | 0.927551425 |
| ENSMUSG0000004056 | chr9 | Plppr2 | -0.042760835 | 0.927551425 |
| ENSMUSG0000002413 | chr17 | E4f1 | -0.035873628 | 0.927551425 |
| ENSMUSG0000003074 | chr7 | Lat | 0.22700833 | 0.927551425 |
| ENSMUSG0000002419 | chr17 | Plin3 | 0.056951657 | 0.927551425 |
| ENSMUSG0000003551 | chr4 | Tdrd7 | 0.041952615 | 0.927551425 |
| ENSMUSG0000003851 | chr13 | Jarid2 | 0.058875873 | 0.927551425 |
| ENSMUSG0000005427 | chr15 | Arfgap3 | 0.046247002 | 0.927551425 |
| ENSMUSG0000001947 | chr8 | Rab4a | 0.044655651 | 0.927551425 |
| ENSMUSG0000002250 | chr16 | Nubp1 | 0.058265409 | 0.927551425 |
| ENSMUSG0000003860 | chr6 | Atp6v0a4 | -0.28385214 | 0.927688908 |
| ENSMUSG0000003733 | chr7 | Map4k1 | -0.150398422 | 0.927688908 |
| ENSMUSG0000007503 | chr13 | H2bc3 | -0.243491515 | 0.927688908 |

| | | | | |
|---|---|---|---|---|
| ENSMUSG0000004821 | chr15 | Amigo2 | -0.056155902 | 0.927688908 |
| ENSMUSG0000005441 | chr3 | Slc30a7 | -0.035076531 | 0.927688908 |
| ENSMUSG0000002002 | chr10 | Nudt4 | -0.031883447 | 0.927688908 |
| ENSMUSG0000004383 | chr6 | Clec4a3 | 0.098342457 | 0.927688908 |
| ENSMUSG0000003144 | chr8 | Atp11a | 0.035789262 | 0.927688908 |
| ENSMUSG0000002813 | chr3 | Riiad1 | 0.085630429 | 0.927904251 |
| ENSMUSG0000002172 | chr13 | Emb | 0.049835465 | 0.92809683 |
| ENSMUSG0000000782 | chr6 | Ankrd26 | 0.042872419 | 0.928187702 |
| ENSMUSG0000005195 | chr1 | Xkr4 | -0.097111186 | 0.928193781 |
| ENSMUSG0000005897 | chr7 | Kcnc1 | -0.061880669 | 0.928193781 |
| ENSMUSG0000003829 | chr11 | Smg6 | -0.047738862 | 0.928193781 |
| ENSMUSG0000004406 | chr11 | Cep68 | -0.041665278 | 0.928193781 |
| ENSMUSG0000004251 | chrX | Pwwp3b | 0.095737987 | 0.928193781 |
| ENSMUSG0000004822 | chr2 | Mfap1b | 0.04689604 | 0.928193781 |
| ENSMUSG0000002701 | chr2 | Hat1 | 0.050658898 | 0.928193781 |
| ENSMUSG0000004018 | chr6 | Bmal2 | 0.058577176 | 0.928193781 |
| ENSMUSG0000002781 | chr3 | 4930579G24Rik | 0.141612013 | 0.928228461 |
| ENSMUSG0000005879 | chr2 | Cds2 | -0.047139613 | 0.928495246 |
| ENSMUSG0000000651 | chr8 | Mvd | 0.102148111 | 0.928529352 |
| ENSMUSG0000001988 | chr10 | Echdc1 | -0.075007542 | 0.928693007 |
| ENSMUSG0000007643 | chr2 | Selenoh | -0.07060375 | 0.928693007 |
| ENSMUSG0000003784 | chr9 | Fdxacb1 | -0.064831913 | 0.928693007 |
| ENSMUSG0000002441 | chr18 | Rmc1 | -0.04673542 | 0.928693007 |
| ENSMUSG0000005734 | chr7 | Sphk2 | -0.046798322 | 0.928693007 |
| ENSMUSG0000003741 | chr18 | Dmxl1 | -0.047255766 | 0.928693007 |
| ENSMUSG0000007474 | chr19 | Pdzd8 | -0.037374397 | 0.928693007 |
| ENSMUSG0000003995 | chr4 | Clstn1 | -0.033996589 | 0.928693007 |
| ENSMUSG0000002271 | chr16 | Usp7 | -0.027985657 | 0.928693007 |
| ENSMUSG0000002222 | chr14 | Ripk3 | 0.143314246 | 0.928693007 |
| ENSMUSG0000006189 | chr4 | Zscan20 | 0.049620556 | 0.928693007 |
| ENSMUSG0000002776 | chr3 | Mbnl1 | 0.04057328 | 0.928693007 |
| ENSMUSG0000005548 | chr2 | Mtcl2 | 0.064164144 | 0.928693007 |
| ENSMUSG0000007835 | chr4 | Smim1 | 0.072822558 | 0.928693007 |
| ENSMUSG0000004167 | chr8 | Fbxl9 | -0.126545826 | 0.928768788 |
| ENSMUSG0000002742 | chr2 | Sec23b | -0.035895859 | 0.928808112 |
| ENSMUSG0000003489 | chr14 | Cog3 | -0.02838862 | 0.928928196 |
| ENSMUSG0000003627 | chr8 | Edc4 | 0.034269291 | 0.928928196 |
| ENSMUSG0000003855 | chr3 | Ciart | -0.431381001 | 0.928992668 |
| ENSMUSG0000005585 | chr6 | Rnf181 | -0.036682786 | 0.929086771 |
| ENSMUSG0000006114 | chr3 | Maml3 | -0.051667298 | 0.929153379 |
| ENSMUSG0000000319 | chr17 | Zfp959 | -0.062472702 | 0.929153379 |
| ENSMUSG0000002746 | chr2 | Rbck1 | 0.029022906 | 0.929153379 |
| ENSMUSG0000011853 | chr13 | Shld3 | 0.122765257 | 0.929153379 |
| ENSMUSG0000002901 | chr5 | Dnajc2 | 0.040154471 | 0.929153379 |
| ENSMUSG0000005613 | chr18 | Ticam2 | 0.152570799 | 0.929153379 |
| ENSMUSG0000004690 | chr14 | Ltb4r1 | 0.207279734 | 0.929153379 |
| ENSMUSG0000004932 | chr11 | Smcr8 | -0.04315567 | 0.929346641 |
| ENSMUSG0000001833 | chr11 | Ksr1 | 0.039085844 | 0.92942997 |
| ENSMUSG0000004877 | chr4 | Tmem53 | 0.066814233 | 0.929649975 |
| ENSMUSG0000007861 | chr11 | Smarcd2 | -0.057841893 | 0.929667466 |

| | | | | |
|---|---|---|---|---|
| ENSMUSG0000005143 | chr4 | Fhad1 | 0.075371511 | 0.929694492 |
| ENSMUSG0000002190 | chr14 | Galnt15 | -0.092208588 | 0.929712549 |
| ENSMUSG0000002488 | chr19 | Rin1 | -0.120601798 | 0.929712549 |
| ENSMUSG0000002429 | chr17 | Adamts10 | -0.073120898 | 0.929712549 |
| ENSMUSG0000005563 | chr14 | Dach1 | -0.086888028 | 0.929712549 |
| ENSMUSG0000003657 | chrX | Upf3b | -0.066854381 | 0.929712549 |
| ENSMUSG0000002215 | chr15 | Dab2 | -0.060482879 | 0.929712549 |
| ENSMUSG0000001834 | chr11 | Zkscan6 | -0.055854041 | 0.929712549 |
| ENSMUSG0000003526 | chr10 | Amh | -0.193804549 | 0.929712549 |
| ENSMUSG0000003833 | chr11 | Tsr1 | -0.042795875 | 0.929712549 |
| ENSMUSG0000006067 | chr13 | H4c3 | -0.212557571 | 0.929712549 |
| ENSMUSG0000001571 | chr3 | Cers2 | 0.031091461 | 0.929712549 |
| ENSMUSG0000003941 | chr4 | Prdm16 | 0.04818328 | 0.929712549 |
| ENSMUSG0000002649 | chr1 | Ahctf1 | 0.062917507 | 0.929712549 |
| ENSMUSG0000006925 | chr13 | Dusp22 | 0.037057247 | 0.929712549 |
| ENSMUSG0000002517 | chr11 | Rab40b | 0.071008174 | 0.929712549 |
| ENSMUSG0000004088 | chr17 | Gfer | 0.065485772 | 0.929712549 |
| ENSMUSG0000003988 | chr5 | Lrrc17 | 0.136866126 | 0.929768819 |
| ENSMUSG0000003119 | chrX | Vbp1 | 0.043453695 | 0.929768819 |
| ENSMUSG0000002010 | chr10 | Slc16a7 | 0.042081885 | 0.929768819 |
| ENSMUSG0000002036 | chr11 | Zfp354a | 0.072151155 | 0.929768819 |
| ENSMUSG0000007441 | chr7 | Pira12 | 0.231092375 | 0.929844087 |
| ENSMUSG0000006409 | chr11 | Vrk2 | 0.065669651 | 0.929844087 |
| ENSMUSG0000003815 | chr7 | Spon1 | -0.067882742 | 0.929956624 |
| ENSMUSG0000002542 | chr18 | Katnal2 | -0.106557949 | 0.929956624 |
| ENSMUSG0000003777 | chr2 | Pced1a | -0.045003908 | 0.929956624 |
| ENSMUSG0000004550 | chr5 | Hcar2 | -0.221404394 | 0.929956624 |
| ENSMUSG0000003936 | chr14 | Sec24c | 0.023930435 | 0.929956624 |
| ENSMUSG0000011325 | chr13 | Tes3-ps | -0.15708388 | 0.93003731 |
| ENSMUSG0000003136 | chrX | Grpr | -0.134201209 | 0.930049743 |
| ENSMUSG0000004600 | chr13 | Gapt | -0.240586044 | 0.930049743 |
| ENSMUSG0000003567 | chr10 | Sbno2 | -0.055297786 | 0.930049743 |
| ENSMUSG0000003159 | chr8 | Frg1 | -0.044784915 | 0.930049743 |
| ENSMUSG0000002917 | chr5 | Klf3 | -0.041371199 | 0.930049743 |
| ENSMUSG0000001816 | chr11 | Med1 | -0.052119666 | 0.930049743 |
| ENSMUSG0000002680 | chr2 | Ttf1 | 0.040328965 | 0.930049743 |
| ENSMUSG0000007560 | chr15 | Zc3h3 | 0.0517745 | 0.930049743 |
| ENSMUSG0000003722 | chr5 | Mospd3 | 0.052008071 | 0.930049743 |
| ENSMUSG0000003910 | chr13 | F13a1 | 0.094703552 | 0.930049743 |
| ENSMUSG0000001595 | chr5 | Ncf1 | 0.091870136 | 0.930049743 |
| ENSMUSG0000002086 | chr11 | Cacna1g | 0.10420146 | 0.930063389 |
| ENSMUSG0000003848 | chr11 | Socs7 | 0.041094504 | 0.930063389 |
| ENSMUSG0000006009 | chrX | Rp2 | 0.044008647 | 0.930145848 |
| ENSMUSG0000002566 | chrX | Tmem47 | -0.046654719 | 0.930219288 |
| ENSMUSG0000002167 | chr13 | Poc5 | -0.041238021 | 0.930219288 |
| ENSMUSG0000003045 | chr7 | Herc2 | 0.046255183 | 0.930219288 |
| ENSMUSG0000004449 | chr4 | 2510039O18Rik | 0.036699518 | 0.930219288 |
| ENSMUSG0000011656 | chr17 | Riok2 | 0.053532242 | 0.930219288 |
| ENSMUSG0000002023 | chr10 | Dip2a | -0.041450343 | 0.930246052 |
| ENSMUSG0000004160 | chr5 | Bicdl1 | -0.073591284 | 0.930328247 |

| | | | | |
|---|---|---|---|---|
| ENSMUSG0000003020 | chr6 | Lrp6 | 0.040633846 | 0.930328247 |
| ENSMUSG0000002843 | chr4 | Kif24 | 0.10263076 | 0.930328247 |
| ENSMUSG0000007398 | chr4 | Ttpa | 0.094380618 | 0.930328247 |
| ENSMUSG0000002981 | chr6 | Igf2bp3 | 0.116243268 | 0.930328247 |
| ENSMUSG0000003951 | chr17 | Bltp3a | 0.047766522 | 0.93078174 |
| ENSMUSG0000007451 | chr3 | Arfip1 | 0.053688479 | 0.930828241 |
| ENSMUSG0000000641 | chr2 | Rnf114 | -0.038737949 | 0.931061575 |
| ENSMUSG0000009195 | chr7 | Tmsb10b | 0.191473702 | 0.931061575 |
| ENSMUSG0000003006 | chr6 | Hmces | 0.047310767 | 0.931061575 |
| ENSMUSG0000003122 | chrX | Igbp1 | 0.057798703 | 0.931061575 |
| ENSMUSG0000000250 | chr17 | Rpl3l | 0.177102755 | 0.931221753 |
| ENSMUSG0000002168 | chr13 | Pde8b | -0.079297612 | 0.931283471 |
| ENSMUSG0000005402 | chr13 | Sirt5 | -0.052511816 | 0.931283471 |
| ENSMUSG0000005940 | chr17 | Ppp2r5d | -0.033358417 | 0.931283471 |
| ENSMUSG0000001785 | chr2 | Ift52 | -0.032946595 | 0.931283471 |
| ENSMUSG0000003926 | chr2 | Prrc2b | -0.053693 | 0.931285759 |
| ENSMUSG0000003232 | chr9 | Tspan3 | -0.044202695 | 0.931285759 |
| ENSMUSG0000004134 | chr11 | Wrap53 | -0.058948821 | 0.931285759 |
| ENSMUSG0000005071 | chr2 | Zbtb26 | 0.051353194 | 0.931294275 |
| ENSMUSG0000003779 | chr5 | N4bp2 | -0.056426496 | 0.931364208 |
| ENSMUSG0000003318 | chr14 | Mzt1 | -0.037070152 | 0.931364208 |
| ENSMUSG0000003967 | chr2 | Zmynd8 | -0.03077741 | 0.931678789 |
| ENSMUSG0000007356 | chr18 | Prr16 | 0.089748783 | 0.931685953 |
| ENSMUSG0000002191 | chr14 | Glt8d1 | -0.029112081 | 0.931719332 |
| ENSMUSG0000002954 | chr5 | Ints1 | 0.030720598 | 0.931719332 |
| ENSMUSG0000003735 | chr5 | Uvssa | 0.06135158 | 0.931719332 |
| ENSMUSG0000009065 | chr13 | Zfp493 | 0.085394807 | 0.931719332 |
| ENSMUSG0000002448 | chr18 | Ankhd1 | -0.056013815 | 0.931819306 |
| ENSMUSG0000003192 | chr9 | Cep57 | -0.049208548 | 0.931819306 |
| ENSMUSG0000003614 | chr12 | Meox2 | -0.128508221 | 0.931822861 |
| ENSMUSG0000002084 | chr11 | Rflnb | -0.074152766 | 0.931822861 |
| ENSMUSG0000005805 | chr15 | Tmt1a3 | -0.256405299 | 0.931822861 |
| ENSMUSG0000001973 | chr7 | Syne4 | -0.084493903 | 0.931822861 |
| ENSMUSG0000006047 | chr8 | Adgrg3 | -0.066799443 | 0.931822861 |
| ENSMUSG0000000002 | chr16 | Cdc45 | -0.111400594 | 0.931822861 |
| ENSMUSG0000009147 | chr7 | 2610021A01Rik | -0.04687862 | 0.931822861 |
| ENSMUSG0000004737 | chr7 | Zfp768 | -0.042775753 | 0.931822861 |
| ENSMUSG0000006976 | chr11 | Msi2 | -0.043236896 | 0.931822861 |
| ENSMUSG0000004699 | chr3 | 1110032F04Rik | -0.144013627 | 0.931822861 |
| ENSMUSG0000011027 | chr18 | Gm45871 | -0.046133506 | 0.931822861 |
| ENSMUSG0000003610 | chr1 | Rab3gap1 | 0.026666709 | 0.931822861 |
| ENSMUSG0000002118 | chr12 | Ccdc88c | 0.061324406 | 0.931822861 |
| ENSMUSG0000009833 | chr9 | Pigbos1 | 0.113107891 | 0.931822861 |
| ENSMUSG0000002722 | chr2 | Pex16 | 0.050575457 | 0.931822861 |
| ENSMUSG0000002934 | chr5 | Tfip11 | 0.036374868 | 0.931822861 |
| ENSMUSG0000005945 | chr14 | Ptk2b | 0.055819834 | 0.931822861 |
| ENSMUSG0000005583 | chr8 | Zfp1 | 0.043971317 | 0.931822861 |
| ENSMUSG0000003398 | chr11 | Dnah17 | 0.109828053 | 0.931822861 |
| ENSMUSG0000002624 | chr1 | Cops7b | 0.050601594 | 0.931822861 |
| ENSMUSG0000007347 | chr17 | Rsph3a | 0.062221558 | 0.931822861 |

| | | | | |
|---|---|---|---|---|
| ENSMUSG0000007046 | chr7 | Adamtsl3 | 0.077031613 | 0.931822861 |
| ENSMUSG0000007154 | chr14 | Nt5dc2 | 0.084616725 | 0.931822861 |
| ENSMUSG0000006175 | chr10 | Armt1 | 0.044909758 | 0.931865135 |
| ENSMUSG0000002200 | chr14 | Slc25a30 | -0.059855714 | 0.931914615 |
| ENSMUSG0000000909 | chr11 | Ap1b1 | -0.030834754 | 0.931914615 |
| ENSMUSG0000002604 | chr1 | Tex30 | 0.099095571 | 0.931914615 |
| ENSMUSG0000002550 | chr7 | Pnpla2 | 0.058911531 | 0.932086033 |
| ENSMUSG0000004956 | chr3 | Aknad1 | -0.101794648 | 0.93221141 |
| ENSMUSG0000009029 | chr19 | Lrrc10b | -0.080236016 | 0.93221141 |
| ENSMUSG0000002159 | chr13 | Spata9 | -0.126856329 | 0.93221141 |
| ENSMUSG0000006437 | chr15 | Selenop | -0.068101844 | 0.93221141 |
| ENSMUSG0000011834 | chr19 | Tmem179b | -0.053624138 | 0.93221141 |
| ENSMUSG0000006855 | chr6 | Zfp467 | -0.042103092 | 0.93221141 |
| ENSMUSG0000003983 | chr2 | Zfp335 | -0.047939917 | 0.93221141 |
| ENSMUSG0000002958 | chr5 | Fscn1 | -0.054793675 | 0.93221141 |
| ENSMUSG0000003930 | chr11 | Hexd | -0.044538085 | 0.93221141 |
| ENSMUSG0000002105 | chr12 | Sgpp1 | -0.039756473 | 0.93221141 |
| ENSMUSG0000005360 | chr6 | Rpia | -0.051404204 | 0.93221141 |
| ENSMUSG0000002803 | chr3 | Tbck | -0.062733887 | 0.93221141 |
| ENSMUSG0000006063 | chr13 | H4c9 | 0.171699599 | 0.93221141 |
| ENSMUSG0000000214 | chr10 | Stat6 | 0.028153647 | 0.93221141 |
| ENSMUSG0000000637 | chrX | Pgrmc1 | 0.039801806 | 0.93221141 |
| ENSMUSG0000000568 | chr10 | Pan2 | 0.062962642 | 0.93221141 |
| ENSMUSG0000005796 | chr11 | Fgf18 | 0.110601264 | 0.93221141 |
| ENSMUSG0000000456 | chr14 | Mettl17 | 0.03715082 | 0.93221141 |
| ENSMUSG0000002774 | chr3 | Cog6 | 0.034196608 | 0.93221141 |
| ENSMUSG0000003760 | chr11 | Nme1 | 0.063569099 | 0.93221141 |
| ENSMUSG0000002596 | chr1 | Fastkd2 | 0.044629324 | 0.93221141 |
| ENSMUSG0000004995 | chr2 | Rpp38 | 0.059920081 | 0.93221141 |
| ENSMUSG0000005451 | chr11 | Trim65 | 0.055797326 | 0.93221141 |
| ENSMUSG0000004211 | chr4 | Vwa1 | 0.050824539 | 0.93221141 |
| ENSMUSG0000004703 | chr7 | Nipa1 | 0.049897278 | 0.93221141 |
| ENSMUSG0000005611 | chr17 | H2-T22 | 0.041641372 | 0.93221141 |
| ENSMUSG0000005001 | chr5 | Shisa3 | 0.12366685 | 0.93221141 |
| ENSMUSG0000006847 | chr2 | Mfap1a | 0.05233516 | 0.93221141 |
| ENSMUSG0000001987 | chr10 | Pkib | 0.066357328 | 0.93221141 |
| ENSMUSG0000002042 | chr7 | Zfp607a | 0.084298512 | 0.93221141 |
| ENSMUSG0000004687 | chr13 | Atxn1 | 0.066978242 | 0.93221141 |
| ENSMUSG0000007893 | chr15 | Cpt1b | 0.24921317 | 0.93221141 |
| ENSMUSG0000007171 | chr15 | Csf2rb2 | -0.143031874 | 0.932211856 |
| ENSMUSG0000002203 | chr14 | Clu | -0.064814939 | 0.932212421 |
| ENSMUSG0000002807 | chr3 | Pear1 | 0.083472464 | 0.932212421 |
| ENSMUSG0000010347 | chr18 | Pcdhga7 | -0.085282618 | 0.932355204 |
| ENSMUSG0000003238 | chr9 | Dapk2 | -0.08565875 | 0.932355204 |
| ENSMUSG0000002028 | chr11 | Mpg | -0.060224553 | 0.932355204 |
| ENSMUSG0000003643 | chr5 | Exoc1 | -0.03638174 | 0.932355204 |
| ENSMUSG0000007464 | chr2 | Mmp24os1 | 0.030358116 | 0.932355204 |
| ENSMUSG0000002778 | chr3 | Ppm1l | 0.047570745 | 0.932355204 |
| ENSMUSG0000002849 | chr4 | Mllt3 | 0.046648027 | 0.932355204 |
| ENSMUSG0000002991 | chr6 | Prdm5 | 0.070708546 | 0.932355204 |

| | | | | |
|---|---|---|---|---|
| ENSMUSG0000002278 | chr16 | Pak2 | 0.028404284 | 0.932397901 |
| ENSMUSG0000002843 | chr4 | Ubap1 | 0.028637049 | 0.932397901 |
| ENSMUSG0000002772 | chr3 | Afg2a | 0.056690043 | 0.932397901 |
| ENSMUSG0000002028 | chr11 | Ahsa2 | 0.062166979 | 0.932397901 |
| ENSMUSG0000000221 | chr7 | Smg9 | 0.046923463 | 0.932397901 |
| ENSMUSG0000000956 | chr16 | Mrtfb | 0.063086131 | 0.932417135 |
| ENSMUSG0000003359 | chr8 | Rfwd3 | 0.030897309 | 0.932475671 |
| ENSMUSG0000004910 | chr6 | Ntf3 | -0.106454714 | 0.932745988 |
| ENSMUSG0000007865 | chr11 | Cntd1 | -0.187437773 | 0.932785157 |
| ENSMUSG0000002851 | chr4 | Dab1 | -0.120876734 | 0.932855297 |
| ENSMUSG0000003056 | chr7 | Nox4 | -0.097867136 | 0.933076247 |
| ENSMUSG0000002037 | chr11 | Rasgef1c | -0.075653836 | 0.933076247 |
| ENSMUSG0000009441 | chr1 | Zbed6 | 0.060331237 | 0.933076247 |
| ENSMUSG0000003116 | chrX | Porcn | 0.042072885 | 0.933076247 |
| ENSMUSG0000005513 | chr13 | Sugct | -0.131669685 | 0.933117469 |
| ENSMUSG0000002437 | chr18 | Epb41l4a | 0.073478161 | 0.933117469 |
| ENSMUSG0000002043 | chr11 | Osbp2 | -0.058705748 | 0.933124968 |
| ENSMUSG0000003402 | chr15 | Cpsf1 | 0.026947999 | 0.933124968 |
| ENSMUSG0000001476 | chr17 | Tbp | 0.033948452 | 0.933124968 |
| ENSMUSG0000004048 | chr7 | Ltbp4 | 0.045648487 | 0.933124968 |
| ENSMUSG0000007120 | chr17 | Ccdc78 | 0.151397785 | 0.933124968 |
| ENSMUSG0000002525 | chrX | Ribc1 | -0.196378808 | 0.933382136 |
| ENSMUSG0000000431 | chrX | Clcn5 | -0.074588165 | 0.933382136 |
| ENSMUSG0000004266 | chr2 | Dusp15 | -0.16355459 | 0.933382136 |
| ENSMUSG0000003852 | chr6 | C1rl | -0.07389644 | 0.933382136 |
| ENSMUSG0000004085 | chr11 | Psme4 | -0.03293209 | 0.933382136 |
| ENSMUSG0000002133 | chr13 | Mrs2 | -0.035300714 | 0.933382136 |
| ENSMUSG0000003107 | chr7 | Cttn | -0.028802332 | 0.933382136 |
| ENSMUSG0000003081 | chr7 | Rnf40 | 0.027802791 | 0.933382136 |
| ENSMUSG0000003850 | chr8 | Gdf15 | 0.162192639 | 0.933382136 |
| ENSMUSG0000004640 | chr8 | 1700067K01Rik | 0.214207034 | 0.933382136 |
| ENSMUSG0000009493 | chr19 | Rbm4 | -0.084106592 | 0.933431775 |
| ENSMUSG0000002685 | chr2 | BC005624 | -0.033260648 | 0.933431775 |
| ENSMUSG0000002812 | chr3 | Abca4 | 0.124194155 | 0.933431775 |
| ENSMUSG0000002056 | chr12 | Polr1f | 0.035191295 | 0.933431775 |
| ENSMUSG0000008990 | chr14 | Gm8113 | 0.316709975 | 0.933431775 |
| ENSMUSG0000004470 | chr11 | Ccnjl | 0.116001251 | 0.933562249 |
| ENSMUSG0000000498 | chr6 | Hnrnpa2b1 | 0.03753219 | 0.933587447 |
| ENSMUSG0000006067 | chr3 | Atp8b2 | -0.042576708 | 0.93380761 |
| ENSMUSG0000002565 | chrX | Cnksr2 | -0.086836742 | 0.933847006 |
| ENSMUSG0000005502 | chr15 | Ep300 | -0.064583494 | 0.933847006 |
| ENSMUSG0000001985 | chr10 | Perp | -0.149965063 | 0.933847006 |
| ENSMUSG0000003744 | chr4 | Cep85 | -0.054373188 | 0.933847006 |
| ENSMUSG0000007397 | chr7 | Or51r1 | -0.139616148 | 0.933847006 |
| ENSMUSG0000003736 | chr8 | Letm2 | -0.05746953 | 0.933847006 |
| ENSMUSG0000010225 | chr7 | Snrpn | -0.047490378 | 0.933847006 |
| ENSMUSG0000006673 | chr5 | Vkorc1l1 | -0.039282146 | 0.933847006 |
| ENSMUSG0000000211 | chr2 | Spi1 | -0.082245285 | 0.933847006 |
| ENSMUSG0000003404 | chr4 | Gpbp1l1 | -0.048329696 | 0.933847006 |
| ENSMUSG0000006051 | chr9 | Zfp266 | -0.037329566 | 0.933847006 |

| | | | | |
|---|---|---|---|---|
| ENSMUSG0000002514 | chr11 | Aspscr1 | -0.037720877 | 0.933847006 |
| ENSMUSG0000002010 | chr10 | Lrig3 | -0.050626498 | 0.933847006 |
| ENSMUSG0000007409 | chr7 | Svip | -0.038480591 | 0.933847006 |
| ENSMUSG0000006079 | chr7 | Gmfg | -0.064402828 | 0.933847006 |
| ENSMUSG0000003223 | chr9 | Rp9 | -0.035059202 | 0.933847006 |
| ENSMUSG0000001997 | chr10 | Epb41l2 | 0.029349656 | 0.933847006 |
| ENSMUSG0000005255 | chr8 | Gan | 0.035604026 | 0.933847006 |
| ENSMUSG0000002666 | chr2 | Sephs1 | 0.029847829 | 0.933847006 |
| ENSMUSG0000006276 | chr5 | Zfp512 | 0.031843588 | 0.933847006 |
| ENSMUSG0000002730 | chr2 | Ptpra | 0.032250997 | 0.933847006 |
| ENSMUSG0000002604 | chr1 | Ercc5 | 0.032185332 | 0.933847006 |
| ENSMUSG0000007541 | chr2 | Fnbp1 | 0.031872977 | 0.933847006 |
| ENSMUSG0000004947 | chr11 | Aff4 | 0.038584738 | 0.933847006 |
| ENSMUSG0000003552 | chr17 | Gnptg | 0.034359371 | 0.933847006 |
| ENSMUSG0000004018 | chr1 | Fmo1 | 0.172053995 | 0.933847006 |
| ENSMUSG0000002907 | chr4 | Mxra8 | 0.048914831 | 0.933847006 |
| ENSMUSG0000002684 | chr2 | Tor1a | 0.050553791 | 0.933847006 |
| ENSMUSG0000003289 | chr5 | Fbxo21 | 0.035824429 | 0.933847006 |
| ENSMUSG0000006792 | chrX | Rtl8a | 0.038299842 | 0.933847006 |
| ENSMUSG0000003030 | chr6 | Ccdc91 | 0.045000702 | 0.933847006 |
| ENSMUSG0000003737 | chr2 | Trmt6 | 0.046017073 | 0.933847006 |
| ENSMUSG0000006623 | chr9 | Tmem42 | 0.080696662 | 0.933847006 |
| ENSMUSG0000003537 | chr11 | Ccl7 | 0.205068718 | 0.933847006 |
| ENSMUSG0000002219 | chr14 | Pabpn1 | 0.051167468 | 0.933847006 |
| ENSMUSG0000002836 | chr4 | Svep1 | 0.087495944 | 0.933847006 |
| ENSMUSG0000006384 | chr9 | Ppcdc | 0.037903591 | 0.933960522 |
| ENSMUSG0000003221 | chr9 | Rsl24d1 | 0.058928864 | 0.933960522 |
| ENSMUSG0000007165 | chr19 | Lrrn4cl | 0.078349602 | 0.933960522 |
| ENSMUSG0000005630 | chr6 | Usp39 | 0.042192224 | 0.934006048 |
| ENSMUSG0000001466 | chr5 | Chfr | 0.032137723 | 0.934006048 |
| ENSMUSG0000001675 | chr15 | Bik | 0.061766076 | 0.934006048 |
| ENSMUSG0000005700 | chrX | Nxf3 | 0.212902957 | 0.934006048 |
| ENSMUSG0000001914 | chr15 | Cacng2 | 0.148453804 | 0.934006048 |
| ENSMUSG0000002713 | chr2 | Slc12a6 | -0.039436242 | 0.93460217 |
| ENSMUSG0000004035 | chr17 | Skic2 | -0.034658456 | 0.934667073 |
| ENSMUSG0000003271 | chr6 | Resf1 | 0.02737226 | 0.934667073 |
| ENSMUSG0000001675 | chr15 | Ttll12 | -0.050983015 | 0.93475139 |
| ENSMUSG0000003887 | chr8 | Zfhx3 | 0.081443774 | 0.934775324 |
| ENSMUSG0000004282 | chr7 | Trim72 | -0.216643233 | 0.934918718 |
| ENSMUSG0000002804 | chr3 | Cks1b | 0.067998737 | 0.934918718 |
| ENSMUSG0000004672 | chr18 | Cystm1 | -0.053669684 | 0.93502379 |
| ENSMUSG0000003257 | chr9 | Col6a4 | -0.189542043 | 0.935111642 |
| ENSMUSG0000002084 | chr11 | Doc2b | -0.06663572 | 0.935111642 |
| ENSMUSG0000003033 | chr6 | Cd27 | -0.150013803 | 0.935111642 |
| ENSMUSG0000005871 | chr1 | Fcer1g | -0.068941813 | 0.935111642 |
| ENSMUSG0000003736 | chrX | Kdm6a | -0.038970208 | 0.935111642 |
| ENSMUSG0000000558 | chr16 | Adcy9 | -0.047573007 | 0.935111642 |
| ENSMUSG0000001807 | chr5 | Med13l | -0.038506551 | 0.935111642 |
| ENSMUSG0000002660 | chr1 | Cenpf | 0.184998739 | 0.935111642 |
| ENSMUSG0000000156 | chr5 | Nom1 | 0.027126934 | 0.935111642 |

| | | | | |
|---|---|---|---|---|
| ENSMUSG0000000929 | chr10 | Pttg1ip | 0.031285063 | 0.935111642 |
| ENSMUSG0000003082 | chr7 | Prr14 | 0.036446082 | 0.935111642 |
| ENSMUSG0000003609 | chr19 | Myrf | 0.139920029 | 0.935111642 |
| ENSMUSG0000004737 | chr19 | B4gat1 | -0.277796169 | 0.935115536 |
| ENSMUSG0000002519 | chr19 | Cyp2c23 | -0.333177522 | 0.935115536 |
| ENSMUSG0000003431 | chrX | Kif4 | -0.24544553 | 0.935115536 |
| ENSMUSG0000002042 | chr11 | Igfbp3 | -0.075939842 | 0.935115536 |
| ENSMUSG0000003446 | chr8 | Dynlrb2 | -0.107059999 | 0.935115536 |
| ENSMUSG0000002070 | chr11 | Rnf135 | -0.083505501 | 0.935115536 |
| ENSMUSG0000003218 | chr9 | Yipf2 | -0.060715676 | 0.935115536 |
| ENSMUSG0000002841 | chr4 | B4galt1 | -0.05548933 | 0.935115536 |
| ENSMUSG0000004841 | chr18 | Zfp407 | -0.058506157 | 0.935115536 |
| ENSMUSG0000003388 | chr14 | Pxk | -0.036865081 | 0.935115536 |
| ENSMUSG0000006344 | chr16 | Nmral1 | -0.045165068 | 0.935115536 |
| ENSMUSG0000002887 | chr4 | Cdca8 | -0.13197377 | 0.935115536 |
| ENSMUSG0000000075 | chr8 | Tubgcp3 | -0.022479232 | 0.935115536 |
| ENSMUSG0000003186 | chr8 | Ints10 | 0.029171769 | 0.935115536 |
| ENSMUSG0000005384 | chr4 | Txlna | 0.030490138 | 0.935115536 |
| ENSMUSG0000003100 | chrX | Atp6ap2 | 0.031738323 | 0.935115536 |
| ENSMUSG0000003170 | chr8 | Tbc1d9 | 0.051548902 | 0.935115536 |
| ENSMUSG0000003207 | chr9 | Bud13 | 0.046985374 | 0.935115536 |
| ENSMUSG0000003579 | chr12 | Twist1 | 0.049568875 | 0.935115536 |
| ENSMUSG0000009568 | chr17 | Rnaset2a | 0.234037445 | 0.935115536 |
| ENSMUSG0000003561 | chr4 | Frmpd1 | 0.06235436 | 0.935245099 |
| ENSMUSG0000002049 | chr11 | Ska2 | 0.048735517 | 0.935678185 |
| ENSMUSG0000002442 | chr17 | Dhx16 | 0.041751132 | 0.935745201 |
| ENSMUSG0000004960 | chr5 | Zfp644 | 0.056775069 | 0.935745201 |
| ENSMUSG0000003058 | chr7 | Rasgrp4 | -0.121728543 | 0.935878314 |
| ENSMUSG0000007122 | chr6 | Cecr2 | -0.112175486 | 0.935878314 |
| ENSMUSG0000002672 | chr1 | Tnn | -0.12415979 | 0.935878314 |
| ENSMUSG0000002031 | chr10 | Shc2 | -0.048464609 | 0.935878314 |
| ENSMUSG0000003780 | chr9 | Iqch | -0.217144061 | 0.935878314 |
| ENSMUSG0000000778 | chr7 | Cpt1c | -0.05156174 | 0.935878314 |
| ENSMUSG0000002059 | chr11 | Rgs9 | -0.039423586 | 0.935878314 |
| ENSMUSG0000004364 | chr11 | Pld6 | -0.152572288 | 0.935878314 |
| ENSMUSG0000005928 | chr13 | Cdyl | -0.045984879 | 0.935878314 |
| ENSMUSG0000004050 | chr2 | Ambra1 | -0.032363584 | 0.935878314 |
| ENSMUSG0000007165 | chr19 | Hnrnpul2 | 0.022263737 | 0.935878314 |
| ENSMUSG0000006212 | chr3 | Cttnbp2nl | 0.047553552 | 0.935878314 |
| ENSMUSG0000005918 | chr6 | Skap2 | 0.037395759 | 0.935878314 |
| ENSMUSG0000005167 | chr18 | Pcdhb6 | 0.114875374 | 0.935878314 |
| ENSMUSG0000000703 | chr17 | Hspa1l | 0.164687343 | 0.935878314 |
| ENSMUSG0000001714 | chr2 | Rnd3 | 0.038627274 | 0.935878314 |
| ENSMUSG0000002432 | chr17 | Ring1 | 0.033659322 | 0.935878314 |
| ENSMUSG0000003517 | chr11 | Plekhh3 | 0.055828267 | 0.935878314 |
| ENSMUSG0000002439 | chr17 | Aif1 | 0.145921253 | 0.935878314 |
| ENSMUSG0000004640 | chr9 | Rbp1 | 0.047933661 | 0.935878314 |
| ENSMUSG0000005019 | chr19 | Scd4 | -0.099392527 | 0.935905141 |
| ENSMUSG0000002841 | chr4 | Fktn | -0.031370387 | 0.935905141 |
| ENSMUSG0000002633 | chr1 | Ccdc93 | -0.030584159 | 0.935905141 |

| | | | | |
|---|---|---|---|---|
| ENSMUSG0000001575 | chr10 | Ppil4 | 0.025612648 | 0.935905141 |
| ENSMUSG0000002720 | chr2 | Myef2 | 0.02646874 | 0.935905141 |
| ENSMUSG0000006024 | chr1 | Alyref2 | 0.124241423 | 0.935905141 |
| ENSMUSG0000003233 | chr9 | Hcn4 | 0.109376718 | 0.935905141 |
| ENSMUSG0000003079 | chr7 | Cd37 | 0.096093907 | 0.935905141 |
| ENSMUSG0000003113 | chrX | Rbmx | 0.049708738 | 0.935905141 |
| ENSMUSG0000006839 | chr2 | Cep152 | 0.051927334 | 0.935905141 |
| ENSMUSG0000005529 | chr4 | Tmem245 | 0.054294529 | 0.935945807 |
| ENSMUSG0000005596 | chr4 | Triqk | 0.052223798 | 0.936038631 |
| ENSMUSG0000003863 | chr4 | Zkscan16 | -0.123332899 | 0.936060237 |
| ENSMUSG0000003681 | chr1 | Dpp10 | -0.073359022 | 0.936060237 |
| ENSMUSG0000003225 | chr9 | Phip | -0.043318834 | 0.936060237 |
| ENSMUSG0000006136 | chr15 | Phf5a | 0.053706137 | 0.936060237 |
| ENSMUSG0000002401 | chr17 | Mtch1 | 0.036859656 | 0.936060237 |
| ENSMUSG0000002562 | chrX | Phf6 | 0.027825468 | 0.936126733 |
| ENSMUSG0000005858 | chr15 | Serhl | 0.053749773 | 0.936299705 |
| ENSMUSG0000005472 | chr17 | Vmac | -0.048162259 | 0.936359317 |
| ENSMUSG0000002244 | chr15 | Myh9 | -0.028300456 | 0.93638442 |
| ENSMUSG0000002299 | chr15 | Wnt10b | 0.192653176 | 0.93638442 |
| ENSMUSG0000003175 | chr8 | Amfr | 0.034379991 | 0.93638442 |
| ENSMUSG0000002049 | chr11 | 2810021J22Rik | 0.059362595 | 0.93638442 |
| ENSMUSG0000007881 | chr7 | Prkcg | 0.109297474 | 0.93638442 |
| ENSMUSG0000003180 | chr8 | Jak3 | -0.064308423 | 0.936740464 |
| ENSMUSG0000004230 | chr7 | Setd1a | 0.053256299 | 0.936740464 |
| ENSMUSG0000004786 | chr8 | Lonp2 | 0.032918932 | 0.936785856 |
| ENSMUSG0000007955 | chr17 | Kifc1 | 0.169726355 | 0.936785856 |
| ENSMUSG0000004873 | chr17 | Ggnbp1 | 0.073811254 | 0.936785856 |
| ENSMUSG0000005376 | chr3 | Lysmd1 | 0.07620886 | 0.936789838 |
| ENSMUSG0000006131 | chr8 | Ddhd2 | -0.033494361 | 0.936859973 |
| ENSMUSG0000003412 | chr12 | Pomt2 | -0.032466597 | 0.936859973 |
| ENSMUSG0000003421 | chr9 | Atm | -0.065971319 | 0.936887761 |
| ENSMUSG0000005252 | chr17 | Spdya | -0.208305321 | 0.936887761 |
| ENSMUSG0000005520 | chr5 | Ankrd17 | -0.047671658 | 0.936887761 |
| ENSMUSG0000002900 | chr4 | Agtrap | -0.039043491 | 0.936887761 |
| ENSMUSG0000006668 | chr5 | Pilrb1 | -0.18840336 | 0.936887761 |
| ENSMUSG0000002054 | chr11 | Elac2 | -0.033235869 | 0.936887761 |
| ENSMUSG0000004140 | chr14 | Wapl | -0.035782428 | 0.936887761 |
| ENSMUSG0000004265 | chr11 | Alkbh5 | -0.031361426 | 0.936887761 |
| ENSMUSG0000006412 | chrX | Med14 | 0.042249233 | 0.936887761 |
| ENSMUSG0000000087 | chr2 | Pxmp4 | 0.041136163 | 0.936887761 |
| ENSMUSG0000002005 | chr10 | Igf1 | 0.054431178 | 0.936887761 |
| ENSMUSG0000000279 | chr12 | Jag2 | -0.064950604 | 0.937112787 |
| ENSMUSG0000004691 | chr10 | Myct1 | -0.069029649 | 0.937112787 |
| ENSMUSG0000003871 | chr3 | Mindy1 | -0.046842474 | 0.937112787 |
| ENSMUSG0000001594 | chr5 | Gtf2ird2 | -0.048590603 | 0.937112787 |
| ENSMUSG0000002982 | chr6 | Luc7l2 | -0.033968667 | 0.937112787 |
| ENSMUSG0000004618 | chr7 | Zfp84 | -0.043924201 | 0.937112787 |
| ENSMUSG0000005422 | chr6 | Tprkb | -0.031808999 | 0.937112787 |
| ENSMUSG0000007346 | chr17 | Pnldc1 | 0.172301008 | 0.937112787 |
| ENSMUSG0000002553 | chrX | Chm | 0.034922197 | 0.937112787 |

| | | | | |
|---|---|---|---|---|
| ENSMUSG0000003437 | chr19 | Tkfc | 0.044661045 | 0.937112787 |
| ENSMUSG0000002490 | chr19 | Ppp6r3 | -0.0228979 | 0.937258016 |
| ENSMUSG0000001602 | chr15 | Celsr1 | -0.086536799 | 0.937357876 |
| ENSMUSG0000004576 | chr15 | Basp1 | -0.042295893 | 0.937357876 |
| ENSMUSG0000005423 | chr19 | Fra10ac1 | -0.032893051 | 0.937357876 |
| ENSMUSG0000003262 | chr5 | Oas1d | 0.095412005 | 0.937357876 |
| ENSMUSG0000002919 | chr5 | D5Ertd579e | 0.030762576 | 0.937357876 |
| ENSMUSG0000003320 | chr5 | Ttc28 | 0.04314044 | 0.937357876 |
| ENSMUSG0000002268 | chr16 | Bfar | 0.042785279 | 0.93744421 |
| ENSMUSG0000001575 | chr3 | Aph1a | 0.041504052 | 0.93744421 |
| ENSMUSG0000002089 | chr11 | Pfas | 0.033207942 | 0.937731651 |
| ENSMUSG0000003040 | chr7 | Vasp | 0.037144746 | 0.937731651 |
| ENSMUSG0000002235 | chr15 | Klhl38 | 0.118162998 | 0.937742572 |
| ENSMUSG0000002491 | chr19 | Ctsw | 0.113024713 | 0.937742572 |
| ENSMUSG0000003735 | chrX | Dipk2b | 0.116486419 | 0.937742572 |
| ENSMUSG0000002065 | chr12 | Dnajc27 | -0.031236598 | 0.937765142 |
| ENSMUSG0000004299 | chr6 | Borcs5 | 0.061463267 | 0.937811282 |
| ENSMUSG0000002335 | chr5 | Agap3 | 0.034345409 | 0.93782786 |
| ENSMUSG0000003013 | chr6 | Bms1 | 0.031398082 | 0.937842406 |
| ENSMUSG0000007064 | chr1 | Sox13 | 0.051519635 | 0.937885204 |
| ENSMUSG0000006223 | chr3 | Rapgef2 | 0.042394608 | 0.937953988 |
| ENSMUSG0000004505 | chr17 | Kcng3 | -0.082140416 | 0.937962961 |
| ENSMUSG0000007456 | chr2 | Gcnt7 | -0.149122062 | 0.938177655 |
| ENSMUSG0000005353 | chr19 | Cstf2t | 0.032862731 | 0.938177655 |
| ENSMUSG0000002522 | chr19 | Kcnip2 | -0.066927372 | 0.938725285 |
| ENSMUSG0000001929 | chr5 | Tmem129 | 0.039776271 | 0.938746538 |
| ENSMUSG0000002028 | chr10 | Cfap410 | -0.031681013 | 0.938760524 |
| ENSMUSG0000004199 | chr12 | Rapgef5 | 0.080404434 | 0.938760524 |
| ENSMUSG0000002828 | chr4 | Epha7 | 0.056017652 | 0.938760524 |
| ENSMUSG0000004940 | chr2 | Ogfr | -0.042857208 | 0.938808576 |
| ENSMUSG0000002630 | chr1 | Ilkap | 0.031207954 | 0.938808576 |
| ENSMUSG0000003963 | chr9 | Odad3 | -0.100236522 | 0.938870932 |
| ENSMUSG0000003163 | chr8 | Cfap97 | -0.034450221 | 0.938870932 |
| ENSMUSG0000007876 | chr7 | U2af1l4 | -0.21479824 | 0.938870932 |
| ENSMUSG0000002854 | chr4 | Dph2 | -0.050969012 | 0.938870932 |
| ENSMUSG0000002194 | chr14 | Zmym2 | -0.027692297 | 0.938870932 |
| ENSMUSG0000004218 | chr9 | Nfrkb | 0.037863637 | 0.938870932 |
| ENSMUSG0000004698 | chr5 | Tapt1 | 0.033111306 | 0.938870932 |
| ENSMUSG0000004828 | chr14 | Dleu7 | -0.093331528 | 0.938939947 |
| ENSMUSG0000002879 | chr4 | A3galt2 | 0.161493109 | 0.938977171 |
| ENSMUSG0000004056 | chr19 | Btaf1 | 0.05801914 | 0.938977171 |
| ENSMUSG0000004516 | chr7 | AI467606 | -0.064292789 | 0.93906265 |
| ENSMUSG0000004491 | chr12 | Syt16 | 0.083611817 | 0.939074355 |
| ENSMUSG0000006963 | chr11 | Strada | 0.032495581 | 0.939274241 |
| ENSMUSG0000003240 | chr9 | Smad3 | 0.043477583 | 0.93928381 |
| ENSMUSG0000002106 | chr12 | Atl1 | -0.033133092 | 0.939330272 |
| ENSMUSG0000009226 | chr8 | Zfp963 | 0.074362405 | 0.939466513 |
| ENSMUSG0000004344 | chr11 | Gjc2 | 0.081879039 | 0.939504994 |
| ENSMUSG0000003812 | chr5 | Mphosph9 | -0.060770889 | 0.939705616 |
| ENSMUSG0000005010 | chr7 | Hmx2 | -0.076450051 | 0.939705616 |

| | | | | |
|---|---|---|---|---|
| ENSMUSG0000002497 | chr19 | Pdcd4 | -0.058509142 | 0.939705616 |
| ENSMUSG0000006432 | chr2 | Scn1a | -0.089142241 | 0.939705616 |
| ENSMUSG0000002522 | chr19 | Ldb1 | -0.04276338 | 0.939705616 |
| ENSMUSG0000003818 | chr11 | Scarf1 | -0.089917385 | 0.939705616 |
| ENSMUSG0000010274 | chr18 | Pcdhgb2 | -0.072126501 | 0.939705616 |
| ENSMUSG0000001196 | chr15 | Ccnt1 | -0.033683382 | 0.939705616 |
| ENSMUSG0000002283 | chr16 | Hcls1 | -0.074770703 | 0.939705616 |
| ENSMUSG0000003217 | chr9 | Tyk2 | -0.0329429 | 0.939705616 |
| ENSMUSG0000004046 | chr10 | Os9 | -0.039216997 | 0.939705616 |
| ENSMUSG0000005626 | chr9 | Cep70 | -0.048536723 | 0.939705616 |
| ENSMUSG0000003087 | chr7 | Mettl9 | -0.027634269 | 0.939705616 |
| ENSMUSG0000006045 | chr18 | Rnf14 | -0.027944678 | 0.939705616 |
| ENSMUSG0000004041 | chr10 | Dtx3 | -0.036028696 | 0.939705616 |
| ENSMUSG0000004113 | chr17 | Zfp598 | -0.029261594 | 0.939705616 |
| ENSMUSG0000008364 | chr7 | Rasl2-9 | 0.20188362 | 0.939705616 |
| ENSMUSG0000010509 | chr5 | Gbp10 | -0.430854198 | 0.939705616 |
| ENSMUSG0000002096 | chr12 | Sel1l | 0.023741589 | 0.939705616 |
| ENSMUSG0000003919 | chr14 | Adk | 0.040135128 | 0.939705616 |
| ENSMUSG0000003145 | chr8 | Coprs | 0.056756631 | 0.939705616 |
| ENSMUSG0000000580 | chr2 | Bloc1s6 | 0.026565014 | 0.939705616 |
| ENSMUSG0000004967 | chr17 | Zbtb14 | 0.052422988 | 0.939705616 |
| ENSMUSG0000003437 | chr16 | Wdr5b | 0.04744384 | 0.939705616 |
| ENSMUSG0000005334 | chr17 | Zfp943 | 0.037629169 | 0.939705616 |
| ENSMUSG0000006332 | chr10 | Uqcc6 | 0.112084711 | 0.939705616 |
| ENSMUSG0000003456 | chr9 | Ccpg1 | 0.108417008 | 0.939705616 |
| ENSMUSG0000004975 | chr12 | Rpl36al | 0.168738114 | 0.939705616 |
| ENSMUSG0000003876 | chr2 | Lmx1b | 0.097952975 | 0.939705616 |
| ENSMUSG0000004767 | chrX | Gpr82 | 0.210681496 | 0.939993441 |
| ENSMUSG0000003884 | chr2 | Ythdf1 | -0.028214646 | 0.940064211 |
| ENSMUSG0000006118 | chr2 | Sfmbt2 | 0.131778005 | 0.940131028 |
| ENSMUSG0000005067 | chr11 | Gp1ba | 0.094539827 | 0.940131028 |
| ENSMUSG0000005370 | chr2 | Nebl | 0.068576802 | 0.940131028 |
| ENSMUSG0000001769 | chr2 | Ada | -0.104762051 | 0.940176685 |
| ENSMUSG0000002478 | chr19 | Jak2 | 0.034020755 | 0.940436455 |
| ENSMUSG0000006211 | chr7 | Zfp954 | 0.045596125 | 0.940436455 |
| ENSMUSG0000003506 | chr4 | Oma1 | 0.052555551 | 0.940639606 |
| ENSMUSG0000002171 | chr13 | Nln | -0.049611968 | 0.940787544 |
| ENSMUSG0000003158 | chr8 | Wrn | 0.046743545 | 0.940787544 |
| ENSMUSG0000007845 | chr10 | Ppil6 | -0.042935024 | 0.941190955 |
| ENSMUSG0000004787 | chr18 | Usp14 | 0.028825109 | 0.941190955 |
| ENSMUSG0000006420 | chr15 | Krt2 | 0.180551406 | 0.941190955 |
| ENSMUSG0000003073 | chr7 | Atp2a1 | 0.386631136 | 0.941190955 |
| ENSMUSG0000000345 | chr1 | Ncstn | 0.029783995 | 0.941207387 |
| ENSMUSG0000002283 | chr16 | Ccdc14 | -0.067538167 | 0.941224173 |
| ENSMUSG0000000798 | chr5 | Ift22 | -0.042803725 | 0.941224173 |
| ENSMUSG0000003980 | chr15 | Cplane1 | -0.049146105 | 0.941224173 |
| ENSMUSG0000001971 | chr3 | Mrpl24 | 0.043967212 | 0.941258055 |
| ENSMUSG0000002867 | chr4 | Rnf220 | 0.028826101 | 0.941267303 |
| ENSMUSG0000002907 | chr4 | Tnfrsf4 | -0.134652969 | 0.941274605 |
| ENSMUSG0000002932 | chr5 | Enoph1 | 0.048369097 | 0.941274605 |

| | | | | |
|---|---|---|---|---|
| ENSMUSG0000003289 | chr4 | Nfyc | 0.036141452 | 0.941274605 |
| ENSMUSG0000006627 | chr5 | Vps37b | 0.039398501 | 0.941274605 |
| ENSMUSG0000005687 | chr1 | Gulp1 | 0.033114033 | 0.941275806 |
| ENSMUSG0000003123 | chrX | Itm2a | -0.032310687 | 0.941280986 |
| ENSMUSG0000007422 | chr7 | Zfp382 | 0.066268624 | 0.941280986 |
| ENSMUSG0000002052 | chr11 | Fam114a2 | 0.030668597 | 0.941310513 |
| ENSMUSG0000002945 | chr5 | Tmem116 | -0.096690465 | 0.941655101 |
| ENSMUSG0000001009 | chr19 | Slc3a2 | 0.026652993 | 0.941655101 |
| ENSMUSG0000005404 | chr7 | Klk13 | 0.167543214 | 0.941655101 |
| ENSMUSG0000010434 | chr18 | Pcdhga3 | -0.0759142 | 0.941704926 |
| ENSMUSG0000004114 | chr4 | Tmco4 | -0.059818401 | 0.941704926 |
| ENSMUSG0000004699 | chr17 | Wdr27 | 0.085399965 | 0.941704926 |
| ENSMUSG0000003126 | chrX | Btk | 0.137412557 | 0.941704926 |
| ENSMUSG0000007891 | chr2 | Dpm1 | 0.051218549 | 0.941704926 |
| ENSMUSG0000003845 | chr6 | Spsb2 | -0.058793664 | 0.941733562 |
| ENSMUSG0000002266 | chr16 | Slc35a5 | -0.0272146 | 0.941733562 |
| ENSMUSG0000002505 | chrX | Tasl | 0.184580521 | 0.941733562 |
| ENSMUSG0000004654 | chr7 | Zfp526 | 0.048017256 | 0.941733562 |
| ENSMUSG0000001776 | chr2 | Zswim1 | 0.051508813 | 0.941733562 |
| ENSMUSG0000003138 | chrX | Arhgap4 | -0.067205376 | 0.941749155 |
| ENSMUSG0000009114 | chr14 | Phf11c | 0.086527813 | 0.941749155 |
| ENSMUSG0000002193 | chr14 | Spryd7 | -0.032819329 | 0.941852991 |
| ENSMUSG0000005123 | chr12 | Gen1 | -0.099034513 | 0.941852991 |
| ENSMUSG0000002424 | chr17 | Tmem178 | 0.112365857 | 0.941852991 |
| ENSMUSG0000003562 | chr10 | Ric8b | 0.028701357 | 0.941855035 |
| ENSMUSG0000002102 | chr12 | Prorp | 0.041450125 | 0.941855035 |
| ENSMUSG0000002018 | chr10 | Cpm | -0.086354803 | 0.942183311 |
| ENSMUSG0000003589 | chr10 | Rnf126 | 0.030738136 | 0.942273569 |
| ENSMUSG0000004703 | chr18 | Pcdhb15 | -0.085385594 | 0.942337748 |
| ENSMUSG0000000526 | chr11 | Zfp287 | -0.045448687 | 0.942337748 |
| ENSMUSG0000002291 | chr16 | Brwd1 | -0.038002287 | 0.942337748 |
| ENSMUSG0000003832 | chr1 | 1700066M21Rik | -0.043125158 | 0.942337748 |
| ENSMUSG0000004076 | chr14 | Appl1 | -0.027381642 | 0.942337748 |
| ENSMUSG0000002307 | chr5 | Gtf2ird1 | -0.035176221 | 0.942337748 |
| ENSMUSG0000002547 | chr7 | Tubgcp2 | 0.027990119 | 0.942337748 |
| ENSMUSG0000000176 | chr2 | Rin2 | 0.027660042 | 0.942337748 |
| ENSMUSG0000003490 | chr9 | Sidt2 | 0.037364471 | 0.942337748 |
| ENSMUSG0000003696 | chr5 | Cnpy4 | 0.037452592 | 0.942337748 |
| ENSMUSG0000002176 | chr14 | Kat6b | 0.05416163 | 0.942337748 |
| ENSMUSG0000003345 | chr12 | Zbtb1 | 0.036079348 | 0.942337748 |
| ENSMUSG0000004183 | chr17 | Sytl3 | 0.161127994 | 0.942409365 |
| ENSMUSG0000005866 | chr12 | Nkx2-9 | -0.219691921 | 0.942673507 |
| ENSMUSG0000003509 | chr2 | Secisbp2l | -0.035866756 | 0.942673507 |
| ENSMUSG0000002900 | chr4 | Mthfr | -0.041442954 | 0.942673507 |
| ENSMUSG0000002579 | chr11 | Hgs | -0.025566194 | 0.942673507 |
| ENSMUSG0000003310 | chr10 | Lss | -0.073824056 | 0.942850603 |
| ENSMUSG0000003691 | chrX | Zfp280c | -0.057075243 | 0.942850603 |
| ENSMUSG0000003464 | chr17 | Ankrd12 | -0.060328821 | 0.942850603 |
| ENSMUSG0000004431 | chr7 | Gpr4 | -0.062730142 | 0.942850603 |
| ENSMUSG0000005817 | chr9 | Smco4 | -0.077994558 | 0.942850603 |

| | | | | |
|---|---|---|---|---|
| ENSMUSG0000002773 | chr3 | Slc7a11 | -0.205065479 | 0.942850603 |
| ENSMUSG0000002654 | chr1 | Tagln2 | -0.046663516 | 0.942850603 |
| ENSMUSG0000005862 | chr2 | Adra2b | -0.088902392 | 0.942850603 |
| ENSMUSG0000004019 | chr10 | Nemp1 | -0.050637359 | 0.942850603 |
| ENSMUSG0000003424 | chr14 | Slc25a37 | -0.060901026 | 0.942850603 |
| ENSMUSG0000005042 | chr7 | Fbxo46 | -0.041355794 | 0.942850603 |
| ENSMUSG0000000090 | chr10 | Mmp11 | -0.041413042 | 0.942850603 |
| ENSMUSG0000007070 | chr7 | Eid2b | -0.03786194 | 0.942850603 |
| ENSMUSG0000002296 | chr16 | Ifngr2 | -0.034266163 | 0.942850603 |
| ENSMUSG0000004471 | chr5 | Dok7 | -0.068776924 | 0.942850603 |
| ENSMUSG0000007949 | chr6 | Nat8f5 | -0.159930347 | 0.942850603 |
| ENSMUSG0000003162 | chr8 | Casp3 | -0.031729027 | 0.942850603 |
| ENSMUSG0000003322 | chr1 | Wnt6 | -0.098169374 | 0.942850603 |
| ENSMUSG0000005543 | chr3 | Srsf11 | -0.033009452 | 0.942850603 |
| ENSMUSG0000002677 | chr2 | Yme1l1 | -0.027706971 | 0.942850603 |
| ENSMUSG0000002766 | chr3 | Mfn1 | -0.026031608 | 0.942850603 |
| ENSMUSG0000003110 | chrX | Rab33a | -0.113404806 | 0.942850603 |
| ENSMUSG0000000642 | chrX | Steep1 | -0.023419708 | 0.942850603 |
| ENSMUSG0000003448 | chr9 | Poglut3 | 0.024313183 | 0.942850603 |
| ENSMUSG0000003062 | chr7 | Zfand6 | 0.033016892 | 0.942850603 |
| ENSMUSG0000005986 | chr5 | Tnip2 | 0.045501097 | 0.942850603 |
| ENSMUSG0000002444 | chr18 | Dele1 | 0.027992373 | 0.942850603 |
| ENSMUSG0000004060 | chr16 | Bace2 | 0.071515687 | 0.942850603 |
| ENSMUSG0000003563 | chr7 | Cnot3 | 0.036140388 | 0.942850603 |
| ENSMUSG0000003522 | chr7 | Ccdc106 | 0.071192126 | 0.942850603 |
| ENSMUSG0000003468 | chr11 | Fam171a2 | 0.029313024 | 0.942850603 |
| ENSMUSG0000000589 | chr10 | Nr2c1 | 0.041750965 | 0.942850603 |
| ENSMUSG0000002548 | chr7 | Sirt3 | 0.039468578 | 0.942850603 |
| ENSMUSG0000004162 | chr11 | Mtnap1 | 0.044221175 | 0.942850603 |
| ENSMUSG0000003921 | chr1 | Gpatch2 | 0.031639845 | 0.942850603 |
| ENSMUSG0000007867 | chr7 | Fam174b | 0.051178945 | 0.942850603 |
| ENSMUSG0000003093 | chr7 | Chst15 | 0.040419073 | 0.942850603 |
| ENSMUSG0000007558 | chr11 | Hoxb2 | 0.063565477 | 0.942850603 |
| ENSMUSG0000003812 | chr18 | Fam210a | 0.039576451 | 0.942850603 |
| ENSMUSG0000002716 | chr2 | Pax6 | 0.19734729 | 0.942850603 |
| ENSMUSG0000003894 | chr7 | Prc1 | 0.085365143 | 0.942850603 |
| ENSMUSG0000003107 | chr7 | Mrgprf | 0.07065401 | 0.942850603 |
| ENSMUSG0000000498 | chr6 | Fxyd4 | 0.188089695 | 0.942850603 |
| ENSMUSG0000006417 | chr7 | Tnnt1 | 0.093270653 | 0.942850603 |
| ENSMUSG0000005006 | chr3 | Zfp697 | -0.077436929 | 0.943162539 |
| ENSMUSG0000003470 | chr11 | Dnai2 | -0.066501045 | 0.943457708 |
| ENSMUSG0000002285 | chr16 | Senp2 | 0.028681863 | 0.943457708 |
| ENSMUSG0000003557 | chr4 | Dcaf10 | 0.030439871 | 0.943457708 |
| ENSMUSG0000002118 | chr12 | Dglucy | -0.060581096 | 0.943465883 |
| ENSMUSG0000002425 | chr17 | Thada | 0.039829429 | 0.943465883 |
| ENSMUSG0000004403 | chr2 | Ccdc141 | -0.057133451 | 0.943748193 |
| ENSMUSG0000005659 | chr7 | Zfp175 | 0.047139487 | 0.943901229 |
| ENSMUSG0000002971 | chr5 | Gigyf1 | -0.045537097 | 0.943913822 |
| ENSMUSG0000006387 | chr6 | Chd4 | 0.03752221 | 0.943913822 |
| ENSMUSG0000003801 | chr13 | Fam120a | 0.02706731 | 0.943916177 |

| | | | | |
|---|---|---|---|---|
| ENSMUSG0000004950 | chr3 | Proser1 | 0.045340233 | 0.943916177 |
| ENSMUSG0000007450 | chr9 | Fat3 | 0.059022581 | 0.943916177 |
| ENSMUSG0000002180 | chr14 | Nid2 | 0.059156526 | 0.944087592 |
| ENSMUSG0000001570 | chr3 | Anxa9 | -0.114433623 | 0.944124316 |
| ENSMUSG0000003117 | chrX | Rpgr | -0.047349728 | 0.944124316 |
| ENSMUSG0000002089 | chr11 | Tmem107 | -0.065862739 | 0.944124316 |
| ENSMUSG0000002157 | chr13 | Pdcd6 | -0.036205831 | 0.944124316 |
| ENSMUSG0000002399 | chr17 | Foxp4 | -0.034831948 | 0.944124316 |
| ENSMUSG0000003412 | chr11 | Srsf2 | -0.032208598 | 0.944124316 |
| ENSMUSG0000002012 | chr11 | Vps54 | -0.02090901 | 0.944124316 |
| ENSMUSG0000002485 | chr19 | Sf3b2 | 0.020875854 | 0.944124316 |
| ENSMUSG0000007250 | chr15 | Phf20l1 | 0.034385478 | 0.944124316 |
| ENSMUSG0000002053 | chr11 | Acaca | 0.040794672 | 0.944124316 |
| ENSMUSG0000003767 | chr12 | Inf2 | 0.052076587 | 0.944124316 |
| ENSMUSG0000003102 | chr7 | Stk33 | 0.109440315 | 0.944124316 |
| ENSMUSG0000002575 | chr3 | Plk4 | -0.059112912 | 0.944184518 |
| ENSMUSG0000006132 | chr4 | Dnai1 | -0.117672397 | 0.94419264 |
| ENSMUSG0000002637 | chr1 | Tsn | 0.027729073 | 0.94419264 |
| ENSMUSG0000005829 | chr5 | Zfp68 | 0.040748841 | 0.94419264 |
| ENSMUSG0000004142 | chr17 | Nthl1 | -0.04485394 | 0.944238842 |
| ENSMUSG0000002007 | chr10 | Vps26a | 0.035017271 | 0.944258673 |
| ENSMUSG0000002300 | chr15 | Prpf40b | 0.034105096 | 0.944258673 |
| ENSMUSG0000005238 | chr19 | Trpm3 | -0.052041897 | 0.944289155 |
| ENSMUSG0000002325 | chr9 | Slc26a6 | -0.054751523 | 0.944313733 |
| ENSMUSG0000002687 | chr2 | Phf19 | 0.108921224 | 0.944313733 |
| ENSMUSG0000004195 | chr11 | Pigs | 0.026536755 | 0.944313733 |
| ENSMUSG0000002561 | chr16 | Bach1 | 0.04421179 | 0.944313733 |
| ENSMUSG0000002603 | chr1 | Ppil3 | 0.043324279 | 0.944313733 |
| ENSMUSG0000002978 | chr6 | Nt5c3 | 0.025318288 | 0.944332627 |
| ENSMUSG0000004225 | chr13 | Isl1 | -0.115178737 | 0.94442578 |
| ENSMUSG0000000450 | chr7 | Gab2 | -0.044980159 | 0.94442578 |
| ENSMUSG0000003243 | chr9 | Cmtm6 | -0.045237892 | 0.94442578 |
| ENSMUSG0000003197 | chr8 | Abcb10 | -0.037519563 | 0.94442578 |
| ENSMUSG0000003339 | chr2 | Spg11 | -0.029055087 | 0.94442578 |
| ENSMUSG0000003153 | chr8 | Ikbkb | -0.019881852 | 0.94442578 |
| ENSMUSG0000006316 | chr7 | Numbl | 0.03781153 | 0.94442578 |
| ENSMUSG0000003741 | chr19 | Best1 | 0.098901811 | 0.94442578 |
| ENSMUSG0000000106 | chr8 | Zfp276 | 0.030747281 | 0.94442578 |
| ENSMUSG0000005622 | chr8 | Cars2 | 0.040906989 | 0.94442578 |
| ENSMUSG0000002009 | chr10 | Pcbd1 | 0.044670337 | 0.94442578 |
| ENSMUSG0000004230 | chr1 | Tmem183a | 0.035989663 | 0.94442578 |
| ENSMUSG0000003445 | chr10 | Polr3b | 0.053824763 | 0.94442578 |
| ENSMUSG0000002579 | chr4 | Pgm1 | 0.033547634 | 0.94442578 |
| ENSMUSG0000002918 | chr5 | Cimip2c | 0.222691555 | 0.94442578 |
| ENSMUSG0000002833 | chr4 | Trmo | -0.073849514 | 0.944492261 |
| ENSMUSG0000000232 | chr14 | Mdp1 | -0.037055066 | 0.944492261 |
| ENSMUSG0000000205 | chr11 | Supt6 | -0.026673898 | 0.944492261 |
| ENSMUSG0000004896 | chr1 | Prex2 | -0.065673409 | 0.944584186 |
| ENSMUSG0000002171 | chr13 | Rgs7bp | -0.053591704 | 0.944584186 |
| ENSMUSG0000002795 | chr3 | Tmem144 | -0.152184864 | 0.944584186 |

| | | | | |
|---|---|---|---|---|
| ENSMUSG0000011866 | chr7 | Muc6 | -0.115057325 | 0.944584186 |
| ENSMUSG0000006727 | chrX | Capn6 | -0.062826226 | 0.944584186 |
| ENSMUSG0000004992 | chr11 | Glp2r | -0.051215695 | 0.944584186 |
| ENSMUSG0000005425 | chr5 | Fgfr3 | -0.082224845 | 0.944584186 |
| ENSMUSG0000002501 | chr19 | Tll2 | -0.062163899 | 0.944584186 |
| ENSMUSG0000001842 | chr11 | Akap1 | -0.042474656 | 0.944584186 |
| ENSMUSG0000004456 | chr7 | Rasip1 | -0.070421995 | 0.944584186 |
| ENSMUSG0000001547 | chr17 | Ppt2 | -0.043028557 | 0.944584186 |
| ENSMUSG0000000274 | chr5 | Baz1b | -0.034614612 | 0.944584186 |
| ENSMUSG0000003732 | chr17 | Capn15 | -0.035246274 | 0.944584186 |
| ENSMUSG0000003825 | chr3 | Bcl9 | -0.036488054 | 0.944584186 |
| ENSMUSG0000004835 | chr4 | Coa7 | -0.04106262 | 0.944584186 |
| ENSMUSG0000002733 | chr2 | Rassf2 | -0.03923388 | 0.944584186 |
| ENSMUSG0000002801 | chr3 | Gar1 | -0.04054362 | 0.944584186 |
| ENSMUSG0000001771 | chr11 | Tha1 | -0.077053003 | 0.944584186 |
| ENSMUSG0000000182 | chr7 | Clpb | -0.031948552 | 0.944584186 |
| ENSMUSG0000002842 | chr4 | Nfx1 | -0.026494893 | 0.944584186 |
| ENSMUSG0000002011 | chr10 | Micu1 | -0.024234677 | 0.944584186 |
| ENSMUSG0000002077 | chr11 | Fbf1 | -0.028111116 | 0.944584186 |
| ENSMUSG0000004775 | chrX | Fancb | -0.084543214 | 0.944584186 |
| ENSMUSG0000002259 | chr15 | Slurp1 | 0.293783541 | 0.944584186 |
| ENSMUSG0000002300 | chr15 | Dhh | 0.125981545 | 0.944584186 |
| ENSMUSG0000002824 | chr4 | Coq3 | 0.050636931 | 0.944584186 |
| ENSMUSG0000002749 | chr2 | Fam210b | 0.046614498 | 0.944584186 |
| ENSMUSG0000005062 | chr9 | Gpd1l | 0.027227923 | 0.944584186 |
| ENSMUSG0000004184 | chr15 | Card6 | 0.043066646 | 0.944584186 |
| ENSMUSG0000003639 | chr11 | Sec24a | 0.039433216 | 0.944584186 |
| ENSMUSG0000005721 | chr11 | Armc7 | 0.053830409 | 0.944584186 |
| ENSMUSG0000003529 | chr12 | G2e3 | 0.044128082 | 0.944584186 |
| ENSMUSG0000003302 | chr15 | 1700088E04Rik | 0.061273404 | 0.944584186 |
| ENSMUSG0000002678 | chr2 | Zbtb43 | 0.034025078 | 0.944584186 |
| ENSMUSG0000002941 | chr5 | Sdad1 | 0.045026633 | 0.944584186 |
| ENSMUSG0000002161 | chr13 | Xrcc4 | 0.049436894 | 0.944584186 |
| ENSMUSG0000003954 | chr4 | Ajap1 | 0.066076991 | 0.944584186 |
| ENSMUSG0000004163 | chr5 | Gcn1 | 0.055571083 | 0.944584186 |
| ENSMUSG0000002693 | chr2 | Fcna | 0.074891082 | 0.944584186 |
| ENSMUSG0000004565 | chr14 | Fam216b | -0.230700719 | 0.944684395 |
| ENSMUSG0000001477 | chr8 | Fhod1 | -0.049915984 | 0.944684395 |
| ENSMUSG0000005472 | chr13 | Phactr1 | -0.048613349 | 0.944684395 |
| ENSMUSG0000003192 | chr9 | Mre11a | -0.027892246 | 0.944684395 |
| ENSMUSG0000002643 | chr1 | Elk4 | -0.052512087 | 0.944891806 |
| ENSMUSG0000002456 | chr18 | Smad2 | -0.033899276 | 0.944917372 |
| ENSMUSG0000005943 | chr11 | Bcas3 | 0.053469397 | 0.944917372 |
| ENSMUSG0000002558 | chr11 | Rptor | 0.031551409 | 0.944917372 |
| ENSMUSG0000002708 | chr2 | Fastkd1 | 0.035022461 | 0.944917372 |
| ENSMUSG0000006792 | chrX | Rtl8b | -0.035977882 | 0.945137745 |
| ENSMUSG0000003027 | chr6 | Cidec | -0.152646389 | 0.945239042 |
| ENSMUSG0000000758 | chr14 | Tinf2 | -0.047653472 | 0.945239042 |
| ENSMUSG0000003379 | chr1 | Atp6v1h | 0.030554886 | 0.945552849 |
| ENSMUSG0000003319 | chr4 | Tie1 | 0.049113384 | 0.945793157 |

| | | | | |
|---|---|---|---|---|
| ENSMUSG0000004407 | chr10 | Tafa2 | -0.042676603 | 0.945861546 |
| ENSMUSG0000002936 | chr5 | Dck | -0.038111373 | 0.946021216 |
| ENSMUSG0000004116 | chr11 | Zmiz2 | 0.03560515 | 0.946021216 |
| ENSMUSG0000004319 | chr13 | Rfesd | 0.047111256 | 0.946109132 |
| ENSMUSG0000004275 | chr1 | Nmnat2 | 0.044784905 | 0.946141789 |
| ENSMUSG0000003699 | chr2 | Nxt1 | -0.070566033 | 0.9463549 |
| ENSMUSG0000003780 | chr9 | Fam76b | -0.038138167 | 0.946396649 |
| ENSMUSG0000002808 | chr3 | Chd1l | -0.035749732 | 0.946396649 |
| ENSMUSG0000003315 | chr16 | Abhd10 | -0.028455334 | 0.946396649 |
| ENSMUSG0000002523 | chr19 | Sufu | 0.030036682 | 0.946437071 |
| ENSMUSG0000004877 | chr7 | P2ry6 | 0.043837662 | 0.946437071 |
| ENSMUSG0000002745 | chr2 | Gins1 | -0.113932474 | 0.9464922 |
| ENSMUSG0000002879 | chr4 | Khdrbs1 | -0.019593265 | 0.946594283 |
| ENSMUSG0000005361 | chr19 | Sh3pxd2a | 0.032525764 | 0.946594324 |
| ENSMUSG0000006092 | chr11 | Acyp2 | -0.049008813 | 0.946607752 |
| ENSMUSG0000000783 | chr13 | Hnrnpa0 | 0.038955216 | 0.946706017 |
| ENSMUSG0000002752 | chr2 | Phactr3 | -0.063136914 | 0.946940297 |
| ENSMUSG0000004275 | chr9 | Tmem108 | -0.054401583 | 0.946940297 |
| ENSMUSG0000003390 | chr11 | Usp36 | 0.044171729 | 0.946940297 |
| ENSMUSG0000001998 | chr10 | Enpp3 | 0.053093723 | 0.947032966 |
| ENSMUSG0000003206 | chr9 | Pts | -0.040466752 | 0.947048649 |
| ENSMUSG0000009450 | chr8 | Smim18 | -0.148786153 | 0.947048649 |
| ENSMUSG0000004430 | chr2 | Ubr3 | -0.035451728 | 0.947048649 |
| ENSMUSG0000002857 | chr4 | Fggy | -0.043869816 | 0.947048649 |
| ENSMUSG0000006793 | chr17 | Zfp948 | -0.039519236 | 0.947048649 |
| ENSMUSG0000007532 | chr10 | Zbtb2 | -0.035489844 | 0.947048649 |
| ENSMUSG0000002111 | chr12 | Snapc1 | -0.036303669 | 0.947048649 |
| ENSMUSG0000003154 | chr8 | Gpat4 | 0.032572272 | 0.947048649 |
| ENSMUSG0000000907 | chr11 | Nf2 | 0.023020762 | 0.947048649 |
| ENSMUSG0000002830 | chr4 | C9orf72 | 0.029907384 | 0.947048649 |
| ENSMUSG0000009890 | chr13 | Zfp953 | 0.064366954 | 0.947048649 |
| ENSMUSG0000003575 | chr15 | Selenoo | 0.034960569 | 0.947048649 |
| ENSMUSG0000001512 | chr17 | Unkl | 0.045268626 | 0.947048649 |
| ENSMUSG0000000761 | chr8 | Gtpbp3 | 0.03181556 | 0.947137235 |
| ENSMUSG0000002967 | chr5 | Limk1 | 0.045055789 | 0.947147994 |
| ENSMUSG0000003260 | chr9 | Prkar2a | 0.028739023 | 0.947232648 |
| ENSMUSG0000002578 | chr2 | Itih5 | 0.040758462 | 0.947685047 |
| ENSMUSG0000002875 | chr4 | Pink1 | 0.041558328 | 0.947685047 |
| ENSMUSG0000003903 | chr10 | Arhgap18 | -0.081970593 | 0.947710816 |
| ENSMUSG0000002812 | chr3 | Bcar3 | -0.044225536 | 0.947710816 |
| ENSMUSG0000004873 | chr11 | Klhl11 | -0.041884078 | 0.947710816 |
| ENSMUSG0000001327 | chr1 | Slc41a1 | -0.035317341 | 0.947710816 |
| ENSMUSG0000009164 | chr3 | C2cd4d | -0.124676637 | 0.947710816 |
| ENSMUSG0000002437 | chr18 | Tslp | -0.070075771 | 0.947710816 |
| ENSMUSG0000000097 | chr11 | Heatr6 | -0.025875189 | 0.947710816 |
| ENSMUSG0000006188 | chr4 | Ssbp3 | -0.024674968 | 0.947710816 |
| ENSMUSG0000001800 | chr15 | Cyth4 | -0.051637144 | 0.947710816 |
| ENSMUSG0000003473 | chr14 | Dgkh | 0.049028622 | 0.947710816 |
| ENSMUSG0000003153 | chr8 | Saraf | 0.044079173 | 0.947710816 |
| ENSMUSG0000002639 | chr1 | Nek7 | 0.023875781 | 0.947710816 |

| | | | | |
|---|---|---|---|---|
| ENSMUSG0000006430 | chr1 | Clasp1 | 0.031961486 | 0.947710816 |
| ENSMUSG0000003686 | chr9 | Smad6 | 0.050447606 | 0.947710816 |
| ENSMUSG0000001383 | chr10 | Med16 | 0.038990419 | 0.947710816 |
| ENSMUSG0000000623 | chr9 | Epor | 0.100807845 | 0.947710816 |
| ENSMUSG0000000182 | chr2 | Hoxd12 | 0.179959285 | 0.947710816 |
| ENSMUSG0000002950 | chr5 | Ankle2 | 0.027326857 | 0.947710816 |
| ENSMUSG0000003739 | chr1 | Rcor3 | 0.043358605 | 0.947710816 |
| ENSMUSG0000002585 | chr5 | Pdgfa | 0.030063536 | 0.947710816 |
| ENSMUSG0000001984 | chr10 | Traf3ip2 | 0.02923669 | 0.947710816 |
| ENSMUSG0000005723 | chr6 | Aak1 | 0.033473381 | 0.947710816 |
| ENSMUSG0000007865 | chr11 | Aoc2 | 0.047936139 | 0.947710816 |
| ENSMUSG0000002819 | chr3 | Cryz | 0.033025044 | 0.947710816 |
| ENSMUSG0000004464 | chr18 | Zbtb7c | 0.056316076 | 0.947710816 |
| ENSMUSG0000002241 | chr15 | Tab1 | 0.02874382 | 0.947710816 |
| ENSMUSG0000007258 | chr11 | Ptrh2 | 0.040338646 | 0.947710816 |
| ENSMUSG0000012195 | chrX | ENSMUSG000001 | 0.088083455 | 0.947710816 |
| ENSMUSG0000007341 | chr17 | Lst1 | 0.10210605 | 0.947710816 |
| ENSMUSG0000002884 | chr4 | Gpn2 | 0.051769477 | 0.947710816 |
| ENSMUSG0000002288 | chr16 | Rfc4 | 0.073614653 | 0.947710816 |
| ENSMUSG0000007171 | chr15 | Csf2rb | -0.07206472 | 0.947720011 |
| ENSMUSG0000002069 | chr11 | Mettl2 | -0.035575018 | 0.947720011 |
| ENSMUSG0000003240 | chr9 | Zwilch | 0.093309805 | 0.947720011 |
| ENSMUSG0000002514 | chr11 | Myadml2 | -0.194568995 | 0.947842315 |
| ENSMUSG0000002935 | chr5 | Crybb3 | -0.119578799 | 0.947842315 |
| ENSMUSG0000002425 | chr17 | Abcg8 | -0.178859027 | 0.947842315 |
| ENSMUSG0000005112 | chr6 | Gimap9 | -0.113405578 | 0.947893506 |
| ENSMUSG0000003600 | chr10 | Mettl25 | 0.047190672 | 0.947927033 |
| ENSMUSG0000003620 | chr16 | Nepro | 0.048287723 | 0.948008273 |
| ENSMUSG0000000601 | chr7 | Dhx34 | -0.03779694 | 0.948039599 |
| ENSMUSG0000004318 | chr13 | Simc1 | -0.040273534 | 0.948039599 |
| ENSMUSG0000002728 | chr2 | Cdan1 | -0.033965068 | 0.948039599 |
| ENSMUSG0000007559 | chr11 | Zfp652 | -0.034294682 | 0.948039599 |
| ENSMUSG0000003884 | chr19 | Gcnt1 | -0.087186483 | 0.948039599 |
| ENSMUSG0000004823 | chr4 | Fbxo10 | 0.02236445 | 0.948039599 |
| ENSMUSG0000002304 | chr15 | Csad | 0.028182007 | 0.948039599 |
| ENSMUSG0000004723 | chrX | Mageh1 | -0.038321506 | 0.948040229 |
| ENSMUSG0000003156 | chr8 | Wwc2 | -0.026187027 | 0.948040229 |
| ENSMUSG0000005392 | chr15 | Zftraf1 | -0.033770603 | 0.948040229 |
| ENSMUSG0000002400 | chr17 | Stk38 | 0.035992126 | 0.948040229 |
| ENSMUSG0000005918 | chr6 | Tafa1 | 0.048018279 | 0.948040229 |
| ENSMUSG0000005476 | chr14 | Defb42 | 0.121888122 | 0.948040229 |
| ENSMUSG0000001892 | chr11 | Alox15 | 0.10880172 | 0.948040229 |
| ENSMUSG0000000446 | chr16 | Tbccd1 | -0.032139685 | 0.948046806 |
| ENSMUSG0000003415 | chr12 | Cipc | -0.043563798 | 0.948046806 |
| ENSMUSG0000006037 | chr14 | Hnrnpc | 0.050064041 | 0.948046806 |
| ENSMUSG0000003323 | chr15 | Arid2 | 0.046278539 | 0.948046806 |
| ENSMUSG0000002110 | chr12 | Prkch | -0.064271734 | 0.948071074 |
| ENSMUSG0000004066 | chr11 | Nup88 | 0.029412286 | 0.948171823 |
| ENSMUSG0000000953 | chr18 | Rnmt | 0.030326531 | 0.948359989 |
| ENSMUSG0000003468 | chr13 | Prr7 | 0.052359657 | 0.948535352 |

| | | | | |
|---|---|---|---|---|
| ENSMUSG0000002209 | chr14 | Fhip2b | -0.047002507 | 0.948580517 |
| ENSMUSG0000003702 | chr7 | Wdr62 | -0.106590153 | 0.948720171 |
| ENSMUSG0000001858 | chr12 | Dnah11 | -0.08324131 | 0.948720171 |
| ENSMUSG0000002590 | chr1 | Sox17 | -0.088716037 | 0.948720171 |
| ENSMUSG0000001186 | chr8 | Tusc3 | -0.038920618 | 0.948720171 |
| ENSMUSG0000002800 | chr3 | Gucy1b1 | 0.030419267 | 0.948720171 |
| ENSMUSG0000004708 | chr7 | Ngrn | 0.030849416 | 0.948720171 |
| ENSMUSG0000007000 | chr8 | Ssbp4 | 0.030941861 | 0.948720171 |
| ENSMUSG0000007570 | chr12 | Dio3 | 0.138958348 | 0.948720171 |
| ENSMUSG0000006891 | chr3 | Clk2 | 0.03343222 | 0.948805202 |
| ENSMUSG0000004943 | chr16 | Upk1b | 0.149537246 | 0.948815681 |
| ENSMUSG0000003790 | chr11 | Ankrd13b | 0.033404487 | 0.948815681 |
| ENSMUSG0000007903 | chr12 | Alkbh1 | 0.040466584 | 0.948815681 |
| ENSMUSG0000009543 | chr13 | Zfp748 | -0.035455238 | 0.949006386 |
| ENSMUSG0000004464 | chr2 | Csrnp3 | -0.044062956 | 0.949056156 |
| ENSMUSG0000005863 | chr7 | Zfp110 | -0.034762482 | 0.949056156 |
| ENSMUSG0000004029 | chr1 | Suco | -0.031376962 | 0.949112295 |
| ENSMUSG0000004134 | chr7 | Ankrd42 | -0.059415104 | 0.949115221 |
| ENSMUSG0000003050 | chr7 | Asb7 | 0.03071951 | 0.949119369 |
| ENSMUSG0000002028 | chr11 | 1700093K21Rik | 0.305153806 | 0.949199412 |
| ENSMUSG0000010064 | chr2 | Gm28230 | -0.080238545 | 0.949199412 |
| ENSMUSG0000002882 | chr4 | Syf2 | 0.036532355 | 0.949199412 |
| ENSMUSG0000004734 | chr1 | Mettl21c | 0.180705266 | 0.949199412 |
| ENSMUSG0000005328 | chr1 | Trmt1l | 0.029637757 | 0.949199412 |
| ENSMUSG0000002437 | chr17 | C2 | -0.106195874 | 0.949205457 |
| ENSMUSG0000003129 | chrX | Phka2 | -0.061368846 | 0.949205457 |
| ENSMUSG0000002759 | chr2 | Ahcy | -0.067447645 | 0.949205457 |
| ENSMUSG0000005111 | chr7 | Garin5a | -0.080793687 | 0.949205457 |
| ENSMUSG0000002065 | chr12 | Grhl1 | -0.084497546 | 0.949205457 |
| ENSMUSG0000004037 | chr4 | Gpr63 | -0.063933134 | 0.949205457 |
| ENSMUSG0000002177 | chr14 | Samd8 | -0.037273554 | 0.949205457 |
| ENSMUSG0000007480 | chr10 | Gas2l3 | 0.118173328 | 0.949205457 |
| ENSMUSG0000008984 | chr7 | Timm10b | -0.049721508 | 0.949205457 |
| ENSMUSG0000003564 | chr4 | Zcchc7 | -0.033139415 | 0.949205457 |
| ENSMUSG0000003489 | chr13 | Sncb | -0.053332565 | 0.949205457 |
| ENSMUSG0000003217 | chr9 | Ilf3 | -0.022475292 | 0.949205457 |
| ENSMUSG0000005586 | chr1 | Per2 | 0.239547344 | 0.949205457 |
| ENSMUSG0000011633 | chr15 | Krt85 | 0.190465794 | 0.949205457 |
| ENSMUSG0000003895 | chr9 | Edc3 | 0.025397252 | 0.949205457 |
| ENSMUSG0000000201 | chrX | Pnck | 0.032583724 | 0.949205457 |
| ENSMUSG0000003067 | chr7 | Mylpf | 0.112876643 | 0.949205457 |
| ENSMUSG0000001207 | chr12 | Brms1l | 0.024051468 | 0.949205457 |
| ENSMUSG0000004126 | chr5 | Uspl1 | 0.031418182 | 0.949205457 |
| ENSMUSG0000007537 | chr2 | Rc3h2 | 0.039615985 | 0.949205457 |
| ENSMUSG0000003258 | chr9 | Mon1a | 0.028865496 | 0.949205457 |
| ENSMUSG0000003436 | chr8 | Cpne2 | 0.036688078 | 0.949205457 |
| ENSMUSG0000003479 | chr8 | Cpne7 | 0.080913129 | 0.949205457 |
| ENSMUSG0000012158 | chr15 | Tmem276 | 0.072579445 | 0.949205457 |
| ENSMUSG0000002416 | chr17 | C3 | -0.140220329 | 0.94942776 |
| ENSMUSG0000007087 | chr7 | Lilra5 | -0.105859797 | 0.94942776 |

| | | | | |
|---|---|---|---|---|
| ENSMUSG0000007928 | chr1 | 2310009B15Rik | -0.052226567 | 0.94942776 |
| ENSMUSG0000000296 | chr7 | Med25 | -0.022491717 | 0.94942776 |
| ENSMUSG0000002062 | chr12 | Rdh14 | 0.044725227 | 0.94942776 |
| ENSMUSG0000002895 | chr4 | Tas1r1 | 0.121093184 | 0.94942776 |
| ENSMUSG0000000067 | chr17 | Haao | 0.153215273 | 0.94942776 |
| ENSMUSG0000002963 | chr5 | Wasf3 | 0.033180203 | 0.949440201 |
| ENSMUSG0000007886 | chr2 | Gm14326 | 0.037389126 | 0.949546302 |
| ENSMUSG0000005239 | chr17 | Ezr | -0.061407217 | 0.949564033 |
| ENSMUSG0000007277 | chr6 | Acrbp | -0.074650497 | 0.949564033 |
| ENSMUSG0000003206 | chr9 | 2310030G06Rik | -0.063519206 | 0.949564033 |
| ENSMUSG0000002618 | chr16 | Ppm1f | -0.035261037 | 0.949564033 |
| ENSMUSG0000001775 | chr2 | Pltp | -0.039356463 | 0.949564033 |
| ENSMUSG0000003687 | chr8 | Phkb | -0.019615126 | 0.949564033 |
| ENSMUSG0000006355 | chr7 | Nup98 | 0.031227919 | 0.949564033 |
| ENSMUSG0000008615 | chr9 | Pierce2 | 0.042525979 | 0.949564033 |
| ENSMUSG0000003690 | chr4 | C1qb | 0.051302935 | 0.949564033 |
| ENSMUSG0000002955 | chr6 | Tfec | 0.198882201 | 0.949564033 |
| ENSMUSG0000002541 | chr4 | Ttc4 | -0.024119233 | 0.949601857 |
| ENSMUSG0000002230 | chr15 | Oxr1 | -0.023466253 | 0.949601857 |
| ENSMUSG0000003164 | chr8 | Mfap3l | 0.051249306 | 0.949601857 |
| ENSMUSG0000003007 | chr6 | Gxylt2 | 0.101940082 | 0.949601857 |
| ENSMUSG0000008761 | chr3 | Gm16253 | 0.105474557 | 0.949601857 |
| ENSMUSG0000004234 | chr13 | Arl15 | 0.029623444 | 0.949601857 |
| ENSMUSG0000002194 | chr14 | Cryl1 | 0.046460075 | 0.949601857 |
| ENSMUSG0000002473 | chr19 | Prpf19 | 0.032475592 | 0.949683811 |
| ENSMUSG0000002537 | chr11 | Chmp6 | -0.031335681 | 0.949803547 |
| ENSMUSG0000005927 | chr7 | Zc3h4 | 0.036448727 | 0.949814937 |
| ENSMUSG0000003967 | chr15 | Capsl | -0.18107417 | 0.94989235 |
| ENSMUSG0000003659 | chr5 | Cpz | 0.175429782 | 0.949909351 |
| ENSMUSG0000000537 | chr17 | Msh6 | -0.033125024 | 0.949919829 |
| ENSMUSG0000002621 | chr1 | Trip12 | -0.029870846 | 0.949936054 |
| ENSMUSG0000006792 | chr17 | Zfp760 | -0.033380511 | 0.949936054 |
| ENSMUSG0000004259 | chr13 | Ipo11 | 0.028464512 | 0.949936054 |
| ENSMUSG0000003403 | chr2 | Rpap1 | 0.036931111 | 0.949936054 |
| ENSMUSG0000003407 | chr7 | Zfp551 | 0.059120557 | 0.949936054 |
| ENSMUSG0000000857 | chr4 | Nfib | 0.042900442 | 0.949958497 |
| ENSMUSG0000002706 | chr2 | Ssrp1 | 0.023109996 | 0.95022399 |
| ENSMUSG0000003868 | chr2 | Rtel1 | -0.046148388 | 0.950340532 |
| ENSMUSG0000002907 | chr4 | Sdf4 | 0.021856166 | 0.950340532 |
| ENSMUSG0000001844 | chr11 | Derl2 | 0.030352255 | 0.950340532 |
| ENSMUSG0000002777 | chr3 | Schip1 | 0.071202718 | 0.950340532 |
| ENSMUSG0000008579 | chr12 | Lin52 | 0.037839364 | 0.950342824 |
| ENSMUSG0000003484 | chr2 | Ttc21b | 0.039959539 | 0.950342824 |
| ENSMUSG0000002303 | chr15 | Scn8a | 0.070508703 | 0.950479307 |
| ENSMUSG0000003714 | chr2 | Cfap61 | 0.097964534 | 0.950479307 |
| ENSMUSG0000002560 | chr6 | Podxl | 0.041757211 | 0.950479307 |
| ENSMUSG0000003838 | chr5 | Setd1b | -0.05278662 | 0.950605433 |
| ENSMUSG0000002304 | chr15 | Soat2 | -0.211981249 | 0.950605433 |
| ENSMUSG0000005149 | chr5 | Tlr6 | -0.115449125 | 0.950605433 |
| ENSMUSG0000002861 | chr4 | Lrp8 | -0.063711276 | 0.950605433 |

| | | | | |
|---|---|---|---|---|
| ENSMUSG0000006708 | chr1 | Asb18 | 0.199843362 | 0.950605433 |
| ENSMUSG0000002435 | chr18 | Spata24 | -0.107419227 | 0.950605433 |
| ENSMUSG0000004541 | chr13 | Akr1e1 | -0.041940239 | 0.950605433 |
| ENSMUSG0000001035 | chr11 | Ifi35 | -0.048447024 | 0.950605433 |
| ENSMUSG0000003736 | chr4 | Pafah2 | -0.042075025 | 0.950605433 |
| ENSMUSG0000000311 | chr11 | Cdk12 | -0.044200865 | 0.950605433 |
| ENSMUSG0000003110 | chrX | Rbmx2 | -0.054940544 | 0.950605433 |
| ENSMUSG0000007060 | chr7 | Vsig10l | -0.043048704 | 0.950605433 |
| ENSMUSG0000004141 | chr12 | Dicer1 | -0.034728003 | 0.950605433 |
| ENSMUSG0000002854 | chr4 | Bend5 | -0.035577261 | 0.950605433 |
| ENSMUSG0000003479 | chr14 | Ccdc122 | -0.071138338 | 0.950605433 |
| ENSMUSG0000002777 | chr3 | Mfsd1 | -0.03461755 | 0.950605433 |
| ENSMUSG0000006813 | chr2 | Zfp120 | -0.033987282 | 0.950605433 |
| ENSMUSG0000002226 | chr15 | Dnah5 | -0.074391438 | 0.950605433 |
| ENSMUSG0000006974 | chr17 | Zfp820 | -0.065963339 | 0.950605433 |
| ENSMUSG0000003275 | chr6 | Bet1 | -0.031648967 | 0.950605433 |
| ENSMUSG0000001159 | chr15 | Vps28 | -0.027562851 | 0.950605433 |
| ENSMUSG0000002824 | chr4 | Nsmaf | -0.026899161 | 0.950605433 |
| ENSMUSG0000004165 | chr14 | Pcca | -0.022224087 | 0.950605433 |
| ENSMUSG0000007092 | chr4 | Klhl9 | -0.02152679 | 0.950605433 |
| ENSMUSG0000001890 | chr7 | Arrb1 | -0.026943816 | 0.950605433 |
| ENSMUSG0000002864 | chr4 | Tmem269 | -0.154308579 | 0.950605433 |
| ENSMUSG0000003241 | chr9 | Atp1b3 | 0.038852538 | 0.950605433 |
| ENSMUSG0000002397 | chr17 | Ubr2 | 0.039522485 | 0.950605433 |
| ENSMUSG0000000191 | chr8 | Nfix | 0.037975239 | 0.950605433 |
| ENSMUSG0000009172 | chr14 | Siah3 | 0.122318442 | 0.950605433 |
| ENSMUSG0000006079 | chr3 | Intu | 0.028620618 | 0.950605433 |
| ENSMUSG0000003781 | chr3 | Abhd18 | 0.034811275 | 0.950605433 |
| ENSMUSG0000003137 | chrX | Abcd1 | 0.03004057 | 0.950605433 |
| ENSMUSG0000003661 | chr9 | Eepd1 | 0.037046679 | 0.950605433 |
| ENSMUSG0000002603 | chr1 | Clk1 | 0.028401593 | 0.950605433 |
| ENSMUSG0000003395 | chr2 | Tnks1bp1 | 0.028658155 | 0.950605433 |
| ENSMUSG0000000199 | chr8 | Sipa1l2 | 0.03225617 | 0.950605433 |
| ENSMUSG0000003071 | chr7 | Sgf29 | 0.040677045 | 0.950605433 |
| ENSMUSG0000003039 | chr7 | Mark4 | 0.034907741 | 0.950605433 |
| ENSMUSG0000004810 | chr3 | Taf13 | 0.037512531 | 0.950605433 |
| ENSMUSG0000000508 | chr2 | Cd44 | 0.048627142 | 0.950605433 |
| ENSMUSG0000002235 | chr15 | Mtss1 | 0.027707275 | 0.950605433 |
| ENSMUSG0000003728 | chr9 | Stag1 | 0.031173856 | 0.950605433 |
| ENSMUSG0000002504 | chr8 | Msr1 | 0.064943837 | 0.950605433 |
| ENSMUSG0000003953 | chr10 | Zup1 | 0.035610812 | 0.950605433 |
| ENSMUSG0000002870 | chr4 | Nsun4 | 0.038049588 | 0.950605433 |
| ENSMUSG0000002236 | chr15 | Tbc1d31 | 0.038051756 | 0.950605433 |
| ENSMUSG0000003021 | chr6 | Art4 | 0.07689138 | 0.950605433 |
| ENSMUSG0000002109 | chr12 | Serpina3n | 0.156048439 | 0.950605433 |
| ENSMUSG0000003198 | chr8 | Agt | 0.089095584 | 0.950605433 |
| ENSMUSG0000002689 | chr2 | Morn5 | 0.145913484 | 0.950615774 |
| ENSMUSG0000008976 | chr2 | Ier5l | 0.036292013 | 0.950615774 |
| ENSMUSG0000002111 | chr12 | Vrk1 | 0.049583053 | 0.950703219 |
| ENSMUSG0000004832 | chr2 | Ckap2l | 0.139057987 | 0.950769827 |

| | | | | |
|---|---|---|---|---|
| ENSMUSG0000002675 | chr2 | Ppp6c | 0.020218834 | 0.950769827 |
| ENSMUSG0000004987 | chr16 | Nlrc3 | -0.078462317 | 0.95081581 |
| ENSMUSG0000002288 | chr16 | Robo1 | -0.034288094 | 0.95081581 |
| ENSMUSG0000003792 | chr3 | Bank1 | -0.08652628 | 0.950823602 |
| ENSMUSG0000004119 | chr4 | Pla2g5 | -0.052367509 | 0.950924346 |
| ENSMUSG0000007298 | chr2 | Oip5 | 0.126613103 | 0.950924346 |
| ENSMUSG0000002486 | chr19 | Gm49405 | 0.105674453 | 0.95109542 |
| ENSMUSG0000001996 | chr10 | Dusp6 | -0.047173004 | 0.951102209 |
| ENSMUSG0000003076 | chr7 | Myo7a | -0.044991608 | 0.951203323 |
| ENSMUSG0000002735 | chr2 | Spred1 | -0.019768761 | 0.951270637 |
| ENSMUSG0000003928 | chr9 | Azi2 | 0.0246359 | 0.951270637 |
| ENSMUSG0000002323 | chr2 | Scg5 | -0.031893264 | 0.951484546 |
| ENSMUSG0000002519 | chr19 | Abcc2 | -0.111691254 | 0.951553385 |
| ENSMUSG0000003032 | chr6 | Mbd4 | -0.051386263 | 0.951553385 |
| ENSMUSG0000002932 | chr5 | Hnrnpdl | -0.033122185 | 0.951553385 |
| ENSMUSG0000004549 | chr18 | Pcdhb3 | 0.083749889 | 0.951553385 |
| ENSMUSG0000005210 | chr17 | Mtcl1 | 0.028327926 | 0.951553385 |
| ENSMUSG0000002137 | chr13 | Id4 | 0.044002802 | 0.951553385 |
| ENSMUSG0000002538 | chr11 | Fscn2 | 0.135861624 | 0.951553385 |
| ENSMUSG0000004045 | chr8 | Arglu1 | -0.038525491 | 0.951941312 |
| ENSMUSG0000005719 | chr9 | Slc44a2 | -0.040158966 | 0.951987777 |
| ENSMUSG0000000048 | chr15 | Pdgfb | -0.037992262 | 0.951991351 |
| ENSMUSG0000003689 | chr5 | Zfp157 | -0.028566791 | 0.951991351 |
| ENSMUSG0000006820 | chr2 | Macrod2 | -0.035009223 | 0.951991351 |
| ENSMUSG0000003948 | chr17 | Nrtn | 0.047179412 | 0.951991351 |
| ENSMUSG0000003242 | chr9 | Snx14 | 0.020474328 | 0.951991351 |
| ENSMUSG0000004874 | chr4 | E130114P18Rik | -0.061997592 | 0.951994431 |
| ENSMUSG0000005918 | chr9 | Mtfmt | 0.038883272 | 0.951994431 |
| ENSMUSG0000002673 | chr2 | Pip4k2a | 0.02256451 | 0.952284552 |
| ENSMUSG0000003666 | chr15 | Dennd3 | -0.049923605 | 0.952350729 |
| ENSMUSG0000002670 | chr1 | Cenpl | -0.061505468 | 0.952350729 |
| ENSMUSG0000004192 | chr1 | Rnpep | -0.032455097 | 0.952350729 |
| ENSMUSG0000003177 | chr8 | Cntnap4 | -0.127162512 | 0.952350729 |
| ENSMUSG0000002218 | chr14 | Slc7a8 | -0.034806761 | 0.952350729 |
| ENSMUSG0000004239 | chr3 | Gatad2b | -0.030816888 | 0.952350729 |
| ENSMUSG0000002672 | chr1 | Rabgap1l | -0.024055953 | 0.952350729 |
| ENSMUSG0000001534 | chr8 | Golga7 | -0.022026318 | 0.952350729 |
| ENSMUSG0000000101 | chr3 | Chtop | -0.021754601 | 0.952350729 |
| ENSMUSG0000009811 | chr15 | Bin2 | -0.077004513 | 0.952350729 |
| ENSMUSG0000003699 | chr4 | Asap3 | -0.036078885 | 0.952350729 |
| ENSMUSG0000003224 | chr9 | Anp32a | 0.032180613 | 0.952350729 |
| ENSMUSG0000003000 | chr6 | Dusp11 | 0.023830827 | 0.952350729 |
| ENSMUSG0000001899 | chr6 | Slc35b4 | 0.020101368 | 0.952350729 |
| ENSMUSG0000003908 | chr10 | L3mbtl3 | 0.02902173 | 0.952350729 |
| ENSMUSG0000002796 | chr3 | Extl2 | 0.033738155 | 0.952350729 |
| ENSMUSG0000002896 | chr4 | Errfi1 | 0.027669376 | 0.952350729 |
| ENSMUSG0000001694 | chr11 | Kctd2 | 0.030514635 | 0.952350729 |
| ENSMUSG0000003930 | chr14 | Ndst2 | 0.028540769 | 0.952350729 |
| ENSMUSG0000002208 | chr14 | Bin3 | 0.035720365 | 0.952350729 |
| ENSMUSG0000002485 | chr19 | Pold4 | 0.046155246 | 0.952350729 |

| | | | | |
|---|---|---|---|---|
| ENSMUSG0000003203 | chr9 | St3gal4 | 0.046433189 | 0.952350729 |
| ENSMUSG0000004183 | chr7 | Ptpre | 0.058148238 | 0.952627247 |
| ENSMUSG0000002118 | chr12 | Trip11 | -0.030654359 | 0.952647711 |
| ENSMUSG0000002397 | chr17 | Rrp36 | 0.04989401 | 0.952676231 |
| ENSMUSG0000002607 | chr1 | Il18r1 | 0.187141671 | 0.952909419 |
| ENSMUSG0000009867 | chr15 | Mroh6 | -0.176881373 | 0.952909419 |
| ENSMUSG0000002782 | chr3 | Kcnab1 | 0.059787531 | 0.952909419 |
| ENSMUSG0000002613 | chr1 | Dst | 0.035983336 | 0.952909419 |
| ENSMUSG0000004010 | chr6 | Klhl42 | 0.033640248 | 0.952909419 |
| ENSMUSG0000003895 | chr17 | Supt3 | 0.041430285 | 0.952909419 |
| ENSMUSG0000002253 | chr16 | Glyr1 | -0.01963806 | 0.952921254 |
| ENSMUSG0000001867 | chr11 | Sp2 | 0.031797527 | 0.952921254 |
| ENSMUSG0000009011 | chrX | Cmc4 | 0.061082932 | 0.953182779 |
| ENSMUSG0000006644 | chr12 | Zfyve26 | 0.042747798 | 0.953324306 |
| ENSMUSG0000002478 | chr19 | Ak3 | -0.026736699 | 0.953403483 |
| ENSMUSG0000002945 | chr5 | Acad10 | 0.066757867 | 0.953403483 |
| ENSMUSG0000007349 | chr1 | Ifi207 | 0.079169758 | 0.953403483 |
| ENSMUSG0000005302 | chr1 | Cntn2 | -0.059595422 | 0.95348322 |
| ENSMUSG0000007865 | chr11 | Psme3 | -0.020821223 | 0.95348322 |
| ENSMUSG0000002430 | chr17 | Myo1f | -0.07609389 | 0.95348322 |
| ENSMUSG0000003751 | chr7 | Ppfia1 | 0.022644454 | 0.95348322 |
| ENSMUSG0000002625 | chr1 | Eif4e2 | 0.024274538 | 0.95348322 |
| ENSMUSG0000001897 | chr5 | Sart3 | 0.025121573 | 0.95348322 |
| ENSMUSG0000003088 | chr7 | Arfip2 | 0.019035126 | 0.95348322 |
| ENSMUSG0000000747 | chr2 | Lrrc8a | 0.029571852 | 0.95348322 |
| ENSMUSG0000000667 | chrX | Pola1 | 0.031255078 | 0.95348322 |
| ENSMUSG0000005620 | chr8 | Pgpep1 | 0.023394111 | 0.95348322 |
| ENSMUSG0000004298 | chr5 | Upk3b | 0.122119516 | 0.95348322 |
| ENSMUSG0000005536 | chr8 | Slc6a2 | 0.11152457 | 0.95348322 |
| ENSMUSG0000003558 | chr17 | Kcnh8 | -0.07718733 | 0.953537547 |
| ENSMUSG0000000515 | chr8 | Wdr83 | -0.028275523 | 0.953537547 |
| ENSMUSG0000000612 | chr11 | Inpp5k | -0.048127144 | 0.953817195 |
| ENSMUSG0000007885 | chr11 | Igtp | 0.056953578 | 0.95392476 |
| ENSMUSG0000002591 | chr1 | Adhfe1 | -0.065389051 | 0.954039924 |
| ENSMUSG0000003055 | chr7 | Nr2f2 | -0.034506074 | 0.954039924 |
| ENSMUSG0000002069 | chr11 | Lig3 | -0.027656457 | 0.954039924 |
| ENSMUSG0000002796 | chr3 | Larp7 | 0.022760212 | 0.954039924 |
| ENSMUSG0000005533 | chr9 | Snupn | 0.040051628 | 0.954039924 |
| ENSMUSG0000006376 | chr3 | S100a7a | 0.077856929 | 0.954039924 |
| ENSMUSG0000004886 | chr1 | Arhgap30 | -0.058456999 | 0.954189334 |
| ENSMUSG0000002907 | chr4 | Cptp | -0.034442859 | 0.954380771 |
| ENSMUSG0000000889 | chr8 | Vdac3 | 0.037686613 | 0.95450832 |
| ENSMUSG0000003633 | chr12 | Kidins220 | 0.021778806 | 0.95450832 |
| ENSMUSG0000002422 | chr17 | Zfp523 | 0.021436819 | 0.95450832 |
| ENSMUSG0000002157 | chr13 | Tppp | 0.036666214 | 0.954605523 |
| ENSMUSG0000001417 | chr11 | Tvp23b | -0.04316477 | 0.954707565 |
| ENSMUSG0000006314 | chr14 | Kcnma1 | -0.045771464 | 0.954707565 |
| ENSMUSG0000001545 | chr17 | C4a | -0.157937091 | 0.954750448 |
| ENSMUSG0000000021 | chr7 | Th | -0.107176065 | 0.954750448 |
| ENSMUSG0000007093 | chr1 | Tgfbrap1 | -0.025028146 | 0.954965266 |

| | | | | |
|---|---|---|---|---|
| ENSMUSG0000002065 | chr12 | Cbll1 | -0.026429117 | 0.955111855 |
| ENSMUSG0000006745 | chr13 | H4c11 | -0.104571942 | 0.955180259 |
| ENSMUSG0000006274 | chr17 | Zfp677 | -0.037484192 | 0.955180259 |
| ENSMUSG0000003683 | chr3 | Plch1 | -0.078591579 | 0.955180259 |
| ENSMUSG0000006645 | chr9 | Hmgn3 | -0.034487683 | 0.955180259 |
| ENSMUSG0000003143 | chrX | Morc4 | 0.04300661 | 0.955180259 |
| ENSMUSG0000000594 | chr11 | Ctns | -0.042458062 | 0.955182452 |
| ENSMUSG0000000028 | chr11 | Mnt | -0.043269509 | 0.955226484 |
| ENSMUSG0000002794 | chr3 | Tpm3 | 0.02552683 | 0.955336828 |
| ENSMUSG0000001683 | chr14 | Tox4 | 0.020665057 | 0.955336828 |
| ENSMUSG0000007408 | chr4 | Snrnp40 | -0.029136357 | 0.955682183 |
| ENSMUSG0000003621 | chr17 | Polr1has | 0.045685012 | 0.955682183 |
| ENSMUSG0000002884 | chr4 | Tekt2 | -0.039199729 | 0.955696847 |
| ENSMUSG0000010331 | chr18 | Pcdha12 | 0.091837861 | 0.955696847 |
| ENSMUSG0000003303 | chr16 | Cip2a | 0.074335281 | 0.955696847 |
| ENSMUSG0000002893 | chr5 | Xrcc2 | 0.048237649 | 0.955696847 |
| ENSMUSG0000003067 | chr7 | Pde3b | 0.033026102 | 0.955696847 |
| ENSMUSG0000005309 | chr7 | Lins1 | -0.037709736 | 0.955711403 |
| ENSMUSG0000004020 | chr3 | Zfp704 | -0.043163892 | 0.955768365 |
| ENSMUSG0000002692 | chr2 | Sec16a | -0.025705287 | 0.955768365 |
| ENSMUSG0000005598 | chr1 | Irs1 | -0.035104739 | 0.955768365 |
| ENSMUSG0000004631 | chr12 | Stxbp6 | 0.019520397 | 0.955768365 |
| ENSMUSG0000000280 | chr12 | Btbd6 | 0.029080881 | 0.955768365 |
| ENSMUSG0000005649 | chr5 | Foxk1 | 0.030306291 | 0.9558101 |
| ENSMUSG0000002060 | chr12 | Lratd1 | -0.129823882 | 0.955923888 |
| ENSMUSG0000003278 | chr11 | Cntrob | -0.039938673 | 0.955923888 |
| ENSMUSG0000000327 | chr17 | Dlgap1 | -0.033649834 | 0.955923888 |
| ENSMUSG0000004668 | chr3 | Tifa | -0.047070043 | 0.955923888 |
| ENSMUSG0000009043 | chr10 | Gm17455 | 0.134935401 | 0.955923888 |
| ENSMUSG0000004599 | chr18 | Onecut2 | 0.09742699 | 0.955923888 |
| ENSMUSG0000003292 | chr14 | Itgbl1 | 0.076450895 | 0.955923888 |
| ENSMUSG0000003405 | chrX | Phka1 | 0.034590037 | 0.955923888 |
| ENSMUSG0000003667 | chr8 | Cenpt | 0.042539328 | 0.955923888 |
| ENSMUSG0000000703 | chr17 | Msh5 | 0.068284706 | 0.955923888 |
| ENSMUSG0000002620 | chr1 | Slc23a3 | 0.124860624 | 0.955923888 |
| ENSMUSG0000003961 | chr18 | Mocos | 0.045263689 | 0.955944674 |
| ENSMUSG0000003859 | chr10 | Cep85l | -0.03552688 | 0.95594665 |
| ENSMUSG0000004113 | chr5 | N4bp2l1 | 0.045183058 | 0.95594665 |
| ENSMUSG0000003049 | chr7 | Slc7a10 | -0.142504283 | 0.956106058 |
| ENSMUSG0000003949 | chr4 | Ccdc27 | -0.138100745 | 0.956106058 |
| ENSMUSG0000003137 | chrX | Bmx | -0.076179815 | 0.956106058 |
| ENSMUSG0000003606 | chr4 | Phf24 | -0.039694469 | 0.956106058 |
| ENSMUSG0000006294 | chr7 | Zfp696 | -0.041035313 | 0.956106058 |
| ENSMUSG0000007905 | chr8 | Cyp4v3 | -0.034999419 | 0.956106058 |
| ENSMUSG0000003698 | chr17 | Tfb1m | -0.050337137 | 0.956106058 |
| ENSMUSG0000004493 | chr10 | Ttc41 | -0.049317149 | 0.956106058 |
| ENSMUSG0000002150 | chr13 | Ddx46 | -0.023498303 | 0.956106058 |
| ENSMUSG0000003664 | chr11 | Tbc1d9b | -0.018613185 | 0.956106058 |
| ENSMUSG0000002189 | chr14 | Eaf1 | -0.019604288 | 0.956106058 |
| ENSMUSG0000002644 | chr1 | Lrrn2 | 0.051475023 | 0.956106058 |

| | | | | |
|---|---|---|---|---|
| ENSMUSG0000000903 | chr2 | Pdcl | 0.026411477 | 0.956106058 |
| ENSMUSG0000002217 | chr14 | Dad1 | 0.03737751 | 0.956106058 |
| ENSMUSG0000002143 | chr13 | Snrnp48 | 0.027233852 | 0.956106058 |
| ENSMUSG0000000102 | chr3 | S100a4 | 0.086280194 | 0.956106058 |
| ENSMUSG0000002518 | chr19 | R3hcc1l | 0.025887854 | 0.956106058 |
| ENSMUSG0000002719 | chr2 | Ttc17 | 0.02920608 | 0.956106058 |
| ENSMUSG0000009668 | chr10 | Mfsd4b4 | 0.03525312 | 0.956106058 |
| ENSMUSG0000007486 | chr13 | Platr25 | 0.037587766 | 0.956106058 |
| ENSMUSG0000005420 | chr1 | Ifi205 | 0.049602762 | 0.956106058 |
| ENSMUSG0000004605 | chr7 | Eid2 | 0.034504261 | 0.956106058 |
| ENSMUSG0000000255 | chr2 | Uck1 | -0.030155572 | 0.95620382 |
| ENSMUSG0000001435 | chr2 | Anapc1 | -0.023275532 | 0.956250367 |
| ENSMUSG0000002391 | chr17 | Flywch2 | -0.05737804 | 0.956250367 |
| ENSMUSG0000003825 | chr6 | Ncapd2 | -0.061543853 | 0.956306645 |
| ENSMUSG0000004260 | chr2 | Kcna4 | -0.069898461 | 0.956306645 |
| ENSMUSG0000000830 | chr5 | 1700109H08Rik | -0.062447326 | 0.956306645 |
| ENSMUSG0000003076 | chr7 | Slc5a11 | -0.144713167 | 0.956306645 |
| ENSMUSG0000009487 | chr13 | Zfp131 | -0.037291797 | 0.956306645 |
| ENSMUSG0000003596 | chrX | Ints6l | -0.041542251 | 0.956306645 |
| ENSMUSG0000004255 | chr4 | Adprs | -0.031343307 | 0.956306645 |
| ENSMUSG0000003790 | chr12 | Ankrd9 | -0.042683318 | 0.956306645 |
| ENSMUSG0000005461 | chr19 | Kdm2a | -0.027767136 | 0.956306645 |
| ENSMUSG0000001854 | chr11 | Pip4k2b | -0.026843377 | 0.956306645 |
| ENSMUSG0000009691 | chr17 | Zfp955b | -0.023605799 | 0.956306645 |
| ENSMUSG0000001977 | chr10 | Fbxo5 | -0.05329841 | 0.956306645 |
| ENSMUSG0000005554 | chr7 | Lair1 | -0.052033152 | 0.956306645 |
| ENSMUSG0000003240 | chr9 | Atr | -0.025282501 | 0.956306645 |
| ENSMUSG0000006276 | chr9 | Ei24 | -0.025549337 | 0.956306645 |
| ENSMUSG0000002252 | chr16 | Crebbp | -0.04387754 | 0.956306645 |
| ENSMUSG0000002535 | chr10 | Pmel | -0.074873796 | 0.956306645 |
| ENSMUSG0000002225 | chr15 | Mtdh | 0.018906778 | 0.956306645 |
| ENSMUSG0000007541 | chr2 | Dolk | 0.033985827 | 0.956306645 |
| ENSMUSG0000004761 | chr2 | Paxx | 0.035692901 | 0.956306645 |
| ENSMUSG0000005630 | chr3 | Sertm1 | 0.066657765 | 0.956306645 |
| ENSMUSG0000003081 | chr7 | Phkg2 | 0.023602135 | 0.956306645 |
| ENSMUSG0000003169 | chr8 | Tnpo2 | 0.025725389 | 0.956306645 |
| ENSMUSG0000004675 | chr14 | Ccdc66 | 0.02752699 | 0.956306645 |
| ENSMUSG0000004315 | chr14 | Arl11 | 0.121639008 | 0.956306645 |
| ENSMUSG0000004644 | chr8 | Cmtr2 | 0.036770603 | 0.956306645 |
| ENSMUSG0000000018 | chr6 | Ccnd2 | 0.03397275 | 0.956306645 |
| ENSMUSG0000003803 | chr16 | Socs1 | -0.099721472 | 0.956306645 |
| ENSMUSG0000004041 | chr19 | Slc25a28 | 0.027375657 | 0.956306645 |
| ENSMUSG0000002064 | chr12 | Rrm2 | 0.099962304 | 0.956306645 |
| ENSMUSG0000003602 | chrX | Pabir2 | 0.039298126 | 0.956306645 |
| ENSMUSG0000000562 | chr3 | Tmod4 | 0.148775807 | 0.956306645 |
| ENSMUSG0000002382 | chr17 | Prkn | 0.057340406 | 0.956306645 |
| ENSMUSG0000000361 | chr3 | Cp | 0.04837388 | 0.956306645 |
| ENSMUSG0000004519 | chr10 | Cirbp | 0.050922037 | 0.956306645 |
| ENSMUSG0000000394 | chr11 | Hlf | 0.144622222 | 0.956306645 |
| ENSMUSG0000007126 | chr17 | Zfp942 | -0.030882856 | 0.956315961 |

| | | | | |
|---|---|---|---|---|
| ENSMUSG0000004142 | chr7 | Meis3 | -0.025235231 | 0.956426711 |
| ENSMUSG0000003866 | chr9 | Herc1 | -0.035349873 | 0.956562878 |
| ENSMUSG0000002059 | chr12 | Pum2 | 0.019459476 | 0.956577356 |
| ENSMUSG0000005947 | chr9 | Zfp426 | 0.022284899 | 0.957119067 |
| ENSMUSG0000003083 | chr7 | Sergef | -0.046189089 | 0.957233774 |
| ENSMUSG0000000208 | chr7 | Bbc3 | -0.039374341 | 0.957233774 |
| ENSMUSG0000003020 | chr6 | Dusp16 | -0.036663456 | 0.957233774 |
| ENSMUSG0000001986 | chr10 | Gopc | -0.02077141 | 0.957233774 |
| ENSMUSG0000002256 | chr15 | Scrib | -0.017708466 | 0.957233774 |
| ENSMUSG0000000041 | chr6 | Cttnbp2 | -0.048699621 | 0.957233774 |
| ENSMUSG0000002059 | chr12 | Lpin1 | 0.035693432 | 0.957233774 |
| ENSMUSG0000002754 | chr2 | Atp9a | 0.026993996 | 0.957233774 |
| ENSMUSG0000004973 | chr7 | Zfp646 | 0.026524341 | 0.957233774 |
| ENSMUSG0000003804 | chr4 | Haus6 | 0.046238726 | 0.957233774 |
| ENSMUSG0000004405 | chr11 | Efcab9 | 0.063962239 | 0.957233774 |
| ENSMUSG0000006832 | chr6 | Htra2 | 0.037475478 | 0.957233774 |
| ENSMUSG0000003688 | chr4 | C1qa | 0.036885931 | 0.957233774 |
| ENSMUSG0000000384 | chr8 | Nfat5 | -0.039554186 | 0.957479405 |
| ENSMUSG0000005087 | chr18 | Minar2 | -0.03145303 | 0.957479405 |
| ENSMUSG0000004219 | chr9 | Slc35f2 | -0.039075876 | 0.957479405 |
| ENSMUSG0000004035 | chr4 | Ufl1 | 0.021074278 | 0.957479405 |
| ENSMUSG0000005070 | chr17 | 2310061I04Rik | 0.030377024 | 0.957479405 |
| ENSMUSG0000003474 | chr10 | Sirt6 | 0.033357958 | 0.957479405 |
| ENSMUSG0000004774 | chr6 | Zc3hav1l | -0.026043977 | 0.957724417 |
| ENSMUSG0000003775 | chr2 | Ppp1r16b | -0.046359709 | 0.957752622 |
| ENSMUSG0000007482 | chr13 | Rslcan18 | -0.095369303 | 0.957752622 |
| ENSMUSG0000004099 | chr14 | Abhd4 | -0.025613259 | 0.957752622 |
| ENSMUSG0000007086 | chr4 | Trabd2b | -0.030696075 | 0.957752622 |
| ENSMUSG0000003775 | chr4 | Xkr8 | -0.034392155 | 0.957752622 |
| ENSMUSG0000002677 | chr2 | Il2ra | -0.110629364 | 0.957752622 |
| ENSMUSG0000004683 | chr1 | Brox | -0.019700553 | 0.957752622 |
| ENSMUSG0000002747 | chr2 | Kif3b | -0.02007479 | 0.957752622 |
| ENSMUSG0000002393 | chr17 | Efhb | 0.04277178 | 0.957752622 |
| ENSMUSG0000004576 | chr17 | Togaram2 | 0.049467444 | 0.957752622 |
| ENSMUSG0000005377 | chr16 | Ubxn7 | 0.030094221 | 0.957752622 |
| ENSMUSG0000003811 | chr9 | Cdon | -0.032668106 | 0.957793379 |
| ENSMUSG0000005250 | chr16 | Epha3 | -0.047246743 | 0.957793379 |
| ENSMUSG0000010291 | chr18 | Pcdhgc3 | 0.026621271 | 0.957793379 |
| ENSMUSG0000004085 | chr4 | Bsdc1 | -0.025939757 | 0.957944841 |
| ENSMUSG0000001537 | chr15 | Dennd6b | -0.052093237 | 0.957990629 |
| ENSMUSG0000002241 | chr15 | Deptor | 0.050091872 | 0.957990629 |
| ENSMUSG0000006960 | chr10 | Ank3 | 0.029685289 | 0.957990629 |
| ENSMUSG0000003146 | chr8 | Angpt2 | 0.060181072 | 0.957990629 |
| ENSMUSG0000004306 | chr12 | Tmem18 | 0.026365768 | 0.957990629 |
| ENSMUSG0000000226 | chr7 | Zim1 | 0.078220409 | 0.958197491 |
| ENSMUSG0000008986 | chr6 | Umad1 | -0.035829634 | 0.958197491 |
| ENSMUSG0000004714 | chr16 | Zfp654 | -0.025090899 | 0.958197491 |
| ENSMUSG0000002242 | chr15 | Dmc1 | -0.113849363 | 0.958197491 |
| ENSMUSG0000003734 | chr5 | Slc12a9 | -0.031775244 | 0.958197491 |
| ENSMUSG0000004191 | chr18 | Ammecr1l | 0.034837103 | 0.958197491 |

| | | | | |
|---|---|---|---|---|
| ENSMUSG0000000510 | chr2 | Eif2ak4 | 0.026656921 | 0.958197491 |
| ENSMUSG0000004095 | chr11 | Mgl2 | 0.046224927 | 0.958197491 |
| ENSMUSG0000002221 | chr14 | Tgm1 | -0.120333188 | 0.958213209 |
| ENSMUSG0000005368 | chr10 | Kics2 | -0.042525616 | 0.958213209 |
| ENSMUSG0000007145 | chr12 | Dtnb | -0.027448583 | 0.958213209 |
| ENSMUSG0000006644 | chr12 | Rdh11 | -0.029208627 | 0.958213209 |
| ENSMUSG0000002885 | chr4 | Csf3r | -0.139113221 | 0.958213209 |
| ENSMUSG0000002148 | chr13 | Prxl2c | 0.037582337 | 0.958213209 |
| ENSMUSG0000004800 | chrX | Timm8a1 | 0.058436109 | 0.958213209 |
| ENSMUSG0000002953 | chr9 | Ccr9 | 0.114926836 | 0.958213209 |
| ENSMUSG0000007266 | chr15 | Spef2 | -0.06305076 | 0.958292092 |
| ENSMUSG0000003976 | chr5 | Cc2d2a | -0.036414965 | 0.958292092 |
| ENSMUSG0000004579 | chr5 | Ccdc149 | -0.030948811 | 0.958292092 |
| ENSMUSG0000001693 | chr2 | Plcg1 | 0.024725244 | 0.958345839 |
| ENSMUSG0000003097 | chr7 | Rrm1 | -0.031875382 | 0.958398981 |
| ENSMUSG0000003429 | chr1 | Traf3ip1 | -0.025241798 | 0.958398981 |
| ENSMUSG0000003535 | chr16 | Nfkbiz | 0.048013164 | 0.958471294 |
| ENSMUSG0000004459 | chr18 | Dnd1 | 0.112191616 | 0.958578931 |
| ENSMUSG0000002956 | chr6 | Foxp2 | 0.031651755 | 0.958578931 |
| ENSMUSG0000002809 | chr3 | Gpr89 | -0.036935388 | 0.958792391 |
| ENSMUSG0000006040 | chr7 | Chst8 | 0.053072332 | 0.958792391 |
| ENSMUSG0000003148 | chr8 | Adgra2 | 0.023091871 | 0.958878812 |
| ENSMUSG0000005322 | chr8 | Dand5 | 0.056235617 | 0.958878812 |
| ENSMUSG0000002741 | chr2 | Vps16 | 0.02552935 | 0.958928157 |
| ENSMUSG0000004955 | chr9 | Lingo1 | -0.049675276 | 0.95900731 |
| ENSMUSG0000002984 | chr6 | Hoxa1 | -0.066716584 | 0.95900731 |
| ENSMUSG0000005560 | chr17 | Tcp10b | -0.148474703 | 0.95900731 |
| ENSMUSG0000006279 | chr9 | Zfp599 | -0.137340325 | 0.95900731 |
| ENSMUSG0000003009 | chr6 | Slc6a6 | -0.038036667 | 0.95900731 |
| ENSMUSG0000003255 | chr9 | Topbp1 | -0.031829713 | 0.95900731 |
| ENSMUSG0000007172 | chrX | Spin4 | -0.042946359 | 0.95900731 |
| ENSMUSG0000004948 | chr11 | Ccnq | -0.029164629 | 0.95900731 |
| ENSMUSG0000003024 | chr6 | Abcc9 | -0.033244019 | 0.95900731 |
| ENSMUSG0000004429 | chr11 | Zfp879 | 0.056722347 | 0.95900731 |
| ENSMUSG0000004751 | chr4 | Slfnl1 | -0.184610036 | 0.95900731 |
| ENSMUSG0000003145 | chr8 | Gas6 | 0.029214934 | 0.95900731 |
| ENSMUSG0000002540 | chr10 | Ddit3 | 0.042615107 | 0.95900731 |
| ENSMUSG0000001982 | chr10 | Mical1 | 0.02612296 | 0.95900731 |
| ENSMUSG0000002082 | chr11 | Pld2 | 0.02538784 | 0.95900731 |
| ENSMUSG0000002886 | chr4 | Cd164l2 | 0.135172332 | 0.95900731 |
| ENSMUSG0000003008 | chr6 | Klf15 | 0.066868886 | 0.95900731 |
| ENSMUSG0000006178 | chr10 | Cfd | -0.140003386 | 0.959028171 |
| ENSMUSG0000003540 | chr2 | Crb2 | -0.06755245 | 0.959028171 |
| ENSMUSG0000005288 | chr7 | Prkcb | -0.041226907 | 0.959028171 |
| ENSMUSG0000008592 | chr12 | Rtl1 | -0.064440879 | 0.959028171 |
| ENSMUSG0000004076 | chr4 | Spen | -0.035598207 | 0.959028171 |
| ENSMUSG0000004264 | chr5 | Acad12 | -0.045753344 | 0.959028171 |
| ENSMUSG0000009006 | chr3 | 1110002E22Rik | 0.098667817 | 0.959028171 |
| ENSMUSG0000007890 | chr2 | Gm14443 | -0.059912744 | 0.959028171 |
| ENSMUSG0000002638 | chr1 | Steap3 | 0.071187415 | 0.959028171 |

| | | | | |
|---|---|---|---|---|
| ENSMUSG0000002057 | chr12 | Nbas | -0.030635288 | 0.959028171 |
| ENSMUSG0000003589 | chr15 | Cerk | -0.029729492 | 0.959028171 |
| ENSMUSG0000003161 | chr8 | Ednra | -0.031529749 | 0.959028171 |
| ENSMUSG0000003362 | chr9 | Hacd3 | -0.028972875 | 0.959028171 |
| ENSMUSG0000003410 | chr3 | Golim4 | -0.024060724 | 0.959028171 |
| ENSMUSG0000004486 | chr3 | Ankrd50 | -0.028162321 | 0.959028171 |
| ENSMUSG0000003323 | chr3 | Trim45 | -0.04075819 | 0.959028171 |
| ENSMUSG0000002693 | chr2 | Gpsm1 | -0.032941486 | 0.959028171 |
| ENSMUSG0000000105 | chr11 | Nhp2 | -0.03723896 | 0.959028171 |
| ENSMUSG0000002258 | chr15 | Rhpn1 | -0.054369186 | 0.959028171 |
| ENSMUSG0000005055 | chr2 | Kcnb1 | -0.027980641 | 0.959028171 |
| ENSMUSG0000004122 | chrX | Phf8 | -0.026089963 | 0.959028171 |
| ENSMUSG0000004042 | chr1 | Rc3h1 | -0.03187505 | 0.959028171 |
| ENSMUSG0000001460 | chr3 | Strip1 | -0.019672128 | 0.959028171 |
| ENSMUSG0000002991 | chr6 | Mrps33 | 0.042353372 | 0.959028171 |
| ENSMUSG0000002011 | chr10 | Cand1 | 0.013220576 | 0.959028171 |
| ENSMUSG0000004503 | chr17 | Tmem232 | 0.084236705 | 0.959028171 |
| ENSMUSG0000002477 | chr19 | Stambpl1 | 0.039904836 | 0.959028171 |
| ENSMUSG0000007164 | chr19 | Mta2 | 0.020872302 | 0.959028171 |
| ENSMUSG0000005281 | chr12 | Atad2b | 0.025345675 | 0.959028171 |
| ENSMUSG0000004107 | chr14 | Grid1 | 0.047783073 | 0.959028171 |
| ENSMUSG0000002252 | chr16 | Hes1 | 0.025608188 | 0.959028171 |
| ENSMUSG0000001837 | chr11 | Cep95 | 0.036694282 | 0.959028171 |
| ENSMUSG0000002071 | chr11 | Pecam1 | 0.040467064 | 0.959028171 |
| ENSMUSG0000004657 | chr5 | Zfp518b | 0.032255607 | 0.959028171 |
| ENSMUSG0000002494 | chr19 | Capn1 | 0.023213604 | 0.959028171 |
| ENSMUSG0000002078 | chr11 | P2rx1 | 0.088048525 | 0.959028171 |
| ENSMUSG0000002065 | chr12 | Cenpo | 0.044551826 | 0.959028171 |
| ENSMUSG0000001893 | chr11 | Map2k3 | 0.02685314 | 0.959028171 |
| ENSMUSG0000005438 | chr7 | Zfp747 | 0.036805591 | 0.959028171 |
| ENSMUSG0000004641 | chr11 | Lrrc75a | 0.037376912 | 0.959028171 |
| ENSMUSG0000004232 | chr5 | Hps4 | 0.027269752 | 0.959028171 |
| ENSMUSG0000002771 | chr3 | 1810062G17Rik | 0.112917323 | 0.959028171 |
| ENSMUSG0000003939 | chr7 | Ccdc81 | 0.096152741 | 0.959028171 |
| ENSMUSG0000001884 | chr11 | Unc45b | 0.077804698 | 0.959028171 |
| ENSMUSG0000003043 | chr7 | Tmem238 | 0.05836112 | 0.959028171 |
| ENSMUSG0000004404 | chr2 | Fmn1 | -0.030878921 | 0.959045833 |
| ENSMUSG0000004044 | chr18 | Rprd1a | -0.023624737 | 0.959045833 |
| ENSMUSG0000001854 | chr11 | Trim37 | -0.021378692 | 0.959045833 |
| ENSMUSG0000003552 | chr10 | Prdm4 | -0.019814654 | 0.959045833 |
| ENSMUSG0000007469 | chr2 | Csnk2a1 | -0.017079921 | 0.959045833 |
| ENSMUSG0000004165 | chr15 | Pnpla3 | 0.060833707 | 0.959045833 |
| ENSMUSG0000002417 | chr17 | Tekt4 | 0.096646497 | 0.959045833 |
| ENSMUSG0000004161 | chr4 | Nppa | -0.139789014 | 0.959066455 |
| ENSMUSG0000002550 | chr7 | Tmem80 | -0.03188197 | 0.959066455 |
| ENSMUSG0000004879 | chr6 | Cfap100 | -0.054302753 | 0.95932683 |
| ENSMUSG0000002646 | chr1 | Xpr1 | -0.023655331 | 0.95932683 |
| ENSMUSG0000001485 | chr8 | E2f4 | 0.018047925 | 0.95932683 |
| ENSMUSG0000002212 | chr14 | Obi1 | -0.030932152 | 0.959634276 |
| ENSMUSG0000002848 | chr4 | Snapc3 | -0.02160057 | 0.959634276 |

| | | | | |
|---|---|---|---|---|
| ENSMUSG0000002661 | chr1 | Spata17 | -0.105276364 | 0.959782559 |
| ENSMUSG0000005872 | chr1 | Lin9 | 0.096415882 | 0.960049555 |
| ENSMUSG0000005553 | chr10 | Cpsf6 | -0.019678029 | 0.960049555 |
| ENSMUSG0000004446 | chr3 | Tent5c | 0.074005271 | 0.960049555 |
| ENSMUSG0000005260 | chr12 | Plekhg3 | 0.024121316 | 0.960049555 |
| ENSMUSG0000005005 | chr8 | Tdrp | -0.031750206 | 0.960095885 |
| ENSMUSG0000003161 | chr8 | Tmem184c | -0.021835892 | 0.960218724 |
| ENSMUSG0000005211 | chr4 | D630039A03Rik | 0.038013998 | 0.960218724 |
| ENSMUSG0000000920 | chr2 | Lnpk | -0.035489998 | 0.960456681 |
| ENSMUSG0000003023 | chr6 | Plekha5 | -0.02923 | 0.960510382 |
| ENSMUSG0000003181 | chr8 | Emc8 | 0.030908359 | 0.960526775 |
| ENSMUSG0000005364 | chr9 | Dennd4a | -0.035501649 | 0.960738474 |
| ENSMUSG0000001908 | chrX | Atp6ap1 | -0.017503332 | 0.960738474 |
| ENSMUSG0000004054 | chr11 | Tex2 | -0.016395343 | 0.960738474 |
| ENSMUSG0000000381 | chr8 | Dnase2a | 0.043762234 | 0.960738474 |
| ENSMUSG0000002142 | chr13 | Ssr1 | 0.021633304 | 0.960738474 |
| ENSMUSG0000007081 | chr7 | Zswim9 | 0.021908783 | 0.960738474 |
| ENSMUSG0000002649 | chr1 | Cdc42bpa | 0.024121188 | 0.960738474 |
| ENSMUSG0000005769 | chr6 | Zfp746 | 0.019363639 | 0.960738474 |
| ENSMUSG0000004953 | chrX | Tceal1 | 0.030476387 | 0.960738474 |
| ENSMUSG0000004689 | chr15 | Zfp740 | 0.024024037 | 0.960738474 |
| ENSMUSG0000004404 | chr18 | Pcdhb14 | 0.031452521 | 0.960738474 |
| ENSMUSG0000002940 | chr5 | Abcb9 | 0.031978405 | 0.960738474 |
| ENSMUSG0000005867 | chr13 | Tubb2a | 0.032883059 | 0.960887459 |
| ENSMUSG0000007125 | chr17 | Zfp213 | -0.042412927 | 0.960991698 |
| ENSMUSG0000002451 | chr18 | Grp | -0.039995883 | 0.960991698 |
| ENSMUSG0000002212 | chr14 | Scel | 0.104096457 | 0.961094722 |
| ENSMUSG0000006983 | chr19 | Ahnak | 0.04392826 | 0.961094722 |
| ENSMUSG0000009096 | chr5 | Gm17655 | -0.084239791 | 0.961210938 |
| ENSMUSG0000007905 | chr12 | Slc8a3 | -0.052174182 | 0.961317904 |
| ENSMUSG0000002327 | chr6 | Cd4 | -0.118698777 | 0.961317904 |
| ENSMUSG0000003079 | chr7 | Adm | -0.057078987 | 0.961317904 |
| ENSMUSG0000003899 | chr11 | H1f9 | -0.108197052 | 0.961317904 |
| ENSMUSG0000005940 | chrX | Mamld1 | -0.039878906 | 0.961317904 |
| ENSMUSG0000004866 | chr10 | Lemd3 | -0.027004052 | 0.961317904 |
| ENSMUSG0000003271 | chr2 | Trib3 | -0.047723931 | 0.961317904 |
| ENSMUSG0000002716 | chr2 | Ccdc34 | -0.035075629 | 0.961317904 |
| ENSMUSG0000001547 | chr17 | Rnf5 | -0.023216132 | 0.961317904 |
| ENSMUSG0000003475 | chr11 | Tmub2 | -0.018993614 | 0.961317904 |
| ENSMUSG0000006822 | chr15 | Il2rb | -0.087879564 | 0.961317904 |
| ENSMUSG0000003032 | chr6 | Efcab12 | -0.065692358 | 0.961317904 |
| ENSMUSG0000004844 | chr11 | Ccdc57 | 0.043706913 | 0.961317904 |
| ENSMUSG0000005292 | chr8 | Rnaseh2a | 0.029399511 | 0.961317904 |
| ENSMUSG0000002858 | chr4 | Pum1 | 0.015308945 | 0.961317904 |
| ENSMUSG0000002509 | chr19 | Hspa12a | 0.02020607 | 0.961317904 |
| ENSMUSG0000001754 | chr11 | Suz12 | 0.021534486 | 0.961317904 |
| ENSMUSG0000002101 | chr16 | Polr2h | 0.033230961 | 0.961317904 |
| ENSMUSG0000002267 | chr16 | Ube2v2 | 0.029802739 | 0.961317904 |
| ENSMUSG0000003925 | chr2 | Pomt1 | 0.021944697 | 0.961317904 |
| ENSMUSG0000002235 | chr15 | Fbxo32 | 0.036040305 | 0.961317904 |

| | | | | |
|---|---|---|---|---|
| ENSMUSG0000001336 | chr7 | Iglon5 | 0.02507553 | 0.961317904 |
| ENSMUSG0000005149 | chr8 | Irf2bp2 | 0.028530972 | 0.961317904 |
| ENSMUSG0000000053 | chr11 | Tom1l2 | -0.021372795 | 0.96133521 |
| ENSMUSG0000003527 | chr5 | Hpse | 0.050376934 | 0.961429045 |
| ENSMUSG0000004457 | chr12 | Acp1 | -0.051247689 | 0.961649655 |
| ENSMUSG0000010093 | chr19 | Nscme3l | -0.07164881 | 0.961649655 |
| ENSMUSG0000000262 | chr17 | Akap8l | -0.028037434 | 0.961649655 |
| ENSMUSG0000003747 | chrX | Thoc2 | -0.027231286 | 0.961649655 |
| ENSMUSG0000003464 | chr3 | Tiparp | -0.0299791 | 0.961649655 |
| ENSMUSG0000000973 | chr15 | Tfcp2 | -0.025724693 | 0.961649655 |
| ENSMUSG0000004181 | chr15 | Poldip3 | -0.019787328 | 0.961649655 |
| ENSMUSG0000006913 | chr17 | Cep43 | -0.023340757 | 0.961649655 |
| ENSMUSG0000002543 | chr7 | Clns1a | 0.029281719 | 0.961649655 |
| ENSMUSG0000003153 | chr8 | Plat | 0.046150421 | 0.961649655 |
| ENSMUSG0000002504 | chrX | Fundc1 | 0.025608915 | 0.961649655 |
| ENSMUSG0000002910 | chr5 | Rgs12 | 0.037060872 | 0.961649655 |
| ENSMUSG0000002094 | chr12 | Fkbp3 | 0.018789825 | 0.961649655 |
| ENSMUSG0000002255 | chr15 | Mroh1 | 0.022124304 | 0.961649655 |
| ENSMUSG0000003305 | chr9 | Npat | 0.031104432 | 0.961649655 |
| ENSMUSG0000003152 | chr8 | Vegfc | 0.038690213 | 0.961649655 |
| ENSMUSG0000002749 | chr3 | Pkia | 0.022230948 | 0.961649655 |
| ENSMUSG0000004111 | chr7 | Ccdc8 | 0.030744235 | 0.961649655 |
| ENSMUSG0000007054 | chr2 | Top1 | 0.029614249 | 0.961649655 |
| ENSMUSG0000003545 | chr7 | Tnni3 | 0.054278095 | 0.961649655 |
| ENSMUSG0000003841 | chr13 | Foxq1 | 0.057225832 | 0.961649655 |
| ENSMUSG0000011794 | chr18 | Maskbp3 | -0.089749397 | 0.961673881 |
| ENSMUSG0000004833 | chr14 | Npy4r | -0.028667641 | 0.961951913 |
| ENSMUSG0000002276 | chr16 | Serpind1 | 0.121106246 | 0.962062231 |
| ENSMUSG0000000203 | chr9 | Cd3g | -0.16372563 | 0.962103966 |
| ENSMUSG0000000374 | chr10 | Man1a | -0.027567423 | 0.962103966 |
| ENSMUSG0000002268 | chr16 | Ntan1 | -0.024464424 | 0.962103966 |
| ENSMUSG0000000634 | chr10 | Ggt5 | -0.044035419 | 0.962103966 |
| ENSMUSG0000003883 | chr2 | Ralgps1 | 0.029133402 | 0.962103966 |
| ENSMUSG0000002043 | chr11 | Adcy1 | 0.040591737 | 0.962103966 |
| ENSMUSG0000003087 | chr7 | Ears2 | 0.042751295 | 0.962103966 |
| ENSMUSG0000003274 | chr13 | Gpbp1 | 0.017993461 | 0.962103966 |
| ENSMUSG0000002419 | chr17 | Bnip1 | 0.030143083 | 0.962103966 |
| ENSMUSG0000003879 | chr1 | Lefty1 | 0.052959527 | 0.962103966 |
| ENSMUSG0000003446 | chr12 | Six4 | -0.095368042 | 0.962140621 |
| ENSMUSG0000001999 | chr10 | Ccn2 | 0.038295011 | 0.962140621 |
| ENSMUSG0000004262 | chr3 | Shc1 | 0.016431821 | 0.962197578 |
| ENSMUSG0000003562 | chr16 | Rubcn | 0.023769933 | 0.962321261 |
| ENSMUSG0000003153 | chr8 | Polb | 0.023267068 | 0.962448748 |
| ENSMUSG0000009556 | chr4 | Noc2l | 0.022609475 | 0.962448748 |
| ENSMUSG0000004065 | chr10 | Ppp1r14c | 0.067152608 | 0.962448748 |
| ENSMUSG0000002256 | chr15 | Gpaa1 | 0.020562011 | 0.962448748 |
| ENSMUSG0000009001 | chr6 | Gimap1 | 0.076042188 | 0.962667083 |
| ENSMUSG0000004259 | chr5 | Sh2b3 | 0.031243758 | 0.962784842 |
| ENSMUSG0000004277 | chr1 | Smg7 | -0.023146961 | 0.96287946 |
| ENSMUSG0000003484 | chr8 | Plvap | 0.052802523 | 0.962892136 |

| | | | | |
|---|---|---|---|---|
| ENSMUSG0000001997 | chr10 | Apaf1 | 0.029436766 | 0.962892136 |
| ENSMUSG0000004680 | chr6 | Cyren | 0.056410958 | 0.962959654 |
| ENSMUSG0000002467 | chr19 | Ms4a6b | -0.053250515 | 0.96317734 |
| ENSMUSG0000003942 | chr16 | Alg1 | -0.022875758 | 0.96317734 |
| ENSMUSG0000003995 | chr6 | Etfbkmt | 0.034127846 | 0.96317734 |
| ENSMUSG0000005159 | chr18 | Pcdhb2 | 0.069486481 | 0.96317734 |
| ENSMUSG0000003454 | chr11 | Morc2a | 0.024717626 | 0.963189562 |
| ENSMUSG0000002972 | chr5 | Mepce | 0.028102535 | 0.963189562 |
| ENSMUSG0000003495 | chr14 | Rubcnl | -0.094394801 | 0.96332407 |
| ENSMUSG0000002548 | chr7 | Cox8b | 0.048651416 | 0.96332407 |
| ENSMUSG0000002595 | chr1 | Creb1 | 0.019429672 | 0.96332407 |
| ENSMUSG0000004263 | chr2 | Xkr7 | 0.085705188 | 0.96332407 |
| ENSMUSG0000003508 | chr11 | 1700020L24Rik | -0.072800196 | 0.963363175 |
| ENSMUSG0000004230 | chr11 | Ehbp1 | 0.022667854 | 0.963363175 |
| ENSMUSG0000002481 | chr19 | Frmd8 | 0.028523276 | 0.963363175 |
| ENSMUSG0000002217 | chr14 | Ajuba | -0.036420751 | 0.963446927 |
| ENSMUSG0000006160 | chr12 | Akap6 | -0.027165792 | 0.963446927 |
| ENSMUSG0000003797 | chr9 | Cfap68 | -0.02549187 | 0.963596028 |
| ENSMUSG0000002084 | chr11 | Ywhae | 0.016353947 | 0.963654486 |
| ENSMUSG0000002189 | chr14 | Sh3bp5 | -0.021129471 | 0.963750513 |
| ENSMUSG0000000063 | chr11 | Sez6 | -0.030788951 | 0.963856045 |
| ENSMUSG0000003266 | chr6 | Pon2 | -0.021382744 | 0.963856045 |
| ENSMUSG0000007941 | chr19 | Ms4a6c | -0.062884041 | 0.96393032 |
| ENSMUSG0000003923 | chr1 | Tgfb2 | 0.031059186 | 0.96393032 |
| ENSMUSG0000000151 | chr6 | Foxm1 | -0.08845752 | 0.9639817 |
| ENSMUSG0000003250 | chr15 | Trib1 | -0.037900942 | 0.9639817 |
| ENSMUSG0000005400 | chr4 | Tusc1 | -0.038440951 | 0.9639817 |
| ENSMUSG0000003462 | chr11 | Gpatch8 | -0.023927579 | 0.9639817 |
| ENSMUSG0000008631 | chrX | Nbdy | -0.024602471 | 0.9639817 |
| ENSMUSG0000002742 | chr2 | Rrbp1 | -0.016880837 | 0.9639817 |
| ENSMUSG0000005838 | chr3 | Phtf1 | -0.016262089 | 0.9639817 |
| ENSMUSG0000004343 | chr7 | Leng9 | -0.042347299 | 0.9639817 |
| ENSMUSG0000002623 | chr1 | Ptma | 0.028224949 | 0.9639817 |
| ENSMUSG0000004680 | chr5 | Atp10d | 0.037166492 | 0.9639817 |
| ENSMUSG0000004771 | chr8 | Champ1 | 0.014789963 | 0.9639817 |
| ENSMUSG0000002089 | chr11 | Ctc1 | 0.033095958 | 0.9639817 |
| ENSMUSG0000004130 | chr1 | Gtf3c3 | 0.020275042 | 0.9639817 |
| ENSMUSG0000004012 | chr14 | Zmym5 | 0.024247168 | 0.9639817 |
| ENSMUSG0000003181 | chr8 | Mthfsd | 0.017676951 | 0.9639817 |
| ENSMUSG0000003998 | chr2 | Ccdc32 | 0.036375693 | 0.9639817 |
| ENSMUSG0000005005 | chr5 | Ccdc158 | 0.063062535 | 0.9639817 |
| ENSMUSG0000003239 | chr9 | Dis3l | 0.021756291 | 0.9639817 |
| ENSMUSG0000007034 | chr7 | Ccnd1 | 0.036177158 | 0.9639817 |
| ENSMUSG0000002136 | chr13 | Mak | 0.070047926 | 0.9639817 |
| ENSMUSG0000005493 | chr16 | Zfp174 | 0.034758377 | 0.9639817 |
| ENSMUSG0000000596 | chr3 | Tuft1 | -0.057201767 | 0.963984117 |
| ENSMUSG0000002212 | chr14 | Cln5 | -0.030232611 | 0.963984117 |
| ENSMUSG0000002107 | chr12 | Bdkrb2 | 0.036252141 | 0.963984117 |
| ENSMUSG0000002200 | chr14 | Zc3h13 | 0.023247167 | 0.963984117 |
| ENSMUSG0000003603 | chr17 | Zfp57 | -0.042288544 | 0.963991146 |

| | | | | |
|---|---|---|---|---|
| ENSMUSG0000000126 | chr5 | Gabrg1 | -0.106019461 | 0.963991146 |
| ENSMUSG0000006117 | chr2 | Slc38a11 | 0.081432091 | 0.963991146 |
| ENSMUSG0000009484 | chr11 | Tmem95 | 0.124875718 | 0.963991146 |
| ENSMUSG0000003464 | chr11 | Cd300ld | 0.060916493 | 0.963991146 |
| ENSMUSG0000003156 | chr8 | Rwdd4a | 0.016645482 | 0.964026315 |
| ENSMUSG0000005010 | chr4 | Fam110d | 0.099611 | 0.964026315 |
| ENSMUSG0000002432 | chr17 | Slc39a7 | 0.049587192 | 0.964026315 |
| ENSMUSG0000002274 | chr16 | St3gal6 | 0.027792578 | 0.964169223 |
| ENSMUSG0000002968 | chr6 | Cul1 | 0.017689914 | 0.964223734 |
| ENSMUSG0000002848 | chr4 | Bnc2 | 0.049441612 | 0.964223734 |
| ENSMUSG0000003918 | chr7 | Fanci | 0.148099887 | 0.964225646 |
| ENSMUSG0000002421 | chr17 | Nudt3 | 0.022238278 | 0.964225646 |
| ENSMUSG0000008973 | chr8 | Tgfbr3l | -0.080342224 | 0.964612644 |
| ENSMUSG0000004315 | chr4 | Hpdl | -0.110231288 | 0.964612644 |
| ENSMUSG0000007049 | chr2 | Ctcfl | -0.058506151 | 0.964765247 |
| ENSMUSG0000005985 | chr3 | Ntng1 | -0.037406624 | 0.964938419 |
| ENSMUSG0000002268 | chr16 | B3gnt5 | 0.034793371 | 0.964938419 |
| ENSMUSG0000007403 | chr8 | Exoc8 | 0.027435796 | 0.964938419 |
| ENSMUSG0000002665 | chr1 | Dusp12 | -0.033737602 | 0.964951579 |
| ENSMUSG0000003151 | chr8 | Leprotl1 | -0.019463052 | 0.964951579 |
| ENSMUSG0000003269 | chr2 | Lmo2 | -0.037722853 | 0.964951579 |
| ENSMUSG0000008558 | chrX | Rtl9 | 0.091353407 | 0.964984614 |
| ENSMUSG0000000721 | chr6 | Zfp775 | -0.034456847 | 0.965047946 |
| ENSMUSG0000002171 | chr13 | Ppwd1 | 0.023993391 | 0.965047946 |
| ENSMUSG0000004061 | chr1 | Ildr2 | -0.059979931 | 0.965267909 |
| ENSMUSG0000002323 | chr8 | Ccl25 | -0.035692304 | 0.965386519 |
| ENSMUSG0000004099 | chrX | Sh3kbp1 | -0.022866136 | 0.965386519 |
| ENSMUSG0000005073 | chr9 | Arhgap42 | -0.026390396 | 0.965386519 |
| ENSMUSG0000000014 | chr11 | Axin2 | -0.021222848 | 0.965386519 |
| ENSMUSG0000002870 | chr4 | Rad54l | 0.075023549 | 0.965386519 |
| ENSMUSG0000000230 | chr17 | Cd320 | 0.049632494 | 0.965386519 |
| ENSMUSG0000010023 | chr13 | Gm28557 | 0.081514325 | 0.965386519 |
| ENSMUSG0000002759 | chr2 | Itch | 0.018293561 | 0.965386519 |
| ENSMUSG0000007463 | chr3 | Sox2 | 0.045230688 | 0.965386519 |
| ENSMUSG0000002282 | chr16 | Ndufb4 | 0.041699673 | 0.965386519 |
| ENSMUSG0000002190 | chr14 | Dph3 | -0.028302748 | 0.965455172 |
| ENSMUSG0000007377 | chr4 | Btbd19 | -0.041538391 | 0.965455172 |
| ENSMUSG0000004084 | chr1 | Tiprl | -0.022052905 | 0.965455172 |
| ENSMUSG0000002646 | chr1 | Tor1aip1 | 0.017286503 | 0.965455172 |
| ENSMUSG0000002682 | chr2 | Gbgt1 | 0.031019673 | 0.965455172 |
| ENSMUSG0000001943 | chr11 | Tlcd1 | 0.030405851 | 0.965553014 |
| ENSMUSG0000003512 | chr4 | Dnai4 | -0.032648957 | 0.965594831 |
| ENSMUSG0000002924 | chr5 | Rest | 0.027082776 | 0.965594831 |
| ENSMUSG0000002448 | chr18 | Lvrn | 0.105115082 | 0.965749905 |
| ENSMUSG0000006404 | chr17 | Trerf1 | -0.035586626 | 0.965749905 |
| ENSMUSG0000003432 | chr11 | Brip1 | -0.094491002 | 0.965749905 |
| ENSMUSG0000005771 | chr6 | Tmem178b | -0.030690656 | 0.965749905 |
| ENSMUSG0000005974 | chr3 | Fdps | 0.049872014 | 0.965749905 |
| ENSMUSG0000004724 | chr13 | H2bc6 | -0.048315422 | 0.965749905 |
| ENSMUSG0000004963 | chr2 | C1ql3 | -0.063944467 | 0.965749905 |

| | | | | |
|---|---|---|---|---|
| ENSMUSG0000002131 | chr13 | Mtr | -0.027062267 | 0.965749905 |
| ENSMUSG0000009062 | chr9 | Tex9 | -0.03824138 | 0.965749905 |
| ENSMUSG0000008198 | chr1 | Dnajb3 | -0.062282153 | 0.965749905 |
| ENSMUSG0000004192 | chr2 | Metap1d | -0.023400284 | 0.965749905 |
| ENSMUSG0000003316 | chr3 | Atp1a1 | -0.027286236 | 0.965749905 |
| ENSMUSG0000002521 | chr19 | Btrc | -0.021212916 | 0.965749905 |
| ENSMUSG0000001130 | chr17 | Plin5 | 0.139764864 | 0.965749905 |
| ENSMUSG0000002141 | chr13 | Fam217a | -0.061615203 | 0.965749905 |
| ENSMUSG0000002210 | chr14 | Xpo7 | -0.01525292 | 0.965749905 |
| ENSMUSG0000008579 | chr8 | Zfp703 | -0.024763202 | 0.965749905 |
| ENSMUSG0000003950 | chr2 | Znfx1 | -0.019836252 | 0.965749905 |
| ENSMUSG0000003341 | chr10 | Gucd1 | -0.025228354 | 0.965749905 |
| ENSMUSG0000004107 | chr16 | Rwdd2b | -0.05093959 | 0.965749905 |
| ENSMUSG0000005100 | chr3 | Fhip1a | 0.047310485 | 0.965749905 |
| ENSMUSG0000002125 | chr12 | Ccnk | 0.026007291 | 0.965749905 |
| ENSMUSG0000002701 | chr2 | Dync1i2 | 0.011581326 | 0.965749905 |
| ENSMUSG0000002074 | chr11 | Gga3 | 0.022658482 | 0.965749905 |
| ENSMUSG0000000589 | chr16 | Smpd4 | 0.022467111 | 0.965749905 |
| ENSMUSG0000003088 | chr7 | Polr3e | 0.023063549 | 0.965749905 |
| ENSMUSG0000003451 | chr8 | Hmgxb4 | 0.01970598 | 0.965749905 |
| ENSMUSG0000003723 | chr18 | Matr3 | 0.01994917 | 0.965749905 |
| ENSMUSG0000002864 | chr4 | Mycbp | 0.023389533 | 0.965749905 |
| ENSMUSG0000002091 | chr11 | Stat5b | 0.015920355 | 0.965749905 |
| ENSMUSG0000000937 | chr6 | Met | 0.017141771 | 0.965749905 |
| ENSMUSG0000002916 | chr5 | Mapre3 | 0.016134245 | 0.965749905 |
| ENSMUSG0000008658 | chr4 | Gm11837 | 0.040970956 | 0.965749905 |
| ENSMUSG0000002765 | chr2 | Tti1 | 0.01839483 | 0.965749905 |
| ENSMUSG0000005823 | chr7 | Arhgap35 | 0.022092346 | 0.965749905 |
| ENSMUSG0000003925 | chr7 | Vstm2b | 0.046528822 | 0.965749905 |
| ENSMUSG0000004224 | chr5 | Grk3 | 0.026322741 | 0.965749905 |
| ENSMUSG0000007373 | chr4 | Cplane2 | 0.066329997 | 0.965749905 |
| ENSMUSG0000007412 | chr7 | Ntf5 | 0.092030898 | 0.965749905 |
| ENSMUSG0000003187 | chr8 | Cdh5 | 0.043126177 | 0.965816695 |
| ENSMUSG0000003692 | chr5 | Stag3 | -0.058743947 | 0.965816695 |
| ENSMUSG0000002887 | chr4 | Stx12 | -0.014955068 | 0.965816695 |
| ENSMUSG0000000288 | chr1 | Nab1 | 0.030931238 | 0.965816695 |
| ENSMUSG0000002475 | chr19 | Cemip2 | -0.020186455 | 0.966331579 |
| ENSMUSG0000006113 | chr2 | Prpf40a | -0.014719022 | 0.966331579 |
| ENSMUSG0000002736 | chr2 | Stard7 | -0.015738255 | 0.966331579 |
| ENSMUSG0000003676 | chr1 | Dner | 0.02746274 | 0.966331579 |
| ENSMUSG0000003953 | chr5 | Mmd2 | -0.034501382 | 0.966405354 |
| ENSMUSG0000003047 | chr7 | Siglece | -0.066068775 | 0.966405354 |
| ENSMUSG0000002875 | chr4 | Hp1bp3 | -0.021691934 | 0.966405354 |
| ENSMUSG0000005124 | chr18 | Pcdhb9 | -0.022154469 | 0.966405354 |
| ENSMUSG0000007262 | chr6 | Zfp9 | 0.019990078 | 0.966405354 |
| ENSMUSG0000003596 | chr14 | Apex1 | 0.025608483 | 0.966405354 |
| ENSMUSG0000000532 | chr13 | Fgfr4 | 0.065543324 | 0.966405354 |
| ENSMUSG0000004252 | chr3 | Ubap2l | 0.018321412 | 0.966474129 |
| ENSMUSG0000004367 | chr7 | Dpy19l3 | -0.040280736 | 0.966501812 |
| ENSMUSG0000003450 | chr17 | Mad2l1bp | -0.038197513 | 0.966501812 |

| | | | | |
|---|---|---|---|---|
| ENSMUSG0000003754 | chr7 | Shank2 | -0.023315411 | 0.966501812 |
| ENSMUSG0000003547 | chrX | Tab3 | -0.021472798 | 0.966501812 |
| ENSMUSG0000007896 | chr18 | Hsbp1l1 | -0.085485313 | 0.966501812 |
| ENSMUSG0000004044 | chr10 | Slc26a10 | -0.112322606 | 0.966501812 |
| ENSMUSG0000010254 | chr18 | Pcdhgc5 | 0.042041828 | 0.966501812 |
| ENSMUSG0000006188 | chr5 | Ccdc62 | 0.060213136 | 0.966501812 |
| ENSMUSG0000000347 | chr6 | Crhr2 | 0.120559213 | 0.966501812 |
| ENSMUSG0000003238 | chr9 | Spg21 | 0.021186219 | 0.966501812 |
| ENSMUSG0000002946 | chr5 | Gpn3 | 0.031922444 | 0.966501812 |
| ENSMUSG0000003018 | chr6 | Kdm5a | 0.023716276 | 0.966501812 |
| ENSMUSG0000004176 | chr1 | Tpp2 | -0.019616039 | 0.966525893 |
| ENSMUSG0000002718 | chr2 | Caprin1 | -0.014210398 | 0.966525893 |
| ENSMUSG0000005079 | chr4 | B3galt6 | 0.04099492 | 0.96653157 |
| ENSMUSG0000002549 | chr7 | Ifitm3 | 0.032303731 | 0.966596628 |
| ENSMUSG0000000489 | chr3 | Nes | 0.037800576 | 0.966770128 |
| ENSMUSG0000004827 | chr5 | Rbm33 | -0.022828066 | 0.966865827 |
| ENSMUSG0000007421 | chr3 | Dnajb14 | -0.019221899 | 0.966875258 |
| ENSMUSG0000002537 | chr11 | Tepsin | 0.027757285 | 0.966875258 |
| ENSMUSG0000003808 | chr2 | Cnbd2 | -0.051825268 | 0.966898424 |
| ENSMUSG0000002232 | chr8 | Shcbp1 | -0.110718848 | 0.96714424 |
| ENSMUSG0000005711 | chr2 | Cntrl | -0.02771779 | 0.967280685 |
| ENSMUSG0000002823 | chr4 | Tgs1 | -0.021089999 | 0.967280685 |
| ENSMUSG0000000644 | chr4 | Epha2 | -0.032671736 | 0.967280685 |
| ENSMUSG0000003331 | chr8 | Fbxl8 | -0.03749587 | 0.967280685 |
| ENSMUSG0000004553 | chr11 | C1ql1 | 0.044959235 | 0.967280685 |
| ENSMUSG0000002589 | chr9 | Cwf19l2 | 0.022749059 | 0.967280685 |
| ENSMUSG0000002843 | chr4 | Dcaf12 | 0.02034228 | 0.967280685 |
| ENSMUSG0000003080 | chr7 | Stx4a | 0.019962893 | 0.967280685 |
| ENSMUSG0000005133 | chr13 | Gfod1 | 0.020426944 | 0.967280685 |
| ENSMUSG0000005316 | chr2 | Gpr21 | 0.094307224 | 0.967280685 |
| ENSMUSG0000003820 | chr5 | Asb10 | 0.131063759 | 0.967280685 |
| ENSMUSG0000002700 | chr2 | Dusp19 | -0.03090042 | 0.967405185 |
| ENSMUSG0000006771 | chr15 | Prkag1 | 0.02670129 | 0.967407374 |
| ENSMUSG0000004285 | chr2 | Zc3h6 | -0.052025326 | 0.967431735 |
| ENSMUSG0000006426 | chr5 | Hvcn1 | -0.092175899 | 0.967431735 |
| ENSMUSG0000006414 | chr9 | Arih2 | -0.013062097 | 0.967431735 |
| ENSMUSG0000003204 | chr2 | Abhd12 | -0.016448501 | 0.967431735 |
| ENSMUSG0000002844 | chr4 | Nudt2 | 0.034862519 | 0.967431735 |
| ENSMUSG0000006380 | chr7 | Ap3s2 | 0.018143724 | 0.967431735 |
| ENSMUSG0000004170 | chr12 | Btbd7 | 0.021262871 | 0.967431735 |
| ENSMUSG0000004866 | chr6 | Rhno1 | 0.023015894 | 0.967431735 |
| ENSMUSG0000009478 | chr2 | Gm14403 | 0.042683588 | 0.967431735 |
| ENSMUSG0000002215 | chr14 | Rab2b | -0.026975552 | 0.967499588 |
| ENSMUSG0000003051 | chr7 | Tjp1 | 0.020881967 | 0.967499588 |
| ENSMUSG0000002828 | chr4 | Casp8ap2 | 0.024248339 | 0.967499588 |
| ENSMUSG0000004247 | chrX | Tbc1d8b | 0.019411162 | 0.967499588 |
| ENSMUSG0000009494 | chr13 | Gm3604 | 0.031835844 | 0.967499588 |
| ENSMUSG0000004527 | chr13 | Cenph | 0.135570856 | 0.967499588 |
| ENSMUSG0000007287 | chr6 | Nup50l | 0.143893898 | 0.967499588 |
| ENSMUSG0000001988 | chr10 | Rspo3 | 0.036058658 | 0.967544058 |

| | | | | |
|---|---|---|---|---|
| ENSMUSG0000007296 | chrX | Armcx5 | -0.039175961 | 0.967594464 |
| ENSMUSG0000002722 | chr2 | Sord | 0.034273248 | 0.967671386 |
| ENSMUSG0000002469 | chr19 | Zfp91 | 0.02028488 | 0.967814753 |
| ENSMUSG0000007931 | chrX | Trappc2 | 0.028835003 | 0.96783159 |
| ENSMUSG0000005260 | chr7 | Isoc2b | -0.043124922 | 0.967845651 |
| ENSMUSG0000002542 | chr18 | Pias2 | -0.02249874 | 0.967845651 |
| ENSMUSG0000002664 | chr1 | Usf1 | -0.021317518 | 0.967845651 |
| ENSMUSG0000005077 | chr1 | Tmem37 | 0.057173707 | 0.967845651 |
| ENSMUSG0000007107 | chr8 | Nr2c2ap | -0.058580458 | 0.967928861 |
| ENSMUSG0000004720 | chr11 | Dusp18 | -0.035860684 | 0.96794499 |
| ENSMUSG0000005311 | chr7 | Fank1 | 0.057447949 | 0.96794499 |
| ENSMUSG0000002320 | chr2 | Il15ra | 0.030684812 | 0.96794499 |
| ENSMUSG0000003923 | chr11 | Tbcd | 0.016889433 | 0.96794499 |
| ENSMUSG0000002396 | chr17 | Cyp39a1 | -0.031694653 | 0.967980071 |
| ENSMUSG0000003509 | chr14 | Fam167a | 0.04275252 | 0.967980071 |
| ENSMUSG0000003332 | chr17 | Tnxb | 0.041433647 | 0.968013591 |
| ENSMUSG0000004562 | chr18 | Sh3tc2 | -0.047080454 | 0.968040733 |
| ENSMUSG0000003691 | chr7 | Kirrel2 | -0.125724298 | 0.968482274 |
| ENSMUSG0000001509 | chr2 | Ptgds | 0.08185135 | 0.968482274 |
| ENSMUSG0000007955 | chr9 | Pou2af3 | 0.060199071 | 0.968486119 |
| ENSMUSG0000004249 | chr2 | Hoxd11 | -0.031026178 | 0.968535695 |
| ENSMUSG0000005653 | chrX | Rlim | -0.019518333 | 0.968535695 |
| ENSMUSG0000005592 | chr5 | Aasdh | -0.025893295 | 0.968573829 |
| ENSMUSG0000005783 | chrX | Xlr3a | -0.122575028 | 0.96863028 |
| ENSMUSG0000002256 | chr15 | Oplah | 0.034985342 | 0.96863028 |
| ENSMUSG0000005755 | chr9 | Zfp317 | -0.015810128 | 0.968903457 |
| ENSMUSG0000007904 | chr2 | Fastkd5 | 0.041936386 | 0.968903457 |
| ENSMUSG0000005471 | chr7 | Zscan22 | 0.020644515 | 0.968903457 |
| ENSMUSG0000001583 | chr2 | Nfe2l2 | 0.032018487 | 0.968940655 |
| ENSMUSG0000003106 | chrX | Cdk16 | 0.016298379 | 0.968980302 |
| ENSMUSG0000002128 | chr12 | Xrcc3 | 0.042182644 | 0.968980302 |
| ENSMUSG0000002194 | chr14 | Ptpn20 | -0.062142993 | 0.969054279 |
| ENSMUSG0000003536 | chr13 | Rmi1 | 0.022887987 | 0.969054279 |
| ENSMUSG0000003155 | chr5 | Slit2 | -0.025493655 | 0.969094774 |
| ENSMUSG0000001579 | chr2 | Surf1 | -0.023220372 | 0.969094774 |
| ENSMUSG0000004008 | chr2 | Bub1b | -0.054518449 | 0.969094774 |
| ENSMUSG0000003170 | chr8 | Tecr | 0.015243948 | 0.969094774 |
| ENSMUSG0000003013 | chr6 | Rasgef1a | -0.039557515 | 0.969277271 |
| ENSMUSG0000002307 | chr7 | Cep89 | 0.024777986 | 0.969299072 |
| ENSMUSG0000007069 | chr4 | Runx3 | -0.09258567 | 0.969344107 |
| ENSMUSG0000004878 | chr7 | Insc | -0.045730729 | 0.969344107 |
| ENSMUSG0000002131 | chr13 | Sfrp4 | 0.079517328 | 0.969344107 |
| ENSMUSG0000002457 | chr18 | Pde6a | 0.076747883 | 0.969344107 |
| ENSMUSG0000002528 | chr14 | Polr3a | 0.019526329 | 0.969344107 |
| ENSMUSG0000004965 | chr13 | Bdp1 | 0.026153858 | 0.969349011 |
| ENSMUSG0000004728 | chr11 | Neurl4 | -0.013379878 | 0.969351169 |
| ENSMUSG0000003593 | chr9 | Pknox2 | -0.03063489 | 0.969414088 |
| ENSMUSG0000002210 | chr14 | Itm2b | 0.017024466 | 0.969414088 |
| ENSMUSG0000001936 | chr11 | Sec14l4 | -0.09086239 | 0.969588817 |
| ENSMUSG0000004002 | chr19 | Rab11fip2 | -0.026646083 | 0.969588817 |

| | | | | |
|---|---|---|---|---|
| ENSMUSG0000002113 | chr12 | Srsf5 | -0.028337734 | 0.969588817 |
| ENSMUSG0000002697 | chr2 | Dph7 | -0.025156633 | 0.969588817 |
| ENSMUSG0000003429 | chr11 | Med13 | -0.021964559 | 0.969588817 |
| ENSMUSG0000001370 | chr3 | Tnfaip8l2 | -0.049392416 | 0.969588817 |
| ENSMUSG0000002328 | chr5 | Zfp605 | -0.033989484 | 0.969588817 |
| ENSMUSG0000002130 | chr13 | Ggps1 | -0.017982725 | 0.969588817 |
| ENSMUSG0000004540 | chr17 | Trim39 | -0.018190715 | 0.969588817 |
| ENSMUSG0000002442 | chr17 | Gnl1 | -0.014581411 | 0.969588817 |
| ENSMUSG0000001918 | chr11 | Rnf145 | 0.025034164 | 0.969588817 |
| ENSMUSG0000004222 | chrX | Ammecr1 | 0.029683255 | 0.969588817 |
| ENSMUSG0000005056 | chr1 | Tor1aip2 | 0.012195016 | 0.969588817 |
| ENSMUSG0000002068 | chr11 | Cdc27 | 0.016350918 | 0.969588817 |
| ENSMUSG0000003966 | chr4 | Icmt | 0.020047463 | 0.969588817 |
| ENSMUSG0000001362 | chr5 | Cad | 0.022470879 | 0.969588817 |
| ENSMUSG0000005104 | chr7 | P4ha3 | 0.035331883 | 0.969588817 |
| ENSMUSG0000009181 | chr7 | Inafm1 | -0.020967396 | 0.969664678 |
| ENSMUSG0000001027 | chr11 | Bltp2 | 0.015494836 | 0.969664678 |
| ENSMUSG0000006187 | chr11 | Sphk1 | 0.027106456 | 0.969664678 |
| ENSMUSG0000006920 | chr13 | Zfp825 | 0.027473758 | 0.969716699 |
| ENSMUSG0000003539 | chr18 | Cfap53 | 0.103157798 | 0.969716699 |
| ENSMUSG0000004436 | chr15 | BC024139 | 0.091252 | 0.969739536 |
| ENSMUSG0000004495 | chr7 | Kctd21 | 0.027985391 | 0.969739536 |
| ENSMUSG0000002719 | chr2 | Hsd17b12 | 0.029981181 | 0.969739536 |
| ENSMUSG0000000113 | chrX | Uxt | -0.040243555 | 0.969762879 |
| ENSMUSG0000006829 | chr2 | Ddrgk1 | 0.019014973 | 0.969762879 |
| ENSMUSG0000003627 | chr15 | Lrrk2 | 0.024126601 | 0.969762879 |
| ENSMUSG0000002022 | chr10 | Helb | -0.032206858 | 0.969930967 |
| ENSMUSG0000001699 | chr2 | Matn4 | 0.055869592 | 0.969930967 |
| ENSMUSG0000002447 | chr18 | Pggt1b | 0.017282766 | 0.969930967 |
| ENSMUSG0000003527 | chr10 | Plekhj1 | 0.024083604 | 0.969930967 |
| ENSMUSG0000000652 | chr14 | Sfmbt1 | 0.022556274 | 0.969930967 |
| ENSMUSG0000002672 | chr2 | Cubn | 0.136309239 | 0.970072547 |
| ENSMUSG0000002912 | chr5 | Nsg1 | -0.025130869 | 0.970072547 |
| ENSMUSG0000005093 | chr3 | Sgms2 | 0.040175143 | 0.970072547 |
| ENSMUSG0000006189 | chr5 | Rbak | -0.024079152 | 0.970072547 |
| ENSMUSG0000005618 | chr19 | Snx32 | -0.019450629 | 0.970072547 |
| ENSMUSG0000004269 | chr1 | Dhx9 | 0.011975275 | 0.970072547 |
| ENSMUSG0000002809 | chr3 | Rnf115 | 0.018402045 | 0.970072547 |
| ENSMUSG0000003262 | chr13 | Srek1 | 0.01527898 | 0.970072547 |
| ENSMUSG0000002006 | chr10 | Hnrnph3 | 0.015768102 | 0.970072547 |
| ENSMUSG0000002686 | chr2 | Gapvd1 | 0.016878324 | 0.970072547 |
| ENSMUSG0000002809 | chr3 | Polr3c | 0.015285414 | 0.970072547 |
| ENSMUSG0000000911 | chr6 | Bcl2l13 | 0.014468697 | 0.970072547 |
| ENSMUSG0000003322 | chr15 | Scaf11 | 0.018055213 | 0.970072547 |
| ENSMUSG0000003034 | chr6 | Rad51ap1 | -0.102442975 | 0.970145152 |
| ENSMUSG0000005132 | chr2 | Nup160 | -0.01568804 | 0.970145152 |
| ENSMUSG0000003213 | chr9 | Mcam | 0.022975243 | 0.970145152 |
| ENSMUSG0000004609 | chr7 | Mosmo | 0.015888527 | 0.970145152 |
| ENSMUSG0000003621 | chr15 | Pdzrn4 | -0.039712801 | 0.970145213 |
| ENSMUSG0000005127 | chr4 | Gdf6 | 0.135055457 | 0.970145213 |

| | | | | |
|---|---|---|---|---|
| ENSMUSG0000000598 | chr16 | Naa60 | -0.028139465 | 0.970145213 |
| ENSMUSG0000003721 | chr8 | Thap1 | -0.033107974 | 0.970145213 |
| ENSMUSG0000002188 | chr14 | Hacl1 | -0.050727001 | 0.970145213 |
| ENSMUSG0000003162 | chr8 | Cenpu | -0.062180749 | 0.970145213 |
| ENSMUSG0000002155 | chr13 | Naa35 | -0.016060235 | 0.970145213 |
| ENSMUSG0000002676 | chr2 | Orc4 | -0.016371013 | 0.970261261 |
| ENSMUSG0000005387 | chr7 | Srcap | -0.043473334 | 0.970529834 |
| ENSMUSG0000002416 | chr17 | Tmem204 | -0.030614649 | 0.970529834 |
| ENSMUSG0000001353 | chr16 | Tango2 | -0.021922951 | 0.970529834 |
| ENSMUSG0000002444 | chr18 | Pcdh12 | -0.053202397 | 0.970529834 |
| ENSMUSG0000003802 | chr13 | Phf2 | 0.01861108 | 0.970529834 |
| ENSMUSG0000005476 | chr2 | Set | 0.016086169 | 0.970529834 |
| ENSMUSG0000002231 | chr15 | Rad21 | 0.013584443 | 0.970529834 |
| ENSMUSG0000002634 | chr1 | Acmsd | 0.106308881 | 0.970529834 |
| ENSMUSG0000002930 | chr5 | Spp1 | 0.061370578 | 0.970529834 |
| ENSMUSG0000003548 | chr10 | Plk5 | -0.251982985 | 0.97059416 |
| ENSMUSG0000002257 | chr15 | Pycr3 | -0.0324645 | 0.97059416 |
| ENSMUSG0000002386 | chr17 | Pde10a | -0.02814731 | 0.97059416 |
| ENSMUSG0000000241 | chr6 | Braf | -0.018893134 | 0.97059416 |
| ENSMUSG0000007933 | chr9 | Naa80 | -0.019531808 | 0.97059416 |
| ENSMUSG0000002716 | chr2 | Traf6 | -0.016227286 | 0.97059416 |
| ENSMUSG0000005449 | chr7 | Dedd2 | -0.027295156 | 0.97059416 |
| ENSMUSG0000000785 | chr11 | Hnrnph1 | 0.011104805 | 0.97059416 |
| ENSMUSG0000003546 | chr14 | Rcbtb1 | 0.016349401 | 0.97059416 |
| ENSMUSG0000001869 | chr11 | Aatf | 0.019447224 | 0.97059416 |
| ENSMUSG0000000353 | chr17 | Ddr1 | 0.024835946 | 0.97059416 |
| ENSMUSG0000003734 | chr4 | Paqr7 | 0.021782349 | 0.97059416 |
| ENSMUSG0000002609 | chr1 | Asnsd1 | 0.040969841 | 0.97059416 |
| ENSMUSG0000006084 | chr10 | Ctnna3 | 0.029817976 | 0.97059416 |
| ENSMUSG0000002255 | chr15 | Hgh1 | -0.036376972 | 0.970616514 |
| ENSMUSG0000010358 | chr18 | Pcdhgb4 | -0.033159493 | 0.970616514 |
| ENSMUSG0000003161 | chr8 | Scrg1 | -0.044214645 | 0.970616514 |
| ENSMUSG0000003343 | chrX | Armcx2 | -0.020175568 | 0.970616514 |
| ENSMUSG0000007073 | chr1 | Dgkd | -0.011499578 | 0.970616514 |
| ENSMUSG0000002897 | chr5 | Nos3 | 0.040448755 | 0.970616514 |
| ENSMUSG0000005014 | chr3 | Slc25a44 | -0.018018233 | 0.970855146 |
| ENSMUSG0000007139 | chr10 | Ect2l | -0.108201828 | 0.970994659 |
| ENSMUSG0000003920 | chr2 | Ciz1 | -0.017907547 | 0.970997242 |
| ENSMUSG0000002056 | chr12 | Atxn7l1 | -0.022249578 | 0.970997242 |
| ENSMUSG0000005064 | chr5 | Tmem150c | -0.028691907 | 0.970997242 |
| ENSMUSG0000002666 | chr2 | Mcm10 | -0.075880617 | 0.970997242 |
| ENSMUSG0000003529 | chrX | Mid1 | 0.051065747 | 0.970997242 |
| ENSMUSG0000001838 | chr11 | Abi3 | -0.056624777 | 0.971408972 |
| ENSMUSG0000003319 | chr8 | Lpcat2 | -0.042530083 | 0.971701285 |
| ENSMUSG0000004026 | chr17 | Daam2 | -0.023591114 | 0.971701285 |
| ENSMUSG0000002037 | chr11 | Rufy1 | 0.017632376 | 0.971701285 |
| ENSMUSG0000004381 | chr16 | Rtn4r | -0.048201817 | 0.971864709 |
| ENSMUSG0000007948 | chrX | Med12 | -0.026299773 | 0.971864709 |
| ENSMUSG0000000899 | chr2 | Bmp7 | -0.027216984 | 0.971864709 |
| ENSMUSG0000002976 | chr6 | Plxna4 | -0.023686109 | 0.97192872 |

| | | | | |
|---|---|---|---|---|
| ENSMUSG0000003492 | chr13 | Rnf44 | -0.019663016 | 0.97192872 |
| ENSMUSG0000004741 | chr10 | Rexo1 | -0.019590508 | 0.97192872 |
| ENSMUSG0000002867 | chr4 | Fuca1 | -0.021016456 | 0.97192872 |
| ENSMUSG0000003040 | chr7 | Ercc2 | -0.018798782 | 0.97192872 |
| ENSMUSG0000004916 | chr19 | Zfp518a | -0.016608742 | 0.97192872 |
| ENSMUSG0000003022 | chr6 | Pik3c2g | 0.107409038 | 0.97192872 |
| ENSMUSG0000003978 | chr5 | Cpeb2 | 0.025188542 | 0.97192872 |
| ENSMUSG0000004892 | chr13 | Ccdc125 | 0.048152473 | 0.97192872 |
| ENSMUSG0000005499 | chr19 | Naaladl1 | -0.1878126 | 0.971966503 |
| ENSMUSG0000002835 | chr4 | Zfp618 | -0.035282181 | 0.971966503 |
| ENSMUSG0000002527 | chr14 | Dnase1l3 | -0.08209459 | 0.971966503 |
| ENSMUSG0000002501 | chr19 | Ccnj | -0.028122083 | 0.971966503 |
| ENSMUSG0000000401 | chr11 | Fancl | -0.033315307 | 0.971966503 |
| ENSMUSG0000004241 | chr2 | Agps | -0.012630527 | 0.971966503 |
| ENSMUSG0000003010 | chr6 | Slc6a12 | -0.082193118 | 0.971966503 |
| ENSMUSG0000010917 | chr15 | Wdr97 | 0.036398634 | 0.971966503 |
| ENSMUSG0000002608 | chr1 | Rev1 | 0.030562302 | 0.971966503 |
| ENSMUSG0000002409 | chr17 | Ralbp1 | 0.01257851 | 0.971966503 |
| ENSMUSG0000003682 | chr4 | Topors | 0.015513747 | 0.971966503 |
| ENSMUSG0000002222 | chr13 | Zscan26 | 0.015051259 | 0.971966503 |
| ENSMUSG0000005776 | chr18 | Ankrd29 | 0.026308668 | 0.971966503 |
| ENSMUSG0000001430 | chr16 | Pam16 | 0.023250357 | 0.971966503 |
| ENSMUSG0000004786 | chr6 | Gimap6 | 0.025524861 | 0.971966503 |
| ENSMUSG0000003746 | chr15 | Klf10 | 0.022250609 | 0.971966503 |
| ENSMUSG0000002852 | chr4 | Pde4b | 0.021751603 | 0.971966503 |
| ENSMUSG0000004626 | chrX | Usp27x | 0.029984886 | 0.971966503 |
| ENSMUSG0000003689 | chr4 | C1qc | 0.03017354 | 0.971966503 |
| ENSMUSG0000002580 | chr9 | Ccr1 | 0.04036004 | 0.972107525 |
| ENSMUSG0000003983 | chr4 | Slc45a1 | -0.028105187 | 0.972113521 |
| ENSMUSG0000002942 | chr5 | Zcchc8 | -0.01705135 | 0.972113521 |
| ENSMUSG0000004709 | chr10 | Tmem198b | 0.021652341 | 0.972113521 |
| ENSMUSG0000005258 | chr14 | Serp2 | 0.031637071 | 0.972113521 |
| ENSMUSG0000004460 | chr8 | Smim7 | 0.022733426 | 0.972169787 |
| ENSMUSG0000009732 | chr11 | Tnfsf12 | 0.022057845 | 0.972169787 |
| ENSMUSG0000001948 | chr17 | Trip10 | 0.021153575 | 0.972169787 |
| ENSMUSG0000005760 | chr14 | Colq | 0.025963504 | 0.972169787 |
| ENSMUSG0000005709 | chr11 | Ebf1 | -0.0377693 | 0.972326227 |
| ENSMUSG0000006653 | chr7 | Vmn2r57 | -0.079298103 | 0.972326227 |
| ENSMUSG0000005727 | chr6 | Snrpg | 0.030871301 | 0.972326227 |
| ENSMUSG0000004566 | chr11 | Smtnl2 | -0.040255987 | 0.972427126 |
| ENSMUSG0000002439 | chr17 | Prrc2a | -0.019725492 | 0.972427126 |
| ENSMUSG0000002860 | chr4 | Echdc2 | -0.025517012 | 0.972427126 |
| ENSMUSG0000004414 | chr12 | Arf6 | -0.017342604 | 0.972427126 |
| ENSMUSG0000000249 | chr17 | Tsc2 | 0.013284628 | 0.972427126 |
| ENSMUSG0000002548 | chr7 | Ric8a | 0.01488053 | 0.972427126 |
| ENSMUSG0000002856 | chr4 | Nfia | 0.02610935 | 0.972427126 |
| ENSMUSG0000002252 | chr16 | Zfp263 | 0.01687893 | 0.972427126 |
| ENSMUSG0000004942 | chr10 | Tmem200a | 0.017476069 | 0.972427126 |
| ENSMUSG0000007878 | chr11 | Dph1 | 0.024323839 | 0.972427126 |
| ENSMUSG0000003261 | chr9 | 1700102P08Rik | 0.068485383 | 0.972427126 |

| | | | | |
|---|---|---|---|---|
| ENSMUSG0000004761 | chr9 | Ddi1 | 0.079534477 | 0.972427126 |
| ENSMUSG0000002740 | chr2 | Tgm3 | -0.058757236 | 0.972474466 |
| ENSMUSG0000003148 | chr8 | Thsd1 | -0.026396541 | 0.972474466 |
| ENSMUSG0000003198 | chr8 | Arv1 | 0.032518903 | 0.972474466 |
| ENSMUSG0000006660 | chr9 | Insyn1 | 0.027156467 | 0.972474466 |
| ENSMUSG0000004049 | chr17 | Lrfn2 | 0.035439021 | 0.972474466 |
| ENSMUSG0000002843 | chr4 | Nol6 | 0.011144628 | 0.972474466 |
| ENSMUSG0000004435 | chr14 | Lacc1 | 0.034416025 | 0.972474466 |
| ENSMUSG0000007309 | chr6 | Lrrc61 | 0.022228977 | 0.972474466 |
| ENSMUSG0000003545 | chr11 | Fignl1 | 0.065926406 | 0.972474466 |
| ENSMUSG0000004025 | chr5 | Sema3d | 0.031164344 | 0.972474466 |
| ENSMUSG0000002498 | chr19 | Cep55 | -0.100014205 | 0.972675428 |
| ENSMUSG0000001947 | chr8 | Xab2 | 0.016452694 | 0.972675428 |
| ENSMUSG0000005722 | chr7 | Dmac2 | -0.027516239 | 0.972830927 |
| ENSMUSG0000007843 | chr10 | AU041133 | 0.031623026 | 0.972830927 |
| ENSMUSG0000005783 | chr17 | Zfp119a | 0.024738173 | 0.972830927 |
| ENSMUSG0000002233 | chr15 | Zfat | -0.034892159 | 0.972999936 |
| ENSMUSG0000002481 | chr19 | Il33 | -0.04291352 | 0.973007324 |
| ENSMUSG0000005469 | chr9 | Adam10 | -0.014647553 | 0.973025742 |
| ENSMUSG0000004367 | chr12 | Kcns3 | -0.03311687 | 0.973355661 |
| ENSMUSG0000007876 | chr7 | Haus5 | -0.066353358 | 0.973395866 |
| ENSMUSG0000007056 | chr1 | Rasal2 | 0.02068792 | 0.973460722 |
| ENSMUSG0000002279 | chr16 | Tnk2 | -0.019436875 | 0.973667334 |
| ENSMUSG0000002738 | chr2 | Bcl2l11 | 0.024844909 | 0.973667334 |
| ENSMUSG0000004887 | chr11 | Hexim1 | -0.018297902 | 0.973765231 |
| ENSMUSG0000003123 | chrX | Fgf16 | 0.088628856 | 0.973906027 |
| ENSMUSG0000003877 | chr18 | Kdm3b | -0.013553073 | 0.973963375 |
| ENSMUSG0000004848 | chr4 | Zbtb8b | 0.118830093 | 0.974048512 |
| ENSMUSG0000005035 | chr8 | Carmil2 | -0.031073775 | 0.974073999 |
| ENSMUSG0000003233 | chr9 | Col12a1 | -0.027353742 | 0.974110668 |
| ENSMUSG0000006148 | chr13 | H4c4 | -0.126987469 | 0.974227812 |
| ENSMUSG0000007166 | chr19 | Zbtb3 | -0.035026263 | 0.974454843 |
| ENSMUSG0000003228 | chr9 | Thsd4 | 0.021543341 | 0.974454843 |
| ENSMUSG0000004779 | chr11 | Cd300lf | 0.068096258 | 0.974454843 |
| ENSMUSG0000002899 | chr5 | Rint1 | 0.021124398 | 0.974454843 |
| ENSMUSG0000003311 | chr14 | Cep15 | -0.014893132 | 0.974488243 |
| ENSMUSG0000003226 | chr9 | Bckdhb | 0.024718257 | 0.974488243 |
| ENSMUSG0000006153 | chr12 | Cep128 | 0.051371499 | 0.974575718 |
| ENSMUSG0000003753 | chr11 | Rapgef6 | -0.020643178 | 0.974575718 |
| ENSMUSG0000004062 | chr10 | Plekhg1 | 0.033110744 | 0.974575718 |
| ENSMUSG0000009459 | chr4 | Fsbp | 0.077393319 | 0.974575718 |
| ENSMUSG0000004323 | chr1 | Fam124b | -0.106577915 | 0.974583342 |
| ENSMUSG0000002679 | chr2 | Odf2 | -0.025384698 | 0.974619795 |
| ENSMUSG0000003512 | chr6 | Gcfc2 | -0.02624609 | 0.974619795 |
| ENSMUSG0000003466 | chr5 | Bmp2k | -0.02095056 | 0.974619795 |
| ENSMUSG0000005599 | chr8 | Nod2 | 0.074887365 | 0.974619795 |
| ENSMUSG0000001912 | chr6 | Scrn1 | -0.017771019 | 0.974619795 |
| ENSMUSG0000003799 | chr8 | Dhx38 | -0.015466371 | 0.974619795 |
| ENSMUSG0000002204 | chr14 | Trim35 | -0.015570529 | 0.974619795 |
| ENSMUSG0000003435 | chr11 | Mtmr3 | -0.012263018 | 0.974619795 |

| | | | | |
|---|---|---|---|---|
| ENSMUSG0000000667 | chr9 | Usp19 | -0.011836946 | 0.974619795 |
| ENSMUSG0000002047 | chr11 | Dbnl | -0.011319179 | 0.974619795 |
| ENSMUSG0000003111 | chrX | Stk26 | 0.040524315 | 0.974619795 |
| ENSMUSG0000003398 | chr4 | Tesk2 | 0.034917716 | 0.974619795 |
| ENSMUSG0000001574 | chr3 | Vps45 | 0.025026079 | 0.974619795 |
| ENSMUSG0000003975 | chr5 | Fbxl5 | 0.011218961 | 0.974619795 |
| ENSMUSG0000002844 | chr4 | Cntfr | 0.029979347 | 0.974619795 |
| ENSMUSG0000003575 | chr10 | Bbs10 | 0.028852983 | 0.974619795 |
| ENSMUSG0000003144 | chr8 | Proz | 0.041051987 | 0.974619795 |
| ENSMUSG0000002023 | chr10 | Fzr1 | 0.011913953 | 0.974619795 |
| ENSMUSG0000009748 | chr11 | Ptges3l | 0.040476056 | 0.974619795 |
| ENSMUSG0000003518 | chr10 | Ano4 | 0.063365448 | 0.974619795 |
| ENSMUSG0000003904 | chr7 | Arpin | 0.012933709 | 0.974659556 |
| ENSMUSG0000002521 | chr19 | Poll | 0.02329886 | 0.974659556 |
| ENSMUSG0000002128 | chr12 | Ppp1r13b | -0.01641473 | 0.974763953 |
| ENSMUSG0000003486 | chr8 | Ano8 | -0.021374833 | 0.974763953 |
| ENSMUSG0000001513 | chr7 | Aldh1a3 | 0.018781143 | 0.974763953 |
| ENSMUSG0000000274 | chr11 | Ykt6 | 0.012762973 | 0.974763953 |
| ENSMUSG0000000124 | chr11 | Ramp2 | 0.018641338 | 0.974764698 |
| ENSMUSG0000005244 | chr8 | Zfp961 | 0.017210032 | 0.974855236 |
| ENSMUSG0000012016 | chr5 | Zfp33b | -0.124397899 | 0.974860097 |
| ENSMUSG0000003998 | chr3 | Ankrd13c | -0.013723758 | 0.974862261 |
| ENSMUSG0000001308 | chr16 | Etv5 | 0.021315305 | 0.97499426 |
| ENSMUSG0000003948 | chr2 | Asb6 | 0.015980687 | 0.975062455 |
| ENSMUSG0000003470 | chr17 | Ppp1r21 | -0.013673224 | 0.975174508 |
| ENSMUSG0000003590 | chr15 | Gramd4 | 0.022005283 | 0.975234967 |
| ENSMUSG0000007029 | chr9 | Trcg1 | 0.071342832 | 0.975290563 |
| ENSMUSG0000003032 | chr6 | Pianp | 0.078128662 | 0.975290563 |
| ENSMUSG0000006154 | chr17 | Zfp229 | -0.018107267 | 0.975450516 |
| ENSMUSG0000002328 | chr4 | Ube2j2 | -0.014497305 | 0.975450516 |
| ENSMUSG0000005649 | chr3 | Tmem154 | 0.039689987 | 0.975450516 |
| ENSMUSG0000007499 | chr2 | Qser1 | 0.016348579 | 0.975450516 |
| ENSMUSG0000007528 | chr19 | Carns1 | 0.027443736 | 0.975529979 |
| ENSMUSG0000002922 | chr5 | Slc30a9 | 0.011756698 | 0.97553313 |
| ENSMUSG0000003322 | chr3 | Ttf2 | -0.026766558 | 0.975533997 |
| ENSMUSG0000003370 | chr6 | Smyd5 | 0.026810382 | 0.975533997 |
| ENSMUSG0000007272 | chr5 | Ccdc121rt2 | -0.074083132 | 0.975731707 |
| ENSMUSG0000003201 | chr9 | Grik4 | -0.027560436 | 0.975731707 |
| ENSMUSG0000003652 | chr15 | Sbf1 | -0.015409287 | 0.975731707 |
| ENSMUSG0000003559 | chr12 | Prpf39 | -0.014095228 | 0.975731707 |
| ENSMUSG0000004002 | chr4 | Ythdf2 | -0.012654877 | 0.975731707 |
| ENSMUSG0000000342 | chr7 | Nosip | 0.01996333 | 0.975731707 |
| ENSMUSG0000005172 | chr12 | Wdcp | 0.017939422 | 0.975731707 |
| ENSMUSG0000002669 | chr1 | Fmo4 | -0.094682242 | 0.975746445 |
| ENSMUSG0000003609 | chr2 | Arl5a | -0.011990732 | 0.975746445 |
| ENSMUSG0000002189 | chr14 | Asb14 | -0.06852979 | 0.975763711 |
| ENSMUSG0000003571 | chr13 | Dok3 | -0.035153605 | 0.975763711 |
| ENSMUSG0000002038 | chr11 | Rad50 | -0.023299175 | 0.975763711 |
| ENSMUSG0000002880 | chr4 | Hdac1 | -0.022304251 | 0.975763711 |
| ENSMUSG0000003766 | chr7 | Cdkn1c | -0.025043145 | 0.975763711 |

| | | | | |
|---|---|---|---|---|
| ENSMUSG0000001767 | chr11 | Wsb1 | -0.026383756 | 0.975763711 |
| ENSMUSG0000002509 | chr19 | Slc18a2 | -0.019585373 | 0.975763711 |
| ENSMUSG0000005279 | chr8 | 1700030K09Rik | -0.021870104 | 0.975763711 |
| ENSMUSG0000009691 | chr7 | Zfp850 | -0.030988189 | 0.975763711 |
| ENSMUSG0000002458 | chr18 | Napg | -0.014173515 | 0.975763711 |
| ENSMUSG0000004125 | chr18 | Zfp236 | -0.014128927 | 0.975763711 |
| ENSMUSG0000000278 | chr7 | Tmem143 | -0.016227434 | 0.975763711 |
| ENSMUSG0000001577 | chr2 | Med22 | -0.015920612 | 0.975763711 |
| ENSMUSG0000004082 | chr16 | Hlcs | -0.022744153 | 0.975763711 |
| ENSMUSG0000002154 | chr13 | Rasa1 | -0.01034491 | 0.975763711 |
| ENSMUSG0000003681 | chr9 | Slc6a20a | 0.178697369 | 0.975763711 |
| ENSMUSG0000000165 | chr15 | Hoxc11 | 0.043958166 | 0.975763711 |
| ENSMUSG0000003701 | chr10 | Hk1 | -0.011412674 | 0.975763711 |
| ENSMUSG0000006917 | chr13 | Adgrv1 | 0.091731196 | 0.975763711 |
| ENSMUSG0000004718 | chr4 | Rab2a | 0.014377684 | 0.975763711 |
| ENSMUSG0000009616 | chr17 | Or10h5 | 0.09432481 | 0.975763711 |
| ENSMUSG0000004052 | chr4 | Manea | 0.011975192 | 0.975763711 |
| ENSMUSG0000000873 | chr3 | Hipk1 | 0.015041395 | 0.975763711 |
| ENSMUSG0000005206 | chr1 | Pard3b | 0.016500384 | 0.975763711 |
| ENSMUSG0000002008 | chr10 | Fam241b | 0.0246801 | 0.975763711 |
| ENSMUSG0000003556 | chr4 | Aldh1b1 | 0.01797319 | 0.975763711 |
| ENSMUSG0000009027 | chr17 | Prr22 | 0.043721892 | 0.975763711 |
| ENSMUSG0000004047 | chr2 | Dgkz | 0.010029957 | 0.975771075 |
| ENSMUSG0000003171 | chr8 | Gab1 | -0.02078803 | 0.976038428 |
| ENSMUSG0000002721 | chr2 | Cd82 | -0.029280846 | 0.976038428 |
| ENSMUSG0000007145 | chr13 | Psmg4 | -0.031843204 | 0.976038428 |
| ENSMUSG0000002863 | chr4 | Kcnq4 | -0.018543214 | 0.976038428 |
| ENSMUSG0000003708 | chr15 | Trmt12 | 0.024481014 | 0.976038428 |
| ENSMUSG0000005594 | chr17 | Prr18 | -0.058602038 | 0.976038428 |
| ENSMUSG0000002619 | chr1 | Zfand2b | 0.016541821 | 0.976038428 |
| ENSMUSG0000003624 | chr2 | Rbm43 | 0.014103767 | 0.976038428 |
| ENSMUSG0000004655 | chr8 | Zfp319 | 0.019801341 | 0.976038428 |
| ENSMUSG0000002896 | chr4 | Ube4b | 0.015494951 | 0.976038428 |
| ENSMUSG0000003842 | chr18 | Poli | 0.019640051 | 0.976038428 |
| ENSMUSG0000003650 | chr2 | Phpt1 | 0.052978628 | 0.976038428 |
| ENSMUSG0000005021 | chr2 | Pla2g4e | 0.079088976 | 0.976038428 |
| ENSMUSG0000003388 | chr3 | Zfp267 | -0.018606253 | 0.976090515 |
| ENSMUSG0000004036 | chrX | Bcor | -0.016491005 | 0.976104246 |
| ENSMUSG0000002970 | chr6 | Rbm28 | 0.015605225 | 0.976282798 |
| ENSMUSG0000003160 | chr8 | Sap30 | 0.023776315 | 0.976320301 |
| ENSMUSG0000005146 | chr18 | Zfp24 | 0.015915648 | 0.976531755 |
| ENSMUSG0000000319 | chr17 | Mpnd | -0.016672938 | 0.97659335 |
| ENSMUSG0000004113 | chrX | Smc1a | -0.011836051 | 0.97659335 |
| ENSMUSG0000003996 | chr8 | Rhou | 0.037774376 | 0.976793237 |
| ENSMUSG0000003139 | chrX | Fam3a | 0.012789014 | 0.976799274 |
| ENSMUSG0000001842 | chr11 | Dhx40 | -0.017897167 | 0.976820034 |
| ENSMUSG0000005517 | chr6 | C1ra | -0.028541771 | 0.976990796 |
| ENSMUSG0000006715 | chr1 | Col4a4 | -0.030514106 | 0.977030785 |
| ENSMUSG0000002460 | chr18 | Isoc1 | -0.025449153 | 0.977035971 |
| ENSMUSG0000007489 | chr2 | Sptbn5 | -0.049705331 | 0.977035971 |

| | | | | |
|---|---|---|---|---|
| ENSMUSG0000002462 | chr18 | Csf1r | -0.025161727 | 0.977035971 |
| ENSMUSG0000003159 | chr8 | Pcm1 | 0.014653972 | 0.977035971 |
| ENSMUSG0000007135 | chr6 | Reg3b | 0.233942889 | 0.977035971 |
| ENSMUSG0000003740 | chr9 | Icam1 | -0.026306466 | 0.977058293 |
| ENSMUSG0000002171 | chr13 | Cenpk | -0.076701119 | 0.977119663 |
| ENSMUSG0000002135 | chr13 | Irf4 | -0.06010496 | 0.977119663 |
| ENSMUSG0000002495 | chr19 | Kcnk4 | -0.050531016 | 0.977119663 |
| ENSMUSG0000002819 | chr3 | Col24a1 | -0.04341856 | 0.977119663 |
| ENSMUSG0000003852 | chr10 | Mfsd4b1 | -0.079370616 | 0.977119663 |
| ENSMUSG0000003859 | chr15 | Colec10 | -0.022268688 | 0.977119663 |
| ENSMUSG0000008071 | chr16 | B230307C23Rik | -0.023240099 | 0.977119663 |
| ENSMUSG0000005870 | chr6 | 0610030E20Rik | -0.021649565 | 0.977119663 |
| ENSMUSG0000003699 | chr8 | Otud4 | -0.016491507 | 0.977119663 |
| ENSMUSG0000002911 | chr5 | Man2b2 | -0.017942652 | 0.977119663 |
| ENSMUSG0000004643 | chrX | Bex3 | -0.017492329 | 0.977119663 |
| ENSMUSG0000002027 | chr11 | Fbxw11 | -0.012041975 | 0.977119663 |
| ENSMUSG0000004217 | chr7 | Armc5 | -0.014072556 | 0.977119663 |
| ENSMUSG0000004930 | chr19 | Syt12 | 0.064612733 | 0.977119663 |
| ENSMUSG0000004221 | chr6 | Zfand4 | 0.031600455 | 0.977119663 |
| ENSMUSG0000004539 | chr2 | Or5m3b | 0.037006926 | 0.977119663 |
| ENSMUSG0000008966 | chr11 | Tnfsf13 | 0.062573725 | 0.977119663 |
| ENSMUSG0000001914 | chr18 | Hars2 | 0.014247301 | 0.977119663 |
| ENSMUSG0000003140 | chrX | Dkc1 | 0.015632366 | 0.977119663 |
| ENSMUSG0000004854 | chr15 | Tob2 | 0.025060074 | 0.977119663 |
| ENSMUSG0000002922 | chr5 | Fip1l1 | 0.013116735 | 0.977119663 |
| ENSMUSG0000004052 | chrX | Tlr8 | 0.050083702 | 0.977119663 |
| ENSMUSG0000006314 | chr2 | Bbs5 | 0.021533109 | 0.977119663 |
| ENSMUSG0000006248 | chr17 | Acat3 | 0.070156001 | 0.977119663 |
| ENSMUSG0000002002 | chr10 | Socs2 | 0.023595219 | 0.977119663 |
| ENSMUSG0000007111 | chr6 | Spx | -0.063960533 | 0.977119663 |
| ENSMUSG0000007949 | chr17 | H2-T10 | 0.044885521 | 0.977119663 |
| ENSMUSG0000010414 | chr18 | Pcdha2 | -0.054462869 | 0.977135836 |
| ENSMUSG0000000961 | chr2 | Sardh | -0.031123881 | 0.977135836 |
| ENSMUSG0000005992 | chr3 | Unc5c | -0.02291117 | 0.977135836 |
| ENSMUSG0000005705 | chr11 | Inca1 | -0.044191992 | 0.977135836 |
| ENSMUSG0000003260 | chr9 | Nicn1 | -0.013655805 | 0.977135836 |
| ENSMUSG0000004065 | chr4 | Efhd2 | -0.015824439 | 0.977135836 |
| ENSMUSG0000004105 | chr4 | Wwp1 | 0.015424052 | 0.977135836 |
| ENSMUSG0000000381 | chr8 | Rad23a | 0.012964583 | 0.977135836 |
| ENSMUSG0000005173 | chr7 | Rinl | -0.044729602 | 0.977135836 |
| ENSMUSG0000007860 | chr11 | 1810010H24Rik | 0.024500621 | 0.977135836 |
| ENSMUSG0000002581 | chr8 | Pard3 | 0.015669373 | 0.977135836 |
| ENSMUSG0000005606 | chr15 | Otulinl | 0.034619793 | 0.977135836 |
| ENSMUSG0000003539 | chr10 | Klf16 | 0.023806708 | 0.977332471 |
| ENSMUSG0000005384 | chr18 | Lipg | 0.049664701 | 0.977378471 |
| ENSMUSG0000002012 | chr11 | Plek | -0.019082791 | 0.977378471 |
| ENSMUSG0000003770 | chr9 | Tecta | 0.032649816 | 0.977378471 |
| ENSMUSG0000002951 | chr5 | Ulk1 | 0.016187034 | 0.977378471 |
| ENSMUSG0000000205 | chr11 | Unc119 | 0.020516739 | 0.977378471 |
| ENSMUSG0000003985 | chr4 | Rere | -0.02548485 | 0.977398491 |

| | | | | |
|---|---|---|---|---|
| ENSMUSG0000010048 | chr14 | Gm4131 | -0.083058662 | 0.977398491 |
| ENSMUSG0000000034 | chr15 | Dazap2 | -0.018451836 | 0.977398491 |
| ENSMUSG0000002824 | chr4 | Sdcbp | -0.013885463 | 0.977398491 |
| ENSMUSG0000002289 | chr16 | Adamts5 | 0.025564242 | 0.977398491 |
| ENSMUSG0000004777 | chr4 | Phf13 | -0.01769177 | 0.977398491 |
| ENSMUSG0000005314 | chr2 | Ptprt | 0.029788564 | 0.977398491 |
| ENSMUSG0000003537 | chr10 | Gm49322 | 0.064050495 | 0.977398491 |
| ENSMUSG0000005336 | chr5 | Rxfp2 | -0.080200338 | 0.977450881 |
| ENSMUSG0000002117 | chr12 | Nrde2 | 0.01998312 | 0.977562094 |
| ENSMUSG0000005186 | chr15 | Tbc1d22a | 0.024720516 | 0.977562094 |
| ENSMUSG0000005446 | chr17 | Lclat1 | -0.017557782 | 0.977666997 |
| ENSMUSG0000002140 | chr13 | Wrnip1 | 0.015621754 | 0.977666997 |
| ENSMUSG0000005432 | chr18 | Taf4b | 0.021933434 | 0.977774189 |
| ENSMUSG0000000486 | chr17 | Srpk1 | 0.013227038 | 0.978010141 |
| ENSMUSG0000007206 | chr13 | Zfp1008 | 0.025671163 | 0.97808883 |
| ENSMUSG0000004061 | chr14 | Pck2 | -0.017429963 | 0.978101623 |
| ENSMUSG0000002252 | chr16 | Fgf12 | -0.026677272 | 0.978101623 |
| ENSMUSG0000003468 | chr9 | Sema3f | -0.014830372 | 0.978101623 |
| ENSMUSG0000002803 | chr3 | Fubp1 | 0.01235752 | 0.978101623 |
| ENSMUSG0000004969 | chr1 | Nckap5 | 0.029161036 | 0.978181918 |
| ENSMUSG0000001891 | chr11 | Tm4sf5 | 0.068574645 | 0.978229289 |
| ENSMUSG0000009573 | chr7 | Vmn2r29 | -0.059775024 | 0.978229289 |
| ENSMUSG0000001694 | chr17 | Kctd5 | -0.011922272 | 0.978229289 |
| ENSMUSG0000002887 | chr4 | Fgr | 0.080354622 | 0.978229289 |
| ENSMUSG0000000155 | chr11 | Jup | -0.015046823 | 0.978229289 |
| ENSMUSG0000003835 | chr11 | Arl5c | 0.042530827 | 0.978229289 |
| ENSMUSG0000003754 | chr17 | H2-DMb2 | -0.070289737 | 0.978333789 |
| ENSMUSG0000002357 | chr2 | Ccndbp1 | -0.015113604 | 0.978338461 |
| ENSMUSG0000003562 | chr7 | Rsf1 | -0.015635582 | 0.978338461 |
| ENSMUSG0000002557 | chr11 | Tnrc6c | 0.019657291 | 0.978338461 |
| ENSMUSG0000002276 | chr16 | Lztr1 | 0.009135652 | 0.978338461 |
| ENSMUSG0000001518 | chr6 | Casd1 | 0.01134118 | 0.978338461 |
| ENSMUSG0000003567 | chr17 | Tnfsf9 | 0.045369234 | 0.978338461 |
| ENSMUSG0000005028 | chr11 | Fzd2 | 0.014318713 | 0.97864755 |
| ENSMUSG0000000513 | chr9 | 4930550C14Rik | -0.037622406 | 0.97873665 |
| ENSMUSG0000003187 | chr8 | Ces2g | -0.052716459 | 0.978832621 |
| ENSMUSG0000004402 | chr18 | Pcdhb21 | -0.034536118 | 0.978832621 |
| ENSMUSG0000003251 | chr9 | Wdr48 | -0.011554284 | 0.978832621 |
| ENSMUSG0000002649 | chr1 | Acbd3 | 0.010825415 | 0.978832621 |
| ENSMUSG0000009012 | chr4 | Pou3f1 | -0.057748006 | 0.978832621 |
| ENSMUSG0000006275 | chr17 | Smim29 | 0.020490895 | 0.978832621 |
| ENSMUSG0000001730 | chr2 | Acot8 | 0.019611351 | 0.978832621 |
| ENSMUSG0000006811 | chr15 | Ccdc134 | 0.021787878 | 0.978832621 |
| ENSMUSG0000004624 | chr5 | Pilra | 0.056919266 | 0.978832621 |
| ENSMUSG0000001530 | chr10 | Sash1 | 0.010156452 | 0.978924145 |
| ENSMUSG0000003964 | chr2 | Kyat1 | -0.021818331 | 0.979056956 |
| ENSMUSG0000002594 | chr1 | Pikfyve | -0.011823292 | 0.979056956 |
| ENSMUSG0000004487 | chr7 | Zfp444 | -0.022451934 | 0.97912471 |
| ENSMUSG0000006873 | chr2 | Trp53i11 | -0.017210232 | 0.97922735 |
| ENSMUSG0000004302 | chr3 | Dnai3 | 0.070162133 | 0.979326266 |

| | | | | |
|---|---|---|---|---|
| ENSMUSG0000001358 | chr9 | Aldh1a2 | 0.06503029 | 0.979345894 |
| ENSMUSG0000002229 | chr15 | Atp6v1c1 | 0.008675931 | 0.979490728 |
| ENSMUSG0000002895 | chr5 | Nub1 | -0.013594806 | 0.979649695 |
| ENSMUSG0000003556 | chr8 | Ankrd11 | 0.014893401 | 0.979670191 |
| ENSMUSG0000002266 | chr16 | Cd200r1 | -0.061022795 | 0.979792238 |
| ENSMUSG0000001502 | chr8 | Galns | -0.022107954 | 0.979792238 |
| ENSMUSG0000005753 | chr4 | Ece1 | -0.013333669 | 0.979792238 |
| ENSMUSG0000002406 | chr17 | Galnt14 | -0.035406156 | 0.979792238 |
| ENSMUSG0000000069 | chr6 | Loxl3 | -0.013208397 | 0.979792238 |
| ENSMUSG0000002957 | chr5 | Wipi2 | 0.011267486 | 0.979792238 |
| ENSMUSG0000001348 | chr11 | Card14 | -0.058030957 | 0.97982733 |
| ENSMUSG0000005373 | chr4 | Tmem39b | 0.021069059 | 0.979848402 |
| ENSMUSG0000007902 | chr15 | Slc45a4 | -0.015865251 | 0.979977751 |
| ENSMUSG0000003607 | chr4 | Galt | -0.014906225 | 0.979977751 |
| ENSMUSG0000002405 | chr17 | Wiz | 0.01331376 | 0.979977751 |
| ENSMUSG0000002844 | chr4 | Spmip6 | 0.048230056 | 0.979977751 |
| ENSMUSG0000003231 | chr9 | Tmem266 | 0.029846387 | 0.979977751 |
| ENSMUSG0000003591 | chr10 | Ptprq | -0.093528774 | 0.98007884 |
| ENSMUSG0000002895 | chr4 | Per3 | -0.116327878 | 0.980083896 |
| ENSMUSG0000003226 | chr9 | Htr3a | -0.015160212 | 0.980083896 |
| ENSMUSG0000010297 | chr1 | Zc3h11a | 0.021985994 | 0.980083896 |
| ENSMUSG0000005415 | chr12 | Syne3 | 0.020593913 | 0.980083896 |
| ENSMUSG0000003422 | chr11 | Foxj1 | 0.02477302 | 0.980083896 |
| ENSMUSG0000007057 | chr5 | Mn1 | 0.019334621 | 0.980083896 |
| ENSMUSG0000002243 | chr15 | Fam118a | 0.014834375 | 0.980135625 |
| ENSMUSG0000005178 | chr15 | Tubgcp6 | 0.015681994 | 0.980155853 |
| ENSMUSG0000002762 | chr2 | Rbm39 | -0.006804806 | 0.980183698 |
| ENSMUSG0000003615 | chr1 | Mgat5 | -0.016294867 | 0.980304052 |
| ENSMUSG0000001984 | chr10 | Fyn | -0.010277211 | 0.980304052 |
| ENSMUSG0000005150 | chr7 | Siglech | 0.042939159 | 0.980304052 |
| ENSMUSG0000002039 | chr11 | Havcr2 | 0.067424716 | 0.980304052 |
| ENSMUSG0000003128 | chr3 | Glt28d2 | 0.038527541 | 0.980314342 |
| ENSMUSG0000001270 | chr8 | Retn | -0.097473185 | 0.980367618 |
| ENSMUSG0000002455 | chr18 | Galr1 | -0.023594496 | 0.980367618 |
| ENSMUSG0000009141 | chr10 | Ak9 | -0.048586117 | 0.980367618 |
| ENSMUSG0000002412 | chr17 | Lrpprc | -0.015074746 | 0.980367618 |
| ENSMUSG0000002091 | chr11 | Adprm | -0.016748689 | 0.980367618 |
| ENSMUSG0000005576 | chr17 | Gemin6 | -0.024763143 | 0.980367618 |
| ENSMUSG0000002845 | chr4 | Fancg | -0.030334611 | 0.980367618 |
| ENSMUSG0000003199 | chr9 | Snx19 | -0.010653683 | 0.980367618 |
| ENSMUSG0000002270 | chr16 | Zbtb20 | 0.027020189 | 0.980367618 |
| ENSMUSG0000002150 | chr13 | Caml | 0.01680159 | 0.980367618 |
| ENSMUSG0000002427 | chr18 | Elp2 | 0.00988655 | 0.980367618 |
| ENSMUSG0000000708 | chr5 | Pole | 0.057427194 | 0.980367618 |
| ENSMUSG0000000182 | chr7 | Folr1 | 0.050889314 | 0.980367618 |
| ENSMUSG0000009207 | chr17 | Dynlt1a | 0.0264319 | 0.980367618 |
| ENSMUSG0000005339 | chr17 | Zfp952 | -0.013576731 | 0.980501642 |
| ENSMUSG0000002761 | chr2 | Eif6 | 0.019173182 | 0.980501642 |
| ENSMUSG0000003835 | chr9 | Camp | -0.074543939 | 0.980501642 |
| ENSMUSG0000003834 | chr1 | Plcl1 | 0.014527378 | 0.980501642 |

| | | | | |
|---|---|---|---|---|
| ENSMUSG0000011866 | chr12 | Ahnak2 | 0.023440744 | 0.980501642 |
| ENSMUSG0000001910 | chr11 | Aldh3a1 | 0.031210493 | 0.980501642 |
| ENSMUSG0000005006 | chr1 | Grem2 | 0.052571813 | 0.980501642 |
| ENSMUSG0000004065 | chr17 | Dnph1 | 0.046397416 | 0.980501642 |
| ENSMUSG0000004114 | chr5 | Brca2 | -0.026483545 | 0.98055408 |
| ENSMUSG0000000937 | chr19 | Slc16a12 | -0.033684416 | 0.98055408 |
| ENSMUSG0000004044 | chr11 | Spns2 | 0.024845445 | 0.980622274 |
| ENSMUSG0000003017 | chr6 | Wnt5b | 0.031422039 | 0.980622274 |
| ENSMUSG0000003905 | chr2 | Hrh3 | -0.044171659 | 0.980622274 |
| ENSMUSG0000004305 | chr8 | Disc1 | 0.036116841 | 0.980622274 |
| ENSMUSG0000005761 | chr5 | Gnai1 | 0.014908643 | 0.981305327 |
| ENSMUSG0000009162 | chr6 | Lsm5 | -0.022809643 | 0.981420116 |
| ENSMUSG0000006017 | chr13 | Kif27 | -0.023096998 | 0.981531092 |
| ENSMUSG0000005236 | chr15 | Tmem106c | 0.015094154 | 0.981531092 |
| ENSMUSG0000003452 | chr14 | Zfp395 | -0.015650192 | 0.981573029 |
| ENSMUSG0000003604 | chr15 | 5031439G07Rik | 0.012474646 | 0.981573029 |
| ENSMUSG0000002205 | chr14 | Bnip3l | 0.016732118 | 0.981645472 |
| ENSMUSG0000004279 | chr1 | Lgr6 | 0.055829876 | 0.981645472 |
| ENSMUSG0000000000 | chr3 | Gnai3 | -0.010360927 | 0.981820045 |
| ENSMUSG0000003432 | chr14 | Kctd9 | -0.017202788 | 0.981978983 |
| ENSMUSG0000003862 | chr15 | Med30 | -0.018567454 | 0.981978983 |
| ENSMUSG0000006987 | chr11 | Irgm2 | -0.025900916 | 0.981978983 |
| ENSMUSG0000000315 | chr6 | Foxj2 | -0.011016001 | 0.981978983 |
| ENSMUSG0000000144 | chr11 | Mrpl10 | -0.013699587 | 0.981978983 |
| ENSMUSG0000003598 | chr18 | Nme5 | -0.015822608 | 0.981978983 |
| ENSMUSG0000000616 | chr11 | Clint1 | -0.008772765 | 0.981978983 |
| ENSMUSG0000002861 | chr4 | Tmem59 | 0.01518257 | 0.981978983 |
| ENSMUSG0000004363 | chr19 | Rbm20 | 0.019176228 | 0.981978983 |
| ENSMUSG0000003239 | chr9 | Parp16 | 0.016231081 | 0.981978983 |
| ENSMUSG0000003027 | chr6 | Ogg1 | 0.012319302 | 0.981978983 |
| ENSMUSG0000004450 | chr17 | Zfp758 | 0.013243001 | 0.981978983 |
| ENSMUSG0000009158 | chr17 | Cyp4f17 | 0.018092975 | 0.981978983 |
| ENSMUSG0000002025 | chr10 | Txnrd1 | 0.011647862 | 0.981978983 |
| ENSMUSG0000007959 | chrX | 4933416I08Rik | -0.025764665 | 0.981978983 |
| ENSMUSG0000006345 | chr14 | Lrmda | -0.055869867 | 0.981999669 |
| ENSMUSG0000001823 | chr11 | Zcchc10 | 0.023074837 | 0.982005937 |
| ENSMUSG0000004565 | chr18 | Pcdhb10 | 0.03278718 | 0.98203433 |
| ENSMUSG0000004033 | chr7 | Nsmce4a | -0.014441919 | 0.98203433 |
| ENSMUSG0000007522 | chr16 | Mix23 | 0.017707694 | 0.98203433 |
| ENSMUSG0000005520 | chr17 | Zfp811 | 0.016973419 | 0.98203433 |
| ENSMUSG0000003240 | chr9 | Pias1 | -0.012255173 | 0.982143556 |
| ENSMUSG0000004290 | chrX | Foxo4 | -0.012832367 | 0.982167076 |
| ENSMUSG0000002663 | chr1 | Tatdn3 | 0.014715792 | 0.982167076 |
| ENSMUSG0000004130 | chr7 | Nkx6-2 | 0.052566149 | 0.982167076 |
| ENSMUSG0000001602 | chr2 | Lbp | 0.014474129 | 0.982167076 |
| ENSMUSG0000006600 | chr4 | Zfp979 | -0.025594795 | 0.98217052 |
| ENSMUSG0000004000 | chr5 | Magi2 | -0.02059296 | 0.98217052 |
| ENSMUSG0000001584 | chr1 | Rxrg | -0.029926513 | 0.98217052 |
| ENSMUSG0000003690 | chr18 | Fzd8 | -0.016172643 | 0.98217052 |
| ENSMUSG0000001999 | chr10 | Mtfr2 | -0.106074641 | 0.98217052 |

| | | | | |
|---|---|---|---|---|
| ENSMUSG0000003698 | chr5 | Taf6 | -0.013732893 | 0.98217052 |
| ENSMUSG0000003963 | chr1 | Hnrnpu | -0.009894412 | 0.98217052 |
| ENSMUSG0000005785 | chr19 | Fam204a | 0.012598115 | 0.98217052 |
| ENSMUSG0000004396 | chr4 | Thrap3 | 0.009262411 | 0.98217052 |
| ENSMUSG0000003925 | chr11 | Fn3krp | 0.013507362 | 0.98217052 |
| ENSMUSG0000003235 | chr9 | Ctsh | 0.017504253 | 0.98217052 |
| ENSMUSG0000005999 | chr5 | Nptx2 | 0.032460706 | 0.98217052 |
| ENSMUSG0000010929 | chr3 | Dcst2 | 0.102954911 | 0.98217052 |
| ENSMUSG0000003805 | chr16 | Dexi | -0.012990174 | 0.982568558 |
| ENSMUSG0000002879 | chr4 | Rnf19b | -0.011137984 | 0.982688878 |
| ENSMUSG0000002178 | chr14 | Dlg5 | -0.012798268 | 0.982701018 |
| ENSMUSG0000002089 | chr11 | Per1 | 0.062686479 | 0.982728782 |
| ENSMUSG0000002868 | chr4 | Mutyh | -0.035554461 | 0.982728782 |
| ENSMUSG0000002289 | chr16 | Dyrk1a | -0.011384144 | 0.982728782 |
| ENSMUSG0000006135 | chr6 | Cxcl12 | -0.010804591 | 0.982728782 |
| ENSMUSG0000002735 | chr2 | Mcm8 | 0.028781746 | 0.982728782 |
| ENSMUSG0000007643 | chr2 | Wfdc3 | -0.062870403 | 0.982728782 |
| ENSMUSG0000005859 | chr2 | Fbh1 | -0.008659304 | 0.982728782 |
| ENSMUSG0000003653 | chrX | Rnf113a1 | -0.020164028 | 0.982728782 |
| ENSMUSG0000004062 | chr11 | Dhx33 | -0.011130882 | 0.982728782 |
| ENSMUSG0000004467 | chr5 | Fzd1 | 0.01299436 | 0.982728782 |
| ENSMUSG0000002810 | chr3 | Pex11b | 0.012121432 | 0.982728782 |
| ENSMUSG0000002459 | chr18 | Prrc1 | 0.011595946 | 0.982728782 |
| ENSMUSG0000001740 | chr11 | Nek8 | 0.016279251 | 0.982728782 |
| ENSMUSG0000002192 | chr14 | Itih4 | 0.05551707 | 0.982728782 |
| ENSMUSG0000002878 | chr4 | Azin2 | 0.01916021 | 0.982734282 |
| ENSMUSG0000003962 | chr8 | Trmt9b | -0.021445464 | 0.982860504 |
| ENSMUSG0000002128 | chr12 | Klc1 | -0.012512861 | 0.982860504 |
| ENSMUSG0000001792 | chr2 | B4galt5 | 0.010134742 | 0.982860504 |
| ENSMUSG0000006967 | chr6 | Pcgf1 | -0.016138951 | 0.982919005 |
| ENSMUSG0000005268 | chr1 | Rab7b | -0.024574515 | 0.982944917 |
| ENSMUSG0000004240 | chr19 | Crtac1 | -0.01990034 | 0.982944917 |
| ENSMUSG0000002982 | chr6 | Spmip4 | 0.028166156 | 0.982944917 |
| ENSMUSG0000009892 | chr1 | Tmem185b | -0.018359471 | 0.98313857 |
| ENSMUSG0000000460 | chr7 | Cd33 | -0.022311409 | 0.983203423 |
| ENSMUSG0000006368 | chr18 | Pcdhb5 | -0.023847701 | 0.983203423 |
| ENSMUSG0000002494 | chr19 | Sf1 | -0.010366428 | 0.983203423 |
| ENSMUSG0000003212 | chr9 | Dpagt1 | -0.011000702 | 0.983203423 |
| ENSMUSG0000003243 | chr9 | Dync1li1 | 0.010208061 | 0.983203423 |
| ENSMUSG0000002029 | chr11 | Xpo1 | 0.009995174 | 0.983203423 |
| ENSMUSG0000003033 | chr6 | Ing4 | 0.035460153 | 0.983238294 |
| ENSMUSG0000002900 | chr5 | Kmt2e | 0.017852672 | 0.983238294 |
| ENSMUSG0000002498 | chr19 | Zdhhc6 | 0.011541089 | 0.983238294 |
| ENSMUSG0000002055 | chr11 | Pctp | -0.019797065 | 0.983289821 |
| ENSMUSG0000003154 | chr8 | Kat6a | -0.013089744 | 0.983289821 |
| ENSMUSG0000002462 | chr18 | Pdgfrb | 0.018874203 | 0.983289821 |
| ENSMUSG0000002207 | chr14 | Rhobtb2 | 0.017381625 | 0.983289821 |
| ENSMUSG0000011866 | chr5 | Tctn2 | 0.013140823 | 0.983289821 |
| ENSMUSG0000002214 | chr15 | Fyb1 | 0.024606133 | 0.983289821 |
| ENSMUSG0000003344 | chr10 | Specc1l | 0.009272232 | 0.983289821 |

| | | | | |
|---|---|---|---|---|
| ENSMUSG0000005364 | chr5 | Gper1 | 0.022066075 | 0.983289821 |
| ENSMUSG0000003702 | chr8 | Mmaa | 0.016259364 | 0.983300833 |
| ENSMUSG0000002917 | chr5 | Pgm2 | -0.013700558 | 0.983300833 |
| ENSMUSG0000002798 | chr3 | Cyp2u1 | 0.018367636 | 0.983300833 |
| ENSMUSG0000003951 | chr17 | Cdsn | -0.05990894 | 0.983469034 |
| ENSMUSG0000005137 | chr2 | Flrt3 | -0.016259737 | 0.983469034 |
| ENSMUSG0000004962 | chr9 | Slc17a5 | -0.011666117 | 0.983469034 |
| ENSMUSG0000009206 | chr5 | Bend4 | -0.020995382 | 0.983469034 |
| ENSMUSG0000006715 | chr17 | Xpo5 | 0.009352487 | 0.983519136 |
| ENSMUSG0000003418 | chr11 | Nsf | -0.008899572 | 0.983700165 |
| ENSMUSG0000007486 | chr13 | Zfp808 | 0.018164605 | 0.983700165 |
| ENSMUSG0000002752 | chr2 | Stx16 | -0.009723564 | 0.983760588 |
| ENSMUSG0000002899 | chr4 | Mtor | -0.009737574 | 0.983760588 |
| ENSMUSG0000002060 | chr12 | Hs1bp3 | 0.011167335 | 0.983760588 |
| ENSMUSG0000009462 | chr10 | Gm3055 | 0.049909779 | 0.983760588 |
| ENSMUSG0000001038C | chr18 | Pcdha8 | 0.055283585 | 0.983772581 |
| ENSMUSG0000003859 | chr5 | Tctn1 | -0.017075858 | 0.983836897 |
| ENSMUSG0000001635 | chr2 | Col20a1 | -0.017774126 | 0.983836897 |
| ENSMUSG0000002044 | chr11 | 2310033P09Rik | -0.016255735 | 0.983836897 |
| ENSMUSG0000004280 | chr1 | Hecw2 | -0.013882948 | 0.983836897 |
| ENSMUSG0000005081 | chr4 | Ecpas | -0.011017454 | 0.983836897 |
| ENSMUSG0000002184 | chr14 | Mapk1ip1l | -0.007437601 | 0.983836897 |
| ENSMUSG0000000245 | chr2 | Rgs19 | 0.019466966 | 0.983836897 |
| ENSMUSG0000002522 | chr19 | Oga | -0.007714639 | 0.983836897 |
| ENSMUSG0000004033 | chr3 | Eeig2 | 0.011200852 | 0.983836897 |
| ENSMUSG0000004236 | chr2 | Rbm45 | -0.014500467 | 0.983893885 |
| ENSMUSG0000005789 | chr7 | Zfp329 | -0.009493599 | 0.983893885 |
| ENSMUSG0000002480 | chr19 | Syvn1 | -0.007372449 | 0.984028657 |
| ENSMUSG0000002142 | chr13 | Riok1 | 0.008999975 | 0.984028657 |
| ENSMUSG0000002203 | chr14 | Esco2 | 0.094938084 | 0.984312144 |
| ENSMUSG0000006145 | chr19 | Tmem151a | -0.027850776 | 0.984312144 |
| ENSMUSG0000002643 | chr1 | Cdk18 | -0.020728381 | 0.984312144 |
| ENSMUSG0000002784 | chr3 | Syt6 | -0.019845705 | 0.984312144 |
| ENSMUSG0000008696 | chr11 | Faxdc2 | 0.068839495 | 0.984312144 |
| ENSMUSG0000007426 | chr9 | Rec114 | -0.039456684 | 0.984312144 |
| ENSMUSG0000000225 | chr17 | Def6 | -0.028422397 | 0.984312144 |
| ENSMUSG0000005276 | chr1 | A630001G21Rik | -0.04200806 | 0.984312144 |
| ENSMUSG0000002427 | chr18 | Mapre2 | -0.013461609 | 0.984312144 |
| ENSMUSG0000002841 | chr4 | Chmp5 | -0.014035176 | 0.984312144 |
| ENSMUSG0000003889 | chr11 | Fam117a | -0.013704147 | 0.984312144 |
| ENSMUSG0000002300 | chr15 | Nckap5l | -0.01593072 | 0.984312144 |
| ENSMUSG0000004118 | chr7 | Prkd2 | -0.014198558 | 0.984312144 |
| ENSMUSG0000001850 | chr11 | Cenpv | -0.015563922 | 0.984312144 |
| ENSMUSG0000003983 | chr10 | Nhsl1 | -0.015521305 | 0.984312144 |
| ENSMUSG0000002217 | chr14 | 4931414P19Rik | -0.021974915 | 0.984312144 |
| ENSMUSG0000002237 | chr15 | Sla | -0.031063921 | 0.984312144 |
| ENSMUSG0000002394 | chr17 | Slc29a1 | -0.011043487 | 0.984312144 |
| ENSMUSG0000005360 | chr17 | Zfp472 | -0.018144608 | 0.984312144 |
| ENSMUSG0000004849 | chr11 | Mmgt2 | -0.010396826 | 0.984312144 |
| ENSMUSG0000002810 | chr3 | Rprd2 | -0.013884083 | 0.984312144 |

| | | | | |
|---|---|---|---|---|
| ENSMUSG0000002732 | chr2 | Rpusd2 | -0.018190251 | 0.984312144 |
| ENSMUSG0000006157 | chr8 | Adgrg5 | -0.052615535 | 0.984312144 |
| ENSMUSG000001039( | chr15 | Tigd5 | -0.015796076 | 0.984312144 |
| ENSMUSG000000602( | chr4 | Zfp462 | -0.016393893 | 0.984312144 |
| ENSMUSG0000004213 | chr9 | Msantd2 | -0.010163486 | 0.984312144 |
| ENSMUSG0000002002 | chr10 | Fgd6 | -0.011351531 | 0.984312144 |
| ENSMUSG000000791€ | chr1 | Tlr5 | 0.022358464 | 0.984312144 |
| ENSMUSG0000004184 | chr12 | Ppp4r3a | -0.010096213 | 0.984312144 |
| ENSMUSG0000004229 | chr11 | Ttc19 | -0.010401391 | 0.984312144 |
| ENSMUSG000000266€ | chr1 | Atf6 | -0.009926014 | 0.984312144 |
| ENSMUSG000000292( | chr5 | Pds5a | -0.008269829 | 0.984312144 |
| ENSMUSG0000000232 | chr14 | Tm9sf1 | -0.012157098 | 0.984312144 |
| ENSMUSG0000004457 | chr5 | Garem2 | 0.0348049 | 0.984312144 |
| ENSMUSG0000000423 | chr3 | Wars2 | -0.016586082 | 0.984312144 |
| ENSMUSG0000005215 | chr2 | Acvr2a | -0.007895679 | 0.984312144 |
| ENSMUSG0000003231 | chr9 | Clk3 | -0.007762134 | 0.984312144 |
| ENSMUSG000000388( | chr1 | Sde2 | -0.008668265 | 0.984312144 |
| ENSMUSG0000002484 | chr19 | Eif1ad | -0.008882741 | 0.984312144 |
| ENSMUSG0000001692 | chr2 | Srsf6 | -0.008609764 | 0.984312144 |
| ENSMUSG0000000147 | chr8 | Tcf25 | -0.006559168 | 0.984312144 |
| ENSMUSG000000376( | chr10 | Bclaf1 | 0.011690486 | 0.984312144 |
| ENSMUSG0000004232 | chr12 | Prox2 | 0.021337672 | 0.984312144 |
| ENSMUSG000000306€ | chr7 | Pik3c2a | 0.011667086 | 0.984312144 |
| ENSMUSG0000004075 | chr5 | Lat2 | 0.037385301 | 0.984312144 |
| ENSMUSG0000009241 | chr7 | Zfp141 | 0.012258164 | 0.984312144 |
| ENSMUSG0000005347 | chr18 | Tcf4 | 0.011419284 | 0.984312144 |
| ENSMUSG0000002628 | chr1 | Ing5 | 0.010676912 | 0.984312144 |
| ENSMUSG000000325€ | chr9 | Dnajc13 | 0.007502418 | 0.984312144 |
| ENSMUSG000000292( | chr5 | Guf1 | 0.012219203 | 0.984312144 |
| ENSMUSG0000002819 | chr3 | Ccn1 | 0.043754058 | 0.984312144 |
| ENSMUSG000000488( | chr11 | Slc35e4 | 0.01123681 | 0.984312144 |
| ENSMUSG0000002201 | chr14 | Tdrd3 | 0.008437726 | 0.984312144 |
| ENSMUSG000000417( | chr14 | Tmem273 | -0.053224449 | 0.984312144 |
| ENSMUSG0000002539 | chr10 | Prim1 | 0.015812177 | 0.984312144 |
| ENSMUSG0000000294 | chr8 | Map2k7 | 0.008721047 | 0.984312144 |
| ENSMUSG0000002589 | chr9 | Alkbh8 | 0.010636192 | 0.984312144 |
| ENSMUSG0000002154 | chr13 | Ccnh | 0.010553055 | 0.984312144 |
| ENSMUSG000000290€ | chr4 | Ccnl2 | 0.011876185 | 0.984312144 |
| ENSMUSG0000004208 | chr13 | Arsb | 0.009148871 | 0.984312144 |
| ENSMUSG0000002287 | chr16 | Adipoq | 0.05055737 | 0.984312144 |
| ENSMUSG0000003934 | chr16 | Mettl22 | 0.01437482 | 0.984312144 |
| ENSMUSG0000006285 | chr17 | Tcp11 | 0.052593083 | 0.984312144 |
| ENSMUSG0000006615 | chr4 | Slc31a1 | 0.012239312 | 0.984312144 |
| ENSMUSG0000002394 | chr17 | Nfkbie | 0.017384369 | 0.984312144 |
| ENSMUSG0000009147 | chr14 | Gm3739 | 0.037085009 | 0.984312144 |
| ENSMUSG0000006983 | chr11 | Sat2 | 0.01832514 | 0.984312144 |
| ENSMUSG0000004028 | chr10 | Stac3 | 0.035681635 | 0.984312144 |
| ENSMUSG0000004714 | chr4 | Dmrta2 | -0.076738986 | 0.984312144 |
| ENSMUSG0000005124 | chr9 | Islr2 | -0.016280408 | 0.984492248 |
| ENSMUSG0000002784 | chr3 | Olfml3 | 0.01397211 | 0.984492248 |

| | | | | |
|---|---|---|---|---|
| ENSMUSG0000003693 | chrX | Aifm1 | 0.011339296 | 0.984492248 |
| ENSMUSG0000003231 | chr9 | Isl2 | 0.028333098 | 0.984492248 |
| ENSMUSG0000010167 | chr5 | Gm29609 | 0.040003787 | 0.984492248 |
| ENSMUSG0000003520 | chr9 | Lars2 | -0.059013706 | 0.984492248 |
| ENSMUSG0000001930 | chr11 | Psmc3ip | 0.017236199 | 0.984635991 |
| ENSMUSG0000010275 | chr3 | Naaladl2 | -0.033322346 | 0.984992981 |
| ENSMUSG0000004532 | chr3 | Cenpe | 0.056215543 | 0.984992981 |
| ENSMUSG0000002520 | chr19 | Mrpl43 | -0.0181209 | 0.984992981 |
| ENSMUSG0000004647 | chr2 | Sox18 | 0.024619024 | 0.984992981 |
| ENSMUSG0000003720 | chr9 | Islr | -0.006902813 | 0.984992981 |
| ENSMUSG0000004526 | chr4 | Zfp691 | 0.017571791 | 0.984992981 |
| ENSMUSG0000004309 | chr8 | Zfp866 | 0.007053796 | 0.985026179 |
| ENSMUSG0000003265 | chr6 | Crebl2 | -0.012962449 | 0.985091503 |
| ENSMUSG0000003984 | chr18 | Epg5 | -0.011342151 | 0.985091503 |
| ENSMUSG0000002467 | chr19 | Ms4a4d | -0.021066932 | 0.985091503 |
| ENSMUSG0000003225 | chr9 | Lca5 | -0.014539825 | 0.985155022 |
| ENSMUSG0000002868 | chr4 | Ccdc163 | -0.018641499 | 0.985155022 |
| ENSMUSG0000002782 | chr3 | Mme | 0.045580866 | 0.985311497 |
| ENSMUSG0000005371 | chr9 | Dusp7 | -0.017845896 | 0.985311497 |
| ENSMUSG0000006141 | chr8 | Zcchc14 | -0.011121074 | 0.985311497 |
| ENSMUSG0000002466 | chr19 | Tmem138 | -0.015856205 | 0.985311497 |
| ENSMUSG0000002708 | chr2 | Med19 | -0.010909218 | 0.985311497 |
| ENSMUSG0000006183 | chr6 | Suclg2 | -0.011929442 | 0.985311497 |
| ENSMUSG0000003425 | chr12 | Flvcr2 | -0.025742455 | 0.985311497 |
| ENSMUSG0000000303 | chr4 | Hmgn2 | 0.016897157 | 0.985311497 |
| ENSMUSG0000002418 | chr17 | Luc7l | -0.006159215 | 0.985311497 |
| ENSMUSG0000002047 | chr11 | Pold2 | -0.008414062 | 0.985311497 |
| ENSMUSG0000002793 | chr3 | Crtc2 | 0.012285278 | 0.985311497 |
| ENSMUSG0000007223 | chr15 | Tuba1a | 0.010449805 | 0.985311497 |
| ENSMUSG0000003456 | chr10 | Washc4 | 0.00762905 | 0.985311497 |
| ENSMUSG0000000783 | chr7 | Aldh16a1 | -0.01610281 | 0.985311497 |
| ENSMUSG0000006395 | chr10 | Amd2 | 0.03812038 | 0.985311497 |
| ENSMUSG0000001508 | chr2 | Entpd2 | 0.012239701 | 0.985311497 |
| ENSMUSG0000003802 | chr6 | Mindy4 | 0.017020106 | 0.985311497 |
| ENSMUSG0000000646 | chr19 | Bbs1 | -0.011746248 | 0.985340741 |
| ENSMUSG0000005714 | chr2 | Dph6 | 0.010123591 | 0.98547479 |
| ENSMUSG0000007954 | chr17 | H2-DMb1 | -0.026731389 | 0.98548084 |
| ENSMUSG0000004436 | chr2 | Tmem74b | 0.048815847 | 0.98548084 |
| ENSMUSG0000002242 | chr15 | Josd1 | -0.009914848 | 0.985534289 |
| ENSMUSG0000007095 | chr2 | Rabepk | 0.011699069 | 0.985534289 |
| ENSMUSG0000007434 | chr3 | I830077J02Rik | -0.069599476 | 0.985605928 |
| ENSMUSG0000003611 | chr4 | Rpp25l | -0.016154707 | 0.985605928 |
| ENSMUSG0000007503 | chr16 | Nxpe3 | -0.014648034 | 0.985605928 |
| ENSMUSG0000004212 | chr6 | Rassf4 | -0.013230704 | 0.985605928 |
| ENSMUSG0000003157 | chr8 | Ash2l | -0.007714248 | 0.985605928 |
| ENSMUSG0000007340 | chr17 | Mucl3 | 0.060033356 | 0.985605928 |
| ENSMUSG0000003820 | chr8 | Fbxo8 | 0.011721711 | 0.985605928 |
| ENSMUSG0000002233 | chr15 | Eny2 | -0.008904301 | 0.985605928 |
| ENSMUSG0000002491 | chr19 | Fibp | 0.008678842 | 0.985605928 |
| ENSMUSG0000002109 | chr12 | Lrrc9 | 0.02065317 | 0.985641475 |

| | | | | |
|---|---|---|---|---|
| ENSMUSG0000004414 | chr2 | 1810024B03Rik | 0.026107249 | 0.985661248 |
| ENSMUSG0000003776 | chr2 | Actr5 | 0.010498997 | 0.985661248 |
| ENSMUSG0000005430 | chr12 | Cpsf3 | 0.007993262 | 0.985661248 |
| ENSMUSG0000002618 | chr1 | Igfbp5 | -0.016174745 | 0.985689627 |
| ENSMUSG0000001645 | chr2 | Wt1 | 0.03505683 | 0.985689627 |
| ENSMUSG0000002398 | chr17 | Gm20517 | -0.042303935 | 0.985689627 |
| ENSMUSG0000007886 | chr2 | Zfp931 | 0.016801063 | 0.985689627 |
| ENSMUSG0000002564 | chr9 | Atrip | -0.010233633 | 0.985689627 |
| ENSMUSG0000002022 | chr10 | Irak3 | -0.015200168 | 0.985689627 |
| ENSMUSG0000003800 | chr17 | Cramp1 | 0.014168481 | 0.985689627 |
| ENSMUSG0000003727 | chr11 | Gemin5 | -0.008146519 | 0.985689627 |
| ENSMUSG0000004567 | chr12 | Slc66a3 | 0.014930776 | 0.985689627 |
| ENSMUSG0000000403 | chr3 | Gstm7 | 0.016184277 | 0.985689627 |
| ENSMUSG0000005140 | chr7 | Ppp1r37 | 0.00816423 | 0.985689627 |
| ENSMUSG0000004794 | chr4 | Marcksl1 | -0.013121255 | 0.985689627 |
| ENSMUSG0000004514 | chr11 | Pigw | -0.017658144 | 0.985689627 |
| ENSMUSG0000001880 | chr11 | Smyd4 | 0.01467494 | 0.985689627 |
| ENSMUSG0000001989 | chr10 | Ccdc59 | 0.010856618 | 0.985689627 |
| ENSMUSG0000004052 | chr9 | Zfp609 | 0.012323914 | 0.985689627 |
| ENSMUSG0000001661 | chr15 | Nup50 | 0.008247153 | 0.985689627 |
| ENSMUSG0000000465 | chr6 | Aqp1 | 0.014457126 | 0.985689627 |
| ENSMUSG0000004969 | chr5 | Nkx3-2 | 0.016735936 | 0.985689627 |
| ENSMUSG0000003878 | chr5 | Smurf1 | -0.010187156 | 0.985840186 |
| ENSMUSG0000006664 | chr5 | Fbxl18 | -0.010313613 | 0.985840186 |
| ENSMUSG0000000463 | chr6 | Chn2 | -0.01442979 | 0.985840186 |
| ENSMUSG0000007955 | chr5 | Haus3 | -0.012892927 | 0.985840186 |
| ENSMUSG0000007040 | chr11 | Hs3st3b1 | -0.028545059 | 0.985840186 |
| ENSMUSG0000003917 | chr7 | Polg | -0.007479095 | 0.985840186 |
| ENSMUSG0000002756 | chr2 | Mrgbp | -0.012523193 | 0.985840186 |
| ENSMUSG0000006093 | chr10 | Tmem263 | 0.009344145 | 0.985840186 |
| ENSMUSG0000002107 | chr12 | Tmx1 | 0.00839094 | 0.985840186 |
| ENSMUSG0000000222 | chr6 | Rmnd5a | -0.00886063 | 0.985974344 |
| ENSMUSG0000003549 | chr4 | Tstd2 | -0.00849261 | 0.985974344 |
| ENSMUSG0000002897 | chr5 | Abcb1b | 0.050232872 | 0.986098455 |
| ENSMUSG0000001346 | chr2 | Nelfb | -0.009840234 | 0.986098455 |
| ENSMUSG0000003051 | chr7 | Apba2 | -0.012078863 | 0.986098455 |
| ENSMUSG0000007877 | chr7 | Fcgbpl1 | -0.052792652 | 0.986098455 |
| ENSMUSG0000002398 | chr17 | Taf8 | 0.008766039 | 0.986098455 |
| ENSMUSG0000006684 | chr1 | Hmcn1 | -0.013988276 | 0.986123803 |
| ENSMUSG0000002510 | chr7 | Bnc1 | 0.039200508 | 0.986123803 |
| ENSMUSG0000003123 | chrX | Magt1 | -0.011997267 | 0.986123803 |
| ENSMUSG0000002450 | chr18 | Dpysl3 | -0.008930105 | 0.986123803 |
| ENSMUSG0000010058 | chr7 | Vmn1r90 | -0.052195328 | 0.986123803 |
| ENSMUSG0000002223 | chr15 | Ankrd33b | 0.043154395 | 0.986123803 |
| ENSMUSG0000004433 | chr5 | Alkbh2 | 0.028494599 | 0.986123803 |
| ENSMUSG0000002904 | chr4 | Morn1 | -0.013259271 | 0.986166768 |
| ENSMUSG0000002657 | chr1 | Ccdc181 | -0.010983658 | 0.986195651 |
| ENSMUSG0000006403 | chr18 | Mro | -0.022294309 | 0.98640842 |
| ENSMUSG0000002461 | chr18 | Camk2a | 0.014317556 | 0.986482172 |
| ENSMUSG0000000193 | chr6 | Vwf | -0.020166618 | 0.986603012 |

| | | | | |
|---|---|---|---|---|
| ENSMUSG0000002231 | chr15 | Utp23 | -0.01502686 | 0.986603012 |
| ENSMUSG0000004215 | chr14 | Dzip1 | 0.011659983 | 0.986603012 |
| ENSMUSG0000006963 | chr8 | Pex11g | 0.019302955 | 0.986603012 |
| ENSMUSG0000005621 | chr1 | R3hdm1 | 0.008705913 | 0.986691332 |
| ENSMUSG0000002611 | chr1 | Tmem131 | -0.009281479 | 0.986720615 |
| ENSMUSG0000003601 | chr10 | Tmtc2 | 0.012440655 | 0.986720615 |
| ENSMUSG0000003963 | chr8 | Lonrf1 | -0.03004445 | 0.986837742 |
| ENSMUSG0000002745 | chr2 | Acss1 | -0.011177257 | 0.986837742 |
| ENSMUSG0000003881 | chr11 | Gngt2 | 0.023548942 | 0.986837742 |
| ENSMUSG0000004278 | chr4 | Cimip2b | 0.032728551 | 0.986837742 |
| ENSMUSG0000004111 | chr13 | Elmo1 | 0.015376533 | 0.986837742 |
| ENSMUSG0000001985 | chr10 | Fam184a | -0.009606428 | 0.986837742 |
| ENSMUSG0000001586 | chr11 | Prpsap1 | 0.010610575 | 0.986837742 |
| ENSMUSG0000004635 | chr13 | Zfp322a | 0.007299074 | 0.986837742 |
| ENSMUSG0000005967 | chr14 | Cdh24 | -0.018269146 | 0.986837742 |
| ENSMUSG0000005003 | chr10 | Fhl4 | 0.037155189 | 0.986837742 |
| ENSMUSG0000002175 | chr14 | Fam107a | -0.009080784 | 0.98687651 |
| ENSMUSG0000004032 | chr17 | Cul9 | 0.01025055 | 0.98687651 |
| ENSMUSG0000003426 | chr12 | Batf | -0.037945032 | 0.986884946 |
| ENSMUSG0000000577 | chr3 | Rfx5 | 0.0124334 | 0.986884946 |
| ENSMUSG0000003866 | chr4 | Lpar1 | -0.005591099 | 0.986884946 |
| ENSMUSG0000003386 | chr8 | Cdk10 | -0.008472672 | 0.986884946 |
| ENSMUSG0000012191 | chrX | Sts | 0.01833664 | 0.986884946 |
| ENSMUSG0000006826 | chr6 | Efcc1 | 0.019313656 | 0.987044596 |
| ENSMUSG0000002738 | chr2 | Zc3h8 | 0.012878042 | 0.987404543 |
| ENSMUSG0000002473 | chr19 | Slc15a3 | 0.027243615 | 0.987426581 |
| ENSMUSG0000006338 | chr17 | Zfp947 | 0.021100358 | 0.987426581 |
| ENSMUSG0000003384 | chr7 | Pla2g4c | 0.054001146 | 0.987426581 |
| ENSMUSG0000003499 | chr17 | Foxn2 | -0.01183669 | 0.987489581 |
| ENSMUSG0000005789 | chr17 | Adgrf2 | 0.040788084 | 0.987551502 |
| ENSMUSG0000009908 | chr15 | Atf7 | -0.010995859 | 0.987697211 |
| ENSMUSG0000003064 | chr7 | Ddias | 0.032061484 | 0.987697211 |
| ENSMUSG0000001397 | chr1 | Dedd | 0.012214975 | 0.987697211 |
| ENSMUSG0000003244 | chr9 | Zcwpw2 | -0.041747532 | 0.987697211 |
| ENSMUSG0000000709 | chr1 | Atp1a2 | -0.015938508 | 0.987713199 |
| ENSMUSG0000003579 | chr16 | Cep19 | -0.011474729 | 0.987713199 |
| ENSMUSG0000002674 | chr2 | Plxdc2 | 0.013861385 | 0.987713199 |
| ENSMUSG0000008560 | chr7 | Meiosin | -0.02413241 | 0.987713199 |
| ENSMUSG0000002906 | chr4 | Gnb1 | -0.00732589 | 0.987713199 |
| ENSMUSG0000002513 | chr11 | Alyref | 0.010210686 | 0.987713199 |
| ENSMUSG0000000508 | chr2 | Slc1a2 | 0.016894017 | 0.987713199 |
| ENSMUSG0000003061 | chr7 | Ccdc90b | 0.009852271 | 0.987713199 |
| ENSMUSG0000006102 | chr1 | Rrs1 | 0.011256173 | 0.987713199 |
| ENSMUSG0000002040 | chr11 | Tnip1 | 0.008290471 | 0.987713199 |
| ENSMUSG0000007902 | chr15 | Col22a1 | -0.032229893 | 0.987843322 |
| ENSMUSG0000006224 | chr13 | Cks2 | -0.036202108 | 0.988028936 |
| ENSMUSG0000005537 | chr2 | Stam2 | 0.008018589 | 0.988028936 |
| ENSMUSG0000006130 | chr11 | Slc38a10 | -0.006894783 | 0.988063786 |
| ENSMUSG0000005709 | chr7 | Zfp607b | -0.015287895 | 0.988063786 |
| ENSMUSG0000000009 | chr11 | Tbx2 | 0.014144707 | 0.988063786 |

| | | | | |
|---|---|---|---|---|
| ENSMUSG000000040 | chr4 | Lck | 0.030974897 | 0.988063786 |
| ENSMUSG000002019 | chr10 | Cabin1 | -0.005817857 | 0.988188087 |
| ENSMUSG000005793 | chr9 | Gsta2 | -0.024404119 | 0.988192741 |
| ENSMUSG000004989 | chr11 | Rasd1 | -0.022952733 | 0.988192741 |
| ENSMUSG000007533 | chr16 | A930003A15Rik | 0.029155699 | 0.988192741 |
| ENSMUSG000007106 | chr8 | Zfp827 | -0.010046961 | 0.988192741 |
| ENSMUSG000002048 | chr11 | Dynll2 | -0.008134866 | 0.988192741 |
| ENSMUSG000003493 | chr11 | Dhx8 | -0.007423468 | 0.988192741 |
| ENSMUSG000003119 | chrX | F8 | 0.018343145 | 0.988192741 |
| ENSMUSG000004619 | chr6 | Iqub | 0.014192705 | 0.988192741 |
| ENSMUSG000002080 | chr11 | Rhbdf2 | 0.01446023 | 0.988192741 |
| ENSMUSG000003873 | chr15 | Nudcd1 | 0.0094145 | 0.988192741 |
| ENSMUSG000002393 | chr17 | Cdc5l | 0.009148235 | 0.988192741 |
| ENSMUSG000003379 | chr13 | Tasor2 | 0.008321788 | 0.988192741 |
| ENSMUSG0000002731 | chr2 | Atrn | -0.008903887 | 0.988288318 |
| ENSMUSG000004038 | chr14 | Cbln3 | 0.066216915 | 0.988423826 |
| ENSMUSG000003536 | chr10 | Parpbp | 0.047302115 | 0.988423826 |
| ENSMUSG000002592 | chr1 | Rdh10 | -0.013863542 | 0.988423826 |
| ENSMUSG0000002471 | chr19 | Pcsk5 | -0.018806291 | 0.988423826 |
| ENSMUSG000002150 | chr13 | Pitx1 | 0.085792752 | 0.988423826 |
| ENSMUSG000003603 | chr9 | Prtg | 0.028892467 | 0.988423826 |
| ENSMUSG000004498 | chr18 | Sft2d3 | -0.011194144 | 0.988423826 |
| ENSMUSG000002908 | chr5 | 5730480H06Rik | -0.010690443 | 0.988423826 |
| ENSMUSG000004254 | chr2 | Asxl1 | 0.007809855 | 0.988423826 |
| ENSMUSG000002545 | chr13 | Nnt | 0.027960288 | 0.988423826 |
| ENSMUSG000002853 | chr4 | Jak1 | 0.004152686 | 0.988423826 |
| ENSMUSG000007416 | chr7 | AW146154 | 0.012716076 | 0.988423826 |
| ENSMUSG000003613 | chr15 | Hoxc9 | -0.028480363 | 0.988423826 |
| ENSMUSG000002113 | chr12 | Susd6 | 0.006176359 | 0.988423826 |
| ENSMUSG000005174 | chr2 | Ttn | 0.028676143 | 0.988423826 |
| ENSMUSG000003500 | chr2 | Dpp4 | 0.030533974 | 0.988423826 |
| ENSMUSG000003737 | chr1 | Hhat | -0.016105482 | 0.988468015 |
| ENSMUSG000002598 | chr1 | Hspd1 | -0.010650551 | 0.988496579 |
| ENSMUSG000002700 | chr2 | Frzb | -0.01520445 | 0.988496579 |
| ENSMUSG000002234 | chr15 | Myc | 0.015046576 | 0.988496579 |
| ENSMUSG000002158 | chr13 | Erap1 | -0.005747367 | 0.988496579 |
| ENSMUSG000002506 | chr19 | Slk | 0.009225175 | 0.988496579 |
| ENSMUSG000001625 | chr2 | Ctsz | 0.008955062 | 0.988496579 |
| ENSMUSG000005570 | chr8 | Klhl26 | 0.006368893 | 0.988496579 |
| ENSMUSG000004896 | chr7 | Mrgpre | 0.010774181 | 0.988496579 |
| ENSMUSG0000004311 | chr2 | Lrrn4 | 0.03416824 | 0.988496579 |
| ENSMUSG000002060 | chr12 | Trib2 | 0.014012591 | 0.988699985 |
| ENSMUSG000003190 | chr8 | Pla2g15 | -0.009395314 | 0.988699985 |
| ENSMUSG000005374 | chr2 | Ptrh1 | 0.018583696 | 0.988699985 |
| ENSMUSG000005675 | chr11 | Dnah9 | 0.023046979 | 0.988699985 |
| ENSMUSG000004388 | chr9 | Slc36a4 | 0.008398332 | 0.988699985 |
| ENSMUSG000006829 | chr6 | Nat8f4 | -0.020418474 | 0.988699985 |
| ENSMUSG000007946 | chr1 | Col4a3 | -0.018960026 | 0.988762971 |
| ENSMUSG000004797 | chr6 | Kcna1 | -0.017410821 | 0.988778752 |
| ENSMUSG000002909 | chr5 | Acox3 | -0.010539028 | 0.988778752 |

| | | | | |
|---|---|---|---|---|
| ENSMUSG0000005407 | chr2 | Skida1 | -0.008221452 | 0.988778752 |
| ENSMUSG0000004969 | chr2 | Tmem239 | 0.043702867 | 0.98885515 |
| ENSMUSG0000000151 | chr6 | Itfg2 | 0.008232658 | 0.98885515 |
| ENSMUSG0000003978 | chr16 | Zfp597 | -0.008572549 | 0.98890153 |
| ENSMUSG0000004930 | chr9 | Ccdc71 | -0.007793904 | 0.989060411 |
| ENSMUSG0000005939 | chr13 | Nkapl | 0.030122122 | 0.989088186 |
| ENSMUSG0000003219 | chr9 | Ldlr | -0.019359333 | 0.98918016 |
| ENSMUSG0000003945 | chr16 | Ppl | -0.028507365 | 0.98918016 |
| ENSMUSG0000002090 | chr11 | Cfap52 | -0.038938194 | 0.98918016 |
| ENSMUSG0000006175 | chr6 | Akr1b10 | -0.012984042 | 0.98918016 |
| ENSMUSG0000007969 | chrX | Ssxb16 | -0.036275437 | 0.98918016 |
| ENSMUSG0000005210 | chr18 | Gnpda1 | -0.012662032 | 0.98918016 |
| ENSMUSG0000004086 | chr12 | Begain | -0.013001237 | 0.98918016 |
| ENSMUSG0000003083 | chr7 | Itgal | -0.028786541 | 0.98918016 |
| ENSMUSG0000002415 | chr17 | Hagh | -0.009842011 | 0.98918016 |
| ENSMUSG0000000630 | chr7 | Kmt2b | 0.011184011 | 0.98918016 |
| ENSMUSG0000000152 | chr6 | Nrip2 | -0.014361555 | 0.98918016 |
| ENSMUSG0000000634 | chr10 | Ggt1 | 0.042504174 | 0.98918016 |
| ENSMUSG0000005482 | chr8 | Nsd3 | -0.007517295 | 0.98918016 |
| ENSMUSG0000007867 | chr7 | Chd2 | -0.009087674 | 0.98918016 |
| ENSMUSG0000002805 | chr3 | Ash1l | -0.009850214 | 0.98918016 |
| ENSMUSG0000005998 | chr7 | Taok2 | -0.006547747 | 0.98918016 |
| ENSMUSG0000003087 | chr7 | Ubfd1 | -0.007538553 | 0.98918016 |
| ENSMUSG0000000013 | chr7 | Xpo6 | -0.006414992 | 0.98918016 |
| ENSMUSG0000002832 | chr4 | Exosc3 | -0.010285804 | 0.98918016 |
| ENSMUSG0000005096 | chr11 | Prkca | 0.008018883 | 0.98918016 |
| ENSMUSG0000003046 | chr7 | Zfp719 | -0.010197871 | 0.98918016 |
| ENSMUSG0000004611 | chr9 | Cep295 | 0.012192518 | 0.98918016 |
| ENSMUSG0000003119 | chrX | Fundc2 | 0.01005066 | 0.98918016 |
| ENSMUSG0000006325 | chr8 | Scoc | -0.005849297 | 0.98918016 |
| ENSMUSG0000003694 | chr5 | Trappc14 | -0.00947868 | 0.98918016 |
| ENSMUSG0000002236 | chr15 | Has2 | 0.026196052 | 0.98918016 |
| ENSMUSG0000005181 | chr7 | Cox6b2 | 0.013108442 | 0.98918016 |
| ENSMUSG0000003258 | chr9 | Rbm6 | -0.005433368 | 0.98918016 |
| ENSMUSG0000004566 | chr19 | Cdc42ep2 | -0.007540501 | 0.98918016 |
| ENSMUSG0000004219 | chr1 | Zfp451 | -0.006525441 | 0.98918016 |
| ENSMUSG0000009605 | chr10 | Syne1 | 0.011491688 | 0.98918016 |
| ENSMUSG0000007910 | chr12 | Prps1l3 | -0.008213587 | 0.98918016 |
| ENSMUSG0000004195 | chr4 | Tnfrsf18 | 0.015377448 | 0.98918016 |
| ENSMUSG0000002271 | chr16 | Dgcr8 | -0.007742274 | 0.98918016 |
| ENSMUSG0000002767 | chr3 | Pex5l | 0.011205219 | 0.98918016 |
| ENSMUSG0000009133 | chr2 | Eid1 | 0.007350561 | 0.98918016 |
| ENSMUSG0000003257 | chr9 | Hemk1 | -0.010811493 | 0.98918016 |
| ENSMUSG0000002855 | chr4 | Dock7 | 0.007772302 | 0.98918016 |
| ENSMUSG0000003153 | chr8 | Ap3m2 | -0.007081327 | 0.98918016 |
| ENSMUSG0000001501 | chr8 | Acsf3 | 0.008255845 | 0.98918016 |
| ENSMUSG0000002039 | chr11 | Ube2b | 0.008467153 | 0.98918016 |
| ENSMUSG0000002612 | chr1 | Imp4 | 0.007288253 | 0.98918016 |
| ENSMUSG0000002957 | chr5 | Mmab | -0.006324256 | 0.98918016 |
| ENSMUSG0000003978 | chr11 | Cep131 | 0.009843731 | 0.98918016 |

| | | | | |
|---|---|---|---|---|
| ENSMUSG0000000190 | chr8 | Trmt1 | 0.004790356 | 0.98918016 |
| ENSMUSG0000004631 | chr11 | Zfp62 | 0.006962567 | 0.98918016 |
| ENSMUSG0000002523 | chr9 | Limd1 | 0.006720595 | 0.98918016 |
| ENSMUSG0000003609 | chr10 | Vezt | 0.007010532 | 0.98918016 |
| ENSMUSG0000003253 | chr9 | Cck | -0.018573636 | 0.98918016 |
| ENSMUSG0000000210 | chr2 | Rapsn | 0.029967039 | 0.98918016 |
| ENSMUSG0000000248 | chr16 | Tiam1 | 0.012161721 | 0.98918016 |
| ENSMUSG0000004052 | chr10 | Tsfm | 0.012665569 | 0.98918016 |
| ENSMUSG0000004593 | chr19 | Ifit2 | 0.008971043 | 0.98918016 |
| ENSMUSG0000002973 | chr6 | Tpk1 | -0.012481252 | 0.98918016 |
| ENSMUSG0000000956 | chr2 | Tor2a | 0.010563577 | 0.98918016 |
| ENSMUSG0000009216 | chr14 | Gm3696 | 0.035619667 | 0.98918016 |
| ENSMUSG0000005459 | chr7 | Oscar | -0.035613897 | 0.989213426 |
| ENSMUSG0000005844 | chr9 | Map2k5 | 0.00826768 | 0.989213426 |
| ENSMUSG0000002861 | chr4 | Tceanc2 | 0.007102742 | 0.989228538 |
| ENSMUSG0000003138 | chrX | Hcfc1 | -0.009213428 | 0.989290801 |
| ENSMUSG0000000569 | chr8 | Ctcf | -0.005666493 | 0.989290801 |
| ENSMUSG0000000030 | chr11 | Pemt | -0.018365481 | 0.989290801 |
| ENSMUSG0000002819 | chr3 | Bcl10 | 0.009524879 | 0.989290801 |
| ENSMUSG0000004665 | chr5 | Zfp316 | -0.006129512 | 0.989290801 |
| ENSMUSG0000003257 | chr9 | Pik3r4 | 0.006099188 | 0.989290801 |
| ENSMUSG0000002059 | chr12 | Snx13 | 0.004782539 | 0.989290801 |
| ENSMUSG0000000499 | chr8 | Mri1 | 0.00763387 | 0.989290801 |
| ENSMUSG0000007479 | chr2 | Slc4a11 | 0.036681692 | 0.989343147 |
| ENSMUSG0000002799 | chr3 | Casp6 | -0.013210585 | 0.989343147 |
| ENSMUSG0000003134 | chrX | Cetn2 | -0.008920097 | 0.989343147 |
| ENSMUSG0000003175 | chr8 | Cdyl2 | -0.012401942 | 0.989343147 |
| ENSMUSG0000003003 | chr6 | Ino80b | -0.010060722 | 0.989343147 |
| ENSMUSG0000005169 | chr6 | Pcbp1 | -0.00878622 | 0.989343147 |
| ENSMUSG0000006604 | chr4 | Phactr4 | -0.007716235 | 0.989343147 |
| ENSMUSG0000005580 | chr7 | Dnaaf3 | -0.013547846 | 0.989343147 |
| ENSMUSG0000005574 | chr15 | Rtl6 | -0.00631435 | 0.989343147 |
| ENSMUSG0000001502 | chr8 | Ddx19a | 0.007893339 | 0.989343147 |
| ENSMUSG0000004968 | chr11 | Urgcp | -0.004778318 | 0.989343147 |
| ENSMUSG0000002169 | chr13 | Jmy | -0.008142337 | 0.989343147 |
| ENSMUSG0000004479 | chr9 | Setd2 | 0.007772762 | 0.989343147 |
| ENSMUSG0000002098 | chr12 | Sec23a | 0.004459982 | 0.989343147 |
| ENSMUSG0000002634 | chr1 | Tmem163 | 0.029594103 | 0.989343147 |
| ENSMUSG0000003109 | chrX | Dock11 | -0.007238119 | 0.989343147 |
| ENSMUSG0000002053 | chr11 | Ggnbp2 | 0.005007513 | 0.989343147 |
| ENSMUSG0000002829 | chr4 | Rars2 | 0.00636726 | 0.989343147 |
| ENSMUSG0000000132 | chr11 | Srr | 0.005259654 | 0.989343147 |
| ENSMUSG0000002121 | chr7 | Tshz3 | 0.007258388 | 0.989343147 |
| ENSMUSG0000000483 | chr11 | Grap | -0.014428572 | 0.989343147 |
| ENSMUSG0000002274 | chr16 | Cpox | -0.007780181 | 0.989519778 |
| ENSMUSG0000002591 | chr1 | Mybl1 | -0.013250314 | 0.989530274 |
| ENSMUSG0000003760 | chr5 | Adgrl3 | -0.010590188 | 0.989530274 |
| ENSMUSG0000002302 | chr15 | Dip2b | -0.012077152 | 0.989530274 |
| ENSMUSG0000003206 | chr9 | Dixdc1 | 0.00918933 | 0.989530274 |
| ENSMUSG0000003057 | chr7 | Tyrobp | -0.011709648 | 0.989530274 |

| | | | | |
|---|---|---|---|---|
| ENSMUSG0000005554 | chr11 | Timd4 | -0.020669778 | 0.989530274 |
| ENSMUSG0000002459 | chr18 | C330018D20Rik | 0.010433605 | 0.989530274 |
| ENSMUSG0000004351 | chrX | Rai2 | -0.008094479 | 0.989530274 |
| ENSMUSG0000003076 | chr7 | Arhgap17 | -0.005887322 | 0.989530274 |
| ENSMUSG0000002423 | chr18 | Cul2 | -0.005883913 | 0.989530274 |
| ENSMUSG0000000780 | chr1 | Twist2 | -0.018604764 | 0.989530274 |
| ENSMUSG0000002725 | chr2 | Adal | -0.006843669 | 0.989530274 |
| ENSMUSG0000002720 | chr2 | Dut | 0.01115253 | 0.989530274 |
| ENSMUSG0000005881 | chr7 | Pirb | -0.01367722 | 0.989530274 |
| ENSMUSG0000002742 | chr2 | Mgme1 | 0.009829189 | 0.989530274 |
| ENSMUSG0000005070 | chr7 | Ftl1 | 0.007473493 | 0.989530274 |
| ENSMUSG0000005581 | chr17 | Mta3 | 0.004711359 | 0.989530274 |
| ENSMUSG0000002784 | chr3 | Ptpn22 | 0.022928784 | 0.989530274 |
| ENSMUSG0000003842 | chr4 | Hdhd3 | -0.012684864 | 0.989530274 |
| ENSMUSG0000004419 | chr10 | S1pr4 | 0.032876967 | 0.989530274 |
| ENSMUSG0000005170 | chr1 | Tmem198 | 0.012324257 | 0.989530274 |
| ENSMUSG0000007297 | chr12 | Adam4 | -0.024725834 | 0.989530274 |
| ENSMUSG0000011867 | chr15 | Eppk1 | -0.018861009 | 0.989530274 |
| ENSMUSG0000004196 | chr2 | Dcaf17 | -0.008057594 | 0.989708713 |
| ENSMUSG0000003398 | chr3 | Gria2 | 0.009846969 | 0.989708713 |
| ENSMUSG0000002798 | chr3 | Rnpc3 | -0.00660702 | 0.989708713 |
| ENSMUSG0000002522 | chr19 | Fbxl15 | 0.012465849 | 0.989708713 |
| ENSMUSG0000003779 | chr11 | Phf12 | 0.006878005 | 0.989708713 |
| ENSMUSG0000002438 | chr18 | Map3k2 | -0.007236037 | 0.989708713 |
| ENSMUSG0000004174 | chr6 | Pde3a | 0.008627595 | 0.989708713 |
| ENSMUSG0000002757 | chr2 | Col9a3 | 0.020419591 | 0.989708713 |
| ENSMUSG0000007323 | chr5 | Fam237b | -0.020145205 | 0.989763345 |
| ENSMUSG0000003876 | chr3 | Gabpb2 | -0.007413851 | 0.989763345 |
| ENSMUSG0000002059 | chr12 | Nrcam | 0.008822949 | 0.989763345 |
| ENSMUSG0000002261 | chr15 | Mapk12 | -0.008930907 | 0.989763345 |
| ENSMUSG0000002453 | chr18 | Sncaip | -0.007198825 | 0.989763345 |
| ENSMUSG0000006102 | chr7 | Clasrp | 0.00907213 | 0.989763345 |
| ENSMUSG0000005844 | chr6 | Znrf2 | -0.005265481 | 0.989763345 |
| ENSMUSG0000003189 | chr8 | Ctrl | -0.027547324 | 0.989763345 |
| ENSMUSG0000003443 | chr6 | Zxdc | 0.005869055 | 0.989763345 |
| ENSMUSG0000001562 | chr2 | Gata5 | -0.020994804 | 0.989763345 |
| ENSMUSG0000000173 | chr5 | Cldn15 | -0.026721416 | 0.989872264 |
| ENSMUSG0000002874 | chr4 | Akr7a5 | -0.009493206 | 0.989872264 |
| ENSMUSG0000002196 | chr14 | Zdhhc20 | 0.004840809 | 0.989872264 |
| ENSMUSG0000004091 | chr1 | Slc19a2 | 0.007923006 | 0.989872264 |
| ENSMUSG0000002730 | chr2 | Nusap1 | -0.015825251 | 0.98995768 |
| ENSMUSG0000003479 | chr10 | Gna15 | -0.029062462 | 0.98995768 |
| ENSMUSG0000006099 | chr15 | Copz1 | -0.005589009 | 0.98995768 |
| ENSMUSG0000002698 | chr2 | Baz2b | 0.007307467 | 0.990159481 |
| ENSMUSG0000002254 | chr16 | Ercc4 | -0.00668761 | 0.990264478 |
| ENSMUSG0000001839 | chr11 | Septin8 | -0.00596956 | 0.990289478 |
| ENSMUSG0000002285 | chr16 | Tmem41a | 0.008962849 | 0.990289478 |
| ENSMUSG0000003136 | chrX | Ap1s2 | -0.004812861 | 0.990519114 |
| ENSMUSG0000002199 | chr14 | Wnt5a | -0.012987957 | 0.990582546 |
| ENSMUSG0000004694 | chr6 | Adck2 | -0.009477627 | 0.990584903 |

| | | | | |
|---|---|---|---|---|
| ENSMUSG0000004348 | chr12 | Acot6 | 0.006677939 | 0.990584903 |
| ENSMUSG0000003517 | chr19 | Saxo4 | 0.046656557 | 0.990584903 |
| ENSMUSG0000007064 | chr1 | Ren1 | -0.036018117 | 0.990613532 |
| ENSMUSG0000004260 | chr5 | Atxn2 | 0.006146264 | 0.990613532 |
| ENSMUSG0000007486 | chr10 | BC025920 | 0.014397941 | 0.990613532 |
| ENSMUSG0000011264 | chr10 | Gm32687 | 0.046927078 | 0.990726759 |
| ENSMUSG0000004250 | chr6 | Sdhaf3 | -0.010738232 | 0.990726759 |
| ENSMUSG0000001598 | chr7 | Lrrc27 | -0.011564309 | 0.990726759 |
| ENSMUSG0000002071 | chr11 | Polg2 | -0.00918449 | 0.990829888 |
| ENSMUSG0000002967 | chr5 | Auts2 | 0.010660464 | 0.990829888 |
| ENSMUSG0000003008 | chr6 | Plxna1 | 0.005585021 | 0.990829888 |
| ENSMUSG0000009173 | chr9 | Gpr62 | 0.021437233 | 0.990829888 |
| ENSMUSG0000006256 | chr12 | Cys1 | -0.027528303 | 0.990997353 |
| ENSMUSG0000002622 | chr1 | Htr2b | -0.008763995 | 0.991029999 |
| ENSMUSG0000003017 | chr6 | Erc1 | -0.008032766 | 0.991029999 |
| ENSMUSG0000006304 | chr8 | Ing2 | 0.007057678 | 0.991029999 |
| ENSMUSG0000000377 | chr18 | Brd8 | -0.004096614 | 0.991029999 |
| ENSMUSG0000002244 | chr15 | Ttll1 | 0.005180936 | 0.991029999 |
| ENSMUSG0000002399 | chr17 | Kif6 | -0.016044082 | 0.991034739 |
| ENSMUSG0000004864 | chr2 | Exd1 | -0.01525066 | 0.991063394 |
| ENSMUSG0000007088 | chr2 | Cfap210 | -0.008826325 | 0.991063394 |
| ENSMUSG0000002542 | chr18 | Hdhd2 | 0.004220891 | 0.991063394 |
| ENSMUSG0000005122 | chrX | Tceanc | 0.014306064 | 0.991071369 |
| ENSMUSG0000004129 | chr17 | Adgrf1 | 0.02806216 | 0.991071369 |
| ENSMUSG0000002533 | chrX | Kdm5c | -0.004842287 | 0.991071369 |
| ENSMUSG0000007135 | chr10 | Tbpl1 | -0.005285814 | 0.991071369 |
| ENSMUSG0000004800 | chr1 | Gigyf2 | 0.004818472 | 0.991071369 |
| ENSMUSG0000002823 | chr4 | Atp6v0d2 | -0.026769683 | 0.991119477 |
| ENSMUSG0000002624 | chr1 | Nppc | -0.028123551 | 0.991119477 |
| ENSMUSG0000005812 | chr14 | Tpm3-rs7 | 0.019545038 | 0.991129457 |
| ENSMUSG0000001817 | chr11 | Vmp1 | -0.006353933 | 0.991129457 |
| ENSMUSG0000003258 | chr9 | Mst1r | -0.008667156 | 0.991129457 |
| ENSMUSG0000004350 | chr15 | Lgals2 | 0.020228393 | 0.991275842 |
| ENSMUSG0000002671 | chr2 | Mrc1 | -0.011935716 | 0.991275842 |
| ENSMUSG0000001884 | chr11 | Fndc8 | -0.027386254 | 0.991275842 |
| ENSMUSG0000004817 | chr1 | Tmem81 | -0.010652111 | 0.991275842 |
| ENSMUSG0000004816 | chr5 | Selplg | -0.013438405 | 0.991275842 |
| ENSMUSG0000002619 | chr1 | Atic | -0.006147618 | 0.991275842 |
| ENSMUSG0000007893 | chr8 | Pdf | 0.022278849 | 0.991275842 |
| ENSMUSG0000001741 | chr2 | Arl5b | -0.005757808 | 0.991275842 |
| ENSMUSG0000003329 | chr4 | Ptprf | -0.005869991 | 0.991275842 |
| ENSMUSG0000002407 | chr17 | Vit | -0.008423463 | 0.991275842 |
| ENSMUSG0000003183 | chr8 | Pik3r2 | -0.005429163 | 0.991275842 |
| ENSMUSG0000002081 | chr11 | Rabep1 | -0.004714406 | 0.991275842 |
| ENSMUSG0000003026 | chr6 | Thumpd3 | -0.005862473 | 0.991275842 |
| ENSMUSG0000001009 | chr19 | Nxf1 | -0.003635534 | 0.991275842 |
| ENSMUSG0000003086 | chr7 | Dctn5 | 0.004139584 | 0.991275842 |
| ENSMUSG0000002677 | chr2 | Spopl | 0.005417805 | 0.991275842 |
| ENSMUSG0000005115 | chrX | Cited1 | -0.017646467 | 0.991275842 |
| ENSMUSG0000002433 | chr17 | Brd2 | -0.004412135 | 0.991321716 |

| | | | | |
|---|---|---|---|---|
| ENSMUSG0000005505 | chr10 | Nfic | 0.008161978 | 0.991631788 |
| ENSMUSG0000005660 | chr8 | Chd9 | -0.005653436 | 0.991652137 |
| ENSMUSG0000005547 | chr17 | Alk | 0.008419858 | 0.99165524 |
| ENSMUSG0000003725 | chr8 | Pomk | -0.005526143 | 0.991661141 |
| ENSMUSG0000003945 | chr16 | Morc3 | -0.004732315 | 0.991661141 |
| ENSMUSG0000000012 | chr11 | Wnt9a | 0.008802093 | 0.991661141 |
| ENSMUSG0000005741 | chr17 | Antkmt | -0.006271596 | 0.991661141 |
| ENSMUSG0000002473 | chr19 | Tmem132a | 0.009968554 | 0.991765184 |
| ENSMUSG0000002965 | chr5 | Slc46a3 | 0.025887196 | 0.991771533 |
| ENSMUSG0000006989 | chr8 | Atxn1l | -0.005940421 | 0.991771533 |
| ENSMUSG0000006733 | chr1 | Bmpr2 | -0.005777541 | 0.991771533 |
| ENSMUSG0000002724 | chr2 | Atg13 | 0.004865926 | 0.991771533 |
| ENSMUSG0000003927 | chr4 | Megf9 | 0.005753101 | 0.991771533 |
| ENSMUSG0000002122 | chr12 | Numb | -0.003873435 | 0.991771533 |
| ENSMUSG0000000340 | chr9 | Prkcsh | 0.004823257 | 0.991771533 |
| ENSMUSG0000008987 | chr1 | Rps6kc1 | 0.005215005 | 0.991771533 |
| ENSMUSG0000002681 | chr2 | Ak1 | 0.004351566 | 0.991771533 |
| ENSMUSG0000003466 | chr11 | Itga2b | 0.011186429 | 0.991771533 |
| ENSMUSG0000002551 | chr7 | Tspan4 | -0.013180619 | 0.991771533 |
| ENSMUSG0000010370 | chr18 | Pcdha6 | -0.025446281 | 0.991771533 |
| ENSMUSG0000004022 | chrX | Gpr34 | -0.008668824 | 0.991807315 |
| ENSMUSG0000003028 | chr6 | Il17rc | 0.006724949 | 0.991807315 |
| ENSMUSG0000003740 | chr8 | Htra4 | -0.040121902 | 0.991809759 |
| ENSMUSG0000003492 | chr4 | Dhcr24 | -0.009813834 | 0.991809759 |
| ENSMUSG0000000174 | chr7 | Il16 | -0.011968519 | 0.991809759 |
| ENSMUSG0000003035 | chr6 | Fkbp4 | 0.007050824 | 0.991809759 |
| ENSMUSG0000002065 | chr12 | Klf11 | -0.009262007 | 0.991809759 |
| ENSMUSG0000004275 | chr7 | Apobr | 0.018477321 | 0.991809759 |
| ENSMUSG0000002224 | chr15 | Amacr | -0.007435903 | 0.991809759 |
| ENSMUSG0000002723 | chr2 | Frmd5 | -0.007725295 | 0.991809759 |
| ENSMUSG0000003461 | chr11 | Pik3ip1 | 0.008625088 | 0.991809759 |
| ENSMUSG0000003163 | chr8 | Lrp2bp | -0.016483127 | 0.991809759 |
| ENSMUSG0000002624 | chr1 | Ecel1 | -0.025866002 | 0.991809759 |
| ENSMUSG0000003680 | chr15 | Fam135b | -0.019265479 | 0.991809759 |
| ENSMUSG0000004262 | chr17 | Safb2 | -0.004681249 | 0.991809759 |
| ENSMUSG0000003388 | chr3 | Rbm46 | -0.020992495 | 0.991809759 |
| ENSMUSG0000007965 | chr6 | Prrt4 | -0.010464433 | 0.991809759 |
| ENSMUSG0000003228 | chr9 | Imp3 | 0.007450649 | 0.991809759 |
| ENSMUSG0000006866 | chr16 | Clec16a | -0.006108447 | 0.991809759 |
| ENSMUSG0000002181 | chr14 | Zswim8 | 0.005317818 | 0.991809759 |
| ENSMUSG0000004341 | chr4 | Usp48 | -0.004693872 | 0.991809759 |
| ENSMUSG0000003237 | chr9 | Aph1b | 0.005728677 | 0.991809759 |
| ENSMUSG0000002641 | chr1 | Tmem9 | -0.004745022 | 0.991809759 |
| ENSMUSG0000004897 | chrX | C1galt1c1 | 0.006672979 | 0.991809759 |
| ENSMUSG0000009679 | chr10 | Zfp433 | 0.015489355 | 0.991809759 |
| ENSMUSG0000001896 | chr5 | Ywhah | 0.005237694 | 0.991809759 |
| ENSMUSG0000000031 | chr11 | Bcl6b | 0.012080235 | 0.991809759 |
| ENSMUSG0000009180 | chr12 | Cox16 | -0.007449544 | 0.991809759 |
| ENSMUSG0000004123 | chr13 | Vps41 | -0.004089684 | 0.991809759 |
| ENSMUSG0000004185 | chr1 | Mcm3 | -0.00904863 | 0.991809759 |

| | | | | |
|---|---|---|---|---|
| ENSMUSG0000002122 | chr12 | Dcaf4 | -0.006194543 | 0.991809759 |
| ENSMUSG0000002941 | chr5 | Slc15a4 | -0.005594268 | 0.991809759 |
| ENSMUSG0000004980 | chr11 | Sertad2 | 0.005496835 | 0.991809759 |
| ENSMUSG0000003031 | chr6 | Tamm41 | -0.006459819 | 0.991809759 |
| ENSMUSG0000007867 | chr11 | Casc3 | -0.003274211 | 0.991809759 |
| ENSMUSG0000005888 | chr18 | Zfp516 | -0.004885689 | 0.991809759 |
| ENSMUSG0000003416 | chr12 | Irf2bpl | 0.00440892 | 0.991809759 |
| ENSMUSG0000003183 | chr8 | Dnaaf1 | 0.037534173 | 0.991809759 |
| ENSMUSG0000000194 | chr9 | Siae | -0.003835908 | 0.991809759 |
| ENSMUSG0000002149 | chr13 | Ddx41 | -0.004909411 | 0.991809759 |
| ENSMUSG0000006111 | chr5 | Dnajc30 | -0.006474374 | 0.991809759 |
| ENSMUSG0000000366 | chr2 | Snrnp200 | 0.004857853 | 0.991809759 |
| ENSMUSG0000003585 | chr5 | Ythdc1 | -0.002880216 | 0.991809759 |
| ENSMUSG0000002198 | chr14 | Mtmr6 | -0.003253228 | 0.991809759 |
| ENSMUSG0000002496 | chr19 | Fermt3 | -0.010754497 | 0.991809759 |
| ENSMUSG0000004812 | chr19 | Entpd1 | -0.004562262 | 0.991809759 |
| ENSMUSG0000007186 | chr18 | Lrrtm2 | -0.006907388 | 0.991809759 |
| ENSMUSG0000003661 | chr12 | Eipr1 | -0.006233534 | 0.991809759 |
| ENSMUSG0000006335 | chr16 | Mapk1 | 0.002857483 | 0.991809759 |
| ENSMUSG0000002914 | chr5 | Slc4a1ap | 0.003717168 | 0.991809759 |
| ENSMUSG0000002424 | chr17 | Map4k3 | 0.003856067 | 0.991809759 |
| ENSMUSG0000002468 | chr19 | Osbp | 0.002548058 | 0.991809759 |
| ENSMUSG0000002297 | chr16 | Cfap298 | 0.008710059 | 0.991809759 |
| ENSMUSG0000002008 | chr10 | Macroh2a2 | 0.005109794 | 0.991809759 |
| ENSMUSG0000004270 | chr4 | Dnali1 | 0.009096584 | 0.991809759 |
| ENSMUSG0000007854 | chr17 | Zfp995 | -0.007197169 | 0.991809759 |
| ENSMUSG0000003068 | chr7 | Cdipt | 0.002674242 | 0.991809759 |
| ENSMUSG0000002691 | chr2 | Brd3 | 0.003476087 | 0.991809759 |
| ENSMUSG0000003312 | chr15 | Gga1 | 0.00283792 | 0.991809759 |
| ENSMUSG0000000426 | chr6 | Atn1 | 0.004782263 | 0.991809759 |
| ENSMUSG0000002091 | chr11 | Kat2a | -0.004479999 | 0.991809759 |
| ENSMUSG0000005031 | chr16 | Nsun3 | 0.006103032 | 0.991809759 |
| ENSMUSG0000000176 | chr6 | Tspan33 | 0.009228244 | 0.991809759 |
| ENSMUSG0000000407 | chr16 | Cdip1 | 0.002774587 | 0.991809759 |
| ENSMUSG0000000230 | chr17 | Daxx | 0.004124637 | 0.991809759 |
| ENSMUSG0000002168 | chr13 | Ap3b1 | -0.005056842 | 0.991809759 |
| ENSMUSG0000002395 | chr17 | Polh | 0.007269633 | 0.991809759 |
| ENSMUSG0000001508 | chr2 | C8g | 0.011511084 | 0.991809759 |
| ENSMUSG0000004478 | chr7 | Zfp36 | -0.005979101 | 0.991809759 |
| ENSMUSG0000004248 | chr4 | Clspn | -0.026787864 | 0.991809759 |
| ENSMUSG0000005999 | chr11 | Atxn7l3 | 0.004991932 | 0.991809759 |
| ENSMUSG0000002707 | chr2 | Lrp2 | 0.017865232 | 0.991809759 |
| ENSMUSG0000002926 | chr5 | Pigg | 0.004500745 | 0.991809759 |
| ENSMUSG0000002054 | chr11 | Cox11 | 0.009545823 | 0.991809759 |
| ENSMUSG0000004964 | chr4 | Gpr3 | 0.020161779 | 0.991809759 |
| ENSMUSG0000003798 | chr6 | Neurod6 | 0.010276561 | 0.991809759 |
| ENSMUSG0000001976 | chr11 | Krt10 | 0.008592247 | 0.991809759 |
| ENSMUSG0000002611 | chr1 | Zap70 | 0.016482718 | 0.991809759 |
| ENSMUSG0000000122 | chr17 | Uhrf1 | 0.019166913 | 0.991809759 |
| ENSMUSG0000006832 | chr6 | Tlx2 | 0.007055285 | 0.991809759 |

| | | | | |
|---|---|---|---|---|
| ENSMUSG0000009397 | chr14 | Gm2237 | 0.013864908 | 0.991809759 |
| ENSMUSG0000003581 | chr19 | Plekhs1 | 0.026514292 | 0.991809759 |
| ENSMUSG0000006737 | chr17 | B3galt4 | 0.010737893 | 0.991841802 |
| ENSMUSG0000003229 | chr9 | Commd4 | -0.003933407 | 0.991844928 |
| ENSMUSG0000002403 | chr17 | Wdr4 | 0.00403846 | 0.991844928 |
| ENSMUSG0000002681 | chr2 | Dpm2 | 0.003861198 | 0.991844928 |
| ENSMUSG0000002272 | chr16 | Trmt2a | -0.003276785 | 0.991869352 |
| ENSMUSG0000005159 | chr1 | Map3k19 | 0.019506558 | 0.992025559 |
| ENSMUSG0000006304 | chr7 | Zfp780b | -0.005889554 | 0.992090296 |
| ENSMUSG0000000103 | chr11 | B9d1 | -0.006727575 | 0.992211391 |
| ENSMUSG0000002197 | chr14 | Extl3 | -0.002667266 | 0.992211391 |
| ENSMUSG0000006098 | chr1 | Tdrd5 | -0.03033187 | 0.992211391 |
| ENSMUSG0000006662 | chr5 | Tecpr1 | 0.004239293 | 0.992221327 |
| ENSMUSG0000003400 | chr9 | Scaper | -0.004101956 | 0.992223975 |
| ENSMUSG0000003305 | chr19 | Cfap95 | 0.024962075 | 0.992232311 |
| ENSMUSG0000007410 | chr9 | Rbm15b | -0.003259757 | 0.992236041 |
| ENSMUSG0000003641 | chr9 | Matcap2 | -0.006336689 | 0.992236041 |
| ENSMUSG0000002025 | chr10 | Nopchap1 | -0.004025189 | 0.992236041 |
| ENSMUSG0000002677 | chr2 | Mastl | -0.015242359 | 0.992246378 |
| ENSMUSG0000002667 | chr1 | Hsd17b7 | 0.008910058 | 0.992246378 |
| ENSMUSG0000007348 | chr1 | Ifi204 | -0.007080013 | 0.992246378 |
| ENSMUSG0000003605 | chr8 | Sugp2 | -0.004124832 | 0.992246378 |
| ENSMUSG0000002698 | chr2 | Hnmt | 0.005684091 | 0.992246378 |
| ENSMUSG0000004036 | chr11 | Trim41 | -0.003963044 | 0.992246378 |
| ENSMUSG0000003517 | chr19 | Ccdc186 | -0.003330007 | 0.992246378 |
| ENSMUSG0000002466 | chr19 | Tmem216 | 0.006331599 | 0.992246378 |
| ENSMUSG0000003272 | chr7 | Folr2 | -0.009996824 | 0.992246378 |
| ENSMUSG0000002801 | chr3 | Ppa2 | 0.00423277 | 0.992246378 |
| ENSMUSG0000001850 | chr11 | Ncor1 | 0.004363105 | 0.992246378 |
| ENSMUSG0000007570 | chr3 | Selenot | 0.00265456 | 0.992246378 |
| ENSMUSG0000005306 | chr16 | Jam2 | -0.006368956 | 0.992246378 |
| ENSMUSG0000003741 | chr8 | Ranbp10 | 0.003441158 | 0.992246378 |
| ENSMUSG0000003427 | chr2 | Dnajc17 | 0.00547481 | 0.992246378 |
| ENSMUSG0000002284 | chr16 | Pdia5 | -0.006104224 | 0.992246378 |
| ENSMUSG0000004016 | chr6 | 1700034J05Rik | 0.017978756 | 0.992246378 |
| ENSMUSG0000005634 | chr11 | Usp34 | -0.004656006 | 0.992258134 |
| ENSMUSG0000003846 | chr2 | Chmp4b | 0.003084908 | 0.992258134 |
| ENSMUSG0000003476 | chr19 | Dusp5 | -0.010292215 | 0.992389581 |
| ENSMUSG0000002168 | chr13 | Aggf1 | -0.00344938 | 0.992389581 |
| ENSMUSG0000003261 | chr9 | Usp4 | 0.002837622 | 0.992389581 |
| ENSMUSG0000005050 | chr9 | Fbxl22 | -0.007128112 | 0.992496725 |
| ENSMUSG0000002517 | chr19 | Hoga1 | -0.010274343 | 0.992496725 |
| ENSMUSG0000001449 | chr10 | Ankrd52 | 0.005164786 | 0.992496725 |
| ENSMUSG0000002540 | chr10 | R3hdm2 | -0.003395519 | 0.992496725 |
| ENSMUSG0000006782 | chr6 | Pex26 | 0.00490597 | 0.992496725 |
| ENSMUSG0000003696 | chr15 | Spryd3 | 0.004610753 | 0.992496725 |
| ENSMUSG0000002036 | chr11 | Cnot6 | -0.003825109 | 0.992496725 |
| ENSMUSG0000002232 | chr15 | Rida | 0.006332251 | 0.992496725 |
| ENSMUSG0000009118 | chr13 | Gm5141 | -0.005347591 | 0.992507052 |
| ENSMUSG0000003973 | chr3 | Fnbp1l | -0.003536736 | 0.992507052 |

| | | | | |
|---|---|---|---|---|
| ENSMUSG0000002916 | chr5 | Khk | -0.005790245 | 0.992688327 |
| ENSMUSG0000002712 | chr2 | Arl14ep | -0.00287129 | 0.992862898 |
| ENSMUSG0000003041 | chr7 | Dmwd | -0.00270792 | 0.992862898 |
| ENSMUSG0000001581 | chr14 | Gnrh1 | 0.021459884 | 0.992879731 |
| ENSMUSG0000002962 | chr5 | Arpc1b | 0.003355948 | 0.992883638 |
| ENSMUSG0000002290 | chr16 | Cd86 | 0.008205331 | 0.993010677 |
| ENSMUSG0000002236 | chr15 | Zhx1 | -0.003462839 | 0.993149911 |
| ENSMUSG0000002003 | chr10 | Nuak1 | 0.006508061 | 0.993234985 |
| ENSMUSG0000004285 | chr2 | Trp53rkb | -0.008289718 | 0.993234985 |
| ENSMUSG0000001998 | chr10 | Med23 | -0.002886529 | 0.993234985 |
| ENSMUSG0000003195 | chr8 | Wdr59 | -0.004142447 | 0.993234985 |
| ENSMUSG0000005744 | chr18 | Mpp7 | 0.003936648 | 0.993234985 |
| ENSMUSG0000003947 | chr16 | Ubn1 | 0.003600705 | 0.993234985 |
| ENSMUSG0000001593 | chr13 | Macroh2a1 | 0.002477623 | 0.993234985 |
| ENSMUSG0000002627 | chr1 | Ppp1r7 | -0.002672791 | 0.993234985 |
| ENSMUSG0000003303 | chr18 | Afap1l1 | -0.005843094 | 0.993234985 |
| ENSMUSG0000005135 | chr15 | Ncald | 0.006017525 | 0.993238702 |
| ENSMUSG0000002238 | chr15 | Brd1 | 0.003869998 | 0.993238702 |
| ENSMUSG0000003114 | chrX | Gpkow | -0.002929589 | 0.993238702 |
| ENSMUSG0000006679 | chr2 | Zbtb6 | 0.002979274 | 0.993238702 |
| ENSMUSG0000003753 | chr14 | Fbxo34 | 0.002510882 | 0.993238702 |
| ENSMUSG0000000038 | chr11 | Tbrg4 | -0.003351256 | 0.993282629 |
| ENSMUSG0000005145 | chr11 | Meioc | -0.015360633 | 0.993743917 |
| ENSMUSG0000003920 | chr5 | Daglb | 0.005550136 | 0.993976036 |
| ENSMUSG0000000376 | chr7 | Coq8b | 0.003897945 | 0.994170761 |
| ENSMUSG0000002817 | chr3 | Depdc1a | 0.026667233 | 0.994481395 |
| ENSMUSG0000009593 | chr13 | Nim1k | -0.006831147 | 0.994481395 |
| ENSMUSG0000007134 | chr14 | C1qtnf9 | 0.006302161 | 0.994481395 |
| ENSMUSG0000002519 | chr19 | Entpd7 | -0.004979199 | 0.994481395 |
| ENSMUSG0000009111 | chr15 | Ccdc152 | 0.009663843 | 0.994481395 |
| ENSMUSG0000002093 | chr11 | Ccdc103 | -0.010277161 | 0.994481395 |
| ENSMUSG0000000027 | chr11 | Dgke | 0.003525992 | 0.994481395 |
| ENSMUSG0000002277 | chr16 | Ppil2 | -0.002508038 | 0.994481395 |
| ENSMUSG0000002092 | chr11 | Ubtf | 0.002086858 | 0.994481395 |
| ENSMUSG0000002688 | chr2 | Dab2ip | -0.002378186 | 0.994481395 |
| ENSMUSG0000005166 | chr16 | AU021092 | -0.010508761 | 0.994501672 |
| ENSMUSG0000004228 | chr14 | Stab1 | 0.004789783 | 0.994501672 |
| ENSMUSG0000005865 | chr15 | Eif4b | 0.004500261 | 0.994501672 |
| ENSMUSG0000003155 | chr8 | Tm2d2 | -0.003227948 | 0.994501672 |
| ENSMUSG0000003191 | chr8 | Has3 | -0.004178598 | 0.994501672 |
| ENSMUSG0000009567 | chr17 | Dynlt1f | 0.004296032 | 0.994501672 |
| ENSMUSG0000002680 | chr2 | Ddx31 | -0.003685409 | 0.994587768 |
| ENSMUSG0000001509 | chr2 | Npdc1 | -0.003030239 | 0.994587768 |
| ENSMUSG0000000787 | chr11 | Tcap | 0.022491801 | 0.994659425 |
| ENSMUSG0000009393 | chr13 | Hmgcs1 | 0.006594721 | 0.994659425 |
| ENSMUSG0000004069 | chr17 | Apobec2 | -0.007997674 | 0.994659425 |
| ENSMUSG0000007028 | chr9 | Gmppb | -0.005049874 | 0.994659425 |
| ENSMUSG0000003852 | chr6 | C1s1 | -0.005224236 | 0.994659425 |
| ENSMUSG0000007172 | chr15 | Smpd5 | -0.006308907 | 0.994659425 |
| ENSMUSG0000002601 | chr1 | Carf | -0.004007909 | 0.994659425 |

| | | | | |
|---|---|---|---|---|
| ENSMUSG0000007438 | chr8 | AI429214 | -0.005190914 | 0.994659425 |
| ENSMUSG0000003328 | chr8 | Rpgrip1l | -0.003105079 | 0.994659425 |
| ENSMUSG0000003457 | chr13 | Tent4a | 0.003215088 | 0.994659425 |
| ENSMUSG0000007950 | chr2 | Cfap77 | -0.008600928 | 0.994659425 |
| ENSMUSG0000004679 | chr7 | Zfp787 | 0.002599563 | 0.994659425 |
| ENSMUSG0000003190 | chr8 | Slc7a6 | -0.003772478 | 0.994659425 |
| ENSMUSG0000004250 | chr7 | Cd2bp2 | -0.002154458 | 0.994668251 |
| ENSMUSG0000002692 | chr2 | Inpp5e | 0.002309897 | 0.994726609 |
| ENSMUSG0000005176 | chr7 | Xrcc1 | 0.00257362 | 0.994733173 |
| ENSMUSG0000007859 | chr7 | Hs3st4 | -0.007625316 | 0.994778954 |
| ENSMUSG0000007465 | chr2 | Myh7b | 0.011284339 | 0.994778954 |
| ENSMUSG0000002946 | chr5 | P2rx7 | 0.003421932 | 0.994778954 |
| ENSMUSG0000003905 | chr11 | Eme1 | 0.015384835 | 0.994778954 |
| ENSMUSG0000002659 | chr1 | Ralgps2 | -0.002496672 | 0.994778954 |
| ENSMUSG0000002991 | chr6 | Hpgds | -0.004666724 | 0.994778954 |
| ENSMUSG0000006614 | chr4 | Gm11232 | -0.010691555 | 0.994778954 |
| ENSMUSG0000004870 | chr5 | Speer4b | -0.010710206 | 0.994778954 |
| ENSMUSG0000003299 | chr1 | Chpf | 0.002692106 | 0.994778954 |
| ENSMUSG0000002504 | chr19 | Pdcd11 | 0.002171016 | 0.994778954 |
| ENSMUSG0000002427 | chr18 | Zfp397 | -0.001852565 | 0.994778954 |
| ENSMUSG0000002622 | chr1 | 4933407L21Rik | -0.007453923 | 0.994778954 |
| ENSMUSG0000002278 | chr16 | Dnm1l | 0.001738406 | 0.994778954 |
| ENSMUSG0000001366 | chr19 | Pten | -0.00313836 | 0.994778954 |
| ENSMUSG0000003201 | chr9 | Pou2f3 | -0.005221928 | 0.994778954 |
| ENSMUSG0000003520 | chr7 | Epn1 | 0.002245982 | 0.994778954 |
| ENSMUSG0000002267 | chr16 | Prkdc | 0.0029742 | 0.994778954 |
| ENSMUSG0000002405 | chr17 | Emilin2 | 0.004510134 | 0.994787094 |
| ENSMUSG0000002857 | chr4 | Ift74 | 0.002180664 | 0.994787094 |
| ENSMUSG0000003870 | chr9 | Golga4 | 0.002131464 | 0.994787094 |
| ENSMUSG0000006207 | chr10 | Lmnb2 | -0.004159234 | 0.994787094 |
| ENSMUSG0000004241 | chr17 | Nfkbil1 | 0.005010608 | 0.994877432 |
| ENSMUSG0000000629 | chr1 | Aamp | 0.002216057 | 0.994877432 |
| ENSMUSG0000002424 | chr17 | Pkdcc | -0.003664795 | 0.994877432 |
| ENSMUSG0000002876 | chr4 | Eif4g3 | 0.002262898 | 0.994928798 |
| ENSMUSG0000002412 | chr17 | Atp6v0c | -0.002042442 | 0.995017264 |
| ENSMUSG0000000091 | chr5 | Hip1r | -0.004654233 | 0.995084564 |
| ENSMUSG0000003734 | chr15 | Taf2 | -0.001788934 | 0.995084564 |
| ENSMUSG0000000627 | chr8 | Eps15l1 | -0.002337751 | 0.995088444 |
| ENSMUSG0000010702 | chr7 | Gm42715 | 0.004918893 | 0.995088444 |
| ENSMUSG0000002572 | chr17 | Pigq | 0.001405043 | 0.995088444 |
| ENSMUSG0000002855 | chr4 | Eps15 | 0.001254572 | 0.995104806 |
| ENSMUSG0000002057 | chr12 | Nampt | 0.002176415 | 0.995150077 |
| ENSMUSG0000003625 | chr12 | Pnpla8 | -0.001389835 | 0.995150077 |
| ENSMUSG0000002685 | chr2 | Usp20 | -0.002342032 | 0.995319442 |
| ENSMUSG0000002229 | chr15 | Rrm2b | 0.002176081 | 0.995815468 |
| ENSMUSG0000012160 | chrX | ENSMUSG000001 | 0.00473839 | 0.995826872 |
| ENSMUSG0000006317 | chr4 | Ift25 | 0.003252143 | 0.995826872 |
| ENSMUSG0000006923 | chr13 | Fam8a1 | -0.001218025 | 0.995826872 |
| ENSMUSG0000002929 | chr5 | Zfp326 | -0.0019976 | 0.995826872 |
| ENSMUSG0000003728 | chr15 | Galnt6 | 0.0085396 | 0.995829323 |

| | | | | |
|---|---|---|---|---|
| ENSMUSG0000003370 | chr2 | Stard9 | -0.004375429 | 0.995845452 |
| ENSMUSG0000004562 | chr16 | Pigz | -0.005235838 | 0.995845452 |
| ENSMUSG0000004579 | chr7 | Whamm | -0.001889267 | 0.995845452 |
| ENSMUSG0000002212 | chr14 | Ednrb | 0.00406447 | 0.995845452 |
| ENSMUSG0000002931 | chr5 | Coq2 | 0.002518743 | 0.995880751 |
| ENSMUSG0000005268 | chr4 | Jun | -0.003124 | 0.995890403 |
| ENSMUSG0000003825 | chr8 | Usp38 | 0.002128307 | 0.995962051 |
| ENSMUSG0000002665 | chr2 | Frmd4a | 0.001977351 | 0.995973152 |
| ENSMUSG0000003938 | chr1 | Dusp10 | 0.003509475 | 0.995997308 |
| ENSMUSG0000002397 | chr17 | Cnpy3 | -0.002237553 | 0.996149281 |
| ENSMUSG0000004581 | chr17 | Zfp36l2 | 0.002395118 | 0.996149281 |
| ENSMUSG0000005089 | chr15 | Tatdn1 | 0.00245085 | 0.996149281 |
| ENSMUSG0000007570 | chr5 | Selenoi | 0.002316462 | 0.996149281 |
| ENSMUSG0000002689 | chr2 | Lhx6 | -0.00822388 | 0.996223232 |
| ENSMUSG0000000523 | chr2 | Spc25 | -0.006431423 | 0.996228386 |
| ENSMUSG0000002217 | chr14 | Haus4 | 0.002969413 | 0.996228386 |
| ENSMUSG0000002023 | chr10 | Prmt2 | 0.002189087 | 0.996228386 |
| ENSMUSG0000002897 | chr4 | Casz1 | -0.001994659 | 0.996228386 |
| ENSMUSG0000004314 | chr16 | Tmem186 | -0.001832003 | 0.996228386 |
| ENSMUSG0000004357 | chr4 | Pars2 | 0.00236484 | 0.996228386 |
| ENSMUSG0000002034 | chr11 | Mgat1 | -0.001186219 | 0.996228386 |
| ENSMUSG0000003861 | chr7 | Phrf1 | 0.001239069 | 0.996228386 |
| ENSMUSG0000001876 | chr11 | Fxr2 | -0.000885277 | 0.996228386 |
| ENSMUSG0000002549 | chr7 | Ptdss2 | 0.001496395 | 0.996228386 |
| ENSMUSG0000004923 | chr6 | Tigd2 | -0.001857885 | 0.996228386 |
| ENSMUSG0000006107 | chr7 | Zfp143 | 0.002434018 | 0.996228386 |
| ENSMUSG0000007457 | chr2 | Cass4 | -0.005089713 | 0.996228386 |
| ENSMUSG0000000290 | chr8 | Kcnn1 | 0.002702069 | 0.996228386 |
| ENSMUSG0000004517 | chr11 | Borcs6 | -0.002583432 | 0.996228386 |
| ENSMUSG0000005086 | chr11 | Phospho1 | 0.005257814 | 0.996228386 |
| ENSMUSG0000003852 | chr7 | Tbc1d17 | -0.002011243 | 0.996665168 |
| ENSMUSG0000000234 | chr8 | Ncan | -0.003009628 | 0.996665168 |
| ENSMUSG0000005489 | chr7 | Zfp667 | -0.001430527 | 0.996665168 |
| ENSMUSG0000007276 | chr5 | Zfp1007 | -0.003357724 | 0.996665168 |
| ENSMUSG0000003736 | chr5 | Srrt | -0.001300046 | 0.996665168 |
| ENSMUSG0000003240 | chr9 | 2300009A05Rik | -0.003025632 | 0.996665168 |
| ENSMUSG0000001908 | chr7 | Slc25a22 | 0.001400778 | 0.996665168 |
| ENSMUSG0000006290 | chr15 | Hdac10 | -0.002244592 | 0.996665168 |
| ENSMUSG0000002854 | chr4 | Itgb3bp | -0.002635205 | 0.996714648 |
| ENSMUSG0000004147 | chr14 | Shld2 | -0.002157543 | 0.996854988 |
| ENSMUSG0000004177 | chr7 | Mapk1ip1 | -0.001699571 | 0.996872288 |
| ENSMUSG0000001984 | chr10 | Popdc3 | -0.002144584 | 0.996872288 |
| ENSMUSG0000001799 | chr2 | Ddx27 | -0.001431075 | 0.996872288 |
| ENSMUSG0000003185 | chr8 | Mau2 | 0.001039276 | 0.996872288 |
| ENSMUSG0000001784 | chr12 | Ppp2r5c | -0.000828831 | 0.996872288 |
| ENSMUSG0000002014 | chr11 | Dock2 | -0.003539127 | 0.996872288 |
| ENSMUSG0000003085 | chr7 | Ate1 | 0.000916449 | 0.996979017 |
| ENSMUSG0000002887 | chr4 | Rspo1 | -0.005603178 | 0.997068405 |
| ENSMUSG0000002028 | chr11 | Pex13 | 0.001277263 | 0.997068405 |
| ENSMUSG0000002670 | chr1 | Klhl20 | 0.00103114 | 0.997068405 |

| | | | | |
|---|---|---|---|---|
| ENSMUSG0000004928 | chr5 | Mblac1 | 0.002498694 | 0.997068405 |
| ENSMUSG0000003816 | chr10 | Atg5 | 0.001390579 | 0.99712565 |
| ENSMUSG0000005823 | chr7 | Usf2 | 0.00115204 | 0.99712565 |
| ENSMUSG0000007440 | chr7 | Zfp865 | 0.00132516 | 0.997281082 |
| ENSMUSG0000007435 | chr7 | Ccdc61 | 0.001597995 | 0.997281082 |
| ENSMUSG0000002163 | chr13 | Gtf2h2 | 0.001159055 | 0.997281082 |
| ENSMUSG0000002679 | chr2 | Slc2a8 | -0.00138149 | 0.997641885 |
| ENSMUSG0000002731 | chr2 | Chac1 | 0.003793582 | 0.997704333 |
| ENSMUSG0000003896 | chr11 | Pdk2 | -0.001913927 | 0.997704333 |
| ENSMUSG0000002941 | chr5 | Kntc1 | -0.005980922 | 0.997704333 |
| ENSMUSG0000004264 | chr1 | Rgsl1 | -0.004616542 | 0.997704333 |
| ENSMUSG0000003504 | chrX | Zc3h12b | 0.001471676 | 0.997704333 |
| ENSMUSG0000000056 | chr3 | Wdr77 | 0.001409698 | 0.997704333 |
| ENSMUSG0000002801 | chr3 | Egf | -0.003069893 | 0.997704333 |
| ENSMUSG0000001535 | chr1 | Cd48 | -0.001888035 | 0.997704333 |
| ENSMUSG0000007923 | chr9 | Ccdc13 | 0.002900727 | 0.997704333 |
| ENSMUSG0000004037 | chr6 | Etfrf1 | -0.001221579 | 0.997704333 |
| ENSMUSG0000003572 | chr14 | Supt16 | 0.00089337 | 0.997704333 |
| ENSMUSG0000002400 | chr17 | Cpne5 | 0.002668908 | 0.997704333 |
| ENSMUSG0000003076 | chr7 | Lcmt1 | 0.000928225 | 0.997704333 |
| ENSMUSG0000003513 | chr1 | Brinp3 | 0.002077176 | 0.997704333 |
| ENSMUSG0000006047 | chr6 | Irak2 | 0.00141359 | 0.997704333 |
| ENSMUSG0000007968 | chr10 | Ulbp1 | 0.003294582 | 0.997733806 |
| ENSMUSG0000009685 | chr10 | Gm10778 | -0.004454847 | 0.997733806 |
| ENSMUSG0000004628 | chr3 | She | 0.002146671 | 0.997733806 |
| ENSMUSG0000001700 | chr2 | Rbpjl | -0.003949419 | 0.997733806 |
| ENSMUSG0000002896 | chr5 | Cdk5 | 0.000786859 | 0.997733806 |
| ENSMUSG0000002429 | chr18 | Mib1 | -0.000863014 | 0.997733806 |
| ENSMUSG0000003186 | chr8 | Atp13a1 | -0.000661843 | 0.997733806 |
| ENSMUSG0000003074 | chr7 | Dgat2 | -0.00333127 | 0.997780729 |
| ENSMUSG0000003151 | chr8 | Gpm6a | -0.003103904 | 0.997780729 |
| ENSMUSG0000001981 | chr10 | Cep57l1 | -0.001357846 | 0.997780729 |
| ENSMUSG0000002839 | chr4 | Kdm4c | 0.000840762 | 0.997780729 |
| ENSMUSG0000003068 | chr7 | Kctd13 | 0.000729089 | 0.997780729 |
| ENSMUSG0000011762 | chr18 | Hspe1-rs1 | 0.00397175 | 0.997780729 |
| ENSMUSG0000002766 | chr2 | Slc2a10 | -0.00150004 | 0.997780729 |
| ENSMUSG0000007125 | chr13 | 2210408I21Rik | -0.000987262 | 0.998057155 |
| ENSMUSG0000000103 | chr11 | Mapk7 | 0.000947266 | 0.998096821 |
| ENSMUSG0000002132 | chr13 | Aoah | 0.002452158 | 0.998117114 |
| ENSMUSG0000002238 | chr15 | Wnt7b | -0.002737144 | 0.998135943 |
| ENSMUSG0000000227 | chr17 | Metrn | 0.001313046 | 0.99815317 |
| ENSMUSG0000003044 | chr7 | Zfp583 | 0.000933475 | 0.998721241 |
| ENSMUSG0000001833 | chr11 | Gpx3 | 0.000919545 | 0.998725229 |
| ENSMUSG0000007104 | chr17 | Rasgrp3 | -0.001301297 | 0.998820113 |
| ENSMUSG0000003015 | chr6 | Klrb1f | -0.003062148 | 0.99884761 |
| ENSMUSG0000001951 | chr5 | Ap4m1 | 0.00053252 | 0.999079279 |
| ENSMUSG0000003996 | chr4 | Zfp292 | -0.000572796 | 0.999079279 |
| ENSMUSG0000003037 | chr7 | Slc8a2 | -0.000493191 | 0.999127094 |
| ENSMUSG0000009472 | chr17 | Rnaset2b | -0.001460304 | 0.999440514 |
| ENSMUSG0000002088 | chr11 | Fbxl20 | 0.000474919 | 0.999440514 |

| | | | | |
|---|---|---|---|---|
| ENSMUSG0000009614 | chr1 | Dnah7a | -0.001164778 | 0.999440514 |
| ENSMUSG0000003957 | chr4 | Nphp4 | 0.000454296 | 0.999440514 |
| ENSMUSG0000003998 | chr11 | Cbx4 | 0.000301008 | 0.999440514 |
| ENSMUSG0000002435 | chr18 | Dnajc18 | -0.000299024 | 0.999440514 |
| ENSMUSG0000001651 | chr5 | Mtif3 | 0.000384328 | 0.999452809 |
| ENSMUSG0000003273 | chr9 | Snx33 | -0.000386385 | 0.999467424 |
| ENSMUSG0000006027 | chr7 | Ap2a1 | 0.000217812 | 0.999467424 |
| ENSMUSG0000002282 | chr16 | Rabl3 | -0.000409476 | 0.999467424 |
| ENSMUSG0000003072 | chr7 | Lipt2 | 0.000668269 | 0.999467424 |
| ENSMUSG0000007416 | chr7 | Zfp788 | -0.000301491 | 0.999626711 |
| ENSMUSG0000004331 | chr17 | D17H6S53E | -0.000334162 | 0.999626711 |
| ENSMUSG0000003135 | chrX | Zfp185 | 0.00032976 | 0.999690255 |
| ENSMUSG0000002392 | chr17 | Tbc1d5 | 0.000164339 | 0.999690255 |
| ENSMUSG0000000185 | chr2 | Nup214 | 0.000184771 | 0.999690255 |
| ENSMUSG0000004491 | chr2 | Frey1 | 0.000371727 | 0.999690255 |
| ENSMUSG0000000822 | chr2 | Scrn3 | 0.000148335 | 0.999690255 |
| ENSMUSG0000006832 | chr6 | Aup1 | 0.000139656 | 0.999690255 |
| ENSMUSG0000007876 | chr7 | Zfp566 | 0.00031125 | 0.999690255 |
| ENSMUSG0000001859 | chr11 | Mief2 | -0.000143253 | 0.999690255 |
| ENSMUSG0000002125 | chr12 | Erg28 | 0.000143146 | 0.999690255 |
| ENSMUSG0000004145 | chr4 | Tardbp | -0.000144954 | 0.999690255 |
| ENSMUSG0000003591 | chr9 | Cd276 | 0.000283197 | 0.999690255 |
| ENSMUSG0000004273 | chr12 | Ttc9 | 0.000200608 | 0.999690255 |
| ENSMUSG0000004582 | chr2 | Zswim3 | 0.000161994 | 0.999852189 |
| ENSMUSG0000000194 | chr9 | Esam | 0.000108788 | 0.999983719 |
| ENSMUSG0000002001 | chr10 | Hal | 0.000746381 | 0.999983719 |
| ENSMUSG0000002973 | chr5 | Mcm7 | 2.90E-06 | 0.999988385 |
| ENSMUSG0000003999 | chr7 | Edrf1 | 5.58E-06 | 0.999988385 |
| ENSMUSG0000002606 | chr1 | Il1rl1 | -5.84E-05 | 0.999988385 |
